# The full-length transcriptome of *Spartina alterniflora* reveals the complexity of high salt tolerance in monocotyledonous halophyte

**DOI:** 10.1101/680819

**Authors:** Wenbin Ye, Taotao Wang, Wei Wei, Shuaitong Lou, Faxiu Lan, Sheng Zhu, Qinzhen Li, Guoli Ji, Chentao Lin, Xiaohui Wu, Liuyin Ma

## Abstract

*Spartina alterniflora* (*Spartina*) is the only halophyte in the salt marsh. However, the molecular basis of its high salt tolerance remains elusive. In this study, we used PacBio full-length single molecule long-read sequencing and RNA-seq to elucidate the transcriptome dynamics of high salt tolerance in *Spartina* by salt-gradient experiments (0, 350, 500 and 800 mM NaCl). We systematically analyzed the gene expression diversity and deciphered possible roles of ion transporters, protein kinases and photosynthesis in salt tolerance. Moreover, the co-expression network analysis revealed several hub genes in salt stress regulatory networks, including protein kinases such as *SaOST1*, *SaCIPK10* and three *SaLRRs.* Furthermore, high salt stress affected the gene expression of photosynthesis through down-regulation at the transcription level and alternative splicing at the post-transcriptional level. In addition, overexpression of two *Spartina* salt-tolerant genes *SaHSP70-I* and *SaAF2* in Arabidopsis significantly promoted the salt tolerance of transgenic lines. Finally, we built the SAPacBio website for visualizing the full-length transcriptome sequences, transcription factors, ncRNAs, salt-tolerant genes, and alternative splicing events in *Spartina*. Overall, this study sheds light on the high salt tolerance mechanisms of monocotyledonous-halophyte and demonstrates the potential of *Spartina* genes for engineering salt-tolerant plants.

## INTRODUCTION

Over 20% of the world’s irrigated land suffers from soil salinization, and salt stress challenges the survival of plants through affecting their development and reproduction (Deinlein et al., 2014; Roy, Negrao, & Tester, 2014). However, only ∼2% of terrestrial plants are salt-tolerant halophytes with strong ability to survive in concentrations of NaCl over 200 mM (Mangu et al., 2015). *Spartina alterniflora* (*Spartina*) is the only halophytic plant that can survive in highly saline environment of salt marshes (Subudhi & Baisakh, 2011). Intraspecific variation studies reveal that the strong saline environmental adaptation of *Spartina alterniflora* may be largely due to the greater ion selectivity to the uptake of potassium and excluding sodium (Bradley & Morris, 1991; Smart & Barko, 1980), which results in a lower Na^+^/K^+^ ratio in *Spartina alterniflora* compared to the brackish water plant-*Spartina patens* (Hester, Mendelssohn, & McKee, 2001). Moreover, *Spartina alternifora* also develops specialized salt glands to secret ions (Nestler, 1977; Subudhi & Baisakh, 2011). Furthermore, cell membranes actively exclude salt out of cells to avoid the toxic effects of them inside cells (Subudhi & Baisakh, 2011). Therefore, *Spartina alterniflora* exhibits a delicate ion exclusion system from physiology aspects to survive hash saline environment and it is a good model plant to study high salt tolerance. However, the molecular basis of salt tolerance in *Spartina alterniflora* remains unexplored, and the post-transcriptional regulation of high salt tolerance has not been characterized in plants.

Ion transporters play important roles in response to salt stress (J. K. Zhu, 2002, 2016). SOS1 (Salt Overlay Sensitive 1) is the most well-known Na^+^/H^+^ antiporter that is used to extrude Na ^+^ from root cells into the soil and load Na^+^ into the xylem for long-distance transportation from roots to leaves via transpiration systems (Yang & Guo, 2018; Y. Zhang et al., 2018; J. K. Zhu, 2002, 2016). Briefly, excessive salt stress triggers cytoplasmic Ca^2+^ signaling, which is then sensed by SOS3 (J. K. Zhu, 2016). SOS3 then interacts and activates the CBL-interacting protein kinase SOS2 (J. K. Zhu, 2016). The activated SOS2 then phosphorylates and activates the Na^+^/H^+^ antiporter SOS1 (J. K. Zhu, 2016). HKT1 has an antagonistic function with SOS1 on long-distance Na^+^ transport and prevents Na^+^ transport into leaves (J. K. Zhu, 2016). Lower Na^+^/K^+^ ratio is very important in plant salt tolerance (G. Chen et al., 2015; Y. Zhang et al., 2018; X. Zhu et al., 2018) and ion transporters from Shaker family are responsible for potassium uptake and transport (Barragan et al., 2012; Lacombe et al., 2000). AKT1 and AKT2 are two important potassium transporters from the Shaker family that function for both potassium influx and efflux (Y. Zhang et al., 2018). KAT2 and KAT1 interacted with each other to form inward K^+^ channels (Nieves-Cordones et al., 2014). SKOR plays a role in delivering K^+^ from the root stelar cells toward shoots (Gaymard et al., 1998). The K^+^/H^+^ antiporter NHX1 and NHX2 mediate the K^+^ uptake from cytosol to vacuolar and regulate reservoir dynamics of K^+^ (Barragan et al., 2012). However, none of these ion transporters has been characterized in *Spartina* and their regulatory roles in high salt tolerance are largely unexplored.

Protein phosphorylation is one important post-translational regulation mechanism and is regulated by protein kinases (Y. Ding et al., 2015). SOS2 represents one of the Ca^2+^ dependent protein kinases, including 10 SnRK2 and 25 SnRK3 in Arabidopsis (J. K. Zhu, 2016). In the SnRK2 family, SnRK2.10 is rapidly activated in Arabidopsis roots under salt stress (McLoughlin et al., 2012) and OST1 regulates cold stress by phosphorylating C-repeat binding factors such as ICE1 under cold stress (Y. Ding et al., 2018; Fujii, Verslues, & Zhu, 2011). In the SnRK3 family, the loss of Arabidopsis SnRK3.16/CIPK1 and SnRK3.14/CIPK6 causes osmotic or salt stress sensitive phenotype (D’Angelo et al., 2006; Tripathi, Parasuraman, Laxmi, & Chattopadhyay, 2009). In addition, the leucine-rich repeat receptor protein kinase (LRRs) was also reported in response to environment stresses. For example, MIK2 is correlated with mild salt tolerance by the natural variation study (Van der Does et al., 2017). The overexpression of PnLRR-RLK27 from Antarctic moss confers salt tolerance (J. Wang et al., 2017). On the contrary, Arabidopsis LRR-like kinase AtRPK1 and OsRPK1 are negatively correlated with salt tolerance (Shi et al., 2014). However, it remains elusive whether protein kinases involve in high salt stress associated regulatory network.

Transcriptome sequencing has been widely adopted to explore the gene expression dynamics in response to salt stress from different species (Bushman, Amundsen, Warnke, Robins, & Johnson, 2016; Liu et al., 2019; Mutwakil et al., 2017; Qiu et al., 2017). However, the transcriptome studies in high salt tolerance from halophyte are relative limited and most of them mainly focus on dicotyledonous halophytes (Lee et al., 2013; Subudhi & Baisakh, 2011; Y. Zhang et al., 2008). Moreover, although the genome-wide transcriptome reprogramming under salt stress has been reported (Liu et al., 2019), the reprogramming under different salinity stress conditions remains unexplored. In addition, alternative splicing (AS) is one important post-transcriptional regulation that plays critical roles in eukaryotic pre-mRNA processing and contributes to diversifying the transcriptome and proteome (Deng & Cao, 2017). AS is affected by cold and heat stress in plants, and more importantly, ∼10% of AS events are associated with salt stress in Arabidopsis (F. Ding et al., 2014; Srivastava, Lu, Zinta, Lang, & Zhu, 2018). Mutations of splicing factors such as SKIP, Sm-like protein 5, and SKB1 lead to the increase of AS events, especially intron retention (IR), under salt stress (Cui, Zhang, Ding, Ali, & Xiong, 2014; Feng et al., 2015; Z. Zhang et al., 2011), indicating the important function of AS in response to salt stress. However, how AS responds to salt stress in halophytes remains unclear. The main difficulty in analyzing transcriptome in *Spartina* is largely attributed to its complex genome with aneu-hexaploid (2n = 6x = 62) and the lack of high quality reference genome (Renny-Byfield et al., 2010). Fortunately, the recently developed single-molecule long-read sequencing technology (SMRT) by the Pacific Biosciences (PacBio) platform can sequence full-length mRNA isoforms without *de novo* assembly, which has been widely adopted to profile full-length transcripts in *Salvia miltiorrhiza*, *Sorghum bicolor*, *Phyllostachys edulis*, *Drynaria roosii*, maize, wheat and strawberry (Abdel-Ghany et al., 2016; An, Cao, Li, Humbeck, & Wang, 2018; Dong et al., 2015; Y. Li, Dai, Hu, Liu, & Kang, 2017; Sun et al., 2018; B. Wang et al., 2016; T. Wang et al., 2017; Xu et al., 2015). These pioneering studies provide elegant solutions for the study of biological mechanisms in poorly-annotated species.

In this study, we used PacBio SMRT sequencing and RNA-seq to get insight into the high salt tolerance mechanism of *Spartina alterniflora.* Full-length reference transcript sequences of *Spartina* were obtained. We uncovered the crucial role of ion transporters associated with the high salt tolerance and identified hub genes in the high salt tolerant regulatory network. The involvement of alternative splicing in high salt tolerance was also documented. Furthermore, candidate transcripts for salt tolerance of *Spartina* were functionally analyzed in Arabidopsis. We also built the SAPacBio website (http://plantpolya.org/SAPacBio/) for visualization and query of full-length transcriptome sequences in *Spartina*. Overall, our results provide new insights into the high salt tolerance regulatory mechanism of halophytes and demonstrate the great potential of *Spartina* to provide genetic resources for engineering salt-tolerant plants.

## MATERIALS AND METHODS

### Plant Materials

*Spartina alterniflora* seeds were collected in Ganyu County (E119°16’; N34°46’), Jiangsu Province, China. *Spartina* seedlings were grown on half-strength Hoagland liquid medium for 6 weeks in a growth chamber under a 16-hour light (80 µmol m^-2^s^-1^)/8-hour dark regime at 26°C (day) and 16°C (night). Plants with similar vigor were selected and salinized for 24 hours, in triplicate, at different NaCl concentrations (0, 350, 500, and 800 mM). These concentrations were selected to mimic non-salt-stressed conditions, low, medium, and high-salinity stress, since the lethal salt concentration for *Spartina* germination is ∼1000 mM NaCl (Subudhi & Baisakh, 2011), and the average salinity level of the sea is ∼599 mM (3.5%) NaCl. Total RNAs were isolated (Tiangen, Cat.DP441) and high-quality samples (RNA integrity number > 8) were used to prepare libraries for experimental validation.

### Library construction and sequencing

Pacbio SMRT and RNA-seq (Levin et al., 2010) were performed using whole seedlings grown at different salt concentrations (0, 350, 500, and 800 mM NaCl), as described previously (T. Wang et al., 2017). These libraries were sequenced on the PacBio RSII platform (Iso-seq) and the HiSeq 2500 sequencing platform (RNA-seq, paired-ends 125 bp).

### Bioinformatics analysis

*Spartina* full-length non-chimeric (FLNC) transcripts were identified from PacBio SMRT reads using the PacBio’s SMRT Analysis pipeline (v2.3.0) (Gordon et al., 2015), and were further corrected by proovread (Hackl, Hedrich, Schultz, & Forster, 2014) using Illumina RNA-seq data. These FLNC transcripts were clustered into unique transcript clusters by a two-rounds clustering incorporating CD-HIT-EST (v4.6) (Fu, Niu, Zhu, Wu, & Li, 2012) and GMAP (v2016-06-09) (T. D. Wu & Watanabe, 2005), and the longest isoform was defined as the unigene for each cluster.

The 5′ UTR, open reading frame (ORF), and 3′ UTR of each FLNC transcript were obtained using TransDecode (v3.0.1, https://transdecoder.github.io/). Functional annotations were generated using Blast2GO (blastx 2.2.26+) (Conesa et al., 2005) with the National Center for Biotechnology Information’s (NCBI) non-redundant database (e-value < 1e-5). Transcript datasets of *Oryza sativa* (IRSGP1.0) and *Arabidopsis thaliana* (TAIR10) were downloaded from EnsemblPlants (release34, http://plants.ensembl.org). All isoforms of *Spartina* were aligned to these datasets using blastn (e-value < 1e-5).

Gene ontology (GO) enrichment analysis (p-value < 0.05) was performed by BiNGO (Maere, Heymans, & Kuiper, 2005). Non-coding RNAs (ncRNAs) were predicted using CPAT (coding score < 0.298) (L. Wang et al., 2013) and the plant non-coding RNA database (PNRD) (e-value < 1e-5) (Yi, Zhang, Ling, Xu, & Su, 2015). Transcription factors were identified using the Plant Transcription Factor Database (PTFD, v4.0) with default parameters (Jin et al., 2017).

### Co-expression network analysis and alternative splicing

Differentially expressed (DE) unigenes were identified from RNA-seq data by RSEM (v1.2.25) combined with bowtie2 (v2.2.9) and EBSeq (v1.20.0) (Leng et al., 2013; B. Li & Dewey, 2011) based on the following criteria: false discovery rate (FDR)<0.05, and |fold change|≥2. All DE genes were used to construct the co-expression network by WGCNA (v1.66) (Langfelder & Horvath, 2008). The GO enrichment analysis of genes in each module was conducted by topGO (V2.32.0)(Alexa, Rahnenfuhrer, & Lengauer, 2006). Hub genes from the salt stress network were identified using Cytoscape plugin cytoHubba and ranked according to the maximal clique centrality (MCC) scores (Chin et al., 2014). AS events were identified using AStrap (Ji et al., 2018) and differentially expressed alternative splicing events (DEAS) were also characterized by using RNA-seq data (see Supplemental Methods).

### Experimental validation

RT-PCR and qRT-PCR experiments were conducted as described in a previous study (T. Wang et al., 2017). Primers used are listed in Table S1. Results from RT-PCR were visualized in 1% agarose gels. Coding sequences of *SaAF2* and *SaHSP70-I* (without stop codons) were cloned into modified pCambia3301 constructs (*pACT2::gene-GFP*), and transformed into the gene slicing suppression *rdr6-11* Arabidopsis mutants (Peragine, Yoshikawa, Wu, Albrecht, & Poethig, 2004). Two independent T_3_ transgenic lines were grown on plates containing half-strength Murashige and Skoog medium (1/2 MS) for 10 days under different salinity stress conditions (0, 120, and 150 mM NaCl). The primary root length was measured using ImageJ (Schneider, Rasband, & Eliceiri, 2012).

### Availability and accession numbers

Datasets from Illumina HiSeq 2500 and PacBio SMRT sequencing have been deposited at the NCBI website under the Bioproject accession number PRJNA413596. The SAPacBio website was built to query and visualize full-length isoforms, DE genes and AS in *Spartina* (http://plantpolya.org/SAPacBio/).

## RESULTS

### Characterization of the full-length transcriptome in *Spartina alterniflora*

We used PacBio SMRT sequencing to obtain full-length reference transcripts of high-confidence in *Spartina* to avoid potential bias attributed to its complex aneu-hexaploid genome (2n = 6x = 62) (Bedre, Mangu, Srivastava, Sanchez, & Baisakh, 2016). An overview of the experimental procedure is illustrated in Fig. 1. Briefly, 6-week-old whole *Spartina* plants were treated with 0, 350, 500, and 800 mM NaCl for 24 hours to simulate non-salt stressed conditions, low, medium, and high salt-stressed conditions. Total RNA was isolated from each salt stress condition and mixed equally to prepare three different size-selected libraries (1-2 kb, 2-3 kb, and > 3 kb). Seven cells were sequenced on the PacBio RSII real-time (RT) sequencing platform. The length distribution of reads of inserts (ROIs) in each library is consistent with the selected size (Fig. 2a and Table **S2**).

**Fig. 1.**
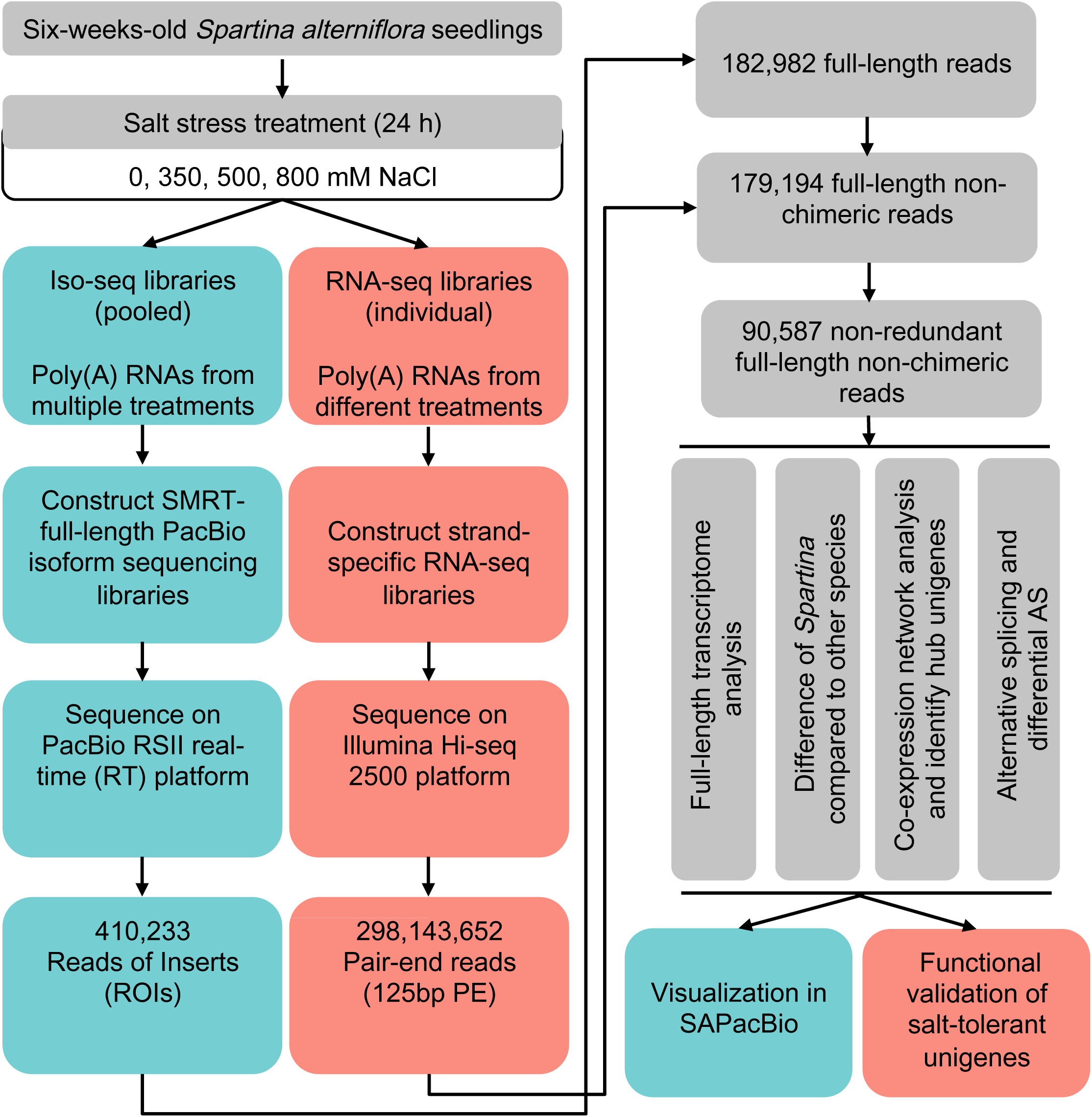
Overview of experimental and bioinformatics procedures in this study.

**Fig. 2.**
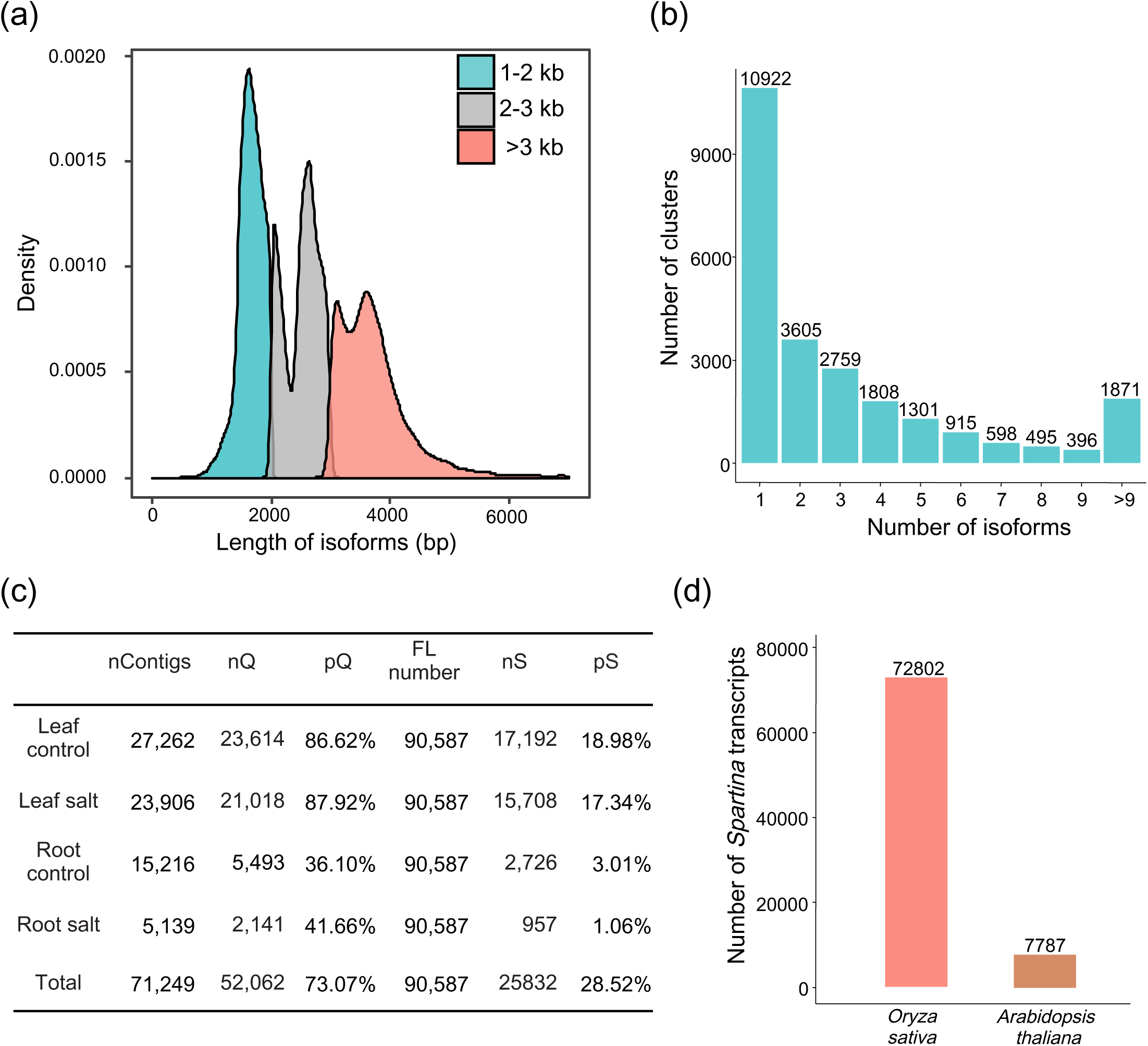
Characterization of the full-length transcriptome of *Spartina alterniflora*. (**a**) Distribution of reads of insert (ROI) lengths in each size-selected library. (**b**) Distribution of unique transcript clusters supported by different numbers of isoforms. (**c**) Comparison of transcripts identified from RNA-seq and PacBio data. Contigs from *de novo* assembly of RNA-seq data (Bedre *et al*., 2016) were mapped to PacBio full-length transcripts (FLs), and vice versa. nContigs: the number of *de novo*-assembled contigs from RNA-seq data; nQ: the number of contigs mapped to PacBio FLs; pQ: the percentage of contigs mapped to FLs; nS: the number of FLs mapped to contigs; pS: the percentage of FLs mapped to contigs. (**d**) Sequence similarity analysis of *Spartina* FLNC transcripts compared to rice and Arabidopsis using BLAST (e-value < 1e-5).

After the quality control by SMRT analysis (Gordon et al., 2015) and further correction with ∼250 millions Illumina RNA-seq reads by proovread (Hackl et al., 2014), a total of 179,194 FLNC reads (including a poly(A) tail, 5′ UTR and 3′ UTR) were obtained, 90,587 of which are non-redundant FLNC reads (Table **S2**). Over 90% of FLNC transcripts are ORF-containing transcripts with an average coding sequence (CDS) length of 1601 bp. These FLNC transcripts cover a wide variety of functional annotations such as catalytic activity, metabolic processes, and transporter activity (Fig. **S1**). These transcripts were further grouped into 24,670 unique transcript clusters (unigenes). Among them, 55.72% contained multiple isoforms (Fig. 2b), the percentage of which is comparable to that from previous studies (M. Wang et al., 2018; T. Wang et al., 2017). The number of isoforms in each cluster is correlated with their expression; single isoform-containing clusters tend to have lower expression levels than clusters with multiple isoforms (Fig. **S2**). Additionally, 3494 ncRNAs (Table **S3**) were identified. Furthermore, we used the plant transcription factor database (Jin *et al*., 2017) to identify transcription factors (TFs) in *Spartina*. In total, 4997 transcripts from 1323 unigenes were identified as encoding functional domains of TF families (Tables **S4-5**).

We further compared FLNC transcripts from PacBio with those obtained from RNA-seq data (Bedre et al., 2016). The average length of FLNC transcripts (2405 bp) is significantly longer than that from RNA-seq data (324 bp) (Bedre et al., 2016). Over 73% of RNA-seq contigs can be mapped to FLNC transcripts, while 64,755 FLNC transcripts are newly discovered and not present in RNA-seq data (Fig. 2c). Eight FLNC transcripts were randomly selected and all were further validated by RT-PCR (Fig. **S3**). Overall, our PacBio FLNC transcripts provide qualified full-length reference sequences for comprehensive profiling of the *Spartina* transcriptome.

### Genome-wide comparison of *Spartina* full-length transcriptome to other species

As *Spartina* is a monocot halophyte (Subudhi & Baisakh, 2011), we examined whether its transcripts are homologous to other plants. To this end, FLNC transcripts of *Spartina* were mapped to Arabidopsis and rice. Less than 9% of transcripts were mapped to Arabidopsis, while over 80% are highly similar to transcripts in rice (Fig. 2d). These results indicated that *Spartina* is highly homologous to the monocotyledonous-crop rice, and thus has the potential to provide genetic resources for engineering salt-tolerant rice. To understand the potential difference between *Spartina* and other plant species, we conducted a comparison of GO term percentages that was previously used to compare *Arabidopsis thaliana* and *Thellungiella salsuginea* (H. J. Wu et al., 2012). GO terms were mapped to 19,093 unigenes in *Spartina* using Blast2GO (Conesa et al., 2005), and were assigned to 30,821 annotated genes in *Arabidopsis thaliana* and 30,241 annotated genes in *Oryza sativa* using AgriGO (Tian et al., 2017). Detailed analysis revealed that the proportion of genes in the “membrane” (35.8% for *Spartina*, 27% for Arabidopsis, 9.5% for rice, with Fisher’s exact test p-value=1.94e-95 for the comparison between *Spartina* and Arabidopsis, and p-value=0 for *Spartina* and rice), “localization” (13.4% for *Spartina*, 8.8% for Arabidopsis, 7.3% for rice, with Fisher’s test p-value=4.70e-56 for *Spartina* and Arabidopsis, and p-value=6.18e-108 for *Spartina* and rice) and “transporter activity” (6.6% for *Spartina*, 4.6% for Arabidopsis, 4.8% for rice, with Fisher’s test p-value=2.06e-20 for *Spartina* and Arabidopsis, and p-value=7.19e-17 for *Spartina* and rice) categories were significantly increased in *Spartina* compared to both rice and Arabidopsis (Figs. 3c-d). As transporters are mainly membrane proteins and modulate cellular localization, the increasing percentage of transporter coding genes might contribute to the high salt tolerance phenotype in *Spartina*. Furthermore, 1577 *Spartina* transcripts differ from transcripts in other plant species, suggesting that these transcripts might be specifically expressed in *Spartina* and play unique roles in salt adaptation (**Table S6**).

**Fig. 3.**
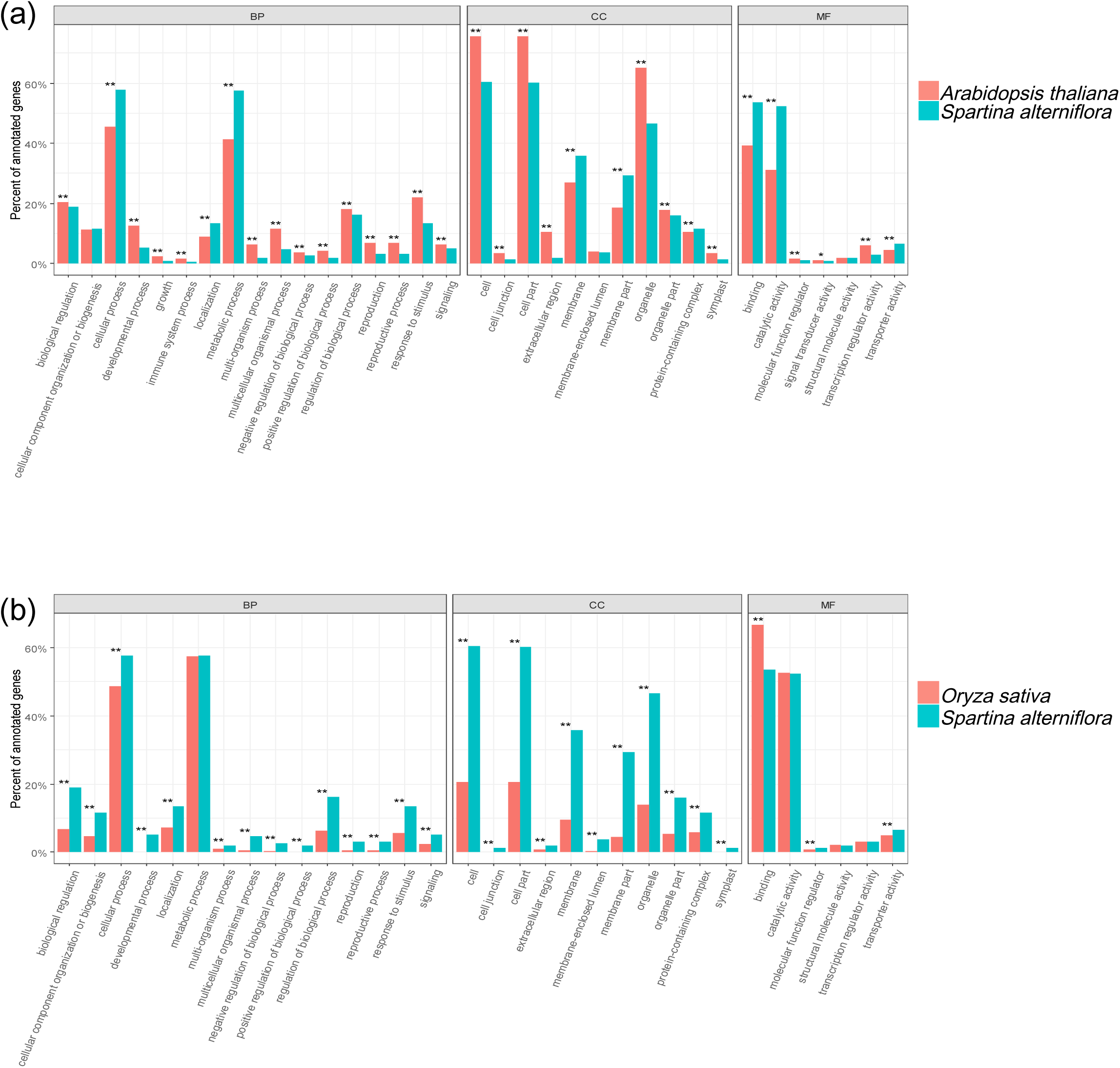
Comparison of gene ontology category percentage among *Spartina alterniflora*, rice and Arabidopsis. Blast2GO results of annotated genes or unigenes from these three species were mapped to categories in the second level of GO terms. The p-value was calculated using Fisher’s exact test and double stars represent p-value below 0.01, while a single star represents p-value between 0.01 and 0.05.

### Genome-wide dynamics of gene expression profile under salt stress in *Spartina*

To identify salt-responsive transcripts and explore salt tolerance mechanisms in *Spartina*, twelve RNA-seq libraries were constructed from plants grown under four salt stress gradients with three biological replicates. RNA-seq samples from the same conditions were clustered together, indicating the high reproducibility of our RNA-seq data (Fig. **S4**). A total of 3875 unigenes are differentially expressed (FDR < 0.05 and |log_2_FC| ≥ 1) among the salt-gradient experiments (Table **S7**). Moreover, 5108 FLNC transcripts, corresponding to 3375 unigenes, are differentially expressed between salinity stresses and control conditions. The number of DE transcripts increases with the salinity gradient (Fig. **S5a**). Moreover, the number of up-regulated transcripts is greater than that of down-regulated transcripts, particularly under high salt stress (Fig. **S5b-c**). The same trend was observed for DE unigenes (Figs. 4a, **S5d-f**), indicating that salt stress significantly affects the expression of *Spartina* unigenes.

**Fig. 4.**
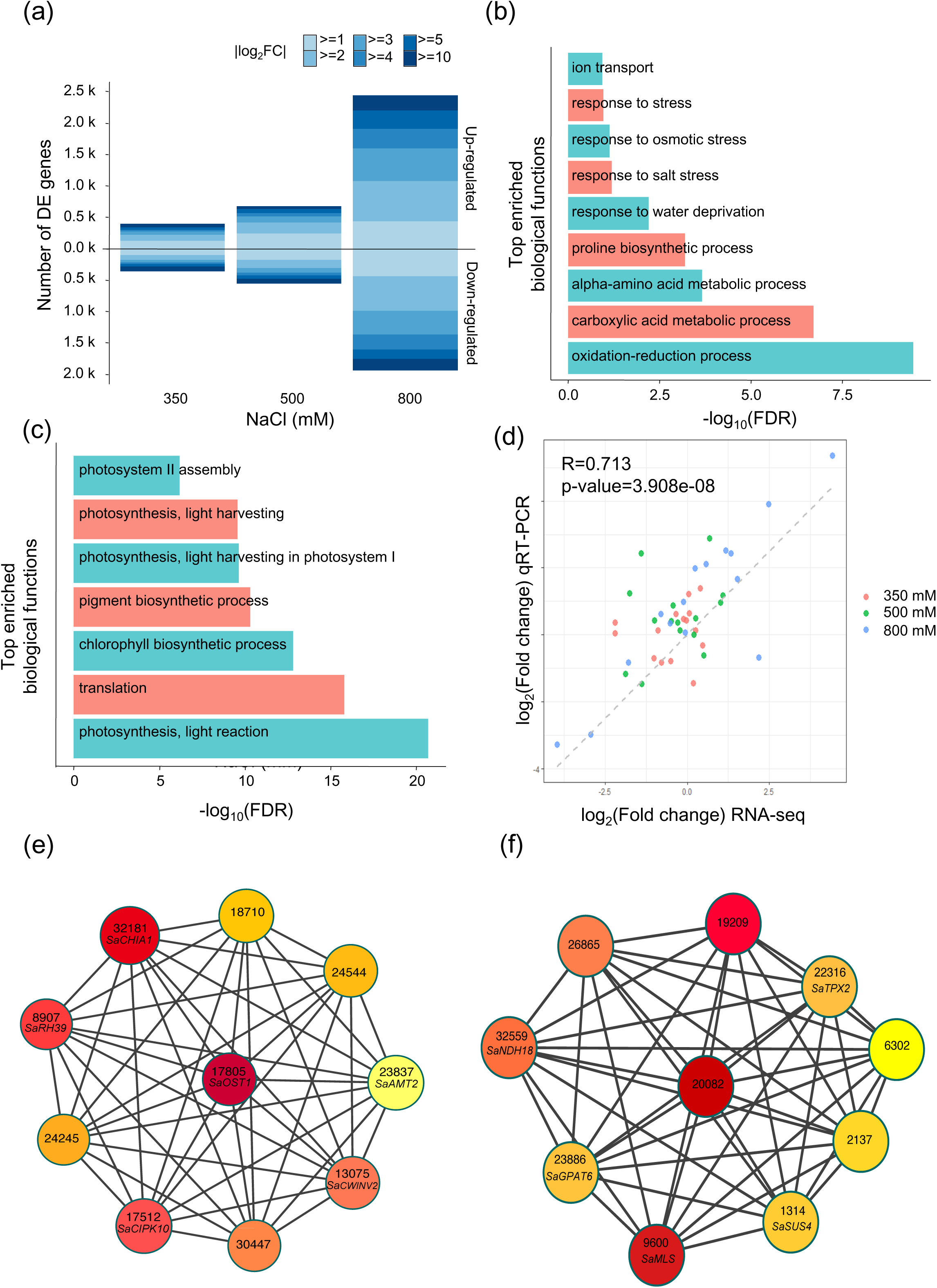
Identification of salt stress related transcripts and regulatory hub genes. (**a**) Number of differentially expressed (DE) genes in 24 hours under different NaCl stress conditions compared to control conditions. FC: fold change; up-regulated: FDR < 0.05 and log_2_FC ≥ 1; down-regulated: FDR < 0.05 and log_2_FC ≤ –1. (**b-c**) Gene ontology enrichment of high salt stress associated unigenes. (**b**) GO results for up-regulated unigenes; (**c**) GO results for down-regulated unigenes. (**d**) Plot of relative expression values of 15 DE genes from qRT-PCR and RNA-seq experiments. Each point denotes the log_2_(fold change) of expression levels between the respective treated condition and the control condition. The Pearson correlation is 0.713 and the p-value is 3.908e-08. (e) Top ten hub genes in the turquoise module of salt stress regulatory network. Hub genes were ranked according to the maximal clique centrality (MCC) scores with colors indicating MCC scores from high (red) to low (white). (f) Top ten hub genes in the blue module of the salt stress regulatory network.

Next, high salt stress associated unigenes (DE unigenes between 800 mM NaCl and control) were used to identify enriched pathways. Up-regulated unigenes are enriched in pathways including ion transport, response to stress, and amino acid metabolic processes (Fig. 4b), while photosynthesis and translation are significantly enriched in down-regulated unigenes (Fig. 4c). Importantly, these pathways are crucial components of salt stress-responsive mechanisms in plants (Deinlein et al., 2014; Munns & Tester, 2008; Soni et al., 2015), suggesting that high salt stress greatly affects the gene expression of salt-responsive pathways in *Spartina*.

To validate high salinity stress-associated unigenes, 15 DE unigenes were randomly selected for qRT-PCR experiments. The relative transcript levels measured by qRT-PCR and RNA-seq correlated significantly (Pearson correlation = 0.713, p-value = 3.908e-08, Fig. 4d). In addition, 10 DE unigenes were randomly selected for RT-PCR experiments and results confirmed the gene expression pattern from RNA-seq (Fig. **S6**).

### Co-expression network analysis reveals regulatory hub genes under high salt stress

To unveil the interrelationships among high-salt-stress responsive genes, we performed the weighted gene co-expression networks analysis (WGCNA) (Liu et al., 2019) with all 3875 DE unigenes from varying degrees of salt stress (Table **S7**). The co-expression network was built and DE unigenes were classified into five modules with hierarchical clustering. Different colors (blue, brown, green, turquoise, yellow) were used to represent distinct modules.

The largest module turquoise contains 2380 DE unigenes, showing an increasing expression trend especially under high salt stress (800 mM NaCl) (Table **S8**). We next performed gene ontology analysis to decipher potential functions of DE unigenes in the turquoise module. These genes are significantly enriched in “cellular homeostasis”, “chemical homeostasis” and “carbohydrate homeostasis” (Table **S8**). Specially, we observed that ion homeostasis processes such as metal ion homeostasis, cellular ion homeostasis, cellular metal ion homeostasis and transition metal ion homeostasis are enriched in DE genes of the turquoise module (Table **S8**). Consistently, transporters response to these homeostasis processes are also over-represented, which is further supported by the enrichment of “plasma membrane” under cellular components category and “transporter activity” under molecular function category (Table **S8**). These results suggested that transporters are involved in response to high salt stress.

The second largest module (blue) contained 1261 DE unigenes and showed a decreasing expression trends along with salt stress gradient, which was opposite with the turquoise module (Table **S8**). DE genes of the blue module are mainly enriched in “translation” and “photosynthesis” (Table **S8**). The module brown (97 DE genes) was induced under salt stress of 350 mM and 500 mM NaCl and is enriched in “small molecule metabolic process”, “methylation” and “amino acid metabolic process” (Table **S8**).

To uncover potential key regulators under high salt stress, we identified top-ten hub genes from each module. Several genes related to environmental stress regulation were identified (Fig. 4e-f). In the turquoise up-regulation module (Fig. 4e), *OST1* (*Cluster17805*, also known as *SnRK2.6* or *SnRK2E*) encodes an ABA activated protein kinase, playing vital roles in the ionic osmotic stress by regulating the stomatal closure (Boudsocq, Barbier-Brygoo, & Lauriere, 2004; Fujii et al., 2011). *CHIA* (*Cluster32181*) encodes a chitinase, which was reported not expressed at normal conditions but exclusively expressed under environmental stresses especially salt and wound stress in Arabidopsis (Takenaka, Nakano, Tamoi, Sakuda, & Fukamizo, 2009). *CIPK10* (*Cluster17512*, also known as *PKS2, SIP1,* and *SnRK3.8*) encodes a CBL-like protein kinase which was expressed higher in roots and showed increased expression under 200 mM NaCl treatment (Guo, Halfter, Ishitani, & Zhu, 2001). In the blue down-regulation module (Fig. 4f), *SUS4* (*Cluster1314*) encodes sucrose synthase 4, which was involved in response to hypoxia stress (Bieniawska et al., 2007). Therefore, these newly discovered hub genes might play crucial roles in the high salt stress associated regulatory network of *Spartina*.

### Complex gene expression dynamics of transporters and associated protein kinases

Genome-wide comparison analysis and co-expression network analysis showed that ion transporters play important roles in high salt tolerance of *Spartina* (Fig. 3 and Table **S8**). To further explore the function of ion transporters, we investigated the gene expression dynamics of sodium and potassium transporters under high salt stress. Gene expression levels of each ion transporter for sodium and potassium transport and related protein kinases are listed in Table **S9**. In general, under high salt stress, most of the sodium and potassium transporters showed gene expression variation under high salt stress.

Genes involved in SOS pathways including Na^+^/H^+^ antiporter *SaSOS1* (*Cluster1424*) and the Na^+^-K^+^ homeostasis modulator *SaSOS2* (*Cluster19661*) were significantly induced under high salt stress. HKT1 restricts long-distance transport of Na^+^ from root to leave and therefore has antagonistic functions with SOS1 (J. K. Zhu, 2016). However, *SaHKT1* is not induced under high salt stress at the transcription level (Table **S9**). In *Spartina*, seven *SaAHAs* were identified and *SaAHA2 (Cluster4145)* showed a significant up-regulation under high salt stress (Table **S9**). These results indicated that SaAHA2 might be involved in the high salt tolerance of *Spartina*.

SOS2 represents a large number of similar protein kinases, which include 10 SnRK2 and 25 SnRK3 in Arabidopsis (J. K. Zhu, 2016). In *Spartina*, *SaSnRK2.10/SnRK2B (Cluster27491)* and *SaOST1/SnRK2.6/SRK2E (Cluster17805)* from SnRK2 family are significantly up-regulated under high salt stress (Table **S9**). However, *SaASK1/SnRK2.4/SRK2A (Cluster27812)* is not induced under high salt stress (Table **S9**). Ten SnRK3s showed differential expression in salt stress by the co-expression network analysis. Eight of them including *SaSnRK3.8/CIPK10, SaSnRK3.9/CIPK12 (Cluster4217), SaSnRK3.11/SOS2 (Cluster19661), SaSnRK3.14/CIPK6/SIP3 (Cluster23013), SaSnRK3.16/CIPK1 (Cluster23029), SaSnRK3.22/CIPK11/SIP4 (Cluster25001), SaSnRK3.24/CIPK5 (Cluster21691)* and *SaSnRK3.25/CIPK25 (Cluster23892)* are significantly up-regulated under high salt stress (Table **S9**). In addition, the expression of *SaSnRK3.12/CIPK9 (Cluster4692)* is increased under mild salt stress but decreased under high salt stress (Table **S9**). However, *SaSnRK3.3/CIPK4* (*Cluster28689*) is the only down-regulated *SnRK3s* under high salt stress (Table **S9**).

Ion transporters from Shaker family are responsible for potassium uptake and transport (Barragan et al., 2012; Lacombe et al., 2000). In *Spartina*, *SaAKT1* did not show any significant expression change under high salt stress, while *SaAKT2* showed a significant down-regulation (Table **S9**). The inward K^+^ channel *SaKAT2* (*Cluster24848*) was significantly up-regulated under high salt stress, while *SaKAT1* (*Cluster13882*) did not have significant change (Table **S9**). The K^+^ transport related *SaSKOR* (*Cluster12966*) gene was also up-regulated under high salt stress (Table **S9**). We also found that the expression of tonoplast localized K^+^/H^+^ antiporter *SaNHX2 (Cluster17097)* was dramatically decreased at lower salt stress condition, while significant up-regulation was observed under high salt stress (Table **S9**).

### High salt stress induces global alternative splicing in *Spartina*

To investigate whether high salt stress affects AS in *Spartina*, 14,058 non-redundant AS events corresponding to 20,431 transcripts were identified using our bioinformatics pipeline. Approximately 39% (5328/13,748) of unigenes with multiple isoforms are associated with AS events. Nearly half (2857/5328) of AS-containing unigenes have more than two associated AS events (Fig. 5a). The distribution of different types of AS in *Spartina* is comparable to that of other species (Reddy, Marquez, Kalyna, & Barta, 2013; Shen et al., 2014). Intron retention (IR) is the major AS type (56%). Most IR-AS events (93%) contain the canonical GT-AG splice site; the percentage of AS using alternative acceptor sites (AltA) is two times of that of alternative donor sites (AltD) (Fig. 5b).

**Fig. 5.**
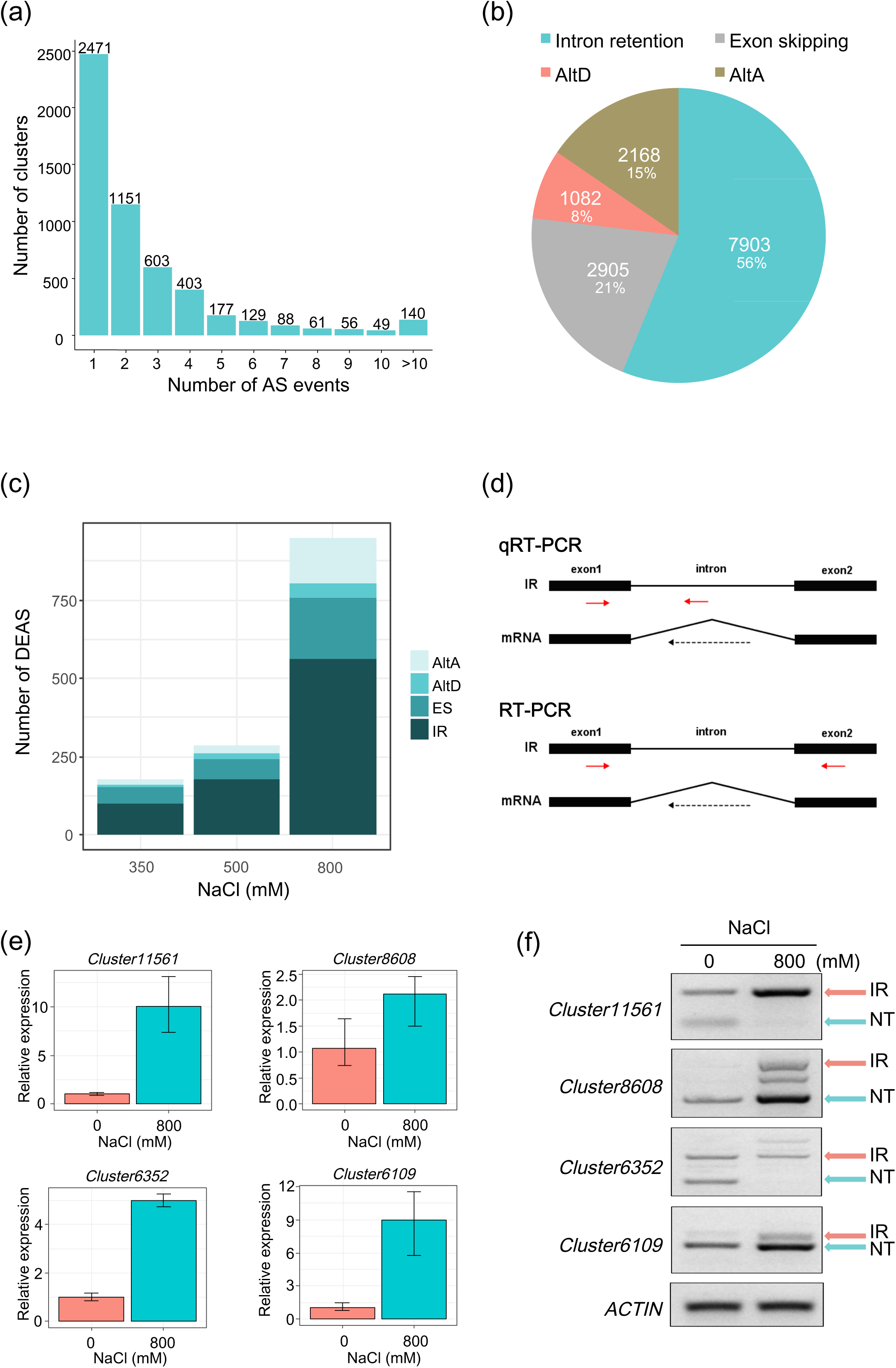
Alternative splicing (AS) and high salt stress-associated AS in *Spartina*. (**a**) Distribution of unique transcript clusters supported by different numbers of AS events. Proportion of different types of AS in *Spartina*. (**c**) Number of differentially expressed AS (DEAS) events associated with salinity stress. (**d**) Schematic showing the strategy used to validate intron-retention (IR) AS events. Primers were designed to specifically detect the expression of IR-containing transcripts by qRT-PCR under high salt stress, and a pair of primers flanking the IR region was used to validate the AS events by RT-PCR. (**e**) Relative expression of IR-containing transcripts measured by qRT-PCR. (**f**) Expression of IR transcripts and normal transcripts (NT) by RT-PCR.

Importantly, 1154 AS events corresponding to 1627 transcripts are associated with salinity stress in *Spartina*, accounting for ∼11% (578/5328) of AS-containing unigenes. The number of AS events is increased with growing salinity stress (Fig. 5c), suggesting that high salt stress induces AS events in *Spartina*. In particular, IR remains the major AS type associated with salt stress (Fig. 5c), and 233 IR events were identified in response to high salt stress (Table **S10**). According to the GO analysis, these AS events mainly affect photosynthesis-related unigenes (Fig. **S7**). Six IR events were selected for validation (Fig. 5d). Levels of transcripts associated with these IR events were indeed increased under high salt stress when examined by both RT-PCR and qRT-PCR (Fig. 5e, f). Overall, these results suggest that AS responds to high salt stress in *Spartina*.

### Functional validation of *Spartina* salt-tolerant unigenes in Arabidopsis

To test whether high salt stress responsive transcripts could be candidate gene resources for engineering salt-tolerant plants, we constitutively overexpressed heat shock protein coding gene-*SaHSP70-I (Cluster1653-042)* and NAC transcription factor-*SaAF2 (Cluster30510-002)* in Arabidopsis. Two independent T_3_ generation transgenic lines of overexpressed *SaHSP70-I* were grown at 120 and 150 mM NaCl. They showed better salt tolerance and had longer primary roots than control plants under salt stress (Fig. 5a-b). Similarly, the salt tolerance analysis in Arabidopsis showed that plants with overexpressed *SaAF2* also exhibited increased salt tolerance (Fig. 5c-d), indicating that *AF2* is a salt-tolerant candidate gene. The identification of many salt-responsive transcripts in *Spartina* and their functional validation in Arabidopsis further demonstrate the potential of using *Spartina* genes to engineer salt-tolerant plants.

## DISCUSSION

### Overview of the *Spartina alterniflora* full-length transcriptome

Salt stress is one of the most important environmental stresses in crop growth and production. High salt stress severely delays crop growth and significantly reduces its yield. *Spartina alterniflora* is a highly salt-tolerant monocotyledonous halophyte that belongs to the *Poaceae* family. In this study, we performed comprehensive transcriptome analysis in *Spartina* exposed to different salt stresses using the hybrids of SMRT full-length transcriptome sequencing and RNA-seq. A total of 90,587 non-redundant full-length transcripts, 24,670 unigenes, 3494 ncRNAs, 1323 TFs and 1577 specific transcripts in *Spartina* were identified (Tables. **S3-6**). More importantly, the sequence similarity analysis showed that approximately 80% of the full-length transcripts in *Spartina* could be aligned to rice transcripts (Fig. 2d), indicating that *Spartina* is evolutionarily similar to rice. As *Spartina* is very similar to rice from evolutionary aspects (Fig. 2d), we also attempted to apply the *Spartina* unigenes for engineering salt-tolerance plants. We over-expressed two *Spartina* salt stress-responsive genes *SaHSP70-I* and *SaAF2* in Arabidopsis and observed that their transgenic lines confer salt-tolerance phenotypes (Fig. 6). Previous studies showed that the over-expression of *Spartina* vacuolar ATPase subunit c1 (*SaVHAc1*) and actin-depolymerizing factor (*SaADF2*) genes in rice promoted the salt tolerance of transgenic rice (Baisakh et al., 2012; Sengupta et al., 2019). Therefore, these results indicate that *Spartina* could be an ideal species for engineering salt tolerance in crops. Overall, the full-length transcriptome of *Spartina* potentially provides valuable resources for improving the salt tolerance of plants.

**Fig. 6.**
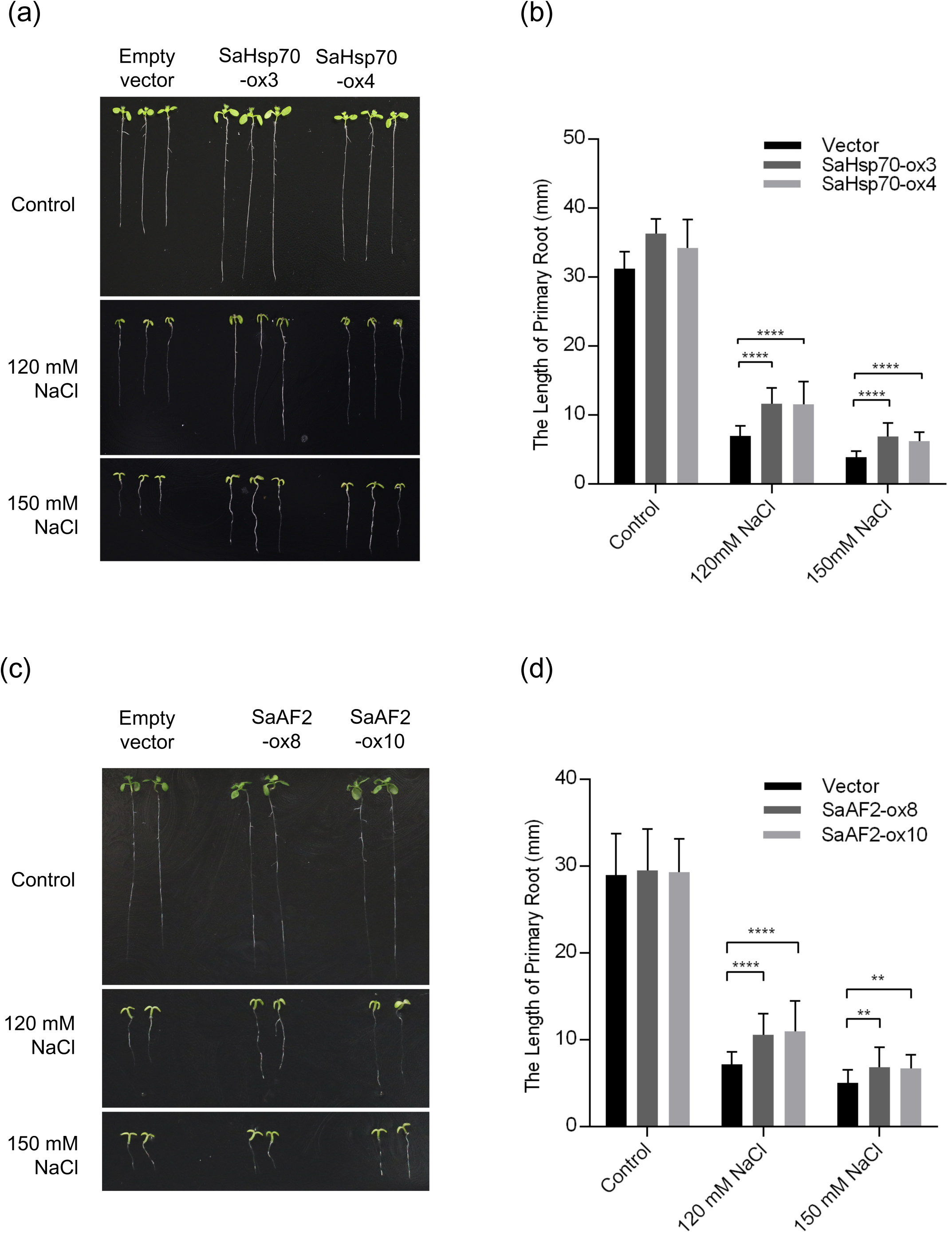
Functional validation of salt-responsive transcripts. (**a**) The salt tolerance phenotype of *SaHSP70-I* transgenic lines in Arabidopsis: The T_3_ generation seedlings of *SaHSP70-I* overexpression lines were grown on half-strength Murashige and Skoog medium after 10 days of salinity stress treatment. (b) Overexpression of *SaHSP70-I* confers better salt tolerance in Arabidopsis transgenic line. The primary root lengths (n > 20) were measured with ImageJ. (**c**) The salt tolerance phenotype of the *SaAF2* transgenic lines in Arabidopsis. (**d**) Overexpression of *SaAF2* improved salt tolerance of Arabidopsis transgenic line (n > 20).

### Gene expression dynamics of ion transporters under high salt stress

Genome-wide analysis showed that *Spartina* has a higher proportion of transporter related genes compared to rice and Arabidopsis (Fig. 3) and the co-expression network analysis also found that ion transporters are significantly enriched in the high salt stress associated module (Table **S8**). These results indicate that ion transporters may play a key role in the high salt tolerance of *Spartina*.

The most well known ion transporter under salt stress is SOS1 (Salt Overlay Sensitive 1), a Na^+^/H^+^ antiporter that is used to extrude Na ^+^ from root cells into the soil and load Na^+^ into the xylem for long-distance transportation from roots to leaves via transpiration systems (Yang & Guo, 2018; Y. Zhang et al., 2018; J. K. Zhu, 2002, 2016). Another important ion transporter is HKT1, which plays important roles in long-distance Na^+^ transport and has an antagonistic functions with SOS1 to restrict long-distance transport of Na^+^ from root to leave (J. K. Zhu, 2016). It is proposed that plants can activate *SOS1* under mild salt stress to transfer Na^+^ from root to leave, while HKT1 is activated under high salt stress to limit the transport of Na^+^ from roots because Na^+^ accumulation in leaves exceeds its storage capacity (J. K. Zhu, 2016). However, in *Spartina*, we showed that the expression level of *SaSOS1* is significantly higher under high salt stress, while no significant gene expression change was observed for *SaHKT1* (Table **S9**). These results suggest that the long-distance transport of Na^+^ from root to leave may continue to be active under high salt stress in *Spartina*. One of the explanations is that leaves of *Spartina* have salt glands (Nestler, 1977; Subudhi & Baisakh, 2011) and can secrete salt, so that high expression of *SaHKT1* may not be required to shut down Na^+^ transport compared to other plants (Jaime-Perez et al., 2017; M. Zhang et al., 2018).

It has been reported that the Na^+^/K^+^ ratio is very important in plant salt tolerance, and salt stress increases Na^+^ but leads to loss of K^+^ (G. Chen et al., 2015; Y. Zhang et al., 2018; X. Zhu et al., 2018). In general, *Spartina* maintained a higher K+ accumulation under high salt stress via the active gene expression dynamics of *SaAKT2*, *SaKAT2*, *SaSKOR* and *SaNHX2*. It is suggested that AKT1 is a potassium transporter under lower K^+^ conditions for uptake of K^+^, while AKT2 is one K^+^ efflux carrier under higher K^+^ conditions (Y. Zhang et al., 2018). In *Spartina*, the reduction in expression of *SaAKT2* but no change of *SaAKT1* (Table **S9**) may limit the K^+^ efflux under high salt stress and the results supported the physiologic observation that *Spartina* has a higher K^+^ accumulation level (Hester et al., 2001; Subudhi & Baisakh, 2011). Moreover, the down-regulation of *SaAKT2* may also reduce the Na^+^ uptake root cells initially through the K^+^ channel via *SaAKT2* (Salvador-Recatala, 2016). KAT2 and KAT1 have been reported to form inward K^+^ channels in the form of heterodimers and KAT2 can also form active homotrimeric channels in the plasma membrane (Nieves-Cordones et al., 2014). We found *SaKAT2* rather than *SaKAT1* is significantly up-regulated under high salt stress (Table **S9**), suggesting that *SaKAT2* may play a role in K^+^ uptake and maintain a higher K^+^ level through its homotrimeric channels in *Spartina* under high salt stress. SKOR is involved in delivering of K^+^ from stelar cells of roots to xylem toward the shoot (Gaymard et al., 1998). *SaSKOR* is also up-regulated under high salt stress (Table **S9**), indicating higher K^+^ transport from root to leaf under high salt stress. More importantly, the result is consistent with the observation that the lower Na^+^ / K^+^ ratio is present in *Spartina* leaves (Bradley & Morris, 1991; Smart & Barko, 1980). Higher Na^+^/K^+^ ratio is considered to be the major physiological reason for high salt tolerance of *Spartina* (Subudhi & Baisakh, 2011). The vacuolar is the largest dynamics reservoir of K^+^ in plant cells and the tonoplast localized K^+^/H^+^ antiporter, NHX1 and NHX2, are reported to mediate the K^+^ uptake from cytosol to vacuolar (Barragan et al., 2012). We found that *SaNHX2 (Cluster17097)* showed significant up-regulation under high salt stress (Table **S9**), indicating that *SaNHX2* may involve in the K^+^ reservoir in *Spartina* (Hester et al., 2001).

Overall, these results indicate that the reduction of Na^+^ exclusion, the activation of Na^+^ long-distant transport to secret them through salt gland and the higher K^+^ accumulation under high salt stress may be several important high salt tolerance mechanisms in *Spartina*.

### Protein kinase plays hub roles in the salt stress regulatory network

Protein phosphorylation plays important regulatory roles in plants in response to environmental stresses at the post-translational level (Y. Ding et al., 2015). The CBL-interacting protein kinase SOS2 involves in the SOS regulatory pathway to phosphorylate and activate the Na^+^/H^+^ antiporter SOS1 (J. K. Zhu, 2016). In *Spartina*, the expression level of *SaSOS2* (*Cluster19661*) is significantly increased under high salt stress (Table **S9**), indicating that *SaSOS2* is involved in high salt tolerance of *Spartina*.

Moreover, three SnRK2s and 16 SnRK3s were identified in *Spartina* (Table **S9**). Among them, two of *SnRK2s* and ten of *SnRK3s* showed significantly gene expression dynamics under high salt stress (Table **S9**). In particular, *SaOST1* from *SnRK2s* and *SaCIPK10* from *SnRK3s* belong to the top ten hub genes in the high salt stress associated up-regulation modules (Fig. 4e). The OST1 is triggered by ABA signaling under stress to phosphorylate the ABREs-binding proteins and activate the expression of stress responsive genes (Furihata et al., 2006). For example, the OST1 interacts and phosphorylates the ICE1 from C-repeat binding factors dependent pathways under cold stress to stable their proteins (Y. Ding et al., 2015). However, how OST1 involves in the high salt stress remains elusive. *SaOST1* is present in the top ten hub genes under high salt stress (Fig. 4e), suggesting that OST1 may also play crucial roles in the high salt tolerance of *Spartina* via post-translational regulation of salt stress related genes. CIPKs from many species including rice, maize and wheat have been reported to play important roles in salt tolerance and the overexpression of *CIPKs* significant improved salt tolerance of transgenic plants (X. Chen et al., 2014; Deng et al., 2013; Xiang, Huang, & Xiong, 2007). However, it is unexplored whether CIPKs play regulatory roles in salt tolerance. In this study, we showed that *SaCIPK10* is among the top ten hubs genes of high salt stress associated up-regulation modules in *Spartina* (Fig. 4e). These results suggest that SaCIPK10 may play regulatory roles in high salt tolerance. Interestingly, three of the top ten hub genes in the high salt stress associated down-regulation module are *leucine-rich repeat receptor kinases (LRRs)* (Fig. 4f), indicating that LRRs may also involve in the regulation of high salt tolerance.

In summary, over one fourth of the hub genes from high salt stress associated regulator network are protein kinases from SnRKs or LRRs families, indicating that kinases may play important regulatory roles in high salt tolerance of *Spartina*.

### The transcriptional and post-transcriptional regulation of photosynthesis related genes in *Spartina*

Photosynthesis is impaired under salt stress and it has been reported that *Spartina* saturates photosynthesis at relatively low CO_2_ level (Bedre et al., 2016). At the molecular level, we showed that photosynthesis genes, especially those associated with photosynthetic pigment content and the performance of photosystem II (Brzezowski, Richter, & Grimm, 2015; Chaves, Flexas, & Pinheiro, 2009), are also down-regulated under high salt stress (Fig. 4c), indicating that photosynthesis is regulated at transcriptional level.

AS is one post-transcriptional regulatory mechanism used by plants to respond to different environmental stresses (Deng & Cao, 2017; F. Ding et al., 2014; Feng et al., 2015). In this study, transcriptome-wide analysis indicated that high salt stress affects global AS (Fig. 5, Table **S10**). In particular, high salt stress induces a higher number of AS events along salinity stress gradients (Fig. 5c). In addition, the percentage of salt stress-associated AS genes (11%) is very close to that found in cotton and Arabidopsis (10%) (F. Ding et al., 2014; J. Zhang et al., 2014). Although salt stress increased IR events in several key salt-responsive genes in Arabidopsis (F. Ding et al., 2014; Feng et al., 2015), we did not observe this phenomenon in *Spartina* (Fig. **S7**). However, interestingly, photosynthesis-related genes are significantly enriched in the high salt stress associated AS, indicating extensive regulation of this process (Fig, **S7**). Overall, our results suggest that photosynthesis is highly regulated at both transcriptional and post-transcriptional level in *Spartina* under high salt stress.

In summary, our results provide a glimpse of the high salt tolerance mechanism of *Spartina alterniflora*. In addition, the full-length reference sequences, splice isoforms, transcription factors, ncRNAs and high salt stress associated hub genes in *Spartina* identified in this study will provide useful information for future research on halophytes and engineering salt tolerant plants.

## Acknowledgements

The authors thank Wenwen Liu from Xiamen University for his kind help in collecting *Spartina alterniflora* seeds. This work was supported by the National Natural Science Foundation of China (Nos. 31500258 and 31741025 to L.M., 61871463 and 61673323 to X.W., and 61573296 to G.J.), the Natural Science Foundation of Fujian Province of China (No. 2017J01068 to X.W.), the Outstanding Youth Research Talents Development Program in Fujian Province University to Liuyin Ma, the Outstanding Youth Research Talents Program of Fujian Agriculture and Forestry University (No. KXJQ17011 to L.M.), and the Scientific Research Foundation of the Graduate School of Fujian Agriculture and Forestry University (to T.W.).

## Author contributions

L.M. conceived the ideas. L.M., X.W., G.J., and C.L. designed experiments. T.W., W.W., and F.L. performed the experiments. X.W., W.Y., S.L., S.Z., T.W. and Q.L. contributed to data analysis and L.M. wrote the manuscript.

## SUPPORTING INFORMATION

### Supplemental Figures

**Fig. S1** Gene ontology analysis of unigenes in *Spartina alterniflora*.

**Fig. S2** Clusters with multiple isoforms showed higher expression levels than those with a single isoform.

**Fig. S3** RT-PCR validation of randomly selected transcripts in *Spartina alterniflora*.

**Fig. S4** Heatmap showing the Pearson correlation of unigene expression levels in different samples and repeated experiments.

**Fig. S5** Differentially expressed transcripts and unigenes in different conditions.

**Fig. S6** RT-PCR experimental validation of salt-responsive unigenes.

**Fig. S7** GO enrichment analysis using transcripts involved in differentially expressed alternative splicing events.

### Supplemental Tables

**Table S1.** List of primers used in this study.

**Table S2.** Summary of PacBio single-molecule long-read sequencing data.

**Table S3.** List of non-coding RNAs in *Spartina alterniflora*.

**Table S4.** List of transcription factor coding transcripts in *Spartina alterniflora*.

**Table S5.** Summary of transcription factor families in *Spartina alterniflora*.

**Table S6.** List of *Spartina*-specific transcripts.

**Table S7.** List of salt stress-responsive unigenes in *Spartina alterniflora*.

**Table S8.** Selective modules with diverse patterns in salt gradient experiment

**Table S9.** List of ion transporters and related protein kinases

**Table S10.** List of high salt stress associated intron retention events.

## Notes

http://plantpolya.org/SAPacBio/

