## Supplemental Figures for "The full-length transcriptome of *Spartina alterniflora* reveals the complexity of high salt tolerance in monocotyledonous halophyte"

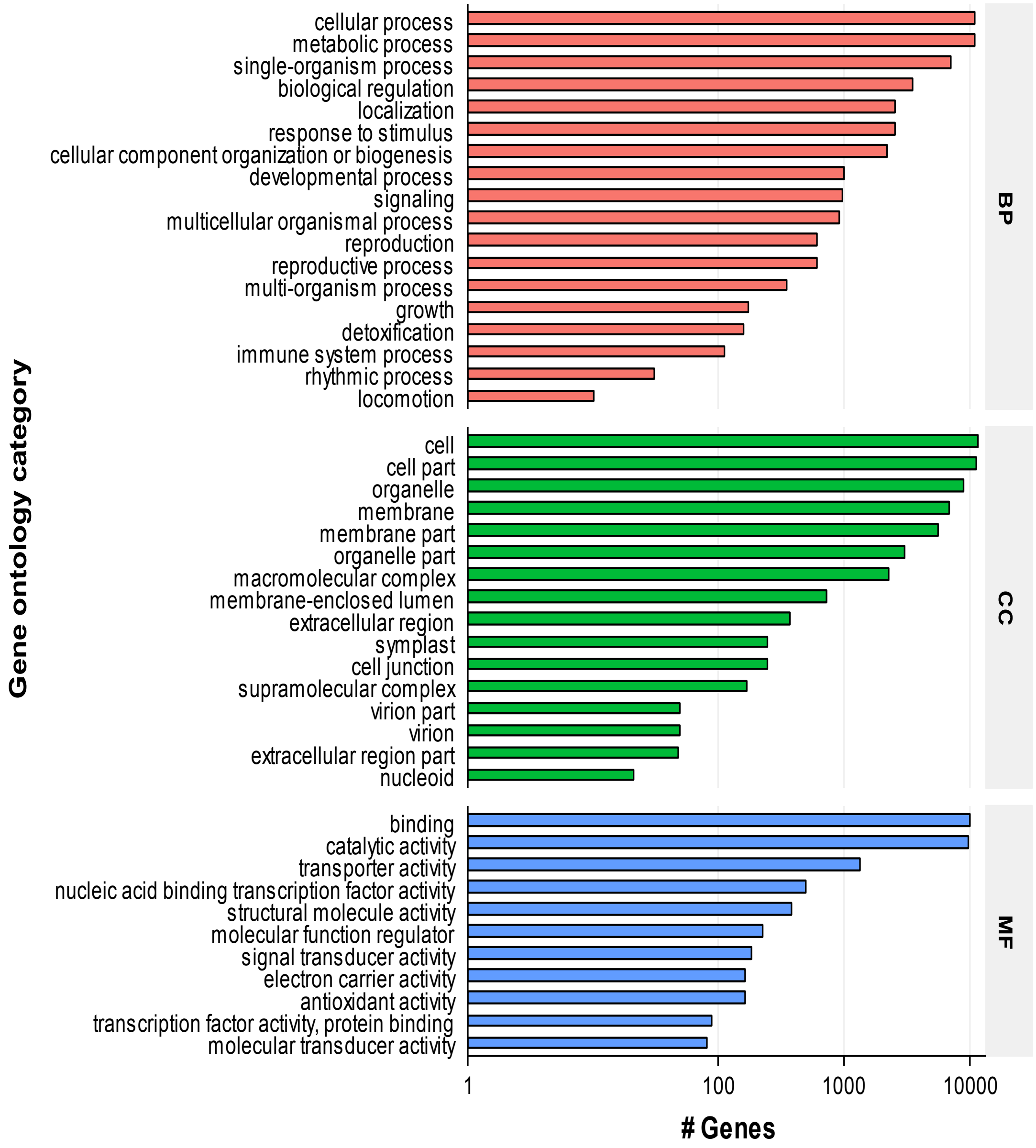


Fig. S1 Gene ontology analysis of unigenes in *Spartina*

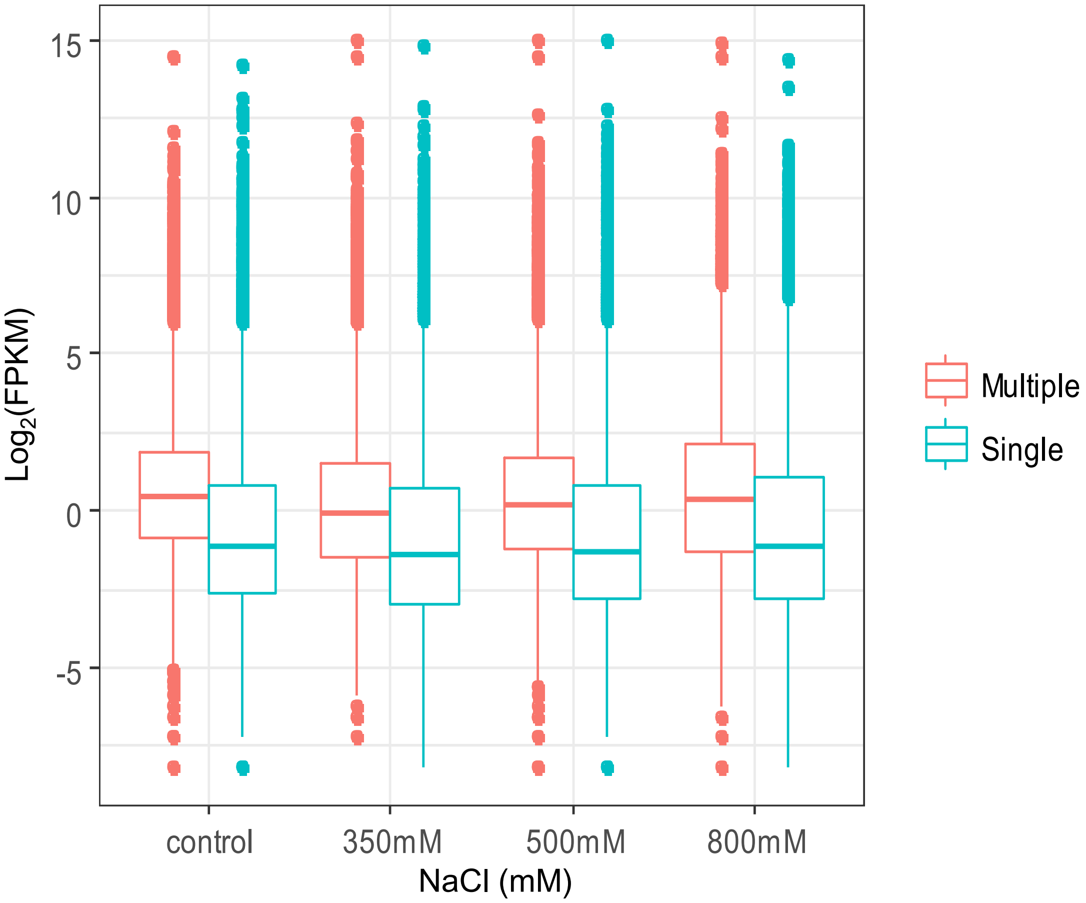


Fig. S2. Clusters with multiple isoforms showed higher expression levels than those with a single isoform. Expression level is denoted by FPKM (fragments per kilobase of exon per million fragments mapped).

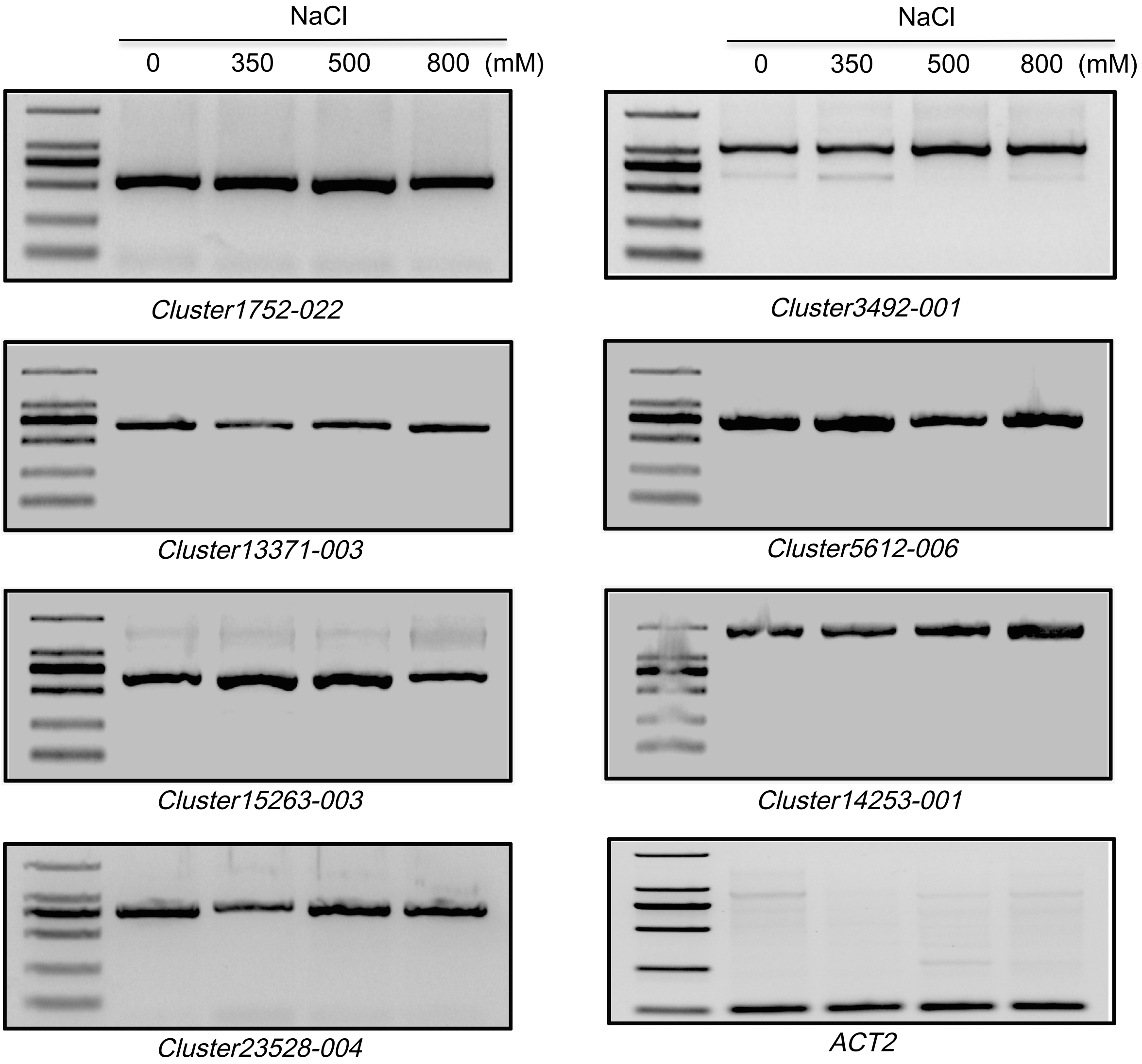


Fig. S3 RT-PCR validation of randomly selected transcripts in *Spartina.*

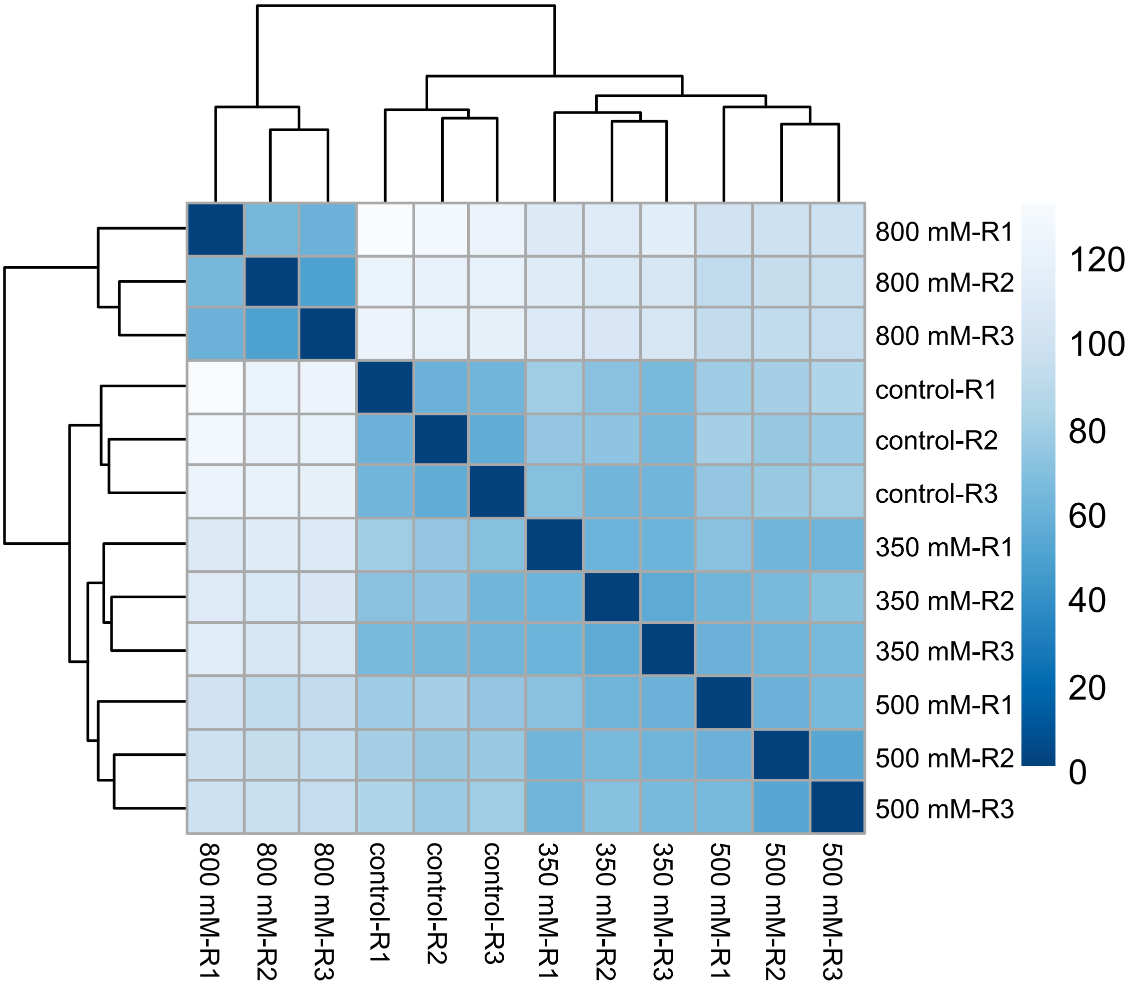


Fig. S4 Heatmap showing the Pearson correlation of unigene expression levels in different samples and repeated experiments.

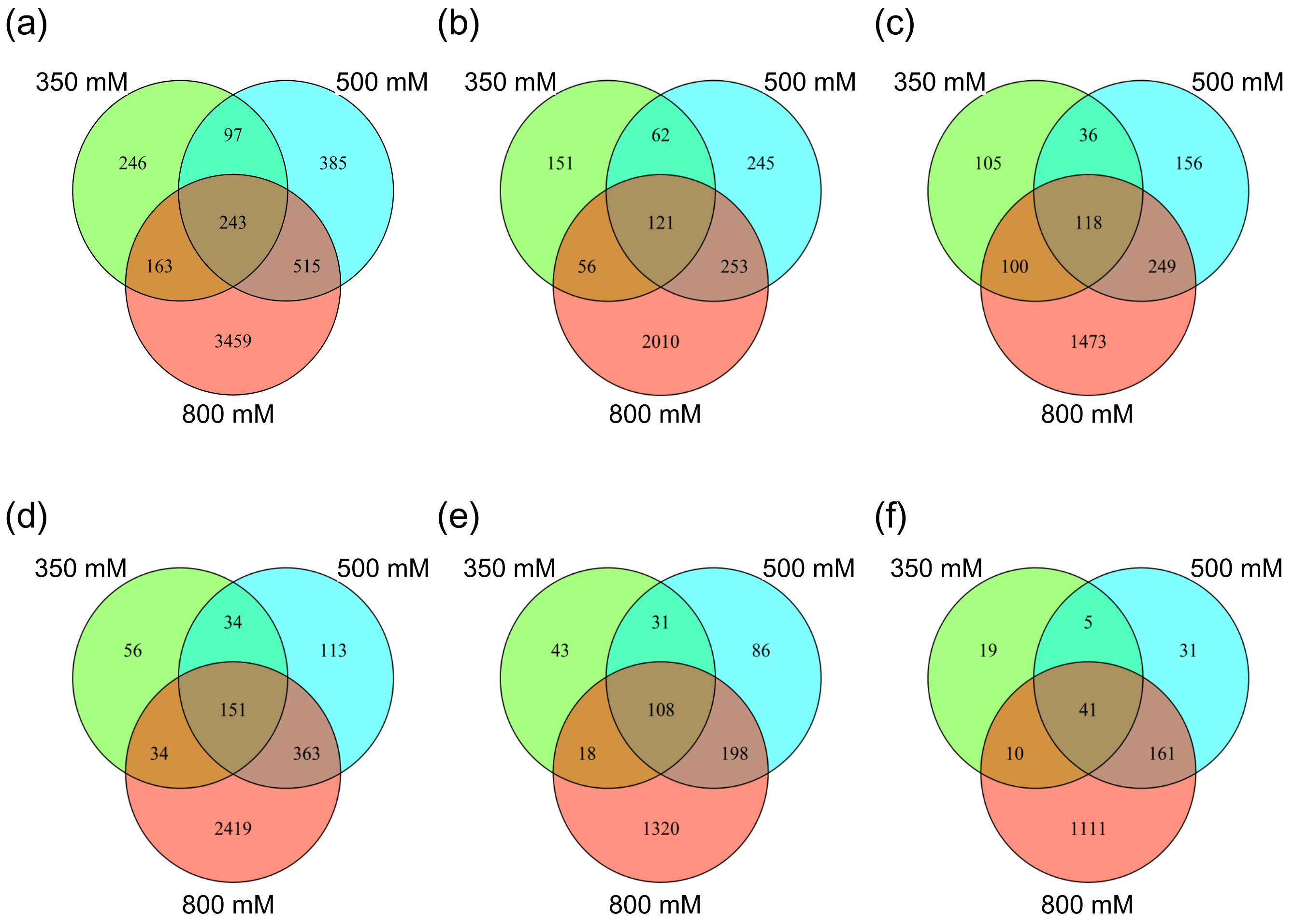


Fig. S5 Differentially expressed transcripts and unigenes in different conditions. Venn diagram showing the overlap between DE transcripts (a), up-regulated DE transcripts (b), down-regulated DE transcripts (c), DE genes (d), up-regulated DE genes (e), and down-regulated DE genes (f).

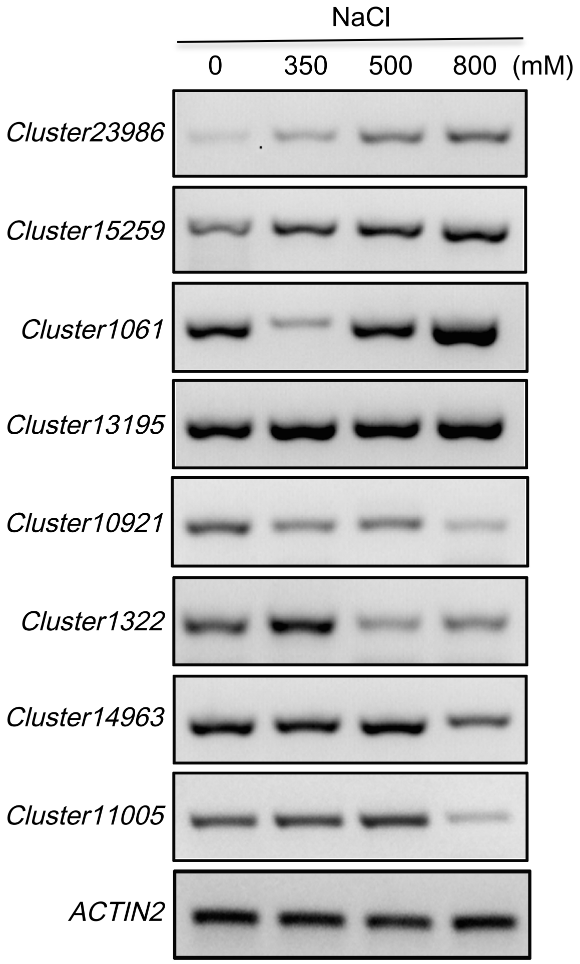


Fig. S6 RT-PCR experimental validation of salt-responsive unigenes

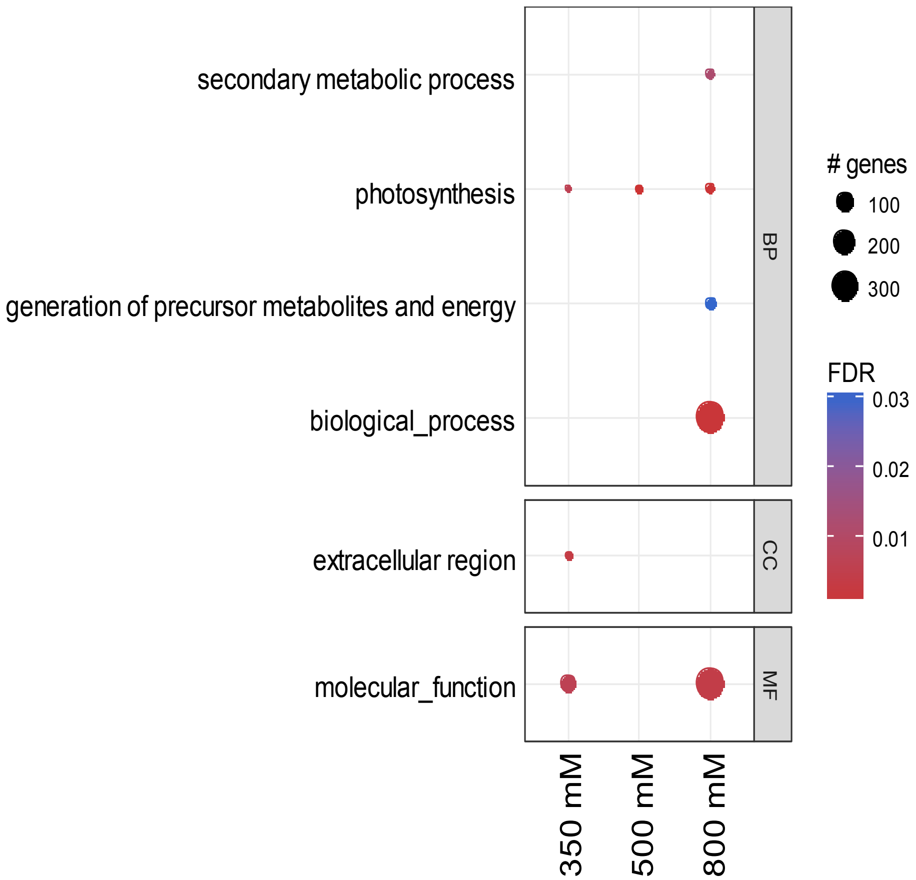


Fig. S7 GO enrichment analysis using transcripts involved in differentially expressed alternative splicing events. Shown are the enriched GO terms:biological process (BP), cellular component (CC) categories, and molecular function (MF).
