## Supplemental Table 1 for "The full-length transcriptome of *Spartina alterniflora* reveals the complexity of high salt tolerance in monocotyledonous halophyte"

### Table S1. List of primers used in this study

| Usage | Name | Primer sequence(5'-3') |
|---|---|---|
| **House keeping control** | Actin2 Cluster4283-F | 5'-AGGGCAGTTTTCCCTAGCAT-3' |
| | Actin2 Cluster4283-R | 5'-CTCTCTTGGACTGTGCCTCA-3' |
| **Transcripts validation RT-PCR** | Cluster1752-022-F | 5'-GCGCCTTCGATCTCTTTGTACT-3' |
| | Cluster1752-022-R | 5'-TTGGAGTACGAACTGCACGCTA-3' |
| | Cluster23528-004-F | 5'-CGTGCTCCTTCATTTTTCTTCCT-3' |
| | Cluster23528-004-R | 5'-ACAACATCAACAGTGGACTTCAG-3' |
| | Cluster15263-003-F | 5'-CACAAAGCTAATCCTGAGCATCAG-3' |
| | Cluster15263-003-R | 5'-TGAAGTGATTTAGCAGCCCATAGG-3' |
| | Cluster5612-006-F | 5'-CGTATTCGCACACGCATCCTT-3' |
| | Cluster5612-006-R | 5'-GCGGGCCCAAAATATCATCT-3' |
| | Cluster13371-003-F | 5'-CTTTGCCGTCTTTTCTTCTTTCC-3' |
| | Cluster13371-003-R | 5'-CGGGTAAACATTTACCATGAAGAC-3' |
| | Cluster14253-001-F | 5'-CCTGCATCTATAGTACCCATCTCA-3' |
| | Cluster14253-001-R | 5'-TCCATCCTCCAGATGGTGCATC-3' |
| | Cluster28222-001-F | 5'-ACGTACCCTGCATCTTCGGTGTA-3' |
| | Cluster28222-001-R | 5'-GAAGCTCCATCATTGTTGCCCT-3' |
| | Cluster3492-001-F | 5'-TTTTGACCCACGAGAGAACCG-3' |
| | Cluster3492-001-R | 5'-ACATGCTACGAGATCAAGTGAGC-3' |
| **RNA-seq DE qPCR** | Cluster27289-F | 5'-GAGTACTACGACGAGCAGCT-3' |
| | Cluster27289-R | 5'-CAGCTTGTTCTCCTCCTCGA-3' |
| | Cluster17941-F | 5'-GTGTTGAAGAGGCAGCGAAA-3' |
| | Cluster17941-R | 5'-AGTAGCCAGGAACCTTGTCC-3' |
| | Cluster1730-F | 5'-AGAGAGACCCCTAACGACCT-3' |
| | Cluster1730-R | 5'-AACTTTAAACGGCAGGGCAG-3' |
| | Cluster25106-F | 5'-AAGAACACGGCACAGCATTT-3' |
| | Cluster25106-R | 5'-TCCCTCATTCTTTCCACCCC-3' |
| | Cluster22263-F | 5'-CAAGTAGCCAGGAAAGTGCG-3' |
| | Cluster22263-R | 5'-CACCACAAAAGCCATCCCTC-3' |
| | Cluster18689-F | 5'-TCTCCCCACTTCACCATGAC-3' |
| | Cluster18689-R | 5'-CACACACTTTTGGCCCATGA-3' |
| | Cluster10346-F | 5'-AAGACCAGAGCCAGTTCCTC-3' |
| | Cluster10346-R | 5'-TAAGGAGGATGCAGCCAACA-3' |
| | Cluster1683-F | 5'-GGTCCTGCTTTGATCCTTGC-3' |
| | Cluster1683-R | 5'-TCTTGCCAGTCATCCAGCAT-3' |
| | Cluster3495-F | 5'-GAGACGGTACCAGGTGTTCA-3' |
| | Cluster3495-R | 5'-GACCTTCCTCTCTCTCTGCG-3' |
| | Cluster5236-F | 5'-TGGACATTGTTCCCAGGGAT-3' |
| | Cluster5236-R | 5'-CCACTTTCATGCAGGGATCG-3' |
| | Cluster7806-F | 5'-CATGTGTCAACTGTTCCGCA-3' |
| | Cluster7806-R | 5'-CCAAGTTGATCTCGCTGTGG-3' |
| | Cluster17652-F | 5'-ATCTCGTTGAAAGCCCTGGA-3' |
| | Cluster17652-R | 5'-ACAGCCCAACATGACTCTGA-3' |
| | Cluster19406-F | 5'-CCTATTCGATCGTTGCAGCC-3' |
| | Cluster19406-R | 5'-TTGCGGGACTCCAAGAAGAT-3' |
| | Cluster23489-F | 5'-AGGAACAGCTTCTCTACCGC-3' |
| | Cluster23489-R | 5'-CGAATAAGCTGGCGTGGC-3' |
| | Cluster23511-F | 5'-TAACTGTCTCCCTGTGGCTG-3' |
| | Cluster23511-R | 5'-CACCCCACAGCATCTTTTCC-3' |
| | Cluster18301-F | 5'-CTGCAATGGCTGTGCCTAA-3' |
| | Cluster18301-R | 5'-CTCACGCGCATATGATATGC-3' |

| | | |
|---|---|---|
| **DE RT-PCR** | Cluster13759-F | 5'-TACCGCTATCCTTTCTCTCTG-3' |
| | Cluster13759-R | 5'-TTTCGCGAATGCTCTCCAGG-3' |
| | Cluster23986-F | 5'-ACCGTCTTGTGGTCTCGGAT-3' |
| | Cluster23986-R | 5'-AGCTGCACGAGATCCACAGA-3' |
| | Cluster15259-F | 5'-TGCTGAACCTGGTCCCAGT-3' |
| | Cluster15259-R | 5'-ATCTTGGTGAGCGTGTGCT-3' |
| | Cluster1061-F | 5'-TGTACCTGGTGGACTTCGC-3' |
| | Cluster1061-R | 5'-CCGATGTCCTTGGGCTTGA-3' |
| | Cluster13195-F | 5'-ATAATTGGACCCAGGGTCGT-3' |
| | Cluster13195-R | 5'-TGGTGAAGGGTGGGAAGATC-3' |
| | Cluster10921-F | 5'-TTCTTGGCGTGGTTTGCTA-3' |
| | Cluster10921-R | 5'-ATGGTGTGCCTCGTTCAGT-3' |
| | Cluster1322-F | 5'-TGAACATGAAGGTGGAGCC-3' |
| | Cluster1322-R | 5'-GCACGGTCCTAGAACTCCAT-3' |
| | Cluster14963-F | 5'-TTGCAGCTATGGCTACGATC-3' |
| | Cluster14963-R | 5'-TAGTTGAGGATGTTGCCGTCC-3' |
| | Cluster11005-F | 5'-ACCCGGTACAGGATGAGGAT-3' |
| | Cluster11005-R | 5'-AAGCTTTGATGCTCCCCAAG-3' |
| **AS RT-PCR** | Cluster11561-F | 5'-CATGCTGCCTGAGACCAAT-3' |
| | Cluster11561-R | 5'-GCATCGTCATCGTCAGGTA-3' |
| | Cluster8608-F | 5'-GCAGGTATGGCAGTTCAGG-3' |
| | Cluster8608-R | 5'-TCAGTGACTGGAACAAGCG-3' |
| | Cluster6352-F | 5'-TCACCGACCGAATCGACAA-3' |
| | Cluster6352-R | 5'-CCAGCTCAAATACCAAGCGC-3' |
| | Cluster6109-F | 5'-CACACACGAAATCCTCTGTG-3' |
| | Cluster6109-R | 5'-TCCAGCATCAGCTCGTCGTT-3' |
| **AS qPCR** | Cluster11561-F | 5'-CATGCTGCCTGAGACCAAT-3' |
| | Cluster11561-R | 5'-TTGCTTGCGAACTGACCGG-3' |
| | Cluster8608-F | 5'-GCAGGTATGGCAGTTCAGG-3' |
| | Cluster8608-R | 5'-TCCGGAAAGCACGAACAAC-3' |
| | Cluster6352-F | 5'-TCACCGACCGAATCGACAA-3' |
| | Cluster6352-R | 5'-ACCAAGGGAAACCGTTCGG-3' |
| | Cluster6109-F | 5'-CACACACGAAATCCTCTGTG-3' |
| | Cluster6109-R | 5'-AGGAAAACGGGAGCCTCGT-3' |
| **Overexpressing genes cloning** | SaAF2-F | 5'-TGACCTCGAGACTAGTATGACGACAGGAGCGATGG-3' |
| | SaAF2-R | 5'-AGGTGGAGGTCCCCCGGGACCAGAGATAAGAGGGAACCAGA-3' |
| | SaHSP70-I-F | 5'-TGACCTCGAGACTAGTATGGTCAACCACTTTGTCCA-3' |
| | SaHSP70-I-R | 5'-AGGTGGAGGTCCCCCGGGGTCGACCTCCTCGATCTTGG-3' |
