## Supplemental Table 2 for "The full-length transcriptome of *Spartina alterniflora* reveals the complexity of high salt tolerance in monocotyledonous halophyte"

**Table S2. Summary of PacBio single-molecule long-read sequencing**

| | 1-2kb | 2-3kb | >3kb |
|---|---|---|---|
| No. of Reads of Inserts | 210,733 | 123,537 | 75,963 |
| No. of 5' reads | 137,472 | 72,105 | 36,278 |
| No. of 3' reads | 143,546 | 75,330 | 41,454 |
| No. of poly(A) reads | 136,459 | 71,914 | 39,543 |
| No. of filtered short reads | 10,043 | 4,402 | 1,545 |
| No. of non-full-length reads | 93,457 | 67,355 | 50,449 |
| No. of full-length reads | 107,233 | 51,780 | 23,969 |
| No. of full-length chimeric reads | 2,823 | 678 | 287 |
| No. of full-length non-chimeric reads | 104,410 | 51,102 | 23,682 |
| Average FLNC read length(bp) | 1,740 | 2,759 | 3,928 |
