## Supplemental Table 3 for "The full-length transcriptome of *Spartina alterniflora* reveals the complexity of high salt tolerance in monocotyledonous halophyte"

**Table S3. List of non-coding RNAs in *Spartina alterniflora.***

ncRNA: CPAT coding score<0.298 or PNRD e-value<1e-05

| Seqname | CPAT | PNRD | length(bp) |
|---|---|---|---|
| SALT00000085214_Cluster5-001 | 0.003 | NA | 16,659 |
| SALT00000079246_Cluster21-001 | 0.110 | NA | 15,016 |
| SALT00000086347_Cluster24-081 | 1.000 | NONATHT003785\|lincRNA493_PMID_24635777\| | 14,665 |
| SALT00000069783_Cluster25-001 | 1.000 | lincRNA372_PMID_24635777\| | 2,891 |
| SALT00000075022_Cluster25-002 | 1.000 | lincRNA372_PMID_24635777\| | 3,345 |
| SALT00000082777_Cluster25-004 | 1.000 | lincRNA372_PMID_24635777\| | 2,735 |
| SALT00000090954_Cluster25-005 | 1.000 | lincRNA372_PMID_24635777\| | 3,612 |
| SALT00000089194_Cluster25-006 | 1.000 | lincRNA372_PMID_24635777\| | 4,223 |
| SALT00000083573_Cluster25-007 | 1.000 | lincRNA372_PMID_24635777\| | 4,121 |
| SALT00000085320_Cluster25-008 | 1.000 | lincRNA372_PMID_24635777\| | 3,625 |
| SALT00000079046_Cluster25-009 | 1.000 | lincRNA372_PMID_24635777\| | 2,479 |
| SALT00000032820_Cluster25-010 | 1.000 | lincRNA372_PMID_24635777\| | 3,861 |
| SALT00000081268_Cluster25-011 | 0.999 | sit-MIR95-npr\|lincRNA372_PMID_24635777\| | 3,223 |
| SALT00000090416_Cluster25-012 | 1.000 | lincRNA372_PMID_24635777\| | 3,334 |
| SALT00000016805_Cluster37-001 | 1.000 | lincRNA304_PMID_24635777\| | 1,543 |
| SALT00000086025_Cluster38-001 | 0.032 | NA | 14,026 |
| SALT00000002367_Cluster43-001 | 0.349 | NONATHT003847\|sit-MIR121-1-npr\|sit-MIR122-1-r | 1,677 |
| SALT00000004540_Cluster43-002 | 0.382 | gma-MIR4995\|NONATHT003847\|lincRNA493_PMI | 1,476 |
| SALT00000004842_Cluster43-003 | 0.779 | ghr-MIR5368b\|NONATHT003839\|lincRNA12_PMII | 1,453 |
| SALT00000011573_Cluster43-004 | 0.318 | NONATHT003839\|lincRNA493_PMID_24635777\|gı | 1,868 |
| SALT00000013199_Cluster43-005 | 0.712 | ghr-MIR5368*\|sit-MIR121-1-npr\|lincRNA12_PMID_ | 1,957 |
| SALT00000016636_Cluster43-006 | 0.720 | TCONS_00018605\|sit-MIR122-1-npr\|lincRNA372_F | 1,899 |
| SALT00000019983_Cluster43-007 | 0.722 | TCONS_00018605\|lincRNA372_PMID_24635777\|si | 1,887 |
| SALT00000020523_Cluster43-008 | 0.705 | NONATHT003847\|lincRNA12_PMID_24635777\|sit- | 2,007 |
| SALT00000020568_Cluster43-009 | 0.610 | lincRNA372_PMID_24635777\|TCONS_00018605\|N | 3,218 |
| SALT00000023449_Cluster43-010 | 0.176 | TCONS_00018605\|lincRNA372_PMID_24635777\|si | 2,967 |
| SALT00000024050_Cluster43-011 | 0.121 | sit-MIR122-1-npr\|lincRNA372_PMID_24635777\|TC | 2,415 |
| SALT00000025079_Cluster43-012 | 0.608 | lincRNA372_PMID_24635777\|sit-MIR122-1-npr\|TC | 3,230 |
| SALT00000025160_Cluster43-013 | 0.656 | NONATHT003839\|TCONS_00018605\|ghr-MIR5368 | 1,318 |
| SALT00000025181_Cluster43-014 | 0.170 | lincRNA372_PMID_24635777\|ghr-MIR5368*\|NON | 1,824 |
| SALT00000025596_Cluster43-015 | 0.955 | sit-MIR121-1-npr\|lincRNA12_PMID_24635777\|NOI | 3,359 |
| SALT00000027220_Cluster43-016 | 0.881 | NONATHT003839\|lincRNA493_PMID_24635777\|N | 1,026 |
| SALT00000027275_Cluster43-017 | 0.594 | ghr-MIR5368b\|lincRNA493_PMID_24635777\|NON | 2,696 |
| SALT00000027372_Cluster43-018 | 0.661 | lincRNA372_PMID_24635777\|sit-MIR122-1-npr\|TC | 2,290 |
| SALT00000027475_Cluster43-019 | 0.635 | lincRNA372_PMID_24635777\|sit-MIR122-1-npr\|TC | 2,452 |
| SALT00000028498_Cluster43-020 | 0.608 | ghr-MIR5368*\|gma-MIR4995\|NONATHT003847\|lin | 3,232 |
| SALT00000030174_Cluster43-021 | 0.172 | sit-MIR122-1-npr\|lincRNA372_PMID_24635777\|TC | 3,109 |
| SALT00000030295_Cluster43-022 | 0.804 | NONATHT003847\|lincRNA12_PMID_24635777\|sit- | 3,224 |
| SALT00000031375_Cluster43-023 | 0.595 | lincRNA372_PMID_24635777\|sit-MIR122-1-npr\|TC | 3,305 |
| SALT00000031439_Cluster43-024 | 0.199 | TCONS_00018605\|sit-MIR122-1-npr\|lincRNA372_F | 2,758 |
| SALT00000031699_Cluster43-025 | 0.805 | ghr-MIR5368b\|TCONS_00018605\|NONATHT00383 | 1,235 |
| SALT00000031712_Cluster43-026 | 0.667 | NONATHT003847\|lincRNA12_PMID_24635777\|sit- | 2,253 |
| SALT00000031730_Cluster43-027 | 0.741 | ghr-MIR5368*\|lincRNA12_PMID_24635777\|NONA | 1,752 |
| SALT00000031757_Cluster43-028 | 0.382 | TCONS_00018605\|sit-MIR122-1-npr\|lincRNA372_F | 2,906 |
| SALT00000031780_Cluster43-029 | 0.773 | TCONS_00018605\|ghr-MIR5368b\|NONATHT00383 | 1,509 |
| SALT00000031781_Cluster43-030 | 0.398 | ghr-MIR5368b\|NONATHT003839\|lincRNA493_PMI | 2,816 |
| SALT00000031858_Cluster43-031 | 0.947 | ghr-MIR5368b\|NONATHT003839\|lincRNA493_PMI | 3,618 |
| SALT00000031957_Cluster43-032 | 0.787 | NONATHT003839\|TCONS_00018605\|ghr-MIR5368 | 1,394 |
| SALT00000032076_Cluster43-033 | 0.758 | NONATHT003847\|lincRNA12_PMID_24635777\|sit- | 1,621 |
| SALT00000033147_Cluster43-034 | 0.608 | ghr-MIR5368b\|NONATHT003839\|lincRNA493_PMI | 3,228 |
| SALT00000033608_Cluster43-036 | 0.608 | ghr-MIR5368b\|lincRNA493_PMID_24635777\|NON | 3,229 |

| | | | |
|---|---|---|---|
| SALT00000033737_Cluster43-037 | 0.502 | ghr-MIR5368*\|gma-MIR4995\|NONATHT003847\|lin | 3,219 |
| SALT00000033839_Cluster43-038 | 0.952 | sit-MIR122-1-npr\|lincRNA372_PMID_24635777\|TC | 3,466 |
| SALT00000033904_Cluster43-039 | 0.527 | NONATHT003839\|ghr-MIR5368b\|ghr-MIR5368*\|sit | 2,079 |
| SALT00000034627_Cluster43-040 | 0.582 | ghr-MIR5368b\|lincRNA493_PMID_24635777\|NON | 2,765 |
| SALT00000034992_Cluster43-041 | 0.635 | NONATHT003839\|ghr-MIR5368b\|ghr-MIR5368*\|lin | 2,455 |
| SALT00000035038_Cluster43-042 | 0.347 | ghr-MIR5368b\|NONATHT003839\|lincRNA493_PM | 3,119 |
| SALT00000035088_Cluster43-043 | 0.425 | lincRNA12_PMID_24635777\|NONATHT003847\|sit- | 2,655 |
| SALT00000035331_Cluster43-044 | 0.349 | TCONS_00018605\|sit-MIR122-1-npr\|lincRNA372_P | 3,108 |
| SALT00000038520_Cluster43-045 | 0.672 | lincRNA372_PMID_24635777\|ghr-MIR5368*\|NON | 1,426 |
| SALT00000041214_Cluster43-046 | 0.688 | TCONS_00018605\|lincRNA372_PMID_24635777\|si | 2,119 |
| SALT00000042958_Cluster43-047 | 0.824 | NONATHT003839\|ghr-MIR5368b\|TCONS_0001860 | 1,223 |
| SALT00000043139_Cluster43-048 | 0.555 | lincRNA372_PMID_24635777\|sit-MIR122-1-npr\|TC | 1,918 |
| SALT00000047377_Cluster43-049 | 0.755 | TCONS_00018605\|lincRNA372_PMID_24635777\|si | 1,646 |
| SALT00000049947_Cluster43-050 | 0.361 | NONATHT003847\|sit-MIR121-1-npr\|gma-MIR4995\| | 1,598 |
| SALT00000051421_Cluster43-051 | 0.347 | sit-MIR121-1-npr\|NONATHT003847\|gma-MIR4995\| | 1,685 |
| SALT00000054402_Cluster43-052 | 1.000 | ghr-MIR5368b\|TCONS_00018605\|NONATHT00383 | 1,995 |
| SALT00000054741_Cluster43-053 | 0.347 | sit-MIR121-1-npr\|NONATHT003847\|sit-MIR122-1-r | 1,686 |
| SALT00000055775_Cluster43-054 | 0.694 | NONATHT003847\|lincRNA12_PMID_24635777\|gh | 2,081 |
| SALT00000059422_Cluster43-055 | 0.718 | TCONS_00018605\|lincRNA372_PMID_24635777\|si | 1,918 |
| SALT00000061389_Cluster43-056 | 0.871 | NONATHT003839\|lincRNA493_PMID_24635777\|g | 1,661 |
| SALT00000062285_Cluster43-057 | 0.508 | TCONS_00018605\|NONATHT003785\|lincRNA372_ | 3,222 |
| SALT00000062959_Cluster43-058 | 0.609 | TCONS_00018605\|sit-MIR122-1-npr\|NONATHT00 | 3,225 |
| SALT00000063048_Cluster43-059 | 0.818 | NONATHT003839\|lincRNA493_PMID_24635777\|g | 1,690 |
| SALT00000063338_Cluster43-060 | 0.609 | TCONS_00018605\|lincRNA372_PMID_24635777\|si | 3,222 |
| SALT00000063564_Cluster43-061 | 0.968 | sit-MIR122-1-npr\|lincRNA372_PMID_24635777\|TC | 3,447 |
| SALT00000063879_Cluster43-062 | 0.608 | lincRNA372_PMID_24635777\|sit-MIR122-1-npr\|TC | 3,228 |
| SALT00000064260_Cluster43-063 | 0.851 | TCONS_00018605\|lincRNA372_PMID_24635777\|si | 3,231 |
| SALT00000064497_Cluster43-064 | 0.291 | sit-MIR122-1-npr\|TCONS_00018605\|gma-MIR4995\| | 3,217 |
| SALT00000064694_Cluster43-065 | 0.152 | TCONS_00018605\|lincRNA372_PMID_24635777\|si | 3,222 |
| SALT00000064733_Cluster43-066 | 0.609 | sit-MIR121-1-npr\|NONATHT003847\|lincRNA12_PI | 3,223 |
| SALT00000064874_Cluster43-067 | 0.612 | NONATHT003785\|lincRNA372_PMID_24635777\|T | 3,206 |
| SALT00000065802_Cluster43-069 | 0.587 | ghr-MIR5368b\|NONATHT003839\|lincRNA493_PM | 3,354 |
| SALT00000066224_Cluster43-070 | 0.593 | lincRNA372_PMID_24635777\|sit-MIR122-1-npr\|TC | 2,700 |
| SALT00000066991_Cluster43-071 | 0.581 | sit-MIR121-1-npr\|NONATHT003847\|lincRNA12_PI | 2,773 |
| SALT00000067101_Cluster43-072 | 0.609 | sit-MIR121-1-npr\|NONATHT003847\|lincRNA12_PI | 3,221 |
| SALT00000067253_Cluster43-073 | 0.581 | sit-MIR122-1-npr\|lincRNA372_PMID_24635777\|TC | 2,771 |
| SALT00000067358_Cluster43-074 | 0.478 | ghr-MIR5368b\|NONATHT003839\|lincRNA493_PM | 2,125 |
| SALT00000067400_Cluster43-075 | 0.481 | ghr-MIR5368*\|gma-MIR4995\|NONATHT003847\|lin | 3,339 |
| SALT00000067613_Cluster43-076 | 0.502 | sit-MIR122-1-npr\|TCONS_00018605\|ghr-MIR5368* | 3,225 |
| SALT00000067648_Cluster43-077 | 0.501 | sit-MIR121-1-npr\|NONATHT003847\|lincRNA12_PI | 3,227 |
| SALT00000068064_Cluster43-078 | 0.500 | TCONS_00018605\|lincRNA372_PMID_24635777\|si | 3,232 |
| SALT00000068733_Cluster43-079 | 0.526 | sit-MIR122-1-npr\|ghr-MIR5368*\|gma-MIR4995\|linc | 3,088 |
| SALT00000068904_Cluster43-080 | 0.609 | NONATHT003847\|lincRNA12_PMID_24635777\|sit- | 2,607 |
| SALT00000069066_Cluster43-081 | 0.606 | NONATHT003839\|lincRNA493_PMID_24635777\|g | 3,240 |
| SALT00000071142_Cluster43-082 | 0.590 | ghr-MIR5368*\|gma-MIR4995\|NONATHT003847\|lin | 3,330 |
| SALT00000071406_Cluster43-083 | 0.225 | ghr-MIR5368*\|gma-MIR4995\|lincRNA12_PMID_24 | 3,253 |
| SALT00000072065_Cluster43-084 | 0.480 | ghr-MIR5368b\|lincRNA493_PMID_24635777\|NON | 2,873 |
| SALT00000072786_Cluster43-085 | 0.609 | sit-MIR122-1-npr\|lincRNA372_PMID_24635777\|TC | 3,221 |
| SALT00000073063_Cluster43-086 | 0.858 | ghr-MIR5368b\|NONATHT003839\|lincRNA493_PM | 3,227 |
| SALT00000073437_Cluster43-087 | 0.608 | ghr-MIR5368b\|NONATHT003839\|lincRNA493_PM | 3,229 |
| SALT00000073707_Cluster43-088 | 0.618 | TCONS_00018605\|lincRNA372_PMID_24635777\|si | 3,170 |
| SALT00000073761_Cluster43-089 | 0.607 | sit-MIR122-1-npr\|lincRNA372_PMID_24635777\|TC | 3,237 |
| SALT00000073769_Cluster43-090 | 0.563 | lincRNA12_PMID_24635777\|NONATHT003847\|sit- | 2,878 |
| SALT00000074651_Cluster43-091 | 0.610 | sit-MIR122-1-npr\|lincRNA372_PMID_24635777\|TC | 3,218 |
| SALT00000075719_Cluster43-092 | 0.619 | sit-MIR122-1-npr\|lincRNA372_PMID_24635777\|TC | 2,547 |
| SALT00000075737_Cluster43-093 | 0.615 | sit-MIR122-1-npr\|lincRNA372_PMID_24635777\|TC | 3,030 |
| SALT00000075827_Cluster43-094 | 0.925 | NONATHT003839\|lincRNA493_PMID_24635777\|g | 3,451 |
| SALT00000076336_Cluster43-095 | 0.609 | gma-MIR4995\|ghr-MIR5368*\|sit-MIR121-1-npr\|NO | 3,224 |

| | | | |
|---|---|---|---|
| SALT00000077190_Cluster43-096 | 0.613 | ghr-MIR5368b\|lincRNA493_PMID_24635777\|NON. | 3,202 |
| SALT00000077938_Cluster43-098 | 0.978 | sit-MIR122-1-npr\|lincRNA372_PMID_24635777\|TC | 2,579 |
| SALT00000078014_Cluster43-099 | 0.510 | sit-MIR121-1-npr\|NONATHT003847\|lincRNA12_PN | 3,215 |
| SALT00000078290_Cluster43-100 | 0.593 | TCONS_00018605\|sit-MIR122-1-npr\|lincRNA372_P | 3,223 |
| SALT00000080000_Cluster43-101 | 0.595 | ghr-MIR5368b\|NONATHT003839\|lincRNA493_PMI | 3,306 |
| SALT00000080070_Cluster43-102 | 0.609 | gma-MIR4995\|ghr-MIR5368*\|sit-MIR121-1-npr\|linc | 3,221 |
| SALT00000080076_Cluster43-103 | 0.607 | TCONS_00018605\|sit-MIR122-1-npr\|lincRNA372_P | 3,234 |
| SALT00000080469_Cluster43-104 | 0.607 | TCONS_00018605\|sit-MIR122-1-npr\|lincRNA372_P | 3,233 |
| SALT00000080656_Cluster43-105 | 0.631 | NONATHT003839\|ghr-MIR5368b\|ghr-MIR5368*\|sit | 2,475 |
| SALT00000080779_Cluster43-106 | 0.611 | ghr-MIR5368b\|NONATHT003839\|lincRNA493_PMI | 3,213 |
| SALT00000080920_Cluster43-107 | 0.532 | TCONS_00018605\|lincRNA372_PMID_24635777\|g | 2,838 |
| SALT00000081451_Cluster43-108 | 0.500 | lincRNA493_PMID_24635777\|NONATHT003839\|gl | 3,230 |
| SALT00000081927_Cluster43-109 | 0.451 | TCONS_00018605\|sit-MIR122-1-npr\|lincRNA372_P | 3,510 |
| SALT00000082679_Cluster43-110 | 0.446 | lincRNA372_PMID_24635777\|sit-MIR122-1-npr\|TC | 3,537 |
| SALT00000082811_Cluster43-111 | 0.396 | sit-MIR121-1-npr\|NONATHT003847\|lincRNA12_PN | 2,824 |
| SALT00000082848_Cluster43-112 | 0.499 | TCONS_00018605\|sit-MIR122-1-npr\|lincRNA372_P | 3,241 |
| SALT00000082852_Cluster43-113 | 0.488 | lincRNA372_PMID_24635777\|sit-MIR122-1-npr\|TC | 3,302 |
| SALT00000083799_Cluster43-114 | 0.480 | sit-MIR122-1-npr\|lincRNA372_PMID_24635777\|TC | 3,348 |
| SALT00000083914_Cluster43-115 | 0.631 | lincRNA372_PMID_24635777\|sit-MIR122-1-npr\|TC | 2,479 |
| SALT00000084603_Cluster43-116 | 0.500 | lincRNA493_PMID_24635777\|NONATHT003839\|gl | 3,231 |
| SALT00000084922_Cluster43-117 | 0.482 | sit-MIR122-1-npr\|lincRNA372_PMID_24635777\|TC | 3,214 |
| SALT00000085236_Cluster43-118 | 0.607 | TCONS_00018605\|lincRNA372_PMID_24635777\|si | 3,237 |
| SALT00000085558_Cluster43-119 | 0.502 | NONATHT003847\|lincRNA12_PMID_24635777\|sit- | 3,222 |
| SALT00000085669_Cluster43-120 | 0.604 | TCONS_00018605\|lincRNA372_PMID_24635777\|si | 3,252 |
| SALT00000085757_Cluster43-121 | 0.502 | TCONS_00018605\|sit-MIR122-1-npr\|lincRNA493_P | 3,220 |
| SALT00000086137_Cluster43-123 | 0.608 | ghr-MIR5368b\|NONATHT003839\|lincRNA493_PMI | 3,232 |
| SALT00000086308_Cluster43-124 | 0.607 | ghr-MIR5368b\|NONATHT003839\|lincRNA493_PMI | 3,234 |
| SALT00000086325_Cluster43-125 | 0.608 | TCONS_00018605\|lincRNA372_PMID_24635777\|si | 3,231 |
| SALT00000086697_Cluster43-126 | 0.500 | lincRNA372_PMID_24635777\|TCONS_00018605\|li | 3,235 |
| SALT00000086902_Cluster43-127 | 0.608 | sit-MIR121-1-npr\|NONATHT003847\|lincRNA12_PN | 3,230 |
| SALT00000087164_Cluster43-128 | 0.609 | lincRNA493_PMID_24635777\|NONATHT003839\|gl | 3,223 |
| SALT00000087462_Cluster43-129 | 0.502 | sit-MIR121-1-npr\|lincRNA12_PMID_24635777\|NOI | 3,222 |
| SALT00000087472_Cluster43-130 | 0.607 | TCONS_00018605\|lincRNA372_PMID_24635777\|si | 3,236 |
| SALT00000087810_Cluster43-131 | 0.500 | NONATHT003839\|lincRNA493_PMID_24635777\|gl | 3,234 |
| SALT00000088374_Cluster43-132 | 0.602 | lincRNA372_PMID_24635777\|sit-MIR122-1-npr\|TC | 3,264 |
| SALT00000088944_Cluster43-133 | 0.497 | gma-MIR4995\|ghr-MIR5368*\|sit-MIR121-1-npr\|linc | 3,247 |
| SALT00000088977_Cluster43-134 | 0.490 | sit-MIR122-1-npr\|lincRNA372_PMID_24635777\|TC | 3,324 |
| SALT00000089460_Cluster43-135 | 0.606 | gma-MIR4995\|ghr-MIR5368*\|sit-MIR121-1-npr\|NOI | 3,239 |
| SALT00000089530_Cluster43-136 | 0.497 | TCONS_00018605\|sit-MIR122-1-npr\|lincRNA372_P | 3,249 |
| SALT00000089847_Cluster43-137 | 0.606 | TCONS_00018605\|sit-MIR122-1-npr\|lincRNA372_P | 3,243 |
| SALT00000090171_Cluster43-138 | 0.597 | NONATHT003847\|lincRNA12_PMID_24635777\|sit- | 3,293 |
| SALT00000090499_Cluster43-139 | 0.599 | TCONS_00018605\|sit-MIR122-1-npr\|lincRNA372_P | 3,285 |
| SALT00000090558_Cluster43-140 | 0.152 | sit-MIR122-1-npr\|lincRNA372_PMID_24635777\|TC | 3,223 |
| SALT00000091047_Cluster43-141 | 0.608 | lincRNA372_PMID_24635777\|sit-MIR122-1-npr\|TC | 3,232 |
| SALT00000091366_Cluster43-143 | 0.151 | TCONS_00018605\|sit-MIR122-1-npr\|NONATHT003 | 3,233 |
| SALT00000091656_Cluster43-144 | 0.504 | lincRNA372_PMID_24635777\|sit-MIR122-1-npr\|TC | 3,212 |
| SALT00000092427_Cluster43-145 | 0.966 | ghr-MIR5368*\|gma-MIR4995\|lincRNA12_PMID_24 | 3,306 |
| SALT00000092632_Cluster43-146 | 0.998 | ghr-MIR5368*\|gma-MIR4995\|lincRNA12_PMID_24 | 3,653 |
| SALT00000093066_Cluster43-147 | 0.606 | ghr-MIR5368b\|lincRNA493_PMID_24635777\|NON. | 3,241 |
| SALT00000093549_Cluster43-148 | 0.499 | TCONS_00018605\|lincRNA372_PMID_24635777\|si | 3,240 |
| SALT00000093906_Cluster43-149 | 0.540 | lincRNA372_PMID_24635777\|sit-MIR122-1-npr\|TC | 3,005 |
| SALT00000081158_Cluster43-150 | 0.990 | lincRNA372_PMID_24635777\|sit-MIR122-1-npr\|TC | 3,401 |
| SALT00000085349_Cluster43-151 | 0.592 | NONATHT003839\|lincRNA493_PMID_24635777\|si | 3,321 |
| SALT00000092934_Cluster43-153 | 0.938 | gma-MIR4995\|ghr-MIR5368*\|sit-MIR121-1-npr\|linc | 3,748 |
| SALT00000075997_Cluster43-154 | 0.512 | TCONS_00018605\|lincRNA372_PMID_24635777\|g | 3,167 |
| SALT00000079178_Cluster43-155 | 0.248 | sit-MIR122-1-npr\|TCONS_00018605\|ghr-MIR5368* | 3,521 |
| SALT00000090344_Cluster55-001 | 0.094 | NA | 13,036 |
| SALT00000086318_Cluster57-028 | 0.040 | NA | 9,393 |

| | | | |
|---|---|---|---|
| SALT00000009459_Cluster80-001 | 1.000 | lincRNA11_PMID_24635777\| | 1,577 |
| SALT00000009488_Cluster80-002 | 1.000 | lincRNA11_PMID_24635777\| | 1,767 |
| SALT00000012212_Cluster80-003 | 1.000 | lincRNA11_PMID_24635777\| | 1,275 |
| SALT00000012393_Cluster80-004 | 1.000 | lincRNA11_PMID_24635777\| | 1,899 |
| SALT00000012735_Cluster80-005 | 1.000 | lincRNA11_PMID_24635777\| | 2,050 |
| SALT00000014777_Cluster80-006 | 1.000 | lincRNA11_PMID_24635777\| | 1,799 |
| SALT00000017656_Cluster80-007 | 1.000 | lincRNA11_PMID_24635777\| | 1,062 |
| SALT00000017778_Cluster80-008 | 1.000 | lincRNA11_PMID_24635777\| | 1,878 |
| SALT00000020473_Cluster80-009 | 1.000 | lincRNA11_PMID_24635777\| | 2,287 |
| SALT00000020524_Cluster80-010 | 1.000 | lincRNA11_PMID_24635777\| | 1,713 |
| SALT00000020587_Cluster80-011 | 1.000 | lincRNA11_PMID_24635777\| | 2,812 |
| SALT00000020594_Cluster80-012 | 1.000 | lincRNA11_PMID_24635777\| | 2,963 |
| SALT00000020728_Cluster80-013 | 1.000 | lincRNA11_PMID_24635777\| | 3,551 |
| SALT00000023740_Cluster80-014 | 1.000 | lincRNA11_PMID_24635777\| | 2,554 |
| SALT00000023996_Cluster80-015 | 1.000 | lincRNA11_PMID_24635777\| | 2,851 |
| SALT00000025029_Cluster80-016 | 1.000 | lincRNA11_PMID_24635777\| | 2,497 |
| SALT00000025215_Cluster80-017 | 1.000 | lincRNA11_PMID_24635777\| | 2,747 |
| SALT00000028127_Cluster80-018 | 1.000 | lincRNA11_PMID_24635777\| | 3,317 |
| SALT00000028349_Cluster80-019 | 1.000 | lincRNA11_PMID_24635777\| | 3,170 |
| SALT00000028357_Cluster80-020 | 1.000 | lincRNA11_PMID_24635777\| | 2,631 |
| SALT00000028500_Cluster80-021 | 1.000 | lincRNA11_PMID_24635777\| | 2,152 |
| SALT00000028529_Cluster80-022 | 1.000 | lincRNA11_PMID_24635777\| | 4,890 |
| SALT00000028569_Cluster80-023 | 1.000 | lincRNA11_PMID_24635777\| | 4,121 |
| SALT00000028623_Cluster80-024 | 1.000 | lincRNA11_PMID_24635777\| | 3,839 |
| SALT00000028718_Cluster80-025 | 1.000 | lincRNA11_PMID_24635777\| | 4,262 |
| SALT00000029945_Cluster80-026 | 1.000 | lincRNA11_PMID_24635777\| | 4,319 |
| SALT00000030421_Cluster80-027 | 1.000 | lincRNA11_PMID_24635777\| | 4,660 |
| SALT00000031064_Cluster80-028 | 1.000 | lincRNA11_PMID_24635777\| | 4,734 |
| SALT00000031222_Cluster80-029 | 1.000 | lincRNA11_PMID_24635777\| | 2,697 |
| SALT00000031250_Cluster80-030 | 1.000 | lincRNA11_PMID_24635777\| | 5,111 |
| SALT00000031441_Cluster80-031 | 1.000 | lincRNA11_PMID_24635777\| | 1,606 |
| SALT00000031452_Cluster80-032 | 1.000 | lincRNA11_PMID_24635777\| | 2,019 |
| SALT00000031472_Cluster80-033 | 1.000 | lincRNA11_PMID_24635777\| | 2,679 |
| SALT00000031491_Cluster80-034 | 1.000 | lincRNA11_PMID_24635777\| | 1,501 |
| SALT00000031567_Cluster80-035 | 1.000 | lincRNA11_PMID_24635777\| | 3,933 |
| SALT00000031876_Cluster80-037 | 1.000 | lincRNA11_PMID_24635777\| | 4,813 |
| SALT00000032027_Cluster80-038 | 1.000 | lincRNA11_PMID_24635777\| | 4,929 |
| SALT00000032875_Cluster80-039 | 1.000 | lincRNA11_PMID_24635777\| | 4,868 |
| SALT00000033088_Cluster80-040 | 1.000 | lincRNA11_PMID_24635777\| | 4,175 |
| SALT00000034467_Cluster80-041 | 1.000 | lincRNA11_PMID_24635777\| | 4,378 |
| SALT00000034626_Cluster80-042 | 1.000 | lincRNA11_PMID_24635777\| | 3,318 |
| SALT00000034739_Cluster80-043 | 1.000 | lincRNA11_PMID_24635777\| | 3,603 |
| SALT00000034759_Cluster80-044 | 1.000 | lincRNA11_PMID_24635777\| | 3,798 |
| SALT00000034817_Cluster80-045 | 1.000 | lincRNA11_PMID_24635777\| | 4,710 |
| SALT00000034879_Cluster80-046 | 1.000 | lincRNA11_PMID_24635777\| | 3,727 |
| SALT00000034914_Cluster80-047 | 1.000 | lincRNA11_PMID_24635777\| | 1,537 |
| SALT00000034943_Cluster80-048 | 1.000 | lincRNA11_PMID_24635777\| | 3,711 |
| SALT00000034944_Cluster80-049 | 1.000 | lincRNA11_PMID_24635777\| | 1,985 |
| SALT00000034973_Cluster80-050 | 1.000 | lincRNA11_PMID_24635777\| | 4,561 |
| SALT00000035164_Cluster80-051 | 1.000 | lincRNA11_PMID_24635777\| | 4,149 |
| SALT00000035539_Cluster80-052 | 1.000 | lincRNA11_PMID_24635777\| | 4,043 |
| SALT00000035575_Cluster80-053 | 1.000 | lincRNA11_PMID_24635777\| | 3,528 |
| SALT00000039439_Cluster80-054 | 1.000 | lincRNA11_PMID_24635777\| | 2,187 |
| SALT00000039630_Cluster80-055 | 0.947 | lincRNA11_PMID_24635777\| | 1,910 |
| SALT00000041620_Cluster80-056 | 1.000 | lincRNA11_PMID_24635777\| | 2,206 |
| SALT00000043494_Cluster80-057 | 0.996 | lincRNA11_PMID_24635777\| | 1,753 |
| SALT00000053265_Cluster80-058 | 0.999 | lincRNA11_PMID_24635777\| | 1,531 |
| SALT00000053614_Cluster80-059 | 0.999 | lincRNA11_PMID_24635777\| | 1,807 |

| | | | |
|---|---|---|---|
| SALT00000055324_Cluster80-060 | 0.999 | lincRNA11_PMID_24635777\| | 2,377 |
| SALT00000057152_Cluster80-061 | 1.000 | lincRNA11_PMID_24635777\| | 1,772 |
| SALT00000062674_Cluster80-062 | 1.000 | lincRNA11_PMID_24635777\| | 1,973 |
| SALT00000063154_Cluster80-063 | 1.000 | lincRNA11_PMID_24635777\| | 2,831 |
| SALT00000066895_Cluster80-064 | 1.000 | lincRNA11_PMID_24635777\| | 2,895 |
| SALT00000066907_Cluster80-065 | 1.000 | lincRNA11_PMID_24635777\| | 3,463 |
| SALT00000068579_Cluster80-067 | 1.000 | lincRNA11_PMID_24635777\| | 2,680 |
| SALT00000071125_Cluster80-068 | 1.000 | lincRNA11_PMID_24635777\| | 3,035 |
| SALT00000071736_Cluster80-069 | 1.000 | lincRNA11_PMID_24635777\| | 4,174 |
| SALT00000071842_Cluster80-070 | 1.000 | lincRNA11_PMID_24635777\| | 1,166 |
| SALT00000074209_Cluster80-071 | 1.000 | lincRNA11_PMID_24635777\| | 3,134 |
| SALT00000074987_Cluster80-072 | 1.000 | lincRNA11_PMID_24635777\| | 3,126 |
| SALT00000076652_Cluster80-073 | 1.000 | lincRNA11_PMID_24635777\| | 2,025 |
| SALT00000076674_Cluster80-074 | 1.000 | lincRNA11_PMID_24635777\| | 1,532 |
| SALT00000076677_Cluster80-075 | 1.000 | lincRNA11_PMID_24635777\| | 3,227 |
| SALT00000076693_Cluster80-076 | 1.000 | lincRNA11_PMID_24635777\| | 4,248 |
| SALT00000077526_Cluster80-077 | 1.000 | lincRNA11_PMID_24635777\| | 3,251 |
| SALT00000079165_Cluster80-078 | 1.000 | lincRNA11_PMID_24635777\| | 3,210 |
| SALT00000079815_Cluster80-079 | 1.000 | lincRNA11_PMID_24635777\| | 3,371 |
| SALT00000081530_Cluster80-080 | 1.000 | lincRNA11_PMID_24635777\| | 3,874 |
| SALT00000082137_Cluster80-081 | 1.000 | lincRNA11_PMID_24635777\| | 4,291 |
| SALT00000082150_Cluster80-082 | 1.000 | lincRNA11_PMID_24635777\| | 3,939 |
| SALT00000082677_Cluster80-083 | 1.000 | lincRNA11_PMID_24635777\| | 3,749 |
| SALT00000083887_Cluster80-084 | 1.000 | lincRNA11_PMID_24635777\| | 2,934 |
| SALT00000084313_Cluster80-085 | 1.000 | lincRNA11_PMID_24635777\| | 3,790 |
| SALT00000084446_Cluster80-086 | 1.000 | lincRNA11_PMID_24635777\| | 5,024 |
| SALT00000084549_Cluster80-087 | 1.000 | lincRNA11_PMID_24635777\| | 3,283 |
| SALT00000084698_Cluster80-088 | 1.000 | lincRNA11_PMID_24635777\| | 3,349 |
| SALT00000085035_Cluster80-089 | 1.000 | lincRNA11_PMID_24635777\| | 3,955 |
| SALT00000085232_Cluster80-090 | 1.000 | lincRNA11_PMID_24635777\| | 4,873 |
| SALT00000085412_Cluster80-091 | 1.000 | lincRNA11_PMID_24635777\| | 3,829 |
| SALT00000085490_Cluster80-092 | 1.000 | lincRNA11_PMID_24635777\| | 5,092 |
| SALT00000085562_Cluster80-093 | 1.000 | lincRNA11_PMID_24635777\| | 4,923 |
| SALT00000085595_Cluster80-094 | 1.000 | lincRNA11_PMID_24635777\| | 4,940 |
| SALT00000085710_Cluster80-095 | 1.000 | lincRNA11_PMID_24635777\| | 5,039 |
| SALT00000085830_Cluster80-096 | 1.000 | lincRNA11_PMID_24635777\| | 4,916 |
| SALT00000086149_Cluster80-098 | 1.000 | lincRNA11_PMID_24635777\| | 5,098 |
| SALT00000087046_Cluster80-099 | 1.000 | lincRNA11_PMID_24635777\| | 4,669 |
| SALT00000087051_Cluster80-100 | 1.000 | lincRNA11_PMID_24635777\| | 5,535 |
| SALT00000087300_Cluster80-101 | 1.000 | lincRNA11_PMID_24635777\| | 4,164 |
| SALT00000087582_Cluster80-102 | 1.000 | lincRNA11_PMID_24635777\| | 4,865 |
| SALT00000087655_Cluster80-103 | 1.000 | lincRNA11_PMID_24635777\| | 3,753 |
| SALT00000088054_Cluster80-104 | 1.000 | lincRNA11_PMID_24635777\| | 4,910 |
| SALT00000088164_Cluster80-105 | 1.000 | lincRNA11_PMID_24635777\| | 4,874 |
| SALT00000088244_Cluster80-106 | 1.000 | lincRNA11_PMID_24635777\| | 5,127 |
| SALT00000088625_Cluster80-107 | 1.000 | lincRNA11_PMID_24635777\| | 4,962 |
| SALT00000088759_Cluster80-108 | 1.000 | lincRNA11_PMID_24635777\| | 4,331 |
| SALT00000088830_Cluster80-109 | 1.000 | lincRNA11_PMID_24635777\| | 4,146 |
| SALT00000089061_Cluster80-110 | 1.000 | lincRNA11_PMID_24635777\| | 4,877 |
| SALT00000089271_Cluster80-111 | 1.000 | lincRNA11_PMID_24635777\| | 4,574 |
| SALT00000089340_Cluster80-112 | 1.000 | lincRNA11_PMID_24635777\| | 4,879 |
| SALT00000089578_Cluster80-113 | 1.000 | lincRNA11_PMID_24635777\| | 5,978 |
| SALT00000089693_Cluster80-114 | 1.000 | lincRNA11_PMID_24635777\| | 4,888 |
| SALT00000089840_Cluster80-115 | 1.000 | lincRNA11_PMID_24635777\| | 4,837 |
| SALT00000090204_Cluster80-116 | 1.000 | lincRNA11_PMID_24635777\| | 4,897 |
| SALT00000090784_Cluster80-117 | 1.000 | lincRNA11_PMID_24635777\| | 4,889 |
| SALT00000090878_Cluster80-118 | 1.000 | lincRNA11_PMID_24635777\| | 4,570 |
| SALT00000090888_Cluster80-119 | 1.000 | lincRNA11_PMID_24635777\| | 4,864 |

| | | | |
|---|---|---|---|
| SALT00000090973_Cluster80-120 | 1.000 | lincRNA11_PMID_24635777\| | 4,894 |
| SALT00000091509_Cluster80-121 | 1.000 | lincRNA11_PMID_24635777\| | 3,403 |
| SALT00000091724_Cluster80-122 | 0.061 | lincRNA11_PMID_24635777\| | 4,254 |
| SALT00000091733_Cluster80-123 | 1.000 | lincRNA11_PMID_24635777\| | 4,865 |
| SALT00000092359_Cluster80-125 | 1.000 | lincRNA11_PMID_24635777\| | 5,038 |
| SALT00000092627_Cluster80-126 | 1.000 | lincRNA11_PMID_24635777\| | 5,085 |
| SALT00000093001_Cluster80-127 | 1.000 | lincRNA11_PMID_24635777\| | 3,639 |
| SALT00000093022_Cluster80-128 | 1.000 | lincRNA11_PMID_24635777\| | 3,828 |
| SALT00000093309_Cluster80-129 | 1.000 | lincRNA11_PMID_24635777\| | 4,896 |
| SALT00000093623_Cluster80-130 | 1.000 | lincRNA11_PMID_24635777\| | 3,349 |
| SALT00000093672_Cluster80-131 | 1.000 | lincRNA11_PMID_24635777\| | 3,982 |
| SALT00000093679_Cluster80-132 | 1.000 | lincRNA11_PMID_24635777\| | 1,475 |
| SALT00000093706_Cluster80-133 | 1.000 | lincRNA11_PMID_24635777\| | 2,956 |
| SALT00000093713_Cluster80-134 | 1.000 | lincRNA11_PMID_24635777\| | 1,491 |
| SALT00000093718_Cluster80-135 | 1.000 | lincRNA11_PMID_24635777\| | 4,528 |
| SALT00000093797_Cluster80-136 | 1.000 | lincRNA11_PMID_24635777\| | 3,722 |
| SALT00000093885_Cluster80-137 | 0.970 | lincRNA11_PMID_24635777\| | 1,383 |
| SALT00000070248_Cluster80-138 | 0.121 | lincRNA11_PMID_24635777\| | 4,299 |
| SALT00000093274_Cluster80-139 | 1.000 | lincRNA11_PMID_24635777\| | 4,917 |
| SALT00000091443_Cluster80-140 | 1.000 | lincRNA11_PMID_24635777\| | 4,670 |
| SALT00000086600_Cluster80-141 | 1.000 | lincRNA11_PMID_24635777\| | 4,363 |
| SALT00000088949_Cluster80-142 | 1.000 | lincRNA11_PMID_24635777\| | 6,661 |
| SALT00000070652_Cluster80-145 | 0.996 | lincRNA11_PMID_24635777\| | 4,707 |
| SALT00000077014_Cluster80-147 | 1.000 | lincRNA11_PMID_24635777\| | 4,811 |
| SALT00000083686_Cluster95-001 | 0.153 | NA | 2,944 |
| SALT00000006720_Cluster117-001 | 0.519 | sit-MIR121-1-npr\|NONATHT003847\|lincRNA12_PM | 2,126 |
| SALT00000028509_Cluster117-002 | 0.562 | lincRNA372_PMID_24635777\|sit-MIR122-1-npr\|TC | 1,877 |
| SALT00000028678_Cluster117-003 | 0.919 | lincRNA372_PMID_24635777\|sit-MIR122-1-npr\|TC | 4,252 |
| SALT00000031088_Cluster117-004 | 0.929 | TCONS_00018605\|sit-MIR122-1-npr\|lincRNA372_P | 4,043 |
| SALT00000031886_Cluster117-005 | 0.905 | lincRNA12_PMID_24635777\|NONATHT003847\|sit- | 4,494 |
| SALT00000033881_Cluster117-006 | 0.938 | sit-MIR121-1-npr\|lincRNA12_PMID_24635777\|NON | 3,836 |
| SALT00000034559_Cluster117-007 | 0.870 | gma-MIR4995\|ghr-MIR5368*\|sit-MIR121-1-npr\|lincl | 4,994 |
| SALT00000034698_Cluster117-008 | 0.758 | TCONS_00018605\|sit-MIR122-1-npr\|lincRNA372_P | 1,623 |
| SALT00000035641_Cluster117-009 | 0.472 | sit-MIR122-1-npr\|lincRNA372_PMID_24635777\|TC | 2,390 |
| SALT00000064615_Cluster117-010 | 0.986 | ghr-MIR5368b\|NONATHT003839\|lincRNA493_PMI | 5,003 |
| SALT00000070195_Cluster117-011 | 0.613 | ghr-MIR5368b\|NONATHT003839\|NONATHT00384 | 2,585 |
| SALT00000073189_Cluster117-012 | 0.228 | sit-MIR122-1-npr\|lincRNA372_PMID_24635777\|TC | 2,513 |
| SALT00000076918_Cluster117-013 | 0.929 | TCONS_00018605\|lincRNA372_PMID_24635777\|si | 4,054 |
| SALT00000077034_Cluster117-014 | 0.128 | lincRNA372_PMID_24635777\|TCONS_00018605\|g | 4,947 |
| SALT00000081969_Cluster117-015 | 0.924 | NONATHT003839\|lincRNA493_PMID_24635777\|gl | 4,159 |
| SALT00000083585_Cluster117-016 | 0.749 | sit-MIR121-1-npr\|NONATHT003847\|lincRNA12_PM | 2,610 |
| SALT00000086209_Cluster117-017 | 0.137 | ghr-MIR5368b\|NONATHT003839\|lincRNA493_PMI | 10,999 |
| SALT00000087019_Cluster117-018 | 0.934 | lincRNA372_PMID_24635777\|sit-MIR122-1-npr\|TC | 3,940 |
| SALT00000088579_Cluster117-019 | 1.000 | TCONS_00018605\|sit-MIR122-1-npr\|NONATHT003 | 4,695 |
| SALT00000092084_Cluster117-020 | 0.455 | sit-MIR122-1-npr\|TCONS_00018605\|lincRNA12_PM | 3,486 |
| SALT00000072610_Cluster117-023 | 0.418 | NONATHT003839\|lincRNA493_PMID_24635777\|gl | 4,044 |
| SALT00000081481_Cluster117-024 | 0.578 | lincRNA372_PMID_24635777\|TCONS_00018605\|li | 3,403 |
| SALT00000066763_Cluster117-025 | 0.655 | sit-MIR122-1-npr\|lincRNA372_PMID_24635777\|TC | 2,945 |
| SALT00000092454_Cluster117-026 | 0.878 | lincRNA493_PMID_24635777\|NONATHT003839\|g | 4,889 |
| SALT00000092544_Cluster119-003 | 0.083 | NA | 4,271 |
| SALT00000058184_Cluster155-001 | 1.000 | lincRNA11_PMID_24635777\| | 1,576 |
| SALT00000083257_Cluster155-002 | 1.000 | lincRNA11_PMID_24635777\| | 10,172 |
| SALT00000050681_Cluster161-005 | 1.000 | lincRNA688_PMID_24635777\| | 1,791 |
| SALT00000000696_Cluster183-001 | 0.997 | lincRNA676_PMID_24635777\| | 1,624 |
| SALT00000002323_Cluster183-002 | 0.999 | lincRNA676_PMID_24635777\| | 1,692 |
| SALT00000008400_Cluster183-003 | 1.000 | pnrd_Mtr_chr1.trna88_HisGTG\|pnrd_Gma_scaffold_ | 2,010 |
| SALT00000009297_Cluster183-004 | 1.000 | pnrd_Mtr_chr1.trna2_HisGTG\|pnrd_Mtr_chr8.trna33 | 1,864 |
| SALT00000011978_Cluster183-005 | 0.997 | pnrd_Mtr_chr8.trna33_HisGTG\|pnrd_Gma_scaffold_ | 1,759 |

| | | | |
|---|---|---|---|
| SALT00000019964_Cluster183-006 | 0.998 | pnrd_Osa_chr4.trna91_HisGTG\|pnrd_Mtr_chr7.trna7 | 1,695 |
| SALT00000020091_Cluster183-007 | 1.000 | pnrd_Osa_chr10.trna13_HisGTG\|pnrd_Gma_scaffolc | 2,002 |
| SALT00000021114_Cluster183-008 | 1.000 | pnrd_Gma_chr13.trna72_HisGTG\|pnrd_Mtr_chr8.trn | 3,069 |
| SALT00000025378_Cluster183-009 | 1.000 | pnrd_Osa_chr8.trna27_HisGTG\|pnrd_Osa_chr9.trna3 | 3,041 |
| SALT00000027484_Cluster183-010 | 1.000 | pnrd_Gma_scaffold_828.trna2_HisGTG\|pnrd_Osa_cl | 2,903 |
| SALT00000027877_Cluster183-011 | 1.000 | pnrd_Osa_chr10.trna27_HisGTG\|pnrd_Gma_scaffolc | 2,915 |
| SALT00000028983_Cluster183-012 | 1.000 | pnrd_Gma_scaffold_931.trna1_HisGTG\|pnrd_Gma_s | 2,762 |
| SALT00000031944_Cluster183-013 | 1.000 | pnrd_Mtr_chr1.trna88_HisGTG\|pnrd_Gma_scaffold_ | 3,581 |
| SALT00000033274_Cluster183-014 | 1.000 | pnrd_Gma_scaffold_998.trna2_HisGTG\|pnrd_Osa_cl | 2,938 |
| SALT00000036041_Cluster183-015 | 0.998 | pnrd_Osa_chr4.trna41_HisGTG\|pnrd_Osa_chr11.trna | 1,845 |
| SALT00000043211_Cluster183-016 | 1.000 | pnrd_Mtr_chr5.trna1_HisGTG\|pnrd_Osa_chr12.trna1 | 2,336 |
| SALT00000046052_Cluster183-017 | 0.935 | lincRNA676_PMID_24635777\| | 1,456 |
| SALT00000046985_Cluster183-018 | 0.994 | pnrd_Gma_scaffold_998.trna2_HisGTG\|pnrd_Osa_cl | 1,605 |
| SALT00000053438_Cluster183-019 | 0.995 | lincRNA676_PMID_24635777\| | 1,216 |
| SALT00000054124_Cluster183-020 | 0.997 | pnrd_Osa_chr8.trna46_HisGTG\|pnrd_Gma_scaffold_ | 2,208 |
| SALT00000057487_Cluster183-021 | 1.000 | pnrd_Osa_chr11.trna7_HisGTG\|pnrd_Osa_chr4.trna4 | 2,139 |
| SALT00000062070_Cluster183-022 | 0.997 | pnrd_Mtr_chr1.trna88_HisGTG\|pnrd_Osa_chr10.trna | 2,460 |
| SALT00000065755_Cluster183-023 | 0.916 | lincRNA676_PMID_24635777\| | 1,870 |
| SALT00000067754_Cluster183-024 | 1.000 | osa-MIR5538\|pnrd_Mtr_chr8.trna33_HisGTG\|pnrd_( | 3,047 |
| SALT00000080340_Cluster183-025 | 1.000 | pnrd_Osa_chr4.trna41_HisGTG\|pnrd_Osa_chr11.trna | 2,926 |
| SALT00000082895_Cluster183-026 | 1.000 | pnrd_Mtr_chr1.trna88_HisGTG\|pnrd_Gma_scaffold_ | 3,068 |
| SALT00000091281_Cluster183-027 | 0.997 | pnrd_Mtr_chr8.trna33_HisGTG\|pnrd_Gma_scaffold_ | 9,656 |
| SALT00000051815_Cluster194-005 | 1.000 | NONATHT002169\|tae-MIR170b_npr\|tae-MIR170a_r | 9,571 |
| SALT00000085313_Cluster202-001 | 0.702 | osa-MIR444c\|osa-MIR444d\|oru-MIR444\|bdi-MIR44 | 4,756 |
| SALT00000017168_Cluster202-003 | 0.250 | oru-MIR444\|osa-MIR444d\|osa-MIR444c\|bdi-MIR44 | 1,504 |
| SALT00000090349_Cluster211-006 | 1.000 | lincRNA12_PMID_24635777\|NONATHT003847\|gm | 9,356 |
| SALT00000008933_Cluster220-006 | 0.101 | pde-MIR1310\|han-MIR1310\|ghr-MIR4370\|cln-MIR1 | 1,548 |
| SALT00000049394_Cluster220-035 | 0.232 | pta-MIR1310\|ghr-MIR4370\|cln-MIR1310\|pde-MIR1 | 1,653 |
| SALT00000050979_Cluster220-036 | 0.852 | han-MIR1310\|pde-MIR1310\|pta-MIR1310\|cln-MIR1 | 1,881 |
| SALT00000064378_Cluster220-041 | 0.164 | pta-MIR1310\|ghr-MIR4370\|cln-MIR1310\|pde-MIR1 | 2,446 |
| SALT00000031125_Cluster220-045 | 0.995 | han-MIR1310\|pde-MIR1310\|cln-MIR1310\|pta-MIR1 | 2,765 |
| SALT00000001026_Cluster238-001 | 1.000 | lincRNA304_PMID_24635777\| | 2,124 |
| SALT00000001641_Cluster238-002 | 1.000 | lincRNA304_PMID_24635777\| | 1,662 |
| SALT00000001711_Cluster238-003 | 1.000 | lincRNA304_PMID_24635777\| | 2,021 |
| SALT00000003298_Cluster238-004 | 1.000 | lincRNA304_PMID_24635777\| | 1,563 |
| SALT00000006779_Cluster238-005 | 1.000 | lincRNA304_PMID_24635777\| | 1,953 |
| SALT00000011232_Cluster238-006 | 1.000 | lincRNA304_PMID_24635777\| | 1,765 |
| SALT00000018027_Cluster238-007 | 1.000 | lincRNA304_PMID_24635777\| | 1,583 |
| SALT00000023702_Cluster238-008 | 1.000 | lincRNA304_PMID_24635777\| | 2,289 |
| SALT00000028451_Cluster238-009 | 1.000 | lincRNA304_PMID_24635777\| | 1,756 |
| SALT00000032094_Cluster238-010 | 1.000 | lincRNA304_PMID_24635777\| | 1,710 |
| SALT00000034905_Cluster238-011 | 1.000 | lincRNA304_PMID_24635777\| | 1,608 |
| SALT00000035583_Cluster238-012 | 1.000 | lincRNA304_PMID_24635777\| | 1,484 |
| SALT00000037994_Cluster238-013 | 1.000 | lincRNA304_PMID_24635777\| | 1,510 |
| SALT00000076655_Cluster238-014 | 1.000 | lincRNA304_PMID_24635777\| | 1,820 |
| SALT00000084574_Cluster238-016 | 1.000 | lincRNA304_PMID_24635777\| | 1,401 |
| SALT00000087776_Cluster240-030 | 1.000 | lincRNA688_PMID_24635777\| | 1,740 |
| SALT00000023973_Cluster241-001 | 1.000 | pnrd_Mtr_chr3.trna109_MetCAT\|pnrd_Osa_chr12.trn | 2,894 |
| SALT00000028535_Cluster241-002 | 1.000 | pnrd_Osa_chr12.trna61_MetCAT\|pnrd_Mtr_chr3.trna | 3,652 |
| SALT00000031594_Cluster241-003 | 1.000 | pnrd_Osa_chr3.trna31_MetCAT\|pnrd_Osa_chr12.trna | 2,838 |
| SALT00000032305_Cluster241-004 | 1.000 | pnrd_Osa_chr12.trna15_MetCAT\|pnrd_Osa_chr12.trn | 3,482 |
| SALT00000033423_Cluster241-005 | 1.000 | pnrd_Mtr_chr3.trna72_MetCAT\|pnrd_Gma_scaffold_ | 3,413 |
| SALT00000034911_Cluster241-006 | 1.000 | pnrd_Osa_chr3.trna31_MetCAT\|pnrd_Osa_chr12.trna | 2,690 |
| SALT00000061048_Cluster241-007 | 0.151 | NA | 1,079 |
| SALT00000079443_Cluster241-008 | 1.000 | pnrd_Mtr_chr3.trna109_MetCAT\|pnrd_Osa_chr12.trn | 3,642 |
| SALT00000081809_Cluster241-010 | 1.000 | pnrd_Mtr_chr3.trna72_MetCAT\|pnrd_Gma_scaffold_ | 2,519 |
| SALT00000085585_Cluster241-011 | 1.000 | pnrd_Osa_chr6.trna3_MetCAT\|pnrd_Mtr_chr3.trna67 | 3,637 |
| SALT00000086302_Cluster241-012 | 1.000 | pnrd_Osa_chr6.trna3_MetCAT\|pnrd_Gma_scaffold_1 | 3,643 |

| | | | |
|---|---|---|---|
| SALT00000089114_Cluster241-013 | 1.000 | pnrd_Osa_chr12.trna61_MetCAT\|pnrd_Mtr_chr3.trna | 3,653 |
| SALT00000091480_Cluster241-014 | 1.000 | pnrd_Mtr_chr3.trna109_MetCAT\|pnrd_Osa_chr12.trn | 3,616 |
| SALT00000092905_Cluster241-015 | 1.000 | pnrd_Ara_chr2.trna90_MetCAT\|pnrd_Gma_scaffold_ | 3,644 |
| SALT00000090063_Cluster241-016 | 1.000 | pnrd_Mtr_chr3.trna67_MetCAT\|pnrd_Gma_scaffold_ | 4,141 |
| SALT00000087232_Cluster241-017 | 1.000 | pnrd_Osa_chr6.trna3_MetCAT\|pnrd_Gma_scaffold_1 | 3,571 |
| SALT00000047645_Cluster272-001 | 0.027 | NA | 8,829 |
| SALT00000088886_Cluster273-001 | 0.024 | NA | 8,828 |
| SALT00000085367_Cluster281-009 | 0.293 | NA | 3,736 |
| SALT00000029069_Cluster281-010 | 0.092 | NA | 2,455 |
| SALT00000000255_Cluster283-002 | 1.000 | NONATHT000919\| | 1,386 |
| SALT00000000324_Cluster283-003 | 1.000 | NONATHT000919\| | 1,352 |
| SALT00000012354_Cluster283-007 | 1.000 | NONATHT000919\| | 1,188 |
| SALT00000014323_Cluster283-009 | 1.000 | NONATHT000919\| | 1,266 |
| SALT00000022128_Cluster283-015 | 1.000 | NONATHT000919\| | 1,309 |
| SALT00000036250_Cluster283-017 | 1.000 | NONATHT000919\| | 1,337 |
| SALT00000048833_Cluster283-026 | 1.000 | NONATHT000919\| | 1,305 |
| SALT00000018372_Cluster283-036 | 1.000 | NONATHT000919\| | 1,581 |
| SALT00000002995_Cluster300-001 | 0.620 | lincRNA11_PMID_24635777\| | 1,866 |
| SALT00000005947_Cluster300-002 | 0.122 | lincRNA11_PMID_24635777\| | 1,499 |
| SALT00000008645_Cluster300-003 | 0.661 | lincRNA11_PMID_24635777\| | 1,614 |
| SALT00000019511_Cluster300-004 | 0.822 | pnrd_Mtr_chr4.trna1_SerGCT\|pnrd_Gma_chr15.trna | 2,378 |
| SALT00000021187_Cluster300-005 | 0.477 | lincRNA11_PMID_24635777\| | 2,686 |
| SALT00000023217_Cluster300-006 | 1.000 | pnrd_Gma_scaffold_1736.trna1_SerGCT\|pnrd_Osa_ | 2,679 |
| SALT00000027274_Cluster300-007 | 1.000 | lincRNA11_PMID_24635777\|pnrd_Osa_chr2.trna31_ | 2,795 |
| SALT00000027493_Cluster300-008 | 0.452 | lincRNA11_PMID_24635777\| | 2,814 |
| SALT00000028809_Cluster300-009 | 0.514 | lincRNA11_PMID_24635777\| | 2,478 |
| SALT00000047050_Cluster300-010 | 0.280 | lincRNA11_PMID_24635777\| | 1,577 |
| SALT00000052839_Cluster300-011 | 0.961 | pnrd_Osa_chr10.trna36_SerGGA\|pnrd_Osa_chr4.trna | 1,234 |
| SALT00000061398_Cluster300-012 | 0.594 | pnrd_Mtr_chr3.trna65_SerGGA\|pnrd_Mtr_chr8.trna1 | 1,899 |
| SALT00000068688_Cluster300-013 | 0.511 | lincRNA11_PMID_24635777\| | 2,481 |
| SALT00000073341_Cluster300-014 | 0.482 | lincRNA11_PMID_24635777\| | 2,660 |
| SALT00000079237_Cluster300-015 | 0.425 | lincRNA11_PMID_24635777\| | 2,969 |
| SALT00000084694_Cluster300-017 | 0.473 | lincRNA11_PMID_24635777\| | 2,711 |
| SALT00000093816_Cluster300-019 | 0.531 | lincRNA11_PMID_24635777\| | 2,379 |
| SALT00000067824_Cluster349-001 | 1.000 | lincRNA11_PMID_24635777\| | 8,433 |
| SALT00000085874_Cluster364-092 | 1.000 | lincRNA11_PMID_24635777\| | 8,369 |
| SALT00000026918_Cluster400-001 | 0.111 | NA | 2,655 |
| SALT00000030691_Cluster402-013 | 0.285 | NA | 3,548 |
| SALT00000088044_Cluster406-002 | 1.000 | lincRNA11_PMID_24635777\| | 4,853 |
| SALT00000090201_Cluster406-003 | 1.000 | lincRNA11_PMID_24635777\| | 5,143 |
| SALT00000092263_Cluster406-004 | 1.000 | lincRNA11_PMID_24635777\| | 4,018 |
| SALT00000093587_Cluster406-005 | 0.867 | lincRNA11_PMID_24635777\| | 3,016 |
| SALT00000023187_Cluster414-001 | 1.000 | TCONS_00084148\| | 3,124 |
| SALT00000082940_Cluster455-005 | 0.256 | NA | 4,454 |
| SALT00000012301_Cluster462-004 | 1.000 | lincRNA304_PMID_24635777\| | 1,742 |
| SALT00000019011_Cluster462-006 | 1.000 | lincRNA304_PMID_24635777\| | 1,602 |
| SALT00000041449_Cluster462-024 | 1.000 | lincRNA304_PMID_24635777\| | 1,753 |
| SALT00000055512_Cluster462-025 | 1.000 | lincRNA304_PMID_24635777\| | 1,698 |
| SALT00000059511_Cluster462-026 | 1.000 | lincRNA304_PMID_24635777\| | 1,713 |
| SALT00000022403_Cluster463-001 | 1.000 | TCONS_00047640\| | 2,698 |
| SALT00000029615_Cluster463-002 | 1.000 | TCONS_00047640\| | 4,086 |
| SALT00000031807_Cluster463-003 | 1.000 | TCONS_00047640\| | 4,046 |
| SALT00000032176_Cluster463-005 | 1.000 | TCONS_00047640\| | 3,682 |
| SALT00000035040_Cluster463-006 | 1.000 | TCONS_00047640\| | 4,178 |
| SALT00000035323_Cluster463-007 | 1.000 | TCONS_00047640\| | 4,145 |
| SALT00000067501_Cluster463-010 | 1.000 | TCONS_00047640\| | 2,911 |
| SALT00000081702_Cluster463-011 | 1.000 | TCONS_00047640\| | 3,635 |
| SALT00000083089_Cluster463-013 | 1.000 | TCONS_00047640\| | 3,220 |

| | | | |
|---|---|---|---|
| SALT00000084392_Cluster463-015 | 1.000 | TCONS_00047640\| | 4,070 |
| SALT00000085908_Cluster463-016 | 1.000 | TCONS_00047640\| | 3,801 |
| SALT00000086254_Cluster463-017 | 1.000 | TCONS_00047640\| | 4,056 |
| SALT00000087326_Cluster463-018 | 1.000 | TCONS_00047640\| | 4,128 |
| SALT00000088519_Cluster463-019 | 1.000 | TCONS_00047640\| | 3,679 |
| SALT00000088591_Cluster463-020 | 1.000 | TCONS_00047640\| | 3,984 |
| SALT00000089688_Cluster463-022 | 1.000 | TCONS_00047640\| | 3,542 |
| SALT00000090620_Cluster463-023 | 1.000 | TCONS_00047640\| | 3,895 |
| SALT00000093532_Cluster463-025 | 1.000 | TCONS_00047640\| | 4,752 |
| SALT00000093690_Cluster463-026 | 1.000 | TCONS_00047640\| | 4,021 |
| SALT00000090451_Cluster463-027 | 1.000 | TCONS_00047640\| | 4,218 |
| SALT00000092288_Cluster463-028 | 1.000 | TCONS_00047640\| | 4,806 |
| SALT00000032101_Cluster518-001 | 1.000 | TCONS_00084148\| | 4,484 |
| SALT00000083699_Cluster518-008 | 1.000 | TCONS_00084148\| | 4,591 |
| SALT00000087209_Cluster518-010 | 1.000 | TCONS_00084148\| | 4,575 |
| SALT00000087834_Cluster518-011 | 1.000 | TCONS_00084148\| | 4,622 |
| SALT00000088235_Cluster518-012 | 1.000 | TCONS_00084148\| | 4,693 |
| SALT00000088842_Cluster518-014 | 1.000 | TCONS_00084148\| | 4,448 |
| SALT00000091112_Cluster518-017 | 1.000 | TCONS_00084148\| | 4,616 |
| SALT00000091556_Cluster518-018 | 1.000 | TCONS_00084148\| | 4,625 |
| SALT00000092271_Cluster518-019 | 1.000 | TCONS_00084148\| | 4,585 |
| SALT00000089089_Cluster518-020 | 1.000 | TCONS_00084148\| | 4,741 |
| SALT00000092250_Cluster605-008 | 1.000 | cln-MIR166\|far-MIR166\|pta-MIR166c\| | 3,207 |
| SALT00000009927_Cluster629-002 | 1.000 | lincRNA302_PMID_24635777\|lincRNA246_PMID_ | 1,508 |
| SALT00000020416_Cluster629-003 | 1.000 | lincRNA302_PMID_24635777\|lincRNA246_PMID_ | 1,554 |
| SALT00000020507_Cluster629-004 | 1.000 | lincRNA246_PMID_24635777\|lincRNA302_PMID_ | 1,588 |
| SALT00000025180_Cluster629-005 | 1.000 | lincRNA246_PMID_24635777\|lincRNA302_PMID_ | 1,598 |
| SALT00000036024_Cluster629-006 | 1.000 | lincRNA302_PMID_24635777\|lincRNA246_PMID_ | 7,524 |
| SALT00000043373_Cluster629-007 | 1.000 | lincRNA246_PMID_24635777\|lincRNA302_PMID_ | 1,558 |
| SALT00000045058_Cluster629-008 | 1.000 | lincRNA302_PMID_24635777\|lincRNA246_PMID_ | 1,410 |
| SALT00000059487_Cluster629-010 | 1.000 | lincRNA302_PMID_24635777\|lincRNA246_PMID_ | 1,555 |
| SALT00000087253_Cluster642-001 | 1.000 | lincRNA11_PMID_24635777\| | 7,503 |
| SALT00000085638_Cluster687-004 | 1.000 | pnrd_Mtr_chr7.trna73_MetCAT\|pnrd_Osa_chr6.trna2 | 4,449 |
| SALT00000087550_Cluster687-006 | 1.000 | pnrd_Gma_scaffold_1114.trna1_MetCAT\|pnrd_Ara_ | 4,692 |
| SALT00000033186_Cluster687-007 | 1.000 | pnrd_Osa_chr10.trna12_MetCAT\|pnrd_Mtr_chr1.trna | 3,639 |
| SALT00000082139_Cluster687-008 | 1.000 | pnrd_Gma_scaffold_1114.trna1_MetCAT\|pnrd_Ara_ | 6,161 |
| SALT00000072921_Cluster693-001 | 0.222 | NA | 2,795 |
| SALT00000081260_Cluster693-002 | 0.287 | NA | 2,634 |
| SALT00000047253_Cluster698-001 | 0.252 | NA | 1,725 |
| SALT00000071662_Cluster721-001 | 0.196 | NA | 7,345 |
| SALT00000070328_Cluster775-004 | 1.000 | far-MIR166\| | 7,217 |
| SALT00000039124_Cluster801-001 | 0.964 | osa-MIR5538\| | 1,176 |
| SALT00000086594_Cluster801-002 | 0.876 | pnrd_Mtr_chr8.trna33_HisGTG\|pnrd_Gma_scaffold_ | 7,167 |
| SALT00000088270_Cluster801-003 | 1.000 | pnrd_Mtr_chr8.trna33_HisGTG\|pnrd_Gma_scaffold_ | 3,903 |
| SALT00000027588_Cluster801-005 | 0.248 | lincRNA676_PMID_24635777\| | 2,657 |
| SALT00000078649_Cluster810-001 | 1.000 | lincRNA11_PMID_24635777\| | 7,151 |
| SALT00000079063_Cluster857-008 | 1.000 | lincRNA623_PMID_24635777\|NONATHT003785\|li | 7,067 |
| SALT00000092861_Cluster908-001 | 0.127 | NA | 6,964 |
| SALT00000064369_Cluster913-001 | 0.205 | NA | 6,951 |
| SALT00000032661_Cluster916-004 | 0.932 | lincRNA688_PMID_24635777\|NONATHT003847\|N | 3,986 |
| SALT00000034362_Cluster916-005 | 0.626 | NONATHT003839\|lincRNA493_PMID_24635777\|li | 6,945 |
| SALT00000000030_Cluster992-001 | 0.267 | NONATHT003785\|lincRNA493_PMID_24635777\| | 1,591 |
| SALT00000000861_Cluster992-002 | 0.286 | NONATHT003785\|lincRNA493_PMID_24635777\| | 1,453 |
| SALT00000002184_Cluster992-003 | 0.990 | NONATHT003785\|lincRNA493_PMID_24635777\| | 2,232 |
| SALT00000005385_Cluster992-004 | 0.931 | lincRNA493_PMID_24635777\|NONATHT003785\| | 1,739 |
| SALT00000006469_Cluster992-005 | 0.326 | lincRNA493_PMID_24635777\|NONATHT003785\| | 1,188 |
| SALT00000009301_Cluster992-006 | 0.932 | lincRNA12_PMID_24635777\|NONATHT003785\|lin | 1,720 |
| SALT00000009607_Cluster992-007 | 0.909 | NONATHT003785\|lincRNA493_PMID_24635777\| | 2,169 |

| | | | |
|---|---|---|---|
| SALT00000010034_Cluster992-008 | 0.064 | NONATHT003785\|lincRNA12_PMID_24635777\|lin | 1,170 |
| SALT00000012097_Cluster992-009 | 0.895 | lincRNA493_PMID_24635777\|lincRNA623_PMID_ | 2,390 |
| SALT00000012459_Cluster992-010 | 0.186 | NONATHT003785\|lincRNA493_PMID_24635777\| | 1,657 |
| SALT00000012849_Cluster992-011 | 0.921 | lincRNA493_PMID_24635777\|lincRNA12_PMID_2 | 1,965 |
| SALT00000012979_Cluster992-012 | 0.996 | NONATHT003785\|lincRNA493_PMID_24635777\| | 2,177 |
| SALT00000013988_Cluster992-013 | 0.931 | NONATHT003785\|lincRNA493_PMID_24635777\| | 1,752 |
| SALT00000016579_Cluster992-014 | 0.995 | lincRNA493_PMID_24635777\|NONATHT003785\| | 1,222 |
| SALT00000017752_Cluster992-015 | 0.907 | lincRNA623_PMID_24635777\|lincRNA12_PMID_2 | 2,199 |
| SALT00000018469_Cluster992-016 | 0.913 | NONATHT003785\|lincRNA493_PMID_24635777\| | 2,104 |
| SALT00000018502_Cluster992-017 | 0.955 | lincRNA493_PMID_24635777\|NONATHT003785\| | 1,112 |
| SALT00000020419_Cluster992-018 | 0.919 | NONATHT003785\|lincRNA493_PMID_24635777\| | 2,000 |
| SALT00000020502_Cluster992-019 | 0.931 | lincRNA493_PMID_24635777\|lincRNA623_PMID_ | 1,748 |
| SALT00000020747_Cluster992-020 | 0.833 | lincRNA623_PMID_24635777\|NONATHT003785\|li | 3,153 |
| SALT00000020924_Cluster992-021 | 0.821 | lincRNA493_PMID_24635777\|lincRNA623_PMID_ | 3,267 |
| SALT00000021135_Cluster992-022 | 0.844 | lincRNA493_PMID_24635777\|lincRNA623_PMID_ | 3,042 |
| SALT00000021794_Cluster992-023 | 0.886 | lincRNA493_PMID_24635777\|lincRNA623_PMID_ | 2,527 |
| SALT00000022464_Cluster992-024 | 0.946 | lincRNA623_PMID_24635777\|NONATHT003785\|li | 2,687 |
| SALT00000024192_Cluster992-025 | 0.942 | lincRNA493_PMID_24635777\|NONATHT003785\| | 1,485 |
| SALT00000025114_Cluster992-026 | 0.911 | lincRNA493_PMID_24635777\|lincRNA12_PMID_2 | 2,135 |
| SALT00000025138_Cluster992-027 | 0.924 | lincRNA493_PMID_24635777\|NONATHT003785\| | 1,887 |
| SALT00000025143_Cluster992-028 | 0.905 | NONATHT003785\|lincRNA493_PMID_24635777\| | 2,238 |
| SALT00000025177_Cluster992-029 | 0.930 | NONATHT003785\|lincRNA493_PMID_24635777\| | 1,775 |
| SALT00000025228_Cluster992-030 | 0.934 | lincRNA493_PMID_24635777\|NONATHT003785\| | 1,679 |
| SALT00000025597_Cluster992-031 | 0.852 | NONATHT003785\|lincRNA12_PMID_24635777\|lin | 2,956 |
| SALT00000026105_Cluster992-032 | 0.847 | lincRNA493_PMID_24635777\|NONATHT003785\|li | 3,007 |
| SALT00000027055_Cluster992-033 | 0.845 | lincRNA12_PMID_24635777\|NONATHT003785\|lin | 3,028 |
| SALT00000027059_Cluster992-034 | 0.795 | lincRNA12_PMID_24635777\|NONATHT003785\|lin | 3,505 |
| SALT00000027242_Cluster992-035 | 0.913 | lincRNA493_PMID_24635777\|NONATHT003785\| | 2,106 |
| SALT00000028233_Cluster992-036 | 0.048 | lincRNA623_PMID_24635777\|lincRNA12_PMID_2 | 1,608 |
| SALT00000028368_Cluster992-037 | 0.048 | lincRNA12_PMID_24635777\|NONATHT003785\|lin | 1,614 |
| SALT00000028424_Cluster992-038 | 0.886 | NONATHT003785\|lincRNA12_PMID_24635777\|lin | 2,532 |
| SALT00000028427_Cluster992-039 | 0.050 | lincRNA12_PMID_24635777\|NONATHT003785\|lin | 1,534 |
| SALT00000028645_Cluster992-040 | 0.930 | lincRNA493_PMID_24635777\|NONATHT003785\| | 1,764 |
| SALT00000028716_Cluster992-041 | 0.796 | lincRNA493_PMID_24635777\|lincRNA623_PMID_ | 3,502 |
| SALT00000028851_Cluster992-042 | 0.917 | lincRNA493_PMID_24635777\|lincRNA623_PMID_ | 2,029 |
| SALT00000031590_Cluster992-043 | 0.054 | lincRNA493_PMID_24635777\|lincRNA12_PMID_2 | 1,420 |
| SALT00000031821_Cluster992-044 | 0.769 | lincRNA623_PMID_24635777\|lincRNA12_PMID_2 | 3,719 |
| SALT00000032471_Cluster992-045 | 0.741 | NONATHT003785\|lincRNA12_PMID_24635777\|lin | 3,935 |
| SALT00000033533_Cluster992-046 | 0.762 | lincRNA623_PMID_24635777\|lincRNA12_PMID_2 | 2,597 |
| SALT00000034037_Cluster992-047 | 0.796 | NONATHT003785\|lincRNA12_PMID_24635777\|lin | 3,500 |
| SALT00000034658_Cluster992-048 | 0.812 | NONATHT003785\|lincRNA12_PMID_24635777\|lin | 3,358 |
| SALT00000034753_Cluster992-049 | 0.922 | lincRNA623_PMID_24635777\|lincRNA12_PMID_2 | 1,945 |
| SALT00000035225_Cluster992-050 | 0.860 | lincRNA493_PMID_24635777\|NONATHT003785\|li | 2,854 |
| SALT00000035296_Cluster992-051 | 0.869 | lincRNA493_PMID_24635777\|lincRNA623_PMID_ | 2,752 |
| SALT00000035522_Cluster992-052 | 0.937 | lincRNA493_PMID_24635777\|NONATHT003785\| | 1,622 |
| SALT00000035813_Cluster992-053 | 0.235 | NONATHT003785\|lincRNA493_PMID_24635777\| | 1,582 |
| SALT00000037507_Cluster992-054 | 0.905 | lincRNA493_PMID_24635777\|NONATHT003785\| | 2,240 |
| SALT00000038215_Cluster992-055 | 0.192 | lincRNA493_PMID_24635777\|NONATHT003785\| | 1,596 |
| SALT00000038622_Cluster992-056 | 0.256 | NONATHT003785\|lincRNA493_PMID_24635777\| | 1,669 |
| SALT00000039084_Cluster992-057 | 0.045 | lincRNA12_PMID_24635777\|NONATHT003785\|lin | 1,710 |
| SALT00000040292_Cluster992-058 | 0.915 | lincRNA493_PMID_24635777\|NONATHT003785\| | 2,077 |
| SALT00000040970_Cluster992-059 | 0.303 | NONATHT003785\|lincRNA493_PMID_24635777\| | 1,338 |
| SALT00000042414_Cluster992-060 | 0.851 | lincRNA493_PMID_24635777\|NONATHT003785\| | 1,778 |
| SALT00000042586_Cluster992-061 | 0.905 | NONATHT003785\|lincRNA493_PMID_24635777\| | 2,237 |
| SALT00000043072_Cluster992-062 | 0.273 | NONATHT003785\|lincRNA493_PMID_24635777\| | 1,548 |
| SALT00000043257_Cluster992-063 | 0.185 | lincRNA493_PMID_24635777\|NONATHT003785\| | 1,667 |
| SALT00000046379_Cluster992-064 | 0.929 | lincRNA493_PMID_24635777\|lincRNA12_PMID_2 | 1,790 |
| SALT00000046977_Cluster992-065 | 0.896 | lincRNA493_PMID_24635777\|NONATHT003785\| | 2,381 |

| | | | |
|---|---|---|---|
| SALT00000047237_Cluster992-066 | 0.191 | lincRNA493_PMID_24635777\|NONATHT003785\| | 1,611 |
| SALT00000051659_Cluster992-068 | 0.051 | lincRNA623_PMID_24635777\|lincRNA12_PMID_2 | 1,515 |
| SALT00000053221_Cluster992-069 | 0.864 | NONATHT003785\|lincRNA493_PMID_24635777\| | 1,471 |
| SALT00000053308_Cluster992-070 | 0.263 | NONATHT003785\|lincRNA493_PMID_24635777\| | 1,622 |
| SALT00000053353_Cluster992-071 | 0.934 | NONATHT003785\|lincRNA493_PMID_24635777\| | 1,723 |
| SALT00000054585_Cluster992-072 | 0.245 | NONATHT003785\|lincRNA493_PMID_24635777\| | 1,159 |
| SALT00000057094_Cluster992-074 | 0.287 | lincRNA493_PMID_24635777\|NONATHT003785\| | 1,451 |
| SALT00000057116_Cluster992-075 | 0.999 | lincRNA493_PMID_24635777\|NONATHT003785\| | 1,200 |
| SALT00000057143_Cluster992-076 | 0.048 | lincRNA493_PMID_24635777\|lincRNA623_PMID_ | 1,620 |
| SALT00000057153_Cluster992-077 | 0.270 | NONATHT003785\|lincRNA493_PMID_24635777\| | 1,570 |
| SALT00000058269_Cluster992-078 | 0.910 | lincRNA493_PMID_24635777\|NONATHT003785\| | 2,160 |
| SALT00000059380_Cluster992-079 | 0.051 | lincRNA623_PMID_24635777\|NONATHT003785\|N | 1,531 |
| SALT00000060530_Cluster992-080 | 0.053 | lincRNA493_PMID_24635777\|lincRNA623_PMID_ | 1,474 |
| SALT00000060624_Cluster992-081 | 0.927 | lincRNA493_PMID_24635777\|NONATHT003785\| | 1,835 |
| SALT00000061349_Cluster992-082 | 0.230 | lincRNA493_PMID_24635777\|lincRNA623_PMID_ | 1,685 |
| SALT00000061566_Cluster992-083 | 0.048 | lincRNA623_PMID_24635777\|lincRNA12_PMID_2 | 1,615 |
| SALT00000061952_Cluster992-084 | 1.000 | lincRNA493_PMID_24635777\|lincRNA623_PMID_ | 2,975 |
| SALT00000069123_Cluster992-085 | 0.993 | lincRNA493_PMID_24635777\|lincRNA623_PMID_ | 2,870 |
| SALT00000071855_Cluster992-086 | 0.875 | lincRNA493_PMID_24635777\|NONATHT003839\|li | 2,675 |
| SALT00000071982_Cluster992-087 | 0.829 | lincRNA623_PMID_24635777\|lincRNA12_PMID_2 | 3,189 |
| SALT00000073108_Cluster992-088 | 0.887 | NONATHT003785\|lincRNA493_PMID_24635777\| | 2,511 |
| SALT00000075943_Cluster992-089 | 0.881 | lincRNA493_PMID_24635777\|lincRNA623_PMID_ | 2,597 |
| SALT00000076243_Cluster992-090 | 0.890 | NONATHT003785\|lincRNA12_PMID_24635777\|lin | 2,472 |
| SALT00000076246_Cluster992-091 | 0.823 | lincRNA493_PMID_24635777\|lincRNA623_PMID_ | 3,252 |
| SALT00000078955_Cluster992-093 | 0.217 | NONATHT003785\|lincRNA493_PMID_24635777\| | 1,377 |
| SALT00000079631_Cluster992-094 | 0.860 | lincRNA493_PMID_24635777\|NONATHT003839\|li | 2,862 |
| SALT00000081087_Cluster992-096 | 0.904 | lincRNA12_PMID_24635777\|NONATHT003785\|lin | 2,258 |
| SALT00000081308_Cluster992-097 | 0.771 | lincRNA493_PMID_24635777\|lincRNA623_PMID_ | 2,803 |
| SALT00000082270_Cluster992-098 | 0.795 | lincRNA493_PMID_24635777\|NONATHT003839\|li | 3,507 |
| SALT00000082393_Cluster992-099 | 0.840 | lincRNA623_PMID_24635777\|lincRNA12_PMID_2 | 3,082 |
| SALT00000082765_Cluster992-100 | 0.795 | lincRNA12_PMID_24635777\|NONATHT003785\|lin | 3,506 |
| SALT00000083505_Cluster992-102 | 0.854 | lincRNA623_PMID_24635777\|NONATHT003785\|li | 2,927 |
| SALT00000085382_Cluster992-104 | 0.818 | NONATHT003847\|lincRNA12_PMID_24635777\|NC | 3,304 |
| SALT00000085915_Cluster992-107 | 0.795 | lincRNA12_PMID_24635777\|NONATHT003785\|lin | 3,505 |
| SALT00000085998_Cluster992-108 | 0.795 | TCONS_00018605\|NONATHT003839\|lincRNA493_ | 3,506 |
| SALT00000086974_Cluster992-109 | 0.769 | lincRNA12_PMID_24635777\|NONATHT003785\|lin | 3,720 |
| SALT00000087131_Cluster992-110 | 0.791 | lincRNA12_PMID_24635777\|NONATHT003785\|lin | 3,538 |
| SALT00000087621_Cluster992-111 | 0.795 | lincRNA12_PMID_24635777\|NONATHT003785\|lin | 3,505 |
| SALT00000087676_Cluster992-112 | 0.915 | NONATHT003785\|lincRNA12_PMID_24635777\|lin | 3,676 |
| SALT00000087801_Cluster992-113 | 0.791 | lincRNA623_PMID_24635777\|lincRNA12_PMID_2 | 3,538 |
| SALT00000088061_Cluster992-114 | 0.793 | lincRNA12_PMID_24635777\|NONATHT003847\|NC | 3,522 |
| SALT00000088066_Cluster992-115 | 0.796 | lincRNA493_PMID_24635777\|NONATHT003785\|li | 3,502 |
| SALT00000088307_Cluster992-116 | 0.745 | lincRNA12_PMID_24635777\|NONATHT003785\|lin | 3,908 |
| SALT00000088360_Cluster992-117 | 0.796 | lincRNA623_PMID_24635777\|lincRNA12_PMID_2 | 3,501 |
| SALT00000089647_Cluster992-118 | 0.781 | lincRNA623_PMID_24635777\|NONATHT003785\|N | 3,626 |
| SALT00000089960_Cluster992-120 | 0.741 | NONATHT003785\|lincRNA12_PMID_24635777\|lin | 3,933 |
| SALT00000090198_Cluster992-121 | 0.820 | lincRNA493_PMID_24635777\|NONATHT003785\| | 2,093 |
| SALT00000091405_Cluster992-122 | 0.796 | lincRNA493_PMID_24635777\|NONATHT003785\|li | 3,502 |
| SALT00000091742_Cluster992-123 | 0.999 | NONATHT003839\|lincRNA493_PMID_24635777\|li | 3,500 |
| SALT00000092024_Cluster992-124 | 0.967 | NONATHT003847\|NONATHT003785\|lincRNA12_P | 3,507 |
| SALT00000093271_Cluster992-125 | 0.796 | lincRNA12_PMID_24635777\|NONATHT003847\|NC | 3,503 |
| SALT00000093375_Cluster992-126 | 0.796 | NONATHT003785\|lincRNA12_PMID_24635777\|lin | 3,500 |
| SALT00000078698_Cluster992-127 | 1.000 | lincRNA493_PMID_24635777\|lincRNA623_PMID_ | 3,095 |
| SALT00000086479_Cluster992-128 | 1.000 | lincRNA493_PMID_24635777\|NONATHT003785\|li | 3,956 |
| SALT00000088826_Cluster992-129 | 0.632 | lincRNA493_PMID_24635777\|lincRNA12_PMID_2 | 3,474 |
| SALT00000088006_Cluster992-131 | 0.178 | lincRNA493_PMID_24635777\|lincRNA623_PMID_ | 3,607 |
| SALT00000029878_Cluster1000-001 | 1.000 | hvu-MIR6182\| | 3,719 |
| SALT00000030497_Cluster1000-002 | 1.000 | hvu-MIR6182\| | 3,766 |

| | | | |
|---|---|---|---|
| SALT00000031874_Cluster1000-003 | 1.000 | hvu-MIR6182| | 3,829 |
| SALT00000033565_Cluster1000-004 | 1.000 | hvu-MIR6182| | 3,499 |
| SALT00000036469_Cluster1000-005 | 1.000 | hvu-MIR6182| | 1,621 |
| SALT00000064589_Cluster1000-006 | 1.000 | hvu-MIR6182| | 2,806 |
| SALT00000066190_Cluster1000-007 | 1.000 | hvu-MIR6182| | 2,576 |
| SALT00000075182_Cluster1000-008 | 1.000 | hvu-MIR6182| | 3,357 |
| SALT00000079532_Cluster1000-009 | 1.000 | hvu-MIR6182| | 2,692 |
| SALT00000086973_Cluster1000-010 | 1.000 | hvu-MIR6182| | 3,554 |
| SALT00000087773_Cluster1000-011 | 1.000 | hvu-MIR6182| | 3,932 |
| SALT00000089129_Cluster1000-013 | 1.000 | hvu-MIR6182| | 3,827 |
| SALT00000093772_Cluster1000-014 | 1.000 | hvu-MIR6182| | 3,796 |
| SALT00000067379_Cluster1028-042 | 1.000 | lincRNA11_PMID_24635777| | 3,868 |
| SALT00000042273_Cluster1037-001 | 1.000 | lincRNA688_PMID_24635777| | 6,701 |
| SALT00000042882_Cluster1074-001 | 1.000 | lincRNA376_PMID_24635777| | 2,030 |
| SALT00000048726_Cluster1074-002 | 0.971 | lincRNA376_PMID_24635777| | 1,865 |
| SALT00000062607_Cluster1074-003 | 1.000 | lincRNA376_PMID_24635777| | 2,667 |
| SALT00000073067_Cluster1074-004 | 1.000 | lincRNA376_PMID_24635777| | 2,724 |
| SALT00000073892_Cluster1074-005 | 1.000 | lincRNA376_PMID_24635777| | 3,037 |
| SALT00000076715_Cluster1074-006 | 1.000 | lincRNA376_PMID_24635777| | 2,656 |
| SALT00000080266_Cluster1074-007 | 1.000 | lincRNA376_PMID_24635777| | 3,594 |
| SALT00000083358_Cluster1074-008 | 1.000 | lincRNA376_PMID_24635777| | 3,148 |
| SALT00000083794_Cluster1074-009 | 1.000 | lincRNA376_PMID_24635777| | 2,837 |
| SALT00000084562_Cluster1074-010 | 1.000 | lincRNA376_PMID_24635777| | 3,583 |
| SALT00000085390_Cluster1074-011 | 1.000 | lincRNA376_PMID_24635777| | 3,516 |
| SALT00000092537_Cluster1074-012 | 1.000 | lincRNA376_PMID_24635777| | 3,557 |
| SALT00000092850_Cluster1074-013 | 1.000 | lincRNA376_PMID_24635777| | 2,780 |
| SALT00000093500_Cluster1074-015 | 1.000 | lincRNA376_PMID_24635777| | 3,532 |
| SALT00000089749_Cluster1097-016 | 1.000 | sit-MIR122-1-npr|TCONS_00018605|ghr-MIR5368* | 6,526 |
| SALT00000087973_Cluster1106-001 | 1.000 | lincRNA11_PMID_24635777| | 6,495 |
| SALT00000093307_Cluster1130-003 | 1.000 | lincRNA11_PMID_24635777| | 6,437 |
| SALT00000064705_Cluster1133-009 | 1.000 | lincRNA11_PMID_24635777| | 6,423 |
| SALT00000027906_Cluster1139-001 | 1.000 | lincRNA376_PMID_24635777| | 3,168 |
| SALT00000029622_Cluster1139-002 | 1.000 | lincRNA376_PMID_24635777| | 3,584 |
| SALT00000030489_Cluster1139-003 | 1.000 | lincRNA376_PMID_24635777| | 3,635 |
| SALT00000034718_Cluster1139-004 | 1.000 | lincRNA376_PMID_24635777| | 3,638 |
| SALT00000035258_Cluster1139-005 | 1.000 | lincRNA376_PMID_24635777| | 3,537 |
| SALT00000035510_Cluster1139-006 | 1.000 | lincRNA376_PMID_24635777| | 3,647 |
| SALT00000035627_Cluster1139-007 | 1.000 | lincRNA376_PMID_24635777| | 3,447 |
| SALT00000076900_Cluster1139-008 | 1.000 | lincRNA376_PMID_24635777| | 3,598 |
| SALT00000079057_Cluster1139-009 | 1.000 | lincRNA376_PMID_24635777| | 3,733 |
| SALT00000084811_Cluster1139-010 | 1.000 | lincRNA376_PMID_24635777| | 3,504 |
| SALT00000086249_Cluster1139-011 | 1.000 | lincRNA376_PMID_24635777| | 3,628 |
| SALT00000091410_Cluster1139-013 | 1.000 | lincRNA376_PMID_24635777| | 3,563 |
| SALT00000093593_Cluster1139-014 | 1.000 | lincRNA376_PMID_24635777| | 3,619 |
| SALT00000091274_Cluster1139-015 | 1.000 | lincRNA376_PMID_24635777| | 3,590 |
| SALT00000092418_Cluster1139-016 | 1.000 | lincRNA376_PMID_24635777| | 3,621 |
| SALT00000093019_Cluster1139-017 | 1.000 | lincRNA376_PMID_24635777| | 4,708 |
| SALT00000093493_Cluster1149-001 | 1.000 | lincRNA11_PMID_24635777| | 6,380 |
| SALT00000062412_Cluster1193-002 | 0.807 | sit-MIR122-1-npr|lincRNA372_PMID_24635777|TC | 4,344 |
| SALT00000002546_Cluster1243-001 | 1.000 | osa-MIR5079b|osa-MIR5079a| | 2,172 |
| SALT00000007346_Cluster1243-002 | 1.000 | osa-MIR5079a|osa-MIR5079b| | 1,549 |
| SALT00000011388_Cluster1243-003 | 1.000 | osa-MIR5079b|osa-MIR5079a| | 1,675 |
| SALT00000014659_Cluster1243-004 | 1.000 | osa-MIR5079b|osa-MIR5079a| | 2,043 |
| SALT00000022309_Cluster1243-005 | 1.000 | osa-MIR5079a|osa-MIR5079b| | 2,564 |
| SALT00000025518_Cluster1243-006 | 1.000 | osa-MIR5079a|osa-MIR5079b| | 2,901 |
| SALT00000033948_Cluster1243-007 | 1.000 | osa-MIR5079b|osa-MIR5079a| | 3,457 |
| SALT00000034218_Cluster1243-008 | 1.000 | osa-MIR5079b|osa-MIR5079a| | 3,828 |
| SALT00000034407_Cluster1243-009 | 1.000 | osa-MIR5079a|osa-MIR5079b| | 3,693 |

| | | | |
|---|---|---|---|
| SALT00000053547_Cluster1243-011 | 1.000 | osa-MIR5079b\|osa-MIR5079a\| | 1,877 |
| SALT00000064939_Cluster1243-013 | 1.000 | osa-MIR5079a\|osa-MIR5079b\| | 2,345 |
| SALT00000071545_Cluster1243-014 | 1.000 | osa-MIR5079a\|osa-MIR5079b\| | 2,329 |
| SALT00000079051_Cluster1243-016 | 1.000 | osa-MIR5079b\|osa-MIR5079a\| | 2,683 |
| SALT00000081148_Cluster1243-017 | 1.000 | osa-MIR5079a\|osa-MIR5079b\| | 4,032 |
| SALT00000081738_Cluster1243-018 | 1.000 | osa-MIR5079b\|osa-MIR5079a\| | 3,336 |
| SALT00000086401_Cluster1243-019 | 1.000 | osa-MIR5079b\|osa-MIR5079a\| | 3,325 |
| SALT00000088121_Cluster1243-020 | 1.000 | osa-MIR5079a\|osa-MIR5079b\| | 3,495 |
| SALT00000089404_Cluster1243-021 | 1.000 | osa-MIR5079b\|osa-MIR5079a\| | 6,125 |
| SALT00000090428_Cluster1243-023 | 1.000 | osa-MIR5079b\|osa-MIR5079a\| | 5,388 |
| SALT00000072681_Cluster1244-001 | 0.061 | NA | 6,124 |
| SALT00000088107_Cluster1249-016 | 1.000 | NONATHT003839\|NONATHT003847\| | 3,741 |
| SALT00000008931_Cluster1256-001 | 0.998 | lincRNA623_PMID_24635777\|NONATHT003785\|li | 1,982 |
| SALT00000027861_Cluster1256-002 | 0.847 | lincRNA493_PMID_24635777\|lincRNA12_PMID_2 | 3,008 |
| SALT00000034535_Cluster1256-003 | 0.992 | lincRNA493_PMID_24635777\|lincRNA623_PMID_ | 3,821 |
| SALT00000034665_Cluster1256-004 | 0.990 | lincRNA493_PMID_24635777\|lincRNA12_PMID_2 | 4,054 |
| SALT00000039070_Cluster1256-005 | 0.841 | NONATHT003785\|lincRNA12_PMID_24635777\|lin | 1,563 |
| SALT00000077011_Cluster1256-007 | 0.967 | lincRNA493_PMID_24635777\|lincRNA623_PMID_ | 3,969 |
| SALT00000077980_Cluster1256-008 | 0.991 | lincRNA493_PMID_24635777\|lincRNA12_PMID_2 | 4,015 |
| SALT00000078975_Cluster1256-009 | 0.992 | lincRNA623_PMID_24635777\|lincRNA12_PMID_2 | 3,697 |
| SALT00000079479_Cluster1256-010 | 0.997 | NONATHT003785\|lincRNA12_PMID_24635777\|lin | 2,548 |
| SALT00000081069_Cluster1256-011 | 0.637 | lincRNA493_PMID_24635777\|lincRNA623_PMID_ | 4,625 |
| SALT00000084862_Cluster1256-012 | 0.994 | lincRNA493_PMID_24635777\|lincRNA623_PMID_ | 3,420 |
| SALT00000085089_Cluster1256-013 | 0.993 | lincRNA623_PMID_24635777\|lincRNA12_PMID_2 | 3,670 |
| SALT00000086517_Cluster1256-014 | 1.000 | NONATHT003785\|lincRNA12_PMID_24635777\|lin | 6,099 |
| SALT00000089889_Cluster1256-015 | 0.610 | lincRNA12_PMID_24635777\|NONATHT003785\|lin | 4,787 |
| SALT00000090463_Cluster1256-016 | 0.582 | lincRNA493_PMID_24635777\|lincRNA623_PMID_ | 4,949 |
| SALT00000032575_Cluster1285-001 | 1.000 | hvu-MIR6182\| | 3,566 |
| SALT00000033685_Cluster1285-002 | 1.000 | hvu-MIR6182\| | 3,833 |
| SALT00000045417_Cluster1285-003 | 1.000 | hvu-MIR6182\| | 1,670 |
| SALT00000056251_Cluster1285-004 | 1.000 | hvu-MIR6182\| | 1,458 |
| SALT00000067359_Cluster1285-005 | 1.000 | hvu-MIR6182\| | 3,016 |
| SALT00000081867_Cluster1285-006 | 1.000 | hvu-MIR6182\| | 2,817 |
| SALT00000082196_Cluster1285-007 | 1.000 | hvu-MIR6182\| | 2,655 |
| SALT00000042470_Cluster1310-001 | 0.251 | NA | 5,953 |
| SALT00000006539_Cluster1314-001 | 1.000 | lincRNA304_PMID_24635777\| | 2,002 |
| SALT00000008773_Cluster1314-002 | 1.000 | lincRNA304_PMID_24635777\| | 2,204 |
| SALT00000020567_Cluster1314-003 | 1.000 | lincRNA304_PMID_24635777\| | 2,893 |
| SALT00000020595_Cluster1314-004 | 1.000 | lincRNA304_PMID_24635777\| | 2,894 |
| SALT00000020835_Cluster1314-005 | 1.000 | lincRNA304_PMID_24635777\| | 2,628 |
| SALT00000022784_Cluster1314-006 | 1.000 | lincRNA304_PMID_24635777\| | 2,498 |
| SALT00000022808_Cluster1314-007 | 1.000 | lincRNA304_PMID_24635777\| | 2,575 |
| SALT00000023889_Cluster1314-008 | 1.000 | lincRNA304_PMID_24635777\| | 2,772 |
| SALT00000023972_Cluster1314-009 | 1.000 | lincRNA304_PMID_24635777\| | 2,866 |
| SALT00000024764_Cluster1314-010 | 1.000 | lincRNA304_PMID_24635777\| | 2,784 |
| SALT00000025149_Cluster1314-011 | 1.000 | lincRNA304_PMID_24635777\| | 2,825 |
| SALT00000025807_Cluster1314-012 | 1.000 | lincRNA304_PMID_24635777\| | 2,602 |
| SALT00000026540_Cluster1314-013 | 1.000 | lincRNA304_PMID_24635777\| | 2,831 |
| SALT00000026981_Cluster1314-014 | 1.000 | lincRNA304_PMID_24635777\| | 2,606 |
| SALT00000027010_Cluster1314-015 | 1.000 | lincRNA304_PMID_24635777\| | 2,505 |
| SALT00000027137_Cluster1314-016 | 1.000 | lincRNA304_PMID_24635777\| | 2,897 |
| SALT00000029191_Cluster1314-017 | 1.000 | lincRNA304_PMID_24635777\| | 2,795 |
| SALT00000032446_Cluster1314-018 | 1.000 | lincRNA304_PMID_24635777\| | 3,720 |
| SALT00000044096_Cluster1314-019 | 1.000 | lincRNA304_PMID_24635777\| | 1,762 |
| SALT00000051992_Cluster1314-020 | 1.000 | lincRNA304_PMID_24635777\| | 2,062 |
| SALT00000061948_Cluster1314-021 | 1.000 | lincRNA304_PMID_24635777\| | 2,844 |
| SALT00000062373_Cluster1314-022 | 1.000 | lincRNA304_PMID_24635777\| | 2,873 |
| SALT00000062519_Cluster1314-023 | 1.000 | lincRNA304_PMID_24635777\| | 2,678 |

| | | | |
|---|---|---|---|
| SALT00000064268_Cluster1314-024 | 1.000 | lincRNA304_PMID_24635777\| | 2,958 |
| SALT00000066435_Cluster1314-025 | 1.000 | lincRNA304_PMID_24635777\| | 2,811 |
| SALT00000068308_Cluster1314-026 | 1.000 | lincRNA304_PMID_24635777\| | 2,785 |
| SALT00000071152_Cluster1314-027 | 1.000 | lincRNA304_PMID_24635777\| | 2,892 |
| SALT00000071367_Cluster1314-028 | 1.000 | lincRNA304_PMID_24635777\| | 2,880 |
| SALT00000071392_Cluster1314-029 | 1.000 | lincRNA304_PMID_24635777\| | 2,774 |
| SALT00000071629_Cluster1314-030 | 1.000 | lincRNA304_PMID_24635777\| | 2,822 |
| SALT00000071967_Cluster1314-031 | 1.000 | lincRNA304_PMID_24635777\| | 2,740 |
| SALT00000072206_Cluster1314-032 | 1.000 | lincRNA304_PMID_24635777\| | 2,872 |
| SALT00000072658_Cluster1314-033 | 1.000 | lincRNA304_PMID_24635777\| | 2,855 |
| SALT00000073637_Cluster1314-034 | 1.000 | lincRNA304_PMID_24635777\| | 2,846 |
| SALT00000074593_Cluster1314-035 | 1.000 | lincRNA304_PMID_24635777\| | 2,919 |
| SALT00000074991_Cluster1314-036 | 1.000 | lincRNA304_PMID_24635777\| | 2,868 |
| SALT00000075990_Cluster1314-037 | 1.000 | lincRNA304_PMID_24635777\| | 3,146 |
| SALT00000076650_Cluster1314-038 | 1.000 | lincRNA304_PMID_24635777\| | 2,867 |
| SALT00000077796_Cluster1314-039 | 1.000 | lincRNA304_PMID_24635777\| | 2,863 |
| SALT00000080622_Cluster1314-040 | 1.000 | lincRNA304_PMID_24635777\| | 2,956 |
| SALT00000082893_Cluster1314-041 | 1.000 | lincRNA304_PMID_24635777\| | 2,712 |
| SALT00000083299_Cluster1314-042 | 1.000 | lincRNA304_PMID_24635777\| | 2,836 |
| SALT00000083362_Cluster1314-043 | 1.000 | lincRNA304_PMID_24635777\| | 2,932 |
| SALT00000083684_Cluster1314-044 | 1.000 | lincRNA304_PMID_24635777\| | 2,829 |
| SALT00000084386_Cluster1314-045 | 1.000 | lincRNA304_PMID_24635777\| | 2,830 |
| SALT00000070614_Cluster1314-047 | 1.000 | lincRNA304_PMID_24635777\| | 2,862 |
| SALT00000087036_Cluster1314-050 | 1.000 | lincRNA304_PMID_24635777\| | 5,221 |
| SALT00000070872_Cluster1314-051 | 1.000 | lincRNA304_PMID_24635777\| | 2,961 |
| SALT00000079408_Cluster1320-001 | 1.000 | lincRNA304_PMID_24635777\| | 5,935 |
| SALT00000085385_Cluster1338-001 | 1.000 | lincRNA376_PMID_24635777\| | 5,887 |
| SALT00000089009_Cluster1339-001 | 1.000 | lincRNA372_PMID_24635777\| | 5,887 |
| SALT00000077889_Cluster1340-009 | 1.000 | pta-MIR166c\|cln-MIR166\|far-MIR166\| | 5,886 |
| SALT00000086051_Cluster1346-001 | 1.000 | lincRNA11_PMID_24635777\| | 4,749 |
| SALT00000088776_Cluster1346-002 | 1.000 | lincRNA11_PMID_24635777\| | 5,881 |
| SALT00000089568_Cluster1346-003 | 1.000 | lincRNA11_PMID_24635777\| | 5,693 |
| SALT00000090682_Cluster1346-004 | 1.000 | lincRNA11_PMID_24635777\| | 4,759 |
| SALT00000085169_Cluster1358-002 | 0.952 | lincRNA686_PMID_24635777\|lincRNA616_PMID_ | 5,853 |
| SALT00000087379_Cluster1365-010 | 1.000 | lincRNA372_PMID_24635777\| | 3,679 |
| SALT00000005717_Cluster1376-001 | 1.000 | ace-MIR170c\|bdi-MIR171a\| | 1,731 |
| SALT00000021836_Cluster1376-002 | 1.000 | ace-MIR170c\| | 2,599 |
| SALT00000022014_Cluster1376-003 | 1.000 | ace-MIR170c\| | 2,728 |
| SALT00000022150_Cluster1376-004 | 1.000 | bdi-MIR171a\| | 2,650 |
| SALT00000022402_Cluster1376-005 | 1.000 | bdi-MIR171a\| | 2,742 |
| SALT00000022752_Cluster1376-006 | 1.000 | ace-MIR170c\|bdi-MIR171a\| | 2,655 |
| SALT00000025885_Cluster1376-007 | 1.000 | bdi-MIR171a\| | 2,566 |
| SALT00000028995_Cluster1376-008 | 1.000 | bdi-MIR171a\|ace-MIR170c\| | 2,678 |
| SALT00000072931_Cluster1376-009 | 1.000 | ace-MIR170c\| | 5,819 |
| SALT00000073489_Cluster1376-010 | 1.000 | bdi-MIR171a\| | 2,683 |
| SALT00000075767_Cluster1376-011 | 1.000 | bdi-MIR171a\|ace-MIR170c\| | 2,665 |
| SALT00000079084_Cluster1376-012 | 1.000 | bdi-MIR171a\|ace-MIR170c\| | 2,583 |
| SALT00000053574_Cluster1377-001 | 0.027 | NA | 5,816 |
| SALT00000053583_Cluster1411-016 | 1.000 | pta-MIR1310\|cln-MIR1310\|pde-MIR1310\|han-MIR1 | 5,731 |
| SALT00000090601_Cluster1430-001 | 1.000 | hvu-MIR6182\| | 5,713 |
| SALT00000085159_Cluster1441-001 | 0.125 | NA | 5,700 |
| SALT00000085627_Cluster1443-001 | 0.038 | NA | 5,698 |
| SALT00000092241_Cluster1461-001 | 1.000 | lincRNA11_PMID_24635777\| | 5,673 |
| SALT00000076019_Cluster1496-001 | 0.506 | NONATHT003785\|lincRNA493_PMID_24635777\| | 5,624 |
| SALT00000023305_Cluster1504-001 | 1.000 | lincRNA681_PMID_24635777\| | 3,007 |
| SALT00000025237_Cluster1504-002 | 1.000 | lincRNA681_PMID_24635777\| | 3,019 |
| SALT00000037273_Cluster1504-003 | 1.000 | lincRNA681_PMID_24635777\| | 2,155 |
| SALT00000062349_Cluster1504-005 | 1.000 | lincRNA681_PMID_24635777\| | 2,847 |

| | | | |
|---|---|---|---|
| SALT00000068362_Cluster1504-006 | 1.000 | lincRNA681_PMID_24635777\| | 3,050 |
| SALT00000068977_Cluster1504-008 | 1.000 | lincRNA681_PMID_24635777\| | 3,067 |
| SALT00000071457_Cluster1504-009 | 1.000 | lincRNA681_PMID_24635777\| | 2,606 |
| SALT00000082812_Cluster1504-010 | 1.000 | lincRNA681_PMID_24635777\| | 2,998 |
| SALT00000092778_Cluster1522-001 | 0.995 | pnrd_Osa_chr2.trna14_SerTGA\|pnrd_Osa_chr10.trna | 5,595 |
| SALT00000075003_Cluster1551-004 | 1.000 | NONATHT003839\|lincRNA493_PMID_24635777\|gl | 5,558 |
| SALT00000067008_Cluster1592-001 | 1.000 | lincRNA688_PMID_24635777\|NONATHT003836\|N | 3,191 |
| SALT00000079561_Cluster1607-001 | 1.000 | lincRNA11_PMID_24635777\| | 5,480 |
| SALT00000091259_Cluster1619-001 | 1.000 | lincRNA11_PMID_24635777\| | 5,470 |
| SALT00000059262_Cluster1636-001 | 0.231 | NA | 5,451 |
| SALT00000076318_Cluster1668-001 | 1.000 | lincRNA11_PMID_24635777\| | 2,949 |
| SALT00000090470_Cluster1668-002 | 1.000 | lincRNA11_PMID_24635777\| | 5,417 |
| SALT00000068887_Cluster1676-001 | 0.805 | lincRNA304_PMID_24635777\| | 5,409 |
| SALT00000062163_Cluster1696-001 | 0.166 | NA | 5,393 |
| SALT00000075213_Cluster1698-001 | 1.000 | lincRNA493_PMID_24635777\|NONATHT003785\| | 5,391 |
| SALT00000087097_Cluster1712-003 | 1.000 | pnrd_Gma_scaffold_1114.trna1_MetCAT\|pnrd_Ara_ | 5,375 |
| SALT00000088790_Cluster1713-001 | 1.000 | lincRNA11_PMID_24635777\| | 5,375 |
| SALT00000021408_Cluster1715-001 | 1.000 | TCONS_00028487\| | 2,968 |
| SALT00000024587_Cluster1715-002 | 1.000 | TCONS_00028487\| | 2,567 |
| SALT00000026502_Cluster1715-003 | 1.000 | TCONS_00028487\| | 2,514 |
| SALT00000028139_Cluster1715-005 | 1.000 | TCONS_00028487\| | 2,591 |
| SALT00000078482_Cluster1715-009 | 1.000 | TCONS_00028487\| | 3,328 |
| SALT00000088974_Cluster1715-010 | 1.000 | TCONS_00028487\| | 4,547 |
| SALT00000076892_Cluster1731-001 | 1.000 | lincRNA11_PMID_24635777\| | 4,485 |
| SALT00000078451_Cluster1731-002 | 1.000 | lincRNA11_PMID_24635777\| | 5,362 |
| SALT00000091000_Cluster1736-001 | 1.000 | lincRNA11_PMID_24635777\| | 5,358 |
| SALT00000027704_Cluster1743-001 | 0.089 | NA | 2,872 |
| SALT00000083731_Cluster1743-002 | 0.024 | NA | 3,506 |
| SALT00000086662_Cluster1743-003 | 0.058 | NA | 5,194 |
| SALT00000092128_Cluster1743-004 | 0.006 | NA | 5,352 |
| SALT00000087913_Cluster1743-006 | 0.120 | NA | 3,616 |
| SALT00000029381_Cluster1796-006 | 1.000 | GRMZM5G835418_T01\| | 3,316 |
| SALT00000083645_Cluster1796-012 | 1.000 | GRMZM5G835418_T01\| | 3,237 |
| SALT00000091860_Cluster1798-001 | 1.000 | lincRNA11_PMID_24635777\| | 5,310 |
| SALT00000092509_Cluster1811-001 | 1.000 | pnrd_Gma_chr13.trna72_HisGTG\|pnrd_Gma_scaffol | 5,301 |
| SALT00000090842_Cluster1812-001 | 1.000 | lincRNA376_PMID_24635777\| | 5,300 |
| SALT00000064314_Cluster1819-002 | 1.000 | cln-MIR1310\|pta-MIR1310\|pde-MIR1310\|han-MIR1 | 5,291 |
| SALT00000084072_Cluster1825-001 | 1.000 | lincRNA11_PMID_24635777\| | 5,284 |
| SALT00000064389_Cluster1839-001 | 0.229 | NA | 2,736 |
| SALT00000075460_Cluster1839-003 | 0.247 | NA | 2,649 |
| SALT00000091822_Cluster1851-003 | 0.213 | NA | 5,262 |
| SALT00000054699_Cluster1866-001 | 0.208 | NA | 1,544 |
| SALT00000076024_Cluster1867-001 | 0.063 | NA | 5,245 |
| SALT00000088401_Cluster1880-001 | 0.099 | NA | 5,238 |
| SALT00000091681_Cluster1896-003 | 1.000 | lincRNA483_PMID_24635777\| | 5,228 |
| SALT00000062683_Cluster1904-006 | 1.000 | TCONS_00018618\| | 2,457 |
| SALT00000026431_Cluster1905-002 | 0.248 | NA | 2,996 |
| SALT00000078617_Cluster1905-003 | 0.266 | NA | 5,224 |
| SALT00000080275_Cluster1905-004 | 0.246 | NA | 3,002 |
| SALT00000048508_Cluster1911-001 | 0.183 | NA | 1,192 |
| SALT00000088837_Cluster1916-001 | 0.998 | lincRNA11_PMID_24635777\| | 5,213 |
| SALT00000091086_Cluster1928-004 | 1.000 | lincRNA344_PMID_24635777\| | 5,207 |
| SALT00000093102_Cluster1947-001 | 0.232 | NA | 5,194 |
| SALT00000088777_Cluster2020-018 | 0.101 | NA | 3,414 |
| SALT00000067261_Cluster2027-001 | 1.000 | lincRNA11_PMID_24635777\| | 3,407 |
| SALT00000087074_Cluster2027-002 | 1.000 | lincRNA11_PMID_24635777\| | 5,135 |
| SALT00000093064_Cluster2053-001 | 1.000 | pnrd_Osa_chr4.trna83_SerGCT\|pnrd_Osa_chr10.trna | 5,110 |
| SALT00000086900_Cluster2085-001 | 1.000 | lincRNA11_PMID_24635777\| | 5,088 |

| | | | |
|---|---|---|---|
| SALT00000092282_Cluster2085-002 | 1.000 | lincRNA11_PMID_24635777\| | 3,779 |
| SALT00000000001_Cluster2099-001 | 1.000 | lincRNA688_PMID_24635777\| | 1,771 |
| SALT00000003827_Cluster2099-002 | 1.000 | lincRNA688_PMID_24635777\| | 1,575 |
| SALT00000006818_Cluster2099-003 | 1.000 | lincRNA688_PMID_24635777\| | 1,993 |
| SALT00000007970_Cluster2099-004 | 1.000 | lincRNA688_PMID_24635777\| | 1,767 |
| SALT00000009938_Cluster2099-005 | 1.000 | lincRNA688_PMID_24635777\| | 1,769 |
| SALT00000013413_Cluster2099-006 | 1.000 | lincRNA688_PMID_24635777\| | 1,894 |
| SALT00000018951_Cluster2099-008 | 1.000 | lincRNA688_PMID_24635777\| | 2,092 |
| SALT00000021366_Cluster2099-010 | 1.000 | lincRNA688_PMID_24635777\|tae-MIR2028a_1_npr | 2,683 |
| SALT00000021766_Cluster2099-011 | 1.000 | lincRNA688_PMID_24635777\| | 2,304 |
| SALT00000025176_Cluster2099-012 | 1.000 | lincRNA688_PMID_24635777\| | 1,773 |
| SALT00000026192_Cluster2099-013 | 1.000 | lincRNA688_PMID_24635777\| | 2,532 |
| SALT00000031751_Cluster2099-015 | 1.000 | lincRNA688_PMID_24635777\| | 1,622 |
| SALT00000035760_Cluster2099-017 | 1.000 | lincRNA688_PMID_24635777\| | 1,768 |
| SALT00000036968_Cluster2099-019 | 1.000 | lincRNA688_PMID_24635777\| | 1,763 |
| SALT00000040309_Cluster2099-022 | 1.000 | lincRNA688_PMID_24635777\| | 1,773 |
| SALT00000040953_Cluster2099-023 | 1.000 | lincRNA688_PMID_24635777\| | 5,081 |
| SALT00000042489_Cluster2099-026 | 1.000 | lincRNA688_PMID_24635777\| | 1,637 |
| SALT00000042602_Cluster2099-027 | 1.000 | lincRNA688_PMID_24635777\| | 1,762 |
| SALT00000043078_Cluster2099-028 | 1.000 | lincRNA688_PMID_24635777\| | 1,762 |
| SALT00000043106_Cluster2099-030 | 1.000 | lincRNA688_PMID_24635777\| | 1,775 |
| SALT00000043448_Cluster2099-031 | 1.000 | lincRNA688_PMID_24635777\| | 1,625 |
| SALT00000044186_Cluster2099-032 | 1.000 | lincRNA688_PMID_24635777\| | 1,707 |
| SALT00000044763_Cluster2099-034 | 1.000 | lincRNA688_PMID_24635777\| | 1,760 |
| SALT00000045212_Cluster2099-036 | 1.000 | lincRNA688_PMID_24635777\| | 2,043 |
| SALT00000045220_Cluster2099-037 | 1.000 | lincRNA688_PMID_24635777\| | 1,764 |
| SALT00000045386_Cluster2099-038 | 1.000 | lincRNA688_PMID_24635777\| | 1,774 |
| SALT00000045684_Cluster2099-039 | 1.000 | lincRNA688_PMID_24635777\| | 1,887 |
| SALT00000046112_Cluster2099-040 | 1.000 | lincRNA688_PMID_24635777\| | 1,779 |
| SALT00000046888_Cluster2099-041 | 1.000 | lincRNA688_PMID_24635777\| | 1,769 |
| SALT00000047402_Cluster2099-042 | 1.000 | lincRNA688_PMID_24635777\| | 1,765 |
| SALT00000047977_Cluster2099-043 | 1.000 | lincRNA688_PMID_24635777\| | 1,743 |
| SALT00000048931_Cluster2099-044 | 1.000 | lincRNA688_PMID_24635777\| | 1,771 |
| SALT00000049307_Cluster2099-045 | 1.000 | lincRNA688_PMID_24635777\| | 1,755 |
| SALT00000049667_Cluster2099-046 | 1.000 | lincRNA688_PMID_24635777\| | 1,732 |
| SALT00000050646_Cluster2099-048 | 1.000 | lincRNA688_PMID_24635777\| | 1,806 |
| SALT00000050925_Cluster2099-049 | 1.000 | lincRNA688_PMID_24635777\| | 1,809 |
| SALT00000051496_Cluster2099-051 | 1.000 | lincRNA688_PMID_24635777\| | 1,947 |
| SALT00000052456_Cluster2099-053 | 1.000 | lincRNA688_PMID_24635777\| | 1,763 |
| SALT00000053046_Cluster2099-054 | 1.000 | lincRNA688_PMID_24635777\| | 1,771 |
| SALT00000053232_Cluster2099-055 | 1.000 | lincRNA688_PMID_24635777\| | 1,771 |
| SALT00000053292_Cluster2099-056 | 1.000 | lincRNA688_PMID_24635777\| | 1,771 |
| SALT00000054222_Cluster2099-057 | 1.000 | lincRNA688_PMID_24635777\| | 1,619 |
| SALT00000054721_Cluster2099-059 | 1.000 | lincRNA688_PMID_24635777\| | 1,626 |
| SALT00000055603_Cluster2099-061 | 1.000 | lincRNA688_PMID_24635777\| | 1,760 |
| SALT00000055697_Cluster2099-063 | 1.000 | lincRNA688_PMID_24635777\| | 1,845 |
| SALT00000055860_Cluster2099-064 | 1.000 | lincRNA688_PMID_24635777\| | 1,765 |
| SALT00000058267_Cluster2099-068 | 1.000 | lincRNA688_PMID_24635777\| | 1,762 |
| SALT00000058439_Cluster2099-069 | 1.000 | lincRNA688_PMID_24635777\| | 1,625 |
| SALT00000058480_Cluster2099-070 | 1.000 | lincRNA688_PMID_24635777\| | 1,622 |
| SALT00000058529_Cluster2099-072 | 1.000 | lincRNA688_PMID_24635777\| | 1,817 |
| SALT00000058836_Cluster2099-075 | 1.000 | lincRNA688_PMID_24635777\| | 1,796 |
| SALT00000059138_Cluster2099-077 | 1.000 | lincRNA688_PMID_24635777\| | 1,769 |
| SALT00000059607_Cluster2099-078 | 1.000 | lincRNA688_PMID_24635777\| | 1,793 |
| SALT00000060383_Cluster2099-080 | 1.000 | lincRNA688_PMID_24635777\| | 1,730 |
| SALT00000060987_Cluster2099-082 | 1.000 | lincRNA688_PMID_24635777\| | 1,765 |
| SALT00000061073_Cluster2099-083 | 1.000 | lincRNA688_PMID_24635777\| | 1,627 |
| SALT00000061671_Cluster2099-087 | 1.000 | lincRNA688_PMID_24635777\| | 1,627 |

| | | | |
|---|---|---|---|
| SALT00000062262_Cluster2099-088 | 1.000 | lincRNA688_PMID_24635777\| | 1,784 |
| SALT00000067371_Cluster2099-089 | 1.000 | lincRNA688_PMID_24635777\| | 1,773 |
| SALT00000068032_Cluster2099-091 | 1.000 | lincRNA688_PMID_24635777\| | 1,772 |
| SALT00000070331_Cluster2099-092 | 1.000 | lincRNA688_PMID_24635777\| | 1,625 |
| SALT00000037315_Cluster2099-094 | 1.000 | lincRNA688_PMID_24635777\| | 2,325 |
| SALT00000004708_Cluster2101-001 | 1.000 | lincRNA304_PMID_24635777\| | 1,899 |
| SALT00000005618_Cluster2101-002 | 1.000 | lincRNA304_PMID_24635777\| | 2,036 |
| SALT00000008531_Cluster2101-003 | 1.000 | lincRNA304_PMID_24635777\| | 2,233 |
| SALT00000009882_Cluster2101-004 | 1.000 | lincRNA304_PMID_24635777\| | 1,905 |
| SALT00000020589_Cluster2101-005 | 1.000 | lincRNA304_PMID_24635777\| | 2,750 |
| SALT00000020626_Cluster2101-006 | 1.000 | lincRNA304_PMID_24635777\| | 2,662 |
| SALT00000020647_Cluster2101-007 | 1.000 | lincRNA304_PMID_24635777\| | 2,767 |
| SALT00000020665_Cluster2101-008 | 1.000 | lincRNA304_PMID_24635777\| | 2,598 |
| SALT00000020820_Cluster2101-009 | 1.000 | lincRNA304_PMID_24635777\| | 2,738 |
| SALT00000021354_Cluster2101-010 | 1.000 | lincRNA304_PMID_24635777\| | 2,803 |
| SALT00000021562_Cluster2101-011 | 1.000 | lincRNA304_PMID_24635777\| | 2,853 |
| SALT00000021704_Cluster2101-012 | 1.000 | lincRNA304_PMID_24635777\| | 2,779 |
| SALT00000023799_Cluster2101-013 | 1.000 | lincRNA304_PMID_24635777\| | 2,716 |
| SALT00000024286_Cluster2101-014 | 1.000 | lincRNA304_PMID_24635777\| | 2,541 |
| SALT00000025014_Cluster2101-015 | 1.000 | lincRNA304_PMID_24635777\| | 1,922 |
| SALT00000025255_Cluster2101-016 | 1.000 | lincRNA304_PMID_24635777\| | 2,920 |
| SALT00000026156_Cluster2101-017 | 1.000 | lincRNA304_PMID_24635777\| | 2,881 |
| SALT00000026428_Cluster2101-018 | 1.000 | lincRNA304_PMID_24635777\| | 2,849 |
| SALT00000027286_Cluster2101-019 | 1.000 | lincRNA304_PMID_24635777\| | 2,716 |
| SALT00000028307_Cluster2101-020 | 1.000 | lincRNA304_PMID_24635777\| | 2,678 |
| SALT00000028486_Cluster2101-021 | 1.000 | lincRNA304_PMID_24635777\| | 2,870 |
| SALT00000028505_Cluster2101-022 | 1.000 | lincRNA304_PMID_24635777\| | 2,833 |
| SALT00000028630_Cluster2101-023 | 1.000 | lincRNA304_PMID_24635777\| | 2,848 |
| SALT00000029469_Cluster2101-024 | 1.000 | lincRNA304_PMID_24635777\| | 2,621 |
| SALT00000029837_Cluster2101-025 | 1.000 | lincRNA304_PMID_24635777\| | 2,758 |
| SALT00000030790_Cluster2101-026 | 1.000 | lincRNA304_PMID_24635777\| | 2,865 |
| SALT00000031015_Cluster2101-027 | 1.000 | lincRNA304_PMID_24635777\| | 2,599 |
| SALT00000031077_Cluster2101-028 | 1.000 | lincRNA304_PMID_24635777\| | 2,797 |
| SALT00000031542_Cluster2101-029 | 1.000 | lincRNA304_PMID_24635777\| | 2,564 |
| SALT00000031763_Cluster2101-030 | 1.000 | lincRNA304_PMID_24635777\| | 2,830 |
| SALT00000062060_Cluster2101-031 | 1.000 | lincRNA304_PMID_24635777\| | 2,846 |
| SALT00000064821_Cluster2101-032 | 1.000 | lincRNA304_PMID_24635777\| | 2,663 |
| SALT00000065977_Cluster2101-033 | 1.000 | lincRNA304_PMID_24635777\| | 2,758 |
| SALT00000065989_Cluster2101-034 | 1.000 | lincRNA304_PMID_24635777\| | 2,702 |
| SALT00000066196_Cluster2101-035 | 1.000 | lincRNA304_PMID_24635777\| | 2,729 |
| SALT00000066739_Cluster2101-036 | 1.000 | lincRNA304_PMID_24635777\| | 2,730 |
| SALT00000067396_Cluster2101-037 | 1.000 | lincRNA304_PMID_24635777\| | 2,551 |
| SALT00000067944_Cluster2101-038 | 1.000 | lincRNA304_PMID_24635777\| | 5,080 |
| SALT00000070367_Cluster2101-039 | 1.000 | lincRNA304_PMID_24635777\| | 2,790 |
| SALT00000071522_Cluster2101-040 | 1.000 | lincRNA304_PMID_24635777\| | 2,826 |
| SALT00000071625_Cluster2101-041 | 1.000 | lincRNA304_PMID_24635777\| | 2,794 |
| SALT00000072779_Cluster2101-042 | 1.000 | lincRNA304_PMID_24635777\| | 2,848 |
| SALT00000074281_Cluster2101-043 | 1.000 | lincRNA304_PMID_24635777\| | 2,907 |
| SALT00000074476_Cluster2101-044 | 1.000 | lincRNA304_PMID_24635777\| | 2,769 |
| SALT00000077008_Cluster2101-045 | 1.000 | lincRNA304_PMID_24635777\| | 2,637 |
| SALT00000078146_Cluster2101-046 | 1.000 | lincRNA304_PMID_24635777\| | 2,797 |
| SALT00000080229_Cluster2101-047 | 1.000 | lincRNA304_PMID_24635777\| | 2,796 |
| SALT00000083937_Cluster2101-048 | 1.000 | lincRNA304_PMID_24635777\| | 2,749 |
| SALT00000084852_Cluster2101-049 | 1.000 | lincRNA304_PMID_24635777\| | 2,802 |
| SALT00000093746_Cluster2101-050 | 1.000 | lincRNA304_PMID_24635777\| | 2,871 |
| SALT00000093798_Cluster2101-051 | 1.000 | lincRNA304_PMID_24635777\| | 2,898 |
| SALT00000072559_Cluster2101-052 | 1.000 | lincRNA304_PMID_24635777\| | 2,771 |
| SALT00000027257_Cluster2101-053 | 1.000 | lincRNA304_PMID_24635777\| | 2,581 |

| | | | |
|---|---|---|---|
| SALT00000063428_Cluster2101-054 | 1.000 | lincRNA304_PMID_24635777\| | 2,617 |
| SALT00000016329_Cluster2170-001 | 0.153 | NA | 1,649 |
| SALT00000008723_Cluster2170-003 | 0.111 | NA | 1,644 |
| SALT00000007826_Cluster2172-002 | 0.090 | NA | 1,771 |
| SALT00000023581_Cluster2172-003 | 1.000 | osa-MIR5523\| | 2,631 |
| SALT00000026341_Cluster2172-004 | 1.000 | sit-MIR35-npr\|osa-MIR5523\| | 2,799 |
| SALT00000087656_Cluster2172-008 | 0.995 | sit-MIR35-npr\| | 5,036 |
| SALT00000044115_Cluster2197-001 | 1.000 | lincRNA688_PMID_24635777\| | 1,596 |
| SALT00000085534_Cluster2251-001 | 1.000 | lincRNA11_PMID_24635777\| | 4,252 |
| SALT00000087798_Cluster2251-002 | 1.000 | lincRNA11_PMID_24635777\| | 4,948 |
| SALT00000091993_Cluster2251-003 | 1.000 | lincRNA11_PMID_24635777\| | 4,979 |
| SALT00000092435_Cluster2251-004 | 1.000 | lincRNA11_PMID_24635777\| | 4,986 |
| SALT00000080365_Cluster2258-001 | 0.096 | NONATHT002123\| | 4,983 |
| SALT00000082027_Cluster2297-001 | 0.969 | lincRNA11_PMID_24635777\| | 4,487 |
| SALT00000089753_Cluster2297-002 | 1.000 | lincRNA11_PMID_24635777\| | 4,966 |
| SALT00000078427_Cluster2331-001 | 1.000 | lincRNA11_PMID_24635777\| | 4,948 |
| SALT00000085743_Cluster2331-002 | 1.000 | lincRNA11_PMID_24635777\| | 4,680 |
| SALT00000092874_Cluster2331-003 | 1.000 | lincRNA11_PMID_24635777\| | 4,519 |
| SALT00000049359_Cluster2347-001 | 0.177 | NA | 4,940 |
| SALT00000021025_Cluster2353-001 | 1.000 | lincRNA462_PMID_24635777\| | 3,063 |
| SALT00000032654_Cluster2353-007 | 1.000 | lincRNA462_PMID_24635777\| | 3,201 |
| SALT00000075974_Cluster2369-001 | 0.265 | NA | 4,925 |
| SALT00000038677_Cluster2375-001 | 0.599 | ghr-MIR5368*\|NONATHT003847\|lincRNA12_PMII | 2,101 |
| SALT00000072587_Cluster2375-002 | 0.599 | lincRNA372_PMID_24635777\|sit-MIR122-1-npr\|TC | 4,922 |
| SALT00000092660_Cluster2379-001 | 1.000 | TCONS_00047640\| | 4,920 |
| SALT00000063647_Cluster2410-001 | 1.000 | ghr-MIR5368b\|lincRNA493_PMID_24635777\|NON | 4,893 |
| SALT00000065300_Cluster2418-009 | 1.000 | lincRNA304_PMID_24635777\| | 4,885 |
| SALT00000090876_Cluster2431-001 | 0.194 | NA | 4,881 |
| SALT00000027587_Cluster2434-002 | 1.000 | lincRNA11_PMID_24635777\| | 2,516 |
| SALT00000032687_Cluster2434-003 | 1.000 | lincRNA11_PMID_24635777\| | 3,526 |
| SALT00000074341_Cluster2434-007 | 1.000 | lincRNA11_PMID_24635777\| | 2,759 |
| SALT00000051477_Cluster2475-011 | 1.000 | NONATHT000930\|peu-MIR2910\|NONATHT00216 | 1,766 |
| SALT00000078241_Cluster2538-001 | 0.073 | NA | 4,828 |
| SALT00000029449_Cluster2555-005 | 1.000 | NONATHT002169\|NONATHT000930\| | 3,770 |
| SALT00000086770_Cluster2560-001 | 1.000 | lincRNA11_PMID_24635777\| | 4,809 |
| SALT00000078957_Cluster2564-001 | 0.247 | lincRNA375_PMID_24635777\|lincRNA395_PMID_ | 4,804 |
| SALT00000086908_Cluster2564-002 | 0.254 | lincRNA395_PMID_24635777\|lincRNA375_PMID_ | 4,752 |
| SALT00000092063_Cluster2564-003 | 0.921 | lincRNA395_PMID_24635777\|lincRNA375_PMID_ | 4,715 |
| SALT00000000127_Cluster2573-001 | 1.000 | NONATHT002123\| | 1,445 |
| SALT00000001079_Cluster2573-002 | 1.000 | NONATHT002123\| | 1,487 |
| SALT00000003041_Cluster2573-003 | 1.000 | NONATHT002123\| | 1,595 |
| SALT00000003459_Cluster2573-004 | 1.000 | NONATHT002123\| | 1,790 |
| SALT00000007907_Cluster2573-005 | 1.000 | NONATHT002123\| | 2,196 |
| SALT00000009364_Cluster2573-006 | 1.000 | NONATHT002123\| | 2,308 |
| SALT00000013638_Cluster2573-007 | 1.000 | NONATHT002123\| | 1,983 |
| SALT00000021961_Cluster2573-008 | 1.000 | NONATHT002123\| | 2,937 |
| SALT00000023795_Cluster2573-009 | 1.000 | NONATHT002123\| | 2,576 |
| SALT00000032352_Cluster2573-012 | 1.000 | NONATHT002123\| | 3,597 |
| SALT00000032816_Cluster2573-013 | 1.000 | NONATHT002123\| | 3,718 |
| SALT00000038044_Cluster2573-014 | 1.000 | NONATHT002123\| | 1,834 |
| SALT00000042256_Cluster2573-015 | 1.000 | NONATHT002123\| | 1,835 |
| SALT00000044992_Cluster2573-016 | 1.000 | NONATHT002123\| | 1,489 |
| SALT00000049027_Cluster2573-017 | 0.997 | NONATHT002123\| | 1,256 |
| SALT00000063467_Cluster2573-018 | 1.000 | NONATHT002123\| | 2,617 |
| SALT00000064228_Cluster2573-019 | 1.000 | NONATHT002123\| | 2,593 |
| SALT00000071500_Cluster2573-020 | 1.000 | NONATHT002123\| | 2,547 |
| SALT00000074812_Cluster2573-021 | 1.000 | NONATHT002123\| | 2,429 |
| SALT00000076736_Cluster2573-022 | 1.000 | NONATHT002123\| | 3,327 |

| | | | |
|---|---|---|---|
| SALT00000080142_Cluster2573-023 | 1.000 | NONATHT002123\| | 3,069 |
| SALT00000081994_Cluster2573-024 | 1.000 | NONATHT002123\| | 4,798 |
| SALT00000052511_Cluster2583-001 | 0.213 | NA | 1,991 |
| SALT00000069495_Cluster2583-002 | 0.202 | NA | 3,028 |
| SALT00000086996_Cluster2583-005 | 0.197 | NA | 3,595 |
| SALT00000090051_Cluster2583-006 | 0.067 | NA | 4,794 |
| SALT00000088007_Cluster2585-001 | 1.000 | lincRNA376_PMID_24635777\| | 4,793 |
| SALT00000065698_Cluster2629-003 | 0.241 | NA | 2,602 |
| SALT00000071297_Cluster2629-004 | 0.262 | NA | 2,446 |
| SALT00000075224_Cluster2629-005 | 0.252 | NA | 2,521 |
| SALT00000075984_Cluster2629-006 | 0.255 | NA | 2,497 |
| SALT00000093359_Cluster2659-001 | 1.000 | lincRNA616_PMID_24635777\|osa-MIRf11419-npr\|c | 4,756 |
| SALT00000064401_Cluster2660-001 | 0.148 | NA | 3,090 |
| SALT00000093432_Cluster2660-002 | 1.000 | lincRNA372_PMID_24635777\| | 4,756 |
| SALT00000042739_Cluster2708-001 | 0.043 | NA | 2,028 |
| SALT00000057697_Cluster2708-002 | 0.049 | NA | 1,858 |
| SALT00000050966_Cluster2831-001 | 0.793 | lincRNA493_PMID_24635777\|NONATHT003785\| | 4,677 |
| SALT00000029656_Cluster2846-001 | 1.000 | TCONS_00011371\| | 2,703 |
| SALT00000087202_Cluster2846-002 | 1.000 | TCONS_00011371\| | 4,673 |
| SALT00000091260_Cluster2846-003 | 1.000 | TCONS_00011371\| | 3,991 |
| SALT00000046656_Cluster2860-001 | 0.058 | NA | 4,666 |
| SALT00000000447_Cluster2918-001 | 0.998 | osa-MIR5538\|lincRNA691_PMID_24635777\| | 1,804 |
| SALT00000005761_Cluster2918-002 | 0.998 | osa-MIR5538\|lincRNA691_PMID_24635777\| | 1,657 |
| SALT00000012253_Cluster2918-003 | 0.924 | pnrd_Gma_chr13.trna72_HisGTG\|pnrd_Mtr_chr1.trn | 1,822 |
| SALT00000025247_Cluster2918-004 | 0.998 | pnrd_Gma_scaffold_931.trna1_HisGTG\|pnrd_Osa_cl | 2,950 |
| SALT00000025712_Cluster2918-005 | 0.999 | pnrd_Gma_scaffold_828.trna2_HisGTG\|pnrd_Osa_cl | 2,805 |
| SALT00000030060_Cluster2918-006 | 0.998 | pnrd_Osa_chr12.trna16_HisGTG\|pnrd_Osa_chr10.trı | 3,127 |
| SALT00000030556_Cluster2918-007 | 1.000 | pnrd_Gma_scaffold_931.trna1_HisGTG\|pnrd_Osa_cl | 3,848 |
| SALT00000030734_Cluster2918-008 | 1.000 | pnrd_Osa_chr4.trna41_HisGTG\|pnrd_Osa_chr11.trna | 3,729 |
| SALT00000031869_Cluster2918-009 | 1.000 | pnrd_Mtr_chr8.trna33_HisGTG\|pnrd_Gma_scaffold_ | 3,705 |
| SALT00000036382_Cluster2918-010 | 0.999 | pnrd_Osa_chr8.trna46_HisGTG\|pnrd_Gma_scaffold_ | 1,963 |
| SALT00000040302_Cluster2918-011 | 1.000 | osa-MIR5538\|pnrd_Mtr_chr8.trna33_HisGTG\|pnrd_( | 2,109 |
| SALT00000064316_Cluster2918-012 | 0.999 | pnrd_Mtr_chr1.trna2_HisGTG\|pnrd_Mtr_chr8.trna33 | 2,552 |
| SALT00000067202_Cluster2918-013 | 0.630 | osa-MIR5538\|pnrd_Mtr_chr1.trna2_HisGTG\|pnrd_M | 2,957 |
| SALT00000069245_Cluster2918-014 | 0.994 | pnrd_Gma_scaffold_931.trna1_HisGTG\|pnrd_Osa_cl | 2,713 |
| SALT00000085559_Cluster2918-015 | 1.000 | pnrd_Osa_chr10.trna27_HisGTG\|pnrd_Gma_scaffolc | 3,708 |
| SALT00000091048_Cluster2918-016 | 1.000 | pnrd_Osa_chr11.trna7_HisGTG\|pnrd_Osa_chr4.trna4 | 4,639 |
| SALT00000092611_Cluster2918-017 | 1.000 | pnrd_Gma_chr13.trna72_HisGTG\|pnrd_Mtr_chr8.trn | 3,726 |
| SALT00000093074_Cluster2918-018 | 1.000 | pnrd_Gma_scaffold_828.trna2_HisGTG\|pnrd_Osa_cl | 3,704 |
| SALT00000079191_Cluster2918-019 | 0.993 | pnrd_Osa_chr10.trna27_HisGTG\|pnrd_Gma_scaffolc | 2,927 |
| SALT00000092915_Cluster2918-020 | 1.000 | pnrd_Osa_chr4.trna85_HisGTG\|pnrd_Osa_chr9.trna3 | 4,385 |
| SALT00000081835_Cluster2998-043 | 1.000 | NONATHT003839\|ghr-MIR5368b\|ghr-MIR5368*\|sit | 4,089 |
| SALT00000026686_Cluster3007-002 | 1.000 | lincRNA499_PMID_24635777\| | 3,104 |
| SALT00000048805_Cluster3007-003 | 1.000 | lincRNA499_PMID_24635777\|bdi-MIR5056\| | 1,288 |
| SALT00000049905_Cluster3007-004 | 1.000 | lincRNA499_PMID_24635777\|bdi-MIR5056\| | 1,780 |
| SALT00000086897_Cluster3007-007 | 1.000 | lincRNA499_PMID_24635777\| | 3,455 |
| SALT00000087055_Cluster3007-008 | 1.000 | lincRNA499_PMID_24635777\|bdi-MIR5056\| | 4,596 |
| SALT00000085747_Cluster3023-001 | 0.761 | NONATHT003839\|lincRNA12_PMID_24635777\|NC | 4,589 |
| SALT00000032805_Cluster3039-001 | 1.000 | GRMZM2G072760_T02\| | 3,747 |
| SALT00000087877_Cluster3039-002 | 1.000 | GRMZM2G072760_T02\| | 4,582 |
| SALT00000088205_Cluster3039-003 | 1.000 | GRMZM2G072760_T02\| | 4,573 |
| SALT00000089490_Cluster3039-004 | 1.000 | GRMZM2G072760_T02\| | 3,983 |
| SALT00000007223_Cluster3048-006 | 0.144 | NA | 1,412 |
| SALT00000011017_Cluster3048-008 | 0.741 | NONATHT002123\| | 1,546 |
| SALT00000026130_Cluster3048-010 | 0.566 | NONATHT002123\| | 2,654 |
| SALT00000026263_Cluster3048-011 | 1.000 | NONATHT002123\| | 2,625 |
| SALT00000027173_Cluster3048-012 | 0.536 | NONATHT002123\| | 2,825 |
| SALT00000028704_Cluster3048-013 | 1.000 | NONATHT002123\| | 3,269 |

| | | | |
|---|---|---|---|
| SALT00000028813_Cluster3048-014 | 1.000 | NONATHT002123\| | 3,227 |
| SALT00000033758_Cluster3048-015 | 1.000 | NONATHT002123\| | 3,601 |
| SALT00000034604_Cluster3048-016 | 1.000 | NONATHT002123\| | 4,011 |
| SALT00000051138_Cluster3048-021 | 0.281 | NA | 1,510 |
| SALT00000064410_Cluster3048-025 | 0.085 | NA | 2,251 |
| SALT00000069823_Cluster3048-028 | 1.000 | NONATHT002123\| | 2,968 |
| SALT00000070472_Cluster3048-029 | 0.999 | NONATHT002123\| | 2,864 |
| SALT00000077956_Cluster3048-030 | 1.000 | NONATHT002123\| | 3,002 |
| SALT00000078083_Cluster3048-031 | 1.000 | NONATHT002123\| | 4,577 |
| SALT00000080483_Cluster3048-032 | 1.000 | NONATHT002123\| | 2,546 |
| SALT00000084802_Cluster3048-033 | 1.000 | NONATHT002123\| | 4,203 |
| SALT00000085260_Cluster3048-034 | 1.000 | NONATHT002123\| | 3,271 |
| SALT00000086041_Cluster3048-035 | 1.000 | NONATHT002123\| | 3,766 |
| SALT00000090316_Cluster3048-036 | 1.000 | NONATHT002123\| | 3,886 |
| SALT00000031836_Cluster3058-001 | 1.000 | hvu-MIR6182\| | 3,956 |
| SALT00000032006_Cluster3058-002 | 1.000 | hvu-MIR6182\| | 3,936 |
| SALT00000081814_Cluster3058-007 | 1.000 | hvu-MIR6182\| | 3,695 |
| SALT00000081543_Cluster3062-001 | 0.157 | NA | 4,569 |
| SALT00000027727_Cluster3066-001 | 0.046 | NA | 2,527 |
| SALT00000086366_Cluster3066-003 | 0.026 | lincRNA686_PMID_24635777\|lincRNA616_PMID_ | 3,646 |
| SALT00000091731_Cluster3066-004 | 0.995 | lincRNA686_PMID_24635777\|lincRNA616_PMID_ | 3,608 |
| SALT00000071285_Cluster3077-001 | 0.996 | lincRNA304_PMID_24635777\| | 4,563 |
| SALT00000009050_Cluster3102-001 | 1.000 | far-MIR166\| | 2,290 |
| SALT00000023876_Cluster3102-003 | 1.000 | far-MIR166\| | 2,971 |
| SALT00000024372_Cluster3102-004 | 1.000 | far-MIR166\| | 2,913 |
| SALT00000024482_Cluster3102-005 | 1.000 | far-MIR166\| | 2,929 |
| SALT00000025207_Cluster3102-006 | 1.000 | far-MIR166\| | 2,932 |
| SALT00000025322_Cluster3102-007 | 1.000 | far-MIR166\| | 2,993 |
| SALT00000026307_Cluster3102-008 | 1.000 | far-MIR166\| | 2,883 |
| SALT00000048524_Cluster3102-011 | 1.000 | far-MIR166\| | 2,166 |
| SALT00000072156_Cluster3102-013 | 1.000 | far-MIR166\| | 3,222 |
| SALT00000075715_Cluster3102-014 | 1.000 | far-MIR166\| | 2,983 |
| SALT00000078331_Cluster3102-015 | 1.000 | far-MIR166\| | 2,991 |
| SALT00000078430_Cluster3102-016 | 1.000 | far-MIR166\| | 3,040 |
| SALT00000078926_Cluster3102-017 | 1.000 | far-MIR166\| | 4,553 |
| SALT00000084543_Cluster3102-018 | 1.000 | far-MIR166\| | 3,042 |
| SALT00000092756_Cluster3102-019 | 1.000 | far-MIR166\| | 3,811 |
| SALT00000056748_Cluster3133-004 | 0.125 | NA | 1,831 |
| SALT00000084927_Cluster3185-002 | 0.117 | NA | 4,521 |
| SALT00000093490_Cluster3185-008 | 0.117 | NA | 3,981 |
| SALT00000043665_Cluster3193-001 | 0.997 | pde-MIR1310\|han-MIR1310\|pta-MIR1310\|cln-MIR1 | 2,543 |
| SALT00000092200_Cluster3235-003 | 0.999 | osa-MIR5339\| | 4,502 |
| SALT00000078123_Cluster3266-001 | 0.999 | NONATHT003847\|lincRNA12_PMID_24635777\|NC | 4,489 |
| SALT00000051442_Cluster3267-001 | 0.136 | NA | 4,488 |
| SALT00000033847_Cluster3274-001 | 0.936 | tae-MIR141a_npr\| | 3,787 |
| SALT00000083413_Cluster3274-004 | 0.954 | tae-MIR141a_npr\| | 3,290 |
| SALT00000087509_Cluster3274-005 | 0.936 | tae-MIR141a_npr\| | 3,772 |
| SALT00000091620_Cluster3274-006 | 0.999 | tae-MIR141a_npr\| | 4,487 |
| SALT00000091630_Cluster3274-007 | 0.939 | tae-MIR141a_npr\| | 3,715 |
| SALT00000009150_Cluster3287-003 | 0.436 | NONATHT002169\|tae-MIR170a_npr\|tae-MIR170b_1 | 1,414 |
| SALT00000019812_Cluster3287-011 | 0.857 | peu-MIR2916\|peu-MIR2914\|tae-MIR170a_npr\|tae-M | 1,695 |
| SALT00000028956_Cluster3287-018 | 0.206 | NONATHT002169\|peu-MIR2910\|NONATHT00093C | 2,607 |
| SALT00000036536_Cluster3287-024 | 0.443 | peu-MIR2914\|NONATHT000930\|tae-MIR170b_npr\|t | 1,116 |
| SALT00000037827_Cluster3287-025 | 0.102 | tae-MIR170b_npr\|tae-MIR170a_npr\|NONATHT0009 | 1,076 |
| SALT00000072976_Cluster3287-041 | 0.143 | NONATHT000930\|NONATHT002169\|peu-MIR2914 | 833 |
| SALT00000071023_Cluster3287-049 | 1.000 | tae-MIR170b_npr\|tae-MIR170a_npr\|peu-MIR2910\|N | 3,807 |
| SALT00000022539_Cluster3297-002 | 0.270 | NA | 2,612 |
| SALT00000001123_Cluster3310-001 | 0.991 | lincRNA186_PMID_24635777\|lincRNA592_PMID_ | 1,730 |

| | | | |
|---|---|---|---|
| SALT00000001454_Cluster3310-002 | 0.988 | lincRNA92_PMID_24635777\|osa-MIR5503\|lincRNA | 2,052 |
| SALT00000001552_Cluster3310-003 | 0.997 | lincRNA186_PMID_24635777\|lincRNA592_PMID_ | 1,573 |
| SALT00000018212_Cluster3310-004 | 1.000 | pnrd_Mtr_chr1.trna88_HisGTG\|pnrd_Gma_scaffold_ | 1,414 |
| SALT00000029693_Cluster3310-005 | 1.000 | lincRNA186_PMID_24635777\|lincRNA92_PMID_2 | 2,911 |
| SALT00000029852_Cluster3310-006 | 1.000 | pnrd_Osa_chr4.trna41_HisGTG\|pnrd_Osa_chr11.trna | 3,887 |
| SALT00000034771_Cluster3310-008 | 0.984 | lincRNA592_PMID_24635777\|osa-MIR5503\|lincRN | 2,476 |
| SALT00000037898_Cluster3310-009 | 0.982 | lincRNA186_PMID_24635777\|lincRNA592_PMID_ | 1,550 |
| SALT00000043713_Cluster3310-010 | 0.991 | lincRNA92_PMID_24635777\|osa-MIR5503\|lincRNA | 1,618 |
| SALT00000049611_Cluster3310-011 | 0.992 | lincRNA592_PMID_24635777\|lincRNA92_PMID_2 | 1,453 |
| SALT00000055058_Cluster3310-012 | 0.982 | lincRNA186_PMID_24635777\|lincRNA592_PMID_ | 1,555 |
| SALT00000059749_Cluster3310-013 | 0.987 | lincRNA92_PMID_24635777\|osa-MIR5503\|lincRNA | 2,256 |
| SALT00000059902_Cluster3310-014 | 0.998 | pnrd_Mtr_chr1.trna88_HisGTG\|pnrd_Gma_scaffold_ | 1,446 |
| SALT00000081556_Cluster3310-015 | 0.995 | lincRNA691_PMID_24635777\| | 1,139 |
| SALT00000088769_Cluster3310-016 | 1.000 | pnrd_Osa_chr10.trna27_HisGTG\|pnrd_Osa_chr9.trna | 4,472 |
| SALT00000091391_Cluster3339-011 | 1.000 | NONATHT003839\|lincRNA493_PMID_24635777\|gı | 4,461 |
| SALT00000068730_Cluster3340-001 | 0.626 | sit-MIR122-1-npr\|TCONS_00018605\|sit-MIR121-1-ı | 3,016 |
| SALT00000088806_Cluster3340-002 | 0.907 | TCONS_00018605\|sit-MIR122-1-npr\|lincRNA372_P | 4,460 |
| SALT00000014284_Cluster3340-003 | 0.357 | ghr-MIR5368*\|sit-MIR121-1-npr\|NONATHT003847 | 1,842 |
| SALT00000076546_Cluster3346-001 | 0.094 | NA | 3,093 |
| SALT00000003919_Cluster3350-001 | 1.000 | NONATHT001142\|lincRNA11_PMID_24635777\| | 2,093 |
| SALT00000022578_Cluster3350-002 | 0.500 | NONATHT001142\| | 3,141 |
| SALT00000031243_Cluster3350-003 | 1.000 | NONATHT001142\| | 4,264 |
| SALT00000033945_Cluster3350-004 | 1.000 | NONATHT001142\| | 3,465 |
| SALT00000040517_Cluster3350-005 | 1.000 | NONATHT001142\|lincRNA11_PMID_24635777\| | 1,758 |
| SALT00000085497_Cluster3350-006 | 1.000 | lincRNA11_PMID_24635777\|NONATHT001142\| | 4,455 |
| SALT00000050089_Cluster3350-007 | 1.000 | NONATHT001142\|lincRNA11_PMID_24635777\| | 1,753 |
| SALT00000089898_Cluster3358-001 | 0.250 | NA | 4,451 |
| SALT00000092059_Cluster3364-004 | 1.000 | pnrd_Pop_scaffold_8.trna40_ProAGG\|pnrd_Gma_ch | 4,448 |
| SALT00000090091_Cluster3381-016 | 1.000 | lincRNA11_PMID_24635777\| | 4,440 |
| SALT00000092673_Cluster3395-001 | 0.225 | NA | 4,437 |
| SALT00000079806_Cluster3416-003 | 0.045 | NA | 2,728 |
| SALT00000022130_Cluster3455-010 | 0.906 | ghr-MIR4370\| | 1,597 |
| SALT00000040986_Cluster3455-039 | 0.213 | ghr-MIR4370\|cln-MIR1310\|pta-MIR1310\|pde-MIR1: | 1,984 |
| SALT00000050594_Cluster3455-042 | 0.861 | ghr-MIR4370\| | 1,575 |
| SALT00000085189_Cluster3457-001 | 0.833 | lincRNA372_PMID_24635777\|lincRNA12_PMID_2 | 4,410 |
| SALT00000085801_Cluster43-122 | 0.584 | sit-MIR122-1-npr\|lincRNA372_PMID_24635777\|TC | 3,265 |
| SALT00000082075_Cluster66-001 | 0.068 | NA | 12,566 |
| SALT00000031685_Cluster80-036 | 1.000 | lincRNA11_PMID_24635777\| | 2,043 |
| SALT00000054670_Cluster167-001 | 0.030 | NA | 9,940 |
| SALT00000093860_Cluster194-006 | 0.450 | peu-MIR2916\|NONATHT002169\|peu-MIR2910\|NON | 1,335 |
| SALT00000001723_Cluster220-001 | 0.743 | pde-MIR1310\|han-MIR1310\|pta-MIR1310\|cln-MIR1 | 1,860 |
| SALT00000004236_Cluster220-002 | 0.620 | pta-MIR1310\|cln-MIR1310\|pde-MIR1310\|han-MIR1 | 1,570 |
| SALT00000005093_Cluster220-003 | 0.755 | cln-MIR1310\|pta-MIR1310\|pde-MIR1310\|han-MIR1 | 1,706 |
| SALT00000005916_Cluster220-004 | 0.762 | cln-MIR1310\|pta-MIR1310\|pde-MIR1310\|han-MIR1 | 1,712 |
| SALT00000007713_Cluster220-005 | 0.273 | cln-MIR1310\|ghr-MIR4370\|pta-MIR1310\|han-MIR1: | 1,692 |
| SALT00000009845_Cluster220-007 | 0.120 | pde-MIR1310\|han-MIR1310\|pta-MIR1310\|cln-MIR1 | 1,271 |
| SALT00000013776_Cluster220-008 | 0.815 | han-MIR1310\|pde-MIR1310\|pta-MIR1310\|cln-MIR1 | 1,262 |
| SALT00000013806_Cluster220-009 | 0.290 | ghr-MIR4370\| | 1,404 |
| SALT00000015447_Cluster220-010 | 0.265 | ghr-MIR4370\| | 1,582 |
| SALT00000015542_Cluster220-011 | 0.112 | cln-MIR1310\|pta-MIR1310\|han-MIR1310\|pde-MIR1 | 1,375 |
| SALT00000016114_Cluster220-012 | 0.860 | cln-MIR1310\|pta-MIR1310\|han-MIR1310\|pde-MIR1 | 1,404 |
| SALT00000016529_Cluster220-013 | 0.137 | pde-MIR1310\|han-MIR1310\|cln-MIR1310\|pta-MIR1 | 1,054 |
| SALT00000016662_Cluster220-014 | 0.513 | ghr-MIR4370\|cln-MIR1310\|pta-MIR1310\|pde-MIR1: | 1,908 |
| SALT00000019305_Cluster220-015 | 0.180 | han-MIR1310\|pde-MIR1310\|cln-MIR1310\|pta-MIR1 | 1,244 |
| SALT00000019535_Cluster220-016 | 0.146 | pta-MIR1310\|cln-MIR1310\|han-MIR1310\|pde-MIR1 | 1,604 |
| SALT00000020418_Cluster220-017 | 0.741 | cln-MIR1310\|pta-MIR1310\|pde-MIR1310\|han-MIR1 | 1,930 |
| SALT00000021073_Cluster220-018 | 0.616 | pta-MIR1310\|cln-MIR1310\|ghr-MIR4370\|han-MIR1: | 2,692 |
| SALT00000024058_Cluster220-019 | 0.635 | han-MIR1310\|pde-MIR1310\|ghr-MIR4370\|cln-MIR1 | 2,577 |

| | | | |
|---|---|---|---|
| SALT00000025053_Cluster220-020 | 0.754 | han-MIR1310\|pde-MIR1310\|pta-MIR1310\|cln-MIR1 | 1,717 |
| SALT00000025482_Cluster220-021 | 0.161 | pta-MIR1310\|cln-MIR1310\|ghr-MIR4370\|han-MIR1: | 2,475 |
| SALT00000025780_Cluster220-022 | 0.692 | pde-MIR1310\|han-MIR1310\|pta-MIR1310\|cln-MIR1 | 2,216 |
| SALT00000025917_Cluster220-023 | 0.623 | pta-MIR1310\|cln-MIR1310\|ghr-MIR4370\|han-MIR1: | 2,647 |
| SALT00000029635_Cluster220-024 | 0.564 | cln-MIR1310\|pta-MIR1310\|han-MIR1310\|pde-MIR1 | 2,255 |
| SALT00000031548_Cluster220-025 | 0.664 | pde-MIR1310\|han-MIR1310\|pta-MIR1310\|cln-MIR1 | 1,604 |
| SALT00000031686_Cluster220-026 | 0.683 | pta-MIR1310\|cln-MIR1310\|han-MIR1310\|pde-MIR1 | 1,538 |
| SALT00000034741_Cluster220-027 | 0.759 | cln-MIR1310\|pta-MIR1310\|pde-MIR1310\|han-MIR1 | 1,741 |
| SALT00000034940_Cluster220-028 | 0.686 | pde-MIR1310\|han-MIR1310\|cln-MIR1310\|pta-MIR1 | 1,514 |
| SALT00000037959_Cluster220-029 | 0.671 | ghr-MIR4370\|cln-MIR1310\|pta-MIR1310\|han-MIR1: | 2,351 |
| SALT00000041788_Cluster220-031 | 0.224 | han-MIR1310\|pde-MIR1310\|ghr-MIR4370\|cln-MIR1 | 1,900 |
| SALT00000043112_Cluster220-032 | 0.739 | cln-MIR1310\|pta-MIR1310\|han-MIR1310\|pde-MIR1 | 1,888 |
| SALT00000044751_Cluster220-033 | 0.915 | pta-MIR1310\|cln-MIR1310\|ghr-MIR4370\|han-MIR1: | 2,061 |
| SALT00000047663_Cluster220-034 | 0.919 | han-MIR1310\|pde-MIR1310\|pta-MIR1310\|ghr-MIR4 | 1,976 |
| SALT00000051103_Cluster220-037 | 0.156 | pta-MIR1310\|cln-MIR1310\|han-MIR1310\|pde-MIR1 | 1,602 |
| SALT00000055548_Cluster220-038 | 0.301 | han-MIR1310\|pde-MIR1310\|cln-MIR1310\|pta-MIR1 | 1,523 |
| SALT00000057612_Cluster220-039 | 0.826 | pde-MIR1310\|han-MIR1310\|pta-MIR1310\|cln-MIR1 | 1,169 |
| SALT00000059886_Cluster220-040 | 0.225 | han-MIR1310\|pde-MIR1310\|cln-MIR1310\|ghr-MIR4 | 1,892 |
| SALT00000073038_Cluster220-042 | 0.663 | pde-MIR1310\|han-MIR1310\|ghr-MIR4370\|cln-MIR1 | 2,462 |
| SALT00000080195_Cluster220-043 | 0.658 | han-MIR1310\|ghr-MIR4370\| | 2,433 |
| SALT00000029098_Cluster322-004 | 1.000 | hvu-MIR6183\| | 3,413 |
| SALT00000030289_Cluster322-005 | 1.000 | hvu-MIR6183\| | 3,178 |
| SALT00000033503_Cluster322-012 | 1.000 | hvu-MIR6183\| | 3,786 |
| SALT00000083594_Cluster322-019 | 1.000 | hvu-MIR6183\| | 3,769 |
| SALT00000086648_Cluster322-020 | 1.000 | hvu-MIR6183\| | 3,778 |
| SALT00000088116_Cluster322-022 | 1.000 | hvu-MIR6183\| | 3,532 |
| SALT00000030512_Cluster474-001 | 1.000 | tae-MIR2028a_1_npr\| | 3,872 |
| SALT00000030523_Cluster474-002 | 1.000 | tae-MIR2028a_1_npr\| | 3,800 |
| SALT00000031203_Cluster474-003 | 1.000 | tae-MIR2028a_1_npr\| | 3,997 |
| SALT00000032189_Cluster474-004 | 1.000 | tae-MIR2028a_1_npr\| | 3,772 |
| SALT00000032252_Cluster474-005 | 1.000 | tae-MIR2028a_1_npr\| | 3,893 |
| SALT00000032384_Cluster474-006 | 1.000 | tae-MIR2028a_1_npr\| | 3,930 |
| SALT00000033200_Cluster474-007 | 1.000 | tae-MIR2028a_1_npr\| | 3,789 |
| SALT00000033486_Cluster474-008 | 1.000 | tae-MIR2028a_1_npr\| | 3,561 |
| SALT00000087975_Cluster474-010 | 1.000 | tae-MIR2028a_1_npr\| | 3,718 |
| SALT00000089283_Cluster474-011 | 1.000 | tae-MIR2028a_1_npr\| | 3,958 |
| SALT00000091092_Cluster474-012 | 1.000 | tae-MIR2028a_1_npr\| | 3,394 |
| SALT00000091734_Cluster474-013 | 1.000 | tae-MIR2028a_1_npr\| | 3,461 |
| SALT00000093304_Cluster474-014 | 1.000 | tae-MIR2028a_1_npr\| | 3,798 |
| SALT00000015554_Cluster567-001 | 1.000 | lincRNA442_PMID_24635777\|lincRNA482_PMID_ | 2,274 |
| SALT00000026801_Cluster620-001 | 1.000 | hvu-MIR6183\| | 2,787 |
| SALT00000028702_Cluster620-002 | 1.000 | hvu-MIR6183\| | 3,842 |
| SALT00000068864_Cluster620-006 | 1.000 | hvu-MIR6183\| | 2,230 |
| SALT00000078080_Cluster620-007 | 1.000 | hvu-MIR6183\| | 2,620 |
| SALT00000088739_Cluster620-008 | 1.000 | hvu-MIR6183\| | 3,821 |
| SALT00000046911_Cluster620-012 | 1.000 | hvu-MIR6183\| | 2,227 |
| SALT00000011588_Cluster916-001 | 0.272 | lincRNA688_PMID_24635777\|NONATHT003850\|gl | 1,766 |
| SALT00000022963_Cluster916-002 | 0.113 | osa-MIR5076\|NONATHT003836\|NONATHT003850 | 3,276 |
| SALT00000027312_Cluster916-003 | 0.161 | NONATHT003836\|osa-MIR5076\|ghr-MIR6424e\|NO | 2,701 |
| SALT00000057794_Cluster916-006 | 0.304 | NONATHT003836\|NONATHT003850\|ghr-MIR6424 | 1,548 |
| SALT00000050510_Cluster992-067 | 0.549 | lincRNA493_PMID_24635777\|NONATHT003785\| | 1,740 |
| SALT00000056770_Cluster992-073 | 0.196 | NONATHT003785\|lincRNA493_PMID_24635777\| | 1,561 |
| SALT00000085903_Cluster992-106 | 0.795 | NONATHT003785\|NONATHT003847\|lincRNA12_P | 3,504 |
| SALT00000082565_Cluster1158-001 | 1.000 | NONATHT003839\|lincRNA12_PMID_24635777\|NC | 6,353 |
| SALT00000063575_Cluster1189-007 | 1.000 | NONATHT003839\|lincRNA493_PMID_24635777\|T | 6,263 |
| SALT00000091856_Cluster1360-001 | 0.768 | NONATHT003268\| | 5,849 |
| SALT00000082283_Cluster1446-001 | 0.221 | sit-MIR122-1-npr\|lincRNA372_PMID_24635777\|NC | 5,695 |
| SALT00000079049_Cluster1559-007 | 1.000 | lincRNA493_PMID_24635777\|NONATHT003839\|gi | 4,014 |

| | | | |
|---|---|---|---|
| SALT00000086235_Cluster1590-001 | 1.000 | lincRNA616_PMID_24635777\| | 5,506 |
| SALT00000068041_Cluster1777-001 | 0.256 | NA | 2,792 |
| SALT00000086178_Cluster1777-002 | 0.149 | NA | 3,393 |
| SALT00000086314_Cluster1777-003 | 0.043 | NA | 5,324 |
| SALT00000036729_Cluster1819-001 | 1.000 | pde-MIR1310\|han-MIR1310\|cln-MIR1310\|pta-MIR1 | 1,904 |
| SALT00000007550_Cluster2057-005 | 0.277 | NA | 1,958 |
| SALT00000010969_Cluster2057-008 | 0.264 | NA | 2,053 |
| SALT00000027371_Cluster2057-015 | 0.128 | NA | 3,308 |
| SALT00000033890_Cluster2057-016 | 0.122 | NA | 3,350 |
| SALT00000043527_Cluster2057-020 | 0.294 | NA | 1,842 |
| SALT00000049576_Cluster2057-022 | 0.293 | NA | 1,493 |
| SALT00000068359_Cluster2057-028 | 0.196 | NA | 2,555 |
| SALT00000089503_Cluster2057-031 | 0.096 | NA | 3,762 |
| SALT00000091842_Cluster2057-032 | 0.090 | NA | 5,107 |
| SALT00000080284_Cluster2454-001 | 1.000 | pde-MIR1310\|han-MIR1310\|pta-MIR1310\|cln-MIR1 | 4,869 |
| SALT00000014082_Cluster2475-001 | 0.321 | ghr-MIR4370\| | 1,202 |
| SALT00000078288_Cluster2581-001 | 0.185 | NA | 4,796 |
| SALT00000016702_Cluster2656-001 | 1.000 | lincRNA369_PMID_24635777\| | 2,069 |
| SALT00000033019_Cluster2656-011 | 1.000 | lincRNA369_PMID_24635777\| | 3,998 |
| SALT00000071215_Cluster2656-012 | 1.000 | lincRNA369_PMID_24635777\| | 2,842 |
| SALT00000090936_Cluster2940-005 | 1.000 | lincRNA493_PMID_24635777\|NONATHT003785\| | 4,625 |
| SALT00000047089_Cluster3193-002 | 0.986 | pta-MIR1310\|cln-MIR1310\|han-MIR1310\|pde-MIR1 | 4,520 |
| SALT00000048191_Cluster3193-003 | 0.991 | cln-MIR1310\|pta-MIR1310\|han-MIR1310\|pde-MIR1 | 2,721 |
| SALT00000060943_Cluster3271-002 | 0.195 | NA | 2,249 |
| SALT00000085171_Cluster3271-004 | 0.250 | NA | 3,775 |
| SALT00000087771_Cluster3271-005 | 0.168 | NA | 4,488 |
| SALT00000022304_Cluster3273-001 | 1.000 | tae-MIR2028a_1_npr\| | 3,226 |
| SALT00000043239_Cluster3274-002 | 0.984 | tae-MIR141a_npr\| | 1,743 |
| SALT00000079175_Cluster3274-003 | 0.989 | tae-MIR141a_npr\| | 2,309 |
| SALT00000003717_Cluster3287-001 | 0.355 | peu-MIR2914\|peu-MIR2916\|tae-MIR170b_npr\|tae-M | 2,280 |
| SALT00000005398_Cluster3287-002 | 0.425 | peu-MIR2916\|peu-MIR2910\|NONATHT000930\|tae-I | 1,480 |
| SALT00000012401_Cluster3287-004 | 0.400 | NONATHT002169\|peu-MIR2910\|NONATHT00093C | 1,621 |
| SALT00000013787_Cluster3287-005 | 0.266 | peu-MIR2916\|NONATHT002169\|peu-MIR2910\|NOI | 2,140 |
| SALT00000014704_Cluster3287-006 | 0.390 | NONATHT002169\|tae-MIR170b_npr\|tae-MIR170a_ı | 1,680 |
| SALT00000015079_Cluster3287-007 | 0.336 | tae-MIR170a_npr\|tae-MIR170b_npr\|NONATHT0009 | 2,013 |
| SALT00000017684_Cluster3287-008 | 0.408 | peu-MIR2916\|peu-MIR2914\|NONATHT002169\|tae-I | 1,578 |
| SALT00000017762_Cluster3287-009 | 0.384 | peu-MIR2916\|peu-MIR2914\|peu-MIR2910\|NONATI | 2,104 |
| SALT00000019316_Cluster3287-010 | 0.390 | peu-MIR2916\|peu-MIR2914\|NONATHT000930\|peu- | 2,068 |
| SALT00000020493_Cluster3287-012 | 0.367 | peu-MIR2916\|peu-MIR2914\|peu-MIR2910\|NONATI | 1,821 |
| SALT00000021050_Cluster3287-013 | 0.764 | peu-MIR2916\|peu-MIR2914\|NONATHT002169\|peu- | 2,489 |
| SALT00000024036_Cluster3287-014 | 0.282 | NONATHT002169\|tae-MIR170b_npr\|tae-MIR170a_ı | 2,754 |
| SALT00000025033_Cluster3287-015 | 0.356 | NONATHT002169\|tae-MIR170b_npr\|tae-MIR170a_ı | 1,891 |
| SALT00000025318_Cluster3287-016 | 0.181 | peu-MIR2910\|NONATHT000930\|tae-MIR170a_npr\|t | 2,837 |
| SALT00000027223_Cluster3287-017 | 0.298 | tae-MIR170a_npr\|tae-MIR170b_npr\|NONATHT0009 | 2,642 |
| SALT00000031306_Cluster3287-019 | 0.718 | peu-MIR2914\|peu-MIR2916\|NONATHT002169\|tae-I | 2,829 |
| SALT00000032733_Cluster3287-020 | 0.399 | peu-MIR2916\|peu-MIR2914\|NONATHT002169\|tae-I | 1,628 |
| SALT00000034822_Cluster3287-021 | 0.166 | peu-MIR2914\|peu-MIR2916\|NONATHT002169\|tae-I | 1,675 |
| SALT00000035353_Cluster3287-022 | 0.173 | peu-MIR2914\|peu-MIR2916\|NONATHT002169\|peu- | 1,607 |
| SALT00000036093_Cluster3287-023 | 0.364 | peu-MIR2910\|NONATHT000930\|tae-MIR170a_npr\|t | 1,836 |
| SALT00000039095_Cluster3287-026 | 0.160 | NONATHT002169\|tae-MIR170a_npr\|tae-MIR170b_ı | 1,734 |
| SALT00000041164_Cluster3287-027 | 0.174 | tae-MIR170a_npr\|tae-MIR170b_npr\|NONATHT0009 | 1,593 |
| SALT00000044894_Cluster3287-028 | 0.192 | tae-MIR170a_npr\|tae-MIR170b_npr\|peu-MIR2910\|N | 1,423 |
| SALT00000048315_Cluster3287-029 | 0.158 | NONATHT002169\|tae-MIR170a_npr\|tae-MIR170b_ı | 1,752 |
| SALT00000048528_Cluster3287-030 | 0.177 | tae-MIR170b_npr\|tae-MIR170a_npr\|NONATHT0009 | 1,563 |
| SALT00000051747_Cluster3287-031 | 0.346 | peu-MIR2914\|peu-MIR2916\|peu-MIR2910\|NONATI | 1,950 |
| SALT00000053520_Cluster3287-032 | 0.417 | NONATHT002169\|tae-MIR170a_npr\|tae-MIR170b_ı | 1,525 |
| SALT00000055035_Cluster3287-033 | 0.596 | tae-MIR170a_npr\|tae-MIR170b_npr\|peu-MIR2910\|N | 2,022 |
| SALT00000055372_Cluster3287-034 | 0.903 | NONATHT002169\|NONATHT000930\|tae-MIR170a_ | 1,622 |

| | | | |
|---|---|---|---|
| SALT00000057538_Cluster3287-035 | 0.172 | peu-MIR2914\|peu-MIR2916\|NONATHT000930\|peu- | 1,615 |
| SALT00000057602_Cluster3287-036 | 0.369 | peu-MIR2914\|peu-MIR2916\|NONATHT002169\|tae-I | 1,808 |
| SALT00000060799_Cluster3287-037 | 0.320 | peu-MIR2914\|peu-MIR2916\|NONATHT002169\|peu- | 2,116 |
| SALT00000060800_Cluster3287-038 | 0.368 | peu-MIR2910\|NONATHT000930\|tae-MIR170b_npr\|t | 1,814 |
| SALT00000064353_Cluster3287-039 | 0.318 | NONATHT002169\|tae-MIR170b_npr\|tae-MIR170a_ı | 2,513 |
| SALT00000065800_Cluster3287-040 | 0.306 | peu-MIR2916\|NONATHT002169\|tae-MIR170b_npr\|t | 1,859 |
| SALT00000073264_Cluster3287-042 | 0.356 | peu-MIR2914\|peu-MIR2916\|NONATHT002169\|tae-I | 2,272 |
| SALT00000074688_Cluster3287-043 | 0.800 | peu-MIR2916\|peu-MIR2914\|peu-MIR2910\|NONATI | 2,195 |
| SALT00000076673_Cluster3287-044 | 0.393 | NONATHT002169\|NONATHT000930\|peu-MIR2910 | 1,667 |
| SALT00000089269_Cluster3287-045 | 0.441 | peu-MIR2916\|peu-MIR2914\|tae-MIR170a_npr\|tae-M | 4,482 |
| SALT00000089358_Cluster3287-046 | 0.640 | NONATHT002169\|NONATHT000930\|peu-MIR2910 | 3,290 |
| SALT00000093574_Cluster3287-047 | 0.441 | tae-MIR170a_npr\|tae-MIR170b_npr\|NONATHT0009 | 1,387 |
| SALT00000072143_Cluster3287-050 | 0.884 | NONATHT002169\|tae-MIR170a_npr\|tae-MIR170b_ı | 2,747 |
| SALT00000048001_Cluster3364-002 | 1.000 | pnrd_Ara_chr1.trna194_ProAGG\|pnrd_Osa_chr11.trı | 1,641 |
| SALT00000048820_Cluster3364-003 | 1.000 | pnrd_Osa_chr12.trna31_ProAGG\|pnrd_Ara_chr1.trna | 1,669 |
| SALT00000085981_Cluster3371-001 | 0.981 | lincRNA225_PMID_24635777\|lincRNA56_PMID_2 | 4,445 |
| SALT00000000380_Cluster3455-001 | 0.688 | cln-MIR1310\|pta-MIR1310\|han-MIR1310\|pde-MIR1 | 1,501 |
| SALT00000000956_Cluster3455-002 | 0.869 | ghr-MIR4370\| | 1,666 |
| SALT00000001315_Cluster3455-003 | 0.745 | han-MIR1310\|pde-MIR1310\|cln-MIR1310\|pta-MIR1 | 1,849 |
| SALT00000002249_Cluster3455-004 | 0.242 | ghr-MIR4370\| | 1,757 |
| SALT00000002931_Cluster3455-005 | 0.943 | ghr-MIR4370\| | 1,433 |
| SALT00000011369_Cluster3455-006 | 0.317 | ghr-MIR4370\| | 1,228 |
| SALT00000019544_Cluster3455-007 | 0.287 | ghr-MIR4370\| | 1,426 |
| SALT00000019981_Cluster3455-008 | 0.533 | pde-MIR1310\|han-MIR1310\|ghr-MIR4370\|cln-MIR1 | 3,169 |
| SALT00000021781_Cluster3455-009 | 0.727 | pta-MIR1310\|cln-MIR1310\|ghr-MIR4370\|pde-MIR1 | 2,869 |
| SALT00000022291_Cluster3455-011 | 0.444 | pta-MIR1310\|cln-MIR1310\|ghr-MIR4370\|han-MIR1 | 2,934 |
| SALT00000022981_Cluster3455-012 | 0.537 | ghr-MIR4370\|cln-MIR1310\|pta-MIR1310\|han-MIR1 | 3,147 |
| SALT00000023150_Cluster3455-013 | 0.708 | pta-MIR1310\|ghr-MIR4370\|cln-MIR1310\|han-MIR1 | 3,002 |
| SALT00000024882_Cluster3455-014 | 0.367 | han-MIR1310\|pde-MIR1310\|pta-MIR1310\|cln-MIR1 | 3,388 |
| SALT00000024974_Cluster3455-015 | 0.308 | ghr-MIR4370\| | 1,284 |
| SALT00000026882_Cluster3455-016 | 0.678 | han-MIR1310\|pde-MIR1310\|ghr-MIR4370\|cln-MIR1 | 3,282 |
| SALT00000027469_Cluster3455-017 | 0.144 | han-MIR1310\|pde-MIR1310\|pta-MIR1310\|cln-MIR1 | 2,658 |
| SALT00000028252_Cluster3455-018 | 0.151 | han-MIR1310\|pde-MIR1310\|pta-MIR1310\|cln-MIR1 | 2,583 |
| SALT00000028514_Cluster3455-019 | 0.106 | han-MIR1310\|pde-MIR1310\|cln-MIR1310\|ghr-MIR4 | 2,713 |
| SALT00000028652_Cluster3455-020 | 0.308 | han-MIR1310\|pde-MIR1310\|pta-MIR1310\|ghr-MIR4 | 3,760 |
| SALT00000029339_Cluster3455-021 | 0.322 | pde-MIR1310\|han-MIR1310\|pta-MIR1310\|ghr-MIR4 | 3,612 |
| SALT00000030654_Cluster3455-022 | 0.120 | han-MIR1310\|pde-MIR1310\|cln-MIR1310\|ghr-MIR4 | 2,958 |
| SALT00000031459_Cluster3455-023 | 0.242 | ghr-MIR4370\| | 1,752 |
| SALT00000031465_Cluster3455-024 | 0.395 | pde-MIR1310\|han-MIR1310\|pta-MIR1310\|ghr-MIR4 | 2,587 |
| SALT00000032572_Cluster3455-025 | 0.286 | han-MIR1310\|pde-MIR1310\|pta-MIR1310\|ghr-MIR4 | 3,910 |
| SALT00000033031_Cluster3455-026 | 0.219 | ghr-MIR4370\|cln-MIR1310\|NONATHT002169\|pta-M | 4,410 |
| SALT00000033248_Cluster3455-027 | 0.761 | pta-MIR1310\|ghr-MIR4370\|cln-MIR1310\|pde-MIR1 | 3,687 |
| SALT00000033713_Cluster3455-028 | 0.448 | pde-MIR1310\|han-MIR1310\|pta-MIR1310\|ghr-MIR4 | 3,650 |
| SALT00000034630_Cluster3455-029 | 0.184 | ghr-MIR4370\| | 2,245 |
| SALT00000034756_Cluster3455-030 | 0.849 | ghr-MIR4370\| | 1,900 |
| SALT00000034893_Cluster3455-031 | 0.224 | ghr-MIR4370\| | 1,893 |
| SALT00000034935_Cluster3455-032 | 0.232 | ghr-MIR4370\| | 1,944 |
| SALT00000035033_Cluster3455-033 | 0.180 | ghr-MIR4370\| | 2,283 |
| SALT00000035259_Cluster3455-034 | 0.211 | ghr-MIR4370\| | 2,006 |
| SALT00000035469_Cluster3455-035 | 0.241 | ghr-MIR4370\| | 1,763 |
| SALT00000035637_Cluster3455-036 | 0.261 | ghr-MIR4370\| | 1,614 |
| SALT00000035649_Cluster3455-037 | 0.194 | ghr-MIR4370\| | 2,153 |
| SALT00000037275_Cluster3455-038 | 0.252 | ghr-MIR4370\| | 1,679 |
| SALT00000041949_Cluster3455-040 | 0.698 | han-MIR1310\| | 2,239 |
| SALT00000044740_Cluster3455-041 | 0.256 | ghr-MIR4370\| | 1,652 |
| SALT00000052365_Cluster3455-043 | 0.190 | ghr-MIR4370\|cln-MIR1310\|pta-MIR1310\|han-MIR1 | 2,189 |
| SALT00000053156_Cluster3455-044 | 0.211 | ghr-MIR4370\| | 2,001 |
| SALT00000053784_Cluster3455-045 | 0.380 | ghr-MIR4370\| | 2,199 |

| | | | |
|---|---|---|---|
| SALT00000056522_Cluster3455-046 | 0.284 | ghr-MIR4370\| | 1,446 |
| SALT00000057195_Cluster3455-047 | 0.994 | han-MIR1310\|pde-MIR1310\|pta-MIR1310\|cln-MIR1 | 2,301 |
| SALT00000057393_Cluster3455-048 | 0.973 | ghr-MIR4370\| | 1,906 |
| SALT00000058634_Cluster3455-049 | 0.849 | ghr-MIR4370\| | 1,788 |
| SALT00000063131_Cluster3455-050 | 0.914 | ghr-MIR4370\| | 2,056 |
| SALT00000063720_Cluster3455-051 | 0.142 | cln-MIR1310\|ghr-MIR4370\|pta-MIR1310\|han-MIR1 | 2,679 |
| SALT00000069208_Cluster3455-053 | 0.798 | pde-MIR1310\|han-MIR1310\|pta-MIR1310\|ghr-MIR4 | 2,396 |
| SALT00000070856_Cluster3455-054 | 0.979 | han-MIR1310\|pde-MIR1310\|cln-MIR1310\|ghr-MIR4 | 2,490 |
| SALT00000073692_Cluster3455-055 | 0.609 | cln-MIR1310\|ghr-MIR4370\|pta-MIR1310\|pde-MIR1 | 2,732 |
| SALT00000074091_Cluster3455-056 | 0.660 | pta-MIR1310\|ghr-MIR4370\|cln-MIR1310\|pde-MIR1 | 3,319 |
| SALT00000075222_Cluster3455-057 | 0.148 | cln-MIR1310\|ghr-MIR4370\|pta-MIR1310\|han-MIR1 | 2,612 |
| SALT00000079348_Cluster3455-058 | 0.776 | pde-MIR1310\|han-MIR1310\|pta-MIR1310\|ghr-MIR4 | 4,349 |
| SALT00000079514_Cluster3455-059 | 0.234 | ghr-MIR4370\| | 1,816 |
| SALT00000081961_Cluster3455-060 | 0.399 | han-MIR1310\|pde-MIR1310\|cln-MIR1310\|ghr-MIR4 | 3,149 |
| SALT00000082220_Cluster3455-061 | 0.149 | ghr-MIR4370\|cln-MIR1310\|pta-MIR1310\|pde-MIR1 | 2,603 |
| SALT00000082961_Cluster3455-062 | 0.607 | pde-MIR1310\|han-MIR1310\|peu-MIR2910\|NONATI | 2,747 |
| SALT00000084318_Cluster3455-063 | 0.415 | ghr-MIR4370\|cln-MIR1310\|pta-MIR1310\|pde-MIR1 | 3,191 |
| SALT00000084720_Cluster3455-064 | 0.951 | ghr-MIR4370\| | 1,652 |
| SALT00000086583_Cluster3455-065 | 0.479 | pta-MIR1310\|ghr-MIR4370\|cln-MIR1310\|han-MIR1 | 3,476 |
| SALT00000088628_Cluster3455-066 | 0.332 | ghr-MIR4370\| | 1,130 |
| SALT00000090934_Cluster3455-067 | 0.579 | han-MIR1310\|pde-MIR1310\|ghr-MIR4370\|cln-MIR1 | 3,802 |
| SALT00000093393_Cluster3455-068 | 0.528 | pta-MIR1310\|ghr-MIR4370\|cln-MIR1310\|pde-MIR1 | 3,197 |
| SALT00000093540_Cluster3455-069 | 0.347 | pta-MIR1310\|ghr-MIR4370\|cln-MIR1310\|han-MIR1 | 3,509 |
| SALT00000093777_Cluster3455-070 | 0.180 | pde-MIR1310\|han-MIR1310\|pta-MIR1310\|ghr-MIR4 | 2,476 |
| SALT00000026853_Cluster3455-072 | 0.147 | han-MIR1310\|pde-MIR1310\|pta-MIR1310\|cln-MIR1 | 2,628 |
| SALT00000069724_Cluster3545-005 | 1.000 | lincRNA304_PMID_24635777\| | 4,376 |
| SALT00000070630_Cluster3545-006 | 1.000 | lincRNA304_PMID_24635777\| | 1,217 |
| SALT00000085723_Cluster3552-001 | 1.000 | GRMZM5G835418_T01\| | 4,375 |
| SALT00000086305_Cluster3552-002 | 1.000 | GRMZM5G835418_T01\| | 3,766 |
| SALT00000015698_Cluster3552-003 | 0.916 | GRMZM5G835418_T01\| | 2,187 |
| SALT00000032182_Cluster3554-002 | 1.000 | bdi-MIR7757\| | 3,802 |
| SALT00000088143_Cluster3554-009 | 0.991 | bdi-MIR7757\| | 3,945 |
| SALT00000029907_Cluster3562-001 | 0.269 | NA | 3,766 |
| SALT00000016650_Cluster3564-001 | 0.076 | NA | 2,031 |
| SALT00000086462_Cluster3592-001 | 0.100 | NA | 4,362 |
| SALT00000005991_Cluster3610-002 | 1.000 | TCONS_00055850\| | 2,163 |
| SALT00000011645_Cluster3610-003 | 1.000 | TCONS_00055850\|TCONS_00055849\| | 1,597 |
| SALT00000013991_Cluster3610-004 | 1.000 | TCONS_00055850\| | 1,825 |
| SALT00000017242_Cluster3610-005 | 1.000 | TCONS_00055850\|TCONS_00055849\| | 1,979 |
| SALT00000018529_Cluster3610-006 | 1.000 | TCONS_00055850\| | 1,839 |
| SALT00000020804_Cluster3610-007 | 1.000 | TCONS_00055850\|TCONS_00055849\| | 2,541 |
| SALT00000020852_Cluster3610-008 | 1.000 | TCONS_00055849\|TCONS_00055850\| | 2,867 |
| SALT00000021991_Cluster3610-009 | 1.000 | TCONS_00055850\| | 2,431 |
| SALT00000026133_Cluster3610-010 | 1.000 | TCONS_00055850\| | 2,048 |
| SALT00000029170_Cluster3610-011 | 1.000 | TCONS_00055850\|TCONS_00055849\| | 4,355 |
| SALT00000036643_Cluster3610-012 | 1.000 | TCONS_00055850\|TCONS_00055849\| | 2,258 |
| SALT00000036825_Cluster3610-013 | 1.000 | TCONS_00055850\|TCONS_00055849\| | 1,794 |
| SALT00000070189_Cluster3610-014 | 1.000 | TCONS_00055850\|TCONS_00055849\| | 2,536 |
| SALT00000074820_Cluster3610-015 | 1.000 | TCONS_00055849\|TCONS_00055850\| | 2,630 |
| SALT00000078708_Cluster3610-016 | 0.981 | TCONS_00055849\|TCONS_00055850\| | 2,792 |
| SALT00000079473_Cluster3610-017 | 1.000 | TCONS_00055850\|TCONS_00055849\| | 2,724 |
| SALT00000046568_Cluster3632-001 | 0.065 | NA | 1,819 |
| SALT00000052063_Cluster3635-001 | 1.000 | lincRNA688_PMID_24635777\| | 4,347 |
| SALT00000024314_Cluster3640-003 | 1.000 | osa-MIR5503\|lincRNA92_PMID_24635777\|lincRN | 3,199 |
| SALT00000031655_Cluster3640-006 | 0.983 | lincRNA186_PMID_24635777\|osa-MIR5503\|lincRN | 2,628 |
| SALT00000031731_Cluster3640-007 | 0.982 | lincRNA186_PMID_24635777\|lincRNA592_PMID_ | 2,684 |
| SALT00000061951_Cluster3640-009 | 0.982 | lincRNA592_PMID_24635777\|osa-MIR5503\|lincRN | 2,682 |
| SALT00000066114_Cluster3640-010 | 0.981 | osa-MIR5503\|lincRNA92_PMID_24635777\|lincRN | 2,762 |

| | | | |
|---|---|---|---|
| SALT00000073730_Cluster3640-011 | 1.000 | lincRNA92_PMID_24635777\|lincRNA186_PMID_2 | 2,835 |
| SALT00000074770_Cluster3640-012 | 0.980 | lincRNA592_PMID_24635777\|osa-MIR5503\|lincRN | 2,822 |
| SALT00000083946_Cluster3640-014 | 0.985 | osa-MIR5503\|lincRNA92_PMID_24635777\|lincRNA | 2,429 |
| SALT00000089836_Cluster3640-015 | 1.000 | lincRNA186_PMID_24635777\|osa-MIR5503\|lincRN | 4,346 |
| SALT00000081053_Cluster3655-001 | 0.682 | lincRNA623_PMID_24635777\|lincRNA12_PMID_2 | 4,340 |
| SALT00000071448_Cluster3687-001 | 1.000 | NONATHT003839\|NONATHT003847\| | 4,328 |
| SALT00000079569_Cluster3700-013 | 1.000 | TCONS_00055850\|TCONS_00055849\| | 4,325 |
| SALT00000077337_Cluster3703-001 | 0.031 | NA | 4,169 |
| SALT00000087966_Cluster3703-002 | 0.103 | NA | 4,325 |
| SALT00000027696_Cluster3732-001 | 0.144 | NONATHT002123\| | 2,561 |
| SALT00000010527_Cluster3732-003 | 0.253 | NA | 1,577 |
| SALT00000087274_Cluster3748-002 | 0.268 | NONATHT003839\|lincRNA493_PMID_24635777\|N | 4,310 |
| SALT00000079101_Cluster3763-010 | 0.257 | NA | 3,804 |
| SALT00000003793_Cluster3769-001 | 1.000 | lincRNA672_PMID_24635777\| | 2,024 |
| SALT00000022872_Cluster3769-002 | 1.000 | lincRNA672_PMID_24635777\| | 2,090 |
| SALT00000037832_Cluster3769-003 | 1.000 | lincRNA672_PMID_24635777\| | 1,466 |
| SALT00000056868_Cluster3769-005 | 1.000 | lincRNA672_PMID_24635777\| | 1,984 |
| SALT00000058029_Cluster3769-006 | 1.000 | lincRNA672_PMID_24635777\| | 1,950 |
| SALT00000059189_Cluster3769-007 | 1.000 | lincRNA672_PMID_24635777\| | 1,489 |
| SALT00000062806_Cluster3769-008 | 1.000 | lincRNA672_PMID_24635777\| | 2,042 |
| SALT00000079048_Cluster3785-001 | 0.935 | lincRNA304_PMID_24635777\| | 4,296 |
| SALT00000087114_Cluster3790-001 | 0.283 | NA | 4,295 |
| SALT00000013117_Cluster3835-001 | 0.671 | NONATHT000930\|NONATHT002169\| | 1,616 |
| SALT00000031333_Cluster3835-002 | 0.236 | cln-MIR1310\|NONATHT000930\|peu-MIR2910\|pde-M | 4,277 |
| SALT00000077052_Cluster3835-003 | 0.420 | cln-MIR1310\|peu-MIR2910\|NONATHT000930\|pde-M | 3,871 |
| SALT00000054000_Cluster3835-004 | 0.892 | pde-MIR1310\|han-MIR1310\|cln-MIR1310\|pta-MIR1 | 2,077 |
| SALT00000085797_Cluster3840-001 | 1.000 | sit-MIR95-npr\| | 4,277 |
| SALT00000022425_Cluster3866-001 | 0.977 | osa-MIR5503\|lincRNA92_PMID_24635777\|lincRNA | 2,995 |
| SALT00000022894_Cluster3866-002 | 0.974 | lincRNA592_PMID_24635777\|osa-MIR5503\|lincRN | 3,207 |
| SALT00000030117_Cluster3866-003 | 0.982 | lincRNA676_PMID_24635777\|osa-MIR5503\|lincRN | 4,270 |
| SALT00000032200_Cluster3866-004 | 0.956 | lincRNA186_PMID_24635777\|lincRNA676_PMID_ | 3,987 |
| SALT00000033684_Cluster3866-005 | 1.000 | lincRNA186_PMID_24635777\|lincRNA592_PMID_ | 3,890 |
| SALT00000080026_Cluster3866-006 | 0.950 | lincRNA186_PMID_24635777\|lincRNA592_PMID_ | 4,157 |
| SALT00000086625_Cluster3866-007 | 1.000 | lincRNA186_PMID_24635777\|osa-MIR5503\|lincRN | 4,269 |
| SALT00000088834_Cluster3866-008 | 0.969 | lincRNA92_PMID_24635777\|osa-MIR5503\|lincRNA | 3,448 |
| SALT00000092066_Cluster3866-009 | 1.000 | lincRNA186_PMID_24635777\|lincRNA676_PMID_ | 4,047 |
| SALT00000066965_Cluster3866-010 | 0.950 | lincRNA186_PMID_24635777\|lincRNA592_PMID_ | 4,146 |
| SALT00000085462_Cluster3887-003 | 1.000 | lincRNA131_PMID_24635777\| | 3,766 |
| SALT00000091839_Cluster3967-002 | 0.264 | NA | 3,316 |
| SALT00000032191_Cluster3999-001 | 1.000 | lincRNA304_PMID_24635777\| | 3,787 |
| SALT00000091012_Cluster3999-002 | 1.000 | lincRNA304_PMID_24635777\| | 4,224 |
| SALT00000003536_Cluster4006-001 | 1.000 | TCONS_00046663\| | 2,037 |
| SALT00000006808_Cluster4006-002 | 1.000 | TCONS_00046663\| | 2,111 |
| SALT00000089173_Cluster4006-003 | 1.000 | TCONS_00046663\| | 4,221 |
| SALT00000021168_Cluster4014-001 | 1.000 | cln-MIR166\|far-MIR166\|pta-MIR166c\| | 3,059 |
| SALT00000026993_Cluster4014-002 | 1.000 | pta-MIR166c\|cln-MIR166\|far-MIR166\| | 2,964 |
| SALT00000028366_Cluster4014-003 | 1.000 | cln-MIR166\|far-MIR166\|pta-MIR166c\| | 3,026 |
| SALT00000065850_Cluster4014-004 | 1.000 | pta-MIR166c\|far-MIR166\|cln-MIR166\| | 3,152 |
| SALT00000076204_Cluster4014-005 | 1.000 | pta-MIR166c\|cln-MIR166\|far-MIR166\| | 2,991 |
| SALT00000077372_Cluster4014-006 | 1.000 | cln-MIR166\|far-MIR166\|pta-MIR166c\| | 2,547 |
| SALT00000079977_Cluster4014-007 | 1.000 | cln-MIR166\|far-MIR166\|pta-MIR166c\| | 3,142 |
| SALT00000081264_Cluster4014-008 | 1.000 | pta-MIR166c\|far-MIR166\|cln-MIR166\| | 4,219 |
| SALT00000089924_Cluster4014-009 | 1.000 | pta-MIR166c\|far-MIR166\|cln-MIR166\| | 3,714 |
| SALT00000016023_Cluster4067-001 | 0.115 | NA | 1,910 |
| SALT00000033748_Cluster4067-002 | 0.097 | NA | 4,200 |
| SALT00000088504_Cluster4067-003 | 0.142 | NA | 3,591 |
| SALT00000043010_Cluster4076-001 | 1.000 | hvu-MIR6196\| | 2,257 |
| SALT00000067204_Cluster4076-003 | 1.000 | hvu-MIR6196\| | 2,343 |

| | | | |
|---|---|---|---|
| SALT00000073029_Cluster4076-004 | 1.000 | hvu-MIR6196\| | 4,196 |
| SALT00000079138_Cluster4076-005 | 1.000 | hvu-MIR6196\| | 2,321 |
| SALT00000079519_Cluster4076-006 | 1.000 | hvu-MIR6196\| | 2,537 |
| SALT00000025929_Cluster4076-007 | 1.000 | hvu-MIR6196\| | 2,661 |
| SALT00000033846_Cluster4095-001 | 1.000 | GRMZM5G835418_T01\| | 3,364 |
| SALT00000034025_Cluster4095-002 | 1.000 | GRMZM5G835418_T01\| | 3,525 |
| SALT00000071947_Cluster4095-003 | 1.000 | GRMZM5G835418_T01\| | 3,130 |
| SALT00000074449_Cluster4095-004 | 1.000 | GRMZM5G835418_T01\| | 3,191 |
| SALT00000078119_Cluster4095-005 | 1.000 | GRMZM5G835418_T01\| | 2,767 |
| SALT00000078690_Cluster4095-006 | 1.000 | GRMZM5G835418_T01\| | 3,972 |
| SALT00000088624_Cluster4095-007 | 1.000 | GRMZM5G835418_T01\| | 4,194 |
| SALT00000032638_Cluster4106-001 | 0.973 | tae-MIR141a_npr\| | 4,097 |
| SALT00000085804_Cluster4106-002 | 0.971 | tae-MIR141a_npr\| | 4,190 |
| SALT00000082582_Cluster4119-002 | 0.212 | NA | 4,188 |
| SALT00000086102_Cluster4135-001 | 0.049 | NA | 4,183 |
| SALT00000086737_Cluster4136-001 | 0.966 | lincRNA56_PMID_24635777\| | 4,183 |
| SALT00000086210_Cluster4142-001 | 1.000 | tae-MIR2028a_1_npr\|lincRNA688_PMID_24635777 | 4,182 |
| SALT00000079412_Cluster4162-001 | 1.000 | lincRNA372_PMID_24635777\| | 3,074 |
| SALT00000086475_Cluster4162-002 | 1.000 | lincRNA372_PMID_24635777\| | 4,175 |
| SALT00000091450_Cluster4162-003 | 1.000 | lincRNA372_PMID_24635777\| | 3,721 |
| SALT00000083785_Cluster4201-002 | 1.000 | NONATHT000373\| | 4,164 |
| SALT00000088093_Cluster4235-015 | 1.000 | lincRNA372_PMID_24635777\| | 4,156 |
| SALT00000003171_Cluster4236-006 | 1.000 | lincRNA246_PMID_24635777\|lincRNA302_PMID_ | 1,426 |
| SALT00000007361_Cluster4236-010 | 1.000 | lincRNA246_PMID_24635777\|lincRNA302_PMID_ | 1,497 |
| SALT00000007574_Cluster4236-011 | 1.000 | lincRNA302_PMID_24635777\|lincRNA246_PMID_ | 1,454 |
| SALT00000008083_Cluster4236-013 | 1.000 | lincRNA302_PMID_24635777\|lincRNA246_PMID_ | 1,567 |
| SALT00000009154_Cluster4236-014 | 1.000 | lincRNA246_PMID_24635777\|lincRNA302_PMID_ | 1,611 |
| SALT00000010313_Cluster4236-015 | 1.000 | lincRNA246_PMID_24635777\|lincRNA302_PMID_ | 1,521 |
| SALT00000011227_Cluster4236-016 | 1.000 | lincRNA246_PMID_24635777\|lincRNA302_PMID_ | 1,580 |
| SALT00000012223_Cluster4236-017 | 1.000 | lincRNA302_PMID_24635777\|lincRNA246_PMID_ | 1,437 |
| SALT00000013010_Cluster4236-018 | 1.000 | lincRNA246_PMID_24635777\|lincRNA302_PMID_ | 1,322 |
| SALT00000013393_Cluster4236-019 | 1.000 | lincRNA246_PMID_24635777\|lincRNA302_PMID_ | 1,574 |
| SALT00000014350_Cluster4236-020 | 1.000 | lincRNA246_PMID_24635777\|lincRNA302_PMID_ | 1,630 |
| SALT00000015233_Cluster4236-021 | 1.000 | lincRNA302_PMID_24635777\|lincRNA246_PMID_ | 1,520 |
| SALT00000017565_Cluster4236-023 | 1.000 | lincRNA246_PMID_24635777\|lincRNA302_PMID_ | 1,737 |
| SALT00000019863_Cluster4236-024 | 1.000 | lincRNA246_PMID_24635777\|lincRNA302_PMID_ | 1,547 |
| SALT00000020312_Cluster4236-025 | 1.000 | lincRNA246_PMID_24635777\|lincRNA302_PMID_ | 1,367 |
| SALT00000023517_Cluster4236-026 | 1.000 | lincRNA302_PMID_24635777\|lincRNA246_PMID_ | 1,454 |
| SALT00000023883_Cluster4236-027 | 1.000 | lincRNA246_PMID_24635777\|lincRNA302_PMID_ | 1,257 |
| SALT00000025104_Cluster4236-028 | 1.000 | lincRNA246_PMID_24635777\|lincRNA302_PMID_ | 1,700 |
| SALT00000025131_Cluster4236-029 | 1.000 | lincRNA302_PMID_24635777\|lincRNA246_PMID_ | 1,640 |
| SALT00000025132_Cluster4236-030 | 1.000 | lincRNA246_PMID_24635777\|lincRNA302_PMID_ | 1,557 |
| SALT00000031662_Cluster4236-035 | 1.000 | lincRNA302_PMID_24635777\|lincRNA246_PMID_ | 1,525 |
| SALT00000034668_Cluster4236-036 | 1.000 | lincRNA246_PMID_24635777\|lincRNA302_PMID_ | 1,497 |
| SALT00000034811_Cluster4236-037 | 1.000 | lincRNA246_PMID_24635777\|lincRNA302_PMID_ | 1,565 |
| SALT00000034909_Cluster4236-038 | 1.000 | lincRNA302_PMID_24635777\|lincRNA246_PMID_ | 1,476 |
| SALT00000035212_Cluster4236-039 | 1.000 | lincRNA302_PMID_24635777\|lincRNA246_PMID_ | 1,521 |
| SALT00000035357_Cluster4236-040 | 1.000 | lincRNA302_PMID_24635777\|lincRNA246_PMID_ | 1,515 |
| SALT00000035547_Cluster4236-041 | 1.000 | lincRNA302_PMID_24635777\|lincRNA246_PMID_ | 1,445 |
| SALT00000035744_Cluster4236-042 | 1.000 | lincRNA246_PMID_24635777\|lincRNA302_PMID_ | 1,578 |
| SALT00000035918_Cluster4236-043 | 1.000 | lincRNA302_PMID_24635777\|lincRNA246_PMID_ | 1,616 |
| SALT00000036735_Cluster4236-045 | 1.000 | lincRNA302_PMID_24635777\|lincRNA246_PMID_ | 1,633 |
| SALT00000036844_Cluster4236-046 | 1.000 | lincRNA302_PMID_24635777\|lincRNA246_PMID_ | 1,620 |
| SALT00000036912_Cluster4236-047 | 1.000 | lincRNA246_PMID_24635777\|lincRNA302_PMID_ | 1,579 |
| SALT00000037816_Cluster4236-048 | 1.000 | lincRNA246_PMID_24635777\|lincRNA302_PMID_ | 1,533 |
| SALT00000037872_Cluster4236-049 | 1.000 | lincRNA302_PMID_24635777\|lincRNA246_PMID_ | 1,506 |
| SALT00000037973_Cluster4236-050 | 1.000 | lincRNA302_PMID_24635777\|lincRNA246_PMID_ | 4,155 |
| SALT00000038605_Cluster4236-051 | 1.000 | lincRNA246_PMID_24635777\|lincRNA302_PMID_ | 1,593 |

| | | | |
|---|---|---|---|
| SALT00000038726_Cluster4236-052 | 1.000 | lincRNA302_PMID_24635777\|lincRNA246_PMID_ | 1,567 |
| SALT00000039979_Cluster4236-055 | 1.000 | lincRNA302_PMID_24635777\|lincRNA246_PMID_ | 2,031 |
| SALT00000040603_Cluster4236-056 | 1.000 | lincRNA302_PMID_24635777\|lincRNA246_PMID_ | 1,515 |
| SALT00000041939_Cluster4236-057 | 1.000 | lincRNA302_PMID_24635777\|lincRNA246_PMID_ | 1,596 |
| SALT00000042175_Cluster4236-058 | 1.000 | lincRNA302_PMID_24635777\|lincRNA246_PMID_ | 1,554 |
| SALT00000043015_Cluster4236-059 | 1.000 | lincRNA302_PMID_24635777\|lincRNA246_PMID_ | 1,614 |
| SALT00000043627_Cluster4236-060 | 1.000 | lincRNA302_PMID_24635777\|lincRNA246_PMID_ | 1,529 |
| SALT00000045627_Cluster4236-061 | 1.000 | lincRNA302_PMID_24635777\|lincRNA246_PMID_ | 1,614 |
| SALT00000046132_Cluster4236-064 | 1.000 | lincRNA246_PMID_24635777\|lincRNA302_PMID_ | 1,598 |
| SALT00000046855_Cluster4236-065 | 1.000 | lincRNA246_PMID_24635777\|lincRNA302_PMID_ | 1,648 |
| SALT00000047788_Cluster4236-066 | 1.000 | lincRNA302_PMID_24635777\|lincRNA246_PMID_ | 1,574 |
| SALT00000048784_Cluster4236-067 | 1.000 | lincRNA302_PMID_24635777\|lincRNA246_PMID_ | 1,536 |
| SALT00000049140_Cluster4236-068 | 1.000 | lincRNA246_PMID_24635777\|lincRNA302_PMID_ | 1,606 |
| SALT00000050027_Cluster4236-069 | 1.000 | lincRNA246_PMID_24635777\|lincRNA302_PMID_ | 1,588 |
| SALT00000051179_Cluster4236-070 | 1.000 | lincRNA302_PMID_24635777\|lincRNA246_PMID_ | 1,575 |
| SALT00000051441_Cluster4236-071 | 1.000 | lincRNA302_PMID_24635777\|lincRNA246_PMID_ | 1,569 |
| SALT00000051733_Cluster4236-072 | 1.000 | lincRNA246_PMID_24635777\|lincRNA302_PMID_ | 1,643 |
| SALT00000052563_Cluster4236-073 | 1.000 | lincRNA302_PMID_24635777\|lincRNA246_PMID_ | 1,596 |
| SALT00000052742_Cluster4236-074 | 1.000 | lincRNA246_PMID_24635777\|lincRNA302_PMID_ | 1,461 |
| SALT00000052834_Cluster4236-075 | 1.000 | lincRNA302_PMID_24635777\|lincRNA246_PMID_ | 1,599 |
| SALT00000053994_Cluster4236-076 | 1.000 | lincRNA302_PMID_24635777\|lincRNA246_PMID_ | 1,607 |
| SALT00000054622_Cluster4236-077 | 1.000 | lincRNA246_PMID_24635777\|lincRNA302_PMID_ | 1,604 |
| SALT00000056569_Cluster4236-078 | 1.000 | lincRNA246_PMID_24635777\|lincRNA302_PMID_ | 1,436 |
| SALT00000057189_Cluster4236-079 | 1.000 | lincRNA302_PMID_24635777\|lincRNA246_PMID_ | 1,570 |
| SALT00000057514_Cluster4236-080 | 1.000 | lincRNA246_PMID_24635777\|lincRNA302_PMID_ | 1,419 |
| SALT00000057522_Cluster4236-081 | 1.000 | lincRNA246_PMID_24635777\|lincRNA302_PMID_ | 1,532 |
| SALT00000060565_Cluster4236-082 | 1.000 | lincRNA302_PMID_24635777\|lincRNA246_PMID_ | 1,604 |
| SALT00000061694_Cluster4236-084 | 1.000 | lincRNA246_PMID_24635777\|lincRNA302_PMID_ | 1,528 |
| SALT00000066871_Cluster4236-086 | 1.000 | lincRNA302_PMID_24635777\|lincRNA246_PMID_ | 1,642 |
| SALT00000066963_Cluster4236-087 | 1.000 | lincRNA302_PMID_24635777\|lincRNA246_PMID_ | 1,509 |
| SALT00000071326_Cluster4236-088 | 1.000 | lincRNA302_PMID_24635777\|lincRNA246_PMID_ | 1,631 |
| SALT00000071931_Cluster4236-089 | 1.000 | lincRNA302_PMID_24635777\|lincRNA246_PMID_ | 1,570 |
| SALT00000053941_Cluster4236-090 | 1.000 | lincRNA246_PMID_24635777\|lincRNA302_PMID_ | 2,263 |
| SALT00000057869_Cluster4236-093 | 1.000 | lincRNA246_PMID_24635777\|lincRNA302_PMID_ | 1,922 |
| SALT00000060460_Cluster4236-094 | 1.000 | lincRNA302_PMID_24635777\|lincRNA246_PMID_ | 2,621 |
| SALT00000081829_Cluster4246-001 | 1.000 | lincRNA628_PMID_24635777\| | 4,152 |
| SALT00000088565_Cluster4253-002 | 1.000 | hvu-MIR6182\| | 4,151 |
| SALT00000085779_Cluster4262-001 | 0.991 | GRMZM2G042948_T01\| | 4,149 |
| SALT00000008402_Cluster4299-006 | 1.000 | lincRNA302_PMID_24635777\|lincRNA246_PMID_ | 1,311 |
| SALT00000015812_Cluster4299-011 | 1.000 | lincRNA302_PMID_24635777\|lincRNA246_PMID_ | 1,034 |
| SALT00000023347_Cluster4299-016 | 1.000 | lincRNA302_PMID_24635777\|lincRNA246_PMID_ | 1,485 |
| SALT00000045158_Cluster4299-025 | 1.000 | lincRNA302_PMID_24635777\|lincRNA246_PMID_ | 1,415 |
| SALT00000047926_Cluster4299-028 | 1.000 | lincRNA246_PMID_24635777\|lincRNA302_PMID_ | 1,623 |
| SALT00000054596_Cluster4299-031 | 1.000 | lincRNA302_PMID_24635777\|lincRNA246_PMID_ | 4,140 |
| SALT00000062655_Cluster4321-001 | 0.148 | NA | 1,601 |
| SALT00000077875_Cluster4340-001 | 0.056 | NA | 4,130 |
| SALT00000073242_Cluster4345-001 | 0.231 | NA | 2,909 |
| SALT00000032159_Cluster4347-001 | 1.000 | JNnc_loci0077\| | 4,090 |
| SALT00000093802_Cluster4347-002 | 1.000 | JNnc_loci0077\| | 4,130 |
| SALT00000077524_Cluster4363-001 | 0.027 | NA | 3,406 |
| SALT00000090626_Cluster4363-002 | 0.016 | NA | 4,125 |
| SALT00000086264_Cluster4413-001 | 0.051 | NA | 4,113 |
| SALT00000030759_Cluster4418-001 | 1.000 | hvu-MIR6182\| | 3,973 |
| SALT00000031294_Cluster4418-002 | 1.000 | hvu-MIR6182\| | 3,982 |
| SALT00000032037_Cluster4418-003 | 1.000 | hvu-MIR6182\| | 4,046 |
| SALT00000033221_Cluster4418-004 | 1.000 | hvu-MIR6182\| | 4,073 |
| SALT00000034035_Cluster4418-005 | 1.000 | hvu-MIR6182\| | 4,111 |
| SALT00000077494_Cluster4418-006 | 1.000 | hvu-MIR6182\| | 4,074 |

| | | | |
|---|---|---|---|
| SALT00000085481_Cluster4418-007 | 1.000 | hvu-MIR6182\| | 4,024 |
| SALT00000087879_Cluster4418-008 | 1.000 | hvu-MIR6182\| | 4,074 |
| SALT00000088999_Cluster4418-009 | 1.000 | hvu-MIR6182\| | 4,022 |
| SALT00000088136_Cluster4488-001 | 0.194 | NA | 4,093 |
| SALT00000061425_Cluster4507-001 | 1.000 | egu-MIR172d\| | 4,086 |
| SALT00000001577_Cluster4542-001 | 0.988 | lincRNA11_PMID_24635777\| | 1,808 |
| SALT00000017014_Cluster4542-002 | 0.649 | lincRNA11_PMID_24635777\| | 1,687 |
| SALT00000022978_Cluster4542-003 | 0.976 | pnrd_Osa_chr2.trna15_SerGCT\|pnrd_Gma_scaffold_ | 2,779 |
| SALT00000024645_Cluster4542-004 | 0.384 | pnrd_Gma_chr15.trna36_SerGCT\|pnrd_Osa_chr10.tr | 3,211 |
| SALT00000033181_Cluster4542-005 | 0.319 | pnrd_Osa_chr2.trna31_SerGCT\|pnrd_Gma_scaffold_ | 3,626 |
| SALT00000033618_Cluster4542-006 | 1.000 | pnrd_Gma_chr11.trna17_SerGCT\|pnrd_Osa_chr4.trn | 4,076 |
| SALT00000034223_Cluster4542-007 | 0.791 | pnrd_Osa_chr2.trna26_SerGCT\|pnrd_Mtr_chr4.trna9 | 3,885 |
| SALT00000068570_Cluster4542-008 | 0.385 | pnrd_Osa_chr10.trna36_SerGGA\|pnrd_Osa_chr4.trna | 3,217 |
| SALT00000073218_Cluster4542-009 | 0.373 | pnrd_Osa_chr10.trna62_SerGCT\|pnrd_Gma_scaffold | 3,286 |
| SALT00000077067_Cluster4542-010 | 0.386 | pnrd_Gma_chr15.trna20_SerGCT\|pnrd_Gma_scaffol | 3,212 |
| SALT00000088674_Cluster4542-011 | 0.314 | pnrd_Osa_chr10.trna62_SerGCT\|pnrd_Ara_chr2.trna | 3,647 |
| SALT00000064742_Cluster4542-012 | 0.939 | pnrd_Gma_chr15.trna20_SerGCT\|pnrd_Gma_scaffol | 2,440 |
| SALT00000092790_Cluster4569-001 | 0.193 | NA | 4,071 |
| SALT00000070403_Cluster4575-001 | 0.236 | NA | 4,069 |
| SALT00000034083_Cluster4583-001 | 0.277 | tae-MIR141a_npr\| | 4,067 |
| SALT00000077370_Cluster4583-002 | 0.453 | tae-MIR141a_npr\| | 2,981 |
| SALT00000085472_Cluster4583-003 | 0.937 | tae-MIR141a_npr\| | 3,762 |
| SALT00000086343_Cluster4583-004 | 0.306 | tae-MIR141a_npr\| | 3,871 |
| SALT00000086435_Cluster4583-005 | 0.938 | tae-MIR141a_npr\| | 3,746 |
| SALT00000088242_Cluster4583-006 | 0.932 | tae-MIR141a_npr\| | 3,886 |
| SALT00000089796_Cluster4583-007 | 0.356 | tae-MIR141a_npr\| | 3,554 |
| SALT00000000937_Cluster4596-001 | 1.000 | NONATHT003850\|NONATHT003836\|lincRNA493_ | 1,967 |
| SALT00000001298_Cluster4596-002 | 0.412 | lincRNA493_PMID_24635777\|NONATHT003850\|N | 1,504 |
| SALT00000003334_Cluster4596-003 | 1.000 | NONATHT003836\|NONATHT003850\|lincRNA493_ | 2,272 |
| SALT00000010349_Cluster4596-004 | 0.942 | lincRNA493_PMID_24635777\|NONATHT003850\|N | 1,687 |
| SALT00000018510_Cluster4596-005 | 1.000 | lincRNA688_PMID_24635777\|tae-MIR2028a_1_npr | 1,730 |
| SALT00000022966_Cluster4596-006 | 1.000 | lincRNA688_PMID_24635777\|tae-MIR2028a_1_npr | 2,504 |
| SALT00000025368_Cluster4596-007 | 1.000 | lincRNA493_PMID_24635777\|NONATHT003850\|N | 2,999 |
| SALT00000028563_Cluster4596-008 | 1.000 | lincRNA493_PMID_24635777\|NONATHT003850\|N | 3,854 |
| SALT00000028871_Cluster4596-009 | 1.000 | lincRNA493_PMID_24635777\|NONATHT003836\|N | 3,467 |
| SALT00000032542_Cluster4596-010 | 1.000 | NONATHT003836\|NONATHT003850\|lincRNA493_ | 3,743 |
| SALT00000034875_Cluster4596-011 | 1.000 | lincRNA688_PMID_24635777\|tae-MIR2028a_1_npr | 3,594 |
| SALT00000039716_Cluster4596-012 | 0.412 | lincRNA493_PMID_24635777\|NONATHT003850\|N | 1,503 |
| SALT00000044651_Cluster4596-013 | 1.000 | lincRNA493_PMID_24635777\|NONATHT003836\|N | 2,173 |
| SALT00000062849_Cluster4596-014 | 1.000 | NONATHT003850\|NONATHT003836\|lincRNA493_ | 1,955 |
| SALT00000069733_Cluster4596-015 | 1.000 | lincRNA688_PMID_24635777\|tae-MIR2028a_1_npr | 3,092 |
| SALT00000070410_Cluster4596-016 | 1.000 | tae-MIR2028a_1_npr\|lincRNA688_PMID_24635777 | 2,969 |
| SALT00000090707_Cluster4596-017 | 1.000 | NONATHT003850\|NONATHT003836\|lincRNA493_ | 4,064 |
| SALT00000052384_Cluster4596-018 | 0.992 | lincRNA688_PMID_24635777\| | 1,948 |
| SALT00000086089_Cluster4596-019 | 1.000 | lincRNA493_PMID_24635777\|NONATHT003850\|N | 3,898 |
| SALT00000058409_Cluster4596-020 | 1.000 | lincRNA688_PMID_24635777\| | 2,060 |
| SALT00000030125_Cluster4615-001 | 0.205 | NA | 3,781 |
| SALT00000083018_Cluster4615-002 | 0.175 | NA | 4,059 |
| SALT00000032614_Cluster4615-003 | 0.215 | NA | 3,702 |
| SALT00000077357_Cluster4640-001 | 1.000 | lincRNA482_PMID_24635777\|lincRNA442_PMID_ | 3,531 |
| SALT00000086564_Cluster4640-003 | 1.000 | lincRNA442_PMID_24635777\|lincRNA482_PMID_ | 3,609 |
| SALT00000087413_Cluster4640-004 | 1.000 | lincRNA442_PMID_24635777\|lincRNA482_PMID_ | 3,552 |
| SALT00000090224_Cluster4640-007 | 1.000 | lincRNA442_PMID_24635777\|lincRNA482_PMID_ | 4,053 |
| SALT00000092419_Cluster4648-001 | 1.000 | lincRNA372_PMID_24635777\| | 4,051 |
| SALT00000080926_Cluster4660-001 | 0.418 | ghr-MIR5368*\|lincRNA12_PMID_24635777\|NONA | 4,047 |
| SALT00000069897_Cluster4665-001 | 0.976 | tae-MIR141a_npr\| | 2,761 |
| SALT00000087598_Cluster4665-002 | 0.816 | tae-MIR141a_npr\| | 3,966 |
| SALT00000089545_Cluster4665-003 | 0.807 | tae-MIR141a_npr\| | 4,046 |

| | | | |
|---|---|---|---|
| SALT00000081517_Cluster4666-003 | 1.000 | GRMZM5G835418_T01\| | 3,126 |
| SALT00000082896_Cluster4666-004 | 1.000 | GRMZM5G835418_T01\| | 2,993 |
| SALT00000092946_Cluster4666-006 | 1.000 | GRMZM5G835418_T01\| | 3,391 |
| SALT00000086495_Cluster4682-001 | 0.780 | NONATHT003785\|lincRNA12_PMID_24635777\|lin | 3,636 |
| SALT00000092327_Cluster4682-002 | 0.726 | lincRNA493_PMID_24635777\|lincRNA12_PMID_2 | 4,043 |
| SALT00000033663_Cluster4712-001 | 0.238 | NA | 4,037 |
| SALT00000033690_Cluster4717-001 | 1.000 | hvu-MIR6182\| | 3,865 |
| SALT00000081500_Cluster4717-002 | 1.000 | hvu-MIR6182\| | 3,954 |
| SALT00000091252_Cluster4743-001 | 0.095 | NA | 4,030 |
| SALT00000032095_Cluster4830-001 | 1.000 | tae-MIR2028a_1_npr\| | 3,897 |
| SALT00000087461_Cluster4830-002 | 1.000 | tae-MIR2028a_1_npr\| | 3,747 |
| SALT00000093417_Cluster4830-003 | 1.000 | tae-MIR2028a_1_npr\| | 4,011 |
| SALT00000085773_Cluster4867-003 | 0.203 | NA | 4,004 |
| SALT00000080578_Cluster4877-001 | 1.000 | lincRNA676_PMID_24635777\|osa-MIR5538\|lincRN | 4,002 |
| SALT00000004648_Cluster4924-001 | 0.913 | lincRNA520_PMID_24635777\| | 1,843 |
| SALT00000066835_Cluster4924-002 | 0.789 | lincRNA520_PMID_24635777\| | 3,302 |
| SALT00000086937_Cluster4924-003 | 0.698 | lincRNA520_PMID_24635777\| | 3,992 |
| SALT00000078943_Cluster4924-004 | 0.800 | lincRNA520_PMID_24635777\| | 3,212 |
| SALT00000091992_Cluster4936-001 | 0.239 | NA | 3,990 |
| SALT00000034597_Cluster4942-003 | 1.000 | osa-MIR5523\|sit-MIR35-npr\| | 3,988 |
| SALT00000027665_Cluster4942-004 | 1.000 | sit-MIR35-npr\| | 2,671 |
| SALT00000030356_Cluster4942-005 | 1.000 | sit-MIR35-npr\| | 3,426 |
| SALT00000045840_Cluster4961-001 | 0.365 | NONATHT001131\| | 1,731 |
| SALT00000081670_Cluster4961-002 | 0.992 | NONATHT001131\| | 3,982 |
| SALT00000088969_Cluster4961-003 | 0.994 | NONATHT001131\| | 3,689 |
| SALT00000003706_Cluster4966-001 | 1.000 | pnrd_Osa_chr2.trna26_SerGCT\|pnrd_Mtr_chr8.trna1 | 1,944 |
| SALT00000010601_Cluster4966-002 | 1.000 | pnrd_Osa_chr4.trna88_SerTGA\|pnrd_Osa_chr3.trna4 | 1,648 |
| SALT00000024522_Cluster4966-005 | 1.000 | pnrd_Gma_chr2.trna23_SerTGA\|pnrd_Osa_chr2.trna | 3,038 |
| SALT00000026248_Cluster4966-006 | 1.000 | pnrd_Osa_chr4.trna88_SerTGA\|pnrd_Osa_chr3.trna4 | 3,305 |
| SALT00000029739_Cluster4966-007 | 1.000 | pnrd_Osa_chr2.trna14_SerTGA\|pnrd_Gma_chr2.trna | 2,689 |
| SALT00000049566_Cluster4966-008 | 1.000 | pnrd_Osa_chr4.trna80_SerTGA\|pnrd_Osa_chr5.trna5 | 1,465 |
| SALT00000061503_Cluster4966-009 | 0.992 | pnrd_Osa_chr12.trna17_SerTGA\|pnrd_Osa_chr1.trna | 1,654 |
| SALT00000064747_Cluster4966-010 | 0.992 | pnrd_Osa_chr3.trna45_SerTGA\|pnrd_Osa_chr4.trna8 | 1,560 |
| SALT00000090172_Cluster4966-011 | 1.000 | pnrd_Osa_chr4.trna80_SerTGA\|pnrd_Osa_chr5.trna5 | 3,981 |
| SALT00000033242_Cluster4982-001 | 0.285 | NA | 3,585 |
| SALT00000092484_Cluster4986-004 | 0.274 | NA | 3,486 |
| SALT00000088424_Cluster4994-001 | 0.184 | NA | 3,977 |
| SALT00000089355_Cluster4995-001 | 0.074 | NA | 3,977 |
| SALT00000087181_Cluster5023-001 | 0.982 | lincRNA493_PMID_24635777\|NONATHT003839\|g | 3,971 |
| SALT00000087821_Cluster5035-003 | 0.170 | NA | 3,969 |
| SALT00000090294_Cluster5042-001 | 0.252 | NA | 3,968 |
| SALT00000070307_Cluster5048-002 | 0.202 | NA | 2,596 |
| SALT00000073177_Cluster5048-004 | 0.257 | NA | 2,616 |
| SALT00000037666_Cluster5048-006 | 0.216 | NA | 1,897 |
| SALT00000041255_Cluster5061-010 | 0.087 | NA | 1,637 |
| SALT00000089632_Cluster5077-001 | 0.066 | NA | 3,961 |
| SALT00000045271_Cluster5083-005 | 1.000 | hvu-MIR6199\| | 2,149 |
| SALT00000090169_Cluster5083-008 | 1.000 | hvu-MIR6199\| | 3,612 |
| SALT00000030442_Cluster5083-013 | 1.000 | hvu-MIR6199\| | 3,867 |
| SALT00000088773_Cluster5090-001 | 0.029 | NA | 3,958 |
| SALT00000090513_Cluster5108-002 | 1.000 | TCONS_00054575\| | 3,954 |
| SALT00000082020_Cluster5108-003 | 1.000 | TCONS_00054575\| | 3,749 |
| SALT00000087456_Cluster5114-001 | 0.963 | lincRNA566_PMID_24635777\| | 3,953 |
| SALT00000037418_Cluster5142-001 | 0.265 | NA | 3,948 |
| SALT00000087594_Cluster5156-001 | 1.000 | lincRNA616_PMID_24635777\| | 3,946 |
| SALT00000032046_Cluster5172-001 | 0.225 | NA | 3,781 |
| SALT00000034008_Cluster5172-002 | 0.212 | NA | 3,869 |
| SALT00000089180_Cluster5210-001 | 0.437 | lincRNA344_PMID_24635777\| | 3,935 |

| | | | |
|---|---|---|---|
| SALT00000031167_Cluster5219-001 | 0.192 | NA | 2,582 |
| SALT00000088096_Cluster5243-001 | 0.142 | NA | 3,928 |
| SALT00000091874_Cluster5260-001 | 0.066 | NA | 3,925 |
| SALT00000087374_Cluster5270-001 | 0.999 | lincRNA366_PMID_24635777\|lincRNA229_PMID_ | 3,923 |
| SALT00000070408_Cluster5302-002 | 1.000 | osa-MIRf10781-npr\| | 2,256 |
| SALT00000083271_Cluster5310-002 | 0.249 | NA | 3,743 |
| SALT00000091617_Cluster5310-003 | 0.227 | NA | 3,915 |
| SALT00000047993_Cluster5345-002 | 1.000 | lincRNA688_PMID_24635777\| | 3,908 |
| SALT00000061604_Cluster5345-003 | 1.000 | lincRNA688_PMID_24635777\| | 1,741 |
| SALT00000049149_Cluster5378-001 | 0.999 | egu-MIR172d\| | 3,900 |
| SALT00000030142_Cluster5413-001 | 0.116 | NA | 3,894 |
| SALT00000081178_Cluster5427-001 | 1.000 | lincRNA375_PMID_24635777\| | 3,890 |
| SALT00000083034_Cluster5468-001 | 0.099 | NA | 3,882 |
| SALT00000032296_Cluster5474-001 | 0.170 | NA | 3,881 |
| SALT00000032525_Cluster5474-002 | 0.226 | NA | 3,380 |
| SALT00000034094_Cluster5474-003 | 0.207 | NA | 3,543 |
| SALT00000064621_Cluster5474-004 | 0.253 | NA | 3,174 |
| SALT00000078040_Cluster5474-005 | 0.216 | NA | 3,464 |
| SALT00000080912_Cluster5474-006 | 0.199 | NA | 2,687 |
| SALT00000089609_Cluster5474-008 | 0.200 | NA | 3,599 |
| SALT00000091777_Cluster5474-009 | 0.223 | NA | 3,409 |
| SALT00000067491_Cluster5487-001 | 0.231 | NA | 3,880 |
| SALT00000051670_Cluster5509-003 | 0.200 | NA | 1,833 |
| SALT00000090540_Cluster5541-001 | 0.998 | lincRNA595_PMID_24635777\| | 3,873 |
| SALT00000033375_Cluster5607-001 | 0.245 | NA | 3,777 |
| SALT00000090347_Cluster5607-004 | 0.268 | NA | 3,603 |
| SALT00000032029_Cluster5626-001 | 0.999 | tae-MIR141a_npr\| | 3,644 |
| SALT00000082578_Cluster5626-002 | 0.999 | tae-MIR141a_npr\| | 3,857 |
| SALT00000089863_Cluster5626-003 | 0.999 | tae-MIR141a_npr\| | 3,818 |
| SALT00000088552_Cluster5638-002 | 0.196 | NA | 3,855 |
| SALT00000056141_Cluster5669-001 | 0.057 | NA | 3,849 |
| SALT00000084894_Cluster5688-002 | 0.125 | NA | 3,846 |
| SALT00000086548_Cluster5711-001 | 0.180 | NA | 3,843 |
| SALT00000088297_Cluster5731-001 | 0.144 | NA | 3,839 |
| SALT00000033419_Cluster5744-001 | 1.000 | lincRNA304_PMID_24635777\| | 3,520 |
| SALT00000089446_Cluster5744-002 | 1.000 | lincRNA304_PMID_24635777\| | 3,836 |
| SALT00000092700_Cluster5744-003 | 1.000 | lincRNA304_PMID_24635777\| | 3,674 |
| SALT00000075851_Cluster5744-004 | 1.000 | lincRNA304_PMID_24635777\| | 4,925 |
| SALT00000088422_Cluster5744-005 | 1.000 | lincRNA304_PMID_24635777\| | 3,910 |
| SALT00000089708_Cluster5745-001 | 0.754 | lincRNA493_PMID_24635777\|lincRNA623_PMID_ | 3,836 |
| SALT00000092866_Cluster5747-001 | 0.060 | NA | 3,836 |
| SALT00000086358_Cluster5763-001 | 0.267 | NA | 3,833 |
| SALT00000069352_Cluster5769-001 | 0.946 | lincRNA582_PMID_24635777\| | 3,045 |
| SALT00000081723_Cluster5769-002 | 1.000 | lincRNA582_PMID_24635777\| | 3,832 |
| SALT00000015363_Cluster5769-003 | 1.000 | lincRNA582_PMID_24635777\| | 1,745 |
| SALT00000090970_Cluster5773-002 | 0.097 | NA | 3,832 |
| SALT00000057890_Cluster5794-001 | 0.126 | lincRNA493_PMID_24635777\| | 1,599 |
| SALT00000074636_Cluster5794-002 | 0.009 | TCONS_00018605\|NONATHT003839\|lincRNA493_ | 3,828 |
| SALT00000073297_Cluster5807-001 | 1.000 | lincRNA304_PMID_24635777\| | 3,826 |
| SALT00000092631_Cluster5847-001 | 0.195 | NA | 3,819 |
| SALT00000032255_Cluster5859-001 | 1.000 | TCONS_00054575\| | 3,815 |
| SALT00000032821_Cluster5859-002 | 1.000 | TCONS_00054575\| | 3,739 |
| SALT00000085030_Cluster5859-004 | 1.000 | TCONS_00054575\| | 3,589 |
| SALT00000085218_Cluster5859-005 | 1.000 | TCONS_00054575\| | 3,802 |
| SALT00000041865_Cluster5908-001 | 1.000 | lincRNA688_PMID_24635777\|cln-MIR1310\|pde-MI | 3,806 |
| SALT00000088746_Cluster5936-001 | 1.000 | NONATHT003847\|NONATHT003839\| | 3,804 |
| SALT00000091746_Cluster6013-001 | 0.552 | sit-MIR95-npr\| | 3,791 |
| SALT00000059458_Cluster6025-001 | 0.097 | NA | 1,355 |

| | | | |
|---|---|---|---|
| SALT00000093286_Cluster6035-001 | 0.245 | NA | 3,787 |
| SALT00000013820_Cluster6053-001 | 1.000 | lincRNA478_PMID_24635777\| | 1,633 |
| SALT00000025620_Cluster6053-002 | 1.000 | lincRNA478_PMID_24635777\| | 2,898 |
| SALT00000027112_Cluster6053-003 | 1.000 | lincRNA478_PMID_24635777\| | 2,690 |
| SALT00000027351_Cluster6053-004 | 1.000 | lincRNA478_PMID_24635777\| | 2,916 |
| SALT00000027867_Cluster6053-005 | 1.000 | lincRNA478_PMID_24635777\| | 2,911 |
| SALT00000029347_Cluster6053-006 | 1.000 | lincRNA478_PMID_24635777\| | 2,771 |
| SALT00000068343_Cluster6053-007 | 1.000 | lincRNA478_PMID_24635777\| | 2,703 |
| SALT00000090273_Cluster6053-008 | 1.000 | lincRNA478_PMID_24635777\| | 3,783 |
| SALT00000088259_Cluster6106-001 | 1.000 | lincRNA413_PMID_24635777\| | 3,776 |
| SALT00000086384_Cluster6114-001 | 1.000 | lincRNA309_PMID_24635777\|lincRNA276_PMID_ | 3,774 |
| SALT00000086741_Cluster6115-001 | 0.185 | NA | 3,774 |
| SALT00000001677_Cluster6125-001 | 1.000 | lincRNA672_PMID_24635777\| | 1,909 |
| SALT00000004542_Cluster6125-002 | 1.000 | lincRNA672_PMID_24635777\| | 2,331 |
| SALT00000005540_Cluster6125-003 | 1.000 | lincRNA672_PMID_24635777\| | 1,995 |
| SALT00000007619_Cluster6125-004 | 1.000 | lincRNA672_PMID_24635777\| | 2,060 |
| SALT00000011397_Cluster6125-005 | 1.000 | lincRNA672_PMID_24635777\| | 1,994 |
| SALT00000019341_Cluster6125-006 | 1.000 | lincRNA672_PMID_24635777\| | 1,947 |
| SALT00000030553_Cluster6125-007 | 1.000 | lincRNA672_PMID_24635777\| | 3,772 |
| SALT00000035913_Cluster6125-008 | 1.000 | lincRNA672_PMID_24635777\| | 2,101 |
| SALT00000045893_Cluster6125-009 | 1.000 | lincRNA672_PMID_24635777\| | 1,955 |
| SALT00000047358_Cluster6125-010 | 1.000 | lincRNA672_PMID_24635777\| | 1,974 |
| SALT00000050208_Cluster6125-011 | 1.000 | lincRNA672_PMID_24635777\| | 1,858 |
| SALT00000051470_Cluster6125-012 | 1.000 | lincRNA672_PMID_24635777\| | 2,150 |
| SALT00000090475_Cluster6162-001 | 1.000 | NONATHT001126\|lincRNA37_PMID_24635777\| | 3,767 |
| SALT00000022081_Cluster6193-004 | 0.997 | pnrd_Gma_scaffold_1210.trna1_ProTGG\|pnrd_Gma_ | 2,609 |
| SALT00000024279_Cluster6193-005 | 0.997 | pnrd_Gma_scaffold_1346.trna1_TrpCCA\|pnrd_Mtr_ | 2,542 |
| SALT00000033833_Cluster6193-006 | 0.994 | pnrd_Osa_chr1.trna94_TrpCCA\|pnrd_Mtr_chr7.trna4 | 3,762 |
| SALT00000041080_Cluster6193-007 | 0.262 | NA | 2,045 |
| SALT00000074802_Cluster6193-009 | 0.066 | pnrd_Mtr_chr1.trna6_TrpCCA\|pnrd_Mtr_chr7.trna41 | 2,491 |
| SALT00000091944_Cluster6200-001 | 0.153 | NA | 3,762 |
| SALT00000084444_Cluster6246-001 | 0.105 | NA | 3,753 |
| SALT00000086375_Cluster6264-002 | 1.000 | lincRNA88_PMID_24635777\| | 3,750 |
| SALT00000089595_Cluster6277-001 | 0.942 | sit-MIR122-1-npr\|lincRNA372_PMID_24635777\|TC | 3,749 |
| SALT00000031986_Cluster6283-001 | 1.000 | lincRNA304_PMID_24635777\| | 3,636 |
| SALT00000032798_Cluster6283-002 | 1.000 | lincRNA304_PMID_24635777\| | 3,708 |
| SALT00000033378_Cluster6283-003 | 1.000 | lincRNA304_PMID_24635777\| | 3,747 |
| SALT00000070855_Cluster6283-004 | 1.000 | lincRNA304_PMID_24635777\| | 3,359 |
| SALT00000076769_Cluster6283-005 | 1.000 | lincRNA304_PMID_24635777\| | 3,667 |
| SALT00000087683_Cluster6283-006 | 1.000 | lincRNA304_PMID_24635777\| | 3,659 |
| SALT00000091071_Cluster6283-007 | 1.000 | lincRNA304_PMID_24635777\| | 3,633 |
| SALT00000092533_Cluster6283-008 | 1.000 | lincRNA304_PMID_24635777\| | 3,522 |
| SALT00000091610_Cluster6292-001 | 0.214 | NA | 3,746 |
| SALT00000085324_Cluster6327-001 | 1.000 | lincRNA11_PMID_24635777\| | 3,740 |
| SALT00000034060_Cluster6343-001 | 1.000 | lincRNA1_PMID_24635777\| | 3,309 |
| SALT00000081691_Cluster6343-002 | 0.998 | lincRNA1_PMID_24635777\| | 3,696 |
| SALT00000087342_Cluster6343-003 | 0.998 | lincRNA1_PMID_24635777\| | 3,738 |
| SALT00000090744_Cluster6352-003 | 0.042 | NA | 3,737 |
| SALT00000084121_Cluster6376-001 | 0.094 | NA | 3,732 |
| SALT00000066289_Cluster6383-002 | 0.665 | JNnc_loci0896\|osa-MIR6253\| | 3,034 |
| SALT00000079304_Cluster6383-003 | 0.311 | osa-MIR6253\|JNnc_loci0896\| | 3,731 |
| SALT00000049687_Cluster6405-007 | 1.000 | NONATHT002169\|NONATHT000930\| | 1,776 |
| SALT00000083215_Cluster6437-001 | 1.000 | GRMZM5G835418_T01\| | 3,624 |
| SALT00000086022_Cluster6437-003 | 1.000 | GRMZM5G835418_T01\| | 3,361 |
| SALT00000086929_Cluster6468-001 | 0.039 | NA | 3,718 |
| SALT00000091694_Cluster6479-001 | 0.092 | NA | 3,716 |
| SALT00000092242_Cluster6487-001 | 0.068 | NA | 3,715 |
| SALT00000086515_Cluster6502-001 | 0.262 | NA | 3,712 |

| | | | |
|---|---|---|---|
| SALT00000069457_Cluster6517-003 | 0.262 | NA | 3,155 |
| SALT00000093070_Cluster6534-001 | 0.222 | NA | 3,708 |
| SALT00000067734_Cluster6544-001 | 0.228 | NA | 2,727 |
| SALT00000072293_Cluster6544-002 | 0.204 | NA | 2,929 |
| SALT00000086075_Cluster6544-003 | 0.763 | lincRNA292_PMID_24635777\|lincRNA314_PMID_ | 3,478 |
| SALT00000093476_Cluster6544-004 | 0.732 | lincRNA314_PMID_24635777\|lincRNA292_PMID_ | 3,707 |
| SALT00000087874_Cluster6547-001 | 1.000 | GRMZM5G835418_T01\| | 3,706 |
| SALT00000093465_Cluster6547-002 | 1.000 | GRMZM5G835418_T01\| | 3,448 |
| SALT00000065969_Cluster6557-001 | 0.170 | NA | 3,704 |
| SALT00000065891_Cluster6574-002 | 0.078 | NA | 2,996 |
| SALT00000085745_Cluster6593-001 | 0.558 | lincRNA344_PMID_24635777\| | 3,699 |
| SALT00000082979_Cluster6612-005 | 1.000 | lincRNA304_PMID_24635777\| | 3,696 |
| SALT00000057708_Cluster6630-003 | 0.168 | NA | 1,654 |
| SALT00000079891_Cluster6636-001 | 0.039 | NA | 3,692 |
| SALT00000091355_Cluster6637-001 | 0.279 | NA | 3,692 |
| SALT00000083340_Cluster6652-001 | 0.292 | NA | 3,689 |
| SALT00000088048_Cluster6657-001 | 1.000 | osa-MIR396d\| | 3,688 |
| SALT00000088523_Cluster6659-001 | 1.000 | lincRNA225_PMID_24635777\|lincRNA696_PMID_ | 3,688 |
| SALT00000091353_Cluster6672-001 | 0.740 | lincRNA520_PMID_24635777\| | 3,686 |
| SALT00000079747_Cluster6689-001 | 0.762 | TCONS_00018605\|pde-MIR1310\|lincRNA372_PMI | 3,683 |
| SALT00000033243_Cluster6716-001 | 0.110 | NA | 3,679 |
| SALT00000010781_Cluster6717-001 | 0.140 | NA | 1,890 |
| SALT00000089784_Cluster6766-001 | 0.295 | NA | 3,672 |
| SALT00000019612_Cluster6780-002 | 1.000 | lincRNA691_PMID_24635777\|lincRNA756_PMID_ | 1,721 |
| SALT00000089005_Cluster6780-006 | 1.000 | lincRNA691_PMID_24635777\|lincRNA756_PMID_ | 3,669 |
| SALT00000091446_Cluster6785-001 | 1.000 | lincRNA301_PMID_24635777\|lincRNA634_PMID_ | 3,668 |
| SALT00000061643_Cluster6788-006 | 1.000 | lincRNA186_PMID_24635777\|osa-MIR5503\|lincRN | 3,667 |
| SALT00000089737_Cluster6819-001 | 0.187 | NA | 3,663 |
| SALT00000076216_Cluster6874-005 | 0.052 | NA | 2,791 |
| SALT00000093201_Cluster6878-001 | 0.187 | NA | 3,655 |
| SALT00000022469_Cluster6898-001 | 1.000 | bdi-MIR5056\|lincRNA499_PMID_24635777\| | 2,738 |
| SALT00000033407_Cluster6898-002 | 1.000 | lincRNA499_PMID_24635777\|bdi-MIR5056\| | 3,651 |
| SALT00000056298_Cluster6898-003 | 1.000 | bdi-MIR5056\|lincRNA499_PMID_24635777\| | 1,767 |
| SALT00000066605_Cluster6898-004 | 1.000 | lincRNA499_PMID_24635777\|bdi-MIR5056\| | 2,767 |
| SALT00000084532_Cluster6898-005 | 1.000 | lincRNA499_PMID_24635777\|bdi-MIR5056\| | 3,059 |
| SALT00000071922_Cluster6917-001 | 0.110 | NONATHT000930\|peu-MIR2910\|NONATHT00216 | 3,648 |
| SALT00000087100_Cluster6928-001 | 0.993 | TCONS_00018618\| | 3,647 |
| SALT00000089221_Cluster6944-001 | 0.269 | NA | 3,645 |
| SALT00000075829_Cluster6955-001 | 1.000 | lincRNA618_PMID_24635777\| | 3,642 |
| SALT00000087029_Cluster6998-001 | 0.234 | NA | 3,636 |
| SALT00000033605_Cluster7020-001 | 0.283 | NA | 3,633 |
| SALT00000004112_Cluster7037-007 | 1.000 | lincRNA672_PMID_24635777\| | 2,116 |
| SALT00000017395_Cluster7037-016 | 1.000 | lincRNA672_PMID_24635777\| | 1,983 |
| SALT00000044720_Cluster7037-024 | 1.000 | lincRNA672_PMID_24635777\| | 3,631 |
| SALT00000053776_Cluster7037-027 | 1.000 | lincRNA672_PMID_24635777\| | 1,879 |
| SALT00000019036_Cluster7037-030 | 1.000 | lincRNA672_PMID_24635777\| | 1,927 |
| SALT00000045818_Cluster7037-031 | 1.000 | lincRNA672_PMID_24635777\| | 2,041 |
| SALT00000052836_Cluster7043-016 | 1.000 | pde-MIR1310\|han-MIR1310\|cln-MIR1310\|pta-MIR1 | 3,630 |
| SALT00000012601_Cluster7086-001 | 0.230 | NA | 1,725 |
| SALT00000030482_Cluster7088-001 | 1.000 | lincRNA688_PMID_24635777\|NONATHT003785\|li | 3,623 |
| SALT00000055503_Cluster7092-017 | 1.000 | ghr-MIR5368b\|NONATHT003839\|NONATHT00384 | 3,622 |
| SALT00000044132_Cluster7092-022 | 1.000 | osa-MIR5079a\|osa-MIR5079b\| | 2,656 |
| SALT00000083014_Cluster7146-001 | 0.656 | lincRNA623_PMID_24635777\|lincRNA12_PMID_2 | 3,616 |
| SALT00000034470_Cluster7163-001 | 1.000 | lincRNA11_PMID_24635777\| | 3,613 |
| SALT00000085027_Cluster7164-018 | 1.000 | pnrd_Ara_chr2.trna6_TrpCCA\| | 3,613 |
| SALT00000088794_Cluster7205-001 | 1.000 | lincRNA369_PMID_24635777\| | 3,607 |
| SALT00000088515_Cluster7217-001 | 0.829 | lincRNA372_PMID_24635777\| | 3,606 |
| SALT00000010172_Cluster7227-001 | 1.000 | TCONS_00077817\| | 2,135 |

| | | | |
|---|---|---|---|
| SALT00000083745_Cluster7227-002 | 1.000 | TCONS_00077817\| | 3,485 |
| SALT00000088313_Cluster7227-003 | 1.000 | TCONS_00077817\| | 3,424 |
| SALT00000088478_Cluster7227-004 | 1.000 | TCONS_00077817\| | 3,605 |
| SALT00000056647_Cluster7306-006 | 1.000 | lincRNA266_PMID_24635777\|lincRNA588_PMID_ | 1,578 |
| SALT00000091719_Cluster7321-001 | 0.998 | osa-MIRf10280-npr\|lincRNA153_PMID_24635777\|l | 3,591 |
| SALT00000033861_Cluster7356-001 | 1.000 | lincRNA413_PMID_24635777\|lincRNA616_PMID_ | 3,583 |
| SALT00000087183_Cluster7381-001 | 0.027 | NA | 3,579 |
| SALT00000081424_Cluster7408-005 | 1.000 | pnrd_Gma_chr12.trna15_GlnCTG\|pnrd_Gma_chr11. | 3,574 |
| SALT00000083020_Cluster7438-001 | 0.566 | lincRNA493_PMID_24635777\|NONATHT003839\|si | 3,569 |
| SALT00000033177_Cluster7458-001 | 0.190 | NA | 3,488 |
| SALT00000034012_Cluster7458-002 | 0.182 | NA | 3,566 |
| SALT00000088636_Cluster7460-001 | 0.284 | NA | 3,566 |
| SALT00000072417_Cluster7543-004 | 0.247 | NA | 2,783 |
| SALT00000003917_Cluster7585-001 | 1.000 | hvu-MIR6207\| | 1,130 |
| SALT00000028388_Cluster7585-002 | 1.000 | hvu-MIR6207\| | 1,432 |
| SALT00000084679_Cluster7585-004 | 1.000 | hvu-MIR6207\| | 1,466 |
| SALT00000003583_Cluster7585-005 | 1.000 | hvu-MIR6207\| | 1,707 |
| SALT00000091843_Cluster7611-001 | 0.459 | lincRNA691_PMID_24635777\|lincRNA756_PMID_ | 3,543 |
| SALT00000044443_Cluster7646-001 | 0.170 | NA | 1,760 |
| SALT00000087691_Cluster7655-001 | 0.239 | NA | 3,536 |
| SALT00000003184_Cluster7709-001 | 1.000 | rgl-MIR5141\|lincRNA430_PMID_24635777\| | 2,174 |
| SALT00000003237_Cluster7709-002 | 1.000 | rgl-MIR5141\|lincRNA430_PMID_24635777\| | 1,506 |
| SALT00000006399_Cluster7709-003 | 1.000 | rgl-MIR5141\|lincRNA430_PMID_24635777\| | 2,290 |
| SALT00000006807_Cluster7709-004 | 1.000 | rgl-MIR5141\|lincRNA430_PMID_24635777\| | 2,047 |
| SALT00000008872_Cluster7709-005 | 1.000 | rgl-MIR5141\|lincRNA430_PMID_24635777\| | 1,827 |
| SALT00000014726_Cluster7709-006 | 1.000 | lincRNA430_PMID_24635777\|rgl-MIR5141\| | 1,647 |
| SALT00000027509_Cluster7709-007 | 1.000 | lincRNA430_PMID_24635777\|rgl-MIR5141\| | 2,352 |
| SALT00000041934_Cluster7709-008 | 1.000 | rgl-MIR5141\|lincRNA430_PMID_24635777\| | 1,605 |
| SALT00000045254_Cluster7709-010 | 1.000 | rgl-MIR5141\|lincRNA430_PMID_24635777\| | 2,028 |
| SALT00000060247_Cluster7709-011 | 1.000 | lincRNA430_PMID_24635777\|rgl-MIR5141\| | 1,586 |
| SALT00000087748_Cluster7735-001 | 1.000 | lincRNA11_PMID_24635777\| | 3,523 |
| SALT00000029004_Cluster7784-001 | 0.271 | NA | 3,516 |
| SALT00000048657_Cluster7784-003 | 0.246 | NA | 1,561 |
| SALT00000053950_Cluster7784-004 | 0.248 | NA | 1,549 |
| SALT00000047127_Cluster7784-007 | 0.196 | NA | 1,973 |
| SALT00000091475_Cluster7797-001 | 0.240 | NA | 3,515 |
| SALT00000066071_Cluster7827-001 | 1.000 | lincRNA11_PMID_24635777\| | 3,510 |
| SALT00000092207_Cluster7867-001 | 0.253 | NA | 3,504 |
| SALT00000021384_Cluster7890-003 | 0.999 | lincRNA526_PMID_24635777\|lincRNA334_PMID_ | 2,728 |
| SALT00000083814_Cluster7912-001 | 0.796 | lincRNA623_PMID_24635777\|lincRNA12_PMID_2 | 3,496 |
| SALT00000091983_Cluster7936-001 | 0.113 | NA | 3,492 |
| SALT00000010701_Cluster7948-001 | 0.023 | NA | 1,443 |
| SALT00000046900_Cluster7985-001 | 0.173 | NA | 3,485 |
| SALT00000038662_Cluster8007-006 | 0.249 | NA | 1,534 |
| SALT00000086801_Cluster8010-001 | 0.081 | NA | 3,480 |
| SALT00000034386_Cluster8041-001 | 0.124 | NA | 3,473 |
| SALT00000085040_Cluster8061-002 | 0.183 | NA | 3,469 |
| SALT00000085789_Cluster8091-001 | 0.052 | NA | 3,463 |
| SALT00000045327_Cluster8143-004 | 0.999 | peu-MIR2914\|NONATHT002169\|NONATHT00093C | 1,808 |
| SALT00000042992_Cluster8190-003 | 1.000 | NONATHT003839\|NONATHT003847\| | 1,640 |
| SALT00000090213_Cluster8215-001 | 0.987 | lincRNA616_PMID_24635777\|lincRNA153_PMID_ | 3,441 |
| SALT00000017240_Cluster8227-001 | 0.998 | JNnc_loci0077\| | 1,808 |
| SALT00000032057_Cluster8227-002 | 0.994 | JNnc_loci0077\| | 3,438 |
| SALT00000090426_Cluster8239-001 | 0.080 | NA | 3,436 |
| SALT00000093126_Cluster8246-001 | 0.174 | NA | 3,435 |
| SALT00000065218_Cluster8301-003 | 1.000 | pta-MIR1310\|cln-MIR1310\|pde-MIR1310\|han-MIR1. | 3,424 |
| SALT00000065127_Cluster8315-001 | 0.290 | NA | 2,791 |
| SALT00000087839_Cluster8315-002 | 0.115 | NA | 3,421 |

| | | | |
|---|---|---|---|
| SALT00000040161_Cluster8334-004 | 0.152 | NA | 1,765 |
| SALT00000034088_Cluster8397-001 | 0.040 | NA | 3,406 |
| SALT00000052250_Cluster8412-001 | 0.541 | TCONS_00018605\|sit-MIR122-1-npr\|lincRNA372_P | 1,788 |
| SALT00000067840_Cluster8412-002 | 0.470 | TCONS_00018605\|lincRNA372_PMID_24635777\|si | 3,403 |
| SALT00000078070_Cluster8445-008 | 0.220 | NA | 3,395 |
| SALT00000092033_Cluster8461-001 | 1.000 | lincRNA37_PMID_24635777\| | 3,393 |
| SALT00000069570_Cluster8501-001 | 1.000 | sof-MIR172a\| | 3,385 |
| SALT00000066757_Cluster8586-001 | 0.613 | JNnc_loci0246\|JNnc_loci0427\|JNnc_loci0581\|JNnc_ | 3,367 |
| SALT00000064748_Cluster8591-001 | 0.132 | pnrd_Osa_chr10.trna43_CysGCA\|pnrd_Osa_chr4.trn | 1,655 |
| SALT00000076911_Cluster8591-002 | 0.299 | pnrd_Osa_chr4.trna86_CysGCA\|pnrd_Osa_chr10.trn | 3,366 |
| SALT00000089685_Cluster8639-001 | 0.057 | NA | 3,357 |
| SALT00000065079_Cluster8702-001 | 0.086 | NA | 3,346 |
| SALT00000001318_Cluster8733-001 | 1.000 | lincRNA246_PMID_24635777\|lincRNA302_PMID_ | 1,650 |
| SALT00000003582_Cluster8733-002 | 1.000 | lincRNA246_PMID_24635777\|lincRNA302_PMID_ | 1,483 |
| SALT00000005840_Cluster8733-003 | 1.000 | lincRNA302_PMID_24635777\|lincRNA246_PMID_ | 1,535 |
| SALT00000018302_Cluster8733-004 | 1.000 | lincRNA246_PMID_24635777\|lincRNA302_PMID_ | 1,447 |
| SALT00000020422_Cluster8733-005 | 1.000 | lincRNA246_PMID_24635777\|lincRNA302_PMID_ | 1,486 |
| SALT00000025186_Cluster8733-006 | 1.000 | lincRNA302_PMID_24635777\|lincRNA246_PMID_ | 1,617 |
| SALT00000035279_Cluster8733-007 | 1.000 | lincRNA246_PMID_24635777\|lincRNA302_PMID_ | 1,562 |
| SALT00000035367_Cluster8733-008 | 1.000 | lincRNA246_PMID_24635777\|lincRNA302_PMID_ | 1,565 |
| SALT00000035600_Cluster8733-009 | 1.000 | lincRNA246_PMID_24635777\|lincRNA302_PMID_ | 1,563 |
| SALT00000037480_Cluster8733-011 | 1.000 | lincRNA246_PMID_24635777\|lincRNA302_PMID_ | 1,535 |
| SALT00000038764_Cluster8733-012 | 1.000 | lincRNA302_PMID_24635777\|lincRNA246_PMID_ | 3,340 |
| SALT00000041914_Cluster8733-013 | 1.000 | lincRNA246_PMID_24635777\|lincRNA302_PMID_ | 1,477 |
| SALT00000042715_Cluster8733-014 | 1.000 | lincRNA246_PMID_24635777\|lincRNA302_PMID_ | 1,571 |
| SALT00000045598_Cluster8733-015 | 1.000 | lincRNA302_PMID_24635777\|lincRNA246_PMID_ | 1,502 |
| SALT00000051882_Cluster8733-016 | 1.000 | lincRNA302_PMID_24635777\|lincRNA246_PMID_ | 1,476 |
| SALT00000053412_Cluster8733-017 | 1.000 | lincRNA302_PMID_24635777\|lincRNA246_PMID_ | 1,483 |
| SALT00000055804_Cluster8733-018 | 1.000 | lincRNA246_PMID_24635777\|lincRNA302_PMID_ | 1,536 |
| SALT00000093720_Cluster8733-019 | 1.000 | lincRNA246_PMID_24635777\|lincRNA302_PMID_ | 1,557 |
| SALT00000061201_Cluster8733-020 | 1.000 | lincRNA302_PMID_24635777\|lincRNA246_PMID_ | 1,615 |
| SALT00000059821_Cluster8733-021 | 1.000 | lincRNA302_PMID_24635777\|lincRNA246_PMID_ | 2,146 |
| SALT00000037120_Cluster8733-022 | 1.000 | lincRNA302_PMID_24635777\|lincRNA246_PMID_ | 2,426 |
| SALT00000047353_Cluster8748-001 | 1.000 | cln-MIR1310\|pta-MIR1310\|han-MIR1310\|pde-MIR1 | 3,337 |
| SALT00000080661_Cluster8791-001 | 0.148 | NA | 3,329 |
| SALT00000075718_Cluster8830-001 | 0.272 | NA | 3,320 |
| SALT00000065832_Cluster8840-001 | 0.205 | NA | 3,318 |
| SALT00000082251_Cluster8862-001 | 0.385 | lincRNA225_PMID_24635777\| | 3,313 |
| SALT00000071600_Cluster8902-002 | 1.000 | lincRNA769_PMID_24635777\| | 2,970 |
| SALT00000073074_Cluster8902-003 | 1.000 | lincRNA769_PMID_24635777\| | 2,710 |
| SALT00000078280_Cluster8902-004 | 1.000 | lincRNA769_PMID_24635777\| | 3,306 |
| SALT00000000572_Cluster8905-001 | 1.000 | osa-MIR5075\| | 1,533 |
| SALT00000004346_Cluster8905-002 | 1.000 | osa-MIR5075\| | 1,493 |
| SALT00000006716_Cluster8905-003 | 1.000 | osa-MIR5075\| | 1,624 |
| SALT00000011143_Cluster8905-004 | 1.000 | osa-MIR5075\| | 1,454 |
| SALT00000011263_Cluster8905-005 | 1.000 | osa-MIR5075\| | 1,472 |
| SALT00000014728_Cluster8905-006 | 1.000 | osa-MIR5075\| | 1,575 |
| SALT00000014778_Cluster8905-007 | 1.000 | osa-MIR5075\| | 1,541 |
| SALT00000017738_Cluster8905-008 | 1.000 | osa-MIR5075\| | 1,562 |
| SALT00000036731_Cluster8905-009 | 1.000 | osa-MIR5075\| | 1,627 |
| SALT00000047405_Cluster8905-010 | 1.000 | osa-MIR5075\| | 1,636 |
| SALT00000051469_Cluster8905-011 | 1.000 | osa-MIR5075\| | 3,305 |
| SALT00000059685_Cluster8905-012 | 1.000 | osa-MIR5075\| | 1,542 |
| SALT00000060124_Cluster8905-013 | 1.000 | osa-MIR5075\| | 1,690 |
| SALT00000065612_Cluster8905-014 | 1.000 | osa-MIR5075\| | 1,496 |
| SALT00000035873_Cluster8905-016 | 1.000 | osa-MIR5075\| | 2,994 |
| SALT00000073694_Cluster8951-001 | 0.745 | lincRNA11_PMID_24635777\| | 3,296 |
| SALT00000089950_Cluster8954-001 | 0.265 | NA | 3,296 |

| | | | |
|---|---|---|---|
| SALT00000087545_Cluster8991-002 | 0.785 | pnrd_Gma_chr18.trna29_LysTTT\|pnrd_Osa_chr6.trn | 3,289 |
| SALT00000007616_Cluster9022-002 | 1.000 | tae-MIR081a_npr\| | 1,588 |
| SALT00000073539_Cluster9024-001 | 0.985 | hvu-MIR6182\| | 3,282 |
| SALT00000066957_Cluster9041-001 | 0.272 | NA | 3,279 |
| SALT00000022729_Cluster9067-001 | 0.998 | lincRNA483_PMID_24635777\|lincRNA444_PMID_ | 2,759 |
| SALT00000074599_Cluster9067-002 | 1.000 | lincRNA483_PMID_24635777\|lincRNA444_PMID_ | 3,274 |
| SALT00000087780_Cluster9091-015 | 1.000 | lincRNA616_PMID_24635777\| | 3,269 |
| SALT00000091401_Cluster9157-001 | 0.114 | NA | 3,259 |
| SALT00000065257_Cluster9181-001 | 1.000 | lincRNA295_PMID_24635777\| | 3,255 |
| SALT00000007999_Cluster9273-001 | 1.000 | osa-MIR5075\| | 1,491 |
| SALT00000017722_Cluster9273-002 | 1.000 | osa-MIR5075\| | 1,565 |
| SALT00000017764_Cluster9273-003 | 1.000 | osa-MIR5075\| | 1,597 |
| SALT00000019082_Cluster9273-004 | 1.000 | osa-MIR5075\| | 1,465 |
| SALT00000035479_Cluster9273-005 | 1.000 | osa-MIR5075\| | 1,457 |
| SALT00000087430_Cluster9273-006 | 1.000 | osa-MIR5075\| | 3,237 |
| SALT00000044397_Cluster9291-001 | 0.048 | NA | 1,506 |
| SALT00000002969_Cluster9297-002 | 0.276 | NA | 1,763 |
| SALT00000059851_Cluster9297-003 | 0.296 | NA | 1,627 |
| SALT00000071305_Cluster9327-001 | 0.249 | NA | 3,223 |
| SALT00000052620_Cluster9346-001 | 0.999 | lincRNA11_PMID_24635777\| | 3,218 |
| SALT00000023867_Cluster9489-001 | 0.178 | NA | 2,897 |
| SALT00000062281_Cluster9489-002 | 0.149 | NA | 3,194 |
| SALT00000089306_Cluster9489-003 | 0.153 | NA | 3,145 |
| SALT00000064249_Cluster9495-004 | 0.159 | NA | 3,051 |
| SALT00000018954_Cluster9506-001 | 0.133 | NA | 1,752 |
| SALT00000026070_Cluster9509-001 | 1.000 | TCONS_00063431\| | 3,190 |
| SALT00000081276_Cluster9509-002 | 1.000 | TCONS_00063431\| | 3,174 |
| SALT00000038558_Cluster9539-001 | 0.202 | NA | 2,228 |
| SALT00000081344_Cluster9576-001 | 0.217 | NA | 3,177 |
| SALT00000010481_Cluster9669-001 | 0.206 | han-MIR1310\|pde-MIR1310\|pta-MIR1310\|cln-MIR1 | 1,608 |
| SALT00000031267_Cluster9669-002 | 0.105 | ghr-MIR4370\| | 3,162 |
| SALT00000063528_Cluster9669-003 | 0.155 | NA | 1,609 |
| SALT00000045515_Cluster9670-001 | 0.991 | NONATHT002169\|peu-MIR2910\|NONATHT00093C | 3,162 |
| SALT00000075093_Cluster9673-003 | 1.000 | pnrd_Mtr_chr1.trna73_LeuAAG\| | 2,576 |
| SALT00000063764_Cluster9699-001 | 0.025 | NA | 3,157 |
| SALT00000076472_Cluster9760-001 | 0.205 | NA | 3,147 |
| SALT00000084036_Cluster9833-001 | 0.995 | lincRNA254_PMID_24635777\|lincRNA649_PMID_ | 3,135 |
| SALT00000088512_Cluster9863-001 | 1.000 | lincRNA11_PMID_24635777\|hvu-MIR6183\| | 3,130 |
| SALT00000082827_Cluster9909-005 | 0.196 | NA | 2,593 |
| SALT00000091855_Cluster9946-003 | 0.260 | NA | 3,119 |
| SALT00000072394_Cluster9947-001 | 1.000 | osa-MIR6253\|JNnc_loci0896\| | 3,118 |
| SALT00000070415_Cluster10066-001 | 0.918 | GRMZM2G150286_T04\| | 3,099 |
| SALT00000063772_Cluster10104-001 | 0.164 | NA | 2,935 |
| SALT00000084029_Cluster10104-003 | 0.191 | NA | 2,693 |
| SALT00000075191_Cluster10105-001 | 0.989 | TCONS_00018605\|NONATHT003839\|NONATHT0( | 3,094 |
| SALT00000006400_Cluster10108-005 | 0.113 | NA | 1,481 |
| SALT00000076106_Cluster10153-001 | 1.000 | osa-MIRf12017-npr\| | 3,088 |
| SALT00000029035_Cluster10182-001 | 0.076 | NA | 3,083 |
| SALT00000051916_Cluster10182-002 | 0.183 | NA | 1,680 |
| SALT00000075959_Cluster10182-003 | 0.099 | NA | 2,754 |
| SALT00000061793_Cluster10182-004 | 0.201 | NA | 1,515 |
| SALT00000064252_Cluster10182-005 | 0.115 | NA | 2,444 |
| SALT00000074038_Cluster10234-002 | 0.297 | NA | 2,117 |
| SALT00000083746_Cluster10257-002 | 0.257 | NA | 3,072 |
| SALT00000053596_Cluster10330-001 | 1.000 | lincRNA147_PMID_24635777\| | 1,614 |
| SALT00000065086_Cluster10330-002 | 1.000 | lincRNA147_PMID_24635777\| | 3,060 |
| SALT00000030884_Cluster10349-002 | 1.000 | far-MIR166\| | 2,948 |
| SALT00000031047_Cluster10349-003 | 1.000 | far-MIR166\| | 3,010 |

| | | | |
|---|---|---|---|
| SALT00000067422_Cluster10349-004 | 1.000 | far-MIR166\| | 2,856 |
| SALT00000072256_Cluster10349-005 | 1.000 | far-MIR166\| | 2,977 |
| SALT00000073675_Cluster10349-006 | 1.000 | far-MIR166\| | 2,985 |
| SALT00000074300_Cluster10349-007 | 1.000 | far-MIR166\| | 3,058 |
| SALT00000076026_Cluster10349-008 | 1.000 | far-MIR166\| | 3,005 |
| SALT00000078539_Cluster10349-009 | 1.000 | far-MIR166\| | 2,467 |
| SALT00000081571_Cluster10360-001 | 1.000 | tae-MIR172a_1_npr\| | 3,056 |
| SALT00000083654_Cluster10381-002 | 0.253 | NA | 3,053 |
| SALT00000024325_Cluster10420-001 | 0.242 | NA | 3,048 |
| SALT00000041284_Cluster10445-001 | 1.000 | lincRNA613_PMID_24635777\| | 1,491 |
| SALT00000052235_Cluster10445-002 | 1.000 | lincRNA613_PMID_24635777\| | 1,629 |
| SALT00000067174_Cluster10445-003 | 1.000 | lincRNA613_PMID_24635777\| | 3,046 |
| SALT00000082333_Cluster10445-004 | 1.000 | lincRNA613_PMID_24635777\| | 3,006 |
| SALT00000068507_Cluster10492-004 | 1.000 | lincRNA482_PMID_24635777\|lincRNA442_PMID_ | 3,039 |
| SALT00000069468_Cluster10586-005 | 1.000 | TCONS_00056274\| | 2,820 |
| SALT00000072766_Cluster10586-006 | 1.000 | TCONS_00056274\| | 2,659 |
| SALT00000022228_Cluster10586-008 | 1.000 | TCONS_00056274\| | 2,993 |
| SALT00000076184_Cluster10603-001 | 0.059 | NA | 3,022 |
| SALT00000053233_Cluster10611-013 | 0.022 | NA | 555 |
| SALT00000062185_Cluster10619-001 | 0.781 | zma-MIR444b\|ssp-MIR444c\|bdi-MIR444d\|bdi-MIR4 | 3,019 |
| SALT00000072219_Cluster10619-002 | 0.786 | zma-MIR444a\|osa-MIR444b\|pvi-MIR444\|bdi-MIR44 | 2,983 |
| SALT00000058584_Cluster10657-004 | 0.123 | NA | 1,493 |
| SALT00000030675_Cluster10686-001 | 0.853 | lincRNA301_PMID_24635777\|lincRNA634_PMID_ | 3,011 |
| SALT00000000836_Cluster10734-001 | 0.802 | pnrd_Ara_chr2.trna8_TyrGTA\|lincRNA11_PMID_24 | 1,666 |
| SALT00000027597_Cluster10734-002 | 0.761 | pnrd_Pop_scaffold_8.trna16_TyrGTA\|pnrd_Mtr_chr1 | 3,005 |
| SALT00000064064_Cluster10761-001 | 0.127 | NA | 3,002 |
| SALT00000063123_Cluster10777-001 | 0.212 | NA | 3,000 |
| SALT00000068407_Cluster10780-001 | 0.291 | NA | 3,000 |
| SALT00000064808_Cluster10791-006 | 1.000 | lincRNA65_PMID_24635777\|lincRNA76_PMID_24 | 1,637 |
| SALT00000021790_Cluster10804-001 | 1.000 | lincRNA769_PMID_24635777\| | 2,997 |
| SALT00000073098_Cluster10811-001 | 1.000 | lincRNA769_PMID_24635777\| | 2,996 |
| SALT00000077282_Cluster10840-001 | 0.239 | NA | 2,993 |
| SALT00000072308_Cluster10858-001 | 0.200 | NA | 2,991 |
| SALT00000072966_Cluster10865-001 | 0.326 | ghr-MIR4370\| | 2,990 |
| SALT00000050112_Cluster10893-001 | 0.133 | NONATHT003839\|TCONS_00018605\|NONATHT0( | 1,855 |
| SALT00000063644_Cluster10893-003 | 0.114 | TCONS_00018605\|NONATHT003839\|NONATHT0( | 2,409 |
| SALT00000063219_Cluster10984-001 | 0.241 | NA | 2,973 |
| SALT00000064117_Cluster10984-002 | 0.277 | NA | 2,884 |
| SALT00000052979_Cluster10995-001 | 1.000 | lincRNA11_PMID_24635777\| | 1,753 |
| SALT00000079465_Cluster10995-002 | 1.000 | lincRNA11_PMID_24635777\| | 2,972 |
| SALT00000055182_Cluster11107-001 | 1.000 | ptr-MIR156\|pts-MIR156b\|pts-MIR157\| | 2,150 |
| SALT00000063352_Cluster11107-002 | 1.000 | pts-MIR157\|ptr-MIR156\|pts-MIR156b\| | 2,406 |
| SALT00000075451_Cluster11107-003 | 1.000 | pts-MIR156b\|ptr-MIR156\|pts-MIR157\| | 2,959 |
| SALT00000028119_Cluster11150-001 | 0.128 | NA | 2,951 |
| SALT00000063748_Cluster11209-001 | 0.244 | osa-MIR444a\|osa-MIR444e\| | 2,845 |
| SALT00000068962_Cluster11209-002 | 0.231 | osa-MIR444e\|osa-MIR444a\| | 2,945 |
| SALT00000073412_Cluster11213-001 | 0.118 | NA | 2,945 |
| SALT00000069528_Cluster11218-001 | 0.243 | NA | 2,944 |
| SALT00000045310_Cluster11220-001 | 0.125 | NA | 1,661 |
| SALT00000064875_Cluster11247-001 | 0.999 | pnrd_Pop_scaffold_1.trna11_ThrTGT\|pnrd_Osa_chr3 | 2,941 |
| SALT00000067758_Cluster11265-001 | 0.117 | NA | 2,939 |
| SALT00000021541_Cluster11313-001 | 1.000 | TCONS_00063431\| | 2,838 |
| SALT00000023756_Cluster11313-002 | 1.000 | TCONS_00063431\| | 2,933 |
| SALT00000068528_Cluster11365-001 | 0.104 | NA | 2,928 |
| SALT00000020716_Cluster11414-001 | 1.000 | lincRNA369_PMID_24635777\| | 2,784 |
| SALT00000021678_Cluster11414-002 | 1.000 | lincRNA369_PMID_24635777\| | 2,796 |
| SALT00000024860_Cluster11414-003 | 1.000 | lincRNA369_PMID_24635777\| | 2,922 |
| SALT00000027872_Cluster11414-004 | 1.000 | lincRNA369_PMID_24635777\| | 2,911 |

| | | | |
|---|---|---|---|
| SALT00000028358_Cluster11414-005 | 1.000 | lincRNA369_PMID_24635777\| | 2,744 |
| SALT00000028426_Cluster11414-006 | 1.000 | lincRNA369_PMID_24635777\| | 2,753 |
| SALT00000031644_Cluster11414-007 | 1.000 | lincRNA369_PMID_24635777\| | 2,819 |
| SALT00000062809_Cluster11414-008 | 1.000 | lincRNA369_PMID_24635777\| | 2,656 |
| SALT00000063639_Cluster11414-009 | 1.000 | lincRNA369_PMID_24635777\| | 2,854 |
| SALT00000065372_Cluster11414-010 | 1.000 | lincRNA369_PMID_24635777\| | 2,640 |
| SALT00000070560_Cluster11414-011 | 1.000 | lincRNA369_PMID_24635777\| | 2,770 |
| SALT00000073288_Cluster11414-012 | 1.000 | lincRNA369_PMID_24635777\| | 2,647 |
| SALT00000062518_Cluster11429-001 | 1.000 | osa-MIRf10144-npr\|osa-MIRf10132-npr\| | 2,921 |
| SALT00000038067_Cluster11553-001 | 0.229 | NA | 2,046 |
| SALT00000062406_Cluster11553-002 | 0.185 | NA | 2,739 |
| SALT00000065551_Cluster11553-003 | 0.167 | NA | 2,909 |
| SALT00000044361_Cluster11562-005 | 1.000 | ghr-MIR4370\| | 2,908 |
| SALT00000019989_Cluster11583-001 | 0.399 | lincRNA541_PMID_24635777\| | 2,120 |
| SALT00000057305_Cluster11583-002 | 0.157 | NA | 1,957 |
| SALT00000069176_Cluster11583-003 | 0.292 | NA | 2,906 |
| SALT00000074903_Cluster11583-004 | 0.087 | lincRNA541_PMID_24635777\| | 2,906 |
| SALT00000075810_Cluster11583-005 | 0.564 | lincRNA541_PMID_24635777\| | 2,802 |
| SALT00000084016_Cluster11583-006 | 0.092 | lincRNA541_PMID_24635777\| | 2,819 |
| SALT00000079810_Cluster11585-001 | 0.258 | NA | 2,906 |
| SALT00000063820_Cluster11592-001 | 0.218 | NA | 2,905 |
| SALT00000069562_Cluster11594-001 | 0.037 | NA | 2,905 |
| SALT00000053615_Cluster11607-001 | 0.157 | NONATHT003847\|NONATHT003839\|TCONS_0001 | 1,915 |
| SALT00000068467_Cluster11607-002 | 0.672 | NONATHT003847\|lincRNA12_PMID_24635777\|NC | 2,695 |
| SALT00000080524_Cluster11607-003 | 0.695 | TCONS_00018605\|NONATHT003839\|lincRNA493_ | 2,717 |
| SALT00000081958_Cluster11607-004 | 0.645 | NONATHT003839\|lincRNA493_PMID_24635777\|T | 2,904 |
| SALT00000075296_Cluster11659-001 | 0.258 | NA | 2,898 |
| SALT00000074993_Cluster11676-001 | 1.000 | lincRNA304_PMID_24635777\| | 2,896 |
| SALT00000076525_Cluster11676-002 | 1.000 | lincRNA304_PMID_24635777\| | 2,888 |
| SALT00000071554_Cluster11687-001 | 0.240 | NA | 2,895 |
| SALT00000074185_Cluster11687-002 | 0.241 | NA | 2,888 |
| SALT00000057547_Cluster11727-003 | 0.966 | TCONS_00044268\| | 2,041 |
| SALT00000026071_Cluster11742-001 | 1.000 | lincRNA298_PMID_24635777\| | 2,647 |
| SALT00000027473_Cluster11742-002 | 1.000 | lincRNA298_PMID_24635777\| | 2,557 |
| SALT00000062024_Cluster11742-003 | 1.000 | lincRNA298_PMID_24635777\| | 2,734 |
| SALT00000064368_Cluster11742-004 | 1.000 | lincRNA298_PMID_24635777\| | 2,577 |
| SALT00000065932_Cluster11742-005 | 1.000 | lincRNA298_PMID_24635777\| | 2,558 |
| SALT00000066335_Cluster11742-006 | 1.000 | lincRNA298_PMID_24635777\| | 2,523 |
| SALT00000074181_Cluster11742-008 | 1.000 | lincRNA298_PMID_24635777\| | 2,389 |
| SALT00000079737_Cluster11742-009 | 1.000 | lincRNA298_PMID_24635777\| | 2,833 |
| SALT00000080923_Cluster11742-010 | 1.000 | lincRNA298_PMID_24635777\| | 2,891 |
| SALT00000083981_Cluster11742-011 | 1.000 | lincRNA298_PMID_24635777\| | 2,646 |
| SALT00000064576_Cluster11757-002 | 0.731 | osa-MIR444e\|osa-MIR444a\| | 2,889 |
| SALT00000028793_Cluster11783-001 | 0.247 | NA | 2,887 |
| SALT00000045824_Cluster11786-001 | 0.991 | NONATHT003839\|ghr-MIR5368b\|ghr-MIR5368*\|sit | 2,887 |
| SALT00000030724_Cluster11792-002 | 0.089 | NA | 2,886 |
| SALT00000071271_Cluster11812-007 | 1.000 | pnrd_Gma_chr12.trna36_ArgTCG\|pnrd_Mtr_chr2.trr | 2,885 |
| SALT00000014195_Cluster11827-001 | 1.000 | lincRNA1_PMID_24635777\| | 2,178 |
| SALT00000078937_Cluster11827-002 | 1.000 | lincRNA1_PMID_24635777\| | 2,884 |
| SALT00000080794_Cluster11827-003 | 1.000 | lincRNA1_PMID_24635777\| | 2,702 |
| SALT00000077244_Cluster11915-014 | 1.000 | osa-MIR6246\| | 2,751 |
| SALT00000022903_Cluster11937-001 | 1.000 | pta-MIR166c\|cln-MIR166\|far-MIR166\| | 2,748 |
| SALT00000065644_Cluster11937-002 | 1.000 | far-MIR166\|cln-MIR166\|pta-MIR166c\| | 2,874 |
| SALT00000063610_Cluster12048-001 | 0.273 | NA | 2,739 |
| SALT00000071080_Cluster12071-001 | 0.249 | NA | 2,861 |
| SALT00000069011_Cluster12085-001 | 0.162 | NA | 2,859 |
| SALT00000074442_Cluster12124-004 | 0.999 | osa-MIRf10211-npr\| | 2,856 |
| SALT00000016624_Cluster12149-001 | 0.012 | NA | 1,626 |

| | | | |
|---|---|---|---|
| SALT00000027186_Cluster12149-002 | 0.078 | NONATHT002123\| | 2,853 |
| SALT00000040796_Cluster12149-003 | 0.047 | NA | 1,959 |
| SALT00000043849_Cluster12149-004 | 0.058 | NA | 1,641 |
| SALT00000055221_Cluster12149-005 | 0.076 | NA | 2,116 |
| SALT00000067143_Cluster12187-004 | 0.080 | NA | 2,587 |
| SALT00000084051_Cluster12261-001 | 0.159 | NA | 2,843 |
| SALT00000022733_Cluster12287-001 | 0.066 | NA | 2,553 |
| SALT00000064995_Cluster12287-002 | 0.054 | NA | 2,839 |
| SALT00000070097_Cluster12287-003 | 0.027 | NA | 2,703 |
| SALT00000009291_Cluster12296-001 | 1.000 | lincRNA613_PMID_24635777\| | 1,809 |
| SALT00000011864_Cluster12296-002 | 1.000 | lincRNA613_PMID_24635777\| | 1,575 |
| SALT00000024539_Cluster12296-003 | 1.000 | lincRNA613_PMID_24635777\| | 2,838 |
| SALT00000063843_Cluster12296-004 | 0.997 | lincRNA613_PMID_24635777\| | 2,772 |
| SALT00000072535_Cluster12308-001 | 0.283 | NA | 2,838 |
| SALT00000072080_Cluster12330-001 | 0.106 | NA | 2,836 |
| SALT00000030074_Cluster12359-001 | 0.163 | osa-MIR5523\| | 2,833 |
| SALT00000070955_Cluster12382-001 | 0.134 | NA | 2,831 |
| SALT00000081721_Cluster12393-001 | 0.596 | lincRNA503_PMID_24635777\| | 2,830 |
| SALT00000020968_Cluster12401-002 | 1.000 | GRMZM2G125392_T01\| | 2,728 |
| SALT00000021148_Cluster12401-003 | 1.000 | GRMZM2G125392_T01\| | 2,722 |
| SALT00000022973_Cluster12401-004 | 1.000 | GRMZM2G125392_T01\| | 2,653 |
| SALT00000023221_Cluster12401-005 | 1.000 | GRMZM2G125392_T01\| | 2,470 |
| SALT00000072870_Cluster12401-013 | 1.000 | GRMZM2G125392_T01\| | 2,663 |
| SALT00000036926_Cluster12458-001 | 1.000 | lincRNA688_PMID_24635777\| | 2,823 |
| SALT00000002807_Cluster12461-001 | 1.000 | bdi-MIR171a\| | 1,876 |
| SALT00000021369_Cluster12461-002 | 1.000 | bdi-MIR171a\| | 2,703 |
| SALT00000022110_Cluster12461-003 | 1.000 | bdi-MIR171a\| | 2,635 |
| SALT00000023213_Cluster12461-004 | 1.000 | bdi-MIR171a\| | 2,643 |
| SALT00000024560_Cluster12461-005 | 1.000 | bdi-MIR171a\| | 2,540 |
| SALT00000060475_Cluster12461-006 | 1.000 | bdi-MIR171a\| | 1,887 |
| SALT00000064776_Cluster12461-007 | 1.000 | bdi-MIR171a\| | 2,823 |
| SALT00000065347_Cluster12461-008 | 1.000 | bdi-MIR171a\| | 2,680 |
| SALT00000070091_Cluster12461-009 | 1.000 | bdi-MIR171a\| | 2,655 |
| SALT00000077427_Cluster12461-010 | 1.000 | bdi-MIR171a\| | 2,573 |
| SALT00000083403_Cluster12461-011 | 1.000 | bdi-MIR171a\| | 2,525 |
| SALT00000063531_Cluster12472-001 | 0.103 | NA | 2,536 |
| SALT00000068259_Cluster12472-002 | 0.086 | NA | 2,822 |
| SALT00000074402_Cluster12560-001 | 1.000 | lincRNA372_PMID_24635777\| | 2,814 |
| SALT00000062743_Cluster12608-001 | 0.135 | NA | 2,809 |
| SALT00000070894_Cluster12640-001 | 1.000 | lincRNA304_PMID_24635777\| | 2,807 |
| SALT00000000402_Cluster12647-003 | 1.000 | egu-MIR172d\| | 1,580 |
| SALT00000007244_Cluster12647-013 | 1.000 | egu-MIR172d\| | 1,614 |
| SALT00000010211_Cluster12647-015 | 1.000 | egu-MIR172d\| | 1,570 |
| SALT00000012969_Cluster12647-021 | 1.000 | egu-MIR172d\| | 1,467 |
| SALT00000013286_Cluster12647-022 | 1.000 | egu-MIR172d\| | 1,575 |
| SALT00000014832_Cluster12647-025 | 1.000 | egu-MIR172d\| | 1,590 |
| SALT00000020389_Cluster12647-040 | 1.000 | egu-MIR172d\| | 1,653 |
| SALT00000027036_Cluster12647-053 | 1.000 | egu-MIR172d\| | 1,539 |
| SALT00000037867_Cluster12647-065 | 1.000 | egu-MIR172d\| | 1,616 |
| SALT00000041191_Cluster12647-069 | 1.000 | egu-MIR172d\| | 1,631 |
| SALT00000042712_Cluster12647-072 | 1.000 | egu-MIR172d\| | 1,659 |
| SALT00000044807_Cluster12647-076 | 1.000 | egu-MIR172d\| | 1,539 |
| SALT00000052668_Cluster12647-084 | 1.000 | egu-MIR172d\| | 1,557 |
| SALT00000054365_Cluster12647-087 | 1.000 | egu-MIR172d\| | 1,547 |
| SALT00000069269_Cluster12653-001 | 0.220 | NA | 2,806 |
| SALT00000076052_Cluster12690-001 | 1.000 | lincRNA616_PMID_24635777\| | 2,803 |
| SALT00000067662_Cluster12695-001 | 0.128 | NA | 2,802 |
| SALT00000066274_Cluster12802-001 | 1.000 | ttu-MIR160b\|sce-MIR160\| | 2,721 |

| | | | |
|---|---|---|---|
| SALT00000072272_Cluster12802-003 | 1.000 | sce-MIR160\|ttu-MIR160b\| | 2,749 |
| SALT00000079129_Cluster12802-004 | 1.000 | sce-MIR160\|ttu-MIR160b\| | 2,792 |
| SALT00000014125_Cluster12810-001 | 1.000 | lincRNA11_PMID_24635777\| | 1,884 |
| SALT00000029474_Cluster12810-002 | 1.000 | lincRNA11_PMID_24635777\| | 2,509 |
| SALT00000072478_Cluster12810-003 | 1.000 | lincRNA11_PMID_24635777\| | 2,791 |
| SALT00000011235_Cluster12819-001 | 1.000 | NONATHT001141\| | 1,185 |
| SALT00000029097_Cluster12819-002 | 1.000 | lincRNA688_PMID_24635777\|NONATHT001141\| | 2,790 |
| SALT00000025439_Cluster12942-001 | 0.335 | NONATHT003850\|NONATHT003836\|lincRNA688_ | 2,778 |
| SALT00000042158_Cluster12942-002 | 0.195 | NONATHT003850\|NONATHT003836\|lincRNA688_ | 1,592 |
| SALT00000061515_Cluster12942-003 | 0.435 | lincRNA688_PMID_24635777\|NONATHT003850\|N | 1,554 |
| SALT00000074994_Cluster12948-001 | 0.203 | NA | 2,778 |
| SALT00000019129_Cluster12974-001 | 1.000 | lincRNA595_PMID_24635777\| | 1,793 |
| SALT00000041386_Cluster12974-002 | 1.000 | lincRNA595_PMID_24635777\| | 1,967 |
| SALT00000060037_Cluster12974-003 | 1.000 | lincRNA595_PMID_24635777\| | 1,907 |
| SALT00000067517_Cluster12974-004 | 1.000 | lincRNA595_PMID_24635777\| | 2,775 |
| SALT00000070717_Cluster12974-005 | 1.000 | lincRNA595_PMID_24635777\| | 2,559 |
| SALT00000026427_Cluster13012-001 | 1.000 | lincRNA720_PMID_24635777\| | 2,765 |
| SALT00000026443_Cluster13012-002 | 1.000 | lincRNA720_PMID_24635777\| | 2,643 |
| SALT00000029388_Cluster13012-003 | 1.000 | lincRNA720_PMID_24635777\| | 2,487 |
| SALT00000063986_Cluster13012-004 | 1.000 | lincRNA720_PMID_24635777\| | 2,772 |
| SALT00000075052_Cluster13012-005 | 1.000 | lincRNA720_PMID_24635777\| | 2,769 |
| SALT00000076354_Cluster13012-006 | 1.000 | lincRNA720_PMID_24635777\| | 2,703 |
| SALT00000035953_Cluster13012-007 | 1.000 | lincRNA720_PMID_24635777\| | 2,227 |
| SALT00000019723_Cluster13012-008 | 1.000 | lincRNA720_PMID_24635777\| | 1,912 |
| SALT00000003870_Cluster13034-001 | 0.097 | NA | 1,567 |
| SALT00000021982_Cluster13034-002 | 0.111 | NA | 1,508 |
| SALT00000052564_Cluster13034-003 | 0.093 | NA | 1,645 |
| SALT00000000271_Cluster13039-001 | 1.000 | lincRNA613_PMID_24635777\| | 1,896 |
| SALT00000000990_Cluster13039-002 | 1.000 | lincRNA613_PMID_24635777\| | 2,048 |
| SALT00000003282_Cluster13039-003 | 1.000 | lincRNA613_PMID_24635777\| | 1,610 |
| SALT00000003525_Cluster13039-004 | 1.000 | lincRNA613_PMID_24635777\| | 2,008 |
| SALT00000004847_Cluster13039-005 | 1.000 | lincRNA613_PMID_24635777\| | 1,938 |
| SALT00000008267_Cluster13039-006 | 1.000 | lincRNA613_PMID_24635777\| | 1,510 |
| SALT00000018410_Cluster13039-007 | 1.000 | lincRNA613_PMID_24635777\| | 2,371 |
| SALT00000020225_Cluster13039-008 | 1.000 | lincRNA613_PMID_24635777\| | 1,599 |
| SALT00000024238_Cluster13039-009 | 1.000 | lincRNA613_PMID_24635777\| | 1,716 |
| SALT00000061889_Cluster13039-010 | 1.000 | lincRNA613_PMID_24635777\| | 1,906 |
| SALT00000075157_Cluster13039-011 | 1.000 | lincRNA613_PMID_24635777\| | 2,770 |
| SALT00000040992_Cluster13039-012 | 1.000 | lincRNA613_PMID_24635777\| | 2,136 |
| SALT00000080805_Cluster13056-008 | 1.000 | lincRNA493_PMID_24635777\|NONATHT003839\|N | 2,769 |
| SALT00000022566_Cluster13057-001 | 1.000 | lincRNA37_PMID_24635777\| | 2,768 |
| SALT00000057227_Cluster13057-002 | 1.000 | lincRNA37_PMID_24635777\| | 2,207 |
| SALT00000057931_Cluster13057-003 | 1.000 | lincRNA37_PMID_24635777\| | 2,361 |
| SALT00000080307_Cluster13057-004 | 1.000 | lincRNA37_PMID_24635777\| | 2,426 |
| SALT00000076067_Cluster13057-005 | 1.000 | lincRNA37_PMID_24635777\| | 2,857 |
| SALT00000056865_Cluster13068-001 | 1.000 | lincRNA672_PMID_24635777\| | 1,512 |
| SALT00000081505_Cluster13068-002 | 1.000 | lincRNA672_PMID_24635777\| | 2,768 |
| SALT00000069004_Cluster13103-001 | 0.152 | TCONS_00084091\| | 2,765 |
| SALT00000038889_Cluster13104-002 | 0.978 | lincRNA376_PMID_24635777\| | 1,614 |
| SALT00000060293_Cluster13104-003 | 0.990 | lincRNA376_PMID_24635777\| | 1,659 |
| SALT00000065786_Cluster13104-004 | 1.000 | lincRNA376_PMID_24635777\| | 2,430 |
| SALT00000066960_Cluster13104-005 | 1.000 | lincRNA376_PMID_24635777\| | 2,625 |
| SALT00000069232_Cluster13104-006 | 1.000 | lincRNA376_PMID_24635777\| | 2,765 |
| SALT00000070081_Cluster13104-007 | 1.000 | lincRNA376_PMID_24635777\| | 2,470 |
| SALT00000074640_Cluster13162-005 | 1.000 | ghr-MIR4370\| | 2,761 |
| SALT00000072758_Cluster13174-001 | 1.000 | lincRNA613_PMID_24635777\| | 2,760 |
| SALT00000079105_Cluster13174-002 | 1.000 | lincRNA613_PMID_24635777\| | 2,709 |
| SALT00000074706_Cluster13175-001 | 1.000 | NONATHT001131\| | 2,760 |

| | | | |
|---|---|---|---|
| SALT00000047117_Cluster13213-006 | 1.000 | tae-MIR172a_1_npr\| | 2,756 |
| SALT00000079319_Cluster13219-001 | 0.035 | NA | 2,756 |
| SALT00000062594_Cluster13244-001 | 0.292 | NA | 2,753 |
| SALT00000064441_Cluster13346-001 | 1.000 | lincRNA613_PMID_24635777\| | 2,745 |
| SALT00000082038_Cluster13346-002 | 1.000 | lincRNA613_PMID_24635777\| | 2,523 |
| SALT00000066987_Cluster13358-001 | 1.000 | hvu-MIR6196\| | 2,744 |
| SALT00000070959_Cluster13358-002 | 1.000 | hvu-MIR6196\| | 2,606 |
| SALT00000078262_Cluster13358-003 | 1.000 | hvu-MIR6196\| | 2,629 |
| SALT00000075877_Cluster13358-004 | 1.000 | hvu-MIR6196\| | 2,580 |
| SALT00000076047_Cluster13358-005 | 1.000 | hvu-MIR6196\| | 2,781 |
| SALT00000079015_Cluster13364-002 | 0.187 | NA | 2,744 |
| SALT00000082973_Cluster13366-001 | 1.000 | pnrd_Osa_chr10.trna25_HisGTG\|pnrd_Gma_chr3.trn | 2,744 |
| SALT00000021564_Cluster13408-001 | 0.242 | NA | 2,579 |
| SALT00000028056_Cluster13408-003 | 0.140 | NA | 2,727 |
| SALT00000016547_Cluster13421-001 | 0.113 | NA | 1,691 |
| SALT00000045831_Cluster13421-002 | 0.155 | lincRNA493_PMID_24635777\|NONATHT003785\| | 1,677 |
| SALT00000080699_Cluster13421-003 | 0.244 | TCONS_00018605\|NONATHT003839\|NONATHT0( | 2,739 |
| SALT00000080996_Cluster13421-004 | 0.039 | NONATHT003839\|TCONS_00018605\|NONATHT0( | 1,649 |
| SALT00000068063_Cluster13464-001 | 0.196 | NA | 2,735 |
| SALT00000077031_Cluster13481-001 | 0.184 | NA | 2,734 |
| SALT00000083189_Cluster13531-004 | 1.000 | ghr-MIR4370\| | 2,730 |
| SALT00000068463_Cluster13570-001 | 0.719 | NONATHT002169\|NONATHT000930\| | 2,727 |
| SALT00000072691_Cluster13575-002 | 0.238 | NA | 2,543 |
| SALT00000074486_Cluster13575-003 | 0.225 | NA | 2,727 |
| SALT00000076766_Cluster13575-004 | 0.237 | NA | 2,548 |
| SALT00000007753_Cluster13580-001 | 1.000 | lincRNA613_PMID_24635777\| | 1,999 |
| SALT00000027820_Cluster13580-002 | 1.000 | lincRNA613_PMID_24635777\| | 2,726 |
| SALT00000028935_Cluster13580-003 | 1.000 | lincRNA613_PMID_24635777\| | 2,699 |
| SALT00000043396_Cluster13580-004 | 1.000 | lincRNA613_PMID_24635777\| | 1,665 |
| SALT00000075763_Cluster13580-005 | 1.000 | lincRNA613_PMID_24635777\| | 2,692 |
| SALT00000049402_Cluster13580-006 | 1.000 | lincRNA613_PMID_24635777\| | 1,685 |
| SALT00000063641_Cluster13580-007 | 1.000 | lincRNA613_PMID_24635777\| | 2,641 |
| SALT00000072000_Cluster13600-001 | 1.000 | bdi-MIR171a\| | 2,603 |
| SALT00000073295_Cluster13600-002 | 1.000 | bdi-MIR171a\| | 2,725 |
| SALT00000080141_Cluster13600-003 | 1.000 | bdi-MIR171a\| | 2,687 |
| SALT00000082858_Cluster13600-004 | 1.000 | bdi-MIR171a\| | 2,643 |
| SALT00000029212_Cluster13624-001 | 0.226 | NA | 2,723 |
| SALT00000042519_Cluster13671-001 | 0.209 | NA | 2,054 |
| SALT00000070184_Cluster13671-002 | 0.129 | NA | 2,718 |
| SALT00000020002_Cluster13706-001 | 0.097 | NA | 1,701 |
| SALT00000074915_Cluster13807-001 | 1.000 | ace-MIR171a\|ace-MIR170a\| | 2,708 |
| SALT00000024344_Cluster13876-001 | 0.285 | NA | 2,544 |
| SALT00000025869_Cluster13876-002 | 0.273 | NA | 2,634 |
| SALT00000073014_Cluster13876-003 | 0.263 | NA | 2,704 |
| SALT00000067107_Cluster13893-004 | 0.200 | NA | 1,786 |
| SALT00000069559_Cluster13921-001 | 0.944 | NONATHT003839\|NONATHT003847\| | 2,700 |
| SALT00000002200_Cluster13925-001 | 0.063 | NA | 1,890 |
| SALT00000075425_Cluster13925-002 | 0.223 | NA | 2,700 |
| SALT00000067766_Cluster13980-001 | 0.956 | NONATHT001142\| | 2,696 |
| SALT00000000473_Cluster13994-001 | 1.000 | osa-MIR2927\| | 1,677 |
| SALT00000001252_Cluster13994-002 | 1.000 | osa-MIR2927\| | 1,755 |
| SALT00000001479_Cluster13994-003 | 1.000 | osa-MIR2927\| | 1,754 |
| SALT00000010874_Cluster13994-004 | 1.000 | osa-MIR2927\| | 1,761 |
| SALT00000011221_Cluster13994-005 | 1.000 | osa-MIR2927\| | 1,754 |
| SALT00000016782_Cluster13994-006 | 1.000 | osa-MIR2927\| | 1,680 |
| SALT00000017506_Cluster13994-007 | 1.000 | osa-MIR2927\| | 1,873 |
| SALT00000041264_Cluster13994-008 | 1.000 | osa-MIR2927\| | 1,652 |
| SALT00000047599_Cluster13994-009 | 1.000 | osa-MIR2927\| | 1,789 |

| | | | |
|---|---|---|---|
| SALT00000049317_Cluster13994-010 | 1.000 | osa-MIR2927\| | 1,765 |
| SALT00000054913_Cluster13994-011 | 1.000 | osa-MIR2927\| | 1,956 |
| SALT00000059321_Cluster13994-012 | 1.000 | osa-MIR2927\| | 2,013 |
| SALT00000063223_Cluster13994-013 | 1.000 | osa-MIR2927\| | 2,502 |
| SALT00000069488_Cluster13994-014 | 1.000 | osa-MIR2927\| | 2,695 |
| SALT00000074909_Cluster13994-015 | 1.000 | osa-MIR2927\| | 2,492 |
| SALT00000077858_Cluster14000-001 | 0.096 | pnrd_Osa_chr10.trna51_AsnGTT\|pnrd_Gma_scaffol | 2,695 |
| SALT00000023744_Cluster14045-001 | 0.166 | NA | 2,558 |
| SALT00000075088_Cluster14045-002 | 0.119 | NA | 2,692 |
| SALT00000037353_Cluster14076-001 | 1.000 | lincRNA11_PMID_24635777\| | 2,689 |
| SALT00000063386_Cluster14092-001 | 0.151 | NA | 2,626 |
| SALT00000027543_Cluster14115-001 | 0.055 | NA | 2,686 |
| SALT00000037454_Cluster14115-002 | 0.204 | NA | 1,504 |
| SALT00000051907_Cluster14115-003 | 0.117 | NA | 1,515 |
| SALT00000074799_Cluster14146-002 | 1.000 | lincRNA594_PMID_24635777\| | 2,684 |
| SALT00000070686_Cluster14154-001 | 0.815 | lincRNA107_PMID_24635777\|lincRNA98_PMID_2 | 2,612 |
| SALT00000072157_Cluster14154-002 | 0.807 | lincRNA98_PMID_24635777\|lincRNA107_PMID_2 | 2,683 |
| SALT00000073612_Cluster14167-001 | 0.284 | NA | 2,682 |
| SALT00000079590_Cluster14168-001 | 0.108 | NA | 2,682 |
| SALT00000070470_Cluster14178-001 | 0.166 | NA | 2,681 |
| SALT00000081155_Cluster14182-004 | 0.292 | NA | 2,681 |
| SALT00000005705_Cluster14292-001 | 1.000 | NONATHT001141\| | 1,648 |
| SALT00000014983_Cluster14292-002 | 1.000 | NONATHT001141\| | 1,791 |
| SALT00000050221_Cluster14292-004 | 1.000 | rgl-MIR5141\|NONATHT001141\|lincRNA430_PMID | 1,605 |
| SALT00000000074_Cluster14319-001 | 1.000 | egu-MIR172d\| | 1,644 |
| SALT00000000233_Cluster14319-004 | 1.000 | egu-MIR172d\| | 1,682 |
| SALT00000000275_Cluster14319-005 | 1.000 | egu-MIR172d\| | 1,627 |
| SALT00000005380_Cluster14319-008 | 1.000 | egu-MIR172d\| | 1,672 |
| SALT00000008312_Cluster14319-010 | 1.000 | egu-MIR172d\| | 1,647 |
| SALT00000008688_Cluster14319-011 | 1.000 | egu-MIR172d\| | 1,702 |
| SALT00000010243_Cluster14319-012 | 1.000 | egu-MIR172d\| | 1,684 |
| SALT00000014577_Cluster14319-014 | 1.000 | egu-MIR172d\| | 1,648 |
| SALT00000018715_Cluster14319-023 | 1.000 | egu-MIR172d\| | 1,636 |
| SALT00000020506_Cluster14319-025 | 1.000 | egu-MIR172d\| | 1,695 |
| SALT00000021948_Cluster14319-027 | 1.000 | egu-MIR172d\| | 1,724 |
| SALT00000025140_Cluster14319-029 | 1.000 | egu-MIR172d\| | 1,700 |
| SALT00000052709_Cluster14319-038 | 1.000 | egu-MIR172d\| | 1,650 |
| SALT00000056750_Cluster14319-050 | 1.000 | egu-MIR172d\| | 1,920 |
| SALT00000053622_Cluster14319-051 | 1.000 | egu-MIR172d\| | 2,030 |
| SALT00000069006_Cluster14368-001 | 0.876 | lincRNA12_PMID_24635777\|NONATHT003785\|lin | 2,667 |
| SALT00000005138_Cluster14423-001 | 0.211 | NA | 1,685 |
| SALT00000021530_Cluster14458-001 | 1.000 | bdi-MIR5064\| | 2,661 |
| SALT00000068175_Cluster14468-002 | 0.193 | NA | 2,661 |
| SALT00000068793_Cluster14468-003 | 0.204 | NA | 2,564 |
| SALT00000075279_Cluster14518-001 | 1.000 | lincRNA11_PMID_24635777\| | 2,658 |
| SALT00000059163_Cluster14577-001 | 0.166 | NA | 2,654 |
| SALT00000001707_Cluster14605-001 | 1.000 | NONATHT001127\| | 1,788 |
| SALT00000051260_Cluster14605-002 | 1.000 | NONATHT001127\| | 1,646 |
| SALT00000064341_Cluster14605-003 | 1.000 | NONATHT001127\|pnrd_lincRNA2850\| | 2,652 |
| SALT00000063907_Cluster14644-001 | 1.000 | pnrd_lincRNA2558\|lincRNA430_PMID_24635777\| | 2,649 |
| SALT00000028212_Cluster14687-001 | 0.999 | JNnc_loci0375\|JNnc_loci0374\| | 2,646 |
| SALT00000082749_Cluster14705-003 | 0.248 | NA | 2,551 |
| SALT00000065183_Cluster14757-002 | 0.222 | NA | 2,642 |
| SALT00000071711_Cluster14781-001 | 0.218 | NA | 2,641 |
| SALT00000049025_Cluster14841-001 | 1.000 | peu-MIR2916\|NONATHT002169\|NONATHT00093 | 2,637 |
| SALT00000081414_Cluster14844-003 | 0.196 | NA | 2,474 |
| SALT00000081390_Cluster14876-001 | 0.454 | tae-MIR156e_1_npr\|tae-MIR156e_2_npr\| | 2,635 |
| SALT00000080793_Cluster14891-001 | 0.388 | tae-MIR444a\|zma-MIR444a\|osa-MIR444c\|pvi-MIR4 | 2,634 |

| | | | |
|---|---|---|---|
| SALT00000065647_Cluster14899-001 | 0.290 | NA | 2,633 |
| SALT00000067559_Cluster14912-002 | 0.083 | NA | 2,632 |
| SALT00000066609_Cluster14923-001 | 0.076 | NA | 2,631 |
| SALT00000079776_Cluster14941-001 | 0.064 | NA | 2,630 |
| SALT00000076587_Cluster15046-001 | 0.116 | NA | 2,622 |
| SALT00000082064_Cluster15046-002 | 0.138 | NA | 2,335 |
| SALT00000082779_Cluster15090-001 | 0.139 | NA | 2,619 |
| SALT00000067023_Cluster15114-001 | 0.105 | NA | 2,617 |
| SALT00000068869_Cluster15157-001 | 0.094 | NA | 2,614 |
| SALT00000053162_Cluster15180-001 | 1.000 | cln-MIR1310\|pta-MIR1310\|han-MIR1310\|pde-MIR1 | 2,612 |
| SALT00000060769_Cluster15189-005 | 0.990 | JNnc_loci0271\|JNnc_loci0136\|JNnc_loci0132\|JNnc_ | 1,743 |
| SALT00000079396_Cluster15193-001 | 0.249 | NA | 2,612 |
| SALT00000035933_Cluster15194-001 | 0.900 | lincRNA520_PMID_24635777\| | 2,072 |
| SALT00000079516_Cluster15194-002 | 0.859 | lincRNA520_PMID_24635777\| | 2,612 |
| SALT00000022351_Cluster15227-001 | 1.000 | pnrd_Mtr_chr1.trna73_LeuAAG\| | 2,561 |
| SALT00000081338_Cluster15277-001 | 0.204 | NA | 2,607 |
| SALT00000074039_Cluster15282-001 | 0.224 | NA | 2,606 |
| SALT00000067696_Cluster15303-002 | 0.063 | NA | 869 |
| SALT00000039498_Cluster15360-001 | 0.219 | NA | 2,111 |
| SALT00000026645_Cluster15360-003 | 0.169 | NA | 2,585 |
| SALT00000082718_Cluster15365-001 | 0.184 | NA | 2,600 |
| SALT00000059583_Cluster15488-003 | 0.204 | NA | 2,086 |
| SALT00000072040_Cluster15494-002 | 0.219 | NA | 2,590 |
| SALT00000073456_Cluster15513-001 | 0.297 | NA | 2,589 |
| SALT00000015933_Cluster15556-001 | 1.000 | lincRNA595_PMID_24635777\| | 1,836 |
| SALT00000039600_Cluster15556-002 | 1.000 | lincRNA595_PMID_24635777\| | 1,677 |
| SALT00000055567_Cluster15556-003 | 1.000 | lincRNA595_PMID_24635777\| | 1,790 |
| SALT00000081032_Cluster15556-004 | 1.000 | lincRNA595_PMID_24635777\| | 2,585 |
| SALT00000019872_Cluster15556-005 | 1.000 | lincRNA595_PMID_24635777\| | 1,980 |
| SALT00000082362_Cluster15571-001 | 0.106 | NA | 2,584 |
| SALT00000052471_Cluster15659-001 | 1.000 | lincRNA246_PMID_24635777\|lincRNA302_PMID_ | 2,577 |
| SALT00000079305_Cluster15683-013 | 1.000 | tae-MIR170a_npr\|tae-MIR170b_npr\|NONATHT0009 | 2,576 |
| SALT00000019762_Cluster15690-001 | 0.190 | NA | 2,126 |
| SALT00000050180_Cluster15690-002 | 0.188 | NA | 2,143 |
| SALT00000072146_Cluster15690-003 | 0.146 | NA | 2,575 |
| SALT00000067561_Cluster15704-001 | 0.273 | NA | 2,574 |
| SALT00000006800_Cluster15708-001 | 1.000 | lincRNA672_PMID_24635777\| | 2,149 |
| SALT00000011478_Cluster15708-002 | 1.000 | lincRNA672_PMID_24635777\| | 1,995 |
| SALT00000011748_Cluster15708-003 | 1.000 | lincRNA672_PMID_24635777\| | 2,136 |
| SALT00000037915_Cluster15708-004 | 1.000 | lincRNA672_PMID_24635777\| | 1,933 |
| SALT00000038092_Cluster15708-005 | 1.000 | lincRNA672_PMID_24635777\| | 2,062 |
| SALT00000063200_Cluster15708-006 | 1.000 | lincRNA672_PMID_24635777\| | 2,148 |
| SALT00000071690_Cluster15708-007 | 1.000 | lincRNA672_PMID_24635777\| | 2,479 |
| SALT00000077510_Cluster15708-008 | 1.000 | lincRNA672_PMID_24635777\| | 2,434 |
| SALT00000084290_Cluster15708-009 | 1.000 | lincRNA672_PMID_24635777\| | 2,574 |
| SALT00000048968_Cluster15708-010 | 1.000 | lincRNA672_PMID_24635777\| | 1,542 |
| SALT00000050918_Cluster15708-011 | 1.000 | lincRNA672_PMID_24635777\| | 1,600 |
| SALT00000018848_Cluster15728-001 | 1.000 | lincRNA672_PMID_24635777\| | 1,946 |
| SALT00000026904_Cluster15728-002 | 1.000 | lincRNA672_PMID_24635777\| | 2,571 |
| SALT00000048895_Cluster15728-003 | 1.000 | lincRNA672_PMID_24635777\| | 2,305 |
| SALT00000068486_Cluster15728-004 | 1.000 | lincRNA672_PMID_24635777\| | 2,453 |
| SALT00000072826_Cluster15728-005 | 1.000 | lincRNA672_PMID_24635777\| | 2,501 |
| SALT00000024303_Cluster15728-006 | 1.000 | lincRNA672_PMID_24635777\| | 2,539 |
| SALT00000030024_Cluster15774-001 | 0.015 | NA | 2,568 |
| SALT00000065719_Cluster15808-001 | 0.241 | NA | 2,566 |
| SALT00000083751_Cluster15815-001 | 0.088 | NA | 2,566 |
| SALT00000072846_Cluster16062-003 | 1.000 | peu-MIR2914\|peu-MIR2916\|NONATHT002169\|peu- | 2,547 |
| SALT00000081931_Cluster16066-001 | 0.007 | NA | 2,547 |

| | | | |
|---|---|---|---|
| SALT00000048663_Cluster16070-001 | 0.122 | TCONS_00018605\|NONATHT003839\|NONATHT0( | 2,546 |
| SALT00000059573_Cluster16142-001 | 0.222 | NA | 1,545 |
| SALT00000052736_Cluster16183-003 | 0.103 | NA | 1,539 |
| SALT00000030040_Cluster16200-001 | 0.066 | NA | 2,536 |
| SALT00000082081_Cluster16207-001 | 0.011 | zma-MIR444a\|bdi-MIR444c\|zma-MIR444b\|ssp-MIR | 2,536 |
| SALT00000063098_Cluster16229-001 | 0.200 | NA | 2,534 |
| SALT00000022720_Cluster16246-001 | 0.188 | pde-MIR1310\|pta-MIR1310\|ghr-MIR4370\|cln-MIR1 | 2,533 |
| SALT00000025756_Cluster16265-001 | 0.135 | NA | 2,532 |
| SALT00000026313_Cluster16273-001 | 0.942 | gba-MIR156c\|mdm-MIR156k\| | 2,531 |
| SALT00000024870_Cluster16296-001 | 0.154 | NA | 2,530 |
| SALT00000024401_Cluster16371-001 | 1.000 | ace-MIR170a\|ace-MIR171a\| | 2,490 |
| SALT00000070939_Cluster16371-002 | 1.000 | ace-MIR171a\|ace-MIR170a\| | 2,221 |
| SALT00000083792_Cluster16371-003 | 1.000 | ace-MIR170a\|ace-MIR171a\| | 2,526 |
| SALT00000062751_Cluster16375-001 | 0.057 | NA | 2,525 |
| SALT00000065328_Cluster16441-001 | 0.171 | NA | 2,520 |
| SALT00000049523_Cluster16477-001 | 1.000 | lincRNA246_PMID_24635777\|lincRNA302_PMID_ | 2,517 |
| SALT00000073879_Cluster16580-001 | 0.022 | NA | 2,508 |
| SALT00000082152_Cluster16584-001 | 0.155 | NA | 2,508 |
| SALT00000026982_Cluster16622-001 | 1.000 | JNnc_loci0332\| | 2,503 |
| SALT00000063480_Cluster16622-003 | 1.000 | JNnc_loci0332\| | 2,339 |
| SALT00000060912_Cluster16708-005 | 0.185 | NA | 1,787 |
| SALT00000045525_Cluster16708-006 | 0.168 | NA | 1,954 |
| SALT00000073887_Cluster16760-001 | 1.000 | lincRNA246_PMID_24635777\|lincRNA302_PMID_ | 2,494 |
| SALT00000078664_Cluster16795-001 | 1.000 | sof-MIR319c\| | 2,490 |
| SALT00000058600_Cluster16866-001 | 0.996 | ghr-MIR5368*\|NONATHT003847\|lincRNA12_PMII | 2,483 |
| SALT00000025803_Cluster16875-001 | 0.157 | NA | 2,482 |
| SALT00000074804_Cluster16932-001 | 0.953 | sit-MIR95-npr\| | 2,477 |
| SALT00000076378_Cluster16933-001 | 0.057 | NA | 2,477 |
| SALT00000029629_Cluster16953-001 | 0.111 | NA | 2,474 |
| SALT00000022321_Cluster16979-001 | 0.149 | TCONS_00018605\|NONATHT003839\|lincRNA12_F | 2,472 |
| SALT00000021911_Cluster17061-002 | 1.000 | TCONS_00012946\| | 2,464 |
| SALT00000054349_Cluster17083-001 | 1.000 | TCONS_00077884\| | 2,462 |
| SALT00000042749_Cluster17097-015 | 1.000 | NONATHT003785\|lincRNA493_PMID_24635777\| | 2,207 |
| SALT00000039920_Cluster17118-004 | 0.111 | NA | 1,990 |
| SALT00000027113_Cluster17306-001 | 0.968 | GRMZM2G150286_T04\| | 2,436 |
| SALT00000038551_Cluster17323-001 | 0.999 | lincRNA11_PMID_24635777\| | 2,433 |
| SALT00000048641_Cluster17331-005 | 0.232 | NA | 1,604 |
| SALT00000072897_Cluster17360-001 | 0.145 | NA | 2,429 |
| SALT00000056033_Cluster17396-001 | 1.000 | lincRNA688_PMID_24635777\| | 2,423 |
| SALT00000012385_Cluster17427-001 | 0.132 | NA | 1,622 |
| SALT00000022984_Cluster17427-002 | 0.084 | NA | 1,910 |
| SALT00000027689_Cluster17441-001 | 0.268 | NA | 2,416 |
| SALT00000038838_Cluster17463-001 | 1.000 | NONATHT001888\| | 1,973 |
| SALT00000067738_Cluster17497-001 | 0.729 | lincRNA65_PMID_24635777\|lincRNA76_PMID_24 | 2,407 |
| SALT00000030972_Cluster17511-001 | 1.000 | GRMZM5G816602_T01\| | 2,404 |
| SALT00000069685_Cluster17543-001 | 0.911 | lincRNA325_PMID_24635777\| | 2,397 |
| SALT00000001704_Cluster17559-001 | 1.000 | lincRNA720_PMID_24635777\| | 2,190 |
| SALT00000006679_Cluster17559-002 | 1.000 | lincRNA720_PMID_24635777\| | 1,984 |
| SALT00000006915_Cluster17559-003 | 1.000 | lincRNA720_PMID_24635777\| | 2,016 |
| SALT00000009978_Cluster17559-004 | 1.000 | lincRNA720_PMID_24635777\| | 2,231 |
| SALT00000010807_Cluster17559-005 | 1.000 | lincRNA720_PMID_24635777\| | 1,820 |
| SALT00000014766_Cluster17559-006 | 1.000 | lincRNA720_PMID_24635777\| | 2,094 |
| SALT00000019599_Cluster17559-007 | 1.000 | lincRNA720_PMID_24635777\| | 2,152 |
| SALT00000020027_Cluster17559-008 | 1.000 | lincRNA720_PMID_24635777\| | 1,501 |
| SALT00000022817_Cluster17559-009 | 1.000 | lincRNA720_PMID_24635777\| | 2,316 |
| SALT00000024236_Cluster17559-010 | 1.000 | lincRNA720_PMID_24635777\| | 2,393 |
| SALT00000028441_Cluster17559-011 | 1.000 | lincRNA720_PMID_24635777\| | 2,012 |
| SALT00000028590_Cluster17559-012 | 1.000 | lincRNA720_PMID_24635777\| | 2,047 |

| | | | |
|---|---|---|---|
| SALT00000035018_Cluster17559-013 | 1.000 | lincRNA720_PMID_24635777\| | 2,015 |
| SALT00000053237_Cluster17559-014 | 1.000 | lincRNA720_PMID_24635777\| | 2,077 |
| SALT00000059062_Cluster17627-003 | 1.000 | NONATHT003839\|NONATHT003847\|lincRNA12_P | 2,381 |
| SALT00000066783_Cluster17679-002 | 0.280 | NA | 2,371 |
| SALT00000063519_Cluster17684-001 | 0.166 | NA | 2,370 |
| SALT00000008591_Cluster17798-001 | 0.241 | NA | 1,968 |
| SALT00000023313_Cluster17896-002 | 1.000 | lincRNA707_PMID_24635777\| | 2,324 |
| SALT00000074187_Cluster17909-001 | 0.243 | NA | 2,322 |
| SALT00000041831_Cluster17914-001 | 1.000 | lincRNA691_PMID_24635777\|lincRNA756_PMID_ | 1,775 |
| SALT00000063672_Cluster17914-002 | 1.000 | lincRNA256_PMID_24635777\|lincRNA691_PMID_ | 1,783 |
| SALT00000076028_Cluster17914-003 | 1.000 | lincRNA691_PMID_24635777\|lincRNA756_PMID_ | 2,320 |
| SALT00000009786_Cluster17914-004 | 1.000 | lincRNA691_PMID_24635777\|lincRNA756_PMID_ | 1,488 |
| SALT00000059862_Cluster17914-005 | 1.000 | lincRNA256_PMID_24635777\|lincRNA756_PMID_ | 2,174 |
| SALT00000048253_Cluster17917-001 | 0.105 | NA | 2,319 |
| SALT00000060372_Cluster17930-001 | 0.999 | lincRNA372_PMID_24635777\| | 2,317 |
| SALT00000064608_Cluster17948-001 | 0.197 | NA | 2,311 |
| SALT00000001024_Cluster17978-001 | 1.000 | lincRNA672_PMID_24635777\| | 2,125 |
| SALT00000018760_Cluster17978-002 | 1.000 | lincRNA672_PMID_24635777\| | 2,052 |
| SALT00000041287_Cluster17978-003 | 1.000 | lincRNA672_PMID_24635777\| | 2,168 |
| SALT00000047840_Cluster17978-004 | 1.000 | lincRNA672_PMID_24635777\| | 2,075 |
| SALT00000056954_Cluster17978-005 | 1.000 | lincRNA672_PMID_24635777\| | 2,305 |
| SALT00000019726_Cluster17998-001 | 0.128 | NA | 1,899 |
| SALT00000078679_Cluster18019-001 | 0.111 | NA | 2,298 |
| SALT00000060873_Cluster18033-001 | 1.000 | lincRNA688_PMID_24635777\| | 2,296 |
| SALT00000057783_Cluster18157-004 | 0.207 | NA | 1,887 |
| SALT00000082051_Cluster18199-004 | 0.231 | lincRNA151_PMID_24635777\| | 2,222 |
| SALT00000055414_Cluster18281-001 | 0.268 | NA | 2,251 |
| SALT00000060827_Cluster18283-001 | 1.000 | lincRNA179_PMID_24635777\| | 2,251 |
| SALT00000030690_Cluster18292-004 | 1.000 | ghr-MIR4370\| | 2,250 |
| SALT00000018540_Cluster18327-001 | 0.234 | NA | 2,243 |
| SALT00000055176_Cluster18353-001 | 1.000 | pts-MIR157\|ptr-MIR156\|pts-MIR156b\| | 1,753 |
| SALT00000072009_Cluster18353-002 | 1.000 | pts-MIR156b\|ptr-MIR156\|pts-MIR157\| | 2,240 |
| SALT00000079376_Cluster18359-001 | 1.000 | JNnc_loci0735\|JNnc_loci0076\|JNnc_loci0732\|JNnc_ | 2,239 |
| SALT00000041084_Cluster18389-001 | 0.200 | NA | 2,233 |
| SALT00000052615_Cluster18480-001 | 0.906 | lincRNA493_PMID_24635777\|NONATHT003785\| | 2,222 |
| SALT00000039405_Cluster18520-001 | 0.288 | NA | 2,216 |
| SALT00000051546_Cluster18523-001 | 1.000 | sof-MIR319c\| | 2,216 |
| SALT00000079547_Cluster18535-001 | 1.000 | lincRNA25_PMID_24635777\| | 2,215 |
| SALT00000045482_Cluster18678-001 | 0.200 | osa-MIR444d\|ssp-MIR444b\|bdi-MIR444b\|ssp-MIR4 | 2,199 |
| SALT00000055823_Cluster18689-001 | 0.159 | NA | 2,198 |
| SALT00000051579_Cluster18734-001 | 1.000 | pnrd_Osa_chr4.trna23_ThrGGT\|pnrd_Osa_chr4.trna | 2,192 |
| SALT00000055343_Cluster18742-002 | 1.000 | lincRNA634_PMID_24635777\| | 2,191 |
| SALT00000027011_Cluster18742-003 | 0.109 | lincRNA634_PMID_24635777\| | 2,504 |
| SALT00000040273_Cluster18797-001 | 1.000 | lincRNA688_PMID_24635777\| | 2,185 |
| SALT00000060289_Cluster18797-002 | 1.000 | lincRNA688_PMID_24635777\| | 2,133 |
| SALT00000001706_Cluster18833-001 | 1.000 | lincRNA65_PMID_24635777\|lincRNA76_PMID_24 | 2,181 |
| SALT00000019088_Cluster18833-002 | 1.000 | lincRNA76_PMID_24635777\|lincRNA65_PMID_24 | 1,526 |
| SALT00000068236_Cluster18833-003 | 1.000 | lincRNA65_PMID_24635777\|lincRNA76_PMID_24 | 1,796 |
| SALT00000056717_Cluster18856-001 | 0.180 | NA | 2,180 |
| SALT00000042832_Cluster19021-001 | 0.131 | NA | 2,164 |
| SALT00000017806_Cluster19048-001 | 1.000 | osa-MIRf10760-npr\| | 2,108 |
| SALT00000017852_Cluster19048-002 | 1.000 | osa-MIRf10760-npr\| | 2,070 |
| SALT00000051322_Cluster19048-003 | 1.000 | osa-MIRf10760-npr\| | 2,027 |
| SALT00000053442_Cluster19048-004 | 1.000 | osa-MIRf10760-npr\| | 2,162 |
| SALT00000054642_Cluster19048-005 | 1.000 | osa-MIRf10760-npr\| | 1,868 |
| SALT00000056064_Cluster19048-006 | 1.000 | osa-MIRf10760-npr\| | 1,928 |
| SALT00000040490_Cluster19057-002 | 0.149 | NA | 1,488 |
| SALT00000009377_Cluster19067-001 | 1.000 | tae-MIR172a_1_npr\| | 1,627 |

| | | | |
|---|---|---|---|
| SALT00000016313_Cluster19067-002 | 1.000 | tae-MIR172a_1_npr\| | 1,892 |
| SALT00000016980_Cluster19067-003 | 1.000 | tae-MIR172a_1_npr\| | 1,899 |
| SALT00000019051_Cluster19067-004 | 1.000 | tae-MIR172a_1_npr\| | 2,044 |
| SALT00000020188_Cluster19067-005 | 1.000 | tae-MIR172a_1_npr\| | 1,983 |
| SALT00000038737_Cluster19067-006 | 1.000 | tae-MIR172a_1_npr\| | 2,161 |
| SALT00000047763_Cluster19067-007 | 1.000 | tae-MIR172a_1_npr\| | 1,927 |
| SALT00000052554_Cluster19067-008 | 1.000 | tae-MIR172a_1_npr\| | 1,469 |
| SALT00000059471_Cluster19067-009 | 1.000 | tae-MIR172a_1_npr\| | 2,104 |
| SALT00000048252_Cluster19067-010 | 1.000 | tae-MIR172a_1_npr\| | 2,016 |
| SALT00000052932_Cluster19093-001 | 1.000 | osa-MIR6256\| | 2,159 |
| SALT00000069448_Cluster19135-001 | 0.144 | NA | 2,155 |
| SALT00000036868_Cluster19350-001 | 0.048 | NA | 2,133 |
| SALT00000002966_Cluster19370-001 | 1.000 | TCONS_00077884\| | 1,883 |
| SALT00000004707_Cluster19370-002 | 1.000 | TCONS_00077884\| | 1,974 |
| SALT00000012678_Cluster19370-003 | 1.000 | TCONS_00077884\| | 1,861 |
| SALT00000019626_Cluster19370-004 | 1.000 | TCONS_00077884\| | 1,899 |
| SALT00000035920_Cluster19370-005 | 1.000 | TCONS_00077884\| | 1,927 |
| SALT00000051500_Cluster19370-006 | 1.000 | TCONS_00077884\| | 1,818 |
| SALT00000055045_Cluster19370-007 | 1.000 | TCONS_00077884\| | 2,132 |
| SALT00000057750_Cluster19370-008 | 1.000 | TCONS_00077884\| | 1,949 |
| SALT00000015779_Cluster19395-001 | 1.000 | lincRNA541_PMID_24635777\|lincRNA616_PMID_ | 2,128 |
| SALT00000054080_Cluster19443-001 | 0.289 | NA | 2,124 |
| SALT00000046605_Cluster19511-001 | 0.190 | NA | 2,119 |
| SALT00000011033_Cluster19549-001 | 0.297 | NA | 2,116 |
| SALT00000074115_Cluster19558-001 | 0.214 | NA | 2,116 |
| SALT00000012973_Cluster19574-001 | 0.068 | NA | 2,114 |
| SALT00000045304_Cluster19629-001 | 1.000 | lincRNA516_PMID_24635777\| | 1,466 |
| SALT00000059241_Cluster19629-002 | 1.000 | lincRNA516_PMID_24635777\| | 2,110 |
| SALT00000060132_Cluster19629-003 | 1.000 | lincRNA516_PMID_24635777\| | 1,802 |
| SALT00000063554_Cluster19629-004 | 1.000 | lincRNA516_PMID_24635777\| | 1,965 |
| SALT00000040530_Cluster19635-002 | 0.796 | NONATHT003839\|NONATHT003847\| | 1,993 |
| SALT00000037199_Cluster19643-001 | 0.656 | pnrd_Osa_chr2.trna64_GlnTTG\|pnrd_Osa_chr2.trna6 | 2,108 |
| SALT00000066686_Cluster19800-010 | 1.000 | NONATHT003839\|NONATHT003847\| | 1,792 |
| SALT00000061763_Cluster19825-001 | 0.043 | NA | 2,094 |
| SALT00000039916_Cluster19862-001 | 0.377 | NONATHT001126\| | 1,679 |
| SALT00000040959_Cluster19862-002 | 0.811 | NONATHT001126\|lincRNA37_PMID_24635777\| | 2,091 |
| SALT00000000837_Cluster19916-001 | 1.000 | lincRNA310_PMID_24635777\| | 2,086 |
| SALT00000005621_Cluster19916-002 | 1.000 | lincRNA310_PMID_24635777\| | 1,876 |
| SALT00000017445_Cluster19916-003 | 1.000 | lincRNA310_PMID_24635777\| | 1,980 |
| SALT00000038599_Cluster19919-001 | 1.000 | NONATHT000930\|tae-MIR170a_npr\|tae-MIR170b_ı | 2,086 |
| SALT00000042787_Cluster19941-001 | 0.105 | NA | 2,084 |
| SALT00000013291_Cluster19981-003 | 0.249 | NA | 1,977 |
| SALT00000024065_Cluster19993-002 | 0.992 | han-MIR1310\|pde-MIR1310\|cln-MIR1310\|pta-MIR1 | 2,080 |
| SALT00000047479_Cluster20116-001 | 1.000 | pnrd_Osa_chr9.trna37_LeuAAG\|pnrd_Pop_scaffold_ | 2,070 |
| SALT00000018143_Cluster20225-001 | 0.073 | NA | 2,005 |
| SALT00000059479_Cluster20276-001 | 0.282 | NA | 2,057 |
| SALT00000054324_Cluster20349-001 | 0.974 | lincRNA493_PMID_24635777\|NONATHT003785\| | 2,051 |
| SALT00000068029_Cluster20378-001 | 0.087 | NA | 2,049 |
| SALT00000018450_Cluster20383-001 | 0.237 | NA | 2,048 |
| SALT00000006755_Cluster20449-002 | 1.000 | hvu-MIR6177\| | 1,615 |
| SALT00000024561_Cluster20449-005 | 1.000 | hvu-MIR6177\| | 1,659 |
| SALT00000038387_Cluster20449-006 | 1.000 | hvu-MIR6177\| | 1,645 |
| SALT00000040362_Cluster20449-008 | 1.000 | hvu-MIR6177\| | 1,733 |
| SALT00000044940_Cluster20449-009 | 1.000 | hvu-MIR6177\| | 1,718 |
| SALT00000049887_Cluster20458-001 | 1.000 | JNnc_loci0349\| | 2,043 |
| SALT00000005695_Cluster20515-001 | 0.115 | pnrd_Osa_chr4.trna24_GluTTC\|pnrd_Osa_chr12.trna | 2,039 |
| SALT00000008778_Cluster20527-001 | 0.063 | NA | 1,956 |
| SALT00000009099_Cluster20527-002 | 0.068 | NA | 1,841 |

| | | | |
|---|---|---|---|
| SALT00000012198_Cluster20527-003 | 0.071 | NA | 1,791 |
| SALT00000036263_Cluster20527-004 | 0.069 | NA | 1,830 |
| SALT00000044400_Cluster20527-005 | 0.102 | NA | 1,362 |
| SALT00000045730_Cluster20549-001 | 0.577 | TCONS_00011371\| | 2,037 |
| SALT00000019753_Cluster20589-001 | 0.189 | NA | 2,034 |
| SALT00000056679_Cluster20589-002 | 0.195 | NA | 1,986 |
| SALT00000038106_Cluster20591-001 | 0.235 | NA | 2,034 |
| SALT00000018058_Cluster20617-001 | 0.109 | NONATHT003847\|lincRNA493_PMID_24635777\|N | 2,032 |
| SALT00000063275_Cluster20695-001 | 0.939 | lincRNA616_PMID_24635777\| | 2,027 |
| SALT00000023030_Cluster20816-001 | 0.246 | NA | 2,001 |
| SALT00000041428_Cluster20816-002 | 0.244 | NA | 2,018 |
| SALT00000005815_Cluster20848-001 | 1.000 | lincRNA185_PMID_24635777\|lincRNA166_PMID_ | 2,015 |
| SALT00000051891_Cluster20874-001 | 0.336 | NONATHT000930\|peu-MIR2910\|tae-MIR170a_npr\|t | 2,013 |
| SALT00000052706_Cluster20883-013 | 1.000 | lincRNA493_PMID_24635777\| | 1,700 |
| SALT00000013845_Cluster20950-001 | 0.181 | NA | 1,867 |
| SALT00000013854_Cluster20950-002 | 0.167 | NA | 2,008 |
| SALT00000039284_Cluster20984-001 | 0.189 | NA | 2,006 |
| SALT00000017167_Cluster21048-001 | 1.000 | TCONS_00077817\| | 2,001 |
| SALT00000037101_Cluster21062-001 | 1.000 | pts-MIR156c\|sof-MIR156g\| | 1,606 |
| SALT00000040573_Cluster21062-002 | 1.000 | pts-MIR156c\|sof-MIR156g\| | 1,812 |
| SALT00000047800_Cluster21062-003 | 1.000 | pts-MIR156c\|sof-MIR156g\| | 2,000 |
| SALT00000052114_Cluster21062-004 | 1.000 | pts-MIR156c\|sof-MIR156g\| | 1,954 |
| SALT00000003897_Cluster21063-001 | 1.000 | lincRNA618_PMID_24635777\| | 1,929 |
| SALT00000007740_Cluster21063-002 | 1.000 | lincRNA618_PMID_24635777\| | 1,938 |
| SALT00000010112_Cluster21063-003 | 1.000 | lincRNA618_PMID_24635777\| | 1,983 |
| SALT00000013141_Cluster21063-004 | 1.000 | lincRNA618_PMID_24635777\| | 1,932 |
| SALT00000019485_Cluster21063-005 | 1.000 | lincRNA618_PMID_24635777\| | 1,824 |
| SALT00000044287_Cluster21063-006 | 1.000 | lincRNA618_PMID_24635777\| | 1,823 |
| SALT00000051783_Cluster21063-007 | 1.000 | lincRNA618_PMID_24635777\| | 1,584 |
| SALT00000053750_Cluster21063-008 | 1.000 | lincRNA618_PMID_24635777\| | 2,000 |
| SALT00000060701_Cluster21063-009 | 1.000 | lincRNA618_PMID_24635777\| | 1,933 |
| SALT00000013613_Cluster21063-010 | 1.000 | lincRNA618_PMID_24635777\| | 1,927 |
| SALT00000047329_Cluster21063-011 | 1.000 | lincRNA618_PMID_24635777\| | 2,124 |
| SALT00000052492_Cluster21179-001 | 0.057 | NA | 1,993 |
| SALT00000010114_Cluster21183-001 | 1.000 | lincRNA20_PMID_24635777\| | 1,992 |
| SALT00000011556_Cluster21183-002 | 1.000 | lincRNA20_PMID_24635777\| | 1,895 |
| SALT00000059203_Cluster21232-001 | 0.228 | NA | 1,989 |
| SALT00000042150_Cluster21244-001 | 0.010 | NONATHT003847\|lincRNA12_PMID_24635777\|NC | 1,988 |
| SALT00000046667_Cluster21269-001 | 1.000 | egu-MIR172d\| | 1,987 |
| SALT00000059887_Cluster21269-002 | 1.000 | egu-MIR172d\| | 1,850 |
| SALT00000037718_Cluster21307-002 | 1.000 | egu-MIR172d\| | 1,706 |
| SALT00000060713_Cluster21307-003 | 1.000 | egu-MIR172d\| | 1,743 |
| SALT00000000391_Cluster21343-001 | 0.128 | lincRNA11_PMID_24635777\| | 1,981 |
| SALT00000007206_Cluster21343-002 | 0.172 | lincRNA11_PMID_24635777\| | 1,487 |
| SALT00000013026_Cluster21343-003 | 0.141 | lincRNA11_PMID_24635777\| | 1,824 |
| SALT00000049202_Cluster21343-004 | 0.134 | lincRNA11_PMID_24635777\| | 1,899 |
| SALT00000011506_Cluster21350-001 | 0.081 | NA | 1,948 |
| SALT00000036200_Cluster21350-002 | 0.144 | NA | 1,981 |
| SALT00000056292_Cluster21358-001 | 1.000 | osa-MIRf10781-npr\| | 1,981 |
| SALT00000038123_Cluster21387-001 | 0.281 | NA | 1,978 |
| SALT00000001567_Cluster21434-001 | 0.997 | peu-MIR2914\|tae-MIR170b_npr\|tae-MIR170a_npr\|N | 1,974 |
| SALT00000013376_Cluster21434-002 | 0.393 | tae-MIR170b_npr\|tae-MIR170a_npr\|NONATHT0009 | 1,665 |
| SALT00000038892_Cluster21434-003 | 0.179 | peu-MIR2914\|tae-MIR170b_npr\|tae-MIR170a_npr\|N | 1,608 |
| SALT00000006715_Cluster21669-001 | 1.000 | osa-MIR2927\| | 1,945 |
| SALT00000010182_Cluster21669-002 | 1.000 | osa-MIR2927\| | 1,765 |
| SALT00000013821_Cluster21669-003 | 1.000 | osa-MIR2927\| | 1,961 |
| SALT00000051459_Cluster21775-001 | 1.000 | JNnc_loci0732\|JNnc_loci0250\|JNnc_loci0735\|JNnc_ | 1,954 |
| SALT00000049407_Cluster21787-001 | 0.372 | pnrd_Osa_chr2.trna65_GlnTTG\|pnrd_Osa_chr2.trna6 | 1,953 |

| | | | |
|---|---|---|---|
| SALT00000044370_Cluster21801-001 | 0.217 | NA | 1,952 |
| SALT00000053972_Cluster21840-001 | 1.000 | lincRNA688_PMID_24635777\| | 1,950 |
| SALT00000043210_Cluster21854-001 | 0.214 | NA | 1,949 |
| SALT00000017093_Cluster21895-004 | 1.000 | NONATHT000930\|NONATHT002169\| | 1,888 |
| SALT00000043063_Cluster21933-001 | 0.224 | NA | 1,944 |
| SALT00000043166_Cluster21933-002 | 0.209 | NA | 1,727 |
| SALT00000058978_Cluster21937-001 | 1.000 | sof-MIR172a\| | 1,944 |
| SALT00000045521_Cluster21955-001 | 0.112 | NA | 1,942 |
| SALT00000015230_Cluster21982-001 | 1.000 | ptc-MIRf10179-akr\| | 1,940 |
| SALT00000018695_Cluster21982-002 | 1.000 | ptc-MIRf10179-akr\| | 1,880 |
| SALT00000052267_Cluster21982-003 | 1.000 | ptc-MIRf10179-akr\| | 1,674 |
| SALT00000056170_Cluster21982-004 | 1.000 | ptc-MIRf10179-akr\| | 1,626 |
| SALT00000056524_Cluster21982-005 | 1.000 | ptc-MIRf10179-akr\| | 1,800 |
| SALT00000030777_Cluster21984-003 | 1.000 | osa-MIRf10576-npr\| | 1,386 |
| SALT00000000738_Cluster22001-001 | 1.000 | lincRNA172_PMID_24635777\| | 1,853 |
| SALT00000011312_Cluster22001-002 | 1.000 | lincRNA172_PMID_24635777\| | 1,872 |
| SALT00000051210_Cluster22001-003 | 1.000 | lincRNA172_PMID_24635777\| | 1,747 |
| SALT00000051972_Cluster22001-004 | 1.000 | lincRNA172_PMID_24635777\| | 1,939 |
| SALT00000059463_Cluster22001-005 | 1.000 | lincRNA172_PMID_24635777\| | 1,180 |
| SALT00000059859_Cluster22001-006 | 1.000 | lincRNA172_PMID_24635777\| | 1,622 |
| SALT00000003052_Cluster22096-001 | 1.000 | lincRNA65_PMID_24635777\|lincRNA76_PMID_24 | 1,890 |
| SALT00000004249_Cluster22096-002 | 1.000 | lincRNA76_PMID_24635777\|lincRNA65_PMID_24 | 1,817 |
| SALT00000014271_Cluster22096-003 | 1.000 | lincRNA65_PMID_24635777\|lincRNA76_PMID_24 | 1,779 |
| SALT00000037893_Cluster22096-004 | 1.000 | lincRNA65_PMID_24635777\|lincRNA76_PMID_24 | 1,932 |
| SALT00000041644_Cluster22096-005 | 1.000 | lincRNA76_PMID_24635777\|lincRNA65_PMID_24 | 1,797 |
| SALT00000048573_Cluster22096-006 | 1.000 | lincRNA65_PMID_24635777\|lincRNA76_PMID_24 | 1,854 |
| SALT00000049877_Cluster22096-007 | 1.000 | lincRNA65_PMID_24635777\|lincRNA76_PMID_24 | 1,887 |
| SALT00000059670_Cluster22096-008 | 1.000 | lincRNA76_PMID_24635777\|lincRNA65_PMID_24 | 1,713 |
| SALT00000060537_Cluster22096-009 | 1.000 | lincRNA65_PMID_24635777\|lincRNA76_PMID_24 | 1,933 |
| SALT00000061624_Cluster22096-010 | 1.000 | lincRNA76_PMID_24635777\|lincRNA65_PMID_24 | 1,751 |
| SALT00000003941_Cluster22096-011 | 1.000 | lincRNA65_PMID_24635777\|lincRNA76_PMID_24 | 1,884 |
| SALT00000052772_Cluster22108-001 | 0.173 | NA | 1,932 |
| SALT00000036796_Cluster22217-001 | 1.000 | peu-MIR2914\|peu-MIR2916\|tae-MIR170a_npr\|tae-M | 1,925 |
| SALT00000038594_Cluster22218-001 | 0.988 | lincRNA623_PMID_24635777\|NONATHT003785\|li | 1,925 |
| SALT00000054985_Cluster22228-001 | 1.000 | hvu-MIR6188\| | 1,925 |
| SALT00000055242_Cluster22239-001 | 0.993 | lincRNA623_PMID_24635777\|lincRNA12_PMID_2 | 1,924 |
| SALT00000058297_Cluster22268-001 | 0.996 | NONATHT002123\| | 1,922 |
| SALT00000012790_Cluster22271-001 | 0.995 | JNnc_loci0423\|JNnc_loci0422\|JNnc_loci0408\|TCON | 1,921 |
| SALT00000002713_Cluster22321-001 | 0.144 | NA | 1,918 |
| SALT00000043250_Cluster22396-001 | 1.000 | lincRNA672_PMID_24635777\| | 1,914 |
| SALT00000063259_Cluster22400-001 | 0.146 | NA | 1,914 |
| SALT00000051721_Cluster22440-002 | 0.098 | NA | 1,911 |
| SALT00000057510_Cluster22443-001 | 0.058 | NA | 1,911 |
| SALT00000018467_Cluster22496-001 | 1.000 | lincRNA185_PMID_24635777\|lincRNA166_PMID_ | 1,854 |
| SALT00000038638_Cluster22496-002 | 1.000 | lincRNA166_PMID_24635777\|lincRNA185_PMID_ | 1,908 |
| SALT00000042426_Cluster22496-003 | 1.000 | lincRNA185_PMID_24635777\|lincRNA166_PMID_ | 1,719 |
| SALT00000047428_Cluster22501-001 | 0.200 | NA | 1,908 |
| SALT00000053098_Cluster22507-001 | 0.213 | NONATHT003839\|lincRNA493_PMID_24635777\|N | 1,908 |
| SALT00000056987_Cluster22533-001 | 0.160 | NA | 1,907 |
| SALT00000042772_Cluster22579-001 | 0.207 | NA | 1,904 |
| SALT00000016889_Cluster22595-001 | 0.169 | NA | 1,903 |
| SALT00000055669_Cluster22602-001 | 0.186 | NA | 1,903 |
| SALT00000037012_Cluster22709-001 | 0.230 | NA | 1,896 |
| SALT00000008474_Cluster22801-001 | 0.249 | NA | 1,890 |
| SALT00000048561_Cluster22823-001 | 0.182 | NA | 1,889 |
| SALT00000041820_Cluster22839-004 | 1.000 | pnrd_Pop_scaffold_13.trna17_PheGAA\|pnrd_Pop_sc | 1,888 |
| SALT00000012469_Cluster22856-001 | 0.126 | NA | 1,886 |
| SALT00000007935_Cluster22911-001 | 1.000 | bdi-MIR9494\| | 1,848 |

| | | | |
|---|---|---|---|
| SALT00000009578_Cluster22911-002 | 1.000 | bdi-MIR9494\| | 1,883 |
| SALT00000060629_Cluster22911-003 | 1.000 | bdi-MIR9494\| | 1,740 |
| SALT00000016096_Cluster22968-001 | 1.000 | lincRNA1_PMID_24635777\| | 1,838 |
| SALT00000052592_Cluster22968-002 | 1.000 | lincRNA1_PMID_24635777\| | 1,879 |
| SALT00000041779_Cluster23026-005 | 1.000 | lincRNA499_PMID_24635777\| | 1,829 |
| SALT00000003977_Cluster23060-001 | 1.000 | osa-MIR5523\| | 1,527 |
| SALT00000005348_Cluster23060-002 | 1.000 | osa-MIR5523\| | 1,873 |
| SALT00000038986_Cluster23060-003 | 1.000 | osa-MIR5523\| | 1,570 |
| SALT00000045351_Cluster23060-004 | 1.000 | osa-MIR5523\| | 1,491 |
| SALT00000059674_Cluster23060-005 | 1.000 | osa-MIR5523\| | 1,467 |
| SALT00000037204_Cluster23124-001 | 0.142 | NA | 1,869 |
| SALT00000003942_Cluster23205-001 | 0.097 | NA | 1,864 |
| SALT00000007641_Cluster23219-001 | 1.000 | lincRNA377_PMID_24635777\| | 1,725 |
| SALT00000010793_Cluster23219-002 | 1.000 | lincRNA377_PMID_24635777\| | 1,756 |
| SALT00000035221_Cluster23219-003 | 1.000 | lincRNA377_PMID_24635777\| | 1,683 |
| SALT00000049622_Cluster23219-004 | 1.000 | lincRNA377_PMID_24635777\| | 1,594 |
| SALT00000053702_Cluster23219-005 | 1.000 | lincRNA377_PMID_24635777\| | 1,864 |
| SALT00000093880_Cluster23219-006 | 1.000 | lincRNA377_PMID_24635777\| | 1,654 |
| SALT00000009681_Cluster23292-001 | 1.000 | tae-MIR172a_1_npr\| | 1,859 |
| SALT00000001891_Cluster23306-001 | 1.000 | lincRNA88_PMID_24635777\| | 1,526 |
| SALT00000041835_Cluster23306-002 | 1.000 | lincRNA88_PMID_24635777\| | 1,575 |
| SALT00000049484_Cluster23306-003 | 1.000 | lincRNA88_PMID_24635777\| | 1,858 |
| SALT00000060013_Cluster23306-004 | 1.000 | lincRNA88_PMID_24635777\| | 1,430 |
| SALT00000020057_Cluster23342-001 | 0.368 | pnrd_Osa_chr2.trna64_GlnTTG\|pnrd_Osa_chr2.trna6 | 1,856 |
| SALT00000002889_Cluster23415-001 | 1.000 | lincRNA20_PMID_24635777\| | 1,762 |
| SALT00000005064_Cluster23415-002 | 1.000 | lincRNA20_PMID_24635777\| | 1,852 |
| SALT00000039480_Cluster23415-003 | 1.000 | lincRNA20_PMID_24635777\| | 1,830 |
| SALT00000042872_Cluster23415-004 | 1.000 | lincRNA20_PMID_24635777\| | 1,709 |
| SALT00000055091_Cluster23415-005 | 1.000 | lincRNA20_PMID_24635777\| | 1,716 |
| SALT00000059101_Cluster23443-001 | 0.237 | NA | 1,851 |
| SALT00000042017_Cluster23519-001 | 0.991 | osa-MIRf11621-npr\| | 1,845 |
| SALT00000015163_Cluster23533-001 | 0.179 | NA | 1,844 |
| SALT00000003185_Cluster23600-001 | 1.000 | lincRNA750_PMID_24635777\|lincRNA677_PMID_ | 1,840 |
| SALT00000009575_Cluster23600-002 | 1.000 | lincRNA750_PMID_24635777\|lincRNA677_PMID_ | 1,672 |
| SALT00000046313_Cluster23600-003 | 1.000 | lincRNA750_PMID_24635777\|lincRNA677_PMID_ | 1,741 |
| SALT00000060902_Cluster23600-004 | 1.000 | lincRNA677_PMID_24635777\|lincRNA750_PMID_ | 1,627 |
| SALT00000037998_Cluster23651-001 | 0.174 | NA | 1,837 |
| SALT00000044014_Cluster23688-001 | 0.105 | NA | 1,835 |
| SALT00000046429_Cluster23689-001 | 1.000 | lincRNA688_PMID_24635777\| | 1,835 |
| SALT00000018611_Cluster23723-001 | 0.238 | NA | 1,833 |
| SALT00000045503_Cluster23750-001 | 1.000 | osa-MIR396d\| | 1,832 |
| SALT00000048246_Cluster23750-002 | 1.000 | osa-MIR396d\| | 1,567 |
| SALT00000048779_Cluster23753-001 | 0.173 | NA | 1,832 |
| SALT00000052394_Cluster23787-001 | 1.000 | lincRNA688_PMID_24635777\| | 1,830 |
| SALT00000042118_Cluster23795-001 | 1.000 | osa-MIRf11620-npr\| | 1,813 |
| SALT00000057070_Cluster23795-002 | 1.000 | osa-MIRf11620-npr\| | 1,830 |
| SALT00000008647_Cluster23802-001 | 0.284 | NA | 1,829 |
| SALT00000046998_Cluster23820-001 | 0.136 | NA | 1,828 |
| SALT00000017321_Cluster23836-001 | 0.016 | NA | 1,827 |
| SALT00000059756_Cluster23836-002 | 0.011 | NA | 1,807 |
| SALT00000036077_Cluster23837-001 | 1.000 | osa-MIRf10185-npr\| | 1,827 |
| SALT00000042604_Cluster23837-002 | 1.000 | osa-MIRf10185-npr\| | 1,711 |
| SALT00000051731_Cluster23837-003 | 1.000 | osa-MIRf10185-npr\| | 1,744 |
| SALT00000060312_Cluster23837-004 | 1.000 | osa-MIRf10185-npr\| | 1,773 |
| SALT00000067006_Cluster23837-005 | 1.000 | osa-MIRf10185-npr\| | 1,691 |
| SALT00000012045_Cluster23873-001 | 1.000 | lincRNA613_PMID_24635777\| | 1,760 |
| SALT00000016628_Cluster23873-002 | 1.000 | lincRNA613_PMID_24635777\| | 1,822 |
| SALT00000038083_Cluster23873-003 | 1.000 | lincRNA613_PMID_24635777\| | 1,806 |

| | | | |
|---|---|---|---|
| SALT00000060649_Cluster23873-004 | 1.000 | lincRNA613_PMID_24635777\| | 1,826 |
| SALT00000048414_Cluster24087-001 | 0.998 | sit-MIR156b-1\|bdi-MIR156c\|sit-MIR156a-1\|sit-MIR1 | 1,814 |
| SALT00000041088_Cluster24159-001 | 1.000 | lincRNA166_PMID_24635777\|lincRNA185_PMID_ | 1,632 |
| SALT00000041921_Cluster24159-002 | 1.000 | lincRNA185_PMID_24635777\|lincRNA166_PMID_ | 1,810 |
| SALT00000044986_Cluster24159-003 | 1.000 | lincRNA185_PMID_24635777\|lincRNA166_PMID_ | 1,347 |
| SALT00000040635_Cluster24217-001 | 0.018 | osa-MIR444c\|osa-MIR444b\|pvi-MIR444\|bdi-MIR44 | 1,807 |
| SALT00000043841_Cluster24257-001 | 0.178 | NA | 1,805 |
| SALT00000042634_Cluster24290-001 | 1.000 | NONATHT001713\| | 1,803 |
| SALT00000004114_Cluster24461-001 | 0.294 | NA | 1,792 |
| SALT00000005612_Cluster24480-001 | 1.000 | TCONS_00077884\| | 1,747 |
| SALT00000013606_Cluster24480-002 | 1.000 | TCONS_00077884\| | 1,747 |
| SALT00000016106_Cluster24480-003 | 1.000 | TCONS_00077884\| | 1,668 |
| SALT00000023351_Cluster24480-004 | 1.000 | TCONS_00077884\| | 1,791 |
| SALT00000041015_Cluster24535-001 | 0.093 | NA | 1,788 |
| SALT00000036544_Cluster24540-001 | 1.000 | sof-MIR172a\| | 1,444 |
| SALT00000045635_Cluster24540-002 | 1.000 | sof-MIR172a\| | 1,588 |
| SALT00000056550_Cluster24540-003 | 1.000 | sof-MIR172a\| | 1,788 |
| SALT00000057721_Cluster24540-004 | 1.000 | sof-MIR172a\| | 1,538 |
| SALT00000038905_Cluster24540-005 | 1.000 | sof-MIR172a\| | 1,697 |
| SALT00000055326_Cluster24593-005 | 1.000 | NONATHT000930\|NONATHT002169\| | 1,785 |
| SALT00000005542_Cluster24617-001 | 0.260 | pnrd_Osa_chr10.trna63_GlnTTG\|GRMZM2G15966 | 1,783 |
| SALT00000010935_Cluster24617-002 | 0.288 | pnrd_Osa_chr10.trna63_GlnTTG\|GRMZM2G15966 | 1,585 |
| SALT00000059959_Cluster24711-001 | 0.814 | pnrd_Osa_chr2.trna70_GlnTTG\|pnrd_Osa_chr2.trna | 1,779 |
| SALT00000076196_Cluster24754-002 | 0.184 | NA | 1,777 |
| SALT00000041862_Cluster24764-001 | 0.094 | NA | 1,776 |
| SALT00000015031_Cluster24787-001 | 1.000 | NONATHT000919\| | 1,473 |
| SALT00000051349_Cluster24806-001 | 0.187 | NA | 1,774 |
| SALT00000039766_Cluster24827-001 | 0.217 | NA | 1,772 |
| SALT00000000277_Cluster24838-001 | 1.000 | TCONS_00082580\| | 1,771 |
| SALT00000007644_Cluster24838-002 | 1.000 | TCONS_00082580\| | 1,543 |
| SALT00000007920_Cluster24838-003 | 1.000 | TCONS_00082580\| | 1,649 |
| SALT00000012157_Cluster24838-004 | 1.000 | TCONS_00082580\| | 1,533 |
| SALT00000014613_Cluster24838-005 | 1.000 | TCONS_00082580\| | 1,731 |
| SALT00000040849_Cluster24845-002 | 0.116 | NA | 1,771 |
| SALT00000048668_Cluster24908-001 | 0.254 | NA | 1,768 |
| SALT00000053595_Cluster24983-001 | 1.000 | GRMZM2G165119_T01\| | 1,764 |
| SALT00000056077_Cluster24983-002 | 1.000 | GRMZM2G165119_T01\| | 1,680 |
| SALT00000049533_Cluster24983-003 | 1.000 | GRMZM2G165119_T01\| | 1,759 |
| SALT00000059609_Cluster25000-001 | 0.221 | NA | 1,763 |
| SALT00000018812_Cluster25062-001 | 0.183 | NA | 1,759 |
| SALT00000041170_Cluster25087-001 | 0.119 | NA | 1,758 |
| SALT00000060538_Cluster25142-001 | 0.074 | NA | 1,756 |
| SALT00000004182_Cluster25146-001 | 1.000 | lincRNA11_PMID_24635777\| | 1,755 |
| SALT00000037568_Cluster25146-002 | 1.000 | lincRNA11_PMID_24635777\| | 1,504 |
| SALT00000050386_Cluster25146-003 | 1.000 | lincRNA11_PMID_24635777\| | 1,344 |
| SALT00000058541_Cluster25146-004 | 1.000 | lincRNA11_PMID_24635777\| | 1,647 |
| SALT00000036750_Cluster25192-003 | 1.000 | TCONS_00065704\| | 1,617 |
| SALT00000043442_Cluster25192-004 | 1.000 | TCONS_00065704\| | 1,534 |
| SALT00000052266_Cluster25299-001 | 1.000 | lincRNA29_PMID_24635777\| | 1,747 |
| SALT00000056922_Cluster25299-002 | 1.000 | lincRNA29_PMID_24635777\| | 1,384 |
| SALT00000038072_Cluster25382-001 | 0.660 | lincRNA279_PMID_24635777\|lincRNA310_PMID_ | 1,632 |
| SALT00000047062_Cluster25382-002 | 0.642 | lincRNA279_PMID_24635777\|lincRNA310_PMID_ | 1,743 |
| SALT00000045682_Cluster25398-001 | 0.115 | NA | 1,742 |
| SALT00000037954_Cluster25412-001 | 1.000 | lincRNA246_PMID_24635777\|lincRNA302_PMID_ | 1,741 |
| SALT00000052269_Cluster25439-001 | 1.000 | pnrd_Pop_scaffold_3.trna1_ValAAC\| | 1,740 |
| SALT00000057665_Cluster25550-001 | 0.180 | NA | 1,735 |
| SALT00000041396_Cluster25566-001 | 0.132 | NA | 1,734 |
| SALT00000055015_Cluster25638-001 | 0.118 | NA | 1,731 |

| | | | |
|---|---|---|---|
| SALT00000056590_Cluster25664-001 | 0.089 | NA | 1,729 |
| SALT00000054419_Cluster25731-001 | 0.102 | NA | 1,726 |
| SALT00000004277_Cluster25739-001 | 1.000 | GRMZM2G047709_T01\| | 1,546 |
| SALT00000005485_Cluster25739-002 | 1.000 | GRMZM2G047709_T01\| | 1,518 |
| SALT00000007282_Cluster25739-003 | 1.000 | GRMZM2G047709_T01\| | 1,605 |
| SALT00000008053_Cluster25739-004 | 1.000 | GRMZM2G047709_T01\| | 1,528 |
| SALT00000013865_Cluster25739-005 | 1.000 | GRMZM2G047709_T01\| | 1,636 |
| SALT00000017247_Cluster25739-006 | 1.000 | GRMZM2G047709_T01\| | 1,677 |
| SALT00000018112_Cluster25739-007 | 1.000 | GRMZM2G047709_T01\| | 1,509 |
| SALT00000020034_Cluster25739-008 | 1.000 | GRMZM2G047709_T01\| | 1,725 |
| SALT00000047817_Cluster25739-009 | 1.000 | GRMZM2G047709_T01\| | 1,701 |
| SALT00000049470_Cluster25739-010 | 1.000 | GRMZM2G047709_T01\| | 1,464 |
| SALT00000055958_Cluster25739-011 | 1.000 | GRMZM2G047709_T01\| | 1,623 |
| SALT00000061149_Cluster25739-012 | 1.000 | GRMZM2G047709_T01\| | 1,681 |
| SALT00000075280_Cluster25739-013 | 1.000 | GRMZM2G047709_T01\| | 1,266 |
| SALT00000059898_Cluster25739-014 | 1.000 | GRMZM2G047709_T01\| | 1,676 |
| SALT00000010285_Cluster25800-001 | 1.000 | lincRNA376_PMID_24635777\| | 1,722 |
| SALT00000009378_Cluster25808-001 | 1.000 | lincRNA316_PMID_24635777\| | 1,644 |
| SALT00000039041_Cluster25808-002 | 1.000 | lincRNA316_PMID_24635777\| | 1,722 |
| SALT00000041215_Cluster25808-003 | 1.000 | lincRNA316_PMID_24635777\| | 1,443 |
| SALT00000061133_Cluster25808-004 | 1.000 | lincRNA316_PMID_24635777\| | 1,538 |
| SALT00000041118_Cluster25817-001 | 1.000 | sof-MIR172a\| | 1,663 |
| SALT00000055907_Cluster25817-002 | 1.000 | sof-MIR172a\| | 1,722 |
| SALT00000011965_Cluster25817-003 | 1.000 | sof-MIR172a\| | 1,650 |
| SALT00000039592_Cluster25832-001 | 0.123 | NA | 1,721 |
| SALT00000047257_Cluster25904-009 | 0.114 | NA | 664 |
| SALT00000058446_Cluster25950-001 | 0.040 | NA | 1,715 |
| SALT00000048905_Cluster25967-001 | 0.179 | NONATHT003847\|TCONS_00018605\|NONATHT0( | 1,714 |
| SALT00000001611_Cluster26079-001 | 0.250 | NA | 1,682 |
| SALT00000055157_Cluster26079-002 | 0.247 | NA | 1,707 |
| SALT00000060919_Cluster26083-009 | 1.000 | oni-MIR812\| | 1,681 |
| SALT00000038693_Cluster26087-001 | 1.000 | osa-MIR2918\| | 1,706 |
| SALT00000003406_Cluster26097-001 | 1.000 | lincRNA117_PMID_24635777\| | 1,490 |
| SALT00000048284_Cluster26097-003 | 1.000 | lincRNA117_PMID_24635777\| | 1,599 |
| SALT00000018179_Cluster26148-002 | 1.000 | peu-MIR2914\|NONATHT002169\|tae-MIR170a_npr\|t | 1,702 |
| SALT00000037306_Cluster26159-001 | 0.984 | lincRNA56_PMID_24635777\|lincRNA696_PMID_2 | 1,701 |
| SALT00000001156_Cluster26171-001 | 1.000 | lincRNA29_PMID_24635777\| | 1,656 |
| SALT00000020429_Cluster26171-002 | 1.000 | lincRNA29_PMID_24635777\| | 1,700 |
| SALT00000045821_Cluster26195-004 | 1.000 | pnrd_Pop_scaffold_4.trna26_TrpCCA\|pnrd_Gma_chi | 1,699 |
| SALT00000017984_Cluster26199-001 | 1.000 | tae-MIR093a_npr\| | 1,694 |
| SALT00000019910_Cluster26199-002 | 1.000 | tae-MIR093a_npr\| | 1,578 |
| SALT00000051688_Cluster26199-003 | 1.000 | tae-MIR093a_npr\| | 1,699 |
| SALT00000039859_Cluster26199-004 | 1.000 | tae-MIR093a_npr\| | 1,715 |
| SALT00000038070_Cluster26211-001 | 0.978 | NONATHT001128\| | 1,698 |
| SALT00000050970_Cluster26267-001 | 0.255 | NA | 1,696 |
| SALT00000036933_Cluster26276-001 | 0.164 | NA | 1,623 |
| SALT00000062719_Cluster26276-002 | 0.157 | NA | 1,696 |
| SALT00000002554_Cluster26289-001 | 0.240 | NA | 1,694 |
| SALT00000019724_Cluster26312-001 | 0.188 | pnrd_Osa_chr4.trna15_AspGTC\|pnrd_Osa_chr2.trna: | 1,693 |
| SALT00000004434_Cluster26353-001 | 1.000 | NONATHT003839\|ghr-MIR5368b\|TCONS_0001860 | 1,691 |
| SALT00000049216_Cluster26370-001 | 0.242 | NA | 1,691 |
| SALT00000056072_Cluster26489-002 | 0.250 | NA | 1,685 |
| SALT00000058849_Cluster26491-001 | 1.000 | NONATHT001713\| | 1,685 |
| SALT00000059534_Cluster26561-001 | 0.907 | pnrd_Osa_chr2.trna63_GlnTTG\|pnrd_Osa_chr2.trna( | 1,682 |
| SALT00000060069_Cluster26651-001 | 0.117 | NA | 1,678 |
| SALT00000012627_Cluster26688-001 | 0.516 | NONATHT001126\| | 1,675 |
| SALT00000036451_Cluster26770-001 | 1.000 | osa-MIR396d\| | 1,671 |
| SALT00000041507_Cluster26888-002 | 1.000 | hvu-MIR6188\| | 1,658 |

| | | | |
|---|---|---|---|
| SALT00000039130_Cluster26888-003 | 1.000 | hvu-MIR6188\| | 1,285 |
| SALT00000036959_Cluster26888-005 | 1.000 | hvu-MIR6188\| | 1,712 |
| SALT00000069007_Cluster26922-002 | 0.206 | NA | 1,525 |
| SALT00000037624_Cluster26989-001 | 0.256 | TCONS_00063443\| | 1,660 |
| SALT00000012482_Cluster27004-001 | 0.237 | NA | 1,659 |
| SALT00000040360_Cluster27048-001 | 0.236 | NA | 1,657 |
| SALT00000000210_Cluster27052-001 | 1.000 | lincRNA556_PMID_24635777\| | 1,483 |
| SALT00000004205_Cluster27052-002 | 1.000 | lincRNA556_PMID_24635777\| | 1,614 |
| SALT00000013312_Cluster27052-003 | 1.000 | lincRNA556_PMID_24635777\| | 1,570 |
| SALT00000044949_Cluster27052-004 | 1.000 | lincRNA556_PMID_24635777\| | 1,657 |
| SALT00000047098_Cluster27052-005 | 1.000 | lincRNA556_PMID_24635777\| | 1,595 |
| SALT00000052371_Cluster27052-006 | 1.000 | lincRNA556_PMID_24635777\| | 1,545 |
| SALT00000076309_Cluster27052-007 | 1.000 | lincRNA556_PMID_24635777\| | 2,445 |
| SALT00000056671_Cluster27062-001 | 0.212 | NA | 1,479 |
| SALT00000069891_Cluster27062-002 | 0.185 | NA | 1,657 |
| SALT00000049682_Cluster27078-001 | 0.276 | NA | 1,656 |
| SALT00000007382_Cluster27166-001 | 0.030 | NA | 1,505 |
| SALT00000016686_Cluster27166-002 | 0.027 | NA | 1,652 |
| SALT00000020169_Cluster27166-003 | 0.028 | NA | 1,649 |
| SALT00000001624_Cluster27169-001 | 1.000 | osa-MIR396d\| | 1,592 |
| SALT00000024315_Cluster27169-002 | 1.000 | osa-MIR396d\| | 1,652 |
| SALT00000067198_Cluster27169-003 | 1.000 | osa-MIR396d\| | 1,427 |
| SALT00000057004_Cluster27179-001 | 1.000 | lincRNA29_PMID_24635777\| | 1,652 |
| SALT00000052056_Cluster27228-001 | 1.000 | osa-MIR396g\|osa-MIR396h\| | 1,650 |
| SALT00000061600_Cluster27228-002 | 1.000 | osa-MIR396g\|osa-MIR396h\| | 1,387 |
| SALT00000064585_Cluster27269-001 | 0.160 | NA | 1,648 |
| SALT00000002586_Cluster27339-001 | 1.000 | JNnc_loci0076\|JNnc_loci0735\|JNnc_loci0732\|JNnc_ | 1,643 |
| SALT00000004318_Cluster27339-002 | 1.000 | JNnc_loci0076\|JNnc_loci0732\| | 1,610 |
| SALT00000035911_Cluster27431-001 | 0.165 | NA | 1,639 |
| SALT00000055381_Cluster27480-001 | 0.178 | NA | 1,637 |
| SALT00000016728_Cluster27582-001 | 1.000 | TCONS_00037642\| | 1,575 |
| SALT00000039984_Cluster27582-002 | 1.000 | TCONS_00037642\| | 1,526 |
| SALT00000070173_Cluster27583-001 | 0.280 | NA | 1,632 |
| SALT00000017976_Cluster27593-001 | 0.087 | NA | 1,631 |
| SALT00000015371_Cluster27649-001 | 0.987 | egu-MIR172d\| | 1,628 |
| SALT00000009501_Cluster27690-001 | 1.000 | bdi-MIR156a\|far-MIR156a\|far-MIR156b\|sof-MIR15 | 1,626 |
| SALT00000054147_Cluster27825-001 | 0.999 | lincRNA165_PMID_24635777\| | 1,620 |
| SALT00000054229_Cluster27940-001 | 0.194 | NA | 1,614 |
| SALT00000017017_Cluster28012-001 | 0.999 | pnrd_Mtr_chr0.trna21_ValAAC\|pnrd_Pop_scaffold_ | 1,610 |
| SALT00000054323_Cluster28124-001 | 0.246 | NA | 1,604 |
| SALT00000018007_Cluster28161-001 | 0.255 | NA | 1,602 |
| SALT00000056774_Cluster28161-002 | 0.285 | NA | 1,456 |
| SALT00000038281_Cluster28165-001 | 0.157 | JNnc_loci1052\| | 1,602 |
| SALT00000066699_Cluster28165-002 | 0.408 | JNnc_loci1052\| | 1,562 |
| SALT00000004769_Cluster28229-001 | 0.284 | NA | 1,598 |
| SALT00000048264_Cluster28462-001 | 0.092 | NA | 1,586 |
| SALT00000050786_Cluster28466-001 | 0.063 | pnrd_Osa_chr1.trna36_ThrTGT\|pnrd_Osa_chr12.trna | 1,586 |
| SALT00000004805_Cluster28481-001 | 0.037 | TCONS_00097716\| | 1,226 |
| SALT00000013968_Cluster28481-002 | 0.029 | TCONS_00097716\|GRMZM2G129587_T01\|GRMZI | 1,585 |
| SALT00000051989_Cluster28677-001 | 0.068 | NA | 1,575 |
| SALT00000002433_Cluster28703-001 | 0.998 | TCONS_00020521\| | 1,573 |
| SALT00000039059_Cluster28768-001 | 0.536 | ghr-MIR4370\| | 1,570 |
| SALT00000044501_Cluster28771-001 | 0.267 | ghr-MIR4370\| | 1,570 |
| SALT00000051609_Cluster28910-001 | 0.268 | NA | 1,564 |
| SALT00000017336_Cluster28956-001 | 0.198 | NA | 1,561 |
| SALT00000010492_Cluster28987-001 | 0.312 | ghr-MIR4370\| | 1,559 |
| SALT00000012180_Cluster28987-002 | 0.383 | ghr-MIR4370\| | 1,150 |
| SALT00000060734_Cluster28987-003 | 0.309 | ghr-MIR4370\| | 1,598 |

| | | | |
|---|---|---|---|
| SALT00000040569_Cluster29011-001 | 1.000 | lincRNA590_PMID_24635777\| | 1,558 |
| SALT00000051592_Cluster29011-002 | 1.000 | lincRNA590_PMID_24635777\| | 1,512 |
| SALT00000013904_Cluster29068-001 | 0.041 | NA | 1,555 |
| SALT00000049132_Cluster29073-001 | 0.020 | NA | 1,555 |
| SALT00000057834_Cluster29095-001 | 0.261 | NA | 1,554 |
| SALT00000049931_Cluster29117-001 | 1.000 | XLOC_057324\| | 1,553 |
| SALT00000010367_Cluster29210-001 | 0.267 | NA | 1,548 |
| SALT00000010775_Cluster29210-002 | 0.275 | NA | 1,487 |
| SALT00000014509_Cluster29212-001 | 1.000 | lincRNA88_PMID_24635777\| | 1,548 |
| SALT00000041475_Cluster29212-002 | 1.000 | lincRNA88_PMID_24635777\| | 1,064 |
| SALT00000058579_Cluster29226-005 | 0.060 | NA | 1,344 |
| SALT00000045982_Cluster29312-003 | 1.000 | pnrd_Mtr_chr5.trna48_GlyGCC\|pnrd_Osa_chr3.trna[illegible] | 1,473 |
| SALT00000050379_Cluster29384-002 | 0.854 | peu-MIR2914\|NONATHT000930\|tae-MIR170b_npr\|[illegible] | 1,539 |
| SALT00000036795_Cluster29398-001 | 0.110 | TCONS_00055850\| | 1,538 |
| SALT00000036749_Cluster29495-001 | 0.131 | NA | 1,533 |
| SALT00000013802_Cluster29509-001 | 0.272 | NA | 1,532 |
| SALT00000051925_Cluster29537-001 | 0.992 | NONATHT001130\| | 1,531 |
| SALT00000049358_Cluster29564-001 | 1.000 | NONATHT001713\| | 1,529 |
| SALT00000015587_Cluster29619-001 | 0.290 | NA | 1,525 |
| SALT00000018459_Cluster29718-001 | 0.221 | NA | 1,519 |
| SALT00000045579_Cluster29741-001 | 0.197 | NA | 1,493 |
| SALT00000058687_Cluster29741-002 | 0.109 | NA | 1,518 |
| SALT00000010072_Cluster29745-001 | 0.115 | NA | 1,517 |
| SALT00000043673_Cluster29761-001 | 0.212 | NA | 1,516 |
| SALT00000037242_Cluster29945-001 | 1.000 | JNnc_loci0849\| | 1,505 |
| SALT00000059915_Cluster29976-001 | 0.062 | NA | 1,504 |
| SALT00000037623_Cluster30002-001 | 1.000 | lincRNA244_PMID_24635777\| | 1,502 |
| SALT00000042886_Cluster30029-001 | 0.232 | tae-MIR444a\|zma-MIR444a\|osa-MIR444c\|pvi-MIR4[illegible] | 1,500 |
| SALT00000044967_Cluster30030-001 | 1.000 | XLOC_057324\| | 1,500 |
| SALT00000055732_Cluster30060-001 | 0.999 | TCONS_00063443\| | 1,498 |
| SALT00000019518_Cluster30140-002 | 1.000 | osa-MIRf10576-npr\| | 1,455 |
| SALT00000041789_Cluster30157-001 | 0.127 | NA | 1,491 |
| SALT00000061033_Cluster30192-001 | 0.016 | NA | 1,489 |
| SALT00000019336_Cluster30251-001 | 0.140 | NA | 1,485 |
| SALT00000041770_Cluster30267-001 | 0.200 | NA | 1,484 |
| SALT00000058736_Cluster30501-001 | 1.000 | XLOC_057324\| | 1,468 |
| SALT00000016938_Cluster30521-001 | 1.000 | TCONS_00063488\| | 1,466 |
| SALT00000060539_Cluster30560-001 | 0.214 | NA | 1,463 |
| SALT00000071667_Cluster30593-001 | 0.237 | NA | 1,461 |
| SALT00000047209_Cluster30605-001 | 1.000 | lincRNA376_PMID_24635777\| | 1,460 |
| SALT00000060219_Cluster30610-001 | 0.997 | hvu-MIR6196\| | 1,460 |
| SALT00000049648_Cluster30679-001 | 0.287 | NA | 1,454 |
| SALT00000062958_Cluster30694-001 | 0.092 | NA | 1,453 |
| SALT00000060691_Cluster30743-002 | 0.073 | NA | 1,249 |
| SALT00000059623_Cluster30813-001 | 0.247 | NA | 1,441 |
| SALT00000049074_Cluster30878-001 | 1.000 | NONATHT001713\| | 1,433 |
| SALT00000060587_Cluster30879-001 | 1.000 | lincRNA396_PMID_24635777\| | 1,433 |
| SALT00000048604_Cluster30986-001 | 1.000 | NONATHT001713\| | 1,422 |
| SALT00000038945_Cluster31000-001 | 0.228 | NA | 1,420 |
| SALT00000023046_Cluster31087-001 | 0.201 | NA | 1,410 |
| SALT00000059133_Cluster31139-001 | 1.000 | lincRNA33_PMID_24635777\|lincRNA73_PMID_24 | 1,402 |
| SALT00000016739_Cluster31151-001 | 1.000 | TCONS_00063488\| | 1,399 |
| SALT00000065971_Cluster31178-001 | 1.000 | NONATHT001713\| | 1,397 |
| SALT00000059729_Cluster31247-001 | 0.961 | peu-MIR2910\|NONATHT000930\|NONATHT00216[illegible] | 1,386 |
| SALT00000008695_Cluster31311-001 | 0.875 | pnrd_Osa_chr4.trna89_SerGCT\|pnrd_Osa_chr4.trna6 | 1,374 |
| SALT00000040371_Cluster31316-001 | 0.248 | NA | 1,373 |
| SALT00000051417_Cluster31478-001 | 1.000 | pnrd_Ara_chr5.trna98_CysGCA\|pnrd_Ara_chr5.trna[illegible] | 1,337 |
| SALT00000053737_Cluster31488-001 | 0.259 | NA | 1,335 |

| | | | |
|---|---|---|---|
| SALT00000059694_Cluster31850-001 | 0.227 | JNnc_loci0363\|JNnc_loci0813\|JNnc_loci0814\|JNnc_ | 1,236 |
| SALT00000003391_Cluster32196-001 | 1.000 | osa-MIRf11040-npr\| | 1,041 |
| SALT00000059643_Cluster32226-002 | 1.000 | TCONS_00044708\|TCONS_00044707\| | 1,103 |
| SALT00000042826_Cluster32236-001 | 0.896 | GRMZM2G073934_T04\|GRMZM2G073934_T05\| | 1,098 |
| SALT00000042725_Cluster32390-001 | 0.194 | pnrd_Osa_chr2.trna13_MetCAT\|pnrd_Osa_chr12.trna | 1,007 |
| SALT00000012029_Cluster32578-001 | 0.153 | NA | 665 |
| SALT00000059722_Cluster32584-001 | 0.269 | NA | 612 |
| SALT00000063356_Cluster293-001 | 0.005 | NA | 8,717 |
| SALT00000082117_Cluster300-016 | 0.022 | lincRNA11_PMID_24635777\| | 831 |
| SALT00000015342_Cluster413-001 | 0.248 | NA | 1,637 |
| SALT00000022825_Cluster413-002 | 0.192 | NA | 2,595 |
| SALT00000024336_Cluster413-003 | 0.164 | NA | 2,865 |
| SALT00000029276_Cluster413-004 | 0.105 | NA | 2,745 |
| SALT00000029588_Cluster413-005 | 0.056 | NA | 4,547 |
| SALT00000033371_Cluster413-006 | 0.125 | NA | 3,318 |
| SALT00000034028_Cluster413-007 | 0.055 | NA | 4,577 |
| SALT00000052429_Cluster413-008 | 0.245 | NA | 1,657 |
| SALT00000066381_Cluster413-009 | 0.132 | NA | 3,224 |
| SALT00000069942_Cluster413-010 | 0.064 | NA | 2,889 |
| SALT00000080872_Cluster413-011 | 0.071 | NA | 2,722 |
| SALT00000085486_Cluster413-012 | 0.036 | NA | 4,383 |
| SALT00000085740_Cluster413-013 | 0.103 | NA | 3,617 |
| SALT00000093171_Cluster413-015 | 0.056 | NA | 4,292 |
| SALT00000073214_Cluster529-007 | 0.229 | NA | 2,568 |
| SALT00000022091_Cluster1001-001 | 0.295 | NA | 2,688 |
| SALT00000063004_Cluster1001-004 | 0.281 | NA | 2,787 |
| SALT00000080671_Cluster1001-007 | 0.280 | NA | 2,792 |
| SALT00000092574_Cluster1005-001 | 0.023 | NA | 6,801 |
| SALT00000088134_Cluster1170-001 | 0.164 | NA | 6,307 |
| SALT00000062143_Cluster1395-037 | 0.211 | NA | 1,936 |
| SALT00000047692_Cluster1516-016 | 0.280 | NA | 1,968 |
| SALT00000007916_Cluster1723-001 | 0.291 | NA | 1,968 |
| SALT00000084955_Cluster1732-001 | 0.047 | NA | 5,361 |
| SALT00000031276_Cluster1752-023 | 0.066 | NA | 2,346 |
| SALT00000041260_Cluster1752-029 | 0.081 | NA | 2,026 |
| SALT00000064695_Cluster1771-001 | 0.173 | NA | 3,074 |
| SALT00000085707_Cluster1776-002 | 0.113 | NA | 5,325 |
| SALT00000047775_Cluster1855-001 | 0.128 | NA | 5,257 |
| SALT00000013535_Cluster1905-001 | 0.044 | NA | 1,793 |
| SALT00000056793_Cluster1905-005 | 0.084 | NA | 1,826 |
| SALT00000079069_Cluster2012-002 | 0.192 | NA | 5,148 |
| SALT00000084547_Cluster2012-003 | 0.100 | NA | 3,806 |
| SALT00000093092_Cluster2012-004 | 0.059 | NA | 4,475 |
| SALT00000081399_Cluster2025-002 | 0.095 | NA | 2,658 |
| SALT00000034178_Cluster2057-017 | 0.151 | NA | 4,280 |
| SALT00000073130_Cluster2140-001 | 0.196 | NA | 3,267 |
| SALT00000091647_Cluster2140-003 | 0.025 | NA | 3,838 |
| SALT00000003670_Cluster2206-001 | 0.172 | NA | 2,066 |
| SALT00000034275_Cluster2271-001 | 0.062 | NA | 4,388 |
| SALT00000067519_Cluster2271-002 | 0.168 | NA | 2,827 |
| SALT00000083598_Cluster2271-004 | 0.098 | NA | 3,705 |
| SALT00000088955_Cluster2271-005 | 0.064 | NA | 4,343 |
| SALT00000091009_Cluster2271-006 | 0.055 | NA | 4,584 |
| SALT00000091380_Cluster2271-007 | 0.125 | NA | 3,309 |
| SALT00000091442_Cluster2271-008 | 0.042 | NA | 4,978 |
| SALT00000091708_Cluster2271-009 | 0.043 | NA | 4,677 |
| SALT00000090329_Cluster2271-010 | 0.076 | NA | 4,088 |
| SALT00000075250_Cluster2271-012 | 0.184 | NA | 2,673 |

| | | | |
|---|---|---|---|
| SALT00000014809_Cluster2278-001 | 0.037 | NA | 1,643 |
| SALT00000039004_Cluster2296-003 | 0.073 | NA | 2,009 |
| SALT00000039056_Cluster2296-010 | 0.257 | NA | 1,898 |
| SALT00000029096_Cluster2312-001 | 0.160 | NA | 3,714 |
| SALT00000077691_Cluster2312-003 | 0.231 | NA | 2,905 |
| SALT00000085561_Cluster2312-004 | 0.066 | NA | 4,957 |
| SALT00000015916_Cluster2485-002 | 0.085 | NA | 2,003 |
| SALT00000041114_Cluster2485-003 | 0.091 | NA | 1,899 |
| SALT00000090612_Cluster2588-002 | 0.079 | NA | 4,792 |
| SALT00000036719_Cluster2588-003 | 0.277 | NA | 1,468 |
| SALT00000031948_Cluster2631-002 | 0.112 | NA | 3,600 |
| SALT00000032786_Cluster2631-003 | 0.093 | NA | 3,803 |
| SALT00000067059_Cluster2631-008 | 0.184 | NA | 2,783 |
| SALT00000091541_Cluster2631-013 | 0.113 | NA | 3,587 |
| SALT00000023676_Cluster2631-023 | 0.080 | NA | 2,615 |
| SALT00000023831_Cluster2631-026 | 0.131 | NA | 2,640 |
| SALT00000051596_Cluster2631-027 | 0.195 | NA | 1,373 |
| SALT00000040203_Cluster2652-001 | 0.195 | NA | 1,872 |
| SALT00000074672_Cluster2652-004 | 0.120 | NA | 2,903 |
| SALT00000081457_Cluster2744-001 | 0.056 | NA | 4,720 |
| SALT00000091166_Cluster2924-001 | 0.364 | pnrd_Osa_chr2.trna64_GlnTTG\|pnrd_Osa_chr2.trna6 | 4,636 |
| SALT00000080217_Cluster3019-001 | 0.153 | NA | 3,633 |
| SALT00000087452_Cluster3100-001 | 0.206 | NA | 4,555 |
| SALT00000078670_Cluster3124-001 | 0.250 | NA | 2,813 |
| SALT00000086363_Cluster3167-009 | 0.252 | NA | 3,860 |
| SALT00000054673_Cluster3185-007 | 0.052 | NA | 1,993 |
| SALT00000074327_Cluster3185-009 | 0.251 | NA | 2,365 |
| SALT00000087706_Cluster3436-002 | 0.169 | NA | 4,420 |
| SALT00000033360_Cluster3492-001 | 0.085 | NA | 4,396 |
| SALT00000076886_Cluster3492-002 | 0.101 | NA | 4,127 |
| SALT00000093058_Cluster3539-001 | 0.279 | TCONS_00084421\| | 4,379 |
| SALT00000060433_Cluster3692-001 | 0.131 | NA | 2,223 |
| SALT00000056455_Cluster3704-002 | 0.073 | NA | 1,797 |
| SALT00000016598_Cluster3931-002 | 0.117 | NA | 1,611 |
| SALT00000076533_Cluster3931-007 | 0.122 | NA | 2,913 |
| SALT00000005426_Cluster3951-001 | 0.157 | NA | 1,608 |
| SALT00000022329_Cluster4041-001 | 0.280 | NA | 2,120 |
| SALT00000056278_Cluster4041-002 | 0.283 | NA | 2,100 |
| SALT00000092270_Cluster4104-002 | 0.263 | NA | 3,760 |
| SALT00000066029_Cluster4217-004 | 0.114 | NA | 2,518 |
| SALT00000039350_Cluster4521-001 | 0.222 | NA | 4,081 |
| SALT00000080322_Cluster4753-002 | 0.234 | NA | 2,958 |
| SALT00000081349_Cluster4753-003 | 0.174 | NA | 4,027 |
| SALT00000089949_Cluster4753-004 | 0.141 | NA | 3,706 |
| SALT00000033424_Cluster4832-001 | 0.074 | NA | 4,010 |
| SALT00000064017_Cluster4832-002 | 0.090 | NA | 2,740 |
| SALT00000086661_Cluster4917-001 | 0.011 | NA | 3,993 |
| SALT00000064616_Cluster4989-003 | 0.144 | NA | 2,910 |
| SALT00000004753_Cluster5035-004 | 0.173 | NA | 1,354 |
| SALT00000063790_Cluster5087-001 | 0.042 | NA | 3,958 |
| SALT00000052386_Cluster5154-004 | 0.070 | NA | 1,782 |
| SALT00000087161_Cluster5660-004 | 0.125 | NA | 3,852 |
| SALT00000078466_Cluster5957-005 | 0.165 | NA | 2,569 |
| SALT00000073184_Cluster6210-001 | 0.240 | NA | 2,787 |
| SALT00000082387_Cluster6210-002 | 0.235 | NA | 2,824 |
| SALT00000092051_Cluster6210-003 | 0.137 | NA | 3,761 |
| SALT00000074055_Cluster6255-001 | 0.262 | NA | 2,306 |
| SALT00000077198_Cluster6255-002 | 0.178 | NA | 3,010 |

| | | | |
|---|---|---|---|
| SALT00000086268_Cluster6255-003 | 0.113 | NA | 3,752 |
| SALT00000085011_Cluster6285-001 | 0.183 | NA | 3,747 |
| SALT00000085371_Cluster6285-002 | 0.198 | NA | 3,651 |
| SALT00000092802_Cluster6285-003 | 0.221 | NA | 3,486 |
| SALT00000077464_Cluster6296-001 | 0.097 | NA | 3,745 |
| SALT00000083298_Cluster6339-002 | 0.053 | NA | 3,738 |
| SALT00000025510_Cluster6352-001 | 0.099 | NA | 2,635 |
| SALT00000064920_Cluster6383-001 | 0.179 | JNnc_loci0896\|osa-MIR6253\| | 3,109 |
| SALT00000074537_Cluster6530-004 | 0.168 | NA | 2,494 |
| SALT00000088994_Cluster6587-003 | 0.260 | NA | 3,700 |
| SALT00000089108_Cluster6594-001 | 0.202 | NA | 3,699 |
| SALT00000092676_Cluster6640-005 | 0.192 | NA | 3,360 |
| SALT00000007469_Cluster6681-001 | 0.131 | NA | 1,443 |
| SALT00000057188_Cluster6681-003 | 0.251 | NA | 1,838 |
| SALT00000092957_Cluster6751-001 | 0.074 | NA | 3,675 |
| SALT00000071526_Cluster6945-001 | 0.011 | NA | 3,050 |
| SALT00000093134_Cluster6945-002 | 0.066 | NA | 3,645 |
| SALT00000061817_Cluster7002-001 | 0.254 | NA | 1,457 |
| SALT00000013501_Cluster7082-001 | 0.114 | NA | 1,595 |
| SALT00000031975_Cluster7082-002 | 0.279 | NA | 3,624 |
| SALT00000012825_Cluster7452-001 | 0.287 | NA | 2,078 |
| SALT00000024775_Cluster7452-003 | 0.224 | NA | 2,547 |
| SALT00000092256_Cluster7462-001 | 0.166 | NA | 3,566 |
| SALT00000085759_Cluster7683-001 | 0.202 | NA | 3,530 |
| SALT00000088644_Cluster7714-001 | 0.290 | NA | 3,526 |
| SALT00000083163_Cluster7754-001 | 0.225 | NA | 3,521 |
| SALT00000009987_Cluster7973-001 | 0.139 | NA | 1,728 |
| SALT00000069645_Cluster8184-003 | 0.165 | NA | 2,643 |
| SALT00000039145_Cluster8184-004 | 0.019 | NA | 1,646 |
| SALT00000011688_Cluster8200-001 | 0.109 | NA | 1,957 |
| SALT00000049745_Cluster8200-002 | 0.098 | NA | 1,625 |
| SALT00000083726_Cluster8322-001 | 0.281 | NA | 3,420 |
| SALT00000069958_Cluster8475-001 | 0.145 | NA | 3,390 |
| SALT00000089611_Cluster8504-002 | 0.712 | osa-MIR444a\|osa-MIR444e\| | 3,385 |
| SALT00000019624_Cluster8509-001 | 0.084 | NA | 1,774 |
| SALT00000092931_Cluster8509-002 | 0.048 | NA | 3,384 |
| SALT00000088731_Cluster8533-001 | 0.233 | NA | 3,378 |
| SALT00000085456_Cluster8719-001 | 0.106 | NA | 3,344 |
| SALT00000078041_Cluster8997-001 | 0.804 | pnrd_Osa_chr4.trna42_IleAAT\|pnrd_Osa_chr11.trna1 | 3,288 |
| SALT00000053153_Cluster9044-001 | 0.234 | NA | 1,743 |
| SALT00000068846_Cluster9044-002 | 0.093 | NA | 3,278 |
| SALT00000068378_Cluster9091-014 | 0.496 | lincRNA616_PMID_24635777\| | 1,550 |
| SALT00000072267_Cluster9212-002 | 0.265 | NA | 1,321 |
| SALT00000057668_Cluster9373-007 | 0.078 | NA | 1,506 |
| SALT00000071850_Cluster9516-001 | 0.192 | NA | 3,189 |
| SALT00000074089_Cluster9516-002 | 0.202 | NA | 3,099 |
| SALT00000089351_Cluster9516-003 | 0.202 | NA | 3,099 |
| SALT00000016479_Cluster9531-003 | 0.102 | NA | 1,654 |
| SALT00000041702_Cluster9606-008 | 0.222 | NA | 1,596 |
| SALT00000074195_Cluster9772-001 | 0.069 | NA | 3,145 |
| SALT00000018658_Cluster9788-002 | 0.207 | NA | 1,643 |
| SALT00000043189_Cluster9788-004 | 0.169 | NA | 1,419 |
| SALT00000000224_Cluster9931-001 | 0.131 | NA | 2,014 |
| SALT00000004048_Cluster9931-002 | 0.129 | NA | 2,039 |
| SALT00000008315_Cluster9931-003 | 0.129 | NA | 1,538 |
| SALT00000009445_Cluster9931-004 | 0.150 | NA | 1,791 |
| SALT00000010547_Cluster9931-005 | 0.124 | NA | 2,103 |
| SALT00000010934_Cluster9931-006 | 0.141 | NA | 1,901 |

| | | | |
|---|---|---|---|
| SALT00000016173_Cluster9931-007 | 0.128 | NA | 2,050 |
| SALT00000019881_Cluster9931-008 | 0.163 | NA | 1,650 |
| SALT00000042740_Cluster9931-009 | 0.146 | NA | 1,843 |
| SALT00000050667_Cluster9931-011 | 0.144 | NA | 1,857 |
| SALT00000055232_Cluster9931-012 | 0.165 | NA | 1,637 |
| SALT00000093547_Cluster9970-001 | 0.108 | NA | 3,114 |
| SALT00000002400_Cluster9972-004 | 0.126 | NA | 1,750 |
| SALT00000057201_Cluster9972-021 | 0.161 | NA | 1,343 |
| SALT00000078224_Cluster10070-001 | 0.035 | NA | 3,098 |
| SALT00000048858_Cluster10096-001 | 0.142 | NA | 1,691 |
| SALT00000077884_Cluster10126-002 | 0.175 | NA | 3,091 |
| SALT00000065487_Cluster10227-002 | 0.136 | NA | 3,077 |
| SALT00000078861_Cluster10282-001 | 0.951 | pnrd_Osa_chr2.trna65_GlnTTG\|pnrd_Osa_chr2.trna( | 3,068 |
| SALT00000022560_Cluster10470-001 | 0.272 | NA | 2,668 |
| SALT00000068345_Cluster10600-001 | 0.099 | NA | 3,022 |
| SALT00000019778_Cluster10619-003 | 0.586 | bdi-MIR444c\|bdi-MIR444d\|ssp-MIR444c\|zma-MIR4 | 1,670 |
| SALT00000027676_Cluster10820-001 | 0.162 | NA | 2,846 |
| SALT00000031089_Cluster10820-002 | 0.116 | NA | 2,986 |
| SALT00000059455_Cluster10820-003 | 0.229 | NA | 2,240 |
| SALT00000078112_Cluster10820-004 | 0.148 | NA | 2,995 |
| SALT00000081014_Cluster10820-005 | 0.220 | NA | 2,312 |
| SALT00000056586_Cluster10871-001 | 0.170 | NA | 1,944 |
| SALT00000076542_Cluster10942-001 | 0.056 | NA | 2,980 |
| SALT00000047709_Cluster11041-001 | 0.182 | NA | 2,966 |
| SALT00000013945_Cluster11095-001 | 0.211 | NA | 1,828 |
| SALT00000027479_Cluster11095-002 | 0.199 | NA | 2,960 |
| SALT00000069757_Cluster11095-003 | 0.155 | NA | 2,177 |
| SALT00000063206_Cluster11132-001 | 0.141 | NA | 2,954 |
| SALT00000005825_Cluster11138-001 | 0.192 | NA | 1,748 |
| SALT00000068049_Cluster11159-001 | 0.254 | NA | 2,950 |
| SALT00000056700_Cluster11428-001 | 0.226 | NA | 2,921 |
| SALT00000027851_Cluster11495-001 | 0.131 | NA | 2,766 |
| SALT00000042588_Cluster11495-002 | 0.241 | NA | 1,722 |
| SALT00000066375_Cluster11495-003 | 0.162 | NA | 2,165 |
| SALT00000078880_Cluster11495-004 | 0.120 | NA | 2,915 |
| SALT00000080004_Cluster11495-005 | 0.120 | NA | 2,911 |
| SALT00000080587_Cluster11799-001 | 0.262 | NA | 2,886 |
| SALT00000020714_Cluster12040-001 | 0.270 | NA | 2,863 |
| SALT00000024075_Cluster12040-002 | 0.280 | NA | 2,791 |
| SALT00000066937_Cluster12040-004 | 0.240 | NA | 3,235 |
| SALT00000073900_Cluster12085-002 | 0.145 | NA | 2,531 |
| SALT00000074884_Cluster12319-001 | 0.121 | NA | 2,837 |
| SALT00000039699_Cluster12357-001 | 0.087 | NA | 1,908 |
| SALT00000068194_Cluster12380-003 | 0.087 | NA | 2,831 |
| SALT00000053658_Cluster12434-001 | 0.138 | NA | 1,772 |
| SALT00000069401_Cluster12434-002 | 0.060 | NA | 2,759 |
| SALT00000076188_Cluster12434-003 | 0.057 | NA | 2,826 |
| SALT00000070702_Cluster12434-004 | 0.061 | NA | 2,727 |
| SALT00000067783_Cluster12543-002 | 0.253 | NA | 2,747 |
| SALT00000031101_Cluster12594-004 | 0.013 | NA | 2,678 |
| SALT00000082114_Cluster12744-001 | 0.133 | NA | 2,797 |
| SALT00000022863_Cluster12806-001 | 0.021 | NA | 2,791 |
| SALT00000052568_Cluster12806-002 | 0.041 | NA | 1,846 |
| SALT00000054357_Cluster12806-003 | 0.042 | NA | 1,797 |
| SALT00000039361_Cluster12976-003 | 0.218 | NA | 1,731 |
| SALT00000076481_Cluster13230-001 | 0.085 | NA | 2,755 |
| SALT00000063950_Cluster13318-001 | 0.188 | NA | 2,747 |
| SALT00000062568_Cluster13473-001 | 0.194 | NA | 2,734 |

| | | | |
|---|---|---|---|
| SALT00000070440_Cluster13562-004 | 0.194 | NA | 2,615 |
| SALT00000081266_Cluster13642-001 | 0.244 | NA | 2,722 |
| SALT00000071349_Cluster13672-001 | 0.028 | NA | 2,718 |
| SALT00000073555_Cluster13682-001 | 0.104 | NA | 2,717 |
| SALT00000064859_Cluster13691-001 | 0.230 | NA | 2,716 |
| SALT00000066502_Cluster13710-001 | 0.267 | NA | 2,590 |
| SALT00000080227_Cluster13710-002 | 0.250 | NA | 2,715 |
| SALT00000068278_Cluster13748-001 | 0.094 | NA | 2,712 |
| SALT00000061442_Cluster13845-001 | 0.184 | NA | 1,854 |
| SALT00000077778_Cluster14192-001 | 0.991 | pnrd_Osa_chr5.trna34_LysCTT\|pnrd_Osa_chr8.trna2 | 2,680 |
| SALT00000070142_Cluster14253-001 | 0.033 | NA | 2,675 |
| SALT00000082773_Cluster14332-001 | 0.262 | NA | 2,670 |
| SALT00000076718_Cluster14342-001 | 0.086 | NA | 2,669 |
| SALT00000083224_Cluster14497-001 | 0.491 | gma-MIR172g\|dpr-MIR172a\|tcc-MIR172c\|nta-MIR1 | 2,660 |
| SALT00000066464_Cluster14511-001 | 0.193 | NA | 2,658 |
| SALT00000051997_Cluster14596-002 | 0.257 | NA | 1,964 |
| SALT00000065528_Cluster14596-003 | 0.178 | NA | 2,580 |
| SALT00000077480_Cluster14596-005 | 0.176 | NA | 2,600 |
| SALT00000073785_Cluster14598-001 | 0.114 | NA | 2,432 |
| SALT00000072961_Cluster15020-001 | 0.266 | NA | 2,623 |
| SALT00000058512_Cluster15034-001 | 0.045 | NA | 2,073 |
| SALT00000052945_Cluster15066-002 | 0.090 | NA | 1,380 |
| SALT00000062218_Cluster15066-003 | 0.242 | NA | 2,620 |
| SALT00000068020_Cluster15145-001 | 0.154 | NA | 2,615 |
| SALT00000024164_Cluster15251-001 | 0.096 | NA | 2,608 |
| SALT00000007764_Cluster15263-001 | 0.140 | NA | 1,697 |
| SALT00000078809_Cluster15422-001 | 0.054 | NA | 2,596 |
| SALT00000042660_Cluster15440-001 | 0.105 | NA | 2,594 |
| SALT00000073036_Cluster15477-001 | 0.036 | NA | 2,591 |
| SALT00000030946_Cluster15523-004 | 0.029 | NA | 2,749 |
| SALT00000000585_Cluster15523-005 | 0.121 | NA | 1,813 |
| SALT00000005046_Cluster15548-001 | 0.066 | NA | 1,591 |
| SALT00000046247_Cluster15548-004 | 0.075 | NA | 1,404 |
| SALT00000082804_Cluster15908-001 | 0.253 | NA | 2,559 |
| SALT00000011996_Cluster16111-001 | 0.121 | NA | 2,052 |
| SALT00000036752_Cluster16208-001 | 0.123 | NA | 2,170 |
| SALT00000083971_Cluster16208-002 | 0.098 | NA | 2,536 |
| SALT00000081950_Cluster16243-001 | 0.101 | NA | 2,534 |
| SALT00000063936_Cluster16573-001 | 0.238 | NA | 2,508 |
| SALT00000016549_Cluster16593-001 | 0.063 | NA | 1,454 |
| SALT00000075538_Cluster16673-002 | 0.032 | NA | 2,500 |
| SALT00000083773_Cluster17088-001 | 0.252 | NA | 2,462 |
| SALT00000078633_Cluster17340-001 | 0.298 | tae-MIR2005a_1_npr\|pnrd_Mtr_chr1.trna82_LysTTT | 2,432 |
| SALT00000073422_Cluster17409-001 | 0.283 | NA | 2,421 |
| SALT00000036778_Cluster17509-001 | 0.224 | NA | 1,910 |
| SALT00000061545_Cluster17509-002 | 0.275 | NA | 1,521 |
| SALT00000073928_Cluster17736-001 | 0.066 | NA | 2,361 |
| SALT00000051362_Cluster18015-001 | 0.996 | osa-MIR444b\|zma-MIR444a\|pvi-MIR444\|bdi-MIR44 | 2,298 |
| SALT00000008165_Cluster18095-001 | 0.151 | NA | 1,202 |
| SALT00000047616_Cluster18196-001 | 0.146 | NA | 2,264 |
| SALT00000017340_Cluster18450-001 | 0.039 | NA | 1,671 |
| SALT00000057258_Cluster18450-002 | 0.039 | NA | 2,226 |
| SALT00000016448_Cluster18450-003 | 0.055 | NA | 1,714 |
| SALT00000079153_Cluster18718-001 | 0.047 | NA | 2,194 |
| SALT00000051147_Cluster18782-001 | 0.147 | NA | 2,084 |
| SALT00000065094_Cluster18782-002 | 0.128 | NA | 2,187 |
| SALT00000037496_Cluster18829-001 | 0.155 | NA | 1,456 |
| SALT00000056678_Cluster18946-001 | 0.068 | NA | 2,172 |

| | | | |
|---|---|---|---|
| SALT00000040430_Cluster19068-001 | 0.120 | NA | 2,161 |
| SALT00000045306_Cluster19114-001 | 0.080 | NA | 2,157 |
| SALT00000045273_Cluster19307-001 | 0.139 | NA | 1,549 |
| SALT00000060007_Cluster19307-002 | 0.096 | NA | 2,138 |
| SALT00000061868_Cluster19307-003 | 0.141 | NA | 1,533 |
| SALT00000016915_Cluster19467-001 | 0.137 | NA | 2,122 |
| SALT00000051702_Cluster19473-001 | 0.123 | NA | 2,122 |
| SALT00000041646_Cluster19591-001 | 0.052 | NA | 2,044 |
| SALT00000046906_Cluster19591-002 | 0.049 | NA | 2,113 |
| SALT00000058654_Cluster19648-003 | 0.077 | NA | 2,108 |
| SALT00000036705_Cluster19691-001 | 0.085 | NA | 2,104 |
| SALT00000004443_Cluster19748-001 | 0.128 | NA | 1,771 |
| SALT00000039896_Cluster19748-002 | 0.115 | NA | 1,941 |
| SALT00000040495_Cluster19748-003 | 0.123 | NA | 1,832 |
| SALT00000042709_Cluster19748-004 | 0.049 | NA | 1,610 |
| SALT00000057202_Cluster19748-005 | 0.104 | NA | 2,099 |
| SALT00000062113_Cluster19748-006 | 0.084 | NA | 1,607 |
| SALT00000061564_Cluster19766-001 | 0.100 | NA | 2,098 |
| SALT00000060648_Cluster19799-001 | 0.066 | NA | 2,096 |
| SALT00000035952_Cluster20100-001 | 0.138 | NA | 1,930 |
| SALT00000053970_Cluster20100-002 | 0.127 | NA | 2,072 |
| SALT00000058681_Cluster20100-003 | 0.166 | NA | 1,622 |
| SALT00000060808_Cluster20109-001 | 0.078 | NA | 2,071 |
| SALT00000005536_Cluster20585-001 | 0.082 | NA | 2,034 |
| SALT00000051686_Cluster21032-002 | 0.171 | NA | 1,803 |
| SALT00000067843_Cluster21054-001 | 0.261 | NA | 2,001 |
| SALT00000009464_Cluster21169-001 | 0.068 | NA | 1,993 |
| SALT00000065843_Cluster21287-001 | 0.064 | NA | 1,986 |
| SALT00000014864_Cluster21316-001 | 0.108 | NA | 1,983 |
| SALT00000045188_Cluster21322-002 | 0.205 | NA | 1,983 |
| SALT00000067224_Cluster21362-003 | 0.266 | NA | 1,981 |
| SALT00000008741_Cluster21538-001 | 0.172 | NA | 1,968 |
| SALT00000037943_Cluster21782-001 | 0.136 | NA | 1,953 |
| SALT00000041485_Cluster21822-001 | 0.122 | NA | 1,808 |
| SALT00000053661_Cluster22176-001 | 0.108 | NA | 1,928 |
| SALT00000061186_Cluster22509-002 | 0.053 | NA | 1,351 |
| SALT00000039270_Cluster22710-001 | 0.162 | NA | 1,896 |
| SALT00000041017_Cluster22874-001 | 0.146 | NA | 1,885 |
| SALT00000037553_Cluster22899-001 | 0.151 | NA | 1,884 |
| SALT00000047272_Cluster23003-001 | 0.134 | NA | 1,877 |
| SALT00000058339_Cluster23103-001 | 0.293 | NA | 1,871 |
| SALT00000047369_Cluster23110-001 | 0.067 | NA | 1,870 |
| SALT00000002247_Cluster23203-001 | 0.197 | NA | 1,448 |
| SALT00000010153_Cluster23203-002 | 0.197 | NA | 1,444 |
| SALT00000018375_Cluster23227-001 | 0.133 | NA | 1,863 |
| SALT00000036026_Cluster23324-001 | 0.091 | NA | 1,857 |
| SALT00000010331_Cluster23429-001 | 0.202 | NA | 1,851 |
| SALT00000045354_Cluster23458-001 | 0.150 | NA | 1,850 |
| SALT00000007957_Cluster23470-001 | 0.243 | NA | 1,849 |
| SALT00000038276_Cluster23606-001 | 0.058 | NA | 1,840 |
| SALT00000010632_Cluster23647-001 | 0.191 | NA | 1,837 |
| SALT00000047313_Cluster23647-002 | 0.194 | NA | 1,566 |
| SALT00000036588_Cluster23647-003 | 0.285 | NA | 1,684 |
| SALT00000048457_Cluster23690-001 | 0.130 | NA | 1,835 |
| SALT00000046264_Cluster23882-001 | 0.811 | bdi-MIR160c\|sit-MIR160-2\|osa-MIR160a\|sbi-MIR16 | 1,825 |
| SALT00000041712_Cluster24100-001 | 0.104 | NA | 1,813 |
| SALT00000035956_Cluster24135-001 | 0.117 | NA | 1,811 |
| SALT00000058375_Cluster24144-001 | 0.043 | NA | 1,811 |

| | | | |
|---|---|---|---|
| SALT00000047189_Cluster24160-001 | 0.277 | NA | 1,810 |
| SALT00000052480_Cluster24391-001 | 0.165 | NA | 1,797 |
| SALT00000010663_Cluster24794-001 | 0.279 | NA | 1,774 |
| SALT00000001608_Cluster24813-001 | 0.070 | NA | 1,773 |
| SALT00000036607_Cluster24845-001 | 0.134 | NA | 1,541 |
| SALT00000020152_Cluster25495-001 | 0.167 | hvu-MIR168\|sof-MIR168a\|sof-MIR168b\|zma-MIR16 | 1,737 |
| SALT00000054098_Cluster25495-002 | 0.346 | osa-MIR168a\|zma-MIR168a\|sof-MIR168a\|zma-MIR | 1,579 |
| SALT00000051058_Cluster25495-003 | 0.572 | zma-MIR168b\|sof-MIR168a\|zma-MIR168a\|osa-MIR | 1,479 |
| SALT00000016360_Cluster25504-001 | 0.969 | lincRNA556_PMID_24635777\|lincRNA602_PMID_ | 1,527 |
| SALT00000053113_Cluster25504-002 | 0.828 | lincRNA556_PMID_24635777\|lincRNA602_PMID_ | 1,737 |
| SALT00000036287_Cluster25536-004 | 0.022 | NA | 1,732 |
| SALT00000002487_Cluster25694-001 | 0.040 | NA | 1,727 |
| SALT00000036925_Cluster25723-001 | 0.186 | NA | 1,726 |
| SALT00000013066_Cluster25840-001 | 0.282 | NA | 1,720 |
| SALT00000055608_Cluster26033-001 | 0.088 | NA | 1,710 |
| SALT00000057581_Cluster26034-001 | 0.152 | NA | 1,710 |
| SALT00000058357_Cluster26050-001 | 0.240 | NA | 1,709 |
| SALT00000036198_Cluster26056-001 | 0.009 | NA | 1,113 |
| SALT00000051057_Cluster26239-001 | 0.172 | NA | 1,697 |
| SALT00000017473_Cluster26405-001 | 0.192 | NA | 1,516 |
| SALT00000039855_Cluster26509-001 | 0.147 | NA | 1,684 |
| SALT00000012752_Cluster26856-001 | 0.276 | NA | 1,666 |
| SALT00000001166_Cluster27216-001 | 0.179 | NA | 1,648 |
| SALT00000013947_Cluster27241-001 | 0.139 | NA | 1,649 |
| SALT00000013261_Cluster27321-001 | 0.141 | NA | 1,644 |
| SALT00000043055_Cluster27374-001 | 0.210 | NA | 1,642 |
| SALT00000061314_Cluster27506-001 | 0.090 | NA | 1,636 |
| SALT00000043344_Cluster27534-001 | 0.054 | NA | 1,634 |
| SALT00000041289_Cluster27719-001 | 0.142 | NA | 1,625 |
| SALT00000047744_Cluster27719-002 | 0.219 | NA | 1,578 |
| SALT00000053588_Cluster27914-001 | 0.241 | NA | 1,615 |
| SALT00000040005_Cluster27933-001 | 0.230 | NA | 1,614 |
| SALT00000067911_Cluster28193-001 | 0.208 | NA | 1,601 |
| SALT00000004204_Cluster28209-001 | 0.282 | NA | 1,599 |
| SALT00000048490_Cluster28222-001 | 0.079 | NA | 1,599 |
| SALT00000008881_Cluster28267-001 | 0.248 | NA | 1,596 |
| SALT00000005019_Cluster28340-001 | 0.271 | NA | 1,592 |
| SALT00000057164_Cluster28409-001 | 0.129 | NA | 1,589 |
| SALT00000037126_Cluster28488-001 | 0.197 | NA | 1,585 |
| SALT00000044998_Cluster28503-001 | 0.282 | NA | 1,584 |
| SALT00000049380_Cluster28520-001 | 0.116 | NA | 1,583 |
| SALT00000013209_Cluster28550-001 | 0.170 | NA | 1,581 |
| SALT00000044472_Cluster28550-002 | 0.175 | NA | 1,538 |
| SALT00000047904_Cluster28655-001 | 0.296 | NA | 1,576 |
| SALT00000016519_Cluster28724-001 | 0.088 | NA | 1,572 |
| SALT00000004349_Cluster28758-001 | 0.016 | NA | 1,570 |
| SALT00000060950_Cluster28884-001 | 0.072 | NA | 1,565 |
| SALT00000006158_Cluster28890-001 | 0.167 | NA | 1,564 |
| SALT00000042655_Cluster28906-001 | 0.125 | NA | 1,564 |
| SALT00000008415_Cluster29045-001 | 0.129 | NA | 1,556 |
| SALT00000055235_Cluster29189-001 | 0.166 | NA | 1,550 |
| SALT00000002849_Cluster29244-001 | 0.140 | NA | 1,546 |
| SALT00000049840_Cluster29275-001 | 0.081 | NA | 1,545 |
| SALT00000044496_Cluster29330-001 | 0.110 | NA | 1,542 |
| SALT00000059813_Cluster29338-001 | 0.207 | NA | 1,542 |
| SALT00000053054_Cluster29501-001 | 0.119 | NA | 1,533 |
| SALT00000014010_Cluster29858-001 | 0.023 | NA | 1,511 |
| SALT00000016309_Cluster29914-001 | 0.008 | NA | 1,507 |

| | | | |
|---|---|---|---|
| SALT00000005094_Cluster29939-001 | 0.153 | NA | 1,505 |
| SALT00000056024_Cluster30006-001 | 0.144 | NA | 1,502 |
| SALT00000053342_Cluster30187-001 | 0.071 | NA | 1,489 |
| SALT00000051924_Cluster30386-001 | 0.182 | NA | 1,476 |
| SALT00000056929_Cluster30469-001 | 0.058 | NA | 1,470 |
| SALT00000012101_Cluster30614-001 | 0.121 | NA | 1,459 |
| SALT00000007549_Cluster30642-001 | 0.046 | NA | 1,456 |
| SALT00000051809_Cluster30649-001 | 0.048 | NA | 1,456 |
| SALT00000059078_Cluster30727-001 | 0.125 | NA | 1,450 |
| SALT00000050991_Cluster30758-001 | 0.041 | NA | 1,447 |
| SALT00000014511_Cluster30901-001 | 0.150 | NA | 1,430 |
| SALT00000057358_Cluster30909-001 | 0.049 | NA | 1,430 |
| SALT00000043082_Cluster31107-001 | 0.058 | NA | 1,407 |
| SALT00000040527_Cluster31205-001 | 0.284 | NA | 1,392 |
| SALT00000043752_Cluster31256-001 | 0.138 | NA | 1,384 |
| SALT00000053349_Cluster31577-001 | 0.202 | NA | 1,313 |
| SALT00000040103_Cluster32310-001 | 0.251 | NA | 1,062 |
| SALT00000049660_Cluster32312-001 | 0.774 | ppe-MIR166e\|osa-MIR166b\|bdi-MIR166b\| | 1,062 |
| SALT00000001762_Cluster32518-001 | 0.104 | NA | 865 |
| SALT00000063036_Cluster32577-001 | 0.056 | NA | 674 |
| SALT00000079288_Cluster32594-001 | 0.258 | NA | 549 |
