## Supplemental Table 4 for "The full-length transcriptome of *Spartina alterniflora* reveals the complexity of high salt tolerance in monocotyledonous halophyte"

### Table S4. List of Transcription factor coding transcripts in *Spartina alterniflora*.

*Spartina alterniflora* sujected to predict transcription factor functional domains using Plant Transcription Factor Database (PTFD, v4.0) with default parameters.

Detr: differential expression of transcripts compared to control conditions, the crateria of DE: FDR<0.05, |FC|>2.

| **Transcription factor transcripts** | **TF family** | **Description** | **Length (bp)** | **DEtr350** | **DEtr500** | **DEtr800** |
|---|---|---|---|---|---|---|
| SALT00000091208_Cluster3126-003 | AP2 | AP2-like ethylene-respons | 4545 | 0 | 0 | 0 |
| SALT00000081571_Cluster10360-001 | AP2 | transcription factor AP2 | 3056 | 0 | 0 | 0 |
| SALT00000063728_Cluster13745-001 | AP2 | partial [Oryza sativa Japon | 2712 | 0 | 0 | 0 |
| SALT00000066191_Cluster15664-001 | AP2 | 2-A [Triticum turgidum du | 2577 | 0 | 0 | 0 |
| SALT00000047845_Cluster16219-001 | AP2 | transcription factor AP2 | 2098 | 0 | 0 | 0 |
| SALT00000072959_Cluster16219-002 | AP2 | transcription factor AP2 | 2535 | 0 | 0 | 0 |
| SALT00000002521_Cluster16219-003 | AP2 | transcription factor AP2 | 1939 | 0 | 0 | 0 |
| SALT00000029618_Cluster17135-001 | AP2 | AP2-like ethylene-respons | 2455 | 0 | 0 | 0 |
| SALT00000041127_Cluster17135-002 | AP2 | AP2-like ethylene-respons | 1700 | 0 | 0 | 0 |
| SALT00000064511_Cluster17135-003 | AP2 | AP2-like ethylene-respons | 2414 | 0 | 0 | 0 |
| SALT00000079426_Cluster17135-004 | AP2 | AP2-like ethylene-respons | 2457 | 0 | 0 | 0 |
| SALT00000009377_Cluster19067-001 | AP2 | transcription factor AP2 | 1627 | 0 | 0 | 0 |
| SALT00000016313_Cluster19067-002 | AP2 | transcription factor AP2 | 1892 | 0 | down | 0 |
| SALT00000016980_Cluster19067-003 | AP2 | transcription factor AP2 | 1899 | 0 | 0 | 0 |
| SALT00000019051_Cluster19067-004 | AP2 | transcription factor AP2 | 2044 | 0 | 0 | 0 |
| SALT00000020188_Cluster19067-005 | AP2 | transcription factor AP2 | 1983 | 0 | 0 | 0 |
| SALT00000038737_Cluster19067-006 | AP2 | transcription factor AP2 | 2161 | 0 | 0 | 0 |
| SALT00000047763_Cluster19067-007 | AP2 | transcription factor AP2 | 1927 | up | up | 0 |
| SALT00000052554_Cluster19067-008 | AP2 | transcription factor AP2 | 1469 | 0 | 0 | 0 |
| SALT00000059471_Cluster19067-009 | AP2 | transcription factor AP2 | 2104 | 0 | 0 | 0 |
| SALT00000048252_Cluster19067-010 | AP2 | transcription factor AP2 | 2016 | 0 | 0 | 0 |
| SALT00000047484_Cluster19863-001 | AP2 | AP2-like ethylene-respons | 2091 | 0 | 0 | 0 |
| SALT00000062061_Cluster20895-001 | AP2 | Os04g0653600 [Oryza sati | 2012 | 0 | 0 | 0 |
| SALT00000050131_Cluster21204-001 | AP2 | ethylene-responsive transc | 1991 | 0 | 0 | 0 |
| SALT00000058978_Cluster21937-001 | AP2 | floral homeotic APETALA | 1944 | 0 | 0 | 0 |
| SALT00000009681_Cluster23292-001 | AP2 | transcription factor AP2 | 1859 | 0 | 0 | 0 |
| SALT00000018785_Cluster24288-001 | AP2 | AP2-like ethylene-respons | 1562 | 0 | 0 | 0 |
| SALT00000019495_Cluster24288-002 | AP2 | AP2-like ethylene-respons | 1803 | 0 | 0 | 0 |
| SALT00000018560_Cluster24288-003 | AP2 | AP2-like ethylene-respons | 1801 | 0 | 0 | 0 |
| SALT00000036544_Cluster24540-001 | AP2 | floral homeotic | 1444 | 0 | 0 | 0 |
| SALT00000045635_Cluster24540-002 | AP2 | AP2-like ethylene-respons | 1588 | 0 | 0 | 0 |
| SALT00000056550_Cluster24540-003 | AP2 | AP2-like ethylene-respons | 1788 | 0 | 0 | 0 |
| SALT00000057721_Cluster24540-004 | AP2 | spelt factor [Triticum aesti | 1538 | 0 | 0 | 0 |
| SALT00000038905_Cluster24540-005 | AP2 | AP2-like ethylene-respons | 1697 | 0 | 0 | 0 |
| SALT00000041118_Cluster25817-001 | AP2 | AP2-like ethylene-respons | 1663 | 0 | 0 | 0 |
| SALT00000055907_Cluster25817-002 | AP2 | AP2-like ethylene-respons | 1722 | 0 | 0 | 0 |
| SALT00000011965_Cluster25817-003 | AP2 | floral homeotic | 1650 | 0 | 0 | 0 |
| SALT00000060875_Cluster26185-001 | AP2 | AP2-like ethylene-respons | 1700 | 0 | 0 | 0 |
| SALT00000003546_Cluster27112-001 | AP2 | ethylene-responsive transc | 1654 | 0 | 0 | 0 |
| SALT00000049125_Cluster27476-001 | AP2 | AP2-like ethylene-respons | 1637 | 0 | 0 | 0 |
| SALT00000037563_Cluster27516-001 | AP2 | TPA: AP2 EREBP transcri | 1635 | 0 | 0 | 0 |
| SALT00000049364_Cluster27516-002 | AP2 | TPA: AP2 EREBP transcri | 1518 | 0 | 0 | 0 |
| SALT00000067445_Cluster27516-003 | AP2 | TPA: AP2 EREBP transcri | 1633 | 0 | 0 | 0 |
| SALT00000042173_Cluster28458-001 | AP2 | BABY BOOM 1 [Zea may | 1586 | 0 | 0 | 0 |

| | | | | | | |
|---|---|---|---|---|---|---|
| SALT00000066353_Cluster160-001 | ARF | Os04g0442000 [Oryza sati | 10077 | 0 | 0 | down |
| SALT00000031379_Cluster352-001 | ARF | auxin response factor 16 is | 3532 | 0 | 0 | 0 |
| SALT00000069202_Cluster352-003 | ARF | auxin response factor 16 is | 2960 | 0 | 0 | 0 |
| SALT00000074337_Cluster352-004 | ARF | auxin response factor 16 is | 3195 | 0 | 0 | 0 |
| SALT00000077945_Cluster352-005 | ARF | auxin response factor 16 is | 3941 | 0 | 0 | 0 |
| SALT00000086620_Cluster352-006 | ARF | auxin response factor 16 is | 3622 | 0 | 0 | 0 |
| SALT00000091699_Cluster352-008 | ARF | auxin response factor 16 is | 3526 | 0 | 0 | 0 |
| SALT00000092400_Cluster352-009 | ARF | auxin response factor 16 is | 3806 | 0 | 0 | 0 |
| SALT00000093076_Cluster352-010 | ARF | auxin response factor 16 is | 4403 | 0 | 0 | 0 |
| SALT00000091823_Cluster542-001 | ARF | auxin response factor 16 is | 4106 | 0 | 0 | 0 |
| SALT00000093282_Cluster542-002 | ARF | auxin response factor 16 is | 3971 | 0 | 0 | 0 |
| SALT00000028663_Cluster566-001 | ARF | auxin response factor 7a [( | 3943 | 0 | 0 | 0 |
| SALT00000029364_Cluster566-002 | ARF | auxin response factor 19-li | 3737 | 0 | 0 | 0 |
| SALT00000032067_Cluster566-003 | ARF | auxin response factor 19-li | 4059 | 0 | 0 | 0 |
| SALT00000032124_Cluster566-004 | ARF | auxin response factor 7a [( | 3865 | 0 | 0 | 0 |
| SALT00000032211_Cluster566-005 | ARF | auxin response factor 7a [( | 3979 | 0 | 0 | 0 |
| SALT00000032666_Cluster566-006 | ARF | auxin response factor 7a [( | 3936 | 0 | 0 | 0 |
| SALT00000032830_Cluster566-007 | ARF | auxin response factor 7a [( | 3684 | 0 | 0 | 0 |
| SALT00000033267_Cluster566-008 | ARF | auxin response factor 7a [( | 3615 | 0 | 0 | 0 |
| SALT00000033578_Cluster566-009 | ARF | auxin response factor 7a [( | 3811 | 0 | 0 | 0 |
| SALT00000034541_Cluster566-010 | ARF | auxin response factor 7a [( | 3748 | 0 | 0 | 0 |
| SALT00000065935_Cluster566-011 | ARF | auxin response factor 19-li | 3242 | 0 | 0 | 0 |
| SALT00000080460_Cluster566-014 | ARF | auxin response factor 7a [( | 3925 | 0 | 0 | 0 |
| SALT00000082481_Cluster566-015 | ARF | auxin response factor 19-li | 3749 | 0 | 0 | 0 |
| SALT00000083405_Cluster566-016 | ARF | auxin response factor 7a [( | 3785 | 0 | 0 | 0 |
| SALT00000085489_Cluster566-017 | ARF | auxin response factor 7a [( | 3831 | 0 | 0 | 0 |
| SALT00000087626_Cluster566-018 | ARF | auxin response factor 7a [( | 3873 | 0 | 0 | 0 |
| SALT00000089943_Cluster566-020 | ARF | auxin response factor 7a [( | 4015 | 0 | 0 | 0 |
| SALT00000093241_Cluster566-023 | ARF | auxin response factor 7a [( | 3605 | 0 | 0 | 0 |
| SALT00000093360_Cluster566-024 | ARF | auxin response factor 7a [( | 3998 | 0 | 0 | 0 |
| SALT00000033207_Cluster643-001 | ARF | auxin response factor 19-li | 4164 | 0 | 0 | 0 |
| SALT00000085750_Cluster643-002 | ARF | auxin response factor 7a [( | 3856 | 0 | 0 | 0 |
| SALT00000087214_Cluster643-003 | ARF | auxin response factor 19-li | 3926 | 0 | 0 | 0 |
| SALT00000087377_Cluster643-005 | ARF | auxin response factor 19-li | 4200 | 0 | 0 | 0 |
| SALT00000021090_Cluster679-001 | ARF | auxin response factor 24 is | 3041 | 0 | 0 | 0 |
| SALT00000022960_Cluster679-002 | ARF | auxin response factor 24 is | 3049 | 0 | 0 | 0 |
| SALT00000024632_Cluster679-003 | ARF | auxin response factor 24 is | 2910 | 0 | 0 | 0 |
| SALT00000031474_Cluster679-004 | ARF | Os12g0479400 [Oryza sati | 2932 | 0 | 0 | 0 |
| SALT00000062493_Cluster679-005 | ARF | Os12g0479400 [Oryza sati | 2996 | 0 | 0 | 0 |
| SALT00000067910_Cluster679-006 | ARF | auxin response factor 24 is | 3036 | 0 | 0 | 0 |
| SALT00000077555_Cluster679-008 | ARF | ARF transcription partial [ | 2540 | 0 | 0 | 0 |
| SALT00000080563_Cluster679-009 | ARF | auxin response factor 24 is | 3002 | 0 | 0 | 0 |
| SALT00000075855_Cluster679-011 | ARF | Os12g0479400 [Oryza sati | 3729 | 0 | 0 | 0 |
| SALT00000032370_Cluster1171-006 | ARF | auxin response factor 5 iso | 4080 | 0 | 0 | 0 |
| SALT00000034299_Cluster1171-007 | ARF | auxin response factor 5 iso | 4063 | 0 | 0 | 0 |
| SALT00000077663_Cluster1299-003 | ARF | auxin response factor 8-lik | 2494 | 0 | 0 | 0 |
| SALT00000072177_Cluster1299-004 | ARF | auxin response factor 8-lik | 2539 | 0 | 0 | 0 |
| SALT00000020693_Cluster1364-003 | ARF | Auxin response factor expı | 3158 | 0 | 0 | 0 |
| SALT00000020807_Cluster1364-004 | ARF | Auxin response factor expı | 3060 | 0 | 0 | 0 |
| SALT00000022951_Cluster1364-005 | ARF | Auxin response factor expı | 2963 | 0 | 0 | 0 |
| SALT00000023965_Cluster1364-006 | ARF | Auxin response factor expı | 2685 | 0 | 0 | 0 |
| SALT00000024464_Cluster1364-007 | ARF | Auxin response factor expı | 3141 | 0 | 0 | 0 |
| SALT00000027430_Cluster1364-008 | ARF | Auxin response factor expı | 3174 | 0 | 0 | 0 |
| SALT00000027582_Cluster1364-009 | ARF | Auxin response factor expı | 2862 | 0 | 0 | 0 |
| SALT00000028203_Cluster1364-010 | ARF | Auxin response factor expı | 2905 | 0 | 0 | 0 |
| SALT00000029045_Cluster1364-011 | ARF | Auxin response factor expı | 2947 | 0 | 0 | 0 |
| SALT00000030036_Cluster1364-012 | ARF | Auxin response factor expı | 3173 | 0 | 0 | 0 |
| SALT00000070170_Cluster1364-013 | ARF | Auxin response factor expı | 3091 | 0 | 0 | 0 |

| | | | | | | |
|---|---|---|---|---|---|---|
| SALT00000071341_Cluster1364-014 | ARF | Auxin response factor expı | 3090 | 0 | 0 | 0 |
| SALT00000073858_Cluster1364-015 | ARF | Auxin response factor expı | 3137 | 0 | 0 | 0 |
| SALT00000082628_Cluster1364-017 | ARF | Auxin response factor expı | 3159 | 0 | 0 | 0 |
| SALT00000084171_Cluster1364-018 | ARF | Auxin response factor expı | 3001 | 0 | 0 | 0 |
| SALT00000086582_Cluster1364-019 | ARF | Auxin response factor expı | 3285 | 0 | 0 | 0 |
| SALT00000092752_Cluster1364-020 | ARF | auxin response factor 23-li | 3220 | 0 | 0 | 0 |
| SALT00000092803_Cluster1364-021 | ARF | auxin response factor 23-li | 3475 | 0 | 0 | 0 |
| SALT00000025446_Cluster1665-001 | ARF | auxin response factor 24 is | 3087 | 0 | 0 | 0 |
| SALT00000027385_Cluster1665-002 | ARF | auxin response factor 24 is | 3066 | 0 | 0 | 0 |
| SALT00000069602_Cluster1665-006 | ARF | auxin response factor 24 is | 3400 | 0 | 0 | 0 |
| SALT00000078279_Cluster1665-008 | ARF | auxin response factor 24 is | 3066 | 0 | 0 | 0 |
| SALT00000091888_Cluster1665-009 | ARF | auxin response factor 24 is | 3811 | 0 | 0 | 0 |
| SALT00000092476_Cluster1665-010 | ARF | auxin response factor 24 is | 3206 | 0 | 0 | 0 |
| SALT00000074306_Cluster1665-011 | ARF | auxin response factor 24-li | 3347 | 0 | 0 | 0 |
| SALT00000022393_Cluster2284-002 | ARF | Os01g0927600 [Oryza sati | 2801 | 0 | 0 | 0 |
| SALT00000023630_Cluster2284-003 | ARF | auxin response factor 4-lik | 2943 | 0 | 0 | 0 |
| SALT00000023849_Cluster2284-004 | ARF | auxin response factor 4-lik | 2583 | 0 | 0 | 0 |
| SALT00000025055_Cluster2284-005 | ARF | auxin response factor 4-lik | 2953 | 0 | 0 | 0 |
| SALT00000025664_Cluster2284-006 | ARF | auxin response factor 4-lik | 2917 | 0 | 0 | 0 |
| SALT00000027352_Cluster2284-007 | ARF | Os01g0927600 [Oryza sati | 3063 | 0 | 0 | 0 |
| SALT00000028085_Cluster2284-008 | ARF | auxin response factor 4-lik | 2925 | 0 | 0 | 0 |
| SALT00000029166_Cluster2284-009 | ARF | auxin response factor 4-lik | 2940 | 0 | 0 | 0 |
| SALT00000030824_Cluster2284-010 | ARF | auxin response factor 4-lik | 2531 | 0 | 0 | 0 |
| SALT00000034651_Cluster2284-011 | ARF | auxin response factor 4-lik | 3002 | 0 | 0 | up |
| SALT00000034867_Cluster2284-012 | ARF | auxin response factor 4-lik | 2996 | 0 | 0 | up |
| SALT00000035060_Cluster2284-013 | ARF | auxin response factor 4-lik | 3036 | 0 | 0 | 0 |
| SALT00000035082_Cluster2284-014 | ARF | auxin response factor 4-lik | 3017 | 0 | 0 | 0 |
| SALT00000035534_Cluster2284-015 | ARF | auxin response factor 4-lik | 2975 | 0 | 0 | 0 |
| SALT00000063295_Cluster2284-017 | ARF | Os01g0927600 [Oryza sati | 3153 | 0 | 0 | 0 |
| SALT00000064091_Cluster2284-018 | ARF | auxin response factor 4-lik | 2963 | 0 | 0 | 0 |
| SALT00000067268_Cluster2284-019 | ARF | auxin response factor 4-lik | 2958 | 0 | 0 | 0 |
| SALT00000067535_Cluster2284-020 | ARF | auxin response factor 4-lik | 2969 | 0 | 0 | 0 |
| SALT00000069453_Cluster2284-021 | ARF | auxin response factor 4-lik | 2851 | 0 | 0 | 0 |
| SALT00000075788_Cluster2284-022 | ARF | auxin response factor 4-lik | 2921 | 0 | 0 | 0 |
| SALT00000076733_Cluster2284-024 | ARF | auxin response factor 4-lik | 2944 | 0 | 0 | 0 |
| SALT00000077677_Cluster2284-025 | ARF | auxin response factor 4-lik | 2927 | 0 | 0 | 0 |
| SALT00000078371_Cluster2284-026 | ARF | auxin response factor 4-lik | 2753 | 0 | 0 | 0 |
| SALT00000079589_Cluster2284-027 | ARF | auxin response factor 4-lik | 2643 | 0 | 0 | 0 |
| SALT00000082398_Cluster2284-028 | ARF | auxin response factor 4-lik | 2960 | 0 | 0 | 0 |
| SALT00000082758_Cluster2284-029 | ARF | auxin response factor 4-lik | 2983 | 0 | 0 | 0 |
| SALT00000082883_Cluster2284-030 | ARF | auxin response factor 4-lik | 3002 | 0 | 0 | 0 |
| SALT00000093917_Cluster2284-031 | ARF | auxin response factor 4-lik | 2975 | 0 | 0 | 0 |
| SALT00000084389_Cluster2284-033 | ARF | auxin response factor 4-lik | 2569 | 0 | 0 | 0 |
| SALT00000092198_Cluster2447-002 | ARF | auxin response factor 4-lik | 3268 | 0 | 0 | 0 |
| SALT00000020718_Cluster2610-001 | ARF | Os04g0442000 [Oryza sati | 2773 | 0 | 0 | 0 |
| SALT00000020824_Cluster2610-002 | ARF | Os04g0442000 [Oryza sati | 2650 | 0 | 0 | 0 |
| SALT00000022674_Cluster2610-003 | ARF | Os04g0442000 [Oryza sati | 2800 | 0 | 0 | 0 |
| SALT00000023055_Cluster2610-004 | ARF | Os04g0442000 [Oryza sati | 2640 | 0 | 0 | 0 |
| SALT00000023153_Cluster2610-005 | ARF | Os04g0442000 [Oryza sati | 2912 | 0 | 0 | 0 |
| SALT00000023580_Cluster2610-006 | ARF | Os04g0442000 [Oryza sati | 2977 | 0 | 0 | 0 |
| SALT00000026795_Cluster2610-007 | ARF | Os04g0442000 [Oryza sati | 2533 | 0 | 0 | 0 |
| SALT00000028002_Cluster2610-008 | ARF | Os04g0442000 [Oryza sati | 2849 | 0 | 0 | 0 |
| SALT00000028852_Cluster2610-009 | ARF | Os04g0442000 [Oryza sati | 2534 | 0 | 0 | 0 |
| SALT00000062440_Cluster2610-010 | ARF | Os04g0442000 [Oryza sati | 2610 | 0 | 0 | 0 |
| SALT00000074366_Cluster2610-011 | ARF | Os04g0442000 [Oryza sati | 2883 | 0 | 0 | 0 |
| SALT00000077540_Cluster2610-013 | ARF | Os04g0442000 [Oryza sati | 2837 | 0 | 0 | 0 |
| SALT00000080551_Cluster2610-014 | ARF | Os04g0442000 [Oryza sati | 2745 | 0 | 0 | 0 |
| SALT00000026114_Cluster2610-015 | ARF | Os04g0442000 [Oryza sati | 3124 | 0 | 0 | 0 |

| | | | | | | |
|---|---|---|---|---|---|---|
| SALT00000018800_Cluster2630-002 | ARF | Os02g0557200 [Oryza sati | 2099 | 0 | 0 | 0 |
| SALT00000021014_Cluster2630-003 | ARF | Os02g0557200 [Oryza sati | 2874 | 0 | 0 | 0 |
| SALT00000021249_Cluster2630-004 | ARF | Os02g0557200 [Oryza sati | 2794 | 0 | 0 | 0 |
| SALT00000021424_Cluster2630-005 | ARF | Os02g0557200 [Oryza sati | 2682 | 0 | 0 | 0 |
| SALT00000021524_Cluster2630-006 | ARF | Os02g0557200 [Oryza sati | 2882 | 0 | 0 | 0 |
| SALT00000022856_Cluster2630-007 | ARF | Os02g0557200 [Oryza sati | 2898 | 0 | 0 | 0 |
| SALT00000022961_Cluster2630-008 | ARF | Os02g0557200 [Oryza sati | 2813 | 0 | 0 | 0 |
| SALT00000022977_Cluster2630-009 | ARF | Os02g0557200 [Oryza sati | 2608 | 0 | 0 | 0 |
| SALT00000024056_Cluster2630-010 | ARF | Os02g0557200 [Oryza sati | 2806 | 0 | 0 | 0 |
| SALT00000024596_Cluster2630-011 | ARF | Os02g0557200 [Oryza sati | 2817 | 0 | 0 | 0 |
| SALT00000024656_Cluster2630-012 | ARF | Os02g0557200 [Oryza sati | 2902 | 0 | 0 | 0 |
| SALT00000024889_Cluster2630-013 | ARF | Os02g0557200 [Oryza sati | 2750 | 0 | 0 | 0 |
| SALT00000024991_Cluster2630-014 | ARF | Os02g0557200 [Oryza sati | 2842 | 0 | 0 | 0 |
| SALT00000026168_Cluster2630-015 | ARF | Os02g0557200 [Oryza sati | 2743 | 0 | 0 | 0 |
| SALT00000026593_Cluster2630-016 | ARF | Os02g0557200 [Oryza sati | 2801 | 0 | 0 | 0 |
| SALT00000027073_Cluster2630-017 | ARF | Os02g0557200 [Oryza sati | 2748 | 0 | 0 | 0 |
| SALT00000027144_Cluster2630-018 | ARF | Os02g0557200 [Oryza sati | 2706 | 0 | 0 | 0 |
| SALT00000027513_Cluster2630-019 | ARF | Os02g0557200 [Oryza sati | 2653 | 0 | 0 | 0 |
| SALT00000027922_Cluster2630-020 | ARF | Os02g0557200 [Oryza sati | 2810 | 0 | 0 | 0 |
| SALT00000027994_Cluster2630-021 | ARF | Os02g0557200 [Oryza sati | 2754 | 0 | 0 | 0 |
| SALT00000028808_Cluster2630-022 | ARF | Os02g0557200 [Oryza sati | 2803 | 0 | 0 | 0 |
| SALT00000029832_Cluster2630-023 | ARF | Os02g0557200 [Oryza sati | 2928 | 0 | 0 | 0 |
| SALT00000035305_Cluster2630-024 | ARF | Os02g0557200 [Oryza sati | 2713 | 0 | 0 | 0 |
| SALT00000035647_Cluster2630-025 | ARF | Os02g0557200 [Oryza sati | 2660 | 0 | 0 | 0 |
| SALT00000048994_Cluster2630-026 | ARF | Os02g0557200 [Oryza sati | 1079 | 0 | 0 | 0 |
| SALT00000062864_Cluster2630-027 | ARF | Os02g0557200 [Oryza sati | 2817 | 0 | 0 | 0 |
| SALT00000063593_Cluster2630-028 | ARF | Os02g0557200 [Oryza sati | 2888 | 0 | 0 | 0 |
| SALT00000065893_Cluster2630-029 | ARF | Os02g0557200 [Oryza sati | 2749 | 0 | 0 | 0 |
| SALT00000067281_Cluster2630-030 | ARF | Os02g0557200 [Oryza sati | 2491 | 0 | 0 | 0 |
| SALT00000069715_Cluster2630-031 | ARF | Os02g0557200 [Oryza sati | 2489 | 0 | 0 | 0 |
| SALT00000071375_Cluster2630-032 | ARF | Os02g0557200 [Oryza sati | 2772 | 0 | 0 | 0 |
| SALT00000079019_Cluster2630-033 | ARF | Os02g0557200 [Oryza sati | 2775 | 0 | 0 | 0 |
| SALT00000080812_Cluster2630-034 | ARF | Os02g0557200 [Oryza sati | 2792 | 0 | 0 | 0 |
| SALT00000090960_Cluster2630-036 | ARF | Os02g0557200 [Oryza sati | 3369 | 0 | 0 | 0 |
| SALT00000024685_Cluster2630-037 | ARF | Os02g0557200 [Oryza sati | 2527 | 0 | 0 | 0 |
| SALT00000079809_Cluster2630-038 | ARF | Os02g0557200 [Oryza sati | 2710 | 0 | 0 | 0 |
| SALT00000032589_Cluster2728-001 | ARF | Auxin response factor 5 | 3885 | 0 | 0 | 0 |
| SALT00000030827_Cluster2967-011 | ARF | Os04g0442000 [Oryza sati | 2945 | 0 | 0 | 0 |
| SALT00000064730_Cluster2967-016 | ARF | Os04g0442000 [Oryza sati | 3260 | 0 | 0 | 0 |
| SALT00000084224_Cluster2967-021 | ARF | transmembrane 9 superfan | 4611 | 0 | 0 | 0 |
| SALT00000030046_Cluster3043-001 | ARF | Auxin response factor 5 | 3863 | 0 | 0 | 0 |
| SALT00000032449_Cluster3043-002 | ARF | Auxin response factor 5 | 4056 | 0 | 0 | 0 |
| SALT00000086590_Cluster3043-004 | ARF | Auxin response factor 5 | 4102 | 0 | 0 | 0 |
| SALT00000089731_Cluster3043-005 | ARF | auxin response factor 5 iso | 4128 | 0 | 0 | 0 |
| SALT00000063345_Cluster3214-001 | ARF | Os04g0442000 [Oryza sati | 3097 | 0 | 0 | 0 |
| SALT00000065190_Cluster3214-003 | ARF | Os04g0442000 [Oryza sati | 2741 | 0 | 0 | 0 |
| SALT00000029056_Cluster3633-001 | ARF | Os12g0613700 [Oryza sati | 3869 | 0 | 0 | 0 |
| SALT00000030499_Cluster3633-002 | ARF | auxin response factor 25 | 2614 | 0 | 0 | 0 |
| SALT00000084258_Cluster3633-005 | ARF | Os12g0613700 [Oryza sati | 3726 | 0 | 0 | 0 |
| SALT00000084834_Cluster3633-006 | ARF | ARF transcription partial | 3894 | 0 | 0 | 0 |
| SALT00000087159_Cluster3633-007 | ARF | Os12g0613700 [Oryza sati | 4348 | 0 | 0 | 0 |
| SALT00000088539_Cluster3633-008 | ARF | Os12g0613700 [Oryza sati | 3915 | 0 | 0 | 0 |
| SALT00000022830_Cluster3664-001 | ARF | Os05g0563400 [Oryza sati | 2719 | 0 | 0 | 0 |
| SALT00000064215_Cluster3664-003 | ARF | Os05g0563400 [Oryza sati | 2654 | 0 | 0 | 0 |
| SALT00000082938_Cluster3664-004 | ARF | Os05g0563400 [Oryza sati | 2624 | 0 | 0 | 0 |
| SALT00000075593_Cluster3664-005 | ARF | Os05g0563400 [Oryza sati | 2790 | 0 | 0 | 0 |
| SALT00000032609_Cluster4483-001 | ARF | Os12g0613700 [Oryza sati | 3897 | 0 | 0 | 0 |
| SALT00000033247_Cluster4483-002 | ARF | Os12g0613700 [Oryza sati | 3980 | 0 | 0 | 0 |

| | | | | | | |
|---|---|---|---|---|---|---|
| SALT00000033674_Cluster4483-003 | ARF | Os12g0613700 [Oryza sati | 4035 | 0 | 0 | 0 |
| SALT00000034500_Cluster4483-004 | ARF | Os12g0613700 [Oryza sati | 4007 | 0 | 0 | 0 |
| SALT00000066911_Cluster4483-005 | ARF | Os12g0613700 [Oryza sati | 3133 | 0 | 0 | 0 |
| SALT00000086339_Cluster4483-006 | ARF | Os12g0613700 [Oryza sati | 4094 | 0 | 0 | 0 |
| SALT00000090684_Cluster4483-007 | ARF | ARF transcription partial | 3925 | 0 | 0 | 0 |
| SALT00000093008_Cluster4483-008 | ARF | Os12g0613700 [Oryza sati | 3979 | 0 | 0 | 0 |
| SALT00000090833_Cluster4802-001 | ARF | auxin response factor 4-lik | 3607 | 0 | 0 | 0 |
| SALT00000068557_Cluster4802-004 | ARF | auxin response factor 4-lik | 3147 | 0 | 0 | 0 |
| SALT00000034053_Cluster4919-001 | ARF | auxin response factor 5-lik | 3992 | 0 | 0 | 0 |
| SALT00000080734_Cluster4919-002 | ARF | auxin response factor 5-lik | 3944 | 0 | 0 | 0 |
| SALT00000086382_Cluster4919-003 | ARF | auxin response factor 5-lik | 3947 | 0 | 0 | 0 |
| SALT00000086636_Cluster4919-004 | ARF | auxin response factor 5-lik | 3818 | 0 | 0 | 0 |
| SALT00000090235_Cluster4919-005 | ARF | auxin response factor 5-lik | 3875 | 0 | 0 | 0 |
| SALT00000031109_Cluster5580-001 | ARF | auxin response transcriptio | 3611 | 0 | 0 | 0 |
| SALT00000033599_Cluster5580-002 | ARF | auxin response transcriptio | 3637 | 0 | 0 | 0 |
| SALT00000084829_Cluster5580-003 | ARF | auxin response transcriptio | 3650 | 0 | 0 | 0 |
| SALT00000085231_Cluster5580-004 | ARF | auxin response transcriptio | 3746 | 0 | 0 | 0 |
| SALT00000085553_Cluster5580-005 | ARF | auxin response transcriptio | 3648 | 0 | 0 | 0 |
| SALT00000087969_Cluster5580-006 | ARF | auxin response transcriptio | 3867 | 0 | 0 | 0 |
| SALT00000090097_Cluster5580-007 | ARF | auxin response transcriptio | 3646 | 0 | 0 | down |
| SALT00000023688_Cluster5791-002 | ARF | auxin response factor [Ory | 2503 | 0 | 0 | 0 |
| SALT00000030171_Cluster5791-003 | ARF | auxin response factor [Ory | 3416 | 0 | 0 | 0 |
| SALT00000030464_Cluster5791-004 | ARF | auxin response factor [Ory | 3635 | 0 | 0 | 0 |
| SALT00000032049_Cluster5791-005 | ARF | auxin response factor [Ory | 3602 | 0 | 0 | 0 |
| SALT00000032905_Cluster5791-006 | ARF | auxin response factor [Ory | 3504 | 0 | 0 | 0 |
| SALT00000033161_Cluster5791-007 | ARF | auxin response factor [Ory | 3741 | 0 | 0 | 0 |
| SALT00000066100_Cluster5791-008 | ARF | auxin response factor [Ory | 3369 | 0 | 0 | 0 |
| SALT00000074719_Cluster5791-009 | ARF | auxin response factor [Ory | 2634 | 0 | 0 | 0 |
| SALT00000075663_Cluster5791-010 | ARF | auxin response factor [Ory | 2392 | 0 | 0 | 0 |
| SALT00000077394_Cluster5791-011 | ARF | auxin response factor [Ory | 3505 | 0 | 0 | 0 |
| SALT00000088325_Cluster5791-012 | ARF | auxin response factor [Ory | 3556 | 0 | 0 | 0 |
| SALT00000088452_Cluster5791-013 | ARF | auxin response factor [Ory | 3787 | 0 | 0 | 0 |
| SALT00000089138_Cluster5791-014 | ARF | auxin response factor [Ory | 3590 | 0 | 0 | 0 |
| SALT00000089870_Cluster5791-015 | ARF | auxin response factor [Ory | 3829 | 0 | 0 | 0 |
| SALT00000090320_Cluster5791-016 | ARF | auxin response factor [Ory | 3388 | 0 | 0 | 0 |
| SALT00000091508_Cluster5791-017 | ARF | auxin response factor [Ory | 3687 | 0 | 0 | 0 |
| SALT00000092045_Cluster5791-018 | ARF | auxin response factor [Ory | 3519 | 0 | 0 | 0 |
| SALT00000077821_Cluster5791-019 | ARF | auxin response factor [Ory | 3523 | 0 | 0 | 0 |
| SALT00000033609_Cluster5817-001 | ARF | auxin response transcriptio | 3657 | 0 | 0 | 0 |
| SALT00000086512_Cluster5817-002 | ARF | auxin response transcriptio | 3824 | 0 | 0 | 0 |
| SALT00000091302_Cluster5817-003 | ARF | auxin response transcriptio | 3771 | 0 | 0 | 0 |
| SALT00000087564_Cluster5995-001 | ARF | auxin response factor 2 [Ze | 3741 | 0 | 0 | 0 |
| SALT00000093439_Cluster5995-002 | ARF | auxin response factor 2 [Ze | 3794 | 0 | 0 | 0 |
| SALT00000030199_Cluster6201-002 | ARF | auxin response factor | 3761 | 0 | 0 | 0 |
| SALT00000052015_Cluster6201-003 | ARF | auxin response factor | 2312 | 0 | 0 | 0 |
| SALT00000021295_Cluster6201-004 | ARF | auxin response factor | 3014 | down | 0 | down |
| SALT00000032974_Cluster6614-001 | ARF | auxin response factor [Ory | 3532 | 0 | 0 | 0 |
| SALT00000083555_Cluster6614-003 | ARF | auxin response factor [Ory | 3626 | 0 | 0 | 0 |
| SALT00000089126_Cluster6614-004 | ARF | auxin response factor [Ory | 3696 | 0 | 0 | 0 |
| SALT00000093694_Cluster6614-005 | ARF | auxin response factor [Ory | 3555 | 0 | 0 | 0 |
| SALT00000031917_Cluster6688-001 | ARF | auxin response factor [Ory | 3576 | 0 | 0 | 0 |
| SALT00000034161_Cluster6688-002 | ARF | auxin response factor [Ory | 3683 | 0 | 0 | 0 |
| SALT00000084756_Cluster6688-005 | ARF | auxin response factor [Ory | 3647 | 0 | 0 | 0 |
| SALT00000075533_Cluster7298-002 | ARF | auxin response factor 3-lik | 3138 | 0 | 0 | 0 |
| SALT00000063562_Cluster7298-003 | ARF | auxin response factor 3-lik | 2933 | 0 | 0 | 0 |
| SALT00000064988_Cluster8135-001 | ARF | Os04g0442000 [Oryza sati | 3454 | 0 | 0 | 0 |
| SALT00000023002_Cluster9595-001 | ARF | auxin response factor 3-lik | 2989 | 0 | 0 | 0 |
| SALT00000031010_Cluster9595-002 | ARF | auxin response factor 3-lik | 3032 | 0 | 0 | 0 |

| | | | | | | |
|---|---|---|---|---|---|---|
| SALT00000065060_Cluster9595-003 | ARF | auxin response factor 3-lik | 2931 | 0 | 0 | 0 |
| SALT00000078936_Cluster9595-004 | ARF | auxin response factor 3-lik | 3174 | 0 | 0 | 0 |
| SALT00000021746_Cluster9674-002 | ARF | auxin response factor 4-lik | 3068 | 0 | 0 | 0 |
| SALT00000065543_Cluster9674-003 | ARF | auxin response factor 4-lik | 3162 | 0 | 0 | 0 |
| SALT00000065687_Cluster9674-004 | ARF | auxin response factor 4-lik | 2991 | 0 | 0 | 0 |
| SALT00000070546_Cluster9674-005 | ARF | auxin response factor 4-lik | 3022 | 0 | 0 | 0 |
| SALT00000072252_Cluster9674-006 | ARF | auxin response factor 4-lik | 3049 | 0 | 0 | 0 |
| SALT00000084333_Cluster9674-007 | ARF | auxin response factor 4-lik | 3026 | 0 | 0 | 0 |
| SALT00000062375_Cluster10040-001 | ARF | auxin response factor | 2995 | 0 | 0 | 0 |
| SALT00000070488_Cluster10040-002 | ARF | Os04g0671900 [Oryza sati | 3103 | 0 | 0 | 0 |
| SALT00000071362_Cluster10040-003 | ARF | Os04g0671900 [Oryza sati | 2932 | 0 | 0 | 0 |
| SALT00000084119_Cluster10040-004 | ARF | auxin response factor | 2485 | 0 | 0 | 0 |
| SALT00000024054_Cluster10046-001 | ARF | auxin response factor 2 [Ze | 2325 | 0 | 0 | 0 |
| SALT00000065091_Cluster10046-003 | ARF | auxin response factor 2 [Ze | 2468 | 0 | 0 | 0 |
| SALT00000067978_Cluster10046-004 | ARF | auxin response factor 2 [Ze | 2715 | 0 | 0 | 0 |
| SALT00000072853_Cluster10046-005 | ARF | auxin response factor 2 [Ze | 2589 | 0 | 0 | 0 |
| SALT00000074585_Cluster10125-001 | ARF | Os02g0557200 [Oryza sati | 3091 | 0 | 0 | 0 |
| SALT00000075651_Cluster10125-002 | ARF | Os02g0557200 [Oryza sati | 3014 | 0 | 0 | 0 |
| SALT00000084037_Cluster10125-003 | ARF | Os02g0557200 [Oryza sati | 3058 | 0 | 0 | 0 |
| SALT00000063604_Cluster12111-001 | ARF | Os02g0557200 [Oryza sati | 2857 | 0 | 0 | 0 |
| SALT00000066274_Cluster12802-001 | ARF | auxin response factor 21 is | 2721 | 0 | 0 | 0 |
| SALT00000070647_Cluster12802-002 | ARF | auxin response factor 21 is | 2515 | 0 | 0 | 0 |
| SALT00000072272_Cluster12802-003 | ARF | auxin response factor 21 is | 2749 | 0 | 0 | 0 |
| SALT00000079129_Cluster12802-004 | ARF | auxin response factor 21 is | 2792 | 0 | 0 | 0 |
| SALT00000022263_Cluster13038-001 | ARF | auxin response factor 2 [Ze | 2641 | 0 | 0 | 0 |
| SALT00000029481_Cluster13038-002 | ARF | auxin response factor 2 [Ze | 2616 | 0 | 0 | 0 |
| SALT00000066293_Cluster13038-003 | ARF | auxin response factor 2 [Ze | 2631 | 0 | 0 | 0 |
| SALT00000073745_Cluster13038-004 | ARF | auxin response factor 2 [Ze | 2770 | 0 | 0 | 0 |
| SALT00000028342_Cluster13233-003 | ARF | Os01g0236300 [Oryza sati | 2644 | 0 | 0 | 0 |
| SALT00000064746_Cluster13233-004 | ARF | Os01g0236300 [Oryza sati | 2602 | 0 | 0 | 0 |
| SALT00000068338_Cluster13233-005 | ARF | Os01g0236300 [Oryza sati | 2528 | 0 | 0 | 0 |
| SALT00000071587_Cluster13233-006 | ARF | Os01g0236300 [Oryza sati | 2528 | 0 | 0 | 0 |
| SALT00000079343_Cluster13233-007 | ARF | Os01g0236300 [Oryza sati | 2755 | 0 | 0 | 0 |
| SALT00000062557_Cluster14483-001 | ARF | auxin response factor 21 is | 2602 | 0 | 0 | 0 |
| SALT00000064879_Cluster14483-002 | ARF | auxin response factor 21 is | 2660 | 0 | 0 | 0 |
| SALT00000025191_Cluster15150-001 | ARF | [Oryza sativa Japonica Grc | 2484 | 0 | 0 | 0 |
| SALT00000031225_Cluster15150-002 | ARF | [Oryza sativa Japonica Grc | 2588 | 0 | 0 | 0 |
| SALT00000039320_Cluster15150-003 | ARF | [Oryza sativa Japonica Grc | 2072 | 0 | 0 | 0 |
| SALT00000077325_Cluster15150-004 | ARF | [Oryza sativa Japonica Grc | 2405 | 0 | 0 | 0 |
| SALT00000081860_Cluster15150-005 | ARF | [Oryza sativa Japonica Grc | 2615 | 0 | 0 | 0 |
| SALT00000064869_Cluster15150-006 | ARF | auxin response factor 8-lik | 2571 | 0 | 0 | 0 |
| SALT00000064991_Cluster15150-007 | ARF | [Oryza sativa Japonica Grc | 2697 | 0 | 0 | 0 |
| SALT00000021159_Cluster15758-001 | ARF | Os01g0236300 [Oryza sati | 2554 | 0 | 0 | 0 |
| SALT00000062996_Cluster15758-002 | ARF | Os01g0236300 [Oryza sati | 2569 | 0 | 0 | 0 |
| SALT00000069546_Cluster15758-003 | ARF | Os01g0236300 [Oryza sati | 2509 | 0 | 0 | 0 |
| SALT00000023558_Cluster16239-001 | ARF | auxin response factor 8-lik | 2493 | 0 | 0 | 0 |
| SALT00000029443_Cluster16239-002 | ARF | auxin response factor 8-lik | 2522 | 0 | 0 | 0 |
| SALT00000056080_Cluster16239-003 | ARF | auxin response factor 8-lik | 2299 | 0 | 0 | 0 |
| SALT00000077143_Cluster16239-004 | ARF | auxin response factor 8-lik | 2534 | 0 | 0 | 0 |
| SALT00000068873_Cluster17255-002 | ARF | auxin response factor 23 [Z | 2442 | 0 | 0 | 0 |
| SALT00000041200_Cluster21932-001 | ARF | B3 DNA binding domain c | 1944 | 0 | 0 | 0 |
| SALT00000040158_Cluster7076-001 | ARR-B | two-component response r | 2162 | 0 | 0 | 0 |
| SALT00000068844_Cluster7076-003 | ARR-B | two-component response r | 2697 | 0 | 0 | 0 |
| SALT00000075674_Cluster7076-005 | ARR-B | two-component response r | 2715 | 0 | 0 | 0 |
| SALT00000059213_Cluster9096-001 | ARR-B | two-component response r | 2042 | 0 | 0 | 0 |
| SALT00000067333_Cluster9096-002 | ARR-B | two-component response r | 2703 | 0 | 0 | 0 |
| SALT00000068628_Cluster9096-003 | ARR-B | two-component response r | 2880 | 0 | 0 | 0 |
| SALT00000073668_Cluster9096-004 | ARR-B | two-component response r | 3268 | 0 | 0 | 0 |

| | | | | | | |
|---|---|---|---|---|---|---|
| SALT00000015760_Cluster10783-001 | ARR-B | two-component response r | 2059 | 0 | 0 | 0 |
| SALT00000031266_Cluster10783-002 | ARR-B | two-component response r | 2988 | 0 | 0 | 0 |
| SALT00000076701_Cluster10783-003 | ARR-B | two-component response r | 3000 | 0 | 0 | 0 |
| SALT00000073523_Cluster10783-004 | ARR-B | two-component response r | 2709 | 0 | 0 | 0 |
| SALT00000063634_Cluster11277-001 | ARR-B | two-component response r | 2385 | 0 | 0 | 0 |
| SALT00000069941_Cluster11277-002 | ARR-B | two-component response r | 2938 | 0 | 0 | 0 |
| SALT00000023503_Cluster11857-001 | ARR-B | Os06g0183100 [Oryza sati | 2881 | 0 | 0 | 0 |
| SALT00000026674_Cluster11857-002 | ARR-B | Os06g0183100 [Oryza sati | 2739 | 0 | 0 | 0 |
| SALT00000026757_Cluster11857-003 | ARR-B | Os06g0183100 [Oryza sati | 2635 | 0 | 0 | 0 |
| SALT00000069515_Cluster11857-004 | ARR-B | Os06g0183100 [Oryza sati | 2685 | 0 | 0 | 0 |
| SALT00000073507_Cluster11857-005 | ARR-B | Os06g0183100 [Oryza sati | 2528 | 0 | 0 | 0 |
| SALT00000093704_Cluster11857-006 | ARR-B | Os06g0183100 [Oryza sati | 2631 | 0 | 0 | 0 |
| SALT00000010284_Cluster12663-001 | ARR-B | myb-like DNA-binding SH | 2002 | 0 | 0 | 0 |
| SALT00000022453_Cluster12663-002 | ARR-B | myb-like DNA-binding SH | 2459 | 0 | 0 | 0 |
| SALT00000049051_Cluster12663-003 | ARR-B | myb-like DNA-binding SH | 2333 | 0 | 0 | 0 |
| SALT00000064326_Cluster12663-004 | ARR-B | myb-like DNA-binding SH | 2615 | 0 | 0 | 0 |
| SALT00000069182_Cluster12663-005 | ARR-B | myb-like DNA-binding SH | 2805 | 0 | 0 | 0 |
| SALT00000071008_Cluster12663-006 | ARR-B | myb-like DNA-binding SH | 2364 | 0 | 0 | 0 |
| SALT00000073330_Cluster12663-007 | ARR-B | myb-like DNA-binding SH | 2542 | 0 | 0 | 0 |
| SALT00000076250_Cluster12663-008 | ARR-B | myb-like DNA-binding SH | 2624 | 0 | 0 | 0 |
| SALT00000077820_Cluster12663-009 | ARR-B | myb-like DNA-binding SH | 2668 | 0 | 0 | 0 |
| SALT00000083498_Cluster12663-010 | ARR-B | myb-like DNA-binding SH | 2582 | 0 | 0 | 0 |
| SALT00000083646_Cluster12663-011 | ARR-B | myb-like DNA-binding SH | 2482 | 0 | 0 | 0 |
| SALT00000071943_Cluster12663-012 | ARR-B | response regulator 8 [Zea r | 2607 | 0 | 0 | 0 |
| SALT00000072167_Cluster12663-013 | ARR-B | myb-like DNA-binding SH | 2501 | 0 | 0 | 0 |
| SALT00000021468_Cluster14614-001 | ARR-B | response regulator 9 [Oryz | 2651 | 0 | 0 | 0 |
| SALT00000069377_Cluster14614-002 | ARR-B | response regulator 9 [Oryz | 2651 | 0 | 0 | 0 |
| SALT00000084621_Cluster14614-003 | ARR-B | response regulator 9 [Oryz | 2598 | 0 | 0 | 0 |
| SALT00000022233_Cluster16352-001 | ARR-B | two-component response r | 2371 | 0 | 0 | 0 |
| SALT00000023908_Cluster16352-002 | ARR-B | two-component response r | 2405 | 0 | 0 | 0 |
| SALT00000068782_Cluster16352-003 | ARR-B | two-component response r | 2429 | 0 | 0 | 0 |
| SALT00000069036_Cluster16352-004 | ARR-B | two-component response r | 2527 | 0 | 0 | 0 |
| SALT00000072098_Cluster16352-005 | ARR-B | two-component response r | 2634 | 0 | 0 | 0 |
| SALT00000077955_Cluster16424-003 | ARR-B | two-component response r | 2522 | 0 | 0 | 0 |
| SALT00000025461_Cluster1439-001 | B3 | B3 domain-containing OsC | 3259 | 0 | 0 | 0 |
| SALT00000028791_Cluster1439-002 | B3 | B3 domain-containing OsC | 2779 | 0 | 0 | 0 |
| SALT00000028796_Cluster1439-003 | B3 | B3 domain-containing OsC | 3285 | 0 | 0 | 0 |
| SALT00000030111_Cluster1439-004 | B3 | B3 domain-containing OsC | 3293 | 0 | 0 | 0 |
| SALT00000033455_Cluster1439-005 | B3 | B3 domain-containing OsC | 3329 | 0 | 0 | 0 |
| SALT00000073545_Cluster1439-007 | B3 | B3 domain-containing OsC | 3389 | 0 | 0 | 0 |
| SALT00000081995_Cluster1439-008 | B3 | B3 domain-containing OsC | 3098 | 0 | 0 | 0 |
| SALT00000082486_Cluster1439-009 | B3 | B3 domain-containing OsC | 2614 | 0 | 0 | 0 |
| SALT00000084238_Cluster1439-010 | B3 | B3 domain-containing OsC | 3393 | 0 | 0 | 0 |
| SALT00000024117_Cluster2196-001 | B3 | B3 domain-containing OsC | 2708 | 0 | 0 | 0 |
| SALT00000024350_Cluster2196-002 | B3 | B3 domain-containing OsC | 3085 | 0 | 0 | 0 |
| SALT00000063213_Cluster2196-003 | B3 | B3 domain-containing OsC | 2712 | 0 | 0 | 0 |
| SALT00000064770_Cluster2196-004 | B3 | B3 domain-containing OsC | 2621 | 0 | 0 | 0 |
| SALT00000064810_Cluster2196-005 | B3 | B3 domain-containing OsC | 2722 | 0 | 0 | 0 |
| SALT00000074520_Cluster2196-006 | B3 | B3 domain-containing OsC | 2506 | 0 | 0 | 0 |
| SALT00000074744_Cluster2196-007 | B3 | B3 domain-containing OsC | 2581 | 0 | 0 | 0 |
| SALT00000023798_Cluster2196-009 | B3 | B3 domain-containing OsC | 4585 | 0 | 0 | down |
| SALT00000018873_Cluster2555-001 | B3 | Os07g0679700 [Oryza sati | 2000 | 0 | 0 | 0 |
| SALT00000023429_Cluster2555-002 | B3 | Os07g0679700 [Oryza sati | 3232 | 0 | 0 | 0 |
| SALT00000026380_Cluster2555-003 | B3 | Os07g0679700 [Oryza sati | 3015 | 0 | 0 | 0 |
| SALT00000026647_Cluster2555-004 | B3 | Os07g0679700 [Oryza sati | 2965 | 0 | 0 | 0 |
| SALT00000029449_Cluster2555-005 | B3 | Os07g0679700 [Oryza sati | 3770 | 0 | 0 | 0 |
| SALT00000032648_Cluster2555-006 | B3 | Os07g0679700 [Oryza sati | 3404 | 0 | 0 | 0 |
| SALT00000046225_Cluster2555-008 | B3 | B3 domain-containing OsC | 2086 | 0 | 0 | 0 |

| | | | | | | |
|---|---|---|---|---|---|---|
| SALT00000082074_Cluster2555-010 | B3 | Os07g0679700 [Oryza sati | 3024 | 0 | 0 | 0 |
| SALT00000086971_Cluster2555-011 | B3 | Os07g0679700 [Oryza sati | 3690 | 0 | 0 | 0 |
| SALT00000077830_Cluster2797-001 | B3 | B3 domain-containing Os0 | 3217 | 0 | 0 | 0 |
| SALT00000087845_Cluster2797-003 | B3 | B3 domain-containing Os0 | 4103 | 0 | 0 | 0 |
| SALT00000067621_Cluster3779-001 | B3 | B3 domain-containing Os0 | 4297 | 0 | 0 | down |
| SALT00000090014_Cluster6067-001 | B3 | F-box LRR-repeat 17-like | 3781 | 0 | 0 | 0 |
| SALT00000009242_Cluster6470-001 | B3 | B3 domain-containing Os0 | 2065 | 0 | 0 | 0 |
| SALT00000084010_Cluster6470-002 | B3 | B3 domain-containing Os0 | 3717 | 0 | 0 | 0 |
| SALT00000091795_Cluster6470-003 | B3 | B3 domain-containing Os0 | 3277 | 0 | 0 | 0 |
| SALT00000074835_Cluster6747-001 | B3 | Os02g0557200 [Oryza sati | 3675 | 0 | 0 | 0 |
| SALT00000021433_Cluster8813-001 | B3 | Os07g0679700 [Oryza sati | 3173 | 0 | 0 | 0 |
| SALT00000033702_Cluster8813-002 | B3 | Os07g0679700 [Oryza sati | 3323 | 0 | 0 | 0 |
| SALT00000089801_Cluster8813-003 | B3 | Os07g0679700 [Oryza sati | 3316 | 0 | 0 | 0 |
| SALT00000029271_Cluster9688-001 | B3 | B3 domain-containing Os0 | 2911 | 0 | 0 | 0 |
| SALT00000071686_Cluster9688-002 | B3 | B3 domain-containing Os0 | 3032 | 0 | 0 | 0 |
| SALT00000076384_Cluster9688-003 | B3 | B3 domain-containing Os0 | 3058 | 0 | 0 | 0 |
| SALT00000082324_Cluster9688-004 | B3 | B3 domain-containing Os0 | 3161 | 0 | 0 | 0 |
| SALT00000083239_Cluster9688-005 | B3 | B3 domain-containing Os0 | 2924 | 0 | 0 | 0 |
| SALT00000021847_Cluster10515-001 | B3 | Os07g0679700 [Oryza sati | 3034 | 0 | 0 | 0 |
| SALT00000068843_Cluster10515-002 | B3 | Os07g0679700 [Oryza sati | 2910 | 0 | 0 | 0 |
| SALT00000069251_Cluster10515-003 | B3 | Os07g0679700 [Oryza sati | 3024 | 0 | 0 | 0 |
| SALT00000073438_Cluster10515-004 | B3 | Os07g0679700 [Oryza sati | 3021 | 0 | 0 | 0 |
| SALT00000069085_Cluster11711-001 | B3 | Os07g0679700 [Oryza sati | 2893 | 0 | 0 | 0 |
| SALT00000010545_Cluster20962-001 | B3 | Os05g0481400 [Oryza sati | 2007 | 0 | 0 | 0 |
| SALT00000015250_Cluster22088-001 | B3 | Os02g0683500 [Oryza sati | 1743 | 0 | 0 | 0 |
| SALT00000043912_Cluster22088-002 | B3 | B3 domain-containing Os0 | 1933 | 0 | 0 | 0 |
| SALT00000008376_Cluster24186-001 | B3 | B3 domain-containing Os0 | 1747 | 0 | 0 | 0 |
| SALT00000010166_Cluster24186-002 | B3 | B3 domain-containing Os0 | 1808 | 0 | 0 | 0 |
| SALT00000019385_Cluster24252-001 | B3 | B3 domain-containing Os0 | 1805 | 0 | 0 | 0 |
| SALT00000059457_Cluster24459-005 | B3 | AP2 EREBP transcription | 1033 | 0 | 0 | 0 |
| SALT00000003491_Cluster24640-001 | B3 | B3 DNA binding domain f | 1385 | 0 | 0 | 0 |
| SALT00000019096_Cluster25008-001 | B3 | B3 domain-containing Os0 | 1762 | 0 | 0 | 0 |
| SALT00000036303_Cluster25047-001 | B3 | ABI3VP1 transcription pa | 1760 | 0 | 0 | 0 |
| SALT00000038078_Cluster25047-002 | B3 | ABI3VP1 transcription pa | 1561 | 0 | 0 | 0 |
| SALT00000042396_Cluster25336-001 | B3 | B3 domain-containing Os0 | 1745 | 0 | 0 | 0 |
| SALT00000007824_Cluster26433-001 | B3 | B3 domain-containing Os0 | 1687 | 0 | 0 | up |
| SALT00000010714_Cluster26499-001 | B3 | Os11g0197600 [Oryza sati | 1684 | 0 | 0 | 0 |
| SALT00000012783_Cluster26499-002 | B3 | Os11g0197600 [Oryza sati | 1525 | 0 | 0 | 0 |
| SALT00000005117_Cluster29341-001 | B3 | B3 DNA binding domain f | 1541 | 0 | 0 | 0 |
| SALT00000019859_Cluster29341-002 | B3 | B3 DNA binding domain f | 1507 | 0 | 0 | 0 |
| SALT00000045372_Cluster31063-001 | B3 | B3 domain-containing Os0 | 1413 | 0 | 0 | 0 |
| SALT00000009141_Cluster24398-001 | BBR-BP | GAGA-binding [Zea mays | 1536 | 0 | 0 | 0 |
| SALT00000010133_Cluster24398-002 | BBR-BP | GAGA-binding [Zea mays | 1556 | 0 | 0 | 0 |
| SALT00000040179_Cluster24398-003 | BBR-BP | GAGA-binding [Zea mays | 1796 | 0 | 0 | 0 |
| SALT00000041226_Cluster24398-004 | BBR-BP | GAGA-binding [Zea mays | 1542 | 0 | 0 | 0 |
| SALT00000046791_Cluster24398-005 | BBR-BP | GAGA-binding [Zea mays | 1539 | 0 | 0 | 0 |
| SALT00000054117_Cluster24398-006 | BBR-BP | GAGA-binding [Zea mays | 1532 | 0 | 0 | 0 |
| SALT00000054239_Cluster24398-007 | BBR-BP | GAGA-binding [Zea mays | 1511 | 0 | 0 | 0 |
| SALT00000059870_Cluster24398-008 | BBR-BP | GAGA-binding [Zea mays | 1607 | 0 | 0 | 0 |
| SALT00000052240_Cluster24398-009 | BBR-BP | GAGA-binding [Zea mays | 1466 | 0 | 0 | 0 |
| SALT00000004239_Cluster24398-010 | BBR-BP | GAGA-binding [Zea mays | 1535 | 0 | 0 | 0 |
| SALT00000043714_Cluster24398-011 | BBR-BP | barley B recombinant A [C | 1577 | 0 | 0 | 0 |
| SALT00000039627_Cluster25223-001 | BBR-BP | barley B recombinant D [C | 1751 | 0 | 0 | 0 |
| SALT00000041826_Cluster25223-002 | BBR-BP | barley B recombinant D [C | 1546 | 0 | 0 | 0 |
| SALT00000012891_Cluster25223-003 | BBR-BP | barley B recombinant D [C | 1689 | 0 | 0 | 0 |
| SALT00000002527_Cluster3903-001 | BES1 | partial [Oryza sativa Japon | 1564 | 0 | 0 | 0 |
| SALT00000020275_Cluster3903-002 | BES1 | partial [Oryza sativa Japon | 1402 | 0 | 0 | 0 |
| SALT00000067428_Cluster9829-001 | BES1 | Os02g0129600 [Oryza sati | 3135 | 0 | 0 | 0 |

| | | | | | | |
|---|---|---|---|---|---|---|
| SALT00000006778_Cluster12933-001 | BES1 | partial [Oryza sativa Japon | 1575 | 0 | 0 | 0 |
| SALT00000002255_Cluster12933-003 | BES1 | partial [Oryza sativa Japon | 1421 | 0 | 0 | 0 |
| SALT00000003533_Cluster14718-001 | BES1 | brassinazole-resistant 1 [Z | 1543 | 0 | 0 | 0 |
| SALT00000042999_Cluster14718-002 | BES1 | brassinazole-resistant 1 [Z | 1493 | 0 | 0 | 0 |
| SALT00000058061_Cluster14718-003 | BES1 | brassinazole-resistant 1 [Z | 1639 | 0 | 0 | 0 |
| SALT00000073420_Cluster14718-004 | BES1 | brassinazole-resistant 1 [Z | 2595 | 0 | 0 | 0 |
| SALT00000074430_Cluster14718-005 | BES1 | brassinazole-resistant 1 [Z | 2597 | 0 | 0 | 0 |
| SALT00000083267_Cluster14718-006 | BES1 | brassinazole-resistant 1 [Z | 2645 | 0 | 0 | 0 |
| SALT00000009376_Cluster17889-001 | BES1 | beta-amylase 8 [Setaria ita | 2257 | 0 | 0 | 0 |
| SALT00000030506_Cluster17889-002 | BES1 | beta-amylase 8 [Setaria ita | 2325 | 0 | 0 | 0 |
| SALT00000015897_Cluster25098-001 | BES1 | Os02g0129600 [Oryza sati | 1757 | 0 | 0 | 0 |
| SALT00000039363_Cluster30829-001 | BES1 | [Hordeum vulgare vulgare | 1439 | 0 | 0 | 0 |
| SALT00000051749_Cluster30976-001 | BES1 | [Hordeum vulgare vulgare | 1423 | 0 | 0 | 0 |
| SALT00000057844_Cluster31672-001 | BES1 | BES1 BZR1 [Zea mays] | 1289 | 0 | 0 | 0 |
| SALT00000012511_Cluster31728-001 | BES1 | aquaporin NIP1-2-like | 1271 | 0 | 0 | 0 |
| SALT00000016210_Cluster976-001 | bHLH | transcription factor LHW-l | 1659 | 0 | 0 | 0 |
| SALT00000036314_Cluster976-004 | bHLH | transcription factor LHW [ | 2152 | 0 | 0 | 0 |
| SALT00000033798_Cluster1035-001 | bHLH | transcription factor LHW-l | 3459 | 0 | 0 | 0 |
| SALT00000073117_Cluster1035-002 | bHLH | transcription factor LHW-l | 2687 | 0 | 0 | 0 |
| SALT00000088750_Cluster1035-003 | bHLH | transcription factor LHW-l | 3340 | 0 | 0 | 0 |
| SALT00000089971_Cluster1035-004 | bHLH | Os12g0160400 [Oryza sati | 3545 | 0 | 0 | 0 |
| SALT00000092519_Cluster1035-006 | bHLH | Os12g0160400 [Oryza sati | 3424 | 0 | 0 | 0 |
| SALT00000005316_Cluster1065-001 | bHLH | [Hordeum vulgare vulgare | 1921 | 0 | 0 | 0 |
| SALT00000007528_Cluster1065-002 | bHLH | Os10g0575000 [Oryza sati | 2267 | 0 | 0 | 0 |
| SALT00000010165_Cluster1065-003 | bHLH | [Hordeum vulgare vulgare | 2463 | 0 | 0 | 0 |
| SALT00000015469_Cluster1065-004 | bHLH | [Hordeum vulgare vulgare | 1607 | 0 | 0 | 0 |
| SALT00000016024_Cluster1065-005 | bHLH | [Hordeum vulgare vulgare | 1908 | 0 | 0 | 0 |
| SALT00000016109_Cluster1065-006 | bHLH | [Hordeum vulgare vulgare | 1848 | 0 | 0 | 0 |
| SALT00000021002_Cluster1065-007 | bHLH | Os10g0575000 [Oryza sati | 2583 | 0 | 0 | 0 |
| SALT00000024005_Cluster1065-008 | bHLH | Os10g0575000 [Oryza sati | 2374 | 0 | 0 | 0 |
| SALT00000025022_Cluster1065-009 | bHLH | Os10g0575000 [Oryza sati | 2565 | 0 | 0 | 0 |
| SALT00000026339_Cluster1065-010 | bHLH | Os10g0575000 [Oryza sati | 2556 | 0 | 0 | 0 |
| SALT00000027590_Cluster1065-011 | bHLH | Os10g0575000 [Oryza sati | 2572 | 0 | 0 | 0 |
| SALT00000028525_Cluster1065-012 | bHLH | Os10g0575000 [Oryza sati | 2554 | 0 | 0 | 0 |
| SALT00000028571_Cluster1065-013 | bHLH | Os10g0575000 [Oryza sati | 2494 | 0 | 0 | 0 |
| SALT00000028984_Cluster1065-014 | bHLH | Os10g0575000 [Oryza sati | 2530 | 0 | 0 | 0 |
| SALT00000035942_Cluster1065-015 | bHLH | Os10g0575000 [Oryza sati | 2233 | 0 | 0 | 0 |
| SALT00000040438_Cluster1065-016 | bHLH | Os10g0575000 [Oryza sati | 2280 | 0 | 0 | 0 |
| SALT00000044614_Cluster1065-017 | bHLH | [Hordeum vulgare vulgare | 1938 | 0 | 0 | 0 |
| SALT00000054951_Cluster1065-018 | bHLH | Os10g0575000 [Oryza sati | 2032 | 0 | 0 | 0 |
| SALT00000056967_Cluster1065-019 | bHLH | [Hordeum vulgare vulgare | 1681 | 0 | 0 | 0 |
| SALT00000058557_Cluster1065-020 | bHLH | Os10g0575000 [Oryza sati | 2197 | 0 | 0 | 0 |
| SALT00000060214_Cluster1065-021 | bHLH | Os10g0575000 [Oryza sati | 2109 | 0 | 0 | 0 |
| SALT00000062295_Cluster1065-023 | bHLH | [Hordeum vulgare vulgare | 2470 | 0 | 0 | 0 |
| SALT00000063661_Cluster1065-024 | bHLH | [Hordeum vulgare vulgare | 1183 | 0 | 0 | 0 |
| SALT00000066553_Cluster1065-025 | bHLH | Os10g0575000 [Oryza sati | 2408 | 0 | 0 | 0 |
| SALT00000068746_Cluster1065-027 | bHLH | Os10g0575000 [Oryza sati | 2591 | 0 | 0 | 0 |
| SALT00000071576_Cluster1065-028 | bHLH | Os10g0575000 [Oryza sati | 2443 | 0 | 0 | 0 |
| SALT00000071797_Cluster1065-029 | bHLH | [Hordeum vulgare vulgare | 2528 | 0 | 0 | 0 |
| SALT00000078153_Cluster1065-030 | bHLH | Os10g0575000 [Oryza sati | 2849 | 0 | 0 | 0 |
| SALT00000081874_Cluster1065-031 | bHLH | Os10g0575000 [Oryza sati | 2463 | 0 | 0 | 0 |
| SALT00000084513_Cluster1065-032 | bHLH | Os10g0575000 [Oryza sati | 2541 | 0 | 0 | 0 |
| SALT00000072973_Cluster1065-034 | bHLH | Os10g0575000 [Oryza sati | 2540 | 0 | 0 | 0 |
| SALT00000088218_Cluster4226-001 | bHLH | transcription factor PIF1-li | 4158 | 0 | 0 | 0 |
| SALT00000001469_Cluster4394-001 | bHLH | transcription factor bHLH1 | 1539 | 0 | 0 | 0 |
| SALT00000042235_Cluster4394-002 | bHLH | transcription factor bHLH1 | 1611 | 0 | 0 | 0 |
| SALT00000066673_Cluster4394-003 | bHLH | transcription factor bHLH1 | 1538 | 0 | 0 | 0 |
| SALT00000091148_Cluster4394-004 | bHLH | transcription factor bHLH1 | 4119 | 0 | 0 | 0 |

| | | | | | | |
|---|---|---|---|---|---|---|
| SALT00000042792_Cluster4394-005 | bHLH | transcription factor bHLH1 | 1386 | 0 | 0 | 0 |
| SALT00000038508_Cluster6457-007 | bHLH | transcription factor UNE12 | 1415 | 0 | 0 | 0 |
| SALT00000030840_Cluster7602-001 | bHLH | transcription factor LHW-l | 3457 | 0 | 0 | 0 |
| SALT00000033921_Cluster7602-002 | bHLH | transcription factor LHW-l | 3323 | 0 | 0 | 0 |
| SALT00000088582_Cluster7602-003 | bHLH | transcription factor LHW-l | 3544 | 0 | 0 | 0 |
| SALT00000087312_Cluster7789-001 | bHLH | Os01g0915600 [Oryza sati | 3516 | 0 | 0 | 0 |
| SALT00000014857_Cluster10336-001 | bHLH | transcription factor bHLH8 | 1783 | 0 | 0 | 0 |
| SALT00000084067_Cluster10336-002 | bHLH | transcription factor bHLH8 | 3060 | 0 | 0 | 0 |
| SALT00000005899_Cluster10430-001 | bHLH | Os08g0490000 [Oryza sati | 1426 | 0 | 0 | 0 |
| SALT00000007315_Cluster10430-002 | bHLH | Os08g0490000 [Oryza sati | 1953 | 0 | 0 | 0 |
| SALT00000015765_Cluster10430-003 | bHLH | Os08g0490000 [Oryza sati | 1964 | 0 | 0 | 0 |
| SALT00000030531_Cluster10430-004 | bHLH | Os08g0490000 [Oryza sati | 3047 | 0 | 0 | 0 |
| SALT00000074322_Cluster10430-005 | bHLH | Os08g0490000 [Oryza sati | 2944 | 0 | 0 | 0 |
| SALT00000022520_Cluster10430-006 | bHLH | bHLH family-like [Oryza s | 2266 | 0 | 0 | 0 |
| SALT00000071952_Cluster10430-007 | bHLH | Os08g0490000 [Oryza sati | 2622 | 0 | 0 | 0 |
| SALT00000092342_Cluster10443-001 | bHLH | Transcription factor bHLH | 3047 | 0 | 0 | 0 |
| SALT00000019387_Cluster10800-001 | bHLH | transcription factor PIF1-li | 1743 | 0 | 0 | 0 |
| SALT00000076102_Cluster10800-002 | bHLH | transcription factor PIF1-li | 2998 | 0 | 0 | 0 |
| SALT00000008264_Cluster10800-003 | bHLH | transcription factor PIF1-li | 1806 | 0 | 0 | 0 |
| SALT00000059389_Cluster10800-004 | bHLH | transcription factor PIF1-li | 1976 | 0 | 0 | 0 |
| SALT00000053347_Cluster10800-005 | bHLH | transcription factor PIF1-li | 2024 | 0 | 0 | 0 |
| SALT00000036611_Cluster10800-006 | bHLH | transcription factor PIF1-li | 1717 | 0 | 0 | 0 |
| SALT00000068453_Cluster11473-001 | bHLH | transcription factor APG-li | 2917 | 0 | 0 | 0 |
| SALT00000070945_Cluster11473-002 | bHLH | transcription factor APG-li | 2518 | 0 | 0 | 0 |
| SALT00000036766_Cluster11473-003 | bHLH | transcription factor APG-li | 2555 | 0 | 0 | 0 |
| SALT00000018958_Cluster12421-001 | bHLH | transcription factor MYC3 | 1721 | 0 | 0 | 0 |
| SALT00000046816_Cluster12421-003 | bHLH | transcription factor bHLH1 | 1628 | 0 | 0 | 0 |
| SALT00000007514_Cluster12769-001 | bHLH | Os08g0490000 [Oryza sati | 1591 | 0 | 0 | 0 |
| SALT00000008026_Cluster12769-002 | bHLH | Os08g0490000 [Oryza sati | 1547 | 0 | 0 | 0 |
| SALT00000041642_Cluster12769-003 | bHLH | Os08g0490000 [Oryza sati | 2057 | 0 | 0 | 0 |
| SALT00000048255_Cluster12769-004 | bHLH | TPA: tubulin alpha-1 chain | 2794 | 0 | 0 | 0 |
| SALT00000002992_Cluster12805-001 | bHLH | TPA: HLH DNA-binding c | 1364 | 0 | 0 | 0 |
| SALT00000047483_Cluster12805-002 | bHLH | TPA: HLH DNA-binding c | 1474 | 0 | 0 | 0 |
| SALT00000079687_Cluster12805-003 | bHLH | TPA: HLH DNA-binding c | 2792 | 0 | 0 | 0 |
| SALT00000019214_Cluster12805-004 | bHLH | TPA: HLH DNA-binding c | 1404 | 0 | 0 | 0 |
| SALT00000039899_Cluster13252-004 | bHLH | TPA: HLH DNA-binding c | 1666 | 0 | 0 | 0 |
| SALT00000012819_Cluster13585-001 | bHLH | transcription factor UNE1( | 1762 | 0 | 0 | 0 |
| SALT00000019259_Cluster13585-002 | bHLH | transcription factor UNE1( | 1835 | 0 | 0 | down |
| SALT00000039197_Cluster13585-003 | bHLH | transcription factor UNE1( | 1761 | 0 | 0 | 0 |
| SALT00000050835_Cluster13585-004 | bHLH | transcription factor UNE1( | 1767 | 0 | 0 | 0 |
| SALT00000070841_Cluster14090-001 | bHLH | transcription factor bHLH7 | 2660 | 0 | 0 | 0 |
| SALT00000072083_Cluster14090-002 | bHLH | transcription factor bHLH7 | 2688 | 0 | 0 | 0 |
| SALT00000013711_Cluster14190-001 | bHLH | transcription factor bHLH8 | 1830 | 0 | 0 | 0 |
| SALT00000050763_Cluster14190-002 | bHLH | transcription factor bHLH8 | 1281 | 0 | 0 | 0 |
| SALT00000052587_Cluster14190-003 | bHLH | transcription factor bHLH8 | 1805 | 0 | 0 | 0 |
| SALT00000058301_Cluster14190-004 | bHLH | transcription factor bHLH8 | 1820 | 0 | 0 | 0 |
| SALT00000076381_Cluster14190-005 | bHLH | transcription factor bHLH8 | 2680 | 0 | 0 | 0 |
| SALT00000057810_Cluster14190-006 | bHLH | transcription factor bHLH8 | 1662 | 0 | 0 | 0 |
| SALT00000043529_Cluster14569-001 | bHLH | transcription factor bHLH( | 1555 | 0 | 0 | 0 |
| SALT00000074513_Cluster14569-002 | bHLH | transcription factor bHLH( | 2655 | down | 0 | down |
| SALT00000064381_Cluster14676-001 | bHLH | transcription factor UNE12 | 2647 | 0 | 0 | 0 |
| SALT00000077758_Cluster14676-002 | bHLH | transcription factor UNE12 | 2616 | 0 | 0 | 0 |
| SALT00000023234_Cluster14735-001 | bHLH | [Hordeum vulgare vulgare] | 2507 | 0 | 0 | 0 |
| SALT00000027078_Cluster14735-002 | bHLH | [Hordeum vulgare vulgare] | 2541 | 0 | 0 | 0 |
| SALT00000079361_Cluster14735-003 | bHLH | [Hordeum vulgare vulgare] | 2644 | 0 | 0 | 0 |
| SALT00000027557_Cluster15505-001 | bHLH | TPA: HLH DNA-binding c | 2589 | 0 | 0 | 0 |
| SALT00000065404_Cluster15976-001 | bHLH | transcription factor PIF4 is | 1941 | 0 | 0 | 0 |
| SALT00000067011_Cluster15976-002 | bHLH | transcription factor PIF4 is | 2554 | 0 | 0 | 0 |

| | | | | | | |
|---|---|---|---|---|---|---|
| SALT00000043238_Cluster15976-003 | bHLH | transcription factor PIF4 is | 1733 | 0 | 0 | 0 |
| SALT00000016984_Cluster15987-001 | bHLH | transcription factor APG-li | 1753 | 0 | 0 | 0 |
| SALT00000066550_Cluster15987-002 | bHLH | transcription factor APG-li | 2321 | 0 | 0 | 0 |
| SALT00000069739_Cluster15987-003 | bHLH | transcription factor APG-li | 2553 | 0 | 0 | 0 |
| SALT00000003386_Cluster16315-001 | bHLH | transcription factor APG-li | 2294 | 0 | 0 | 0 |
| SALT00000014418_Cluster16315-002 | bHLH | transcription factor APG-li | 1678 | 0 | 0 | 0 |
| SALT00000022042_Cluster16315-003 | bHLH | transcription factor APG-li | 2529 | 0 | 0 | 0 |
| SALT00000025328_Cluster16315-004 | bHLH | transcription factor APG-li | 2481 | 0 | 0 | 0 |
| SALT00000027940_Cluster16315-005 | bHLH | transcription factor APG-li | 2431 | 0 | 0 | 0 |
| SALT00000076502_Cluster16630-001 | bHLH | transcription factor APG-li | 2503 | 0 | 0 | 0 |
| SALT00000082970_Cluster16630-002 | bHLH | transcription factor APG-li | 2436 | 0 | 0 | 0 |
| SALT00000003734_Cluster16868-001 | bHLH | Os11g0523700 [Oryza sati | 1946 | 0 | 0 | 0 |
| SALT00000041665_Cluster16868-002 | bHLH | Os11g0523700 [Oryza sati | 1938 | 0 | 0 | 0 |
| SALT00000069898_Cluster16868-003 | bHLH | Os11g0523700 [Oryza sati | 2483 | 0 | 0 | 0 |
| SALT00000006425_Cluster17127-001 | bHLH | transcription factor bHLH1 | 2421 | 0 | 0 | 0 |
| SALT00000021228_Cluster17127-002 | bHLH | transcription factor bHLH1 | 2426 | 0 | 0 | 0 |
| SALT00000026310_Cluster17127-003 | bHLH | transcription factor bHLH1 | 2327 | 0 | 0 | 0 |
| SALT00000079501_Cluster17127-004 | bHLH | transcription factor bHLH1 | 2458 | 0 | 0 | 0 |
| SALT00000044106_Cluster17566-001 | bHLH | transcription factor APG-li | 1939 | 0 | 0 | 0 |
| SALT00000076371_Cluster17566-002 | bHLH | transcription factor APG-li | 2392 | 0 | 0 | 0 |
| SALT00000009164_Cluster17566-003 | bHLH | transcription factor APG-li | 1929 | 0 | 0 | 0 |
| SALT00000008620_Cluster17833-001 | bHLH | Os01g0915600 [Oryza sati | 2028 | 0 | 0 | 0 |
| SALT00000045287_Cluster17833-002 | bHLH | HLH DNA-binding domai | 2055 | 0 | 0 | 0 |
| SALT00000045752_Cluster17833-003 | bHLH | transcription factor bHLH7 | 1903 | 0 | 0 | 0 |
| SALT00000050247_Cluster17833-004 | bHLH | Os01g0915600 [Oryza sati | 2151 | 0 | 0 | 0 |
| SALT00000054051_Cluster17833-005 | bHLH | Os01g0915600 [Oryza sati | 2195 | 0 | 0 | 0 |
| SALT00000067446_Cluster17833-006 | bHLH | Os01g0915600 [Oryza sati | 2204 | 0 | 0 | 0 |
| SALT00000074978_Cluster17833-007 | bHLH | Os01g0915600 [Oryza sati | 2337 | down | down | down |
| SALT00000036878_Cluster17833-008 | bHLH | Os01g0915600 [Oryza sati | 2258 | 0 | 0 | 0 |
| SALT00000010888_Cluster18596-001 | bHLH | transcription factor bHLH1 | 2207 | 0 | 0 | 0 |
| SALT00000047521_Cluster18596-002 | bHLH | transcription factor bHLH1 | 2131 | 0 | 0 | 0 |
| SALT00000010067_Cluster19058-001 | bHLH | HLH DNA-binding domai | 2161 | 0 | 0 | 0 |
| SALT00000015737_Cluster19058-002 | bHLH | HLH DNA-binding domai | 1970 | 0 | 0 | 0 |
| SALT00000037749_Cluster19507-001 | bHLH | helix-loop-helix DNA-bin | 2119 | 0 | 0 | 0 |
| SALT00000055792_Cluster19942-004 | bHLH | HLH DNA-binding domai | 1126 | 0 | 0 | 0 |
| SALT00000010746_Cluster20033-001 | bHLH | transcription factor bHLH1 | 2051 | 0 | 0 | up |
| SALT00000011286_Cluster20033-002 | bHLH | transcription factor bHLH1 | 1916 | 0 | 0 | up |
| SALT00000015086_Cluster20033-003 | bHLH | transcription factor bHLH1 | 1888 | 0 | 0 | 0 |
| SALT00000017668_Cluster20033-004 | bHLH | transcription factor bHLH1 | 2018 | 0 | 0 | 0 |
| SALT00000042888_Cluster20033-005 | bHLH | transcription factor bHLH1 | 2077 | 0 | 0 | 0 |
| SALT00000004741_Cluster20047-001 | bHLH | transcription factor PIF5 is | 1741 | 0 | 0 | 0 |
| SALT00000045499_Cluster20047-002 | bHLH | transcription factor PIF5 is | 2076 | 0 | down | down |
| SALT00000062922_Cluster20047-003 | bHLH | transcription factor PIF5 is | 1747 | 0 | 0 | 0 |
| SALT00000051699_Cluster20177-001 | bHLH | helix-loop-helix DNA-bin | 2065 | 0 | 0 | 0 |
| SALT00000009065_Cluster20402-001 | bHLH | bHLH transcription partial | 1875 | 0 | 0 | 0 |
| SALT00000014870_Cluster20402-002 | bHLH | bHLH transcription partial | 1926 | 0 | 0 | 0 |
| SALT00000039300_Cluster20402-003 | bHLH | bHLH transcription factor | 1895 | 0 | 0 | 0 |
| SALT00000043463_Cluster20402-004 | bHLH | bHLH transcription partial | 1936 | 0 | 0 | 0 |
| SALT00000050574_Cluster20402-005 | bHLH | bHLH transcription partial | 1977 | 0 | 0 | 0 |
| SALT00000056489_Cluster20402-006 | bHLH | bHLH transcription partial | 2047 | 0 | 0 | 0 |
| SALT00000002366_Cluster20968-001 | bHLH | transcription factor bHLH1 | 1732 | 0 | 0 | 0 |
| SALT00000003886_Cluster20968-002 | bHLH | transcription factor bHLH1 | 1996 | 0 | 0 | 0 |
| SALT00000004005_Cluster20968-003 | bHLH | transcription factor bHLH1 | 1550 | 0 | 0 | 0 |
| SALT00000037536_Cluster20968-004 | bHLH | transcription factor bHLH1 | 2007 | 0 | 0 | 0 |
| SALT00000040659_Cluster20968-005 | bHLH | transcription factor bHLH1 | 1990 | 0 | 0 | 0 |
| SALT00000012122_Cluster21005-001 | bHLH | transcription factor bHLH1 | 2004 | down | 0 | down |
| SALT00000014367_Cluster21005-002 | bHLH | transcription factor bHLH1 | 1925 | 0 | 0 | 0 |
| SALT00000020291_Cluster21005-003 | bHLH | transcription factor bHLH1 | 1897 | 0 | 0 | 0 |

| | | | | | | |
|---|---|---|---|---|---|---|
| SALT00000044023_Cluster21005-004 | bHLH | transcription factor bHLH1 | 1950 | 0 | 0 | 0 |
| SALT00000046348_Cluster21005-005 | bHLH | transcription factor bHLH1 | 1700 | 0 | 0 | 0 |
| SALT00000048390_Cluster21005-006 | bHLH | transcription factor bHLH1 | 1930 | 0 | 0 | 0 |
| SALT00000015840_Cluster21305-001 | bHLH | Os01g0196300 [Oryza sati | 1786 | 0 | 0 | 0 |
| SALT00000016292_Cluster21305-002 | bHLH | Os01g0196300 [Oryza sati | 1984 | 0 | 0 | 0 |
| SALT00000053923_Cluster21356-001 | bHLH | transcription factor UNE1( | 1981 | 0 | 0 | 0 |
| SALT00000059283_Cluster21375-001 | bHLH | Os03g0188400 [Oryza sati | 1980 | 0 | 0 | 0 |
| SALT00000036616_Cluster21522-001 | bHLH | transcription factor UNE12 | 1969 | 0 | 0 | 0 |
| SALT00000060235_Cluster21522-002 | bHLH | transcription factor UNE12 | 1645 | 0 | 0 | down |
| SALT00000039216_Cluster21725-001 | bHLH | transcription factor bHLH4 | 1873 | 0 | 0 | 0 |
| SALT00000049048_Cluster21725-002 | bHLH | transcription factor bHLH4 | 1957 | 0 | 0 | 0 |
| SALT00000007397_Cluster21807-001 | bHLH | [Hordeum vulgare vulgare] | 1749 | 0 | 0 | 0 |
| SALT00000014023_Cluster21807-002 | bHLH | [Hordeum vulgare vulgare] | 1609 | 0 | 0 | 0 |
| SALT00000016008_Cluster21807-003 | bHLH | [Hordeum vulgare vulgare] | 1697 | 0 | 0 | 0 |
| SALT00000036246_Cluster21946-001 | bHLH | transcription factor bHLH9 | 1943 | 0 | 0 | 0 |
| SALT00000060269_Cluster21946-002 | bHLH | Os03g0188400 [Oryza sati | 1903 | 0 | 0 | 0 |
| SALT00000037577_Cluster22234-001 | bHLH | symbiotic ammonium trans | 1924 | 0 | 0 | 0 |
| SALT00000061068_Cluster22234-002 | bHLH | symbiotic ammonium trans | 1909 | 0 | 0 | 0 |
| SALT00000040982_Cluster22245-001 | bHLH | transcription factor bHLH( | 1923 | 0 | 0 | 0 |
| SALT00000003523_Cluster22323-001 | bHLH | transcription factor MYC3 | 1918 | 0 | 0 | 0 |
| SALT00000016409_Cluster22323-002 | bHLH | transcription factor MYC3 | 1880 | 0 | 0 | 0 |
| SALT00000062048_Cluster22323-003 | bHLH | transcription factor MYC3 | 1737 | 0 | 0 | 0 |
| SALT00000039012_Cluster23094-001 | bHLH | transcription factor bHLH3 | 1871 | 0 | 0 | 0 |
| SALT00000012709_Cluster23356-001 | bHLH | HLH DNA-binding domai | 1855 | 0 | 0 | 0 |
| SALT00000041682_Cluster23356-002 | bHLH | HLH DNA-binding domai | 1693 | 0 | 0 | 0 |
| SALT00000038050_Cluster23508-001 | bHLH | HLH DNA-binding domai | 1846 | 0 | 0 | 0 |
| SALT00000001241_Cluster23624-001 | bHLH | BHLH transcription factor | 1623 | 0 | 0 | 0 |
| SALT00000007107_Cluster23624-002 | bHLH | BHLH transcription factor | 1784 | 0 | 0 | 0 |
| SALT00000011697_Cluster23624-003 | bHLH | BHLH transcription factor | 1734 | 0 | 0 | 0 |
| SALT00000015610_Cluster23624-004 | bHLH | BHLH transcription factor | 1839 | 0 | 0 | 0 |
| SALT00000017298_Cluster23624-005 | bHLH | BHLH transcription factor | 1509 | 0 | 0 | 0 |
| SALT00000017879_Cluster23624-006 | bHLH | BHLH transcription factor | 1504 | 0 | 0 | 0 |
| SALT00000035183_Cluster23624-007 | bHLH | BHLH transcription factor | 1583 | 0 | 0 | 0 |
| SALT00000035371_Cluster23624-008 | bHLH | BHLH transcription factor | 1614 | 0 | 0 | 0 |
| SALT00000058817_Cluster23695-001 | bHLH | Os03g0188400 [Oryza sati | 1835 | 0 | 0 | 0 |
| SALT00000062088_Cluster23695-002 | bHLH | Os03g0188400 [Oryza sati | 1651 | 0 | 0 | 0 |
| SALT00000018949_Cluster23766-001 | bHLH | transcription factor bHLH1 | 1567 | 0 | 0 | 0 |
| SALT00000003920_Cluster24231-001 | bHLH | helix-loop-helix DNA-binc | 1603 | 0 | 0 | up |
| SALT00000011416_Cluster24231-002 | bHLH | helix-loop-helix DNA-binc | 1572 | 0 | 0 | 0 |
| SALT00000012785_Cluster24231-003 | bHLH | helix-loop-helix DNA-binc | 1806 | 0 | 0 | 0 |
| SALT00000022293_Cluster24231-004 | bHLH | helix-loop-helix DNA-binc | 1611 | 0 | 0 | 0 |
| SALT00000060209_Cluster24231-005 | bHLH | helix-loop-helix DNA-binc | 1950 | 0 | 0 | 0 |
| SALT00000043862_Cluster24231-006 | bHLH | helix-loop-helix DNA-binc | 1664 | 0 | 0 | 0 |
| SALT00000006392_Cluster24462-001 | bHLH | DNA binding [Zea mays] | 1792 | 0 | 0 | 0 |
| SALT00000016846_Cluster24478-001 | bHLH | HLH DNA-binding domai | 1791 | 0 | 0 | 0 |
| SALT00000038126_Cluster24478-002 | bHLH | HLH DNA-binding domai | 1662 | 0 | 0 | 0 |
| SALT00000039715_Cluster24478-003 | bHLH | HLH DNA-binding domai | 1783 | 0 | 0 | 0 |
| SALT00000053132_Cluster24478-004 | bHLH | HLH DNA-binding domai | 1612 | 0 | 0 | 0 |
| SALT00000054097_Cluster24478-005 | bHLH | transcription factor bHLH8 | 1697 | 0 | 0 | 0 |
| SALT00000010074_Cluster24503-001 | bHLH | HLH DNA-binding domai | 1705 | 0 | 0 | 0 |
| SALT00000052161_Cluster24503-002 | bHLH | HLH DNA-binding domai | 1790 | 0 | 0 | 0 |
| SALT00000051365_Cluster25054-001 | bHLH | transcription factor ABOR | 1760 | 0 | 0 | 0 |
| SALT00000004618_Cluster25370-001 | bHLH | DNA binding [Zea mays] | 1743 | 0 | 0 | 0 |
| SALT00000001287_Cluster25380-001 | bHLH | DNA binding [Zea mays] | 1665 | 0 | 0 | 0 |
| SALT00000019650_Cluster25380-002 | bHLH | DNA binding [Zea mays] | 1593 | 0 | 0 | 0 |
| SALT00000039546_Cluster25380-003 | bHLH | DNA binding [Zea mays] | 1743 | 0 | 0 | up |
| SALT00000056811_Cluster25380-004 | bHLH | DNA binding [Zea mays] | 1349 | 0 | 0 | 0 |
| SALT00000001833_Cluster25605-001 | bHLH | transcription factor SPATU | 1673 | 0 | 0 | 0 |

| | | | | | | |
|---|---|---|---|---|---|---|
| SALT00000036162_Cluster25605-002 | bHLH | transcription factor SPATU | 1732 | 0 | 0 | 0 |
| SALT00000006527_Cluster26008-001 | bHLH | BHLH transcription factor | 1661 | 0 | 0 | 0 |
| SALT00000056557_Cluster26008-002 | bHLH | BHLH transcription factor | 1712 | 0 | 0 | 0 |
| SALT00000014066_Cluster26038-001 | bHLH | TPA: HLH DNA-binding c | 1709 | 0 | 0 | 0 |
| SALT00000042148_Cluster26260-001 | bHLH | transcription factor bHLH9 | 1696 | 0 | 0 | 0 |
| SALT00000006960_Cluster26327-001 | bHLH | transcription factor bHLH3 | 1692 | 0 | 0 | 0 |
| SALT00000050399_Cluster26384-001 | bHLH | HLH DNA-binding domai | 1690 | 0 | 0 | 0 |
| SALT00000057507_Cluster26384-002 | bHLH | HLH DNA-binding domai | 1357 | 0 | 0 | 0 |
| SALT00000040689_Cluster26384-003 | bHLH | transcription factor bHLH9 | 1523 | 0 | 0 | 0 |
| SALT00000004428_Cluster26455-001 | bHLH | Os03g0188400 [Oryza sati | 1366 | 0 | 0 | 0 |
| SALT00000010915_Cluster26455-002 | bHLH | Os03g0188400 [Oryza sati | 1686 | 0 | 0 | 0 |
| SALT00000045411_Cluster26455-004 | bHLH | Os03g0188400 [Oryza sati | 1642 | 0 | 0 | 0 |
| SALT00000053516_Cluster26455-005 | bHLH | Os03g0188400 [Oryza sati | 1218 | 0 | 0 | 0 |
| SALT00000036233_Cluster26476-001 | bHLH | helix-loop-helix DNA-binc | 1685 | 0 | 0 | 0 |
| SALT00000001837_Cluster26706-001 | bHLH | BHLH transcription factor | 1506 | down | down | 0 |
| SALT00000009888_Cluster26706-002 | bHLH | BHLH transcription factor | 1576 | 0 | 0 | 0 |
| SALT00000017723_Cluster26706-003 | bHLH | BHLH transcription factor | 1466 | 0 | 0 | 0 |
| SALT00000042354_Cluster26706-004 | bHLH | BHLH transcription factor | 1551 | 0 | 0 | 0 |
| SALT00000054371_Cluster26706-005 | bHLH | BHLH transcription factor | 1675 | 0 | 0 | 0 |
| SALT00000036808_Cluster27262-001 | bHLH | transcription factor bHLH1 | 1260 | 0 | 0 | 0 |
| SALT00000006390_Cluster27645-001 | bHLH | bHLH transcription partial | 1628 | 0 | 0 | 0 |
| SALT00000043647_Cluster27645-002 | bHLH | BHLH transcription factor | 1529 | 0 | 0 | down |
| SALT00000004990_Cluster27645-003 | bHLH | bHLH transcription partial | 1597 | 0 | 0 | 0 |
| SALT00000007031_Cluster27645-004 | bHLH | bHLH transcription partial | 1551 | 0 | 0 | 0 |
| SALT00000045956_Cluster27657-001 | bHLH | TPA: HLH DNA-binding c | 1628 | 0 | 0 | 0 |
| SALT00000044620_Cluster27780-001 | bHLH | transcription factor bHLH9 | 1622 | 0 | 0 | 0 |
| SALT00000056014_Cluster27780-002 | bHLH | transcription factor bHLH9 | 1371 | 0 | 0 | 0 |
| SALT00000053668_Cluster27915-001 | bHLH | transcription factor BHLH' | 1615 | 0 | 0 | 0 |
| SALT00000013423_Cluster27924-001 | bHLH | [Hordeum vulgare vulgare] | 1614 | 0 | 0 | 0 |
| SALT00000009025_Cluster28319-001 | bHLH | HLH DNA-binding domai | 1593 | 0 | 0 | 0 |
| SALT00000054539_Cluster28470-001 | bHLH | transcription factor bHLH6 | 1586 | 0 | 0 | 0 |
| SALT00000057291_Cluster28470-002 | bHLH | transcription factor bHLH6 | 1260 | 0 | 0 | 0 |
| SALT00000059217_Cluster28470-003 | bHLH | transcription factor bHLH6 | 1435 | 0 | 0 | 0 |
| SALT00000055239_Cluster28470-004 | bHLH | transcription factor bHLH6 | 1516 | 0 | 0 | 0 |
| SALT00000010484_Cluster28470-005 | bHLH | transcription factor bHLH6 | 1564 | 0 | 0 | 0 |
| SALT00000015483_Cluster28668-001 | bHLH | transcription factor bHLH3 | 1575 | 0 | 0 | 0 |
| SALT00000048638_Cluster29115-001 | bHLH | TPA: HLH DNA-binding c | 1553 | 0 | 0 | 0 |
| SALT00000002962_Cluster29130-001 | bHLH | transcription factor bHLH6 | 1443 | 0 | 0 | 0 |
| SALT00000007624_Cluster29130-002 | bHLH | transcription factor bHLH6 | 1459 | 0 | 0 | 0 |
| SALT00000015721_Cluster29130-003 | bHLH | transcription factor bHLH6 | 1527 | 0 | 0 | 0 |
| SALT00000019909_Cluster29130-004 | bHLH | transcription factor bHLH6 | 1552 | 0 | 0 | 0 |
| SALT00000059414_Cluster29193-001 | bHLH | TPA: HLH DNA-binding c | 1518 | 0 | 0 | 0 |
| SALT00000060171_Cluster29193-002 | bHLH | TPA: HLH DNA-binding c | 1550 | 0 | 0 | 0 |
| SALT00000039885_Cluster29193-003 | bHLH | TPA: HLH DNA-binding c | 1760 | 0 | 0 | 0 |
| SALT00000045185_Cluster29315-001 | bHLH | HLH DNA-binding domai | 1543 | 0 | 0 | 0 |
| SALT00000045321_Cluster29725-001 | bHLH | BHLH transcription factor | 1519 | 0 | 0 | 0 |
| SALT00000017932_Cluster29860-001 | bHLH | Os01g0900800 [Oryza sati | 1511 | 0 | 0 | 0 |
| SALT00000039052_Cluster30130-001 | bHLH | HLH DNA-binding domai | 1493 | 0 | 0 | 0 |
| SALT00000048006_Cluster30255-001 | bHLH | Os04g0350700 [Oryza sati | 1485 | 0 | 0 | 0 |
| SALT00000064240_Cluster30272-001 | bHLH | transcription factor bHLH7 | 1484 | 0 | 0 | 0 |
| SALT00000018788_Cluster30597-001 | bHLH | transcription factor UNE12 | 1460 | 0 | 0 | 0 |
| SALT00000061167_Cluster30597-002 | bHLH | transcription factor UNE12 | 1457 | 0 | 0 | 0 |
| SALT00000003506_Cluster31337-001 | bHLH | transcription factor bHLH4 | 1369 | 0 | 0 | 0 |
| SALT00000037556_Cluster31568-001 | bHLH | transcription factor bHLH3 | 1315 | 0 | 0 | up |
| SALT00000068478_Cluster32023-001 | bHLH | transcription factor bHLH3 | 1182 | 0 | 0 | 0 |
| SALT00000046295_Cluster32269-001 | bHLH | transcription factor bHLH9 | 1084 | 0 | 0 | 0 |
| SALT00000051482_Cluster21300-001 | bHLH | ---NA--- | 1985 | 0 | 0 | 0 |
| SALT00000002003_Cluster1113-001 | bZIP | transcription factor HBP-1 | 1625 | 0 | 0 | 0 |

| | | | | | | |
|---|---|---|---|---|---|---|
| SALT00000002770_Cluster1113-002 | bZIP | transcription factor HBP-1 | 1711 | 0 | 0 | 0 |
| SALT00000003751_Cluster1113-003 | bZIP | transcription factor HBP-1 | 1570 | 0 | 0 | 0 |
| SALT00000005300_Cluster1113-004 | bZIP | transcription factor HBP-1 | 1787 | 0 | 0 | 0 |
| SALT00000007354_Cluster1113-005 | bZIP | transcription factor HBP-1 | 1654 | 0 | 0 | 0 |
| SALT00000009465_Cluster1113-006 | bZIP | transcription factor HBP-1 | 1523 | 0 | 0 | 0 |
| SALT00000014148_Cluster1113-007 | bZIP | transcription factor HBP-1 | 1641 | 0 | 0 | 0 |
| SALT00000017902_Cluster1113-008 | bZIP | transcription factor HBP-1 | 1695 | 0 | 0 | 0 |
| SALT00000024934_Cluster1113-009 | bZIP | transcription factor HBP-1 | 1582 | 0 | 0 | 0 |
| SALT00000036966_Cluster1113-010 | bZIP | transcription factor HBP-1 | 1694 | 0 | 0 | 0 |
| SALT00000044288_Cluster1113-011 | bZIP | transcription factor HBP-1 | 1702 | 0 | 0 | 0 |
| SALT00000049792_Cluster1113-012 | bZIP | transcription factor HBP-1 | 6465 | 0 | 0 | 0 |
| SALT00000051466_Cluster1113-013 | bZIP | transcription factor HBP-1 | 1646 | 0 | 0 | 0 |
| SALT00000051881_Cluster1113-014 | bZIP | transcription factor HBP-1 | 1799 | 0 | 0 | 0 |
| SALT00000052750_Cluster1113-015 | bZIP | transcription factor HBP-1 | 1862 | 0 | 0 | 0 |
| SALT00000053689_Cluster1113-017 | bZIP | transcription factor TGA2 | 1834 | 0 | 0 | 0 |
| SALT00000016837_Cluster2949-001 | bZIP | transcription factor HBP-1 | 1885 | 0 | 0 | 0 |
| SALT00000002053_Cluster3551-001 | bZIP | bZIP family transcription f | 2096 | 0 | 0 | up |
| SALT00000005204_Cluster3551-002 | bZIP | bZIP family transcription f | 1926 | 0 | 0 | 0 |
| SALT00000051251_Cluster3551-003 | bZIP | bZIP family transcription f | 1894 | 0 | 0 | 0 |
| SALT00000063675_Cluster3551-004 | bZIP | bZIP family transcription f | 3205 | 0 | 0 | 0 |
| SALT00000085178_Cluster3551-005 | bZIP | bZIP family transcription f | 4375 | 0 | 0 | 0 |
| SALT00000090140_Cluster3551-006 | bZIP | bZIP family transcription f | 4367 | 0 | 0 | 0 |
| SALT00000080129_Cluster3572-001 | bZIP | Os01g0542700 [Oryza sati | 4368 | 0 | 0 | 0 |
| SALT00000071760_Cluster4255-001 | bZIP | Light-inducible CPRF2 | 2651 | 0 | 0 | down |
| SALT00000002013_Cluster4496-001 | bZIP | transcription factor HBP-1 | 1659 | 0 | 0 | 0 |
| SALT00000002922_Cluster4496-002 | bZIP | transcription factor HBP-1 | 1645 | 0 | 0 | 0 |
| SALT00000010691_Cluster4496-003 | bZIP | transcription factor HBP-1 | 1650 | 0 | 0 | 0 |
| SALT00000018376_Cluster4496-004 | bZIP | transcription factor HBP-1 | 1902 | 0 | 0 | 0 |
| SALT00000021823_Cluster4496-005 | bZIP | transcription factor HBP-1 | 2297 | 0 | 0 | 0 |
| SALT00000024252_Cluster4496-006 | bZIP | transcription factor HBP-1 | 1688 | 0 | 0 | 0 |
| SALT00000033345_Cluster4496-007 | bZIP | transcription factor HBP-1 | 3015 | 0 | 0 | 0 |
| SALT00000041598_Cluster4496-008 | bZIP | transcription factor HBP-1 | 1580 | 0 | 0 | 0 |
| SALT00000056041_Cluster4496-009 | bZIP | transcription factor HBP-1 | 1589 | 0 | 0 | 0 |
| SALT00000067391_Cluster4496-010 | bZIP | transcription factor HBP-1 | 1560 | 0 | 0 | 0 |
| SALT00000092865_Cluster4496-011 | bZIP | transcription factor HBP-1 | 4091 | 0 | 0 | 0 |
| SALT00000060828_Cluster4496-012 | bZIP | transcription factor HBP-1 | 1741 | 0 | 0 | 0 |
| SALT00000031130_Cluster5227-001 | bZIP | TPA: bZIP transcription fa | 3699 | 0 | 0 | 0 |
| SALT00000033425_Cluster5227-002 | bZIP | TPA: bZIP transcription fa | 3931 | 0 | 0 | 0 |
| SALT00000063527_Cluster5227-003 | bZIP | Light-inducible CPRF2 | 2575 | 0 | 0 | 0 |
| SALT00000050415_Cluster5247-001 | bZIP | LOC100282457 isoform X | 1703 | 0 | 0 | 0 |
| SALT00000089436_Cluster5247-002 | bZIP | LOC100282457 isoform X | 3927 | 0 | 0 | 0 |
| SALT00000009666_Cluster5247-003 | bZIP | LOC100282457 isoform X | 1499 | 0 | 0 | 0 |
| SALT00000086863_Cluster5360-001 | bZIP | transcription factor VIP1-l | 3904 | 0 | 0 | 0 |
| SALT00000088440_Cluster5805-001 | bZIP | transcription factor 21 | 3823 | 0 | 0 | 0 |
| SALT00000093036_Cluster5805-002 | bZIP | transcription factor 21 | 3735 | 0 | 0 | 0 |
| SALT00000093197_Cluster5805-003 | bZIP | transcription factor 21 | 3827 | 0 | 0 | 0 |
| SALT00000061242_Cluster5805-005 | bZIP | transcription factor 21 [Zea | 1407 | 0 | 0 | 0 |
| SALT00000082886_Cluster5810-001 | bZIP | light-inducible CPRF2-like | 3826 | 0 | 0 | 0 |
| SALT00000038352_Cluster6286-001 | bZIP | transcription factor TGA4 | 1704 | 0 | 0 | 0 |
| SALT00000058265_Cluster6286-002 | bZIP | transcription factor TGA4 | 1682 | 0 | 0 | 0 |
| SALT00000050507_Cluster6286-004 | bZIP | transcription factor TGA4 | 1574 | 0 | 0 | 0 |
| SALT00000002838_Cluster6738-001 | bZIP | bZIP transcription factor su | 1697 | 0 | 0 | 0 |
| SALT00000046488_Cluster6738-003 | bZIP | bZIP transcription factor su | 1749 | 0 | 0 | 0 |
| SALT00000055772_Cluster6738-004 | bZIP | bZIP transcription factor su | 1825 | 0 | 0 | 0 |
| SALT00000091446_Cluster6785-001 | bZIP | transcription factor TGA2- | 3668 | 0 | 0 | 0 |
| SALT00000005469_Cluster6863-001 | bZIP | transcription factor TGA4 | 1705 | 0 | 0 | 0 |
| SALT00000005553_Cluster6863-002 | bZIP | TPA: bZIP transcription fa | 1888 | 0 | 0 | 0 |
| SALT00000013905_Cluster6863-003 | bZIP | TPA: bZIP transcription fa | 1672 | 0 | 0 | 0 |

| | | | | | | |
|---|---|---|---|---|---|---|
| SALT00000016081_Cluster6863-004 | bZIP | transcription factor TGA4 | 1838 | 0 | 0 | 0 |
| SALT00000040173_Cluster6863-005 | bZIP | TPA: bZIP transcription fa | 1684 | 0 | 0 | 0 |
| SALT00000040504_Cluster6863-006 | bZIP | transcription factor TGA4 | 1772 | 0 | 0 | 0 |
| SALT00000045001_Cluster6863-007 | bZIP | transcription factor TGA4 | 1605 | 0 | 0 | 0 |
| SALT00000048126_Cluster6863-008 | bZIP | TPA: bZIP transcription fa | 1744 | 0 | 0 | 0 |
| SALT00000061805_Cluster6863-009 | bZIP | transcription factor TGA4 | 1561 | 0 | 0 | 0 |
| SALT00000087308_Cluster6863-010 | bZIP | TPA: bZIP transcription fa | 3656 | 0 | 0 | 0 |
| SALT00000005777_Cluster6863-011 | bZIP | transcription factor TGA4 | 1669 | 0 | 0 | 0 |
| SALT00000005217_Cluster9910-002 | bZIP | bZIP transcription factor A | 1564 | 0 | 0 | 0 |
| SALT00000027285_Cluster9910-003 | bZIP | bZIP transcription factor A | 2975 | 0 | 0 | 0 |
| SALT00000074691_Cluster9910-004 | bZIP | bZIP transcription factor A | 3124 | 0 | 0 | 0 |
| SALT00000081007_Cluster9914-001 | bZIP | Os11g0218000 [Oryza sati | 3124 | 0 | 0 | 0 |
| SALT00000071556_Cluster10340-001 | bZIP | bZIP transcription factor su | 3059 | 0 | 0 | 0 |
| SALT00000005573_Cluster10351-001 | bZIP | opaque2 heterodimerizing | 1554 | 0 | 0 | 0 |
| SALT00000045405_Cluster10351-002 | bZIP | fructose-bisphosphate aldo | 3057 | 0 | 0 | 0 |
| SALT00000050833_Cluster10351-003 | bZIP | opaque2 heterodimerizing | 1444 | 0 | 0 | 0 |
| SALT00000061570_Cluster10351-004 | bZIP | opaque2 heterodimerizing | 1531 | 0 | 0 | 0 |
| SALT00000046442_Cluster10367-001 | bZIP | TPA: bZIP transcription fa | 3054 | 0 | 0 | down |
| SALT00000031407_Cluster11744-001 | bZIP | TPA: bZIP transcription fa | 2890 | 0 | 0 | 0 |
| SALT00000045534_Cluster11744-002 | bZIP | TPA: bZIP transcription fa | 1541 | 0 | 0 | 0 |
| SALT00000067771_Cluster11744-003 | bZIP | TPA: bZIP transcription fa | 2871 | 0 | 0 | 0 |
| SALT00000001631_Cluster12336-001 | bZIP | Transcription factor HBP- | 1708 | 0 | 0 | 0 |
| SALT00000083834_Cluster12336-002 | bZIP | Transcription factor HBP- | 2836 | 0 | 0 | 0 |
| SALT00000079851_Cluster12707-001 | bZIP | transcription factor HBP-1 | 2801 | 0 | 0 | 0 |
| SALT00000014300_Cluster12757-001 | bZIP | basic leucine zipper 9-like | 1261 | 0 | 0 | 0 |
| SALT00000023979_Cluster12757-002 | bZIP | poly [Oryza sativa Japonic | 2795 | 0 | 0 | 0 |
| SALT00000024603_Cluster12757-003 | bZIP | basic leucine zipper 9-like | 1381 | 0 | 0 | 0 |
| SALT00000049103_Cluster12777-001 | bZIP | LOC100282457 isoform X | 1345 | 0 | 0 | 0 |
| SALT00000076372_Cluster12777-002 | bZIP | LOC100282457 isoform X | 2794 | 0 | 0 | 0 |
| SALT00000024502_Cluster13424-004 | bZIP | senescence-associated 12 [ | 2738 | 0 | 0 | 0 |
| SALT00000072463_Cluster15619-001 | bZIP | bZIP transcription factor su | 2580 | 0 | 0 | 0 |
| SALT00000023385_Cluster15693-001 | bZIP | DNA binding [Zea mays] | 2475 | 0 | 0 | 0 |
| SALT00000025308_Cluster15693-002 | bZIP | DNA binding [Zea mays] | 2507 | 0 | 0 | 0 |
| SALT00000027559_Cluster15693-003 | bZIP | DNA binding [Zea mays] | 2219 | 0 | 0 | 0 |
| SALT00000028430_Cluster15693-004 | bZIP | DNA binding [Zea mays] | 2413 | 0 | 0 | 0 |
| SALT00000031704_Cluster15693-005 | bZIP | DNA binding [Zea mays] | 2560 | 0 | 0 | 0 |
| SALT00000061955_Cluster15693-007 | bZIP | DNA binding [Zea mays] | 2537 | 0 | 0 | 0 |
| SALT00000062124_Cluster15693-008 | bZIP | DNA binding [Zea mays] | 2443 | 0 | 0 | 0 |
| SALT00000075935_Cluster15693-009 | bZIP | DNA binding [Zea mays] | 2575 | 0 | 0 | 0 |
| SALT00000083439_Cluster15693-010 | bZIP | DNA binding [Zea mays] | 2695 | 0 | 0 | 0 |
| SALT00000026124_Cluster15693-011 | bZIP | DNA binding [Zea mays] | 2435 | 0 | 0 | 0 |
| SALT00000074243_Cluster15946-001 | bZIP | transcription factor HBP-1 | 2556 | 0 | 0 | 0 |
| SALT00000080904_Cluster16405-001 | bZIP | TPA: bZIP transcription fa | 2524 | 0 | 0 | 0 |
| SALT00000004770_Cluster17069-001 | bZIP | bZIP transcription factor su | 1776 | 0 | 0 | 0 |
| SALT00000005589_Cluster17069-002 | bZIP | bZIP transcription factor su | 1898 | 0 | 0 | 0 |
| SALT00000008982_Cluster17069-003 | bZIP | bZIP transcription factor su | 1952 | 0 | 0 | 0 |
| SALT00000054075_Cluster17069-004 | bZIP | bZIP transcription factor su | 1885 | 0 | 0 | 0 |
| SALT00000074658_Cluster17069-005 | bZIP | transcription factor HBP-1 | 2464 | 0 | 0 | 0 |
| SALT00000077633_Cluster17533-001 | bZIP | bzip transcription factor [E | 2400 | 0 | 0 | 0 |
| SALT00000046594_Cluster18295-001 | bZIP | transcription factor HBP-1 | 1717 | 0 | 0 | 0 |
| SALT00000049637_Cluster18295-002 | bZIP | transcription factor HBP-1 | 2230 | 0 | 0 | 0 |
| SALT00000058233_Cluster18295-003 | bZIP | transcription factor HBP-1 | 2250 | 0 | 0 | 0 |
| SALT00000004012_Cluster18748-001 | bZIP | transcription factor TGA2- | 2190 | 0 | 0 | 0 |
| SALT00000006106_Cluster18961-001 | bZIP | ABA response element bin | 2170 | 0 | 0 | 0 |
| SALT00000060678_Cluster18961-002 | bZIP | ABA response element bin | 1831 | 0 | 0 | 0 |
| SALT00000044876_Cluster18961-003 | bZIP | ABA response element bin | 1668 | 0 | 0 | 0 |
| SALT00000042821_Cluster19720-001 | bZIP | transcription factor HBP-1 | 1517 | 0 | 0 | 0 |
| SALT00000043416_Cluster19720-002 | bZIP | transcription factor HBP-1 | 1532 | 0 | 0 | 0 |

| | | | | | | |
|---|---|---|---|---|---|---|
| SALT00000060750_Cluster19720-003 | bZIP | transcription factor HBP-1 | 2102 | 0 | 0 | 0 |
| SALT00000061893_Cluster19720-004 | bZIP | transcription factor HBP-1 | 1522 | 0 | 0 | 0 |
| SALT00000023211_Cluster19740-001 | bZIP | transcription factor TGA2- | 2099 | 0 | 0 | 0 |
| SALT00000049871_Cluster19740-002 | bZIP | TPA: liguleless2 | 1595 | 0 | 0 | 0 |
| SALT00000053074_Cluster19740-003 | bZIP | transcription factor TGA2- | 1970 | 0 | 0 | 0 |
| SALT00000012065_Cluster20119-001 | bZIP | Transcription factor HBP- | 1739 | 0 | 0 | 0 |
| SALT00000051728_Cluster20119-002 | bZIP | Transcription factor HBP- | 2070 | 0 | 0 | 0 |
| SALT00000001320_Cluster20200-001 | bZIP | Transcription factor HBP- | 1826 | 0 | 0 | 0 |
| SALT00000015125_Cluster20200-002 | bZIP | Transcription factor HBP- | 1769 | 0 | 0 | 0 |
| SALT00000047424_Cluster20200-003 | bZIP | Transcription factor HBP- | 2063 | 0 | 0 | 0 |
| SALT00000055360_Cluster20200-004 | bZIP | Transcription factor HBP- | 1770 | 0 | 0 | 0 |
| SALT00000064984_Cluster20200-005 | bZIP | Transcription factor HBP- | 1666 | 0 | 0 | 0 |
| SALT00000042242_Cluster20200-006 | bZIP | Transcription factor HBP- | 1757 | 0 | 0 | 0 |
| SALT00000005741_Cluster20292-001 | bZIP | ABSCISIC ACID-INSENS | 1746 | 0 | 0 | 0 |
| SALT00000009909_Cluster20292-002 | bZIP | ABSCISIC ACID-INSENS | 1990 | 0 | 0 | 0 |
| SALT00000043581_Cluster20292-003 | bZIP | ABSCISIC ACID-INSENS | 2055 | 0 | 0 | 0 |
| SALT00000039489_Cluster20292-004 | bZIP | ABSCISIC ACID-INSENS | 1704 | 0 | 0 | 0 |
| SALT00000055276_Cluster20308-001 | bZIP | transcription factor HBP-1 | 2054 | 0 | 0 | 0 |
| SALT00000044396_Cluster20324-001 | bZIP | ABA responsive element b | 1505 | up | up | up |
| SALT00000058554_Cluster20324-002 | bZIP | ABA responsive element b | 2053 | 0 | 0 | 0 |
| SALT00000041252_Cluster20324-003 | bZIP | ABA responsive element b | 1650 | 0 | 0 | 0 |
| SALT00000014808_Cluster20403-001 | bZIP | transcription factor TGA2- | 2019 | 0 | 0 | 0 |
| SALT00000054896_Cluster20403-002 | bZIP | transcription factor TGA2- | 1935 | 0 | 0 | 0 |
| SALT00000060145_Cluster20403-003 | bZIP | transcription factor TGA2- | 2047 | 0 | 0 | 0 |
| SALT00000016048_Cluster20860-001 | bZIP | bZIP transcription factor su | 1970 | 0 | 0 | 0 |
| SALT00000045237_Cluster20860-002 | bZIP | bZIP transcription factor su | 2014 | 0 | 0 | 0 |
| SALT00000058528_Cluster20860-003 | bZIP | bZIP transcription factor su | 1833 | 0 | 0 | 0 |
| SALT00000000842_Cluster20941-001 | bZIP | DNA binding [Zea mays] | 1645 | 0 | 0 | 0 |
| SALT00000001257_Cluster20941-002 | bZIP | DNA binding [Zea mays] | 1741 | 0 | 0 | 0 |
| SALT00000009687_Cluster20941-003 | bZIP | DNA binding [Zea mays] | 1692 | 0 | 0 | 0 |
| SALT00000011305_Cluster20941-004 | bZIP | DNA binding [Zea mays] | 1561 | 0 | 0 | 0 |
| SALT00000043815_Cluster20941-005 | bZIP | DNA binding [Zea mays] | 2009 | 0 | 0 | 0 |
| SALT00000020156_Cluster21078-001 | bZIP | bZIP transcription factor su | 1999 | 0 | 0 | 0 |
| SALT00000018496_Cluster21194-001 | bZIP | transcription factor TGA2- | 1621 | 0 | 0 | 0 |
| SALT00000040213_Cluster21194-002 | bZIP | TPA: liguleless2 | 1514 | 0 | 0 | 0 |
| SALT00000047520_Cluster21194-003 | bZIP | TPA: liguleless2 | 1992 | 0 | 0 | 0 |
| SALT00000016892_Cluster22264-001 | bZIP | transcription factor HBP-1 | 1715 | 0 | 0 | 0 |
| SALT00000037641_Cluster22264-002 | bZIP | transcription factor HBP-1 | 1636 | 0 | 0 | 0 |
| SALT00000048318_Cluster22264-003 | bZIP | transcription factor HBP-1 | 1577 | 0 | 0 | 0 |
| SALT00000048486_Cluster22264-004 | bZIP | transcription factor HBP-1 | 1579 | 0 | 0 | 0 |
| SALT00000053186_Cluster22264-005 | bZIP | transcription factor HBP-1 | 1638 | 0 | 0 | 0 |
| SALT00000055217_Cluster22264-006 | bZIP | transcription factor HBP-1 | 1703 | 0 | 0 | 0 |
| SALT00000055300_Cluster22264-007 | bZIP | transcription factor HBP-1 | 1763 | 0 | 0 | 0 |
| SALT00000055375_Cluster22264-008 | bZIP | transcription factor TGA4- | 1922 | 0 | 0 | 0 |
| SALT00000060703_Cluster22264-009 | bZIP | transcription factor HBP-1 | 1777 | 0 | 0 | 0 |
| SALT00000049762_Cluster22347-001 | bZIP | bZIP transcription partial [ | 1917 | 0 | 0 | 0 |
| SALT00000058727_Cluster22347-002 | bZIP | Os08g0543900 [Oryza sati | 1690 | 0 | 0 | 0 |
| SALT00000037034_Cluster22575-001 | bZIP | transcription factor TGA2- | 1750 | 0 | 0 | 0 |
| SALT00000037686_Cluster22575-002 | bZIP | transcription factor TGA2- | 1904 | 0 | 0 | 0 |
| SALT00000037918_Cluster22575-003 | bZIP | transcription factor TGA2- | 1639 | 0 | 0 | 0 |
| SALT00000010966_Cluster22787-001 | bZIP | bZIP transcription factor su | 1791 | 0 | 0 | 0 |
| SALT00000011278_Cluster22787-002 | bZIP | bZIP transcription factor su | 1891 | 0 | 0 | 0 |
| SALT00000024935_Cluster22787-003 | bZIP | bZIP transcription factor su | 1855 | 0 | 0 | 0 |
| SALT00000046726_Cluster22787-004 | bZIP | bZIP transcription factor su | 1801 | 0 | 0 | 0 |
| SALT00000051735_Cluster22787-005 | bZIP | bZIP transcription factor su | 1704 | 0 | 0 | 0 |
| SALT00000093710_Cluster22787-006 | bZIP | bZIP transcription factor su | 1776 | 0 | 0 | 0 |
| SALT00000055274_Cluster22787-008 | bZIP | bZIP transcription factor su | 1852 | 0 | 0 | 0 |
| SALT00000010154_Cluster23211-001 | bZIP | Transcription factor HBP- | 1615 | 0 | 0 | 0 |

| | | | | | | |
|---|---|---|---|---|---|---|
| SALT00000039177_Cluster23211-002 | bZIP | Transcription factor HBP- | 1864 | 0 | 0 | 0 |
| SALT00000042865_Cluster23211-003 | bZIP | Transcription factor HBP- | 1642 | 0 | 0 | 0 |
| SALT00000048441_Cluster23252-001 | bZIP | light-inducible CPRF-2 [Z | 1721 | 0 | 0 | 0 |
| SALT00000054572_Cluster23252-002 | bZIP | ocs element-binding factor | 1862 | 0 | 0 | 0 |
| SALT00000012055_Cluster23252-003 | bZIP | light-inducible CPRF-2 [Z | 1542 | 0 | 0 | 0 |
| SALT00000001297_Cluster23289-001 | bZIP | transcription factor HBP-1 | 1859 | 0 | 0 | 0 |
| SALT00000008707_Cluster23289-002 | bZIP | transcription factor HBP-1 | 1739 | 0 | 0 | 0 |
| SALT00000005246_Cluster23406-001 | bZIP | DNA binding [Zea mays] | 1759 | 0 | 0 | 0 |
| SALT00000017074_Cluster23406-002 | bZIP | DNA binding [Zea mays] | 1763 | 0 | 0 | 0 |
| SALT00000053407_Cluster23406-003 | bZIP | DNA binding [Zea mays] | 1853 | 0 | 0 | up |
| SALT00000002889_Cluster23415-001 | bZIP | transcription factor HBP-1 | 1762 | 0 | 0 | 0 |
| SALT00000005064_Cluster23415-002 | bZIP | transcription factor HBP-1 | 1852 | 0 | 0 | 0 |
| SALT00000039480_Cluster23415-003 | bZIP | transcription factor HBP-1 | 1830 | 0 | 0 | 0 |
| SALT00000042872_Cluster23415-004 | bZIP | transcription factor HBP-1 | 1709 | 0 | 0 | 0 |
| SALT00000055091_Cluster23415-005 | bZIP | transcription factor HBP-1 | 1716 | 0 | 0 | 0 |
| SALT00000003810_Cluster23856-001 | bZIP | ABA responsive element b | 1700 | up | 0 | 0 |
| SALT00000004666_Cluster23856-002 | bZIP | ABA responsive element b | 1826 | 0 | 0 | up |
| SALT00000005397_Cluster23856-003 | bZIP | ABA responsive element b | 1604 | 0 | up | up |
| SALT00000006609_Cluster23856-004 | bZIP | ABA responsive element b | 1664 | 0 | 0 | 0 |
| SALT00000011677_Cluster23856-005 | bZIP | ABA responsive element b | 1674 | 0 | 0 | up |
| SALT00000014609_Cluster23856-006 | bZIP | ABA responsive element b | 1606 | 0 | 0 | 0 |
| SALT00000037603_Cluster23856-007 | bZIP | ABA responsive element b | 1681 | 0 | up | up |
| SALT00000048267_Cluster23856-008 | bZIP | ABA responsive element b | 1573 | up | up | up |
| SALT00000049473_Cluster23856-009 | bZIP | ABA responsive element b | 1652 | 0 | 0 | 0 |
| SALT00000007427_Cluster23860-001 | bZIP | transcription factor RF2a-l | 1550 | 0 | 0 | 0 |
| SALT00000016029_Cluster23860-002 | bZIP | Os09g0516200 [Oryza sati | 1826 | 0 | 0 | 0 |
| SALT00000004851_Cluster23942-001 | bZIP | Os11g0218000 [Oryza sati | 1697 | 0 | 0 | 0 |
| SALT00000005757_Cluster23942-002 | bZIP | Os11g0218000 [Oryza sati | 1645 | 0 | 0 | 0 |
| SALT00000015602_Cluster23942-003 | bZIP | Os11g0218000 [Oryza sati | 1821 | 0 | 0 | 0 |
| SALT00000018383_Cluster23942-004 | bZIP | Os11g0218000 [Oryza sati | 1499 | 0 | 0 | 0 |
| SALT00000006254_Cluster24042-001 | bZIP | transcription factor TGA4 | 1722 | 0 | up | 0 |
| SALT00000009192_Cluster24042-002 | bZIP | TPA: bZIP transcription fa | 1365 | 0 | 0 | 0 |
| SALT00000014007_Cluster24042-003 | bZIP | transcription factor TGA4 | 1598 | 0 | 0 | 0 |
| SALT00000039166_Cluster24042-004 | bZIP | TPA: bZIP transcription fa | 1816 | 0 | 0 | 0 |
| SALT00000049005_Cluster24042-005 | bZIP | transcription factor TGA4 | 1692 | 0 | 0 | 0 |
| SALT00000003058_Cluster24116-001 | bZIP | G-box binding factor 1 iso | 1557 | 0 | 0 | 0 |
| SALT00000011311_Cluster24116-002 | bZIP | G-box binding factor 1 iso | 1552 | 0 | 0 | 0 |
| SALT00000012856_Cluster24116-003 | bZIP | G-box binding factor 1 iso | 1682 | 0 | 0 | 0 |
| SALT00000013006_Cluster24116-004 | bZIP | G-box binding factor 1 iso | 1600 | 0 | 0 | 0 |
| SALT00000013298_Cluster24116-005 | bZIP | G-box binding factor 1 iso | 1719 | 0 | 0 | 0 |
| SALT00000013619_Cluster24116-006 | bZIP | G-box binding factor 1 iso | 1812 | 0 | 0 | 0 |
| SALT00000059912_Cluster24116-007 | bZIP | G-box binding factor 1 iso | 1715 | 0 | 0 | 0 |
| SALT00000008992_Cluster24116-008 | bZIP | G-box binding factor 1 iso | 1811 | 0 | 0 | 0 |
| SALT00000016986_Cluster24116-009 | bZIP | G-box binding factor 1 iso | 1613 | 0 | 0 | 0 |
| SALT00000042708_Cluster24116-010 | bZIP | G-box binding factor 1 iso | 1541 | 0 | 0 | 0 |
| SALT00000005379_Cluster24425-001 | bZIP | cross-pathway control 1 | 1794 | 0 | 0 | 0 |
| SALT00000047854_Cluster24726-001 | bZIP | transcription factor HBP-1 | 1623 | 0 | 0 | 0 |
| SALT00000048552_Cluster24726-002 | bZIP | transcription factor HBP-1 | 1778 | 0 | 0 | 0 |
| SALT00000002136_Cluster25143-001 | bZIP | DNA binding [Zea mays] | 1731 | 0 | 0 | 0 |
| SALT00000065319_Cluster25143-002 | bZIP | DNA binding [Zea mays] | 1756 | 0 | 0 | 0 |
| SALT00000019737_Cluster25538-001 | bZIP | ABSCISIC ACID-INSENS | 1675 | 0 | 0 | 0 |
| SALT00000040154_Cluster25538-002 | bZIP | ABSCISIC ACID-INSENS | 1735 | 0 | 0 | 0 |
| SALT00000041597_Cluster25538-003 | bZIP | ABSCISIC ACID-INSENS | 1732 | 0 | 0 | 0 |
| SALT00000045157_Cluster25538-004 | bZIP | ABSCISIC ACID-INSENS | 1728 | 0 | 0 | 0 |
| SALT00000010770_Cluster25573-001 | bZIP | TPA: bZIP transcription fa | 1635 | 0 | 0 | 0 |
| SALT00000039826_Cluster25573-002 | bZIP | maize Em binding -1a [Zea | 1493 | 0 | 0 | 0 |
| SALT00000054618_Cluster25573-003 | bZIP | maize Em binding -1a [Zea | 1734 | 0 | 0 | 0 |
| SALT00000057467_Cluster25573-004 | bZIP | TPA: bZIP transcription fa | 1610 | 0 | 0 | 0 |

| | | | | | | |
|---|---|---|---|---|---|---|
| SALT00000042031_Cluster25914-001 | bZIP | basic leucine zipper 2-like | 1716 | 0 | 0 | 0 |
| SALT00000048732_Cluster25914-002 | bZIP | basic leucine zipper 2-like | 1596 | 0 | 0 | 0 |
| SALT00000012268_Cluster25914-003 | bZIP | basic leucine zipper 2-like | 1541 | 0 | down | down |
| SALT00000053068_Cluster26795-001 | bZIP | bZIP transcription factor A | 1614 | 0 | 0 | 0 |
| SALT00000055664_Cluster26795-002 | bZIP | bZIP transcription factor A | 1670 | 0 | 0 | 0 |
| SALT00000003969_Cluster26820-001 | bZIP | [Hordeum vulgare vulgare] | 1388 | 0 | 0 | 0 |
| SALT00000007322_Cluster26820-002 | bZIP | ABSCISIC ACID-INSENS | 1494 | 0 | 0 | 0 |
| SALT00000012502_Cluster26820-003 | bZIP | ABSCISIC ACID-INSENS | 1668 | 0 | 0 | 0 |
| SALT00000016525_Cluster26820-004 | bZIP | ABSCISIC ACID-INSENS | 1528 | 0 | 0 | 0 |
| SALT00000018026_Cluster26820-005 | bZIP | ABSCISIC ACID-INSENS | 1661 | 0 | 0 | 0 |
| SALT00000038260_Cluster26820-006 | bZIP | ABSCISIC ACID-INSENS | 1436 | 0 | 0 | 0 |
| SALT00000009314_Cluster27530-001 | bZIP | basic region leucine zipper | 1281 | 0 | 0 | 0 |
| SALT00000019247_Cluster27530-002 | bZIP | basic region leucine zipper | 1634 | 0 | 0 | 0 |
| SALT00000009429_Cluster27557-001 | bZIP | transcription factor HBP-1 | 1583 | 0 | 0 | 0 |
| SALT00000042853_Cluster27557-002 | bZIP | transcription factor HBP-1 | 1633 | 0 | 0 | 0 |
| SALT00000004538_Cluster27615-001 | bZIP | DNA-binding EMBP-1-lik | 1446 | 0 | 0 | 0 |
| SALT00000017082_Cluster27615-002 | bZIP | DNA-binding EMBP-1-lik | 1522 | 0 | 0 | 0 |
| SALT00000037176_Cluster27615-003 | bZIP | DNA-binding EMBP-1-lik | 1630 | 0 | 0 | 0 |
| SALT00000052531_Cluster27615-004 | bZIP | DNA-binding EMBP-1-lik | 1489 | 0 | 0 | 0 |
| SALT00000050517_Cluster27615-005 | bZIP | DNA-binding EMBP-1-lik | 1570 | 0 | 0 | 0 |
| SALT00000005158_Cluster27668-001 | bZIP | transcription factor RF2a-l | 1627 | 0 | 0 | 0 |
| SALT00000057736_Cluster27668-002 | bZIP | transcription factor RF2a-l | 1448 | 0 | 0 | 0 |
| SALT00000017729_Cluster27760-001 | bZIP | bZIP transcription factor su | 1591 | 0 | 0 | 0 |
| SALT00000046087_Cluster27760-002 | bZIP | bZIP transcription factor su | 1623 | 0 | 0 | 0 |
| SALT00000061666_Cluster27760-003 | bZIP | bZIP transcription factor su | 1493 | 0 | 0 | 0 |
| SALT00000003217_Cluster27957-001 | bZIP | transcription factor RF2a-l | 1434 | 0 | 0 | 0 |
| SALT00000043570_Cluster27957-002 | bZIP | transcription factor RF2a-l | 1544 | 0 | 0 | 0 |
| SALT00000045288_Cluster27957-003 | bZIP | transcription factor RF2a-l | 1595 | 0 | 0 | 0 |
| SALT00000047465_Cluster27957-004 | bZIP | transcription factor RF2a-l | 1613 | 0 | 0 | 0 |
| SALT00000003232_Cluster28279-001 | bZIP | ocs element-binding factor | 1595 | 0 | 0 | 0 |
| SALT00000007212_Cluster28279-002 | bZIP | ocs element-binding factor | 1528 | 0 | 0 | 0 |
| SALT00000008286_Cluster28279-003 | bZIP | ocs element-binding factor | 1534 | 0 | 0 | 0 |
| SALT00000037002_Cluster28279-004 | bZIP | ocs element-binding factor | 1549 | 0 | 0 | 0 |
| SALT00000050301_Cluster28279-005 | bZIP | ocs element-binding factor | 1496 | 0 | 0 | 0 |
| SALT00000059992_Cluster28279-006 | bZIP | ocs element-binding factor | 1557 | 0 | 0 | 0 |
| SALT00000054383_Cluster28679-001 | bZIP | G-box binding factor 1 iso | 1575 | 0 | 0 | 0 |
| SALT00000009900_Cluster28700-001 | bZIP | bZIP transcription factor su | 1500 | 0 | 0 | 0 |
| SALT00000056559_Cluster28700-002 | bZIP | bZIP transcription factor su | 1573 | 0 | 0 | 0 |
| SALT00000066154_Cluster28700-003 | bZIP | Os09g0306400 [Oryza sati | 1574 | 0 | 0 | 0 |
| SALT00000012796_Cluster28967-001 | bZIP | Os01g0756200 [Oryza sati | 1560 | 0 | 0 | 0 |
| SALT00000004516_Cluster29260-001 | bZIP | transcription factor RF2b [ | 1465 | 0 | 0 | 0 |
| SALT00000056648_Cluster29260-002 | bZIP | transcription factor RF2b [ | 1546 | 0 | 0 | 0 |
| SALT00000011524_Cluster29306-001 | bZIP | Os01g0867300 [Oryza sati | 1543 | 0 | 0 | 0 |
| SALT00000043891_Cluster29306-002 | bZIP | Os01g0867300 [Oryza sati | 1393 | 0 | 0 | 0 |
| SALT00000049257_Cluster29306-003 | bZIP | Os01g0867300 [Oryza sati | 1452 | 0 | 0 | up |
| SALT00000052880_Cluster29306-004 | bZIP | Os01g0867300 [Oryza sati | 1536 | 0 | 0 | up |
| SALT00000006933_Cluster29360-001 | bZIP | bZIP transcription factor fa | 1540 | 0 | 0 | 0 |
| SALT00000042276_Cluster29641-001 | bZIP | bZIP transcription factor su | 1381 | 0 | 0 | 0 |
| SALT00000043085_Cluster29641-002 | bZIP | transcription factor VIP1-l | 1357 | 0 | 0 | 0 |
| SALT00000049372_Cluster29641-003 | bZIP | TPA: bZIP transcription fa | 1524 | 0 | 0 | 0 |
| SALT00000049501_Cluster29641-004 | bZIP | TPA: bZIP transcription fa | 1294 | 0 | 0 | 0 |
| SALT00000052133_Cluster29729-001 | bZIP | ocs element-binding factor | 1470 | 0 | 0 | 0 |
| SALT00000060029_Cluster29729-002 | bZIP | ocs element-binding factor | 1519 | 0 | 0 | 0 |
| SALT00000011679_Cluster29963-001 | bZIP | opaque2 heterodimerizing | 1504 | 0 | 0 | 0 |
| SALT00000046844_Cluster30132-001 | bZIP | Os10g0531900 [Oryza sati | 1493 | 0 | 0 | 0 |
| SALT00000009936_Cluster30206-001 | bZIP | ocs element-binding factor | 1356 | 0 | 0 | 0 |
| SALT00000016203_Cluster30206-002 | bZIP | ocs element-binding factor | 1325 | 0 | 0 | 0 |
| SALT00000058830_Cluster30206-003 | bZIP | ocs element-binding factor | 1446 | 0 | 0 | up |

| | | | | | | |
|---|---|---|---|---|---|---|
| SALT00000060297_Cluster30206-004 | bZIP | ocs element-binding factor | 1488 | 0 | 0 | 0 |
| SALT00000056824_Cluster30243-001 | bZIP | ocs element-binding factor | 1486 | 0 | 0 | 0 |
| SALT00000045478_Cluster31746-001 | bZIP | ocs element-binding factor | 1268 | 0 | 0 | 0 |
| SALT00000024912_Cluster31832-001 | bZIP | light-inducible CPRF-2 [Z | 1041 | 0 | 0 | up |
| SALT00000039680_Cluster31832-002 | bZIP | light-inducible CPRF-2 [Z | 1241 | down | down | 0 |
| SALT00000010555_Cluster31848-001 | bZIP | Os05g0129300 [Oryza sati | 1236 | 0 | 0 | up |
| SALT00000047381_Cluster49-005 | C2H2 | Os01g0534800 [Oryza sati | 1980 | 0 | 0 | 0 |
| SALT00000032729_Cluster570-002 | C2H2 | Os09g0566100 [Oryza sati | 3542 | 0 | 0 | 0 |
| SALT00000032742_Cluster570-003 | C2H2 | Os09g0566100 [Oryza sati | 3581 | 0 | 0 | 0 |
| SALT00000066113_Cluster570-004 | C2H2 | Os09g0566100 [Oryza sati | 2518 | 0 | 0 | 0 |
| SALT00000088876_Cluster570-005 | C2H2 | Os09g0566100 [Oryza sati | 3656 | 0 | 0 | 0 |
| SALT00000087836_Cluster570-007 | C2H2 | Os09g0566100 [Oryza sati | 3836 | 0 | 0 | 0 |
| SALT00000090553_Cluster927-004 | C2H2 | probable E3 ubiquitin ligas | 5572 | 0 | 0 | 0 |
| SALT00000034296_Cluster1225-001 | C2H2 | lysine-specific demethylas | 4615 | 0 | 0 | 0 |
| SALT00000076110_Cluster1225-002 | C2H2 | Lysine-specific demethylas | 2547 | 0 | 0 | 0 |
| SALT00000084218_Cluster1225-003 | C2H2 | lysine-specific demethylas | 4513 | 0 | 0 | 0 |
| SALT00000087789_Cluster1225-005 | C2H2 | lysine-specific demethylas | 4399 | 0 | 0 | 0 |
| SALT00000092261_Cluster1225-007 | C2H2 | lysine-specific demethylas | 4507 | 0 | 0 | 0 |
| SALT00000092909_Cluster1225-008 | C2H2 | lysine-specific demethylas | 3956 | 0 | 0 | 0 |
| SALT00000090248_Cluster1225-010 | C2H2 | Lysine-specific demethylas | 4388 | 0 | 0 | 0 |
| SALT00000001865_Cluster2289-001 | C2H2 | SUPPRESSOR OF FRI 4 [ | 1629 | 0 | 0 | 0 |
| SALT00000026208_Cluster2289-002 | C2H2 | Os09g0560900 [Oryza sati | 2499 | 0 | 0 | 0 |
| SALT00000026721_Cluster2289-003 | C2H2 | Os09g0560900 [Oryza sati | 2446 | 0 | 0 | 0 |
| SALT00000030527_Cluster2289-004 | C2H2 | Os09g0560900 [Oryza sati | 2564 | 0 | 0 | 0 |
| SALT00000030950_Cluster2289-005 | C2H2 | Os09g0560900 [Oryza sati | 2682 | 0 | 0 | 0 |
| SALT00000031350_Cluster2289-006 | C2H2 | Os09g0560900 [Oryza sati | 2580 | 0 | 0 | 0 |
| SALT00000056620_Cluster2289-007 | C2H2 | Os09g0560900 [Oryza sati | 1470 | 0 | 0 | 0 |
| SALT00000060402_Cluster2289-008 | C2H2 | SUPPRESSOR OF FRI 4 [ | 1242 | 0 | 0 | 0 |
| SALT00000064759_Cluster2289-009 | C2H2 | Os09g0560900 [Oryza sati | 2464 | 0 | 0 | 0 |
| SALT00000066968_Cluster2289-010 | C2H2 | Os09g0560900 [Oryza sati | 2380 | 0 | 0 | 0 |
| SALT00000070806_Cluster2289-011 | C2H2 | Os09g0560900 [Oryza sati | 2521 | 0 | 0 | 0 |
| SALT00000073697_Cluster2289-013 | C2H2 | Os09g0560900 [Oryza sati | 2534 | 0 | 0 | 0 |
| SALT00000076535_Cluster2289-014 | C2H2 | Os09g0560900 [Oryza sati | 2515 | 0 | 0 | 0 |
| SALT00000035816_Cluster2289-015 | C2H2 | SUPPRESSOR OF FRI 4 [ | 1551 | 0 | 0 | 0 |
| SALT00000004914_Cluster2604-001 | C2H2 | [Hordeum vulgare vulgare | 2121 | 0 | 0 | 0 |
| SALT00000023968_Cluster2604-002 | C2H2 | [Hordeum vulgare vulgare | 2523 | 0 | 0 | 0 |
| SALT00000025966_Cluster2604-003 | C2H2 | [Hordeum vulgare vulgare | 2584 | 0 | 0 | 0 |
| SALT00000056615_Cluster2604-005 | C2H2 | [Hordeum vulgare vulgare | 2303 | 0 | 0 | 0 |
| SALT00000063812_Cluster2604-007 | C2H2 | zinc finger [Oryza sativa J | 2583 | 0 | 0 | 0 |
| SALT00000064910_Cluster2604-008 | C2H2 | [Hordeum vulgare vulgare | 2323 | 0 | 0 | 0 |
| SALT00000066959_Cluster2604-009 | C2H2 | [Hordeum vulgare vulgare | 2617 | 0 | 0 | 0 |
| SALT00000072551_Cluster2604-011 | C2H2 | [Hordeum vulgare vulgare | 2633 | 0 | 0 | 0 |
| SALT00000081849_Cluster2604-012 | C2H2 | [Hordeum vulgare vulgare | 2381 | 0 | 0 | 0 |
| SALT00000064322_Cluster2604-013 | C2H2 | zinc finger [Oryza sativa J | 2791 | 0 | 0 | 0 |
| SALT00000074842_Cluster2604-014 | C2H2 | zinc finger [Oryza sativa J | 2629 | 0 | 0 | 0 |
| SALT00000000157_Cluster3494-001 | C2H2 | LOC100282635 isoform X | 1751 | 0 | 0 | 0 |
| SALT00000002812_Cluster3494-002 | C2H2 | LOC100282635 isoform X | 1542 | 0 | 0 | 0 |
| SALT00000006124_Cluster3494-003 | C2H2 | LOC100282635 isoform X | 1691 | 0 | 0 | 0 |
| SALT00000011519_Cluster3494-004 | C2H2 | LOC100282635 isoform X | 1900 | 0 | 0 | 0 |
| SALT00000011876_Cluster3494-005 | C2H2 | LOC100282635 isoform X | 1355 | 0 | 0 | 0 |
| SALT00000045120_Cluster3494-006 | C2H2 | LOC100282635 isoform X | 1681 | 0 | 0 | 0 |
| SALT00000093688_Cluster3494-008 | C2H2 | LOC100282635 isoform X | 1671 | 0 | 0 | 0 |
| SALT00000037422_Cluster4047-001 | C2H2 | LOC100282635 isoform X | 4208 | 0 | 0 | up |
| SALT00000002832_Cluster4767-001 | C2H2 | LOC542199 isoform X1 [Z | 2299 | 0 | 0 | 0 |
| SALT00000020855_Cluster4767-003 | C2H2 | LOC542199 isoform X1 [Z | 2610 | 0 | 0 | 0 |
| SALT00000024146_Cluster4767-004 | C2H2 | LOC542199 isoform X1 [Z | 2397 | 0 | 0 | 0 |
| SALT00000028259_Cluster4767-005 | C2H2 | LOC542199 isoform X1 [Z | 2450 | 0 | 0 | 0 |
| SALT00000055928_Cluster4767-006 | C2H2 | LOC542199 isoform X1 [Z | 2152 | 0 | 0 | 0 |

| | | | | | | |
|---|---|---|---|---|---|---|
| SALT00000065115_Cluster4767-007 | C2H2 | LOC542199 isoform X1 [Z | 2529 | 0 | 0 | 0 |
| SALT00000073453_Cluster4767-008 | C2H2 | LOC542199 isoform X1 [Z | 2466 | 0 | 0 | 0 |
| SALT00000074303_Cluster4767-009 | C2H2 | LOC542199 isoform X1 [Z | 2967 | 0 | 0 | 0 |
| SALT00000081531_Cluster4767-010 | C2H2 | LOC542199 isoform X1 [Z | 2426 | 0 | 0 | 0 |
| SALT00000087318_Cluster4767-011 | C2H2 | LOC542199 isoform X1 [Z | 4024 | 0 | 0 | 0 |
| SALT00000091123_Cluster4767-012 | C2H2 | LOC542199 isoform X1 [Z | 3798 | 0 | 0 | 0 |
| SALT00000063132_Cluster4767-013 | C2H2 | LOC542199 isoform X1 [Z | 2473 | 0 | 0 | 0 |
| SALT00000019583_Cluster7065-002 | C2H2 | zinc finger MAGPIE-like i | 1766 | 0 | 0 | 0 |
| SALT00000044177_Cluster7065-003 | C2H2 | zinc finger MAGPIE-like i | 1644 | 0 | 0 | 0 |
| SALT00000051859_Cluster7065-004 | C2H2 | zinc finger MAGPIE-like i | 2177 | 0 | 0 | 0 |
| SALT00000087936_Cluster7065-005 | C2H2 | zinc finger MAGPIE-like i | 3627 | 0 | 0 | 0 |
| SALT00000039877_Cluster7065-006 | C2H2 | zinc finger MAGPIE-like i | 1949 | 0 | 0 | 0 |
| SALT00000075553_Cluster7362-001 | C2H2 | lysine-specific demethylas | 2755 | 0 | 0 | 0 |
| SALT00000089124_Cluster7362-002 | C2H2 | lysine-specific demethylas | 3582 | 0 | 0 | 0 |
| SALT00000005830_Cluster7691-001 | C2H2 | indeterminate-domain chlo | 1810 | 0 | 0 | 0 |
| SALT00000027569_Cluster7691-003 | C2H2 | indeterminate-domain chlo | 3298 | 0 | 0 | 0 |
| SALT00000067043_Cluster7691-004 | C2H2 | indeterminate-domain chlo | 2521 | 0 | 0 | 0 |
| SALT00000086633_Cluster7691-005 | C2H2 | indeterminate-domain chlo | 3529 | 0 | 0 | 0 |
| SALT00000063914_Cluster8578-003 | C2H2 | lysine-specific demethylas | 2956 | 0 | 0 | 0 |
| SALT00000002480_Cluster9608-001 | C2H2 | zinc finger STOP1 homolo | 1991 | 0 | 0 | 0 |
| SALT00000002703_Cluster9608-002 | C2H2 | nucleic acid binding [Zea r | 1805 | 0 | 0 | 0 |
| SALT00000004863_Cluster9608-003 | C2H2 | nucleic acid binding isofor | 1846 | 0 | 0 | 0 |
| SALT00000005456_Cluster9608-004 | C2H2 | nucleic acid binding [Zea r | 1679 | 0 | 0 | 0 |
| SALT00000011123_Cluster9608-005 | C2H2 | zinc finger STOP1 homolo | 1971 | 0 | 0 | 0 |
| SALT00000011301_Cluster9608-006 | C2H2 | zinc finger STOP1 homolo | 1812 | 0 | 0 | 0 |
| SALT00000018939_Cluster9608-007 | C2H2 | nucleic acid binding [Zea r | 1793 | 0 | 0 | 0 |
| SALT00000037011_Cluster9608-008 | C2H2 | nucleic acid binding isofor | 1848 | 0 | 0 | 0 |
| SALT00000043505_Cluster9608-010 | C2H2 | zinc finger STOP1 homolo | 2008 | 0 | 0 | 0 |
| SALT00000048228_Cluster9608-012 | C2H2 | nucleic acid binding [Zea r | 3171 | 0 | 0 | 0 |
| SALT00000050159_Cluster9608-014 | C2H2 | zinc finger STOP1 homolo | 1880 | 0 | 0 | 0 |
| SALT00000056204_Cluster9608-015 | C2H2 | zinc finger STOP1 homolo | 2040 | 0 | 0 | 0 |
| SALT00000056502_Cluster9608-016 | C2H2 | zinc finger STOP1 homolo | 1960 | 0 | 0 | 0 |
| SALT00000001502_Cluster10141-001 | C2H2 | zinc finger STOP1 homolo | 2037 | 0 | 0 | 0 |
| SALT00000004766_Cluster10141-002 | C2H2 | zinc finger STOP1 homolo | 2078 | 0 | 0 | 0 |
| SALT00000005041_Cluster10141-003 | C2H2 | zinc finger STOP1 homolo | 2020 | 0 | 0 | 0 |
| SALT00000007247_Cluster10141-004 | C2H2 | zinc finger STOP1 homolo | 2124 | 0 | 0 | 0 |
| SALT00000009159_Cluster10141-005 | C2H2 | zinc finger STOP1 homolo | 2135 | 0 | 0 | 0 |
| SALT00000012425_Cluster10141-006 | C2H2 | zinc finger STOP1 homolo | 2167 | 0 | 0 | 0 |
| SALT00000016381_Cluster10141-007 | C2H2 | nucleic acid binding [Zea r | 1725 | 0 | 0 | 0 |
| SALT00000023330_Cluster10141-008 | C2H2 | zinc finger STOP1 homolo | 2615 | 0 | 0 | 0 |
| SALT00000027895_Cluster10141-009 | C2H2 | zinc finger STOP1 homolo | 2424 | 0 | 0 | 0 |
| SALT00000031705_Cluster10141-010 | C2H2 | zinc finger STOP1 homolo | 2086 | 0 | 0 | 0 |
| SALT00000034868_Cluster10141-011 | C2H2 | zinc finger STOP1 homolo | 2007 | 0 | 0 | 0 |
| SALT00000036467_Cluster10141-012 | C2H2 | zinc finger STOP1 homolo | 1804 | 0 | 0 | 0 |
| SALT00000037065_Cluster10141-013 | C2H2 | zinc finger STOP1 homolo | 1897 | 0 | 0 | 0 |
| SALT00000041435_Cluster10141-014 | C2H2 | nucleic acid binding [Zea r | 1618 | 0 | 0 | 0 |
| SALT00000042064_Cluster10141-015 | C2H2 | zinc finger STOP1 homolo | 2096 | 0 | 0 | down |
| SALT00000045543_Cluster10141-016 | C2H2 | zinc finger STOP1 homolo | 2056 | 0 | 0 | 0 |
| SALT00000046175_Cluster10141-017 | C2H2 | zinc finger STOP1 homolo | 1969 | 0 | 0 | 0 |
| SALT00000049177_Cluster10141-018 | C2H2 | zinc finger STOP1 homolo | 1974 | 0 | 0 | 0 |
| SALT00000049398_Cluster10141-019 | C2H2 | zinc finger STOP1 homolo | 2048 | 0 | 0 | 0 |
| SALT00000050982_Cluster10141-020 | C2H2 | zinc finger STOP1 homolo | 2085 | 0 | 0 | 0 |
| SALT00000056562_Cluster10141-021 | C2H2 | zinc finger STOP1 homolo | 2070 | 0 | 0 | 0 |
| SALT00000058602_Cluster10141-022 | C2H2 | zinc finger STOP1 homolo | 1909 | 0 | 0 | 0 |
| SALT00000061439_Cluster10141-023 | C2H2 | zinc finger STOP1 homolo | 1968 | 0 | 0 | 0 |
| SALT00000073062_Cluster10141-024 | C2H2 | zinc finger STOP1 homolo | 3089 | 0 | 0 | 0 |
| SALT00000073385_Cluster10141-025 | C2H2 | zinc finger STOP1 homolo | 2234 | 0 | 0 | 0 |
| SALT00000041135_Cluster10141-026 | C2H2 | STOP1 [Triticum aestivum | 2343 | 0 | 0 | 0 |

| | | | | | | |
|---|---|---|---|---|---|---|
| SALT00000058069_Cluster10833-013 | C2H2 | Os01g0140100 [Oryza sati | 2993 | 0 | 0 | 0 |
| SALT00000038547_Cluster11420-001 | C2H2 | indeterminate-domain 1-lil | 1472 | 0 | 0 | 0 |
| SALT00000080048_Cluster11420-002 | C2H2 | zinc finger MAGPIE-like [ | 2922 | 0 | 0 | 0 |
| SALT00000008153_Cluster11777-001 | C2H2 | Os09g0560900 [Oryza sati | 1628 | 0 | 0 | 0 |
| SALT00000025279_Cluster11777-002 | C2H2 | SUPPRESSOR OF FRI 4 [ | 2618 | 0 | 0 | 0 |
| SALT00000030478_Cluster11777-003 | C2H2 | SUPPRESSOR OF FRI 4 [ | 2586 | 0 | 0 | 0 |
| SALT00000043522_Cluster11777-004 | C2H2 | Os09g0560900 [Oryza sati | 1685 | 0 | 0 | 0 |
| SALT00000060059_Cluster11777-006 | C2H2 | SUPPRESSOR OF FRI 4 [ | 1765 | 0 | 0 | 0 |
| SALT00000060266_Cluster11777-007 | C2H2 | SUPPRESSOR OF FRI 4 [ | 1721 | 0 | 0 | 0 |
| SALT00000077025_Cluster11777-008 | C2H2 | Os09g0560900 [Oryza sati | 2888 | 0 | 0 | 0 |
| SALT00000059666_Cluster11777-009 | C2H2 | SUPPRESSOR OF FRI 4 [ | 1685 | 0 | 0 | 0 |
| SALT00000082659_Cluster12564-002 | C2H2 | zinc finger MSN2 4 | 2814 | 0 | 0 | 0 |
| SALT00000067682_Cluster12822-001 | C2H2 | Os05g0106000 [Oryza sati | 2790 | 0 | 0 | 0 |
| SALT00000035929_Cluster13078-001 | C2H2 | zinc finger X-linked ZXD | 1549 | 0 | 0 | 0 |
| SALT00000036982_Cluster13078-002 | C2H2 | transcription factor IIIA-lil | 1714 | 0 | 0 | 0 |
| SALT00000048988_Cluster13078-003 | C2H2 | zinc finger X-linked ZXD | 1530 | 0 | 0 | 0 |
| SALT00000075468_Cluster13078-004 | C2H2 | transcription factor IIIA-lil | 2767 | 0 | 0 | 0 |
| SALT00000078453_Cluster13078-005 | C2H2 | transcription factor IIIA-lil | 2839 | 0 | 0 | 0 |
| SALT00000061969_Cluster13078-006 | C2H2 | pentatricopeptide repeat-cc | 2440 | 0 | 0 | 0 |
| SALT00000067916_Cluster13247-001 | C2H2 | zinc finger transcription fa | 2753 | 0 | 0 | 0 |
| SALT00000000965_Cluster13321-001 | C2H2 | LOC100282635 isoform X | 1855 | 0 | 0 | 0 |
| SALT00000014643_Cluster13321-002 | C2H2 | LOC100282635 isoform X | 1750 | 0 | 0 | 0 |
| SALT00000020459_Cluster13321-003 | C2H2 | LOC100282635 isoform X | 1761 | 0 | 0 | 0 |
| SALT00000020548_Cluster13321-004 | C2H2 | LOC100282635 isoform X | 1661 | 0 | 0 | 0 |
| SALT00000055924_Cluster13321-005 | C2H2 | LOC100282635 isoform X | 1692 | 0 | 0 | 0 |
| SALT00000070591_Cluster13321-006 | C2H2 | LOC100282635 isoform X | 2747 | 0 | 0 | 0 |
| SALT00000028131_Cluster13659-001 | C2H2 | [Hordeum vulgare vulgare | 2720 | 0 | 0 | 0 |
| SALT00000040951_Cluster13659-002 | C2H2 | [Hordeum vulgare vulgare | 1899 | 0 | 0 | 0 |
| SALT00000019546_Cluster13780-001 | C2H2 | LOC542199 isoform X1 [Z | 2031 | 0 | 0 | 0 |
| SALT00000021098_Cluster13780-002 | C2H2 | LOC542199 isoform X1 [Z | 2427 | 0 | 0 | 0 |
| SALT00000064207_Cluster13780-003 | C2H2 | LOC542199 isoform X1 [Z | 2413 | 0 | 0 | 0 |
| SALT00000065662_Cluster13780-004 | C2H2 | LOC542199 isoform X1 [Z | 2615 | 0 | 0 | 0 |
| SALT00000070842_Cluster13780-005 | C2H2 | LOC542199 isoform X1 [Z | 2479 | 0 | 0 | 0 |
| SALT00000083401_Cluster13780-006 | C2H2 | LOC542199 isoform X1 [Z | 2710 | 0 | 0 | 0 |
| SALT00000067864_Cluster13998-001 | C2H2 | zinc finger 7 [Zea mays] | 2645 | 0 | 0 | 0 |
| SALT00000073408_Cluster13998-002 | C2H2 | zinc finger 7 [Zea mays] | 2695 | 0 | 0 | 0 |
| SALT00000050026_Cluster13998-003 | C2H2 | zinc finger 7 [Zea mays] | 1286 | 0 | 0 | 0 |
| SALT00000081168_Cluster14358-001 | C2H2 | zinc finger msn2 4 | 2668 | 0 | 0 | 0 |
| SALT00000001721_Cluster14433-001 | C2H2 | indeterminate-domain chlo | 2117 | 0 | 0 | 0 |
| SALT00000001991_Cluster14433-002 | C2H2 | indeterminate-domain chlo | 2007 | 0 | 0 | 0 |
| SALT00000015826_Cluster14433-003 | C2H2 | indeterminate-domain chlo | 1885 | 0 | 0 | 0 |
| SALT00000029420_Cluster14433-004 | C2H2 | indeterminate-domain chlo | 2663 | 0 | 0 | 0 |
| SALT00000069606_Cluster14433-005 | C2H2 | indeterminate-domain chlo | 2443 | 0 | 0 | 0 |
| SALT00000027984_Cluster14723-001 | C2H2 | LOC542697 isoform X4 [Z | 2644 | 0 | 0 | 0 |
| SALT00000051070_Cluster14723-002 | C2H2 | INDETERMINATE-relate | 1580 | 0 | 0 | 0 |
| SALT00000072376_Cluster14723-003 | C2H2 | LOC542697 isoform X4 [Z | 2599 | 0 | 0 | down |
| SALT00000077892_Cluster14723-004 | C2H2 | LOC542697 isoform X4 [Z | 2736 | 0 | 0 | 0 |
| SALT00000053352_Cluster14723-005 | C2H2 | INDETERMINATE-relate | 2254 | 0 | 0 | 0 |
| SALT00000021428_Cluster14846-001 | C2H2 | indeterminate-domain chlo | 2542 | 0 | 0 | 0 |
| SALT00000021697_Cluster14846-002 | C2H2 | indeterminate-domain chlo | 2435 | 0 | 0 | 0 |
| SALT00000023430_Cluster14846-003 | C2H2 | indeterminate-domain chlo | 2527 | 0 | 0 | 0 |
| SALT00000062085_Cluster14846-004 | C2H2 | indeterminate-domain chlo | 2496 | 0 | 0 | 0 |
| SALT00000062618_Cluster14846-005 | C2H2 | indeterminate-domain chlo | 2332 | 0 | 0 | 0 |
| SALT00000071200_Cluster14846-006 | C2H2 | indeterminate-domain chlo | 2515 | 0 | 0 | 0 |
| SALT00000076861_Cluster14846-007 | C2H2 | indeterminate-domain chlo | 2637 | 0 | 0 | 0 |
| SALT00000078879_Cluster14846-008 | C2H2 | indeterminate-domain chlo | 2587 | 0 | 0 | 0 |
| SALT00000080434_Cluster14846-009 | C2H2 | indeterminate-domain chlo | 2470 | 0 | 0 | 0 |
| SALT00000080707_Cluster14846-010 | C2H2 | indeterminate-domain chlo | 2592 | 0 | 0 | 0 |

| | | | | | | |
|---|---|---|---|---|---|---|
| SALT00000029961_Cluster14846-011 | C2H2 | indeterminate-domain chlo | 2353 | 0 | 0 | 0 |
| SALT00000023416_Cluster15014-001 | C2H2 | Os09g0560900 [Oryza sati | 2623 | 0 | 0 | 0 |
| SALT00000074315_Cluster15014-002 | C2H2 | Os09g0560900 [Oryza sati | 2581 | 0 | 0 | 0 |
| SALT00000066866_Cluster15014-003 | C2H2 | SUPPRESSOR OF FRI 4 [ | 1623 | 0 | 0 | 0 |
| SALT00000029587_Cluster15093-001 | C2H2 | zinc finger 343-like [Setari | 2618 | 0 | 0 | 0 |
| SALT00000030826_Cluster15805-001 | C2H2 | lysine-specific demethylas | 2526 | 0 | 0 | 0 |
| SALT00000063674_Cluster15805-002 | C2H2 | lysine-specific demethylas | 2566 | 0 | 0 | 0 |
| SALT00000017020_Cluster16005-001 | C2H2 | PREDICTED: uncharacter | 2056 | 0 | 0 | 0 |
| SALT00000056021_Cluster16005-003 | C2H2 | PREDICTED: uncharacter | 2420 | 0 | 0 | 0 |
| SALT00000080954_Cluster16005-004 | C2H2 | PREDICTED: hypothetica | 2552 | 0 | 0 | 0 |
| SALT00000022461_Cluster16054-001 | C2H2 | SUPPRESSOR OF FRI 4 [ | 2547 | 0 | 0 | 0 |
| SALT00000063799_Cluster16054-002 | C2H2 | SUPPRESSOR OF FRI 4 [ | 2453 | 0 | 0 | 0 |
| SALT00000080642_Cluster16054-003 | C2H2 | SUPPRESSOR OF FRI 4 [ | 2506 | 0 | 0 | 0 |
| SALT00000024758_Cluster16980-001 | C2H2 | LOC542199 isoform X1 [Z | 2472 | 0 | 0 | 0 |
| SALT00000026813_Cluster16980-002 | C2H2 | LOC542199 isoform X1 [Z | 2421 | 0 | 0 | 0 |
| SALT00000042077_Cluster18953-001 | C2H2 | [Hordeum vulgare vulgare] | 1932 | 0 | 0 | 0 |
| SALT00000045365_Cluster18953-002 | C2H2 | [Hordeum vulgare vulgare] | 2171 | 0 | 0 | 0 |
| SALT00000036306_Cluster18953-003 | C2H2 | [Hordeum vulgare vulgare] | 2139 | 0 | 0 | 0 |
| SALT00000002144_Cluster19008-001 | C2H2 | zinc finger MAGPIE-like i | 2165 | 0 | 0 | 0 |
| SALT00000010795_Cluster19008-002 | C2H2 | zinc finger MAGPIE-like i | 2101 | 0 | 0 | 0 |
| SALT00000039062_Cluster19008-003 | C2H2 | zinc finger MAGPIE-like i | 2145 | 0 | 0 | 0 |
| SALT00000012786_Cluster19107-001 | C2H2 | Os12g0502700 [Oryza sati | 1951 | 0 | 0 | 0 |
| SALT00000016883_Cluster19107-002 | C2H2 | Os12g0502700 [Oryza sati | 1992 | 0 | 0 | 0 |
| SALT00000051506_Cluster19107-003 | C2H2 | Os12g0502700 [Oryza sati | 2158 | 0 | 0 | 0 |
| SALT00000036707_Cluster19107-004 | C2H2 | Os12g0502700 [Oryza sati | 2092 | 0 | 0 | 0 |
| SALT00000008181_Cluster19315-001 | C2H2 | indeterminate-domain chlo | 1653 | 0 | 0 | 0 |
| SALT00000011736_Cluster19315-002 | C2H2 | indeterminate-domain chlo | 1964 | 0 | 0 | 0 |
| SALT00000015257_Cluster19315-003 | C2H2 | indeterminate-domain chlo | 2090 | 0 | 0 | 0 |
| SALT00000047356_Cluster19315-004 | C2H2 | indeterminate-domain chlo | 1980 | 0 | 0 | 0 |
| SALT00000050602_Cluster19315-005 | C2H2 | indeterminate-domain chlo | 1859 | 0 | 0 | 0 |
| SALT00000058188_Cluster19315-006 | C2H2 | indeterminate-domain chlo | 1765 | 0 | 0 | 0 |
| SALT00000060135_Cluster19315-007 | C2H2 | indeterminate-domain chlo | 1839 | 0 | 0 | 0 |
| SALT00000061528_Cluster19315-008 | C2H2 | indeterminate-domain chlo | 2137 | 0 | 0 | 0 |
| SALT00000016370_Cluster19465-001 | C2H2 | indeterminate-domain chlo | 2122 | 0 | 0 | 0 |
| SALT00000044694_Cluster19465-002 | C2H2 | indeterminate-domain chlo | 1577 | 0 | 0 | 0 |
| SALT00000054480_Cluster19465-003 | C2H2 | indeterminate-domain chlo | 1950 | 0 | 0 | down |
| SALT00000057310_Cluster19465-004 | C2H2 | indeterminate-domain chlo | 1976 | 0 | 0 | 0 |
| SALT00000059051_Cluster19465-005 | C2H2 | indeterminate-domain chlo | 2022 | 0 | 0 | 0 |
| SALT00000006787_Cluster19611-001 | C2H2 | zinc finger MAGPIE-like [ | 2111 | 0 | 0 | 0 |
| SALT00000007924_Cluster19611-002 | C2H2 | zinc finger MAGPIE-like [ | 1767 | 0 | 0 | 0 |
| SALT00000016163_Cluster19611-003 | C2H2 | zinc finger MAGPIE-like [ | 2013 | 0 | 0 | 0 |
| SALT00000060680_Cluster19611-004 | C2H2 | zinc finger MAGPIE-like [ | 2043 | 0 | 0 | up |
| SALT00000061464_Cluster19611-005 | C2H2 | zinc finger MAGPIE-like [ | 1834 | 0 | 0 | 0 |
| SALT00000016033_Cluster20350-001 | C2H2 | Transcription factor IIIA | 1797 | 0 | 0 | 0 |
| SALT00000057527_Cluster20350-002 | C2H2 | Transcription factor IIIA | 2051 | 0 | 0 | 0 |
| SALT00000060435_Cluster20350-003 | C2H2 | Transcription factor IIIA | 1906 | 0 | 0 | 0 |
| SALT00000063567_Cluster20350-004 | C2H2 | Transcription factor IIIA | 1963 | 0 | 0 | 0 |
| SALT00000062812_Cluster20350-005 | C2H2 | Transcription factor IIIA | 2657 | 0 | 0 | 0 |
| SALT00000015407_Cluster21584-001 | C2H2 | transcription factor IIIA-lil | 1615 | 0 | 0 | 0 |
| SALT00000036869_Cluster21584-002 | C2H2 | transcription factor IIIA-lil | 1966 | 0 | 0 | 0 |
| SALT00000039292_Cluster21584-003 | C2H2 | transcription factor IIIA-lil | 1625 | 0 | 0 | 0 |
| SALT00000056241_Cluster21584-004 | C2H2 | transcription factor IIIA-lil | 1768 | 0 | 0 | 0 |
| SALT00000017000_Cluster21929-001 | C2H2 | transcription factor IIIA-lil | 1944 | 0 | 0 | 0 |
| SALT00000052259_Cluster21929-002 | C2H2 | zinc finger X-linked ZXD/ | 1471 | 0 | 0 | 0 |
| SALT00000046498_Cluster22797-001 | C2H2 | zinc finger 511 [Zea mays] | 1891 | 0 | 0 | 0 |
| SALT00000013609_Cluster25330-001 | C2H2 | Os04g0115500 [Oryza sati | 1745 | up | 0 | 0 |
| SALT00000050760_Cluster25863-001 | C2H2 | zinc C2H2 type family [Ze | 1719 | 0 | 0 | 0 |
| SALT00000018286_Cluster27272-001 | C2H2 | zinc finger [Oryza sativa Ja | 1647 | 0 | 0 | 0 |

| | | | | | | |
|---|---|---|---|---|---|---|
| SALT00000056447_Cluster27272-002 | C2H2 | zinc finger [Oryza sativa Ja | 1584 | 0 | 0 | 0 |
| SALT00000049337_Cluster28797-001 | C2H2 | indeterminate-domain 16 [ | 1569 | 0 | 0 | 0 |
| SALT00000003239_Cluster28888-001 | C2H2 | Os04g0115500 [Oryza sati | 1564 | 0 | 0 | 0 |
| SALT00000012910_Cluster29026-001 | C2H2 | transcription repressor TRI | 1557 | 0 | 0 | 0 |
| SALT00000042034_Cluster29026-002 | C2H2 | transcription repressor TRI | 1540 | 0 | 0 | 0 |
| SALT00000052989_Cluster29026-003 | C2H2 | transcription repressor TRI | 1426 | 0 | 0 | 0 |
| SALT00000047274_Cluster29026-004 | C2H2 | transcription repressor TRI | 1318 | 0 | 0 | 0 |
| SALT00000063958_Cluster29370-002 | C2H2 | SUPPRESSOR OF FRI 4 [ | 1540 | 0 | 0 | 0 |
| SALT00000046021_Cluster29867-001 | C2H2 | Os05g0106000 [Oryza sati | 1511 | 0 | 0 | 0 |
| SALT00000038937_Cluster30113-001 | C2H2 | partial [Oryza sativa Japon | 1494 | 0 | 0 | 0 |
| SALT00000008990_Cluster30210-001 | C2H2 | 26s proteasome regulatory | 1487 | 0 | 0 | 0 |
| SALT00000061752_Cluster30473-002 | C2H2 | Os05g0597100 [Oryza sati | 1470 | 0 | 0 | 0 |
| SALT00000014111_Cluster30571-001 | C2H2 | zinc C2H2 type family [Ze | 1452 | 0 | 0 | 0 |
| SALT00000039573_Cluster30571-002 | C2H2 | zinc C2H2 type family [Ze | 1462 | 0 | 0 | 0 |
| SALT00000054247_Cluster30571-003 | C2H2 | zinc C2H2 type family [Ze | 1437 | 0 | 0 | 0 |
| SALT00000059587_Cluster31045-001 | C2H2 | zinc finger [Oryza sativa Ja | 1415 | 0 | 0 | 0 |
| SALT00000056150_Cluster31200-001 | C2H2 | histone deacetylase HDT2- | 1393 | 0 | 0 | 0 |
| SALT00000049195_Cluster31308-001 | C2H2 | histone deacetylase HDT2- | 1375 | 0 | 0 | 0 |
| SALT00000015838_Cluster31556-001 | C2H2 | zinc finger ID1 | 1319 | 0 | 0 | 0 |
| SALT00000047771_Cluster31609-001 | C2H2 | [Oryza sativa Japonica Gro | 1306 | 0 | 0 | 0 |
| SALT00000012464_Cluster31794-001 | C2H2 | zinc-finger WZF1 [Triticu | 1201 | 0 | 0 | 0 |
| SALT00000018598_Cluster31794-002 | C2H2 | zinc-finger WZF1 [Triticu | 1251 | 0 | 0 | up |
| SALT00000027391_Cluster31794-003 | C2H2 | zinc-finger WZF1 [Triticu | 1183 | 0 | 0 | up |
| SALT00000001517_Cluster31873-001 | C2H2 | zinc-finger WZF1 [Triticu | 1228 | 0 | 0 | up |
| SALT00000003479_Cluster31873-002 | C2H2 | zinc-finger WZF1 [Triticu | 1204 | 0 | 0 | up |
| SALT00000005411_Cluster1386-001 | C3H | Os03g0112700 [Oryza sati | 1624 | 0 | 0 | 0 |
| SALT00000016258_Cluster1386-002 | C3H | Os03g0112700 [Oryza sati | 1998 | 0 | 0 | 0 |
| SALT00000044246_Cluster1386-003 | C3H | hypothetical protein SETI1 | 1845 | 0 | 0 | 0 |
| SALT00000087375_Cluster1386-004 | C3H | Os03g0112700 [Oryza sati | 3612 | 0 | 0 | 0 |
| SALT00000085397_Cluster1386-006 | C3H | zinc finger CCCH domain- | 5335 | 0 | 0 | 0 |
| SALT00000020884_Cluster2221-001 | C3H | zinc finger C-x8-C-x5-C-x | 2556 | 0 | 0 | 0 |
| SALT00000021517_Cluster2221-002 | C3H | zinc finger C-x8-C-x5-C-x | 2661 | 0 | 0 | 0 |
| SALT00000022174_Cluster2221-003 | C3H | zinc finger C-x8-C-x5-C-x | 2645 | 0 | 0 | 0 |
| SALT00000022759_Cluster2221-004 | C3H | zinc finger C-x8-C-x5-C-x | 2642 | 0 | 0 | 0 |
| SALT00000025195_Cluster2221-006 | C3H | zinc finger C-x8-C-x5-C-x | 2512 | 0 | 0 | 0 |
| SALT00000027608_Cluster2221-007 | C3H | zinc finger C-x8-C-x5-C-x | 2585 | 0 | 0 | 0 |
| SALT00000063926_Cluster2221-008 | C3H | zinc finger C-x8-C-x5-C-x | 2544 | 0 | 0 | 0 |
| SALT00000065019_Cluster2221-009 | C3H | zinc finger C-x8-C-x5-C-x | 2560 | 0 | 0 | 0 |
| SALT00000067148_Cluster2221-010 | C3H | zinc finger C-x8-C-x5-C-x | 2646 | 0 | 0 | 0 |
| SALT00000067955_Cluster2221-011 | C3H | zinc finger C-x8-C-x5-C-x | 2814 | 0 | 0 | 0 |
| SALT00000068270_Cluster2221-012 | C3H | zinc finger C-x8-C-x5-C-x | 2742 | 0 | 0 | 0 |
| SALT00000068674_Cluster2221-013 | C3H | zinc finger C-x8-C-x5-C-x | 2516 | 0 | 0 | 0 |
| SALT00000074677_Cluster2221-014 | C3H | zinc finger C-x8-C-x5-C-x | 2773 | 0 | 0 | 0 |
| SALT00000075037_Cluster2221-015 | C3H | zinc finger C-x8-C-x5-C-x | 2429 | 0 | 0 | 0 |
| SALT00000078475_Cluster2221-016 | C3H | zinc finger C-x8-C-x5-C-x | 2547 | 0 | 0 | 0 |
| SALT00000081641_Cluster2221-017 | C3H | zinc finger C-x8-C-x5-C-x | 2585 | 0 | 0 | 0 |
| SALT00000082443_Cluster2221-018 | C3H | zinc finger C-x8-C-x5-C-x | 2622 | 0 | 0 | 0 |
| SALT00000083475_Cluster2221-019 | C3H | zinc finger C-x8-C-x5-C-x | 2697 | 0 | 0 | 0 |
| SALT00000064257_Cluster2221-021 | C3H | zinc finger C-x8-C-x5-C-x | 2714 | 0 | 0 | 0 |
| SALT00000067342_Cluster4757-001 | C3H | zinc finger C-x8-C-x5-C-x | 4026 | 0 | 0 | 0 |
| SALT00000030988_Cluster4941-001 | C3H | zinc finger CCCH domain- | 2432 | 0 | 0 | 0 |
| SALT00000033534_Cluster4941-002 | C3H | zinc finger CCCH domain- | 3988 | 0 | 0 | 0 |
| SALT00000037698_Cluster4941-003 | C3H | zinc finger CCCH domain- | 2286 | 0 | 0 | 0 |
| SALT00000065034_Cluster4941-004 | C3H | zinc finger CCCH domain- | 2688 | 0 | 0 | 0 |
| SALT00000093501_Cluster4941-005 | C3H | zinc finger CCCH domain- | 3854 | 0 | 0 | 0 |
| SALT00000040512_Cluster4947-001 | C3H | zinc finger CCCH domain- | 3987 | 0 | 0 | 0 |
| SALT00000083109_Cluster5279-001 | C3H | zinc finger CCCH domain- | 2592 | 0 | 0 | 0 |
| SALT00000087385_Cluster5279-002 | C3H | zinc finger CCCH domain- | 3921 | 0 | 0 | 0 |

| | | | | | | |
|---|---|---|---|---|---|---|
| SALT00000050199_Cluster5337-001 | C3H | zinc finger CCCH type doı | 1706 | 0 | 0 | 0 |
| SALT00000087662_Cluster5337-002 | C3H | zinc finger CCCH type doı | 3909 | 0 | 0 | 0 |
| SALT00000087950_Cluster5337-003 | C3H | zinc finger CCCH type doı | 3720 | 0 | 0 | 0 |
| SALT00000012220_Cluster5337-004 | C3H | zinc finger CCCH type doı | 1979 | 0 | 0 | up |
| SALT00000090540_Cluster5541-001 | C3H | zinc finger CCCH domain- | 3873 | 0 | 0 | 0 |
| SALT00000025621_Cluster5782-001 | C3H | zinc finger CCCH domain- | 3337 | 0 | 0 | 0 |
| SALT00000031940_Cluster5782-002 | C3H | zinc finger CCCH domain- | 3505 | 0 | 0 | 0 |
| SALT00000033244_Cluster5782-003 | C3H | zinc finger CCCH domain- | 3398 | 0 | 0 | 0 |
| SALT00000033547_Cluster5782-004 | C3H | zinc finger CCCH domain- | 3542 | 0 | 0 | 0 |
| SALT00000066788_Cluster5782-007 | C3H | zinc finger CCCH domain- | 3482 | 0 | 0 | 0 |
| SALT00000086440_Cluster5782-008 | C3H | zinc finger CCCH domain- | 3830 | 0 | 0 | 0 |
| SALT00000027771_Cluster5974-003 | C3H | zinc finger CCCH domain- | 2914 | 0 | 0 | 0 |
| SALT00000028600_Cluster5974-004 | C3H | zinc finger CCCH domain- | 3704 | 0 | 0 | 0 |
| SALT00000028639_Cluster5974-005 | C3H | zinc finger CCCH domain- | 3778 | 0 | 0 | 0 |
| SALT00000030644_Cluster5974-006 | C3H | zinc finger CCCH domain- | 3084 | 0 | 0 | 0 |
| SALT00000031224_Cluster5974-007 | C3H | zinc finger CCCH domain- | 3120 | 0 | 0 | 0 |
| SALT00000031793_Cluster5974-008 | C3H | zinc finger CCCH domain- | 3593 | 0 | 0 | 0 |
| SALT00000031796_Cluster5974-009 | C3H | zinc finger CCCH domain- | 3610 | 0 | 0 | 0 |
| SALT00000033723_Cluster5974-010 | C3H | zinc finger CCCH domain- | 3567 | 0 | 0 | 0 |
| SALT00000034295_Cluster5974-011 | C3H | zinc finger CCCH domain- | 3511 | 0 | 0 | 0 |
| SALT00000073782_Cluster5974-012 | C3H | zinc finger CCCH domain- | 2602 | 0 | 0 | 0 |
| SALT00000080092_Cluster5974-014 | C3H | zinc finger CCCH domain- | 2473 | 0 | 0 | 0 |
| SALT00000081573_Cluster5974-015 | C3H | zinc finger CCCH domain- | 3660 | 0 | 0 | 0 |
| SALT00000086509_Cluster5974-016 | C3H | zinc finger CCCH domain- | 3796 | 0 | 0 | 0 |
| SALT00000087160_Cluster5974-017 | C3H | zinc finger CCCH domain- | 3604 | 0 | 0 | 0 |
| SALT00000050651_Cluster6350-001 | C3H | Zinc finger CCCH domain | 1714 | 0 | 0 | 0 |
| SALT00000086346_Cluster6350-002 | C3H | Zinc finger CCCH domain | 3737 | 0 | 0 | 0 |
| SALT00000050144_Cluster7266-005 | C3H | splicing factor U2af small | 1942 | 0 | 0 | 0 |
| SALT00000092367_Cluster7847-001 | C3H | retrotransposon expressed | 3507 | 0 | 0 | 0 |
| SALT00000036739_Cluster9125-001 | C3H | TPA: RNA-binding zinc fi | 1569 | 0 | 0 | 0 |
| SALT00000004217_Cluster9852-001 | C3H | zinc finger CCCH domain- | 1651 | 0 | 0 | 0 |
| SALT00000010677_Cluster9852-002 | C3H | zinc finger CCCH domain- | 1624 | 0 | 0 | 0 |
| SALT00000021457_Cluster10586-001 | C3H | zinc finger CCCH domain- | 2807 | 0 | 0 | 0 |
| SALT00000021646_Cluster10586-002 | C3H | zinc finger CCCH domain- | 3024 | 0 | 0 | 0 |
| SALT00000055341_Cluster10586-003 | C3H | zinc finger CCCH domain- | 1966 | 0 | 0 | 0 |
| SALT00000063287_Cluster10586-004 | C3H | zinc finger CCCH domain- | 2921 | 0 | 0 | 0 |
| SALT00000069468_Cluster10586-005 | C3H | zinc finger CCCH domain- | 2820 | 0 | 0 | 0 |
| SALT00000072766_Cluster10586-006 | C3H | zinc finger CCCH domain- | 2659 | 0 | 0 | 0 |
| SALT00000074665_Cluster10586-007 | C3H | zinc finger CCCH domain- | 2842 | 0 | 0 | 0 |
| SALT00000022228_Cluster10586-008 | C3H | zinc finger CCCH domain- | 2993 | 0 | 0 | 0 |
| SALT00000000471_Cluster10654-001 | C3H | Zinc finger CCCH type do | 1912 | 0 | 0 | 0 |
| SALT00000005826_Cluster10654-002 | C3H | Zinc finger CCCH type do | 1900 | 0 | 0 | 0 |
| SALT00000016766_Cluster10654-003 | C3H | Zinc finger CCCH type do | 1854 | 0 | 0 | 0 |
| SALT00000017678_Cluster10654-004 | C3H | Zinc finger CCCH type do | 1861 | 0 | 0 | 0 |
| SALT00000050048_Cluster10654-005 | C3H | Zinc finger CCCH type do | 1553 | 0 | 0 | 0 |
| SALT00000057835_Cluster10654-006 | C3H | Zinc finger CCCH type do | 2106 | 0 | 0 | 0 |
| SALT00000068641_Cluster10654-007 | C3H | Zinc finger CCCH type do | 2574 | 0 | 0 | 0 |
| SALT00000070216_Cluster10654-008 | C3H | Zinc finger CCCH type do | 3015 | 0 | 0 | 0 |
| SALT00000073060_Cluster10838-005 | C3H | tRNA-dihydrouridine(47) : | 2427 | 0 | 0 | 0 |
| SALT00000016895_Cluster11197-001 | C3H | Os01g0258700 [Oryza sati | 2040 | 0 | 0 | 0 |
| SALT00000037808_Cluster11197-002 | C3H | Os01g0258700 [Oryza sati | 1995 | 0 | 0 | 0 |
| SALT00000040050_Cluster11197-003 | C3H | Os01g0258700 [Oryza sati | 1569 | 0 | 0 | 0 |
| SALT00000052238_Cluster11197-004 | C3H | Os01g0258700 [Oryza sati | 2064 | 0 | 0 | 0 |
| SALT00000052725_Cluster11197-005 | C3H | Os01g0258700 [Oryza sati | 2104 | 0 | 0 | 0 |
| SALT00000068231_Cluster11197-006 | C3H | Os01g0258700 [Oryza sati | 1839 | 0 | 0 | 0 |
| SALT00000072823_Cluster11197-007 | C3H | Os01g0258700 [Oryza sati | 2947 | 0 | 0 | 0 |
| SALT00000008491_Cluster11400-001 | C3H | zinc finger CCCH type doı | 1637 | 0 | 0 | 0 |
| SALT00000054166_Cluster11400-002 | C3H | zinc finger CCCH type doı | 1560 | 0 | 0 | 0 |

| | | | | | | |
|---|---|---|---|---|---|---|
| SALT00000066140_Cluster11400-003 | C3H | zinc finger CCCH type do | 2924 | 0 | 0 | 0 |
| SALT00000012207_Cluster12031-001 | C3H | [Hordeum vulgare vulgare | 1697 | 0 | 0 | 0 |
| SALT00000013411_Cluster12031-002 | C3H | zinc finger CCCH domain- | 1457 | 0 | 0 | 0 |
| SALT00000013887_Cluster12031-003 | C3H | [Hordeum vulgare vulgare | 1737 | 0 | 0 | 0 |
| SALT00000016001_Cluster12031-004 | C3H | [Hordeum vulgare vulgare | 1637 | 0 | 0 | 0 |
| SALT00000039593_Cluster12031-005 | C3H | [Hordeum vulgare vulgare | 2864 | 0 | 0 | 0 |
| SALT00000052009_Cluster12031-006 | C3H | [Hordeum vulgare vulgare | 1598 | 0 | 0 | 0 |
| SALT00000029673_Cluster12268-002 | C3H | zinc finger CCCH domain- | 2726 | 0 | 0 | 0 |
| SALT00000065238_Cluster12268-003 | C3H | zinc finger CCCH domain- | 2810 | 0 | 0 | 0 |
| SALT00000019129_Cluster12974-001 | C3H | partial [Oryza sativa Japon | 1793 | 0 | 0 | 0 |
| SALT00000041386_Cluster12974-002 | C3H | partial [Oryza sativa Japon | 1967 | 0 | 0 | 0 |
| SALT00000060037_Cluster12974-003 | C3H | partial [Oryza sativa Japon | 1907 | 0 | 0 | 0 |
| SALT00000067517_Cluster12974-004 | C3H | Zinc finger CCCH type do | 2775 | 0 | 0 | 0 |
| SALT00000070717_Cluster12974-005 | C3H | partial [Oryza sativa Japon | 2559 | 0 | 0 | 0 |
| SALT00000076360_Cluster13311-001 | C3H | zinc finger CCCH domain- | 2748 | 0 | 0 | 0 |
| SALT00000072603_Cluster13674-001 | C3H | zinc finger CCCH domain- | 2428 | 0 | 0 | 0 |
| SALT00000053239_Cluster13674-003 | C3H | zinc finger CCCH domain- | 1488 | 0 | 0 | 0 |
| SALT00000021051_Cluster14013-001 | C3H | [Hordeum vulgare vulgare | 2546 | 0 | 0 | 0 |
| SALT00000021733_Cluster14013-002 | C3H | [Hordeum vulgare vulgare | 2589 | 0 | 0 | 0 |
| SALT00000023950_Cluster14013-003 | C3H | [Hordeum vulgare vulgare | 2666 | 0 | 0 | 0 |
| SALT00000026177_Cluster14013-004 | C3H | [Hordeum vulgare vulgare | 2693 | 0 | 0 | 0 |
| SALT00000028218_Cluster14013-005 | C3H | [Hordeum vulgare vulgare | 2580 | 0 | 0 | 0 |
| SALT00000028944_Cluster14013-006 | C3H | [Hordeum vulgare vulgare | 2514 | 0 | 0 | 0 |
| SALT00000064844_Cluster14013-007 | C3H | [Hordeum vulgare vulgare | 2613 | 0 | 0 | 0 |
| SALT00000069237_Cluster14013-008 | C3H | [Hordeum vulgare vulgare | 2466 | 0 | 0 | 0 |
| SALT00000074106_Cluster14013-009 | C3H | [Hordeum vulgare vulgare | 2574 | 0 | 0 | 0 |
| SALT00000079697_Cluster14013-010 | C3H | [Hordeum vulgare vulgare | 2631 | 0 | 0 | 0 |
| SALT00000075337_Cluster14013-011 | C3H | [Hordeum vulgare vulgare | 2467 | 0 | 0 | 0 |
| SALT00000079345_Cluster14411-001 | C3H | zinc finger CCCH domain- | 2665 | 0 | 0 | 0 |
| SALT00000015933_Cluster15556-001 | C3H | zinc finger CCCH domain- | 1836 | 0 | 0 | 0 |
| SALT00000039600_Cluster15556-002 | C3H | partial [Oryza sativa Japon | 1677 | 0 | 0 | 0 |
| SALT00000055567_Cluster15556-003 | C3H | zinc finger CCCH domain- | 1790 | 0 | 0 | 0 |
| SALT00000081032_Cluster15556-004 | C3H | zinc finger CCCH domain- | 2585 | 0 | 0 | 0 |
| SALT00000019872_Cluster15556-005 | C3H | Zinc finger CCCH type do | 1980 | 0 | 0 | 0 |
| SALT00000039995_Cluster15839-001 | C3H | nucleic acid binding [Zea r | 1917 | 0 | 0 | 0 |
| SALT00000041750_Cluster15839-002 | C3H | nucleic acid binding [Zea r | 1877 | 0 | 0 | 0 |
| SALT00000063291_Cluster15839-003 | C3H | nucleic acid binding [Zea r | 2554 | 0 | 0 | 0 |
| SALT00000064113_Cluster15839-004 | C3H | nucleic acid binding [Zea r | 2473 | 0 | 0 | 0 |
| SALT00000081459_Cluster15839-005 | C3H | nucleic acid binding [Zea r | 2564 | 0 | 0 | 0 |
| SALT00000066689_Cluster15901-001 | C3H | Os01g0258700 [Oryza sati | 2559 | 0 | 0 | 0 |
| SALT00000036935_Cluster16559-001 | C3H | zinc finger C-x8-C-x5-C-x | 1322 | 0 | 0 | 0 |
| SALT00000041184_Cluster16559-002 | C3H | zinc finger C-x8-C-x5-C-x | 1357 | 0 | 0 | 0 |
| SALT00000044548_Cluster16559-003 | C3H | zinc finger C-x8-C-x5-C-x | 1415 | 0 | 0 | 0 |
| SALT00000047576_Cluster16559-004 | C3H | zinc finger C-x8-C-x5-C-x | 1231 | 0 | 0 | 0 |
| SALT00000059375_Cluster16559-005 | C3H | zinc finger C-x8-C-x5-C-x | 1271 | 0 | 0 | 0 |
| SALT00000061635_Cluster16559-006 | C3H | zinc finger C-x8-C-x5-C-x | 1290 | 0 | 0 | 0 |
| SALT00000061895_Cluster16559-007 | C3H | zinc finger C-x8-C-x5-C-x | 1421 | 0 | 0 | 0 |
| SALT00000082559_Cluster16559-008 | C3H | zinc finger C-x8-C-x5-C-x | 2510 | 0 | 0 | 0 |
| SALT00000001573_Cluster16771-001 | C3H | nucleic acid binding [Zea r | 1862 | 0 | 0 | 0 |
| SALT00000013644_Cluster16771-002 | C3H | nucleic acid binding [Zea r | 1957 | 0 | 0 | 0 |
| SALT00000043934_Cluster16771-003 | C3H | nucleic acid binding [Zea r | 1868 | 0 | 0 | 0 |
| SALT00000072325_Cluster16771-004 | C3H | Os06g0704300 [Oryza sati | 2492 | 0 | 0 | 0 |
| SALT00000077937_Cluster16771-005 | C3H | Os06g0704300 [Oryza sati | 2438 | 0 | 0 | 0 |
| SALT00000050951_Cluster16771-006 | C3H | nucleic acid binding [Zea r | 1811 | 0 | 0 | 0 |
| SALT00000075406_Cluster17115-002 | C3H | nucleic acid binding [Zea r | 2459 | 0 | 0 | 0 |
| SALT00000026896_Cluster17461-001 | C3H | zinc finger C-x8-C-x5-C-x | 2412 | 0 | 0 | 0 |
| SALT00000040501_Cluster17461-002 | C3H | zinc finger C-x8-C-x5-C-x | 2273 | 0 | 0 | 0 |
| SALT00000041593_Cluster17461-003 | C3H | zinc finger C-x8-C-x5-C-x | 1586 | 0 | 0 | 0 |

| | | | | | | |
|---|---|---|---|---|---|---|
| SALT00000058826_Cluster17461-004 | C3H | zinc finger C-x8-C-x5-C-x | 1948 | 0 | 0 | 0 |
| SALT00000065270_Cluster17461-005 | C3H | zinc finger C-x8-C-x5-C-x | 2250 | 0 | 0 | 0 |
| SALT00000053380_Cluster19389-002 | C3H | zinc finger (C3HC4-type R | 2129 | 0 | 0 | 0 |
| SALT00000058735_Cluster19628-001 | C3H | nucleic acid binding [Zea r | 2110 | 0 | 0 | 0 |
| SALT00000059223_Cluster19628-002 | C3H | nucleic acid binding [Zea r | 1646 | 0 | 0 | 0 |
| SALT00000007887_Cluster19758-001 | C3H | zinc finger CCCH domain- | 1429 | 0 | 0 | down |
| SALT00000045481_Cluster19758-002 | C3H | zinc finger CCCH domain- | 2098 | 0 | 0 | 0 |
| SALT00000004425_Cluster19988-001 | C3H | zinc finger CCCH type do | 1930 | 0 | 0 | 0 |
| SALT00000015453_Cluster19988-002 | C3H | zinc finger CCCH type do | 1627 | 0 | 0 | 0 |
| SALT00000049856_Cluster19988-003 | C3H | zinc finger CCCH type do | 2030 | 0 | 0 | 0 |
| SALT00000052177_Cluster19988-004 | C3H | zinc finger CCCH type do | 1887 | 0 | 0 | 0 |
| SALT00000058304_Cluster19988-005 | C3H | zinc finger CCCH type do | 2081 | 0 | 0 | 0 |
| SALT00000051694_Cluster19988-006 | C3H | zinc finger CCCH type do | 1774 | 0 | 0 | 0 |
| SALT00000019401_Cluster20597-001 | C3H | zinc finger CCCH type do | 1796 | 0 | 0 | 0 |
| SALT00000040355_Cluster20597-002 | C3H | zinc finger CCCH type do | 1880 | 0 | 0 | 0 |
| SALT00000048433_Cluster20597-003 | C3H | zinc finger CCCH type do | 2034 | 0 | 0 | 0 |
| SALT00000002040_Cluster20602-001 | C3H | [Hordeum vulgare vulgare] | 1825 | 0 | 0 | 0 |
| SALT00000003238_Cluster20602-002 | C3H | [Hordeum vulgare vulgare] | 1526 | down | 0 | down |
| SALT00000007484_Cluster20602-003 | C3H | nucleic acid binding [Zea r | 1747 | 0 | 0 | 0 |
| SALT00000007699_Cluster20602-004 | C3H | Os01g0192000 [Oryza sati | 1794 | 0 | 0 | 0 |
| SALT00000008600_Cluster20602-005 | C3H | nucleic acid binding [Zea r | 1850 | 0 | 0 | 0 |
| SALT00000008887_Cluster20602-006 | C3H | [Hordeum vulgare vulgare] | 2033 | 0 | 0 | 0 |
| SALT00000012242_Cluster20602-007 | C3H | Os01g0192000 [Oryza sati | 1661 | 0 | 0 | 0 |
| SALT00000013306_Cluster20602-008 | C3H | nucleic acid binding [Zea r | 2017 | 0 | 0 | 0 |
| SALT00000018564_Cluster20602-009 | C3H | nucleic acid binding [Zea r | 1803 | 0 | 0 | 0 |
| SALT00000039248_Cluster20602-010 | C3H | nucleic acid binding [Zea r | 2004 | 0 | 0 | 0 |
| SALT00000040556_Cluster20602-011 | C3H | Os01g0192000 [Oryza sati | 1434 | 0 | 0 | 0 |
| SALT00000043165_Cluster20602-012 | C3H | [Hordeum vulgare vulgare] | 1923 | 0 | 0 | 0 |
| SALT00000052174_Cluster20602-013 | C3H | nucleic acid binding [Zea r | 1748 | 0 | 0 | 0 |
| SALT00000054746_Cluster20602-014 | C3H | nucleic acid binding [Zea r | 1999 | 0 | 0 | 0 |
| SALT00000058057_Cluster20602-015 | C3H | Os01g0192000 [Oryza sati | 1601 | 0 | 0 | 0 |
| SALT00000002571_Cluster20752-001 | C3H | nucleic acid binding [Zea r | 1937 | 0 | 0 | 0 |
| SALT00000005028_Cluster20752-002 | C3H | nucleic acid binding [Zea r | 1811 | 0 | 0 | 0 |
| SALT00000006494_Cluster20752-003 | C3H | nucleic acid binding [Zea r | 1887 | 0 | 0 | 0 |
| SALT00000009593_Cluster20752-004 | C3H | nucleic acid binding [Zea r | 1720 | 0 | 0 | 0 |
| SALT00000012148_Cluster20752-005 | C3H | nucleic acid binding [Zea r | 1997 | 0 | 0 | 0 |
| SALT00000035802_Cluster20752-006 | C3H | nucleic acid binding [Zea r | 1906 | 0 | 0 | 0 |
| SALT00000037226_Cluster20752-007 | C3H | nucleic acid binding [Zea r | 1627 | 0 | 0 | 0 |
| SALT00000049832_Cluster20752-008 | C3H | nucleic acid binding [Zea r | 1863 | 0 | 0 | 0 |
| SALT00000051429_Cluster20752-009 | C3H | nucleic acid binding [Zea r | 1668 | 0 | 0 | 0 |
| SALT00000051848_Cluster20752-010 | C3H | nucleic acid binding [Zea r | 2023 | 0 | 0 | 0 |
| SALT00000056143_Cluster20752-011 | C3H | nucleic acid binding [Zea r | 1943 | 0 | 0 | 0 |
| SALT00000056632_Cluster20752-012 | C3H | nucleic acid binding [Zea r | 1773 | 0 | 0 | 0 |
| SALT00000056902_Cluster20752-013 | C3H | nucleic acid binding [Zea r | 1956 | 0 | 0 | 0 |
| SALT00000057686_Cluster20752-014 | C3H | nucleic acid binding [Zea r | 1850 | 0 | 0 | 0 |
| SALT00000059289_Cluster20752-015 | C3H | nucleic acid binding [Zea r | 1908 | 0 | 0 | 0 |
| SALT00000040589_Cluster21025-001 | C3H | Os09g0364000 [Oryza sati | 1926 | 0 | 0 | 0 |
| SALT00000043364_Cluster21025-002 | C3H | Os09g0364000 [Oryza sati | 1864 | 0 | 0 | 0 |
| SALT00000054261_Cluster21025-003 | C3H | Os09g0364000 [Oryza sati | 2003 | 0 | 0 | 0 |
| SALT00000059941_Cluster21025-004 | C3H | zinc finger CCCH domain- | 1310 | 0 | 0 | 0 |
| SALT00000035822_Cluster21264-001 | C3H | Os05g0128200 [Oryza sati | 1987 | 0 | 0 | up |
| SALT00000045978_Cluster21264-002 | C3H | Os05g0128200 [Oryza sati | 1653 | down | 0 | 0 |
| SALT00000052689_Cluster21264-003 | C3H | Os05g0128200 [Oryza sati | 1710 | 0 | 0 | 0 |
| SALT00000067888_Cluster21264-004 | C3H | Os05g0128200 [Oryza sati | 1807 | 0 | 0 | 0 |
| SALT00000007770_Cluster21264-005 | C3H | Os05g0128200 [Oryza sati | 1820 | 0 | 0 | 0 |
| SALT00000015132_Cluster21264-006 | C3H | zinc finger CCCH domain- | 1776 | 0 | 0 | 0 |
| SALT00000000677_Cluster21668-001 | C3H | [Hordeum vulgare vulgare] | 1821 | 0 | 0 | 0 |
| SALT00000000771_Cluster21668-002 | C3H | [Hordeum vulgare vulgare] | 1643 | 0 | 0 | 0 |

| | | | | | | |
|---|---|---|---|---|---|---|
| SALT00000000977_Cluster21668-003 | C3H | [Hordeum vulgare vulgare | 1958 | 0 | 0 | 0 |
| SALT00000003601_Cluster21668-004 | C3H | [Hordeum vulgare vulgare | 1490 | 0 | 0 | 0 |
| SALT00000006013_Cluster21668-005 | C3H | [Hordeum vulgare vulgare | 1686 | 0 | 0 | 0 |
| SALT00000006638_Cluster21668-006 | C3H | [Hordeum vulgare vulgare | 1820 | 0 | 0 | 0 |
| SALT00000006751_Cluster21668-007 | C3H | [Hordeum vulgare vulgare | 1847 | 0 | 0 | 0 |
| SALT00000009300_Cluster21668-008 | C3H | [Hordeum vulgare vulgare | 1720 | 0 | 0 | 0 |
| SALT00000012843_Cluster21668-009 | C3H | [Hordeum vulgare vulgare | 1546 | 0 | 0 | 0 |
| SALT00000013817_Cluster21668-010 | C3H | [Hordeum vulgare vulgare | 1961 | 0 | 0 | 0 |
| SALT00000016580_Cluster21668-011 | C3H | [Hordeum vulgare vulgare | 1541 | 0 | 0 | 0 |
| SALT00000030413_Cluster21668-012 | C3H | [Hordeum vulgare vulgare | 1942 | 0 | 0 | 0 |
| SALT00000034996_Cluster21668-013 | C3H | [Hordeum vulgare vulgare | 1908 | 0 | 0 | 0 |
| SALT00000035085_Cluster21668-014 | C3H | [Hordeum vulgare vulgare | 1936 | 0 | 0 | 0 |
| SALT00000035525_Cluster21668-015 | C3H | [Hordeum vulgare vulgare | 1896 | 0 | 0 | 0 |
| SALT00000035642_Cluster21668-016 | C3H | [Hordeum vulgare vulgare | 1912 | 0 | 0 | 0 |
| SALT00000037620_Cluster21668-017 | C3H | [Hordeum vulgare vulgare | 1665 | down | 0 | down |
| SALT00000043651_Cluster21668-018 | C3H | [Hordeum vulgare vulgare | 1773 | 0 | 0 | 0 |
| SALT00000048716_Cluster21668-019 | C3H | [Hordeum vulgare vulgare | 1678 | 0 | 0 | 0 |
| SALT00000061342_Cluster21668-020 | C3H | [Hordeum vulgare vulgare | 1777 | 0 | 0 | 0 |
| SALT00000061865_Cluster21668-021 | C3H | [Hordeum vulgare vulgare | 1820 | 0 | 0 | 0 |
| SALT00000076692_Cluster21668-022 | C3H | [Hordeum vulgare vulgare | 1941 | 0 | 0 | 0 |
| SALT00000080593_Cluster21668-023 | C3H | [Hordeum vulgare vulgare | 1908 | 0 | 0 | 0 |
| SALT00000084639_Cluster21668-024 | C3H | [Hordeum vulgare vulgare | 1931 | 0 | 0 | 0 |
| SALT00000006991_Cluster21968-001 | C3H | [Hordeum vulgare vulgare | 1941 | down | 0 | 0 |
| SALT00000018405_Cluster21968-002 | C3H | [Hordeum vulgare vulgare | 1838 | 0 | 0 | 0 |
| SALT00000043661_Cluster21968-003 | C3H | [Hordeum vulgare vulgare | 1840 | 0 | 0 | 0 |
| SALT00000048875_Cluster22755-001 | C3H | zinc finger C-x8-C-x5-C-x | 1893 | 0 | 0 | 0 |
| SALT00000004039_Cluster22826-001 | C3H | Os05g0195200 [Oryza sati | 1888 | 0 | up | 0 |
| SALT00000004439_Cluster23275-001 | C3H | zinc finger CCCH type doı | 1860 | 0 | 0 | 0 |
| SALT00000002653_Cluster23531-001 | C3H | zinc finger C-x8-C-x5-C-x | 1764 | 0 | 0 | 0 |
| SALT00000005603_Cluster23531-002 | C3H | zinc finger C-x8-C-x5-C-x | 1808 | 0 | 0 | 0 |
| SALT00000012480_Cluster23531-003 | C3H | zinc finger C-x8-C-x5-C-x | 1844 | 0 | 0 | 0 |
| SALT00000044593_Cluster23531-004 | C3H | zinc finger C-x8-C-x5-C-x | 1663 | 0 | 0 | 0 |
| SALT00000051018_Cluster23531-005 | C3H | zinc finger C-x8-C-x5-C-x | 1813 | 0 | 0 | 0 |
| SALT00000045141_Cluster23531-006 | C3H | zinc finger C-x8-C-x5-C-x | 1714 | 0 | 0 | 0 |
| SALT00000006189_Cluster23623-001 | C3H | zinc finger CCCH type doı | 1839 | 0 | 0 | 0 |
| SALT00000006535_Cluster23623-002 | C3H | zinc finger CCCH type doı | 1587 | 0 | 0 | 0 |
| SALT00000006953_Cluster23623-003 | C3H | zinc finger CCCH type doı | 1527 | 0 | 0 | 0 |
| SALT00000007094_Cluster23623-004 | C3H | zinc finger CCCH type doı | 1817 | 0 | 0 | 0 |
| SALT00000008580_Cluster23623-005 | C3H | zinc finger CCCH type doı | 1507 | 0 | 0 | 0 |
| SALT00000014241_Cluster23623-006 | C3H | zinc finger CCCH type doı | 1681 | 0 | 0 | down |
| SALT00000017623_Cluster23623-007 | C3H | zinc finger CCCH type doı | 1547 | 0 | 0 | 0 |
| SALT00000018028_Cluster23623-008 | C3H | zinc finger CCCH type doı | 1637 | 0 | 0 | 0 |
| SALT00000020077_Cluster23623-009 | C3H | zinc finger CCCH type doı | 1615 | 0 | 0 | 0 |
| SALT00000042500_Cluster23623-010 | C3H | zinc finger CCCH type doı | 1676 | 0 | 0 | 0 |
| SALT00000049553_Cluster23623-011 | C3H | zinc finger CCCH type doı | 1590 | 0 | 0 | 0 |
| SALT00000057662_Cluster23623-012 | C3H | zinc finger CCCH type doı | 1699 | 0 | 0 | 0 |
| SALT00000059447_Cluster23623-013 | C3H | zinc finger CCCH type doı | 1828 | 0 | 0 | 0 |
| SALT00000046942_Cluster23623-014 | C3H | Os01g0917400 [Oryza sati | 1626 | 0 | 0 | 0 |
| SALT00000000854_Cluster24015-001 | C3H | E3 ubiquitin- ligase makor | 1610 | 0 | 0 | 0 |
| SALT00000015523_Cluster24015-002 | C3H | E3 ubiquitin- ligase makor | 1817 | 0 | 0 | 0 |
| SALT00000036877_Cluster24015-003 | C3H | E3 ubiquitin- ligase makor | 1565 | 0 | 0 | 0 |
| SALT00000058168_Cluster24015-004 | C3H | E3 ubiquitin- ligase makor | 1779 | 0 | 0 | 0 |
| SALT00000053498_Cluster24522-001 | C3H | zinc finger CCCH domain- | 1789 | 0 | 0 | 0 |
| SALT00000007336_Cluster24525-001 | C3H | Os01g0616400 [Oryza sati | 1636 | 0 | 0 | 0 |
| SALT00000007739_Cluster24525-002 | C3H | Os01g0616400 [Oryza sati | 1788 | 0 | 0 | 0 |
| SALT00000044430_Cluster24525-003 | C3H | zinc finger CCCH domain- | 1663 | 0 | 0 | 0 |
| SALT00000052550_Cluster24525-004 | C3H | Os01g0616400 [Oryza sati | 1741 | 0 | 0 | 0 |
| SALT00000061891_Cluster24525-005 | C3H | Os01g0616400 [Oryza sati | 1726 | 0 | 0 | 0 |

| | | | | | | |
|---|---|---|---|---|---|---|
| SALT00000012527_Cluster24525-006 | C3H | zinc finger CCCH domain- | 1550 | 0 | 0 | 0 |
| SALT00000057695_Cluster24525-007 | C3H | Os01g0616400 [Oryza sati | 1644 | 0 | 0 | 0 |
| SALT00000063622_Cluster26082-001 | C3H | TPA: MYB DNA-binding | 1707 | 0 | 0 | 0 |
| SALT00000000924_Cluster26381-001 | C3H | zinc finger C-x8-C-x5-C-x | 1549 | 0 | 0 | 0 |
| SALT00000008022_Cluster26381-002 | C3H | zinc finger C-x8-C-x5-C-x | 1634 | 0 | 0 | 0 |
| SALT00000016711_Cluster26381-003 | C3H | zinc finger C-x8-C-x5-C-x | 1509 | 0 | 0 | 0 |
| SALT00000024479_Cluster26381-004 | C3H | zinc finger C-x8-C-x5-C-x | 864 | 0 | 0 | 0 |
| SALT00000047372_Cluster26381-005 | C3H | zinc finger C-x8-C-x5-C-x | 1690 | 0 | 0 | 0 |
| SALT00000047931_Cluster26381-006 | C3H | zinc finger C-x8-C-x5-C-x | 1424 | 0 | 0 | 0 |
| SALT00000051726_Cluster26381-007 | C3H | zinc finger C-x8-C-x5-C-x | 1297 | 0 | 0 | 0 |
| SALT00000063380_Cluster26745-001 | C3H | zinc finger CCCH domain- | 1673 | 0 | 0 | 0 |
| SALT00000002082_Cluster27968-001 | C3H | zinc finger CCCH domain- | 1612 | 0 | 0 | 0 |
| SALT00000012226_Cluster28079-001 | C3H | Os05g0195200 [Oryza sati | 1145 | 0 | 0 | 0 |
| SALT00000013396_Cluster28079-002 | C3H | Os05g0195200 [Oryza sati | 1606 | 0 | 0 | 0 |
| SALT00000023299_Cluster28079-003 | C3H | Os05g0195200 [Oryza sati | 1540 | 0 | 0 | 0 |
| SALT00000055678_Cluster28079-004 | C3H | [Hordeum vulgare vulgare] | 1486 | 0 | 0 | 0 |
| SALT00000016795_Cluster28803-001 | C3H | AGAP010954-PA [Anophe | 1568 | 0 | 0 | 0 |
| SALT00000009418_Cluster29618-001 | C3H | zinc finger C-x8-C-x5-C-x | 1525 | 0 | 0 | 0 |
| SALT00000013328_Cluster29618-002 | C3H | zinc finger C-x8-C-x5-C-x | 1387 | 0 | 0 | 0 |
| SALT00000056652_Cluster30558-001 | C3H | splicing factor U2af 38 kD | 1463 | 0 | 0 | 0 |
| SALT00000025678_Cluster14-001 | CAMTA | calmodulin-binding transcı | 3324 | 0 | 0 | up |
| SALT00000028561_Cluster14-003 | CAMTA | calmodulin-binding transcı | 3438 | 0 | 0 | 0 |
| SALT00000031557_Cluster14-004 | CAMTA | calmodulin-binding transcı | 3467 | 0 | 0 | 0 |
| SALT00000032042_Cluster14-005 | CAMTA | calmodulin-binding transcı | 3318 | 0 | 0 | 0 |
| SALT00000032458_Cluster14-006 | CAMTA | calmodulin-binding transcı | 3471 | 0 | 0 | 0 |
| SALT00000032762_Cluster14-007 | CAMTA | calmodulin-binding transcı | 3432 | 0 | 0 | 0 |
| SALT00000033907_Cluster14-008 | CAMTA | calmodulin-binding transcı | 3566 | 0 | 0 | 0 |
| SALT00000035086_Cluster14-009 | CAMTA | calmodulin-binding transcı | 3467 | 0 | 0 | 0 |
| SALT00000035497_Cluster14-010 | CAMTA | calmodulin-binding transcı | 3354 | 0 | 0 | 0 |
| SALT00000079450_Cluster14-012 | CAMTA | calmodulin-binding transcı | 3620 | 0 | 0 | 0 |
| SALT00000080293_Cluster14-013 | CAMTA | calmodulin-binding transcı | 3449 | 0 | 0 | 0 |
| SALT00000085306_Cluster14-016 | CAMTA | calmodulin-binding transcı | 3578 | 0 | 0 | 0 |
| SALT00000085440_Cluster14-017 | CAMTA | calmodulin-binding transcı | 3468 | 0 | 0 | 0 |
| SALT00000085546_Cluster14-018 | CAMTA | calmodulin-binding transcı | 3473 | 0 | 0 | 0 |
| SALT00000085974_Cluster14-019 | CAMTA | calmodulin-binding transcı | 3551 | 0 | 0 | 0 |
| SALT00000086016_Cluster14-020 | CAMTA | calmodulin-binding transcı | 3602 | 0 | 0 | 0 |
| SALT00000090737_Cluster14-021 | CAMTA | calmodulin-binding transcı | 3364 | 0 | 0 | 0 |
| SALT00000092108_Cluster14-022 | CAMTA | calmodulin-binding transcı | 3458 | 0 | 0 | 0 |
| SALT00000092518_Cluster14-023 | CAMTA | calmodulin-binding transcı | 3458 | 0 | 0 | 0 |
| SALT00000092722_Cluster14-024 | CAMTA | calmodulin-binding transcı | 3588 | 0 | 0 | 0 |
| SALT00000028878_Cluster486-003 | CAMTA | Os10g0375600 [Oryza sati | 3514 | 0 | 0 | 0 |
| SALT00000033817_Cluster486-004 | CAMTA | Os10g0375600 [Oryza sati | 3537 | 0 | 0 | 0 |
| SALT00000087220_Cluster486-006 | CAMTA | Os10g0375600 [Oryza sati | 3564 | 0 | 0 | 0 |
| SALT00000028958_Cluster958-001 | CAMTA | calmodulin-binding transcı | 3519 | 0 | 0 | 0 |
| SALT00000030436_Cluster958-002 | CAMTA | calmodulin-binding transcı | 3594 | 0 | 0 | 0 |
| SALT00000031853_Cluster958-003 | CAMTA | calmodulin-binding transcı | 3609 | 0 | 0 | 0 |
| SALT00000032021_Cluster958-004 | CAMTA | calmodulin-binding transcı | 3570 | 0 | 0 | 0 |
| SALT00000032394_Cluster958-005 | CAMTA | calmodulin-binding transcı | 3578 | 0 | 0 | 0 |
| SALT00000033885_Cluster958-006 | CAMTA | calmodulin-binding transcı | 3652 | 0 | 0 | 0 |
| SALT00000078062_Cluster958-007 | CAMTA | calmodulin-binding transcı | 3653 | 0 | 0 | 0 |
| SALT00000084932_Cluster958-008 | CAMTA | calmodulin-binding transcı | 3625 | 0 | 0 | 0 |
| SALT00000089324_Cluster958-010 | CAMTA | calmodulin-binding transcı | 3645 | 0 | 0 | 0 |
| SALT00000089908_Cluster958-011 | CAMTA | calmodulin-binding transcı | 3672 | 0 | 0 | 0 |
| SALT00000091898_Cluster958-012 | CAMTA | calmodulin-binding transcı | 3550 | 0 | 0 | 0 |
| SALT00000092597_Cluster958-013 | CAMTA | calmodulin-binding transcı | 3660 | 0 | 0 | 0 |
| SALT00000092975_Cluster958-014 | CAMTA | calmodulin-binding transcı | 3576 | 0 | 0 | 0 |
| SALT00000088482_Cluster958-016 | CAMTA | calmodulin-binding transcı | 4322 | 0 | 0 | 0 |
| SALT00000025590_Cluster1680-001 | CAMTA | Calmodulin-binding transc | 3485 | 0 | 0 | 0 |

| | | | | | | |
|---|---|---|---|---|---|---|
| SALT00000030000_Cluster1680-002 | CAMTA | calmodulin-binding transc | 3463 | 0 | 0 | 0 |
| SALT00000031626_Cluster1680-003 | CAMTA | Calmodulin-binding transc | 3336 | 0 | 0 | 0 |
| SALT00000032430_Cluster1680-004 | CAMTA | calmodulin-binding transc | 3648 | 0 | 0 | 0 |
| SALT00000034228_Cluster1680-005 | CAMTA | calmodulin-binding transc | 3476 | 0 | 0 | 0 |
| SALT00000034287_Cluster1680-006 | CAMTA | Calmodulin-binding transc | 3541 | 0 | 0 | 0 |
| SALT00000084727_Cluster1680-008 | CAMTA | calmodulin-binding transc | 3566 | 0 | 0 | 0 |
| SALT00000070705_Cluster1881-002 | CAMTA | calmodulin-binding transc | 2491 | 0 | 0 | 0 |
| SALT00000079095_Cluster1956-002 | CAMTA | ethylene-induced calmodu | 3017 | 0 | 0 | 0 |
| SALT00000021059_Cluster2141-002 | CAMTA | Os07g0490200 [Oryza sati | 3104 | 0 | 0 | 0 |
| SALT00000023245_Cluster2141-003 | CAMTA | Os07g0490200 [Oryza sati | 3055 | 0 | 0 | 0 |
| SALT00000027170_Cluster2141-004 | CAMTA | Os07g0490200 [Oryza sati | 2714 | 0 | 0 | 0 |
| SALT00000030061_Cluster2141-005 | CAMTA | Os07g0490200 [Oryza sati | 3011 | 0 | 0 | 0 |
| SALT00000065567_Cluster2141-007 | CAMTA | Os07g0490200 [Oryza sati | 2876 | 0 | 0 | 0 |
| SALT00000068218_Cluster2141-009 | CAMTA | Os07g0490200 [Oryza sati | 3082 | 0 | 0 | 0 |
| SALT00000080095_Cluster2141-011 | CAMTA | Os07g0490200 [Oryza sati | 2899 | 0 | 0 | 0 |
| SALT00000050046_Cluster2141-012 | CAMTA | Os07g0490200 [Oryza sati | 1904 | 0 | 0 | 0 |
| SALT00000088762_Cluster2264-001 | CAMTA | calmodulin-binding transc | 4981 | 0 | 0 | 0 |
| SALT00000084456_Cluster2779-001 | CAMTA | calmodulin-binding transc | 3759 | 0 | 0 | 0 |
| SALT00000085866_Cluster2779-002 | CAMTA | calmodulin-binding transc | 4703 | 0 | 0 | 0 |
| SALT00000092119_Cluster2779-003 | CAMTA | calmodulin-binding transc | 4036 | 0 | 0 | 0 |
| SALT00000022100_Cluster4614-003 | CAMTA | Os03g0191000 [Oryza sati | 3273 | 0 | 0 | 0 |
| SALT00000028954_Cluster4614-004 | CAMTA | Os03g0191000 [Oryza sati | 3604 | 0 | 0 | 0 |
| SALT00000030682_Cluster4614-005 | CAMTA | Os03g0191000 [Oryza sati | 3356 | 0 | 0 | 0 |
| SALT00000031806_Cluster4614-006 | CAMTA | Os03g0191000 [Oryza sati | 3494 | 0 | 0 | 0 |
| SALT00000031838_Cluster4614-007 | CAMTA | Os03g0191000 [Oryza sati | 3590 | 0 | 0 | 0 |
| SALT00000031969_Cluster4614-008 | CAMTA | Os03g0191000 [Oryza sati | 3560 | 0 | 0 | 0 |
| SALT00000032240_Cluster4614-009 | CAMTA | Os03g0191000 [Oryza sati | 3573 | 0 | 0 | 0 |
| SALT00000033137_Cluster4614-010 | CAMTA | Os03g0191000 [Oryza sati | 3523 | 0 | 0 | 0 |
| SALT00000033634_Cluster4614-011 | CAMTA | Os03g0191000 [Oryza sati | 3638 | 0 | 0 | 0 |
| SALT00000033978_Cluster4614-012 | CAMTA | Os03g0191000 [Oryza sati | 3440 | 0 | 0 | 0 |
| SALT00000034527_Cluster4614-013 | CAMTA | Os03g0191000 [Oryza sati | 3587 | 0 | 0 | 0 |
| SALT00000064022_Cluster4614-019 | CAMTA | Os03g0191000 [Oryza sati | 2828 | 0 | 0 | 0 |
| SALT00000077199_Cluster4614-020 | CAMTA | Os03g0191000 [Oryza sati | 3532 | 0 | 0 | 0 |
| SALT00000084098_Cluster4614-023 | CAMTA | Os03g0191000 [Oryza sati | 3579 | 0 | 0 | 0 |
| SALT00000084994_Cluster4614-024 | CAMTA | Os03g0191000 [Oryza sati | 3488 | 0 | 0 | 0 |
| SALT00000090207_Cluster4614-025 | CAMTA | Os03g0191000 [Oryza sati | 3536 | 0 | 0 | 0 |
| SALT00000090502_Cluster4614-026 | CAMTA | Os03g0191000 [Oryza sati | 3533 | 0 | 0 | 0 |
| SALT00000091954_Cluster4614-027 | CAMTA | Os03g0191000 [Oryza sati | 3381 | 0 | 0 | 0 |
| SALT00000092635_Cluster4614-028 | CAMTA | Os03g0191000 [Oryza sati | 3563 | 0 | 0 | 0 |
| SALT00000092672_Cluster4614-029 | CAMTA | Os03g0191000 [Oryza sati | 3914 | 0 | 0 | 0 |
| SALT00000092037_Cluster4614-031 | CAMTA | Os03g0191000 [Oryza sati | 4055 | 0 | 0 | 0 |
| SALT00000086623_Cluster4614-032 | CAMTA | Os03g0191000 [Oryza sati | 3667 | 0 | 0 | 0 |
| SALT00000079782_Cluster4614-033 | CAMTA | Os03g0191000 [Oryza sati | 4032 | 0 | 0 | 0 |
| SALT00000085428_Cluster4614-034 | CAMTA | Os03g0191000 [Oryza sati | 3541 | 0 | 0 | 0 |
| SALT00000043201_Cluster4614-035 | CAMTA | Os03g0191000 [Oryza sati | 1696 | 0 | 0 | 0 |
| SALT00000032516_Cluster5664-001 | CAMTA | calmodulin-binding transc | 3708 | 0 | 0 | 0 |
| SALT00000082019_Cluster5664-002 | CAMTA | calmodulin-binding transc | 3600 | 0 | 0 | 0 |
| SALT00000086796_Cluster5664-003 | CAMTA | calmodulin-binding transc | 3850 | 0 | 0 | 0 |
| SALT00000033440_Cluster5664-005 | CAMTA | calmodulin-binding transc | 3593 | 0 | up | 0 |
| SALT00000033474_Cluster6596-001 | CAMTA | calmodulin-binding transc | 3417 | 0 | 0 | 0 |
| SALT00000092740_Cluster6596-002 | CAMTA | calmodulin-binding transc | 3699 | 0 | 0 | 0 |
| SALT00000068987_Cluster6834-001 | CAMTA | Os07g0490200 [Oryza sati | 3008 | 0 | 0 | 0 |
| SALT00000024211_Cluster7270-001 | CAMTA | ethylene-induced calmodu | 2937 | 0 | 0 | 0 |
| SALT00000026287_Cluster7270-002 | CAMTA | ethylene-induced calmodu | 2891 | 0 | 0 | 0 |
| SALT00000081142_Cluster7270-003 | CAMTA | ethylene-induced calmodu | 3598 | 0 | 0 | 0 |
| SALT00000082001_Cluster7270-004 | CAMTA | ethylene-induced calmodu | 3019 | 0 | 0 | 0 |
| SALT00000089341_Cluster7270-005 | CAMTA | calmodulin-binding transc | 3924 | 0 | 0 | 0 |
| SALT00000021061_Cluster7534-001 | CAMTA | Os07g0490200 [Oryza sati | 3147 | 0 | 0 | 0 |

| | | | | | | |
|---|---|---|---|---|---|---|
| SALT00000021235_Cluster7534-002 | CAMTA | Os07g0490200 [Oryza sati | 3116 | 0 | 0 | 0 |
| SALT00000026176_Cluster7534-003 | CAMTA | Os07g0490200 [Oryza sati | 2556 | 0 | 0 | 0 |
| SALT00000026690_Cluster7534-004 | CAMTA | Os07g0490200 [Oryza sati | 2789 | 0 | 0 | 0 |
| SALT00000029876_Cluster7534-005 | CAMTA | Os07g0490200 [Oryza sati | 3556 | 0 | 0 | 0 |
| SALT00000024636_Cluster7952-001 | CAMTA | calmodulin-binding transcı | 3373 | 0 | 0 | 0 |
| SALT00000030018_Cluster7952-002 | CAMTA | calmodulin-binding transcı | 3489 | 0 | 0 | 0 |
| SALT00000034327_Cluster7952-003 | CAMTA | calmodulin-binding transcı | 3085 | 0 | 0 | 0 |
| SALT00000034659_Cluster7952-004 | CAMTA | calmodulin-binding transcı | 3403 | 0 | 0 | 0 |
| SALT00000006515_Cluster9354-001 | CO-like | zinc finger CONSTANS-L | 1715 | 0 | 0 | 0 |
| SALT00000001213_Cluster12906-001 | CO-like | zinc finger -like [Oryza sat | 1677 | 0 | 0 | 0 |
| SALT00000009834_Cluster14919-001 | CO-like | zinc finger CONSTANS-L | 1950 | 0 | 0 | 0 |
| SALT00000015428_Cluster14919-002 | CO-like | zinc finger CONSTANS-L | 2014 | 0 | 0 | 0 |
| SALT00000017977_Cluster14919-003 | CO-like | zinc finger CONSTANS-L | 2115 | 0 | 0 | 0 |
| SALT00000085425_Cluster14919-005 | CO-like | zinc finger CONSTANS-L | 2044 | 0 | 0 | 0 |
| SALT00000019226_Cluster14919-006 | CO-like | zinc finger CONSTANS-L | 2029 | 0 | 0 | 0 |
| SALT00000042817_Cluster18606-001 | CO-like | zinc finger CONSTANS-L | 2083 | 0 | 0 | 0 |
| SALT00000044012_Cluster18606-002 | CO-like | zinc finger CONSTANS-L | 2027 | 0 | 0 | 0 |
| SALT00000060377_Cluster18606-003 | CO-like | zinc finger CONSTANS-L | 2207 | 0 | 0 | 0 |
| SALT00000063753_Cluster18606-004 | CO-like | zinc finger CONSTANS-L | 2141 | 0 | 0 | 0 |
| SALT00000073179_Cluster18606-005 | CO-like | zinc finger CONSTANS-L | 2189 | 0 | 0 | 0 |
| SALT00000004824_Cluster20857-001 | CO-like | zinc finger CONSTANS-L | 1783 | 0 | 0 | 0 |
| SALT00000025072_Cluster20857-002 | CO-like | zinc finger CONSTANS-L | 1760 | 0 | 0 | 0 |
| SALT00000035663_Cluster20857-003 | CO-like | zinc finger CONSTANS-L | 1663 | 0 | 0 | 0 |
| SALT00000053201_Cluster20857-005 | CO-like | zinc finger CONSTANS-L | 1750 | 0 | 0 | 0 |
| SALT00000058460_Cluster20857-006 | CO-like | zinc finger CONSTANS-L | 1638 | 0 | 0 | down |
| SALT00000041526_Cluster20857-008 | CO-like | C2C2-CO-like transcriptio | 1654 | 0 | 0 | 0 |
| SALT00000010184_Cluster21095-001 | CO-like | Os06g0264200 [Oryza sati | 1641 | 0 | 0 | 0 |
| SALT00000014332_Cluster21095-002 | CO-like | Os06g0264200 [Oryza sati | 1998 | 0 | 0 | 0 |
| SALT00000002956_Cluster23393-001 | CO-like | CCT motif family [Zea ma | 1769 | 0 | 0 | 0 |
| SALT00000006541_Cluster23393-002 | CO-like | CCT motif family [Zea ma | 1816 | 0 | 0 | down |
| SALT00000006904_Cluster23393-003 | CO-like | CCT motif family [Zea ma | 1708 | 0 | 0 | 0 |
| SALT00000012436_Cluster23393-004 | CO-like | CCT motif family [Zea ma | 1579 | 0 | 0 | 0 |
| SALT00000013620_Cluster23393-005 | CO-like | CCT motif family [Zea ma | 1853 | up | 0 | up |
| SALT00000035005_Cluster23393-006 | CO-like | CCT motif family [Zea ma | 1747 | 0 | 0 | down |
| SALT00000035422_Cluster23393-007 | CO-like | CCT motif family [Zea ma | 1724 | 0 | 0 | 0 |
| SALT00000045369_Cluster23393-008 | CO-like | CCT motif family [Zea ma | 1828 | 0 | 0 | 0 |
| SALT00000054322_Cluster23393-009 | CO-like | CCT motif family [Zea ma | 1534 | 0 | 0 | up |
| SALT00000069539_Cluster23393-010 | CO-like | CCT motif family [Zea ma | 1691 | 0 | 0 | 0 |
| SALT00000001675_Cluster23489-001 | CO-like | Os06g0264200 [Oryza sati | 1682 | 0 | 0 | down |
| SALT00000004202_Cluster23489-002 | CO-like | Os06g0264200 [Oryza sati | 1847 | 0 | 0 | 0 |
| SALT00000007701_Cluster23489-003 | CO-like | Os06g0264200 [Oryza sati | 1825 | 0 | 0 | 0 |
| SALT00000021927_Cluster23489-004 | CO-like | Os06g0264200 [Oryza sati | 1604 | 0 | 0 | 0 |
| SALT00000047827_Cluster23489-005 | CO-like | Os06g0264200 [Oryza sati | 1674 | 0 | 0 | 0 |
| SALT00000050279_Cluster23489-006 | CO-like | Os06g0264200 [Oryza sati | 1747 | 0 | 0 | 0 |
| SALT00000065571_Cluster23489-007 | CO-like | Os06g0264200 [Oryza sati | 1796 | 0 | 0 | 0 |
| SALT00000061576_Cluster23489-008 | CO-like | Os06g0264200 [Oryza sati | 1827 | 0 | 0 | 0 |
| SALT00000001463_Cluster24524-001 | CO-like | zinc finger CONSTANS-L | 1788 | 0 | 0 | 0 |
| SALT00000047239_Cluster24524-002 | CO-like | zinc finger CONSTANS-L | 1610 | 0 | 0 | 0 |
| SALT00000037094_Cluster28371-001 | CO-like | CONSTANS CO6 [Zea ma | 1523 | 0 | 0 | 0 |
| SALT00000040524_Cluster28371-002 | CO-like | CONSTANS CO6 [Zea ma | 1591 | 0 | 0 | 0 |
| SALT00000044450_Cluster28371-004 | CO-like | CONSTANS CO6 [Zea ma | 1479 | up | 0 | 0 |
| SALT00000069953_Cluster28371-005 | CO-like | CONSTANS CO6 [Zea ma | 1421 | 0 | 0 | 0 |
| SALT00000050420_Cluster28371-006 | CO-like | mitochondrial phosphate tı | 2703 | 0 | up | 0 |
| SALT00000010369_Cluster30364-001 | CO-like | CONSTANS CO6 [Zea ma | 1300 | 0 | 0 | down |
| SALT00000041923_Cluster30364-002 | CO-like | CONSTANS CO6 [Zea ma | 1477 | 0 | 0 | 0 |
| SALT00000048984_Cluster30429-001 | CO-like | CONSTANS CO6 [Zea ma | 1473 | 0 | 0 | 0 |
| SALT00000050697_Cluster30429-002 | CO-like | CONSTANS CO6 [Zea ma | 1456 | 0 | 0 | 0 |
| SALT00000055393_Cluster30429-003 | CO-like | CONSTANS CO6 [Zea ma | 1445 | 0 | 0 | 0 |

| | | | | | | |
|---|---|---|---|---|---|---|
| SALT00000003088_Cluster31773-001 | CO-like | Os04g0497700 [Oryza sati | 1259 | 0 | 0 | 0 |
| SALT00000015383_Cluster1610-001 | CPP | tesmin TSO1-like CXC 2 i | 1714 | 0 | 0 | 0 |
| SALT00000019242_Cluster1610-002 | CPP | tesmin TSO1-like CXC 2 | 2159 | 0 | 0 | 0 |
| SALT00000021071_Cluster1610-003 | CPP | tesmin TSO1-like CXC 2 | 2931 | 0 | 0 | 0 |
| SALT00000021598_Cluster1610-004 | CPP | tesmin TSO1-like CXC 2 i | 2786 | 0 | 0 | 0 |
| SALT00000023408_Cluster1610-005 | CPP | tesmin TSO1-like CXC 2 i | 2846 | 0 | 0 | 0 |
| SALT00000025192_Cluster1610-006 | CPP | tesmin TSO1-like CXC 2 | 2908 | 0 | 0 | 0 |
| SALT00000025240_Cluster1610-007 | CPP | tesmin TSO1-like CXC 2 | 2767 | 0 | 0 | 0 |
| SALT00000026075_Cluster1610-008 | CPP | tesmin TSO1-like CXC 2 | 2795 | 0 | 0 | 0 |
| SALT00000028213_Cluster1610-009 | CPP | tesmin TSO1-like CXC 2 i | 2885 | 0 | 0 | 0 |
| SALT00000031485_Cluster1610-010 | CPP | tesmin TSO1-like CXC 2 i | 2765 | 0 | 0 | 0 |
| SALT00000033123_Cluster1610-011 | CPP | tesmin TSO1-like CXC 2 | 2748 | 0 | 0 | 0 |
| SALT00000035052_Cluster1610-012 | CPP | tesmin TSO1-like CXC 2 | 2805 | 0 | 0 | 0 |
| SALT00000039684_Cluster1610-013 | CPP | tesmin TSO1-like CXC 2 i | 1756 | 0 | 0 | 0 |
| SALT00000057338_Cluster1610-014 | CPP | tesmin TSO1-like CXC 2 i | 1635 | 0 | 0 | 0 |
| SALT00000071211_Cluster1610-015 | CPP | tesmin TSO1-like CXC 2 | 2820 | 0 | 0 | 0 |
| SALT00000075315_Cluster1610-016 | CPP | tesmin TSO1-like CXC 2 | 2913 | 0 | 0 | 0 |
| SALT00000076150_Cluster1610-017 | CPP | tesmin TSO1-like CXC 2 | 2905 | 0 | 0 | 0 |
| SALT00000081903_Cluster1610-020 | CPP | tesmin TSO1-like CXC 2 i | 2859 | 0 | 0 | 0 |
| SALT00000093884_Cluster1610-021 | CPP | tesmin TSO1-like CXC 2 | 2749 | 0 | 0 | 0 |
| SALT00000023010_Cluster1745-001 | CPP | tesmin TSO1-like CXC 2 | 2826 | 0 | 0 | 0 |
| SALT00000028397_Cluster1745-002 | CPP | tesmin TSO1-like CXC 2 i | 2773 | 0 | 0 | 0 |
| SALT00000080876_Cluster1745-004 | CPP | tesmin TSO1-like CXC 2 i | 4832 | 0 | 0 | down |
| SALT00000073647_Cluster1753-002 | CPP | tesmin TSO1-like CXC 2 i | 2754 | 0 | 0 | 0 |
| SALT00000081660_Cluster6425-002 | CPP | pentatricopeptide repeat-cc | 3724 | 0 | 0 | 0 |
| SALT00000038759_Cluster9499-001 | CPP | tesmin TSO1-like CXC 7 i | 1677 | 0 | 0 | 0 |
| SALT00000050059_Cluster9499-002 | CPP | tesmin TSO1-like CXC 7 i | 1574 | 0 | 0 | 0 |
| SALT00000018334_Cluster15416-001 | CPP | Os07g0176200 [Oryza sati | 1017 | 0 | 0 | 0 |
| SALT00000071399_Cluster15416-002 | CPP | tesmin TSO1-like CXC 3 i | 2596 | 0 | 0 | 0 |
| SALT00000026433_Cluster15435-001 | CPP | Os07g0176200 [Oryza sati | 2531 | 0 | 0 | 0 |
| SALT00000083620_Cluster15435-002 | CPP | Os07g0176200 [Oryza sati | 2595 | 0 | 0 | down |
| SALT00000063031_Cluster15998-014 | CPP | PREDICTED: uncharacter | 2552 | 0 | 0 | 0 |
| SALT00000080477_Cluster15998-016 | CPP | PREDICTED: uncharacter | 2522 | 0 | 0 | 0 |
| SALT00000044142_Cluster18447-001 | CPP | tesmin TSO1-like CXC 5 [ | 2122 | 0 | 0 | 0 |
| SALT00000046868_Cluster18447-002 | CPP | tesmin TSO1-like CXC 5 [ | 2070 | 0 | 0 | 0 |
| SALT00000050220_Cluster18447-003 | CPP | tesmin TSO1-like CXC 5 [ | 2226 | 0 | 0 | 0 |
| SALT00000064391_Cluster18447-004 | CPP | tesmin TSO1-like CXC 5 [ | 1917 | 0 | 0 | 0 |
| SALT00000051624_Cluster21205-001 | CPP | tesmin TSO1-like CXC 7 i | 1991 | 0 | 0 | 0 |
| SALT00000001398_Cluster30034-001 | CPP | Os12g0605300 [Oryza sati | 1499 | 0 | 0 | 0 |
| SALT00000010635_Cluster30034-002 | CPP | Os12g0605300 [Oryza sati | 1463 | 0 | 0 | 0 |
| SALT00000036864_Cluster30313-001 | CPP | CXC domain containing T | 1250 | 0 | 0 | 0 |
| SALT00000051870_Cluster30313-002 | CPP | CXC domain containing T | 1481 | 0 | 0 | 0 |
| SALT00000012663_Cluster30313-003 | CPP | CXC domain containing T | 1608 | 0 | 0 | 0 |
| SALT00000007176_Cluster475-001 | DBB | zinc-finger R2931 [Oryza : | 1528 | 0 | 0 | 0 |
| SALT00000008047_Cluster475-002 | DBB | zinc-finger R2931 [Oryza : | 1560 | 0 | 0 | 0 |
| SALT00000011686_Cluster475-003 | DBB | zinc-finger R2931 [Oryza : | 1528 | 0 | 0 | 0 |
| SALT00000028494_Cluster475-004 | DBB | zinc-finger R2931 [Oryza : | 1492 | 0 | 0 | 0 |
| SALT00000031605_Cluster475-006 | DBB | zinc-finger R2931 [Oryza : | 1467 | 0 | 0 | 0 |
| SALT00000034859_Cluster475-008 | DBB | zinc-finger R2931 [Oryza : | 1435 | 0 | 0 | 0 |
| SALT00000043848_Cluster475-010 | DBB | zinc-finger R2931 [Oryza : | 1438 | 0 | 0 | 0 |
| SALT00000047947_Cluster475-019 | DBB | orphans transcription partia | 1541 | 0 | 0 | 0 |
| SALT00000070848_Cluster11111-001 | DBB | zinc finger CONSTANS-L | 2958 | 0 | 0 | 0 |
| SALT00000000378_Cluster26748-001 | DBB | zinc-finger R2931 [Oryza : | 1620 | 0 | 0 | 0 |
| SALT00000005203_Cluster26748-002 | DBB | zinc-finger R2931 [Oryza : | 1647 | 0 | 0 | 0 |
| SALT00000005513_Cluster26748-003 | DBB | zinc-finger R2931 [Oryza : | 1513 | 0 | 0 | 0 |
| SALT00000012589_Cluster26748-004 | DBB | zinc-finger R2931 [Oryza : | 1577 | 0 | 0 | 0 |
| SALT00000014786_Cluster26748-005 | DBB | zinc-finger R2931 [Oryza : | 1467 | 0 | 0 | 0 |
| SALT00000050983_Cluster26748-006 | DBB | zinc-finger R2931 [Oryza : | 1560 | 0 | 0 | 0 |

| | | | | | | |
|---|---|---|---|---|---|---|
| SALT00000093845_Cluster26748-007 | DBB | zinc-finger R2931 [Oryza s | 1673 | 0 | 0 | 0 |
| SALT00000007257_Cluster30875-001 | DBB | salt tolerance [Zea mays] | 1106 | 0 | 0 | 0 |
| SALT00000010087_Cluster30875-002 | DBB | salt tolerance [Zea mays] | 1433 | 0 | 0 | 0 |
| SALT00000037450_Cluster30875-003 | DBB | zinc finger [Oryza sativa Ja | 1249 | 0 | 0 | 0 |
| SALT00000015206_Cluster30875-004 | DBB | zinc finger [Oryza sativa Ja | 1197 | 0 | 0 | down |
| SALT00000061767_Cluster31356-001 | DBB | Os04g0540200 [Oryza sati | 1365 | 0 | 0 | 0 |
| SALT00000017469_Cluster32170-001 | DBB | zinc finger | 1128 | 0 | 0 | 0 |
| SALT00000000201_Cluster9231-001 | Dof | Os07g0685000 [Oryza sati | 1758 | 0 | 0 | 0 |
| SALT00000001840_Cluster9231-002 | Dof | Os07g0685000 [Oryza sati | 1683 | 0 | 0 | 0 |
| SALT00000002775_Cluster9231-003 | Dof | Os07g0685000 [Oryza sati | 1662 | 0 | 0 | 0 |
| SALT00000016901_Cluster9231-004 | Dof | Os07g0685000 [Oryza sati | 1694 | 0 | 0 | 0 |
| SALT00000017171_Cluster9231-005 | Dof | Os07g0685000 [Oryza sati | 1533 | 0 | 0 | 0 |
| SALT00000060102_Cluster9231-006 | Dof | Os07g0685000 [Oryza sati | 1704 | 0 | 0 | 0 |
| SALT00000060779_Cluster9231-007 | Dof | Os07g0685000 [Oryza sati | 1757 | 0 | 0 | 0 |
| SALT00000080995_Cluster9231-008 | Dof | Os07g0685000 [Oryza sati | 3245 | 0 | 0 | 0 |
| SALT00000052575_Cluster9231-009 | Dof | Os07g0685000 [Oryza sati | 1637 | 0 | 0 | 0 |
| SALT00000042730_Cluster9231-010 | Dof | Os07g0685000 [Oryza sati | 1562 | 0 | 0 | 0 |
| SALT00000070446_Cluster10781-001 | Dof | TMV-MP30 binding 2C [Z | 3000 | 0 | 0 | 0 |
| SALT00000075428_Cluster11795-001 | Dof | dof zinc finger PBF-like | 2886 | 0 | 0 | 0 |
| SALT00000000926_Cluster15554-001 | Dof | cyclic dof factor 1-like [Se | 1654 | 0 | 0 | up |
| SALT00000001082_Cluster15554-002 | Dof | cyclic dof factor 1-like [Se | 1673 | 0 | 0 | 0 |
| SALT00000002481_Cluster15554-003 | Dof | cyclic dof factor 1-like [Se | 1818 | 0 | 0 | 0 |
| SALT00000003728_Cluster15554-004 | Dof | cyclic dof factor 1-like [Se | 1683 | 0 | 0 | 0 |
| SALT00000004253_Cluster15554-005 | Dof | cyclic dof factor 1-like [Se | 1659 | 0 | 0 | up |
| SALT00000004981_Cluster15554-006 | Dof | cyclic dof factor 1-like [Se | 1723 | 0 | 0 | 0 |
| SALT00000012160_Cluster15554-007 | Dof | cyclic dof factor 1-like [Se | 1768 | 0 | 0 | 0 |
| SALT00000014579_Cluster15554-008 | Dof | cyclic dof factor 1-like [Se | 1590 | 0 | 0 | 0 |
| SALT00000020337_Cluster15554-009 | Dof | cyclic dof factor 1-like [Se | 1696 | 0 | 0 | 0 |
| SALT00000040788_Cluster15554-011 | Dof | cyclic dof factor 1-like [Se | 1692 | 0 | 0 | 0 |
| SALT00000047489_Cluster15554-012 | Dof | cyclic dof factor 1-like [Se | 1831 | 0 | 0 | 0 |
| SALT00000047506_Cluster15554-013 | Dof | cyclic dof factor 1-like [Se | 1584 | 0 | 0 | 0 |
| SALT00000048290_Cluster15554-014 | Dof | cyclic dof factor 1-like [Se | 1676 | 0 | 0 | 0 |
| SALT00000058879_Cluster15554-015 | Dof | cyclic dof factor 1-like [Se | 1761 | 0 | 0 | 0 |
| SALT00000071225_Cluster15554-016 | Dof | cyclic dof factor 1-like [Se | 2546 | 0 | 0 | 0 |
| SALT00000078459_Cluster15554-017 | Dof | cyclic dof factor 1-like [Se | 2585 | 0 | 0 | 0 |
| SALT00000004388_Cluster19902-001 | Dof | dof zinc finger MNB1A-lil | 2087 | 0 | 0 | 0 |
| SALT00000052844_Cluster20015-001 | Dof | Os03g0821200 [Oryza sati | 1570 | 0 | 0 | 0 |
| SALT00000061554_Cluster20015-002 | Dof | Os03g0821200 [Oryza sati | 2079 | 0 | 0 | 0 |
| SALT00000040463_Cluster22032-001 | Dof | dof zinc finger [Oryza sativ | 1742 | 0 | 0 | 0 |
| SALT00000056099_Cluster22032-002 | Dof | dof zinc finger [Oryza sativ | 1937 | 0 | 0 | 0 |
| SALT00000006268_Cluster22072-001 | Dof | cyclic dof factor 2-like | 1934 | 0 | 0 | 0 |
| SALT00000003321_Cluster23111-001 | Dof | dof zinc finger [Zea mays] | 1613 | 0 | 0 | 0 |
| SALT00000014058_Cluster23111-002 | Dof | dof zinc finger [Zea mays] | 1594 | 0 | 0 | 0 |
| SALT00000044241_Cluster23111-003 | Dof | dof zinc finger [Zea mays] | 1540 | 0 | 0 | 0 |
| SALT00000047876_Cluster23111-004 | Dof | dof zinc finger [Zea mays] | 1870 | 0 | 0 | 0 |
| SALT00000067603_Cluster23111-005 | Dof | dof zinc finger [Zea mays] | 1625 | 0 | 0 | 0 |
| SALT00000007477_Cluster23909-001 | Dof | dof zinc finger [Oryza sativ | 1823 | 0 | 0 | 0 |
| SALT00000044195_Cluster24374-001 | Dof | dof zinc finger -like [Zea m | 1798 | 0 | 0 | 0 |
| SALT00000045193_Cluster24442-001 | Dof | cyclic dof factor 1-like [Se | 1743 | 0 | 0 | 0 |
| SALT00000057411_Cluster24442-002 | Dof | cyclic dof factor 1-like [Se | 1794 | 0 | 0 | 0 |
| SALT00000043970_Cluster24605-001 | Dof | dof zinc finger -like | 1784 | 0 | 0 | 0 |
| SALT00000014632_Cluster24610-001 | Dof | dof zinc finger -like | 1516 | 0 | 0 | 0 |
| SALT00000046088_Cluster24610-002 | Dof | dof zinc finger -like | 1494 | 0 | 0 | 0 |
| SALT00000046407_Cluster24610-003 | Dof | dof zinc finger -like | 1618 | 0 | 0 | 0 |
| SALT00000053906_Cluster24610-004 | Dof | dof zinc finger -like | 1505 | 0 | 0 | 0 |
| SALT00000055674_Cluster24610-005 | Dof | dof zinc finger -like | 1784 | 0 | 0 | 0 |
| SALT00000053595_Cluster24983-001 | Dof | dof zinc finger isoform X1 | 1764 | 0 | 0 | 0 |
| SALT00000056077_Cluster24983-002 | Dof | dof zinc finger isoform X1 | 1680 | 0 | 0 | 0 |

| | | | | | | |
|---|---|---|---|---|---|---|
| SALT00000049533_Cluster24983-003 | Dof | dof zinc finger isoform X2 | 1759 | 0 | 0 | 0 |
| SALT00000009135_Cluster25274-001 | Dof | TPA_inf: Dof-type zinc fin | 1748 | 0 | 0 | 0 |
| SALT00000054631_Cluster25274-002 | Dof | TPA_inf: Dof-type zinc fin | 1520 | 0 | 0 | 0 |
| SALT00000043209_Cluster25881-001 | Dof | cyclic dof factor 1-like [Se | 1682 | 0 | 0 | 0 |
| SALT00000049791_Cluster25881-002 | Dof | cyclic dof factor 1-like [Se | 1718 | 0 | 0 | 0 |
| SALT00000044427_Cluster26264-001 | Dof | cyclic dof factor 1-like [Se | 1696 | 0 | 0 | 0 |
| SALT00000056955_Cluster26264-002 | Dof | cyclic dof factor 1-like [Se | 1606 | 0 | 0 | 0 |
| SALT00000037177_Cluster27097-001 | Dof | Os03g0821200 [Oryza sati | 1655 | 0 | 0 | 0 |
| SALT00000061055_Cluster27097-002 | Dof | Os03g0821200 [Oryza sati | 1587 | 0 | 0 | 0 |
| SALT00000036337_Cluster28587-001 | Dof | cyclic dof factor 1-like [Se | 1579 | 0 | 0 | 0 |
| SALT00000046122_Cluster28587-002 | Dof | cyclic dof factor 1-like [Se | 1520 | 0 | 0 | 0 |
| SALT00000061199_Cluster28587-003 | Dof | cyclic dof factor 1-like [Se | 1514 | 0 | 0 | 0 |
| SALT00000013812_Cluster28626-001 | Dof | DOF29 transcription partia | 1577 | 0 | 0 | 0 |
| SALT00000023518_Cluster29438-001 | Dof | Os10g0496000 [Oryza sati | 1536 | 0 | 0 | 0 |
| SALT00000054859_Cluster29788-001 | Dof | Os05g0112200 [Oryza sati | 1515 | 0 | 0 | 0 |
| SALT00000002273_Cluster31401-001 | Dof | dof zinc finger MNB1A [Z | 1242 | 0 | 0 | 0 |
| SALT00000005730_Cluster31401-002 | Dof | dof zinc finger MNB1A [Z | 1354 | 0 | 0 | 0 |
| SALT00000015479_Cluster31401-003 | Dof | dof zinc finger MNB1A [Z | 1140 | 0 | 0 | 0 |
| SALT00000061162_Cluster31583-001 | Dof | DNA-binding Dof3 [Oryza | 1312 | 0 | 0 | 0 |
| SALT00000054808_Cluster31705-001 | Dof | dof zinc finger -like | 1278 | 0 | 0 | 0 |
| SALT00000061124_Cluster8281-001 | E2F/DP | transcription factor E2F3 [ | 1996 | 0 | 0 | 0 |
| SALT00000088255_Cluster8281-002 | E2F/DP | transcription factor E2F3 [ | 3428 | 0 | 0 | 0 |
| SALT00000047975_Cluster8281-003 | E2F/DP | transcription factor E2F3 [ | 2183 | 0 | 0 | 0 |
| SALT00000081443_Cluster12365-001 | E2F/DP | E2F-DP transcription parti | 2833 | 0 | 0 | 0 |
| SALT00000082799_Cluster16496-001 | E2F/DP | transcription factor E2F3 [ | 2516 | 0 | 0 | 0 |
| SALT00000062423_Cluster20421-001 | E2F/DP | E2F-related [Zea mays] | 2046 | 0 | 0 | 0 |
| SALT00000001802_Cluster21856-001 | E2F/DP | transcription factor E2F2 [ | 1895 | 0 | 0 | 0 |
| SALT00000036506_Cluster21856-002 | E2F/DP | transcription factor E2F2 [ | 1819 | 0 | 0 | 0 |
| SALT00000041071_Cluster21856-003 | E2F/DP | transcription factor E2F2 [ | 1908 | 0 | 0 | 0 |
| SALT00000044643_Cluster21856-004 | E2F/DP | transcription factor E2F2 [ | 1949 | 0 | 0 | 0 |
| SALT00000057847_Cluster21856-005 | E2F/DP | transcription factor E2F2 [ | 1560 | 0 | 0 | 0 |
| SALT00000047056_Cluster27934-001 | E2F/DP | [Oryza sativa Japonica Gro | 1614 | 0 | 0 | 0 |
| SALT00000040906_Cluster28115-001 | E2F/DP | LOC100283679 isoform X | 1569 | 0 | 0 | 0 |
| SALT00000045184_Cluster28115-002 | E2F/DP | LOC100283679 isoform X | 1604 | 0 | 0 | 0 |
| SALT00000061237_Cluster28115-003 | E2F/DP | LOC100283679 isoform X | 1476 | 0 | 0 | 0 |
| SALT00000006511_Cluster28379-001 | E2F/DP | E2F transcription factor-lik | 1548 | 0 | 0 | 0 |
| SALT00000007955_Cluster28379-002 | E2F/DP | E2F transcription factor-lik | 1590 | 0 | 0 | 0 |
| SALT00000037277_Cluster29134-001 | E2F/DP | Os10g0440100 [Oryza sati | 1552 | 0 | 0 | 0 |
| SALT00000003864_Cluster1479-001 | EIL | ETHYLENE-INSENSITIV | 1631 | 0 | 0 | down |
| SALT00000005844_Cluster1479-002 | EIL | ETHYLENE-INSENSITIV | 1635 | 0 | 0 | 0 |
| SALT00000039587_Cluster1479-004 | EIL | ETHYLENE-INSENSITIV | 1817 | 0 | 0 | up |
| SALT00000042965_Cluster1479-005 | EIL | ETHYLENE-INSENSITIV | 1279 | 0 | 0 | 0 |
| SALT00000064149_Cluster1479-006 | EIL | ETHYLENE-INSENSITIV | 1017 | 0 | 0 | 0 |
| SALT00000011695_Cluster4554-001 | EIL | ETHYLENE-INSENSITIV | 2184 | 0 | 0 | 0 |
| SALT00000024757_Cluster4554-002 | EIL | ETHYLENE-INSENSITIV | 2598 | 0 | 0 | 0 |
| SALT00000026003_Cluster4554-003 | EIL | ETHYLENE-INSENSITIV | 2717 | 0 | 0 | 0 |
| SALT00000028294_Cluster4554-005 | EIL | ETHYLENE-INSENSITIV | 2655 | 0 | 0 | 0 |
| SALT00000069148_Cluster4554-006 | EIL | ETHYLENE-INSENSITIV | 2672 | 0 | 0 | 0 |
| SALT00000071776_Cluster4554-008 | EIL | ETHYLENE-INSENSITIV | 4073 | 0 | 0 | 0 |
| SALT00000002753_Cluster7441-001 | EIL | ETHYLENE-INSENSITIV | 1924 | 0 | 0 | 0 |
| SALT00000003429_Cluster7441-002 | EIL | ETHYLENE-INSENSITIV | 1839 | 0 | 0 | 0 |
| SALT00000008420_Cluster7441-003 | EIL | ETHYLENE-INSENSITIV | 1578 | 0 | 0 | 0 |
| SALT00000011014_Cluster7441-004 | EIL | ETHYLENE-INSENSITIV | 1816 | 0 | 0 | 0 |
| SALT00000011421_Cluster7441-005 | EIL | ETHYLENE-INSENSITIV | 2325 | 0 | 0 | 0 |
| SALT00000011863_Cluster7441-006 | EIL | ETHYLENE-INSENSITIV | 2270 | 0 | 0 | 0 |
| SALT00000012411_Cluster7441-007 | EIL | ETHYLENE-INSENSITIV | 1717 | 0 | 0 | 0 |
| SALT00000012576_Cluster7441-008 | EIL | ETHYLENE-INSENSITIV | 1716 | 0 | 0 | 0 |
| SALT00000012739_Cluster7441-009 | EIL | ETHYLENE-INSENSITIV | 1714 | 0 | 0 | 0 |

| | | | | | | |
|---|---|---|---|---|---|---|
| SALT00000018843_Cluster7441-010 | EIL | ETHYLENE-INSENSITIV | 2054 | 0 | 0 | 0 |
| SALT00000019352_Cluster7441-011 | EIL | ETHYLENE-INSENSITIV | 2146 | 0 | 0 | 0 |
| SALT00000020848_Cluster7441-012 | EIL | ETHYLENE-INSENSITIV | 2476 | 0 | 0 | 0 |
| SALT00000022658_Cluster7441-013 | EIL | ETHYLENE-INSENSITIV | 2485 | 0 | 0 | 0 |
| SALT00000024265_Cluster7441-014 | EIL | ETHYLENE-INSENSITIV | 3035 | 0 | 0 | 0 |
| SALT00000024968_Cluster7441-015 | EIL | ETHYLENE-INSENSITIV | 2790 | 0 | 0 | 0 |
| SALT00000026879_Cluster7441-016 | EIL | ETHYLENE-INSENSITIV | 2369 | 0 | 0 | 0 |
| SALT00000028905_Cluster7441-017 | EIL | ETHYLENE-INSENSITIV | 3449 | 0 | 0 | 0 |
| SALT00000039882_Cluster7441-018 | EIL | ETHYLENE-INSENSITIV | 1993 | 0 | 0 | up |
| SALT00000040432_Cluster7441-019 | EIL | ETHYLENE-INSENSITIV | 1769 | 0 | 0 | 0 |
| SALT00000043903_Cluster7441-020 | EIL | ETHYLENE-INSENSITIV | 1913 | 0 | 0 | 0 |
| SALT00000066025_Cluster7441-021 | EIL | ETHYLENE-INSENSITIV | 2602 | 0 | 0 | 0 |
| SALT00000069551_Cluster7441-022 | EIL | ETHYLENE-INSENSITIV | 2388 | 0 | 0 | 0 |
| SALT00000070572_Cluster7441-023 | EIL | ETHYLENE-INSENSITIV | 2399 | 0 | 0 | 0 |
| SALT00000077311_Cluster7441-024 | EIL | ETHYLENE-INSENSITIV | 2743 | 0 | 0 | 0 |
| SALT00000081518_Cluster7441-025 | EIL | ETHYLENE-INSENSITIV | 2481 | 0 | 0 | 0 |
| SALT00000093637_Cluster7441-026 | EIL | ETHYLENE-INSENSITIV | 3569 | 0 | 0 | 0 |
| SALT00000020023_Cluster7441-027 | EIL | ETHYLENE-INSENSITIV | 2333 | 0 | 0 | up |
| SALT00000053846_Cluster7441-028 | EIL | ETHYLENE-INSENSITIV | 2069 | down | 0 | 0 |
| SALT00000006151_Cluster7992-001 | EIL | ETHYLENE-INSENSITIV | 1842 | 0 | 0 | 0 |
| SALT00000021204_Cluster7992-002 | EIL | ETHYLENE-INSENSITIV | 2438 | 0 | 0 | 0 |
| SALT00000021578_Cluster7992-003 | EIL | ETHYLENE-INSENSITIV | 2471 | up | up | up |
| SALT00000023444_Cluster7992-004 | EIL | ETHYLENE-INSENSITIV | 2394 | 0 | 0 | 0 |
| SALT00000026189_Cluster7992-005 | EIL | ETHYLENE-INSENSITIV | 2507 | 0 | 0 | 0 |
| SALT00000029432_Cluster7992-006 | EIL | ETHYLENE-INSENSITIV | 2598 | 0 | 0 | 0 |
| SALT00000031055_Cluster7992-007 | EIL | ETHYLENE-INSENSITIV | 3483 | 0 | 0 | 0 |
| SALT00000041193_Cluster7992-008 | EIL | ETHYLENE-INSENSITIV | 1614 | 0 | 0 | 0 |
| SALT00000083067_Cluster7992-009 | EIL | ETHYLENE-INSENSITIV | 2414 | 0 | 0 | 0 |
| SALT00000010366_Cluster13136-001 | EIL | Os09g0490200 [Oryza sati | 2360 | 0 | 0 | 0 |
| SALT00000025550_Cluster13136-002 | EIL | Os09g0490200 [Oryza sati | 2535 | 0 | 0 | 0 |
| SALT00000047928_Cluster13136-003 | EIL | ETHYLENE INSENSITIV | 1598 | 0 | 0 | 0 |
| SALT00000052134_Cluster13136-004 | EIL | Os09g0490200 [Oryza sati | 2097 | 0 | 0 | 0 |
| SALT00000066300_Cluster13136-005 | EIL | Os09g0490200 [Oryza sati | 2634 | 0 | 0 | 0 |
| SALT00000067979_Cluster13136-006 | EIL | Os09g0490200 [Oryza sati | 2763 | 0 | 0 | 0 |
| SALT00000024752_Cluster16256-001 | EIL | Os09g0490200 [Oryza sati | 2319 | 0 | 0 | 0 |
| SALT00000070904_Cluster16256-002 | EIL | Os09g0490200 [Oryza sati | 2533 | 0 | 0 | 0 |
| SALT00000072354_Cluster16256-003 | EIL | Os09g0490200 [Oryza sati | 2481 | 0 | 0 | 0 |
| SALT00000025746_Cluster17129-001 | EIL | ETHYLENE INSENSITIV | 2457 | 0 | 0 | 0 |
| SALT00000067600_Cluster87-001 | ERF | DRE binding factor 1 [Zea | 11622 | up | up | up |
| SALT00000005050_Cluster629-001 | ERF | EREBP [Oryza sativa Japo | 1524 | 0 | 0 | 0 |
| SALT00000009927_Cluster629-002 | ERF | EREBP [Oryza sativa Japo | 1508 | 0 | 0 | 0 |
| SALT00000020416_Cluster629-003 | ERF | EREBP [Oryza sativa Japo | 1554 | 0 | 0 | 0 |
| SALT00000020507_Cluster629-004 | ERF | EREBP [Oryza sativa Japo | 1588 | 0 | 0 | 0 |
| SALT00000025180_Cluster629-005 | ERF | EREBP [Oryza sativa Japo | 1598 | 0 | 0 | 0 |
| SALT00000036024_Cluster629-006 | ERF | EREBP [Oryza sativa Japo | 7524 | 0 | 0 | 0 |
| SALT00000043373_Cluster629-007 | ERF | EREBP [Oryza sativa Japo | 1558 | 0 | 0 | 0 |
| SALT00000045058_Cluster629-008 | ERF | EREBP [Oryza sativa Japo | 1410 | 0 | 0 | 0 |
| SALT00000059487_Cluster629-010 | ERF | EREBP [Oryza sativa Japo | 1555 | 0 | 0 | 0 |
| SALT00000003816_Cluster1577-001 | ERF | EREBP transcription facto | 1510 | 0 | 0 | 0 |
| SALT00000006694_Cluster1577-002 | ERF | EREBP transcription facto | 1602 | 0 | 0 | 0 |
| SALT00000017210_Cluster1577-003 | ERF | EREBP transcription facto | 1625 | 0 | 0 | 0 |
| SALT00000018205_Cluster1577-004 | ERF | EREBP transcription facto | 1566 | 0 | 0 | 0 |
| SALT00000020323_Cluster1577-005 | ERF | EREBP transcription facto | 1454 | 0 | 0 | 0 |
| SALT00000020550_Cluster1577-006 | ERF | EREBP transcription facto | 1646 | 0 | 0 | 0 |
| SALT00000023900_Cluster1577-007 | ERF | EREBP transcription facto | 1470 | 0 | 0 | 0 |
| SALT00000025847_Cluster1577-008 | ERF | EREBP transcription facto | 1604 | 0 | 0 | 0 |
| SALT00000037250_Cluster1577-009 | ERF | EREBP transcription facto | 1603 | 0 | 0 | 0 |
| SALT00000039436_Cluster1577-011 | ERF | EREBP transcription facto | 1719 | 0 | 0 | 0 |

| | | | | | | |
|---|---|---|---|---|---|---|
| SALT00000049312_Cluster1577-012 | ERF | EREBP transcription facto | 1616 | 0 | 0 | 0 |
| SALT00000053847_Cluster1577-013 | ERF | EREBP transcription facto | 1693 | 0 | 0 | 0 |
| SALT00000054965_Cluster1577-014 | ERF | EREBP transcription facto | 1725 | 0 | 0 | 0 |
| SALT00000055814_Cluster1577-015 | ERF | EREBP transcription facto | 1718 | 0 | 0 | 0 |
| SALT00000058989_Cluster1577-016 | ERF | EREBP transcription facto | 1576 | 0 | 0 | 0 |
| SALT00000061504_Cluster1577-017 | ERF | EREBP transcription facto | 1572 | 0 | 0 | 0 |
| SALT00000003171_Cluster4236-006 | ERF | EREBP [Oryza sativa Japo | 1426 | 0 | 0 | 0 |
| SALT00000007361_Cluster4236-010 | ERF | EREBP [Oryza sativa Japo | 1497 | 0 | 0 | 0 |
| SALT00000007574_Cluster4236-011 | ERF | EREBP [Oryza sativa Japo | 1454 | 0 | 0 | 0 |
| SALT00000008083_Cluster4236-013 | ERF | EREBP [Oryza sativa Japo | 1567 | 0 | 0 | 0 |
| SALT00000009154_Cluster4236-014 | ERF | EREBP [Oryza sativa Japo | 1611 | 0 | 0 | 0 |
| SALT00000010313_Cluster4236-015 | ERF | EREBP [Oryza sativa Japo | 1521 | 0 | 0 | 0 |
| SALT00000011227_Cluster4236-016 | ERF | [Hordeum vulgare vulgare | 1580 | 0 | 0 | 0 |
| SALT00000012223_Cluster4236-017 | ERF | EREBP [Oryza sativa Japo | 1437 | 0 | 0 | 0 |
| SALT00000013010_Cluster4236-018 | ERF | EREBP [Oryza sativa Japo | 1322 | 0 | 0 | 0 |
| SALT00000013393_Cluster4236-019 | ERF | EREBP [Oryza sativa Japo | 1574 | 0 | 0 | 0 |
| SALT00000014350_Cluster4236-020 | ERF | EREBP [Oryza sativa Japo | 1630 | 0 | 0 | 0 |
| SALT00000015233_Cluster4236-021 | ERF | EREBP [Oryza sativa Japo | 1520 | 0 | 0 | 0 |
| SALT00000017565_Cluster4236-023 | ERF | EREBP [Oryza sativa Japo | 1737 | 0 | 0 | 0 |
| SALT00000019863_Cluster4236-024 | ERF | EREBP [Oryza sativa Japo | 1547 | 0 | 0 | 0 |
| SALT00000020312_Cluster4236-025 | ERF | EREBP [Oryza sativa Japo | 1367 | 0 | 0 | 0 |
| SALT00000023517_Cluster4236-026 | ERF | EREBP [Oryza sativa Japo | 1454 | 0 | 0 | 0 |
| SALT00000023883_Cluster4236-027 | ERF | EREBP [Oryza sativa Japo | 1257 | 0 | 0 | 0 |
| SALT00000025104_Cluster4236-028 | ERF | EREBP [Oryza sativa Japo | 1700 | 0 | 0 | 0 |
| SALT00000025131_Cluster4236-029 | ERF | EREBP [Oryza sativa Japo | 1640 | 0 | 0 | 0 |
| SALT00000025132_Cluster4236-030 | ERF | EREBP [Oryza sativa Japo | 1557 | 0 | 0 | 0 |
| SALT00000031662_Cluster4236-035 | ERF | EREBP [Oryza sativa Japo | 1525 | 0 | 0 | 0 |
| SALT00000034668_Cluster4236-036 | ERF | EREBP [Oryza sativa Japo | 1497 | 0 | 0 | 0 |
| SALT00000034811_Cluster4236-037 | ERF | EREBP [Oryza sativa Japo | 1565 | 0 | 0 | 0 |
| SALT00000034909_Cluster4236-038 | ERF | EREBP [Oryza sativa Japo | 1476 | 0 | 0 | 0 |
| SALT00000035212_Cluster4236-039 | ERF | EREBP [Oryza sativa Japo | 1521 | 0 | 0 | 0 |
| SALT00000035357_Cluster4236-040 | ERF | EREBP [Oryza sativa Japo | 1515 | 0 | 0 | 0 |
| SALT00000035547_Cluster4236-041 | ERF | EREBP [Oryza sativa Japo | 1445 | 0 | 0 | 0 |
| SALT00000035744_Cluster4236-042 | ERF | EREBP [Oryza sativa Japo | 1578 | 0 | 0 | 0 |
| SALT00000035918_Cluster4236-043 | ERF | EREBP [Oryza sativa Japo | 1616 | 0 | 0 | 0 |
| SALT00000036735_Cluster4236-045 | ERF | EREBP [Oryza sativa Japo | 1633 | 0 | 0 | 0 |
| SALT00000036844_Cluster4236-046 | ERF | EREBP [Oryza sativa Japo | 1620 | 0 | 0 | 0 |
| SALT00000036912_Cluster4236-047 | ERF | EREBP [Oryza sativa Japo | 1579 | 0 | 0 | 0 |
| SALT00000037816_Cluster4236-048 | ERF | EREBP [Oryza sativa Japo | 1533 | 0 | 0 | 0 |
| SALT00000037872_Cluster4236-049 | ERF | EREBP [Oryza sativa Japo | 1506 | 0 | 0 | 0 |
| SALT00000037973_Cluster4236-050 | ERF | Tubulin alpha-3 chain | 4155 | 0 | down | down |
| SALT00000038605_Cluster4236-051 | ERF | EREBP [Oryza sativa Japo | 1593 | 0 | 0 | 0 |
| SALT00000038726_Cluster4236-052 | ERF | EREBP [Oryza sativa Japo | 1567 | 0 | down | down |
| SALT00000039979_Cluster4236-055 | ERF | EREBP [Oryza sativa Japo | 2031 | 0 | 0 | 0 |
| SALT00000040603_Cluster4236-056 | ERF | EREBP [Oryza sativa Japo | 1515 | 0 | 0 | 0 |
| SALT00000041939_Cluster4236-057 | ERF | EREBP [Oryza sativa Japo | 1596 | 0 | 0 | 0 |
| SALT00000042175_Cluster4236-058 | ERF | EREBP [Oryza sativa Japo | 1554 | 0 | 0 | 0 |
| SALT00000043015_Cluster4236-059 | ERF | EREBP [Oryza sativa Japo | 1614 | 0 | 0 | 0 |
| SALT00000043627_Cluster4236-060 | ERF | EREBP [Oryza sativa Japo | 1529 | 0 | 0 | 0 |
| SALT00000045627_Cluster4236-061 | ERF | EREBP [Oryza sativa Japo | 1614 | 0 | 0 | 0 |
| SALT00000045974_Cluster4236-063 | ERF | EREBP [Oryza sativa Japo | 1536 | 0 | 0 | 0 |
| SALT00000046132_Cluster4236-064 | ERF | EREBP [Oryza sativa Japo | 1598 | 0 | 0 | 0 |
| SALT00000046855_Cluster4236-065 | ERF | EREBP [Oryza sativa Japo | 1648 | 0 | 0 | 0 |
| SALT00000047788_Cluster4236-066 | ERF | EREBP [Oryza sativa Japo | 1574 | 0 | 0 | 0 |
| SALT00000048784_Cluster4236-067 | ERF | EREBP [Oryza sativa Japo | 1536 | 0 | 0 | 0 |
| SALT00000049140_Cluster4236-068 | ERF | EREBP [Oryza sativa Japo | 1606 | 0 | 0 | 0 |
| SALT00000050027_Cluster4236-069 | ERF | EREBP [Oryza sativa Japo | 1588 | 0 | 0 | 0 |
| SALT00000051179_Cluster4236-070 | ERF | EREBP [Oryza sativa Japo | 1575 | 0 | 0 | 0 |

| | | | | | | |
|---|---|---|---|---|---|---|
| SALT00000051441_Cluster4236-071 | ERF | EREBP [Oryza sativa Japo | 1569 | 0 | 0 | 0 |
| SALT00000051733_Cluster4236-072 | ERF | EREBP [Oryza sativa Japo | 1643 | 0 | up | up |
| SALT00000052563_Cluster4236-073 | ERF | EREBP [Oryza sativa Japo | 1596 | 0 | 0 | 0 |
| SALT00000052742_Cluster4236-074 | ERF | AP2 EREBP partial | 1461 | 0 | 0 | 0 |
| SALT00000052834_Cluster4236-075 | ERF | EREBP [Oryza sativa Japo | 1599 | 0 | 0 | 0 |
| SALT00000053994_Cluster4236-076 | ERF | EREBP [Oryza sativa Japo | 1607 | 0 | 0 | 0 |
| SALT00000054622_Cluster4236-077 | ERF | EREBP [Oryza sativa Japo | 1604 | 0 | 0 | 0 |
| SALT00000056569_Cluster4236-078 | ERF | EREBP [Oryza sativa Japo | 1436 | 0 | 0 | 0 |
| SALT00000057189_Cluster4236-079 | ERF | EREBP [Oryza sativa Japo | 1570 | 0 | 0 | 0 |
| SALT00000057514_Cluster4236-080 | ERF | EREBP [Oryza sativa Japo | 1419 | 0 | 0 | 0 |
| SALT00000057522_Cluster4236-081 | ERF | EREBP [Oryza sativa Japo | 1532 | 0 | 0 | 0 |
| SALT00000060565_Cluster4236-082 | ERF | EREBP [Oryza sativa Japo | 1604 | 0 | 0 | 0 |
| SALT00000061694_Cluster4236-084 | ERF | EREBP [Oryza sativa Japo | 1528 | 0 | 0 | 0 |
| SALT00000066871_Cluster4236-086 | ERF | EREBP [Oryza sativa Japo | 1642 | 0 | 0 | 0 |
| SALT00000066963_Cluster4236-087 | ERF | EREBP [Oryza sativa Japo | 1509 | 0 | 0 | 0 |
| SALT00000071326_Cluster4236-088 | ERF | EREBP [Oryza sativa Japo | 1631 | 0 | 0 | 0 |
| SALT00000071931_Cluster4236-089 | ERF | EREBP [Oryza sativa Japo | 1570 | 0 | 0 | 0 |
| SALT00000053941_Cluster4236-090 | ERF | EREBP [Oryza sativa Japo | 2263 | 0 | up | 0 |
| SALT00000057869_Cluster4236-093 | ERF | EREBP [Oryza sativa Japo | 1922 | 0 | 0 | 0 |
| SALT00000060460_Cluster4236-094 | ERF | fructose-bisphosphate aldo | 2621 | 0 | 0 | 0 |
| SALT00000008402_Cluster4299-006 | ERF | EREBP [Oryza sativa Japo | 1311 | 0 | 0 | 0 |
| SALT00000015812_Cluster4299-011 | ERF | AP2 EREBP partial | 1034 | 0 | 0 | 0 |
| SALT00000023347_Cluster4299-016 | ERF | ethylene-responsive transc | 1485 | 0 | 0 | 0 |
| SALT00000045158_Cluster4299-025 | ERF | EREBP [Oryza sativa Japo | 1415 | 0 | 0 | 0 |
| SALT00000047926_Cluster4299-028 | ERF | EREBP [Oryza sativa Japo | 1623 | 0 | 0 | 0 |
| SALT00000054596_Cluster4299-031 | ERF | serine mitochondrial [Setar | 4140 | 0 | 0 | down |
| SALT00000082495_Cluster7361-001 | ERF | DREB 1 [Zoysia japonica] | 3453 | 0 | 0 | 0 |
| SALT00000084917_Cluster7361-002 | ERF | Os01g0165100 [Oryza sati | 3582 | up | 0 | up |
| SALT00000087001_Cluster7361-003 | ERF | Os01g0165100 [Oryza sati | 3544 | up | up | 0 |
| SALT00000078197_Cluster7361-004 | ERF | Os01g0165100 [Oryza sati | 3209 | up | up | up |
| SALT00000016748_Cluster7627-003 | ERF | AP2-EREBP transcription | 1626 | 0 | 0 | up |
| SALT00000037681_Cluster7627-007 | ERF | AP2-EREBP transcription | 1626 | 0 | 0 | 0 |
| SALT00000048901_Cluster7627-008 | ERF | AP2-EREBP transcription | 1522 | 0 | 0 | up |
| SALT00000055547_Cluster7627-009 | ERF | AP2-EREBP transcription | 1325 | 0 | 0 | up |
| SALT00000090829_Cluster7627-010 | ERF | glycine--tRNA mitochondr | 3541 | 0 | 0 | up |
| SALT00000008441_Cluster7627-011 | ERF | AP2-EREBP transcription | 1976 | 0 | 0 | 0 |
| SALT00000061810_Cluster7627-013 | ERF | AP2-EREBP transcription | 1614 | 0 | 0 | 0 |
| SALT00000085405_Cluster8631-001 | ERF | ethylene-responsive transc | 3358 | 0 | 0 | 0 |
| SALT00000001318_Cluster8733-001 | ERF | EREBP [Oryza sativa Japo | 1650 | 0 | 0 | 0 |
| SALT00000003582_Cluster8733-002 | ERF | EREBP [Oryza sativa Japo | 1483 | 0 | 0 | 0 |
| SALT00000005840_Cluster8733-003 | ERF | EREBP [Oryza sativa Japo | 1535 | 0 | 0 | 0 |
| SALT00000018302_Cluster8733-004 | ERF | EREBP [Oryza sativa Japo | 1447 | 0 | 0 | 0 |
| SALT00000020422_Cluster8733-005 | ERF | EREBP [Oryza sativa Japo | 1486 | 0 | 0 | 0 |
| SALT00000025186_Cluster8733-006 | ERF | EREBP [Oryza sativa Japo | 1617 | 0 | 0 | 0 |
| SALT00000035279_Cluster8733-007 | ERF | EREBP [Oryza sativa Japo | 1562 | 0 | 0 | 0 |
| SALT00000035367_Cluster8733-008 | ERF | EREBP [Oryza sativa Japo | 1565 | 0 | 0 | 0 |
| SALT00000035600_Cluster8733-009 | ERF | EREBP [Oryza sativa Japo | 1563 | 0 | 0 | 0 |
| SALT00000037480_Cluster8733-011 | ERF | EREBP [Oryza sativa Japo | 1535 | 0 | 0 | 0 |
| SALT00000038764_Cluster8733-012 | ERF | peroxisomal membrane PE | 3340 | 0 | 0 | 0 |
| SALT00000041914_Cluster8733-013 | ERF | EREBP [Oryza sativa Japo | 1477 | 0 | 0 | 0 |
| SALT00000042715_Cluster8733-014 | ERF | EREBP [Oryza sativa Japo | 1571 | 0 | 0 | down |
| SALT00000045598_Cluster8733-015 | ERF | EREBP [Oryza sativa Japo | 1502 | 0 | 0 | 0 |
| SALT00000051882_Cluster8733-016 | ERF | EREBP [Oryza sativa Japo | 1476 | 0 | 0 | 0 |
| SALT00000053412_Cluster8733-017 | ERF | EREBP [Oryza sativa Japo | 1483 | 0 | 0 | 0 |
| SALT00000055804_Cluster8733-018 | ERF | EREBP [Oryza sativa Japo | 1536 | 0 | 0 | 0 |
| SALT00000093720_Cluster8733-019 | ERF | EREBP [Oryza sativa Japo | 1557 | 0 | 0 | 0 |
| SALT00000061201_Cluster8733-020 | ERF | EREBP [Oryza sativa Japo | 1615 | 0 | 0 | 0 |
| SALT00000059821_Cluster8733-021 | ERF | EREBP [Oryza sativa Japo | 2146 | 0 | 0 | 0 |

| | | | | | | |
|---|---|---|---|---|---|---|
| SALT00000037120_Cluster8733-022 | ERF | Ribulose bisphosphate carl | 2426 | 0 | 0 | up |
| SALT00000006164_Cluster9190-001 | ERF | dehydration responsive ele | 1421 | 0 | 0 | 0 |
| SALT00000013894_Cluster9190-002 | ERF | dehydration responsive ele | 1471 | 0 | 0 | 0 |
| SALT00000033145_Cluster9190-003 | ERF | dehydration responsive ele | 3253 | 0 | 0 | 0 |
| SALT00000041387_Cluster9190-004 | ERF | dehydration responsive ele | 1521 | 0 | 0 | 0 |
| SALT00000078276_Cluster9190-005 | ERF | dehydration responsive ele | 2468 | 0 | 0 | 0 |
| SALT00000087086_Cluster9467-001 | ERF | 7 [Hordeum vulgare vulgai | 3198 | 0 | 0 | up |
| SALT00000037554_Cluster11439-001 | ERF | dehydration responsive ele | 1364 | 0 | 0 | 0 |
| SALT00000047186_Cluster11439-002 | ERF | dehydration responsive ele | 1405 | 0 | 0 | 0 |
| SALT00000054194_Cluster11439-003 | ERF | dehydration responsive ele | 1390 | 0 | 0 | 0 |
| SALT00000056661_Cluster11439-004 | ERF | dehydration responsive fac | 1542 | 0 | 0 | 0 |
| SALT00000058703_Cluster11439-005 | ERF | dehydration responsive ele | 2180 | 0 | 0 | 0 |
| SALT00000061324_Cluster11439-006 | ERF | dehydration responsive ele | 2153 | 0 | 0 | 0 |
| SALT00000061403_Cluster11439-007 | ERF | dehydration responsive ele | 1437 | 0 | 0 | 0 |
| SALT00000065162_Cluster11439-008 | ERF | dehydration responsive ele | 2284 | 0 | 0 | 0 |
| SALT00000082096_Cluster11439-009 | ERF | dehydration responsive ele | 2921 | 0 | 0 | 0 |
| SALT00000050185_Cluster12141-006 | ERF | glyceraldehyde-3-phospha | 2854 | 0 | 0 | 0 |
| SALT00000021738_Cluster13637-001 | ERF | ethylene response element | 1362 | 0 | 0 | 0 |
| SALT00000035512_Cluster13637-002 | ERF | ethylene response element | 1426 | 0 | 0 | 0 |
| SALT00000044102_Cluster13637-003 | ERF | ethylene response element | 1413 | 0 | 0 | 0 |
| SALT00000046314_Cluster13637-004 | ERF | ethylene response element | 1414 | 0 | 0 | 0 |
| SALT00000062975_Cluster13637-005 | ERF | ethylene response element | 2722 | 0 | 0 | 0 |
| SALT00000067510_Cluster13637-006 | ERF | ethylene response element | 2662 | 0 | 0 | 0 |
| SALT00000049523_Cluster16477-001 | ERF | EREBP [Oryza sativa Japo | 2517 | 0 | 0 | 0 |
| SALT00000073887_Cluster16760-001 | ERF | EREBP [Oryza sativa Japo | 2494 | 0 | 0 | 0 |
| SALT00000038454_Cluster19508-001 | ERF | ethylene-responsive transc | 2119 | 0 | 0 | 0 |
| SALT00000012510_Cluster19521-001 | ERF | ethylene-responsive transc | 2118 | 0 | 0 | 0 |
| SALT00000017421_Cluster19521-002 | ERF | ethylene-responsive transc | 2062 | 0 | 0 | 0 |
| SALT00000044376_Cluster19521-003 | ERF | ethylene-responsive transc | 1718 | 0 | 0 | up |
| SALT00000027345_Cluster19521-004 | ERF | ethylene-responsive transc | 2078 | 0 | 0 | 0 |
| SALT00000009337_Cluster20172-001 | ERF | AP2 EREBP transcription | 1942 | 0 | 0 | 0 |
| SALT00000014110_Cluster20172-002 | ERF | AP2 EREBP transcription | 2065 | 0 | 0 | 0 |
| SALT00000042312_Cluster20172-003 | ERF | AP2 EREBP transcription | 1812 | 0 | 0 | 0 |
| SALT00000002224_Cluster20172-004 | ERF | AP2 EREBP transcription | 1894 | 0 | 0 | 0 |
| SALT00000012422_Cluster20172-005 | ERF | AP2 EREBP transcription | 1835 | 0 | 0 | 0 |
| SALT00000000672_Cluster20354-001 | ERF | ethylene response element | 1419 | 0 | 0 | 0 |
| SALT00000004571_Cluster20354-002 | ERF | ethylene response element | 1506 | 0 | 0 | 0 |
| SALT00000004817_Cluster20354-003 | ERF | ethylene response element | 2050 | 0 | 0 | 0 |
| SALT00000014945_Cluster20354-004 | ERF | ethylene response element | 1660 | 0 | 0 | 0 |
| SALT00000019919_Cluster20354-005 | ERF | ethylene response element | 1492 | 0 | 0 | 0 |
| SALT00000035267_Cluster20354-006 | ERF | ethylene response element | 1486 | 0 | 0 | 0 |
| SALT00000035645_Cluster20354-007 | ERF | ethylene response element | 1459 | 0 | 0 | up |
| SALT00000037391_Cluster20354-008 | ERF | ethylene response element | 1834 | 0 | 0 | 0 |
| SALT00000038647_Cluster20354-009 | ERF | ethylene response element | 1560 | 0 | 0 | 0 |
| SALT00000045084_Cluster20354-010 | ERF | ethylene response element | 1678 | 0 | 0 | 0 |
| SALT00000051161_Cluster20354-011 | ERF | ethylene response element | 1766 | 0 | 0 | 0 |
| SALT00000055105_Cluster20354-012 | ERF | ethylene response element | 1708 | 0 | 0 | 0 |
| SALT00000059425_Cluster20354-013 | ERF | ethylene response element | 1510 | 0 | 0 | 0 |
| SALT00000000082_Cluster20659-001 | ERF | AP2-EREBP transcription | 1884 | 0 | 0 | down |
| SALT00000002195_Cluster20659-002 | ERF | AP2-EREBP transcription | 1813 | 0 | 0 | 0 |
| SALT00000013225_Cluster20659-003 | ERF | AP2-EREBP transcription | 1761 | 0 | 0 | 0 |
| SALT00000014496_Cluster20659-004 | ERF | AP2-EREBP transcription | 1567 | 0 | 0 | down |
| SALT00000037332_Cluster20659-005 | ERF | AP2-EREBP transcription | 2029 | 0 | 0 | 0 |
| SALT00000052732_Cluster20659-006 | ERF | AP2-EREBP transcription | 1877 | 0 | 0 | 0 |
| SALT00000054798_Cluster20659-007 | ERF | AP2-EREBP transcription | 1887 | 0 | 0 | 0 |
| SALT00000058210_Cluster20659-008 | ERF | ethylene-responsive transc | 1632 | 0 | 0 | 0 |
| SALT00000050000_Cluster20659-009 | ERF | AP2-EREBP transcription | 1932 | 0 | 0 | 0 |
| SALT00000048411_Cluster20828-001 | ERF | ethylene-responsive transc | 2017 | 0 | 0 | 0 |

| | | | | | | |
|---|---|---|---|---|---|---|
| SALT00000011310_Cluster21485-001 | ERF | AP2 domain containing [Z | 1819 | 0 | 0 | 0 |
| SALT00000018940_Cluster21485-002 | ERF | AP2 domain containing [Z | 1971 | 0 | 0 | 0 |
| SALT00000016245_Cluster22099-001 | ERF | AP2 domain containing [Z | 1932 | 0 | 0 | 0 |
| SALT00000053544_Cluster22718-001 | ERF | ethylene-responsive transc | 1840 | 0 | 0 | 0 |
| SALT00000061122_Cluster22718-002 | ERF | ethylene-responsive transc | 1896 | 0 | 0 | 0 |
| SALT00000000377_Cluster22777-001 | ERF | DRE binding factor 2 [Ory | 1778 | 0 | 0 | 0 |
| SALT00000003224_Cluster22777-002 | ERF | DRE binding factor 2 [Ory | 1730 | 0 | 0 | 0 |
| SALT00000009678_Cluster22777-003 | ERF | DRE binding factor 2 [Ory | 1701 | 0 | 0 | down |
| SALT00000015464_Cluster22777-004 | ERF | DRE binding factor 2 [Ory | 1849 | 0 | 0 | 0 |
| SALT00000046760_Cluster22777-005 | ERF | DRE binding factor 2 [Ory | 1778 | 0 | 0 | 0 |
| SALT00000049073_Cluster22777-006 | ERF | DRE binding factor 2 [Ory | 1892 | 0 | 0 | 0 |
| SALT00000001375_Cluster23150-001 | ERF | AP2 EREBP transcription | 1862 | 0 | 0 | 0 |
| SALT00000011489_Cluster23150-002 | ERF | AP2 EREBP transcription | 1729 | 0 | 0 | 0 |
| SALT00000045189_Cluster23150-003 | ERF | AP2 EREBP transcription | 1868 | 0 | 0 | 0 |
| SALT00000007552_Cluster23381-001 | ERF | pathogenesis-related transc | 1730 | 0 | 0 | 0 |
| SALT00000010162_Cluster23381-002 | ERF | pathogenesis-related transc | 1308 | 0 | 0 | 0 |
| SALT00000014734_Cluster23381-003 | ERF | pathogenesis-related transc | 1854 | 0 | 0 | 0 |
| SALT00000018397_Cluster23381-004 | ERF | pathogenesis-related transc | 1835 | 0 | 0 | 0 |
| SALT00000059810_Cluster23381-005 | ERF | pathogenesis-related transc | 1726 | 0 | down | down |
| SALT00000060499_Cluster23582-001 | ERF | Os01g0657400 [Oryza sati | 1842 | 0 | 0 | 0 |
| SALT00000001188_Cluster23635-001 | ERF | Os02g0752800 [Oryza sati | 1838 | 0 | 0 | 0 |
| SALT00000006798_Cluster23635-002 | ERF | AP2 EREBP transcription | 1816 | 0 | 0 | 0 |
| SALT00000007462_Cluster23635-003 | ERF | Os02g0752800 [Oryza sati | 1627 | 0 | 0 | 0 |
| SALT00000010640_Cluster23635-004 | ERF | AP2 EREBP transcription | 1549 | 0 | 0 | up |
| SALT00000017056_Cluster23635-005 | ERF | Os02g0752800 [Oryza sati | 1533 | 0 | down | 0 |
| SALT00000036893_Cluster23635-006 | ERF | Os02g0752800 [Oryza sati | 1721 | 0 | 0 | 0 |
| SALT00000042483_Cluster23635-007 | ERF | Os02g0752800 [Oryza sati | 1808 | 0 | 0 | 0 |
| SALT00000000327_Cluster23656-001 | ERF | AP2 EREBP transcription | 1828 | 0 | up | 0 |
| SALT00000004661_Cluster23656-002 | ERF | AP2 EREBP transcription | 1716 | 0 | 0 | 0 |
| SALT00000040121_Cluster23656-003 | ERF | AP2 EREBP transcription | 1675 | 0 | 0 | 0 |
| SALT00000046967_Cluster23656-004 | ERF | AP2 EREBP transcription | 1837 | 0 | 0 | 0 |
| SALT00000052710_Cluster23656-005 | ERF | AP2 EREBP transcription | 1837 | 0 | down | down |
| SALT00000000066_Cluster23716-001 | ERF | AP2-domain DRE binding | 1747 | 0 | 0 | down |
| SALT00000000521_Cluster23716-002 | ERF | AP2-domain DRE binding | 1833 | 0 | 0 | 0 |
| SALT00000008370_Cluster23716-003 | ERF | AP2-domain DRE binding | 1695 | 0 | 0 | 0 |
| SALT00000017062_Cluster23716-004 | ERF | AP2-domain DRE binding | 1436 | up | 0 | up |
| SALT00000018718_Cluster23716-005 | ERF | AP2-domain DRE binding | 1465 | 0 | 0 | 0 |
| SALT00000038843_Cluster23716-006 | ERF | AP2-domain DRE binding | 1739 | 0 | 0 | 0 |
| SALT00000039314_Cluster23716-007 | ERF | AP2-domain DRE binding | 1771 | 0 | 0 | 0 |
| SALT00000042381_Cluster23716-008 | ERF | AP2-domain DRE binding | 1833 | 0 | 0 | 0 |
| SALT00000043579_Cluster23716-009 | ERF | AP2-domain DRE binding | 1744 | 0 | 0 | 0 |
| SALT00000045520_Cluster23716-010 | ERF | AP2-domain DRE binding | 1801 | 0 | 0 | 0 |
| SALT00000046608_Cluster23716-011 | ERF | AP2-domain DRE binding | 1799 | 0 | 0 | 0 |
| SALT00000061177_Cluster23716-012 | ERF | AP2-domain DRE binding | 1729 | 0 | 0 | 0 |
| SALT00000003029_Cluster24204-001 | ERF | Os05g0316800 [Oryza sati | 1583 | 0 | 0 | 0 |
| SALT00000004400_Cluster24204-002 | ERF | Os05g0316800 [Oryza sati | 1785 | 0 | 0 | 0 |
| SALT00000014940_Cluster24204-003 | ERF | Os05g0316800 [Oryza sati | 1644 | 0 | 0 | 0 |
| SALT00000054688_Cluster24204-004 | ERF | Os05g0316800 [Oryza sati | 1808 | 0 | 0 | 0 |
| SALT00000019551_Cluster24204-005 | ERF | Os05g0316800 [Oryza sati | 1568 | 0 | 0 | 0 |
| SALT00000050198_Cluster25297-001 | ERF | Os01g0657400 [Oryza sati | 1747 | 0 | 0 | 0 |
| SALT00000037954_Cluster25412-001 | ERF | EREBP [Oryza sativa Japo | 1741 | 0 | 0 | 0 |
| SALT00000004983_Cluster25995-001 | ERF | ethylene response element | 1629 | 0 | 0 | up |
| SALT00000007621_Cluster25995-002 | ERF | ethylene response element | 1458 | 0 | 0 | 0 |
| SALT00000016965_Cluster25995-003 | ERF | ethylene response element | 1712 | 0 | 0 | 0 |
| SALT00000078990_Cluster25995-004 | ERF | ethylene response element | 1429 | 0 | 0 | 0 |
| SALT00000001793_Cluster26193-001 | ERF | ethylene-responsive transc | 1551 | 0 | 0 | 0 |
| SALT00000002945_Cluster26193-002 | ERF | ethylene-responsive transc | 1513 | 0 | 0 | 0 |
| SALT00000004560_Cluster26193-003 | ERF | ethylene-responsive transc | 1447 | 0 | 0 | 0 |

| | | | | | | |
|---|---|---|---|---|---|---|
| SALT00000006339_Cluster26193-004 | ERF | ethylene-responsive transc | 1503 | 0 | 0 | 0 |
| SALT00000007226_Cluster26193-005 | ERF | ethylene-responsive transc | 1517 | 0 | 0 | 0 |
| SALT00000008224_Cluster26193-006 | ERF | ethylene-responsive transc | 1362 | 0 | 0 | 0 |
| SALT00000008480_Cluster26193-007 | ERF | ethylene-responsive transc | 1382 | 0 | 0 | 0 |
| SALT00000011515_Cluster26193-008 | ERF | ethylene-responsive transc | 1360 | 0 | 0 | 0 |
| SALT00000026243_Cluster26193-009 | ERF | ethylene-responsive transc | 1518 | 0 | 0 | 0 |
| SALT00000042239_Cluster26193-010 | ERF | ethylene-responsive transc | 1481 | 0 | 0 | 0 |
| SALT00000042955_Cluster26193-011 | ERF | ethylene-responsive transc | 1699 | 0 | 0 | 0 |
| SALT00000044056_Cluster26193-012 | ERF | ethylene-responsive transc | 1569 | 0 | 0 | 0 |
| SALT00000044475_Cluster26193-013 | ERF | ethylene-responsive transc | 1563 | 0 | 0 | 0 |
| SALT00000056978_Cluster26193-014 | ERF | ethylene-responsive transc | 1427 | 0 | 0 | 0 |
| SALT00000061849_Cluster26193-015 | ERF | ethylene-responsive transc | 1520 | 0 | 0 | 0 |
| SALT00000092762_Cluster26193-016 | ERF | ethylene-responsive transc | 1226 | 0 | 0 | 0 |
| SALT00000057611_Cluster26193-017 | ERF | ethylene-responsive transc | 1507 | 0 | 0 | 0 |
| SALT00000061654_Cluster26351-001 | ERF | Os01g0657400 [Oryza sati | 1692 | 0 | 0 | 0 |
| SALT00000002665_Cluster27382-001 | ERF | dehydration responsive ele | 1641 | 0 | 0 | 0 |
| SALT00000020349_Cluster27382-002 | ERF | dehydration responsive ele | 1493 | 0 | 0 | 0 |
| SALT00000036421_Cluster27382-003 | ERF | dehydration responsive ele | 1574 | 0 | 0 | 0 |
| SALT00000055131_Cluster27382-004 | ERF | dehydration responsive fac | 1577 | 0 | 0 | 0 |
| SALT00000060631_Cluster28208-001 | ERF | ethylene-responsive transc | 1600 | 0 | 0 | 0 |
| SALT00000006321_Cluster28599-001 | ERF | DRE binding factor 1 [Zea | 1508 | 0 | 0 | 0 |
| SALT00000006703_Cluster28599-002 | ERF | DRE binding factor 1 [Zea | 1544 | 0 | 0 | 0 |
| SALT00000012350_Cluster28599-003 | ERF | DRE binding factor 1 [Zea | 1578 | up | 0 | 0 |
| SALT00000019776_Cluster28599-004 | ERF | DRE binding factor 1 [Zea | 1483 | 0 | 0 | 0 |
| SALT00000054619_Cluster28599-005 | ERF | DRE binding factor 1 [Zea | 1474 | 0 | 0 | 0 |
| SALT00000042977_Cluster28876-001 | ERF | dehydration responsive ele | 1526 | 0 | 0 | 0 |
| SALT00000045493_Cluster28876-002 | ERF | dehydration responsive ele | 1565 | 0 | 0 | 0 |
| SALT00000054822_Cluster28876-003 | ERF | dehydration responsive ele | 1480 | 0 | 0 | 0 |
| SALT00000035853_Cluster29326-001 | ERF | ethylene-responsive transc | 1542 | 0 | 0 | up |
| SALT00000037070_Cluster29489-001 | ERF | Os02g0654700 [Oryza sati | 1448 | 0 | 0 | 0 |
| SALT00000039468_Cluster29489-002 | ERF | Os02g0654700 [Oryza sati | 1483 | 0 | 0 | 0 |
| SALT00000059224_Cluster29489-003 | ERF | Os02g0654700 [Oryza sati | 1534 | 0 | 0 | 0 |
| SALT00000060198_Cluster29489-004 | ERF | Os02g0654700 [Oryza sati | 1123 | 0 | 0 | up |
| SALT00000042710_Cluster29489-005 | ERF | Os02g0654700 [Oryza sati | 1325 | 0 | 0 | up |
| SALT00000059843_Cluster29666-001 | ERF | AP2 EREBP transcription | 1523 | 0 | 0 | 0 |
| SALT00000044826_Cluster29904-001 | ERF | AP2 EREBP transcription | 1508 | 0 | 0 | 0 |
| SALT00000059057_Cluster29935-001 | ERF | Ethylene-responsive transc | 1506 | 0 | 0 | 0 |
| SALT00000002932_Cluster30718-001 | ERF | pathogenesis-related genes | 1450 | 0 | 0 | 0 |
| SALT00000005998_Cluster31032-001 | ERF | AP2-EREBP transcription | 1249 | 0 | 0 | 0 |
| SALT00000044327_Cluster31032-002 | ERF | ethylene-responsive transc | 1354 | 0 | 0 | 0 |
| SALT00000059416_Cluster31032-003 | ERF | AP2-EREBP transcription | 1416 | 0 | 0 | up |
| SALT00000042193_Cluster31606-001 | ERF | ethylene-responsive transc | 1307 | 0 | 0 | 0 |
| SALT00000023657_Cluster31801-001 | ERF | AP2 domain-containing tra | 1250 | 0 | 0 | 0 |
| SALT00000057145_Cluster31801-002 | ERF | AP2 domain-containing tra | 1210 | 0 | 0 | 0 |
| SALT00000012542_Cluster31821-001 | ERF | ethylene-responsive eleme | 1194 | 0 | 0 | 0 |
| SALT00000039804_Cluster31821-002 | ERF | ethylene-responsive eleme | 1146 | 0 | 0 | down |
| SALT00000040704_Cluster31821-003 | ERF | ethylene-responsive eleme | 1243 | 0 | 0 | up |
| SALT00000037593_Cluster31978-001 | ERF | [Hordeum vulgare vulgare | 1195 | 0 | 0 | down |
| SALT00000053642_Cluster31978-002 | ERF | dehydration responsive ele | 1191 | 0 | 0 | up |
| SALT00000036384_Cluster32082-001 | ERF | Os05g0361700 [Oryza sati | 1157 | 0 | 0 | 0 |
| SALT00000057228_Cluster32194-001 | ERF | Os03g0815800 [Oryza sati | 1118 | 0 | 0 | 0 |
| SALT00000061426_Cluster32198-001 | ERF | 7 [Hordeum vulgare vulgai | 1116 | 0 | 0 | 0 |
| SALT00000027370_Cluster32263-001 | ERF | partial [Oryza sativa Japon | 1087 | 0 | 0 | 0 |
| SALT00000071723_Cluster32412-001 | ERF | ethylene-responsive transc | 991 | 0 | 0 | 0 |
| SALT00000060612_Cluster32455-001 | ERF | AP2 EREBP transcription | 957 | 0 | 0 | 0 |
| SALT00000071879_Cluster32476-001 | ERF | ethylene-responsive transc | 938 | 0 | 0 | 0 |
| SALT00000073054_Cluster409-001 | FAR1 | Os03g0255400 [Oryza sati | 8159 | 0 | 0 | 0 |
| SALT00000032367_Cluster628-003 | FAR1 | FAR1-RELATED SEQUE | 3772 | 0 | 0 | 0 |

| | | | | | | |
|---|---|---|---|---|---|---|
| SALT00000033457_Cluster628-004 | FAR1 | FAR1-RELATED SEQUE | 3967 | 0 | 0 | 0 |
| SALT00000034484_Cluster628-005 | FAR1 | FAR1-RELATED SEQUE | 3885 | 0 | 0 | 0 |
| SALT00000086213_Cluster628-007 | FAR1 | FAR1-RELATED SEQUE | 3486 | 0 | 0 | 0 |
| SALT00000086408_Cluster628-008 | FAR1 | FAR1-RELATED SEQUE | 3550 | 0 | 0 | 0 |
| SALT00000090623_Cluster628-009 | FAR1 | FAR1-RELATED SEQUE | 3929 | 0 | 0 | 0 |
| SALT00000092053_Cluster628-011 | FAR1 | FAR1-RELATED SEQUE | 4159 | 0 | 0 | 0 |
| SALT00000028014_Cluster1006-001 | FAR1 | FAR1-RELATED SEQUE | 2996 | 0 | 0 | 0 |
| SALT00000063389_Cluster1006-002 | FAR1 | FAR1-RELATED SEQUE | 2917 | 0 | 0 | 0 |
| SALT00000076409_Cluster1006-004 | FAR1 | FAR1-RELATED SEQUE | 2884 | 0 | 0 | 0 |
| SALT00000021408_Cluster1715-001 | FAR1 | FAR1-RELATED SEQUE | 2968 | 0 | 0 | 0 |
| SALT00000024587_Cluster1715-002 | FAR1 | FAR1-RELATED SEQUE | 2567 | 0 | 0 | 0 |
| SALT00000026502_Cluster1715-003 | FAR1 | FAR1-RELATED SEQUE | 2514 | 0 | 0 | 0 |
| SALT00000027964_Cluster1715-004 | FAR1 | FAR1-RELATED SEQUE | 2911 | 0 | 0 | 0 |
| SALT00000028139_Cluster1715-005 | FAR1 | FAR1-RELATED SEQUE | 2591 | 0 | 0 | 0 |
| SALT00000078482_Cluster1715-009 | FAR1 | FAR1-RELATED SEQUE | 3328 | 0 | 0 | 0 |
| SALT00000088974_Cluster1715-010 | FAR1 | FAR1-RELATED SEQUE | 4547 | 0 | 0 | 0 |
| SALT00000021094_Cluster1856-001 | FAR1 | Os07g0615900 [Oryza sati | 2556 | 0 | 0 | 0 |
| SALT00000022543_Cluster1856-002 | FAR1 | Os07g0615900 [Oryza sati | 2480 | 0 | 0 | 0 |
| SALT00000023590_Cluster1856-003 | FAR1 | Os07g0615900 [Oryza sati | 2871 | 0 | 0 | 0 |
| SALT00000030184_Cluster1856-004 | FAR1 | Os07g0615900 [Oryza sati | 2813 | 0 | 0 | 0 |
| SALT00000079927_Cluster1856-005 | FAR1 | Os07g0615900 [Oryza sati | 2558 | 0 | 0 | 0 |
| SALT00000084032_Cluster1856-007 | FAR1 | Os07g0615900 [Oryza sati | 2618 | 0 | 0 | 0 |
| SALT00000029436_Cluster2162-001 | FAR1 | FAR1-RELATED SEQUE | 3202 | 0 | 0 | 0 |
| SALT00000089606_Cluster2162-002 | FAR1 | FAR1-RELATED SEQUE | 3859 | 0 | 0 | 0 |
| SALT00000090096_Cluster2162-003 | FAR1 | FAR1-RELATED SEQUE | 5041 | 0 | 0 | 0 |
| SALT00000030822_Cluster2400-001 | FAR1 | Os07g0615900 [Oryza sati | 3011 | 0 | 0 | 0 |
| SALT00000027961_Cluster2468-002 | FAR1 | FAR1-RELATED SEQUE | 3450 | 0 | 0 | 0 |
| SALT00000032591_Cluster2468-003 | FAR1 | FAR1-RELATED SEQUE | 3515 | 0 | 0 | 0 |
| SALT00000033290_Cluster2468-004 | FAR1 | FAR1-RELATED SEQUE | 3608 | 0 | 0 | 0 |
| SALT00000034040_Cluster2468-005 | FAR1 | FAR1-RELATED SEQUE | 3440 | 0 | 0 | 0 |
| SALT00000034221_Cluster2468-006 | FAR1 | FAR1-RELATED SEQUE | 3458 | 0 | 0 | 0 |
| SALT00000063839_Cluster2468-007 | FAR1 | FAR1-RELATED SEQUE | 3316 | 0 | 0 | 0 |
| SALT00000086823_Cluster2468-008 | FAR1 | FAR1-RELATED SEQUE | 4863 | 0 | 0 | 0 |
| SALT00000088002_Cluster2468-009 | FAR1 | FAR1-RELATED SEQUE | 3521 | 0 | 0 | 0 |
| SALT00000089832_Cluster2468-010 | FAR1 | FAR1-RELATED SEQUE | 3651 | 0 | 0 | 0 |
| SALT00000090646_Cluster2468-011 | FAR1 | FAR1-RELATED SEQUE | 3513 | 0 | 0 | 0 |
| SALT00000092801_Cluster2468-012 | FAR1 | FAR1-RELATED SEQUE | 3509 | 0 | 0 | 0 |
| SALT00000066910_Cluster3083-001 | FAR1 | FAR1-RELATED SEQUE | 2723 | 0 | 0 | 0 |
| SALT00000023233_Cluster3206-001 | FAR1 | far-red impaired response [ | 2816 | 0 | 0 | 0 |
| SALT00000026602_Cluster3206-002 | FAR1 | FAR1-RELATED SEQUE | 2665 | 0 | 0 | 0 |
| SALT00000026716_Cluster3206-003 | FAR1 | FAR1-RELATED SEQUE | 2548 | 0 | 0 | 0 |
| SALT00000082266_Cluster3206-005 | FAR1 | far-red impaired response [ | 2897 | 0 | 0 | 0 |
| SALT00000034192_Cluster3397-001 | FAR1 | FAR1-RELATED SEQUE | 4435 | 0 | 0 | 0 |
| SALT00000044799_Cluster3397-002 | FAR1 | FAR1-RELATED SEQUE | 2149 | 0 | 0 | 0 |
| SALT00000080473_Cluster3397-004 | FAR1 | FAR1-RELATED SEQUE | 3054 | 0 | 0 | 0 |
| SALT00000021311_Cluster3600-001 | FAR1 | Os03g0255400 [Oryza sati | 2918 | 0 | 0 | 0 |
| SALT00000026402_Cluster3600-002 | FAR1 | Os03g0255400 [Oryza sati | 2719 | 0 | 0 | 0 |
| SALT00000073349_Cluster3600-003 | FAR1 | Os03g0255400 [Oryza sati | 2714 | 0 | 0 | 0 |
| SALT00000076544_Cluster3600-005 | FAR1 | Os03g0255400 [Oryza sati | 2858 | 0 | 0 | 0 |
| SALT00000077617_Cluster3600-006 | FAR1 | Os03g0255400 [Oryza sati | 2853 | 0 | 0 | 0 |
| SALT00000078102_Cluster3600-007 | FAR1 | Os03g0255400 [Oryza sati | 2795 | 0 | 0 | 0 |
| SALT00000084323_Cluster3890-002 | FAR1 | FAR1-RELATED SEQUE | 4261 | 0 | 0 | 0 |
| SALT00000029285_Cluster5444-001 | FAR1 | FAR1-RELATED SEQUE | 3857 | 0 | 0 | 0 |
| SALT00000033762_Cluster5444-002 | FAR1 | FAR1-RELATED SEQUE | 3887 | 0 | 0 | 0 |
| SALT00000021873_Cluster5818-001 | FAR1 | FAR1-RELATED SEQUE | 3160 | 0 | 0 | 0 |
| SALT00000032435_Cluster5818-002 | FAR1 | FAR1-RELATED SEQUE | 3419 | 0 | 0 | 0 |
| SALT00000077789_Cluster5818-003 | FAR1 | FAR1-RELATED SEQUE | 3065 | 0 | 0 | 0 |
| SALT00000087654_Cluster5818-004 | FAR1 | FAR1-RELATED SEQUE | 3824 | 0 | 0 | 0 |

| | | | | | | |
|---|---|---|---|---|---|---|
| SALT00000088932_Cluster5818-005 | FAR1 | FAR1-RELATED SEQUE | 3438 | 0 | 0 | 0 |
| SALT00000032454_Cluster5952-001 | FAR1 | FAR1-domain family sequ | 3529 | 0 | 0 | 0 |
| SALT00000090400_Cluster5952-002 | FAR1 | Os07g0563600 [Oryza sati | 3800 | 0 | 0 | 0 |
| SALT00000023105_Cluster8477-001 | FAR1 | Os07g0496300 [Oryza sati | 2575 | 0 | 0 | 0 |
| SALT00000024747_Cluster8477-002 | FAR1 | Os07g0496300 [Oryza sati | 2924 | 0 | 0 | 0 |
| SALT00000068091_Cluster8477-003 | FAR1 | Os07g0496300 [Oryza sati | 2821 | 0 | 0 | 0 |
| SALT00000078769_Cluster8477-004 | FAR1 | Os07g0496300 [Oryza sati | 2672 | 0 | 0 | 0 |
| SALT00000090495_Cluster8477-005 | FAR1 | Os07g0496300 [Oryza sati | 3390 | 0 | 0 | 0 |
| SALT00000078669_Cluster8477-006 | FAR1 | Os07g0496300 [Oryza sati | 2732 | 0 | 0 | 0 |
| SALT00000080743_Cluster8823-001 | FAR1 | FAR1-RELATED SEQUE | 3322 | 0 | 0 | 0 |
| SALT00000022521_Cluster9226-002 | FAR1 | Os12g0160900 [Oryza sati | 2645 | 0 | 0 | 0 |
| SALT00000026840_Cluster9226-003 | FAR1 | Os12g0160900 [Oryza sati | 3245 | 0 | 0 | 0 |
| SALT00000034273_Cluster9337-001 | FAR1 | FAR1-RELATED SEQUE | 3175 | 0 | 0 | 0 |
| SALT00000067550_Cluster9337-003 | FAR1 | FAR1-RELATED SEQUE | 3221 | 0 | 0 | 0 |
| SALT00000069381_Cluster9337-004 | FAR1 | FAR1-RELATED SEQUE | 2753 | 0 | 0 | 0 |
| SALT00000080571_Cluster9337-005 | FAR1 | FAR1-RELATED SEQUE | 2934 | 0 | 0 | 0 |
| SALT00000028078_Cluster10497-001 | FAR1 | FAR1-RELATED SEQUE | 2835 | 0 | 0 | 0 |
| SALT00000067123_Cluster10497-002 | FAR1 | FAR1-RELATED SEQUE | 2921 | 0 | 0 | 0 |
| SALT00000067194_Cluster10497-003 | FAR1 | FAR1-RELATED SEQUE | 3038 | 0 | 0 | 0 |
| SALT00000069705_Cluster10497-004 | FAR1 | FAR1-RELATED SEQUE | 2968 | 0 | 0 | 0 |
| SALT00000071342_Cluster10497-005 | FAR1 | FAR1-RELATED SEQUE | 2896 | 0 | 0 | 0 |
| SALT00000073544_Cluster10497-006 | FAR1 | FAR1-RELATED SEQUE | 2863 | 0 | 0 | 0 |
| SALT00000075437_Cluster10497-007 | FAR1 | FAR1-RELATED SEQUE | 2929 | 0 | 0 | 0 |
| SALT00000075892_Cluster10497-008 | FAR1 | FAR1-RELATED SEQUE | 2743 | 0 | 0 | 0 |
| SALT00000077366_Cluster10497-009 | FAR1 | FAR1-RELATED SEQUE | 2997 | 0 | 0 | 0 |
| SALT00000078284_Cluster10497-010 | FAR1 | FAR1-RELATED SEQUE | 3007 | 0 | 0 | 0 |
| SALT00000079745_Cluster10497-011 | FAR1 | FAR1-RELATED SEQUE | 2948 | 0 | 0 | 0 |
| SALT00000084297_Cluster10497-012 | FAR1 | FAR1-RELATED SEQUE | 2967 | 0 | 0 | 0 |
| SALT00000091312_Cluster10497-013 | FAR1 | FAR1-RELATED SEQUE | 2989 | 0 | 0 | 0 |
| SALT00000068721_Cluster10548-001 | FAR1 | FAR1-RELATED SEQUE | 3031 | 0 | 0 | 0 |
| SALT00000025931_Cluster10591-001 | FAR1 | FAR1-RELATED SEQUE | 2999 | 0 | 0 | 0 |
| SALT00000077294_Cluster10591-002 | FAR1 | far-red impaired response [ | 3024 | 0 | 0 | 0 |
| SALT00000022862_Cluster11720-001 | FAR1 | FAR1-RELATED SEQUE | 2740 | 0 | 0 | 0 |
| SALT00000024725_Cluster11720-002 | FAR1 | FAR1-RELATED SEQUE | 2599 | 0 | 0 | 0 |
| SALT00000067430_Cluster11720-003 | FAR1 | FAR1-RELATED SEQUE | 2840 | 0 | 0 | 0 |
| SALT00000070684_Cluster11720-004 | FAR1 | FAR1-RELATED SEQUE | 2675 | 0 | 0 | 0 |
| SALT00000079663_Cluster11720-005 | FAR1 | FAR1-RELATED SEQUE | 2893 | 0 | 0 | 0 |
| SALT00000029039_Cluster11998-001 | FAR1 | Os03g0255400 [Oryza sati | 2867 | 0 | 0 | 0 |
| SALT00000082067_Cluster11998-002 | FAR1 | Os03g0255400 [Oryza sati | 2701 | 0 | 0 | 0 |
| SALT00000022268_Cluster13209-001 | FAR1 | FAR1-RELATED SEQUE | 2756 | 0 | 0 | 0 |
| SALT00000023147_Cluster13444-001 | FAR1 | FAR1-RELATED SEQUE | 2723 | 0 | 0 | 0 |
| SALT00000026011_Cluster13444-002 | FAR1 | FAR1-RELATED SEQUE | 2736 | 0 | 0 | 0 |
| SALT00000026417_Cluster13510-001 | FAR1 | far-red impaired response [ | 2731 | 0 | 0 | 0 |
| SALT00000078529_Cluster15570-001 | FAR1 | FAR-RED IMPAIRED RE | 2584 | 0 | 0 | 0 |
| SALT00000024478_Cluster15628-001 | FAR1 | FAR1-RELATED SEQUE | 2492 | 0 | 0 | 0 |
| SALT00000024567_Cluster15628-002 | FAR1 | FAR1-RELATED SEQUE | 2546 | 0 | 0 | 0 |
| SALT00000026638_Cluster15628-003 | FAR1 | FAR1-RELATED SEQUE | 2579 | 0 | 0 | 0 |
| SALT00000027156_Cluster15628-004 | FAR1 | FAR1-RELATED SEQUE | 2577 | 0 | 0 | 0 |
| SALT00000071153_Cluster16127-001 | FAR1 | FAR1-RELATED SEQUE | 2542 | 0 | 0 | up |
| SALT00000062442_Cluster16929-001 | FAR1 | FAR1-RELATED SEQUE | 2477 | 0 | 0 | 0 |
| SALT00000030366_Cluster19348-001 | FAR1 | far-red impaired response [ | 2133 | 0 | 0 | 0 |
| SALT00000067205_Cluster22672-001 | FAR1 | FAR1-RELATED SEQUE | 1899 | 0 | 0 | 0 |
| SALT00000019736_Cluster26210-001 | FAR1 | FAR1-RELATED SEQUE | 1698 | 0 | 0 | 0 |
| SALT00000059359_Cluster263-002 | G2-like | transfactor [Oryza sativa Ja | 1448 | 0 | 0 | 0 |
| SALT00000088174_Cluster6168-001 | G2-like | two-component response r | 3766 | 0 | 0 | 0 |
| SALT00000092100_Cluster6867-001 | G2-like | transcription factor | 3656 | 0 | 0 | 0 |
| SALT00000065899_Cluster7076-004 | G2-like | two-component response r | 2140 | 0 | 0 | 0 |
| SALT00000036105_Cluster7180-001 | G2-like | KANADI1 [Zea mays] | 1797 | 0 | 0 | 0 |

| | | | | | | |
|---|---|---|---|---|---|---|
| SALT00000037748_Cluster7180-002 | G2-like | KANADI1 [Zea mays] | 1692 | 0 | 0 | 0 |
| SALT00000091425_Cluster7180-003 | G2-like | KANADI1 [Zea mays] | 3611 | 0 | 0 | 0 |
| SALT00000043655_Cluster7180-004 | G2-like | KANADI3 [Zea mays] | 1658 | 0 | 0 | 0 |
| SALT00000012997_Cluster7180-005 | G2-like | KANADI1 [Zea mays] | 1709 | 0 | 0 | 0 |
| SALT00000047353_Cluster8748-001 | G2-like | partial [Micromonas pusill | 3337 | 0 | 0 | 0 |
| SALT00000054769_Cluster8748-002 | G2-like | myb-like DNA-binding SF | 1483 | 0 | 0 | 0 |
| SALT00000091545_Cluster8974-001 | G2-like | Os02g0814600 [Oryza sati | 3292 | 0 | 0 | 0 |
| SALT00000005447_Cluster9318-001 | G2-like | PHR1-LIKE 1-like [Brach | 1877 | 0 | 0 | 0 |
| SALT00000008250_Cluster9318-002 | G2-like | PHR1-LIKE 1-like [Brach | 1854 | 0 | 0 | 0 |
| SALT00000013122_Cluster9318-003 | G2-like | PHR1-LIKE 1-like [Brach | 1857 | 0 | 0 | 0 |
| SALT00000073867_Cluster9318-004 | G2-like | PHR1-LIKE 1-like [Brach | 3226 | 0 | 0 | 0 |
| SALT00000084367_Cluster9318-005 | G2-like | ribonuclease H At1g65750 | 2569 | 0 | 0 | 0 |
| SALT00000057715_Cluster10755-004 | G2-like | myb family transcription fa | 1381 | 0 | 0 | 0 |
| SALT00000071670_Cluster11421-002 | G2-like | ribonuclease H At1g65750 | 2841 | 0 | 0 | 0 |
| SALT00000082444_Cluster11421-003 | G2-like | ribonuclease H At1g65750 | 2922 | 0 | 0 | 0 |
| SALT00000043051_Cluster11421-004 | G2-like | PHR1-LIKE 1-like [Brach | 1682 | 0 | 0 | 0 |
| SALT00000072024_Cluster14623-001 | G2-like | PHR1-LIKE 1-like isoform | 2651 | 0 | 0 | 0 |
| SALT00000082293_Cluster14768-001 | G2-like | CDPK substrate 1 CSP1 [( | 2642 | 0 | 0 | 0 |
| SALT00000013636_Cluster16424-001 | G2-like | Two-component response r | 1957 | 0 | 0 | 0 |
| SALT00000056863_Cluster16424-002 | G2-like | two-component response r | 1884 | 0 | 0 | 0 |
| SALT00000039815_Cluster19785-001 | G2-like | [Hordeum vulgare vulgare] | 2096 | 0 | 0 | 0 |
| SALT00000008032_Cluster20285-001 | G2-like | phosphate starvation respo | 1927 | 0 | 0 | 0 |
| SALT00000011254_Cluster20285-002 | G2-like | phosphate starvation respo | 1954 | 0 | 0 | 0 |
| SALT00000013624_Cluster20285-003 | G2-like | PHR1-LIKE 1-like [Setaria | 1747 | 0 | 0 | 0 |
| SALT00000055123_Cluster20285-004 | G2-like | phosphate starvation respo | 2056 | 0 | 0 | 0 |
| SALT00000093753_Cluster20285-005 | G2-like | phosphate starvation respo | 1928 | 0 | 0 | 0 |
| SALT00000054299_Cluster20285-006 | G2-like | phosphate starvation respo | 1797 | 0 | 0 | 0 |
| SALT00000003134_Cluster20547-001 | G2-like | PHR1-LIKE 1-like [Brach | 1672 | 0 | 0 | 0 |
| SALT00000008230_Cluster20547-002 | G2-like | PHR1-LIKE 1-like [Brach | 1836 | 0 | 0 | 0 |
| SALT00000009981_Cluster20547-003 | G2-like | PHR1-LIKE 1-like [Brach | 1699 | 0 | 0 | 0 |
| SALT00000010668_Cluster20547-004 | G2-like | PHR1-LIKE 1-like [Brach | 1840 | 0 | 0 | 0 |
| SALT00000011789_Cluster20547-005 | G2-like | PHR1-LIKE 1-like [Brach | 1705 | 0 | 0 | 0 |
| SALT00000041821_Cluster20547-006 | G2-like | PHR1-LIKE 1-like [Brach | 2037 | 0 | 0 | 0 |
| SALT00000051241_Cluster20547-007 | G2-like | PHR1-LIKE 1-like [Brach | 1814 | 0 | 0 | 0 |
| SALT00000044059_Cluster20547-008 | G2-like | PHR1-LIKE 1-like [Brach | 1741 | 0 | 0 | 0 |
| SALT00000008650_Cluster21111-001 | G2-like | probable transcription facto | 1997 | 0 | 0 | 0 |
| SALT00000015568_Cluster21348-001 | G2-like | myb-like DNA-binding SF | 1981 | 0 | 0 | 0 |
| SALT00000054584_Cluster21348-002 | G2-like | myb-like DNA-binding SF | 1694 | 0 | 0 | 0 |
| SALT00000044597_Cluster21348-003 | G2-like | myb-like DNA-binding SF | 1488 | 0 | 0 | down |
| SALT00000058037_Cluster21616-001 | G2-like | Os04g0665600 [Oryza sati | 1956 | 0 | 0 | 0 |
| SALT00000059376_Cluster21616-002 | G2-like | Os04g0665600 [Oryza sati | 1965 | 0 | 0 | 0 |
| SALT00000000918_Cluster21665-001 | G2-like | PHR1-LIKE 1-like [Brach | 1961 | 0 | 0 | 0 |
| SALT00000010159_Cluster21665-002 | G2-like | PHR1-LIKE 1-like [Brach | 1808 | 0 | 0 | 0 |
| SALT00000013725_Cluster21665-003 | G2-like | PHR1-LIKE 1-like [Brach | 1952 | 0 | 0 | 0 |
| SALT00000013929_Cluster21665-004 | G2-like | PHR1-LIKE 1-like [Brach | 1824 | 0 | 0 | 0 |
| SALT00000018543_Cluster21665-005 | G2-like | PHR1-LIKE 1-like [Brach | 1907 | 0 | 0 | 0 |
| SALT00000049410_Cluster21665-006 | G2-like | PHR1-LIKE 1-like [Brach | 1526 | 0 | 0 | 0 |
| SALT00000061542_Cluster21665-007 | G2-like | PHR1-LIKE 1-like [Brach | 1739 | 0 | 0 | 0 |
| SALT00000037169_Cluster21898-001 | G2-like | MYB DNA-binding domai | 1946 | 0 | 0 | 0 |
| SALT00000053463_Cluster21898-002 | G2-like | Os06g0543200 [Oryza sati | 1661 | 0 | 0 | 0 |
| SALT00000020166_Cluster21898-003 | G2-like | Os06g0543200 [Oryza sati | 1855 | 0 | 0 | 0 |
| SALT00000039991_Cluster22115-001 | G2-like | KANADI1 [Zea mays] | 1851 | 0 | 0 | 0 |
| SALT00000040105_Cluster22115-002 | G2-like | KANADI1 [Zea mays] | 1931 | 0 | 0 | 0 |
| SALT00000012172_Cluster22115-003 | G2-like | KANADI1 [Zea mays] | 1742 | 0 | 0 | 0 |
| SALT00000009280_Cluster22161-001 | G2-like | probable transcription facto | 1762 | 0 | 0 | 0 |
| SALT00000047678_Cluster22161-002 | G2-like | probable transcription facto | 1929 | 0 | 0 | 0 |
| SALT00000047684_Cluster22161-003 | G2-like | probable transcription facto | 1747 | 0 | 0 | 0 |
| SALT00000053128_Cluster22161-004 | G2-like | probable transcription facto | 1598 | 0 | 0 | 0 |

| | | | | | | |
|---|---|---|---|---|---|---|
| SALT00000048836_Cluster22439-001 | G2-like | transfactor [Oryza sativa Ja | 1911 | 0 | 0 | 0 |
| SALT00000057632_Cluster22617-001 | G2-like | [Hordeum vulgare vulgare] | 1902 | 0 | 0 | 0 |
| SALT00000011927_Cluster23177-001 | G2-like | MYB DNA-binding domai | 1866 | 0 | 0 | 0 |
| SALT00000043744_Cluster23177-002 | G2-like | Os06g0543200 [Oryza sati | 1499 | 0 | 0 | 0 |
| SALT00000009330_Cluster23905-001 | G2-like | Os06g0348800 [Oryza sati | 1689 | 0 | 0 | 0 |
| SALT00000016359_Cluster23905-002 | G2-like | Os06g0348800 [Oryza sati | 1732 | 0 | 0 | 0 |
| SALT00000048852_Cluster23905-003 | G2-like | Os06g0348800 [Oryza sati | 1795 | 0 | 0 | 0 |
| SALT00000055103_Cluster23905-004 | G2-like | MYB DNA-binding domai | 1824 | 0 | 0 | 0 |
| SALT00000012416_Cluster24485-001 | G2-like | myb family transcription fa | 1570 | 0 | 0 | 0 |
| SALT00000023628_Cluster24485-002 | G2-like | myb family transcription fa | 1647 | 0 | 0 | 0 |
| SALT00000040593_Cluster24485-003 | G2-like | myb family transcription fa | 1571 | 0 | 0 | 0 |
| SALT00000043084_Cluster24485-004 | G2-like | myb family transcription fa | 1574 | 0 | 0 | 0 |
| SALT00000048446_Cluster24485-005 | G2-like | myb family transcription fa | 1791 | 0 | 0 | 0 |
| SALT00000014355_Cluster24873-001 | G2-like | PHR1-LIKE 1 [Oryza brac | 1769 | 0 | 0 | 0 |
| SALT00000007004_Cluster25579-001 | G2-like | Os02g0325600 [Oryza sati | 1733 | 0 | 0 | 0 |
| SALT00000000597_Cluster25742-001 | G2-like | Os02g0325600 [Oryza sati | 1594 | up | 0 | 0 |
| SALT00000015877_Cluster25742-002 | G2-like | Os02g0325600 [Oryza sati | 1556 | 0 | 0 | 0 |
| SALT00000042442_Cluster25742-003 | G2-like | Os02g0325600 [Oryza sati | 1725 | 0 | 0 | 0 |
| SALT00000044449_Cluster25742-004 | G2-like | Os02g0325600 [Oryza sati | 1683 | 0 | 0 | 0 |
| SALT00000047437_Cluster25742-005 | G2-like | Os02g0325600 [Oryza sati | 1586 | 0 | 0 | 0 |
| SALT00000048397_Cluster25742-006 | G2-like | Os02g0325600 [Oryza sati | 1480 | 0 | 0 | 0 |
| SALT00000039277_Cluster25742-007 | G2-like | Os02g0325600 [Oryza sati | 1513 | 0 | 0 | down |
| SALT00000054987_Cluster25792-001 | G2-like | PHR1-LIKE 1 [Oryza brac | 1723 | 0 | 0 | 0 |
| SALT00000010859_Cluster25915-001 | G2-like | transcription factor 1 | 1616 | 0 | 0 | 0 |
| SALT00000016671_Cluster25915-002 | G2-like | transcription factor 1 | 1675 | 0 | 0 | 0 |
| SALT00000043071_Cluster25915-003 | G2-like | transcription factor 1 | 1642 | 0 | 0 | 0 |
| SALT00000045547_Cluster25915-004 | G2-like | transcription factor 1 | 1716 | 0 | 0 | 0 |
| SALT00000093755_Cluster25915-005 | G2-like | transcription factor 1 | 1661 | 0 | 0 | 0 |
| SALT00000010919_Cluster26072-001 | G2-like | MYB DNA-binding domai | 1707 | 0 | 0 | 0 |
| SALT00000017528_Cluster26072-002 | G2-like | MYB DNA-binding domai | 1646 | 0 | 0 | 0 |
| SALT00000044547_Cluster26443-001 | G2-like | Os12g0586300 [Oryza sati | 1687 | 0 | 0 | 0 |
| SALT00000059022_Cluster26450-001 | G2-like | MYB-CC type transfactor | 1687 | 0 | 0 | 0 |
| SALT00000056066_Cluster26883-001 | G2-like | transcription factor PCL1-l | 1665 | 0 | 0 | 0 |
| SALT00000015742_Cluster27011-001 | G2-like | MYB DNA-binding domai | 1521 | 0 | 0 | 0 |
| SALT00000018704_Cluster27011-002 | G2-like | MYB DNA-binding domai | 1659 | 0 | 0 | 0 |
| SALT00000008642_Cluster27527-001 | G2-like | Myb family transcription f | 1500 | 0 | 0 | 0 |
| SALT00000013592_Cluster27527-002 | G2-like | Myb family transcription f | 1634 | 0 | 0 | 0 |
| SALT00000037088_Cluster27527-003 | G2-like | myb family transcription fa | 1560 | 0 | 0 | 0 |
| SALT00000039637_Cluster27527-004 | G2-like | Myb family transcription f | 1500 | 0 | 0 | 0 |
| SALT00000045866_Cluster27527-005 | G2-like | myb family transcription fa | 1425 | 0 | 0 | 0 |
| SALT00000052413_Cluster27527-006 | G2-like | Myb family transcription f | 1517 | 0 | 0 | 0 |
| SALT00000043399_Cluster27678-001 | G2-like | Os03g0766500 [Oryza sati | 1627 | 0 | 0 | 0 |
| SALT00000052319_Cluster28189-001 | G2-like | Os06g0670300 [Oryza sati | 1601 | 0 | 0 | up |
| SALT00000013288_Cluster28417-001 | G2-like | transfactor [Oryza sativa Ja | 1588 | 0 | 0 | 0 |
| SALT00000050087_Cluster28417-002 | G2-like | transfactor [Oryza sativa Ja | 1458 | 0 | 0 | 0 |
| SALT00000041587_Cluster28426-001 | G2-like | MYB DNA-binding domai | 1588 | 0 | 0 | 0 |
| SALT00000043478_Cluster28652-001 | G2-like | DNA binding [Zea mays] | 1461 | 0 | 0 | up |
| SALT00000046686_Cluster28652-002 | G2-like | DNA binding [Zea mays] | 1576 | 0 | 0 | 0 |
| SALT00000051951_Cluster28799-001 | G2-like | partial [Oryza sativa Japon | 1446 | 0 | 0 | 0 |
| SALT00000055071_Cluster28799-002 | G2-like | partial [Oryza sativa Japon | 1569 | 0 | 0 | 0 |
| SALT00000057098_Cluster28799-003 | G2-like | partial [Oryza sativa Japon | 1432 | 0 | 0 | 0 |
| SALT00000051504_Cluster28799-004 | G2-like | partial [Oryza sativa Japon | 1528 | 0 | 0 | 0 |
| SALT00000050601_Cluster29075-001 | G2-like | PHR1-LIKE 1 [Oryza brac | 1555 | 0 | 0 | 0 |
| SALT00000013409_Cluster29346-001 | G2-like | Os03g0766500 [Oryza sati | 1541 | 0 | 0 | 0 |
| SALT00000045787_Cluster30546-001 | G2-like | transcription factor PCL1-l | 1390 | 0 | 0 | 0 |
| SALT00000064625_Cluster30546-002 | G2-like | transcription factor PCL1-l | 1465 | 0 | 0 | 0 |
| SALT00000036427_Cluster30689-001 | G2-like | PHR1-LIKE 1-like isoforn | 1412 | 0 | 0 | 0 |
| SALT00000044307_Cluster30689-002 | G2-like | PHR1-LIKE 1-like isoforn | 1453 | 0 | 0 | 0 |

| | | | | | | |
|---|---|---|---|---|---|---|
| SALT00000017030_Cluster31413-001 | G2-like | Os12g0586300 [Oryza sati | 1184 | 0 | up | 0 |
| SALT00000035727_Cluster31413-002 | G2-like | Os12g0586300 [Oryza sati | 1350 | 0 | 0 | 0 |
| SALT00000045566_Cluster31413-003 | G2-like | Os12g0586300 [Oryza sati | 1521 | 0 | 0 | 0 |
| SALT00000054995_Cluster31413-004 | G2-like | Os12g0586300 [Oryza sati | 1191 | 0 | 0 | down |
| SALT00000017342_Cluster31413-005 | G2-like | myb-like DNA-binding SH | 1589 | 0 | 0 | 0 |
| SALT00000004694_Cluster3908-001 | GATA | GATA transcription factor | 1731 | 0 | 0 | 0 |
| SALT00000005399_Cluster3908-002 | GATA | GATA transcription factor | 1719 | 0 | 0 | 0 |
| SALT00000014197_Cluster3908-003 | GATA | GATA transcription factor | 1686 | 0 | 0 | 0 |
| SALT00000017771_Cluster3908-004 | GATA | GATA transcription factor | 1623 | 0 | 0 | 0 |
| SALT00000020333_Cluster3908-005 | GATA | GATA transcription factor | 1502 | 0 | 0 | 0 |
| SALT00000046810_Cluster3908-007 | GATA | GATA transcription factor | 1490 | 0 | 0 | 0 |
| SALT00000011666_Cluster8940-001 | GATA | GATA transcription factor | 1703 | 0 | 0 | 0 |
| SALT00000016640_Cluster8940-002 | GATA | GATA transcription factor | 1965 | 0 | 0 | 0 |
| SALT00000047416_Cluster8940-003 | GATA | GATA transcription factor | 2033 | 0 | 0 | 0 |
| SALT00000059267_Cluster8940-004 | GATA | GATA transcription factor | 1709 | 0 | 0 | 0 |
| SALT00000071374_Cluster8940-006 | GATA | GATA transcription factor | 3298 | 0 | 0 | 0 |
| SALT00000035889_Cluster9461-001 | GATA | GATA transcription factor | 1611 | 0 | 0 | 0 |
| SALT00000039723_Cluster9461-002 | GATA | GATA transcription factor | 1618 | 0 | 0 | 0 |
| SALT00000050627_Cluster9461-003 | GATA | GATA transcription factor | 1624 | 0 | 0 | 0 |
| SALT00000059354_Cluster9461-004 | GATA | GATA transcription factor | 1601 | 0 | 0 | up |
| SALT00000074889_Cluster9461-005 | GATA | GATA transcription factor | 3199 | 0 | 0 | up |
| SALT00000003143_Cluster10613-001 | GATA | GATA transcription factor | 1689 | 0 | 0 | 0 |
| SALT00000003783_Cluster10613-002 | GATA | GATA transcription factor | 1948 | 0 | 0 | 0 |
| SALT00000006598_Cluster10613-003 | GATA | GATA transcription factor | 1503 | 0 | 0 | 0 |
| SALT00000008335_Cluster10613-004 | GATA | GATA transcription factor | 1497 | 0 | 0 | 0 |
| SALT00000008641_Cluster10613-005 | GATA | GATA transcription factor | 1797 | 0 | 0 | 0 |
| SALT00000014634_Cluster10613-006 | GATA | GATA transcription factor | 1907 | 0 | 0 | 0 |
| SALT00000015465_Cluster10613-007 | GATA | GATA transcription factor | 2065 | 0 | 0 | 0 |
| SALT00000018395_Cluster10613-008 | GATA | GATA transcription factor | 1955 | 0 | 0 | 0 |
| SALT00000018989_Cluster10613-009 | GATA | GATA transcription factor | 1973 | 0 | 0 | 0 |
| SALT00000038133_Cluster10613-010 | GATA | GATA transcription factor | 2010 | 0 | 0 | 0 |
| SALT00000038978_Cluster10613-011 | GATA | GATA transcription factor | 1584 | 0 | 0 | 0 |
| SALT00000039531_Cluster10613-012 | GATA | GATA transcription factor | 2015 | 0 | 0 | 0 |
| SALT00000044416_Cluster10613-013 | GATA | [Hordeum vulgare vulgare] | 1929 | 0 | 0 | 0 |
| SALT00000047151_Cluster10613-014 | GATA | GATA transcription factor | 1841 | 0 | 0 | 0 |
| SALT00000048454_Cluster10613-015 | GATA | GATA transcription factor | 1808 | 0 | 0 | 0 |
| SALT00000048666_Cluster10613-016 | GATA | GATA transcription factor | 1407 | 0 | 0 | 0 |
| SALT00000048723_Cluster10613-017 | GATA | GATA transcription factor | 1827 | 0 | 0 | 0 |
| SALT00000050333_Cluster10613-018 | GATA | GATA transcription factor | 1986 | 0 | 0 | 0 |
| SALT00000053007_Cluster10613-019 | GATA | GATA transcription factor | 1679 | 0 | 0 | 0 |
| SALT00000064943_Cluster10613-020 | GATA | GATA transcription factor | 3020 | 0 | 0 | 0 |
| SALT00000074815_Cluster10613-021 | GATA | GATA transcription factor | 2380 | 0 | 0 | 0 |
| SALT00000079088_Cluster10613-022 | GATA | GATA transcription factor | 2859 | 0 | 0 | 0 |
| SALT00000072218_Cluster10613-023 | GATA | GATA transcription factor | 2557 | 0 | 0 | 0 |
| SALT00000001230_Cluster11042-001 | GATA | GATA transcription factor | 1734 | 0 | 0 | 0 |
| SALT00000001918_Cluster11042-002 | GATA | GATA transcription factor | 1521 | 0 | 0 | 0 |
| SALT00000011521_Cluster11042-003 | GATA | [Hordeum vulgare vulgare] | 1645 | 0 | 0 | 0 |
| SALT00000035667_Cluster11042-004 | GATA | GATA transcription factor | 1617 | 0 | 0 | 0 |
| SALT00000053543_Cluster11042-005 | GATA | GATA transcription factor | 1578 | 0 | 0 | 0 |
| SALT00000056510_Cluster11042-006 | GATA | [Hordeum vulgare vulgare] | 1651 | 0 | 0 | 0 |
| SALT00000005450_Cluster11147-001 | GATA | GATA transcription factor | 2083 | 0 | 0 | 0 |
| SALT00000005946_Cluster11147-002 | GATA | GATA transcription factor | 2132 | 0 | down | 0 |
| SALT00000042991_Cluster11147-003 | GATA | GATA transcription factor | 1924 | 0 | 0 | 0 |
| SALT00000067595_Cluster11147-004 | GATA | GATA transcription factor | 2952 | 0 | 0 | 0 |
| SALT00000063018_Cluster11398-001 | GATA | GATA transcription factor | 2924 | 0 | 0 | 0 |
| SALT00000068702_Cluster13585-005 | GATA | transcription factor UNE1( | 2726 | 0 | 0 | 0 |
| SALT00000026082_Cluster15674-001 | GATA | GATA transcription factor | 2570 | 0 | up | up |
| SALT00000027185_Cluster15674-002 | GATA | GATA transcription factor | 2576 | 0 | 0 | 0 |

| | | | | | | |
|---|---|---|---|---|---|---|
| SALT00000013554_Cluster15674-003 | GATA | GATA transcription factor | 1952 | 0 | 0 | 0 |
| SALT00000001778_Cluster15742-001 | GATA | GATA zinc finger family [ | 1635 | 0 | 0 | 0 |
| SALT00000007291_Cluster15742-002 | GATA | GATA zinc finger family [ | 1504 | 0 | 0 | 0 |
| SALT00000021974_Cluster15742-003 | GATA | GATA zinc finger family [ | 1474 | 0 | 0 | 0 |
| SALT00000051052_Cluster15742-004 | GATA | GATA zinc finger family [ | 1412 | 0 | 0 | 0 |
| SALT00000051184_Cluster15742-005 | GATA | GATA zinc finger family [ | 1609 | 0 | 0 | down |
| SALT00000064813_Cluster15742-006 | GATA | GATA zinc finger family [ | 2570 | 0 | 0 | 0 |
| SALT00000055798_Cluster19159-001 | GATA | GATA transcription factor | 2153 | 0 | 0 | 0 |
| SALT00000001219_Cluster22126-001 | GATA | TPA: GATA transcription f | 1642 | 0 | 0 | 0 |
| SALT00000013247_Cluster22126-002 | GATA | TPA: GATA transcription f | 1789 | 0 | 0 | 0 |
| SALT00000013830_Cluster22126-003 | GATA | GATA zinc finger family [ | 1607 | 0 | 0 | 0 |
| SALT00000014269_Cluster22126-004 | GATA | TPA: GATA transcription f | 1930 | 0 | 0 | 0 |
| SALT00000039812_Cluster22126-005 | GATA | TPA: GATA transcription f | 1565 | 0 | 0 | 0 |
| SALT00000000867_Cluster22192-001 | GATA | GATA zinc finger family [ | 1652 | 0 | 0 | 0 |
| SALT00000017161_Cluster22192-002 | GATA | GATA zinc finger family [ | 1391 | 0 | 0 | 0 |
| SALT00000019348_Cluster22192-003 | GATA | GATA zinc finger family [ | 1570 | 0 | 0 | 0 |
| SALT00000039834_Cluster22192-004 | GATA | C2C2-GATA transcription | 1579 | 0 | 0 | 0 |
| SALT00000045544_Cluster22192-005 | GATA | GATA zinc finger family [ | 1927 | 0 | 0 | 0 |
| SALT00000050550_Cluster22192-006 | GATA | GATA zinc finger family [ | 1716 | 0 | 0 | 0 |
| SALT00000011952_Cluster22912-001 | GATA | zinc-finger [Oryza sativa J | 1883 | 0 | 0 | 0 |
| SALT00000011958_Cluster22912-002 | GATA | zinc-finger [Oryza sativa J | 1436 | 0 | 0 | 0 |
| SALT00000046182_Cluster22912-003 | GATA | zinc-finger [Oryza sativa J | 1537 | 0 | 0 | 0 |
| SALT00000046342_Cluster22912-004 | GATA | zinc-finger [Oryza sativa J | 1660 | 0 | 0 | 0 |
| SALT00000002660_Cluster24308-001 | GATA | AG-motif binding -4 [Ory | 1470 | 0 | 0 | 0 |
| SALT00000004625_Cluster24308-002 | GATA | AG-motif binding -4 [Ory | 1801 | 0 | 0 | 0 |
| SALT00000006242_Cluster24308-003 | GATA | C2C2-GATA transcription | 1714 | 0 | 0 | 0 |
| SALT00000036174_Cluster24308-004 | GATA | C2C2-GATA transcription | 1659 | 0 | 0 | 0 |
| SALT00000051711_Cluster24308-005 | GATA | GATA zinc finger family [ | 1725 | 0 | 0 | 0 |
| SALT00000015953_Cluster25016-001 | GATA | GATA zinc finger family [ | 1441 | 0 | 0 | up |
| SALT00000048741_Cluster25016-002 | GATA | GATA zinc finger family [ | 1571 | 0 | 0 | 0 |
| SALT00000056218_Cluster25016-003 | GATA | GATA zinc finger family [ | 1762 | 0 | 0 | 0 |
| SALT00000035795_Cluster25376-001 | GATA | C2C2-GATA transcription | 1743 | 0 | 0 | 0 |
| SALT00000037530_Cluster25376-002 | GATA | GATA zinc finger family [ | 1529 | 0 | 0 | 0 |
| SALT00000039854_Cluster25376-003 | GATA | C2C2-GATA transcription | 1653 | 0 | 0 | 0 |
| SALT00000045813_Cluster25376-004 | GATA | C2C2-GATA transcription | 1653 | 0 | 0 | 0 |
| SALT00000051043_Cluster25376-005 | GATA | GATA zinc finger family [ | 1602 | 0 | 0 | 0 |
| SALT00000052367_Cluster25376-006 | GATA | GATA transcription factor | 1725 | 0 | 0 | 0 |
| SALT00000052433_Cluster25376-007 | GATA | C2C2-GATA transcription | 1703 | 0 | 0 | 0 |
| SALT00000052532_Cluster25376-008 | GATA | GATA zinc finger family [ | 1501 | 0 | 0 | 0 |
| SALT00000054739_Cluster25376-009 | GATA | GATA transcription factor | 1684 | 0 | 0 | 0 |
| SALT00000059927_Cluster25376-010 | GATA | C2C2-GATA transcription | 1734 | 0 | 0 | 0 |
| SALT00000064993_Cluster25376-011 | GATA | GATA zinc finger family [ | 1681 | 0 | 0 | 0 |
| SALT00000001749_Cluster25663-001 | GATA | GATA transcription factor | 1722 | 0 | 0 | 0 |
| SALT00000047778_Cluster25663-002 | GATA | GATA transcription factor | 1729 | 0 | 0 | 0 |
| SALT00000016187_Cluster26601-001 | GATA | C2C2-GATA transcription | 1506 | 0 | 0 | 0 |
| SALT00000051164_Cluster26601-002 | GATA | GATA zinc finger family [ | 1680 | 0 | 0 | 0 |
| SALT00000009338_Cluster28358-001 | GATA | GATA transcription factor | 1451 | 0 | 0 | 0 |
| SALT00000015174_Cluster28358-002 | GATA | GATA transcription factor | 1489 | 0 | 0 | 0 |
| SALT00000043204_Cluster28358-003 | GATA | GATA transcription factor | 1479 | 0 | 0 | 0 |
| SALT00000056962_Cluster28358-004 | GATA | GATA transcription factor | 1592 | 0 | 0 | 0 |
| SALT00000007055_Cluster28805-001 | GATA | GATA transcription factor | 1513 | 0 | 0 | 0 |
| SALT00000036204_Cluster28805-002 | GATA | GATA transcription factor | 1568 | 0 | 0 | 0 |
| SALT00000012840_Cluster29128-001 | GATA | Os01g0745700 [Oryza sati | 1510 | 0 | 0 | 0 |
| SALT00000019327_Cluster29128-002 | GATA | Os01g0745700 [Oryza sati | 1552 | 0 | 0 | 0 |
| SALT00000055931_Cluster29128-003 | GATA | Os01g0745700 [Oryza sati | 1471 | 0 | 0 | 0 |
| SALT00000046996_Cluster29140-001 | GATA | Os03g0684000 [Oryza sati | 1552 | 0 | 0 | 0 |
| SALT00000009819_Cluster29327-001 | GATA | GATA zinc finger family [ | 1485 | 0 | 0 | 0 |
| SALT00000040881_Cluster29327-002 | GATA | GATA zinc finger family [ | 1542 | 0 | 0 | 0 |

| | | | | | | |
|---|---|---|---|---|---|---|
| SALT00000003083_Cluster29873-001 | GATA | GATA transcription factor | 1389 | 0 | 0 | 0 |
| SALT00000003841_Cluster29873-002 | GATA | GATA transcription factor | 1510 | 0 | down | down |
| SALT00000013579_Cluster29873-003 | GATA | GATA transcription factor | 1440 | 0 | 0 | down |
| SALT00000039590_Cluster29873-004 | GATA | GATA transcription factor | 1479 | 0 | 0 | 0 |
| SALT00000056906_Cluster29873-005 | GATA | GATA transcription factor | 1480 | 0 | 0 | 0 |
| SALT00000051352_Cluster30057-001 | GATA | GATA transcription factor | 1498 | 0 | 0 | 0 |
| SALT00000012829_Cluster30333-001 | GATA | C2C2-GATA transcription | 1479 | 0 | 0 | 0 |
| SALT00000006359_Cluster31125-001 | GATA | zinc-finger [Oryza sativa J | 1396 | 0 | 0 | 0 |
| SALT00000049387_Cluster31125-002 | GATA | zinc-finger [Oryza sativa J | 1351 | 0 | 0 | 0 |
| SALT00000059323_Cluster31125-003 | GATA | zinc-finger [Oryza sativa J | 1405 | 0 | 0 | 0 |
| SALT00000058625_Cluster31359-001 | GATA | Os03g0684000 [Oryza sati | 1364 | 0 | 0 | 0 |
| SALT00000053230_Cluster17346-002 | GeBP | Os02g0771200 [Oryza sati | 1561 | 0 | 0 | 0 |
| SALT00000058530_Cluster17346-003 | GeBP | Os02g0771200 [Oryza sati | 1617 | 0 | 0 | 0 |
| SALT00000039898_Cluster17346-004 | GeBP | Os02g0771200 [Oryza sati | 1674 | 0 | 0 | 0 |
| SALT00000014922_Cluster21096-001 | GeBP | hepatoma-derived growth | 1998 | 0 | 0 | 0 |
| SALT00000039076_Cluster21096-002 | GeBP | hepatoma-derived growth | 1826 | 0 | 0 | 0 |
| SALT00000055392_Cluster21096-003 | GeBP | hepatoma-derived growth | 1763 | 0 | 0 | 0 |
| SALT00000045767_Cluster21096-005 | GeBP | -type transcription partial [ | 2032 | 0 | 0 | 0 |
| SALT00000008864_Cluster22665-001 | GeBP | hepatoma-derived growth | 1833 | 0 | 0 | 0 |
| SALT00000016538_Cluster22665-002 | GeBP | hepatoma-derived growth | 1837 | 0 | 0 | 0 |
| SALT00000046811_Cluster22665-003 | GeBP | hepatoma-derived growth | 1899 | up | 0 | 0 |
| SALT00000011223_Cluster23810-001 | GeBP | serine arginine repetitive m | 1719 | 0 | 0 | 0 |
| SALT00000049354_Cluster23810-002 | GeBP | serine arginine repetitive m | 1608 | 0 | 0 | 0 |
| SALT00000059505_Cluster23810-003 | GeBP | serine arginine repetitive m | 1829 | 0 | 0 | 0 |
| SALT00000022662_Cluster24967-001 | GeBP | Os02g0771200 [Oryza sati | 1564 | 0 | 0 | 0 |
| SALT00000081023_Cluster24967-002 | GeBP | Os02g0771200 [Oryza sati | 1765 | 0 | 0 | 0 |
| SALT00000003946_Cluster25611-001 | GeBP | storekeeper [Oryza sativa J | 1595 | 0 | 0 | 0 |
| SALT00000013682_Cluster25611-002 | GeBP | storekeeper [Oryza sativa J | 1623 | 0 | 0 | 0 |
| SALT00000017953_Cluster25611-003 | GeBP | storekeeper [Oryza sativa J | 1608 | 0 | 0 | 0 |
| SALT00000055106_Cluster25611-004 | GeBP | storekeeper [Oryza sativa J | 1732 | 0 | 0 | 0 |
| SALT00000053573_Cluster25611-005 | GeBP | storekeeper [Oryza sativa J | 1715 | 0 | 0 | 0 |
| SALT00000019625_Cluster27868-001 | GeBP | transcription partial [Zea m | 1617 | 0 | 0 | 0 |
| SALT00000047980_Cluster28505-001 | GeBP | GBP18 type transcription p | 1584 | 0 | 0 | 0 |
| SALT00000052850_Cluster28818-001 | GeBP | [Hordeum vulgare vulgare] | 1568 | 0 | 0 | 0 |
| SALT00000009904_Cluster29597-001 | GeBP | [Hordeum vulgare vulgare] | 1526 | 0 | 0 | 0 |
| SALT00000000262_Cluster29730-001 | GeBP | Os02g0771200 [Oryza sati | 1518 | 0 | 0 | 0 |
| SALT00000000974_Cluster29853-001 | GeBP | GBP12 type transcription p | 1511 | 0 | 0 | 0 |
| SALT00000040396_Cluster30291-001 | GeBP | transcription partial [Zea m | 1482 | 0 | 0 | 0 |
| SALT00000012186_Cluster31320-001 | GeBP | GEBP transcription partial | 1372 | 0 | 0 | 0 |
| SALT00000062856_Cluster32115-001 | GeBP | myristoylated alanine-rich | 1146 | 0 | 0 | 0 |
| SALT00000008700_Cluster1174-001 | GRAS | scarecrow 9 [Oryza sativa | 2113 | 0 | 0 | 0 |
| SALT00000021365_Cluster1174-002 | GRAS | scarecrow 9 [Oryza sativa | 2920 | 0 | 0 | 0 |
| SALT00000024183_Cluster1174-003 | GRAS | scarecrow 9 [Oryza sativa | 3027 | 0 | 0 | 0 |
| SALT00000024890_Cluster1174-004 | GRAS | scarecrow 9 [Oryza sativa | 2812 | 0 | 0 | 0 |
| SALT00000028082_Cluster1174-005 | GRAS | scarecrow 9 [Oryza sativa | 2941 | 0 | 0 | 0 |
| SALT00000028103_Cluster1174-006 | GRAS | scarecrow 9 [Oryza sativa | 2880 | 0 | 0 | 0 |
| SALT00000038429_Cluster1174-007 | GRAS | scarecrow 9 [Oryza sativa | 1978 | 0 | 0 | 0 |
| SALT00000057659_Cluster1174-008 | GRAS | scarecrow 9 [Oryza sativa | 1956 | 0 | 0 | 0 |
| SALT00000059890_Cluster1174-009 | GRAS | scarecrow 9 [Oryza sativa | 1639 | 0 | 0 | 0 |
| SALT00000061843_Cluster1174-010 | GRAS | scarecrow 9 [Oryza sativa | 1961 | 0 | 0 | 0 |
| SALT00000064361_Cluster1174-011 | GRAS | scarecrow 9 [Oryza sativa | 3000 | 0 | 0 | 0 |
| SALT00000071569_Cluster1174-013 | GRAS | scarecrow 9 [Oryza sativa | 3054 | 0 | 0 | 0 |
| SALT00000071955_Cluster1174-014 | GRAS | scarecrow 9 [Oryza sativa | 2601 | 0 | 0 | 0 |
| SALT00000072383_Cluster1174-015 | GRAS | scarecrow 9 [Oryza sativa | 2961 | 0 | 0 | 0 |
| SALT00000074363_Cluster1174-016 | GRAS | scarecrow 9 [Oryza sativa | 3074 | 0 | 0 | 0 |
| SALT00000075982_Cluster1174-017 | GRAS | scarecrow 9 [Oryza sativa | 3058 | 0 | 0 | 0 |
| SALT00000077959_Cluster1174-018 | GRAS | scarecrow 9 [Oryza sativa | 3025 | 0 | 0 | 0 |
| SALT00000079956_Cluster1174-019 | GRAS | scarecrow 9 [Oryza sativa | 2936 | 0 | 0 | 0 |

| | | | | | | |
|---|---|---|---|---|---|---|
| SALT00000080650_Cluster1174-020 | GRAS | scarecrow 9 [Oryza sativa | 3164 | 0 | 0 | 0 |
| SALT00000082238_Cluster1174-021 | GRAS | scarecrow 9 [Oryza sativa | 3028 | 0 | 0 | 0 |
| SALT00000083255_Cluster1174-022 | GRAS | scarecrow 9 [Oryza sativa | 3021 | 0 | 0 | 0 |
| SALT00000021836_Cluster1376-002 | GRAS | scarecrow 27 | 2599 | 0 | 0 | 0 |
| SALT00000022014_Cluster1376-003 | GRAS | scarecrow 27 | 2728 | 0 | 0 | 0 |
| SALT00000022150_Cluster1376-004 | GRAS | Scl1 [Oryza sativa Japonic | 2650 | 0 | 0 | 0 |
| SALT00000022402_Cluster1376-005 | GRAS | Scl1 [Oryza sativa Japonic | 2742 | 0 | 0 | 0 |
| SALT00000022752_Cluster1376-006 | GRAS | scarecrow 27 | 2655 | 0 | 0 | up |
| SALT00000025885_Cluster1376-007 | GRAS | scarecrow 27 | 2566 | 0 | 0 | 0 |
| SALT00000028995_Cluster1376-008 | GRAS | scarecrow 27 | 2678 | 0 | 0 | 0 |
| SALT00000072931_Cluster1376-009 | GRAS | cytosolic orthophosphate d | 5819 | 0 | 0 | up |
| SALT00000073489_Cluster1376-010 | GRAS | Scl1 [Oryza sativa Japonic | 2683 | 0 | 0 | 0 |
| SALT00000075767_Cluster1376-011 | GRAS | scarecrow 27 | 2665 | 0 | 0 | 0 |
| SALT00000079084_Cluster1376-012 | GRAS | scarecrow 27 | 2583 | 0 | 0 | 0 |
| SALT00000002761_Cluster1557-001 | GRAS | scarecrow 9 | 1782 | 0 | 0 | 0 |
| SALT00000018313_Cluster1557-002 | GRAS | scarecrow 9 | 1988 | 0 | 0 | 0 |
| SALT00000021236_Cluster1557-003 | GRAS | scarecrow 9 | 3032 | 0 | 0 | 0 |
| SALT00000024400_Cluster1557-006 | GRAS | scarecrow 9 | 2975 | 0 | 0 | 0 |
| SALT00000025710_Cluster1557-008 | GRAS | scarecrow 9 | 2998 | down | down | down |
| SALT00000025991_Cluster1557-009 | GRAS | scarecrow 9 | 2642 | 0 | 0 | 0 |
| SALT00000026141_Cluster1557-010 | GRAS | scarecrow 9 | 2659 | 0 | 0 | 0 |
| SALT00000026852_Cluster1557-011 | GRAS | scarecrow 9 | 2750 | 0 | 0 | 0 |
| SALT00000027611_Cluster1557-013 | GRAS | scarecrow 9 | 2718 | 0 | 0 | 0 |
| SALT00000039023_Cluster1557-015 | GRAS | scarecrow 9 | 1749 | 0 | 0 | 0 |
| SALT00000044929_Cluster1557-017 | GRAS | scarecrow 9 | 1513 | 0 | 0 | 0 |
| SALT00000071014_Cluster1557-018 | GRAS | scarecrow 9 | 3121 | 0 | 0 | 0 |
| SALT00000071834_Cluster1557-019 | GRAS | STRUBBELIG-RECEPTC | 5550 | 0 | 0 | 0 |
| SALT00000026115_Cluster1557-027 | GRAS | scarecrow 9 | 2590 | 0 | 0 | 0 |
| SALT00000015564_Cluster1795-001 | GRAS | scarecrow 34 | 1650 | 0 | 0 | 0 |
| SALT00000018133_Cluster1795-002 | GRAS | scarecrow 34 | 1628 | 0 | 0 | 0 |
| SALT00000020869_Cluster1795-003 | GRAS | scarecrow 34 | 2578 | 0 | 0 | 0 |
| SALT00000020947_Cluster1795-004 | GRAS | scarecrow 34 | 2628 | 0 | 0 | 0 |
| SALT00000021284_Cluster1795-005 | GRAS | scarecrow 34 | 2802 | 0 | 0 | 0 |
| SALT00000024081_Cluster1795-006 | GRAS | scarecrow 34 | 2572 | 0 | 0 | 0 |
| SALT00000024429_Cluster1795-007 | GRAS | scarecrow 34 | 2536 | 0 | 0 | 0 |
| SALT00000024969_Cluster1795-008 | GRAS | scarecrow 34 | 2766 | 0 | 0 | 0 |
| SALT00000029165_Cluster1795-010 | GRAS | scarecrow 34 | 2783 | 0 | 0 | 0 |
| SALT00000061962_Cluster1795-012 | GRAS | Os11g0705200 [Oryza sati | 2499 | 0 | 0 | 0 |
| SALT00000065236_Cluster1795-013 | GRAS | scarecrow 34 | 2669 | 0 | 0 | 0 |
| SALT00000077072_Cluster1795-014 | GRAS | V-type proton ATPase subu | 5312 | 0 | 0 | 0 |
| SALT00000081398_Cluster1795-015 | GRAS | scarecrow 34 | 2638 | 0 | 0 | 0 |
| SALT00000087898_Cluster1795-016 | GRAS | scarecrow 34 | 3111 | 0 | 0 | 0 |
| SALT00000008390_Cluster1836-001 | GRAS | Os12g0573200 [Oryza sati | 1641 | 0 | 0 | 0 |
| SALT00000018884_Cluster1836-002 | GRAS | Os12g0573200 [Oryza sati | 1550 | 0 | 0 | 0 |
| SALT00000022476_Cluster1836-003 | GRAS | Os12g0573200 [Oryza sati | 2687 | 0 | 0 | 0 |
| SALT00000022767_Cluster1836-004 | GRAS | Os12g0573200 [Oryza sati | 2612 | 0 | 0 | 0 |
| SALT00000023114_Cluster1836-005 | GRAS | Os12g0573200 [Oryza sati | 2561 | 0 | 0 | 0 |
| SALT00000023303_Cluster1836-006 | GRAS | Os12g0573200 [Oryza sati | 2662 | down | down | down |
| SALT00000023739_Cluster1836-007 | GRAS | Os12g0573200 [Oryza sati | 2455 | 0 | 0 | 0 |
| SALT00000030559_Cluster1836-008 | GRAS | Os12g0573200 [Oryza sati | 2273 | 0 | 0 | 0 |
| SALT00000051564_Cluster1836-009 | GRAS | Os12g0573200 [Oryza sati | 2229 | 0 | 0 | 0 |
| SALT00000058472_Cluster1836-010 | GRAS | Os12g0573200 [Oryza sati | 1499 | 0 | 0 | 0 |
| SALT00000062807_Cluster1836-011 | GRAS | Os12g0573200 [Oryza sati | 5276 | 0 | 0 | 0 |
| SALT00000068067_Cluster1836-012 | GRAS | Os12g0573200 [Oryza sati | 2581 | 0 | 0 | 0 |
| SALT00000073415_Cluster1836-013 | GRAS | Os12g0573200 [Oryza sati | 2568 | 0 | 0 | 0 |
| SALT00000075885_Cluster1836-014 | GRAS | Os12g0573200 [Oryza sati | 2725 | 0 | 0 | 0 |
| SALT00000081985_Cluster1836-015 | GRAS | Os12g0573200 [Oryza sati | 2677 | 0 | 0 | 0 |
| SALT00000003973_Cluster1903-001 | GRAS | chitin-inducible gibberellir | 2139 | 0 | 0 | 0 |

| | | | | | | |
|---|---|---|---|---|---|---|
| SALT00000004779_Cluster1903-002 | GRAS | chitin-inducible gibberellir | 2236 | 0 | 0 | 0 |
| SALT00000013362_Cluster1903-003 | GRAS | chitin-inducible gibberellir | 2269 | 0 | 0 | 0 |
| SALT00000016764_Cluster1903-004 | GRAS | chitin-inducible gibberellir | 1945 | 0 | 0 | 0 |
| SALT00000018241_Cluster1903-005 | GRAS | scarecrow 1 [Setaria italica | 1550 | 0 | 0 | 0 |
| SALT00000037188_Cluster1903-006 | GRAS | chitin-inducible gibberellir | 2280 | 0 | 0 | 0 |
| SALT00000052110_Cluster1903-007 | GRAS | chitin-inducible gibberellir | 2129 | 0 | 0 | 0 |
| SALT00000056753_Cluster1903-008 | GRAS | chitin-inducible gibberellir | 2146 | 0 | 0 | 0 |
| SALT00000062400_Cluster1903-009 | GRAS | scarecrow 1 [Setaria italica | 1761 | 0 | 0 | 0 |
| SALT00000057631_Cluster1903-012 | GRAS | chitin-inducible gibberellir | 1990 | 0 | 0 | 0 |
| SALT00000021020_Cluster2129-001 | GRAS | Os04g0590400 [Oryza sati | 2506 | 0 | 0 | 0 |
| SALT00000060848_Cluster2129-002 | GRAS | Os04g0590400 [Oryza sati | 2141 | 0 | 0 | 0 |
| SALT00000064636_Cluster2129-003 | GRAS | Os04g0590400 [Oryza sati | 2666 | 0 | 0 | 0 |
| SALT00000070465_Cluster2129-004 | GRAS | Os04g0590400 [Oryza sati | 2718 | 0 | 0 | 0 |
| SALT00000093618_Cluster2129-006 | GRAS | Os04g0590400 [Oryza sati | 2531 | 0 | 0 | 0 |
| SALT00000068213_Cluster3567-001 | GRAS | Os01g0621900 [Oryza sati | 4369 | 0 | 0 | 0 |
| SALT00000021067_Cluster4592-001 | GRAS | scarecrow 9 [Oryza sativa | 3000 | 0 | 0 | 0 |
| SALT00000025194_Cluster4592-002 | GRAS | scarecrow 9 [Oryza sativa | 2880 | 0 | 0 | 0 |
| SALT00000027239_Cluster4592-003 | GRAS | scarecrow 9 [Oryza sativa | 3310 | 0 | 0 | 0 |
| SALT00000028360_Cluster4592-004 | GRAS | scarecrow 9 [Oryza sativa | 2878 | 0 | 0 | 0 |
| SALT00000030267_Cluster4592-005 | GRAS | scarecrow 9 [Oryza sativa | 3087 | 0 | 0 | 0 |
| SALT00000030406_Cluster4592-006 | GRAS | scarecrow 9 [Oryza sativa | 2790 | 0 | 0 | 0 |
| SALT00000053896_Cluster4592-007 | GRAS | scarecrow 9 [Oryza sativa | 1648 | 0 | 0 | 0 |
| SALT00000061986_Cluster4592-008 | GRAS | scarecrow 9 [Oryza sativa | 2965 | 0 | 0 | 0 |
| SALT00000063391_Cluster4592-009 | GRAS | scarecrow 9 [Oryza sativa | 3161 | 0 | 0 | 0 |
| SALT00000064143_Cluster4592-010 | GRAS | scarecrow 9 [Oryza sativa | 2502 | 0 | 0 | 0 |
| SALT00000066430_Cluster4592-011 | GRAS | scarecrow 9 [Oryza sativa | 2744 | 0 | 0 | 0 |
| SALT00000067361_Cluster4592-012 | GRAS | scarecrow 9 [Oryza sativa | 2918 | 0 | 0 | 0 |
| SALT00000071339_Cluster4592-013 | GRAS | scarecrow 9 [Oryza sativa | 2972 | 0 | 0 | 0 |
| SALT00000085765_Cluster4592-014 | GRAS | scarecrow 9 [Oryza sativa | 3067 | 0 | 0 | 0 |
| SALT00000089508_Cluster4592-015 | GRAS | scarecrow 9 [Oryza sativa | 4065 | 0 | 0 | 0 |
| SALT00000008855_Cluster5972-001 | GRAS | scarecrow 21 [Oryza sativa | 2358 | 0 | 0 | 0 |
| SALT00000021522_Cluster5972-002 | GRAS | scarecrow 21 [Oryza sativa | 2467 | 0 | 0 | 0 |
| SALT00000023964_Cluster5972-003 | GRAS | scarecrow 21 [Oryza sativa | 2427 | 0 | 0 | 0 |
| SALT00000027841_Cluster5972-004 | GRAS | scarecrow 21 [Oryza sativa | 2431 | 0 | 0 | 0 |
| SALT00000058814_Cluster5972-005 | GRAS | scarecrow 21 [Oryza sativa | 2188 | 0 | 0 | 0 |
| SALT00000063458_Cluster5972-006 | GRAS | scarecrow 21 [Oryza sativa | 2750 | 0 | 0 | 0 |
| SALT00000068629_Cluster5972-007 | GRAS | scarecrow 21 [Oryza sativa | 2348 | 0 | 0 | 0 |
| SALT00000068837_Cluster5972-008 | GRAS | scarecrow 21 [Oryza sativa | 2683 | 0 | 0 | 0 |
| SALT00000075912_Cluster5972-009 | GRAS | scarecrow 21 [Oryza sativa | 2510 | 0 | 0 | 0 |
| SALT00000080256_Cluster5972-010 | GRAS | scarecrow 21 [Oryza sativa | 2416 | 0 | 0 | 0 |
| SALT00000081869_Cluster5972-011 | GRAS | scarecrow 21 [Oryza sativa | 3071 | 0 | 0 | 0 |
| SALT00000088540_Cluster5972-012 | GRAS | scarecrow 21 [Oryza sativa | 3797 | 0 | 0 | 0 |
| SALT00000057644_Cluster5972-013 | GRAS | scarecrow 21 [Oryza sativa | 1964 | 0 | 0 | 0 |
| SALT00000034030_Cluster5972-014 | GRAS | scarecrow 21 [Oryza sativa | 4024 | 0 | 0 | 0 |
| SALT00000068250_Cluster5972-015 | GRAS | scarecrow 21 [Oryza sativa | 2531 | 0 | 0 | 0 |
| SALT00000008956_Cluster7039-001 | GRAS | scarecrow 21 [Brachypodiu | 1571 | 0 | 0 | 0 |
| SALT00000031093_Cluster7039-002 | GRAS | scarecrow 21 [Brachypodiu | 2160 | 0 | 0 | 0 |
| SALT00000031348_Cluster7039-003 | GRAS | scarecrow 21 [Brachypodiu | 2122 | 0 | 0 | 0 |
| SALT00000032981_Cluster7039-004 | GRAS | scarecrow 21 [Brachypodiu | 3381 | 0 | 0 | 0 |
| SALT00000035405_Cluster7039-005 | GRAS | scarecrow 21 [Brachypodiu | 2031 | 0 | 0 | 0 |
| SALT00000036852_Cluster7039-006 | GRAS | scarecrow 21 [Brachypodiu | 1584 | 0 | 0 | 0 |
| SALT00000050579_Cluster7039-007 | GRAS | scarecrow 21 [Brachypodiu | 2312 | 0 | 0 | 0 |
| SALT00000051174_Cluster7039-008 | GRAS | scarecrow 21 [Brachypodiu | 2076 | 0 | 0 | 0 |
| SALT00000060173_Cluster7039-009 | GRAS | scarecrow 21 [Brachypodiu | 2176 | 0 | 0 | 0 |
| SALT00000060580_Cluster7039-010 | GRAS | scarecrow 21 [Brachypodiu | 2256 | 0 | 0 | 0 |
| SALT00000085972_Cluster7039-011 | GRAS | scarecrow 21 [Brachypodiu | 3631 | 0 | 0 | 0 |
| SALT00000063178_Cluster7039-012 | GRAS | scarecrow 21 [Brachypodiu | 3114 | 0 | 0 | 0 |
| SALT00000023369_Cluster8659-001 | GRAS | Os12g0573200 [Oryza sati | 2407 | 0 | 0 | 0 |

| | | | | | | |
|---|---|---|---|---|---|---|
| SALT00000071714_Cluster8659-002 | GRAS | Os12g0573200 [Oryza sati | 3352 | 0 | 0 | 0 |
| SALT00000082874_Cluster8736-001 | GRAS | Os10g0551200 [Oryza sati | 3340 | 0 | 0 | 0 |
| SALT00000004998_Cluster8746-001 | GRAS | chitin-inducible gibberellin | 2178 | 0 | 0 | 0 |
| SALT00000012259_Cluster8746-002 | GRAS | chitin-inducible gibberellin | 1964 | 0 | 0 | 0 |
| SALT00000014291_Cluster8746-003 | GRAS | chitin-inducible gibberellin | 2177 | 0 | 0 | 0 |
| SALT00000016450_Cluster8746-004 | GRAS | chitin-inducible gibberellin | 1764 | 0 | 0 | 0 |
| SALT00000042143_Cluster8746-005 | GRAS | chitin-inducible gibberellin | 2207 | 0 | 0 | 0 |
| SALT00000059717_Cluster8746-006 | GRAS | chitin-inducible gibberellin | 1324 | down | 0 | down |
| SALT00000070266_Cluster8746-007 | GRAS | chitin-inducible gibberellin | 3338 | 0 | 0 | 0 |
| SALT00000081112_Cluster8746-008 | GRAS | chitin-inducible gibberellin | 2206 | 0 | 0 | 0 |
| SALT00000005611_Cluster10985-001 | GRAS | reduced height-2 [Eragrost | 2088 | 0 | 0 | 0 |
| SALT00000016855_Cluster10985-002 | GRAS | reduced height-2 [Eragrost | 1676 | 0 | 0 | 0 |
| SALT00000020809_Cluster10985-003 | GRAS | reduced height-2 [Eragrost | 2470 | 0 | 0 | 0 |
| SALT00000021882_Cluster10985-004 | GRAS | reduced height-2 [Eragrost | 2505 | 0 | 0 | 0 |
| SALT00000024007_Cluster10985-005 | GRAS | reduced height-2 [Eragrost | 2626 | 0 | 0 | down |
| SALT00000024414_Cluster10985-006 | GRAS | reduced height-2 [Eragrost | 2513 | 0 | 0 | 0 |
| SALT00000025417_Cluster10985-007 | GRAS | reduced height-2 [Eragrost | 2596 | 0 | 0 | 0 |
| SALT00000027020_Cluster10985-008 | GRAS | reduced height-2 [Eragrost | 2447 | 0 | 0 | 0 |
| SALT00000028230_Cluster10985-009 | GRAS | reduced height-2 [Eragrost | 2324 | 0 | 0 | 0 |
| SALT00000028700_Cluster10985-010 | GRAS | reduced height-2 [Eragrost | 2498 | 0 | 0 | 0 |
| SALT00000030433_Cluster10985-011 | GRAS | reduced height-2 [Eragrost | 2416 | 0 | up | up |
| SALT00000038944_Cluster10985-012 | GRAS | reduced height-2 [Eragrost | 2097 | 0 | 0 | 0 |
| SALT00000072029_Cluster10985-014 | GRAS | reduced height-2 [Eragrost | 2500 | 0 | 0 | down |
| SALT00000072831_Cluster10985-015 | GRAS | reduced height-2 [Eragrost | 2473 | 0 | 0 | 0 |
| SALT00000074507_Cluster10985-016 | GRAS | reduced height-2 [Eragrost | 2509 | 0 | 0 | 0 |
| SALT00000024229_Cluster11052-001 | GRAS | scarecrow 9 | 2892 | 0 | 0 | 0 |
| SALT00000025335_Cluster11052-002 | GRAS | scarecrow 9 | 2653 | 0 | 0 | 0 |
| SALT00000025616_Cluster11052-003 | GRAS | scarecrow 9 | 2928 | 0 | 0 | 0 |
| SALT00000026088_Cluster11052-004 | GRAS | scarecrow 9 | 2934 | 0 | 0 | 0 |
| SALT00000052822_Cluster11052-005 | GRAS | scarecrow 9 | 2004 | 0 | 0 | 0 |
| SALT00000063351_Cluster11052-006 | GRAS | scarecrow 9 | 2615 | 0 | 0 | 0 |
| SALT00000071175_Cluster11052-007 | GRAS | scarecrow 9 | 2925 | 0 | 0 | 0 |
| SALT00000072110_Cluster11052-008 | GRAS | scarecrow 9 | 2643 | 0 | 0 | 0 |
| SALT00000073863_Cluster11052-009 | GRAS | scarecrow 9 | 2965 | 0 | 0 | 0 |
| SALT00000074104_Cluster11052-010 | GRAS | scarecrow 9 | 2542 | 0 | 0 | 0 |
| SALT00000074865_Cluster11052-011 | GRAS | scarecrow 9 | 2675 | 0 | 0 | 0 |
| SALT00000075568_Cluster11052-012 | GRAS | scarecrow 9 | 2688 | 0 | 0 | 0 |
| SALT00000077678_Cluster11052-013 | GRAS | scarecrow 9 | 2882 | 0 | 0 | 0 |
| SALT00000078177_Cluster11052-014 | GRAS | scarecrow 9 | 2766 | 0 | 0 | 0 |
| SALT00000082531_Cluster11052-015 | GRAS | scarecrow 9 | 2582 | 0 | 0 | 0 |
| SALT00000083282_Cluster11052-016 | GRAS | Os01g0842200 [Oryza sati | 2840 | 0 | 0 | 0 |
| SALT00000024313_Cluster11322-001 | GRAS | Os12g0573200 [Oryza sati | 2495 | 0 | 0 | 0 |
| SALT00000074208_Cluster11322-002 | GRAS | Os12g0573200 [Oryza sati | 2933 | 0 | 0 | up |
| SALT00000050409_Cluster11327-001 | GRAS | catalase [Saccharum hybri | 2932 | 0 | 0 | 0 |
| SALT00000074177_Cluster11829-001 | GRAS | scarecrow 28 [Oryza sativa | 2654 | 0 | 0 | 0 |
| SALT00000080380_Cluster11829-002 | GRAS | scarecrow 28 [Oryza sativa | 2884 | 0 | 0 | 0 |
| SALT00000003885_Cluster11983-001 | GRAS | SCARECROW [Zea mays | 2132 | 0 | 0 | 0 |
| SALT00000023882_Cluster11983-002 | GRAS | SCARECROW [Zea mays | 2012 | 0 | 0 | 0 |
| SALT00000038639_Cluster11983-003 | GRAS | SCARECROW [Zea mays | 1941 | 0 | 0 | 0 |
| SALT00000041113_Cluster11983-004 | GRAS | SCARECROW [Zea mays | 1741 | 0 | 0 | down |
| SALT00000050407_Cluster11983-005 | GRAS | SCARECROW [Zea mays | 2156 | 0 | 0 | 0 |
| SALT00000079364_Cluster11983-006 | GRAS | SCARECROW [Zea mays | 2870 | 0 | 0 | 0 |
| SALT00000003473_Cluster11983-007 | GRAS | SCARECROW [Zea mays | 2266 | 0 | 0 | 0 |
| SALT00000002807_Cluster12461-001 | GRAS | GRAS transcription partial | 1876 | 0 | 0 | down |
| SALT00000021369_Cluster12461-002 | GRAS | GRAS transcription partial | 2703 | 0 | down | down |
| SALT00000022110_Cluster12461-003 | GRAS | GRAS transcription partial | 2635 | 0 | 0 | down |
| SALT00000023213_Cluster12461-004 | GRAS | GRAS transcription partial | 2643 | 0 | 0 | 0 |
| SALT00000024560_Cluster12461-005 | GRAS | GRAS transcription partial | 2540 | 0 | 0 | 0 |

| | | | | | | |
|---|---|---|---|---|---|---|
| SALT00000060475_Cluster12461-006 | GRAS | GRAS transcription partial | 1887 | 0 | 0 | 0 |
| SALT00000064776_Cluster12461-007 | GRAS | GRAS transcription partial | 2823 | 0 | 0 | 0 |
| SALT00000065347_Cluster12461-008 | GRAS | GRAS transcription partial | 2680 | 0 | 0 | 0 |
| SALT00000070091_Cluster12461-009 | GRAS | GRAS transcription partial | 2655 | 0 | 0 | 0 |
| SALT00000077427_Cluster12461-010 | GRAS | GRAS transcription partial | 2573 | 0 | down | down |
| SALT00000083403_Cluster12461-011 | GRAS | GRAS transcription partial | 2525 | 0 | 0 | down |
| SALT00000044780_Cluster13439-001 | GRAS | chitin-inducible gibberellin | 2055 | 0 | 0 | 0 |
| SALT00000071697_Cluster13439-002 | GRAS | chitin-inducible gibberellin | 2737 | 0 | 0 | 0 |
| SALT00000084002_Cluster13439-003 | GRAS | chitin-inducible gibberellin | 2183 | 0 | 0 | 0 |
| SALT00000013357_Cluster13439-006 | GRAS | chitin-inducible gibberellin | 2053 | 0 | 0 | 0 |
| SALT00000072000_Cluster13600-001 | GRAS | GRAS transcription partial | 2603 | 0 | 0 | 0 |
| SALT00000073295_Cluster13600-002 | GRAS | GRAS transcription partial | 2725 | 0 | 0 | 0 |
| SALT00000080141_Cluster13600-003 | GRAS | GRAS transcription partial | 2687 | 0 | 0 | 0 |
| SALT00000082858_Cluster13600-004 | GRAS | GRAS transcription partial | 2643 | 0 | 0 | 0 |
| SALT00000026572_Cluster14074-001 | GRAS | Os04g0590400 [Oryza sati | 2579 | 0 | 0 | 0 |
| SALT00000028990_Cluster14074-002 | GRAS | Os04g0590400 [Oryza sati | 2689 | 0 | 0 | 0 |
| SALT00000031288_Cluster14074-003 | GRAS | Os04g0590400 [Oryza sati | 2499 | 0 | 0 | 0 |
| SALT00000062820_Cluster14074-004 | GRAS | Os04g0590400 [Oryza sati | 2578 | 0 | 0 | 0 |
| SALT00000068527_Cluster14074-005 | GRAS | Os04g0590400 [Oryza sati | 2339 | 0 | 0 | 0 |
| SALT00000072840_Cluster14074-006 | GRAS | Os04g0590400 [Oryza sati | 2397 | 0 | 0 | 0 |
| SALT00000074638_Cluster14074-007 | GRAS | Os04g0590400 [Oryza sati | 2506 | 0 | 0 | up |
| SALT00000081764_Cluster14074-008 | GRAS | Os04g0590400 [Oryza sati | 2544 | 0 | 0 | 0 |
| SALT00000073172_Cluster14285-001 | GRAS | reduced height-2 [Eragrost | 2673 | 0 | 0 | 0 |
| SALT00000012478_Cluster14386-001 | GRAS | SCARECROW [Zea mays | 2174 | 0 | 0 | 0 |
| SALT00000014348_Cluster14386-002 | GRAS | Os12g0122000 [Oryza sati | 2076 | 0 | 0 | 0 |
| SALT00000049408_Cluster14386-003 | GRAS | SCARECROW [Zea mays | 2258 | 0 | 0 | 0 |
| SALT00000070876_Cluster14386-004 | GRAS | SCARECROW [Zea mays | 2666 | 0 | 0 | 0 |
| SALT00000078683_Cluster14386-005 | GRAS | SCARECROW [Zea mays | 2649 | 0 | 0 | 0 |
| SALT00000023523_Cluster15152-001 | GRAS | Os12g0573200 [Oryza sati | 2614 | 0 | 0 | 0 |
| SALT00000062919_Cluster15152-002 | GRAS | Os12g0573200 [Oryza sati | 2533 | 0 | 0 | 0 |
| SALT00000068099_Cluster15152-003 | GRAS | Os12g0573200 [Oryza sati | 2613 | 0 | 0 | 0 |
| SALT00000025707_Cluster15449-001 | GRAS | Os10g0551200 [Oryza sati | 2454 | 0 | 0 | 0 |
| SALT00000036235_Cluster15449-002 | GRAS | Os10g0551200 [Oryza sati | 1790 | 0 | 0 | 0 |
| SALT00000038563_Cluster15449-003 | GRAS | Os10g0551200 [Oryza sati | 1287 | 0 | 0 | 0 |
| SALT00000053820_Cluster15449-004 | GRAS | scarecrow 6 [Brachypodiu | 1912 | 0 | 0 | 0 |
| SALT00000084628_Cluster15449-005 | GRAS | Os10g0551200 [Oryza sati | 2594 | 0 | 0 | 0 |
| SALT00000040363_Cluster15940-001 | GRAS | chitin-inducible gibberellin | 2556 | 0 | 0 | 0 |
| SALT00000024401_Cluster16371-001 | GRAS | scarecrow 6 [Brachypodiu | 2490 | 0 | 0 | 0 |
| SALT00000070939_Cluster16371-002 | GRAS | scarecrow 6 [Brachypodiu | 2221 | 0 | 0 | 0 |
| SALT00000083792_Cluster16371-003 | GRAS | scarecrow 6 [Brachypodiu | 2526 | 0 | 0 | 0 |
| SALT00000024381_Cluster16665-001 | GRAS | GRAS transcription partial | 2500 | 0 | 0 | 0 |
| SALT00000046664_Cluster16665-002 | GRAS | GRAS transcription partial | 1788 | 0 | 0 | 0 |
| SALT00000051283_Cluster16665-003 | GRAS | GRAS transcription partial | 2339 | 0 | 0 | 0 |
| SALT00000058079_Cluster16665-004 | GRAS | GRAS transcription partial | 2206 | 0 | 0 | 0 |
| SALT00000082381_Cluster16665-005 | GRAS | GRAS transcription partial | 2453 | 0 | 0 | 0 |
| SALT00000036245_Cluster17447-001 | GRAS | scarecrow 1 [Setaria italica | 1446 | 0 | 0 | 0 |
| SALT00000049248_Cluster17447-002 | GRAS | chitin-inducible gibberellin | 2415 | 0 | 0 | 0 |
| SALT00000006581_Cluster18162-001 | GRAS | scarecrow 8 [Setaria italica | 2136 | 0 | 0 | 0 |
| SALT00000007917_Cluster18162-002 | GRAS | scarecrow 8 [Oryza sativa | 2200 | 0 | 0 | 0 |
| SALT00000013281_Cluster18162-003 | GRAS | scarecrow 8 [Setaria italica | 2097 | 0 | 0 | 0 |
| SALT00000037609_Cluster18162-004 | GRAS | scarecrow 8 [Setaria italica | 1770 | 0 | 0 | 0 |
| SALT00000039271_Cluster18162-005 | GRAS | Os04g0580300 [Oryza sati | 1503 | 0 | 0 | 0 |
| SALT00000049669_Cluster18162-006 | GRAS | scarecrow 8 [Oryza sativa | 2181 | 0 | 0 | 0 |
| SALT00000055522_Cluster18162-007 | GRAS | scarecrow 8 [Setaria italica | 2085 | 0 | 0 | 0 |
| SALT00000060607_Cluster18162-008 | GRAS | scarecrow 8 [Oryza sativa | 2270 | 0 | 0 | 0 |
| SALT00000061592_Cluster18162-009 | GRAS | scarecrow 8 [Setaria italica | 2113 | 0 | 0 | down |
| SALT00000066481_Cluster18162-010 | GRAS | scarecrow 8 [Oryza sativa | 2109 | 0 | 0 | 0 |
| SALT00000050276_Cluster18308-001 | GRAS | partial [Oryza sativa Japon | 2247 | 0 | 0 | 0 |

| | | | | | | |
|---|---|---|---|---|---|---|
| SALT00000004533_Cluster18361-001 | GRAS | chitin-inducible gibberellin | 2176 | 0 | 0 | 0 |
| SALT00000017466_Cluster18361-002 | GRAS | chitin-inducible gibberellin | 2238 | 0 | 0 | 0 |
| SALT00000020467_Cluster18361-003 | GRAS | chitin-inducible gibberellin | 2077 | 0 | 0 | 0 |
| SALT00000023199_Cluster18361-004 | GRAS | chitin-inducible gibberellin | 2068 | 0 | 0 | 0 |
| SALT00000034869_Cluster18361-005 | GRAS | chitin-inducible gibberellin | 2061 | 0 | 0 | 0 |
| SALT00000035491_Cluster18361-006 | GRAS | chitin-inducible gibberellin | 2018 | 0 | 0 | 0 |
| SALT00000036025_Cluster18361-007 | GRAS | chitin-inducible gibberellin | 2142 | 0 | 0 | 0 |
| SALT00000036315_Cluster18361-008 | GRAS | chitin-inducible gibberellin | 1554 | 0 | 0 | 0 |
| SALT00000040563_Cluster18361-009 | GRAS | chitin-inducible gibberellin | 2102 | 0 | 0 | 0 |
| SALT00000044589_Cluster18361-010 | GRAS | chitin-inducible gibberellin | 2079 | 0 | 0 | 0 |
| SALT00000053148_Cluster18361-011 | GRAS | [Hordeum vulgare vulgare] | 1742 | 0 | 0 | 0 |
| SALT00000053289_Cluster18361-012 | GRAS | chitin-inducible gibberellin | 1721 | 0 | 0 | 0 |
| SALT00000058540_Cluster18361-013 | GRAS | chitin-inducible gibberellin | 2086 | 0 | 0 | 0 |
| SALT00000037786_Cluster18483-001 | GRAS | chitin-inducible gibberellin | 1554 | 0 | 0 | 0 |
| SALT00000056208_Cluster18483-002 | GRAS | chitin-inducible gibberellin | 1712 | 0 | 0 | 0 |
| SALT00000058440_Cluster18483-003 | GRAS | chitin-inducible gibberellin | 2222 | 0 | 0 | 0 |
| SALT00000013765_Cluster19392-001 | GRAS | scarecrow 4 [Zea mays] | 1912 | 0 | 0 | 0 |
| SALT00000019477_Cluster19392-002 | GRAS | scarecrow 4 [Zea mays] | 2097 | 0 | 0 | 0 |
| SALT00000036477_Cluster19392-003 | GRAS | scarecrow 4 [Zea mays] | 1846 | 0 | 0 | 0 |
| SALT00000059356_Cluster19392-004 | GRAS | scarecrow 4 [Zea mays] | 2023 | 0 | 0 | 0 |
| SALT00000062473_Cluster19392-005 | GRAS | scarecrow 4 [Zea mays] | 2129 | 0 | 0 | 0 |
| SALT00000010603_Cluster22593-001 | GRAS | scarecrow 23 [Zea mays] | 1903 | 0 | 0 | 0 |
| SALT00000049077_Cluster23500-001 | GRAS | GRAS transcription partial | 1847 | 0 | 0 | 0 |
| SALT00000000312_Cluster24593-001 | GRAS | Os01g0646300 [Oryza sati | 1643 | 0 | 0 | 0 |
| SALT00000006643_Cluster24593-002 | GRAS | Os01g0646300 [Oryza sati | 1625 | 0 | 0 | 0 |
| SALT00000011182_Cluster24593-003 | GRAS | Os01g0646300 [Oryza sati | 1610 | 0 | 0 | 0 |
| SALT00000016591_Cluster24593-004 | GRAS | Os01g0646300 [Oryza sati | 1569 | 0 | 0 | 0 |
| SALT00000055326_Cluster24593-005 | GRAS | Os01g0646300 [Oryza sati | 1785 | 0 | 0 | 0 |
| SALT00000060520_Cluster24593-006 | GRAS | Os01g0646300 [Oryza sati | 1667 | 0 | 0 | 0 |
| SALT00000004490_Cluster24593-007 | GRAS | Os01g0646300 [Oryza sati | 1678 | 0 | 0 | 0 |
| SALT00000041242_Cluster29012-001 | GRAS | gibberellin response modu | 1558 | 0 | 0 | 0 |
| SALT00000066382_Cluster29910-001 | GRAS | scarecrow 23 | 1508 | 0 | 0 | 0 |
| SALT00000069271_Cluster14513-001 | GRF | growth-regulating factor 2 | 2658 | 0 | 0 | 0 |
| SALT00000037754_Cluster19599-001 | GRF | Os03g0729500 [Oryza sati | 2112 | 0 | 0 | 0 |
| SALT00000060939_Cluster19599-003 | GRF | growth-regulating factor 6- | 2098 | 0 | 0 | 0 |
| SALT00000045503_Cluster23750-001 | GRF | growth-regulating factor 5- | 1832 | 0 | 0 | 0 |
| SALT00000048246_Cluster23750-002 | GRF | growth-regulating factor 5- | 1567 | 0 | 0 | down |
| SALT00000001624_Cluster27169-001 | GRF | growth-regulating factor 3- | 1592 | 0 | 0 | 0 |
| SALT00000024315_Cluster27169-002 | GRF | growth-regulating factor 3- | 1652 | 0 | 0 | 0 |
| SALT00000067198_Cluster27169-003 | GRF | growth-regulating factor 3- | 1427 | 0 | 0 | 0 |
| SALT00000052056_Cluster27228-001 | GRF | growth-regulating factor | 1650 | 0 | 0 | 0 |
| SALT00000061600_Cluster27228-002 | GRF | growth-regulating factor | 1387 | 0 | 0 | 0 |
| SALT00000013989_Cluster31213-001 | GRF | growth-regulating factor 1( | 1310 | 0 | 0 | 0 |
| SALT00000026227_Cluster31213-002 | GRF | growth-regulating factor 1( | 1102 | 0 | 0 | 0 |
| SALT00000039756_Cluster31213-003 | GRF | growth-regulating factor 1( | 1391 | 0 | 0 | 0 |
| SALT00000058403_Cluster31213-004 | GRF | growth-regulating factor 1( | 1349 | 0 | 0 | 0 |
| SALT00000030919_Cluster598-004 | HB-other | PREDICTED: uncharacter | 3989 | 0 | 0 | 0 |
| SALT00000032099_Cluster598-005 | HB-other | PREDICTED: uncharacter | 4099 | 0 | 0 | 0 |
| SALT00000032308_Cluster598-006 | HB-other | PREDICTED: uncharacter | 4087 | 0 | 0 | 0 |
| SALT00000033077_Cluster598-007 | HB-other | PREDICTED: uncharacter | 3816 | 0 | 0 | 0 |
| SALT00000079454_Cluster598-008 | HB-other | PREDICTED: uncharacter | 3961 | 0 | 0 | 0 |
| SALT00000087860_Cluster598-010 | HB-other | PREDICTED: uncharacter | 3987 | 0 | 0 | 0 |
| SALT00000087861_Cluster598-011 | HB-other | PREDICTED: uncharacter | 4058 | 0 | 0 | 0 |
| SALT00000090256_Cluster598-012 | HB-other | PREDICTED: uncharacter | 7588 | 0 | 0 | 0 |
| SALT00000090460_Cluster598-013 | HB-other | PREDICTED: uncharacter | 3970 | 0 | 0 | 0 |
| SALT00000092787_Cluster598-014 | HB-other | PREDICTED: uncharacter | 3709 | 0 | 0 | 0 |
| SALT00000081557_Cluster598-016 | HB-other | PREDICTED: uncharacter | 3975 | 0 | 0 | 0 |
| SALT00000077748_Cluster1038-006 | HB-other | homeodomain-like transcri | 6698 | 0 | 0 | 0 |

| | | | | | | |
|---|---|---|---|---|---|---|
| SALT00000031129_Cluster2091-001 | HB-other | flowering-time [Oryza sati | 3631 | 0 | 0 | 0 |
| SALT00000080263_Cluster2091-002 | HB-other | flowering-time [Oryza sati | 3455 | 0 | 0 | 0 |
| SALT00000086152_Cluster2091-003 | HB-other | flowering-time [Oryza sati | 5085 | 0 | 0 | 0 |
| SALT00000085301_Cluster2091-005 | HB-other | homeobox LUMINIDEPE | 3986 | 0 | 0 | 0 |
| SALT00000047580_Cluster3087-001 | HB-other | homeobox3 isoform X1 [Z | 1721 | 0 | 0 | 0 |
| SALT00000086510_Cluster5010-007 | HB-other | flowering-time [Oryza sati | 3808 | 0 | 0 | 0 |
| SALT00000083980_Cluster7732-001 | HB-other | Homeobox-leucine zipper | 3523 | 0 | 0 | 0 |
| SALT00000085529_Cluster7956-001 | HB-other | homeo [Gaeumannomyces | 3489 | 0 | 0 | 0 |
| SALT00000025635_Cluster8478-001 | HB-other | homeodomain-like transcri | 2753 | 0 | 0 | 0 |
| SALT00000025888_Cluster8478-002 | HB-other | homeodomain-like transcri | 2733 | 0 | 0 | 0 |
| SALT00000050429_Cluster8478-003 | HB-other | homeodomain-like transcri | 2158 | 0 | 0 | 0 |
| SALT00000069901_Cluster8478-005 | HB-other | homeodomain-like transcri | 2687 | 0 | 0 | 0 |
| SALT00000091703_Cluster8478-008 | HB-other | homeodomain-like transcri | 3390 | 0 | 0 | 0 |
| SALT00000064399_Cluster8478-010 | HB-other | homeodomain-like transcri | 2544 | 0 | 0 | 0 |
| SALT00000023089_Cluster8834-001 | HB-other | nodulin homeobox isoform | 3274 | 0 | 0 | 0 |
| SALT00000027951_Cluster8834-002 | HB-other | nodulin homeobox isoform | 2913 | 0 | 0 | 0 |
| SALT00000044124_Cluster8834-003 | HB-other | nodulin homeobox isoform | 2030 | 0 | 0 | 0 |
| SALT00000064451_Cluster8834-004 | HB-other | nodulin homeobox isoform | 2886 | 0 | 0 | 0 |
| SALT00000066075_Cluster8834-005 | HB-other | nodulin homeobox isoform | 3198 | 0 | 0 | 0 |
| SALT00000066574_Cluster8834-006 | HB-other | nodulin homeobox isoform | 3207 | 0 | 0 | 0 |
| SALT00000069967_Cluster8834-007 | HB-other | nodulin homeobox isoform | 3281 | 0 | 0 | 0 |
| SALT00000073082_Cluster8834-008 | HB-other | nodulin homeobox isoform | 3293 | 0 | 0 | 0 |
| SALT00000086476_Cluster8834-010 | HB-other | nodulin homeobox isoform | 3209 | 0 | 0 | 0 |
| SALT00000092035_Cluster8834-011 | HB-other | nodulin homeobox isoform | 3241 | 0 | 0 | 0 |
| SALT00000092526_Cluster8834-012 | HB-other | nodulin homeobox isoform | 3320 | 0 | 0 | 0 |
| SALT00000093345_Cluster8834-013 | HB-other | nodulin homeobox isoform | 3309 | 0 | 0 | 0 |
| SALT00000073344_Cluster9137-004 | HB-other | homeobox-leucine zipper I | 2568 | 0 | 0 | 0 |
| SALT00000083406_Cluster10880-010 | HB-other | BEL1-like homeodomain 7 | 2794 | 0 | 0 | 0 |
| SALT00000074134_Cluster16717-001 | HB-other | Homeobox-leucine zipper | 2497 | 0 | 0 | 0 |
| SALT00000000882_Cluster17357-001 | HB-other | SAWADEE HOMEODOM | 1474 | 0 | 0 | 0 |
| SALT00000008945_Cluster17357-002 | HB-other | SAWADEE HOMEODOM | 1423 | 0 | 0 | 0 |
| SALT00000009992_Cluster17357-003 | HB-other | SAWADEE HOMEODOM | 1599 | 0 | 0 | 0 |
| SALT00000010750_Cluster17357-004 | HB-other | SAWADEE HOMEODOM | 1590 | 0 | 0 | 0 |
| SALT00000046033_Cluster17357-005 | HB-other | SAWADEE HOMEODOM | 1430 | 0 | 0 | 0 |
| SALT00000052762_Cluster17357-006 | HB-other | SAWADEE HOMEODOM | 2429 | 0 | 0 | 0 |
| SALT00000055288_Cluster17357-007 | HB-other | SAWADEE HOMEODOM | 1511 | 0 | 0 | 0 |
| SALT00000059914_Cluster19344-005 | HB-other | homeobox knotted-1-like 1 | 1432 | 0 | 0 | 0 |
| SALT00000039185_Cluster19714-001 | HB-other | homeobox3 isoform X1 [Z | 2102 | 0 | 0 | 0 |
| SALT00000059816_Cluster19714-002 | HB-other | homeobox3 isoform X1 [Z | 2081 | 0 | 0 | 0 |
| SALT00000057375_Cluster22299-001 | HB-other | homeobox knotted-1-like 1 | 1920 | 0 | 0 | 0 |
| SALT00000059681_Cluster23486-001 | HB-other | SAWADEE HOMEODOM | 1848 | 0 | 0 | 0 |
| SALT00000039875_Cluster26904-002 | HB-other | TPA: homeobox DNA-bin | 1664 | 0 | 0 | 0 |
| SALT00000001574_Cluster27488-001 | HB-other | SAWADEE HOMEODOM | 1636 | 0 | 0 | 0 |
| SALT00000010303_Cluster27488-002 | HB-other | SAWADEE HOMEODOM | 1568 | 0 | 0 | 0 |
| SALT00000019855_Cluster27488-003 | HB-other | SAWADEE HOMEODOM | 1515 | 0 | 0 | 0 |
| SALT00000039182_Cluster28769-001 | HB-other | homeobox HD1 [Zea mays | 1570 | 0 | 0 | 0 |
| SALT00000060874_Cluster28769-002 | HB-other | homeobox HD1 [Zea mays | 1412 | 0 | 0 | 0 |
| SALT00000049206_Cluster31410-009 | HB-other | homeobox knotted-1-like 1 | 1270 | 0 | 0 | 0 |
| SALT00000022676_Cluster741-001 | HB-PHD | PHD-finger family homeo | 3131 | 0 | 0 | 0 |
| SALT00000030655_Cluster741-002 | HB-PHD | homeobox3 isoform X1 [Z | 2986 | 0 | 0 | 0 |
| SALT00000032341_Cluster741-003 | HB-PHD | PHD-finger family homeo | 4351 | 0 | 0 | 0 |
| SALT00000083608_Cluster741-005 | HB-PHD | PHD-finger family homeo | 4421 | 0 | 0 | 0 |
| SALT00000092984_Cluster741-007 | HB-PHD | homeobox3 isoform X1 [Z | 4163 | 0 | 0 | 0 |
| SALT00000030903_Cluster1924-001 | HB-PHD | homeobox3 isoform X1 [Z | 4591 | 0 | 0 | 0 |
| SALT00000031194_Cluster1924-002 | HB-PHD | homeobox3 isoform X1 [Z | 3199 | 0 | 0 | 0 |
| SALT00000033647_Cluster1924-003 | HB-PHD | homeobox3 isoform X1 [Z | 4762 | 0 | 0 | 0 |
| SALT00000033905_Cluster1924-004 | HB-PHD | homeobox3 isoform X1 [Z | 3823 | 0 | 0 | 0 |
| SALT00000081361_Cluster1924-005 | HB-PHD | homeobox3 isoform X1 [Z | 3281 | 0 | 0 | 0 |

| | | | | | | |
|---|---|---|---|---|---|---|
| SALT00000090098_Cluster1924-007 | HB-PHD | homeobox3 isoform X1 [Z | 4569 | 0 | 0 | 0 |
| SALT00000083482_Cluster3087-002 | HB-PHD | PHD-finger family homeo | 4558 | 0 | 0 | 0 |
| SALT00000090419_Cluster3425-001 | HB-PHD | PHD-finger family homeo | 4423 | 0 | 0 | 0 |
| SALT00000092350_Cluster3425-002 | HB-PHD | PHD-finger family homeo | 4298 | 0 | 0 | 0 |
| SALT00000089823_Cluster741-006 | HB-PHD | DNAJ heat shock N-termi | 7301 | 0 | 0 | down |
| SALT00000065526_Cluster6997-001 | HB-PHD | homeobox HOX1A-like | 2793 | 0 | 0 | 0 |
| SALT00000086118_Cluster6997-002 | HB-PHD | homeobox HOX1A-like | 3636 | 0 | 0 | 0 |
| SALT00000066290_Cluster8478-004 | HB-PHD | homeodomain-like transcri | 2850 | 0 | 0 | 0 |
| SALT00000073846_Cluster8478-006 | HB-PHD | homeodomain-like transcri | 2743 | 0 | 0 | 0 |
| SALT00000077833_Cluster8478-007 | HB-PHD | homeodomain-like transcri | 2735 | 0 | 0 | 0 |
| SALT00000066955_Cluster8478-009 | HB-PHD | homeodomain-like transcri | 2940 | 0 | 0 | 0 |
| SALT00000028779_Cluster9518-001 | HB-PHD | homeodomain-like transcri | 3017 | 0 | 0 | 0 |
| SALT00000069170_Cluster9518-002 | HB-PHD | homeodomain-like transcri | 3085 | 0 | 0 | 0 |
| SALT00000069239_Cluster9518-003 | HB-PHD | homeodomain-like transcri | 3089 | 0 | 0 | 0 |
| SALT00000071470_Cluster9518-004 | HB-PHD | homeodomain-like transcri | 3188 | 0 | 0 | 0 |
| SALT00000073164_Cluster9518-005 | HB-PHD | homeodomain-like transcri | 2918 | 0 | 0 | 0 |
| SALT00000079250_Cluster9518-006 | HB-PHD | homeodomain-like transcri | 2833 | 0 | 0 | 0 |
| SALT00000024141_Cluster605-001 | HD-ZIP | rolled expressed [Oryza sa | 3160 | 0 | 0 | 0 |
| SALT00000063304_Cluster605-002 | HD-ZIP | rolled expressed [Oryza sa | 3207 | 0 | 0 | 0 |
| SALT00000067479_Cluster605-003 | HD-ZIP | rolled expressed [Oryza sa | 2829 | 0 | 0 | 0 |
| SALT00000074011_Cluster605-004 | HD-ZIP | rolled expressed [Oryza sa | 2650 | 0 | 0 | 0 |
| SALT00000075422_Cluster605-005 | HD-ZIP | rolled expressed [Oryza sa | 3147 | 0 | 0 | 0 |
| SALT00000092250_Cluster605-008 | HD-ZIP | rolled expressed [Oryza sa | 3207 | 0 | 0 | 0 |
| SALT00000074852_Cluster605-009 | HD-ZIP | rolled expressed [Oryza sa | 2917 | 0 | 0 | 0 |
| SALT00000088635_Cluster605-010 | HD-ZIP | rolled expressed [Oryza sa | 3242 | 0 | 0 | 0 |
| SALT00000078403_Cluster605-011 | HD-ZIP | rolled expressed [Oryza sa | 3382 | 0 | 0 | 0 |
| SALT00000031817_Cluster2283-004 | HD-ZIP | TPA_exp: homeodomain le | 3520 | 0 | 0 | 0 |
| SALT00000032236_Cluster2283-005 | HD-ZIP | homeobox-leucine zipper I | 3310 | 0 | 0 | 0 |
| SALT00000032692_Cluster2283-006 | HD-ZIP | TPA_exp: homeodomain le | 3464 | 0 | 0 | 0 |
| SALT00000082600_Cluster2283-007 | HD-ZIP | homeobox-leucine zipper I | 4972 | 0 | 0 | 0 |
| SALT00000083437_Cluster2283-008 | HD-ZIP | TPA_exp: homeodomain le | 3464 | 0 | 0 | 0 |
| SALT00000091544_Cluster2283-009 | HD-ZIP | TPA_exp: homeodomain le | 3806 | 0 | 0 | 0 |
| SALT00000081560_Cluster2283-010 | HD-ZIP | TPA_exp: homeodomain le | 2499 | 0 | 0 | 0 |
| SALT00000070489_Cluster2283-011 | HD-ZIP | TPA_exp: homeodomain le | 2704 | 0 | 0 | 0 |
| SALT00000023876_Cluster3102-003 | HD-ZIP | Os03g0640800 [Oryza sati | 2971 | 0 | 0 | 0 |
| SALT00000024372_Cluster3102-004 | HD-ZIP | Os03g0640800 [Oryza sati | 2913 | 0 | 0 | 0 |
| SALT00000024482_Cluster3102-005 | HD-ZIP | Os03g0640800 [Oryza sati | 2929 | 0 | 0 | 0 |
| SALT00000025207_Cluster3102-006 | HD-ZIP | class III HD-Zip expressed | 2932 | 0 | 0 | 0 |
| SALT00000025322_Cluster3102-007 | HD-ZIP | Os03g0640800 [Oryza sati | 2993 | 0 | 0 | 0 |
| SALT00000026307_Cluster3102-008 | HD-ZIP | Os03g0640800 [Oryza sati | 2883 | 0 | 0 | 0 |
| SALT00000072156_Cluster3102-013 | HD-ZIP | class III HD-Zip expressed | 3222 | 0 | 0 | 0 |
| SALT00000075715_Cluster3102-014 | HD-ZIP | Os03g0640800 [Oryza sati | 2983 | 0 | 0 | 0 |
| SALT00000078331_Cluster3102-015 | HD-ZIP | Os03g0640800 [Oryza sati | 2991 | 0 | 0 | 0 |
| SALT00000078430_Cluster3102-016 | HD-ZIP | homeobox-leucine zipper I | 3040 | 0 | 0 | 0 |
| SALT00000078926_Cluster3102-017 | HD-ZIP | Os03g0640800 [Oryza sati | 4553 | 0 | 0 | 0 |
| SALT00000084543_Cluster3102-018 | HD-ZIP | Os03g0640800 [Oryza sati | 3042 | 0 | 0 | 0 |
| SALT00000000310_Cluster3205-001 | HD-ZIP | HB transcription partial [Z | 1555 | 0 | 0 | 0 |
| SALT00000006485_Cluster3205-002 | HD-ZIP | homeodomain leucine zipp | 1461 | 0 | 0 | 0 |
| SALT00000006739_Cluster3205-003 | HD-ZIP | HB transcription partial [Z | 1568 | 0 | up | 0 |
| SALT00000007763_Cluster3205-004 | HD-ZIP | HB transcription partial [Z | 1371 | 0 | 0 | up |
| SALT00000011153_Cluster3205-005 | HD-ZIP | HB transcription partial [Z | 1503 | 0 | 0 | 0 |
| SALT00000053768_Cluster3205-007 | HD-ZIP | HB transcription partial [Z | 1602 | 0 | 0 | 0 |
| SALT00000059740_Cluster3205-008 | HD-ZIP | HB transcription partial [Z | 1437 | 0 | 0 | 0 |
| SALT00000091515_Cluster3963-001 | HD-ZIP | homeobox-leucine zipper I | 4236 | 0 | 0 | 0 |
| SALT00000021168_Cluster4014-001 | HD-ZIP | rolled leaf 2 [Zea mays] | 3059 | 0 | 0 | 0 |
| SALT00000026993_Cluster4014-002 | HD-ZIP | rolled leaf 2 [Zea mays] | 2964 | 0 | 0 | 0 |
| SALT00000028366_Cluster4014-003 | HD-ZIP | rolled leaf 2 [Zea mays] | 3026 | 0 | 0 | 0 |
| SALT00000065850_Cluster4014-004 | HD-ZIP | rolled leaf 2 [Zea mays] | 3152 | 0 | 0 | 0 |

| | | | | | | |
|---|---|---|---|---|---|---|
| SALT00000076204_Cluster4014-005 | HD-ZIP | rolled leaf 2 [Zea mays] | 2991 | 0 | 0 | 0 |
| SALT00000077372_Cluster4014-006 | HD-ZIP | rolled leaf 2 [Zea mays] | 2547 | 0 | 0 | 0 |
| SALT00000079977_Cluster4014-007 | HD-ZIP | rolled leaf 2 [Zea mays] | 3142 | 0 | 0 | 0 |
| SALT00000081264_Cluster4014-008 | HD-ZIP | rolled leaf 2 [Zea mays] | 4219 | 0 | 0 | 0 |
| SALT00000089924_Cluster4014-009 | HD-ZIP | rolled leaf 2 isoform X2 [Z | 3714 | 0 | 0 | 0 |
| SALT00000068604_Cluster4344-001 | HD-ZIP | Os02g0674800 [Oryza sati | 3106 | 0 | 0 | 0 |
| SALT00000073287_Cluster4344-002 | HD-ZIP | Os02g0674800 [Oryza sati | 3037 | 0 | 0 | 0 |
| SALT00000082461_Cluster4344-003 | HD-ZIP | Os02g0674800 [Oryza sati | 2567 | 0 | 0 | 0 |
| SALT00000083250_Cluster4344-004 | HD-ZIP | homeobox-leucine zipper I | 3155 | 0 | down | 0 |
| SALT00000086322_Cluster4344-005 | HD-ZIP | Os02g0674800 [Oryza sati | 4130 | 0 | 0 | 0 |
| SALT00000078363_Cluster7120-001 | HD-ZIP | homeobox-leucine zipper I | 3619 | 0 | 0 | 0 |
| SALT00000088317_Cluster7120-002 | HD-ZIP | homeobox-leucine zipper I | 3482 | 0 | 0 | 0 |
| SALT00000092146_Cluster7120-003 | HD-ZIP | homeobox-leucine zipper I | 3475 | 0 | down | down |
| SALT00000093081_Cluster7120-004 | HD-ZIP | homeobox-leucine zipper I | 3595 | 0 | 0 | down |
| SALT00000091050_Cluster7120-005 | HD-ZIP | homeobox-leucine zipper I | 3418 | 0 | 0 | 0 |
| SALT00000085179_Cluster7887-001 | HD-ZIP | homeobox-leucine zipper I | 3500 | 0 | 0 | down |
| SALT00000022134_Cluster8709-001 | HD-ZIP | homeobox-leucine zipper I | 2972 | 0 | 0 | 0 |
| SALT00000026095_Cluster8709-002 | HD-ZIP | homeobox-leucine zipper I | 2903 | 0 | 0 | 0 |
| SALT00000063066_Cluster8709-003 | HD-ZIP | homeobox-leucine zipper I | 3210 | 0 | 0 | 0 |
| SALT00000075655_Cluster8709-004 | HD-ZIP | homeobox-leucine zipper I | 3234 | 0 | down | down |
| SALT00000083071_Cluster8709-005 | HD-ZIP | homeobox-leucine zipper I | 3345 | 0 | 0 | 0 |
| SALT00000071595_Cluster9137-001 | HD-ZIP | OCL5 [Oryza sativa Japon | 2599 | 0 | 0 | 0 |
| SALT00000084045_Cluster9137-002 | HD-ZIP | OCL5 [Oryza sativa Japon | 2454 | 0 | 0 | 0 |
| SALT00000086497_Cluster9137-003 | HD-ZIP | OCL5 [Oryza sativa Japon | 3262 | 0 | 0 | 0 |
| SALT00000067005_Cluster9519-001 | HD-ZIP | homeobox-leucine zipper I | 3150 | 0 | 0 | 0 |
| SALT00000073644_Cluster9519-002 | HD-ZIP | homeobox-leucine zipper I | 2998 | 0 | 0 | 0 |
| SALT00000073680_Cluster9519-003 | HD-ZIP | homeobox-leucine zipper I | 3188 | down | down | down |
| SALT00000022928_Cluster9746-001 | HD-ZIP | outer cell layer5a [Zea may | 3000 | 0 | 0 | 0 |
| SALT00000027203_Cluster9746-002 | HD-ZIP | outer cell layer5a [Zea may | 3123 | 0 | 0 | 0 |
| SALT00000029466_Cluster9746-003 | HD-ZIP | outer cell layer5a [Zea may | 2595 | 0 | 0 | 0 |
| SALT00000066333_Cluster9746-006 | HD-ZIP | outer cell layer5a [Zea may | 2699 | 0 | 0 | 0 |
| SALT00000066570_Cluster9746-007 | HD-ZIP | outer cell layer5a [Zea may | 3148 | 0 | 0 | 0 |
| SALT00000071018_Cluster9746-008 | HD-ZIP | outer cell layer5a [Zea may | 3101 | 0 | 0 | 0 |
| SALT00000076735_Cluster9746-009 | HD-ZIP | outer cell layer5a [Zea may | 3026 | 0 | 0 | 0 |
| SALT00000078855_Cluster9746-010 | HD-ZIP | outer cell layer5a [Zea may | 2559 | 0 | 0 | 0 |
| SALT00000078915_Cluster9746-011 | HD-ZIP | outer cell layer5a [Zea may | 2835 | 0 | 0 | 0 |
| SALT00000092782_Cluster9746-012 | HD-ZIP | TPA_exp: homeodomain le | 3092 | 0 | 0 | 0 |
| SALT00000083509_Cluster9746-013 | HD-ZIP | TPA_exp: homeodomain le | 2965 | 0 | 0 | 0 |
| SALT00000030884_Cluster10349-002 | HD-ZIP | Os12g0612700 [Oryza sati | 2948 | 0 | 0 | 0 |
| SALT00000031047_Cluster10349-003 | HD-ZIP | Os12g0612700 [Oryza sati | 3010 | 0 | 0 | 0 |
| SALT00000067422_Cluster10349-004 | HD-ZIP | Os12g0612700 [Oryza sati | 2856 | 0 | 0 | 0 |
| SALT00000072256_Cluster10349-005 | HD-ZIP | Os12g0612700 [Oryza sati | 2977 | 0 | 0 | 0 |
| SALT00000073675_Cluster10349-006 | HD-ZIP | Os12g0612700 [Oryza sati | 2985 | 0 | 0 | 0 |
| SALT00000074300_Cluster10349-007 | HD-ZIP | Os12g0612700 [Oryza sati | 3058 | 0 | 0 | 0 |
| SALT00000076026_Cluster10349-008 | HD-ZIP | Os12g0612700 [Oryza sati | 3005 | 0 | 0 | 0 |
| SALT00000017399_Cluster10626-003 | HD-ZIP | Os02g0674800 [Oryza sati | 1860 | 0 | 0 | 0 |
| SALT00000021202_Cluster10626-004 | HD-ZIP | homeobox-leucine zipper I | 3018 | 0 | 0 | 0 |
| SALT00000024678_Cluster10831-001 | HD-ZIP | Os04g0569100 [Oryza sati | 2993 | 0 | 0 | 0 |
| SALT00000064337_Cluster10831-002 | HD-ZIP | Os04g0569100 [Oryza sati | 2775 | 0 | 0 | 0 |
| SALT00000082108_Cluster10831-003 | HD-ZIP | Os04g0569100 [Oryza sati | 2955 | 0 | 0 | 0 |
| SALT00000027536_Cluster11335-001 | HD-ZIP | TPA_exp: homeodomain le | 2931 | 0 | 0 | 0 |
| SALT00000028125_Cluster11335-002 | HD-ZIP | TPA_exp: homeodomain le | 2746 | 0 | 0 | 0 |
| SALT00000022903_Cluster11937-001 | HD-ZIP | homeobox-leucine zipper I | 2748 | 0 | 0 | 0 |
| SALT00000065644_Cluster11937-002 | HD-ZIP | homeobox-leucine zipper I | 2874 | 0 | 0 | 0 |
| SALT00000001948_Cluster12755-001 | HD-ZIP | [Hordeum vulgare vulgare | 1492 | 0 | 0 | 0 |
| SALT00000015757_Cluster12755-002 | HD-ZIP | [Hordeum vulgare vulgare | 1321 | 0 | 0 | 0 |
| SALT00000019270_Cluster12755-003 | HD-ZIP | [Hordeum vulgare vulgare | 1705 | 0 | 0 | 0 |
| SALT00000052112_Cluster12755-004 | HD-ZIP | [Hordeum vulgare vulgare | 1729 | 0 | 0 | 0 |

| | | | | | | |
|---|---|---|---|---|---|---|
| SALT00000084500_Cluster12755-005 | HD-ZIP | [Hordeum vulgare vulgare] | 2796 | 0 | 0 | 0 |
| SALT00000009572_Cluster12755-006 | HD-ZIP | [Hordeum vulgare vulgare] | 1384 | 0 | 0 | 0 |
| SALT00000005865_Cluster20932-001 | HD-ZIP | homeodomain leucine zipp | 2009 | 0 | 0 | 0 |
| SALT00000006047_Cluster20932-002 | HD-ZIP | homeodomain leucine zipp | 1545 | 0 | 0 | 0 |
| SALT00000007536_Cluster20932-003 | HD-ZIP | homeodomain leucine zipp | 1906 | 0 | 0 | 0 |
| SALT00000013488_Cluster20932-004 | HD-ZIP | homeodomain leucine zipp | 1950 | 0 | 0 | 0 |
| SALT00000051656_Cluster20932-005 | HD-ZIP | homeodomain leucine zipp | 1962 | 0 | 0 | 0 |
| SALT00000039174_Cluster21546-001 | HD-ZIP | Os08g0481400 [Oryza sati | 1968 | 0 | 0 | 0 |
| SALT00000041560_Cluster21546-002 | HD-ZIP | Os08g0481400 [Oryza sati | 1708 | 0 | 0 | 0 |
| SALT00000055607_Cluster21546-003 | HD-ZIP | [Hordeum vulgare vulgare] | 1458 | 0 | 0 | 0 |
| SALT00000004662_Cluster21546-004 | HD-ZIP | [Hordeum vulgare vulgare] | 1726 | 0 | 0 | 0 |
| SALT00000011869_Cluster21680-001 | HD-ZIP | HB transcription partial [Z | 1960 | 0 | 0 | 0 |
| SALT00000000251_Cluster21979-001 | HD-ZIP | homeodomain leucine zipp | 1690 | 0 | 0 | 0 |
| SALT00000000693_Cluster21979-002 | HD-ZIP | homeodomain leucine zipp | 1940 | 0 | 0 | 0 |
| SALT00000001267_Cluster21979-003 | HD-ZIP | homeodomain leucine zipp | 1781 | 0 | 0 | 0 |
| SALT00000001547_Cluster21979-004 | HD-ZIP | homeodomain leucine zipp | 1615 | 0 | 0 | 0 |
| SALT00000008277_Cluster21979-005 | HD-ZIP | homeodomain leucine zipp | 1742 | 0 | 0 | 0 |
| SALT00000009424_Cluster21979-006 | HD-ZIP | homeodomain leucine zipp | 1569 | down | down | 0 |
| SALT00000011428_Cluster21979-007 | HD-ZIP | homeodomain leucine zipp | 1517 | 0 | 0 | 0 |
| SALT00000016822_Cluster21979-008 | HD-ZIP | homeodomain leucine zipp | 1704 | 0 | 0 | 0 |
| SALT00000018509_Cluster21979-009 | HD-ZIP | homeodomain leucine zipp | 1582 | 0 | 0 | 0 |
| SALT00000034892_Cluster21979-010 | HD-ZIP | homeodomain leucine zipp | 1904 | 0 | 0 | 0 |
| SALT00000034975_Cluster21979-011 | HD-ZIP | homeodomain leucine zipp | 1897 | 0 | 0 | 0 |
| SALT00000035260_Cluster21979-012 | HD-ZIP | homeodomain leucine zipp | 1898 | 0 | 0 | 0 |
| SALT00000035531_Cluster21979-013 | HD-ZIP | homeodomain leucine zipp | 1866 | 0 | 0 | 0 |
| SALT00000042091_Cluster21979-014 | HD-ZIP | homeodomain leucine zipp | 1501 | 0 | 0 | 0 |
| SALT00000044695_Cluster21979-015 | HD-ZIP | homeodomain leucine zipp | 1786 | 0 | 0 | 0 |
| SALT00000046291_Cluster21979-016 | HD-ZIP | homeodomain leucine zipp | 1634 | 0 | 0 | 0 |
| SALT00000047804_Cluster21979-017 | HD-ZIP | homeodomain leucine zipp | 1666 | 0 | 0 | 0 |
| SALT00000051706_Cluster21979-018 | HD-ZIP | homeodomain leucine zipp | 1687 | 0 | 0 | 0 |
| SALT00000052080_Cluster21979-019 | HD-ZIP | homeodomain leucine zipp | 1657 | 0 | 0 | 0 |
| SALT00000058046_Cluster21979-020 | HD-ZIP | homeodomain leucine zipp | 1861 | 0 | 0 | 0 |
| SALT00000059302_Cluster21979-021 | HD-ZIP | homeodomain leucine zipp | 1859 | 0 | down | 0 |
| SALT00000063498_Cluster21979-022 | HD-ZIP | homeodomain leucine zipp | 1795 | 0 | 0 | 0 |
| SALT00000093919_Cluster21979-023 | HD-ZIP | homeodomain leucine zipp | 1863 | 0 | 0 | 0 |
| SALT00000048383_Cluster21979-024 | HD-ZIP | [Hordeum vulgare vulgare] | 2168 | 0 | down | down |
| SALT00000052293_Cluster21979-025 | HD-ZIP | homeodomain leucine zipp | 1494 | 0 | 0 | 0 |
| SALT00000000047_Cluster22937-001 | HD-ZIP | homeodomain leucine zipp | 1714 | 0 | up | 0 |
| SALT00000000596_Cluster22937-002 | HD-ZIP | homeodomain leucine zipp | 1625 | 0 | 0 | 0 |
| SALT00000002594_Cluster22937-003 | HD-ZIP | homeodomain leucine zipp | 1536 | 0 | 0 | 0 |
| SALT00000005929_Cluster22937-004 | HD-ZIP | homeodomain leucine zipp | 1483 | 0 | 0 | 0 |
| SALT00000007278_Cluster22937-005 | HD-ZIP | homeodomain leucine zipp | 1491 | 0 | 0 | 0 |
| SALT00000011097_Cluster22937-006 | HD-ZIP | homeodomain leucine zipp | 1540 | 0 | 0 | 0 |
| SALT00000013030_Cluster22937-007 | HD-ZIP | homeodomain leucine zipp | 1672 | 0 | 0 | 0 |
| SALT00000015869_Cluster22937-008 | HD-ZIP | homeodomain leucine zipp | 1503 | 0 | 0 | 0 |
| SALT00000017388_Cluster22937-009 | HD-ZIP | homeodomain leucine zipp | 1770 | 0 | 0 | 0 |
| SALT00000025044_Cluster22937-010 | HD-ZIP | homeodomain leucine zipp | 1612 | 0 | 0 | 0 |
| SALT00000025067_Cluster22937-011 | HD-ZIP | homeodomain leucine zipp | 1619 | 0 | 0 | 0 |
| SALT00000025103_Cluster22937-012 | HD-ZIP | homeodomain leucine zipp | 1713 | 0 | 0 | 0 |
| SALT00000040899_Cluster22937-013 | HD-ZIP | homeodomain leucine zipp | 1587 | 0 | 0 | 0 |
| SALT00000041276_Cluster22937-014 | HD-ZIP | homeodomain leucine zipp | 1881 | 0 | 0 | 0 |
| SALT00000042245_Cluster22937-015 | HD-ZIP | homeodomain leucine zipp | 1525 | 0 | 0 | 0 |
| SALT00000045421_Cluster22937-016 | HD-ZIP | homeodomain leucine zipp | 1684 | 0 | 0 | 0 |
| SALT00000045470_Cluster22937-017 | HD-ZIP | homeodomain leucine zipp | 1785 | 0 | 0 | 0 |
| SALT00000059021_Cluster22937-018 | HD-ZIP | homeodomain leucine zipp | 1635 | 0 | 0 | 0 |
| SALT00000066880_Cluster22937-019 | HD-ZIP | homeodomain leucine zipp | 1694 | 0 | down | 0 |
| SALT00000041380_Cluster22937-020 | HD-ZIP | homeodomain leucine zipp | 1757 | 0 | 0 | 0 |
| SALT00000047984_Cluster23423-001 | HD-ZIP | homeodomain leucine zipp | 1811 | 0 | 0 | 0 |

| | | | | | | |
|---|---|---|---|---|---|---|
| SALT00000056017_Cluster23423-002 | HD-ZIP | [Hordeum vulgare vulgare] | 1852 | 0 | 0 | 0 |
| SALT00000049353_Cluster23658-001 | HD-ZIP | homeodomain leucine zipp | 1837 | 0 | 0 | 0 |
| SALT00000057100_Cluster23759-001 | HD-ZIP | homeodomain leucine zipp | 1798 | 0 | 0 | 0 |
| SALT00000068272_Cluster23759-002 | HD-ZIP | [Hordeum vulgare vulgare] | 1832 | 0 | 0 | 0 |
| SALT00000047285_Cluster26702-001 | HD-ZIP | homeodomain-leucine zipp | 1600 | 0 | 0 | 0 |
| SALT00000048871_Cluster26702-002 | HD-ZIP | homeodomain-leucine zipp | 1675 | 0 | 0 | 0 |
| SALT00000060963_Cluster26702-003 | HD-ZIP | homeodomain-leucine zipp | 1434 | 0 | 0 | 0 |
| SALT00000060994_Cluster26702-004 | HD-ZIP | homeodomain-leucine zipp | 1542 | 0 | 0 | 0 |
| SALT00000037683_Cluster26823-001 | HD-ZIP | homeobox-leucine zipper I | 1668 | up | up | up |
| SALT00000002244_Cluster26889-001 | HD-ZIP | homeobox-leucine zipper I | 1664 | up | up | up |
| SALT00000017409_Cluster26889-002 | HD-ZIP | homeobox-leucine zipper I | 1580 | 0 | 0 | 0 |
| SALT00000036367_Cluster26889-003 | HD-ZIP | homeobox-leucine zipper I | 1637 | 0 | 0 | 0 |
| SALT00000036944_Cluster26904-001 | HD-ZIP | homeobox-leucine zipper I | 1311 | 0 | 0 | 0 |
| SALT00000072989_Cluster26904-003 | HD-ZIP | TPA: homeobox DNA-bin | 1507 | 0 | 0 | 0 |
| SALT00000010277_Cluster27289-001 | HD-ZIP | homeodomain leucine zipp | 1563 | 0 | 0 | 0 |
| SALT00000015552_Cluster27289-002 | HD-ZIP | homeodomain leucine zipp | 1646 | 0 | 0 | 0 |
| SALT00000000803_Cluster27305-001 | HD-ZIP | homeodomain leucine zipp | 1595 | 0 | 0 | 0 |
| SALT00000006756_Cluster27305-002 | HD-ZIP | homeodomain leucine zipp | 1635 | 0 | 0 | 0 |
| SALT00000007579_Cluster27305-003 | HD-ZIP | homeodomain leucine zipp | 1513 | 0 | 0 | 0 |
| SALT00000012061_Cluster27305-004 | HD-ZIP | homeodomain leucine zipp | 1553 | 0 | 0 | 0 |
| SALT00000012272_Cluster27305-005 | HD-ZIP | homeodomain leucine zipp | 1645 | 0 | 0 | 0 |
| SALT00000041320_Cluster27305-006 | HD-ZIP | homeodomain leucine zipp | 1602 | 0 | 0 | 0 |
| SALT00000057553_Cluster27305-007 | HD-ZIP | homeodomain leucine zipp | 1570 | 0 | 0 | 0 |
| SALT00000044312_Cluster27305-008 | HD-ZIP | homeodomain leucine zipp | 1565 | 0 | 0 | 0 |
| SALT00000017543_Cluster27791-001 | HD-ZIP | Os10g0404900 [Oryza sati | 1621 | 0 | 0 | 0 |
| SALT00000043995_Cluster28350-001 | HD-ZIP | Os10g0404900 [Oryza sati | 1592 | 0 | 0 | 0 |
| SALT00000011333_Cluster28549-001 | HD-ZIP | [Hordeum vulgare vulgare] | 1581 | 0 | 0 | 0 |
| SALT00000002112_Cluster28716-001 | HD-ZIP | homeodomain-leucine zipp | 1572 | 0 | 0 | 0 |
| SALT00000005888_Cluster28716-002 | HD-ZIP | homeodomain-leucine zipp | 1482 | 0 | 0 | 0 |
| SALT00000013187_Cluster28716-003 | HD-ZIP | homeodomain-leucine zipp | 1521 | 0 | 0 | 0 |
| SALT00000015410_Cluster28716-004 | HD-ZIP | homeodomain-leucine zipp | 1466 | 0 | 0 | 0 |
| SALT00000056501_Cluster28716-005 | HD-ZIP | homeodomain-leucine zipp | 1514 | 0 | 0 | 0 |
| SALT00000046672_Cluster29751-001 | HD-ZIP | [Hordeum vulgare vulgare] | 1517 | 0 | 0 | 0 |
| SALT00000051862_Cluster29751-002 | HD-ZIP | [Hordeum vulgare vulgare] | 1415 | 0 | 0 | 0 |
| SALT00000053874_Cluster30283-001 | HD-ZIP | homeobox-leucine zipper I | 1483 | 0 | 0 | 0 |
| SALT00000057163_Cluster30580-001 | HD-ZIP | [Hordeum vulgare vulgare] | 1280 | 0 | 0 | 0 |
| SALT00000057881_Cluster30580-002 | HD-ZIP | [Hordeum vulgare vulgare] | 1462 | 0 | 0 | 0 |
| SALT00000048525_Cluster30905-001 | HD-ZIP | homeobox-leucine zipper I | 1430 | 0 | 0 | 0 |
| SALT00000038825_Cluster30999-001 | HD-ZIP | homeobox-leucine zipper / | 1420 | up | up | up |
| SALT00000050758_Cluster31065-001 | HD-ZIP | [Hordeum vulgare vulgare] | 1413 | 0 | 0 | 0 |
| SALT00000010598_Cluster31186-001 | HD-ZIP | TPA: homeobox DNA-bin | 1395 | 0 | 0 | up |
| SALT00000037679_Cluster31232-001 | HD-ZIP | homeobox-leucine zipper I | 1257 | 0 | down | down |
| SALT00000060172_Cluster31232-002 | HD-ZIP | homeobox-leucine zipper I | 1389 | 0 | 0 | 0 |
| SALT00000039617_Cluster31464-001 | HD-ZIP | TPA: homeobox DNA-bin | 1340 | 0 | 0 | 0 |
| SALT00000040533_Cluster31714-001 | HD-ZIP | Homeobox-leucine zipper | 1276 | 0 | 0 | 0 |
| SALT00000012123_Cluster31807-001 | HD-ZIP | homeobox-leucine zipper I | 1247 | 0 | up | up |
| SALT00000058648_Cluster2291-003 | HSF | heat shock factor HSF30 is | 1767 | 0 | 0 | 0 |
| SALT00000037932_Cluster18431-001 | HSF | Os03g0854500 [Oryza sati | 2016 | 0 | 0 | 0 |
| SALT00000038633_Cluster18431-002 | HSF | Os03g0854500 [Oryza sati | 2228 | 0 | 0 | up |
| SALT00000039357_Cluster18431-003 | HSF | Os03g0854500 [Oryza sati | 2028 | 0 | 0 | 0 |
| SALT00000009817_Cluster19784-001 | HSF | heat shock factor HSF30 [ | 1639 | 0 | 0 | 0 |
| SALT00000010293_Cluster19784-002 | HSF | heat shock factor HSF30 [ | 1652 | 0 | 0 | 0 |
| SALT00000011663_Cluster19784-003 | HSF | heat shock factor HSF30 [ | 1749 | 0 | 0 | 0 |
| SALT00000015960_Cluster19784-004 | HSF | heat shock factor HSF30 [ | 1700 | 0 | 0 | 0 |
| SALT00000018336_Cluster19784-005 | HSF | heat shock factor HSF30 [ | 1344 | 0 | 0 | 0 |
| SALT00000019032_Cluster19784-006 | HSF | heat shock factor HSF30 [ | 1521 | 0 | 0 | 0 |
| SALT00000039610_Cluster19784-007 | HSF | heat shock factor HSF30 [ | 2096 | 0 | 0 | 0 |
| SALT00000045438_Cluster19784-008 | HSF | heat shock factor HSF30 [ | 1960 | 0 | 0 | 0 |

| | | | | | | |
|---|---|---|---|---|---|---|
| SALT00000046059_Cluster19784-009 | HSF | heat shock factor HSF30 [: | 1852 | 0 | 0 | 0 |
| SALT00000058254_Cluster19784-010 | HSF | heat shock factor HSF30 [: | 1723 | 0 | 0 | 0 |
| SALT00000012162_Cluster20237-001 | HSF | uncharacterized LOC1008: | 1781 | 0 | 0 | 0 |
| SALT00000015102_Cluster20237-002 | HSF | HSF-type DNA-binding do | 1673 | 0 | 0 | 0 |
| SALT00000037572_Cluster20237-003 | HSF | uncharacterized LOC1008: | 2060 | 0 | 0 | 0 |
| SALT00000053174_Cluster20237-004 | HSF | uncharacterized LOC1008: | 1984 | 0 | 0 | 0 |
| SALT00000060204_Cluster20237-005 | HSF | HSF-type DNA-binding do | 1455 | 0 | 0 | 0 |
| SALT00000013013_Cluster20434-001 | HSF | heat shock factor 2 [Zea m | 1719 | 0 | 0 | 0 |
| SALT00000019655_Cluster20434-002 | HSF | heat shock factor 2 [Zea m | 2021 | 0 | 0 | 0 |
| SALT00000044004_Cluster20434-003 | HSF | heat shock factor 2 [Zea m | 1956 | 0 | 0 | 0 |
| SALT00000058872_Cluster20434-004 | HSF | heat shock factor 2 [Zea m | 2045 | 0 | 0 | 0 |
| SALT00000059533_Cluster20434-005 | HSF | heat shock factor 2 [Zea m | 2000 | 0 | 0 | 0 |
| SALT00000037882_Cluster21567-001 | HSF | heat stress transcription fac | 1967 | 0 | 0 | 0 |
| SALT00000056113_Cluster21567-002 | HSF | heat stress transcription fac | 1794 | 0 | 0 | 0 |
| SALT00000013009_Cluster21742-001 | HSF | heat shock factor HSF30 [: | 1947 | 0 | 0 | 0 |
| SALT00000052576_Cluster21742-002 | HSF | heat shock factor HSF30 [: | 1956 | 0 | 0 | 0 |
| SALT00000012950_Cluster22358-001 | HSF | Os03g0854500 [Oryza sati | 1916 | 0 | 0 | 0 |
| SALT00000036228_Cluster22358-002 | HSF | Os03g0854500 [Oryza sati | 1586 | 0 | 0 | 0 |
| SALT00000013095_Cluster22436-001 | HSF | Os03g0854500 [Oryza sati | 1752 | 0 | 0 | 0 |
| SALT00000042010_Cluster22436-002 | HSF | Os03g0854500 [Oryza sati | 1911 | 0 | 0 | 0 |
| SALT00000060453_Cluster23504-001 | HSF | heat stress transcription fac | 1653 | 0 | 0 | 0 |
| SALT00000061137_Cluster23504-002 | HSF | heat stress transcription fac | 1584 | 0 | 0 | 0 |
| SALT00000061276_Cluster23504-003 | HSF | heat stress transcription fac | 1847 | 0 | 0 | 0 |
| SALT00000056921_Cluster24339-001 | HSF | heat shock factor HSF30 is | 1779 | 0 | 0 | 0 |
| SALT00000062775_Cluster24339-002 | HSF | heat shock factor HSF30 is | 1800 | 0 | 0 | 0 |
| SALT00000040433_Cluster24339-003 | HSF | heat shock factor HSF30 is | 1980 | 0 | 0 | 0 |
| SALT00000010152_Cluster24566-001 | HSF | HSF transcription partial [: | 1637 | 0 | 0 | 0 |
| SALT00000042807_Cluster24566-002 | HSF | HSF transcription partial [: | 1786 | 0 | 0 | 0 |
| SALT00000004311_Cluster24712-001 | HSF | Os01g0749300 [Oryza sati | 1778 | 0 | 0 | up |
| SALT00000006893_Cluster24712-002 | HSF | Os01g0749300 [Oryza sati | 1593 | 0 | 0 | 0 |
| SALT00000010806_Cluster24712-003 | HSF | Os01g0749300 [Oryza sati | 1680 | 0 | 0 | 0 |
| SALT00000016209_Cluster24712-004 | HSF | Os01g0749300 [Oryza sati | 1701 | 0 | 0 | 0 |
| SALT00000054793_Cluster26219-001 | HSF | Os02g0496100 [Oryza sati | 1698 | 0 | 0 | 0 |
| SALT00000058865_Cluster26219-002 | HSF | Os02g0496100 [Oryza sati | 1518 | 0 | 0 | 0 |
| SALT00000003354_Cluster26492-001 | HSF | Os09g0526600 [Oryza sati | 1471 | 0 | 0 | up |
| SALT00000003359_Cluster26492-002 | HSF | Os09g0526600 [Oryza sati | 1377 | 0 | 0 | 0 |
| SALT00000011349_Cluster26492-003 | HSF | Os09g0526600 [Oryza sati | 1532 | 0 | 0 | up |
| SALT00000054661_Cluster26492-004 | HSF | Os09g0526600 [Oryza sati | 1464 | 0 | 0 | 0 |
| SALT00000059254_Cluster26492-005 | HSF | Os09g0526600 [Oryza sati | 1685 | 0 | 0 | up |
| SALT00000039736_Cluster26492-006 | HSF | Os09g0526600 [Oryza sati | 1759 | 0 | 0 | 0 |
| SALT00000000890_Cluster27050-001 | HSF | heat shock factor 7 [Zea m | 1570 | 0 | 0 | 0 |
| SALT00000012352_Cluster27050-002 | HSF | heat shock factor 7 [Zea m | 1570 | 0 | 0 | 0 |
| SALT00000036569_Cluster27050-003 | HSF | heat shock factor 7 [Zea m | 1579 | 0 | 0 | 0 |
| SALT00000041297_Cluster27050-004 | HSF | heat shock factor 7 [Zea m | 1657 | 0 | 0 | 0 |
| SALT00000044029_Cluster27050-005 | HSF | heat shock factor 7 [Zea m | 1571 | 0 | 0 | up |
| SALT00000019657_Cluster27442-001 | HSF | [Hordeum vulgare vulgare] | 1477 | 0 | 0 | 0 |
| SALT00000055816_Cluster27442-002 | HSF | Heat shock factor expresse | 1639 | 0 | 0 | 0 |
| SALT00000019887_Cluster27442-003 | HSF | [Hordeum vulgare vulgare] | 1762 | 0 | 0 | 0 |
| SALT00000037979_Cluster27442-004 | HSF | [Hordeum vulgare vulgare] | 1550 | 0 | 0 | 0 |
| SALT00000044552_Cluster27663-001 | HSF | heat shock factor 7 [Zea m | 1579 | 0 | 0 | 0 |
| SALT00000052583_Cluster27663-002 | HSF | heat shock factor 7 [Zea m | 1542 | 0 | 0 | 0 |
| SALT00000054536_Cluster27663-003 | HSF | heat shock factor 7 [Zea m | 1628 | 0 | 0 | up |
| SALT00000056463_Cluster27663-004 | HSF | heat shock factor 7 [Zea m | 1493 | 0 | 0 | up |
| SALT00000052379_Cluster28206-001 | HSF | heat stress transcription fac | 1600 | 0 | 0 | 0 |
| SALT00000000533_Cluster28603-001 | HSF | HSF transcription partial [: | 1543 | 0 | 0 | 0 |
| SALT00000008563_Cluster28603-002 | HSF | HSF transcription partial [: | 1479 | 0 | 0 | up |
| SALT00000040608_Cluster28603-003 | HSF | HSF transcription partial [: | 1578 | 0 | 0 | 0 |
| SALT00000052346_Cluster28603-004 | HSF | HSF transcription partial [: | 1365 | 0 | 0 | 0 |

| | | | | | | |
|---|---|---|---|---|---|---|
| SALT00000008574_Cluster29616-001 | HSF | Os09g0455200 [Oryza sati | 1525 | 0 | 0 | 0 |
| SALT00000065031_Cluster16846-001 | LBD | partial [Oryza sativa Japon | 2485 | 0 | 0 | 0 |
| SALT00000004377_Cluster28578-001 | LBD | LOB domain-containing 1: | 1579 | 0 | 0 | 0 |
| SALT00000041907_Cluster31216-001 | LBD | seed specific Bn15D17A [ | 1154 | 0 | 0 | 0 |
| SALT00000046485_Cluster31216-002 | LBD | seed specific Bn15D17A [ | 1361 | 0 | 0 | 0 |
| SALT00000049321_Cluster31216-003 | LBD | seed specific Bn15D17A [ | 1391 | 0 | 0 | 0 |
| SALT00000038330_Cluster31216-004 | LBD | lateral organ boundaries dc | 1036 | 0 | 0 | 0 |
| SALT00000064074_Cluster31325-001 | LBD | LATERAL ORGAN BOU | 1372 | 0 | 0 | 0 |
| SALT00000029697_Cluster31977-001 | LBD | LOB domain-containing 4 | 1195 | 0 | 0 | 0 |
| SALT00000065074_Cluster32104-001 | LBD | LOB domain-containing 4 | 1149 | 0 | 0 | 0 |
| SALT00000031249_Cluster8077-001 | LSD | proteoglycan 4-like | 3466 | 0 | 0 | 0 |
| SALT00000040943_Cluster8077-002 | LSD | proline-rich 36-like [Brach | 1955 | 0 | 0 | 0 |
| SALT00000088173_Cluster8077-003 | LSD | proteoglycan 4-like | 3453 | 0 | 0 | 0 |
| SALT00000044570_Cluster19085-001 | LSD | Os12g0611000 [Oryza sati | 2160 | 0 | 0 | 0 |
| SALT00000049551_Cluster20830-001 | LSD | LOL3 [Oryza brachyantha | 2017 | 0 | 0 | up |
| SALT00000008052_Cluster32216-001 | LSD | LSD1 [Hordeum vulgare v | 1106 | 0 | 0 | 0 |
| SALT00000036040_Cluster4030-001 | MIKC_M | Os07g0605200 [Oryza sati | 1102 | 0 | 0 | 0 |
| SALT00000054533_Cluster4030-003 | MIKC_M | Os07g0605200 [Oryza sati | 1261 | 0 | 0 | 0 |
| SALT00000013251_Cluster28990-001 | MIKC_M | MADS-box transcription f | 1559 | 0 | 0 | 0 |
| SALT00000050067_Cluster28990-002 | MIKC_M | MADS-box transcription f | 1439 | 0 | 0 | 0 |
| SALT00000004463_Cluster29214-001 | MIKC_M | MADS-box transcription f | 1518 | 0 | 0 | 0 |
| SALT00000018643_Cluster29214-002 | MIKC_M | MADS-box transcription f | 1548 | 0 | 0 | 0 |
| SALT00000038076_Cluster29214-003 | MIKC_M | MADS-box transcription f | 1323 | 0 | 0 | 0 |
| SALT00000047168_Cluster29214-004 | MIKC_M | MADS-box transcription f | 1463 | 0 | 0 | 0 |
| SALT00000005484_Cluster29384-001 | MIKC_M | MADS-box transcription f | 1197 | 0 | 0 | 0 |
| SALT00000050379_Cluster29384-002 | MIKC_M | MADS-box transcription f | 1539 | 0 | 0 | 0 |
| SALT00000055740_Cluster31661-001 | MIKC_M | Os03g0752800 [Oryza sati | 1292 | 0 | 0 | 0 |
| SALT00000069492_Cluster31785-001 | MIKC_M | MADS-box transcription f | 1256 | 0 | 0 | 0 |
| SALT00000012846_Cluster31843-001 | MIKC_M | MADS-box transcription f | 1237 | 0 | 0 | 0 |
| SALT00000047114_Cluster32047-001 | MIKC_M | Os03g0752800 [Oryza sati | 1171 | 0 | 0 | 0 |
| SALT00000072920_Cluster32189-001 | MIKC_M | MADS-box transcription f | 1121 | 0 | 0 | 0 |
| SALT00000003391_Cluster32196-001 | MIKC_M | MADS-box transcription f | 1041 | 0 | 0 | 0 |
| SALT00000037062_Cluster32196-002 | MIKC_M | MADS-box transcription f | 1116 | 0 | 0 | 0 |
| SALT00000041997_Cluster32214-001 | MIKC_M | MADS-box transcription f | 1107 | 0 | 0 | 0 |
| SALT00000018959_Cluster32268-001 | MIKC_M | MADS-domain transcriptic | 931 | 0 | 0 | 0 |
| SALT00000058939_Cluster32268-002 | MIKC_M | MADS-domain transcriptic | 1085 | 0 | 0 | 0 |
| SALT00000087165_Cluster4030-002 | M-type_l | Os07g0605200 [Oryza sati | 4215 | 0 | 0 | 0 |
| SALT00000076487_Cluster9563-001 | M-type_l | agamous-like MADS-box | 3179 | 0 | 0 | 0 |
| SALT00000035970_Cluster894-001 | MYB | transcription factor GAMY | 1918 | 0 | 0 | 0 |
| SALT00000054810_Cluster894-003 | MYB | transcription factor GAMY | 2144 | 0 | 0 | 0 |
| SALT00000041472_Cluster894-005 | MYB | transcription factor GAMY | 2105 | 0 | 0 | 0 |
| SALT00000005223_Cluster954-001 | MYB | cell division cycle 5 | 2140 | 0 | 0 | 0 |
| SALT00000023395_Cluster954-002 | MYB | cell division cycle 5 [Setar | 3359 | 0 | 0 | 0 |
| SALT00000029472_Cluster954-003 | MYB | cell division cycle 5 [Setar | 3381 | 0 | 0 | 0 |
| SALT00000030441_Cluster954-004 | MYB | cell division cycle 5 [Setar | 3445 | 0 | 0 | 0 |
| SALT00000032285_Cluster954-005 | MYB | cell division cycle 5 [Setar | 3547 | 0 | 0 | 0 |
| SALT00000032862_Cluster954-006 | MYB | cell division cycle 5 [Setar | 3446 | 0 | 0 | 0 |
| SALT00000062386_Cluster954-009 | MYB | cell division cycle 5 [Setar | 3420 | 0 | 0 | 0 |
| SALT00000080381_Cluster954-013 | MYB | cell division cycle 5 [Setar | 3524 | 0 | 0 | 0 |
| SALT00000081689_Cluster954-015 | MYB | cell division cycle 5 [Setar | 3298 | 0 | 0 | 0 |
| SALT00000082032_Cluster954-016 | MYB | cell division cycle 5 [Setar | 3775 | 0 | 0 | 0 |
| SALT00000084059_Cluster954-017 | MYB | cell division cycle 5 [Setar | 3368 | 0 | 0 | 0 |
| SALT00000089854_Cluster954-018 | MYB | cell division cycle 5 [Setar | 3488 | 0 | 0 | 0 |
| SALT00000092731_Cluster954-019 | MYB | cell division cycle 5 [Setar | 3510 | 0 | 0 | 0 |
| SALT00000086201_Cluster4445-001 | MYB | MYB DNA-binding domai | 4101 | 0 | up | 0 |
| SALT00000032883_Cluster7374-001 | MYB | myb L isoform X1 [Oryza | 3548 | 0 | 0 | 0 |
| SALT00000078186_Cluster7374-002 | MYB | myb L isoform X1 [Oryza | 3480 | 0 | 0 | 0 |
| SALT00000079895_Cluster7374-003 | MYB | myb L isoform X1 [Oryza | 2696 | 0 | 0 | 0 |

| | | | | | | |
|---|---|---|---|---|---|---|
| SALT00000092665_Cluster7374-004 | MYB | myb L isoform X1 [Oryza | 3580 | 0 | 0 | 0 |
| SALT00000015499_Cluster7584-001 | MYB | snRNA-activating comple: | 1768 | 0 | 0 | 0 |
| SALT00000033234_Cluster7584-002 | MYB | snRNA-activating comple: | 3546 | 0 | 0 | 0 |
| SALT00000012173_Cluster7584-004 | MYB | snRNA-activating comple: | 1923 | 0 | 0 | 0 |
| SALT00000067209_Cluster7584-005 | MYB | snRNA-activating comple: | 2816 | 0 | 0 | 0 |
| SALT00000084846_Cluster8230-001 | MYB | myb-related 3R-1 [Setaria | 3438 | 0 | 0 | 0 |
| SALT00000006577_Cluster9031-003 | MYB | MYB transcription partial | 1231 | 0 | 0 | 0 |
| SALT00000029617_Cluster12402-001 | MYB | myb-related 3R-1-like | 2828 | 0 | 0 | 0 |
| SALT00000021226_Cluster13676-001 | MYB | LOC100285081 isoform X | 2666 | 0 | 0 | 0 |
| SALT00000021977_Cluster13676-002 | MYB | LOC100285081 isoform X | 2587 | 0 | 0 | 0 |
| SALT00000026024_Cluster13676-003 | MYB | LOC100285081 isoform X | 2519 | 0 | 0 | 0 |
| SALT00000026465_Cluster13676-004 | MYB | LOC100285081 isoform X | 2474 | 0 | 0 | 0 |
| SALT00000027462_Cluster13676-005 | MYB | LOC100285081 isoform X | 2668 | 0 | 0 | 0 |
| SALT00000028751_Cluster13676-006 | MYB | LOC100285081 isoform X | 2681 | 0 | 0 | 0 |
| SALT00000031434_Cluster13676-007 | MYB | LOC100285081 isoform X | 2686 | 0 | 0 | 0 |
| SALT00000034959_Cluster13676-008 | MYB | LOC100285081 isoform X | 2647 | 0 | 0 | 0 |
| SALT00000067420_Cluster13676-011 | MYB | LOC100285081 isoform X | 2500 | 0 | 0 | 0 |
| SALT00000071412_Cluster13676-012 | MYB | LOC100285081 isoform X | 2717 | 0 | 0 | 0 |
| SALT00000073284_Cluster13676-013 | MYB | LOC100285081 isoform X | 2608 | 0 | 0 | 0 |
| SALT00000074403_Cluster13676-014 | MYB | LOC100285081 isoform X | 2618 | 0 | 0 | 0 |
| SALT00000084272_Cluster13676-015 | MYB | LOC100285081 isoform X | 2718 | 0 | 0 | 0 |
| SALT00000070984_Cluster13676-016 | MYB | LOC100285081 isoform X | 2469 | 0 | 0 | 0 |
| SALT00000026680_Cluster15197-001 | MYB | myb-related A-like [Setaria | 2611 | 0 | 0 | 0 |
| SALT00000080527_Cluster15197-002 | MYB | myb-related A-like [Setaria | 2472 | 0 | 0 | 0 |
| SALT00000066080_Cluster15197-003 | MYB | myb-related A-like [Setaria | 2783 | 0 | 0 | 0 |
| SALT00000001175_Cluster16583-001 | MYB | transcription factor MYB4 | 1486 | 0 | 0 | 0 |
| SALT00000002814_Cluster16583-002 | MYB | sucrose responsive elemen | 1497 | 0 | 0 | 0 |
| SALT00000023549_Cluster16583-003 | MYB | sucrose responsive elemen | 1638 | 0 | 0 | 0 |
| SALT00000038020_Cluster16583-005 | MYB | sucrose responsive elemen | 1496 | 0 | 0 | 0 |
| SALT00000044428_Cluster16583-006 | MYB | sucrose responsive elemen | 1987 | 0 | 0 | 0 |
| SALT00000049168_Cluster16583-007 | MYB | transcription factor MYB4 | 1499 | 0 | 0 | up |
| SALT00000054289_Cluster16583-008 | MYB | transcription factor MYB4 | 1542 | 0 | 0 | 0 |
| SALT00000080165_Cluster16583-009 | MYB | sucrose responsive elemen | 2508 | 0 | 0 | 0 |
| SALT00000018666_Cluster16583-010 | MYB | sucrose responsive elemen | 1387 | 0 | 0 | up |
| SALT00000078664_Cluster16795-001 | MYB | transcription factor GAMy | 2490 | 0 | 0 | 0 |
| SALT00000004959_Cluster17704-001 | MYB | MYB transcription partial | 1887 | 0 | 0 | 0 |
| SALT00000008282_Cluster17704-002 | MYB | MYB transcription partial | 1634 | 0 | 0 | 0 |
| SALT00000009841_Cluster17704-003 | MYB | MYB transcription partial | 1778 | 0 | 0 | 0 |
| SALT00000010885_Cluster17704-004 | MYB | MYB transcription partial | 2089 | 0 | 0 | 0 |
| SALT00000012403_Cluster17704-005 | MYB | MYB transcription partial | 2049 | 0 | 0 | 0 |
| SALT00000016156_Cluster17704-006 | MYB | MYB transcription partial | 2039 | 0 | 0 | 0 |
| SALT00000020321_Cluster17704-007 | MYB | MYB transcription partial | 1874 | 0 | 0 | 0 |
| SALT00000024060_Cluster17704-008 | MYB | MYB transcription partial | 2366 | 0 | 0 | 0 |
| SALT00000037543_Cluster17704-009 | MYB | MYB transcription partial | 1592 | 0 | 0 | 0 |
| SALT00000039089_Cluster17704-010 | MYB | MYB transcription partial | 1900 | 0 | 0 | 0 |
| SALT00000039471_Cluster17704-011 | MYB | MYB transcription partial | 1871 | 0 | 0 | 0 |
| SALT00000040453_Cluster17704-012 | MYB | MYB transcription partial | 1779 | 0 | 0 | 0 |
| SALT00000041097_Cluster17704-013 | MYB | MYB transcription partial | 1924 | 0 | 0 | 0 |
| SALT00000044532_Cluster17704-014 | MYB | MYB transcription partial | 1584 | 0 | 0 | 0 |
| SALT00000045531_Cluster17704-015 | MYB | MYB transcription partial | 1695 | 0 | 0 | 0 |
| SALT00000050043_Cluster17704-016 | MYB | MYB transcription partial | 1729 | 0 | 0 | 0 |
| SALT00000050618_Cluster17704-017 | MYB | MYB transcription partial | 1718 | 0 | 0 | 0 |
| SALT00000052876_Cluster17704-018 | MYB | MYB transcription partial | 1713 | 0 | 0 | 0 |
| SALT00000059847_Cluster17704-019 | MYB | MYB transcription partial | 1969 | 0 | 0 | 0 |
| SALT00000067538_Cluster17704-020 | MYB | MYB transcription partial | 2311 | 0 | 0 | 0 |
| SALT00000051546_Cluster18523-001 | MYB | transcription factor GAMY | 2216 | 0 | 0 | 0 |
| SALT00000000808_Cluster19998-001 | MYB | R2R3-MYB transcription f | 1835 | 0 | 0 | 0 |
| SALT00000004620_Cluster19998-002 | MYB | R2R3-MYB transcription f | 1866 | 0 | 0 | 0 |

| | | | | | | |
|---|---|---|---|---|---|---|
| SALT00000004884_Cluster19998-003 | MYB | Os01g0285300 [Oryza sati | 1966 | 0 | 0 | 0 |
| SALT00000011800_Cluster19998-004 | MYB | Os01g0285300 [Oryza sati | 1933 | 0 | 0 | 0 |
| SALT00000013159_Cluster19998-005 | MYB | R2R3-MYB transcription f | 1896 | 0 | 0 | 0 |
| SALT00000017057_Cluster19998-006 | MYB | R2R3-MYB transcription f | 1799 | 0 | 0 | 0 |
| SALT00000034987_Cluster19998-007 | MYB | R2R3-MYB transcription f | 1872 | 0 | 0 | 0 |
| SALT00000044200_Cluster19998-008 | MYB | R2R3-MYB transcription f | 1903 | 0 | 0 | 0 |
| SALT00000048715_Cluster19998-010 | MYB | Os01g0285300 [Oryza sati | 2080 | 0 | 0 | 0 |
| SALT00000056062_Cluster19998-011 | MYB | Os01g0285300 [Oryza sati | 1688 | 0 | 0 | 0 |
| SALT00000060014_Cluster19998-013 | MYB | R2R3-MYB transcription f | 2044 | 0 | 0 | 0 |
| SALT00000015592_Cluster20260-001 | MYB | Myb domain [Theobroma c | 1374 | 0 | 0 | 0 |
| SALT00000055118_Cluster20260-003 | MYB | Transcription factor RAX2 | 1494 | 0 | 0 | 0 |
| SALT00000041800_Cluster20269-001 | MYB | tuber-specific and sucrose- | 2057 | 0 | up | 0 |
| SALT00000043394_Cluster20406-003 | MYB | Os05g0449900 [Oryza sati | 1757 | 0 | 0 | 0 |
| SALT00000018925_Cluster20633-001 | MYB | transcription factor MYB4 | 2031 | 0 | 0 | 0 |
| SALT00000044237_Cluster24176-001 | MYB | transcription factor AS1-lil | 1809 | 0 | 0 | 0 |
| SALT00000048127_Cluster24176-002 | MYB | transcription factor AS1-lil | 1708 | 0 | 0 | 0 |
| SALT00000052344_Cluster24176-003 | MYB | transcription factor AS1-lil | 1348 | 0 | 0 | 0 |
| SALT00000041277_Cluster24386-001 | MYB | myb-related Hv33 [Zea ma | 1797 | 0 | 0 | 0 |
| SALT00000002143_Cluster25119-001 | MYB | transcription factor DIVAR | 1756 | 0 | 0 | 0 |
| SALT00000009114_Cluster25119-002 | MYB | transcription factor DIVAR | 1642 | 0 | 0 | 0 |
| SALT00000012625_Cluster25119-003 | MYB | transcription factor DIVAR | 1442 | 0 | 0 | 0 |
| SALT00000023852_Cluster25119-004 | MYB | transcription factor DIVAR | 1491 | 0 | 0 | 0 |
| SALT00000053514_Cluster25119-005 | MYB | transcription factor DIVAR | 1441 | 0 | 0 | 0 |
| SALT00000055433_Cluster25119-006 | MYB | transcription factor DIVAR | 1550 | 0 | 0 | 0 |
| SALT00000018123_Cluster25970-001 | MYB | myb-like DNA-binding do | 1498 | 0 | 0 | 0 |
| SALT00000045168_Cluster25970-002 | MYB | myb-like DNA-binding do | 1561 | 0 | 0 | 0 |
| SALT00000054162_Cluster25970-003 | MYB | myb-like DNA-binding do | 1707 | 0 | 0 | 0 |
| SALT00000055166_Cluster25970-004 | MYB | myb-like DNA-binding do | 1714 | 0 | 0 | 0 |
| SALT00000006725_Cluster26080-001 | MYB | transcription factor DIVAR | 1531 | 0 | 0 | 0 |
| SALT00000015508_Cluster26080-002 | MYB | transcription factor DIVAR | 1667 | 0 | 0 | 0 |
| SALT00000056058_Cluster26080-003 | MYB | transcription factor DIVAR | 1707 | 0 | up | 0 |
| SALT00000007403_Cluster26092-002 | MYB | MYB DNA-binding domai | 1615 | 0 | 0 | down |
| SALT00000012099_Cluster26092-003 | MYB | MYB DNA-binding domai | 1546 | 0 | 0 | 0 |
| SALT00000042764_Cluster26092-007 | MYB | transcription factor MYB5 | 1431 | 0 | 0 | down |
| SALT00000006576_Cluster26166-001 | MYB | transcription factor DIVAR | 1700 | 0 | 0 | 0 |
| SALT00000013922_Cluster26166-002 | MYB | transcription factor DIVAR | 1651 | 0 | 0 | 0 |
| SALT00000014088_Cluster26166-003 | MYB | transcription factor DIVAR | 1573 | 0 | 0 | 0 |
| SALT00000016997_Cluster26166-004 | MYB | transcription factor DIVAR | 1526 | 0 | 0 | 0 |
| SALT00000005684_Cluster26272-001 | MYB | [Hordeum vulgare vulgare] | 1679 | 0 | 0 | 0 |
| SALT00000005707_Cluster26272-002 | MYB | [Hordeum vulgare vulgare] | 1464 | 0 | 0 | 0 |
| SALT00000012290_Cluster26272-003 | MYB | [Hordeum vulgare vulgare] | 1439 | 0 | 0 | 0 |
| SALT00000056610_Cluster26272-004 | MYB | [Hordeum vulgare vulgare] | 1696 | 0 | 0 | 0 |
| SALT00000036648_Cluster26328-002 | MYB | myb-related Hv33-like [Se | 1456 | 0 | 0 | down |
| SALT00000058849_Cluster26491-001 | MYB | myb-related Zm38-like | 1685 | 0 | 0 | down |
| SALT00000006768_Cluster27567-001 | MYB | MYB-like transcription fac | 1632 | 0 | 0 | up |
| SALT00000009000_Cluster27567-002 | MYB | MYB-like transcription fac | 1499 | 0 | 0 | 0 |
| SALT00000020252_Cluster27567-003 | MYB | MYB-like transcription fac | 1497 | 0 | 0 | 0 |
| SALT00000044321_Cluster27567-004 | MYB | MYB-like transcription fac | 1623 | 0 | 0 | 0 |
| SALT00000043589_Cluster27757-001 | MYB | MYB94 [Zea mays] | 1623 | 0 | 0 | 0 |
| SALT00000036305_Cluster27857-001 | MYB | secondary wall-associated | 1618 | 0 | 0 | 0 |
| SALT00000037929_Cluster28218-001 | MYB | transcription factor MYB4 | 1599 | 0 | 0 | 0 |
| SALT00000001906_Cluster28821-001 | MYB | Os03g0315400 [Oryza sati | 1567 | 0 | 0 | 0 |
| SALT00000059183_Cluster28821-002 | MYB | MYB DNA-binding domai | 1500 | 0 | 0 | 0 |
| SALT00000003934_Cluster28952-001 | MYB | MYB-like transcription fac | 1552 | 0 | 0 | 0 |
| SALT00000014009_Cluster28952-002 | MYB | MYB-like transcription fac | 1561 | 0 | 0 | 0 |
| SALT00000012504_Cluster29367-001 | MYB | MYBY1 [Oryza sativa Jap | 1452 | 0 | 0 | 0 |
| SALT00000045689_Cluster29367-002 | MYB | MYBY1 [Oryza sativa Jap | 1540 | 0 | 0 | 0 |
| SALT00000049841_Cluster29367-003 | MYB | MYBY1 [Oryza sativa Jap | 1446 | 0 | 0 | 0 |

| | | | | | | |
|---|---|---|---|---|---|---|
| SALT00000054486_Cluster29367-004 | MYB | MYBY1 [Oryza sativa Jap | 1410 | 0 | 0 | 0 |
| SALT00000061280_Cluster29583-001 | MYB | secondary wall-associated | 1528 | 0 | 0 | 0 |
| SALT00000038084_Cluster30071-001 | MYB | ATMYB4-like [Oryza sativ | 1497 | 0 | 0 | up |
| SALT00000044345_Cluster30310-001 | MYB | myb-related transcription f | 1481 | 0 | 0 | 0 |
| SALT00000036044_Cluster30423-001 | MYB | myb-related MYBAS1-like | 1473 | 0 | 0 | 0 |
| SALT00000040489_Cluster30423-002 | MYB | myb-related MYBAS1-like | 1433 | 0 | 0 | 0 |
| SALT00000007669_Cluster30499-001 | MYB | myb-related Hv33-like [Se | 1265 | 0 | 0 | 0 |
| SALT00000036415_Cluster30769-001 | MYB | Transcription factor MYB3 | 1296 | 0 | 0 | 0 |
| SALT00000046027_Cluster30769-002 | MYB | Transcription factor MYB3 | 1445 | 0 | 0 | 0 |
| SALT00000058116_Cluster30769-003 | MYB | Transcription factor MYB3 | 1296 | 0 | 0 | 0 |
| SALT00000045272_Cluster30779-001 | MYB | Transcription factor MYB3 | 1444 | 0 | 0 | 0 |
| SALT00000036379_Cluster30835-001 | MYB | myb-related Zm1 [Zea may | 1438 | 0 | 0 | 0 |
| SALT00000049074_Cluster30878-001 | MYB | [Hordeum vulgare vulgare] | 1433 | 0 | 0 | 0 |
| SALT00000048604_Cluster30986-001 | MYB | Transcription factor MYB3 | 1422 | 0 | 0 | up |
| SALT00000051073_Cluster30987-001 | MYB | myb-related Zm1-like [Zea | 1422 | 0 | 0 | 0 |
| SALT00000014372_Cluster31081-001 | MYB | Transcription factor MYB8 | 1411 | 0 | 0 | 0 |
| SALT00000055065_Cluster31148-001 | MYB | myb-related 306-like [Zea | 1400 | 0 | 0 | 0 |
| SALT00000065971_Cluster31178-001 | MYB | transcription factor MYB2 | 1397 | 0 | 0 | 0 |
| SALT00000039101_Cluster31204-001 | MYB | Os04g0508500 [Oryza sati | 1392 | 0 | 0 | 0 |
| SALT00000055640_Cluster31380-001 | MYB | transcription factor DIVAR | 1350 | 0 | 0 | 0 |
| SALT00000061207_Cluster31380-002 | MYB | transcription factor DIVAR | 1361 | 0 | 0 | 0 |
| SALT00000053446_Cluster31396-001 | MYB | myb-related Zm1-like [Zea | 1356 | 0 | 0 | 0 |
| SALT00000001019_Cluster31520-001 | MYB | MYB transcription factor 1 | 1329 | 0 | 0 | 0 |
| SALT00000047206_Cluster32037-001 | MYB | transcription factor MYB3 | 1177 | 0 | 0 | 0 |
| SALT00000065113_Cluster32283-001 | MYB | MYB transcription factor [ | 949 | 0 | 0 | 0 |
| SALT00000065578_Cluster32283-002 | MYB | MYB transcription factor [ | 968 | 0 | 0 | 0 |
| SALT00000082169_Cluster32283-003 | MYB | MYB transcription factor [ | 1079 | 0 | 0 | 0 |
| SALT00000033157_Cluster140-003 | MYB_re | ALWAYS EARLY 2 isofor | 3898 | 0 | 0 | 0 |
| SALT00000078530_Cluster140-006 | MYB_re | ALWAYS EARLY 2 isofor | 3795 | 0 | 0 | 0 |
| SALT00000080888_Cluster140-008 | MYB_re | ALWAYS EARLY 2 isofor | 3933 | 0 | 0 | 0 |
| SALT00000087785_Cluster140-010 | MYB_re | ALWAYS EARLY 2 isofor | 3975 | 0 | 0 | 0 |
| SALT00000089822_Cluster140-011 | MYB_re | ALWAYS EARLY 2 isofor | 3830 | 0 | 0 | 0 |
| SALT00000092420_Cluster140-012 | MYB_re | ALWAYS EARLY 2 isofor | 3999 | 0 | 0 | 0 |
| SALT00000030264_Cluster146-001 | MYB_re | MYB DNA-binding domai | 2978 | 0 | 0 | 0 |
| SALT00000011233_Cluster221-001 | MYB_re | transcriptional adapter AD | 1488 | 0 | 0 | 0 |
| SALT00000031004_Cluster221-003 | MYB_re | Transcriptional adapter AD | 3565 | 0 | 0 | 0 |
| SALT00000069394_Cluster221-004 | MYB_re | Transcriptional adapter AD | 3305 | 0 | 0 | 0 |
| SALT00000091460_Cluster221-005 | MYB_re | Transcriptional adapter AD | 4032 | 0 | 0 | down |
| SALT00000001246_Cluster282-001 | MYB_re | single myb histone 6 | 1464 | 0 | 0 | 0 |
| SALT00000002215_Cluster282-002 | MYB_re | single myb histone 5-like [ | 1453 | 0 | 0 | 0 |
| SALT00000006902_Cluster282-003 | MYB_re | single myb histone 6 | 1371 | 0 | 0 | 0 |
| SALT00000041935_Cluster282-004 | MYB_re | single myb histone 6 | 8770 | up | up | up |
| SALT00000052608_Cluster282-005 | MYB_re | single myb histone 6 | 1565 | 0 | 0 | 0 |
| SALT00000063447_Cluster282-006 | MYB_re | single myb histone 6 [Zea | 1599 | 0 | 0 | 0 |
| SALT00000020969_Cluster1845-004 | MYB_re | TSL-kinase interacting 1 [ | 2617 | 0 | 0 | 0 |
| SALT00000021596_Cluster1845-006 | MYB_re | TSL-kinase interacting 1 [ | 2709 | 0 | 0 | 0 |
| SALT00000023200_Cluster1845-007 | MYB_re | TSL-kinase interacting 1 [ | 2583 | 0 | 0 | 0 |
| SALT00000023381_Cluster1845-008 | MYB_re | TSL-kinase interacting 1 [ | 2609 | 0 | 0 | 0 |
| SALT00000028209_Cluster1845-013 | MYB_re | TSL-kinase interacting 1 [ | 2581 | 0 | 0 | 0 |
| SALT00000035104_Cluster1845-017 | MYB_re | TSL-kinase interacting 1 [ | 2579 | 0 | 0 | 0 |
| SALT00000035398_Cluster1845-018 | MYB_re | TSL-kinase interacting 1 [ | 2468 | 0 | 0 | 0 |
| SALT00000070612_Cluster1845-023 | MYB_re | TSL-kinase interacting 1 [ | 2486 | 0 | 0 | 0 |
| SALT00000073403_Cluster1845-025 | MYB_re | TSL-kinase interacting 1 [ | 2580 | 0 | 0 | 0 |
| SALT00000074839_Cluster1845-026 | MYB_re | MYB DNA-binding domai | 2640 | 0 | 0 | 0 |
| SALT00000075004_Cluster1845-027 | MYB_re | TSL-kinase interacting 1 [ | 2671 | 0 | 0 | 0 |
| SALT00000076328_Cluster1845-029 | MYB_re | TSL-kinase interacting 1 [ | 2567 | 0 | 0 | 0 |
| SALT00000076856_Cluster1845-031 | MYB_re | TSL-kinase interacting 1 [ | 5269 | 0 | 0 | 0 |
| SALT00000079744_Cluster1845-032 | MYB_re | TSL-kinase interacting 1 [ | 2729 | 0 | 0 | 0 |

| | | | | | | |
|---|---|---|---|---|---|---|
| SALT00000084470_Cluster1845-036 | MYB_re | MYB DNA-binding domai | 2994 | 0 | 0 | 0 |
| SALT00000036027_Cluster1845-038 | MYB_re | TSL-kinase interacting 1 [( | 1694 | 0 | 0 | 0 |
| SALT00000024204_Cluster2147-001 | MYB_re | MYB DNA-binding domai | 2824 | 0 | 0 | 0 |
| SALT00000030308_Cluster2147-003 | MYB_re | MYB DNA-binding domai | 3144 | 0 | 0 | 0 |
| SALT00000032365_Cluster2147-004 | MYB_re | MYB DNA-binding domai | 4526 | 0 | 0 | 0 |
| SALT00000087620_Cluster2147-008 | MYB_re | MYB DNA-binding domai | 5047 | 0 | 0 | 0 |
| SALT00000089223_Cluster2147-009 | MYB_re | MYB DNA-binding domai | 3881 | 0 | 0 | 0 |
| SALT00000092833_Cluster2147-010 | MYB_re | MYB DNA-binding domai | 3768 | 0 | 0 | 0 |
| SALT00000075217_Cluster2641-001 | MYB_re | LHY isoform X1 [Zea may | 4766 | 0 | 0 | 0 |
| SALT00000079698_Cluster3343-003 | MYB_re | ALWAYS EARLY 2 isofor | 3958 | 0 | 0 | 0 |
| SALT00000088951_Cluster3343-006 | MYB_re | ALWAYS EARLY 2 isofor | 3910 | 0 | 0 | 0 |
| SALT00000026196_Cluster3542-002 | MYB_re | MYB DNA-binding domai | 3077 | 0 | 0 | 0 |
| SALT00000030868_Cluster3542-003 | MYB_re | MYB DNA-binding domai | 4060 | 0 | 0 | 0 |
| SALT00000033227_Cluster3542-004 | MYB_re | MYB DNA-binding domai | 3770 | 0 | 0 | 0 |
| SALT00000034105_Cluster3542-005 | MYB_re | MYB DNA-binding domai | 4377 | 0 | 0 | 0 |
| SALT00000034196_Cluster3542-006 | MYB_re | MYB DNA-binding domai | 3620 | 0 | 0 | 0 |
| SALT00000078522_Cluster3542-008 | MYB_re | MYB DNA-binding domai | 2788 | 0 | 0 | 0 |
| SALT00000092500_Cluster4227-001 | MYB_re | single myb histone 5-like [ | 4158 | 0 | 0 | 0 |
| SALT00000085042_Cluster4922-001 | MYB_re | transcription factor DIVAF | 3992 | 0 | 0 | 0 |
| SALT00000014099_Cluster5436-004 | MYB_re | DNA methyltransferase 1-a | 1628 | 0 | 0 | 0 |
| SALT00000044091_Cluster5436-006 | MYB_re | DNA methyltransferase 1-a | 1609 | 0 | 0 | 0 |
| SALT00000053679_Cluster5436-010 | MYB_re | DNA methyltransferase 1-a | 1621 | 0 | 0 | 0 |
| SALT00000056694_Cluster5927-001 | MYB_re | MYB transcription factor [ | 1773 | 0 | 0 | 0 |
| SALT00000077120_Cluster5927-002 | MYB_re | MYB DNA-binding domai | 3804 | 0 | 0 | 0 |
| SALT00000053688_Cluster5927-003 | MYB_re | MYB-related transcription | 1904 | 0 | 0 | 0 |
| SALT00000048478_Cluster5973-001 | MYB_re | ALWAYS EARLY 2 isofor | 1969 | 0 | 0 | 0 |
| SALT00000073364_Cluster6244-004 | MYB_re | Transcriptional adapter AD | 2716 | 0 | 0 | 0 |
| SALT00000085047_Cluster7584-003 | MYB_re | snRNA-activating complex | 3405 | 0 | 0 | 0 |
| SALT00000014153_Cluster8935-001 | MYB_re | MYB DNA-binding domai | 1406 | 0 | 0 | 0 |
| SALT00000037264_Cluster8935-002 | MYB_re | MYB DNA-binding domai | 1375 | 0 | 0 | up |
| SALT00000038836_Cluster8935-003 | MYB_re | MYB DNA-binding domai | 1283 | 0 | 0 | 0 |
| SALT00000088274_Cluster8935-004 | MYB_re | Os10g0561400 [Oryza sati | 3300 | 0 | 0 | 0 |
| SALT00000080698_Cluster9031-002 | MYB_re | translational activator GCN | 3281 | 0 | 0 | 0 |
| SALT00000028482_Cluster9155-002 | MYB_re | flocculation FLO11-like | 2952 | 0 | 0 | 0 |
| SALT00000028737_Cluster9155-003 | MYB_re | flocculation FLO11-like | 2915 | 0 | 0 | 0 |
| SALT00000082343_Cluster9155-005 | MYB_re | flocculation FLO11-like | 2956 | 0 | 0 | 0 |
| SALT00000026238_Cluster9232-001 | MYB_re | ALWAYS EARLY 2-like [ | 3229 | 0 | 0 | 0 |
| SALT00000062391_Cluster9232-002 | MYB_re | ALWAYS EARLY 2-like [ | 3241 | 0 | 0 | 0 |
| SALT00000089964_Cluster9232-003 | MYB_re | ALWAYS EARLY 2-like [ | 3245 | 0 | 0 | 0 |
| SALT00000025413_Cluster9252-001 | MYB_re | LHY isoform X1 [Zea may | 3141 | 0 | 0 | 0 |
| SALT00000025425_Cluster9252-002 | MYB_re | LHY isoform X1 [Zea may | 2966 | 0 | 0 | 0 |
| SALT00000063795_Cluster9252-005 | MYB_re | LHY isoform X1 [Zea may | 3241 | 0 | 0 | 0 |
| SALT00000064505_Cluster9252-006 | MYB_re | LHY isoform X1 [Zea may | 2973 | 0 | 0 | down |
| SALT00000065839_Cluster9252-007 | MYB_re | LHY isoform X1 [Zea may | 2648 | 0 | 0 | 0 |
| SALT00000066510_Cluster9252-008 | MYB_re | LHY isoform X1 [Zea may | 3050 | 0 | 0 | 0 |
| SALT00000068906_Cluster9252-009 | MYB_re | LHY isoform X1 [Zea may | 2953 | 0 | 0 | 0 |
| SALT00000073342_Cluster9252-010 | MYB_re | LHY isoform X1 [Zea may | 3071 | 0 | 0 | 0 |
| SALT00000074775_Cluster9252-011 | MYB_re | LHY isoform X1 [Zea may | 2747 | 0 | 0 | 0 |
| SALT00000077511_Cluster9252-012 | MYB_re | LHY isoform X1 [Zea may | 3005 | 0 | 0 | 0 |
| SALT00000078346_Cluster9252-013 | MYB_re | LHY isoform X1 [Zea may | 2547 | 0 | 0 | 0 |
| SALT00000079580_Cluster9252-014 | MYB_re | LHY isoform X1 [Zea may | 2917 | 0 | 0 | 0 |
| SALT00000080545_Cluster9252-015 | MYB_re | LHY isoform X1 [Zea may | 3229 | 0 | 0 | 0 |
| SALT00000082104_Cluster9252-016 | MYB_re | LHY isoform X1 [Zea may | 3018 | 0 | 0 | 0 |
| SALT00000083213_Cluster9252-017 | MYB_re | LHY isoform X1 [Zea may | 2967 | 0 | 0 | 0 |
| SALT00000084274_Cluster9252-018 | MYB_re | LHY isoform X1 [Zea may | 3063 | 0 | 0 | 0 |
| SALT00000065514_Cluster10933-001 | MYB_re | RADIALIS-like 3 isoform | 2981 | 0 | 0 | 0 |
| SALT00000074606_Cluster11651-001 | MYB_re | single myb histone 6 | 2899 | 0 | 0 | 0 |
| SALT00000077797_Cluster11740-001 | MYB_re | snRNA-activating complex | 2891 | 0 | 0 | 0 |

| | | | | | | |
|---|---|---|---|---|---|---|
| SALT00000024733_Cluster12055-001 | MYB_re | Os02g0551400 [Oryza sati | 2841 | 0 | 0 | 0 |
| SALT00000025805_Cluster12055-002 | MYB_re | Os02g0551400 [Oryza sati | 2862 | 0 | 0 | 0 |
| SALT00000026772_Cluster12055-003 | MYB_re | Os02g0551400 [Oryza sati | 2749 | 0 | 0 | 0 |
| SALT00000026932_Cluster12055-004 | MYB_re | Os02g0551400 [Oryza sati | 2800 | 0 | 0 | 0 |
| SALT00000007362_Cluster12178-001 | MYB_re | Os11g0183700 [Oryza sati | 1580 | 0 | 0 | 0 |
| SALT00000026689_Cluster12181-001 | MYB_re | MYB-related [Saccharum | 2733 | 0 | 0 | 0 |
| SALT00000029580_Cluster12181-002 | MYB_re | MYB-related [Saccharum | 2671 | 0 | 0 | 0 |
| SALT00000046536_Cluster12181-003 | MYB_re | MYB-related [Saccharum | 1993 | 0 | 0 | 0 |
| SALT00000052789_Cluster12181-004 | MYB_re | MYB-related [Saccharum | 2262 | 0 | 0 | 0 |
| SALT00000062677_Cluster12181-005 | MYB_re | MYB-related [Saccharum | 2301 | 0 | 0 | 0 |
| SALT00000083549_Cluster12181-006 | MYB_re | MYB-related [Saccharum | 2851 | 0 | 0 | 0 |
| SALT00000025975_Cluster12584-001 | MYB_re | nascent polypeptide-associ | 2811 | 0 | 0 | 0 |
| SALT00000062574_Cluster12584-003 | MYB_re | nascent polypeptide-associ | 2768 | 0 | 0 | 0 |
| SALT00000067523_Cluster12592-001 | MYB_re | single myb histone 5-like [ | 2600 | 0 | 0 | 0 |
| SALT00000071293_Cluster12592-002 | MYB_re | single myb histone 5-like [ | 2811 | 0 | 0 | 0 |
| SALT00000067322_Cluster12729-002 | MYB_re | SWI SNF complex subunit | 2624 | 0 | 0 | 0 |
| SALT00000057472_Cluster13676-010 | MYB_re | Myb-related 3R-1 | 1547 | 0 | 0 | 0 |
| SALT00000028980_Cluster13830-001 | MYB_re | TSL-kinase interacting 1 [ | 2577 | 0 | 0 | 0 |
| SALT00000068736_Cluster13830-002 | MYB_re | TSL-kinase interacting 1 [ | 2406 | 0 | 0 | 0 |
| SALT00000075759_Cluster13830-003 | MYB_re | TSL-kinase interacting 1 [ | 2707 | 0 | 0 | 0 |
| SALT00000078952_Cluster13830-004 | MYB_re | TSL-kinase interacting 1 [ | 2658 | 0 | 0 | 0 |
| SALT00000084139_Cluster13830-005 | MYB_re | TSL-kinase interacting 1 [ | 2514 | 0 | 0 | 0 |
| SALT00000006045_Cluster13830-006 | MYB_re | TSL-kinase interacting 1 [ | 2293 | 0 | 0 | 0 |
| SALT00000065705_Cluster14061-001 | MYB_re | snRNA-activating comple | 2690 | 0 | 0 | 0 |
| SALT00000009079_Cluster14773-001 | MYB_re | transcription factor TFIIIB | 1624 | 0 | 0 | 0 |
| SALT00000026345_Cluster14773-002 | MYB_re | Transcription factor TFIIIE | 2641 | 0 | 0 | 0 |
| SALT00000060424_Cluster14773-003 | MYB_re | Transcription factor TFIIIE | 2063 | 0 | 0 | 0 |
| SALT00000064293_Cluster14773-004 | MYB_re | Transcription factor TFIIIE | 2493 | 0 | 0 | 0 |
| SALT00000078390_Cluster14773-005 | MYB_re | Transcription factor TFIIIE | 2554 | 0 | 0 | 0 |
| SALT00000058510_Cluster14773-006 | MYB_re | transcription factor TFIIIB | 2162 | 0 | 0 | 0 |
| SALT00000078840_Cluster14906-001 | MYB_re | single myb histone 3 [Zea | 2633 | 0 | 0 | 0 |
| SALT00000015011_Cluster15293-001 | MYB_re | single myb histone 5-like [ | 1486 | 0 | 0 | 0 |
| SALT00000071756_Cluster15293-002 | MYB_re | single myb histone 5-like [ | 2605 | 0 | 0 | 0 |
| SALT00000041930_Cluster16347-001 | MYB_re | REVEILLE 6-like isoform | 2527 | 0 | 0 | 0 |
| SALT00000000366_Cluster16420-001 | MYB_re | Os04g0583900 [Oryza sati | 1636 | 0 | 0 | 0 |
| SALT00000007687_Cluster16420-006 | MYB_re | Os04g0583900 [Oryza sati | 2006 | 0 | 0 | 0 |
| SALT00000011231_Cluster16420-010 | MYB_re | Os04g0583900 [Oryza sati | 1597 | 0 | 0 | 0 |
| SALT00000036346_Cluster16420-014 | MYB_re | Os04g0583900 [Oryza sati | 1563 | 0 | 0 | 0 |
| SALT00000055706_Cluster16420-016 | MYB_re | Os04g0583900 [Oryza sati | 1607 | 0 | 0 | 0 |
| SALT00000061330_Cluster16420-018 | MYB_re | Os04g0583900 [Oryza sati | 2022 | 0 | 0 | 0 |
| SALT00000027348_Cluster16583-004 | MYB_re | transcription factor MYB4 | 1063 | 0 | 0 | 0 |
| SALT00000020013_Cluster16782-001 | MYB_re | dnaJ homolog subfamily C | 1954 | 0 | 0 | 0 |
| SALT00000021347_Cluster16782-002 | MYB_re | dnaJ homolog subfamily C | 2490 | 0 | 0 | 0 |
| SALT00000027388_Cluster16782-003 | MYB_re | dnaJ homolog subfamily C | 2305 | 0 | 0 | 0 |
| SALT00000028024_Cluster16782-004 | MYB_re | dnaJ homolog subfamily C | 2444 | 0 | 0 | 0 |
| SALT00000028336_Cluster16782-005 | MYB_re | dnaJ homolog subfamily C | 2446 | 0 | 0 | 0 |
| SALT00000048179_Cluster16782-006 | MYB_re | dnaJ homolog subfamily C | 2014 | 0 | 0 | 0 |
| SALT00000051012_Cluster16782-007 | MYB_re | dnaJ homolog subfamily C | 1719 | 0 | 0 | 0 |
| SALT00000078868_Cluster16782-008 | MYB_re | dnaJ homolog subfamily C | 2483 | 0 | 0 | up |
| SALT00000070397_Cluster17067-001 | MYB_re | MYB-like transcription fac | 2464 | 0 | 0 | 0 |
| SALT00000069611_Cluster17242-001 | MYB_re | REVEILLE 6-like isoform | 2444 | 0 | 0 | 0 |
| SALT00000007530_Cluster17647-001 | MYB_re | Transcriptional adapter AD | 2295 | 0 | 0 | 0 |
| SALT00000015855_Cluster17647-002 | MYB_re | Transcriptional adapter AD | 2378 | 0 | 0 | 0 |
| SALT00000024852_Cluster17647-003 | MYB_re | Transcriptional adapter AD | 2228 | 0 | 0 | 0 |
| SALT00000078420_Cluster17647-004 | MYB_re | Transcriptional adapter AD | 2311 | 0 | 0 | 0 |
| SALT00000059128_Cluster18053-001 | MYB_re | Myb-related Myb4 | 1921 | 0 | 0 | 0 |
| SALT00000059558_Cluster18053-002 | MYB_re | Myb-related Myb4 | 2291 | 0 | 0 | 0 |
| SALT00000058382_Cluster18053-003 | MYB_re | Os10g0478300 [Oryza sati | 1715 | 0 | 0 | 0 |

| | | | | | | |
|---|---|---|---|---|---|---|
| SALT00000042051_Cluster18363-001 | MYB_re | TPA: MYB DNA-binding | 2238 | 0 | 0 | 0 |
| SALT00000057244_Cluster18842-001 | MYB_re | TPA: protochlorophyllide r | 2181 | 0 | 0 | down |
| SALT00000045561_Cluster19998-009 | MYB_re | R2R3-MYB transcription f | 1752 | 0 | 0 | 0 |
| SALT00000052175_Cluster20260-002 | MYB_re | Myb domain [Theobroma c | 2058 | 0 | 0 | 0 |
| SALT00000005164_Cluster20406-001 | MYB_re | Os05g0449900 [Oryza sati | 2046 | 0 | 0 | 0 |
| SALT00000009456_Cluster20406-002 | MYB_re | Os05g0449900 [Oryza sati | 1890 | 0 | 0 | 0 |
| SALT00000046794_Cluster20406-004 | MYB_re | Os05g0449900 [Oryza sati | 1427 | 0 | 0 | 0 |
| SALT00000054151_Cluster20406-005 | MYB_re | Os05g0449900 [Oryza sati | 1674 | 0 | 0 | 0 |
| SALT00000005266_Cluster20727-001 | MYB_re | TPA: DNA-binding bromo | 1987 | 0 | 0 | 0 |
| SALT00000010371_Cluster20727-002 | MYB_re | TPA: DNA-binding bromo | 1994 | 0 | 0 | 0 |
| SALT00000043021_Cluster20727-003 | MYB_re | TPA: DNA-binding bromo | 1610 | 0 | 0 | 0 |
| SALT00000054150_Cluster20727-004 | MYB_re | TPA: DNA-binding bromo | 2025 | 0 | 0 | up |
| SALT00000002343_Cluster20959-001 | MYB_re | REVEILLE 2-like isoform | 2007 | 0 | 0 | 0 |
| SALT00000050910_Cluster21324-001 | MYB_re | late elongated hypocoty [C | 1983 | 0 | 0 | 0 |
| SALT00000015848_Cluster21437-001 | MYB_re | late elongated hypocoty [C | 1974 | 0 | 0 | 0 |
| SALT00000044688_Cluster21772-002 | MYB_re | Os04g0583900 [Oryza sati | 1811 | 0 | 0 | 0 |
| SALT00000002230_Cluster22043-001 | MYB_re | Os04g0583900 [Oryza sati | 1838 | 0 | 0 | 0 |
| SALT00000004821_Cluster22043-002 | MYB_re | Os04g0583900 [Oryza sati | 1649 | 0 | 0 | 0 |
| SALT00000015769_Cluster22043-004 | MYB_re | Os04g0583900 [Oryza sati | 1768 | 0 | 0 | 0 |
| SALT00000036507_Cluster22043-005 | MYB_re | Os04g0583900 [Oryza sati | 1796 | 0 | 0 | 0 |
| SALT00000044917_Cluster22043-006 | MYB_re | Os04g0583900 [Oryza sati | 1656 | 0 | 0 | 0 |
| SALT00000042634_Cluster24290-001 | MYB_re | MYB27 [Oryza sativa Japo | 1803 | 0 | 0 | 0 |
| SALT00000002851_Cluster24337-001 | MYB_re | ANTHER INDEHISCENC | 1592 | 0 | 0 | 0 |
| SALT00000013341_Cluster24337-002 | MYB_re | ANTHER INDEHISCENC | 1764 | 0 | 0 | 0 |
| SALT00000059829_Cluster24337-003 | MYB_re | ANTHER INDEHISCENC | 1800 | 0 | 0 | 0 |
| SALT00000047459_Cluster24337-004 | MYB_re | hypothetical protein SETIT | 1994 | 0 | 0 | 0 |
| SALT00000000684_Cluster24577-001 | MYB_re | REVEILLE 6-like [Setaria | 1589 | 0 | 0 | up |
| SALT00000008833_Cluster24577-002 | MYB_re | REVEILLE 6-like [Setaria | 1506 | 0 | 0 | 0 |
| SALT00000014093_Cluster24577-003 | MYB_re | REVEILLE 6-like [Setaria | 1507 | 0 | 0 | 0 |
| SALT00000014319_Cluster24577-004 | MYB_re | REVEILLE 6-like [Setaria | 1629 | 0 | 0 | 0 |
| SALT00000017197_Cluster24577-005 | MYB_re | REVEILLE 6-like [Setaria | 1785 | 0 | 0 | 0 |
| SALT00000043004_Cluster24577-006 | MYB_re | REVEILLE 6-like [Setaria | 1551 | 0 | 0 | 0 |
| SALT00000048301_Cluster24577-007 | MYB_re | REVEILLE 6-like [Setaria | 1426 | 0 | 0 | 0 |
| SALT00000056505_Cluster24577-008 | MYB_re | MYB DNA-binding domai | 1525 | 0 | 0 | 0 |
| SALT00000007499_Cluster24577-009 | MYB_re | REVEILLE 6-like [Setaria | 1678 | 0 | 0 | 0 |
| SALT00000052497_Cluster25187-001 | MYB_re | RNA polymerase I termina | 1753 | 0 | 0 | 0 |
| SALT00000043136_Cluster25446-002 | MYB_re | Os01g0192300 [Oryza sati | 1476 | 0 | 0 | 0 |
| SALT00000056085_Cluster25446-003 | MYB_re | Os01g0192300 [Oryza sati | 1454 | 0 | 0 | 0 |
| SALT00000002078_Cluster25887-001 | MYB_re | REVEILLE 6-like isoform | 1710 | 0 | 0 | 0 |
| SALT00000007171_Cluster25887-002 | MYB_re | REVEILLE 6-like isoform | 1717 | 0 | 0 | 0 |
| SALT00000039923_Cluster25887-003 | MYB_re | REVEILLE 6-like isoform | 1565 | 0 | 0 | 0 |
| SALT00000057135_Cluster25887-004 | MYB_re | REVEILLE 6-like isoform | 1634 | 0 | 0 | 0 |
| SALT00000063923_Cluster25887-005 | MYB_re | REVEILLE 6-like isoform | 1617 | 0 | 0 | 0 |
| SALT00000057333_Cluster26080-004 | MYB_re | transcription factor DIVAR | 1582 | 0 | 0 | 0 |
| SALT00000003571_Cluster26092-001 | MYB_re | transcription factor MYB4 | 1440 | 0 | 0 | 0 |
| SALT00000055634_Cluster26092-006 | MYB_re | transcription factor MYB4 | 1408 | 0 | 0 | 0 |
| SALT00000008191_Cluster26328-001 | MYB_re | transcription repressor MY | 1692 | 0 | 0 | 0 |
| SALT00000010451_Cluster26376-001 | MYB_re | Os01g0192300 [Oryza sati | 1690 | 0 | 0 | 0 |
| SALT00000018992_Cluster27074-001 | MYB_re | MYB DNA-binding domai | 1366 | 0 | 0 | 0 |
| SALT00000041179_Cluster27074-002 | MYB_re | MYB DNA-binding domai | 1656 | 0 | 0 | 0 |
| SALT00000018214_Cluster27074-003 | MYB_re | MYB DNA-binding domai | 1188 | 0 | 0 | 0 |
| SALT00000010987_Cluster27286-001 | MYB_re | Transcription factor MYB1 | 1646 | 0 | 0 | 0 |
| SALT00000050031_Cluster27286-002 | MYB_re | Transcription factor MYB1 | 1513 | 0 | 0 | up |
| SALT00000055366_Cluster27286-003 | MYB_re | Transcription factor MYB1 | 1568 | 0 | 0 | 0 |
| SALT00000017202_Cluster28197-001 | MYB_re | myb [Oryza sativa Japonic | 1600 | 0 | 0 | up |
| SALT00000036276_Cluster28727-001 | MYB_re | MYB DNA-binding domai | 1561 | 0 | 0 | 0 |
| SALT00000037421_Cluster28727-002 | MYB_re | myb-related transcription a | 1572 | 0 | 0 | 0 |
| SALT00000042639_Cluster28727-003 | MYB_re | MYB DNA-binding domai | 1551 | 0 | 0 | 0 |

| | | | | | | |
|---|---|---|---|---|---|---|
| SALT00000058114_Cluster28727-004 | MYB_re | MYB DNA-binding domai | 1512 | 0 | 0 | 0 |
| SALT00000001811_Cluster28742-001 | MYB_re | Os01g0589300 [Oryza sati | 1571 | 0 | 0 | 0 |
| SALT00000015388_Cluster28742-002 | MYB_re | Os01g0589300 [Oryza sati | 1386 | 0 | 0 | 0 |
| SALT00000049358_Cluster29564-001 | MYB_re | myb-related Zm38-like | 1529 | 0 | 0 | 0 |
| SALT00000014198_Cluster30824-001 | MYB_re | LOC100282280 isoform X | 1413 | 0 | 0 | 0 |
| SALT00000060587_Cluster30879-001 | MYB_re | Os08g0144000 [Oryza sati | 1433 | 0 | 0 | 0 |
| SALT00000046661_Cluster31225-001 | MYB_re | Os06g0105800 [Oryza sati | 1390 | 0 | 0 | 0 |
| SALT00000048598_Cluster31225-002 | MYB_re | Os06g0105800 [Oryza sati | 1347 | 0 | 0 | 0 |
| SALT00000023509_Cluster31251-001 | MYB_re | Os01g0192300 [Oryza sati | 1108 | 0 | 0 | 0 |
| SALT00000067770_Cluster31251-002 | MYB_re | Os01g0192300 [Oryza sati | 1211 | 0 | 0 | up |
| SALT00000073880_Cluster31251-003 | MYB_re | Os01g0192300 [Oryza sati | 1385 | 0 | 0 | 0 |
| SALT00000044866_Cluster31251-004 | MYB_re | Os01g0192300 [Oryza sati | 1461 | 0 | 0 | 0 |
| SALT00000053813_Cluster31795-001 | MYB_re | MYB-related transcription | 1251 | 0 | 0 | 0 |
| SALT00000004762_Cluster31871-001 | MYB_re | [Hordeum vulgare vulgare | 1229 | 0 | 0 | down |
| SALT00000017033_Cluster275-001 | NAC | NAC domain-containing 8 | 2249 | 0 | 0 | 0 |
| SALT00000019201_Cluster275-002 | NAC | NAC domain-containing 8 | 1868 | 0 | 0 | 0 |
| SALT00000020887_Cluster275-003 | NAC | NAC domain-containing 8 | 2557 | 0 | 0 | 0 |
| SALT00000029496_Cluster275-005 | NAC | NAC domain-containing 8 | 2535 | 0 | 0 | 0 |
| SALT00000035189_Cluster275-006 | NAC | NAC domain-containing 8 | 2511 | 0 | 0 | 0 |
| SALT00000035385_Cluster275-007 | NAC | NAC domain-containing 8 | 2464 | 0 | 0 | 0 |
| SALT00000039948_Cluster275-008 | NAC | NAC domain-containing 8 | 2012 | 0 | 0 | 0 |
| SALT00000059571_Cluster275-011 | NAC | NAC domain-containing 8 | 2173 | 0 | 0 | 0 |
| SALT00000060946_Cluster275-016 | NAC | NAC domain-containing 8 | 2175 | 0 | 0 | 0 |
| SALT00000010830_Cluster1229-001 | NAC | NAC domain-containing 8 | 1898 | 0 | 0 | 0 |
| SALT00000021171_Cluster2112-001 | NAC | Os08g0157900 [Oryza sati | 2830 | 0 | 0 | 0 |
| SALT00000022939_Cluster2112-005 | NAC | Os08g0157900 [Oryza sati | 2567 | 0 | 0 | 0 |
| SALT00000024673_Cluster2112-006 | NAC | Os08g0157900 [Oryza sati | 2647 | 0 | 0 | 0 |
| SALT00000025571_Cluster2112-008 | NAC | Os08g0157900 [Oryza sati | 2834 | 0 | 0 | 0 |
| SALT00000026257_Cluster2112-010 | NAC | Os08g0157900 [Oryza sati | 2525 | 0 | 0 | 0 |
| SALT00000063921_Cluster2112-023 | NAC | [Oryza sativa Japonica Grc | 5068 | 0 | 0 | 0 |
| SALT00000065461_Cluster2112-024 | NAC | Os08g0157900 [Oryza sati | 2836 | 0 | 0 | 0 |
| SALT00000078736_Cluster2112-032 | NAC | NAC domain transcription | 2878 | 0 | 0 | 0 |
| SALT00000021544_Cluster2829-001 | NAC | hypothetical protein BRAI | 2833 | 0 | 0 | 0 |
| SALT00000021652_Cluster2829-002 | NAC | hypothetical protein BRAI | 2701 | 0 | 0 | 0 |
| SALT00000028190_Cluster2829-003 | NAC | hypothetical protein BRAI | 2748 | 0 | 0 | 0 |
| SALT00000071779_Cluster2829-004 | NAC | hypothetical protein BRAI | 2794 | 0 | 0 | 0 |
| SALT00000025794_Cluster2829-006 | NAC | partial [Oryza sativa Japon | 2690 | 0 | 0 | 0 |
| SALT00000025602_Cluster4261-001 | NAC | NAC transcription partial [ | 2611 | 0 | 0 | 0 |
| SALT00000069397_Cluster4261-002 | NAC | NAC transcription partial [ | 3044 | 0 | 0 | 0 |
| SALT00000086150_Cluster4343-001 | NAC | NAC domain transcription | 4130 | 0 | 0 | 0 |
| SALT00000002174_Cluster5577-001 | NAC | NAC transcription partial [ | 2106 | 0 | 0 | 0 |
| SALT00000008686_Cluster5577-002 | NAC | NAC transcription partial [ | 1727 | 0 | 0 | 0 |
| SALT00000011205_Cluster5577-003 | NAC | NAC transcription partial [ | 2167 | 0 | 0 | 0 |
| SALT00000011980_Cluster5577-004 | NAC | NAC transcription partial [ | 2218 | 0 | 0 | 0 |
| SALT00000048303_Cluster5577-005 | NAC | NAC transcription partial [ | 2253 | 0 | 0 | 0 |
| SALT00000052162_Cluster5577-006 | NAC | NAC transcription partial [ | 2202 | 0 | 0 | 0 |
| SALT00000053732_Cluster5577-007 | NAC | 6-phospho-1-fructokinase | | 3867 | up | up | up |
| SALT00000060009_Cluster5577-008 | NAC | NAC transcription partial [ | 2274 | 0 | 0 | 0 |
| SALT00000061665_Cluster5577-009 | NAC | NAC transcription partial [ | 1802 | 0 | 0 | 0 |
| SALT00000030435_Cluster6349-001 | NAC | NAC domain-containing 8 | 2410 | 0 | 0 | 0 |
| SALT00000073006_Cluster6349-002 | NAC | NAC domain-containing 8 | 3737 | 0 | 0 | 0 |
| SALT00000007630_Cluster7422-001 | NAC | NAC domain-containing 9 | 1652 | 0 | 0 | 0 |
| SALT00000047111_Cluster7422-003 | NAC | NAC domain-containing 9 | 1576 | 0 | 0 | 0 |
| SALT00000051833_Cluster7422-004 | NAC | NAC domain-containing 9 | 1492 | 0 | 0 | 0 |
| SALT00000052094_Cluster7422-005 | NAC | NAC domain-containing 9 | 1575 | 0 | 0 | 0 |
| SALT00000061493_Cluster7422-006 | NAC | NAC domain-containing 9 | 1679 | 0 | 0 | 0 |
| SALT00000012285_Cluster7653-001 | NAC | NAC domain-containing 8 | 1653 | 0 | 0 | 0 |
| SALT00000015917_Cluster7653-002 | NAC | NAC domain-containing 8 | 1676 | 0 | 0 | 0 |

| | | | | | | |
|---|---|---|---|---|---|---|
| SALT00000018277_Cluster7653-003 | NAC | NAC domain-containing 8 | 2206 | 0 | 0 | 0 |
| SALT00000020660_Cluster7653-004 | NAC | NAC domain-containing 8 | 2601 | 0 | 0 | 0 |
| SALT00000020719_Cluster7653-005 | NAC | NAC domain-containing 8 | 2628 | 0 | 0 | 0 |
| SALT00000023246_Cluster7653-006 | NAC | NAC domain-containing 8 | 2419 | 0 | 0 | 0 |
| SALT00000024705_Cluster7653-007 | NAC | NAC domain-containing 8 | 2538 | 0 | 0 | 0 |
| SALT00000025208_Cluster7653-008 | NAC | NAC domain-containing 8 | 2715 | 0 | 0 | 0 |
| SALT00000025265_Cluster7653-009 | NAC | NAC domain-containing 8 | 2668 | 0 | 0 | 0 |
| SALT00000026751_Cluster7653-010 | NAC | NAC domain-containing 8 | 2486 | 0 | 0 | 0 |
| SALT00000030909_Cluster7653-011 | NAC | NAC domain-containing 8 | 2587 | 0 | 0 | 0 |
| SALT00000034380_Cluster7653-012 | NAC | NAC domain-containing 8 | 3536 | 0 | 0 | 0 |
| SALT00000042009_Cluster7653-013 | NAC | NAC domain-containing 8 | 1876 | 0 | 0 | 0 |
| SALT00000050626_Cluster7653-014 | NAC | NAC domain-containing 8 | 1615 | 0 | 0 | 0 |
| SALT00000050801_Cluster7653-015 | NAC | NAC domain-containing 8 | 1584 | 0 | 0 | 0 |
| SALT00000053681_Cluster7653-016 | NAC | NAC domain-containing 8 | 1824 | 0 | 0 | 0 |
| SALT00000060940_Cluster7653-017 | NAC | NAC domain-containing 8 | 1727 | 0 | 0 | 0 |
| SALT00000061306_Cluster7653-018 | NAC | NAC domain-containing 8 | 2017 | 0 | 0 | 0 |
| SALT00000062532_Cluster7653-019 | NAC | NAC domain-containing 8 | 2379 | 0 | 0 | 0 |
| SALT00000064192_Cluster7653-020 | NAC | NAC domain-containing 8 | 2367 | 0 | 0 | 0 |
| SALT00000068511_Cluster7653-021 | NAC | NAC domain-containing 8 | 2521 | 0 | 0 | 0 |
| SALT00000070776_Cluster7653-022 | NAC | NAC domain-containing 8 | 2530 | 0 | 0 | 0 |
| SALT00000071286_Cluster7653-023 | NAC | NAC domain-containing 8 | 2583 | 0 | 0 | 0 |
| SALT00000072749_Cluster7653-024 | NAC | NAC domain-containing 8 | 2701 | 0 | 0 | 0 |
| SALT00000074473_Cluster7653-025 | NAC | NAC domain-containing 8 | 2599 | 0 | 0 | 0 |
| SALT00000077595_Cluster7653-026 | NAC | NAC domain-containing 8 | 3178 | 0 | 0 | 0 |
| SALT00000078637_Cluster7653-027 | NAC | NAC domain-containing 8 | 2697 | 0 | 0 | 0 |
| SALT00000079981_Cluster7653-028 | NAC | NAC domain-containing 8 | 2622 | 0 | 0 | 0 |
| SALT00000080909_Cluster7653-029 | NAC | NAC domain-containing 8 | 2508 | 0 | 0 | 0 |
| SALT00000078168_Cluster7653-030 | NAC | NAC domain-containing 8 | 2508 | 0 | 0 | 0 |
| SALT00000080605_Cluster7653-031 | NAC | NAC domain-containing 8 | 2414 | 0 | 0 | 0 |
| SALT00000047122_Cluster7653-032 | NAC | NAC domain-containing 8 | 1617 | 0 | 0 | 0 |
| SALT00000052503_Cluster8116-003 | NAC | NAC2 [Oryza sativa Japon | 1791 | 0 | 0 | 0 |
| SALT00000027847_Cluster9525-001 | NAC | Os08g0157900 [Oryza sati | 2730 | 0 | 0 | 0 |
| SALT00000028656_Cluster9525-002 | NAC | Os08g0157900 [Oryza sati | 2909 | 0 | 0 | 0 |
| SALT00000075576_Cluster9525-003 | NAC | Os08g0157900 [Oryza sati | 3187 | 0 | 0 | 0 |
| SALT00000075714_Cluster9525-004 | NAC | Os08g0157900 [Oryza sati | 2905 | 0 | 0 | 0 |
| SALT00000065880_Cluster10144-001 | NAC | ANAC075 [Zea mays] | 2648 | 0 | 0 | 0 |
| SALT00000086637_Cluster10144-002 | NAC | ANAC075 [Zea mays] | 3089 | 0 | 0 | 0 |
| SALT00000063668_Cluster11158-001 | NAC | secondary wall NAC maste | 2950 | 0 | 0 | 0 |
| SALT00000031066_Cluster11785-004 | NAC | endonuclease exonuclease | 2887 | 0 | 0 | 0 |
| SALT00000000793_Cluster12101-001 | NAC | NAC transcription partial [ | 1924 | 0 | 0 | 0 |
| SALT00000008689_Cluster12101-002 | NAC | NAC transcription partial [ | 1907 | 0 | 0 | 0 |
| SALT00000018991_Cluster12101-003 | NAC | NAC transcription partial [ | 1387 | 0 | 0 | 0 |
| SALT00000050520_Cluster12101-004 | NAC | NAC transcription partial [ | 1988 | 0 | 0 | 0 |
| SALT00000060948_Cluster12101-005 | NAC | NAC transcription partial [ | 1905 | 0 | 0 | 0 |
| SALT00000066045_Cluster12101-006 | NAC | NAC transcription partial [ | 2858 | 0 | 0 | 0 |
| SALT00000068103_Cluster13102-001 | NAC | NAC transcription partial [ | 2765 | 0 | 0 | 0 |
| SALT00000030671_Cluster13241-001 | NAC | NAC2 [Oryza sativa Japon | 2521 | 0 | 0 | 0 |
| SALT00000063326_Cluster13241-003 | NAC | NAC2 [Oryza sativa Japon | 2243 | 0 | 0 | 0 |
| SALT00000021432_Cluster13328-001 | NAC | PREDICTED: uncharacter | 2713 | 0 | 0 | 0 |
| SALT00000023637_Cluster13328-002 | NAC | PREDICTED: uncharacter | 2746 | 0 | 0 | 0 |
| SALT00000074704_Cluster13328-003 | NAC | hypothetical protein SORE | 2562 | 0 | 0 | 0 |
| SALT00000014052_Cluster13488-001 | NAC | NAC domain-containing 8 | 1866 | 0 | 0 | 0 |
| SALT00000021005_Cluster13488-002 | NAC | NAC domain-containing 8 | 2730 | 0 | 0 | 0 |
| SALT00000062663_Cluster13488-003 | NAC | NAC domain-containing 8 | 2733 | 0 | 0 | 0 |
| SALT00000021007_Cluster14225-001 | NAC | Os08g0157900 [Oryza sati | 2629 | 0 | 0 | 0 |
| SALT00000027567_Cluster14225-002 | NAC | Os08g0157900 [Oryza sati | 2525 | 0 | 0 | 0 |
| SALT00000066039_Cluster14225-004 | NAC | Os08g0157900 [Oryza sati | 2636 | 0 | 0 | 0 |
| SALT00000074240_Cluster14225-005 | NAC | Os08g0157900 [Oryza sati | 2615 | 0 | 0 | 0 |

| | | | | | | |
|---|---|---|---|---|---|---|
| SALT00000080624_Cluster14225-006 | NAC | Os08g0157900 [Oryza sati | 2662 | 0 | 0 | 0 |
| SALT00000080978_Cluster14225-007 | NAC | Os08g0157900 [Oryza sati | 2677 | 0 | 0 | 0 |
| SALT00000083044_Cluster14225-008 | NAC | Os08g0157900 [Oryza sati | 2667 | 0 | 0 | 0 |
| SALT00000076391_Cluster14940-001 | NAC | NAC2 [Oryza sativa Japon | 2528 | 0 | 0 | 0 |
| SALT00000078335_Cluster14940-002 | NAC | NAC2 [Oryza sativa Japon | 2630 | 0 | 0 | 0 |
| SALT00000065915_Cluster15400-001 | NAC | [Oryza sativa Japonica Grc | 2596 | 0 | 0 | 0 |
| SALT00000079570_Cluster15400-002 | NAC | [Oryza sativa Japonica Grc | 2598 | 0 | 0 | 0 |
| SALT00000072121_Cluster15400-003 | NAC | [Oryza sativa Japonica Grc | 1054 | 0 | 0 | 0 |
| SALT00000003444_Cluster15400-004 | NAC | [Oryza sativa Japonica Grc | 1918 | 0 | 0 | 0 |
| SALT00000021203_Cluster15711-001 | NAC | NAC domain transcription | 2536 | 0 | 0 | 0 |
| SALT00000068908_Cluster15711-002 | NAC | NAC domain transcription | 2573 | 0 | 0 | 0 |
| SALT00000080177_Cluster15711-003 | NAC | NAC domain transcription | 2491 | 0 | 0 | 0 |
| SALT00000080970_Cluster15711-004 | NAC | NAC domain transcription | 2513 | 0 | 0 | 0 |
| SALT00000093801_Cluster15711-005 | NAC | NAC domain transcription | 2492 | 0 | 0 | 0 |
| SALT00000021276_Cluster15711-006 | NAC | NAC domain transcription | 2494 | 0 | 0 | 0 |
| SALT00000002627_Cluster16418-001 | NAC | Os11g0184900 [Oryza sati | 1788 | 0 | 0 | 0 |
| SALT00000013230_Cluster16418-002 | NAC | Os11g0184900 [Oryza sati | 1802 | 0 | 0 | 0 |
| SALT00000039544_Cluster16418-003 | NAC | Os11g0184900 [Oryza sati | 1796 | 0 | 0 | 0 |
| SALT00000083502_Cluster16418-004 | NAC | NAC domain transcription | 2523 | 0 | 0 | 0 |
| SALT00000045430_Cluster17526-001 | NAC | NAC2 [Oryza sativa Japon | 2313 | 0 | 0 | 0 |
| SALT00000048040_Cluster17526-002 | NAC | NAC2 [Oryza sativa Japon | 2209 | 0 | 0 | 0 |
| SALT00000065778_Cluster17526-003 | NAC | NAC2 [Oryza sativa Japon | 2275 | 0 | 0 | 0 |
| SALT00000071751_Cluster17526-004 | NAC | NAC2 [Oryza sativa Japon | 2401 | 0 | 0 | 0 |
| SALT00000000482_Cluster17869-001 | NAC | NAC domain transcription | 1790 | 0 | 0 | 0 |
| SALT00000015089_Cluster17869-002 | NAC | NAC domain transcription | 1703 | 0 | 0 | 0 |
| SALT00000028041_Cluster17869-003 | NAC | NAC domain transcription | 2328 | 0 | 0 | 0 |
| SALT00000045049_Cluster17869-005 | NAC | Os11g0184900 [Oryza sati | 1880 | 0 | 0 | 0 |
| SALT00000065761_Cluster17869-006 | NAC | Os11g0184900 [Oryza sati | 1883 | 0 | 0 | up |
| SALT00000076948_Cluster17869-007 | NAC | NAC domain transcription | 2280 | 0 | 0 | 0 |
| SALT00000052874_Cluster17869-008 | NAC | NAC domain transcription | 1842 | 0 | 0 | up |
| SALT00000018382_Cluster18322-001 | NAC | NAC domain-containing 2 | 2244 | 0 | 0 | 0 |
| SALT00000040598_Cluster18322-002 | NAC | NAC domain-containing 2 | 2022 | 0 | 0 | 0 |
| SALT00000058284_Cluster18322-003 | NAC | NAC domain-containing 2 | 2188 | 0 | 0 | 0 |
| SALT00000061340_Cluster18322-004 | NAC | NAC domain-containing 2 | 1112 | 0 | 0 | 0 |
| SALT00000039247_Cluster18506-001 | NAC | NAC2 [Oryza sativa Japon | 2063 | 0 | 0 | 0 |
| SALT00000055415_Cluster18506-002 | NAC | NAC2 [Oryza sativa Japon | 2186 | 0 | 0 | 0 |
| SALT00000059746_Cluster18506-003 | NAC | NAC2 [Oryza sativa Japon | 2219 | 0 | 0 | 0 |
| SALT00000025889_Cluster18506-004 | NAC | NAC2 [Oryza sativa Japon | 2609 | 0 | 0 | 0 |
| SALT00000001555_Cluster18550-001 | NAC | NAC2 [Oryza sativa Japon | 2162 | 0 | 0 | up |
| SALT00000015260_Cluster18550-002 | NAC | NAC2 [Oryza sativa Japon | 2036 | 0 | 0 | 0 |
| SALT00000023096_Cluster18550-003 | NAC | NAC2 [Oryza sativa Japon | 2212 | 0 | 0 | 0 |
| SALT00000004673_Cluster18579-001 | NAC | NAC2 [Oryza sativa Japon | 2128 | 0 | 0 | 0 |
| SALT00000007892_Cluster18579-002 | NAC | NAC2 [Oryza sativa Japon | 2135 | 0 | 0 | 0 |
| SALT00000043605_Cluster18579-004 | NAC | NAC2 [Oryza sativa Japon | 2209 | 0 | 0 | 0 |
| SALT00000075150_Cluster18579-006 | NAC | NAC2 [Oryza sativa Japon | 2181 | 0 | 0 | 0 |
| SALT00000050558_Cluster18579-007 | NAC | NAC2 [Oryza sativa Japon | 2011 | 0 | 0 | up |
| SALT00000004307_Cluster19615-001 | NAC | NAC transcription factor [' | 1415 | up | 0 | up |
| SALT00000005310_Cluster19615-002 | NAC | NAC transcription factor [' | 1420 | 0 | 0 | up |
| SALT00000007307_Cluster19615-003 | NAC | NAC transcription factor [' | 1293 | 0 | 0 | 0 |
| SALT00000011426_Cluster19615-004 | NAC | NAC transcription factor [' | 1493 | down | 0 | up |
| SALT00000038775_Cluster19615-005 | NAC | NAC transcription factor [' | 2111 | 0 | up | 0 |
| SALT00000057157_Cluster19615-006 | NAC | NAC transcription factor [' | 1468 | 0 | up | up |
| SALT00000055238_Cluster20050-002 | NAC | Os01g0816100 [Oryza sati | 2076 | 0 | 0 | 0 |
| SALT00000013118_Cluster20092-001 | NAC | NAC2 [Oryza sativa Japon | 2072 | 0 | 0 | 0 |
| SALT00000059574_Cluster20092-002 | NAC | NAC2 [Oryza sativa Japon | 2020 | 0 | 0 | 0 |
| SALT00000011605_Cluster20280-001 | NAC | NAC domain transcription | 2056 | 0 | 0 | 0 |
| SALT00000017373_Cluster20280-002 | NAC | NAC domain transcription | 1560 | 0 | 0 | 0 |
| SALT00000010727_Cluster21221-001 | NAC | NAC domain transcription | 1989 | 0 | 0 | 0 |

| | | | | | | |
|---|---|---|---|---|---|---|
| SALT00000017739_Cluster21221-003 | NAC | NAC domain transcription | 1986 | 0 | 0 | 0 |
| SALT00000054140_Cluster21221-004 | NAC | NAC domain transcription | 1966 | 0 | 0 | 0 |
| SALT00000005257_Cluster21960-001 | NAC | TPA: NAC domain transcr | 1889 | 0 | 0 | 0 |
| SALT00000060776_Cluster21960-002 | NAC | NAC domain-containing 4 | 1942 | 0 | up | up |
| SALT00000040067_Cluster21960-003 | NAC | NAC domain transcription | 1338 | 0 | up | up |
| SALT00000046140_Cluster22207-001 | NAC | NAC transcription factor | 1926 | 0 | 0 | 0 |
| SALT00000054354_Cluster23285-001 | NAC | ATAF2 isoform X2 [Oryza | 1860 | 0 | 0 | 0 |
| SALT00000049998_Cluster23463-001 | NAC | secondary wall NAC maste | 1850 | 0 | 0 | 0 |
| SALT00000011442_Cluster24200-001 | NAC | ANAC075 [Zea mays] | 1627 | 0 | 0 | 0 |
| SALT00000051633_Cluster24200-002 | NAC | ANAC075 [Zea mays] | 1808 | 0 | 0 | 0 |
| SALT00000052136_Cluster24200-003 | NAC | ANAC075 [Zea mays] | 1656 | 0 | 0 | 0 |
| SALT00000060327_Cluster24200-004 | NAC | ANAC075 [Zea mays] | 1676 | 0 | 0 | 0 |
| SALT00000036529_Cluster24452-001 | NAC | NAC domain-containing 7 | 1793 | 0 | 0 | 0 |
| SALT00000003097_Cluster24596-001 | NAC | NAC domain transcription | 1784 | 0 | 0 | 0 |
| SALT00000013233_Cluster24596-002 | NAC | NAC domain transcription | 1768 | 0 | 0 | 0 |
| SALT00000015069_Cluster24596-003 | NAC | NAC domain transcription | 1679 | 0 | 0 | 0 |
| SALT00000016522_Cluster24596-004 | NAC | NAC domain transcription | 1448 | 0 | 0 | 0 |
| SALT00000017963_Cluster24596-005 | NAC | NAC domain transcription | 1735 | 0 | 0 | 0 |
| SALT00000043628_Cluster24596-006 | NAC | NAC domain transcription | 1721 | 0 | 0 | 0 |
| SALT00000052843_Cluster24596-007 | NAC | NAC domain transcription | 1652 | 0 | 0 | 0 |
| SALT00000001372_Cluster24844-001 | NAC | GRAB2 [Zea mays] | 1615 | 0 | 0 | up |
| SALT00000001759_Cluster24844-002 | NAC | GRAB2 [Zea mays] | 1690 | 0 | 0 | up |
| SALT00000002553_Cluster24844-003 | NAC | GRAB2 [Zea mays] | 1735 | 0 | 0 | up |
| SALT00000016658_Cluster24844-004 | NAC | GRAB2 [Zea mays] | 1654 | 0 | 0 | 0 |
| SALT00000040743_Cluster24844-005 | NAC | GRAB2 [Zea mays] | 1771 | 0 | 0 | up |
| SALT00000053787_Cluster24844-006 | NAC | GRAB2 [Zea mays] | 1567 | 0 | 0 | 0 |
| SALT00000055243_Cluster24844-007 | NAC | GRAB2 [Zea mays] | 1601 | 0 | 0 | up |
| SALT00000008568_Cluster24971-001 | NAC | NAC transcription factor 2 | 1764 | 0 | up | 0 |
| SALT00000060430_Cluster24971-002 | NAC | NAC transcription factor 2 | 1587 | 0 | 0 | up |
| SALT00000016136_Cluster25024-001 | NAC | secondary wall NAC maste | 1719 | 0 | 0 | 0 |
| SALT00000022883_Cluster25024-002 | NAC | secondary wall NAC maste | 1761 | 0 | 0 | 0 |
| SALT00000036687_Cluster25378-001 | NAC | NAC transcription factor [ | 1743 | 0 | 0 | 0 |
| SALT00000004735_Cluster25463-001 | NAC | Os01g0888300 [Oryza sati | 1738 | 0 | 0 | 0 |
| SALT00000016666_Cluster25463-002 | NAC | NAC transcription factor [ | 1116 | 0 | 0 | 0 |
| SALT00000019811_Cluster25463-003 | NAC | NAC transcription factor [ | 1631 | 0 | 0 | 0 |
| SALT00000000895_Cluster25479-001 | NAC | GRAB2 [Zea mays] | 1733 | 0 | 0 | 0 |
| SALT00000003766_Cluster25479-002 | NAC | GRAB2 [Zea mays] | 1353 | 0 | 0 | up |
| SALT00000005022_Cluster25479-003 | NAC | GRAB2 [Zea mays] | 1641 | 0 | 0 | 0 |
| SALT00000009722_Cluster25479-004 | NAC | GRAB2 [Zea mays] | 1632 | 0 | 0 | up |
| SALT00000013637_Cluster25479-005 | NAC | GRAB2 [Zea mays] | 1659 | 0 | 0 | 0 |
| SALT00000047958_Cluster25479-006 | NAC | GRAB2 [Zea mays] | 1738 | 0 | 0 | up |
| SALT00000064975_Cluster25479-007 | NAC | GRAB2 [Zea mays] | 1732 | 0 | 0 | up |
| SALT00000039856_Cluster25519-001 | NAC | secondary wall NAC maste | 1636 | 0 | 0 | 0 |
| SALT00000040828_Cluster25519-002 | NAC | secondary wall NAC maste | 1736 | 0 | 0 | 0 |
| SALT00000049838_Cluster25968-001 | NAC | Os02g0594800 [Oryza sati | 1714 | 0 | 0 | 0 |
| SALT00000001451_Cluster26335-001 | NAC | NAC1 [Sorghum bicolor] | 1347 | 0 | 0 | 0 |
| SALT00000006077_Cluster26335-002 | NAC | NAC1 [Sorghum bicolor] | 1464 | 0 | 0 | 0 |
| SALT00000007873_Cluster26335-003 | NAC | NAC1 [Sorghum bicolor] | 1358 | 0 | 0 | up |
| SALT00000016264_Cluster26335-004 | NAC | NAC1 [Sorghum bicolor] | 1692 | 0 | 0 | 0 |
| SALT00000019494_Cluster26335-005 | NAC | NAC1 [Sorghum bicolor] | 1433 | 0 | 0 | 0 |
| SALT00000061663_Cluster26335-006 | NAC | NAC1 [Sorghum bicolor] | 1649 | 0 | 0 | up |
| SALT00000043772_Cluster26479-001 | NAC | NAC domain-containing 7 | 1685 | 0 | 0 | 0 |
| SALT00000016429_Cluster26767-001 | NAC | hypothetical protein SETI | 1671 | 0 | 0 | 0 |
| SALT00000044365_Cluster26810-001 | NAC | Os01g0816100 [Oryza sati | 1669 | 0 | 0 | 0 |
| SALT00000053707_Cluster26810-002 | NAC | 4 [Oryza sativa Japonica G | 1419 | 0 | 0 | 0 |
| SALT00000019435_Cluster26901-001 | NAC | no apical meristem (NAM | 1454 | 0 | 0 | 0 |
| SALT00000036717_Cluster26901-002 | NAC | no apical meristem (NAM | 1664 | 0 | 0 | 0 |
| SALT00000044708_Cluster26901-003 | NAC | no apical meristem (NAM | 1597 | 0 | 0 | 0 |

| | | | | | | |
|---|---|---|---|---|---|---|
| SALT00000045037_Cluster26901-004 | NAC | no apical meristem (NAM) | 1463 | 0 | 0 | 0 |
| SALT00000048633_Cluster26901-005 | NAC | no apical meristem (NAM) | 1613 | 0 | 0 | 0 |
| SALT00000058189_Cluster26901-006 | NAC | no apical meristem (NAM) | 1456 | 0 | 0 | 0 |
| SALT00000002684_Cluster27315-001 | NAC | NAC domain-containing 4 | 1644 | 0 | 0 | 0 |
| SALT00000004635_Cluster27759-001 | NAC | NAC transcription factor 2 | 1596 | 0 | 0 | up |
| SALT00000044788_Cluster27759-002 | NAC | Os11g0126900 [Oryza sati | 1623 | 0 | 0 | 0 |
| SALT00000048350_Cluster27759-003 | NAC | NAC transcription factor 2 | 1535 | 0 | 0 | 0 |
| SALT00000004080_Cluster27759-004 | NAC | NAC transcription factor 2 | 1547 | 0 | 0 | up |
| SALT00000047254_Cluster27876-001 | NAC | NAC transcription factor 1 | 1617 | 0 | 0 | 0 |
| SALT00000044424_Cluster27938-001 | NAC | NAC domain-containing 8 | 1601 | 0 | 0 | 0 |
| SALT00000051009_Cluster27938-002 | NAC | NAC domain-containing 8 | 1521 | 0 | 0 | 0 |
| SALT00000051242_Cluster27938-003 | NAC | NAC domain-containing 8 | 1614 | 0 | 0 | 0 |
| SALT00000043538_Cluster28031-001 | NAC | NAC domain-containing 9 | 1609 | 0 | 0 | 0 |
| SALT00000041801_Cluster28253-001 | NAC | NAC domain-containing 4 | 1597 | up | up | up |
| SALT00000038763_Cluster28923-001 | NAC | NAC domain-containing 1 | 1563 | 0 | 0 | 0 |
| SALT00000048066_Cluster28941-001 | NAC | NAC domain-containing 4 | 1562 | 0 | 0 | 0 |
| SALT00000001663_Cluster28959-001 | NAC | NAC1 [Zea mays] | 1323 | up | up | up |
| SALT00000043227_Cluster28959-002 | NAC | NAC1 [Zea mays] | 1561 | 0 | 0 | up |
| SALT00000055369_Cluster28959-003 | NAC | NAC1 [Zea mays] | 1310 | 0 | 0 | 0 |
| SALT00000037996_Cluster28982-001 | NAC | Os03g0133000 [Oryza sati | 1320 | 0 | 0 | 0 |
| SALT00000058697_Cluster28982-002 | NAC | TPA: NAC domain transcr | 1560 | 0 | 0 | 0 |
| SALT00000055115_Cluster28982-003 | NAC | Os03g0133000 [Oryza sati | 1209 | 0 | 0 | 0 |
| SALT00000012452_Cluster29067-001 | NAC | NAC transcription factor 2 | 1555 | 0 | 0 | 0 |
| SALT00000050160_Cluster29067-002 | NAC | NAC transcription factor 2 | 1453 | 0 | 0 | 0 |
| SALT00000058724_Cluster29067-003 | NAC | NAC transcription factor 2 | 1399 | 0 | 0 | 0 |
| SALT00000002007_Cluster29067-004 | NAC | NAC transcription factor 2 | 1479 | 0 | 0 | 0 |
| SALT00000006222_Cluster29514-001 | NAC | no apical meristem [Zea m | 1333 | 0 | 0 | up |
| SALT00000040591_Cluster29514-002 | NAC | no apical meristem [Zea m | 1532 | 0 | 0 | up |
| SALT00000010488_Cluster29630-001 | NAC | NAC transcription factor 1 | 1524 | 0 | 0 | 0 |
| SALT00000041765_Cluster29846-001 | NAC | Os01g0816100 [Oryza sati | 1495 | 0 | 0 | 0 |
| SALT00000053055_Cluster29846-002 | NAC | Os01g0816100 [Oryza sati | 1512 | 0 | 0 | 0 |
| SALT00000009971_Cluster30123-001 | NAC | NAC domain-containing 4 | 1356 | 0 | down | 0 |
| SALT00000013838_Cluster30123-002 | NAC | NAC domain-containing 4 | 1493 | 0 | 0 | 0 |
| SALT00000042510_Cluster30123-003 | NAC | NAC domain-containing 4 | 1354 | 0 | 0 | up |
| SALT00000038061_Cluster30128-001 | NAC | ATAF2-like [Setaria italica | 1493 | 0 | 0 | 0 |
| SALT00000054185_Cluster30135-002 | NAC | NAC domain transcription | 1493 | 0 | 0 | 0 |
| SALT00000040896_Cluster30156-001 | NAC | NAC1 [Sorghum bicolor] | 1491 | 0 | 0 | up |
| SALT00000050013_Cluster30156-002 | NAC | NAC1 [Sorghum bicolor] | 1392 | 0 | 0 | 0 |
| SALT00000055786_Cluster30156-003 | NAC | NAC1 [Sorghum bicolor] | 1451 | 0 | 0 | up |
| SALT00000059370_Cluster30326-001 | NAC | NAC domain-containing 6 | 1480 | 0 | 0 | 0 |
| SALT00000079880_Cluster30326-002 | NAC | NAC domain-containing 6 | 1244 | 0 | 0 | 0 |
| SALT00000044545_Cluster30326-003 | NAC | NAC domain-containing 6 | 1340 | 0 | 0 | 0 |
| SALT00000038730_Cluster30465-001 | NAC | NAC domain transcription | 1470 | 0 | 0 | 0 |
| SALT00000010107_Cluster30510-001 | NAC | NAC1 [Zea mays] | 1465 | 0 | 0 | 0 |
| SALT00000036376_Cluster30510-002 | NAC | NAC1 [Sorghum bicolor] | 1467 | 0 | 0 | 0 |
| SALT00000045575_Cluster30510-003 | NAC | NAC1 [Zea mays] | 1459 | 0 | 0 | up |
| SALT00000011262_Cluster30667-001 | NAC | NAC transcription factor 2 | 1405 | 0 | up | up |
| SALT00000011992_Cluster30667-002 | NAC | NAC transcription factor 2 | 1454 | 0 | 0 | up |
| SALT00000046504_Cluster30676-001 | NAC | NAC transcription factor 2 | 1454 | 0 | 0 | up |
| SALT00000042767_Cluster30741-001 | NAC | SNAC1 [Oryza nivara] | 1448 | 0 | 0 | 0 |
| SALT00000044982_Cluster30741-002 | NAC | NAC1 [Zea mays] | 1444 | 0 | 0 | up |
| SALT00000003648_Cluster31570-001 | NAC | NAC domain-containing 6 | 1314 | 0 | 0 | 0 |
| SALT00000035992_Cluster31570-002 | NAC | NAC domain-containing 6 | 1270 | 0 | 0 | 0 |
| SALT00000000770_Cluster31570-003 | NAC | NAC domain-containing 6 | 1323 | 0 | 0 | up |
| SALT00000016093_Cluster32036-001 | NAC | NAC domain transcription | 1177 | 0 | 0 | 0 |
| SALT00000029163_Cluster5749-002 | NF-X1 | NF-X1-type zinc finger NI | 3711 | 0 | 0 | up |
| SALT00000032497_Cluster5749-003 | NF-X1 | NF-X1-type zinc finger NI | 3661 | 0 | 0 | 0 |
| SALT00000080986_Cluster5749-004 | NF-X1 | NF-X1-type zinc finger NI | 2428 | 0 | 0 | 0 |

| | | | | | | |
|---|---|---|---|---|---|---|
| SALT00000084791_Cluster5749-005 | NF-X1 | NF-X1-type zinc finger NI | 3835 | 0 | 0 | 0 |
| SALT00000088449_Cluster5749-006 | NF-X1 | NF-X1-type zinc finger NI | 3759 | 0 | 0 | 0 |
| SALT00000091332_Cluster5749-007 | NF-X1 | NF-X1-type zinc finger NI | 3488 | 0 | 0 | 0 |
| SALT00000093698_Cluster5749-008 | NF-X1 | NF-X1-type zinc finger NI | 3766 | 0 | 0 | 0 |
| SALT00000086251_Cluster6366-001 | NF-X1 | NF-X1-type zinc finger NI | 3734 | 0 | 0 | 0 |
| SALT00000028374_Cluster10564-002 | NF-X1 | NF-X1-type zinc finger NI | 2958 | 0 | 0 | 0 |
| SALT00000069135_Cluster10564-003 | NF-X1 | NF-X1-type zinc finger NI | 3028 | 0 | 0 | 0 |
| SALT00000087172_Cluster2920-001 | NF-YA | CCAAT-binding transcript | 4637 | 0 | 0 | 0 |
| SALT00000071764_Cluster5572-001 | NF-YA | nuclear transcription factor | 3868 | 0 | 0 | 0 |
| SALT00000088233_Cluster6930-001 | NF-YA | nuclear transcription factor | 3647 | 0 | 0 | up |
| SALT00000074489_Cluster15947-002 | NF-YA | nuclear transcription factor | 2556 | 0 | 0 | 0 |
| SALT00000009674_Cluster16262-001 | NF-YA | nuclear transcription factor | 1114 | 0 | 0 | 0 |
| SALT00000081838_Cluster16262-002 | NF-YA | nuclear transcription factor | 2533 | 0 | 0 | 0 |
| SALT00000051082_Cluster16262-003 | NF-YA | [Hordeum vulgare vulgare | 1208 | 0 | 0 | 0 |
| SALT00000044183_Cluster19545-001 | NF-YA | nuclear transcription factor | 1472 | 0 | 0 | 0 |
| SALT00000082231_Cluster19545-002 | NF-YA | nuclear transcription factor | 2117 | 0 | 0 | 0 |
| SALT00000045498_Cluster20227-001 | NF-YA | nuclear transcription factor | 2061 | 0 | 0 | 0 |
| SALT00000057804_Cluster20926-001 | NF-YA | nuclear transcription factor | 2010 | 0 | 0 | 0 |
| SALT00000002843_Cluster21509-001 | NF-YA | nuclear transcription factor | 1731 | 0 | 0 | 0 |
| SALT00000005751_Cluster21509-002 | NF-YA | nuclear transcription factor | 1514 | 0 | 0 | 0 |
| SALT00000006263_Cluster21509-003 | NF-YA | nuclear transcription factor | 1713 | 0 | 0 | 0 |
| SALT00000008396_Cluster21509-004 | NF-YA | nuclear transcription factor | 1585 | 0 | 0 | 0 |
| SALT00000014592_Cluster21509-005 | NF-YA | nuclear transcription factor | 1879 | 0 | 0 | 0 |
| SALT00000016706_Cluster21509-006 | NF-YA | nuclear transcription factor | 1786 | 0 | 0 | 0 |
| SALT00000040149_Cluster21509-007 | NF-YA | nuclear transcription factor | 1667 | 0 | 0 | 0 |
| SALT00000042313_Cluster21509-008 | NF-YA | nuclear transcription factor | 1521 | 0 | 0 | 0 |
| SALT00000043127_Cluster21509-009 | NF-YA | nuclear transcription factor | 1970 | 0 | 0 | 0 |
| SALT00000039126_Cluster21509-010 | NF-YA | nuclear transcription factor | 1657 | 0 | 0 | 0 |
| SALT00000002940_Cluster22544-001 | NF-YA | nuclear transcription factor | 1603 | 0 | 0 | 0 |
| SALT00000049243_Cluster22544-002 | NF-YA | nuclear transcription factor | 1906 | 0 | 0 | 0 |
| SALT00000050501_Cluster22544-003 | NF-YA | nuclear transcription factor | 1654 | 0 | 0 | 0 |
| SALT00000061318_Cluster22544-004 | NF-YA | nuclear transcription factor | 1678 | 0 | 0 | 0 |
| SALT00000038105_Cluster24403-001 | NF-YA | nuclear transcription factor | 1612 | 0 | 0 | 0 |
| SALT00000045686_Cluster24403-002 | NF-YA | nuclear transcription factor | 1403 | 0 | 0 | 0 |
| SALT00000051186_Cluster24403-003 | NF-YA | nuclear transcription factor | 1796 | 0 | 0 | 0 |
| SALT00000016198_Cluster24403-004 | NF-YA | nuclear transcription factor | 1510 | 0 | 0 | 0 |
| SALT00000001819_Cluster24572-001 | NF-YA | nuclear transcription factor | 1785 | 0 | 0 | 0 |
| SALT00000041651_Cluster24572-002 | NF-YA | nuclear transcription factor | 1676 | 0 | 0 | 0 |
| SALT00000043683_Cluster24572-003 | NF-YA | nuclear transcription factor | 1665 | 0 | 0 | 0 |
| SALT00000054267_Cluster24572-004 | NF-YA | nuclear transcription factor | 1639 | 0 | 0 | 0 |
| SALT00000036635_Cluster26549-001 | NF-YA | nuclear transcription factor | 1682 | 0 | 0 | 0 |
| SALT00000048258_Cluster26575-001 | NF-YA | 60S ribosomal mitochondr | 1681 | 0 | 0 | 0 |
| SALT00000053989_Cluster27334-001 | NF-YA | CCAAT-box transcription | 1644 | 0 | 0 | 0 |
| SALT00000041873_Cluster28631-001 | NF-YA | Os12g0613000 [Oryza sati | 1577 | 0 | 0 | 0 |
| SALT00000046406_Cluster29544-001 | NF-YA | nuclear transcription factor | 1530 | 0 | 0 | 0 |
| SALT00000038603_Cluster29850-001 | NF-YA | nuclear transcription factor | 1401 | 0 | 0 | 0 |
| SALT00000046068_Cluster29850-002 | NF-YA | nuclear transcription factor | 1366 | 0 | 0 | 0 |
| SALT00000058035_Cluster29850-003 | NF-YA | nuclear transcription factor | 1512 | 0 | 0 | 0 |
| SALT00000049565_Cluster29850-004 | NF-YA | nuclear transcription factor | 1454 | 0 | 0 | 0 |
| SALT00000047713_Cluster31006-001 | NF-YA | nuclear transcription factor | 1419 | 0 | 0 | 0 |
| SALT00000023465_Cluster21499-001 | NF-YB | repressor [Zea mays] | 1624 | 0 | 0 | 0 |
| SALT00000061310_Cluster21499-002 | NF-YB | Dr1 homolog | 1971 | 0 | 0 | 0 |
| SALT00000062409_Cluster21499-003 | NF-YB | Dr1 homolog | 1743 | 0 | 0 | 0 |
| SALT00000017179_Cluster21850-001 | NF-YB | repressor [Zea mays] | 1676 | 0 | 0 | 0 |
| SALT00000037330_Cluster21850-002 | NF-YB | repressor [Zea mays] | 1949 | 0 | 0 | 0 |
| SALT00000002826_Cluster24447-001 | NF-YB | repressor [Zea mays] | 1412 | 0 | 0 | 0 |
| SALT00000008972_Cluster24447-002 | NF-YB | repressor [Zea mays] | 1793 | 0 | 0 | 0 |
| SALT00000042598_Cluster24447-003 | NF-YB | repressor [Zea mays] | 1368 | 0 | 0 | 0 |

| | | | | | | |
|---|---|---|---|---|---|---|
| SALT00000047497_Cluster30845-001 | NF-YB | Nuclear transcription facto | 1437 | 0 | 0 | 0 |
| SALT00000020141_Cluster32519-001 | NF-YB | Os05g0573500 [Oryza sati | 863 | 0 | 0 | 0 |
| SALT00000037919_Cluster5236-001 | NF-YC | nuclear transcription factor | 1190 | 0 | 0 | 0 |
| SALT00000007434_Cluster10931-001 | NF-YC | nuclear transcription factor | 1372 | 0 | 0 | 0 |
| SALT00000024024_Cluster10931-002 | NF-YC | nuclear transcription factor | 2981 | 0 | 0 | 0 |
| SALT00000043844_Cluster10931-003 | NF-YC | nuclear transcription factor | 1589 | 0 | 0 | 0 |
| SALT00000062758_Cluster11246-001 | NF-YC | nuclear transcription factor | 2941 | 0 | 0 | 0 |
| SALT00000042068_Cluster20162-001 | NF-YC | nuclear transcription factor | 2066 | 0 | 0 | 0 |
| SALT00000055180_Cluster22442-001 | NF-YC | Cytochrome P450 86A2 | 1911 | 0 | 0 | 0 |
| SALT00000042142_Cluster24387-001 | NF-YC | repressor [Zea mays] | 1797 | 0 | 0 | 0 |
| SALT00000046887_Cluster28257-001 | NF-YC | dr1-associated corepressor | 1597 | 0 | 0 | 0 |
| SALT00000033253_Cluster416-001 | Nin-like | NLP1-like [Brachypodium | 3742 | 0 | 0 | 0 |
| SALT00000070471_Cluster416-002 | Nin-like | NLP1-like [Brachypodium | 2647 | 0 | 0 | 0 |
| SALT00000090991_Cluster416-004 | Nin-like | NLP1-like [Brachypodium | 3896 | 0 | 0 | 0 |
| SALT00000030338_Cluster862-001 | Nin-like | NLP1-like [Brachypodium | 4142 | 0 | 0 | 0 |
| SALT00000033796_Cluster862-002 | Nin-like | NLP1-like [Brachypodium | 3755 | 0 | 0 | 0 |
| SALT00000086299_Cluster862-004 | Nin-like | NLP1-like [Brachypodium | 3399 | 0 | 0 | 0 |
| SALT00000089975_Cluster862-005 | Nin-like | NLP1-like [Brachypodium | 3545 | 0 | 0 | 0 |
| SALT00000091357_Cluster862-006 | Nin-like | NLP1-like [Brachypodium | 3621 | 0 | 0 | 0 |
| SALT00000091652_Cluster862-007 | Nin-like | NLP1-like [Brachypodium | 3730 | 0 | 0 | 0 |
| SALT00000089192_Cluster862-008 | Nin-like | NLP1-like [Brachypodium | 3943 | 0 | 0 | 0 |
| SALT00000024151_Cluster2174-001 | Nin-like | NLP1 isoform X2 [Setaria | 2288 | 0 | 0 | 0 |
| SALT00000026044_Cluster2174-002 | Nin-like | NLP1 isoform X2 [Setaria | 2753 | 0 | 0 | 0 |
| SALT00000051190_Cluster2174-003 | Nin-like | NLP1 isoform X2 [Setaria | 1807 | 0 | 0 | 0 |
| SALT00000053523_Cluster2174-004 | Nin-like | TPA: plant regulator RWP- | 2242 | 0 | 0 | 0 |
| SALT00000075993_Cluster2174-005 | Nin-like | NLP1 isoform X2 [Setaria | 2767 | 0 | 0 | 0 |
| SALT00000033715_Cluster3805-001 | Nin-like | NLP2-like [Zea mays] | 3457 | 0 | 0 | 0 |
| SALT00000037866_Cluster3805-002 | Nin-like | hypothetical protein SORE | 1503 | 0 | 0 | 0 |
| SALT00000091606_Cluster3805-003 | Nin-like | NLP2-like [Zea mays] | 4287 | 0 | 0 | 0 |
| SALT00000029777_Cluster3949-001 | Nin-like | NLP1-like [Brachypodium | 4136 | 0 | 0 | 0 |
| SALT00000032201_Cluster3949-002 | Nin-like | NLP1-like [Brachypodium | 4001 | 0 | 0 | 0 |
| SALT00000082063_Cluster3949-003 | Nin-like | NLP1-like [Brachypodium | 4241 | 0 | 0 | 0 |
| SALT00000021845_Cluster4159-001 | Nin-like | NLP1-like [Brachypodium | 3054 | 0 | 0 | 0 |
| SALT00000033868_Cluster4159-002 | Nin-like | NLP1-like [Brachypodium | 4175 | 0 | 0 | 0 |
| SALT00000091369_Cluster4159-003 | Nin-like | NLP1-like [Brachypodium | 3528 | 0 | 0 | 0 |
| SALT00000092570_Cluster4159-004 | Nin-like | NLP1-like [Brachypodium | 4032 | 0 | 0 | 0 |
| SALT00000022398_Cluster4423-001 | Nin-like | NLP2-like [Zea mays] | 2710 | 0 | 0 | 0 |
| SALT00000028927_Cluster4423-002 | Nin-like | NLP2-like [Zea mays] | 4044 | 0 | 0 | 0 |
| SALT00000029325_Cluster4423-003 | Nin-like | NLP2-like [Zea mays] | 3795 | 0 | 0 | 0 |
| SALT00000030546_Cluster4423-004 | Nin-like | NLP2-like [Zea mays] | 3627 | 0 | 0 | 0 |
| SALT00000030785_Cluster4423-005 | Nin-like | NLP2-like [Zea mays] | 3879 | 0 | 0 | 0 |
| SALT00000034214_Cluster4423-006 | Nin-like | NLP2-like [Zea mays] | 3968 | 0 | 0 | 0 |
| SALT00000034281_Cluster4423-007 | Nin-like | NLP2-like [Zea mays] | 3685 | 0 | 0 | 0 |
| SALT00000034517_Cluster4423-008 | Nin-like | NLP2-like [Zea mays] | 3866 | 0 | 0 | 0 |
| SALT00000034556_Cluster4423-009 | Nin-like | NLP2-like [Zea mays] | 3777 | 0 | 0 | 0 |
| SALT00000072793_Cluster4423-010 | Nin-like | NLP2-like [Zea mays] | 2970 | 0 | 0 | 0 |
| SALT00000090196_Cluster4423-011 | Nin-like | NLP2-like [Zea mays] | 3555 | 0 | 0 | 0 |
| SALT00000090298_Cluster4423-012 | Nin-like | NLP2-like [Zea mays] | 3970 | 0 | 0 | 0 |
| SALT00000092328_Cluster4423-013 | Nin-like | NLP2-like [Zea mays] | 4111 | 0 | 0 | 0 |
| SALT00000092976_Cluster4423-014 | Nin-like | NLP2-like [Zea mays] | 3648 | 0 | 0 | 0 |
| SALT00000093045_Cluster4423-015 | Nin-like | NLP2-like [Zea mays] | 3773 | 0 | 0 | 0 |
| SALT00000090528_Cluster4423-016 | Nin-like | NLP2-like [Zea mays] | 3708 | 0 | 0 | 0 |
| SALT00000091127_Cluster4710-001 | Nin-like | NLP1-like [Brachypodium | 4038 | 0 | 0 | 0 |
| SALT00000085702_Cluster5113-001 | Nin-like | TPA: homeodomain-like tr | 3953 | 0 | 0 | 0 |
| SALT00000030015_Cluster7030-001 | Nin-like | NLP1 isoform X2 [Setaria | 3341 | 0 | 0 | 0 |
| SALT00000032041_Cluster7030-002 | Nin-like | NLP1 isoform X2 [Setaria | 3632 | 0 | 0 | 0 |
| SALT00000033110_Cluster7030-003 | Nin-like | NLP1 isoform X2 [Setaria | 3499 | 0 | 0 | 0 |
| SALT00000072742_Cluster7030-004 | Nin-like | NLP1 isoform X3 [Brachy | 3082 | 0 | 0 | 0 |

| | | | | | | |
|---|---|---|---|---|---|---|
| SALT00000077020_Cluster7030-005 | Nin-like | NLP1 isoform X2 [Setaria | 3531 | 0 | 0 | 0 |
| SALT00000082446_Cluster7030-006 | Nin-like | NLP1 isoform X2 [Setaria | 2816 | 0 | 0 | 0 |
| SALT00000084505_Cluster7030-007 | Nin-like | NLP1 isoform X2 [Setaria | 3525 | 0 | 0 | 0 |
| SALT00000070001_Cluster11021-001 | Nin-like | Os01g0236700 [Oryza sati | 2969 | 0 | 0 | 0 |
| SALT00000083643_Cluster11021-002 | Nin-like | Os01g0236700 [Oryza sati | 2936 | 0 | 0 | 0 |
| SALT00000081473_Cluster11856-001 | Nin-like | Os01g0236700 [Oryza sati | 2882 | 0 | 0 | 0 |
| SALT00000081930_Cluster11856-002 | Nin-like | Os01g0236700 [Oryza sati | 2873 | 0 | 0 | 0 |
| SALT00000008404_Cluster24459-001 | RAV | AP2 EREBP transcription | 1467 | 0 | 0 | 0 |
| SALT00000012498_Cluster24459-002 | RAV | AP2 EREBP transcription | 1502 | 0 | 0 | 0 |
| SALT00000023403_Cluster24459-003 | RAV | AP2 EREBP transcription | 1506 | 0 | 0 | 0 |
| SALT00000054039_Cluster24459-004 | RAV | AP2 EREBP transcription | 1793 | 0 | 0 | 0 |
| SALT00000005192_Cluster28939-001 | RAV | DNA-binding RAV1 [Zea | 1525 | 0 | 0 | 0 |
| SALT00000017061_Cluster28939-002 | RAV | Os01g0693400 [Oryza sati | 1524 | 0 | 0 | up |
| SALT00000036848_Cluster28939-003 | RAV | Os01g0693400 [Oryza sati | 1475 | 0 | 0 | 0 |
| SALT00000042947_Cluster28939-004 | RAV | DNA-binding RAV1 [Zea | 1562 | 0 | 0 | 0 |
| SALT00000049370_Cluster28939-005 | RAV | Os01g0693400 [Oryza sati | 1405 | 0 | 0 | 0 |
| SALT00000017094_Cluster29121-001 | RAV | Os01g0141000 [Oryza sati | 1465 | 0 | 0 | 0 |
| SALT00000018929_Cluster29121-002 | RAV | Os01g0141000 [Oryza sati | 1291 | 0 | 0 | 0 |
| SALT00000037437_Cluster29121-003 | RAV | Os01g0141000 [Oryza sati | 1493 | 0 | 0 | 0 |
| SALT00000039618_Cluster29121-004 | RAV | Os01g0141000 [Oryza sati | 1382 | 0 | 0 | down |
| SALT00000054723_Cluster29121-005 | RAV | Os01g0141000 [Oryza sati | 1252 | 0 | 0 | up |
| SALT00000057585_Cluster29121-006 | RAV | Os01g0141000 [Oryza sati | 1553 | 0 | 0 | 0 |
| SALT00000061770_Cluster29121-007 | RAV | Os01g0141000 [Oryza sati | 1528 | 0 | 0 | 0 |
| SALT00000015080_Cluster29232-001 | RAV | Os01g0141000 [Oryza sati | 1547 | 0 | 0 | 0 |
| SALT00000042548_Cluster29232-002 | RAV | Os01g0141000 [Oryza sati | 1410 | 0 | 0 | 0 |
| SALT00000051490_Cluster29232-003 | RAV | Os01g0141000 [Oryza sati | 1499 | 0 | 0 | 0 |
| SALT00000042386_Cluster29499-001 | RAV | Os01g0693400 [Oryza sati | 1533 | 0 | 0 | 0 |
| SALT00000051139_Cluster29499-002 | RAV | Os01g0693400 [Oryza sati | 1390 | 0 | 0 | 0 |
| SALT00000054836_Cluster29499-003 | RAV | Os01g0693400 [Oryza sati | 1493 | 0 | 0 | up |
| SALT00000048999_Cluster30504-001 | RAV | AP2 EREBP transcription | 1447 | 0 | 0 | 0 |
| SALT00000061387_Cluster30504-002 | RAV | AP2 EREBP transcription | 1468 | 0 | 0 | 0 |
| SALT00000028671_Cluster509-009 | SBP | squamosa promoter-bindin | 3922 | 0 | 0 | 0 |
| SALT00000028777_Cluster509-010 | SBP | squamosa promoter-bindin | 4044 | 0 | 0 | 0 |
| SALT00000029515_Cluster509-011 | SBP | squamosa promoter-bindin | 4257 | 0 | 0 | 0 |
| SALT00000030050_Cluster509-012 | SBP | squamosa promoter-bindin | 3611 | 0 | 0 | 0 |
| SALT00000030072_Cluster509-013 | SBP | squamosa promoter-bindin | 4189 | 0 | 0 | 0 |
| SALT00000030329_Cluster509-014 | SBP | squamosa promoter-bindin | 4435 | 0 | 0 | 0 |
| SALT00000030431_Cluster509-015 | SBP | squamosa promoter-bindin | 4229 | 0 | 0 | 0 |
| SALT00000032060_Cluster509-016 | SBP | squamosa promoter-bindin | 4007 | 0 | 0 | 0 |
| SALT00000032164_Cluster509-017 | SBP | squamosa promoter-bindin | 4193 | 0 | 0 | 0 |
| SALT00000032197_Cluster509-018 | SBP | squamosa promoter-bindin | 3376 | 0 | 0 | 0 |
| SALT00000032229_Cluster509-019 | SBP | squamosa promoter-bindin | 4364 | 0 | 0 | 0 |
| SALT00000032490_Cluster509-020 | SBP | squamosa promoter-bindin | 4091 | 0 | 0 | 0 |
| SALT00000032735_Cluster509-021 | SBP | squamosa promoter-bindin | 3761 | 0 | 0 | 0 |
| SALT00000032749_Cluster509-022 | SBP | squamosa promoter-bindin | 3419 | 0 | 0 | 0 |
| SALT00000032822_Cluster509-023 | SBP | squamosa promoter-bindin | 3666 | 0 | 0 | 0 |
| SALT00000032991_Cluster509-024 | SBP | squamosa promoter-bindin | 3860 | 0 | 0 | 0 |
| SALT00000033153_Cluster509-025 | SBP | squamosa promoter-bindin | 3925 | 0 | 0 | 0 |
| SALT00000034109_Cluster509-026 | SBP | squamosa promoter-bindin | 4009 | 0 | 0 | 0 |
| SALT00000034402_Cluster509-027 | SBP | squamosa promoter-bindin | 3878 | 0 | 0 | 0 |
| SALT00000077262_Cluster509-040 | SBP | squamosa promoter-bindin | 4108 | 0 | 0 | 0 |
| SALT00000077984_Cluster509-041 | SBP | squamosa promoter-bindin | 3765 | 0 | 0 | 0 |
| SALT00000079106_Cluster509-042 | SBP | Squamosa promoter-bindir | 3314 | 0 | 0 | 0 |
| SALT00000079197_Cluster509-043 | SBP | squamosa promoter-bindin | 4188 | 0 | 0 | 0 |
| SALT00000079908_Cluster509-044 | SBP | squamosa promoter-bindin | 4295 | 0 | 0 | 0 |
| SALT00000081791_Cluster509-045 | SBP | squamosa promoter-bindin | 4269 | 0 | 0 | 0 |
| SALT00000082274_Cluster509-046 | SBP | squamosa promoter-bindin | 4191 | 0 | 0 | 0 |
| SALT00000083972_Cluster509-047 | SBP | squamosa promoter-bindin | 3444 | 0 | 0 | 0 |

| | | | | | | |
|---|---|---|---|---|---|---|
| SALT00000084095_Cluster509-048 | SBP | Squamosa promoter-bindir | 4164 | 0 | 0 | 0 |
| SALT00000084355_Cluster509-049 | SBP | squamosa promoter-bindin | 4258 | 0 | 0 | 0 |
| SALT00000085809_Cluster509-050 | SBP | squamosa promoter-bindin | 3997 | 0 | 0 | 0 |
| SALT00000085943_Cluster509-051 | SBP | squamosa promoter-bindin | 3419 | 0 | 0 | 0 |
| SALT00000086310_Cluster509-054 | SBP | squamosa promoter-bindin | 3821 | 0 | 0 | 0 |
| SALT00000086991_Cluster509-055 | SBP | squamosa promoter-bindin | 4169 | 0 | 0 | 0 |
| SALT00000087239_Cluster509-056 | SBP | squamosa promoter-bindin | 4311 | 0 | 0 | 0 |
| SALT00000088553_Cluster509-057 | SBP | squamosa promoter-bindin | 4061 | 0 | 0 | 0 |
| SALT00000089747_Cluster509-058 | SBP | squamosa promoter-bindin | 3630 | 0 | 0 | 0 |
| SALT00000090507_Cluster509-059 | SBP | squamosa promoter-bindin | 4185 | 0 | 0 | 0 |
| SALT00000091198_Cluster509-060 | SBP | squamosa promoter-bindin | 4020 | 0 | 0 | 0 |
| SALT00000091908_Cluster509-061 | SBP | squamosa promoter-bindin | 4316 | 0 | 0 | up |
| SALT00000092759_Cluster509-062 | SBP | squamosa promoter-bindin | 4225 | 0 | 0 | 0 |
| SALT00000093049_Cluster509-063 | SBP | squamosa promoter-bindin | 3691 | 0 | 0 | 0 |
| SALT00000093396_Cluster509-064 | SBP | squamosa promoter-bindin | 4378 | 0 | 0 | 0 |
| SALT00000093774_Cluster509-065 | SBP | squamosa promoter-bindin | 4072 | 0 | 0 | 0 |
| SALT00000086616_Cluster509-066 | SBP | squamosa promoter-bindin | 4420 | 0 | 0 | 0 |
| SALT00000085406_Cluster737-001 | SBP | squamosa promoter-bindin | 3754 | 0 | 0 | 0 |
| SALT00000092078_Cluster1886-001 | SBP | squamosa promoter-bindin | 5233 | 0 | 0 | 0 |
| SALT00000032799_Cluster3263-001 | SBP | squamosa promoter-bindin | 3400 | 0 | 0 | 0 |
| SALT00000085448_Cluster3263-002 | SBP | Squamosa promoter-bindir | 4490 | 0 | 0 | 0 |
| SALT00000021558_Cluster4424-001 | SBP | squamosa promoter-bindin | 2965 | 0 | 0 | 0 |
| SALT00000021851_Cluster4424-002 | SBP | squamosa promoter-bindin | 2828 | 0 | 0 | 0 |
| SALT00000023644_Cluster4424-003 | SBP | squamosa promoter-bindin | 2988 | 0 | 0 | 0 |
| SALT00000026833_Cluster4424-004 | SBP | squamosa promoter-bindin | 3057 | 0 | 0 | 0 |
| SALT00000028965_Cluster4424-005 | SBP | squamosa promoter-bindin | 2442 | 0 | 0 | 0 |
| SALT00000063478_Cluster4424-006 | SBP | squamosa promoter-bindin | 2950 | 0 | 0 | 0 |
| SALT00000064156_Cluster4424-007 | SBP | squamosa promoter-bindin | 2994 | 0 | 0 | 0 |
| SALT00000065306_Cluster4424-008 | SBP | squamosa promoter-bindin | 4110 | 0 | 0 | 0 |
| SALT00000067030_Cluster4424-009 | SBP | squamosa promoter-bindin | 3005 | 0 | 0 | 0 |
| SALT00000029446_Cluster5134-001 | SBP | Os03g0833300 [Oryza sati | 3594 | 0 | 0 | 0 |
| SALT00000030031_Cluster5134-002 | SBP | Os03g0833300 [Oryza sati | 3617 | 0 | 0 | 0 |
| SALT00000030159_Cluster5134-003 | SBP | Os03g0833300 [Oryza sati | 3568 | 0 | 0 | 0 |
| SALT00000033551_Cluster5134-004 | SBP | Os03g0833300 [Oryza sati | 3590 | 0 | 0 | 0 |
| SALT00000062866_Cluster5134-005 | SBP | Os03g0833300 [Oryza sati | 2611 | 0 | 0 | 0 |
| SALT00000067190_Cluster5134-006 | SBP | Os03g0833300 [Oryza sati | 3153 | 0 | 0 | 0 |
| SALT00000068165_Cluster5134-007 | SBP | Os03g0833300 [Oryza sati | 2576 | 0 | 0 | 0 |
| SALT00000068811_Cluster5134-009 | SBP | Os03g0833300 [Oryza sati | 2874 | 0 | 0 | 0 |
| SALT00000076197_Cluster5134-010 | SBP | Os03g0833300 [Oryza sati | 2740 | 0 | 0 | 0 |
| SALT00000078007_Cluster5134-011 | SBP | Os03g0833300 [Oryza sati | 3749 | 0 | 0 | 0 |
| SALT00000080955_Cluster5134-012 | SBP | Os03g0833300 [Oryza sati | 2591 | 0 | 0 | 0 |
| SALT00000087238_Cluster5134-013 | SBP | Os03g0833300 [Oryza sati | 3512 | 0 | 0 | 0 |
| SALT00000087324_Cluster5134-014 | SBP | Os03g0833300 [Oryza sati | 3950 | 0 | 0 | 0 |
| SALT00000087415_Cluster5134-015 | SBP | Os03g0833300 [Oryza sati | 3651 | 0 | 0 | 0 |
| SALT00000087852_Cluster5134-016 | SBP | Os03g0833300 [Oryza sati | 3562 | 0 | 0 | 0 |
| SALT00000088585_Cluster5134-017 | SBP | Os03g0833300 [Oryza sati | 3349 | 0 | 0 | 0 |
| SALT00000090025_Cluster5134-018 | SBP | Os03g0833300 [Oryza sati | 3405 | 0 | 0 | 0 |
| SALT00000090194_Cluster5134-019 | SBP | Os03g0833300 [Oryza sati | 3619 | 0 | 0 | 0 |
| SALT00000090745_Cluster5134-020 | SBP | Os03g0833300 [Oryza sati | 3434 | 0 | 0 | 0 |
| SALT00000093619_Cluster5134-021 | SBP | Os03g0833300 [Oryza sati | 3612 | 0 | 0 | 0 |
| SALT00000086087_Cluster5134-022 | SBP | Os03g0833300 [Oryza sati | 3427 | 0 | 0 | 0 |
| SALT00000083867_Cluster5134-023 | SBP | Os03g0833300 [Oryza sati | 3528 | 0 | 0 | 0 |
| SALT00000033032_Cluster5134-024 | SBP | Os03g0833300 [Oryza sati | 3467 | 0 | 0 | 0 |
| SALT00000085434_Cluster5620-002 | SBP | Os03g0833300 [Oryza sati | 3858 | 0 | 0 | 0 |
| SALT00000093546_Cluster7376-001 | SBP | squamosa promoter-bindin | 3580 | 0 | 0 | 0 |
| SALT00000021989_Cluster8646-001 | SBP | squamosa promoter-bindin | 3144 | 0 | 0 | 0 |
| SALT00000027014_Cluster8646-002 | SBP | squamosa promoter-bindin | 2567 | 0 | 0 | 0 |
| SALT00000033784_Cluster8646-003 | SBP | squamosa promoter-bindin | 3354 | 0 | 0 | 0 |

| | | | | | | |
|---|---|---|---|---|---|---|
| SALT00000065711_Cluster8646-005 | SBP | squamosa promoter-bindin | 2976 | 0 | 0 | 0 |
| SALT00000066116_Cluster8646-006 | SBP | squamosa promoter-bindin | 3064 | 0 | 0 | 0 |
| SALT00000068697_Cluster8646-007 | SBP | squamosa promoter-bindin | 2922 | 0 | 0 | 0 |
| SALT00000069741_Cluster8646-008 | SBP | squamosa promoter-bindin | 2953 | 0 | 0 | 0 |
| SALT00000069873_Cluster8646-009 | SBP | squamosa promoter-bindin | 2995 | 0 | 0 | 0 |
| SALT00000070382_Cluster8646-010 | SBP | squamosa promoter-bindin | 3169 | 0 | 0 | 0 |
| SALT00000074029_Cluster8646-011 | SBP | squamosa promoter-bindin | 3082 | 0 | 0 | 0 |
| SALT00000077958_Cluster8646-012 | SBP | squamosa promoter-bindin | 2895 | 0 | 0 | 0 |
| SALT00000082927_Cluster8646-013 | SBP | squamosa promoter-bindin | 2944 | 0 | 0 | 0 |
| SALT00000090598_Cluster8646-014 | SBP | squamosa promoter-bindin | 3104 | 0 | 0 | 0 |
| SALT00000059252_Cluster8646-015 | SBP | squamosa promoter-bindin | 2194 | 0 | 0 | 0 |
| SALT00000032737_Cluster8983-001 | SBP | Os03g0833300 [Oryza sati | 3290 | 0 | 0 | 0 |
| SALT00000022405_Cluster9463-001 | SBP | squamosa promoter-bindin | 3041 | 0 | 0 | 0 |
| SALT00000028450_Cluster9463-002 | SBP | squamosa promoter-bindin | 3063 | 0 | 0 | 0 |
| SALT00000031511_Cluster9463-003 | SBP | squamosa promoter-bindin | 2978 | 0 | 0 | 0 |
| SALT00000063867_Cluster9463-004 | SBP | squamosa promoter-bindin | 3030 | 0 | 0 | 0 |
| SALT00000069327_Cluster9463-005 | SBP | squamosa promoter-bindin | 3198 | 0 | up | 0 |
| SALT00000082650_Cluster9463-006 | SBP | squamosa promoter-bindin | 2843 | 0 | 0 | 0 |
| SALT00000069382_Cluster9556-001 | SBP | Os03g0833300 [Oryza sati | 3180 | 0 | 0 | 0 |
| SALT00000055182_Cluster11107-001 | SBP | squamosa promoter-bindin | 2150 | 0 | 0 | 0 |
| SALT00000063352_Cluster11107-002 | SBP | squamosa promoter-bindin | 2406 | 0 | 0 | 0 |
| SALT00000075451_Cluster11107-003 | SBP | squamosa promoter-bindin | 2959 | 0 | 0 | 0 |
| SALT00000081390_Cluster14876-001 | SBP | squamosa promoter-bindin | 2635 | 0 | 0 | 0 |
| SALT00000083074_Cluster15692-002 | SBP | squamosa promoter-bindin | 1551 | 0 | 0 | 0 |
| SALT00000009252_Cluster17765-001 | SBP | squamosa promoter-bindin | 1650 | 0 | 0 | 0 |
| SALT00000010925_Cluster17765-002 | SBP | squamosa promoter-bindin | 1656 | 0 | 0 | 0 |
| SALT00000027374_Cluster17765-003 | SBP | squamosa promoter-bindin | 2349 | 0 | 0 | 0 |
| SALT00000007083_Cluster17765-006 | SBP | squamosa promoter-bindin | 1475 | 0 | 0 | 0 |
| SALT00000055176_Cluster18353-001 | SBP | squamosa promoter-bindin | 1753 | 0 | 0 | 0 |
| SALT00000072009_Cluster18353-002 | SBP | squamosa promoter-bindin | 2240 | 0 | 0 | 0 |
| SALT00000044835_Cluster20137-001 | SBP | squamosa promoter-bindin | 1860 | 0 | 0 | 0 |
| SALT00000048156_Cluster20137-002 | SBP | squamosa promoter-bindin | 2068 | 0 | 0 | 0 |
| SALT00000037101_Cluster21062-001 | SBP | squamosa promoter-bindin | 1606 | 0 | 0 | 0 |
| SALT00000040573_Cluster21062-002 | SBP | squamosa promoter-bindin | 1812 | 0 | 0 | 0 |
| SALT00000047800_Cluster21062-003 | SBP | squamosa promoter-bindin | 2000 | 0 | 0 | 0 |
| SALT00000052114_Cluster21062-004 | SBP | squamosa promoter-bindin | 1954 | 0 | 0 | 0 |
| SALT00000039576_Cluster22807-001 | SBP | SBP transcription partial [Z | 1646 | 0 | 0 | 0 |
| SALT00000039583_Cluster22807-002 | SBP | SBP transcription partial [Z | 1890 | 0 | 0 | 0 |
| SALT00000043876_Cluster24904-001 | SBP | squamosa promoter-bindin | 1768 | 0 | 0 | 0 |
| SALT00000052770_Cluster24904-002 | SBP | squamosa promoter-bindin | 1732 | 0 | 0 | 0 |
| SALT00000050773_Cluster24928-001 | SBP | SBP-domain 4 [Zea mays] | 1767 | 0 | 0 | 0 |
| SALT00000007121_Cluster25592-001 | SBP | squamosa promoter-bindin | 1698 | 0 | 0 | 0 |
| SALT00000035751_Cluster25592-002 | SBP | squamosa promoter-bindin | 1664 | 0 | 0 | 0 |
| SALT00000039606_Cluster25592-003 | SBP | squamosa promoter-bindin | 1500 | 0 | 0 | 0 |
| SALT00000048576_Cluster25592-004 | SBP | squamosa promoter-bindin | 1733 | 0 | 0 | 0 |
| SALT00000058362_Cluster25592-005 | SBP | squamosa promoter-bindin | 1731 | 0 | 0 | 0 |
| SALT00000009501_Cluster27690-001 | SBP | Squamosa promoter-bindir | 1626 | 0 | 0 | 0 |
| SALT00000056359_Cluster3056-001 | SRS | SRS transcription partial [Z | 1699 | 0 | 0 | 0 |
| SALT00000037413_Cluster21797-001 | SRS | LRP1 [Oryza sativa Japoni | 1952 | 0 | 0 | 0 |
| SALT00000040571_Cluster21797-002 | SRS | LRP1 [Oryza sativa Japoni | 1581 | 0 | 0 | 0 |
| SALT00000041719_Cluster21797-003 | SRS | LRP1 [Oryza sativa Japoni | 1608 | 0 | 0 | up |
| SALT00000002919_Cluster22749-001 | SRS | LRP1 [Oryza sativa Japoni | 1785 | 0 | 0 | 0 |
| SALT00000036511_Cluster22749-002 | SRS | LRP1 [Oryza sativa Japoni | 1893 | 0 | 0 | 0 |
| SALT00000037663_Cluster24893-001 | SRS | LRP1 [Oryza sativa Japoni | 1603 | 0 | 0 | 0 |
| SALT00000060806_Cluster24893-002 | SRS | LRP1 [Oryza sativa Japoni | 1769 | 0 | 0 | 0 |
| SALT00000036221_Cluster25014-001 | SRS | LRP1 [Zea mays] | 1754 | 0 | 0 | 0 |
| SALT00000050478_Cluster25014-002 | SRS | LRP1 [Zea mays] | 1762 | 0 | 0 | 0 |
| SALT00000008401_Cluster25616-001 | SRS | Os08g0129500 [Oryza sati | 1528 | 0 | 0 | 0 |

| | | | | | | |
|---|---|---|---|---|---|---|
| SALT00000018150_Cluster25616-002 | SRS | Os08g0129500 [Oryza sati | 1605 | 0 | 0 | 0 |
| SALT00000038109_Cluster25616-003 | SRS | Os08g0129500 [Oryza sati | 1390 | 0 | 0 | 0 |
| SALT00000045926_Cluster25616-004 | SRS | Os08g0129500 [Oryza sati | 1496 | 0 | 0 | 0 |
| SALT00000058095_Cluster25616-005 | SRS | Os08g0129500 [Oryza sati | 1732 | 0 | 0 | 0 |
| SALT00000059060_Cluster25616-006 | SRS | Os08g0129500 [Oryza sati | 1688 | 0 | 0 | 0 |
| SALT00000075582_Cluster25616-007 | SRS | Os08g0129500 [Oryza sati | 1346 | 0 | 0 | 0 |
| SALT00000058260_Cluster27896-001 | SRS | SHI RELATED SEQUEN( | 1616 | 0 | 0 | 0 |
| SALT00000009857_Cluster28131-001 | SRS | SHI RELATED SEQUEN( | 1603 | 0 | 0 | 0 |
| SALT00000069550_Cluster10388-001 | STAT | PREDICTED: uncharacter | 3052 | 0 | 0 | 0 |
| SALT00000081363_Cluster10388-002 | STAT | PREDICTED: uncharacter | 2416 | 0 | 0 | 0 |
| SALT00000062488_Cluster12427-001 | STAT | PREDICTED: uncharacter | 2826 | 0 | 0 | 0 |
| SALT00000021375_Cluster13988-001 | STAT | PREDICTED: uncharacter | 2531 | 0 | 0 | 0 |
| SALT00000083176_Cluster13988-002 | STAT | PREDICTED: uncharacter | 2696 | 0 | 0 | 0 |
| SALT00000012966_Cluster244-001 | TALE | BEL1-like homeodomain 7 | 1400 | 0 | 0 | 0 |
| SALT00000007866_Cluster1470-001 | TALE | BEL1-like homeodomain 7 | 1299 | 0 | 0 | 0 |
| SALT00000009721_Cluster1470-002 | TALE | Os03g0680800 [Oryza sati | 1574 | 0 | 0 | 0 |
| SALT00000021263_Cluster1470-003 | TALE | Os03g0680800 [Oryza sati | 2545 | 0 | 0 | 0 |
| SALT00000022153_Cluster1470-004 | TALE | BEL1-related homeotic 30 | 2095 | 0 | 0 | 0 |
| SALT00000022916_Cluster1470-005 | TALE | BEL1-related homeotic 30 | 2456 | 0 | 0 | 0 |
| SALT00000023781_Cluster1470-006 | TALE | Os03g0680800 [Oryza sati | 2578 | 0 | 0 | 0 |
| SALT00000025728_Cluster1470-008 | TALE | Os03g0680800 [Oryza sati | 2673 | 0 | 0 | 0 |
| SALT00000025968_Cluster1470-009 | TALE | Os03g0680800 [Oryza sati | 2402 | 0 | 0 | 0 |
| SALT00000026890_Cluster1470-010 | TALE | Os03g0680800 [Oryza sati | 2630 | 0 | 0 | 0 |
| SALT00000027428_Cluster1470-011 | TALE | Os03g0680800 [Oryza sati | 2595 | 0 | 0 | 0 |
| SALT00000027562_Cluster1470-012 | TALE | Os03g0680800 [Oryza sati | 2641 | 0 | 0 | 0 |
| SALT00000029057_Cluster1470-013 | TALE | Os03g0680800 [Oryza sati | 2679 | 0 | 0 | 0 |
| SALT00000060268_Cluster1470-014 | TALE | Os03g0680800 [Oryza sati | 1719 | 0 | 0 | 0 |
| SALT00000065727_Cluster1470-015 | TALE | Os08g0398000 [Oryza sati | 5663 | 0 | 0 | 0 |
| SALT00000072465_Cluster1470-016 | TALE | BEL1-related homeotic 30 | 2501 | 0 | 0 | 0 |
| SALT00000072712_Cluster1470-017 | TALE | Os03g0680800 [Oryza sati | 2670 | 0 | 0 | 0 |
| SALT00000073047_Cluster1470-018 | TALE | Os03g0680800 [Oryza sati | 2845 | 0 | 0 | 0 |
| SALT00000076491_Cluster1470-019 | TALE | Os03g0680800 [Oryza sati | 2512 | 0 | 0 | 0 |
| SALT00000078100_Cluster1470-020 | TALE | Os03g0680800 [Oryza sati | 2706 | 0 | 0 | 0 |
| SALT00000020771_Cluster1986-001 | TALE | BEL1-related homeotic 30 | 2579 | 0 | 0 | 0 |
| SALT00000023621_Cluster1986-002 | TALE | BEL1-related homeotic 30 | 2675 | 0 | 0 | 0 |
| SALT00000024250_Cluster1986-003 | TALE | BEL1-related homeotic 30 | 2524 | 0 | 0 | 0 |
| SALT00000025982_Cluster1986-004 | TALE | BEL1-related homeotic 30 | 2634 | 0 | 0 | 0 |
| SALT00000028179_Cluster1986-005 | TALE | BEL1-related homeotic 30 | 2733 | 0 | 0 | 0 |
| SALT00000029625_Cluster1986-006 | TALE | BEL1-related homeotic 30 | 2672 | 0 | 0 | 0 |
| SALT00000062237_Cluster1986-007 | TALE | BEL1-related homeotic 30 | 2641 | 0 | 0 | 0 |
| SALT00000009076_Cluster3560-001 | TALE | BEL1-like homeodomain 1 | 1738 | 0 | 0 | 0 |
| SALT00000021119_Cluster3560-002 | TALE | BEL1-like homeodomain 1 | 2718 | 0 | 0 | 0 |
| SALT00000023612_Cluster3560-003 | TALE | BEL1-like homeodomain 1 | 2661 | 0 | 0 | 0 |
| SALT00000023890_Cluster3560-004 | TALE | BEL1-like homeodomain 1 | 2437 | 0 | 0 | 0 |
| SALT00000024990_Cluster3560-005 | TALE | BEL1-like homeodomain 1 | 2427 | 0 | 0 | 0 |
| SALT00000025552_Cluster3560-006 | TALE | BEL1-like homeodomain 1 | 2553 | 0 | 0 | 0 |
| SALT00000025714_Cluster3560-007 | TALE | BEL1-like homeodomain 1 | 2515 | 0 | 0 | 0 |
| SALT00000027845_Cluster3560-008 | TALE | BEL1-like homeodomain 1 | 2696 | 0 | 0 | 0 |
| SALT00000027883_Cluster3560-009 | TALE | BEL1-like homeodomain 1 | 2621 | 0 | 0 | 0 |
| SALT00000031065_Cluster3560-010 | TALE | BEL1-like homeodomain 1 | 2467 | 0 | 0 | 0 |
| SALT00000047145_Cluster3560-011 | TALE | BEL1-like homeodomain 1 | 1579 | 0 | 0 | 0 |
| SALT00000064529_Cluster3560-012 | TALE | BEL1-like homeodomain 1 | 3006 | 0 | 0 | 0 |
| SALT00000065350_Cluster3560-013 | TALE | transport inhibitor respons( | 4371 | 0 | 0 | 0 |
| SALT00000068922_Cluster3560-014 | TALE | BEL1-like homeodomain 1 | 2616 | 0 | 0 | 0 |
| SALT00000069365_Cluster3560-015 | TALE | BEL1-like homeodomain 1 | 2573 | 0 | 0 | 0 |
| SALT00000073092_Cluster3560-016 | TALE | BEL1-like homeodomain 1 | 2487 | 0 | 0 | 0 |
| SALT00000073271_Cluster3560-017 | TALE | BEL1-like homeodomain 1 | 2594 | 0 | 0 | 0 |
| SALT00000076656_Cluster3560-018 | TALE | BEL1-like homeodomain 1 | 2407 | 0 | 0 | 0 |

| | | | | | | |
|---|---|---|---|---|---|---|
| SALT00000070165_Cluster3560-019 | TALE | BEL1-like homeodomain 1 | 3004 | 0 | 0 | 0 |
| SALT00000067134_Cluster3639-001 | TALE | BEL1-type homeodomain | 4346 | 0 | 0 | 0 |
| SALT00000078630_Cluster3875-001 | TALE | homeotic knotted-1 [Zea m | 4265 | 0 | 0 | 0 |
| SALT00000021637_Cluster4233-001 | TALE | BEL1-like homeodomain 7 | 2474 | 0 | 0 | 0 |
| SALT00000023562_Cluster4233-002 | TALE | BEL1-like homeodomain 7 | 2558 | 0 | 0 | 0 |
| SALT00000024300_Cluster4233-003 | TALE | BEL1-like homeodomain 7 | 2460 | 0 | 0 | 0 |
| SALT00000034874_Cluster4233-004 | TALE | BEL1-like homeodomain 7 | 2470 | 0 | 0 | 0 |
| SALT00000047587_Cluster4233-005 | TALE | BEL1-like homeodomain 7 | 1845 | 0 | 0 | 0 |
| SALT00000062081_Cluster4233-006 | TALE | BEL1-like homeodomain 7 | 2514 | 0 | 0 | 0 |
| SALT00000068171_Cluster4233-007 | TALE | BEL1-like homeodomain 7 | 2651 | 0 | 0 | 0 |
| SALT00000070098_Cluster4233-008 | TALE | BEL1-like homeodomain 7 | 2461 | 0 | 0 | 0 |
| SALT00000070914_Cluster4233-009 | TALE | BEL1-like homeodomain 7 | 4156 | 0 | 0 | 0 |
| SALT00000074044_Cluster4233-010 | TALE | BEL1-like homeodomain 7 | 2654 | 0 | 0 | 0 |
| SALT00000005859_Cluster5780-001 | TALE | BEL1-like homeodomain 7 | 1717 | 0 | 0 | 0 |
| SALT00000010134_Cluster5780-002 | TALE | BEL1-like homeodomain 7 | 2031 | 0 | 0 | 0 |
| SALT00000010562_Cluster5780-003 | TALE | BEL1-like homeodomain 7 | 1582 | 0 | 0 | 0 |
| SALT00000016539_Cluster5780-004 | TALE | BEL1-like homeodomain 7 | 2037 | 0 | 0 | 0 |
| SALT00000020978_Cluster5780-005 | TALE | BEL1-like homeodomain 7 | 2563 | 0 | 0 | 0 |
| SALT00000022382_Cluster5780-006 | TALE | BEL1-like homeodomain 7 | 2830 | 0 | 0 | 0 |
| SALT00000022812_Cluster5780-007 | TALE | BEL1-like homeodomain 7 | 2672 | 0 | 0 | 0 |
| SALT00000023171_Cluster5780-008 | TALE | BEL1-like homeodomain 7 | 3032 | 0 | 0 | 0 |
| SALT00000023529_Cluster5780-009 | TALE | BEL1-like homeodomain 7 | 2862 | 0 | 0 | 0 |
| SALT00000026265_Cluster5780-010 | TALE | BEL1-like homeodomain 7 | 3087 | 0 | 0 | 0 |
| SALT00000028026_Cluster5780-011 | TALE | BEL1-like homeodomain 7 | 2514 | 0 | 0 | 0 |
| SALT00000028254_Cluster5780-012 | TALE | BEL1-like homeodomain 7 | 2516 | 0 | 0 | 0 |
| SALT00000028303_Cluster5780-013 | TALE | BEL1-like homeodomain 7 | 2608 | 0 | 0 | 0 |
| SALT00000029640_Cluster5780-014 | TALE | BEL1-like homeodomain 7 | 2464 | 0 | 0 | 0 |
| SALT00000030309_Cluster5780-015 | TALE | BEL1-like homeodomain 7 | 2629 | 0 | 0 | 0 |
| SALT00000030907_Cluster5780-016 | TALE | BEL1-like homeodomain 7 | 2568 | 0 | 0 | 0 |
| SALT00000032001_Cluster5780-017 | TALE | BEL1-like homeodomain 7 | 2590 | 0 | 0 | 0 |
| SALT00000034565_Cluster5780-018 | TALE | BEL1-like homeodomain 7 | 2574 | 0 | 0 | 0 |
| SALT00000035172_Cluster5780-019 | TALE | BEL1-like homeodomain 7 | 2680 | 0 | 0 | 0 |
| SALT00000035312_Cluster5780-020 | TALE | BEL1-like homeodomain 7 | 2662 | 0 | 0 | 0 |
| SALT00000035420_Cluster5780-021 | TALE | BEL1-like homeodomain 7 | 2652 | 0 | 0 | 0 |
| SALT00000035486_Cluster5780-022 | TALE | BEL1-like homeodomain 7 | 2540 | 0 | 0 | 0 |
| SALT00000035639_Cluster5780-023 | TALE | BEL1-like homeodomain 7 | 2500 | 0 | 0 | 0 |
| SALT00000039032_Cluster5780-024 | TALE | BEL1-like homeodomain 7 | 1963 | 0 | 0 | 0 |
| SALT00000044231_Cluster5780-025 | TALE | BEL1-like homeodomain 7 | 1435 | 0 | 0 | 0 |
| SALT00000066120_Cluster5780-026 | TALE | BEL1-like homeodomain 7 | 3830 | 0 | 0 | 0 |
| SALT00000066227_Cluster5780-027 | TALE | BEL1-like homeodomain 7 | 2492 | 0 | 0 | 0 |
| SALT00000069330_Cluster5780-028 | TALE | BEL1-like homeodomain 7 | 2689 | 0 | 0 | 0 |
| SALT00000070083_Cluster5780-029 | TALE | BEL1-like homeodomain 7 | 2552 | 0 | 0 | 0 |
| SALT00000070727_Cluster5780-030 | TALE | BEL1-like homeodomain 7 | 2617 | 0 | 0 | 0 |
| SALT00000071046_Cluster5780-031 | TALE | BEL1-like homeodomain 7 | 2553 | 0 | 0 | 0 |
| SALT00000071534_Cluster5780-032 | TALE | BEL1-like homeodomain 7 | 2504 | 0 | 0 | 0 |
| SALT00000072555_Cluster5780-033 | TALE | BEL1-like homeodomain 7 | 2630 | 0 | 0 | 0 |
| SALT00000072710_Cluster5780-034 | TALE | BEL1-like homeodomain 7 | 2604 | 0 | 0 | 0 |
| SALT00000072977_Cluster5780-035 | TALE | BEL1-like homeodomain 7 | 2624 | 0 | 0 | up |
| SALT00000073634_Cluster5780-036 | TALE | BEL1-like homeodomain 7 | 2603 | 0 | 0 | 0 |
| SALT00000073908_Cluster5780-037 | TALE | BEL1-like homeodomain 7 | 2525 | 0 | 0 | 0 |
| SALT00000084151_Cluster5780-038 | TALE | BEL1-like homeodomain 7 | 2649 | 0 | 0 | 0 |
| SALT00000084587_Cluster5780-039 | TALE | BEL1-like homeodomain 7 | 2535 | 0 | 0 | 0 |
| SALT00000079315_Cluster5780-040 | TALE | BEL1-like homeodomain 7 | 2777 | 0 | 0 | 0 |
| SALT00000025245_Cluster5865-001 | TALE | BEL1-related homeotic 30 | 2741 | 0 | 0 | 0 |
| SALT00000030280_Cluster5865-002 | TALE | BEL1-related homeotic 30 | 2552 | 0 | 0 | 0 |
| SALT00000068867_Cluster5865-003 | TALE | BEL1-related homeotic 30 | 3264 | 0 | 0 | 0 |
| SALT00000080937_Cluster5865-004 | TALE | BEL1-related homeotic 30 | 3275 | 0 | 0 | 0 |
| SALT00000084103_Cluster5865-005 | TALE | BEL1-related homeotic 30 | 3814 | 0 | 0 | 0 |

| | | | | | | |
|---|---|---|---|---|---|---|
| SALT00000091338_Cluster5865-006 | TALE | BEL1-related homeotic 30 | 3459 | 0 | 0 | 0 |
| SALT00000046393_Cluster6386-001 | TALE | BEL1-type homeodomain | 1290 | 0 | 0 | 0 |
| SALT00000003309_Cluster6754-001 | TALE | BEL1-like homeodomain 9 | 2265 | 0 | 0 | 0 |
| SALT00000012129_Cluster7188-001 | TALE | BEL1-like homeodomain 7 | 1897 | 0 | 0 | 0 |
| SALT00000044001_Cluster7188-002 | TALE | BEL1-like homeodomain 7 | 1938 | 0 | 0 | 0 |
| SALT00000065039_Cluster7188-003 | TALE | BEL1-like homeodomain 7 | 3609 | 0 | 0 | 0 |
| SALT00000072001_Cluster7378-001 | TALE | BEL1-like homeodomain 7 | 3579 | 0 | 0 | 0 |
| SALT00000022904_Cluster8126-001 | TALE | BEL1-type homeodomain | 2948 | 0 | 0 | 0 |
| SALT00000024162_Cluster8126-002 | TALE | BEL1-type homeodomain | 2980 | 0 | 0 | 0 |
| SALT00000026412_Cluster8126-003 | TALE | BEL1-type homeodomain | 3149 | 0 | 0 | 0 |
| SALT00000026953_Cluster8126-004 | TALE | BEL1-type homeodomain | 2894 | 0 | 0 | 0 |
| SALT00000027197_Cluster8126-005 | TALE | BEL1-type homeodomain | 3146 | 0 | 0 | 0 |
| SALT00000027678_Cluster8126-006 | TALE | BEL1-type homeodomain | 2806 | 0 | 0 | 0 |
| SALT00000028757_Cluster8126-007 | TALE | BEL1-type homeodomain | 2892 | 0 | 0 | 0 |
| SALT00000029441_Cluster8126-008 | TALE | BEL1-type homeodomain | 2558 | 0 | 0 | 0 |
| SALT00000029539_Cluster8126-009 | TALE | BEL1-type homeodomain | 2824 | 0 | 0 | 0 |
| SALT00000029963_Cluster8126-010 | TALE | BEL1-type homeodomain | 2968 | 0 | 0 | 0 |
| SALT00000031150_Cluster8126-011 | TALE | BEL1-type homeodomain | 2683 | 0 | 0 | 0 |
| SALT00000043916_Cluster8126-012 | TALE | BEL1-type homeodomain | 1816 | 0 | 0 | 0 |
| SALT00000064056_Cluster8126-013 | TALE | BEL1-type homeodomain | 3251 | 0 | 0 | 0 |
| SALT00000066437_Cluster8126-014 | TALE | BEL1-type homeodomain | 2767 | 0 | 0 | 0 |
| SALT00000069917_Cluster8126-015 | TALE | BEL1-type homeodomain | 2788 | 0 | 0 | 0 |
| SALT00000071157_Cluster8126-016 | TALE | BEL1-type homeodomain | 2835 | 0 | 0 | 0 |
| SALT00000072038_Cluster8126-017 | TALE | BEL1-type homeodomain | 3041 | 0 | 0 | 0 |
| SALT00000075561_Cluster8126-018 | TALE | BEL1-type homeodomain | 3028 | 0 | 0 | 0 |
| SALT00000076238_Cluster8126-019 | TALE | BEL1-type homeodomain | 3211 | 0 | 0 | down |
| SALT00000078368_Cluster8126-020 | TALE | BEL1-type homeodomain | 2688 | 0 | 0 | 0 |
| SALT00000092941_Cluster8126-021 | TALE | BEL1-type homeodomain | 3456 | 0 | 0 | 0 |
| SALT00000078314_Cluster8126-022 | TALE | BEL1-type homeodomain | 3022 | 0 | 0 | 0 |
| SALT00000021052_Cluster10085-001 | TALE | BEL1-type homeodomain | 2695 | 0 | 0 | 0 |
| SALT00000025391_Cluster10085-002 | TALE | BEL1-type homeodomain | 3070 | 0 | 0 | 0 |
| SALT00000025423_Cluster10085-003 | TALE | BEL1-type homeodomain | 2771 | 0 | 0 | 0 |
| SALT00000027148_Cluster10085-004 | TALE | BEL1-type homeodomain | 2819 | 0 | 0 | 0 |
| SALT00000027270_Cluster10085-005 | TALE | BEL1-type homeodomain | 2907 | 0 | 0 | 0 |
| SALT00000063410_Cluster10085-006 | TALE | BEL1-type homeodomain | 2957 | 0 | 0 | 0 |
| SALT00000063998_Cluster10085-007 | TALE | BEL1-type homeodomain | 2745 | 0 | 0 | 0 |
| SALT00000070047_Cluster10085-008 | TALE | BEL1-type homeodomain | 3074 | 0 | 0 | 0 |
| SALT00000072071_Cluster10085-009 | TALE | BEL1-type homeodomain | 3096 | 0 | 0 | 0 |
| SALT00000072592_Cluster10085-010 | TALE | BEL1-type homeodomain | 2778 | 0 | 0 | 0 |
| SALT00000072796_Cluster10085-011 | TALE | BEL1-type homeodomain | 2557 | 0 | 0 | 0 |
| SALT00000076758_Cluster10085-012 | TALE | BEL1-type homeodomain | 2866 | 0 | 0 | 0 |
| SALT00000077302_Cluster10085-013 | TALE | BEL1-type homeodomain | 2977 | 0 | 0 | 0 |
| SALT00000050424_Cluster10085-014 | TALE | BEL1-type homeodomain | 1803 | 0 | 0 | 0 |
| SALT00000027209_Cluster10085-015 | TALE | BEL1-type homeodomain | 2955 | 0 | 0 | 0 |
| SALT00000021701_Cluster10880-001 | TALE | BEL1-like homeodomain 7 | 2618 | 0 | 0 | 0 |
| SALT00000031743_Cluster10880-002 | TALE | BEL1-like homeodomain 7 | 2659 | 0 | 0 | 0 |
| SALT00000035277_Cluster10880-003 | TALE | BEL1-like homeodomain 7 | 2728 | 0 | 0 | 0 |
| SALT00000062599_Cluster10880-004 | TALE | BEL1-like homeodomain 7 | 2988 | 0 | 0 | 0 |
| SALT00000070054_Cluster10880-005 | TALE | BEL1-like homeodomain 7 | 2792 | 0 | 0 | 0 |
| SALT00000073763_Cluster10880-006 | TALE | BEL1-like homeodomain 7 | 2797 | 0 | 0 | 0 |
| SALT00000078537_Cluster10880-007 | TALE | BEL1-like homeodomain 7 | 2829 | 0 | 0 | 0 |
| SALT00000078680_Cluster10880-008 | TALE | BEL1-like homeodomain 7 | 2728 | 0 | 0 | 0 |
| SALT00000079622_Cluster10880-009 | TALE | BEL1-like homeodomain 7 | 2692 | 0 | 0 | 0 |
| SALT00000013650_Cluster10900-001 | TALE | BEL1-like homeodomain 1 | 1741 | 0 | 0 | 0 |
| SALT00000024501_Cluster10900-002 | TALE | BEL1-like homeodomain 1 | 2560 | 0 | 0 | 0 |
| SALT00000026533_Cluster10900-003 | TALE | BEL1-like homeodomain 1 | 2985 | 0 | 0 | 0 |
| SALT00000040221_Cluster10900-004 | TALE | BEL1-like homeodomain 1 | 1449 | up | 0 | 0 |
| SALT00000075113_Cluster10900-005 | TALE | BEL1-like homeodomain 1 | 2525 | 0 | 0 | 0 |

| | | | | | | |
|---|---|---|---|---|---|---|
| SALT00000077963_Cluster10900-006 | TALE | BEL1-like homeodomain 1 | 2573 | 0 | 0 | 0 |
| SALT00000080522_Cluster10900-007 | TALE | BEL1-like homeodomain 1 | 2557 | 0 | 0 | 0 |
| SALT00000082625_Cluster10900-008 | TALE | BEL1-like homeodomain 1 | 2699 | 0 | 0 | 0 |
| SALT00000079708_Cluster11164-001 | TALE | BEL1-like homeodomain 7 | 2950 | 0 | 0 | 0 |
| SALT00000005269_Cluster11919-001 | TALE | BEL1-related homeotic 14 | 1857 | 0 | 0 | 0 |
| SALT00000012804_Cluster11919-002 | TALE | BEL1-related homeotic 14 | 2165 | 0 | 0 | 0 |
| SALT00000022231_Cluster11919-003 | TALE | BEL1-related homeotic 14 | 2436 | 0 | 0 | 0 |
| SALT00000022855_Cluster11919-004 | TALE | BEL1-related homeotic 14 | 2390 | 0 | 0 | 0 |
| SALT00000044838_Cluster11919-005 | TALE | BEL1-related homeotic 14 | 1476 | 0 | 0 | 0 |
| SALT00000063718_Cluster11919-006 | TALE | BEL1-related homeotic 14 | 2876 | 0 | 0 | 0 |
| SALT00000065081_Cluster11919-007 | TALE | BEL1-related homeotic 14 | 2449 | 0 | 0 | 0 |
| SALT00000075452_Cluster11919-008 | TALE | BEL1-related homeotic 14 | 2717 | 0 | 0 | 0 |
| SALT00000080618_Cluster11919-009 | TALE | BEL1-related homeotic 14 | 2449 | 0 | 0 | 0 |
| SALT00000065506_Cluster12100-002 | TALE | BEL1-type homeodomain | 2858 | 0 | 0 | 0 |
| SALT00000021954_Cluster12269-001 | TALE | BEL1-like homeodomain 7 | 2610 | 0 | 0 | 0 |
| SALT00000080185_Cluster12269-002 | TALE | BEL1-like homeodomain 7 | 2561 | 0 | 0 | 0 |
| SALT00000081758_Cluster12269-003 | TALE | BEL1-like homeodomain 7 | 2842 | 0 | 0 | 0 |
| SALT00000041473_Cluster12269-004 | TALE | BEL1-related homeotic 30 | 2285 | 0 | 0 | 0 |
| SALT00000073101_Cluster13368-001 | TALE | BEL1-like homeodomain 1 | 2743 | 0 | 0 | 0 |
| SALT00000021553_Cluster13423-001 | TALE | BEL1-like homeodomain 7 | 2503 | 0 | 0 | 0 |
| SALT00000022407_Cluster13423-002 | TALE | BEL1-like homeodomain 7 | 2495 | 0 | 0 | 0 |
| SALT00000029840_Cluster13423-003 | TALE | BEL1-like homeodomain 7 | 2459 | 0 | 0 | 0 |
| SALT00000082880_Cluster13423-004 | TALE | BEL1-like homeodomain 7 | 2739 | 0 | 0 | 0 |
| SALT00000021941_Cluster13783-001 | TALE | BEL1-type homeodomain | 2672 | 0 | 0 | 0 |
| SALT00000078379_Cluster14734-001 | TALE | POX domain homeobox D | 2644 | 0 | 0 | 0 |
| SALT00000021798_Cluster15065-001 | TALE | BEL1-like homeodomain 7 | 2620 | 0 | 0 | 0 |
| SALT00000024669_Cluster15065-002 | TALE | BEL1-like homeodomain 7 | 2576 | 0 | 0 | 0 |
| SALT00000071353_Cluster15065-003 | TALE | BEL1-like homeodomain 7 | 2496 | 0 | 0 | up |
| SALT00000073652_Cluster15065-004 | TALE | BEL1-like homeodomain 7 | 2560 | 0 | 0 | 0 |
| SALT00000021105_Cluster15402-001 | TALE | BEL1-related homeotic 14 | 2537 | 0 | 0 | 0 |
| SALT00000022540_Cluster15402-002 | TALE | BEL1-related homeotic 14 | 2496 | 0 | 0 | 0 |
| SALT00000024311_Cluster15402-003 | TALE | BEL1-related homeotic 14 | 2597 | 0 | 0 | 0 |
| SALT00000083391_Cluster15402-004 | TALE | BEL1-related homeotic 14 | 2449 | 0 | 0 | 0 |
| SALT00000003674_Cluster17875-001 | TALE | POX domain homeobox D | 2192 | 0 | 0 | 0 |
| SALT00000043265_Cluster17875-002 | TALE | BEL1-like homeodomain 9 | 2275 | 0 | 0 | 0 |
| SALT00000075592_Cluster17875-003 | TALE | BEL1-like homeodomain 9 | 2328 | 0 | 0 | 0 |
| SALT00000081536_Cluster17875-004 | TALE | POX domain homeobox D | 2317 | 0 | 0 | 0 |
| SALT00000004757_Cluster18775-001 | TALE | transcription factor qSH-1 | 2041 | 0 | 0 | 0 |
| SALT00000009860_Cluster18775-002 | TALE | transcription factor qSH-1 | 2009 | 0 | 0 | 0 |
| SALT00000010879_Cluster18775-003 | TALE | transcription factor qSH-1 | 2187 | 0 | 0 | 0 |
| SALT00000012383_Cluster18775-004 | TALE | transcription factor qSH-1 | 1907 | 0 | 0 | 0 |
| SALT00000019374_Cluster18775-005 | TALE | transcription factor qSH-1 | 1916 | 0 | 0 | 0 |
| SALT00000001481_Cluster19344-001 | TALE | homeobox knotted-1-like 1 | 1654 | 0 | 0 | 0 |
| SALT00000010070_Cluster19344-002 | TALE | homeobox knotted-1-like 1 | 1709 | 0 | 0 | 0 |
| SALT00000020274_Cluster19344-003 | TALE | homeobox knotted-1-like 1 | 1633 | 0 | 0 | 0 |
| SALT00000010076_Cluster20266-001 | TALE | transcription factor qSH-1 | 2057 | 0 | 0 | 0 |
| SALT00000019492_Cluster20266-002 | TALE | transcription factor qSH-1 | 1717 | 0 | 0 | up |
| SALT00000049002_Cluster20266-003 | TALE | transcription factor qSH-1 | 2037 | 0 | 0 | up |
| SALT00000041599_Cluster20266-004 | TALE | transcription factor qSH-1 | 1860 | 0 | 0 | up |
| SALT00000044526_Cluster20266-005 | TALE | transcription factor qSH-1 | 1956 | 0 | 0 | 0 |
| SALT00000053685_Cluster23040-001 | TALE | homeobox OSH1 [Zea ma | 1875 | 0 | 0 | 0 |
| SALT00000002238_Cluster23527-001 | TALE | homeobox knotted-1-like 1 | 1546 | 0 | 0 | 0 |
| SALT00000012024_Cluster23527-002 | TALE | homeobox knotted-1-like 1 | 1733 | 0 | 0 | 0 |
| SALT00000049704_Cluster23527-003 | TALE | homeobox knotted-1-like 1 | 1641 | 0 | 0 | 0 |
| SALT00000049825_Cluster23527-004 | TALE | homeobox knotted-1-like 1 | 1723 | 0 | 0 | 0 |
| SALT00000053311_Cluster23527-005 | TALE | homeobox knotted-1-like 1 | 1845 | 0 | 0 | 0 |
| SALT00000055440_Cluster23527-006 | TALE | homeobox knotted-1-like 1 | 1574 | 0 | 0 | 0 |
| SALT00000001357_Cluster26462-001 | TALE | homeobox OSH1 [Zea ma | 1527 | 0 | 0 | 0 |

| | | | | | | |
|---|---|---|---|---|---|---|
| SALT00000040681_Cluster26462-002 | TALE | homeobox OSH1 [Zea may | 1686 | 0 | 0 | 0 |
| SALT00000001199_Cluster26620-001 | TALE | homeobox rough sheath 1 | 1494 | 0 | 0 | 0 |
| SALT00000043192_Cluster26620-002 | TALE | homeobox rough sheath 1 | 1679 | 0 | 0 | 0 |
| SALT00000055454_Cluster26620-003 | TALE | homeobox rough sheath 1 | 1365 | 0 | 0 | 0 |
| SALT00000083680_Cluster26620-004 | TALE | homeobox rough sheath 1 | 1438 | 0 | 0 | 0 |
| SALT00000035139_Cluster26723-001 | TALE | homeobox OSH1 [Zea may | 1602 | 0 | 0 | 0 |
| SALT00000035411_Cluster26723-002 | TALE | homeobox OSH1 [Zea may | 1617 | 0 | 0 | 0 |
| SALT00000041486_Cluster26723-003 | TALE | homeobox OSH1 [Zea may | 1674 | 0 | 0 | 0 |
| SALT00000054857_Cluster26723-004 | TALE | homeobox OSH1 [Zea may | 1597 | 0 | 0 | 0 |
| SALT00000057901_Cluster26723-005 | TALE | homeobox OSH1 [Zea may | 1474 | 0 | 0 | 0 |
| SALT00000009359_Cluster27319-001 | TALE | homeotic knotted-1 [Zea m | 1644 | 0 | 0 | 0 |
| SALT00000055941_Cluster27319-002 | TALE | homeotic knotted-1 [Zea m | 1477 | 0 | 0 | 0 |
| SALT00000055959_Cluster27319-003 | TALE | homeotic knotted-1 [Zea m | 1550 | 0 | 0 | 0 |
| SALT00000060484_Cluster27319-004 | TALE | homeotic knotted-1 [Zea m | 1545 | 0 | 0 | 0 |
| SALT00000037224_Cluster27885-001 | TALE | Os03g0680700 [Oryza sati | 1616 | 0 | 0 | 0 |
| SALT00000043244_Cluster30254-001 | TALE | homeobox knotted-1-like 1 | 1485 | 0 | 0 | 0 |
| SALT00000052845_Cluster30254-002 | TALE | homeobox knotted-1-like 1 | 1465 | 0 | 0 | 0 |
| SALT00000001457_Cluster31410-001 | TALE | knotted-like transcription f | 1211 | 0 | 0 | 0 |
| SALT00000013689_Cluster31410-002 | TALE | homeobox knotted-1-like 1 | 1350 | 0 | 0 | 0 |
| SALT00000015833_Cluster31410-003 | TALE | homeobox knotted-1-like 1 | 1317 | 0 | 0 | down |
| SALT00000038688_Cluster31410-004 | TALE | homeobox knotted-1-like 1 | 1351 | 0 | 0 | 0 |
| SALT00000042021_Cluster31410-005 | TALE | homeobox knotted-1-like 1 | 1241 | 0 | 0 | 0 |
| SALT00000049293_Cluster31410-006 | TALE | homeobox knotted-1-like 1 | 1349 | 0 | 0 | 0 |
| SALT00000055041_Cluster31410-007 | TALE | knotted-like transcription f | 1371 | 0 | up | up |
| SALT00000046678_Cluster31410-008 | TALE | homeobox knotted-1-like 1 | 1339 | down | 0 | down |
| SALT00000049695_Cluster441-001 | TCP | TCP-domain [Oryza sativa | 8041 | 0 | 0 | down |
| SALT00000023040_Cluster14541-001 | TCP | transcription factor PCF2-l | 2656 | 0 | 0 | 0 |
| SALT00000047924_Cluster18811-001 | TCP | LOC100281134 isoform X | 2184 | 0 | 0 | 0 |
| SALT00000017369_Cluster18955-001 | TCP | TCP-domain [Oryza sativa | 1599 | 0 | 0 | 0 |
| SALT00000039515_Cluster18955-002 | TCP | TCP-domain [Oryza sativa | 1564 | 0 | 0 | 0 |
| SALT00000041108_Cluster18955-003 | TCP | TCP-domain [Oryza sativa | 1677 | 0 | 0 | 0 |
| SALT00000048118_Cluster18955-004 | TCP | TCP-domain [Oryza sativa | 2171 | 0 | 0 | 0 |
| SALT00000048888_Cluster18955-005 | TCP | TCP-domain [Oryza sativa | 1939 | 0 | 0 | 0 |
| SALT00000049306_Cluster18955-006 | TCP | TCP-domain [Oryza sativa | 1329 | 0 | 0 | 0 |
| SALT00000049981_Cluster18955-007 | TCP | TCP-domain [Oryza sativa | 1620 | 0 | 0 | 0 |
| SALT00000050604_Cluster18955-008 | TCP | TCP-domain [Oryza sativa | 1911 | 0 | 0 | 0 |
| SALT00000050814_Cluster18955-009 | TCP | TCP-domain [Oryza sativa | 1537 | 0 | 0 | 0 |
| SALT00000053617_Cluster18955-010 | TCP | TCP-domain [Oryza sativa | 2130 | 0 | 0 | 0 |
| SALT00000061497_Cluster18955-011 | TCP | TCP-domain [Oryza sativa | 1563 | 0 | 0 | 0 |
| SALT00000075293_Cluster18955-012 | TCP | TCP-domain [Oryza sativa | 2014 | 0 | 0 | 0 |
| SALT00000001735_Cluster21666-001 | TCP | transcription factor PCF6 [ | 1961 | 0 | 0 | 0 |
| SALT00000064493_Cluster21666-002 | TCP | transcription factor PCF6 [ | 1686 | 0 | 0 | 0 |
| SALT00000012963_Cluster22695-001 | TCP | TCP transcription partial [ | 1471 | 0 | 0 | 0 |
| SALT00000037682_Cluster22695-002 | TCP | TCP transcription partial [ | 1897 | 0 | 0 | 0 |
| SALT00000042851_Cluster22695-003 | TCP | TCP transcription partial [ | 1711 | 0 | 0 | 0 |
| SALT00000044371_Cluster22695-004 | TCP | TCP transcription partial [ | 1881 | 0 | 0 | 0 |
| SALT00000045195_Cluster22695-005 | TCP | TCP transcription partial [ | 1852 | 0 | 0 | 0 |
| SALT00000045703_Cluster22695-006 | TCP | TCP transcription partial [ | 1802 | 0 | 0 | 0 |
| SALT00000050421_Cluster22695-007 | TCP | TCP transcription partial [ | 1820 | 0 | 0 | down |
| SALT00000039962_Cluster22695-008 | TCP | TCP transcription partial [ | 1856 | 0 | 0 | 0 |
| SALT00000039301_Cluster24191-001 | TCP | TCP transcription partial [ | 1808 | 0 | 0 | 0 |
| SALT00000049900_Cluster24191-002 | TCP | TCP transcription partial [ | 1578 | 0 | 0 | 0 |
| SALT00000067657_Cluster24191-003 | TCP | TCP transcription partial [ | 1603 | 0 | 0 | 0 |
| SALT00000051351_Cluster24407-001 | TCP | TCP transcription partial [ | 1619 | 0 | 0 | 0 |
| SALT00000057288_Cluster24407-002 | TCP | TCP transcription partial [ | 1644 | 0 | 0 | 0 |
| SALT00000057552_Cluster24407-003 | TCP | TCP transcription partial [ | 1796 | 0 | 0 | 0 |
| SALT00000011553_Cluster25313-001 | TCP | TCP transcription partial [ | 1746 | 0 | 0 | 0 |
| SALT00000015339_Cluster25313-002 | TCP | TCP transcription partial [ | 1604 | 0 | 0 | 0 |

| | | | | | | |
|---|---|---|---|---|---|---|
| SALT00000057790_Cluster25313-003 | TCP | TCP transcription partial [ | 1628 | 0 | 0 | 0 |
| SALT00000009165_Cluster25627-001 | TCP | partial [Hordeum vulgare v | 1712 | 0 | 0 | 0 |
| SALT00000014293_Cluster25627-002 | TCP | partial [Hordeum vulgare v | 1731 | 0 | 0 | 0 |
| SALT00000018222_Cluster25627-003 | TCP | partial [Hordeum vulgare v | 1528 | 0 | 0 | 0 |
| SALT00000020025_Cluster25627-004 | TCP | partial [Hordeum vulgare v | 1509 | 0 | 0 | 0 |
| SALT00000039525_Cluster25627-005 | TCP | partial [Hordeum vulgare v | 1577 | 0 | 0 | 0 |
| SALT00000015445_Cluster26259-001 | TCP | TCP-domain [Zea mays] | 1536 | 0 | 0 | 0 |
| SALT00000025017_Cluster26259-002 | TCP | TCP-domain [Zea mays] | 1555 | 0 | 0 | 0 |
| SALT00000036101_Cluster26259-003 | TCP | TCP-domain [Zea mays] | 1642 | 0 | 0 | up |
| SALT00000040211_Cluster26259-004 | TCP | TCP-domain [Zea mays] | 1696 | 0 | 0 | up |
| SALT00000041007_Cluster26259-005 | TCP | TCP-domain [Zea mays] | 1545 | 0 | 0 | 0 |
| SALT00000044737_Cluster26259-006 | TCP | TCP-domain [Zea mays] | 1637 | 0 | 0 | 0 |
| SALT00000048348_Cluster26259-007 | TCP | [Hordeum vulgare vulgare] | 1490 | 0 | 0 | 0 |
| SALT00000003425_Cluster26460-001 | TCP | TCP family transcription fa | 1576 | 0 | 0 | 0 |
| SALT00000036706_Cluster26460-002 | TCP | TCP family transcription fa | 1670 | 0 | 0 | 0 |
| SALT00000038534_Cluster26460-003 | TCP | TCP family transcription fa | 1686 | 0 | 0 | 0 |
| SALT00000013713_Cluster26658-001 | TCP | TCP-domain [Oryza sativa | 1677 | 0 | 0 | 0 |
| SALT00000047731_Cluster26738-001 | TCP | TCP-domain [Oryza sativa | 1673 | 0 | 0 | 0 |
| SALT00000022190_Cluster27884-001 | TCP | transcription factor TCP7-l | 1616 | 0 | 0 | 0 |
| SALT00000005791_Cluster28176-001 | TCP | TCP-domain [Zea mays] | 1530 | 0 | 0 | 0 |
| SALT00000015622_Cluster28176-002 | TCP | TCP-domain [Zea mays] | 1601 | 0 | 0 | 0 |
| SALT00000027925_Cluster28176-003 | TCP | TCP-domain [Zea mays] | 1505 | 0 | 0 | 0 |
| SALT00000057909_Cluster28176-004 | TCP | TCP-domain [Zea mays] | 1479 | down | 0 | 0 |
| SALT00000006831_Cluster29960-001 | TCP | transcription factor TCP13 | 1504 | 0 | 0 | 0 |
| SALT00000037036_Cluster29960-002 | TCP | transcription factor TCP13 | 1450 | 0 | 0 | 0 |
| SALT00000066921_Cluster31858-001 | TCP | transcription factor TCP7-l | 1234 | 0 | 0 | 0 |
| SALT00000026358_Cluster4060-001 | Trihelix | partial [Hordeum vulgare v | 2627 | 0 | 0 | 0 |
| SALT00000056145_Cluster4060-002 | Trihelix | trihelix transcription factor | 1825 | 0 | 0 | 0 |
| SALT00000064137_Cluster4060-003 | Trihelix | partial [Hordeum vulgare v | 2507 | 0 | 0 | 0 |
| SALT00000066890_Cluster4060-004 | Trihelix | partial [Hordeum vulgare v | 3115 | 0 | 0 | 0 |
| SALT00000070426_Cluster4060-005 | Trihelix | partial [Hordeum vulgare v | 4203 | 0 | 0 | 0 |
| SALT00000079071_Cluster4060-006 | Trihelix | partial [Hordeum vulgare v | 2910 | 0 | 0 | 0 |
| SALT00000005625_Cluster5888-001 | Trihelix | 6b-interacting 1 [Zea mays | 1587 | 0 | 0 | 0 |
| SALT00000018360_Cluster5888-003 | Trihelix | 6b-interacting 1 [Zea mays | 1645 | 0 | 0 | 0 |
| SALT00000037621_Cluster5888-004 | Trihelix | 6b-interacting 1 [Zea mays | 3809 | 0 | 0 | 0 |
| SALT00000043046_Cluster5888-005 | Trihelix | 6b-interacting 1 [Zea mays | 1588 | 0 | 0 | 0 |
| SALT00000049017_Cluster5888-006 | Trihelix | 6b-interacting 1 [Zea mays | 1521 | 0 | 0 | 0 |
| SALT00000028046_Cluster6238-001 | Trihelix | partial [Hordeum vulgare v | 2643 | 0 | 0 | 0 |
| SALT00000032771_Cluster6238-002 | Trihelix | trihelix transcription factor | 3313 | 0 | 0 | 0 |
| SALT00000057362_Cluster6238-003 | Trihelix | partial [Hordeum vulgare v | 1858 | 0 | 0 | 0 |
| SALT00000058896_Cluster6238-004 | Trihelix | trihelix transcription factor | 2177 | 0 | 0 | 0 |
| SALT00000083519_Cluster6238-005 | Trihelix | trihelix transcription factor | 2798 | 0 | 0 | 0 |
| SALT00000087357_Cluster6238-006 | Trihelix | partial [Hordeum vulgare v | 3755 | 0 | 0 | 0 |
| SALT00000074970_Cluster6238-007 | Trihelix | partial [Hordeum vulgare v | 3192 | up | 0 | up |
| SALT00000065109_Cluster6238-008 | Trihelix | trihelix transcription factor | 2525 | 0 | 0 | 0 |
| SALT00000007266_Cluster6929-001 | Trihelix | Os05g0560600 [Oryza sati | 1387 | 0 | 0 | 0 |
| SALT00000087355_Cluster6929-002 | Trihelix | Os05g0560600 [Oryza sati | 3647 | 0 | 0 | 0 |
| SALT00000062728_Cluster6929-003 | Trihelix | 6b-interacting 1 [Zea mays | 2524 | 0 | 0 | 0 |
| SALT00000061978_Cluster7003-001 | Trihelix | trihelix transcription factor | 2866 | 0 | 0 | 0 |
| SALT00000063987_Cluster7003-002 | Trihelix | trihelix transcription factor | 2746 | 0 | 0 | 0 |
| SALT00000081535_Cluster7003-003 | Trihelix | trihelix transcription factor | 2997 | 0 | 0 | 0 |
| SALT00000089642_Cluster7003-004 | Trihelix | trihelix transcription factor | 3400 | 0 | 0 | 0 |
| SALT00000092330_Cluster7003-005 | Trihelix | trihelix transcription factor | 3636 | 0 | 0 | 0 |
| SALT00000040515_Cluster7003-006 | Trihelix | trihelix transcription factor | 2112 | 0 | 0 | 0 |
| SALT00000060215_Cluster7003-007 | Trihelix | trihelix transcription factor | 2029 | 0 | 0 | 0 |
| SALT00000038175_Cluster7003-008 | Trihelix | trihelix transcription factor | 1917 | 0 | 0 | 0 |
| SALT00000021929_Cluster8035-001 | Trihelix | ribonuclease J isoform X1 | 3014 | 0 | 0 | 0 |
| SALT00000022004_Cluster8035-002 | Trihelix | ribonuclease J isoform X1 | 3040 | 0 | 0 | 0 |

| | | | | | | |
|---|---|---|---|---|---|---|
| SALT00000023179_Cluster8035-003 | Trihelix | ribonuclease J isoform X1 | 2860 | 0 | 0 | 0 |
| SALT00000030965_Cluster8035-004 | Trihelix | ribonuclease J isoform X2 | 2454 | 0 | 0 | 0 |
| SALT00000032782_Cluster8035-005 | Trihelix | ribonuclease J isoform X1 | 3475 | 0 | 0 | 0 |
| SALT00000062616_Cluster8035-006 | Trihelix | ribonuclease J isoform X2 | 1499 | 0 | 0 | 0 |
| SALT00000064594_Cluster8035-007 | Trihelix | ribonuclease J isoform X1 | 3079 | 0 | 0 | 0 |
| SALT00000064983_Cluster8035-008 | Trihelix | ribonuclease J isoform X1 | 2862 | 0 | 0 | 0 |
| SALT00000093231_Cluster8035-009 | Trihelix | ribonuclease J isoform X2 | 3451 | 0 | 0 | 0 |
| SALT00000008269_Cluster8232-001 | Trihelix | partial [Oryza sativa Japon | 1927 | 0 | 0 | 0 |
| SALT00000026441_Cluster8232-002 | Trihelix | partial [Oryza sativa Japon | 2678 | 0 | 0 | 0 |
| SALT00000041012_Cluster8232-003 | Trihelix | partial [Oryza sativa Japon | 1611 | 0 | 0 | down |
| SALT00000068241_Cluster8232-004 | Trihelix | partial [Oryza sativa Japon | 2776 | 0 | 0 | 0 |
| SALT00000073122_Cluster8232-005 | Trihelix | partial [Oryza sativa Japon | 2823 | 0 | 0 | 0 |
| SALT00000089624_Cluster8232-006 | Trihelix | partial [Oryza sativa Japon | 3438 | 0 | 0 | 0 |
| SALT00000064849_Cluster8232-007 | Trihelix | partial [Oryza sativa Japon | 2988 | 0 | 0 | 0 |
| SALT00000028780_Cluster8853-001 | Trihelix | partial [Hordeum vulgare v | 3315 | 0 | 0 | 0 |
| SALT00000066888_Cluster8853-002 | Trihelix | trihelix transcription factor | 2989 | 0 | 0 | 0 |
| SALT00000071456_Cluster8853-003 | Trihelix | partial [Hordeum vulgare v | 3097 | 0 | 0 | 0 |
| SALT00000074111_Cluster8853-004 | Trihelix | partial [Hordeum vulgare v | 2810 | 0 | 0 | 0 |
| SALT00000020151_Cluster9634-001 | Trihelix | trihelix transcription factor | 2129 | 0 | 0 | 0 |
| SALT00000022107_Cluster9634-002 | Trihelix | partial [Oryza sativa Japon | 2958 | 0 | 0 | 0 |
| SALT00000053511_Cluster9634-003 | Trihelix | trihelix transcription factor | 1957 | 0 | 0 | 0 |
| SALT00000063982_Cluster9634-004 | Trihelix | trihelix transcription factor | 3167 | 0 | 0 | 0 |
| SALT00000083092_Cluster9634-005 | Trihelix | trihelix transcription factor | 2492 | 0 | 0 | 0 |
| SALT00000014905_Cluster10722-001 | Trihelix | partial [Oryza sativa Japon | 2134 | 0 | 0 | 0 |
| SALT00000023497_Cluster10722-002 | Trihelix | partial [Oryza sativa Japon | 2886 | down | 0 | down |
| SALT00000029636_Cluster10722-003 | Trihelix | partial [Oryza sativa Japon | 3006 | 0 | 0 | 0 |
| SALT00000069195_Cluster10722-004 | Trihelix | partial [Oryza sativa Japon | 2975 | 0 | 0 | 0 |
| SALT00000072489_Cluster10722-005 | Trihelix | partial [Oryza sativa Japon | 2700 | 0 | 0 | 0 |
| SALT00000080040_Cluster10990-001 | Trihelix | partial [Oryza sativa Japon | 2927 | 0 | 0 | 0 |
| SALT00000080164_Cluster10990-002 | Trihelix | partial [Oryza sativa Japon | 2973 | 0 | 0 | 0 |
| SALT00000082728_Cluster10990-003 | Trihelix | partial [Oryza sativa Japon | 2441 | 0 | 0 | 0 |
| SALT00000083470_Cluster10990-004 | Trihelix | partial [Oryza sativa Japon | 2960 | 0 | 0 | 0 |
| SALT00000083495_Cluster10990-005 | Trihelix | partial [Oryza sativa Japon | 2952 | 0 | 0 | 0 |
| SALT00000026676_Cluster11457-001 | Trihelix | trihelix transcription factor | 2802 | 0 | 0 | 0 |
| SALT00000045700_Cluster11457-002 | Trihelix | trihelix transcription factor | 2365 | 0 | 0 | 0 |
| SALT00000073611_Cluster11457-003 | Trihelix | trihelix transcription factor | 2533 | 0 | 0 | 0 |
| SALT00000074021_Cluster11457-004 | Trihelix | trihelix transcription factor | 2863 | 0 | 0 | 0 |
| SALT00000077950_Cluster11457-005 | Trihelix | trihelix transcription factor | 2872 | 0 | down | 0 |
| SALT00000083171_Cluster11457-006 | Trihelix | trihelix transcription factor | 2920 | 0 | 0 | 0 |
| SALT00000023165_Cluster11681-001 | Trihelix | trihelix transcription factor | 2895 | 0 | 0 | 0 |
| SALT00000072221_Cluster11681-002 | Trihelix | trihelix transcription factor | 2862 | 0 | 0 | 0 |
| SALT00000075274_Cluster11681-003 | Trihelix | trihelix transcription factor | 2812 | 0 | 0 | 0 |
| SALT00000074809_Cluster12935-008 | Trihelix | polyol transporter 5-like | 2780 | 0 | 0 | 0 |
| SALT00000007016_Cluster15899-001 | Trihelix | trihelix transcription factor | 2016 | 0 | 0 | 0 |
| SALT00000039522_Cluster15899-002 | Trihelix | trihelix transcription factor | 1903 | 0 | 0 | 0 |
| SALT00000043750_Cluster15899-003 | Trihelix | trihelix transcription factor | 1934 | 0 | 0 | 0 |
| SALT00000053035_Cluster15899-004 | Trihelix | trihelix transcription factor | 1959 | 0 | 0 | 0 |
| SALT00000060932_Cluster15899-005 | Trihelix | trihelix transcription factor | 2002 | 0 | 0 | 0 |
| SALT00000062130_Cluster15899-006 | Trihelix | trihelix transcription factor | 2559 | 0 | 0 | 0 |
| SALT00000083105_Cluster15899-007 | Trihelix | trihelix transcription factor | 2209 | 0 | 0 | 0 |
| SALT00000081438_Cluster16474-001 | Trihelix | trihelix transcription factor | 2518 | 0 | 0 | 0 |
| SALT00000082606_Cluster16474-002 | Trihelix | trihelix transcription factor | 2453 | down | 0 | 0 |
| SALT00000002292_Cluster17474-001 | Trihelix | Os05g0560600 [Oryza sati | 1495 | 0 | 0 | 0 |
| SALT00000010038_Cluster17474-002 | Trihelix | 6b-interacting 1 [Zea mays | 1473 | 0 | 0 | 0 |
| SALT00000012163_Cluster17474-003 | Trihelix | 6b-interacting 1 [Zea mays | 1468 | 0 | 0 | 0 |
| SALT00000038085_Cluster17474-004 | Trihelix | Os05g0560600 [Oryza sati | 1568 | 0 | 0 | 0 |
| SALT00000040460_Cluster17474-005 | Trihelix | Os05g0560600 [Oryza sati | 1624 | 0 | 0 | 0 |
| SALT00000045162_Cluster17474-006 | Trihelix | Os05g0560600 [Oryza sati | 1536 | 0 | 0 | 0 |

| | | | | | | |
|---|---|---|---|---|---|---|
| SALT00000081499_Cluster17474-007 | Trihelix | Os05g0560600 [Oryza sati | 2410 | 0 | 0 | 0 |
| SALT00000067580_Cluster18120-001 | Trihelix | trihelix transcription factor | 2278 | 0 | 0 | down |
| SALT00000060656_Cluster19414-001 | Trihelix | [Hordeum vulgare vulgare] | 2127 | 0 | 0 | 0 |
| SALT00000057640_Cluster19914-001 | Trihelix | [Hordeum vulgare vulgare] | 2087 | 0 | 0 | 0 |
| SALT00000001236_Cluster20012-001 | Trihelix | trihelix transcription partia | 1924 | 0 | 0 | 0 |
| SALT00000017598_Cluster20012-002 | Trihelix | trihelix transcription partia | 1816 | 0 | 0 | 0 |
| SALT00000047612_Cluster20012-003 | Trihelix | trihelix transcription partia | 1882 | 0 | 0 | 0 |
| SALT00000050526_Cluster20012-004 | Trihelix | trihelix transcription partia | 1816 | 0 | 0 | 0 |
| SALT00000054447_Cluster20012-005 | Trihelix | trihelix transcription partia | 2079 | 0 | 0 | 0 |
| SALT00000056709_Cluster20012-006 | Trihelix | trihelix transcription partia | 1881 | 0 | 0 | 0 |
| SALT00000066944_Cluster20012-007 | Trihelix | trihelix transcription partia | 1868 | 0 | 0 | 0 |
| SALT00000003485_Cluster20012-008 | Trihelix | trihelix transcription partia | 1550 | 0 | 0 | 0 |
| SALT00000002755_Cluster20025-001 | Trihelix | Os02g0542400 [Oryza sati | 1923 | 0 | 0 | 0 |
| SALT00000010479_Cluster20025-002 | Trihelix | Os02g0542400 [Oryza sati | 1876 | 0 | 0 | up |
| SALT00000015724_Cluster20025-003 | Trihelix | Os02g0542400 [Oryza sati | 1588 | 0 | 0 | 0 |
| SALT00000015810_Cluster20025-004 | Trihelix | Os02g0542400 [Oryza sati | 1818 | 0 | 0 | 0 |
| SALT00000036062_Cluster20025-005 | Trihelix | Os02g0542400 [Oryza sati | 1744 | 0 | 0 | 0 |
| SALT00000038498_Cluster20025-006 | Trihelix | Os02g0542400 [Oryza sati | 1521 | 0 | 0 | 0 |
| SALT00000038578_Cluster20025-007 | Trihelix | Os02g0542400 [Oryza sati | 1807 | 0 | 0 | 0 |
| SALT00000047936_Cluster20025-008 | Trihelix | Os02g0542400 [Oryza sati | 1969 | 0 | 0 | 0 |
| SALT00000049738_Cluster20025-009 | Trihelix | Os02g0542400 [Oryza sati | 2078 | 0 | 0 | 0 |
| SALT00000050879_Cluster20025-010 | Trihelix | Os02g0542400 [Oryza sati | 1955 | 0 | 0 | 0 |
| SALT00000051177_Cluster20025-011 | Trihelix | Os02g0542400 [Oryza sati | 1644 | 0 | 0 | 0 |
| SALT00000053198_Cluster20025-012 | Trihelix | Os02g0542400 [Oryza sati | 1966 | 0 | 0 | 0 |
| SALT00000004224_Cluster20482-001 | Trihelix | Os09g0558200 [Oryza sati | 2041 | 0 | 0 | 0 |
| SALT00000006488_Cluster20482-002 | Trihelix | Os09g0558200 [Oryza sati | 1814 | 0 | 0 | 0 |
| SALT00000007834_Cluster20482-003 | Trihelix | Os09g0558200 [Oryza sati | 1774 | 0 | 0 | 0 |
| SALT00000010236_Cluster20482-004 | Trihelix | Os09g0558200 [Oryza sati | 1930 | 0 | 0 | 0 |
| SALT00000011703_Cluster20482-005 | Trihelix | Os09g0558200 [Oryza sati | 1884 | 0 | 0 | 0 |
| SALT00000012289_Cluster20482-006 | Trihelix | Os09g0558200 [Oryza sati | 1926 | 0 | 0 | 0 |
| SALT00000015284_Cluster20482-007 | Trihelix | Os09g0558200 [Oryza sati | 1909 | 0 | 0 | 0 |
| SALT00000016605_Cluster20482-008 | Trihelix | Os09g0558200 [Oryza sati | 1848 | 0 | 0 | 0 |
| SALT00000044257_Cluster20482-009 | Trihelix | Os09g0558200 [Oryza sati | 2010 | 0 | 0 | 0 |
| SALT00000049290_Cluster20482-010 | Trihelix | Os09g0558200 [Oryza sati | 1924 | 0 | 0 | 0 |
| SALT00000061331_Cluster20482-011 | Trihelix | Os09g0558200 [Oryza sati | 2037 | 0 | 0 | 0 |
| SALT00000059240_Cluster20482-012 | Trihelix | Os09g0558200 [Oryza sati | 1528 | 0 | 0 | 0 |
| SALT00000016049_Cluster22301-001 | Trihelix | Os05g0560600 [Oryza sati | 1609 | 0 | 0 | 0 |
| SALT00000057661_Cluster22301-002 | Trihelix | 6b-interacting 1 [Zea mays | 1920 | 0 | 0 | 0 |
| SALT00000066733_Cluster22301-003 | Trihelix | 6b-interacting 1 [Zea mays | 1642 | 0 | 0 | 0 |
| SALT00000061736_Cluster22301-004 | Trihelix | Os05g0560600 [Oryza sati | 1920 | 0 | 0 | 0 |
| SALT00000001112_Cluster22662-001 | Trihelix | PREDICTED: uncharacter | 1780 | 0 | 0 | 0 |
| SALT00000004084_Cluster22662-002 | Trihelix | PREDICTED: uncharacter | 1635 | 0 | 0 | 0 |
| SALT00000004090_Cluster22662-003 | Trihelix | trihelix transcription factor | 1781 | 0 | 0 | 0 |
| SALT00000011362_Cluster22662-004 | Trihelix | PREDICTED: uncharacter | 1719 | 0 | 0 | 0 |
| SALT00000017479_Cluster22662-005 | Trihelix | PREDICTED: uncharacter | 1684 | 0 | 0 | 0 |
| SALT00000019475_Cluster22662-006 | Trihelix | TPA: hypothetical protein | 1618 | 0 | 0 | 0 |
| SALT00000042584_Cluster22662-007 | Trihelix | trihelix transcription factor | 1899 | 0 | 0 | 0 |
| SALT00000016250_Cluster23307-001 | Trihelix | Os05g0560600 [Oryza sati | 1751 | 0 | 0 | 0 |
| SALT00000053751_Cluster23307-002 | Trihelix | Os05g0560600 [Oryza sati | 1858 | 0 | 0 | 0 |
| SALT00000001322_Cluster24324-001 | Trihelix | TPA: hypothetical protein | 1790 | 0 | 0 | 0 |
| SALT00000033532_Cluster24324-002 | Trihelix | TPA: hypothetical protein | 1800 | 0 | 0 | 0 |
| SALT00000034972_Cluster24324-003 | Trihelix | TPA: hypothetical protein | 1775 | 0 | 0 | 0 |
| SALT00000047710_Cluster24324-004 | Trihelix | PREDICTED: uncharacter | 1705 | 0 | 0 | 0 |
| SALT00000057657_Cluster24324-005 | Trihelix | TPA: hypothetical protein | 1575 | 0 | 0 | 0 |
| SALT00000002619_Cluster24724-001 | Trihelix | PREDICTED: uncharacter | 1748 | 0 | 0 | 0 |
| SALT00000047990_Cluster24724-002 | Trihelix | PREDICTED: uncharacter | 1778 | 0 | 0 | 0 |
| SALT00000051253_Cluster24724-003 | Trihelix | PREDICTED: uncharacter | 1705 | 0 | 0 | 0 |
| SALT00000012869_Cluster25134-001 | Trihelix | Trihelix transcription parti | 1422 | 0 | 0 | 0 |

| | | | | | | |
|---|---|---|---|---|---|---|
| SALT00000039195_Cluster25134-002 | Trihelix | Trihelix transcription parti | 1547 | 0 | 0 | 0 |
| SALT00000040861_Cluster25134-003 | Trihelix | Trihelix transcription parti | 1516 | 0 | 0 | up |
| SALT00000044426_Cluster25134-004 | Trihelix | Trihelix transcription parti | 1375 | 0 | 0 | 0 |
| SALT00000045396_Cluster25134-005 | Trihelix | Trihelix transcription parti | 1476 | 0 | 0 | 0 |
| SALT00000051684_Cluster25134-006 | Trihelix | Trihelix transcription parti | 1756 | 0 | 0 | 0 |
| SALT00000053033_Cluster25134-007 | Trihelix | Trihelix transcription parti | 1520 | 0 | 0 | 0 |
| SALT00000056352_Cluster25134-008 | Trihelix | Trihelix transcription parti | 1526 | 0 | 0 | 0 |
| SALT00000004844_Cluster25408-001 | Trihelix | Os02g0565000 [Oryza sati | 1642 | 0 | 0 | 0 |
| SALT00000005342_Cluster25408-002 | Trihelix | Os02g0565000 [Oryza sati | 1426 | 0 | 0 | 0 |
| SALT00000006965_Cluster25408-003 | Trihelix | Os02g0565000 [Oryza sati | 1741 | 0 | 0 | 0 |
| SALT00000009816_Cluster25408-004 | Trihelix | Os02g0565000 [Oryza sati | 1529 | 0 | 0 | 0 |
| SALT00000010126_Cluster25408-005 | Trihelix | Os02g0565000 [Oryza sati | 1623 | 0 | 0 | 0 |
| SALT00000012090_Cluster25408-006 | Trihelix | Os02g0565000 [Oryza sati | 1603 | 0 | 0 | 0 |
| SALT00000039917_Cluster25408-007 | Trihelix | Os02g0565000 [Oryza sati | 1550 | 0 | 0 | 0 |
| SALT00000056654_Cluster25408-008 | Trihelix | Os02g0565000 [Oryza sati | 1554 | 0 | 0 | 0 |
| SALT00000058307_Cluster25408-009 | Trihelix | Os02g0565000 [Oryza sati | 1502 | 0 | 0 | 0 |
| SALT00000017227_Cluster28153-001 | Trihelix | trihelix transcription factor | 1417 | 0 | 0 | 0 |
| SALT00000061365_Cluster28153-002 | Trihelix | trihelix transcription factor | 1603 | 0 | 0 | 0 |
| SALT00000053487_Cluster29587-002 | Trihelix | trihelix transcription factor | 1526 | 0 | 0 | 0 |
| SALT00000014435_Cluster29838-001 | Trihelix | Os10g0564200 [Oryza sati | 1512 | 0 | 0 | 0 |
| SALT00000054061_Cluster30134-001 | Trihelix | homeodomain-like transcri | 1493 | 0 | 0 | 0 |
| SALT00000049640_Cluster30147-001 | Trihelix | trihelix transcription factor | 1492 | 0 | 0 | 0 |
| SALT00000040679_Cluster890-001 | VOZ | transcription factor VOZ1 | 6998 | 0 | 0 | 0 |
| SALT00000045630_Cluster890-002 | VOZ | transcription factor VOZ1 | 2146 | 0 | 0 | 0 |
| SALT00000046071_Cluster890-003 | VOZ | transcription factor VOZ1 | 2040 | 0 | 0 | 0 |
| SALT00000079960_Cluster890-004 | VOZ | transcription factor VOZ1 | 2061 | 0 | 0 | 0 |
| SALT00000055077_Cluster890-005 | VOZ | transcription factor VOZ1 | 2086 | 0 | 0 | 0 |
| SALT00000003998_Cluster18572-001 | VOZ | Os05g0515700 [Oryza sati | 1964 | 0 | 0 | 0 |
| SALT00000005585_Cluster18572-002 | VOZ | Os05g0515700 [Oryza sati | 1717 | 0 | 0 | 0 |
| SALT00000005664_Cluster18572-003 | VOZ | Os05g0515700 [Oryza sati | 1554 | 0 | 0 | 0 |
| SALT00000006988_Cluster18572-004 | VOZ | Os05g0515700 [Oryza sati | 2057 | 0 | 0 | 0 |
| SALT00000007650_Cluster18572-005 | VOZ | Os05g0515700 [Oryza sati | 1909 | 0 | 0 | 0 |
| SALT00000011947_Cluster18572-006 | VOZ | Os05g0515700 [Oryza sati | 1975 | 0 | 0 | 0 |
| SALT00000016670_Cluster18572-007 | VOZ | Os05g0515700 [Oryza sati | 1451 | 0 | 0 | 0 |
| SALT00000052332_Cluster18572-008 | VOZ | Os05g0515700 [Oryza sati | 1128 | 0 | 0 | 0 |
| SALT00000054605_Cluster18572-009 | VOZ | Os05g0515700 [Oryza sati | 1991 | 0 | 0 | 0 |
| SALT00000057279_Cluster18572-010 | VOZ | Os05g0515700 [Oryza sati | 2210 | 0 | 0 | 0 |
| SALT00000086655_Cluster8812-001 | WOX | TPA: homeobox DNA-bin | 3324 | 0 | 0 | 0 |
| SALT00000010115_Cluster28598-001 | WOX | TPA: homeobox DNA-bin | 1578 | 0 | 0 | 0 |
| SALT00000014886_Cluster28598-002 | WOX | TPA: homeobox DNA-bin | 1434 | 0 | 0 | up |
| SALT00000039279_Cluster28598-003 | WOX | TPA: homeobox DNA-bin | 1321 | 0 | 0 | 0 |
| SALT00000060223_Cluster28598-004 | WOX | TPA: homeobox DNA-bin | 1134 | 0 | 0 | 0 |
| SALT00000046391_Cluster32051-001 | WOX | WUSCHEL-related homeo | 1170 | 0 | 0 | 0 |
| SALT00000004607_Cluster727-001 | WRKY | SPF1-like DNA-binding [ | 1898 | 0 | 0 | 0 |
| SALT00000009703_Cluster727-002 | WRKY | WRKY DNA-binding dom | 1900 | 0 | 0 | 0 |
| SALT00000017291_Cluster727-003 | WRKY | SPF1-like DNA-binding [ | 1978 | 0 | 0 | 0 |
| SALT00000019126_Cluster727-004 | WRKY | WRKY DNA-binding dom | 2043 | 0 | 0 | 0 |
| SALT00000053510_Cluster727-006 | WRKY | WRKY DNA-binding dom | 1992 | 0 | 0 | 0 |
| SALT00000050869_Cluster1938-001 | WRKY | WRKY transcription partia | 2048 | 0 | 0 | 0 |
| SALT00000078779_Cluster3483-001 | WRKY | disease resistance RPP13 2 | 4398 | 0 | 0 | 0 |
| SALT00000049652_Cluster7675-001 | WRKY | 3-N-debenzoyl-2-deoxytax | 3532 | 0 | 0 | 0 |
| SALT00000057714_Cluster7675-002 | WRKY | WRKY transcription partia | 1493 | 0 | 0 | 0 |
| SALT00000058026_Cluster7675-003 | WRKY | WRKY transcription partia | 1238 | 0 | 0 | 0 |
| SALT00000058173_Cluster7675-004 | WRKY | WRKY transcription partia | 1478 | 0 | 0 | 0 |
| SALT00000056440_Cluster7675-005 | WRKY | WRKY DNA-binding dom | 1334 | 0 | 0 | 0 |
| SALT00000053101_Cluster7675-006 | WRKY | WRKY transcription partia | 1600 | 0 | 0 | 0 |
| SALT00000009457_Cluster7787-001 | WRKY | TPA: WRKY DNA-bindin | 1473 | 0 | 0 | 0 |
| SALT00000003109_Cluster7787-003 | WRKY | TPA: WRKY DNA-bindin | 2283 | 0 | 0 | 0 |

| | | | | | | |
|---|---|---|---|---|---|---|
| SALT00000078414_Cluster8016-001 | WRKY | probable WRKY transcript | 3479 | 0 | 0 | 0 |
| SALT00000008473_Cluster9823-001 | WRKY | WRKY transcription partia | 1529 | 0 | 0 | 0 |
| SALT00000049001_Cluster9823-002 | WRKY | WRKY DNA-binding dom | 1583 | 0 | 0 | 0 |
| SALT00000053884_Cluster9823-003 | WRKY | WRKY DNA-binding dom | 1574 | 0 | 0 | 0 |
| SALT00000046023_Cluster9823-005 | WRKY | WRKY transcription partia | 1749 | 0 | 0 | 0 |
| SALT00000076023_Cluster12629-001 | WRKY | probable WRKY transcript | 2664 | 0 | 0 | 0 |
| SALT00000078735_Cluster12629-002 | WRKY | probable WRKY transcript | 2808 | 0 | 0 | 0 |
| SALT00000026660_Cluster12723-001 | WRKY | probable WRKY transcript | 2798 | 0 | 0 | 0 |
| SALT00000002120_Cluster12949-001 | WRKY | Os04g0287400 [Oryza sati | 1197 | 0 | 0 | 0 |
| SALT00000077096_Cluster12949-002 | WRKY | AF140553_1DNA-binding | 2778 | 0 | 0 | 0 |
| SALT00000011889_Cluster12949-004 | WRKY | Os04g0287400 [Oryza sati | 1555 | 0 | up | up |
| SALT00000081184_Cluster14243-001 | WRKY | probable WRKY transcript | 2676 | 0 | 0 | 0 |
| SALT00000063127_Cluster15489-001 | WRKY | probable WRKY transcript | 2590 | 0 | 0 | 0 |
| SALT00000001787_Cluster15959-001 | WRKY | probable WRKY transcript | 2137 | 0 | 0 | 0 |
| SALT00000011546_Cluster15959-002 | WRKY | probable WRKY transcript | 2087 | 0 | 0 | 0 |
| SALT00000063687_Cluster15959-003 | WRKY | probable WRKY transcript | 2555 | 0 | 0 | 0 |
| SALT00000027194_Cluster16064-001 | WRKY | WRKY transcription partia | 2500 | 0 | 0 | 0 |
| SALT00000031476_Cluster16064-002 | WRKY | WRKY transcription partia | 2403 | 0 | 0 | 0 |
| SALT00000067860_Cluster16064-003 | WRKY | WRKY transcription partia | 2518 | 0 | 0 | 0 |
| SALT00000076751_Cluster16064-004 | WRKY | WRKY transcription partia | 2547 | 0 | 0 | 0 |
| SALT00000075731_Cluster16064-005 | WRKY | WRKY transcription partia | 2651 | 0 | 0 | 0 |
| SALT00000039793_Cluster16708-002 | WRKY | uncharacterized protein LC | 1888 | 0 | 0 | 0 |
| SALT00000002798_Cluster16969-001 | WRKY | SPF1-like DNA-binding [Z | 1985 | 0 | 0 | up |
| SALT00000019694_Cluster16969-002 | WRKY | SPF1-like DNA-binding [Z | 1885 | 0 | 0 | 0 |
| SALT00000022826_Cluster16969-003 | WRKY | WRKY DNA-binding dom | 2473 | 0 | 0 | 0 |
| SALT00000049646_Cluster16969-004 | WRKY | WRKY DNA-binding dom | 1632 | 0 | 0 | 0 |
| SALT00000011806_Cluster16969-006 | WRKY | SPF1-like DNA-binding [Z | 1976 | 0 | 0 | 0 |
| SALT00000001076_Cluster18054-001 | WRKY | probable WRKY transcript | 1527 | down | down | down |
| SALT00000015056_Cluster18054-002 | WRKY | probable WRKY transcript | 1626 | 0 | 0 | 0 |
| SALT00000061480_Cluster18054-003 | WRKY | probable WRKY transcript | 2291 | 0 | 0 | 0 |
| SALT00000012082_Cluster18442-001 | WRKY | WRKY transcription factor | 1912 | 0 | 0 | 0 |
| SALT00000017413_Cluster18442-002 | WRKY | WRKY transcription factor | 2091 | 0 | 0 | 0 |
| SALT00000052846_Cluster18442-003 | WRKY | WRKY transcription factor | 2227 | 0 | 0 | 0 |
| SALT00000057373_Cluster18442-004 | WRKY | WRKY transcription factor | 2184 | 0 | 0 | 0 |
| SALT00000011219_Cluster19222-001 | WRKY | Os12g0597700 [Oryza sati | 1797 | 0 | 0 | 0 |
| SALT00000011672_Cluster19222-002 | WRKY | Os12g0597700 [Oryza sati | 1750 | 0 | 0 | 0 |
| SALT00000013301_Cluster19222-003 | WRKY | Os12g0597700 [Oryza sati | 1703 | 0 | 0 | 0 |
| SALT00000015980_Cluster19222-004 | WRKY | Os12g0597700 [Oryza sati | 1672 | 0 | 0 | 0 |
| SALT00000016460_Cluster19222-005 | WRKY | WRKY DNA-binding dom | 1693 | 0 | 0 | 0 |
| SALT00000016745_Cluster19222-006 | WRKY | Os12g0597700 [Oryza sati | 2147 | 0 | 0 | 0 |
| SALT00000038672_Cluster19222-007 | WRKY | WRKY DNA-binding dom | 1770 | 0 | 0 | 0 |
| SALT00000039208_Cluster19222-008 | WRKY | Os12g0597700 [Oryza sati | 1647 | 0 | 0 | 0 |
| SALT00000042217_Cluster19222-009 | WRKY | Os12g0597700 [Oryza sati | 1772 | 0 | 0 | 0 |
| SALT00000044906_Cluster19222-010 | WRKY | Os12g0597700 [Oryza sati | 1777 | 0 | 0 | 0 |
| SALT00000007309_Cluster19737-001 | WRKY | WRKY transcription factor | 2099 | 0 | 0 | 0 |
| SALT00000008863_Cluster19737-002 | WRKY | WRKY transcription factor | 2092 | 0 | 0 | 0 |
| SALT00000015245_Cluster19737-003 | WRKY | WRKY transcription factor | 2045 | 0 | 0 | 0 |
| SALT00000004455_Cluster20184-001 | WRKY | Os01g0826400 [Oryza sati | 2064 | 0 | 0 | up |
| SALT00000058151_Cluster20184-002 | WRKY | Os01g0826400 [Oryza sati | 2025 | 0 | 0 | 0 |
| SALT00000035863_Cluster20184-003 | WRKY | WRKY transcription partia | 1886 | 0 | 0 | 0 |
| SALT00000035722_Cluster20779-001 | WRKY | Os02g0698800 [Oryza sati | 1816 | 0 | 0 | 0 |
| SALT00000039533_Cluster20779-002 | WRKY | Os02g0698800 [Oryza sati | 2021 | 0 | 0 | 0 |
| SALT00000037382_Cluster20803-001 | WRKY | WRKY70 - superfamily of | 2019 | 0 | 0 | 0 |
| SALT00000002512_Cluster21152-001 | WRKY | probable WRKY transcript | 1843 | 0 | 0 | 0 |
| SALT00000006074_Cluster21152-002 | WRKY | probable WRKY transcript | 1994 | 0 | 0 | 0 |
| SALT00000007962_Cluster21152-003 | WRKY | probable WRKY transcript | 1577 | 0 | 0 | 0 |
| SALT00000013279_Cluster21152-004 | WRKY | probable WRKY transcript | 1651 | 0 | 0 | 0 |
| SALT00000038210_Cluster21152-005 | WRKY | probable WRKY transcript | 1793 | 0 | 0 | 0 |

| | | | | | | |
|---|---|---|---|---|---|---|
| SALT00000045500_Cluster21152-006 | WRKY | probable WRKY transcript | 1839 | 0 | 0 | 0 |
| SALT00000053095_Cluster21152-007 | WRKY | probable WRKY transcript | 1571 | 0 | 0 | 0 |
| SALT00000039230_Cluster21152-008 | WRKY | probable WRKY transcript | 1546 | 0 | 0 | 0 |
| SALT00000055149_Cluster21382-001 | WRKY | Os04g0287400 [Oryza sati | 1979 | 0 | 0 | up |
| SALT00000017153_Cluster21425-001 | WRKY | WRKY53 - superfamily of | 1627 | 0 | 0 | up |
| SALT00000019784_Cluster21425-002 | WRKY | WRKY53 - superfamily of | 1864 | 0 | 0 | 0 |
| SALT00000041465_Cluster21425-003 | WRKY | WRKY53 - superfamily of | 1975 | 0 | 0 | up |
| SALT00000006052_Cluster22470-001 | WRKY | Os01g0185900 [Oryza sati | 1760 | 0 | 0 | 0 |
| SALT00000011956_Cluster22470-002 | WRKY | Os01g0185900 [Oryza sati | 1909 | 0 | 0 | 0 |
| SALT00000016191_Cluster22470-003 | WRKY | Os01g0185900 [Oryza sati | 1709 | 0 | 0 | up |
| SALT00000046800_Cluster22470-004 | WRKY | Os01g0185900 [Oryza sati | 1522 | 0 | 0 | 0 |
| SALT00000040365_Cluster22615-001 | WRKY | TPA: WRKY DNA-bindin | 1663 | 0 | 0 | 0 |
| SALT00000051737_Cluster22615-002 | WRKY | TPA: WRKY DNA-bindin | 1885 | 0 | 0 | 0 |
| SALT00000055739_Cluster22615-003 | WRKY | TPA: WRKY DNA-bindin | 1902 | 0 | 0 | 0 |
| SALT00000036123_Cluster22615-004 | WRKY | TPA: WRKY DNA-bindin | 1774 | 0 | 0 | 0 |
| SALT00000000337_Cluster22676-001 | WRKY | WRKY53 - superfamily of | 1759 | 0 | 0 | down |
| SALT00000017567_Cluster22676-002 | WRKY | WRKY53 - superfamily of | 1898 | 0 | 0 | up |
| SALT00000019536_Cluster22676-003 | WRKY | WRKY53 - superfamily of | 1693 | 0 | 0 | up |
| SALT00000024588_Cluster22676-004 | WRKY | WRKY53 - superfamily of | 1604 | 0 | 0 | 0 |
| SALT00000036261_Cluster22676-005 | WRKY | WRKY53 - superfamily of | 1772 | 0 | 0 | 0 |
| SALT00000048412_Cluster22676-006 | WRKY | WRKY53 - superfamily of | 1590 | 0 | 0 | 0 |
| SALT00000050316_Cluster22676-007 | WRKY | WRKY53 - superfamily of | 1850 | 0 | 0 | up |
| SALT00000054490_Cluster22676-008 | WRKY | WRKY53 - superfamily of | 1761 | 0 | 0 | 0 |
| SALT00000036377_Cluster22936-001 | WRKY | [Hordeum vulgare vulgare | 1881 | 0 | 0 | up |
| SALT00000045972_Cluster22936-002 | WRKY | [Hordeum vulgare vulgare | 1745 | 0 | 0 | 0 |
| SALT00000037288_Cluster22936-003 | WRKY | [Hordeum vulgare vulgare | 1696 | 0 | 0 | up |
| SALT00000000740_Cluster23206-001 | WRKY | WRKY53 - superfamily of | 1730 | 0 | 0 | 0 |
| SALT00000006770_Cluster23206-002 | WRKY | WRKY53 - superfamily of | 1798 | 0 | 0 | 0 |
| SALT00000006986_Cluster23206-003 | WRKY | WRKY53 - superfamily of | 1646 | 0 | 0 | 0 |
| SALT00000013131_Cluster23206-004 | WRKY | WRKY53 - superfamily of | 1864 | 0 | 0 | 0 |
| SALT00000058477_Cluster23206-005 | WRKY | WRKY53 - superfamily of | 1795 | 0 | 0 | 0 |
| SALT00000059144_Cluster23206-006 | WRKY | WRKY DNA-binding dom | 1978 | 0 | 0 | 0 |
| SALT00000003192_Cluster23390-001 | WRKY | WRKY53 - superfamily of | 1787 | 0 | 0 | 0 |
| SALT00000003588_Cluster23390-002 | WRKY | WRKY53 - superfamily of | 1848 | 0 | down | 0 |
| SALT00000006704_Cluster23390-003 | WRKY | WRKY53 - superfamily of | 1749 | 0 | 0 | up |
| SALT00000009228_Cluster23390-004 | WRKY | WRKY53 - superfamily of | 1853 | 0 | 0 | 0 |
| SALT00000011152_Cluster23390-005 | WRKY | WRKY53 - superfamily of | 1689 | 0 | 0 | 0 |
| SALT00000037508_Cluster23390-006 | WRKY | WRKY53 - superfamily of | 1676 | 0 | 0 | 0 |
| SALT00000053018_Cluster23390-007 | WRKY | WRKY53 - superfamily of | 1676 | 0 | 0 | up |
| SALT00000000940_Cluster23459-001 | WRKY | WRKY DNA-binding dom | 1719 | 0 | 0 | 0 |
| SALT00000046585_Cluster23459-002 | WRKY | WRKY DNA-binding dom | 1850 | 0 | 0 | 0 |
| SALT00000008393_Cluster23908-001 | WRKY | WRKY DNA-binding dom | 1776 | 0 | 0 | 0 |
| SALT00000045300_Cluster23908-002 | WRKY | WRKY DNA-binding dom | 1588 | 0 | 0 | 0 |
| SALT00000051909_Cluster23908-003 | WRKY | WRKY DNA-binding dom | 1768 | 0 | 0 | 0 |
| SALT00000055800_Cluster23908-004 | WRKY | WRKY DNA-binding dom | 1535 | 0 | 0 | 0 |
| SALT00000061059_Cluster23908-005 | WRKY | WRKY DNA-binding dom | 1682 | 0 | 0 | 0 |
| SALT00000061760_Cluster23908-006 | WRKY | WRKY DNA-binding dom | 1824 | 0 | 0 | 0 |
| SALT00000010779_Cluster23960-001 | WRKY | WRKY transcription factor | 1632 | 0 | 0 | 0 |
| SALT00000011051_Cluster23960-002 | WRKY | WRKY transcription factor | 1656 | 0 | 0 | 0 |
| SALT00000013792_Cluster23960-003 | WRKY | WRKY transcription factor | 1691 | 0 | 0 | 0 |
| SALT00000041514_Cluster23960-004 | WRKY | WRKY transcription factor | 1633 | 0 | 0 | 0 |
| SALT00000043303_Cluster23960-005 | WRKY | WRKY transcription factor | 1820 | 0 | 0 | 0 |
| SALT00000056117_Cluster23960-006 | WRKY | WRKY transcription factor | 1765 | 0 | 0 | 0 |
| SALT00000008585_Cluster24077-001 | WRKY | Os01g0185900 [Oryza sati | 1814 | 0 | 0 | 0 |
| SALT00000004600_Cluster24424-001 | WRKY | probable WRKY transcript | 1794 | 0 | 0 | up |
| SALT00000001348_Cluster24882-001 | WRKY | WRKY transcription factor | 1747 | 0 | 0 | 0 |
| SALT00000007844_Cluster24882-002 | WRKY | WRKY transcription factor | 1713 | 0 | 0 | 0 |
| SALT00000040084_Cluster24882-003 | WRKY | WRKY transcription factor | 1769 | 0 | 0 | 0 |

| | | | | | | |
|---|---|---|---|---|---|---|
| SALT00000044879_Cluster24882-004 | WRKY | WRKY transcription factoi | 1622 | 0 | 0 | 0 |
| SALT00000051116_Cluster25267-001 | WRKY | WRKY family transcriptio | 1749 | 0 | 0 | 0 |
| SALT00000001002_Cluster25493-001 | WRKY | WRKY53 - superfamily of | 1688 | 0 | 0 | 0 |
| SALT00000013463_Cluster25493-002 | WRKY | WRKY53 - superfamily of | 1737 | 0 | 0 | 0 |
| SALT00000044979_Cluster25493-003 | WRKY | WRKY53 - superfamily of | 1660 | 0 | 0 | 0 |
| SALT00000050487_Cluster25493-004 | WRKY | WRKY53 - superfamily of | 1121 | 0 | 0 | up |
| SALT00000007283_Cluster25599-001 | WRKY | WRKY transcription factoi | 1732 | 0 | 0 | 0 |
| SALT00000012850_Cluster25599-002 | WRKY | WRKY transcription factoi | 1660 | 0 | 0 | 0 |
| SALT00000058209_Cluster25599-003 | WRKY | WRKY transcription factoi | 1416 | 0 | 0 | 0 |
| SALT00000040673_Cluster25834-001 | WRKY | WAKY 1 | 1399 | 0 | 0 | 0 |
| SALT00000043407_Cluster25834-002 | WRKY | WAKY 1 | 1721 | 0 | 0 | up |
| SALT00000036923_Cluster25893-001 | WRKY | WRKY transcription factoi | 1717 | 0 | 0 | 0 |
| SALT00000040834_Cluster25893-002 | WRKY | WRKY transcription factoi | 1625 | 0 | 0 | 0 |
| SALT00000067144_Cluster25893-003 | WRKY | WRKY transcription factoi | 1686 | 0 | 0 | 0 |
| SALT00000001459_Cluster26186-001 | WRKY | WRKY DNA-binding dom | 1699 | 0 | 0 | 0 |
| SALT00000046968_Cluster26186-002 | WRKY | WRKY DNA-binding dom | 1679 | 0 | 0 | 0 |
| SALT00000037811_Cluster26186-003 | WRKY | WRKY DNA-binding dom | 1669 | 0 | 0 | 0 |
| SALT00000009662_Cluster26204-001 | WRKY | WRKY DNA-binding dom | 1484 | 0 | 0 | 0 |
| SALT00000059841_Cluster26204-002 | WRKY | WRKY DNA-binding dom | 1699 | 0 | 0 | 0 |
| SALT00000050702_Cluster26814-001 | WRKY | WRKY DNA-binding dom | 1669 | 0 | 0 | 0 |
| SALT00000011815_Cluster27067-001 | WRKY | WRKY transcription partia | 1656 | 0 | 0 | 0 |
| SALT00000017147_Cluster27067-002 | WRKY | WRKY transcription partia | 1596 | 0 | 0 | 0 |
| SALT00000003248_Cluster27423-001 | WRKY | WRKY-type transcription | 1639 | down | 0 | down |
| SALT00000009509_Cluster27423-002 | WRKY | WRKY-type transcription | 1561 | 0 | 0 | 0 |
| SALT00000057323_Cluster27423-003 | WRKY | WRKY-type transcription | 1482 | 0 | 0 | 0 |
| SALT00000059856_Cluster27423-004 | WRKY | WRKY-type transcription | 1588 | 0 | down | 0 |
| SALT00000016512_Cluster27423-005 | WRKY | WRKY-type transcription | 1222 | down | down | down |
| SALT00000010030_Cluster27423-006 | WRKY | WRKY-type transcription | 1565 | 0 | 0 | 0 |
| SALT00000035872_Cluster27700-001 | WRKY | WRKY transcription partia | 1549 | 0 | 0 | up |
| SALT00000042784_Cluster27700-002 | WRKY | WRKY11 - superfamily of | 1626 | 0 | 0 | 0 |
| SALT00000000904_Cluster28800-001 | WRKY | WRKY family transcriptio | 1568 | 0 | 0 | 0 |
| SALT00000038709_Cluster28800-002 | WRKY | WRKY family transcriptio | 1503 | 0 | 0 | 0 |
| SALT00000049053_Cluster28800-003 | WRKY | WRKY family transcriptio | 1239 | 0 | 0 | 0 |
| SALT00000065545_Cluster28963-001 | WRKY | probable WRKY transcript | 1561 | 0 | 0 | 0 |
| SALT00000015440_Cluster29288-001 | WRKY | WRKY transcription partia | 1544 | 0 | 0 | 0 |
| SALT00000036655_Cluster29288-002 | WRKY | WRKY transcription partia | 1383 | 0 | 0 | 0 |
| SALT00000041477_Cluster29288-003 | WRKY | WRKY transcription partia | 1502 | 0 | 0 | 0 |
| SALT00000054691_Cluster29567-001 | WRKY | WRKY7 - superfamily of | 1529 | 0 | 0 | 0 |
| SALT00000004910_Cluster29681-001 | WRKY | probable WRKY transcript | 1521 | 0 | 0 | 0 |
| SALT00000039045_Cluster29801-001 | WRKY | WRKY transcription partia | 1514 | 0 | 0 | 0 |
| SALT00000018114_Cluster29845-001 | WRKY | probable WRKY transcript | 1350 | 0 | 0 | 0 |
| SALT00000049217_Cluster29845-002 | WRKY | probable WRKY transcript | 1512 | 0 | 0 | 0 |
| SALT00000055313_Cluster29845-003 | WRKY | [Hordeum vulgare vulgare | 1541 | 0 | 0 | 0 |
| SALT00000038055_Cluster29845-004 | WRKY | [Hordeum vulgare vulgare | 1625 | 0 | 0 | 0 |
| SALT00000010982_Cluster30067-001 | WRKY | WRKY transcription factoi | 1497 | 0 | 0 | 0 |
| SALT00000018287_Cluster30085-001 | WRKY | WRKY transcription factoi | 1368 | 0 | 0 | up |
| SALT00000028798_Cluster30085-002 | WRKY | WRKY transcription factoi | 1496 | 0 | 0 | 0 |
| SALT00000031617_Cluster30085-003 | WRKY | WRKY transcription factoi | 1495 | 0 | 0 | 0 |
| SALT00000032936_Cluster30085-004 | WRKY | WRKY transcription factoi | 1467 | 0 | 0 | 0 |
| SALT00000035532_Cluster30085-005 | WRKY | WRKY transcription factoi | 1436 | 0 | 0 | up |
| SALT00000039128_Cluster30085-006 | WRKY | WRKY transcription factoi | 1451 | 0 | 0 | 0 |
| SALT00000044470_Cluster30085-007 | WRKY | WRKY transcription factoi | 1488 | 0 | 0 | 0 |
| SALT00000052055_Cluster30085-008 | WRKY | WRKY transcription factoi | 1425 | 0 | 0 | up |
| SALT00000059428_Cluster30085-009 | WRKY | WRKY transcription factoi | 1224 | 0 | 0 | up |
| SALT00000013847_Cluster30095-001 | WRKY | WRKY transcription factoi | 1495 | 0 | 0 | up |
| SALT00000076629_Cluster30095-002 | WRKY | WRKY transcription factoi | 1459 | 0 | 0 | 0 |
| SALT00000006112_Cluster30095-003 | WRKY | WRKY transcription factoi | 1503 | 0 | 0 | up |
| SALT00000066591_Cluster30547-001 | WRKY | WRKY transcription partia | 1465 | 0 | 0 | 0 |

| | | | | | | |
|---|---|---|---|---|---|---|
| SALT00000052918_Cluster31345-001 | WRKY | WRKY DNA-binding isof | 1367 | 0 | 0 | 0 |
| SALT00000048160_Cluster31574-001 | WRKY | WRKY transcription facto | 1313 | 0 | 0 | 0 |
| SALT00000015983_Cluster31655-001 | WRKY | partial [Oryza sativa Japon | 1294 | 0 | 0 | 0 |
| SALT00000045669_Cluster31657-001 | WRKY | WRKY74 - superfamily of | 1293 | 0 | 0 | 0 |
| SALT00000057712_Cluster31657-002 | WRKY | WRKY74 - superfamily of | 1234 | 0 | 0 | 0 |
| SALT00000018528_Cluster31657-003 | WRKY | WRKY74 - superfamily of | 1466 | 0 | 0 | 0 |
| SALT00000065826_Cluster32523-001 | WRKY | WRKY55 - superfamily of | 853 | 0 | 0 | up |
| SALT00000051407_Cluster26202-001 | YABBY | YABBY 4 [Zea mays] | 1575 | 0 | 0 | 0 |
| SALT00000054952_Cluster26202-002 | YABBY | YABBY 4 [Zea mays] | 1699 | 0 | 0 | 0 |
| SALT00000040254_Cluster22227-001 | ZF-HD | zinc finger homeodomain | 1532 | 0 | 0 | 0 |
| SALT00000054461_Cluster22227-002 | ZF-HD | zinc finger homeodomain | 1925 | 0 | 0 | 0 |
| SALT00000058866_Cluster22227-003 | ZF-HD | zinc finger homeodomain | 1313 | 0 | 0 | 0 |
| SALT00000020190_Cluster22449-001 | ZF-HD | ZF-HD homeobox [Zea ma | 1691 | 0 | 0 | 0 |
| SALT00000036710_Cluster22449-002 | ZF-HD | ZF-HD homeobox [Zea ma | 1910 | 0 | 0 | 0 |
| SALT00000054684_Cluster22449-003 | ZF-HD | ZF-HD homeobox [Zea ma | 1499 | 0 | 0 | 0 |
| SALT00000038784_Cluster23115-001 | ZF-HD | zinc finger homeodomain | 1524 | 0 | 0 | 0 |
| SALT00000054370_Cluster23115-002 | ZF-HD | zinc finger homeodomain | 1533 | 0 | 0 | 0 |
| SALT00000055819_Cluster23115-003 | ZF-HD | zinc finger homeodomain | 1870 | 0 | down | 0 |
| SALT00000008753_Cluster25937-001 | ZF-HD | TPA: ZF-HD homeobox [Z | 1494 | 0 | 0 | 0 |
| SALT00000019205_Cluster25937-002 | ZF-HD | TPA: ZF-HD homeobox [Z | 1452 | 0 | 0 | 0 |
| SALT00000042393_Cluster25937-003 | ZF-HD | TPA: ZF-HD homeobox [Z | 1555 | 0 | 0 | 0 |
| SALT00000042682_Cluster25937-004 | ZF-HD | ZF-HD homeobox [Zea ma | 1715 | 0 | 0 | 0 |
| SALT00000005360_Cluster25937-005 | ZF-HD | TPA: ZF-HD homeobox [Z | 1611 | 0 | 0 | 0 |
