## Supplemental Table 5 for "The full-length transcriptome of *Spartina alterniflora* reveals the complexity of high salt tolerance in monocotyledonous halophyte"

### Table S5. Summary of transcription factor families in *Spartina alterniflora*

| TF family | | DNA-binding motif (TF) | Transcripts | Unigenes | DE transcripts | DE unigenes |
|---|---|---|---|---|---|---|
| AP2/ERF | AP2 | AP2 (>=2) (PF00847) | 44 | 20 | 2 | 1 |
| | ERF | AP2 (1) (PF00847) | 308 | 60 | 38 | 22 |
| | RAV | AP2 (PF00847)、B3 (PF( | 24 | 6 | 4 | 3 |
| B3 superfamily | ARF | B3 (PF02362) | 283 | 47 | 5 | 4 |
| | B3 | B3 (PF02362) | 65 | 25 | 3 | 3 |
| BBR-BPC | | GAGA_bind (PF06217) | 14 | 2 | | |
| BES1 | | DUF822 (PF05687) | 18 | 10 | | |
| bHLH | | HLH (PF00010) | 274 | 97 | 13 | 12 |
| bZIP | | bZIP_1 (PF00170) | 297 | 91 | 19 | 12 |
| C2C2 | CO-like | Zf-B_box(PF00643） | 51 | 13 | 9 | 5 |
| | Dof | Zf-Dof (PF02701) | 71 | 26 | 2 | 1 |
| | GATA | GATA-zf (PF00320) | 122 | 29 | 8 | 6 |
| | LSD | Zf-LSD1 (PF06943) | 6 | 4 | 1 | 1 |
| | YABBY | YABBY (PF04690) | 2 | 1 | | |
| C2H2 | | zf-C2H2 (PF00096) | 251 | 65 | 10 | 8 |
| C3H | | Zf-CCCH (PF00642) | 285 | 60 | 9 | 8 |
| CAMTA | | CG1 (PF03859) | 104 | 15 | 2 | 2 |
| CPP | | TCR (PF03638) | 42 | 12 | 2 | 2 |
| DBB | | zf-B_box (>=2) (PF0064[illegible] | 22 | 6 | 1 | 1 |
| E2F/DP | | E2F_TDP (PF02319) | 18 | 9 | | |
| EIL | | EIN3 (PF04873) | 58 | 7 | 6 | 3 |
| FAR1 | | FAR1 (PF03101) | 113 | 35 | 1 | 1 |
| GARP | ARR-B | G2-like (self-build) | 41 | 9 | | |
| | G2-like | G2-like (self-build) | 126 | 50 | 7 | 5 |

| | | | | | | |
|---|---|---|---|---|---|---|
| GeBP | | DUF573 (PF04504) | 29 | 15 | 1 | 1 |
| GRAS | | GRAS (PF03514) | 282 | 42 | 17 | 10 |
| GRF | | WRC (PF08879) | 14 | 6 | 1 | 1 |
| HB | HD-ZIP | Homeobox (PF00046) | 203 | 48 | 19 | 14 |
| | TALE | Homeobox (PF00046) | 244 | 40 | 10 | 6 |
| | WOX | homeobox (PF00046) | 6 | 3 | 1 | 1 |
| | HB-PHD | homeobox (PF00046) | 27 | 8 | 1 | 1 |
| | HB-other | homeobox (PF00046) | 60 | 21 | | |
| HRT-like | | HRT-like (self-build) | / | / | | |
| HSF | | HSF_dna_bind (PF00447 | 71 | 21 | 9 | 6 |
| LBD (AS2/LOB) | | DUF260 (PF03195) | 9 | 6 | | |
| LFY | | FLO_LFY (PF01698) | / | / | | |
| MADS | M_type | SRF-TF (PF00319) | 2 | 2 | | |
| | MIKC | SRF-TF (PF00319) | 20 | 12 | | |
| MYB superfami | MYB | Myb_dna_bind (>=2) (PF | 163 | 56 | 12 | 10 |
| | MYB_rela | Myb_dna_bind (1) (PF00 | 219 | 77 | 12 | 12 |
| NAC | | NAM (PF02365) | 273 | 82 | 42 | 21 |
| NF-X1 | | Zf-NF-X1 (PF01422) | 10 | 3 | 1 | 1 |
| NF-Y | NF-YA | CBFB_NFYA (PF02045) | 43 | 19 | | 1 |
| | NF-YB | NF-YB (self-build) | 10 | 5 | | |
| | NF-YC | NF-YC (self-build) | 9 | 7 | | |
| Nin-like | | RWP-RK (PF02042) | 54 | 12 | | |
| NZZ/SPL | | NOZZLE (PF08744) | / | / | / | / |
| S1Fa-like | | S1FA (PF04689) | / | / | / | / |
| SAP | | SAP (self-build) | / | / | / | / |
| SBP | | SBP (PF03110) | 132 | 24 | 2 | 2 |
| SRS | | DUF702 (PF05142) | 19 | 8 | 1 | 1 |

| | | | | | |
|---|---|---|---|---|---|
| STAT | STAT (self-build) | 5 | 3 | | |
| TCP | TCP (PF03634) | 59 | 18 | 5 | 4 |
| Trihelix | Trihelix (self-build) | 170 | 35 | 8 | 8 |
| VOZ | VOZ (self-build) | 15 | 2 | | |
| Whirly | Whirly (PF08536) | / | / | / | / |
| WRKY | WRKY (PF03106) | 196 | 65 | 31 | 18 |
| ZF-HD | ZF-HD_dimer (PF04770) | 14 | 4 | 1 | 1 |
| | total | | 1343 | 316 | 219 |
