## Supplemental Table 6 for "The full-length transcriptome of *Spartina alterniflora* reveals the complexity of high salt tolerance in monocotyledonous halophyte"

### Table S6. List of *Spartina*-specific transcripts

Definition of *Spartina* specific transcripts: Full-length *Spartina alterniflora* transcripts without a NCBI hits by blast with default parameters

ncRNA: non-coding RNA =1; coding RNA =0.

| Transcripts name | ncRNA | length(bp) | DE350vs0 | DE500vs0 | DE800vs0 |
|---|---|---|---|---|---|
| SALT00000055497_Cluster122-001 | 0 | 1437 | 0 | 0 | 0 |
| SALT00000074078_Cluster122-013 | 0 | 2535 | 0 | 0 | 0 |
| SALT00000064662_Cluster177-001 | 0 | 9792 | 0 | 0 | 0 |
| SALT00000063356_Cluster293-001 | 1 | 8717 | up | up | up |
| SALT00000082117_Cluster300-016 | 1 | 831 | 0 | 0 | 0 |
| SALT00000045796_Cluster358-003 | 0 | 1720 | 0 | 0 | 0 |
| SALT00000051290_Cluster361-004 | 0 | 2067 | 0 | 0 | 0 |
| SALT00000019183_Cluster381-001 | 0 | 1459 | 0 | 0 | 0 |
| SALT00000015342_Cluster413-001 | 1 | 1637 | 0 | 0 | 0 |
| SALT00000022825_Cluster413-002 | 1 | 2595 | 0 | 0 | 0 |
| SALT00000024336_Cluster413-003 | 1 | 2865 | 0 | 0 | 0 |
| SALT00000029276_Cluster413-004 | 1 | 2745 | 0 | 0 | 0 |
| SALT00000029588_Cluster413-005 | 1 | 4547 | 0 | 0 | 0 |
| SALT00000033371_Cluster413-006 | 1 | 3318 | 0 | 0 | 0 |
| SALT00000034028_Cluster413-007 | 1 | 4577 | 0 | 0 | 0 |
| SALT00000052429_Cluster413-008 | 1 | 1657 | 0 | 0 | 0 |
| SALT00000066381_Cluster413-009 | 1 | 3224 | 0 | 0 | 0 |
| SALT00000069942_Cluster413-010 | 1 | 2889 | 0 | 0 | 0 |
| SALT00000080872_Cluster413-011 | 1 | 2722 | 0 | 0 | 0 |
| SALT00000085486_Cluster413-012 | 1 | 4383 | 0 | 0 | 0 |
| SALT00000085740_Cluster413-013 | 1 | 3617 | 0 | 0 | 0 |
| SALT00000093171_Cluster413-015 | 1 | 4292 | 0 | 0 | 0 |
| SALT00000034435_Cluster413-016 | 0 | 4039 | 0 | 0 | 0 |
| SALT00000090875_Cluster445-002 | 0 | 3924 | 0 | 0 | 0 |
| SALT00000009828_Cluster455-001 | 0 | 2012 | 0 | 0 | 0 |
| SALT00000050216_Cluster455-002 | 0 | 1156 | 0 | 0 | 0 |
| SALT00000056621_Cluster455-003 | 0 | 2114 | 0 | 0 | 0 |
| SALT00000049717_Cluster514-001 | 0 | 1867 | 0 | 0 | 0 |
| SALT00000048923_Cluster519-001 | 0 | 1571 | 0 | 0 | 0 |
| SALT00000073214_Cluster529-007 | 1 | 2568 | 0 | 0 | 0 |
| SALT00000002958_Cluster673-001 | 0 | 1988 | 0 | 0 | 0 |
| SALT00000039816_Cluster673-011 | 0 | 1435 | 0 | 0 | 0 |
| SALT00000047005_Cluster673-012 | 0 | 1926 | 0 | 0 | 0 |
| SALT00000064202_Cluster673-017 | 0 | 3225 | 0 | 0 | 0 |
| SALT00000038596_Cluster718-001 | 0 | 1549 | 0 | 0 | 0 |
| SALT00000022091_Cluster1001-001 | 1 | 2688 | 0 | 0 | 0 |
| SALT00000050675_Cluster1001-002 | 0 | 2062 | 0 | 0 | 0 |
| SALT00000059303_Cluster1001-003 | 0 | 1977 | 0 | 0 | 0 |
| SALT00000063004_Cluster1001-004 | 1 | 2787 | 0 | 0 | 0 |
| SALT00000068871_Cluster1001-005 | 0 | 2037 | 0 | 0 | 0 |
| SALT00000080671_Cluster1001-007 | 1 | 2792 | 0 | 0 | 0 |
| SALT00000092574_Cluster1005-001 | 1 | 6801 | 0 | 0 | 0 |
| SALT00000082407_Cluster1046-001 | 0 | 2676 | 0 | 0 | 0 |
| SALT00000085203_Cluster1047-001 | 0 | 6679 | 0 | 0 | 0 |
| SALT00000087351_Cluster1132-001 | 0 | 6424 | 0 | 0 | 0 |
| SALT00000088134_Cluster1170-001 | 1 | 6307 | 0 | 0 | 0 |
| SALT00000051585_Cluster1233-005 | 0 | 1503 | 0 | 0 | 0 |
| SALT00000055950_Cluster1235-001 | 0 | 2107 | 0 | 0 | 0 |
| SALT00000083533_Cluster1316-001 | 0 | 5939 | 0 | 0 | 0 |

| | | | | | |
|---|---|---|---|---|---|
| SALT00000047886_Cluster1358-001 | 0 | 1795 | 0 | 0 | 0 |
| SALT00000062143_Cluster1395-037 | 1 | 1936 | 0 | 0 | 0 |
| SALT00000067113_Cluster1500-001 | 0 | 3288 | 0 | 0 | 0 |
| SALT00000071786_Cluster1500-002 | 0 | 2811 | 0 | 0 | 0 |
| SALT00000043398_Cluster1516-005 | 0 | 1967 | 0 | 0 | 0 |
| SALT00000046296_Cluster1516-010 | 0 | 1722 | 0 | 0 | 0 |
| SALT00000036984_Cluster1516-014 | 0 | 1779 | 0 | 0 | 0 |
| SALT00000047692_Cluster1516-016 | 1 | 1968 | 0 | 0 | 0 |
| SALT00000000969_Cluster1576-001 | 0 | 2072 | 0 | 0 | 0 |
| SALT00000024330_Cluster1576-002 | 0 | 3112 | 0 | 0 | 0 |
| SALT00000037316_Cluster1576-003 | 0 | 1572 | 0 | 0 | 0 |
| SALT00000055876_Cluster1576-004 | 0 | 1873 | 0 | 0 | 0 |
| SALT00000065489_Cluster1576-005 | 0 | 2079 | 0 | 0 | 0 |
| SALT00000076374_Cluster1576-007 | 0 | 3256 | 0 | 0 | 0 |
| SALT00000077099_Cluster1576-008 | 0 | 2846 | 0 | 0 | 0 |
| SALT00000085535_Cluster1576-009 | 0 | 4643 | 0 | 0 | 0 |
| SALT00000074938_Cluster1598-001 | 0 | 5500 | 0 | 0 | 0 |
| SALT00000092010_Cluster1722-001 | 0 | 5369 | 0 | 0 | 0 |
| SALT00000007916_Cluster1723-001 | 1 | 1968 | 0 | 0 | 0 |
| SALT00000084955_Cluster1732-001 | 1 | 5361 | 0 | 0 | 0 |
| SALT00000001370_Cluster1752-001 | 0 | 1911 | 0 | 0 | 0 |
| SALT00000002214_Cluster1752-002 | 0 | 1976 | 0 | 0 | 0 |
| SALT00000002529_Cluster1752-003 | 0 | 2062 | 0 | 0 | 0 |
| SALT00000003884_Cluster1752-004 | 0 | 1208 | 0 | 0 | 0 |
| SALT00000005784_Cluster1752-006 | 0 | 1993 | 0 | 0 | 0 |
| SALT00000006976_Cluster1752-007 | 0 | 1864 | 0 | 0 | 0 |
| SALT00000008120_Cluster1752-008 | 0 | 2199 | 0 | 0 | 0 |
| SALT00000009166_Cluster1752-009 | 0 | 1299 | 0 | 0 | down |
| SALT00000010171_Cluster1752-010 | 0 | 1086 | 0 | 0 | 0 |
| SALT00000010695_Cluster1752-011 | 0 | 1485 | 0 | 0 | 0 |
| SALT00000011134_Cluster1752-012 | 0 | 1902 | 0 | 0 | 0 |
| SALT00000014240_Cluster1752-013 | 0 | 1865 | 0 | 0 | 0 |
| SALT00000016852_Cluster1752-014 | 0 | 1831 | 0 | 0 | 0 |
| SALT00000018209_Cluster1752-015 | 0 | 1655 | 0 | 0 | 0 |
| SALT00000020779_Cluster1752-017 | 0 | 2666 | 0 | 0 | 0 |
| SALT00000022278_Cluster1752-018 | 0 | 2977 | 0 | 0 | 0 |
| SALT00000024996_Cluster1752-019 | 0 | 2621 | 0 | 0 | 0 |
| SALT00000027009_Cluster1752-020 | 0 | 2454 | 0 | 0 | 0 |
| SALT00000030597_Cluster1752-022 | 0 | 2530 | 0 | 0 | 0 |
| SALT00000031276_Cluster1752-023 | 1 | 2346 | 0 | 0 | 0 |
| SALT00000034939_Cluster1752-025 | 0 | 2672 | 0 | 0 | 0 |
| SALT00000035612_Cluster1752-026 | 0 | 2658 | 0 | 0 | 0 |
| SALT00000035856_Cluster1752-027 | 0 | 1531 | 0 | 0 | 0 |
| SALT00000036786_Cluster1752-028 | 0 | 1944 | 0 | 0 | 0 |
| SALT00000041260_Cluster1752-029 | 1 | 2026 | 0 | 0 | 0 |
| SALT00000045403_Cluster1752-030 | 0 | 1917 | 0 | 0 | 0 |
| SALT00000045879_Cluster1752-031 | 0 | 1992 | 0 | 0 | 0 |
| SALT00000053103_Cluster1752-032 | 0 | 1578 | 0 | 0 | 0 |
| SALT00000053713_Cluster1752-033 | 0 | 1740 | 0 | 0 | 0 |
| SALT00000054817_Cluster1752-034 | 0 | 1633 | 0 | 0 | 0 |
| SALT00000057344_Cluster1752-035 | 0 | 1248 | 0 | 0 | 0 |
| SALT00000062832_Cluster1752-036 | 0 | 2698 | 0 | 0 | 0 |
| SALT00000067171_Cluster1752-037 | 0 | 2349 | 0 | 0 | 0 |
| SALT00000069842_Cluster1752-038 | 0 | 5343 | 0 | 0 | 0 |
| SALT00000070758_Cluster1752-039 | 0 | 2714 | 0 | 0 | 0 |
| SALT00000076894_Cluster1752-040 | 0 | 2666 | 0 | 0 | 0 |
| SALT00000079753_Cluster1752-041 | 0 | 2721 | 0 | 0 | 0 |
| SALT00000064695_Cluster1771-001 | 1 | 3074 | 0 | 0 | 0 |

| | | | | | |
|---|---|---|---|---|---|
| SALT00000021682_Cluster1776-001 | 0 | 2579 | 0 | 0 | 0 |
| SALT00000085707_Cluster1776-002 | 1 | 5325 | 0 | 0 | 0 |
| SALT00000047775_Cluster1855-001 | 1 | 5257 | 0 | up | 0 |
| SALT00000043157_Cluster1857-011 | 0 | 1837 | 0 | 0 | 0 |
| SALT00000062828_Cluster1857-015 | 0 | 2584 | 0 | 0 | 0 |
| SALT00000013535_Cluster1905-001 | 1 | 1793 | 0 | 0 | 0 |
| SALT00000056793_Cluster1905-005 | 1 | 1826 | 0 | 0 | 0 |
| SALT00000037652_Cluster1966-002 | 0 | 1479 | 0 | 0 | 0 |
| SALT00000071180_Cluster2005-003 | 0 | 2871 | 0 | 0 | 0 |
| SALT00000077419_Cluster2007-001 | 0 | 2717 | 0 | 0 | 0 |
| SALT00000088052_Cluster2007-002 | 0 | 5151 | 0 | 0 | 0 |
| SALT00000069748_Cluster2012-001 | 0 | 2894 | 0 | 0 | 0 |
| SALT00000079069_Cluster2012-002 | 1 | 5148 | 0 | 0 | 0 |
| SALT00000084547_Cluster2012-003 | 1 | 3806 | 0 | 0 | 0 |
| SALT00000093092_Cluster2012-004 | 1 | 4475 | 0 | 0 | 0 |
| SALT00000081399_Cluster2025-002 | 1 | 2658 | 0 | 0 | 0 |
| SALT00000006886_Cluster2057-004 | 0 | 1709 | 0 | 0 | 0 |
| SALT00000011483_Cluster2057-009 | 0 | 1581 | 0 | 0 | 0 |
| SALT00000011692_Cluster2057-010 | 0 | 1650 | down | 0 | 0 |
| SALT00000020405_Cluster2057-014 | 0 | 1620 | 0 | 0 | 0 |
| SALT00000034178_Cluster2057-017 | 1 | 4280 | 0 | 0 | 0 |
| SALT00000055730_Cluster2057-025 | 0 | 1551 | 0 | 0 | 0 |
| SALT00000075254_Cluster2057-029 | 0 | 2653 | 0 | 0 | 0 |
| SALT00000079947_Cluster2057-030 | 0 | 2585 | 0 | 0 | 0 |
| SALT00000018523_Cluster2131-001 | 0 | 1539 | 0 | 0 | 0 |
| SALT00000050446_Cluster2131-002 | 0 | 1497 | 0 | 0 | 0 |
| SALT00000071189_Cluster2131-003 | 0 | 3278 | 0 | 0 | 0 |
| SALT00000075046_Cluster2131-004 | 0 | 2563 | 0 | 0 | 0 |
| SALT00000084987_Cluster2131-005 | 0 | 5058 | 0 | 0 | 0 |
| SALT00000073130_Cluster2140-001 | 1 | 3267 | 0 | 0 | 0 |
| SALT00000091647_Cluster2140-003 | 1 | 3838 | 0 | 0 | 0 |
| SALT00000003670_Cluster2206-001 | 1 | 2066 | 0 | 0 | 0 |
| SALT00000015637_Cluster2206-006 | 0 | 2093 | 0 | 0 | 0 |
| SALT00000045452_Cluster2206-031 | 0 | 1736 | 0 | 0 | 0 |
| SALT00000049543_Cluster2206-032 | 0 | 1855 | 0 | 0 | 0 |
| SALT00000062007_Cluster2206-036 | 0 | 2817 | 0 | 0 | down |
| SALT00000071106_Cluster2206-053 | 0 | 2662 | 0 | 0 | 0 |
| SALT00000036779_Cluster2206-054 | 0 | 1649 | 0 | 0 | 0 |
| SALT00000022557_Cluster2206-055 | 0 | 2949 | 0 | 0 | 0 |
| SALT00000050287_Cluster2206-056 | 0 | 1819 | 0 | 0 | 0 |
| SALT00000037596_Cluster2206-058 | 0 | 2080 | 0 | 0 | 0 |
| SALT00000034275_Cluster2271-001 | 1 | 4388 | 0 | 0 | 0 |
| SALT00000067519_Cluster2271-002 | 1 | 2827 | 0 | 0 | 0 |
| SALT00000068504_Cluster2271-003 | 0 | 2956 | 0 | 0 | 0 |
| SALT00000083598_Cluster2271-004 | 1 | 3705 | 0 | 0 | 0 |
| SALT00000088955_Cluster2271-005 | 1 | 4343 | 0 | 0 | 0 |
| SALT00000091009_Cluster2271-006 | 1 | 4584 | 0 | 0 | 0 |
| SALT00000091380_Cluster2271-007 | 1 | 3309 | 0 | 0 | 0 |
| SALT00000091442_Cluster2271-008 | 1 | 4978 | 0 | 0 | 0 |
| SALT00000091708_Cluster2271-009 | 1 | 4677 | 0 | 0 | 0 |
| SALT00000090329_Cluster2271-010 | 1 | 4088 | 0 | 0 | 0 |
| SALT00000075250_Cluster2271-012 | 1 | 2673 | 0 | 0 | 0 |
| SALT00000014809_Cluster2278-001 | 1 | 1643 | 0 | 0 | 0 |
| SALT00000019315_Cluster2278-002 | 0 | 1376 | 0 | 0 | 0 |
| SALT00000089440_Cluster2278-003 | 0 | 4974 | 0 | 0 | 0 |
| SALT00000021130_Cluster2296-001 | 0 | 2996 | 0 | 0 | 0 |
| SALT00000024550_Cluster2296-002 | 0 | 3027 | 0 | 0 | 0 |
| SALT00000039004_Cluster2296-003 | 1 | 2009 | 0 | 0 | 0 |

| | | | | | |
|---|---|---|---|---|---|
| SALT00000065261_Cluster2296-004 | 0 | 2973 | 0 | 0 | 0 |
| SALT00000066310_Cluster2296-005 | 0 | 3055 | 0 | 0 | 0 |
| SALT00000073376_Cluster2296-007 | 0 | 2859 | 0 | 0 | 0 |
| SALT00000039056_Cluster2296-010 | 1 | 1898 | 0 | 0 | 0 |
| SALT00000029096_Cluster2312-001 | 1 | 3714 | 0 | 0 | 0 |
| SALT00000057994_Cluster2312-002 | 0 | 1778 | 0 | 0 | 0 |
| SALT00000077691_Cluster2312-003 | 1 | 2905 | 0 | 0 | 0 |
| SALT00000085561_Cluster2312-004 | 1 | 4957 | 0 | 0 | 0 |
| SALT00000090549_Cluster2312-005 | 0 | 3457 | 0 | 0 | 0 |
| SALT00000045178_Cluster2438-001 | 0 | 4875 | 0 | 0 | 0 |
| SALT00000063872_Cluster2451-001 | 0 | 4870 | 0 | 0 | down |
| SALT00000004126_Cluster2476-001 | 0 | 1685 | 0 | 0 | 0 |
| SALT00000008518_Cluster2476-002 | 0 | 1765 | 0 | 0 | 0 |
| SALT00000008582_Cluster2476-003 | 0 | 1789 | 0 | 0 | 0 |
| SALT00000061725_Cluster2476-008 | 0 | 1488 | 0 | 0 | 0 |
| SALT00000072837_Cluster2476-015 | 0 | 3031 | 0 | 0 | 0 |
| SALT00000002093_Cluster2484-001 | 0 | 1777 | 0 | 0 | 0 |
| SALT00000010036_Cluster2484-002 | 0 | 1608 | 0 | 0 | 0 |
| SALT00000024755_Cluster2484-003 | 0 | 2594 | 0 | 0 | 0 |
| SALT00000065465_Cluster2484-004 | 0 | 2562 | 0 | 0 | 0 |
| SALT00000015916_Cluster2485-002 | 1 | 2003 | 0 | 0 | 0 |
| SALT00000041114_Cluster2485-003 | 1 | 1899 | 0 | 0 | 0 |
| SALT00000050963_Cluster2588-001 | 0 | 1964 | 0 | 0 | 0 |
| SALT00000090612_Cluster2588-002 | 1 | 4792 | 0 | 0 | 0 |
| SALT00000036719_Cluster2588-003 | 1 | 1468 | 0 | 0 | 0 |
| SALT00000031190_Cluster2631-001 | 0 | 4399 | 0 | 0 | 0 |
| SALT00000031948_Cluster2631-002 | 1 | 3600 | 0 | 0 | 0 |
| SALT00000032786_Cluster2631-003 | 1 | 3803 | 0 | 0 | 0 |
| SALT00000054948_Cluster2631-005 | 0 | 1604 | 0 | 0 | 0 |
| SALT00000061328_Cluster2631-006 | 0 | 1628 | 0 | 0 | 0 |
| SALT00000067059_Cluster2631-008 | 1 | 2783 | 0 | 0 | 0 |
| SALT00000091541_Cluster2631-013 | 1 | 3587 | 0 | 0 | 0 |
| SALT00000093784_Cluster2631-015 | 0 | 4371 | 0 | 0 | 0 |
| SALT00000028214_Cluster2631-016 | 0 | 2737 | 0 | 0 | 0 |
| SALT00000063086_Cluster2631-017 | 0 | 2839 | 0 | 0 | 0 |
| SALT00000074699_Cluster2631-018 | 0 | 2639 | 0 | 0 | 0 |
| SALT00000074059_Cluster2631-019 | 0 | 2653 | 0 | 0 | 0 |
| SALT00000022088_Cluster2631-020 | 0 | 2656 | 0 | 0 | 0 |
| SALT00000058438_Cluster2631-022 | 0 | 1457 | 0 | 0 | 0 |
| SALT00000023676_Cluster2631-023 | 1 | 2615 | 0 | 0 | 0 |
| SALT00000046513_Cluster2631-024 | 0 | 2140 | 0 | 0 | 0 |
| SALT00000023831_Cluster2631-026 | 1 | 2640 | 0 | 0 | 0 |
| SALT00000051596_Cluster2631-027 | 1 | 1373 | 0 | 0 | 0 |
| SALT00000040203_Cluster2652-001 | 1 | 1872 | 0 | 0 | 0 |
| SALT00000070751_Cluster2652-003 | 0 | 3291 | 0 | 0 | 0 |
| SALT00000074672_Cluster2652-004 | 1 | 2903 | 0 | 0 | 0 |
| SALT00000084565_Cluster2652-005 | 0 | 3070 | 0 | 0 | 0 |
| SALT00000059126_Cluster2698-003 | 0 | 1880 | 0 | 0 | 0 |
| SALT00000040422_Cluster2711-001 | 0 | 1552 | 0 | 0 | 0 |
| SALT00000081457_Cluster2744-001 | 1 | 4720 | 0 | 0 | 0 |
| SALT00000087122_Cluster2750-001 | 0 | 3927 | 0 | 0 | 0 |
| SALT00000091377_Cluster2750-003 | 0 | 4717 | 0 | 0 | 0 |
| SALT00000038521_Cluster2750-004 | 0 | 1550 | 0 | 0 | 0 |
| SALT00000048356_Cluster2873-004 | 0 | 2054 | 0 | 0 | 0 |
| SALT00000003842_Cluster2883-001 | 0 | 1814 | 0 | 0 | 0 |
| SALT00000078253_Cluster2883-002 | 0 | 3428 | 0 | 0 | 0 |
| SALT00000085242_Cluster2883-003 | 0 | 3687 | 0 | 0 | 0 |
| SALT00000088150_Cluster2883-004 | 0 | 3435 | 0 | 0 | 0 |

| | | | | | |
|---|---|---|---|---|---|
| SALT00000089095_Cluster2883-005 | 0 | 4657 | 0 | 0 | 0 |
| SALT00000091166_Cluster2924-001 | 1 | 4636 | 0 | 0 | 0 |
| SALT00000069847_Cluster2958-003 | 0 | 2885 | 0 | 0 | 0 |
| SALT00000080217_Cluster3019-001 | 1 | 3633 | 0 | 0 | 0 |
| SALT00000087081_Cluster3019-002 | 0 | 4590 | 0 | 0 | 0 |
| SALT00000088757_Cluster3019-003 | 0 | 3542 | 0 | 0 | 0 |
| SALT00000087452_Cluster3100-001 | 1 | 4555 | 0 | 0 | 0 |
| SALT00000078670_Cluster3124-001 | 1 | 2813 | 0 | 0 | 0 |
| SALT00000085887_Cluster3124-003 | 0 | 4190 | 0 | 0 | 0 |
| SALT00000071441_Cluster3145-001 | 0 | 2673 | 0 | 0 | 0 |
| SALT00000092334_Cluster3145-002 | 0 | 4535 | 0 | 0 | 0 |
| SALT00000029488_Cluster3167-001 | 0 | 4386 | 0 | 0 | 0 |
| SALT00000032724_Cluster3167-002 | 0 | 4526 | 0 | 0 | 0 |
| SALT00000068240_Cluster3167-003 | 0 | 2898 | 0 | 0 | 0 |
| SALT00000080316_Cluster3167-004 | 0 | 4447 | 0 | 0 | 0 |
| SALT00000084725_Cluster3167-005 | 0 | 4440 | 0 | 0 | 0 |
| SALT00000084943_Cluster3167-006 | 0 | 4435 | 0 | 0 | 0 |
| SALT00000089868_Cluster3167-007 | 0 | 3906 | 0 | 0 | 0 |
| SALT00000092118_Cluster3167-008 | 0 | 3582 | 0 | 0 | 0 |
| SALT00000086363_Cluster3167-009 | 1 | 3860 | 0 | 0 | 0 |
| SALT00000008317_Cluster3185-005 | 0 | 1457 | 0 | 0 | 0 |
| SALT00000054673_Cluster3185-007 | 1 | 1993 | 0 | 0 | 0 |
| SALT00000074327_Cluster3185-009 | 1 | 2365 | 0 | 0 | 0 |
| SALT00000046771_Cluster3297-014 | 0 | 1621 | 0 | 0 | 0 |
| SALT00000003079_Cluster3345-001 | 0 | 1606 | 0 | 0 | 0 |
| SALT00000061266_Cluster3412-019 | 0 | 1737 | 0 | 0 | 0 |
| SALT00000075290_Cluster3424-004 | 0 | 2577 | 0 | 0 | 0 |
| SALT00000048449_Cluster3424-005 | 0 | 1744 | 0 | 0 | 0 |
| SALT00000087706_Cluster3436-002 | 1 | 4420 | 0 | 0 | 0 |
| SALT00000089333_Cluster3436-003 | 0 | 4012 | 0 | 0 | 0 |
| SALT00000074236_Cluster3436-004 | 0 | 2942 | 0 | 0 | 0 |
| SALT00000014166_Cluster3436-005 | 0 | 1627 | 0 | 0 | 0 |
| SALT00000066792_Cluster3446-001 | 0 | 1494 | 0 | 0 | 0 |
| SALT00000087605_Cluster3486-001 | 0 | 4398 | 0 | 0 | 0 |
| SALT00000033360_Cluster3492-001 | 1 | 4396 | 0 | 0 | 0 |
| SALT00000076886_Cluster3492-002 | 1 | 4127 | 0 | 0 | 0 |
| SALT00000079498_Cluster3492-003 | 0 | 5794 | 0 | 0 | 0 |
| SALT00000018344_Cluster3536-005 | 0 | 1659 | 0 | 0 | 0 |
| SALT00000063281_Cluster3536-013 | 0 | 3027 | 0 | 0 | 0 |
| SALT00000093058_Cluster3539-001 | 1 | 4379 | 0 | 0 | 0 |
| SALT00000005803_Cluster3547-001 | 0 | 1560 | 0 | 0 | 0 |
| SALT00000078034_Cluster3547-002 | 0 | 2894 | 0 | 0 | 0 |
| SALT00000060433_Cluster3692-001 | 1 | 2223 | 0 | 0 | 0 |
| SALT00000056455_Cluster3704-002 | 1 | 1797 | 0 | 0 | 0 |
| SALT00000091075_Cluster3856-001 | 0 | 4273 | 0 | 0 | 0 |
| SALT00000004531_Cluster3931-001 | 0 | 2303 | 0 | 0 | 0 |
| SALT00000016598_Cluster3931-002 | 1 | 1611 | 0 | 0 | 0 |
| SALT00000025563_Cluster3931-003 | 0 | 2479 | 0 | 0 | 0 |
| SALT00000041397_Cluster3931-004 | 0 | 2385 | 0 | 0 | 0 |
| SALT00000075223_Cluster3931-005 | 0 | 3029 | 0 | 0 | 0 |
| SALT00000092178_Cluster3931-006 | 0 | 4246 | 0 | 0 | 0 |
| SALT00000076533_Cluster3931-007 | 1 | 2913 | 0 | 0 | 0 |
| SALT00000018068_Cluster3931-008 | 0 | 1675 | 0 | 0 | 0 |
| SALT00000005426_Cluster3951-001 | 1 | 1608 | 0 | 0 | 0 |
| SALT00000044671_Cluster3965-001 | 0 | 1669 | 0 | 0 | 0 |
| SALT00000087133_Cluster3988-001 | 0 | 4227 | 0 | 0 | 0 |
| SALT00000018329_Cluster4023-001 | 0 | 1469 | 0 | 0 | 0 |
| SALT00000022329_Cluster4041-001 | 1 | 2120 | 0 | 0 | 0 |

| | | | | | |
|---|---|---|---|---|---|
| SALT00000056278_Cluster4041-002 | 1 | 2100 | 0 | 0 | 0 |
| SALT00000061079_Cluster4071-001 | 0 | 1821 | 0 | 0 | 0 |
| SALT00000092270_Cluster4104-002 | 1 | 3760 | 0 | 0 | 0 |
| SALT00000082368_Cluster4134-001 | 0 | 3272 | 0 | 0 | 0 |
| SALT00000086411_Cluster4211-001 | 0 | 4162 | 0 | 0 | 0 |
| SALT00000066029_Cluster4217-004 | 1 | 2518 | 0 | 0 | 0 |
| SALT00000079444_Cluster4358-001 | 0 | 4126 | 0 | 0 | 0 |
| SALT00000049197_Cluster4406-005 | 0 | 1851 | 0 | 0 | 0 |
| SALT00000085863_Cluster4409-001 | 0 | 4114 | 0 | 0 | 0 |
| SALT00000034305_Cluster4416-001 | 0 | 3875 | 0 | 0 | 0 |
| SALT00000063508_Cluster4416-002 | 0 | 3144 | 0 | 0 | 0 |
| SALT00000063551_Cluster4416-003 | 0 | 2538 | 0 | 0 | 0 |
| SALT00000088832_Cluster4416-004 | 0 | 4112 | 0 | 0 | 0 |
| SALT00000092741_Cluster4450-001 | 0 | 4100 | 0 | 0 | 0 |
| SALT00000084868_Cluster4487-001 | 0 | 3689 | 0 | 0 | 0 |
| SALT00000086905_Cluster4487-002 | 0 | 4093 | 0 | 0 | 0 |
| SALT00000039350_Cluster4521-001 | 1 | 4081 | 0 | 0 | 0 |
| SALT00000085822_Cluster4699-004 | 0 | 3739 | 0 | 0 | 0 |
| SALT00000089658_Cluster4724-001 | 0 | 4035 | 0 | 0 | 0 |
| SALT00000044560_Cluster4753-001 | 0 | 1724 | 0 | 0 | 0 |
| SALT00000080322_Cluster4753-002 | 1 | 2958 | 0 | 0 | 0 |
| SALT00000081349_Cluster4753-003 | 1 | 4027 | 0 | 0 | 0 |
| SALT00000089949_Cluster4753-004 | 1 | 3706 | 0 | 0 | 0 |
| SALT00000065006_Cluster4753-005 | 0 | 3065 | 0 | 0 | 0 |
| SALT00000068451_Cluster4791-001 | 0 | 2506 | 0 | 0 | 0 |
| SALT00000033424_Cluster4832-001 | 1 | 4010 | 0 | 0 | 0 |
| SALT00000064017_Cluster4832-002 | 1 | 2740 | 0 | 0 | 0 |
| SALT00000086661_Cluster4917-001 | 1 | 3993 | 0 | 0 | 0 |
| SALT00000084930_Cluster4928-001 | 0 | 3991 | 0 | 0 | 0 |
| SALT00000085430_Cluster4932-001 | 0 | 3990 | 0 | up | 0 |
| SALT00000018038_Cluster4988-001 | 0 | 1989 | 0 | 0 | 0 |
| SALT00000025015_Cluster4989-001 | 0 | 2933 | 0 | 0 | 0 |
| SALT00000057341_Cluster4989-002 | 0 | 2103 | 0 | 0 | 0 |
| SALT00000064616_Cluster4989-003 | 1 | 2910 | 0 | 0 | 0 |
| SALT00000093510_Cluster4989-004 | 0 | 3978 | 0 | 0 | 0 |
| SALT00000025736_Cluster4989-007 | 0 | 2585 | 0 | 0 | 0 |
| SALT00000064769_Cluster5035-002 | 0 | 2644 | 0 | 0 | 0 |
| SALT00000004753_Cluster5035-004 | 1 | 1354 | 0 | 0 | 0 |
| SALT00000090522_Cluster5082-001 | 0 | 3960 | 0 | 0 | 0 |
| SALT00000063790_Cluster5087-001 | 1 | 3958 | 0 | 0 | 0 |
| SALT00000052386_Cluster5154-004 | 1 | 1782 | 0 | 0 | 0 |
| SALT00000093489_Cluster5246-009 | 0 | 3411 | 0 | 0 | 0 |
| SALT00000040700_Cluster5254-001 | 0 | 3925 | 0 | 0 | up |
| SALT00000010302_Cluster5262-001 | 0 | 1525 | 0 | 0 | 0 |
| SALT00000090824_Cluster5352-001 | 0 | 3906 | 0 | 0 | 0 |
| SALT00000080579_Cluster5385-001 | 0 | 3899 | 0 | 0 | 0 |
| SALT00000087350_Cluster5441-001 | 0 | 3888 | 0 | 0 | 0 |
| SALT00000089179_Cluster5453-001 | 0 | 3885 | 0 | 0 | 0 |
| SALT00000091334_Cluster5559-001 | 0 | 3870 | 0 | 0 | 0 |
| SALT00000002867_Cluster5612-001 | 0 | 1820 | 0 | 0 | 0 |
| SALT00000040549_Cluster5612-002 | 0 | 2225 | 0 | 0 | 0 |
| SALT00000052512_Cluster5612-003 | 0 | 2156 | 0 | 0 | 0 |
| SALT00000067313_Cluster5612-004 | 0 | 1947 | 0 | 0 | 0 |
| SALT00000071163_Cluster5612-005 | 0 | 2521 | 0 | 0 | 0 |
| SALT00000084402_Cluster5612-006 | 0 | 3274 | 0 | 0 | 0 |
| SALT00000090600_Cluster5612-007 | 0 | 2959 | 0 | 0 | 0 |
| SALT00000091153_Cluster5612-008 | 0 | 3860 | 0 | 0 | 0 |
| SALT00000033092_Cluster5612-009 | 0 | 3536 | 0 | 0 | 0 |

| | | | | | |
|---|---|---|---|---|---|
| SALT00000005395_Cluster5642-001 | 0 | 1682 | 0 | 0 | 0 |
| SALT00000042219_Cluster5642-006 | 0 | 1714 | 0 | 0 | 0 |
| SALT00000024742_Cluster5660-001 | 0 | 2457 | 0 | 0 | 0 |
| SALT00000062068_Cluster5660-002 | 0 | 2900 | 0 | 0 | 0 |
| SALT00000073447_Cluster5660-003 | 0 | 3061 | 0 | 0 | 0 |
| SALT00000087161_Cluster5660-004 | 1 | 3852 | 0 | 0 | 0 |
| SALT00000058696_Cluster5660-005 | 0 | 1722 | 0 | 0 | 0 |
| SALT00000037605_Cluster5675-001 | 0 | 1719 | 0 | 0 | 0 |
| SALT00000058737_Cluster5675-002 | 0 | 1572 | 0 | 0 | 0 |
| SALT00000026489_Cluster5736-005 | 0 | 2629 | 0 | 0 | 0 |
| SALT00000039389_Cluster5760-001 | 0 | 1684 | 0 | 0 | 0 |
| SALT00000027279_Cluster5826-001 | 0 | 2937 | 0 | 0 | 0 |
| SALT00000027713_Cluster5826-002 | 0 | 2894 | 0 | 0 | 0 |
| SALT00000033388_Cluster5826-003 | 0 | 3822 | 0 | 0 | 0 |
| SALT00000063541_Cluster5826-004 | 0 | 2937 | 0 | 0 | 0 |
| SALT00000072980_Cluster5868-004 | 0 | 3234 | 0 | 0 | 0 |
| SALT00000078388_Cluster5868-005 | 0 | 3168 | 0 | 0 | 0 |
| SALT00000085253_Cluster5949-001 | 0 | 3800 | 0 | 0 | 0 |
| SALT00000078466_Cluster5957-005 | 1 | 2569 | 0 | 0 | 0 |
| SALT00000061719_Cluster5968-001 | 0 | 2223 | 0 | 0 | 0 |
| SALT00000087786_Cluster6002-001 | 0 | 3792 | 0 | 0 | 0 |
| SALT00000081760_Cluster6021-001 | 0 | 3789 | 0 | 0 | 0 |
| SALT00000022345_Cluster6108-001 | 0 | 2419 | 0 | 0 | 0 |
| SALT00000011737_Cluster6113-001 | 0 | 1461 | 0 | 0 | 0 |
| SALT00000020130_Cluster6159-009 | 0 | 1746 | 0 | 0 | 0 |
| SALT00000090906_Cluster6163-001 | 0 | 3767 | 0 | 0 | 0 |
| SALT00000080455_Cluster6179-001 | 0 | 3764 | up | 0 | 0 |
| SALT00000073184_Cluster6210-001 | 1 | 2787 | 0 | 0 | 0 |
| SALT00000082387_Cluster6210-002 | 1 | 2824 | 0 | 0 | 0 |
| SALT00000092051_Cluster6210-003 | 1 | 3761 | 0 | 0 | 0 |
| SALT00000075681_Cluster6253-005 | 0 | 2863 | 0 | 0 | 0 |
| SALT00000085602_Cluster6253-006 | 0 | 3752 | 0 | 0 | 0 |
| SALT00000074055_Cluster6255-001 | 1 | 2306 | 0 | 0 | 0 |
| SALT00000077198_Cluster6255-002 | 1 | 3010 | 0 | 0 | 0 |
| SALT00000086268_Cluster6255-003 | 1 | 3752 | 0 | 0 | 0 |
| SALT00000085011_Cluster6285-001 | 1 | 3747 | 0 | 0 | 0 |
| SALT00000085371_Cluster6285-002 | 1 | 3651 | 0 | 0 | 0 |
| SALT00000092802_Cluster6285-003 | 1 | 3486 | 0 | 0 | 0 |
| SALT00000013308_Cluster6285-004 | 0 | 1570 | 0 | 0 | 0 |
| SALT00000090981_Cluster6287-001 | 0 | 3747 | 0 | 0 | 0 |
| SALT00000077464_Cluster6296-001 | 1 | 3745 | 0 | 0 | 0 |
| SALT00000049677_Cluster6339-001 | 0 | 2024 | 0 | 0 | 0 |
| SALT00000083298_Cluster6339-002 | 1 | 3738 | 0 | 0 | 0 |
| SALT00000025510_Cluster6352-001 | 1 | 2635 | 0 | 0 | 0 |
| SALT00000045834_Cluster6352-002 | 0 | 1990 | 0 | 0 | 0 |
| SALT00000093014_Cluster6362-001 | 0 | 3735 | 0 | 0 | 0 |
| SALT00000085447_Cluster6365-001 | 0 | 3734 | 0 | 0 | 0 |
| SALT00000064920_Cluster6383-001 | 1 | 3109 | 0 | 0 | 0 |
| SALT00000072797_Cluster6432-001 | 0 | 2754 | 0 | 0 | 0 |
| SALT00000092497_Cluster6432-002 | 0 | 3724 | 0 | 0 | 0 |
| SALT00000043706_Cluster6432-003 | 0 | 1670 | 0 | 0 | 0 |
| SALT00000073561_Cluster6432-004 | 0 | 2345 | 0 | 0 | 0 |
| SALT00000022509_Cluster6465-014 | 0 | 2661 | 0 | 0 | 0 |
| SALT00000062774_Cluster6530-003 | 0 | 2713 | 0 | 0 | 0 |
| SALT00000074537_Cluster6530-004 | 1 | 2494 | 0 | 0 | 0 |
| SALT00000020182_Cluster6543-001 | 0 | 1519 | 0 | 0 | 0 |
| SALT00000070910_Cluster6543-005 | 0 | 3150 | 0 | 0 | 0 |
| SALT00000092239_Cluster6556-001 | 0 | 3705 | 0 | 0 | up |

| | | | | | |
|---|---|---|---|---|---|
| SALT00000089720_Cluster6562-001 | 0 | 3704 | 0 | 0 | 0 |
| SALT00000012765_Cluster6587-001 | 0 | 1881 | 0 | 0 | 0 |
| SALT00000023856_Cluster6587-002 | 0 | 2592 | 0 | 0 | 0 |
| SALT00000088994_Cluster6587-003 | 1 | 3700 | 0 | 0 | 0 |
| SALT00000089108_Cluster6594-001 | 1 | 3699 | 0 | 0 | 0 |
| SALT00000073901_Cluster6613-001 | 0 | 3011 | 0 | 0 | 0 |
| SALT00000088302_Cluster6613-002 | 0 | 3696 | 0 | 0 | 0 |
| SALT00000033511_Cluster6640-001 | 0 | 3691 | 0 | 0 | 0 |
| SALT00000076969_Cluster6640-002 | 0 | 2692 | 0 | 0 | 0 |
| SALT00000092610_Cluster6640-004 | 0 | 3333 | 0 | 0 | 0 |
| SALT00000092676_Cluster6640-005 | 1 | 3360 | 0 | 0 | 0 |
| SALT00000007469_Cluster6681-001 | 1 | 1443 | 0 | 0 | 0 |
| SALT00000057188_Cluster6681-003 | 1 | 1838 | 0 | 0 | 0 |
| SALT00000093240_Cluster6720-001 | 0 | 3679 | 0 | 0 | 0 |
| SALT00000092957_Cluster6751-001 | 1 | 3675 | 0 | 0 | 0 |
| SALT00000056124_Cluster6804-001 | 0 | 3665 | 0 | 0 | 0 |
| SALT00000070835_Cluster6856-001 | 0 | 2716 | 0 | 0 | 0 |
| SALT00000085526_Cluster6856-002 | 0 | 3637 | 0 | 0 | 0 |
| SALT00000089473_Cluster6856-003 | 0 | 3657 | 0 | 0 | down |
| SALT00000041675_Cluster6856-004 | 0 | 1460 | 0 | 0 | 0 |
| SALT00000010756_Cluster6856-005 | 0 | 1580 | 0 | 0 | 0 |
| SALT00000068244_Cluster6914-001 | 0 | 2558 | 0 | 0 | 0 |
| SALT00000082005_Cluster6914-002 | 0 | 2587 | 0 | 0 | 0 |
| SALT00000071526_Cluster6945-001 | 1 | 3050 | 0 | 0 | 0 |
| SALT00000093134_Cluster6945-002 | 1 | 3645 | 0 | 0 | 0 |
| SALT00000054200_Cluster6951-002 | 0 | 1545 | 0 | 0 | 0 |
| SALT00000061817_Cluster7002-001 | 1 | 1457 | 0 | 0 | 0 |
| SALT00000005904_Cluster7065-001 | 0 | 1552 | 0 | 0 | 0 |
| SALT00000013501_Cluster7082-001 | 1 | 1595 | 0 | 0 | 0 |
| SALT00000031975_Cluster7082-002 | 1 | 3624 | 0 | 0 | 0 |
| SALT00000036974_Cluster7082-003 | 0 | 1858 | 0 | 0 | 0 |
| SALT00000087384_Cluster7124-001 | 0 | 3619 | 0 | 0 | 0 |
| SALT00000069977_Cluster7175-001 | 0 | 3611 | 0 | 0 | 0 |
| SALT00000086539_Cluster7178-001 | 0 | 3611 | 0 | 0 | 0 |
| SALT00000093364_Cluster7297-001 | 0 | 3594 | 0 | 0 | 0 |
| SALT00000082654_Cluster7332-001 | 0 | 3588 | 0 | 0 | 0 |
| SALT00000061785_Cluster7369-001 | 0 | 1899 | 0 | 0 | 0 |
| SALT00000085719_Cluster7451-001 | 0 | 3567 | 0 | 0 | 0 |
| SALT00000012825_Cluster7452-001 | 1 | 2078 | 0 | 0 | 0 |
| SALT00000019052_Cluster7452-002 | 0 | 1723 | 0 | 0 | 0 |
| SALT00000024775_Cluster7452-003 | 1 | 2547 | 0 | 0 | 0 |
| SALT00000087218_Cluster7452-004 | 0 | 3567 | 0 | 0 | 0 |
| SALT00000092860_Cluster7457-001 | 0 | 3567 | 0 | 0 | 0 |
| SALT00000092256_Cluster7462-001 | 1 | 3566 | 0 | 0 | 0 |
| SALT00000086452_Cluster7491-001 | 0 | 3562 | 0 | 0 | 0 |
| SALT00000029043_Cluster7505-001 | 0 | 3560 | 0 | 0 | 0 |
| SALT00000084589_Cluster7505-002 | 0 | 2133 | 0 | 0 | 0 |
| SALT00000038144_Cluster7535-001 | 0 | 3556 | 0 | 0 | 0 |
| SALT00000092779_Cluster7618-001 | 0 | 3542 | 0 | 0 | 0 |
| SALT00000089150_Cluster7626-001 | 0 | 3541 | 0 | 0 | 0 |
| SALT00000027037_Cluster7628-001 | 0 | 3338 | 0 | 0 | 0 |
| SALT00000093372_Cluster7628-002 | 0 | 3541 | 0 | 0 | 0 |
| SALT00000057888_Cluster7678-001 | 0 | 1587 | 0 | 0 | 0 |
| SALT00000085759_Cluster7683-001 | 1 | 3530 | 0 | 0 | 0 |
| SALT00000015108_Cluster7691-002 | 0 | 1249 | 0 | 0 | 0 |
| SALT00000088644_Cluster7714-001 | 1 | 3526 | 0 | 0 | 0 |
| SALT00000083163_Cluster7754-001 | 1 | 3521 | 0 | 0 | 0 |
| SALT00000085756_Cluster7795-001 | 0 | 3515 | 0 | 0 | 0 |

| | | | | | |
|---|---|---|---|---|---|
| SALT00000086211_Cluster7839-001 | 0 | 3508 | 0 | 0 | 0 |
| SALT00000087371_Cluster7921-001 | 0 | 3494 | 0 | 0 | 0 |
| SALT00000009987_Cluster7973-001 | 1 | 1728 | 0 | 0 | 0 |
| SALT00000080124_Cluster8000-001 | 0 | 3482 | 0 | 0 | 0 |
| SALT00000033354_Cluster8127-001 | 0 | 3455 | 0 | 0 | 0 |
| SALT00000022420_Cluster8184-001 | 0 | 2810 | 0 | 0 | 0 |
| SALT00000032867_Cluster8184-002 | 0 | 3446 | 0 | 0 | 0 |
| SALT00000069645_Cluster8184-003 | 1 | 2643 | 0 | 0 | 0 |
| SALT00000039145_Cluster8184-004 | 1 | 1646 | 0 | 0 | 0 |
| SALT00000011688_Cluster8200-001 | 1 | 1957 | 0 | 0 | 0 |
| SALT00000049745_Cluster8200-002 | 1 | 1625 | 0 | 0 | 0 |
| SALT00000091027_Cluster8210-001 | 0 | 3442 | 0 | 0 | 0 |
| SALT00000092194_Cluster8225-001 | 0 | 3439 | 0 | 0 | 0 |
| SALT00000032318_Cluster8310-001 | 0 | 3422 | 0 | 0 | 0 |
| SALT00000083726_Cluster8322-001 | 1 | 3420 | 0 | 0 | 0 |
| SALT00000079958_Cluster8336-001 | 0 | 3418 | 0 | 0 | 0 |
| SALT00000031086_Cluster8396-001 | 0 | 3406 | 0 | 0 | 0 |
| SALT00000070366_Cluster8396-002 | 0 | 2585 | 0 | 0 | 0 |
| SALT00000066770_Cluster8453-001 | 0 | 3394 | 0 | 0 | 0 |
| SALT00000080091_Cluster8458-001 | 0 | 3393 | 0 | 0 | 0 |
| SALT00000019232_Cluster8471-001 | 0 | 1676 | 0 | 0 | 0 |
| SALT00000072817_Cluster8471-002 | 0 | 3391 | 0 | 0 | 0 |
| SALT00000057635_Cluster8474-001 | 0 | 3390 | 0 | 0 | 0 |
| SALT00000069958_Cluster8475-001 | 1 | 3390 | 0 | 0 | 0 |
| SALT00000068578_Cluster8504-001 | 0 | 3005 | 0 | 0 | 0 |
| SALT00000089611_Cluster8504-002 | 1 | 3385 | 0 | 0 | 0 |
| SALT00000019624_Cluster8509-001 | 1 | 1774 | 0 | 0 | 0 |
| SALT00000092931_Cluster8509-002 | 1 | 3384 | 0 | 0 | 0 |
| SALT00000009676_Cluster8525-001 | 0 | 1775 | 0 | 0 | 0 |
| SALT00000088731_Cluster8533-001 | 1 | 3378 | 0 | 0 | 0 |
| SALT00000060479_Cluster8535-001 | 0 | 3377 | 0 | 0 | 0 |
| SALT00000067018_Cluster8540-011 | 0 | 2523 | 0 | 0 | 0 |
| SALT00000031133_Cluster8612-001 | 0 | 3360 | 0 | 0 | 0 |
| SALT00000069055_Cluster8628-001 | 0 | 3358 | 0 | 0 | 0 |
| SALT00000078613_Cluster8629-001 | 0 | 3358 | 0 | 0 | 0 |
| SALT00000079274_Cluster8669-001 | 0 | 3351 | 0 | 0 | 0 |
| SALT00000087466_Cluster8711-001 | 0 | 3345 | 0 | 0 | 0 |
| SALT00000085456_Cluster8719-001 | 1 | 3344 | 0 | 0 | 0 |
| SALT00000065848_Cluster8720-001 | 0 | 3343 | 0 | 0 | 0 |
| SALT00000081054_Cluster8750-001 | 0 | 3337 | 0 | 0 | 0 |
| SALT00000068203_Cluster8787-001 | 0 | 3329 | 0 | 0 | 0 |
| SALT00000088906_Cluster8848-001 | 0 | 3317 | 0 | 0 | 0 |
| SALT00000077467_Cluster8895-001 | 0 | 3307 | 0 | 0 | 0 |
| SALT00000064284_Cluster8922-001 | 0 | 3302 | 0 | 0 | 0 |
| SALT00000082155_Cluster8924-001 | 0 | 3302 | 0 | 0 | 0 |
| SALT00000089814_Cluster8926-001 | 0 | 3302 | 0 | 0 | 0 |
| SALT00000078041_Cluster8997-001 | 1 | 3288 | 0 | 0 | down |
| SALT00000053153_Cluster9044-001 | 1 | 1743 | 0 | 0 | 0 |
| SALT00000068846_Cluster9044-002 | 1 | 3278 | 0 | 0 | 0 |
| SALT00000023267_Cluster9081-001 | 0 | 1960 | 0 | 0 | 0 |
| SALT00000085698_Cluster9081-002 | 0 | 3270 | 0 | 0 | 0 |
| SALT00000068378_Cluster9091-014 | 1 | 1550 | 0 | 0 | 0 |
| SALT00000068797_Cluster9109-001 | 0 | 3266 | 0 | 0 | 0 |
| SALT00000090371_Cluster9167-001 | 0 | 3258 | 0 | 0 | 0 |
| SALT00000087082_Cluster9203-001 | 0 | 3251 | 0 | 0 | 0 |
| SALT00000072267_Cluster9212-002 | 1 | 1321 | 0 | 0 | 0 |
| SALT00000032732_Cluster9227-001 | 0 | 3245 | 0 | 0 | 0 |
| SALT00000079823_Cluster9227-002 | 0 | 2864 | 0 | 0 | 0 |

| | | | | | |
|---|---|---|---|---|---|
| SALT00000092195_Cluster9260-001 | 0 | 3240 | 0 | 0 | 0 |
| SALT00000085713_Cluster9286-001 | 0 | 3234 | 0 | 0 | 0 |
| SALT00000086541_Cluster9299-001 | 0 | 3231 | 0 | 0 | 0 |
| SALT00000019255_Cluster9356-001 | 0 | 1600 | 0 | 0 | 0 |
| SALT00000057668_Cluster9373-007 | 1 | 1506 | 0 | 0 | 0 |
| SALT00000070808_Cluster9502-001 | 0 | 3192 | 0 | 0 | 0 |
| SALT00000071850_Cluster9516-001 | 1 | 3189 | 0 | 0 | 0 |
| SALT00000074089_Cluster9516-002 | 1 | 3099 | 0 | 0 | 0 |
| SALT00000089351_Cluster9516-003 | 1 | 3099 | 0 | 0 | 0 |
| SALT00000077254_Cluster9531-002 | 0 | 3186 | 0 | 0 | 0 |
| SALT00000016479_Cluster9531-003 | 1 | 1654 | 0 | 0 | 0 |
| SALT00000060584_Cluster9591-001 | 0 | 1685 | 0 | 0 | 0 |
| SALT00000082420_Cluster9597-001 | 0 | 3174 | 0 | 0 | 0 |
| SALT00000064532_Cluster9601-001 | 0 | 3173 | 0 | 0 | 0 |
| SALT00000041702_Cluster9606-008 | 1 | 1596 | 0 | 0 | 0 |
| SALT00000076367_Cluster9606-009 | 0 | 2432 | 0 | 0 | 0 |
| SALT00000054749_Cluster9606-013 | 0 | 1578 | 0 | 0 | 0 |
| SALT00000047366_Cluster9628-003 | 0 | 1501 | 0 | 0 | 0 |
| SALT00000029409_Cluster9659-001 | 0 | 3164 | 0 | 0 | 0 |
| SALT00000076089_Cluster9659-002 | 0 | 3046 | 0 | 0 | 0 |
| SALT00000022054_Cluster9676-001 | 0 | 2706 | 0 | 0 | 0 |
| SALT00000023736_Cluster9676-002 | 0 | 2845 | 0 | 0 | 0 |
| SALT00000061970_Cluster9676-003 | 0 | 2654 | 0 | 0 | 0 |
| SALT00000069016_Cluster9676-004 | 0 | 3162 | 0 | 0 | 0 |
| SALT00000075051_Cluster9685-001 | 0 | 3161 | 0 | 0 | 0 |
| SALT00000075082_Cluster9686-001 | 0 | 3161 | 0 | 0 | 0 |
| SALT00000079893_Cluster9702-001 | 0 | 3157 | 0 | 0 | 0 |
| SALT00000047439_Cluster9742-001 | 0 | 2067 | 0 | 0 | 0 |
| SALT00000063395_Cluster9742-002 | 0 | 2474 | 0 | 0 | 0 |
| SALT00000077189_Cluster9742-003 | 0 | 2466 | 0 | 0 | 0 |
| SALT00000082492_Cluster9742-004 | 0 | 3149 | 0 | 0 | 0 |
| SALT00000090783_Cluster9742-005 | 0 | 3124 | 0 | 0 | 0 |
| SALT00000053886_Cluster9742-006 | 0 | 1904 | 0 | 0 | 0 |
| SALT00000075909_Cluster9751-001 | 0 | 3148 | 0 | 0 | 0 |
| SALT00000074195_Cluster9772-001 | 1 | 3145 | 0 | 0 | 0 |
| SALT00000018461_Cluster9788-001 | 0 | 1839 | 0 | 0 | 0 |
| SALT00000018658_Cluster9788-002 | 1 | 1643 | 0 | 0 | 0 |
| SALT00000030003_Cluster9788-003 | 0 | 3141 | 0 | 0 | 0 |
| SALT00000043189_Cluster9788-004 | 1 | 1419 | 0 | 0 | 0 |
| SALT00000079202_Cluster9788-006 | 0 | 2686 | 0 | 0 | 0 |
| SALT00000088057_Cluster9816-001 | 0 | 3139 | 0 | 0 | 0 |
| SALT00000043518_Cluster9886-002 | 0 | 1530 | 0 | 0 | 0 |
| SALT00000068337_Cluster9921-001 | 0 | 2890 | 0 | 0 | 0 |
| SALT00000077723_Cluster9921-003 | 0 | 2773 | 0 | 0 | 0 |
| SALT00000082944_Cluster9921-004 | 0 | 2529 | 0 | 0 | 0 |
| SALT00000084161_Cluster9921-005 | 0 | 2898 | 0 | 0 | 0 |
| SALT00000000224_Cluster9931-001 | 1 | 2014 | 0 | 0 | 0 |
| SALT00000004048_Cluster9931-002 | 1 | 2039 | 0 | 0 | 0 |
| SALT00000008315_Cluster9931-003 | 1 | 1538 | 0 | 0 | 0 |
| SALT00000009445_Cluster9931-004 | 1 | 1791 | 0 | 0 | 0 |
| SALT00000010547_Cluster9931-005 | 1 | 2103 | 0 | 0 | 0 |
| SALT00000010934_Cluster9931-006 | 1 | 1901 | 0 | 0 | 0 |
| SALT00000016173_Cluster9931-007 | 1 | 2050 | 0 | 0 | 0 |
| SALT00000019881_Cluster9931-008 | 1 | 1650 | 0 | 0 | 0 |
| SALT00000042740_Cluster9931-009 | 1 | 1843 | 0 | 0 | 0 |
| SALT00000046756_Cluster9931-010 | 0 | 3121 | 0 | 0 | 0 |
| SALT00000050667_Cluster9931-011 | 1 | 1857 | 0 | 0 | 0 |
| SALT00000055232_Cluster9931-012 | 1 | 1637 | 0 | 0 | 0 |

| | | | | | |
|---|---|---|---|---|---|
| SALT00000038852_Cluster9931-013 | 0 | 1539 | 0 | 0 | 0 |
| SALT00000036865_Cluster9931-015 | 0 | 1739 | 0 | 0 | 0 |
| SALT00000049619_Cluster9959-003 | 0 | 1451 | 0 | 0 | 0 |
| SALT00000023284_Cluster9963-001 | 0 | 3114 | 0 | 0 | 0 |
| SALT00000093547_Cluster9970-001 | 1 | 3114 | 0 | 0 | 0 |
| SALT00000002400_Cluster9972-004 | 1 | 1750 | 0 | 0 | 0 |
| SALT00000057201_Cluster9972-021 | 1 | 1343 | 0 | 0 | 0 |
| SALT00000078224_Cluster10070-00 | 1 | 3098 | 0 | 0 | 0 |
| SALT00000048858_Cluster10096-00 | 1 | 1691 | 0 | 0 | 0 |
| SALT00000014450_Cluster10126-00 | 0 | 1359 | 0 | 0 | 0 |
| SALT00000077884_Cluster10126-00 | 1 | 3091 | 0 | 0 | 0 |
| SALT00000069964_Cluster10163-00 | 0 | 3086 | 0 | 0 | 0 |
| SALT00000077133_Cluster10163-00 | 0 | 2978 | 0 | 0 | 0 |
| SALT00000078356_Cluster10163-00 | 0 | 2773 | 0 | 0 | 0 |
| SALT00000049052_Cluster10227-00 | 0 | 1716 | 0 | 0 | 0 |
| SALT00000065487_Cluster10227-00 | 1 | 3077 | 0 | 0 | 0 |
| SALT00000075805_Cluster10227-00 | 0 | 2899 | 0 | 0 | 0 |
| SALT00000072193_Cluster10227-00 | 0 | 2836 | 0 | 0 | 0 |
| SALT00000007509_Cluster10257-00 | 0 | 1705 | 0 | 0 | 0 |
| SALT00000078861_Cluster10282-00 | 1 | 3068 | 0 | 0 | 0 |
| SALT00000075501_Cluster10320-00 | 0 | 3062 | 0 | 0 | 0 |
| SALT00000080544_Cluster10362-00 | 0 | 3055 | 0 | 0 | 0 |
| SALT00000009690_Cluster10403-00 | 0 | 1587 | 0 | 0 | 0 |
| SALT00000071226_Cluster10413-00 | 0 | 2998 | 0 | 0 | 0 |
| SALT00000071468_Cluster10413-00 | 0 | 2852 | 0 | 0 | 0 |
| SALT00000080118_Cluster10413-00 | 0 | 3050 | 0 | 0 | 0 |
| SALT00000016377_Cluster10424-00 | 0 | 2026 | 0 | 0 | 0 |
| SALT00000022083_Cluster10424-00 | 0 | 2891 | 0 | 0 | 0 |
| SALT00000071089_Cluster10424-00 | 0 | 3048 | 0 | 0 | 0 |
| SALT00000062709_Cluster10461-00 | 0 | 3043 | 0 | 0 | 0 |
| SALT00000022560_Cluster10470-00 | 1 | 2668 | 0 | 0 | 0 |
| SALT00000083703_Cluster10470-00 | 0 | 3042 | 0 | 0 | 0 |
| SALT00000070656_Cluster10470-00 | 0 | 2870 | 0 | 0 | 0 |
| SALT00000059840_Cluster10561-00 | 0 | 3028 | 0 | 0 | 0 |
| SALT00000083126_Cluster10569-00 | 0 | 3028 | 0 | 0 | 0 |
| SALT00000038612_Cluster10582-00 | 0 | 1843 | 0 | 0 | 0 |
| SALT00000045517_Cluster10582-00 | 0 | 1577 | 0 | 0 | 0 |
| SALT00000045728_Cluster10582-00 | 0 | 2363 | 0 | 0 | 0 |
| SALT00000053227_Cluster10582-00 | 0 | 1492 | 0 | 0 | 0 |
| SALT00000024767_Cluster10582-00 | 0 | 2942 | 0 | 0 | 0 |
| SALT00000060516_Cluster10588-00 | 0 | 1505 | 0 | 0 | 0 |
| SALT00000068345_Cluster10600-00 | 1 | 3022 | 0 | 0 | 0 |
| SALT00000019778_Cluster10619-00 | 1 | 1670 | 0 | 0 | 0 |
| SALT00000070458_Cluster10655-00 | 0 | 3015 | 0 | 0 | 0 |
| SALT00000037631_Cluster10681-00 | 0 | 1809 | 0 | 0 | 0 |
| SALT00000052478_Cluster10681-00 | 0 | 1926 | 0 | 0 | 0 |
| SALT00000071238_Cluster10769-00 | 0 | 3002 | 0 | 0 | 0 |
| SALT00000023241_Cluster10798-00 | 0 | 2998 | 0 | 0 | 0 |
| SALT00000027676_Cluster10820-00 | 1 | 2846 | 0 | 0 | 0 |
| SALT00000031089_Cluster10820-00 | 1 | 2986 | 0 | 0 | 0 |
| SALT00000059455_Cluster10820-00 | 1 | 2240 | 0 | 0 | 0 |
| SALT00000078112_Cluster10820-00 | 1 | 2995 | 0 | 0 | 0 |
| SALT00000081014_Cluster10820-00 | 1 | 2312 | 0 | 0 | 0 |
| SALT00000068899_Cluster10870-00 | 0 | 2989 | 0 | 0 | 0 |
| SALT00000070193_Cluster10870-00 | 0 | 2989 | 0 | 0 | 0 |
| SALT00000056586_Cluster10871-00 | 1 | 1944 | 0 | 0 | 0 |
| SALT00000064937_Cluster10883-00 | 0 | 2988 | 0 | 0 | 0 |
| SALT00000073696_Cluster10891-00 | 0 | 2987 | 0 | 0 | 0 |

| | | | | | |
|---|---|---|---|---|---|
| SALT00000076542_Cluster10942-00 | 1 | 2980 | 0 | 0 | 0 |
| SALT00000074867_Cluster10958-00 | 0 | 2978 | 0 | 0 | 0 |
| SALT00000037987_Cluster11009-00 | 0 | 1585 | 0 | 0 | 0 |
| SALT00000047709_Cluster11041-00 | 1 | 2966 | 0 | 0 | 0 |
| SALT00000017561_Cluster11092-00 | 0 | 1362 | 0 | 0 | 0 |
| SALT00000013945_Cluster11095-00 | 1 | 1828 | 0 | 0 | 0 |
| SALT00000027479_Cluster11095-00 | 1 | 2960 | 0 | 0 | 0 |
| SALT00000069757_Cluster11095-00 | 1 | 2177 | 0 | 0 | 0 |
| SALT00000067539_Cluster11115-00 | 0 | 2957 | 0 | 0 | 0 |
| SALT00000063206_Cluster11132-00 | 1 | 2954 | 0 | 0 | 0 |
| SALT00000005825_Cluster11138-00 | 1 | 1748 | 0 | 0 | 0 |
| SALT00000022617_Cluster11138-00 | 0 | 2953 | 0 | 0 | 0 |
| SALT00000025693_Cluster11138-00 | 0 | 2416 | 0 | 0 | 0 |
| SALT00000062746_Cluster11138-00 | 0 | 2711 | 0 | 0 | 0 |
| SALT00000079935_Cluster11138-00 | 0 | 2389 | 0 | 0 | 0 |
| SALT00000068049_Cluster11159-00 | 1 | 2950 | 0 | 0 | 0 |
| SALT00000065655_Cluster11207-00 | 0 | 2945 | 0 | 0 | 0 |
| SALT00000072062_Cluster11278-00 | 0 | 2938 | 0 | 0 | 0 |
| SALT00000079186_Cluster11284-00 | 0 | 2937 | 0 | 0 | 0 |
| SALT00000026157_Cluster11306-00 | 0 | 2934 | 0 | 0 | 0 |
| SALT00000079229_Cluster11333-00 | 0 | 2932 | 0 | 0 | 0 |
| SALT00000063398_Cluster11379-00 | 0 | 2926 | 0 | 0 | 0 |
| SALT00000063484_Cluster11380-00 | 0 | 2926 | 0 | 0 | 0 |
| SALT00000072153_Cluster11410-00 | 0 | 2923 | 0 | 0 | 0 |
| SALT00000056700_Cluster11428-00 | 1 | 2921 | 0 | 0 | 0 |
| SALT00000069408_Cluster11448-00 | 0 | 2920 | 0 | 0 | 0 |
| SALT00000040096_Cluster11459-00 | 0 | 2919 | 0 | 0 | 0 |
| SALT00000028092_Cluster11468-00 | 0 | 2917 | 0 | 0 | 0 |
| SALT00000027851_Cluster11495-00 | 1 | 2766 | 0 | 0 | 0 |
| SALT00000042588_Cluster11495-00 | 1 | 1722 | 0 | 0 | 0 |
| SALT00000066375_Cluster11495-00 | 1 | 2165 | 0 | 0 | 0 |
| SALT00000078880_Cluster11495-00 | 1 | 2915 | 0 | 0 | 0 |
| SALT00000080004_Cluster11495-00 | 1 | 2911 | 0 | 0 | 0 |
| SALT00000092521_Cluster11495-00 | 0 | 3546 | 0 | 0 | 0 |
| SALT00000059490_Cluster11558-00 | 0 | 1656 | 0 | 0 | 0 |
| SALT00000001616_Cluster11575-00 | 0 | 1467 | 0 | 0 | 0 |
| SALT00000073301_Cluster11575-00 | 0 | 2907 | 0 | 0 | 0 |
| SALT00000065678_Cluster11708-00 | 0 | 2893 | 0 | 0 | 0 |
| SALT00000071472_Cluster11715-00 | 0 | 2893 | 0 | 0 | 0 |
| SALT00000079705_Cluster11763-00 | 0 | 2889 | 0 | 0 | 0 |
| SALT00000024989_Cluster11784-00 | 0 | 2755 | 0 | 0 | 0 |
| SALT00000029915_Cluster11784-00 | 0 | 2887 | 0 | 0 | 0 |
| SALT00000075407_Cluster11784-00 | 0 | 2783 | 0 | 0 | 0 |
| SALT00000049327_Cluster11794-00 | 0 | 1615 | 0 | 0 | 0 |
| SALT00000064061_Cluster11794-00 | 0 | 2886 | 0 | 0 | 0 |
| SALT00000070859_Cluster11794-00 | 0 | 2601 | 0 | 0 | 0 |
| SALT00000080587_Cluster11799-00 | 1 | 2886 | 0 | 0 | 0 |
| SALT00000058052_Cluster11862-00 | 0 | 1533 | 0 | 0 | 0 |
| SALT00000062204_Cluster11870-00 | 0 | 2880 | 0 | 0 | 0 |
| SALT00000068993_Cluster11883-00 | 0 | 2879 | 0 | 0 | 0 |
| SALT00000074196_Cluster11912-00 | 0 | 2877 | 0 | 0 | 0 |
| SALT00000079604_Cluster11912-00 | 0 | 2819 | 0 | 0 | 0 |
| SALT00000068076_Cluster11980-00 | 0 | 2870 | 0 | 0 | up |
| SALT00000029270_Cluster12017-00 | 0 | 2865 | 0 | 0 | 0 |
| SALT00000072602_Cluster12026-00 | 0 | 2865 | 0 | 0 | 0 |
| SALT00000020714_Cluster12040-00 | 1 | 2863 | 0 | 0 | 0 |
| SALT00000024075_Cluster12040-00 | 1 | 2791 | 0 | 0 | 0 |
| SALT00000066937_Cluster12040-00 | 1 | 3235 | 0 | 0 | 0 |

| | | | | | |
|---|---|---|---|---|---|
| SALT00000073900_Cluster12085-00 | 1 | 2531 | 0 | 0 | 0 |
| SALT00000021809_Cluster12229-00 | 0 | 2845 | 0 | 0 | 0 |
| SALT00000026240_Cluster12229-00 | 0 | 2813 | 0 | 0 | 0 |
| SALT00000027277_Cluster12229-00 | 0 | 2681 | 0 | 0 | 0 |
| SALT00000024297_Cluster12270-00 | 0 | 2841 | 0 | 0 | 0 |
| SALT00000048762_Cluster12270-00 | 0 | 1826 | 0 | 0 | 0 |
| SALT00000055822_Cluster12270-00 | 0 | 1682 | 0 | 0 | 0 |
| SALT00000074382_Cluster12270-00 | 0 | 2828 | 0 | 0 | 0 |
| SALT00000074884_Cluster12319-00 | 1 | 2837 | 0 | 0 | 0 |
| SALT00000018096_Cluster12343-00 | 0 | 1285 | 0 | 0 | 0 |
| SALT00000039699_Cluster12357-00 | 1 | 1908 | 0 | 0 | 0 |
| SALT00000041684_Cluster12357-00 | 0 | 1972 | 0 | 0 | 0 |
| SALT00000014025_Cluster12380-00 | 0 | 1950 | 0 | 0 | 0 |
| SALT00000068194_Cluster12380-00 | 1 | 2831 | 0 | 0 | 0 |
| SALT00000053658_Cluster12434-00 | 1 | 1772 | 0 | 0 | 0 |
| SALT00000069401_Cluster12434-00 | 1 | 2759 | 0 | 0 | 0 |
| SALT00000076188_Cluster12434-00 | 1 | 2826 | 0 | 0 | 0 |
| SALT00000070702_Cluster12434-00 | 1 | 2727 | 0 | 0 | 0 |
| SALT00000055600_Cluster12434-00 | 0 | 1874 | 0 | 0 | 0 |
| SALT00000016182_Cluster12434-00 | 0 | 1568 | 0 | 0 | 0 |
| SALT00000009709_Cluster12434-00 | 0 | 2149 | 0 | 0 | 0 |
| SALT00000006353_Cluster12446-00 | 0 | 2033 | 0 | 0 | 0 |
| SALT00000084440_Cluster12446-00 | 0 | 2825 | 0 | 0 | 0 |
| SALT00000074685_Cluster12464-00 | 0 | 2823 | 0 | 0 | 0 |
| SALT00000073308_Cluster12525-00 | 0 | 2818 | 0 | 0 | 0 |
| SALT00000067783_Cluster12543-00 | 1 | 2747 | 0 | 0 | 0 |
| SALT00000018907_Cluster12570-00 | 0 | 1269 | 0 | 0 | 0 |
| SALT00000023672_Cluster12594-00 | 0 | 2803 | 0 | 0 | 0 |
| SALT00000080111_Cluster12594-00 | 0 | 2340 | 0 | 0 | 0 |
| SALT00000031101_Cluster12594-00 | 1 | 2678 | 0 | 0 | 0 |
| SALT00000072250_Cluster12600-00 | 0 | 2810 | 0 | 0 | 0 |
| SALT00000074855_Cluster12676-00 | 0 | 2804 | 0 | 0 | 0 |
| SALT00000055948_Cluster12680-00 | 0 | 1861 | 0 | 0 | 0 |
| SALT00000082229_Cluster12680-00 | 0 | 2804 | 0 | 0 | 0 |
| SALT00000055635_Cluster12701-00 | 0 | 2801 | 0 | 0 | 0 |
| SALT00000007138_Cluster12714-00 | 0 | 1991 | 0 | 0 | 0 |
| SALT00000042392_Cluster12714-00 | 0 | 1649 | 0 | 0 | 0 |
| SALT00000072585_Cluster12714-00 | 0 | 1265 | 0 | 0 | 0 |
| SALT00000064375_Cluster12736-00 | 0 | 2797 | 0 | 0 | 0 |
| SALT00000082114_Cluster12744-00 | 1 | 2797 | 0 | 0 | 0 |
| SALT00000022863_Cluster12806-00 | 1 | 2791 | 0 | 0 | 0 |
| SALT00000052568_Cluster12806-00 | 1 | 1846 | 0 | 0 | 0 |
| SALT00000054357_Cluster12806-00 | 1 | 1797 | 0 | 0 | 0 |
| SALT00000077182_Cluster12806-00 | 0 | 2745 | 0 | 0 | 0 |
| SALT00000069499_Cluster12806-00 | 0 | 2838 | 0 | 0 | 0 |
| SALT00000025307_Cluster12806-00 | 0 | 2722 | 0 | 0 | 0 |
| SALT00000073803_Cluster12915-00 | 0 | 2782 | 0 | 0 | 0 |
| SALT00000068897_Cluster12932-00 | 0 | 2780 | 0 | 0 | 0 |
| SALT00000039361_Cluster12976-00 | 1 | 1731 | 0 | 0 | 0 |
| SALT00000016717_Cluster13021-00 | 0 | 1804 | 0 | 0 | 0 |
| SALT00000024866_Cluster13021-00 | 0 | 2771 | 0 | 0 | 0 |
| SALT00000063375_Cluster13021-00 | 0 | 2696 | 0 | 0 | 0 |
| SALT00000081234_Cluster13044-00 | 0 | 2770 | 0 | 0 | 0 |
| SALT00000003075_Cluster13127-00 | 0 | 1696 | 0 | 0 | 0 |
| SALT00000081774_Cluster13127-00 | 0 | 2764 | 0 | 0 | 0 |
| SALT00000029423_Cluster13153-00 | 0 | 2761 | 0 | 0 | 0 |
| SALT00000017133_Cluster13188-00 | 0 | 1861 | 0 | 0 | 0 |
| SALT00000023354_Cluster13188-00 | 0 | 2758 | 0 | 0 | 0 |

| | | | | | |
|---|---|---|---|---|---|
| SALT00000079042_Cluster13188-00 | 0 | 2714 | 0 | 0 | 0 |
| SALT00000072151_Cluster13188-00 | 0 | 2721 | 0 | 0 | 0 |
| SALT00000025498_Cluster13189-00 | 0 | 2758 | 0 | 0 | 0 |
| SALT00000076481_Cluster13230-00 | 1 | 2755 | 0 | 0 | 0 |
| SALT00000084031_Cluster13299-00 | 0 | 2749 | 0 | 0 | 0 |
| SALT00000065440_Cluster13306-00 | 0 | 2748 | 0 | 0 | 0 |
| SALT00000063950_Cluster13318-00 | 1 | 2747 | 0 | 0 | 0 |
| SALT00000059520_Cluster13364-00 | 0 | 1900 | 0 | 0 | 0 |
| SALT00000020765_Cluster13371-00 | 0 | 2742 | 0 | 0 | 0 |
| SALT00000078596_Cluster13371-00 | 0 | 2688 | 0 | 0 | 0 |
| SALT00000025313_Cluster13405-00 | 0 | 2712 | 0 | 0 | 0 |
| SALT00000077184_Cluster13405-00 | 0 | 2740 | 0 | 0 | 0 |
| SALT00000068389_Cluster13451-00 | 0 | 2736 | 0 | 0 | 0 |
| SALT00000062568_Cluster13473-00 | 1 | 2734 | 0 | 0 | 0 |
| SALT00000064208_Cluster13524-00 | 0 | 2730 | 0 | 0 | 0 |
| SALT00000049549_Cluster13562-00 | 0 | 1834 | 0 | 0 | 0 |
| SALT00000080669_Cluster13562-00 | 0 | 2728 | 0 | 0 | 0 |
| SALT00000070422_Cluster13562-00 | 0 | 2689 | 0 | 0 | 0 |
| SALT00000070440_Cluster13562-00 | 1 | 2615 | 0 | 0 | 0 |
| SALT00000059545_Cluster13575-00 | 0 | 1255 | 0 | 0 | 0 |
| SALT00000081452_Cluster13603-00 | 0 | 2725 | 0 | 0 | 0 |
| SALT00000064722_Cluster13619-00 | 0 | 2675 | 0 | 0 | 0 |
| SALT00000073051_Cluster13619-00 | 0 | 2724 | 0 | 0 | 0 |
| SALT00000038045_Cluster13630-00 | 0 | 1670 | 0 | 0 | 0 |
| SALT00000038139_Cluster13630-00 | 0 | 1837 | 0 | 0 | 0 |
| SALT00000081266_Cluster13642-00 | 1 | 2722 | 0 | 0 | 0 |
| SALT00000063601_Cluster13648-00 | 0 | 2721 | 0 | 0 | 0 |
| SALT00000071349_Cluster13672-00 | 1 | 2718 | 0 | 0 | 0 |
| SALT00000073555_Cluster13682-00 | 1 | 2717 | 0 | 0 | 0 |
| SALT00000064859_Cluster13691-00 | 1 | 2716 | 0 | 0 | 0 |
| SALT00000074502_Cluster13707-00 | 0 | 2715 | 0 | 0 | 0 |
| SALT00000066502_Cluster13710-00 | 1 | 2590 | 0 | 0 | 0 |
| SALT00000080227_Cluster13710-00 | 1 | 2715 | 0 | 0 | 0 |
| SALT00000079291_Cluster13736-00 | 0 | 2713 | 0 | 0 | 0 |
| SALT00000068278_Cluster13748-00 | 1 | 2712 | 0 | 0 | 0 |
| SALT00000055350_Cluster13752-00 | 0 | 1249 | 0 | 0 | 0 |
| SALT00000062347_Cluster13752-00 | 0 | 1273 | 0 | 0 | 0 |
| SALT00000080038_Cluster13779-00 | 0 | 2710 | 0 | 0 | 0 |
| SALT00000061442_Cluster13845-00 | 1 | 1854 | 0 | 0 | 0 |
| SALT00000069270_Cluster13845-00 | 0 | 2706 | 0 | 0 | 0 |
| SALT00000047888_Cluster13853-00 | 0 | 1379 | 0 | 0 | 0 |
| SALT00000030084_Cluster13855-00 | 0 | 2705 | 0 | 0 | 0 |
| SALT00000072224_Cluster13875-00 | 0 | 2704 | 0 | 0 | 0 |
| SALT00000053916_Cluster14037-00 | 0 | 1613 | 0 | 0 | 0 |
| SALT00000062041_Cluster14037-00 | 0 | 2692 | 0 | 0 | 0 |
| SALT00000029177_Cluster14050-00 | 0 | 2691 | 0 | 0 | 0 |
| SALT00000028053_Cluster14099-00 | 0 | 2687 | 0 | 0 | 0 |
| SALT00000077694_Cluster14169-00 | 0 | 2614 | 0 | 0 | 0 |
| SALT00000080711_Cluster14169-00 | 0 | 2682 | 0 | 0 | 0 |
| SALT00000073546_Cluster14180-00 | 0 | 2681 | 0 | 0 | 0 |
| SALT00000077778_Cluster14192-00 | 1 | 2680 | 0 | 0 | 0 |
| SALT00000023773_Cluster14196-00 | 0 | 2679 | 0 | 0 | 0 |
| SALT00000070142_Cluster14253-00 | 1 | 2675 | 0 | 0 | 0 |
| SALT00000077707_Cluster14269-00 | 0 | 2674 | 0 | 0 | 0 |
| SALT00000016523_Cluster14270-00 | 0 | 1702 | 0 | 0 | 0 |
| SALT00000052000_Cluster14270-00 | 0 | 1187 | 0 | 0 | 0 |
| SALT00000072180_Cluster14282-00 | 0 | 2673 | 0 | 0 | 0 |
| SALT00000082773_Cluster14332-00 | 1 | 2670 | 0 | 0 | 0 |

| | | | | | |
|---|---|---|---|---|---|
| SALT00000076718_Cluster14342-00 | 1 | 2669 | 0 | 0 | 0 |
| SALT00000068631_Cluster14401-00 | 0 | 2665 | 0 | 0 | 0 |
| SALT00000023775_Cluster14441-00 | 0 | 1611 | 0 | 0 | 0 |
| SALT00000080853_Cluster14455-00 | 0 | 2662 | 0 | 0 | 0 |
| SALT00000031068_Cluster14478-00 | 0 | 2660 | 0 | 0 | 0 |
| SALT00000068415_Cluster14478-00 | 0 | 1614 | 0 | 0 | 0 |
| SALT00000037476_Cluster14480-00 | 0 | 2660 | 0 | 0 | 0 |
| SALT00000083224_Cluster14497-00 | 1 | 2660 | 0 | 0 | 0 |
| SALT00000066464_Cluster14511-00 | 1 | 2658 | 0 | 0 | 0 |
| SALT00000028184_Cluster14573-00 | 0 | 2654 | 0 | 0 | 0 |
| SALT00000049779_Cluster14573-00 | 0 | 1805 | 0 | 0 | 0 |
| SALT00000013336_Cluster14573-00 | 0 | 1874 | 0 | 0 | 0 |
| SALT00000049292_Cluster14596-00 | 0 | 2133 | 0 | 0 | 0 |
| SALT00000051997_Cluster14596-00 | 1 | 1964 | 0 | 0 | 0 |
| SALT00000065528_Cluster14596-00 | 1 | 2580 | 0 | 0 | 0 |
| SALT00000073815_Cluster14596-00 | 0 | 2653 | 0 | 0 | 0 |
| SALT00000077480_Cluster14596-00 | 1 | 2600 | 0 | 0 | 0 |
| SALT00000060139_Cluster14596-00 | 0 | 1504 | 0 | 0 | 0 |
| SALT00000051006_Cluster14596-00 | 0 | 1652 | 0 | 0 | 0 |
| SALT00000073785_Cluster14598-00 | 1 | 2432 | 0 | 0 | 0 |
| SALT00000074009_Cluster14609-00 | 0 | 2652 | 0 | 0 | 0 |
| SALT00000063712_Cluster14725-00 | 0 | 2644 | 0 | 0 | 0 |
| SALT00000068317_Cluster14843-00 | 0 | 2637 | 0 | 0 | 0 |
| SALT00000070891_Cluster14866-00 | 0 | 2635 | 0 | 0 | 0 |
| SALT00000039838_Cluster14957-00 | 0 | 2628 | 0 | 0 | 0 |
| SALT00000065783_Cluster14962-00 | 0 | 2628 | 0 | 0 | 0 |
| SALT00000067411_Cluster14976-00 | 0 | 2627 | 0 | 0 | 0 |
| SALT00000074272_Cluster14976-00 | 0 | 2405 | 0 | 0 | 0 |
| SALT00000072961_Cluster15020-00 | 1 | 2623 | 0 | 0 | 0 |
| SALT00000058512_Cluster15034-00 | 1 | 2073 | 0 | 0 | 0 |
| SALT00000006360_Cluster15066-00 | 0 | 1775 | 0 | 0 | 0 |
| SALT00000052945_Cluster15066-00 | 1 | 1380 | 0 | 0 | 0 |
| SALT00000062218_Cluster15066-00 | 1 | 2620 | 0 | 0 | 0 |
| SALT00000082857_Cluster15105-00 | 0 | 2618 | 0 | 0 | 0 |
| SALT00000068020_Cluster15145-00 | 1 | 2615 | 0 | 0 | 0 |
| SALT00000019908_Cluster15228-00 | 0 | 1449 | up | 0 | 0 |
| SALT00000042553_Cluster15228-00 | 0 | 1635 | 0 | 0 | 0 |
| SALT00000024164_Cluster15251-00 | 1 | 2608 | 0 | 0 | 0 |
| SALT00000007764_Cluster15263-00 | 1 | 1697 | 0 | 0 | 0 |
| SALT00000030001_Cluster15263-00 | 0 | 2607 | 0 | 0 | 0 |
| SALT00000064470_Cluster15263-00 | 0 | 2568 | 0 | 0 | 0 |
| SALT00000069505_Cluster15263-00 | 0 | 2518 | 0 | 0 | 0 |
| SALT00000080534_Cluster15263-00 | 0 | 2597 | 0 | 0 | 0 |
| SALT00000042641_Cluster15308-00 | 0 | 1919 | 0 | 0 | 0 |
| SALT00000071514_Cluster15308-00 | 0 | 2604 | 0 | 0 | 0 |
| SALT00000082655_Cluster15350-00 | 0 | 2601 | 0 | 0 | 0 |
| SALT00000070133_Cluster15357-00 | 0 | 2600 | 0 | 0 | 0 |
| SALT00000082956_Cluster15357-00 | 0 | 2405 | 0 | 0 | 0 |
| SALT00000070582_Cluster15358-00 | 0 | 2600 | 0 | 0 | 0 |
| SALT00000053275_Cluster15389-00 | 0 | 2598 | 0 | 0 | 0 |
| SALT00000069695_Cluster15405-00 | 0 | 2597 | 0 | 0 | 0 |
| SALT00000041459_Cluster15408-00 | 0 | 1673 | 0 | 0 | 0 |
| SALT00000078809_Cluster15422-00 | 1 | 2596 | 0 | 0 | 0 |
| SALT00000042660_Cluster15440-00 | 1 | 2594 | 0 | 0 | 0 |
| SALT00000078720_Cluster15448-00 | 0 | 2594 | 0 | 0 | 0 |
| SALT00000073036_Cluster15477-00 | 1 | 2591 | 0 | 0 | down |
| SALT00000083797_Cluster15481-00 | 0 | 2591 | 0 | 0 | 0 |
| SALT00000070981_Cluster15511-00 | 0 | 2589 | 0 | 0 | 0 |

| | | | | | |
|---|---|---|---|---|---|
| SALT00000025490_Cluster15523-00 | 0 | 2587 | 0 | 0 | 0 |
| SALT00000029070_Cluster15523-00 | 0 | 2513 | 0 | 0 | 0 |
| SALT00000050912_Cluster15523-00 | 0 | 1790 | 0 | 0 | 0 |
| SALT00000030946_Cluster15523-00 | 1 | 2749 | 0 | 0 | 0 |
| SALT00000000585_Cluster15523-00 | 1 | 1813 | 0 | 0 | 0 |
| SALT00000005046_Cluster15548-00 | 1 | 1591 | 0 | 0 | 0 |
| SALT00000046247_Cluster15548-00 | 1 | 1404 | 0 | 0 | 0 |
| SALT00000066640_Cluster15650-00 | 0 | 2578 | 0 | 0 | 0 |
| SALT00000016785_Cluster15690-00 | 0 | 1680 | 0 | 0 | 0 |
| SALT00000053749_Cluster15804-00 | 0 | 1727 | 0 | 0 | 0 |
| SALT00000063309_Cluster15804-00 | 0 | 2566 | 0 | 0 | 0 |
| SALT00000072415_Cluster15824-00 | 0 | 2565 | 0 | 0 | 0 |
| SALT00000084081_Cluster15830-00 | 0 | 2565 | 0 | 0 | 0 |
| SALT00000070271_Cluster15863-00 | 0 | 2562 | 0 | 0 | 0 |
| SALT00000012852_Cluster15905-00 | 0 | 1533 | 0 | 0 | 0 |
| SALT00000082804_Cluster15908-00 | 1 | 2559 | 0 | 0 | 0 |
| SALT00000078387_Cluster16026-00 | 0 | 2550 | 0 | 0 | 0 |
| SALT00000075312_Cluster16046-00 | 0 | 2548 | 0 | 0 | 0 |
| SALT00000081845_Cluster16096-00 | 0 | 2545 | 0 | 0 | 0 |
| SALT00000011996_Cluster16111-00 | 1 | 2052 | 0 | 0 | 0 |
| SALT00000057938_Cluster16111-00 | 0 | 1563 | 0 | 0 | 0 |
| SALT00000082723_Cluster16134-00 | 0 | 2542 | 0 | 0 | 0 |
| SALT00000080684_Cluster16198-00 | 0 | 2537 | 0 | 0 | 0 |
| SALT00000036752_Cluster16208-00 | 1 | 2170 | 0 | 0 | 0 |
| SALT00000083971_Cluster16208-00 | 1 | 2536 | 0 | 0 | 0 |
| SALT00000081950_Cluster16243-00 | 1 | 2534 | 0 | 0 | 0 |
| SALT00000063457_Cluster16268-00 | 0 | 2532 | 0 | 0 | 0 |
| SALT00000069843_Cluster16284-00 | 0 | 2531 | 0 | 0 | 0 |
| SALT00000030167_Cluster16361-00 | 0 | 2526 | 0 | 0 | 0 |
| SALT00000071397_Cluster16361-00 | 0 | 2521 | 0 | 0 | 0 |
| SALT00000069509_Cluster16412-00 | 0 | 2523 | 0 | 0 | 0 |
| SALT00000068876_Cluster16434-00 | 0 | 2521 | 0 | 0 | 0 |
| SALT00000075956_Cluster16449-00 | 0 | 2520 | 0 | 0 | 0 |
| SALT00000079970_Cluster16538-00 | 0 | 2512 | 0 | 0 | 0 |
| SALT00000058236_Cluster16572-00 | 0 | 2508 | 0 | 0 | 0 |
| SALT00000063936_Cluster16573-00 | 1 | 2508 | 0 | 0 | 0 |
| SALT00000016549_Cluster16593-00 | 1 | 1454 | 0 | 0 | 0 |
| SALT00000079130_Cluster16606-00 | 0 | 2505 | 0 | 0 | 0 |
| SALT00000078045_Cluster16631-00 | 0 | 2503 | 0 | 0 | 0 |
| SALT00000003739_Cluster16673-00 | 0 | 1649 | 0 | 0 | 0 |
| SALT00000075538_Cluster16673-00 | 1 | 2500 | 0 | 0 | 0 |
| SALT00000070316_Cluster16713-00 | 0 | 2497 | 0 | 0 | 0 |
| SALT00000075121_Cluster16747-00 | 0 | 2495 | 0 | 0 | 0 |
| SALT00000082953_Cluster16754-00 | 0 | 2495 | 0 | 0 | 0 |
| SALT00000054665_Cluster16808-00 | 0 | 1785 | 0 | 0 | 0 |
| SALT00000027273_Cluster16855-00 | 0 | 2484 | 0 | 0 | 0 |
| SALT00000068152_Cluster16895-00 | 0 | 2480 | 0 | 0 | 0 |
| SALT00000048129_Cluster16913-00 | 0 | 2478 | 0 | 0 | 0 |
| SALT00000075777_Cluster16921-00 | 0 | 2478 | 0 | 0 | 0 |
| SALT00000080351_Cluster16966-00 | 0 | 2474 | 0 | 0 | 0 |
| SALT00000011653_Cluster17002-00 | 0 | 1537 | 0 | 0 | 0 |
| SALT00000040671_Cluster17002-00 | 0 | 2168 | 0 | 0 | 0 |
| SALT00000063385_Cluster17002-00 | 0 | 2470 | 0 | 0 | 0 |
| SALT00000027510_Cluster17043-00 | 0 | 1746 | 0 | 0 | 0 |
| SALT00000073121_Cluster17043-00 | 0 | 2466 | 0 | 0 | 0 |
| SALT00000041014_Cluster17060-00 | 0 | 1735 | up | 0 | 0 |
| SALT00000044384_Cluster17060-00 | 0 | 1711 | 0 | 0 | 0 |
| SALT00000082408_Cluster17060-00 | 0 | 2465 | 0 | 0 | 0 |

| | | | | | |
|---|---|---|---|---|---|
| SALT00000083773_Cluster17088-00 | 1 | 2462 | 0 | 0 | 0 |
| SALT00000082685_Cluster17096-00 | 0 | 2461 | 0 | 0 | 0 |
| SALT00000040390_Cluster17118-00 | 0 | 1315 | 0 | 0 | 0 |
| SALT00000010605_Cluster17161-00 | 0 | 1703 | 0 | 0 | 0 |
| SALT00000037646_Cluster17161-00 | 0 | 2454 | 0 | 0 | 0 |
| SALT00000053548_Cluster17161-00 | 0 | 1601 | 0 | 0 | 0 |
| SALT00000012441_Cluster17161-00 | 0 | 1591 | 0 | 0 | 0 |
| SALT00000073607_Cluster17192-00 | 0 | 2451 | 0 | 0 | 0 |
| SALT00000069577_Cluster17215-00 | 0 | 2448 | 0 | 0 | 0 |
| SALT00000029368_Cluster17245-00 | 0 | 2443 | 0 | 0 | 0 |
| SALT00000038391_Cluster17245-00 | 0 | 1733 | 0 | 0 | 0 |
| SALT00000052031_Cluster17245-00 | 0 | 1791 | 0 | 0 | 0 |
| SALT00000046324_Cluster17253-00 | 0 | 2442 | 0 | 0 | 0 |
| SALT00000024048_Cluster17260-00 | 0 | 2441 | 0 | 0 | 0 |
| SALT00000070459_Cluster17276-00 | 0 | 2440 | 0 | 0 | 0 |
| SALT00000072171_Cluster17282-00 | 0 | 2439 | 0 | 0 | 0 |
| SALT00000076910_Cluster17321-00 | 0 | 2434 | 0 | 0 | 0 |
| SALT00000078633_Cluster17340-00 | 1 | 2432 | 0 | 0 | 0 |
| SALT00000081854_Cluster17392-00 | 0 | 2424 | 0 | 0 | 0 |
| SALT00000073422_Cluster17409-00 | 1 | 2421 | 0 | 0 | 0 |
| SALT00000043851_Cluster17420-00 | 0 | 2419 | 0 | 0 | 0 |
| SALT00000060363_Cluster17435-00 | 0 | 2417 | 0 | 0 | 0 |
| SALT00000036778_Cluster17509-00 | 1 | 1910 | 0 | 0 | 0 |
| SALT00000061545_Cluster17509-00 | 1 | 1521 | 0 | 0 | 0 |
| SALT00000053093_Cluster17541-00 | 0 | 2397 | 0 | 0 | 0 |
| SALT00000066983_Cluster17600-00 | 0 | 2386 | 0 | 0 | 0 |
| SALT00000041124_Cluster17642-00 | 0 | 1680 | 0 | 0 | 0 |
| SALT00000041613_Cluster17644-00 | 0 | 2379 | 0 | 0 | 0 |
| SALT00000073928_Cluster17736-00 | 1 | 2361 | 0 | 0 | 0 |
| SALT00000044270_Cluster17750-00 | 0 | 2206 | 0 | 0 | 0 |
| SALT00000075227_Cluster17750-00 | 0 | 2354 | 0 | 0 | 0 |
| SALT00000063073_Cluster17776-00 | 0 | 2346 | 0 | 0 | 0 |
| SALT00000074673_Cluster17795-00 | 0 | 2343 | 0 | 0 | 0 |
| SALT00000057904_Cluster17901-00 | 0 | 2323 | 0 | 0 | 0 |
| SALT00000063548_Cluster17944-00 | 0 | 2312 | 0 | 0 | 0 |
| SALT00000043946_Cluster17960-00 | 0 | 2308 | 0 | 0 | 0 |
| SALT00000051779_Cluster17961-00 | 0 | 2308 | 0 | 0 | 0 |
| SALT00000051362_Cluster18015-00 | 1 | 2298 | 0 | 0 | 0 |
| SALT00000084220_Cluster18020-00 | 0 | 2298 | 0 | 0 | 0 |
| SALT00000076452_Cluster18075-00 | 0 | 2285 | 0 | 0 | 0 |
| SALT00000078996_Cluster18090-00 | 0 | 2283 | 0 | 0 | 0 |
| SALT00000008165_Cluster18095-00 | 1 | 1202 | 0 | 0 | 0 |
| SALT00000048417_Cluster18095-00 | 0 | 2282 | 0 | 0 | 0 |
| SALT00000063074_Cluster18095-00 | 0 | 2023 | 0 | 0 | 0 |
| SALT00000049058_Cluster18131-00 | 0 | 2275 | 0 | 0 | 0 |
| SALT00000050776_Cluster18131-00 | 0 | 1988 | 0 | 0 | 0 |
| SALT00000045651_Cluster18138-00 | 0 | 2274 | 0 | 0 | 0 |
| SALT00000008093_Cluster18153-00 | 0 | 1492 | 0 | 0 | 0 |
| SALT00000042421_Cluster18153-00 | 0 | 2271 | 0 | 0 | 0 |
| SALT00000037723_Cluster18153-00 | 0 | 1675 | 0 | 0 | 0 |
| SALT00000059634_Cluster18153-00 | 0 | 1892 | 0 | 0 | 0 |
| SALT00000049743_Cluster18157-00 | 0 | 2013 | 0 | 0 | 0 |
| SALT00000038964_Cluster18190-00 | 0 | 2265 | 0 | 0 | 0 |
| SALT00000047616_Cluster18196-00 | 1 | 2264 | 0 | 0 | 0 |
| SALT00000044881_Cluster18215-00 | 0 | 2262 | 0 | 0 | 0 |
| SALT00000048177_Cluster18217-00 | 0 | 2262 | 0 | 0 | 0 |
| SALT00000015348_Cluster18241-00 | 0 | 2258 | 0 | 0 | 0 |
| SALT00000060294_Cluster18286-00 | 0 | 1628 | 0 | 0 | 0 |

| | | | | | |
|---|---|---|---|---|---|
| SALT00000008850_Cluster18407-00 | 0 | 2230 | 0 | 0 | 0 |
| SALT00000038417_Cluster18412-00 | 0 | 2230 | 0 | 0 | 0 |
| SALT00000017340_Cluster18450-00 | 1 | 1671 | 0 | 0 | 0 |
| SALT00000057258_Cluster18450-00 | 1 | 2226 | 0 | 0 | 0 |
| SALT00000016448_Cluster18450-00 | 1 | 1714 | 0 | 0 | 0 |
| SALT00000070425_Cluster18616-00 | 0 | 2206 | 0 | 0 | 0 |
| SALT00000079153_Cluster18718-00 | 1 | 2194 | 0 | 0 | 0 |
| SALT00000043338_Cluster18731-00 | 0 | 2192 | 0 | 0 | 0 |
| SALT00000051147_Cluster18782-00 | 1 | 2084 | 0 | 0 | 0 |
| SALT00000065094_Cluster18782-00 | 1 | 2187 | 0 | 0 | 0 |
| SALT00000046571_Cluster18810-00 | 0 | 2076 | 0 | 0 | 0 |
| SALT00000047634_Cluster18810-00 | 0 | 2184 | 0 | 0 | 0 |
| SALT00000075517_Cluster18810-00 | 0 | 2161 | 0 | 0 | 0 |
| SALT00000012957_Cluster18825-00 | 0 | 2182 | 0 | 0 | 0 |
| SALT00000037496_Cluster18829-00 | 1 | 1456 | 0 | 0 | 0 |
| SALT00000056678_Cluster18946-00 | 1 | 2172 | 0 | 0 | 0 |
| SALT00000017152_Cluster18949-00 | 0 | 2171 | 0 | 0 | 0 |
| SALT00000040430_Cluster19068-00 | 1 | 2161 | 0 | 0 | 0 |
| SALT00000045306_Cluster19114-00 | 1 | 2157 | 0 | 0 | 0 |
| SALT00000050661_Cluster19267-00 | 0 | 2142 | 0 | 0 | 0 |
| SALT00000045273_Cluster19307-00 | 1 | 1549 | 0 | 0 | 0 |
| SALT00000060007_Cluster19307-00 | 1 | 2138 | 0 | 0 | 0 |
| SALT00000061868_Cluster19307-00 | 1 | 1533 | 0 | 0 | 0 |
| SALT00000041493_Cluster19327-00 | 0 | 2135 | 0 | 0 | 0 |
| SALT00000004923_Cluster19406-00 | 0 | 1559 | 0 | 0 | 0 |
| SALT00000048069_Cluster19406-00 | 0 | 2127 | 0 | 0 | 0 |
| SALT00000003186_Cluster19406-00 | 0 | 1429 | 0 | 0 | 0 |
| SALT00000016915_Cluster19467-00 | 1 | 2122 | 0 | 0 | 0 |
| SALT00000010849_Cluster19469-00 | 0 | 1472 | 0 | 0 | 0 |
| SALT00000018422_Cluster19469-00 | 0 | 2122 | 0 | 0 | 0 |
| SALT00000001817_Cluster19469-00 | 0 | 1565 | 0 | 0 | 0 |
| SALT00000051702_Cluster19473-00 | 1 | 2122 | 0 | 0 | 0 |
| SALT00000001572_Cluster19501-00 | 0 | 2119 | 0 | 0 | 0 |
| SALT00000019307_Cluster19550-00 | 0 | 2116 | 0 | 0 | 0 |
| SALT00000041646_Cluster19591-00 | 1 | 2044 | 0 | 0 | 0 |
| SALT00000046906_Cluster19591-00 | 1 | 2113 | 0 | 0 | 0 |
| SALT00000044971_Cluster19644-00 | 0 | 2108 | 0 | 0 | 0 |
| SALT00000042378_Cluster19648-00 | 0 | 2028 | 0 | 0 | 0 |
| SALT00000045755_Cluster19648-00 | 0 | 2097 | 0 | 0 | 0 |
| SALT00000058654_Cluster19648-00 | 1 | 2108 | 0 | 0 | 0 |
| SALT00000051875_Cluster19669-00 | 0 | 2106 | 0 | 0 | 0 |
| SALT00000017375_Cluster19674-00 | 0 | 2105 | 0 | 0 | 0 |
| SALT00000036705_Cluster19691-00 | 1 | 2104 | 0 | 0 | 0 |
| SALT00000060635_Cluster19709-00 | 0 | 2103 | 0 | 0 | 0 |
| SALT00000049528_Cluster19746-00 | 0 | 2099 | 0 | 0 | 0 |
| SALT00000004443_Cluster19748-00 | 1 | 1771 | 0 | 0 | 0 |
| SALT00000039896_Cluster19748-00 | 1 | 1941 | 0 | 0 | 0 |
| SALT00000040495_Cluster19748-00 | 1 | 1832 | 0 | 0 | 0 |
| SALT00000042709_Cluster19748-00 | 1 | 1610 | 0 | 0 | 0 |
| SALT00000057202_Cluster19748-00 | 1 | 2099 | 0 | 0 | 0 |

| | | | | | |
|---|---|---|---|---|---|
| SALT00000062113_Cluster19748-00 | 1 | 1607 | 0 | 0 | 0 |
| SALT00000061564_Cluster19766-00 | 1 | 2098 | 0 | 0 | 0 |
| SALT00000060648_Cluster19799-00 | 1 | 2096 | 0 | 0 | 0 |
| SALT00000006165_Cluster19801-00 | 0 | 2095 | 0 | 0 | 0 |
| SALT00000007332_Cluster19801-00 | 0 | 1811 | 0 | 0 | 0 |
| SALT00000051581_Cluster19801-00 | 0 | 1091 | 0 | 0 | down |
| SALT00000006116_Cluster19893-00 | 0 | 1534 | 0 | up | up |
| SALT00000039821_Cluster19893-00 | 0 | 1729 | 0 | 0 | up |
| SALT00000050513_Cluster19893-00 | 0 | 2088 | 0 | 0 | 0 |
| SALT00000053540_Cluster19893-00 | 0 | 1740 | 0 | 0 | 0 |
| SALT00000000300_Cluster19893-00 | 0 | 1781 | 0 | 0 | 0 |
| SALT00000035982_Cluster20008-00 | 0 | 2079 | 0 | 0 | 0 |
| SALT00000053560_Cluster20057-00 | 0 | 2075 | 0 | 0 | 0 |
| SALT00000043203_Cluster20066-00 | 0 | 2074 | 0 | 0 | 0 |
| SALT00000056238_Cluster20066-00 | 0 | 1775 | 0 | 0 | 0 |
| SALT00000035952_Cluster20100-00 | 1 | 1930 | 0 | 0 | 0 |
| SALT00000053970_Cluster20100-00 | 1 | 2072 | 0 | 0 | 0 |
| SALT00000058681_Cluster20100-00 | 1 | 1622 | 0 | 0 | 0 |
| SALT00000060808_Cluster20109-00 | 1 | 2071 | 0 | 0 | 0 |
| SALT00000015982_Cluster20112-00 | 0 | 2070 | 0 | 0 | 0 |
| SALT00000036965_Cluster20159-00 | 0 | 2066 | 0 | 0 | 0 |
| SALT00000038819_Cluster20225-00 | 0 | 2061 | 0 | 0 | 0 |
| SALT00000035785_Cluster20248-00 | 0 | 2059 | 0 | 0 | 0 |
| SALT00000075135_Cluster20297-00 | 0 | 2055 | 0 | 0 | 0 |
| SALT00000042746_Cluster20372-00 | 0 | 1807 | 0 | 0 | 0 |
| SALT00000012701_Cluster20467-00 | 0 | 2042 | 0 | 0 | 0 |
| SALT00000055509_Cluster20495-00 | 0 | 2041 | 0 | 0 | 0 |
| SALT00000042218_Cluster20559-00 | 0 | 2036 | 0 | 0 | 0 |
| SALT00000005536_Cluster20585-00 | 1 | 2034 | 0 | 0 | 0 |
| SALT00000003986_Cluster20675-00 | 0 | 1874 | 0 | 0 | 0 |
| SALT00000041946_Cluster20680-00 | 0 | 2028 | 0 | 0 | 0 |
| SALT00000050588_Cluster20680-00 | 0 | 1911 | 0 | 0 | 0 |
| SALT00000051146_Cluster20680-00 | 0 | 1432 | 0 | 0 | 0 |
| SALT00000040978_Cluster20688-00 | 0 | 2027 | 0 | 0 | 0 |
| SALT00000052464_Cluster20739-00 | 0 | 2024 | 0 | 0 | down |
| SALT00000054625_Cluster20743-00 | 0 | 2024 | 0 | 0 | 0 |
| SALT00000049004_Cluster20781-00 | 0 | 2021 | 0 | 0 | 0 |
| SALT00000007810_Cluster20850-00 | 0 | 2015 | 0 | 0 | 0 |
| SALT00000019719_Cluster20916-00 | 0 | 2010 | 0 | 0 | 0 |
| SALT00000010586_Cluster20949-00 | 0 | 2008 | 0 | 0 | 0 |
| SALT00000019044_Cluster20949-00 | 0 | 1928 | 0 | 0 | 0 |
| SALT00000055036_Cluster20949-00 | 0 | 1936 | 0 | 0 | 0 |
| SALT00000065198_Cluster20949-00 | 0 | 1811 | up | up | 0 |
| SALT00000021887_Cluster20965-00 | 0 | 1413 | 0 | 0 | 0 |
| SALT00000015327_Cluster21032-00 | 0 | 2002 | 0 | 0 | 0 |
| SALT00000051686_Cluster21032-00 | 1 | 1803 | 0 | 0 | 0 |

| | | | | | |
|---|---|---|---|---|---|
| SALT00000039343_Cluster21034-00 | 0 | 2002 | 0 | 0 | 0 |
| SALT00000067843_Cluster21054-00 | 1 | 2001 | 0 | 0 | 0 |
| SALT00000059616_Cluster21088-00 | 0 | 1999 | 0 | 0 | 0 |
| SALT00000012210_Cluster21116-00 | 0 | 1540 | 0 | 0 | 0 |
| SALT00000035735_Cluster21116-00 | 0 | 1997 | 0 | 0 | 0 |
| SALT00000053937_Cluster21116-00 | 0 | 2148 | 0 | 0 | 0 |
| SALT00000009464_Cluster21169-00 | 1 | 1993 | 0 | 0 | 0 |
| SALT00000038244_Cluster21187-00 | 0 | 1992 | 0 | 0 | 0 |
| SALT00000051949_Cluster21211-00 | 0 | 1990 | 0 | 0 | 0 |
| SALT00000056322_Cluster21252-00 | 0 | 1988 | 0 | 0 | 0 |
| SALT00000065843_Cluster21287-00 | 1 | 1986 | 0 | 0 | 0 |
| SALT00000051482_Cluster21300-00 | 0 | 1985 | 0 | 0 | 0 |
| SALT00000046533_Cluster21314-00 | 0 | 1661 | 0 | 0 | 0 |
| SALT00000052579_Cluster21314-00 | 0 | 1789 | 0 | 0 | 0 |
| SALT00000056242_Cluster21314-00 | 0 | 1814 | 0 | 0 | 0 |
| SALT00000014864_Cluster21316-00 | 1 | 1983 | 0 | 0 | 0 |
| SALT00000012147_Cluster21322-00 | 0 | 1712 | 0 | 0 | 0 |
| SALT00000045188_Cluster21322-00 | 1 | 1983 | 0 | 0 | 0 |
| SALT00000050277_Cluster21323-00 | 0 | 1983 | 0 | 0 | 0 |
| SALT00000067224_Cluster21362-00 | 1 | 1981 | 0 | 0 | 0 |
| SALT00000056295_Cluster21458-00 | 0 | 1973 | 0 | 0 | 0 |
| SALT00000015842_Cluster21505-00 | 0 | 1970 | 0 | 0 | 0 |
| SALT00000069808_Cluster21534-00 | 0 | 1969 | 0 | 0 | 0 |
| SALT00000008741_Cluster21538-00 | 1 | 1968 | 0 | 0 | up |
| SALT00000058609_Cluster21545-00 | 0 | 1455 | 0 | 0 | 0 |
| SALT00000046234_Cluster21589-00 | 0 | 1966 | 0 | 0 | 0 |
| SALT00000048960_Cluster21625-00 | 0 | 1964 | 0 | 0 | 0 |
| SALT00000030850_Cluster21752-00 | 0 | 1955 | 0 | 0 | 0 |
| SALT00000056977_Cluster21752-00 | 0 | 1865 | 0 | 0 | 0 |
| SALT00000051314_Cluster21752-00 | 0 | 1717 | 0 | 0 | 0 |
| SALT00000037943_Cluster21782-00 | 1 | 1953 | 0 | 0 | 0 |
| SALT00000041485_Cluster21822-00 | 1 | 1808 | 0 | 0 | 0 |
| SALT00000046619_Cluster21822-00 | 0 | 1951 | 0 | 0 | 0 |
| SALT00000037552_Cluster21914-00 | 0 | 1945 | 0 | 0 | 0 |
| SALT00000062100_Cluster21961-00 | 0 | 1942 | 0 | 0 | 0 |
| SALT00000049617_Cluster21970-00 | 0 | 1783 | 0 | 0 | 0 |
| SALT00000060869_Cluster21970-00 | 0 | 1825 | 0 | 0 | 0 |
| SALT00000047334_Cluster21970-00 | 0 | 1600 | 0 | 0 | 0 |
| SALT00000012190_Cluster21970-00 | 0 | 1626 | 0 | 0 | 0 |
| SALT00000035806_Cluster22074-00 | 0 | 1934 | 0 | 0 | up |
| SALT00000058631_Cluster22094-00 | 0 | 1933 | 0 | 0 | 0 |
| SALT00000011007_Cluster22124-00 | 0 | 1930 | 0 | 0 | 0 |
| SALT00000037606_Cluster22124-00 | 0 | 1308 | 0 | 0 | 0 |
| SALT00000003913_Cluster22146-00 | 0 | 1929 | 0 | 0 | 0 |
| SALT00000001842_Cluster22163-00 | 0 | 1928 | 0 | 0 | 0 |
| SALT00000053661_Cluster22176-00 | 1 | 1928 | 0 | 0 | 0 |

| | | | | | |
|---|---|---|---|---|---|
| SALT00000059170_Cluster22197-00 | 0 | 1927 | 0 | 0 | 0 |
| SALT00000043426_Cluster22478-00 | 0 | 1909 | 0 | 0 | 0 |
| SALT00000061186_Cluster22509-00 | 1 | 1351 | 0 | 0 | 0 |
| SALT00000036312_Cluster22569-00 | 0 | 1552 | 0 | 0 | 0 |
| SALT00000057544_Cluster22588-00 | 0 | 1904 | 0 | 0 | 0 |
| SALT00000039270_Cluster22710-00 | 1 | 1896 | 0 | 0 | 0 |
| SALT00000041194_Cluster22736-00 | 0 | 1894 | 0 | 0 | 0 |
| SALT00000019582_Cluster22857-00 | 0 | 1886 | 0 | 0 | 0 |
| SALT00000041017_Cluster22874-00 | 1 | 1885 | 0 | 0 | 0 |
| SALT00000037553_Cluster22899-00 | 1 | 1884 | 0 | 0 | 0 |
| SALT00000047272_Cluster23003-00 | 1 | 1877 | 0 | 0 | 0 |
| SALT00000041791_Cluster23082-00 | 0 | 1872 | 0 | 0 | 0 |
| SALT00000058339_Cluster23103-00 | 1 | 1871 | 0 | 0 | 0 |
| SALT00000047369_Cluster23110-00 | 1 | 1870 | 0 | 0 | 0 |
| SALT00000055636_Cluster23202-00 | 0 | 1865 | 0 | 0 | 0 |
| SALT00000002247_Cluster23203-00 | 1 | 1448 | 0 | 0 | 0 |
| SALT00000010153_Cluster23203-00 | 1 | 1444 | 0 | 0 | 0 |
| SALT00000056377_Cluster23203-00 | 0 | 1865 | 0 | 0 | 0 |
| SALT00000018375_Cluster23227-00 | 1 | 1863 | 0 | 0 | 0 |
| SALT00000036026_Cluster23324-00 | 1 | 1857 | 0 | 0 | 0 |
| SALT00000013805_Cluster23364-00 | 0 | 1368 | 0 | 0 | 0 |
| SALT00000043493_Cluster23364-00 | 0 | 1855 | 0 | 0 | 0 |
| SALT00000050577_Cluster23364-00 | 0 | 1749 | 0 | 0 | 0 |
| SALT00000053183_Cluster23364-00 | 0 | 1800 | 0 | 0 | 0 |
| SALT00000015823_Cluster23364-00 | 0 | 1575 | 0 | 0 | 0 |
| SALT00000010331_Cluster23429-00 | 1 | 1851 | 0 | 0 | 0 |
| SALT00000045354_Cluster23458-00 | 1 | 1850 | 0 | 0 | 0 |
| SALT00000007957_Cluster23470-00 | 1 | 1849 | 0 | 0 | 0 |
| SALT00000015322_Cluster23528-00 | 0 | 1509 | 0 | 0 | 0 |
| SALT00000047143_Cluster23528-00 | 0 | 1710 | 0 | 0 | 0 |
| SALT00000050935_Cluster23528-00 | 0 | 1603 | 0 | 0 | 0 |
| SALT00000054320_Cluster23528-00 | 0 | 1845 | 0 | 0 | 0 |
| SALT00000021712_Cluster23589-00 | 0 | 1841 | 0 | 0 | 0 |
| SALT00000037890_Cluster23589-00 | 0 | 1753 | 0 | 0 | 0 |
| SALT00000043286_Cluster23591-00 | 0 | 1841 | 0 | 0 | 0 |
| SALT00000038276_Cluster23606-00 | 1 | 1840 | 0 | 0 | 0 |
| SALT00000058415_Cluster23632-00 | 0 | 1839 | 0 | 0 | 0 |
| SALT00000055827_Cluster23641-00 | 0 | 1884 | 0 | 0 | 0 |
| SALT00000010632_Cluster23647-00 | 1 | 1837 | 0 | 0 | 0 |
| SALT00000047313_Cluster23647-00 | 1 | 1566 | 0 | 0 | 0 |
| SALT00000036588_Cluster23647-00 | 1 | 1684 | 0 | 0 | 0 |
| SALT00000048684_Cluster23657-00 | 0 | 1837 | 0 | 0 | 0 |
| SALT00000048457_Cluster23690-00 | 1 | 1835 | 0 | 0 | 0 |
| SALT00000036442_Cluster23778-00 | 0 | 1830 | 0 | 0 | 0 |
| SALT00000006150_Cluster23840-00 | 0 | 1702 | 0 | 0 | 0 |
| SALT00000041998_Cluster23840-00 | 0 | 1827 | 0 | 0 | 0 |

| | | | | | |
|---|---|---|---|---|---|
| SALT00000045741_Cluster23844-00 | 0 | 1827 | 0 | 0 | 0 |
| SALT00000046264_Cluster23882-00 | 1 | 1825 | 0 | 0 | 0 |
| SALT00000050743_Cluster23884-00 | 0 | 1825 | 0 | 0 | 0 |
| SALT00000030373_Cluster23920-00 | 0 | 1553 | 0 | 0 | 0 |
| SALT00000051160_Cluster23920-00 | 0 | 1823 | 0 | 0 | 0 |
| SALT00000008694_Cluster24078-00 | 0 | 1814 | 0 | 0 | 0 |
| SALT00000041712_Cluster24100-00 | 1 | 1813 | 0 | 0 | 0 |
| SALT00000001884_Cluster24127-00 | 0 | 1811 | 0 | 0 | 0 |
| SALT00000035956_Cluster24135-00 | 1 | 1811 | 0 | 0 | 0 |
| SALT00000058375_Cluster24144-00 | 1 | 1811 | 0 | 0 | 0 |
| SALT00000047189_Cluster24160-00 | 1 | 1810 | 0 | 0 | 0 |
| SALT00000035882_Cluster24179-00 | 0 | 1712 | 0 | 0 | 0 |
| SALT00000044723_Cluster24179-00 | 0 | 1598 | 0 | 0 | 0 |
| SALT00000051329_Cluster24179-00 | 0 | 1809 | 0 | 0 | 0 |
| SALT00000045444_Cluster24221-00 | 0 | 1807 | 0 | 0 | 0 |
| SALT00000013321_Cluster24286-00 | 0 | 1803 | 0 | 0 | 0 |
| SALT00000050078_Cluster24317-00 | 0 | 1801 | 0 | 0 | 0 |
| SALT00000052480_Cluster24391-00 | 1 | 1797 | 0 | 0 | 0 |
| SALT00000036854_Cluster24433-00 | 0 | 1794 | 0 | 0 | 0 |
| SALT00000057733_Cluster24443-00 | 0 | 1794 | 0 | 0 | 0 |
| SALT00000009285_Cluster24494-00 | 0 | 1790 | 0 | 0 | 0 |
| SALT00000016215_Cluster24554-00 | 0 | 1527 | 0 | 0 | 0 |
| SALT00000049369_Cluster24554-00 | 0 | 1787 | 0 | 0 | 0 |
| SALT00000037600_Cluster24636-00 | 0 | 1782 | 0 | 0 | 0 |
| SALT00000046740_Cluster24636-00 | 0 | 1570 | 0 | 0 | 0 |
| SALT00000019636_Cluster24656-00 | 0 | 1781 | 0 | 0 | 0 |
| SALT00000054779_Cluster24687-00 | 0 | 1780 | 0 | 0 | 0 |
| SALT00000044042_Cluster24749-00 | 0 | 1646 | 0 | 0 | 0 |
| SALT00000056093_Cluster24749-00 | 0 | 1777 | 0 | 0 | 0 |
| SALT00000059451_Cluster24749-00 | 0 | 1751 | 0 | 0 | 0 |
| SALT00000057019_Cluster24750-00 | 0 | 1777 | 0 | 0 | 0 |
| SALT00000008108_Cluster24792-00 | 0 | 1774 | 0 | 0 | 0 |
| SALT00000010663_Cluster24794-00 | 1 | 1774 | 0 | 0 | 0 |
| SALT00000046252_Cluster24803-00 | 0 | 1774 | 0 | 0 | 0 |
| SALT00000001608_Cluster24813-00 | 1 | 1773 | 0 | 0 | 0 |
| SALT00000042940_Cluster24828-00 | 0 | 1772 | 0 | 0 | 0 |
| SALT00000019930_Cluster24842-00 | 0 | 1771 | 0 | 0 | 0 |
| SALT00000036607_Cluster24845-00 | 1 | 1541 | 0 | 0 | 0 |
| SALT00000037489_Cluster24855-00 | 0 | 1770 | 0 | 0 | 0 |
| SALT00000037594_Cluster24856-00 | 0 | 1770 | 0 | 0 | 0 |
| SALT00000044921_Cluster24861-00 | 0 | 1770 | 0 | 0 | 0 |
| SALT00000056272_Cluster24912-00 | 0 | 1768 | 0 | 0 | 0 |
| SALT00000036576_Cluster24956-00 | 0 | 1765 | 0 | 0 | 0 |
| SALT00000039580_Cluster24957-00 | 0 | 1765 | 0 | 0 | up |
| SALT00000049850_Cluster24963-00 | 0 | 1765 | 0 | 0 | 0 |
| SALT00000057755_Cluster24985-00 | 0 | 1764 | 0 | 0 | 0 |

| | | | | | |
|---|---|---|---|---|---|
| SALT00000036917_Cluster25102-00 | 0 | 1757 | 0 | 0 | 0 |
| SALT00000059786_Cluster25118-00 | 0 | 1757 | 0 | 0 | 0 |
| SALT00000039477_Cluster25197-00 | 0 | 1752 | 0 | 0 | 0 |
| SALT00000048865_Cluster25206-00 | 0 | 1752 | 0 | 0 | 0 |
| SALT00000040285_Cluster25225-00 | 0 | 1751 | 0 | 0 | 0 |
| SALT00000045094_Cluster25227-00 | 0 | 1751 | 0 | 0 | 0 |
| SALT00000046917_Cluster25279-00 | 0 | 1748 | 0 | 0 | 0 |
| SALT00000045491_Cluster25358-00 | 0 | 1744 | 0 | 0 | 0 |
| SALT00000000831_Cluster25375-00 | 0 | 1509 | 0 | 0 | 0 |
| SALT00000020080_Cluster25375-00 | 0 | 1743 | 0 | 0 | 0 |
| SALT00000055315_Cluster25375-00 | 0 | 1572 | 0 | 0 | 0 |
| SALT00000050575_Cluster25438-00 | 0 | 1740 | 0 | 0 | 0 |
| SALT00000020152_Cluster25495-00 | 1 | 1737 | 0 | 0 | 0 |
| SALT00000054098_Cluster25495-00 | 1 | 1579 | 0 | 0 | 0 |
| SALT00000051058_Cluster25495-00 | 1 | 1479 | 0 | 0 | 0 |
| SALT00000016360_Cluster25504-00 | 1 | 1527 | 0 | 0 | 0 |
| SALT00000053113_Cluster25504-00 | 1 | 1737 | 0 | 0 | 0 |
| SALT00000007350_Cluster25513-00 | 0 | 1736 | 0 | 0 | 0 |
| SALT00000004579_Cluster25536-00 | 0 | 1517 | 0 | 0 | 0 |
| SALT00000012581_Cluster25536-00 | 0 | 1598 | 0 | 0 | 0 |
| SALT00000037364_Cluster25536-00 | 0 | 1735 | 0 | 0 | 0 |
| SALT00000036287_Cluster25536-00 | 1 | 1732 | 0 | 0 | 0 |
| SALT00000006447_Cluster25546-00 | 0 | 1601 | 0 | 0 | 0 |
| SALT00000047844_Cluster25546-00 | 0 | 1661 | 0 | 0 | 0 |
| SALT00000052986_Cluster25546-00 | 0 | 1735 | 0 | 0 | 0 |
| SALT00000016304_Cluster25567-00 | 0 | 1418 | 0 | 0 | 0 |
| SALT00000037355_Cluster25567-00 | 0 | 1459 | 0 | 0 | 0 |
| SALT00000045042_Cluster25567-00 | 0 | 1734 | 0 | 0 | 0 |
| SALT00000053418_Cluster25567-00 | 0 | 1686 | 0 | 0 | 0 |
| SALT00000001668_Cluster25576-00 | 0 | 1733 | 0 | 0 | 0 |
| SALT00000038309_Cluster25589-00 | 0 | 1733 | 0 | 0 | 0 |
| SALT00000007986_Cluster25600-00 | 0 | 1732 | 0 | 0 | 0 |
| SALT00000058492_Cluster25640-00 | 0 | 1731 | 0 | 0 | 0 |
| SALT00000058764_Cluster25689-00 | 0 | 1728 | 0 | 0 | 0 |
| SALT00000064352_Cluster25691-00 | 0 | 1728 | 0 | 0 | 0 |
| SALT00000002487_Cluster25694-00 | 1 | 1727 | 0 | 0 | 0 |
| SALT00000036925_Cluster25723-00 | 1 | 1726 | 0 | 0 | 0 |
| SALT00000057505_Cluster25819-00 | 0 | 1722 | 0 | 0 | 0 |
| SALT00000013066_Cluster25840-00 | 1 | 1720 | 0 | 0 | 0 |
| SALT00000052214_Cluster25849-00 | 0 | 1720 | 0 | 0 | 0 |
| SALT00000018514_Cluster25853-00 | 0 | 1581 | 0 | 0 | 0 |
| SALT00000057285_Cluster25853-00 | 0 | 1720 | 0 | 0 | 0 |
| SALT00000048332_Cluster25853-00 | 0 | 1738 | 0 | 0 | 0 |
| SALT00000037367_Cluster25874-00 | 0 | 1718 | 0 | 0 | 0 |
| SALT00000004806_Cluster25906-00 | 0 | 1716 | 0 | 0 | 0 |
| SALT00000017526_Cluster25909-00 | 0 | 1716 | 0 | 0 | 0 |

| | | | | | |
|---|---|---|---|---|---|
| SALT00000055367_Cluster25947-00 | 0 | 1715 | 0 | 0 | 0 |
| SALT00000015307_Cluster25994-00 | 0 | 1712 | 0 | 0 | down |
| SALT00000047952_Cluster26002-00 | 0 | 1712 | 0 | 0 | 0 |
| SALT00000040235_Cluster26018-00 | 0 | 1711 | 0 | 0 | 0 |
| SALT00000055608_Cluster26033-00 | 1 | 1710 | 0 | 0 | 0 |
| SALT00000057581_Cluster26034-00 | 1 | 1710 | 0 | 0 | 0 |
| SALT00000050809_Cluster26045-00 | 0 | 1709 | 0 | 0 | 0 |
| SALT00000058357_Cluster26050-00 | 1 | 1709 | 0 | 0 | 0 |
| SALT00000036198_Cluster26056-00 | 1 | 1113 | 0 | 0 | 0 |
| SALT00000036636_Cluster26056-00 | 0 | 1708 | 0 | 0 | 0 |
| SALT00000049732_Cluster26056-00 | 0 | 1474 | 0 | 0 | 0 |
| SALT00000036921_Cluster26057-00 | 0 | 1708 | 0 | 0 | 0 |
| SALT00000039305_Cluster26088-00 | 0 | 1706 | 0 | 0 | 0 |
| SALT00000016964_Cluster26148-00 | 0 | 1164 | 0 | up | up |
| SALT00000029841_Cluster26148-00 | 0 | 1308 | 0 | 0 | 0 |
| SALT00000036080_Cluster26191-00 | 0 | 1699 | 0 | 0 | 0 |
| SALT00000051057_Cluster26239-00 | 1 | 1697 | 0 | 0 | 0 |
| SALT00000009994_Cluster26313-00 | 0 | 1589 | 0 | 0 | 0 |
| SALT00000022882_Cluster26313-00 | 0 | 1693 | 0 | 0 | 0 |
| SALT00000011611_Cluster26331-00 | 0 | 1692 | 0 | 0 | 0 |
| SALT00000042274_Cluster26363-00 | 0 | 1691 | 0 | 0 | 0 |
| SALT00000017473_Cluster26405-00 | 1 | 1516 | 0 | 0 | 0 |
| SALT00000036975_Cluster26405-00 | 0 | 1689 | 0 | 0 | 0 |
| SALT00000036534_Cluster26408-00 | 0 | 1647 | 0 | 0 | 0 |
| SALT00000049575_Cluster26408-00 | 0 | 1689 | 0 | 0 | 0 |
| SALT00000054083_Cluster26410-00 | 0 | 1689 | 0 | 0 | 0 |
| SALT00000060179_Cluster26414-00 | 0 | 1689 | 0 | 0 | 0 |
| SALT00000056598_Cluster26448-00 | 0 | 1687 | 0 | 0 | 0 |
| SALT00000039855_Cluster26509-00 | 1 | 1684 | 0 | 0 | 0 |
| SALT00000042307_Cluster26551-00 | 0 | 1682 | 0 | 0 | 0 |
| SALT00000057061_Cluster26579-00 | 0 | 1681 | 0 | 0 | 0 |
| SALT00000061095_Cluster26579-00 | 0 | 1509 | 0 | 0 | 0 |
| SALT00000057420_Cluster26604-00 | 0 | 1680 | 0 | 0 | 0 |
| SALT00000020187_Cluster26617-00 | 0 | 1679 | 0 | 0 | 0 |
| SALT00000053506_Cluster26626-00 | 0 | 1679 | 0 | 0 | 0 |
| SALT00000061555_Cluster26627-00 | 0 | 1679 | 0 | 0 | 0 |
| SALT00000047562_Cluster26645-00 | 0 | 1678 | 0 | 0 | 0 |
| SALT00000051635_Cluster26648-00 | 0 | 1678 | 0 | 0 | 0 |
| SALT00000059082_Cluster26650-00 | 0 | 1678 | 0 | 0 | 0 |
| SALT00000007314_Cluster26673-00 | 0 | 1676 | 0 | 0 | 0 |
| SALT00000059498_Cluster26707-00 | 0 | 1675 | 0 | 0 | 0 |
| SALT00000018145_Cluster26753-00 | 0 | 1672 | 0 | 0 | 0 |
| SALT00000051740_Cluster26758-00 | 0 | 1672 | 0 | 0 | 0 |
| SALT00000008702_Cluster26765-00 | 0 | 1671 | 0 | 0 | 0 |
| SALT00000052053_Cluster26777-00 | 0 | 1671 | 0 | 0 | 0 |
| SALT00000019063_Cluster26804-00 | 0 | 1669 | 0 | 0 | 0 |

| | | | | | |
|---|---|---|---|---|---|
| SALT00000023705_Cluster26805-00 | 0 | 1669 | 0 | 0 | 0 |
| SALT00000016810_Cluster26840-00 | 0 | 1667 | 0 | 0 | 0 |
| SALT00000017164_Cluster26848-00 | 0 | 1589 | 0 | 0 | 0 |
| SALT00000050592_Cluster26848-00 | 0 | 1667 | 0 | 0 | 0 |
| SALT00000052201_Cluster26848-00 | 0 | 1393 | 0 | 0 | 0 |
| SALT00000012752_Cluster26856-00 | 1 | 1666 | 0 | 0 | 0 |
| SALT00000037989_Cluster26861-00 | 0 | 1666 | 0 | 0 | 0 |
| SALT00000051134_Cluster26861-00 | 0 | 1605 | 0 | 0 | 0 |
| SALT00000052193_Cluster26868-00 | 0 | 1666 | 0 | 0 | 0 |
| SALT00000044493_Cluster26880-00 | 0 | 1665 | 0 | 0 | 0 |
| SALT00000045281_Cluster26881-00 | 0 | 1665 | 0 | 0 | 0 |
| SALT00000040424_Cluster26905-00 | 0 | 1664 | 0 | 0 | 0 |
| SALT00000046386_Cluster26970-00 | 0 | 1661 | 0 | 0 | 0 |
| SALT00000059226_Cluster26997-00 | 0 | 1660 | 0 | 0 | 0 |
| SALT00000003651_Cluster27006-00 | 0 | 1585 | 0 | 0 | 0 |
| SALT00000014296_Cluster27006-00 | 0 | 1659 | 0 | 0 | 0 |
| SALT00000049764_Cluster27035-00 | 0 | 1658 | 0 | 0 | 0 |
| SALT00000018495_Cluster27096-00 | 0 | 1655 | 0 | 0 | 0 |
| SALT00000045464_Cluster27103-00 | 0 | 1655 | 0 | 0 | 0 |
| SALT00000051093_Cluster27106-00 | 0 | 1655 | 0 | 0 | 0 |
| SALT00000018860_Cluster27122-00 | 0 | 1654 | 0 | 0 | 0 |
| SALT00000066281_Cluster27157-00 | 0 | 1653 | 0 | 0 | 0 |
| SALT00000039542_Cluster27171-00 | 0 | 1652 | 0 | 0 | 0 |
| SALT00000019337_Cluster27190-00 | 0 | 1651 | 0 | 0 | 0 |
| SALT00000001166_Cluster27216-00 | 1 | 1648 | 0 | 0 | 0 |
| SALT00000035946_Cluster27216-00 | 0 | 1650 | 0 | 0 | 0 |
| SALT00000043680_Cluster27225-00 | 0 | 1650 | 0 | 0 | 0 |
| SALT00000013947_Cluster27241-00 | 1 | 1649 | 0 | 0 | 0 |
| SALT00000043510_Cluster27275-00 | 0 | 1551 | 0 | 0 | 0 |
| SALT00000044519_Cluster27275-00 | 0 | 1488 | 0 | 0 | 0 |
| SALT00000044752_Cluster27275-00 | 0 | 1629 | 0 | 0 | 0 |
| SALT00000049075_Cluster27275-00 | 0 | 1588 | 0 | 0 | 0 |
| SALT00000052209_Cluster27275-00 | 0 | 1647 | 0 | 0 | 0 |
| SALT00000064083_Cluster27275-00 | 0 | 1466 | 0 | 0 | 0 |
| SALT00000041222_Cluster27275-00 | 0 | 1507 | 0 | 0 | 0 |
| SALT00000017236_Cluster27290-00 | 0 | 1646 | 0 | 0 | 0 |
| SALT00000013261_Cluster27321-00 | 1 | 1644 | 0 | 0 | 0 |
| SALT00000043055_Cluster27374-00 | 1 | 1642 | 0 | 0 | 0 |
| SALT00000053427_Cluster27379-00 | 0 | 1642 | 0 | 0 | 0 |
| SALT00000009102_Cluster27405-00 | 0 | 1640 | 0 | 0 | 0 |
| SALT00000002346_Cluster27447-00 | 0 | 1638 | 0 | 0 | 0 |
| SALT00000055386_Cluster27481-00 | 0 | 1637 | 0 | 0 | 0 |
| SALT00000061314_Cluster27506-00 | 1 | 1636 | 0 | 0 | 0 |
| SALT00000043344_Cluster27534-00 | 1 | 1634 | 0 | 0 | 0 |
| SALT00000011847_Cluster27607-00 | 0 | 1630 | 0 | 0 | 0 |
| SALT00000023450_Cluster27612-00 | 0 | 1630 | 0 | 0 | 0 |

| | | | | | |
|---|---|---|---|---|---|
| SALT00000011261_Cluster27712-00 | 0 | 1625 | 0 | 0 | 0 |
| SALT00000041289_Cluster27719-00 | 1 | 1625 | 0 | 0 | 0 |
| SALT00000047744_Cluster27719-00 | 1 | 1578 | 0 | 0 | 0 |
| SALT00000037456_Cluster27736-00 | 0 | 1624 | 0 | 0 | 0 |
| SALT00000003464_Cluster27764-00 | 0 | 1601 | 0 | 0 | 0 |
| SALT00000057947_Cluster27764-00 | 0 | 1623 | 0 | 0 | 0 |
| SALT00000023599_Cluster27764-00 | 0 | 1487 | 0 | 0 | 0 |
| SALT00000040115_Cluster27813-00 | 0 | 1620 | 0 | 0 | 0 |
| SALT00000045275_Cluster27817-00 | 0 | 1620 | 0 | 0 | 0 |
| SALT00000053267_Cluster27823-00 | 0 | 1620 | 0 | 0 | 0 |
| SALT00000010052_Cluster27864-00 | 0 | 1617 | 0 | 0 | 0 |
| SALT00000053588_Cluster27914-00 | 1 | 1615 | 0 | 0 | 0 |
| SALT00000040005_Cluster27933-00 | 1 | 1614 | 0 | 0 | 0 |
| SALT00000036430_Cluster27995-00 | 0 | 1611 | 0 | 0 | 0 |
| SALT00000059095_Cluster28004-00 | 0 | 1611 | 0 | 0 | 0 |
| SALT00000045652_Cluster28089-00 | 0 | 1606 | 0 | 0 | 0 |
| SALT00000055046_Cluster28093-00 | 0 | 1606 | 0 | 0 | 0 |
| SALT00000037905_Cluster28102-00 | 0 | 1605 | 0 | 0 | 0 |
| SALT00000044153_Cluster28106-00 | 0 | 1605 | 0 | 0 | 0 |
| SALT00000010974_Cluster28132-00 | 0 | 1603 | 0 | 0 | 0 |
| SALT00000037319_Cluster28138-00 | 0 | 1603 | 0 | 0 | 0 |
| SALT00000043079_Cluster28140-00 | 0 | 1603 | 0 | 0 | 0 |
| SALT00000049069_Cluster28144-00 | 0 | 1603 | 0 | 0 | 0 |
| SALT00000056746_Cluster28150-00 | 0 | 1603 | 0 | 0 | 0 |
| SALT00000059654_Cluster28152-00 | 0 | 1603 | 0 | 0 | 0 |
| SALT00000061062_Cluster28173-00 | 0 | 1602 | 0 | 0 | 0 |
| SALT00000067911_Cluster28193-00 | 1 | 1601 | 0 | 0 | 0 |
| SALT00000004204_Cluster28209-00 | 1 | 1599 | 0 | 0 | 0 |
| SALT00000048490_Cluster28222-00 | 1 | 1599 | 0 | 0 | 0 |
| SALT00000044500_Cluster28255-00 | 0 | 1597 | 0 | 0 | 0 |
| SALT00000008881_Cluster28267-00 | 1 | 1596 | 0 | 0 | 0 |
| SALT00000005008_Cluster28317-00 | 0 | 1593 | 0 | 0 | 0 |
| SALT00000005019_Cluster28340-00 | 1 | 1592 | 0 | 0 | 0 |
| SALT00000016935_Cluster28347-00 | 0 | 1592 | 0 | 0 | 0 |
| SALT00000053954_Cluster28347-00 | 0 | 1530 | 0 | 0 | 0 |
| SALT00000050147_Cluster28389-00 | 0 | 1590 | 0 | 0 | 0 |
| SALT00000057164_Cluster28409-00 | 1 | 1589 | 0 | 0 | 0 |
| SALT00000057908_Cluster28411-00 | 0 | 1589 | 0 | 0 | 0 |
| SALT00000040535_Cluster28425-00 | 0 | 1588 | 0 | 0 | 0 |
| SALT00000014840_Cluster28454-00 | 0 | 1586 | 0 | 0 | 0 |
| SALT00000002920_Cluster28477-00 | 0 | 1585 | 0 | 0 | 0 |
| SALT00000037126_Cluster28488-00 | 1 | 1585 | 0 | 0 | 0 |
| SALT00000043080_Cluster28489-00 | 0 | 1585 | 0 | 0 | 0 |
| SALT00000044998_Cluster28503-00 | 1 | 1584 | 0 | 0 | 0 |
| SALT00000049380_Cluster28520-00 | 1 | 1583 | 0 | 0 | 0 |
| SALT00000010910_Cluster28536-00 | 0 | 1582 | 0 | 0 | down |

| | | | | | |
|---|---|---|---|---|---|
| SALT00000041250_Cluster28541-00 | 0 | 1582 | 0 | 0 | 0 |
| SALT00000013209_Cluster28550-00 | 1 | 1581 | 0 | 0 | 0 |
| SALT00000044472_Cluster28550-00 | 1 | 1538 | 0 | 0 | 0 |
| SALT00000012119_Cluster28625-00 | 0 | 1577 | 0 | 0 | 0 |
| SALT00000047904_Cluster28655-00 | 1 | 1576 | 0 | 0 | 0 |
| SALT00000048674_Cluster28676-00 | 0 | 1575 | 0 | 0 | 0 |
| SALT00000059687_Cluster28713-00 | 0 | 1573 | 0 | 0 | 0 |
| SALT00000016519_Cluster28724-00 | 1 | 1572 | 0 | 0 | 0 |
| SALT00000055175_Cluster28735-00 | 0 | 1572 | 0 | 0 | 0 |
| SALT00000061356_Cluster28740-00 | 0 | 1572 | 0 | 0 | 0 |
| SALT00000004349_Cluster28758-00 | 1 | 1570 | 0 | 0 | 0 |
| SALT00000012306_Cluster28828-00 | 0 | 1513 | 0 | 0 | down |
| SALT00000020491_Cluster28828-00 | 0 | 1567 | 0 | 0 | 0 |
| SALT00000063603_Cluster28828-00 | 0 | 1482 | 0 | 0 | 0 |
| SALT00000050873_Cluster28837-00 | 0 | 1567 | 0 | 0 | 0 |
| SALT00000060950_Cluster28884-00 | 1 | 1565 | 0 | 0 | 0 |
| SALT00000006158_Cluster28890-00 | 1 | 1564 | 0 | 0 | 0 |
| SALT00000042655_Cluster28906-00 | 1 | 1564 | 0 | 0 | 0 |
| SALT00000046321_Cluster28926-00 | 0 | 1563 | 0 | 0 | down |
| SALT00000053426_Cluster28931-00 | 0 | 1563 | 0 | 0 | 0 |
| SALT00000054026_Cluster28944-00 | 0 | 1562 | 0 | 0 | 0 |
| SALT00000042338_Cluster28958-00 | 0 | 1561 | 0 | 0 | 0 |
| SALT00000008415_Cluster29045-00 | 1 | 1556 | 0 | 0 | 0 |
| SALT00000045395_Cluster29089-00 | 0 | 1554 | 0 | 0 | 0 |
| SALT00000049968_Cluster29187-00 | 0 | 1550 | 0 | 0 | 0 |
| SALT00000055235_Cluster29189-00 | 1 | 1550 | 0 | 0 | 0 |
| SALT00000002849_Cluster29244-00 | 1 | 1546 | 0 | 0 | 0 |
| SALT00000019741_Cluster29248-00 | 0 | 1546 | 0 | 0 | 0 |
| SALT00000052871_Cluster29257-00 | 0 | 1546 | 0 | 0 | 0 |
| SALT00000049840_Cluster29275-00 | 1 | 1545 | 0 | 0 | 0 |
| SALT00000040699_Cluster29311-00 | 0 | 1543 | 0 | 0 | 0 |
| SALT00000044496_Cluster29330-00 | 1 | 1542 | 0 | 0 | 0 |
| SALT00000045233_Cluster29331-00 | 0 | 1542 | 0 | 0 | 0 |
| SALT00000046447_Cluster29332-00 | 0 | 1542 | 0 | 0 | 0 |
| SALT00000059813_Cluster29338-00 | 1 | 1542 | 0 | 0 | 0 |
| SALT00000004079_Cluster29340-00 | 0 | 1541 | 0 | 0 | 0 |
| SALT00000046829_Cluster29419-00 | 0 | 1537 | 0 | 0 | 0 |
| SALT00000049607_Cluster29420-00 | 0 | 1537 | 0 | 0 | 0 |
| SALT00000018175_Cluster29437-00 | 0 | 1536 | 0 | 0 | 0 |
| SALT00000041875_Cluster29443-00 | 0 | 1536 | 0 | 0 | 0 |
| SALT00000053054_Cluster29501-00 | 1 | 1533 | 0 | 0 | 0 |
| SALT00000060897_Cluster29596-00 | 0 | 1527 | 0 | 0 | 0 |
| SALT00000048145_Cluster29639-00 | 0 | 1524 | 0 | 0 | 0 |
| SALT00000057975_Cluster29710-00 | 0 | 1520 | 0 | 0 | down |
| SALT00000012159_Cluster29715-00 | 0 | 1519 | 0 | 0 | 0 |
| SALT00000044711_Cluster29715-00 | 0 | 1448 | 0 | 0 | 0 |

| | | | | | |
|---|---|---|---|---|---|
| SALT00000003246_Cluster29731-00 | 0 | 1518 | 0 | 0 | 0 |
| SALT00000047226_Cluster29737-00 | 0 | 1518 | 0 | 0 | 0 |
| SALT00000049079_Cluster29766-00 | 0 | 1516 | 0 | 0 | 0 |
| SALT00000052703_Cluster29786-00 | 0 | 1515 | 0 | 0 | 0 |
| SALT00000014996_Cluster29787-00 | 0 | 1411 | 0 | 0 | 0 |
| SALT00000053812_Cluster29787-00 | 0 | 1515 | 0 | 0 | 0 |
| SALT00000051771_Cluster29808-00 | 0 | 1514 | 0 | 0 | 0 |
| SALT00000056844_Cluster29817-00 | 0 | 1215 | 0 | 0 | 0 |
| SALT00000019889_Cluster29824-00 | 0 | 1451 | 0 | 0 | 0 |
| SALT00000046470_Cluster29824-00 | 0 | 1513 | 0 | 0 | 0 |
| SALT00000004742_Cluster29856-00 | 0 | 1511 | 0 | 0 | 0 |
| SALT00000014010_Cluster29858-00 | 1 | 1511 | 0 | 0 | 0 |
| SALT00000039725_Cluster29878-00 | 0 | 1510 | 0 | 0 | 0 |
| SALT00000042608_Cluster29879-00 | 0 | 1510 | 0 | 0 | 0 |
| SALT00000003952_Cluster29894-00 | 0 | 1508 | 0 | 0 | 0 |
| SALT00000016309_Cluster29914-00 | 1 | 1507 | 0 | 0 | 0 |
| SALT00000005094_Cluster29939-00 | 1 | 1505 | 0 | 0 | 0 |
| SALT00000056024_Cluster30006-00 | 1 | 1502 | 0 | 0 | 0 |
| SALT00000041808_Cluster30053-00 | 0 | 1498 | 0 | 0 | 0 |
| SALT00000052852_Cluster30076-00 | 0 | 1497 | 0 | 0 | 0 |
| SALT00000054304_Cluster30172-00 | 0 | 1490 | 0 | 0 | 0 |
| SALT00000055648_Cluster30173-00 | 0 | 1490 | 0 | 0 | 0 |
| SALT00000053342_Cluster30187-00 | 1 | 1489 | 0 | 0 | 0 |
| SALT00000015159_Cluster30249-00 | 0 | 1485 | 0 | 0 | 0 |
| SALT00000057297_Cluster30258-00 | 0 | 1485 | 0 | 0 | 0 |
| SALT00000015662_Cluster30286-00 | 0 | 1482 | 0 | 0 | 0 |
| SALT00000037091_Cluster30382-00 | 0 | 1452 | 0 | 0 | 0 |
| SALT00000039322_Cluster30382-00 | 0 | 1476 | 0 | 0 | 0 |
| SALT00000051924_Cluster30386-00 | 1 | 1476 | 0 | 0 | 0 |
| SALT00000052343_Cluster30387-00 | 0 | 1476 | 0 | 0 | 0 |
| SALT00000056941_Cluster30389-00 | 0 | 1476 | 0 | 0 | 0 |
| SALT00000022797_Cluster30407-00 | 0 | 1474 | 0 | 0 | 0 |
| SALT00000046237_Cluster30411-00 | 0 | 1474 | 0 | 0 | 0 |
| SALT00000056929_Cluster30469-00 | 1 | 1470 | 0 | 0 | 0 |
| SALT00000046660_Cluster30513-00 | 0 | 1467 | 0 | 0 | 0 |
| SALT00000003357_Cluster30535-00 | 0 | 1465 | 0 | 0 | 0 |
| SALT00000044859_Cluster30541-00 | 0 | 1465 | 0 | 0 | 0 |
| SALT00000041296_Cluster30549-00 | 0 | 1464 | 0 | up | 0 |
| SALT00000050816_Cluster30549-00 | 0 | 1235 | 0 | 0 | 0 |
| SALT00000014253_Cluster30567-00 | 0 | 1462 | 0 | 0 | 0 |
| SALT00000012101_Cluster30614-00 | 1 | 1459 | 0 | 0 | 0 |
| SALT00000055495_Cluster30629-00 | 0 | 1458 | 0 | 0 | 0 |
| SALT00000004674_Cluster30630-00 | 0 | 1457 | 0 | 0 | 0 |
| SALT00000007549_Cluster30642-00 | 1 | 1456 | 0 | 0 | 0 |
| SALT00000051809_Cluster30649-00 | 1 | 1456 | 0 | 0 | 0 |
| SALT00000030210_Cluster30686-00 | 0 | 1453 | 0 | 0 | 0 |

| | | | | | |
|---|---|---|---|---|---|
| SALT00000045759_Cluster30686-00 | 0 | 1151 | 0 | 0 | 0 |
| SALT00000060540_Cluster30686-00 | 0 | 1185 | 0 | 0 | down |
| SALT00000075729_Cluster30686-00 | 0 | 1184 | down | down | down |
| SALT00000050530_Cluster30704-00 | 0 | 1452 | 0 | 0 | up |
| SALT00000059078_Cluster30727-00 | 1 | 1450 | 0 | 0 | 0 |
| SALT00000038754_Cluster30731-00 | 0 | 1449 | 0 | 0 | 0 |
| SALT00000035810_Cluster30750-00 | 0 | 1447 | 0 | 0 | 0 |
| SALT00000050991_Cluster30758-00 | 1 | 1447 | 0 | 0 | 0 |
| SALT00000009295_Cluster30764-00 | 0 | 1445 | 0 | 0 | 0 |
| SALT00000038724_Cluster30767-00 | 0 | 1445 | 0 | 0 | 0 |
| SALT00000048392_Cluster30790-00 | 0 | 1443 | 0 | 0 | 0 |
| SALT00000015350_Cluster30812-00 | 0 | 1441 | 0 | 0 | 0 |
| SALT00000015915_Cluster30853-00 | 0 | 1436 | 0 | 0 | 0 |
| SALT00000050586_Cluster30896-00 | 0 | 1431 | 0 | 0 | 0 |
| SALT00000014511_Cluster30901-00 | 1 | 1430 | 0 | 0 | 0 |
| SALT00000046999_Cluster30901-00 | 0 | 1189 | 0 | 0 | 0 |
| SALT00000057358_Cluster30909-00 | 1 | 1430 | 0 | 0 | 0 |
| SALT00000018398_Cluster30913-00 | 0 | 1429 | 0 | 0 | 0 |
| SALT00000019779_Cluster30914-00 | 0 | 1429 | 0 | 0 | 0 |
| SALT00000057561_Cluster30950-00 | 0 | 1426 | 0 | 0 | 0 |
| SALT00000052189_Cluster31007-00 | 0 | 1419 | 0 | 0 | 0 |
| SALT00000037858_Cluster31013-00 | 0 | 1418 | 0 | 0 | 0 |
| SALT00000043082_Cluster31107-00 | 1 | 1407 | 0 | 0 | 0 |
| SALT00000057821_Cluster31109-00 | 0 | 1407 | 0 | 0 | 0 |
| SALT00000060389_Cluster31109-00 | 0 | 1269 | 0 | 0 | 0 |
| SALT00000040004_Cluster31114-00 | 0 | 1406 | 0 | 0 | 0 |
| SALT00000061308_Cluster31119-00 | 0 | 1406 | 0 | 0 | 0 |
| SALT00000057626_Cluster31136-00 | 0 | 1403 | 0 | 0 | 0 |
| SALT00000063158_Cluster31156-00 | 0 | 1399 | 0 | 0 | 0 |
| SALT00000036285_Cluster31193-00 | 0 | 1394 | 0 | 0 | 0 |
| SALT00000040527_Cluster31205-00 | 1 | 1392 | 0 | 0 | 0 |
| SALT00000045565_Cluster31207-00 | 0 | 1392 | 0 | 0 | 0 |
| SALT00000044648_Cluster31223-00 | 0 | 1390 | 0 | up | 0 |
| SALT00000043752_Cluster31256-00 | 1 | 1384 | 0 | 0 | 0 |
| SALT00000057256_Cluster31290-00 | 0 | 1379 | 0 | 0 | 0 |
| SALT00000050218_Cluster31329-00 | 0 | 1371 | 0 | 0 | 0 |
| SALT00000061648_Cluster31479-00 | 0 | 1337 | 0 | 0 | 0 |
| SALT00000053349_Cluster31577-00 | 1 | 1313 | 0 | 0 | 0 |
| SALT00000041666_Cluster31669-00 | 0 | 1289 | 0 | 0 | 0 |
| SALT00000016374_Cluster31729-00 | 0 | 1271 | 0 | 0 | 0 |
| SALT00000002468_Cluster31781-00 | 0 | 1257 | 0 | down | down |
| SALT00000003158_Cluster31793-00 | 0 | 1251 | 0 | 0 | 0 |
| SALT00000045392_Cluster31812-00 | 0 | 1246 | 0 | 0 | 0 |
| SALT00000060364_Cluster31861-00 | 0 | 1232 | 0 | 0 | 0 |
| SALT00000010102_Cluster31877-00 | 0 | 1185 | 0 | 0 | 0 |
| SALT00000054784_Cluster31877-00 | 0 | 1228 | 0 | 0 | 0 |

| | | | | | |
|---|---|---|---|---|---|
| SALT00000042737_Cluster31985-00 | 0 | 1193 | 0 | 0 | 0 |
| SALT00000020121_Cluster32033-00 | 0 | 1179 | 0 | 0 | 0 |
| SALT00000081428_Cluster32057-00 | 0 | 1168 | 0 | 0 | 0 |
| SALT00000039250_Cluster32080-00 | 0 | 1160 | 0 | 0 | 0 |
| SALT00000036761_Cluster32092-00 | 0 | 1155 | 0 | 0 | 0 |
| SALT00000039523_Cluster32124-00 | 0 | 1142 | 0 | 0 | 0 |
| SALT00000008378_Cluster32128-00 | 0 | 1141 | 0 | 0 | 0 |
| SALT00000033231_Cluster32132-00 | 0 | 955 | 0 | 0 | 0 |
| SALT00000045563_Cluster32132-00 | 0 | 1140 | 0 | 0 | 0 |
| SALT00000043285_Cluster32132-00 | 0 | 935 | 0 | 0 | 0 |
| SALT00000058431_Cluster32212-00 | 0 | 1108 | 0 | 0 | down |
| SALT00000060597_Cluster32212-00 | 0 | 1107 | 0 | down | down |
| SALT00000086221_Cluster32212-00 | 0 | 1042 | up | 0 | 0 |
| SALT00000001815_Cluster32223-00 | 0 | 1103 | 0 | down | down |
| SALT00000012806_Cluster32242-00 | 0 | 1036 | 0 | 0 | 0 |
| SALT00000013343_Cluster32242-00 | 0 | 1095 | 0 | 0 | up |
| SALT00000029924_Cluster32246-00 | 0 | 1093 | 0 | down | down |
| SALT00000035729_Cluster32246-00 | 0 | 1054 | 0 | 0 | 0 |
| SALT00000018688_Cluster32279-00 | 0 | 882 | 0 | 0 | 0 |
| SALT00000054073_Cluster32279-00 | 0 | 1081 | 0 | 0 | 0 |
| SALT00000057857_Cluster32284-00 | 0 | 1078 | 0 | 0 | 0 |
| SALT00000013746_Cluster32305-00 | 0 | 1064 | 0 | 0 | down |
| SALT00000052108_Cluster32305-00 | 0 | 1058 | 0 | 0 | down |
| SALT00000040103_Cluster32310-00 | 1 | 1062 | 0 | 0 | 0 |
| SALT00000049660_Cluster32312-00 | 1 | 1062 | 0 | 0 | 0 |
| SALT00000018787_Cluster32317-00 | 0 | 1054 | 0 | 0 | 0 |
| SALT00000051729_Cluster32317-00 | 0 | 1057 | 0 | 0 | down |
| SALT00000058142_Cluster32318-00 | 0 | 1056 | 0 | 0 | 0 |
| SALT00000065147_Cluster32339-00 | 0 | 1045 | 0 | 0 | 0 |
| SALT00000021834_Cluster32375-00 | 0 | 1020 | 0 | 0 | up |
| SALT00000067105_Cluster32375-00 | 0 | 962 | 0 | 0 | 0 |
| SALT00000066906_Cluster32398-00 | 0 | 1004 | 0 | 0 | 0 |
| SALT00000037661_Cluster32415-00 | 0 | 987 | 0 | 0 | down |
| SALT00000077438_Cluster32443-00 | 0 | 968 | 0 | 0 | down |
| SALT00000046838_Cluster32451-00 | 0 | 961 | 0 | 0 | 0 |
| SALT00000060729_Cluster32482-00 | 0 | 927 | 0 | up | 0 |
| SALT00000055602_Cluster32491-00 | 0 | 913 | 0 | 0 | 0 |
| SALT00000005983_Cluster32509-00 | 0 | 889 | 0 | 0 | 0 |
| SALT00000001762_Cluster32518-00 | 1 | 865 | 0 | 0 | 0 |
| SALT00000037430_Cluster32524-00 | 0 | 852 | 0 | 0 | 0 |
| SALT00000063036_Cluster32577-00 | 1 | 674 | 0 | 0 | 0 |
| SALT00000041479_Cluster32585-00 | 0 | 611 | 0 | 0 | 0 |
| SALT00000079288_Cluster32594-00 | 1 | 549 | 0 | 0 | 0 |
| SALT00000054269_Cluster32599-00 | 0 | 526 | 0 | 0 | 0 |
