## Supplemental Table 7 for "The full-length transcriptome of *Spartina alterniflora* reveals the complexity of high salt tolerance in monocotyledonous halophyte"

### Table S7. List of salt stress-responsive unigenes in *Spartina alterniflora.*

FC: fold change; Differential expression cutoff: FDR<0.05 and |log2FC| ≥ 1

| Cluster id | 350mM-Control | 500mM-Control | 800mM-Control | 500mM-350mM | 800mM-350mM | 800mM-500mM | annotation | Unigene id |
|---|---|---|---|---|---|---|---|---|
| Cluster100 | - | - | down | - | down | down | ankyrin repeat SKIP35-like [C | SALT00000076165_Cluster100-001 |
| Cluster10004 | - | - | up | - | up | up | probable serine threonine- kin | SALT00000077217_Cluster10004-005 |
| Cluster10012 | - | up | - | - | - | down | phosphatase 2C isoform gamr | SALT00000082631_Cluster10012-004 |
| Cluster10013 | - | - | up | - | up | up | potassium transporter 10 [Zea | SALT00000022902_Cluster10013-002 |
| Cluster1003 | - | - | - | - | up | - | Os02g0748100 [Oryza sativa | SALT00000089107_Cluster1003-007 |
| Cluster10040 | - | - | down | - | - | - | Os04g0671900 [Oryza sativa | SALT00000070488_Cluster10040-002 |
| Cluster10045 | - | - | - | - | down | down | TPA: DUF604-domain contai | SALT00000042590_Cluster10045-001 |
| Cluster10047 | - | - | up | - | - | - | Os06g0635200 [Oryza sativa | SALT00000069263_Cluster10047-001 |
| Cluster10055 | - | - | down | - | down | - | probable mediator of RNA po | SALT00000049775_Cluster10055-007 |
| Cluster10058 | - | - | down | - | - | - | glycosyltransferase 10 [Oryza | SALT00000069308_Cluster10058-004 |
| Cluster10059 | - | - | - | - | down | - | hypothetical protein SORBID | SALT00000069993_Cluster10059-001 |
| Cluster10072 | - | - | down | - | down | down | Lipoxygenase chloroplastic | SALT00000022101_Cluster10072-001 |
| Cluster10076 | - | - | up | - | - | - | partial [Oryza sativa Japonica | SALT00000068613_Cluster10076-002 |
| Cluster10085 | - | - | down | - | - | - | BEL1-type homeodomain | SALT00000072071_Cluster10085-009 |
| Cluster10087 | - | - | up | - | - | - | delta 1-pyrroline-5-carboxylat | SALT00000075096_Cluster10087-004 |
| Cluster10112 | - | - | - | - | - | down | PHD finger [Zea mays] | SALT00000065616_Cluster10112-001 |
| Cluster10130 | - | - | down | - | down | down | Os04g0504800 [Oryza sativa | SALT00000063374_Cluster10130-001 |
| Cluster1014 | - | - | down | - | down | down | pumilio homolog 1-like [Elaei | SALT00000085591_Cluster1014-021 |
| Cluster10141 | down | down | - | - | - | - | zinc finger STOP1 homolog [ | SALT00000073062_Cluster10141-024 |
| Cluster10146 | - | - | up | - | up | up | Disease resistance RPS2 | SALT00000026236_Cluster10146-001 |
| Cluster10168 | - | - | up | - | up | up | probable L-type lectin-domair | SALT00000064049_Cluster10168-005 |
| Cluster10175 | - | up | up | - | - | - | NADP-specific isocitrate deh | SALT00000035989_Cluster10175-002 |
| Cluster10184 | - | - | - | - | down | down | nitrate reductase [NAD(P)H] | | SALT00000065714_Cluster10184-001 |
| Cluster10187 | - | - | - | - | up | - | Os02g0787600 [Oryza sativa | SALT00000073899_Cluster10187-001 |
| Cluster10205 | - | - | down | - | - | - | Os03g0271200 [Oryza sativa | SALT00000071497_Cluster10205-009 |
| Cluster10210 | - | up | up | - | - | - | disulfide isomerase | SALT00000063401_Cluster10210-001 |
| Cluster10215 | - | - | - | - | down | - | probable alpha-glucosidase O | SALT00000082076_Cluster10215-011 |
| Cluster10223 | - | - | down | - | - | - | ribonuclease H At1g65750 | SALT00000082769_Cluster10223-002 |
| Cluster10225 | - | - | down | - | down | down | Os01g0801600 [Oryza sativa | SALT00000061367_Cluster10225-001 |
| Cluster1025 | - | - | down | - | down | - | disease resistance RPM1-like | SALT00000079966_Cluster1025-008 |
| Cluster10253 | - | - | up | - | - | up | cytochrome P450 superfamily | SALT00000065872_Cluster10253-011 |
| Cluster10261 | - | - | up | - | up | - | Os03g0122000 [Oryza sativa | SALT00000063262_Cluster10261-003 |
| Cluster10264 | - | - | up | - | - | up | GAMYB-binding [Oryza sati | SALT00000071915_Cluster10264-003 |
| Cluster10269 | - | - | up | - | up | - | ribulose-1,5-bisphosphate carl | SALT00000049812_Cluster10269-025 |
| Cluster10276 | down | - | - | - | - | - | probable inactive poly [ADP- | SALT00000065838_Cluster10276-002 |
| Cluster1028 | - | up | up | - | up | up | eukaryotic translation initiatio | SALT00000062033_Cluster1028-030 |
| Cluster1029 | - | up | up | - | up | up | Os11g0603200 [Oryza sativa | SALT00000064437_Cluster1029-001 |
| Cluster10299 | - | - | down | - | - | - | pentatricopeptide repeat-conta | SALT00000024630_Cluster10299-001 |
| Cluster103 | - | - | - | - | down | - | phosphoenolpyruvate carboxy | SALT00000078295_Cluster103-001 |
| Cluster1030 | - | - | - | - | down | - | Adaptin N terminal [Oryza sat | SALT00000034414_Cluster1030-014 |
| Cluster10305 | - | - | down | - | - | - | hyperosmolality-gated Ca2+ p | SALT00000072270_Cluster10305-001 |
| Cluster10314 | - | - | - | - | - | up | DEAD-box ATP-dependent R | SALT00000069200_Cluster10314-005 |
| Cluster10331 | - | up | - | up | - | down | CEN 2 | SALT00000066392_Cluster10331-001 |
| Cluster10334 | - | - | down | - | - | - | conserved oligomeric Golgi c | SALT00000079491_Cluster10334-009 |
| Cluster10335 | - | - | - | - | down | - | hypothetical protein SORBID | SALT00000081422_Cluster10335-001 |
| Cluster10341 | - | - | down | - | down | down | small RNA degrading nucleas | SALT00000072964_Cluster10341-002 |
| Cluster10343 | - | - | down | - | - | down | purple acid phosphatase precu | SALT00000080786_Cluster10343-002 |
| Cluster10346 | - | down | down | down | down | down | TPA: tubulin alpha-1 chain [Z | SALT00000061101_Cluster10346-001 |
| Cluster10351 | - | - | up | - | up | - | fructose-bisphosphate aldolas | SALT00000045405_Cluster10351-002 |

| | | | | | | | | |
|---|---|---|---|---|---|---|---|---|
| Cluster10361 | up | - | up | - | up | up | phenylalanine ammonia-lyase | SALT00000079490_Cluster10361-003 |
| Cluster10367 | - | - | down | - | - | - | TPA: bZIP transcription factor | SALT00000046442_Cluster10367-001 |
| Cluster10369 | - | - | up | - | up | up | PREDICTED: uncharacterized | SALT00000079568_Cluster10369-001 |
| Cluster10375 | - | - | down | - | down | - | transmembrane 136-like [Seta | SALT00000069831_Cluster10375-001 |
| Cluster1038 | - | - | - | - | up | - | homeodomain-like transcriptio | SALT00000077748_Cluster1038-006 |
| Cluster104 | - | - | down | - | down | down | clathrin heavy chain 1-like | SALT00000092937_Cluster104-001 |
| Cluster10407 | - | up | up | - | up | up | Os10g0463800 [Oryza sativa | SALT00000052705_Cluster10407-002 |
| Cluster10432 | - | - | up | - | up | up | phosphate transporter [Triticu | SALT00000058162_Cluster10432-009 |
| Cluster10439 | - | - | - | - | - | down | RNA polymerase II C-termina | SALT00000078818_Cluster10439-012 |
| Cluster1045 | - | - | up | - | up | - | [Hordeum vulgare vulgare] | SALT00000083474_Cluster1045-002 |
| Cluster10451 | - | - | up | - | - | - | RCH2 [Oryza sativa Japonica | SALT00000078973_Cluster10451-002 |
| Cluster10455 | - | - | up | - | up | up | Os12g0406100 [Oryza sativa | SALT00000078084_Cluster10455-008 |
| Cluster10475 | - | - | - | - | down | - | probable 1-deoxy-D-xylulose- | SALT00000063147_Cluster10475-004 |
| Cluster10478 | - | - | down | - | down | - | pentatricopeptide repeat-conta | SALT00000074200_Cluster10478-002 |
| Cluster1048 | - | - | up | - | up | - | stress inducible [Zea mays] | SALT00000042979_Cluster1048-001 |
| Cluster10493 | - | - | - | - | - | down | DNA-repair XRCC1 [Oryza s | SALT00000068847_Cluster10493-001 |
| Cluster10495 | - | - | - | - | up | - | cytokinin-O-glucosyltransfera | SALT00000073805_Cluster10495-004 |
| Cluster10512 | - | - | - | - | - | up | Os10g0561200 [Oryza sativa | SALT00000075140_Cluster10512-002 |
| Cluster10528 | - | - | down | - | down | - | transposase Tam3 | SALT00000070514_Cluster10528-001 |
| Cluster10547 | down | - | down | - | - | - | TPA: DUF594 domain contai | SALT00000064843_Cluster10547-002 |
| Cluster1057 | - | - | - | - | - | up | RETICULATA-RELATED ch | SALT00000023464_Cluster1057-005 |
| Cluster10587 | - | - | - | - | up | - | lysine-specific demethylase JI | SALT00000024028_Cluster10587-002 |
| Cluster10598 | - | - | up | - | up | up | Os10g0501500 [Oryza sativa | SALT00000024251_Cluster10598-001 |
| Cluster10599 | - | - | up | - | - | up | lung seven transmembrane rec | SALT00000043261_Cluster10599-001 |
| Cluster10601 | - | - | up | - | up | up | K(+) efflux antiporter chlorop | SALT00000071192_Cluster10601-002 |
| Cluster1061 | - | - | up | - | up | up | fatty acid elongase [Zea mays | SALT00000091658_Cluster1061-003 |
| Cluster10611 | down | down | down | - | - | - | F-box kelch-repeat At5g6057( | SALT00000022843_Cluster10611-009 |
| Cluster10637 | - | - | - | - | up | - | Os03g0726500 [Oryza sativa | SALT00000071620_Cluster10637-001 |
| Cluster1064 | - | - | - | - | down | - | phospholipase D family [Zea | SALT00000084399_Cluster1064-001 |
| Cluster10641 | - | - | down | - | - | - | kDa vesicle transport [Zea ma | SALT00000079379_Cluster10641-002 |
| Cluster10700 | - | - | up | - | up | up | probable receptor-like serine t | SALT00000066513_Cluster10700-008 |
| Cluster10711 | - | - | - | - | - | up | sugar carrier C-like | SALT00000038923_Cluster10711-001 |
| Cluster10726 | - | up | - | - | - | - | GTPase [Oryza sativa Japonic | SALT00000072607_Cluster10726-001 |
| Cluster10734 | - | - | - | - | down | - | NADH partial | SALT00000027597_Cluster10734-002 |
| Cluster10738 | - | - | - | - | down | - | ECERIFERUM 26-like [Setar | SALT00000063075_Cluster10738-001 |
| Cluster1074 | - | - | up | - | - | - | Os07g0672500 [Oryza sativa | SALT00000080266_Cluster1074-007 |
| Cluster10746 | - | - | down | - | - | - | Os02g0826100 [Oryza sativa | SALT00000062770_Cluster10746-002 |
| Cluster1076 | - | - | up | - | up | - | partial [Oryza sativa Japonica | SALT00000077947_Cluster1076-001 |
| Cluster10801 | - | down | down | - | down | down | ring finger 126 isoform 1 [Ory | SALT00000080942_Cluster10801-004 |
| Cluster10803 | - | - | down | - | down | - | TBC1 domain family member | SALT00000021481_Cluster10803-001 |
| Cluster10814 | - | up | down | - | down | down | serine carboxypeptidase precu | SALT00000079326_Cluster10814-007 |
| Cluster10825 | - | - | - | - | up | - | actin-depolymerizing factor 6 | SALT00000067823_Cluster10825-002 |
| Cluster10833 | - | down | - | - | - | - | Os01g0140100 [Oryza sativa | SALT00000058069_Cluster10833-013 |
| Cluster10839 | - | - | down | - | - | down | Os09g0553900 [Oryza sativa | SALT00000073677_Cluster10839-005 |
| Cluster10840 | - | - | down | - | down | down | phosphatidylinositol 4-phosph | SALT00000077282_Cluster10840-001 |
| Cluster10849 | - | - | - | down | down | - | G-type lectin S-receptor-like s | SALT00000065032_Cluster10849-001 |
| Cluster1086 | - | - | - | - | - | up | uncharacterized ATP-depende | SALT00000083152_Cluster1086-003 |
| Cluster10888 | - | - | up | - | up | up | serine threonine- kinase CCR4 | SALT00000062938_Cluster10888-006 |
| Cluster10916 | - | - | down | - | - | - | potassium channel AKT2 3 [Z | SALT00000080543_Cluster10916-002 |
| Cluster10918 | - | - | up | - | up | up | probable methyltransferase PM | SALT00000067271_Cluster10918-002 |
| Cluster10921 | - | down | down | - | down | - | mitochondrial phosphate trans | SALT00000044320_Cluster10921-001 |
| Cluster10934 | - | - | up | - | up | up | ran-binding 1 [Zea mays] | SALT00000068457_Cluster10934-001 |
| Cluster10949 | - | up | - | - | - | - | probable serine threonine- kin | SALT00000073471_Cluster10949-004 |
| Cluster10963 | - | - | up | - | up | up | partial [Oryza sativa Japonica | SALT00000083436_Cluster10963-021 |
| Cluster10967 | - | - | - | - | down | down | subtilisin-like protease | SALT00000071666_Cluster10967-006 |
| Cluster1098 | - | - | down | - | down | down | PREDICTED: uncharacterized | SALT00000090541_Cluster1098-001 |
| Cluster10981 | - | - | down | - | - | - | protease 2 [Zea mays] | SALT00000080412_Cluster10981-001 |
| Cluster10983 | - | - | down | - | - | - | early endosome antigen 1-like | SALT00000029713_Cluster10983-001 |

| | | | | | | | | |
|---|---|---|---|---|---|---|---|---|
| Cluster10999 | - | - | up | - | up | up | kinesin-4 [Zea mays] | SALT00000062268_Cluster10999-001 |
| Cluster11 | - | - | down | - | down | down | LONGIFOLIA 2-like | SALT00000093398_Cluster11-001 |
| Cluster1100 | - | - | down | - | down | down | [Hordeum vulgare vulgare] | SALT00000054765_Cluster1100-001 |
| Cluster11005 | - | - | down | - | - | - | elongation factor 1-beta-like [ | SALT00000082833_Cluster11005-001 |
| Cluster11006 | - | - | up | - | - | - | F-box kelch-repeat SKIP11 [C | SALT00000084022_Cluster11006-011 |
| Cluster11017 | - | - | up | - | up | up | receptor kinase [Oryza sativa | SALT00000028298_Cluster11017-003 |
| Cluster11019 | - | - | - | - | - | down | splicing factor 3B subunit 4 | SALT00000065842_Cluster11019-004 |
| Cluster11030 | - | - | up | - | - | - | ultraviolet-B receptor UVR8 [ | SALT00000043065_Cluster11030-001 |
| Cluster11042 | - | - | up | - | - | - | GATA transcription factor fan | SALT00000001230_Cluster11042-001 |
| Cluster11067 | - | - | down | - | - | - | Os02g0102900 [Oryza sativa | SALT00000069821_Cluster11067-006 |
| Cluster11092 | - | - | down | - | - | - | PREDICTED: LOW QUALIT | SALT00000075866_Cluster11092-004 |
| Cluster11096 | - | - | up | - | - | - | pre-mRNA-splicing factor 38l | SALT00000029292_Cluster11096-002 |
| Cluster11109 | - | - | up | - | up | up | dnaJ subfamily B member 5 [ | SALT00000065448_Cluster11109-008 |
| Cluster11126 | - | - | up | - | up | up | endo-1,3(4)-beta-glucanase 2 | SALT00000074061_Cluster11126-008 |
| Cluster11155 | - | - | up | - | - | - | WD and tetratricopeptide repe | SALT00000022191_Cluster11155-005 |
| Cluster11171 | - | up | up | - | up | up | Os09g0511600 [Oryza sativa | SALT00000038710_Cluster11171-012 |
| Cluster11178 | - | - | - | - | up | - | PREDICTED: uncharacterize | SALT00000084247_Cluster11178-002 |
| Cluster11224 | - | - | up | - | up | up | [Hordeum vulgare vulgare] | SALT00000077792_Cluster11224-008 |
| Cluster11258 | - | - | - | - | down | - | Os11g0592600 [Oryza sativa | SALT00000073339_Cluster11258-007 |
| Cluster11289 | - | - | down | - | down | down | triose phosphate phosphate tra | SALT00000038573_Cluster11289-005 |
| Cluster113 | - | - | up | - | up | up | ubiquitin-activating enzyme E | SALT00000002890_Cluster113-001 |
| Cluster11308 | - | - | down | - | down | down | uncharacterized LOC1002793 | SALT00000061507_Cluster11308-001 |
| Cluster11315 | - | - | up | - | - | - | Os03g0281500 [Oryza sativa | SALT00000031734_Cluster11315-003 |
| Cluster11316 | - | - | down | - | down | down | glyceraldehyde-3-phosphate d | SALT00000055401_Cluster11316-022 |
| Cluster11318 | - | - | up | - | up | up | G-type lectin S-receptor-like s | SALT00000063314_Cluster11318-001 |
| Cluster1133 | - | up | up | up | up | up | photosystem I P700 apo A2 (c | SALT00000064705_Cluster1133-009 |
| Cluster11336 | - | - | down | - | - | - | PREDICTED: uncharacterize | SALT00000066602_Cluster11336-005 |
| Cluster1135 | - | - | up | - | up | up | telomere repeat-binding 5 isof | SALT00000022273_Cluster1135-008 |
| Cluster11350 | down | - | down | - | - | - | Arf GTPase activating | SALT00000027547_Cluster11350-006 |
| Cluster11352 | - | up | up | - | - | - | iron inhibited ABC transporte | SALT00000054629_Cluster11352-001 |
| Cluster11357 | - | - | down | - | down | - | Os10g0567400 [Oryza sativa | SALT00000079784_Cluster11357-001 |
| Cluster1139 | - | - | up | - | - | - | Os07g0672500 [Oryza sativa | SALT00000093019_Cluster1139-017 |
| Cluster11393 | - | - | down | - | down | - | Ulp1 protease C-terminal cata | SALT00000066817_Cluster11393-004 |
| Cluster11398 | - | - | up | - | up | up | GATA transcription factor fan | SALT00000063018_Cluster11398-001 |
| Cluster11405 | - | - | up | - | up | - | PREDICTED: uncharacterize | SALT00000079848_Cluster11405-004 |
| Cluster11443 | - | up | up | - | - | - | AP2 EREBP partial | SALT00000055767_Cluster11443-001 |
| Cluster11444 | - | - | down | - | - | - | probable mediator of RNA po | SALT00000042609_Cluster11444-001 |
| Cluster11457 | - | down | down | - | - | - | trihelix transcription factor G | SALT00000083171_Cluster11457-006 |
| Cluster11460 | up | up | - | - | - | - | partial [Oryza sativa Japonica | SALT00000070597_Cluster11460-001 |
| Cluster11479 | - | - | down | - | - | - | U3 small nucleolar RNA-asso | SALT00000062896_Cluster11479-003 |
| Cluster1148 | - | - | down | - | - | down | partial [Oryza sativa Japonica | SALT00000031958_Cluster1148-003 |
| Cluster1149 | - | - | down | - | down | - | photosystem I P700 apo A1 (c | SALT00000093493_Cluster1149-001 |
| Cluster11490 | - | - | up | - | - | - | pathogen induced 2-4 [Zea ma | SALT00000062195_Cluster11490-001 |
| Cluster11491 | up | up | up | - | down | down | [Hordeum vulgare vulgare] | SALT00000068072_Cluster11491-002 |
| Cluster11510 | - | - | up | - | up | - | Os01g0585600 [Oryza sativa | SALT00000076623_Cluster11510-006 |
| Cluster11514 | - | - | up | - | - | up | AMP-binding [Zea mays] | SALT00000029233_Cluster11514-005 |
| Cluster11517 | down | down | down | - | up | up | lipoxygenase [Zea mays] | SALT00000068949_Cluster11517-007 |
| Cluster11518 | - | - | up | - | - | - | -tyrosine-phosphatase MKP1 | SALT00000071718_Cluster11518-003 |
| Cluster11524 | up | up | - | - | - | - | rhodanese-like domain contai | SALT00000081871_Cluster11524-001 |
| Cluster11535 | - | - | up | - | up | - | CYPRO4 [Zea mays] | SALT00000077108_Cluster11535-006 |
| Cluster11539 | - | - | up | - | up | up | TPA: phospholipase D family | SALT00000063118_Cluster11539-004 |
| Cluster1155 | - | - | - | - | up | up | probable alpha,alpha-trehalos | SALT00000086020_Cluster1155-037 |
| Cluster11554 | - | - | - | - | down | - | Os01g0541900 [Oryza sativa | SALT00000069033_Cluster11554-001 |
| Cluster11561 | up | up | up | - | - | - | [Hordeum vulgare vulgare] | SALT00000043731_Cluster11561-009 |
| Cluster11562 | - | - | up | - | - | - | Os06g0665500 [Oryza sativa | SALT00000044361_Cluster11562-005 |
| Cluster11564 | - | - | up | - | up | up | VTC2 [Zea mays] | SALT00000066357_Cluster11564-001 |
| Cluster11567 | - | - | - | - | - | down | stromal 70 kDa heat shock-rel | SALT00000005966_Cluster11567-001 |
| Cluster1157 | - | - | down | - | - | - | retrotransposon expressed [Or | SALT00000090015_Cluster1157-001 |

| | | | | | | | | |
|---|---|---|---|---|---|---|---|---|
| Cluster11571 | - | - | up | - | up | up | U-box domain-containing 44- | SALT00000063151_Cluster11571-004 |
| Cluster11572 | - | - | down | - | down | - | 60S ribosomal L22-2 [Zea ma | SALT00000067347_Cluster11572-001 |
| Cluster11584 | - | - | down | - | down | - | COP1-interacting 7 (CIP7) [O | SALT00000075615_Cluster11584-001 |
| Cluster11590 | - | down | down | - | - | - | Os03g0245700 [Oryza sativa | SALT00000051854_Cluster11590-008 |
| Cluster11595 | - | - | up | - | up | - | Os12g0618300 [Oryza sativa | SALT00000070203_Cluster11595-004 |
| Cluster1160 | - | - | - | - | up | - | OBERON 4 [Oryza sativa Jap | SALT00000078567_Cluster1160-008 |
| Cluster11607 | - | up | up | - | - | - | cell wall-associated hydrolase | SALT00000081958_Cluster11607-004 |
| Cluster11610 | - | up | up | - | - | - | ppi-phosphofructokinase [Zea | SALT00000010024_Cluster11610-002 |
| Cluster11617 | - | - | - | - | up | - | MYB DNA-binding domain s | SALT00000078018_Cluster11617-009 |
| Cluster1163 | - | - | up | - | - | - | Os02g0216300 [Oryza sativa | SALT00000092279_Cluster1163-001 |
| Cluster11631 | - | - | up | - | up | up | endoplasmin homolog | SALT00000027691_Cluster11631-002 |
| Cluster11637 | - | - | - | - | up | up | Os03g0770800 [Oryza sativa | SALT00000081580_Cluster11637-016 |
| Cluster11638 | - | - | up | - | up | - | multicystatin precursor [Zea n | SALT00000083816_Cluster11638-001 |
| Cluster11640 | up | up | up | - | - | - | carbamoyl-phosphate synthas | SALT00000043322_Cluster11640-010 |
| Cluster11647 | - | - | down | - | - | - | Os12g0566000 [Oryza sativa | SALT00000084441_Cluster11647-009 |
| Cluster11648 | - | up | up | up | up | - | [Hordeum vulgare vulgare] | SALT00000061099_Cluster11648-010 |
| Cluster11652 | - | - | up | - | up | up | Os11g0531700 [Oryza sativa | SALT00000076388_Cluster11652-001 |
| Cluster11661 | - | - | down | - | - | - | Sn1-specific diacylglycerol li | SALT00000079683_Cluster11661-002 |
| Cluster11662 | - | - | up | - | up | up | alkaline alpha galactosidase 2 | SALT00000023258_Cluster11662-003 |
| Cluster11668 | - | up | up | - | up | up | mitochondrial 2-oxoglutarate | SALT00000007879_Cluster11668-004 |
| Cluster11681 | - | down | down | - | - | - | trihelix transcription factor G | SALT00000023165_Cluster11681-001 |
| Cluster11692 | - | - | up | - | up | up | probable alpha,alpha-trehalos | SALT00000083016_Cluster11692-007 |
| Cluster11695 | - | down | - | down | - | up | beta-galactosidase isoform 1 [ | SALT00000070431_Cluster11695-011 |
| Cluster11698 | - | - | up | - | up | up | [Hordeum vulgare vulgare] | SALT00000078724_Cluster11698-002 |
| Cluster11723 | - | - | up | - | up | up | lipoxygenase [Zea mays] | SALT00000027238_Cluster11723-001 |
| Cluster11724 | - | - | down | down | down | down | transmembrane 9 superfamily | SALT00000048038_Cluster11724-002 |
| Cluster11725 | - | - | up | - | - | - | beta-amylase chloroplastic [S | SALT00000062708_Cluster11725-002 |
| Cluster11727 | - | - | down | - | - | - | Double-stranded RNA binding | SALT00000066153_Cluster11727-002 |
| Cluster11728 | - | - | up | - | - | - | Os05g0491200 [Oryza sativa | SALT00000068474_Cluster11728-001 |
| Cluster11739 | - | - | - | - | - | down | Os01g0251100 [Oryza sativa | SALT00000077713_Cluster11739-001 |
| Cluster11754 | - | - | down | down | down | down | Os03g0150800 [Oryza sativa | SALT00000054074_Cluster11754-003 |
| Cluster11774 | - | - | up | - | up | - | probable GTP diphosphokinas | SALT00000072419_Cluster11774-002 |
| Cluster11786 | - | up | up | - | - | - | enolase 2 [Setaria italica] | SALT00000045824_Cluster11786-001 |
| Cluster11813 | - | - | up | - | up | up | FAR1-RELATED SEQUENC | SALT00000071657_Cluster11813-001 |
| Cluster11827 | - | - | down | - | down | - | DAMAGED DNA-BINDING | SALT00000078937_Cluster11827-002 |
| Cluster1183 | - | - | down | - | down | down | [Hordeum vulgare vulgare] | SALT00000077050_Cluster1183-010 |
| Cluster11860 | - | - | up | - | - | - | membrane [Oryza sativa Japo | SALT00000065178_Cluster11860-003 |
| Cluster11861 | - | - | down | - | - | - | retrotransposon unclassified | SALT00000066339_Cluster11861-002 |
| Cluster11864 | - | - | down | - | - | - | Os04g0118700 [Oryza sativa | SALT00000074929_Cluster11864-004 |
| Cluster11865 | - | down | - | - | - | - | endoplasmin precursor [Zea n | SALT00000075215_Cluster11865-002 |
| Cluster11873 | - | down | - | - | - | up | beta-galactosidase isoform 1 [ | SALT00000071685_Cluster11873-004 |
| Cluster11879 | - | up | up | - | - | - | sucrose synthase | SALT00000035061_Cluster11879-004 |
| Cluster11901 | - | - | down | - | - | - | Os06g0294000 [Oryza sativa | SALT00000026654_Cluster11901-002 |
| Cluster11904 | - | - | - | - | - | down | chloroplastic group IIB intron | SALT00000064035_Cluster11904-001 |
| Cluster11921 | - | - | - | - | down | - | calponin homology domain-c | SALT00000070843_Cluster11921-001 |
| Cluster11925 | - | - | - | - | - | down | thioredoxin peroxidase [Oryz | SALT00000075891_Cluster11925-001 |
| Cluster11928 | - | - | up | - | - | - | Leucine Rich Repeat family e | SALT00000068942_Cluster11928-002 |
| Cluster11935 | - | - | up | - | up | up | Os09g0363500 [Oryza sativa | SALT00000058102_Cluster11935-006 |
| Cluster11942 | - | - | - | - | up | - | probable cyclic nucleotide-ga | SALT00000078831_Cluster11942-009 |
| Cluster11944 | - | up | up | - | up | - | laccase 1 [Zea mays] | SALT00000083985_Cluster11944-008 |
| Cluster11953 | - | - | - | - | up | - | oxysterol-binding -related 3C | SALT00000080230_Cluster11953-020 |
| Cluster1196 | - | - | - | - | down | down | catalase [Saccharum hybrid c | SALT00000052030_Cluster1196-001 |
| Cluster11960 | - | - | down | - | - | - | [Hordeum vulgare vulgare] | SALT00000067646_Cluster11960-003 |
| Cluster11962 | - | - | up | - | up | - | receptor-like serine threonine- | SALT00000073643_Cluster11962-001 |
| Cluster11964 | - | - | up | - | up | up | methylmalonate-semialdehyd | SALT00000075031_Cluster11964-014 |
| Cluster11966 | - | up | up | - | up | up | major facilitator superfamily | SALT00000026453_Cluster11966-002 |
| Cluster11972 | up | up | up | - | up | - | long chain acyl- synthetase ch | SALT00000073379_Cluster11972-002 |
| Cluster11980 | - | - | up | - | - | - | ---NA--- | SALT00000068076_Cluster11980-001 |

| | | | | | | | | |
|---|---|---|---|---|---|---|---|---|
| Cluster11981 | - | - | - | - | down | down | Os01g0769900 [Oryza sativa | SALT00000069179_Cluster11981-005 |
| Cluster11983 | - | - | down | - | - | - | SCARECROW [Zea mays] | SALT00000079364_Cluster11983-006 |
| Cluster12009 | - | - | down | - | down | down | 40S ribosomal S7-like [Oryza | SALT00000069169_Cluster12009-002 |
| Cluster12014 | - | - | - | - | - | down | UDP-glucuronate:xylan alpha | SALT00000083697_Cluster12014-002 |
| Cluster12035 | - | - | up | - | up | - | U-box domain-containing 52- | SALT00000071845_Cluster12035-001 |
| Cluster12036 | - | - | down | - | down | - | elongation factor G- chloropla | SALT00000075833_Cluster12036-001 |
| Cluster12045 | - | down | down | down | down | down | Os12g0431300 [Oryza sativa | SALT00000064854_Cluster12045-002 |
| Cluster12060 | - | - | - | - | - | down | pentatricopeptide repeat10 [Ze | SALT00000076130_Cluster12060-002 |
| Cluster12061 | - | - | up | - | - | up | Os01g0784200 [Oryza sativa | SALT00000076157_Cluster12061-003 |
| Cluster12075 | - | - | - | - | up | - | glycerophosphodiester phospl | SALT00000062291_Cluster12075-008 |
| Cluster12082 | - | - | up | - | - | - | ABC transporter subunit-like | SALT00000062225_Cluster12082-002 |
| Cluster12086 | - | - | up | - | - | - | hexaubiquitin [Arabidopsis ly | SALT00000071308_Cluster12086-003 |
| Cluster12095 | - | up | up | - | up | up | pentatricopeptide repeat-conta | SALT00000082217_Cluster12095-001 |
| Cluster12100 | - | - | down | - | - | - | BEL1-type homeodomain | SALT00000065506_Cluster12100-002 |
| Cluster12104 | - | - | down | - | - | - | AF466646_7 poly | SALT00000071609_Cluster12104-001 |
| Cluster1211 | - | - | down | - | down | down | plasma membrane ATPase-lik | SALT00000074269_Cluster1211-001 |
| Cluster12113 | up | - | - | - | down | down | Os06g0472400 [Oryza sativa | SALT00000070177_Cluster12113-001 |
| Cluster1212 | - | - | up | - | up | up | Os05g0363200 [Oryza sativa | SALT00000055879_Cluster1212-043 |
| Cluster12120 | - | - | - | - | up | - | methylcrotonoyl- carboxylase | SALT00000027842_Cluster12120-007 |
| Cluster12122 | - | - | up | - | up | up | Os10g0501500 [Oryza sativa | SALT00000068768_Cluster12122-002 |
| Cluster12141 | down | - | down | - | - | - | glyceraldehyde-3-phosphate d | SALT00000050185_Cluster12141-006 |
| Cluster12142 | - | - | up | - | - | - | dol-P-Man:Man(5) c(2)-PP-D | SALT00000063468_Cluster12142-003 |
| Cluster12149 | - | - | down | - | down | down | ATP synthase CF0 A subunit ( | SALT00000027186_Cluster12149-002 |
| Cluster12159 | - | - | - | - | up | - | ketohexokinase-like isoform X | SALT00000076530_Cluster12159-002 |
| Cluster12177 | - | down | down | - | - | - | shaggy-related kinase kappa-l | SALT00000076259_Cluster12177-001 |
| Cluster122 | - | - | up | - | up | - | retrotransposon unclassified | SALT00000089339_Cluster122-006 |
| Cluster1220 | - | up | up | - | up | up | xylosyltransferase 1-like | SALT00000048028_Cluster1220-005 |
| Cluster12208 | - | - | up | - | - | up | peroxisomal membrane carrie | SALT00000073196_Cluster12208-002 |
| Cluster12211 | - | - | down | - | down | down | protochlorophyllide reductase | SALT00000053379_Cluster12211-009 |
| Cluster12212 | - | - | up | - | up | - | signal recognition particle rec | SALT00000057266_Cluster12212-008 |
| Cluster12223 | down | down | down | - | down | down | aspartic ase nepenthesin-1 pre | SALT00000040033_Cluster12223-013 |
| Cluster12243 | - | - | down | - | - | - | ferrochelatase II [Oryza sativa | SALT00000072137_Cluster12243-006 |
| Cluster12247 | - | - | up | - | up | up | Os08g0110400 [Oryza sativa | SALT00000077577_Cluster12247-006 |
| Cluster12253 | - | - | - | - | - | up | leucine-rich repeat-containing | SALT00000062240_Cluster12253-002 |
| Cluster12255 | - | - | up | - | up | up | OTU domain-containing 5-A- | SALT00000067528_Cluster12255-001 |
| Cluster1226 | - | - | up | - | up | - | calmodulin-binding 60 B-like | SALT00000083779_Cluster1226-016 |
| Cluster12262 | - | - | - | - | down | - | Os08g0345700 [Oryza sativa | SALT00000042562_Cluster12262-011 |
| Cluster12276 | - | - | - | - | - | down | DUF21 domain-containing ch | SALT00000030932_Cluster12276-003 |
| Cluster12295 | - | - | up | - | up | - | Os01g0621900 [Oryza sativa | SALT00000087390_Cluster12295-018 |
| Cluster12300 | - | - | down | - | - | - | aspartic ase nepenthesin-1 pre | SALT00000044136_Cluster12300-001 |
| Cluster12305 | - | - | up | - | up | up | partial [Oryza sativa Japonica | SALT00000066992_Cluster12305-001 |
| Cluster12323 | - | - | up | - | - | up | cytochrome P450 [Echinochlo | SALT00000047558_Cluster12323-002 |
| Cluster12334 | - | up | - | - | - | - | Methyl- binding domain conta | SALT00000079537_Cluster12334-002 |
| Cluster12337 | up | - | - | - | - | - | serine hydroxymethyltransfera | SALT00000045197_Cluster12337-013 |
| Cluster12358 | - | - | up | - | - | - | glycine-rich RNA-binding 7 [ | SALT00000022056_Cluster12358-002 |
| Cluster12359 | - | - | - | - | down | down | cytochrome b6 (chloroplast) [ | SALT00000030074_Cluster12359-001 |
| Cluster12361 | - | - | up | - | up | up | propionyl- carboxylase beta cl | SALT00000073195_Cluster12361-005 |
| Cluster12366 | - | - | up | - | up | up | Os07g0680600 [Oryza sativa | SALT00000022773_Cluster12366-003 |
| Cluster12368 | - | up | - | - | - | - | G-type lectin S-receptor-like s | SALT00000026850_Cluster12368-002 |
| Cluster12375 | - | - | down | - | - | - | G-type lectin S-receptor-like s | SALT00000028045_Cluster12375-001 |
| Cluster12399 | down | - | - | - | - | - | glutathione S-transferase F11- | SALT00000080614_Cluster12399-001 |
| Cluster1240 | - | - | down | - | down | down | Os04g0118700 [Oryza sativa | SALT00000075379_Cluster1240-001 |
| Cluster12405 | - | - | up | - | - | - | universal stress 9303 [Hordeu | SALT00000070262_Cluster12405-002 |
| Cluster12414 | - | - | down | - | down | down | polyamine oxidase-like isofor | SALT00000062932_Cluster12414-005 |
| Cluster12418 | - | - | down | - | down | - | [Hordeum vulgare vulgare] | SALT00000065950_Cluster12418-001 |
| Cluster12424 | up | up | up | - | - | - | transcription elongation factor | SALT00000089641_Cluster12424-001 |
| Cluster12428 | - | - | - | up | - | - | cation H(+) antiporter 19-like | SALT00000065023_Cluster12428-002 |
| Cluster1243 | - | - | - | - | - | down | NADH-plastoquinone oxidore | SALT00000089404_Cluster1243-021 |

| | | | | | | | | |
|---|---|---|---|---|---|---|---|---|
| Cluster12431 | - | - | - | - | down | down | Os08g0407200 [Oryza sativa | SALT00000073518_Cluster12431-003 |
| Cluster12438 | - | - | - | - | - | up | [Hordeum vulgare vulgare] | SALT00000027027_Cluster12438-010 |
| Cluster1244 | - | - | down | - | down | down | zinc finger BED domain-conta | SALT00000072681_Cluster1244-001 |
| Cluster12461 | - | down | down | - | down | down | GRAS transcription partial [Z | SALT00000064776_Cluster12461-007 |
| Cluster12463 | - | - | - | - | - | up | uncharacterized mitochondria | SALT00000070143_Cluster12463-001 |
| Cluster12468 | - | - | up | - | up | up | Os03g0726500 [Oryza sativa | SALT00000024559_Cluster12468-001 |
| Cluster12469 | - | - | - | - | - | down | [Hordeum vulgare vulgare] | SALT00000026741_Cluster12469-003 |
| Cluster12472 | - | - | down | - | down | down | growth-regulating factor 8 [Ze | SALT00000068259_Cluster12472-002 |
| Cluster12473 | - | - | up | - | - | up | beta-amylase chloroplastic [Se | SALT00000069019_Cluster12473-001 |
| Cluster12475 | - | - | up | - | up | - | alpha-glucosidase -like [Oryza | SALT00000072660_Cluster12475-003 |
| Cluster12489 | - | - | up | - | up | - | GTP-binding SAR1A-like [Se | SALT00000065726_Cluster12489-001 |
| Cluster12492 | - | - | - | - | down | down | peptidyl-prolyl cis-trans isom | SALT00000071729_Cluster12492-001 |
| Cluster1250 | - | - | up | - | up | up | Os09g0127800 [Oryza sativa | SALT00000089468_Cluster1250-029 |
| Cluster12503 | - | - | up | - | up | up | Os06g0556000 [Oryza sativa | SALT00000072102_Cluster12503-004 |
| Cluster12519 | - | - | up | - | up | - | pentatricopeptide repeat-conta | SALT00000079862_Cluster12519-001 |
| Cluster12533 | - | - | down | - | - | - | glutathione S-transferase 6 [Z | SALT00000080573_Cluster12533-001 |
| Cluster12536 | - | - | down | - | down | down | 6-phosphogluconate decarbox | SALT00000070719_Cluster12536-011 |
| Cluster12539 | - | - | down | - | down | - | transposon En Spm sub-class | SALT00000075353_Cluster12539-001 |
| Cluster12550 | - | - | up | - | up | up | probable beta-1,3-galactosyltr | SALT00000076755_Cluster12550-004 |
| Cluster1257 | - | - | - | - | up | up | heat shock 70 kDa 15-like [Se | SALT00000084104_Cluster1257-024 |
| Cluster12570 | - | - | up | - | up | up | Os02g0240100 [Oryza sativa | SALT00000072502_Cluster12570-002 |
| Cluster12576 | - | - | - | - | down | down | asparaginyl-tRNA cytoplasmi | SALT00000037141_Cluster12576-007 |
| Cluster12606 | - | - | up | - | up | up | hypersensitive-induced respor | SALT00000026219_Cluster12606-001 |
| Cluster12626 | - | - | up | - | up | up | DETOXIFICATION 33 | SALT00000010213_Cluster12626-001 |
| Cluster1263 | - | - | - | - | down | down | Os02g0195500 [Oryza sativa | SALT00000063422_Cluster1263-008 |
| Cluster12641 | - | - | - | - | - | up | sodium-coupled neutral aminc | SALT00000074644_Cluster12641-002 |
| Cluster12645 | - | - | - | - | - | down | hydroxymethylbutenyl 4-diph | SALT00000027642_Cluster12645-005 |
| Cluster12647 | - | - | down | - | down | - | geranylgeranyl reductase [Ory | SALT00000042735_Cluster12647-073 |
| Cluster12659 | - | - | - | - | up | - | dihydrofolate reductase [Cucu | SALT00000027672_Cluster12659-003 |
| Cluster12663 | - | - | up | - | up | - | myb-like DNA-binding SHA( | SALT00000069182_Cluster12663-005 |
| Cluster12669 | - | - | down | - | - | - | ribonucleotide reductase [Ory | SALT00000021027_Cluster12669-002 |
| Cluster12677 | - | - | down | - | - | - | partial [Brachypodium distach | SALT00000077336_Cluster12677-001 |
| Cluster1268 | - | - | down | - | down | down | cellulose synthase-like 6 [Hor | SALT00000028679_Cluster1268-007 |
| Cluster12684 | - | - | down | - | - | - | uncharacterized membrane At | SALT00000068166_Cluster12684-013 |
| Cluster127 | - | - | - | - | down | - | alpha-L-fucosidase 2 precurso | SALT00000040812_Cluster127-001 |
| Cluster12722 | - | - | - | - | down | - | uncharacterized protein LOC1 | SALT00000083766_Cluster12722-001 |
| Cluster1273 | - | - | - | - | down | - | hypothetical protein SETIT_0 | SALT00000091870_Cluster1273-008 |
| Cluster12748 | - | - | up | - | up | up | cysteine-rich receptor kinase 1 | SALT00000046492_Cluster12748-005 |
| Cluster1275 | up | - | up | - | - | up | TPA: calcium-dependent lipid | SALT00000088677_Cluster1275-006 |
| Cluster12752 | - | - | up | - | up | - | type I inositol-1,4,5-trisphospl | SALT00000066037_Cluster12752-003 |
| Cluster12755 | - | up | - | - | - | down | [Hordeum vulgare vulgare] | SALT00000084500_Cluster12755-005 |
| Cluster12757 | - | up | - | - | - | - | poly [Oryza sativa Japonica G | SALT00000023979_Cluster12757-002 |
| Cluster12773 | - | - | up | - | - | - | uncharacterized LOC1002793 | SALT00000071864_Cluster12773-004 |
| Cluster12780 | - | down | - | - | - | - | fluG [Setaria italica] | SALT00000022564_Cluster12780-002 |
| Cluster12790 | up | - | - | - | down | - | PREDICTED: uncharacterize | SALT00000081942_Cluster12790-001 |
| Cluster128 | - | - | - | - | down | - | Os03g0837100 [Oryza sativa | SALT00000087678_Cluster128-001 |
| Cluster12809 | - | - | up | - | up | up | Os08g0545000 [Oryza sativa | SALT00000064431_Cluster12809-002 |
| Cluster12810 | - | - | down | - | down | down | (chloroplast) [Oryza barthii] | SALT00000072478_Cluster12810-003 |
| Cluster12833 | - | - | down | - | - | - | AF466646_7 poly | SALT00000065631_Cluster12833-001 |
| Cluster12849 | - | - | up | - | up | up | hexokinase [Zea mays] | SALT00000026786_Cluster12849-002 |
| Cluster12858 | - | down | down | - | down | - | fiddlehead [Hordeum vulgare | SALT00000043486_Cluster12858-010 |
| Cluster12860 | - | - | up | - | up | - | ABC transporter subunit-like | SALT00000059053_Cluster12860-009 |
| Cluster12865 | - | - | up | - | up | up | Os01g0110500 [Oryza sativa | SALT00000074179_Cluster12865-002 |
| Cluster12866 | - | - | up | - | - | - | gtk16 [Zea mays] | SALT00000076347_Cluster12866-001 |
| Cluster12870 | - | - | down | - | down | - | ATP synthase CF0 B subunit ( | SALT00000026611_Cluster12870-002 |
| Cluster12871 | - | up | up | - | up | up | ATG8-interacting 1 isoform X | SALT00000031173_Cluster12871-001 |
| Cluster1288 | down | - | down | - | - | - | DNA-directed RNA polymera | SALT00000088473_Cluster1288-003 |
| Cluster12917 | - | - | - | - | down | - | heparanase 3 | SALT00000076784_Cluster12917-002 |

| | | | | | | | | |
|---|---|---|---|---|---|---|---|---|
| Cluster12919 | - | - | down | - | - | - | [Oryza sativa Japonica Group] | SALT00000021331_Cluster12919-001 |
| Cluster12935 | - | - | up | - | up | up | polyol transporter 5-like | SALT00000074809_Cluster12935-008 |
| Cluster12939 | - | down | down | - | - | - | Os05g0476200 [Oryza sativa | SALT00000066461_Cluster12939-004 |
| Cluster12940 | - | - | down | - | - | - | TPX2 isoform X2 | SALT00000065702_Cluster12940-004 |
| Cluster12942 | - | - | up | - | up | - | Uncharacterised [Vibrio chole | SALT00000025439_Cluster12942-001 |
| Cluster12949 | - | - | up | - | - | - | AF140553_1DNA-binding W | SALT00000077096_Cluster12949-002 |
| Cluster1295 | - | down | down | - | - | - | Os07g0252400 [Oryza sativa | SALT00000086225_Cluster1295-045 |
| Cluster12966 | - | - | up | - | up | up | potassium channel KOR1 | SALT00000062980_Cluster12966-002 |
| Cluster12978 | - | - | down | - | down | down | ubiquitin-like-specific proteas | SALT00000078441_Cluster12978-001 |
| Cluster1301 | - | down | - | - | - | - | ramosa 1 enhancer locus 2 [Z | SALT00000068191_Cluster1301-001 |
| Cluster13020 | - | - | down | - | - | - | Os10g0562700 [Oryza sativa | SALT00000023169_Cluster13020-003 |
| Cluster13022 | - | - | up | - | up | up | SLT1 expressed [Oryza sativa | SALT00000026030_Cluster13022-005 |
| Cluster13030 | - | up | up | up | up | up | Os03g0726500 [Oryza sativa | SALT00000006072_Cluster13030-002 |
| Cluster13052 | - | - | up | - | - | - | peroxisomal fatty acid beta-ox | SALT00000073535_Cluster13052-005 |
| Cluster13056 | - | - | down | - | - | - | GTP-binding homolog | SALT00000080805_Cluster13056-008 |
| Cluster13059 | - | - | - | - | up | up | Os09g0344800 [Oryza sativa | SALT00000077914_Cluster13059-012 |
| Cluster13064 | - | - | up | - | up | up | ultraviolet-B receptor UVR8 [ | SALT00000074623_Cluster13064-008 |
| Cluster13068 | - | - | - | - | up | - | calcium-dependent kinase 11 | | SALT00000081505_Cluster13068-002 |
| Cluster13075 | - | - | up | - | up | up | cell wall invertase | SALT00000068026_Cluster13075-002 |
| Cluster13079 | - | - | up | - | - | - | ubiquitin 5 [Cucumis melo] | SALT00000077013_Cluster13079-003 |
| Cluster13081 | - | - | down | - | - | - | serine threonine- kinase recep | SALT00000080555_Cluster13081-005 |
| Cluster13085 | - | - | down | - | - | - | 14-3-3 [Zea mays] | SALT00000049539_Cluster13085-001 |
| Cluster13086 | - | - | down | - | down | down | TPA: oxidoreductase | SALT00000066995_Cluster13086-002 |
| Cluster13091 | - | - | up | - | up | up | F-box kelch-repeat At1g6748( | SALT00000083104_Cluster13091-018 |
| Cluster13103 | - | up | - | - | down | down | ribonuclease H At1g65750 | SALT00000069004_Cluster13103-001 |
| Cluster13117 | - | - | down | - | down | down | LSG [Saccharum hybrid culti | SALT00000065920_Cluster13117-001 |
| Cluster13132 | - | - | - | - | - | down | BURP domain-containing 3 | SALT00000048690_Cluster13132-001 |
| Cluster13133 | - | - | up | - | up | up | nucleotide-binding 1 [Zea ma | SALT00000063618_Cluster13133-006 |
| Cluster13135 | - | - | up | - | up | up | transporter [Zea mays] | SALT00000067392_Cluster13135-005 |
| Cluster1314 | - | down | down | - | down | down | sucrose synthase 1 [Oryza sati | SALT00000087036_Cluster1314-050 |
| Cluster13142 | - | - | up | - | up | - | probable UDP-arabinose 4-ep | SALT00000079605_Cluster13142-002 |
| Cluster13145 | - | - | - | - | down | down | LOC100284908 isoform X1 [ | SALT00000038409_Cluster13145-006 |
| Cluster13157 | - | - | - | - | down | - | hypothetical protein SORBID | SALT00000063909_Cluster13157-003 |
| Cluster13180 | - | - | up | - | up | up | nodulin [Zea mays] | SALT00000027982_Cluster13180-009 |
| Cluster13190 | - | - | up | - | up | up | zinc C3HC4 type family [Zea | SALT00000029407_Cluster13190-005 |
| Cluster13195 | - | up | up | - | - | - | NADP-dependent malic enzy | SALT00000081382_Cluster13195-018 |
| Cluster1320 | - | down | down | down | down | down | sucrose synthase 1 [Oryza sati | SALT00000079408_Cluster1320-001 |
| Cluster13200 | - | - | - | - | up | - | receptor kinase 1 | SALT00000061979_Cluster13200-007 |
| Cluster1322 | - | - | down | - | down | down | mitochondrial 2-oxoglutarate | SALT00000046963_Cluster1322-007 |
| Cluster13235 | down | down | down | - | - | - | [Hordeum vulgare vulgare] | SALT00000080936_Cluster13235-001 |
| Cluster13252 | - | - | up | - | up | up | TPA: HLH DNA-binding dor | SALT00000043048_Cluster13252-005 |
| Cluster13268 | down | down | - | - | - | - | ATP ADP translocator [Oryza | SALT00000064816_Cluster13268-007 |
| Cluster1328 | - | - | - | - | up | up | Os02g0517300 [Oryza sativa | SALT00000089582_Cluster1328-022 |
| Cluster13281 | - | - | up | - | - | - | TPA: ARM repeat-containing | SALT00000067376_Cluster13281-006 |
| Cluster13290 | - | - | up | - | up | up | NRT1 PTR FAMILY -like [Se | SALT00000056925_Cluster13290-001 |
| Cluster13293 | - | - | - | - | down | - | Os08g0401500 [Oryza sativa | SALT00000069432_Cluster13293-001 |
| Cluster13294 | - | - | down | - | down | - | pentatricopeptide repeat-conta | SALT00000069572_Cluster13294-001 |
| Cluster13297 | - | - | up | - | up | up | probable amino acid permease | SALT00000081275_Cluster13297-012 |
| Cluster1330 | - | - | up | - | up | up | chaperone chloroplastic [Setar | SALT00000076137_Cluster1330-022 |
| Cluster13300 | - | - | up | - | - | - | sucrose synthase | SALT00000084146_Cluster13300-002 |
| Cluster13304 | - | - | up | - | up | up | membrane [Zea mays] | SALT00000045118_Cluster13304-009 |
| Cluster1331 | - | down | down | - | - | - | villin 2 [Oryza sativa Japonica | SALT00000085881_Cluster1331-016 |
| Cluster13312 | - | up | up | - | up | up | cysteine protease 1 precursor | | SALT00000077636_Cluster13312-002 |
| Cluster1333 | - | - | down | - | down | down | Na+ H+ antiporter | SALT00000085038_Cluster1333-026 |
| Cluster1334 | - | - | down | - | - | - | LONGIFOLIA 2-like | SALT00000085277_Cluster1334-013 |
| Cluster13366 | - | - | up | - | - | - | Os02g0140300 [Oryza sativa | SALT00000082973_Cluster13366-001 |
| Cluster13390 | - | - | down | - | - | - | MKI67 FHA domain-interacti | SALT00000070304_Cluster13390-002 |
| Cluster13392 | - | - | up | - | up | up | wall-associated receptor kinas | SALT00000076096_Cluster13392-004 |

| | | | | | | | | |
|---|---|---|---|---|---|---|---|---|
| Cluster13399 | - | - | up | - | - | - | [Oryza sativa Indica Group] | SALT00000065941_Cluster13399-001 |
| Cluster134 | - | - | down | - | - | - | cellulose synthase A catalytic | SALT00000086549_Cluster134-031 |
| Cluster13400 | - | - | down | - | - | - | Os05g0566400 [Oryza sativa | SALT00000067846_Cluster13400-002 |
| Cluster13404 | - | - | - | - | up | - | YT521-B-like family [Zea ma | SALT00000076862_Cluster13404-017 |
| Cluster13408 | - | - | down | - | - | down | zinc finger MYM-type 1-like | SALT00000025655_Cluster13408-002 |
| Cluster1341 | up | up | - | - | down | down | nudix -like [Oryza sativa Japo | SALT00000062529_Cluster1341-007 |
| Cluster13410 | - | - | - | - | up | - | Os10g0533600 [Oryza sativa | SALT00000025398_Cluster13410-008 |
| Cluster13414 | - | - | down | - | - | - | tetratricopeptide repeat 28-like | SALT00000063870_Cluster13414-002 |
| Cluster13417 | - | - | up | - | up | up | PALE chloroplastic | SALT00000070130_Cluster13417-002 |
| Cluster13424 | - | - | up | - | up | up | senescence-associated 12 [Zea | SALT00000024502_Cluster13424-004 |
| Cluster13428 | - | down | down | - | - | - | Os05g0164100 [Oryza sativa | SALT00000066975_Cluster13428-005 |
| Cluster1345 | - | - | down | - | down | - | partial [Oryza sativa Japonica | SALT00000078609_Cluster1345-001 |
| Cluster13453 | - | - | up | - | up | up | beta-amylase chloroplastic [Se | SALT00000072522_Cluster13453-010 |
| Cluster1346 | down | - | - | - | up | up | photosystem I P700 apo A1 (c | SALT00000088776_Cluster1346-002 |
| Cluster13495 | - | - | up | - | - | - | ATP ADP translocator [Oryza | SALT00000077615_Cluster13495-002 |
| Cluster13504 | - | - | down | - | - | - | probable LRR receptor-like se | SALT00000072967_Cluster13504-004 |
| Cluster1351 | - | - | up | - | - | - | U-box domain-containing 35 | SALT00000076864_Cluster1351-018 |
| Cluster1354 | - | - | down | - | down | down | cellulose synthase1 | SALT00000086084_Cluster1354-001 |
| Cluster13552 | - | - | down | - | down | - | RITA-2 partial | SALT00000062491_Cluster13552-002 |
| Cluster13577 | - | - | up | - | - | - | phosphatase 2C family [Zea m | SALT00000077214_Cluster13577-003 |
| Cluster13578 | - | - | up | - | up | up | NADPH--cytochrome P450 re | SALT00000081771_Cluster13578-009 |
| Cluster13581 | - | - | up | - | up | up | RCH2 [Oryza sativa Japonica | SALT00000029400_Cluster13581-001 |
| Cluster13585 | - | - | down | - | down | - | transcription factor UNE10 is | SALT00000068702_Cluster13585-005 |
| Cluster13591 | up | up | up | - | - | - | adenosine 5 -phosphosulfate r | SALT00000057418_Cluster13591-009 |
| Cluster13606 | - | - | up | - | up | up | disease resistance At5g47280 | SALT00000026079_Cluster13606-003 |
| Cluster13649 | - | - | up | - | up | up | anthranilate synthase compon | SALT00000065637_Cluster13649-008 |
| Cluster13657 | - | - | - | - | up | - | chloroplastic-like isoform X1 | SALT00000024333_Cluster13657-002 |
| Cluster13660 | - | - | up | - | - | - | pentatricopeptide repeat-conta | SALT00000067952_Cluster13660-001 |
| Cluster13666 | - | - | down | - | - | - | oligosaccharyl transferase ST | SALT00000073027_Cluster13666-014 |
| Cluster13679 | - | - | - | - | - | up | amino acid permease 3-like [S | SALT00000063887_Cluster13679-012 |
| Cluster13700 | - | - | up | - | up | up | wall-associated receptor kinas | SALT00000022827_Cluster13700-002 |
| Cluster13710 | - | - | - | - | down | - | ---NA--- | SALT00000080227_Cluster13710-002 |
| Cluster13712 | - | - | down | - | down | down | F-box At3g25750 [Setaria ital | SALT00000081281_Cluster13712-002 |
| Cluster13759 | - | down | down | - | down | - | CHLOROPLAST ENHANCI | SALT00000062325_Cluster13759-001 |
| Cluster13762 | - | - | down | - | down | - | AF187823_1transposase DOF | SALT00000067472_Cluster13762-002 |
| Cluster13763 | - | - | down | - | down | down | chloride channel CLC-a-like [ | SALT00000071767_Cluster13763-002 |
| Cluster13769 | - | - | up | - | up | up | ankyrin repeat [Oryza sativa J | SALT00000029008_Cluster13769-001 |
| Cluster1377 | - | - | - | - | - | down | serine carboxypeptidase-like | SALT00000053574_Cluster1377-001 |
| Cluster13781 | - | - | up | - | up | up | Os01g0750600 [Oryza sativa | SALT00000022032_Cluster13781-001 |
| Cluster13785 | - | - | down | - | - | - | ketol-acid chloroplastic-like | SALT00000053767_Cluster13785-002 |
| Cluster13804 | - | up | - | - | - | - | DNA repair rhp26 [Zea mays] | SALT00000062763_Cluster13804-001 |
| Cluster13807 | up | - | - | - | - | - | scarecrow 6 [Brachypodium d | SALT00000074915_Cluster13807-001 |
| Cluster13812 | - | - | up | - | up | up | receptor kinase HERK 1 [Ory | SALT00000081725_Cluster13812-002 |
| Cluster13815 | - | - | down | - | - | - | probable isoaspartyl peptidase | SALT00000072362_Cluster13815-004 |
| Cluster13852 | - | - | up | - | - | - | [Hordeum vulgare vulgare] | SALT00000084414_Cluster13852-002 |
| Cluster13866 | - | - | down | - | down | down | Os02g0575500 [Oryza sativa | SALT00000023227_Cluster13866-002 |
| Cluster13877 | - | - | up | - | up | - | filamentation temperature-sen | SALT00000029227_Cluster13877-003 |
| Cluster13889 | - | - | up | - | - | - | zinc finger A20 and AN1 dom | SALT00000062008_Cluster13889-001 |
| Cluster1389 | - | - | down | - | down | - | heat shock HSP70- partial | SALT00000066801_Cluster1389-001 |
| Cluster13899 | - | - | up | - | up | - | digalactosyldiacylglycerol syn | SALT00000021229_Cluster13899-002 |
| Cluster13903 | - | - | down | - | down | - | pentatricopeptide repeat-conta | SALT00000069554_Cluster13903-001 |
| Cluster13911 | - | - | up | - | up | up | Os03g0362200 [Oryza sativa | SALT00000026063_Cluster13911-001 |
| Cluster13914 | - | down | down | down | down | - | [Hordeum vulgare vulgare] | SALT00000030694_Cluster13914-010 |
| Cluster13930 | down | - | - | - | - | - | Os04g0599900 [Oryza sativa | SALT00000081172_Cluster13930-007 |
| Cluster13939 | - | - | down | - | down | - | kinesin-5-like [Setaria italica] | SALT00000067993_Cluster13939-004 |
| Cluster13940 | up | up | up | - | - | - | Os01g0812900 [Oryza sativa | SALT00000069754_Cluster13940-001 |
| Cluster13942 | - | down | down | - | - | - | wall-associated receptor kinas | SALT00000071350_Cluster13942-005 |
| Cluster13947 | - | down | down | - | - | - | long-chain-alcohol oxidase FA | SALT00000082807_Cluster13947-005 |

| | | | | | | | | |
|---|---|---|---|---|---|---|---|---|
| Cluster13953 | - | - | - | - | up | - | axi 1 [Oryza sativa Japonica ( | SALT00000079602_Cluster13953-006 |
| Cluster13955 | - | - | up | - | - | - | Os06g0693700 [Oryza sativa | SALT00000073515_Cluster13955-004 |
| Cluster13961 | - | - | up | - | up | up | Os01g0747400 [Oryza sativa | SALT00000026697_Cluster13961-001 |
| Cluster13969 | - | - | - | - | up | up | E3 ubiquitin- ligase RNF126- | SALT00000072633_Cluster13969-007 |
| Cluster13973 | - | - | - | - | up | - | cactin [Setaria italica] | SALT00000077632_Cluster13973-004 |
| Cluster13980 | - | - | down | - | down | - | orf256 (mitochondrion) [Tritic | SALT00000067766_Cluster13980-001 |
| Cluster13989 | - | - | up | - | up | - | Os01g0621900 [Oryza sativa | SALT00000023466_Cluster13989-001 |
| Cluster13994 | up | - | up | - | - | up | Os01g0858350 [Oryza sativa | SALT00000069488_Cluster13994-014 |
| Cluster14009 | - | - | up | - | up | up | NRT1 PTR FAMILY -like [Se | SALT00000075754_Cluster14009-008 |
| Cluster14028 | up | - | up | - | - | - | ferredoxin-thioredoxin variabl | SALT00000078294_Cluster14028-002 |
| Cluster14034 | - | - | down | down | down | - | cytochrome P450 superfamily | SALT00000047715_Cluster14034-001 |
| Cluster14035 | down | - | down | - | - | - | RING zinc finger domain supe | SALT00000044248_Cluster14035-002 |
| Cluster14049 | up | - | up | - | - | - | L-type lectin-domain containi | SALT00000027776_Cluster14049-001 |
| Cluster14057 | - | - | up | - | - | - | sodium-coupled neutral aminc | SALT00000081165_Cluster14057-004 |
| Cluster14074 | - | - | up | - | - | - | Os04g0590400 [Oryza sativa | SALT00000028990_Cluster14074-002 |
| Cluster14095 | - | - | up | - | up | up | kinase domain containing [Ze | SALT00000083529_Cluster14095-004 |
| Cluster14096 | - | down | down | down | down | - | Os06g0118700 [Oryza sativa | SALT00000084100_Cluster14096-005 |
| Cluster14098 | - | - | up | - | - | - | 14-3-3 [Zea mays] | SALT00000027284_Cluster14098-002 |
| Cluster14102 | - | - | up | - | up | up | ankyrin repeat-containing At5 | SALT00000071540_Cluster14102-006 |
| Cluster14106 | - | - | - | - | down | - | Os02g0595700 [Oryza sativa | SALT00000077119_Cluster14106-003 |
| Cluster1411 | - | - | - | - | down | down | triose phosphate phosphate tra | SALT00000053583_Cluster1411-016 |
| Cluster14112 | - | - | up | - | up | up | glucan endo-1,3-beta-glucosic | SALT00000023956_Cluster14112-001 |
| Cluster14120 | - | - | up | - | up | - | PREDICTED: uncharacterized | SALT00000074270_Cluster14120-003 |
| Cluster14127 | - | - | up | - | - | - | SPX domain-containing meml | SALT00000025706_Cluster14127-004 |
| Cluster14128 | - | - | down | - | down | down | polygalacturonate 4-alpha-gal | SALT00000048711_Cluster14128-002 |
| Cluster14135 | - | - | down | - | down | - | [Oryza sativa Indica Group] | SALT00000074646_Cluster14135-010 |
| Cluster14149 | - | - | - | - | - | up | phospholipase A1- chloroplast | SALT00000081751_Cluster14149-006 |
| Cluster1415 | - | down | down | - | down | - | Os08g0400300 [Oryza sativa | SALT00000090139_Cluster1415-015 |
| Cluster14150 | - | - | - | up | up | - | S-adenosylmethionine synthet | SALT00000025475_Cluster14150-002 |
| Cluster14153 | - | - | down | - | - | - | UPF0160 -like [Setaria italica | SALT00000071221_Cluster14153-001 |
| Cluster14162 | - | - | up | - | up | up | [Hordeum vulgare vulgare] | SALT00000030754_Cluster14162-003 |
| Cluster14176 | - | - | up | - | up | - | ELMO domain-containing A [ | SALT00000067890_Cluster14176-001 |
| Cluster14178 | - | up | up | - | up | up | catalytic hydrolase isoform X | SALT00000070470_Cluster14178-001 |
| Cluster1419 | - | up | up | - | - | - | Os01g0534900 [Oryza sativa | SALT00000081170_Cluster1419-007 |
| Cluster14200 | - | - | up | - | up | up | Os07g0575900 [Oryza sativa | SALT00000065462_Cluster14200-002 |
| Cluster14206 | - | - | - | - | up | - | beta-glucosidase 3B-like | SALT00000080244_Cluster14206-002 |
| Cluster14214 | - | - | down | - | down | - | myosin heavy cardiac muscle | SALT00000062917_Cluster14214-001 |
| Cluster14231 | up | up | up | - | up | up | heat shock cognate 70 kDa -li | SALT00000066104_Cluster14231-008 |
| Cluster14235 | - | - | - | - | down | - | 2-dehydro-3-deoxyphosphooc | SALT00000074144_Cluster14235-001 |
| Cluster1424 | - | up | up | up | up | up | sodium hydrogen exchanger 8 | SALT00000088954_Cluster1424-021 |
| Cluster14241 | - | up | up | - | - | - | alcohol dehydrogenase superf | SALT00000078632_Cluster14241-004 |
| Cluster14247 | up | up | up | - | up | up | LOC100281573 isoform X1 [ | SALT00000023367_Cluster14247-009 |
| Cluster14262 | - | - | up | - | up | - | serine threonine- kinase CCR | SALT00000024009_Cluster14262-001 |
| Cluster14263 | - | - | down | - | down | down | ATP-dependent Clp protease p | SALT00000026314_Cluster14263-003 |
| Cluster14266 | up | up | up | - | - | - | ubiquitin- ligase [Zea mays] | SALT00000069291_Cluster14266-008 |
| Cluster1427 | - | - | down | - | down | down | zinc finger CCCH domain-cor | SALT00000087538_Cluster1427-001 |
| Cluster14301 | - | up | up | - | up | up | cyclic nucleotide-gated ion ch | SALT00000081204_Cluster14301-002 |
| Cluster14333 | - | - | down | - | down | - | probable aldo-keto reductase | SALT00000083843_Cluster14333-001 |
| Cluster14335 | - | - | down | - | - | - | O-Glycosyl hydrolase superfa | SALT00000026842_Cluster14335-003 |
| Cluster14346 | - | - | up | - | up | - | oligopeptide transporter 7 | SALT00000081056_Cluster14346-002 |
| Cluster14363 | - | - | up | - | up | - | filamentation temperature-sen | SALT00000028164_Cluster14363-001 |
| Cluster14365 | - | up | - | - | - | down | serine hydroxymethyltransfera | SALT00000048245_Cluster14365-009 |
| Cluster14381 | - | - | up | - | up | - | submergence induced SI397 [ | SALT00000066608_Cluster14381-002 |
| Cluster14386 | - | - | down | - | down | down | SCARECROW [Zea mays] | SALT00000070876_Cluster14386-004 |
| Cluster1440 | - | - | down | - | - | down | receptor kinase HSL1 | SALT00000087795_Cluster1440-001 |
| Cluster14400 | - | - | up | - | - | - | probable LRR receptor-like se | SALT00000065845_Cluster14400-002 |
| Cluster14403 | - | - | down | - | - | - | Phosphoenolpyruvate carboxy | SALT00000069703_Cluster14403-002 |
| Cluster1441 | - | - | down | - | - | - | retrotransposon expressed [Or | SALT00000085159_Cluster1441-001 |

| | | | | | | | | |
|---|---|---|---|---|---|---|---|---|
| Cluster14416 | - | - | - | - | down | - | PREDICTED: uncharacterize | SALT00000083420_Cluster14416-001 |
| Cluster14445 | - | - | up | - | - | - | 2,3-bisphosphoglycerate-inde | SALT00000047510_Cluster14445-001 |
| Cluster14457 | - | - | up | - | - | up | methylenetetrahydrofolate red | SALT00000082663_Cluster14457-004 |
| Cluster14461 | - | - | up | - | up | up | nuclease HARBI1 | SALT00000026496_Cluster14461-007 |
| Cluster14465 | - | - | - | - | - | down | T-complex 1 subunit beta [Or | SALT00000051957_Cluster14465-001 |
| Cluster14468 | - | up | up | - | up | - | electron transfer flavo subunit | SALT00000068175_Cluster14468-002 |
| Cluster14474 | - | - | down | - | down | - | nucleobase-ascorbate transpor | SALT00000082760_Cluster14474-003 |
| Cluster14483 | - | - | up | - | up | up | auxin response factor 21 isofo | SALT00000064879_Cluster14483-002 |
| Cluster14507 | - | - | - | - | - | down | probable replication factor A 7 | SALT00000077628_Cluster14507-002 |
| Cluster14508 | - | - | up | - | up | up | C2 domain-containing At1g53 | SALT00000084191_Cluster14508-001 |
| Cluster14524 | up | up | up | - | up | - | iron inhibited ABC transporte | SALT00000080184_Cluster14524-006 |
| Cluster14542 | - | - | up | - | up | up | S-type anion channel SLAH2- | SALT00000062179_Cluster14542-005 |
| Cluster14547 | - | - | - | - | - | up | Os05g0419200 [Oryza sativa | SALT00000062258_Cluster14547-006 |
| Cluster1455 | - | - | up | - | up | - | Os09g0454900 [Oryza sativa | SALT00000025478_Cluster1455-002 |
| Cluster14554 | - | up | - | up | - | - | [Hordeum vulgare vulgare] | SALT00000071302_Cluster14554-006 |
| Cluster14564 | - | - | down | - | down | down | eukaryotic translation initiatio | SALT00000053918_Cluster14564-001 |
| Cluster14565 | - | - | up | - | up | up | UDP-glucose 4-epimerase 2 | SALT00000065594_Cluster14565-002 |
| Cluster14569 | down | - | down | - | - | - | transcription factor bHLH041 | SALT00000074513_Cluster14569-002 |
| Cluster14570 | - | - | up | - | up | up | transporter [Zea mays] | SALT00000079669_Cluster14570-001 |
| Cluster14575 | - | - | up | - | up | up | Os03g0305100 [Oryza sativa | SALT00000043645_Cluster14575-008 |
| Cluster14577 | - | - | down | - | down | down | probable serine incorporator | SALT00000059163_Cluster14577-001 |
| Cluster1459 | - | up | up | - | - | - | probable phospholipid-transpo | SALT00000090480_Cluster1459-006 |
| Cluster14594 | - | - | up | - | - | up | Os10g0555100 [Oryza sativa | SALT00000069552_Cluster14594-004 |
| Cluster14621 | - | - | down | - | - | down | probable inactive receptor kin | SALT00000070638_Cluster14621-010 |
| Cluster14631 | - | - | up | - | - | - | Os09g0536700 [Oryza sativa | SALT00000029709_Cluster14631-002 |
| Cluster14640 | - | - | up | - | - | - | [Oryza sativa Japonica Group | SALT00000021162_Cluster14640-002 |
| Cluster14660 | - | - | - | - | up | - | alkaline alpha galactosidase I | SALT00000069360_Cluster14660-003 |
| Cluster1467 | - | up | up | - | - | - | probable transcriptional regula | SALT00000081020_Cluster1467-002 |
| Cluster1468 | - | - | up | - | - | - | serine threonine- kinase STE2 | SALT00000091498_Cluster1468-001 |
| Cluster1469 | - | - | - | - | up | - | [Hordeum vulgare vulgare] | SALT00000073891_Cluster1469-014 |
| Cluster14701 | - | down | down | - | - | - | Os12g0563500 [Oryza sativa | SALT00000073319_Cluster14701-002 |
| Cluster14707 | - | - | down | - | down | down | Os02g0687900 [Oryza sativa | SALT00000080238_Cluster14707-001 |
| Cluster1471 | - | - | down | - | - | - | Os03g0165400 [Oryza sativa | SALT00000068706_Cluster1471-005 |
| Cluster14714 | - | - | - | - | down | - | GDP-mannose 4,6 dehydratas | SALT00000069322_Cluster14714-003 |
| Cluster14721 | - | - | - | - | up | - | target of Myb 1-like [Setaria i | SALT00000020687_Cluster14721-001 |
| Cluster14723 | - | - | down | - | - | - | LOC542697 isoform X4 [Zea | SALT00000077892_Cluster14723-004 |
| Cluster14726 | - | - | - | - | - | up | Os02g0550600 [Oryza sativa | SALT00000064398_Cluster14726-005 |
| Cluster14734 | - | - | - | - | down | - | POX domain homeobox DNA | SALT00000078379_Cluster14734-001 |
| Cluster14778 | - | down | down | - | - | - | long-chain-alcohol oxidase FA | SALT00000070461_Cluster14778-008 |
| Cluster14791 | - | down | - | - | - | - | nicotianamine aminotransfera | SALT00000062027_Cluster14791-005 |
| Cluster1480 | - | - | down | - | - | down | EDR1-like [Oryza sativa Japo | SALT00000053694_Cluster1480-001 |
| Cluster14804 | - | - | down | - | - | - | ribulose-1,5-bisphosphate carl | SALT00000080864_Cluster14804-001 |
| Cluster14810 | - | - | down | - | down | down | amidase 1-like | SALT00000052595_Cluster14810-011 |
| Cluster14827 | up | up | - | - | - | - | PB1 [Oryza sativa Japonica G | SALT00000066821_Cluster14827-003 |
| Cluster14830 | - | - | up | - | up | - | Os01g0880300 [Oryza sativa | SALT00000071475_Cluster14830-001 |
| Cluster14877 | - | - | down | - | - | - | Os07g0682400 [Oryza sativa | SALT00000081605_Cluster14877-002 |
| Cluster14881 | - | - | down | - | - | - | Os01g0241000 [Oryza sativa | SALT00000063182_Cluster14881-001 |
| Cluster14888 | - | - | down | - | down | down | aquaporin NIP1-1 [Zea mays] | SALT00000073134_Cluster14888-001 |
| Cluster149 | up | up | up | - | - | - | phosphatase 2C family [Zea m | SALT00000077665_Cluster149-009 |
| Cluster14917 | - | - | down | - | - | - | fatty aldehyde dehydrogenase | SALT00000026771_Cluster14917-002 |
| Cluster14923 | - | - | down | - | down | - | transposon Ty3-G Gag-Pol po | SALT00000066609_Cluster14923-001 |
| Cluster14933 | - | - | up | - | - | up | stachyose synthase [Zea mays | SALT00000029802_Cluster14933-001 |
| Cluster14936 | - | - | up | - | up | up | Os03g0291200 [Oryza sativa | SALT00000068025_Cluster14936-001 |
| Cluster14941 | up | up | - | - | - | - | transmembrane emp24 domai | SALT00000079776_Cluster14941-001 |
| Cluster14947 | - | - | down | - | - | - | ubiquitin- ligase [Zea mays] | SALT00000067323_Cluster14947-002 |
| Cluster1496 | - | up | up | - | - | - | PREDICTED: uncharacterize | SALT00000076019_Cluster1496-001 |
| Cluster14963 | - | - | down | - | - | - | chlorophyll a b-binding apo C | SALT00000071934_Cluster14963-001 |
| Cluster14972 | - | - | up | - | up | up | acc synthase [Zea mays] | SALT00000025873_Cluster14972-001 |

| | | | | | | | | |
|---|---|---|---|---|---|---|---|---|
| Cluster14973 | - | - | down | - | down | down | [Hordeum vulgare vulgare] | SALT00000030337_Cluster14973-008 |
| Cluster1498 | - | - | - | - | down | - | potassium transporter | SALT00000084335_Cluster1498-013 |
| Cluster14984 | - | - | up | - | up | up | L-type lectin-domain containi | SALT00000063709_Cluster14984-010 |
| Cluster1499 | - | - | down | - | down | - | TSS [Setaria italica] | SALT00000084827_Cluster1499-012 |
| Cluster14995 | - | - | - | - | down | - | pentatricopeptide repeat-conta | SALT00000064503_Cluster14995-004 |
| Cluster14998 | - | - | up | - | - | - | serine arginine repetitive matr | SALT00000067990_Cluster14998-001 |
| Cluster1500 | - | down | down | - | - | - | ---NA--- | SALT00000067113_Cluster1500-001 |
| Cluster15005 | - | up | up | - | up | up | mechanosensitive ion channel | SALT00000084525_Cluster15005-003 |
| Cluster15013 | - | - | down | - | down | down | beta-1,3-glucanase precursor [ | SALT00000081250_Cluster15013-007 |
| Cluster15016 | - | up | up | - | - | - | Os04g0391500 [Oryza sativa | SALT00000053913_Cluster15016-009 |
| Cluster15017 | - | - | down | - | - | down | [Oryza sativa Japonica Group] | SALT00000063065_Cluster15017-001 |
| Cluster15018 | - | - | - | - | - | down | [Hordeum vulgare vulgare] | SALT00000063340_Cluster15018-001 |
| Cluster15031 | up | - | up | - | up | up | lipid phosphate phosphatase e | SALT00000027491_Cluster15031-001 |
| Cluster15050 | - | - | - | - | - | down | retrotransposon unclassified | SALT00000082693_Cluster15050-001 |
| Cluster1506 | - | up | - | - | - | - | RNA-binding 5-B isoform X1 | SALT00000087495_Cluster1506-007 |
| Cluster15065 | - | - | up | - | up | - | BEL1-like homeodomain 7 [S | SALT00000021798_Cluster15065-001 |
| Cluster15069 | - | - | - | - | down | - | Os06g0367900 [Oryza sativa | SALT00000068822_Cluster15069-002 |
| Cluster1507 | - | - | down | - | down | down | UPF0202 At1g10490-like [Se | SALT00000085834_Cluster1507-011 |
| Cluster15072 | - | - | - | - | - | up | subtilisin-like protease [Setari | SALT00000080476_Cluster15072-004 |
| Cluster15077 | - | up | up | - | - | - | serine hydroxymethyltransfera | SALT00000050255_Cluster15077-007 |
| Cluster15081 | - | - | down | - | - | - | reverse transcriptase | SALT00000065296_Cluster15081-001 |
| Cluster1510 | - | down | down | - | - | - | Os12g0277500 [Oryza sativa | SALT00000001302_Cluster1510-001 |
| Cluster15101 | - | - | up | - | up | up | serine carboxypeptidase-like 1 | SALT00000074883_Cluster15101-001 |
| Cluster15118 | - | - | up | - | up | up | Os03g0816800 [Oryza sativa | SALT00000079772_Cluster15118-001 |
| Cluster15121 | - | - | up | - | up | up | heat shock cognate 70 kDa | SALT00000060158_Cluster15121-002 |
| Cluster1513 | - | - | down | - | - | - | Os03g0158500 [Oryza sativa | SALT00000070745_Cluster1513-029 |
| Cluster1514 | - | - | up | - | - | - | Os02g0517300 [Oryza sativa | SALT00000091458_Cluster1514-007 |
| Cluster1515 | - | - | down | - | - | - | Os09g0408000 [Oryza sativa | SALT00000060741_Cluster1515-001 |
| Cluster15154 | - | - | down | - | - | - | soluble acid invertase [Sorghu | SALT00000031177_Cluster15154-001 |
| Cluster15155 | - | - | up | - | - | - | serine threonine- kinase SAPK | SALT00000003866_Cluster15155-001 |
| Cluster1516 | - | - | down | - | down | - | retrotransposon unclassified | SALT00000083859_Cluster1516-004 |
| Cluster15168 | - | - | up | - | up | up | DTM1_ORYSJ ame: Full=Sig | SALT00000028000_Cluster15168-001 |
| Cluster15178 | - | - | down | - | down | down | membrane-associated salt-ind | SALT00000049548_Cluster15178-001 |
| Cluster15181 | - | - | down | - | - | - | hypothetical protein SORBI_( | SALT00000063698_Cluster15181-001 |
| Cluster15183 | - | - | up | - | up | up | ORYA_ORYSJ ame: Full=Or | SALT00000065455_Cluster15183-001 |
| Cluster15207 | - | - | up | - | up | up | pirin At1g50590 | SALT00000070919_Cluster15207-001 |
| Cluster15212 | - | - | down | - | - | down | LUTEIN DEFICIENT chloroj | SALT00000077973_Cluster15212-002 |
| Cluster15219 | - | - | up | - | up | up | [Hordeum vulgare vulgare] | SALT00000084537_Cluster15219-006 |
| Cluster15228 | - | - | up | - | - | - | tubulin alpha chain | SALT00000076134_Cluster15228-003 |
| Cluster15231 | - | up | up | - | up | - | CTP synthase [Zea mays] | SALT00000076747_Cluster15231-002 |
| Cluster15236 | - | - | down | - | - | - | Os04g0528300 [Oryza sativa | SALT00000027225_Cluster15236-003 |
| Cluster15238 | - | - | - | - | up | - | [Hordeum vulgare vulgare] | SALT00000062083_Cluster15238-001 |
| Cluster15254 | - | - | up | - | - | up | Os03g0265600 [Oryza sativa | SALT00000030165_Cluster15254-005 |
| Cluster15259 | up | - | up | - | - | - | arogenate dehydratase prephei | SALT00000084342_Cluster15259-004 |
| Cluster1526 | - | - | - | - | down | - | potassium transporter 12 | SALT00000070438_Cluster1526-013 |
| Cluster15287 | - | - | - | - | - | down | pentatricopeptide repeat-conta | SALT00000077461_Cluster15287-001 |
| Cluster15288 | - | - | down | - | - | - | diphthine expressed [Oryza sa | SALT00000027600_Cluster15288-001 |
| Cluster15289 | - | - | down | - | - | - | transcription factor bHLH144 | SALT00000063250_Cluster15289-002 |
| Cluster15297 | - | - | down | - | down | down | heterogeneous nuclear ribonu | SALT00000076255_Cluster15297-002 |
| Cluster15298 | up | - | down | down | down | - | xyloglucan endotransglucosyl | SALT00000078605_Cluster15298-001 |
| Cluster15302 | - | - | up | - | up | - | Os08g0288500 [Oryza sativa | SALT00000022747_Cluster15302-002 |
| Cluster15314 | - | - | - | - | - | down | pollen-specific C13 [Brachypo | SALT00000079045_Cluster15314-001 |
| Cluster15320 | - | - | up | - | - | - | pre-rRNA-processing ESF1 [( | SALT00000029600_Cluster15320-004 |
| Cluster15334 | - | - | down | - | down | down | hydrolase [Zea mays] | SALT00000072889_Cluster15334-006 |
| Cluster15335 | - | - | down | - | down | down | N-lysine methyltransferase M | SALT00000072910_Cluster15335-003 |
| Cluster15336 | - | - | up | - | up | - | partial [Oryza sativa Japonica | SALT00000078155_Cluster15336-001 |
| Cluster15347 | - | - | down | - | - | - | phospholipid--sterol O-acyltra | SALT00000072527_Cluster15347-004 |
| Cluster1535 | - | - | up | - | up | up | zinc finger CCCH domain-co | SALT00000093089_Cluster1535-006 |

| | | | | | | | | |
|---|---|---|---|---|---|---|---|---|
| Cluster1536 | - | - | down | - | down | - | acetyl-coenzyme A chloroplas | SALT00000063442_Cluster1536-001 |
| Cluster1538 | - | - | down | - | - | - | NRT1 PTR FAMILY -like | SALT00000019697_Cluster1538-008 |
| Cluster15384 | - | - | - | - | - | up | Os03g0376900 [Oryza sativa | SALT00000082225_Cluster15384-002 |
| Cluster1539 | - | - | down | - | - | - | [Hordeum vulgare vulgare] | SALT00000070966_Cluster1539-002 |
| Cluster154 | - | - | down | - | - | - | vacuolar sorting-associated 53 | SALT00000071964_Cluster154-001 |
| Cluster15402 | - | - | down | - | down | - | BEL1-related homeotic 14 iso | SALT00000024311_Cluster15402-003 |
| Cluster15408 | - | - | down | - | - | - | formin 20 [Oryza brachyantha | SALT00000081548_Cluster15408-006 |
| Cluster15409 | - | up | - | - | - | - | glyceraldehyde-3-phosphate p | SALT00000081896_Cluster15409-006 |
| Cluster15412 | - | - | - | - | down | down | jacalin-related lectin 3-like | SALT00000084127_Cluster15412-005 |
| Cluster15417 | - | up | up | - | up | up | pepsin A [Zea mays] | SALT00000071997_Cluster15417-002 |
| Cluster15434 | - | - | down | - | down | - | phototropic-responsive NPH3 | SALT00000074149_Cluster15434-003 |
| Cluster15436 | - | - | - | - | up | - | serine threonine phosphatase r | SALT00000023801_Cluster15436-006 |
| Cluster15437 | - | - | down | - | down | - | cell division control 2 [Zea ma | SALT00000027730_Cluster15437-001 |
| Cluster15451 | - | - | down | - | - | - | retrotransposon Ty3-gypsy sul | SALT00000066467_Cluster15451-001 |
| Cluster15454 | - | - | - | - | down | - | subtilisin-like protease | SALT00000073388_Cluster15454-002 |
| Cluster1546 | - | - | up | - | up | - | receptor kinase 5 [Zea mays] | SALT00000086398_Cluster1546-007 |
| Cluster15469 | - | - | up | - | up | - | C2 domain-containing At1g53 | SALT00000027019_Cluster15469-001 |
| Cluster1547 | - | - | - | - | down | down | transcriptional corepressor LE | SALT00000083736_Cluster1547-018 |
| Cluster15477 | - | - | down | - | down | down | ---NA--- | SALT00000073036_Cluster15477-001 |
| Cluster15478 | - | - | up | - | up | up | NRT1 PTR FAMILY -like [Se | SALT00000061974_Cluster15478-010 |
| Cluster15488 | - | - | down | - | down | - | replication A 70 kDa DNA-bi | SALT00000027000_Cluster15488-001 |
| Cluster155 | - | - | - | - | - | down | photosystem I P700 apo A1 (c | SALT00000083257_Cluster155-002 |
| Cluster15500 | - | - | down | - | - | - | receptor expression-enhancing | SALT00000081992_Cluster15500-001 |
| Cluster15523 | - | - | down | - | - | - | ---NA--- | SALT00000030946_Cluster15523-004 |
| Cluster15534 | - | up | - | up | - | - | probable metal-nicotianamine | SALT00000030801_Cluster15534-002 |
| Cluster15542 | - | - | down | - | - | - | Os03g0304500 [Oryza sativa | SALT00000022047_Cluster15542-001 |
| Cluster15554 | - | - | up | - | up | up | cyclic dof factor 1-like [Setari | SALT00000078459_Cluster15554-017 |
| Cluster15564 | - | - | - | - | up | up | aspartic ase oryzasin-1 [Setari | SALT00000062890_Cluster15564-001 |
| Cluster15572 | - | up | up | - | up | up | [Hordeum vulgare vulgare] | SALT00000023183_Cluster15572-007 |
| Cluster15578 | - | down | down | down | down | - | Os07g0175300 [Oryza sativa | SALT00000073230_Cluster15578-001 |
| Cluster15581 | - | - | - | - | - | down | partial [Oryza sativa Japonica | SALT00000078543_Cluster15581-001 |
| Cluster156 | down | - | - | - | - | - | TSS [Setaria italica] | SALT00000034143_Cluster156-001 |
| Cluster15605 | - | - | down | - | - | - | TPA: DUF594 domain contair | SALT00000063596_Cluster15605-005 |
| Cluster15608 | up | up | - | - | - | - | F-box SKIP23-like | SALT00000076231_Cluster15608-003 |
| Cluster15618 | up | up | - | - | down | down | AC090486_29 retroelement [( | SALT00000072145_Cluster15618-002 |
| Cluster15627 | - | down | down | - | down | - | DEAD-box ATP-dependent R | SALT00000023022_Cluster15627-005 |
| Cluster15634 | - | - | up | - | - | - | B regulatory subunit of PP2A | SALT00000063149_Cluster15634-003 |
| Cluster15644 | - | up | up | - | up | - | ubiquitin carboxyl-terminal hy | SALT00000075364_Cluster15644-001 |
| Cluster15651 | - | - | - | - | - | down | Os11g0580800 [Oryza sativa | SALT00000067670_Cluster15651-002 |
| Cluster15652 | - | - | down | - | down | - | ABC transporter G family me | SALT00000067699_Cluster15652-002 |
| Cluster15674 | - | up | up | - | up | - | GATA transcription factor 8-li | SALT00000027185_Cluster15674-002 |
| Cluster157 | - | - | up | - | - | - | ubiquitin-associated (UBA) [( | SALT00000083432_Cluster157-039 |
| Cluster15704 | - | - | - | - | down | - | arginine N-methyltransferase | SALT00000067561_Cluster15704-001 |
| Cluster1571 | - | - | - | - | up | - | acetyl-coenzyme A carboxylas | SALT00000034112_Cluster1571-001 |
| Cluster15715 | - | - | up | - | up | - | probable esterase KAI2 | SALT00000080047_Cluster15715-001 |
| Cluster15726 | - | - | up | - | up | up | endoglucanase 1 precursor [Ze | SALT00000078740_Cluster15726-008 |
| Cluster15731 | - | down | down | down | - | - | soluble inorganic pyrophospha | SALT00000064178_Cluster15731-001 |
| Cluster15750 | down | down | down | - | - | - | Os03g0628800 [Oryza sativa | SALT00000079811_Cluster15750-004 |
| Cluster15754 | - | - | - | - | down | - | transporter arsB [Nicotiana tal | SALT00000023808_Cluster15754-003 |
| Cluster15772 | - | - | down | - | - | - | chaperone -like isoform X1 [E | SALT00000086194_Cluster15772-001 |
| Cluster15776 | - | - | down | - | - | - | actin-7 [Brachypodium distacl | SALT00000070343_Cluster15776-012 |
| Cluster1578 | - | - | - | - | - | up | U-box domain-containing 6 [S | SALT00000026715_Cluster1578-009 |
| Cluster15802 | - | - | down | - | - | - | actin-97-like [Zea mays] | SALT00000027680_Cluster15802-004 |
| Cluster15827 | - | - | down | - | down | - | 2-hydroxyisoflavanone dehyd | SALT00000075819_Cluster15827-001 |
| Cluster15839 | - | - | up | - | - | - | nucleic acid binding [Zea may | SALT00000081459_Cluster15839-005 |
| Cluster15843 | - | - | up | - | up | up | Os03g0362200 [Oryza sativa | SALT00000066853_Cluster15843-001 |
| Cluster15847 | - | up | up | - | up | up | protease [Oryza sativa Japonic | SALT00000072409_Cluster15847-002 |
| Cluster15852 | - | - | down | - | down | down | partial [Hordeum vulgare vulg | SALT00000077644_Cluster15852-007 |

| | | | | | | | | |
|---|---|---|---|---|---|---|---|---|
| Cluster1586 | - | - | down | - | - | - | E3 ubiquitin- ligase SINA 4 | SALT00000088298_Cluster1586-009 |
| Cluster15862 | - | - | up | - | - | - | Os08g0431100 [Oryza sativa | SALT00000065913_Cluster15862-003 |
| Cluster15864 | - | - | up | - | up | up | Os01g0227100 [Oryza sativa | SALT00000070483_Cluster15864-001 |
| Cluster15873 | - | - | - | - | down | down | macrophage erythroblast attac | SALT00000064812_Cluster15873-001 |
| Cluster15894 | - | - | up | - | - | - | Os01g0203800 [Oryza sativa | SALT00000083045_Cluster15894-006 |
| Cluster15924 | - | - | up | - | up | up | ankycorbin [Oryza sativa Japo | SALT00000024763_Cluster15924-001 |
| Cluster15938 | - | - | down | - | down | - | Os05g0530300 [Oryza sativa | SALT00000082303_Cluster15938-006 |
| Cluster15948 | - | - | - | - | down | - | SUPPRESSOR OF GENE SII | SALT00000076467_Cluster15948-003 |
| Cluster15952 | down | down | down | - | down | down | Os10g0503500 [Oryza sativa | SALT00000022276_Cluster15952-006 |
| Cluster1597 | - | - | down | - | down | - | structural maintenance of chro | SALT00000085603_Cluster1597-001 |
| Cluster15972 | - | up | up | - | up | up | O-succinylhomoserine sulfhyd | SALT00000054392_Cluster15972-004 |
| Cluster15981 | up | - | down | - | down | down | disease resistance response 20 | SALT00000081574_Cluster15981-003 |
| Cluster15988 | - | - | down | - | - | - | transmembrane 9 superfamily | SALT00000070457_Cluster15988-010 |
| Cluster15991 | - | down | down | - | - | - | metal-nicotianamine transport | SALT00000077473_Cluster15991-002 |
| Cluster15998 | - | - | up | - | up | - | PREDICTED: uncharacterized | SALT00000063031_Cluster15998-014 |
| Cluster160 | - | - | down | - | down | down | Os04g0442000 [Oryza sativa | SALT00000066353_Cluster160-001 |
| Cluster16002 | - | - | up | - | - | - | [Hordeum vulgare vulgare] | SALT00000071888_Cluster16002-002 |
| Cluster16004 | - | - | up | - | up | - | nicotianamine aminotransfera | SALT00000076272_Cluster16004-003 |
| Cluster16009 | - | - | up | - | - | - | CYPRO4 [Zea mays] | SALT00000065291_Cluster16009-001 |
| Cluster1601 | - | - | down | - | down | down | GPI-anchored [Setaria italica] | SALT00000087295_Cluster1601-008 |
| Cluster16013 | - | - | up | - | up | up | homeodomain-like transcriptio | SALT00000073099_Cluster16013-001 |
| Cluster1603 | - | - | down | - | down | - | alpha- partial [Populus nigra] | SALT00000048087_Cluster1603-001 |
| Cluster16034 | - | - | up | - | up | up | DEAD-box ATP-dependent R | SALT00000063393_Cluster16034-004 |
| Cluster16041 | - | - | down | - | down | down | UDP-glucose 6-dehydrogenas | SALT00000059064_Cluster16041-001 |
| Cluster16044 | - | up | - | - | - | - | stem-specific TSJT1-like | SALT00000067270_Cluster16044-002 |
| Cluster16060 | - | - | - | - | down | - | peptide transporter PTR2 [Zea | SALT00000071099_Cluster16060-003 |
| Cluster1607 | - | - | - | - | down | down | photosystem I P700 apo A1 (c | SALT00000079561_Cluster1607-001 |
| Cluster16071 | - | up | up | - | up | up | RCH2 [Oryza sativa Japonica | SALT00000058711_Cluster16071-001 |
| Cluster16080 | - | - | down | - | - | - | Os08g0379300 [Oryza sativa | SALT00000074535_Cluster16080-001 |
| Cluster1610 | - | - | down | - | - | - | tesmin TSO1-like CXC 2 | SALT00000021071_Cluster1610-003 |
| Cluster16119 | - | - | - | - | - | up | Os10g0521000 [Oryza sativa | SALT00000069046_Cluster16119-002 |
| Cluster16124 | - | - | up | - | up | - | aminotransferase y4uB [Zea m | SALT00000049446_Cluster16124-004 |
| Cluster16127 | - | up | up | - | up | - | FAR1-RELATED SEQUENC | SALT00000071153_Cluster16127-001 |
| Cluster16134 | - | - | - | - | - | down | ---NA--- | SALT00000082723_Cluster16134-001 |
| Cluster16155 | - | - | down | - | - | - | NRT1 PTR FAMILY -like [Se | SALT00000064738_Cluster16155-005 |
| Cluster16174 | - | - | up | - | - | - | dihydrolipoyllysine-residue su | SALT00000074372_Cluster16174-022 |
| Cluster16209 | - | - | up | - | - | - | citrate synthase family isoforn | SALT00000084442_Cluster16209-022 |
| Cluster1621 | - | - | - | - | - | up | TPA: trehalose phosphatase sy | SALT00000089029_Cluster1621-039 |
| Cluster16212 | - | - | - | - | - | down | PREDICTED: uncharacterized | SALT00000046847_Cluster16212-001 |
| Cluster16213 | - | - | up | - | up | up | equilibrative nucleotide transp | SALT00000053645_Cluster16213-007 |
| Cluster16239 | - | - | down | - | - | - | auxin response factor 8-like [S | SALT00000077143_Cluster16239-004 |
| Cluster16248 | - | - | - | - | down | - | stromal 70 kDa heat shock-rel | SALT00000027869_Cluster16248-004 |
| Cluster16250 | - | down | down | - | down | - | probable mediator of RNA po | SALT00000042147_Cluster16250-003 |
| Cluster16260 | - | - | up | - | up | up | amine oxidase flavin domain- | SALT00000067574_Cluster16260-003 |
| Cluster16267 | - | - | up | - | up | up | ATP binding [Zea mays] | SALT00000060222_Cluster16267-007 |
| Cluster1629 | - | - | down | - | down | down | probable LRR receptor-like se | SALT00000062767_Cluster1629-019 |
| Cluster16290 | - | - | down | - | down | - | deoxyribodipyrimidine photol | SALT00000074415_Cluster16290-001 |
| Cluster16300 | - | - | - | - | - | up | hypothetical protein SORBI_( | SALT00000029167_Cluster16300-002 |
| Cluster16302 | - | - | down | - | down | - | aspartic ase nepenthesin-1 pre | SALT00000057041_Cluster16302-008 |
| Cluster16305 | - | - | down | - | - | - | IQ-DOMAIN 1 [Oryza sativa | SALT00000063169_Cluster16305-004 |
| Cluster16308 | - | - | down | - | - | - | pyrophosphate--fructose 6-pho | SALT00000067585_Cluster16308-005 |
| Cluster16310 | - | - | up | - | up | up | probable inactive receptor kin | SALT00000070566_Cluster16310-002 |
| Cluster16315 | - | - | - | - | - | down | transcription factor APG-like | SALT00000022042_Cluster16315-003 |
| Cluster16333 | - | - | - | - | down | - | betaine aldehyde dehydrogena | SALT00000038744_Cluster16333-003 |
| Cluster16349 | - | - | down | - | down | - | Os01g0285300 [Oryza sativa | SALT00000062328_Cluster16349-001 |
| Cluster16357 | - | - | down | - | - | - | cis-2,3-dihydrobiphenyl-2,3-d | SALT00000081063_Cluster16357-004 |
| Cluster16358 | - | - | up | - | up | up | peptide transporter [Oryza sat | SALT00000082911_Cluster16358-005 |
| Cluster16362 | - | - | down | - | - | - | LMBR1 integral membrane fa | SALT00000039073_Cluster16362-001 |

| | | | | | | | | |
|---|---|---|---|---|---|---|---|---|
| Cluster16374 | - | - | - | - | down | - | fasciclin-like arabinogalactan- | SALT00000049155_Cluster16374-001 |
| Cluster1640 | up | up | - | - | down | - | 5-methyltetrahydropteroyltrig | SALT00000086763_Cluster1640-124 |
| Cluster16400 | - | - | up | - | up | - | beta-glucosidase-like chloropl | SALT00000075394_Cluster16400-002 |
| Cluster16417 | - | - | up | - | - | - | phospholipase A2 | SALT00000076539_Cluster16417-001 |
| Cluster16421 | - | - | - | - | - | up | uncharacterized plant-specific | SALT00000063895_Cluster16421-001 |
| Cluster16429 | - | down | down | - | - | - | Os03g0811100 [Oryza sativa . | SALT00000023916_Cluster16429-001 |
| Cluster16446 | - | - | up | - | - | up | Os02g0111600 [Oryza sativa . | SALT00000067991_Cluster16446-001 |
| Cluster16447 | - | - | down | - | - | down | Os03g0182700 [Oryza sativa | SALT00000069627_Cluster16447-001 |
| Cluster16460 | - | - | up | - | up | - | zinc finger (C3HC4-type RIN | SALT00000068273_Cluster16460-003 |
| Cluster16475 | - | - | up | - | up | up | boron transporter 4-like [Setai | SALT00000028933_Cluster16475-001 |
| Cluster16482 | - | - | up | - | up | up | [Hordeum vulgare vulgare] | SALT00000075159_Cluster16482-001 |
| Cluster165 | - | - | down | - | down | - | Os07g0252400 [Oryza sativa | SALT00000086245_Cluster165-001 |
| Cluster16519 | - | - | up | - | up | - | Os06g0656500 [Oryza sativa | SALT00000026541_Cluster16519-002 |
| Cluster16524 | - | - | - | - | up | - | PREDICTED: uncharacterize | SALT00000072763_Cluster16524-001 |
| Cluster16529 | - | - | up | - | up | up | phospho-2-dehydro-3-deoxyh | SALT00000059624_Cluster16529-020 |
| Cluster1653 | - | - | down | - | - | - | probable mediator of RNA po | SALT00000036513_Cluster1653-045 |
| Cluster16537 | - | - | up | - | - | - | Os05g0408300 [Oryza sativa | SALT00000077452_Cluster16537-005 |
| Cluster16540 | - | - | - | - | down | down | AP-2 complex subunit sigma | SALT00000084457_Cluster16540-002 |
| Cluster16543 | - | - | down | - | down | - | Os01g0913000 [Oryza sativa | SALT00000025875_Cluster16543-001 |
| Cluster16559 | - | - | down | - | - | - | zinc finger C-x8-C-x5-C-x3-I | SALT00000082559_Cluster16559-008 |
| Cluster16560 | - | - | up | - | up | up | PTI1-like tyrosine- kinase At3 | SALT00000082602_Cluster16560-004 |
| Cluster16583 | - | - | up | - | up | up | sucrose responsive element bi | SALT00000080165_Cluster16583-009 |
| Cluster1659 | up | up | up | - | up | up | delta 1-pyrroline-5-carboxylat | SALT00000086316_Cluster1659-032 |
| Cluster16591 | - | down | down | - | down | down | alanine aminotransferase | SALT00000084208_Cluster16591-018 |
| Cluster16599 | - | - | up | - | up | - | receptor homology transmeml | SALT00000023067_Cluster16599-002 |
| Cluster16600 | - | - | up | - | up | up | ribonuclease H At1g65750 | SALT00000063881_Cluster16600-001 |
| Cluster16608 | - | - | down | - | - | - | far upstream element-binding | SALT00000021575_Cluster16608-001 |
| Cluster16636 | - | - | down | - | down | - | partial [Hordeum vulgare vulg | SALT00000026186_Cluster16636-002 |
| Cluster1664 | - | - | up | - | up | up | sulfate transporter [Zea mays] | SALT00000051448_Cluster1664-009 |
| Cluster16643 | - | - | - | - | down | - | [Oryza sativa Indica Group] | SALT00000074292_Cluster16643-001 |
| Cluster16647 | - | - | up | - | - | - | cyclic nucleotide gated chann | SALT00000025691_Cluster16647-001 |
| Cluster16662 | - | up | up | up | up | - | asparagine synthetase [Zea ma | SALT00000022217_Cluster16662-002 |
| Cluster16687 | - | - | up | - | up | up | partial [Oryza sativa Japonica | SALT00000022949_Cluster16687-001 |
| Cluster16693 | - | - | down | - | down | - | Os03g0628800 [Oryza sativa | SALT00000066066_Cluster16693-003 |
| Cluster16700 | - | - | up | - | - | - | unknown [Zea mays] | SALT00000081218_Cluster16700-001 |
| Cluster16703 | - | - | down | - | - | - | partial [Hordeum vulgare vulg | SALT00000084669_Cluster16703-002 |
| Cluster16719 | - | up | up | up | up | up | S-adenosylmethionine decarb | SALT00000082587_Cluster16719-009 |
| Cluster16738 | down | - | up | - | up | up | CBL-interacting kinase 14 [O | SALT00000026818_Cluster16738-006 |
| Cluster16757 | - | - | - | - | up | - | SWI SNF complex subunit SV | SALT00000039487_Cluster16757-003 |
| Cluster1676 | - | - | up | - | up | up | sucrose synthase 1 [Zea mays | SALT00000068887_Cluster1676-001 |
| Cluster16777 | - | - | - | up | - | - | coronatine-insensitive 1 [Zea | SALT00000030265_Cluster16777-004 |
| Cluster16782 | - | - | up | - | - | - | dnaJ homolog subfamily C m | SALT00000021347_Cluster16782-002 |
| Cluster16785 | - | - | up | - | up | up | cytokinin oxidase 3 | SALT00000029646_Cluster16785-001 |
| Cluster16799 | - | down | down | - | down | down | fructokinase-2 [Zea mays] | SALT00000029151_Cluster16799-001 |
| Cluster16804 | - | - | down | - | - | - | Os10g0567400 [Oryza sativa | SALT00000075405_Cluster16804-009 |
| Cluster16810 | - | - | up | - | up | up | ankyrin repeat [Oryza sativa J | SALT00000027159_Cluster16810-001 |
| Cluster16813 | - | - | down | - | - | - | Os09g0258000 [Oryza sativa | SALT00000064234_Cluster16813-001 |
| Cluster16830 | - | - | up | - | up | - | acyltransferase chloroplastic | SALT00000082517_Cluster16830-001 |
| Cluster16832 | - | up | up | - | up | up | viviparous 14 [Sorghum bicol | SALT00000023342_Cluster16832-001 |
| Cluster16833 | down | down | down | - | down | - | oligopeptide transporter 3 | SALT00000024154_Cluster16833-001 |
| Cluster16837 | - | - | - | - | up | - | nucleotide binding [Zea mays | SALT00000060976_Cluster16837-003 |
| Cluster16839 | - | down | down | - | - | - | Os04g0117200 [Oryza sativa . | SALT00000071933_Cluster16839-003 |
| Cluster16850 | - | - | up | - | - | - | [Hordeum vulgare vulgare] | SALT00000067458_Cluster16850-004 |
| Cluster16854 | - | - | down | - | - | - | [Hordeum vulgare vulgare] | SALT00000024491_Cluster16854-002 |
| Cluster16859 | - | - | - | - | down | - | PREDICTED: uncharacterize | SALT00000071848_Cluster16859-001 |
| Cluster16860 | - | - | up | - | - | - | PREDICTED: uncharacterize | SALT00000075603_Cluster16860-001 |
| Cluster16864 | - | - | up | - | up | up | probable LRR receptor-like se | SALT00000025673_Cluster16864-004 |
| Cluster16873 | - | - | up | - | - | up | elicitor inducible beta-1,3-glu | SALT00000078846_Cluster16873-002 |

| | | | | | | | | |
|---|---|---|---|---|---|---|---|---|
| Cluster16888 | - | up | up | - | - | - | Os10g0482900 [Oryza sativa | SALT00000074381_Cluster16888-001 |
| Cluster16902 | - | - | down | - | down | down | [Hordeum vulgare vulgare] | SALT00000082484_Cluster16902-001 |
| Cluster16903 | - | down | down | - | - | - | probable inactive receptor kin | SALT00000030842_Cluster16903-003 |
| Cluster16914 | - | - | down | down | down | down | GDSL esterase lipase At5g37( | SALT00000054212_Cluster16914-003 |
| Cluster16920 | - | - | - | - | up | up | B regulatory subunit of PP2A | SALT00000073824_Cluster16920-003 |
| Cluster16928 | up | - | - | - | down | - | [Hordeum vulgare vulgare] | SALT00000042267_Cluster16928-005 |
| Cluster16930 | - | up | up | - | up | up | acetyl- cytosolic 2 [Zea mays] | SALT00000066687_Cluster16930-017 |
| Cluster16969 | - | - | up | - | up | - | WRKY DNA-binding domain | SALT00000022826_Cluster16969-003 |
| Cluster16995 | - | - | up | - | - | - | F-box kelch-repeat At1g7451( | SALT00000071016_Cluster16995-014 |
| Cluster16999 | - | - | - | - | - | down | N-alpha-acetyltransferase aux | SALT00000079835_Cluster16999-002 |
| Cluster17023 | up | up | up | - | - | - | 5-methyltetrahydropteroyltrig | SALT00000072805_Cluster17023-001 |
| Cluster17025 | - | - | up | - | - | - | cation H(+) antiporter 28 | SALT00000074311_Cluster17025-001 |
| Cluster17039 | - | - | down | - | down | down | COBRA 4 precursor | SALT00000065253_Cluster17039-004 |
| Cluster1704 | - | - | down | - | - | - | importin subunit beta-1-like [S | SALT00000064880_Cluster1704-006 |
| Cluster17048 | - | - | up | - | up | up | probable metal-nicotianamine | SALT00000079158_Cluster17048-004 |
| Cluster17059 | - | - | down | - | down | down | prosaposin-like isoform X1 [S | SALT00000076575_Cluster17059-001 |
| Cluster17060 | - | - | - | - | down | - | ---NA--- | SALT00000082408_Cluster17060-003 |
| Cluster17062 | - | - | down | - | - | - | Os04g0620700 [Oryza sativa | SALT00000025878_Cluster17062-002 |
| Cluster1707 | - | - | down | - | down | down | phosphatidylinositol 4-phosph | SALT00000070695_Cluster1707-011 |
| Cluster17073 | - | - | down | - | down | down | zeaxanthin 7,8(7 ,8 )-cleavage | SALT00000084198_Cluster17073-009 |
| Cluster17074 | - | - | down | - | - | - | formin 14 | SALT00000024473_Cluster17074-001 |
| Cluster17082 | - | up | up | - | - | - | [Hordeum vulgare vulgare] | SALT00000027894_Cluster17082-001 |
| Cluster17094 | - | - | up | - | - | - | PHD finger At1g33420 | SALT00000078270_Cluster17094-001 |
| Cluster17097 | - | - | - | - | up | up | Na+ H+ antiporter | SALT00000083069_Cluster17097-013 |
| Cluster17098 | - | - | up | - | up | up | probable acyl-activating enzy | SALT00000025430_Cluster17098-001 |
| Cluster1710 | - | - | down | - | down | down | serine threonine- kinase Nek5 | SALT00000091040_Cluster1710-001 |
| Cluster17111 | - | - | up | - | - | up | phosphatidylserine synthase 2 | SALT00000039761_Cluster17111-001 |
| Cluster17122 | - | - | up | - | up | up | cysteine-rich receptor kinase 1 | SALT00000066349_Cluster17122-001 |
| Cluster17138 | - | - | up | - | up | - | [Hordeum vulgare vulgare] | SALT00000021009_Cluster17138-004 |
| Cluster17152 | - | - | down | - | - | - | BTB POZ domain-containing | SALT00000064036_Cluster17152-006 |
| Cluster17153 | - | down | down | - | down | down | Os01g0896500 [Oryza sativa | SALT00000064233_Cluster17153-007 |
| Cluster17156 | - | up | up | - | up | up | Os10g0463800 [Oryza sativa | SALT00000070354_Cluster17156-002 |
| Cluster17159 | - | - | - | - | up | - | Os06g0320700 [Oryza sativa | SALT00000081807_Cluster17159-002 |
| Cluster1716 | - | - | down | - | down | - | histone-lysine N- H3 lysine-9 | SALT00000086533_Cluster1716-001 |
| Cluster17170 | - | - | up | - | up | up | holocarboxylase synthetase [Z | SALT00000067488_Cluster17170-002 |
| Cluster17181 | - | - | down | - | - | - | plastidial pyruvate kinase 2 | SALT00000069914_Cluster17181-004 |
| Cluster17191 | - | down | down | - | - | - | T-complex 1 subunit beta | SALT00000072624_Cluster17191-006 |
| Cluster1720 | - | - | up | - | up | - | iron inhibited ABC transporte | SALT00000072371_Cluster1720-033 |
| Cluster17203 | - | - | down | - | down | down | probable LRR receptor-like se | SALT00000084132_Cluster17203-003 |
| Cluster17209 | - | - | up | - | up | up | patellin-3-like isoform X1 [Br | SALT00000026773_Cluster17209-003 |
| Cluster1721 | - | - | up | - | up | up | cytochrome P450 71D7-like [ | SALT00000028521_Cluster1721-004 |
| Cluster17217 | - | - | up | - | - | - | hsp70-Hsp90 organizing -like | SALT00000025389_Cluster17217-007 |
| Cluster17220 | - | - | down | - | down | - | Os03g0831100 [Oryza sativa | SALT00000071032_Cluster17220-002 |
| Cluster17241 | - | - | - | - | - | down | ultraviolet-B receptor UVR8 | SALT00000069493_Cluster17241-004 |
| Cluster17247 | - | - | up | - | up | - | acyl-coenzyme A oxidase perc | SALT00000067255_Cluster17247-002 |
| Cluster17255 | - | - | down | - | - | - | auxin response factor 23 [Zea | SALT00000068873_Cluster17255-002 |
| Cluster17256 | - | - | - | - | - | down | HIPL1 [Nelumbo nucifera] | SALT00000072828_Cluster17256-002 |
| Cluster17264 | - | - | up | - | - | - | Os11g0490300 [Oryza sativa | SALT00000072792_Cluster17264-001 |
| Cluster1728 | up | up | up | - | up | up | xanthine dehydrogenase | SALT00000089298_Cluster1728-001 |
| Cluster17290 | - | - | up | - | up | up | PREDICTED: uncharacterized | SALT00000071818_Cluster17290-001 |
| Cluster17295 | - | - | - | - | - | down | FRIGIDA 4a [Zea mays] | SALT00000050096_Cluster17295-003 |
| Cluster1730 | - | - | up | - | up | up | Os04g0492400 [Oryza sativa | SALT00000038600_Cluster1730-001 |
| Cluster17304 | - | - | down | - | - | down | ATP-dependent Clp protease p | SALT00000024425_Cluster17304-010 |
| Cluster17331 | - | - | up | - | up | up | GDSL esterase lipase At5g45( | SALT00000029679_Cluster17331-003 |
| Cluster17333 | - | - | - | - | up | up | cell death-related SPL11 [Ory | SALT00000063080_Cluster17333-003 |
| Cluster17334 | - | - | down | - | - | - | Os03g0640100 [Oryza sativa | SALT00000027781_Cluster17334-012 |
| Cluster17343 | - | - | - | - | - | up | Os03g0394900 [Oryza sativa | SALT00000084185_Cluster17343-003 |
| Cluster17345 | up | up | up | - | - | - | glucose-6-phosphate isomeras | SALT00000041333_Cluster17345-001 |

| | | | | | | | | |
|---|---|---|---|---|---|---|---|---|
| Cluster17351 | - | - | up | - | up | - | [Hordeum vulgare vulgare] | SALT00000060673_Cluster17351-001 |
| Cluster17370 | - | - | up | - | - | - | alternative oxidase [Oryza sat | SALT00000071759_Cluster17370-001 |
| Cluster17374 | - | - | down | - | - | - | [Oryza sativa Japonica Group] | SALT00000029898_Cluster17374-002 |
| Cluster17386 | up | up | up | - | - | - | Os05g0363200 [Oryza sativa | SALT00000036159_Cluster17386-001 |
| Cluster17418 | - | - | - | - | down | - | guanine nucleotide-binding -li | SALT00000076510_Cluster17418-005 |
| Cluster17427 | - | - | down | - | down | - | ribosomal S7 (chloroplast) [A | SALT00000026684_Cluster17427-003 |
| Cluster17438 | - | up | up | - | - | - | [Hordeum vulgare vulgare] | SALT00000076779_Cluster17438-001 |
| Cluster17442 | - | - | up | - | - | - | 3 -N-debenzoyl-2 -deoxytaxol | SALT00000055968_Cluster17442-009 |
| Cluster1745 | - | - | down | - | down | - | tesmin TSO1-like CXC 2 isof | SALT00000080876_Cluster1745-004 |
| Cluster17451 | - | - | down | - | - | - | nucleolin-like [Brachypodium | SALT00000067587_Cluster17451-002 |
| Cluster17487 | - | - | up | - | up | up | equilibrative nucleotide transp | SALT00000067960_Cluster17487-006 |
| Cluster17503 | - | down | down | - | - | - | serine carboxypeptidase-like 5 | SALT00000062118_Cluster17503-009 |
| Cluster17512 | - | up | up | - | up | up | CBL-interacting kinase 14 [O | SALT00000038008_Cluster17512-001 |
| Cluster17517 | - | - | up | - | up | up | Os02g0709800 [Oryza sativa | SALT00000078134_Cluster17517-005 |
| Cluster17521 | - | - | - | - | up | - | 3-hydroxy-3-methylglutaryl-c | SALT00000066150_Cluster17521-005 |
| Cluster17524 | - | - | up | - | up | up | sugar phosphatase -like [Bracl | SALT00000018595_Cluster17524-002 |
| Cluster17528 | - | - | - | - | down | - | U-box domain-containing 15 | | SALT00000081817_Cluster17528-002 |
| Cluster17535 | - | - | up | - | up | - | alcohol partial | SALT00000058342_Cluster17535-001 |
| Cluster1754 | - | up | - | - | - | down | Glutamate decarboxylase | SALT00000014489_Cluster1754-013 |
| Cluster17548 | - | - | - | - | down | down | cyclin-dependent kinase F-4-l | SALT00000076004_Cluster17548-001 |
| Cluster17556 | - | - | - | - | down | down | Os03g0116500 [Oryza sativa | SALT00000061796_Cluster17556-013 |
| Cluster17569 | - | down | down | - | - | - | beta-1,3-galactosyltransferase | SALT00000057889_Cluster17569-001 |
| Cluster1759 | - | up | up | - | up | up | heat-shock 101 [Zea mays] | SALT00000073440_Cluster1759-025 |
| Cluster17606 | up | up | - | - | down | down | [Hordeum vulgare vulgare] | SALT00000014816_Cluster17606-001 |
| Cluster17610 | - | - | up | - | up | - | Os08g0545000 [Oryza sativa | SALT00000025952_Cluster17610-004 |
| Cluster17616 | - | up | up | - | - | - | signal recognition particle rec | SALT00000027432_Cluster17616-001 |
| Cluster1762 | - | - | - | - | down | - | [Hordeum vulgare vulgare] | SALT00000069305_Cluster1762-001 |
| Cluster17627 | - | down | - | - | - | - | Os03g0245700 [Oryza sativa | SALT00000059062_Cluster17627-003 |
| Cluster1763 | - | up | up | - | up | up | ATP-dependent Clp protease | SALT00000074128_Cluster1763-001 |
| Cluster17631 | - | - | up | - | - | - | Os05g0408300 [Oryza sativa | SALT00000083578_Cluster17631-007 |
| Cluster17634 | up | - | - | - | down | down | type IV inositol polyphosphat | SALT00000055565_Cluster17634-001 |
| Cluster17643 | - | - | up | - | up | up | [Hordeum vulgare vulgare] | SALT00000021287_Cluster17643-003 |
| Cluster17653 | - | - | down | - | down | - | probable inactive receptor kin | SALT00000083125_Cluster17653-002 |
| Cluster17656 | up | up | up | - | - | - | [Hordeum vulgare vulgare] | SALT00000045477_Cluster17656-004 |
| Cluster17665 | - | up | - | - | - | - | ADP-glucose pyrophosphoryl | SALT00000048752_Cluster17665-003 |
| Cluster17675 | - | - | - | - | up | - | serine threonine- kinase SRK2 | SALT00000054711_Cluster17675-001 |
| Cluster17677 | - | - | down | - | down | - | probable mediator of RNA po | SALT00000056354_Cluster17677-001 |
| Cluster17692 | - | - | up | - | - | up | oligopeptide transporter 4-like | SALT00000080010_Cluster17692-001 |
| Cluster17704 | - | - | down | - | - | - | MYB transcription partial [Ze | SALT00000024060_Cluster17704-008 |
| Cluster17710 | - | - | up | - | up | - | Aldehyde dehydrogenase fam | SALT00000040511_Cluster17710-001 |
| Cluster17721 | - | - | up | - | - | - | U-box domain-containing 18- | SALT00000072569_Cluster17721-002 |
| Cluster17761 | - | - | up | - | up | up | Os10g0442000 [Oryza sativa | SALT00000026000_Cluster17761-001 |
| Cluster17767 | up | up | up | - | up | up | [Hordeum vulgare vulgare] | SALT00000038897_Cluster17767-001 |
| Cluster17774 | - | - | up | - | up | up | Os04g0685600 [Oryza sativa | SALT00000071837_Cluster17774-002 |
| Cluster1778 | - | - | up | - | up | - | calcium-transporting ATPase | SALT00000092182_Cluster1778-026 |
| Cluster17780 | - | - | - | - | down | down | sanguinarine reductase [Setari | SALT00000046843_Cluster17780-003 |
| Cluster17781 | - | - | - | down | down | - | NADP-dependant malate part | SALT00000054292_Cluster17781-001 |
| Cluster17782 | up | - | - | - | - | - | Os05g0127200 [Oryza sativa | SALT00000056904_Cluster17782-003 |
| Cluster17788 | - | - | down | - | down | - | auxin efflux carrier [Zea mays | SALT00000045072_Cluster17788-003 |
| Cluster1779 | up | up | up | - | up | up | phosphatase 2C [Oryza sativa | SALT00000007604_Cluster1779-002 |
| Cluster17799 | - | - | - | - | - | down | Regulator of Vps4 activity in | SALT00000082183_Cluster17799-002 |
| Cluster178 | down | - | - | - | - | - | Os04g0660500 [Oryza sativa | SALT00000092172_Cluster178-018 |
| Cluster17805 | - | - | up | - | up | up | serine threonine- kinase SAPK | SALT00000043753_Cluster17805-003 |
| Cluster17816 | - | - | up | - | - | - | Secretory carrier-associated m | SALT00000084183_Cluster17816-001 |
| Cluster17821 | - | - | - | - | down | - | transport inhibitor response 1 | SALT00000065753_Cluster17821-003 |
| Cluster17826 | - | - | up | - | up | up | aminotransferase y4uB [Zea n | SALT00000061459_Cluster17826-001 |
| Cluster17833 | down | down | down | - | down | - | Os01g0915600 [Oryza sativa | SALT00000074978_Cluster17833-007 |
| Cluster17840 | - | - | up | - | - | - | acyl carrier 3 [Zea mays] | SALT00000083464_Cluster17840-003 |

| | | | | | | | | |
|---|---|---|---|---|---|---|---|---|
| Cluster17843 | - | - | - | - | up | up | arm repeat [Oryza sativa Japo | SALT00000067514_Cluster17843-003 |
| Cluster17849 | - | - | down | - | down | down | 4-hydroxyphenylacetaldehyde | SALT00000055307_Cluster17849-007 |
| Cluster17856 | up | - | - | - | down | down | RNA polymerase sigma factor | SALT00000079648_Cluster17856-003 |
| Cluster17869 | - | - | up | - | up | up | NAC domain transcription fac | SALT00000028041_Cluster17869-003 |
| Cluster1787 | - | - | down | - | down | - | beta-adaptin C | SALT00000078883_Cluster1787-019 |
| Cluster17873 | - | - | up | - | - | - | ultraviolet-B receptor UVR8 i | SALT00000049885_Cluster17873-003 |
| Cluster179 | - | - | up | - | up | up | copper methylamine oxidase | SALT00000068765_Cluster179-002 |
| Cluster1790 | - | - | up | - | up | up | Os06g0716000 [Oryza sativa | SALT00000087492_Cluster1790-003 |
| Cluster17913 | - | - | up | - | - | - | cactin [Setaria italica] | SALT00000070484_Cluster17913-002 |
| Cluster17914 | - | - | down | - | down | down | NADH dehydrogenase partial | SALT00000076028_Cluster17914-003 |
| Cluster17920 | - | - | - | - | - | up | peroxisomal membrane PEX1 | SALT00000066378_Cluster17920-001 |
| Cluster17934 | - | - | up | - | up | up | sodium-coupled neutral aminc | SALT00000060127_Cluster17934-009 |
| Cluster17941 | - | up | up | - | up | up | nitronate monooxygenase | SALT00000054457_Cluster17941-001 |
| Cluster17948 | - | - | - | - | down | down | Translation initiation factor eI | SALT00000064608_Cluster17948-001 |
| Cluster17949 | - | - | down | - | down | down | Os04g0602800 [Oryza sativa | SALT00000071646_Cluster17949-001 |
| Cluster17955 | - | - | down | - | - | down | CBS domain-containing [Sacc | SALT00000026377_Cluster17955-001 |
| Cluster17966 | - | up | - | up | - | - | GDA2 [Oryza sativa Japonica | SALT00000024423_Cluster17966-002 |
| Cluster17994 | - | down | down | - | down | - | Os05g0568000 [Oryza sativa | SALT00000055355_Cluster17994-011 |
| Cluster17997 | - | down | down | - | - | - | glycosyltransferase family 64 | SALT00000071424_Cluster17997-001 |
| Cluster18004 | - | - | up | - | up | up | actin-depolymerizing factor 5 | SALT00000088161_Cluster18004-001 |
| Cluster18017 | - | - | down | - | - | - | calreticulin-3 precursor [Zea r | SALT00000061121_Cluster18017-010 |
| Cluster18019 | - | - | up | - | - | - | cytochrome P450 reductase [( | SALT00000078679_Cluster18019-001 |
| Cluster1802 | - | - | down | - | - | - | PREDICTED: uncharacterize | SALT00000080143_Cluster1802-020 |
| Cluster18025 | - | - | up | - | up | up | indole-3-acetic acid amido syi | SALT00000063318_Cluster18025-002 |
| Cluster18032 | - | - | down | - | - | - | 6-phosphogluconate decarbox | SALT00000059523_Cluster18032-002 |
| Cluster18046 | up | - | up | - | - | - | Os04g0628100 [Oryza sativa | SALT00000027013_Cluster18046-004 |
| Cluster18053 | - | - | up | - | - | - | Myb-related Myb4 | SALT00000059558_Cluster18053-002 |
| Cluster18054 | - | - | down | - | down | - | probable WRKY transcription | SALT00000061480_Cluster18054-003 |
| Cluster18059 | - | - | down | - | - | - | LRR receptor-like serine threc | SALT00000037227_Cluster18059-002 |
| Cluster18062 | - | - | - | - | up | - | rac GTPase activating 1 | SALT00000023830_Cluster18062-003 |
| Cluster18067 | - | - | down | - | - | - | [Hordeum vulgare vulgare] | SALT00000055727_Cluster18067-002 |
| Cluster18083 | - | - | up | - | up | up | [Hordeum vulgare vulgare] | SALT00000017698_Cluster18083-001 |
| Cluster18084 | - | - | up | - | up | up | Os06g0143400 [Oryza sativa | SALT00000061753_Cluster18084-010 |
| Cluster18101 | - | - | - | - | down | - | glucan endo-1,3-beta-glucosic | SALT00000052472_Cluster18101-001 |
| Cluster18102 | - | - | up | - | up | up | Os12g0158700 [Oryza sativa | SALT00000054108_Cluster18102-001 |
| Cluster18107 | - | - | up | - | - | - | potassium-dependent sodium- | SALT00000046259_Cluster18107-001 |
| Cluster18108 | down | down | down | - | down | - | Os02g0675700 [Oryza sativa | SALT00000047667_Cluster18108-004 |
| Cluster18117 | - | - | up | - | up | - | [Hordeum vulgare vulgare] | SALT00000052263_Cluster18117-004 |
| Cluster18118 | - | - | - | - | down | - | choline-phosphate cytidylyltra | SALT00000013878_Cluster18118-007 |
| Cluster1812 | - | up | up | - | up | - | Os07g0672500 [Oryza sativa | SALT00000090842_Cluster1812-001 |
| Cluster18120 | - | - | down | - | - | - | trihelix transcription factor G | SALT00000067580_Cluster18120-001 |
| Cluster18133 | - | - | down | - | - | - | Os04g0483600 [Oryza sativa | SALT00000008266_Cluster18133-001 |
| Cluster18139 | - | - | down | - | down | - | ruvB 1 [Oryza sativa Japonica | SALT00000055805_Cluster18139-007 |
| Cluster18142 | - | - | up | - | - | up | lecithin-cholesterol acyltransf | SALT00000012091_Cluster18142-001 |
| Cluster18163 | - | - | up | - | up | up | Os09g0536700 [Oryza sativa | SALT00000070381_Cluster18163-002 |
| Cluster18207 | - | up | up | - | - | - | sucrose transporter | SALT00000037676_Cluster18207-005 |
| Cluster18212 | down | - | - | - | - | - | Os04g0660400 [Oryza sativa | SALT00000054141_Cluster18212-017 |
| Cluster18216 | - | - | up | - | up | - | ATP-dependent RNA helicase | SALT00000045295_Cluster18216-003 |
| Cluster18225 | - | - | up | - | up | - | IST1 [Zea mays] | SALT00000018802_Cluster18225-001 |
| Cluster18233 | - | - | up | - | - | - | LOV domain-containing | SALT00000007425_Cluster18233-004 |
| Cluster18236 | up | up | - | - | down | down | CBL-interacting kinase family | SALT00000049552_Cluster18236-001 |
| Cluster1824 | - | up | - | - | - | - | rab3 GTPase-activating cataly | SALT00000085345_Cluster1824-001 |
| Cluster18240 | - | - | down | - | - | down | microtubule-associated MAP6 | SALT00000013046_Cluster18240-006 |
| Cluster18252 | - | - | - | - | - | up | outer envelope chloroplastic | SALT00000068502_Cluster18252-002 |
| Cluster1826 | - | - | down | - | - | - | Os02g0738900 [Oryza sativa | SALT00000078159_Cluster1826-013 |
| Cluster18260 | - | - | up | - | - | - | Os01g0764900 [Oryza sativa | SALT00000010497_Cluster18260-001 |
| Cluster18263 | - | - | up | - | up | up | copine III-like [Oryza sativa J | SALT00000042113_Cluster18263-017 |
| Cluster18266 | - | - | - | - | up | - | MAC Perforin domain contair | SALT00000007747_Cluster18266-001 |

| | | | | | | | | |
|---|---|---|---|---|---|---|---|---|
| Cluster18273 | - | down | - | - | - | - | ABC transporter G family me | SALT00000037555_Cluster18273-001 |
| Cluster18290 | - | - | up | - | up | - | Os09g0492700 [Oryza sativa | SALT00000012124_Cluster18290-003 |
| Cluster183 | - | down | down | - | down | - | TPA: glycosyl transferase fam | SALT00000091281_Cluster183-027 |
| Cluster18301 | - | up | up | up | up | up | Os10g0471100 [Oryza sativa | SALT00000066217_Cluster18301-003 |
| Cluster18305 | - | - | up | - | up | up | Luminal-binding 5 | SALT00000081089_Cluster18305-001 |
| Cluster18308 | - | - | - | - | down | - | partial [Oryza sativa Japonica | SALT00000050276_Cluster18308-001 |
| Cluster18310 | - | - | up | - | up | - | endoglucanase 1 precursor [Z | SALT00000056639_Cluster18310-004 |
| Cluster18317 | - | - | up | - | up | up | ABA 8 -hydroxylase [Triticun | SALT00000012960_Cluster18317-001 |
| Cluster1832 | - | - | up | - | up | up | arginine decarboxylase 1 | SALT00000062153_Cluster1832-007 |
| Cluster18322 | - | - | up | - | - | - | NAC domain-containing 21 2 | SALT00000018382_Cluster18322-001 |
| Cluster18325 | - | - | - | - | down | - | cinnamoyl- reductase | SALT00000052881_Cluster18325-001 |
| Cluster18326 | - | - | up | - | up | up | signal recognition particle rec | SALT00000001420_Cluster18326-001 |
| Cluster18328 | - | - | up | - | - | - | Os02g0704600 [Oryza sativa | SALT00000033475_Cluster18328-016 |
| Cluster18331 | - | - | up | - | up | up | lysine-specific histone demeth | SALT00000040887_Cluster18331-009 |
| Cluster18337 | - | - | down | - | - | down | mannan endo-1,4-beta-manno | SALT00000075690_Cluster18337-013 |
| Cluster18341 | - | - | - | - | down | - | [Oryza sativa Japonica Group | SALT00000040011_Cluster18341-002 |
| Cluster18344 | - | - | down | - | down | down | ROOT PRIMORDIUM DEFE | SALT00000047343_Cluster18344-001 |
| Cluster18346 | - | - | down | - | - | - | G-type lectin S-receptor-like s | SALT00000060476_Cluster18346-001 |
| Cluster18349 | - | down | down | - | - | - | NRT1 PTR FAMILY -like | SALT00000057282_Cluster18349-001 |
| Cluster1836 | - | - | - | - | up | up | Os12g0573200 [Oryza sativa | SALT00000062807_Cluster1836-011 |
| Cluster18361 | - | - | up | - | up | up | chitin-inducible gibberellin-re | SALT00000017466_Cluster18361-002 |
| Cluster1837 | - | - | up | - | - | - | MODIFIER OF SNC1 1-like | | SALT00000079843_Cluster1837-007 |
| Cluster18372 | - | - | down | - | - | - | Os06g0496000 [Oryza sativa | SALT00000001761_Cluster18372-001 |
| Cluster18397 | - | - | down | - | down | down | Os03g0857400 [Oryza sativa | SALT00000057276_Cluster18397-004 |
| Cluster18398 | - | - | up | - | up | up | coupled to transmembrane mc | SALT00000038632_Cluster18398-001 |
| Cluster18402 | - | - | up | - | up | up | glycosyltransferase [Zea mays | SALT00000066028_Cluster18402-008 |
| Cluster18408 | - | down | down | - | - | - | granule-bound starch synthase | SALT00000010617_Cluster18408-002 |
| Cluster18414 | down | down | down | down | - | up | aminotransferase y4uB [Zea n | SALT00000052192_Cluster18414-009 |
| Cluster18430 | - | - | up | - | - | up | Os07g0130200 [Oryza sativa | SALT00000025112_Cluster18430-002 |
| Cluster18432 | - | - | up | - | up | up | HGA1 [Oryza sativa Japonica | SALT00000039762_Cluster18432-007 |
| Cluster18438 | - | - | up | - | up | up | Os12g0259800 [Oryza sativa | SALT00000036462_Cluster18438-007 |
| Cluster18443 | - | - | down | - | - | - | uncharacterized RNA methylt | SALT00000002353_Cluster18443-001 |
| Cluster1845 | - | - | up | - | - | - | TSL-kinase interacting 1 [Ory | SALT00000076856_Cluster1845-031 |
| Cluster18482 | - | - | - | - | up | - | Os01g0138300 [Oryza sativa | SALT00000056799_Cluster18482-002 |
| Cluster18499 | - | - | - | - | - | up | sodium calcium exchanger fai | SALT00000053800_Cluster18499-001 |
| Cluster1850 | - | - | up | - | up | - | RNA Binding 45 [Oryza sativ | SALT00000022378_Cluster1850-006 |
| Cluster18500 | - | - | down | - | - | - | Os03g0308800 [Oryza sativa | SALT00000054607_Cluster18500-001 |
| Cluster18502 | - | - | down | - | down | - | ru large subunit-binding subu | SALT00000022574_Cluster18502-002 |
| Cluster18506 | - | - | up | - | up | up | NAC2 [Oryza sativa Japonica | SALT00000025889_Cluster18506-004 |
| Cluster18515 | - | - | up | - | up | up | hypersensitive-induced respor | SALT00000040210_Cluster18515-004 |
| Cluster18529 | - | - | - | - | down | - | lysine--tRNA chloroplastic mi | SALT00000041555_Cluster18529-003 |
| Cluster18534 | - | - | up | - | up | up | F-box LRR-repeat 3-like | SALT00000076260_Cluster18534-001 |
| Cluster18536 | up | up | up | - | down | down | S-adenosyl-L-methionine: pho | SALT00000012703_Cluster18536-004 |
| Cluster18537 | - | - | up | - | up | up | probable bifunctional riboflav | SALT00000018627_Cluster18537-004 |
| Cluster1854 | - | - | up | - | up | up | F-box LRR-repeat MAX2 hor | SALT00000070167_Cluster1854-014 |
| Cluster18540 | - | - | up | - | - | - | symbiosis-related [Oryza sativ | SALT00000011370_Cluster18540-004 |
| Cluster18547 | - | - | up | - | up | - | F-box PP2-A13-like [Zea may | SALT00000055229_Cluster18547-006 |
| Cluster1855 | - | up | - | - | - | - | ---NA--- | SALT00000047775_Cluster1855-001 |
| Cluster18550 | - | - | up | - | up | up | NAC2 [Oryza sativa Japonica | SALT00000023096_Cluster18550-003 |
| Cluster18551 | - | down | down | down | down | down | tubulin alpha-1 chain [Miscan | SALT00000036741_Cluster18551-001 |
| Cluster18555 | - | up | - | - | - | - | ATP sulfurylase 2 | SALT00000049420_Cluster18555-001 |
| Cluster1858 | - | - | - | - | down | - | glutathione s-transferase | SALT00000088533_Cluster1858-002 |
| Cluster18580 | up | up | up | - | - | - | S-adenosylmethionine synthas | SALT00000041308_Cluster18580-029 |
| Cluster18581 | - | - | up | - | - | up | ethanolamine-phosphate cytid | SALT00000053394_Cluster18581-009 |
| Cluster18593 | - | up | up | - | - | - | Os01g0742500 [Oryza sativa | SALT00000058241_Cluster18593-002 |
| Cluster18602 | - | - | - | - | - | down | Os05g0366300 [Oryza sativa | SALT00000001126_Cluster18602-001 |
| Cluster18606 | - | - | up | - | up | up | zinc finger CONSTANS-LIKI | SALT00000060377_Cluster18606-003 |
| Cluster18607 | - | - | up | - | - | up | glycosyltransferase [Zea mays | SALT00000066903_Cluster18607-010 |

| | | | | | | | | |
|---|---|---|---|---|---|---|---|---|
| Cluster18618 | up | up | up | up | up | up | low-temperature-induced 65 k | SALT00000008593_Cluster18618-001 |
| Cluster18624 | - | - | down | - | - | - | UDP-N-acetylglucosamine py | SALT00000070850_Cluster18624-013 |
| Cluster18630 | - | - | - | - | - | up | Os03g0809300 [Oryza sativa | SALT00000046975_Cluster18630-015 |
| Cluster18652 | - | - | up | - | - | - | dnaJ ERDJ3A isoform X2 [Br | SALT00000059826_Cluster18652-005 |
| Cluster18656 | - | - | up | - | - | up | NRT1 PTR FAMILY -like [Se | SALT00000003918_Cluster18656-001 |
| Cluster18658 | - | - | down | - | - | - | phospholipase C [Zea mays] | SALT00000037567_Cluster18658-002 |
| Cluster18659 | - | up | up | - | - | - | phosphatase 2C | SALT00000039521_Cluster18659-001 |
| Cluster18666 | - | - | down | - | down | - | NSP-INTERACTING KINAS | SALT00000039928_Cluster18666-001 |
| Cluster1868 | - | - | - | - | down | - | chromomethylase [Zea mays] | SALT00000092001_Cluster1868-005 |
| Cluster18689 | - | - | up | - | - | - | ultraviolet-B receptor UVR8 [ | SALT00000055823_Cluster18689-001 |
| Cluster1870 | - | - | down | - | down | down | COBRA 4 precursor | SALT00000091034_Cluster1870-002 |
| Cluster18703 | - | - | up | - | - | up | Os06g0656500 [Oryza sativa | SALT00000018282_Cluster18703-006 |
| Cluster18706 | up | - | up | - | up | up | Os07g0638100 [Oryza sativa | SALT00000000284_Cluster18706-001 |
| Cluster18710 | - | - | up | - | up | up | [Hordeum vulgare vulgare] | SALT00000062611_Cluster18710-009 |
| Cluster18714 | - | - | up | - | up | - | IWS1 C-terminus family [Zea | SALT00000076138_Cluster18714-015 |
| Cluster18715 | - | - | - | - | up | - | DEAD-box ATP-dependent R | SALT00000039201_Cluster18715-003 |
| Cluster1872 | - | - | - | - | down | - | Os08g0531200 [Oryza sativa | SALT00000078872_Cluster1872-004 |
| Cluster18724 | - | - | up | - | - | - | Os05g0579000 [Oryza sativa | SALT00000045854_Cluster18724-008 |
| Cluster18725 | - | - | - | - | - | up | uncharacterized transporter Y | SALT00000052107_Cluster18725-011 |
| Cluster18726 | - | - | - | - | - | up | Os09g0565400 [Oryza sativa | SALT00000081072_Cluster18726-002 |
| Cluster18733 | - | - | down | - | - | down | chaperonin CPN60- mitochon | SALT00000049858_Cluster18733-004 |
| Cluster18735 | - | - | - | - | down | - | phosphatase 2C containing ex | SALT00000052421_Cluster18735-003 |
| Cluster18737 | - | - | - | - | up | up | MATE transporter 2 [Zea may | SALT00000062512_Cluster18737-014 |
| Cluster18747 | - | - | up | - | - | - | 2-hydroxyphytanoyl- lyase [Z | SALT00000001551_Cluster18747-001 |
| Cluster18754 | - | - | up | - | - | - | heterogeneous nuclear ribonu | SALT00000006364_Cluster18754-001 |
| Cluster18766 | - | - | - | - | up | up | kinase precursor [Zea mays] | SALT00000053194_Cluster18766-001 |
| Cluster18774 | - | - | up | - | up | up | lipase class 3 [Oryza sativa Ja | SALT00000009341_Cluster18774-001 |
| Cluster18776 | - | - | down | - | - | - | chaperonin CPN60- mitochon | SALT00000012631_Cluster18776-002 |
| Cluster18781 | - | up | down | - | down | down | phosphoethanolamine N-meth | SALT00000059494_Cluster18781-006 |
| Cluster18783 | - | - | down | - | - | down | cyp450 [Hordeum vulgare] | SALT00000068622_Cluster18783-005 |
| Cluster18792 | - | up | up | - | up | up | formin 7 | SALT00000007155_Cluster18792-003 |
| Cluster18801 | - | - | down | - | down | down | serine mitochondrial [Setaria i | SALT00000055641_Cluster18801-007 |
| Cluster18822 | - | - | - | - | - | up | Calmodulin [Glycine soja] | SALT00000002168_Cluster18822-001 |
| Cluster18833 | - | - | down | - | - | - | serine threonine- kinase Nek4 | SALT00000001706_Cluster18833-001 |
| Cluster18842 | - | - | down | - | down | down | TPA: protochlorophyllide redu | SALT00000057244_Cluster18842-001 |
| Cluster18847 | - | - | down | - | down | - | cell division cycle cofactor of | SALT00000002518_Cluster18847-001 |
| Cluster18848 | - | - | down | - | - | - | DEAD-box ATP-dependent R | SALT00000011855_Cluster18848-001 |
| Cluster18863 | - | down | down | - | down | - | Os07g0633600 [Oryza sativa | SALT00000042532_Cluster18863-010 |
| Cluster18868 | - | - | up | - | up | up | Os05g0439400 [Oryza sativa | SALT00000061007_Cluster18868-001 |
| Cluster18872 | - | - | up | - | up | up | nodule-specific [Oryza sativa | SALT00000010957_Cluster18872-002 |
| Cluster18874 | - | - | - | - | - | down | ankyrin-kinase [Oryza sativa . | SALT00000043986_Cluster18874-002 |
| Cluster18882 | up | up | up | - | - | - | phosphoethanolamine N-meth | SALT00000019782_Cluster18882-007 |
| Cluster18887 | - | - | up | - | up | - | aldehyde dehydrogenase fami | SALT00000053650_Cluster18887-003 |
| Cluster189 | - | - | down | - | down | down | brassinosteroid biosynthesis [. | SALT00000050494_Cluster189-010 |
| Cluster18904 | - | up | up | up | up | - | aldehyde dehydrogenase fami | SALT00000058988_Cluster18904-001 |
| Cluster18909 | - | - | down | - | down | down | Os03g0352300 [Oryza sativa | SALT00000035701_Cluster18909-001 |
| Cluster18922 | - | - | up | - | up | up | external alternative NAD(P)H | SALT00000058125_Cluster18922-002 |
| Cluster18931 | - | - | - | - | down | - | 3-oxoacyl-[acyl-carrier- ] synt | SALT00000016786_Cluster18931-002 |
| Cluster18933 | - | - | up | - | up | up | pectinesterase [Oryza sativa Ja | SALT00000042443_Cluster18933-005 |
| Cluster18947 | - | - | down | - | - | - | [Hordeum vulgare vulgare] | SALT00000007375_Cluster18947-001 |
| Cluster18952 | - | - | down | - | down | down | ZIP [Oryza sativa Japonica Gr | SALT00000042937_Cluster18952-006 |
| Cluster18959 | - | - | up | - | up | up | nitrate and chloride transporte | SALT00000003642_Cluster18959-001 |
| Cluster18961 | - | - | up | - | up | - | ABA response element bindin | SALT00000006106_Cluster18961-001 |
| Cluster18977 | - | - | down | - | down | down | Os09g0457100 [Oryza sativa | SALT00000070481_Cluster18977-001 |
| Cluster18979 | down | - | - | - | - | - | TPA: O-glycosyl hydrolase fa | SALT00000017410_Cluster18979-005 |
| Cluster18981 | - | up | up | - | up | up | Os08g0114300 [Oryza sativa . | SALT00000042074_Cluster18981-003 |
| Cluster18987 | - | - | up | - | - | up | Os06g0138200 [Oryza sativa | SALT00000006925_Cluster18987-002 |
| Cluster18988 | - | - | - | - | - | up | actin-depolymerizing factor 5 | SALT00000014165_Cluster18988-001 |

| | | | | | | | | |
|---|---|---|---|---|---|---|---|---|
| Cluster18989 | - | - | down | - | down | down | translation initiation factor IF- | SALT00000018172_Cluster18989-001 |
| Cluster18996 | - | - | down | - | down | down | UNC93 1 [Setaria italica] | SALT00000055930_Cluster18996-005 |
| Cluster19002 | - | - | up | - | up | - | Os01g0725600 [Oryza sativa | SALT00000014105_Cluster19002-002 |
| Cluster19014 | - | - | down | - | - | - | 60S ribosomal L12 [Zea mays | SALT00000048503_Cluster19014-001 |
| Cluster19016 | - | - | - | - | down | - | lipase [Oryza sativa Japonica | SALT00000066088_Cluster19016-002 |
| Cluster1902 | - | - | down | - | - | - | Os06g0164000 [Oryza sativa | SALT00000092360_Cluster1902-012 |
| Cluster19027 | - | - | down | - | down | - | Os09g0533900 [Oryza sativa | SALT00000007235_Cluster19027-003 |
| Cluster19030 | - | - | up | - | up | up | VTC2 [Zea mays] | SALT00000028120_Cluster19030-004 |
| Cluster19040 | - | down | down | - | - | - | cytochrome P450 77A3 | SALT00000005632_Cluster19040-003 |
| Cluster19048 | - | - | - | - | - | down | peptide transporter PTR2 [Zea | SALT00000053442_Cluster19048-004 |
| Cluster19059 | - | up | up | - | up | up | Os01g0253900 [Oryza sativa | SALT00000013161_Cluster19059-001 |
| Cluster19071 | - | - | up | - | up | up | aspartic ase 1 | SALT00000046620_Cluster19071-005 |
| Cluster19073 | - | down | down | - | down | down | elongation factor Tu [Zea may | SALT00000058635_Cluster19073-009 |
| Cluster19087 | - | - | - | - | - | down | O-glucosyltransferase rumi-lil | SALT00000056274_Cluster19087-003 |
| Cluster19088 | - | - | - | - | down | - | Os02g0138000 [Oryza sativa | SALT00000064963_Cluster19088-003 |
| Cluster19093 | - | - | up | - | up | up | 2OG-Fe oxygenase family exp | SALT00000052932_Cluster19093-001 |
| Cluster19103 | - | - | - | - | - | down | clustered mitochondria -like is | SALT00000040003_Cluster19103-004 |
| Cluster19105 | - | - | - | - | - | down | ABI3-interacting 2 [Zea mays | SALT00000042714_Cluster19105-016 |
| Cluster19109 | down | down | down | down | down | - | Os01g0655500 [Oryza sativa | SALT00000064575_Cluster19109-002 |
| Cluster19122 | - | - | down | - | - | - | T-complex 1 subunit eta | SALT00000040968_Cluster19122-010 |
| Cluster19128 | - | - | down | - | - | - | Os05g0566500 [Oryza sativa | SALT00000018453_Cluster19128-002 |
| Cluster19137 | up | - | - | - | - | - | Os07g0108300 [Oryza sativa | SALT00000006905_Cluster19137-004 |
| Cluster19150 | - | - | up | - | up | - | Os05g0447700 [Oryza sativa | SALT00000080187_Cluster19150-006 |
| Cluster19161 | - | down | down | - | - | - | adhesion of calyx edges ACE | SALT00000067992_Cluster19161-003 |
| Cluster19162 | - | - | up | - | up | up | probable acyl-activating enzy | SALT00000069673_Cluster19162-010 |
| Cluster19164 | - | - | down | - | down | - | transposon Mutator sub-class | SALT00000007358_Cluster19164-003 |
| Cluster19165 | - | - | up | - | up | up | lipase-like PAD4 [Setaria itali | SALT00000019151_Cluster19165-002 |
| Cluster19187 | - | - | up | - | up | up | regulatory NPR1 [Oryza sativ | SALT00000058144_Cluster19187-007 |
| Cluster19190 | - | - | up | - | up | up | sugar transporter [Oryza sativ | SALT00000007078_Cluster19190-002 |
| Cluster1920 | - | up | up | - | - | - | Os01g0625900 [Oryza sativa | SALT00000047726_Cluster1920-001 |
| Cluster19202 | - | - | up | - | - | - | cytochrome b561 [Zea mays] | SALT00000010199_Cluster19202-002 |
| Cluster19209 | - | down | down | - | down | down | [Oryza sativa Japonica Group] | SALT00000056042_Cluster19209-001 |
| Cluster19210 | - | - | up | - | - | up | polyubiquitin [Brachypodium | SALT00000056342_Cluster19210-003 |
| Cluster19224 | - | - | up | - | - | up | receptor kinase 1-like [Oryza | SALT00000039227_Cluster19224-001 |
| Cluster1925 | - | - | - | - | down | - | Os03g0152800 [Oryza sativa | SALT00000074543_Cluster1925-010 |
| Cluster1928 | - | - | down | - | - | - | transposon unclassified | SALT00000091086_Cluster1928-004 |
| Cluster19286 | - | - | down | - | down | down | methylenetetrahydrofolate red | SALT00000061774_Cluster19286-001 |
| Cluster1930 | - | down | down | - | - | - | respiratory burst oxidase hom | SALT00000068716_Cluster1930-008 |
| Cluster19304 | - | - | up | - | - | - | TOM1 2 [Setaria italica] | SALT00000041256_Cluster19304-011 |
| Cluster19309 | - | up | up | up | up | up | dihydrolipoyllysine-residue ac | SALT00000012470_Cluster19309-005 |
| Cluster1931 | - | - | down | - | - | - | subtilisin-like protease | SALT00000062737_Cluster1931-005 |
| Cluster19314 | - | - | up | - | up | up | [Hordeum vulgare vulgare] | SALT00000057537_Cluster19314-002 |
| Cluster19364 | - | - | - | - | up | - | ubiquilin-1 [Zea mays] | SALT00000036564_Cluster19364-005 |
| Cluster19382 | - | down | down | down | down | - | g14A [Zea mays] | SALT00000007841_Cluster19382-001 |
| Cluster1939 | - | - | - | - | down | - | histidinol chloroplastic isofor | SALT00000018544_Cluster1939-005 |
| Cluster19392 | - | - | up | - | - | - | scarecrow 4 [Zea mays] | SALT00000062473_Cluster19392-005 |
| Cluster19393 | - | - | up | - | up | up | uncharacterized protein LOC1 | SALT00000002021_Cluster19393-001 |
| Cluster19394 | - | - | down | - | - | - | PREDICTED: uncharacterize | SALT00000011514_Cluster19394-002 |
| Cluster19408 | - | - | down | - | down | down | Nucleolar expressed [Oryza sa | SALT00000052184_Cluster19408-008 |
| Cluster19440 | - | - | up | - | up | up | AT-hook motif nuclear-localiz | SALT00000019972_Cluster19440-013 |
| Cluster19441 | down | down | down | - | down | down | esterase [Zea mays] | SALT00000042370_Cluster19441-007 |
| Cluster19450 | - | - | - | - | - | up | Os11g0107400 [Oryza sativa | SALT00000000860_Cluster19450-001 |
| Cluster19451 | - | - | up | - | up | up | allene oxide synthase chloropl | SALT00000002374_Cluster19451-001 |
| Cluster19459 | - | up | up | up | up | - | mitochondrial aldehyde dehyd | SALT00000046696_Cluster19459-009 |
| Cluster19463 | - | - | up | - | up | up | xyloglucan fucosyltransferase | SALT00000011950_Cluster19463-005 |
| Cluster19465 | - | - | down | - | - | down | indeterminate-domain chlorop | SALT00000016370_Cluster19465-001 |
| Cluster19483 | - | - | up | - | - | - | glycerol-3-phosphate acyltran | SALT00000029072_Cluster19483-003 |
| Cluster19494 | - | - | down | - | - | - | Os08g0439600 [Oryza sativa | SALT00000009043_Cluster19494-001 |

| | | | | | | | |
|---|---|---|---|---|---|---|---|
| Cluster1950 | - | - | - | - | down | - | potassium transporter | SALT00000092823_Cluster1950-001 |
| Cluster1952 | - | - | down | - | down | down | probable methionine--tRNA li | SALT00000080228_Cluster1952-009 |
| Cluster19521 | - | - | up | - | up | up | ethylene-responsive transcript | SALT00000012510_Cluster19521-001 |
| Cluster19522 | - | - | up | - | up | up | HGA4 [Zea mays] | SALT00000014711_Cluster19522-004 |
| Cluster19536 | - | - | up | - | up | up | aspartic ase 1 | SALT00000037830_Cluster19536-001 |
| Cluster19553 | - | - | up | - | up | - | anthranilate N-hydroxycinnan | SALT00000038043_Cluster19553-003 |
| Cluster19562 | - | - | down | - | down | - | ABC transporter G family me | SALT00000036374_Cluster19562-004 |
| Cluster19564 | - | - | - | - | - | up | Serine carboxypeptidase III pı | SALT00000040997_Cluster19564-004 |
| Cluster19578 | - | - | - | - | - | up | [Hordeum vulgare vulgare] | SALT00000051269_Cluster19578-002 |
| Cluster1958 | - | up | up | - | up | up | lichenase-2 precursor [Zea ma | SALT00000000145_Cluster1958-001 |
| Cluster19584 | - | - | down | - | - | - | Na(+) H(+) antiporter isoform | SALT00000009057_Cluster19584-003 |
| Cluster19588 | - | - | down | - | down | - | hydroxymuconic semialdehyd | SALT00000051752_Cluster19588-005 |
| Cluster19611 | - | - | - | - | up | - | zinc finger MAGPIE-like [Ela | SALT00000006787_Cluster19611-001 |
| Cluster19613 | - | - | up | - | - | - | cysteine ase 1 [Oryza sativa Ja | SALT00000017474_Cluster19613-003 |
| Cluster19615 | up | up | up | - | up | up | NAC transcription factor [Trit | SALT00000038775_Cluster19615-005 |
| Cluster19620 | - | - | down | - | down | down | Os03g0140100 [Oryza sativa | SALT00000058705_Cluster19620-006 |
| Cluster19624 | - | down | - | - | - | up | Os02g0200300 [Oryza sativa | SALT00000028050_Cluster19624-004 |
| Cluster19631 | - | - | down | - | down | - | [Hordeum vulgare vulgare] | SALT00000003671_Cluster19631-002 |
| Cluster19661 | - | - | up | - | up | - | CBL-interacting kinase family | SALT00000059984_Cluster19661-010 |
| Cluster19665 | - | - | - | - | - | down | hydroxyproline-rich glyco -lik | SALT00000037735_Cluster19665-003 |
| Cluster19669 | - | - | - | - | - | down | ---NA--- | SALT00000051875_Cluster19669-001 |
| Cluster19679 | - | - | down | - | down | - | T-complex 1 subunit theta | SALT00000068124_Cluster19679-013 |
| Cluster19682 | - | - | down | - | down | - | galactoside 2-alpha-L-fucosyl | SALT00000056162_Cluster19682-002 |
| Cluster19690 | - | - | down | - | - | - | signal recognition particle 54 | SALT00000018167_Cluster19690-001 |
| Cluster19700 | - | - | - | - | - | up | dicarboxylate transporter chlo | SALT00000002648_Cluster19700-001 |
| Cluster19701 | - | - | up | - | up | up | probable receptor kinase At3g | SALT00000010965_Cluster19701-002 |
| Cluster19738 | - | - | up | - | up | up | avr9 Cf-9 rapidly elicited 137 | SALT00000010100_Cluster19738-001 |
| Cluster19757 | - | - | down | - | down | down | Os03g0796600 [Oryza sativa | SALT00000038898_Cluster19757-005 |
| Cluster19758 | - | - | down | - | - | - | zinc finger CCCH domain-coı | SALT00000045481_Cluster19758-002 |
| Cluster19760 | - | - | up | - | up | - | galactoside 2-alpha-L-fucosyl | SALT00000046871_Cluster19760-002 |
| Cluster19775 | - | - | down | - | down | - | TPA: laccase family [Zea may | SALT00000050370_Cluster19775-002 |
| Cluster19783 | - | - | up | - | - | - | trafficking particle complex sı | SALT00000038970_Cluster19783-001 |
| Cluster19784 | - | - | up | - | - | - | heat shock factor HSF30 [Zea | SALT00000039610_Cluster19784-007 |
| Cluster19791 | - | - | - | - | - | up | Sugar carrier C | SALT00000046966_Cluster19791-009 |
| Cluster1980 | - | - | up | - | up | up | MRS2-10 [Zea mays] | SALT00000087203_Cluster1980-002 |
| Cluster19801 | - | - | - | - | down | - | ---NA--- | SALT00000006165_Cluster19801-001 |
| Cluster19806 | - | - | - | - | down | - | nucellin-like aspartic protease | SALT00000039148_Cluster19806-003 |
| Cluster19812 | - | - | up | - | - | - | vesicle-associated 1-2-like [Se | SALT00000061109_Cluster19812-001 |
| Cluster19817 | - | - | - | - | up | - | Os09g0525600 [Oryza sativa | SALT00000012938_Cluster19817-005 |
| Cluster19823 | - | - | up | - | up | up | F-box At5g67140 isoform X1 | SALT00000057028_Cluster19823-001 |
| Cluster19827 | - | - | up | - | up | up | interferon-related [Oryza sativ | SALT00000005902_Cluster19827-002 |
| Cluster19829 | - | up | up | - | up | up | beta-amylase [Zea mays] | SALT00000012432_Cluster19829-002 |
| Cluster19871 | - | - | - | - | - | up | Os01g0118400 [Oryza sativa . | SALT00000035348_Cluster19871-007 |
| Cluster19885 | - | - | up | - | up | up | CBL-interacting kinase 10 | SALT00000051345_Cluster19885-004 |
| Cluster19890 | - | - | - | - | up | up | myristoyl-acyl carrier thioeste | SALT00000026810_Cluster19890-008 |
| Cluster19893 | - | up | up | - | up | up | ---NA--- | SALT00000050513_Cluster19893-003 |
| Cluster1990 | - | - | - | - | up | - | phenylalanine ammonia-lyase | SALT00000084873_Cluster1990-113 |
| Cluster19900 | up | up | up | - | up | up | copper transporter [Triticum a | SALT00000002268_Cluster19900-002 |
| Cluster19908 | - | - | down | - | down | - | pectate lyase precursor | SALT00000040037_Cluster19908-005 |
| Cluster19909 | - | - | up | - | up | up | mitochondrial 2-oxoglutarate | SALT00000044565_Cluster19909-001 |
| Cluster19917 | - | - | down | - | - | - | subtilisin-like protease [Brach | SALT00000012507_Cluster19917-001 |
| Cluster1992 | - | - | - | - | down | - | calmodulin-interacting 111 isc | SALT00000092918_Cluster1992-001 |
| Cluster19920 | - | - | - | - | down | - | heparan-alpha-glucosaminide | SALT00000044159_Cluster19920-005 |
| Cluster1993 | - | - | down | - | down | - | TPA: C3HC zinc finger-like fa | SALT00000074861_Cluster1993-005 |
| Cluster19937 | up | up | - | - | down | - | Os07g0108300 [Oryza sativa | SALT00000059116_Cluster19937-001 |
| Cluster19945 | - | - | up | - | up | up | pnFL-2 [Zea mays] | SALT00000062429_Cluster19945-001 |
| Cluster19946 | - | - | up | - | up | - | Na+ dependent neutral amino | SALT00000004574_Cluster19946-001 |
| Cluster19958 | - | - | up | - | up | up | alginate regulatory [Zea mays | SALT00000050653_Cluster19958-003 |

| | | | | | | | | |
|---|---|---|---|---|---|---|---|---|
| Cluster19959 | - | - | up | - | up | - | 6-phosphogluconate decarbox | SALT00000052254_Cluster19959-010 |
| Cluster19968 | - | - | - | - | down | - | external alternative NAD(P)H | SALT00000064166_Cluster19968-007 |
| Cluster1997 | - | - | - | - | - | up | plastid (p)ppGpp synthase [O | SALT00000071498_Cluster1997-002 |
| Cluster1998 | - | - | up | - | up | up | heat shock 1 [Zea mays] | SALT00000072738_Cluster1998-042 |
| Cluster19991 | - | - | up | - | up | - | CBS domain-containing CBS | SALT00000006344_Cluster19991-001 |
| Cluster19998 | - | - | - | - | down | - | Os01g0285300 [Oryza sativa | SALT00000048715_Cluster19998-010 |
| Cluster20005 | - | - | up | - | up | up | probable trehalase | SALT00000015624_Cluster20005-001 |
| Cluster2001 | - | - | down | - | down | down | probable GTP-binding chloro | SALT00000067417_Cluster2001-005 |
| Cluster20010 | - | - | up | - | - | - | BTB POZ domain-containing | SALT00000048091_Cluster20010-002 |
| Cluster20025 | - | - | up | - | - | - | Os02g0542400 [Oryza sativa | SALT00000049738_Cluster20025-009 |
| Cluster20028 | up | - | up | - | up | up | nucleotide pyrophosphatase p | SALT00000000718_Cluster20028-001 |
| Cluster20029 | - | - | down | - | - | - | Os06g0476200 [Oryza sativa | SALT00000006848_Cluster20029-001 |
| Cluster20030 | - | up | - | - | - | - | ubiquitin carboxyl-terminal h | SALT00000012456_Cluster20030-004 |
| Cluster20033 | - | - | up | - | up | up | transcription factor bHLH130 | SALT00000042888_Cluster20033-005 |
| Cluster20035 | - | - | up | - | up | up | hydroxycinnamoyl transferase | SALT00000046982_Cluster20035-005 |
| Cluster20047 | - | - | down | - | down | down | transcription factor PIF5 isofo | SALT00000045499_Cluster20047-002 |
| Cluster20060 | - | - | down | - | - | - | Os03g0412800 [Oryza sativa | SALT00000006137_Cluster20060-001 |
| Cluster20061 | - | - | up | - | - | - | NRT1 PTR FAMILY -like [Se | SALT00000008449_Cluster20061-002 |
| Cluster20072 | - | - | up | - | up | up | polyol transporter 5-like | SALT00000056831_Cluster20072-002 |
| Cluster20075 | - | up | - | - | - | - | white cap1 [Zea mays] | SALT00000023777_Cluster20075-013 |
| Cluster20082 | up | up | up | - | up | up | beta-amylase [Zea mays] | SALT00000051159_Cluster20082-001 |
| Cluster20087 | - | - | down | - | - | - | [Hordeum vulgare vulgare] | SALT00000061096_Cluster20087-006 |
| Cluster20105 | - | - | - | - | - | down | SMR domain-containing At5g | SALT00000015005_Cluster20105-001 |
| Cluster20114 | - | down | down | down | down | down | probable cinnamyl alcohol de | SALT00000044568_Cluster20114-004 |
| Cluster20120 | - | - | - | - | down | - | laccase-11 [Setaria italica] | SALT00000004520_Cluster20120-001 |
| Cluster20134 | - | - | down | - | - | - | Nucleolar GTP-binding expre | SALT00000019339_Cluster20134-004 |
| Cluster20149 | - | - | up | - | - | - | [Hordeum vulgare vulgare] | SALT00000010271_Cluster20149-002 |
| Cluster20158 | - | - | - | - | down | - | Os08g0282500 [Oryza sativa | SALT00000012915_Cluster20158-001 |
| Cluster20162 | - | - | - | down | down | - | nuclear transcription factor Y | SALT00000042068_Cluster20162-001 |
| Cluster20170 | - | - | up | - | up | up | acetylglucosaminyltransferase | SALT00000006335_Cluster20170-002 |
| Cluster20174 | up | up | up | - | - | - | cation calcium exchanger 1-lil | SALT00000041352_Cluster20174-002 |
| Cluster20180 | - | - | up | - | up | up | carbonic anhydrase isoform X | SALT00000007766_Cluster20180-001 |
| Cluster20184 | - | - | up | - | - | - | Os01g0826400 [Oryza sativa | SALT00000004455_Cluster20184-001 |
| Cluster20190 | - | - | down | - | down | down | TPA: TCP-1 cpn60 chaperoni | SALT00000054280_Cluster20190-010 |
| Cluster20199 | - | - | - | - | up | - | Monodehydroascorbate reduc | SALT00000037693_Cluster20199-003 |
| Cluster2020 | - | - | up | - | up | up | Os02g0777100 [Oryza sativa | SALT00000087985_Cluster2020-017 |
| Cluster20218 | - | - | down | - | - | - | H ACA ribonucleo complex s | SALT00000055542_Cluster20218-005 |
| Cluster20222 | - | - | up | - | up | - | Os01g0169900 [Oryza sativa | SALT00000012797_Cluster20222-001 |
| Cluster20234 | - | - | - | - | - | down | serine carboxypeptidase-like 1 | SALT00000003420_Cluster20234-002 |
| Cluster20236 | - | - | - | - | down | - | pentatricopeptide repeat-conta | SALT00000017226_Cluster20236-001 |
| Cluster20239 | - | up | - | - | - | down | fructose-bisphosphate aldolas | SALT00000046941_Cluster20239-001 |
| Cluster20256 | - | - | up | - | up | up | UDP-glucose-4-epimerase [Z | SALT00000061224_Cluster20256-002 |
| Cluster20266 | - | - | up | - | up | up | transcription factor qSH-1 [O | SALT00000010076_Cluster20266-001 |
| Cluster20269 | - | up | - | - | - | - | tuber-specific and sucrose-res | SALT00000041800_Cluster20269-001 |
| Cluster20270 | - | - | up | - | up | up | Os01g0793500 [Oryza sativa | SALT00000042288_Cluster20270-001 |
| Cluster20278 | - | - | down | - | - | - | RNA-binding [Zea mays] | SALT00000002895_Cluster20278-001 |
| Cluster20289 | - | - | up | - | - | up | Os12g0231000 [Oryza sativa | SALT00000061846_Cluster20289-004 |
| Cluster20294 | - | - | down | - | down | - | 60S ribosomal L4-1-like [Seta | SALT00000056722_Cluster20294-021 |
| Cluster20300 | - | - | down | - | - | - | cytosolic 5-nucleotidase III [Z | SALT00000009199_Cluster20300-001 |
| Cluster20313 | - | - | up | - | - | - | uncharacterized LOC1002793 | SALT00000006431_Cluster20313-002 |
| Cluster20324 | up | up | up | - | - | - | ABA responsive element bind | SALT00000058554_Cluster20324-002 |
| Cluster20325 | - | down | down | - | down | down | Calreticulin precursor [Zea m | SALT00000060316_Cluster20325-026 |
| Cluster2033 | - | - | - | - | down | down | Os03g0227000 [Oryza sativa | SALT00000074391_Cluster2033-010 |
| Cluster20340 | - | - | up | - | up | up | avr9 Cf-9 rapidly elicited 137 | SALT00000011949_Cluster20340-002 |
| Cluster20342 | - | down | - | - | - | - | PS60 [Oryza sativa Japonica ( | SALT00000014190_Cluster20342-001 |
| Cluster20345 | - | - | - | - | down | - | calcium-dependent kinase 13- | SALT00000035804_Cluster20345-001 |
| Cluster20348 | - | - | up | - | - | - | neurogenic locus notch precur | SALT00000044893_Cluster20348-001 |
| Cluster20350 | - | - | - | - | down | down | Transcription factor IIIA | SALT00000062812_Cluster20350-005 |

| | | | | | | | | |
|---|---|---|---|---|---|---|---|---|
| Cluster20352 | - | - | up | - | - | - | Os04g0453400 [Oryza sativa | SALT00000076977_Cluster20352-003 |
| Cluster20354 | - | - | up | - | up | - | ethylene response element bin | SALT00000004817_Cluster20354-003 |
| Cluster20361 | - | - | up | - | up | up | endo-1,4-beta-xylanase A-like | SALT00000060500_Cluster20361-002 |
| Cluster20363 | - | - | up | - | - | - | serine arginine repetitive matr | SALT00000001694_Cluster20363-001 |
| Cluster20368 | - | - | up | - | up | up | branched-chain-amino-acid ar | SALT00000043199_Cluster20368-001 |
| Cluster20373 | - | - | up | - | - | - | heparan-alpha-glucosaminide | SALT00000047586_Cluster20373-005 |
| Cluster20381 | - | - | down | - | down | - | AER [Zea mays] | SALT00000007271_Cluster20381-001 |
| Cluster20385 | - | - | up | - | up | up | probable inositol oxygenase [S | SALT00000038364_Cluster20385-001 |
| Cluster20391 | - | - | up | - | up | - | phosphoinositide phospholipa | SALT00000014765_Cluster20391-002 |
| Cluster20393 | - | - | down | - | - | - | nucleolin-like isoform X1 | SALT00000035679_Cluster20393-002 |
| Cluster20406 | - | - | down | - | down | - | Os05g0449900 [Oryza sativa | SALT00000005164_Cluster20406-001 |
| Cluster2042 | - | - | down | - | down | - | Os08g0407200 [Oryza sativa | SALT00000062934_Cluster2042-015 |
| Cluster20422 | - | - | - | - | - | up | Os01g0570800 [Oryza sativa | SALT00000070351_Cluster20422-007 |
| Cluster20438 | - | - | up | - | up | - | trichome birefringence-like 28 | SALT00000012857_Cluster20438-008 |
| Cluster20448 | - | - | down | - | - | - | T-complex 1 subunit delta [Or | SALT00000093783_Cluster20448-010 |
| Cluster20450 | - | - | up | - | up | up | wall-associated receptor kinas | SALT00000005276_Cluster20450-001 |
| Cluster20464 | - | - | up | - | up | up | hexokinase [Zea mays] | SALT00000000164_Cluster20464-001 |
| Cluster20465 | - | - | up | - | up | up | DETOXIFICATION 27-like [ | SALT00000003645_Cluster20465-002 |
| Cluster20469 | - | - | down | - | - | - | peroxidase 18-like | SALT00000015414_Cluster20469-001 |
| Cluster20484 | - | - | down | - | - | - | prolyl-tRNA synthetase | SALT00000011819_Cluster20484-006 |
| Cluster20485 | - | - | up | - | up | up | starmaker-like [Brachypodium | SALT00000012816_Cluster20485-007 |
| Cluster20499 | up | - | up | - | - | up | cation proton exchanger 1a [C | SALT00000004668_Cluster20499-001 |
| Cluster2052 | - | - | down | - | down | down | probable methionine--tRNA li | SALT00000069621_Cluster2052-001 |
| Cluster20526 | - | - | - | - | down | - | mTERF family [Zea mays] | SALT00000048515_Cluster20526-006 |
| Cluster20538 | - | - | up | - | up | - | shikimate O-hydroxycinnamo | SALT00000050541_Cluster20538-012 |
| Cluster2054 | - | - | down | - | - | - | partial [Oryza sativa Japonica | SALT00000087398_Cluster2054-001 |
| Cluster20543 | - | - | - | - | down | - | cytochrome P450 CYP78A53 | SALT00000002981_Cluster20543-001 |
| Cluster20552 | - | - | down | - | - | - | translation initiation factor IF- | SALT00000057320_Cluster20552-001 |
| Cluster20558 | - | - | - | - | - | up | [Hordeum vulgare vulgare] | SALT00000041089_Cluster20558-002 |
| Cluster20565 | - | - | up | - | up | up | Os01g0833000 [Oryza sativa | SALT00000065085_Cluster20565-002 |
| Cluster20567 | - | up | up | - | - | - | Os03g0122300 [Oryza sativa | SALT00000006607_Cluster20567-001 |
| Cluster20571 | - | down | down | - | - | - | Os03g0703100 [Oryza sativa | SALT00000040402_Cluster20571-010 |
| Cluster20584 | - | - | down | - | - | - | thylakoid lumenal kDa chloro | SALT00000003094_Cluster20584-001 |
| Cluster20592 | - | - | up | - | up | up | [Hordeum vulgare vulgare] | SALT00000041967_Cluster20592-002 |
| Cluster20613 | - | - | down | - | - | - | bifunctional aspartokinase hor | SALT00000058494_Cluster20613-006 |
| Cluster20615 | - | - | down | - | down | down | probable anion transporter chl | SALT00000008138_Cluster20615-001 |
| Cluster20622 | - | - | up | - | up | up | probable calcium-binding CM | SALT00000054466_Cluster20622-001 |
| Cluster20634 | - | - | up | - | up | up | Os06g0693700 [Oryza sativa | SALT00000019701_Cluster20634-005 |
| Cluster20651 | - | - | up | - | up | up | DUF1421 domain family [Zea | SALT00000002077_Cluster20651-001 |
| Cluster20669 | - | - | up | - | - | - | endo-1,4-beta-xylanase A-like | SALT00000002729_Cluster20669-001 |
| Cluster2067 | - | - | down | - | down | - | AF466646_7 poly | SALT00000089714_Cluster2067-001 |
| Cluster20670 | - | - | up | - | up | up | MATE efflux family [Zea may | SALT00000004003_Cluster20670-001 |
| Cluster20690 | - | - | up | - | up | up | mpv17 PMP22 family [Zea m | SALT00000044573_Cluster20690-004 |
| Cluster20698 | up | - | - | - | down | - | ras-related Rab11A [Zea mays | SALT00000007734_Cluster20698-001 |
| Cluster20712 | - | - | up | - | up | up | [Hordeum vulgare vulgare] | SALT00000007393_Cluster20712-001 |
| Cluster20733 | - | - | up | - | up | up | CAPIP1 [Zea mays] | SALT00000041219_Cluster20733-001 |
| Cluster20734 | - | - | up | - | up | up | acyl-coenzyme A oxidase pero | SALT00000041363_Cluster20734-006 |
| Cluster20737 | - | - | - | - | down | down | shikimate kinase chloroplastic | SALT00000048298_Cluster20737-002 |
| Cluster20739 | - | - | down | - | - | - | ---NA--- | SALT00000052464_Cluster20739-001 |
| Cluster20765 | - | - | down | down | down | - | fructose-bisphosphate aldolase | SALT00000046329_Cluster20765-001 |
| Cluster20772 | down | down | down | - | - | - | legumain-like protease | SALT00000004254_Cluster20772-002 |
| Cluster20774 | - | - | up | - | up | up | WD-repeat domain phosphoin | SALT00000007231_Cluster20774-002 |
| Cluster20800 | - | - | down | - | - | - | [Hordeum vulgare vulgare] | SALT00000004986_Cluster20800-001 |
| Cluster2081 | - | - | up | - | up | up | Os04g0531500 [Oryza sativa | SALT00000066930_Cluster2081-005 |
| Cluster20814 | - | - | - | - | down | - | Os05g0161200 [Oryza sativa | SALT00000014256_Cluster20814-001 |
| Cluster2083 | - | - | down | - | - | - | rp3-like disease resistance | SALT00000085688_Cluster2083-002 |
| Cluster20830 | - | - | up | - | - | - | LOL3 [Oryza brachyantha] | SALT00000049551_Cluster20830-001 |
| Cluster20836 | - | - | - | - | up | - | Os01g0134700 [Oryza sativa | SALT00000003932_Cluster20836-002 |

| | | | | | | | | |
|---|---|---|---|---|---|---|---|---|
| Cluster20841 | - | - | - | - | up | - | nucellin-like aspartic protease | SALT00000048042_Cluster20841-005 |
| Cluster20848 | - | - | up | - | up | up | amino acid permease [Zea ma | SALT00000005815_Cluster20848-001 |
| Cluster20851 | - | - | up | - | up | up | arm repeat [Oryza sativa Japo | SALT00000055528_Cluster20851-001 |
| Cluster20878 | - | - | - | - | down | - | glycosyltransferase 5 [Zea ma | SALT00000000649_Cluster20878-001 |
| Cluster20883 | - | - | - | - | - | down | cytochrome P450 superfamily | SALT00000011473_Cluster20883-007 |
| Cluster20887 | - | - | up | - | - | - | Serine carboxypeptidase III pı | SALT00000046075_Cluster20887-005 |
| Cluster20890 | up | up | up | - | up | up | Os09g0484900 [Oryza sativa | SALT00000054411_Cluster20890-006 |
| Cluster20900 | - | - | - | - | up | up | electron transfer flavo -ubiqui | SALT00000011999_Cluster20900-001 |
| Cluster20921 | - | - | down | - | down | down | [Hordeum vulgare vulgare] | SALT00000050595_Cluster20921-002 |
| Cluster20927 | - | - | down | - | - | - | Os06g0308000 [Oryza sativa | SALT00000060831_Cluster20927-005 |
| Cluster20936 | - | - | down | - | - | - | PSAG [Oryza sativa Japonica | SALT00000017193_Cluster20936-003 |
| Cluster2094 | - | - | down | - | down | - | PREDICTED: uncharacterize | SALT00000093232_Cluster2094-004 |
| Cluster20946 | - | - | down | - | - | - | Os03g0352300 [Oryza sativa | SALT00000002071_Cluster20946-001 |
| Cluster20949 | up | up | - | - | - | down | ---NA--- | SALT00000010586_Cluster20949-001 |
| Cluster20951 | - | - | up | - | up | up | CBL-interacting kinase 31 | SALT00000013917_Cluster20951-003 |
| Cluster20966 | - | - | down | - | - | - | double-stranded RNA-binding | SALT00000017155_Cluster20966-001 |
| Cluster20973 | - | - | down | - | down | - | chlorophyll a-b binding chlor | SALT00000054919_Cluster20973-003 |
| Cluster2098 | up | up | up | - | up | up | copper transporter [Triticum a | SALT00000028033_Cluster2098-004 |
| Cluster20998 | down | down | down | down | down | - | uncharacterized LOC1008568 | SALT00000052637_Cluster20998-004 |
| Cluster20999 | - | - | down | - | down | down | Os03g0705800 [Oryza sativa | SALT00000052974_Cluster20999-005 |
| Cluster21001 | - | - | - | up | - | - | TPA: legumain-like protease I | SALT00000060028_Cluster21001-003 |
| Cluster21004 | - | - | up | - | - | - | Aspartic ase | SALT00000055417_Cluster21004-018 |
| Cluster2101 | - | - | up | - | up | up | sucrose synthase 1 [Zea mays | SALT00000067944_Cluster2101-038 |
| Cluster21028 | - | - | down | - | - | - | PREDICTED: uncharacterize | SALT00000067785_Cluster21028-001 |
| Cluster21038 | - | - | up | - | up | - | Os03g0729000 [Oryza sativa | SALT00000051096_Cluster21038-002 |
| Cluster21040 | - | - | up | - | up | up | Os02g0717600 [Oryza sativa | SALT00000055541_Cluster21040-005 |
| Cluster21046 | - | - | up | - | up | - | reticuline oxidase-like | SALT00000010763_Cluster21046-003 |
| Cluster21048 | - | - | up | - | up | - | phosphatase 2C | SALT00000017167_Cluster21048-001 |
| Cluster21058 | - | - | down | - | - | - | T-complex 1 subunit zeta [Zea | SALT00000014279_Cluster21058-002 |
| Cluster2106 | - | - | down | - | - | - | Os09g0487200 [Oryza sativa | SALT00000075821_Cluster2106-006 |
| Cluster21067 | - | - | up | - | up | up | glycerol-3-phosphate transpor | SALT00000000651_Cluster21067-001 |
| Cluster21071 | - | - | down | - | - | - | glucan endo-1,3-beta-glucosic | SALT00000006143_Cluster21071-001 |
| Cluster21082 | - | - | down | - | down | - | partial [Oryza sativa Japonica | SALT00000041727_Cluster21082-008 |
| Cluster2111 | - | - | up | - | up | up | phospholipase D delta | SALT00000069974_Cluster2111-025 |
| Cluster21115 | - | - | down | - | - | - | phenylalanyl-tRNA synthetas | SALT00000019083_Cluster21115-005 |
| Cluster21118 | - | down | down | - | - | - | cytochrome P450 [Oryza sativ | SALT00000039235_Cluster21118-002 |
| Cluster2112 | - | - | - | - | up | up | [Oryza sativa Japonica Group | SALT00000063921_Cluster2112-023 |
| Cluster21121 | - | - | down | - | down | down | COBRA 5 | SALT00000051108_Cluster21121-007 |
| Cluster2113 | - | - | down | - | - | - | probable LRR receptor-like se | SALT00000066990_Cluster2113-005 |
| Cluster21139 | - | up | up | - | up | - | aldehyde dehydrogenase MIS | SALT00000000758_Cluster21139-002 |
| Cluster21151 | - | - | up | - | - | up | xylosyltransferase 1 [Zea may | SALT00000061002_Cluster21151-004 |
| Cluster21153 | - | - | up | - | up | - | phosphatase 2C isoform gamr | SALT00000008530_Cluster21153-002 |
| Cluster21154 | - | - | up | - | up | - | NRT1 PTR FAMILY -like | SALT00000014875_Cluster21154-001 |
| Cluster21160 | - | - | up | - | up | up | kinase domain containing [Or | SALT00000050663_Cluster21160-002 |
| Cluster21161 | - | - | down | - | - | - | RNA polymerase sigma factor | SALT00000016107_Cluster21161-005 |
| Cluster21190 | - | - | up | - | up | up | NAC domain transcription fac | SALT00000044900_Cluster21190-004 |
| Cluster21206 | - | - | down | - | - | - | probable carboxylesterase 15 | SALT00000052596_Cluster21206-004 |
| Cluster21211 | - | - | - | down | down | down | ---NA--- | SALT00000051949_Cluster21211-001 |
| Cluster21220 | - | - | up | - | up | - | 3 (2 ),5 -bisphosphate nucleot | SALT00000009288_Cluster21220-002 |
| Cluster21222 | - | - | down | - | down | - | CHLOROPLAST IMPORT A | SALT00000061565_Cluster21222-004 |
| Cluster21236 | - | - | up | - | - | - | dnaJ chaperone family | SALT00000006588_Cluster21236-003 |
| Cluster21251 | - | - | down | - | down | - | Os04g0473400 [Oryza sativa | SALT00000054841_Cluster21251-001 |
| Cluster21254 | - | - | up | - | up | up | glycosyltransferase [Sorghum | SALT00000059750_Cluster21254-004 |
| Cluster21258 | - | - | - | - | up | - | Os08g0359300 [Oryza sativa | SALT00000008628_Cluster21258-001 |
| Cluster21263 | - | up | up | - | up | - | Os05g0540900 [Oryza sativa | SALT00000023008_Cluster21263-003 |
| Cluster21272 | - | - | up | - | - | - | ACT domain-containing ACR | SALT00000051912_Cluster21272-002 |
| Cluster21285 | - | - | down | - | down | - | signal recognition particle 54 | SALT00000049377_Cluster21285-005 |
| Cluster21296 | - | - | - | - | up | - | hemolysin [Zea mays] | SALT00000045232_Cluster21296-005 |

| | | | | | | | | |
|---|---|---|---|---|---|---|---|---|
| Cluster21312 | - | - | up | - | up | - | Os04g0430200 [Oryza sativa | SALT00000048342_Cluster21312-003 |
| Cluster21321 | - | - | up | up | up | - | [Hordeum vulgare vulgare] | SALT00000044602_Cluster21321-004 |
| Cluster21343 | - | down | down | - | down | - | NADH dehydrogenase subuni | SALT00000000391_Cluster21343-001 |
| Cluster21366 | - | up | up | - | up | up | Os06g0635200 [Oryza sativa | SALT00000015923_Cluster21366-003 |
| Cluster2137 | - | down | down | - | down | down | probable LRR receptor-like se | SALT00000089477_Cluster2137-010 |
| Cluster21382 | - | - | up | - | - | - | Os04g0287400 [Oryza sativa | SALT00000055149_Cluster21382-001 |
| Cluster21389 | - | - | up | - | - | - | shikimate kinase chloroplastic | SALT00000038874_Cluster21389-001 |
| Cluster21397 | - | - | up | - | up | - | folate-biopterin transporter ch | SALT00000065758_Cluster21397-003 |
| Cluster21404 | - | up | up | - | - | - | auxin-induced [Oryza sativa J | SALT00000045555_Cluster21404-008 |
| Cluster21423 | down | down | down | - | - | - | cyp450 [Hordeum vulgare] | SALT00000059590_Cluster21423-003 |
| Cluster21425 | up | - | up | - | up | up | WRKY53 - superfamily of TF | SALT00000041465_Cluster21425-003 |
| Cluster21444 | - | - | down | - | - | - | E3 ubiquitin- ligase XB3-like | SALT00000054477_Cluster21444-001 |
| Cluster21447 | - | - | down | - | - | - | TPA: fumarylacetoacetate hyd | SALT00000068553_Cluster21447-001 |
| Cluster2145 | - | - | up | - | up | - | Cell division cycle expressed | SALT00000069325_Cluster2145-029 |
| Cluster21462 | - | - | - | - | down | - | TPA: sigma factor SIG2B [Ze | SALT00000005527_Cluster21462-001 |
| Cluster21464 | - | - | up | - | up | up | [Hordeum vulgare vulgare] | SALT00000019217_Cluster21464-002 |
| Cluster2148 | - | - | - | - | - | up | OBERON 4-like | SALT00000090917_Cluster2148-002 |
| Cluster21500 | - | - | down | - | - | - | E3 ubiquitin- ligase RGLG2-l | SALT00000000788_Cluster21500-002 |
| Cluster21502 | - | - | down | - | - | - | cysteine--tRNA chloroplastic | SALT00000007366_Cluster21502-003 |
| Cluster21504 | - | - | down | - | - | - | Os01g0923900 [Oryza sativa | SALT00000013724_Cluster21504-006 |
| Cluster21512 | - | - | down | - | down | down | UDP-glycosyltransferase 91C | SALT00000053005_Cluster21512-011 |
| Cluster21522 | - | - | down | - | - | - | transcription factor UNE12-lil | SALT00000036616_Cluster21522-001 |
| Cluster21533 | - | - | - | - | - | down | 60S ribosome subunit biogene | SALT00000064172_Cluster21533-001 |
| Cluster21535 | - | - | up | - | up | up | probable mitochondrial chape | SALT00000000431_Cluster21535-001 |
| Cluster21537 | up | - | - | - | down | - | UDP-glycosyltransferase 86A | SALT00000008076_Cluster21537-002 |
| Cluster21538 | - | - | up | - | up | - | ---NA--- | SALT00000008741_Cluster21538-001 |
| Cluster21543 | - | - | - | - | down | - | threonine synthase [Zea mays | SALT00000036089_Cluster21543-005 |
| Cluster21546 | - | - | down | - | down | down | Os08g0481400 [Oryza sativa | SALT00000039174_Cluster21546-001 |
| Cluster21547 | - | - | up | - | up | up | kinase superfamily [Zea mays | SALT00000039253_Cluster21547-003 |
| Cluster2155 | up | up | up | - | up | up | heat-shock 101 [Zea mays] | SALT00000018607_Cluster2155-001 |
| Cluster21552 | - | - | up | - | up | up | [Hordeum vulgare vulgare] | SALT00000051873_Cluster21552-001 |
| Cluster21574 | - | - | down | - | - | - | probable pyridoxal 5 -phospha | SALT00000058156_Cluster21574-001 |
| Cluster2158 | - | - | up | - | up | up | initiation factor (iso)4f p82 su | SALT00000073654_Cluster2158-022 |
| Cluster21585 | - | - | up | - | up | up | allene oxide synthase chloropl | SALT00000040195_Cluster21585-005 |
| Cluster21594 | - | - | down | - | down | down | polyol transporter 4 [Zea may | SALT00000053167_Cluster21594-001 |
| Cluster21597 | - | - | up | - | up | up | sterol C-22 desaturase -like [( | SALT00000060225_Cluster21597-004 |
| Cluster21600 | - | - | down | - | - | - | ribulose-1,5-bisphosphate carl | SALT00000061112_Cluster21600-003 |
| Cluster21618 | - | - | up | - | up | up | Acyl-coenzyme A oxidase per | SALT00000005190_Cluster21618-004 |
| Cluster21622 | - | up | - | - | - | - | Os03g0223100 [Oryza sativa | SALT00000039676_Cluster21622-004 |
| Cluster21623 | - | - | up | - | up | up | Os04g0379900 [Oryza sativa | SALT00000045222_Cluster21623-016 |
| Cluster21626 | - | - | up | - | up | up | plasma membrane H+-ATPase | SALT00000052345_Cluster21626-001 |
| Cluster21636 | - | - | up | - | - | up | Os02g0314600 [Oryza sativa | SALT00000014794_Cluster21636-002 |
| Cluster21659 | - | - | down | down | down | - | DNA-3-methyladenine glycos | SALT00000049578_Cluster21659-003 |
| Cluster21666 | - | - | down | - | - | - | transcription factor PCF6 [Or | SALT00000001735_Cluster21666-001 |
| Cluster21668 | - | - | - | - | down | down | [Hordeum vulgare vulgare] | SALT00000013817_Cluster21668-010 |
| Cluster21681 | - | - | down | - | down | down | peptide-N4-asparagine amidas | SALT00000028202_Cluster21681-003 |
| Cluster21683 | - | - | down | - | - | - | ZINC INDUCED FACILITAT | SALT00000039659_Cluster21683-004 |
| Cluster21691 | up | up | up | - | up | up | CIPK 1 [Zea mays] | SALT00000017140_Cluster21691-007 |
| Cluster21693 | - | - | up | - | up | up | HGA1 [Oryza sativa Japonica | SALT00000042022_Cluster21693-003 |
| Cluster21717 | - | - | up | - | up | up | cystathionine gamma-synthase | SALT00000011853_Cluster21717-004 |
| Cluster21718 | - | - | - | - | down | down | Grx_S17 - glutaredoxin subgr | SALT00000035112_Cluster21718-007 |
| Cluster2172 | - | - | down | - | down | - | RNA polymerase alpha subun | SALT00000087656_Cluster2172-008 |
| Cluster21721 | - | - | up | - | up | up | anthranilate N-benzoyltransfe | SALT00000040604_Cluster21721-005 |
| Cluster21723 | - | - | up | - | - | - | Os01g0259200 [Oryza sativa | SALT00000048254_Cluster21723-002 |
| Cluster21729 | - | down | down | - | - | - | Os02g0733300 [Oryza sativa | SALT00000005162_Cluster21729-001 |
| Cluster21742 | - | - | up | - | up | up | heat shock factor HSF30 [Zea | SALT00000077122_Cluster21742-003 |
| Cluster21763 | - | - | down | - | - | - | Os05g0556400 [Oryza sativa | SALT00000078986_Cluster21763-007 |
| Cluster21779 | down | down | down | - | - | - | metal transporter Nramp3 [Ze | SALT00000014206_Cluster21779-004 |

| | | | | | | | | |
|---|---|---|---|---|---|---|---|---|
| Cluster21780 | - | - | down | - | down | - | eukaryotic translation initiatio | SALT00000037213_Cluster21780-001 |
| Cluster21797 | - | - | up | - | up | up | LRP1 [Oryza sativa Japonica | SALT00000037413_Cluster21797-001 |
| Cluster21798 | - | - | down | - | - | - | probable serine threonine- kin | SALT00000038140_Cluster21798-002 |
| Cluster21799 | - | - | up | - | up | up | kinase superfamily | SALT00000038212_Cluster21799-001 |
| Cluster21801 | - | - | - | - | - | down | Plasma membrane ATPase 1 | SALT00000044370_Cluster21801-001 |
| Cluster21802 | - | - | up | - | up | up | pentatricopeptide repeat-conta | SALT00000047173_Cluster21802-001 |
| Cluster21807 | - | - | up | - | - | - | [Hordeum vulgare vulgare] | SALT00000054309_Cluster21807-005 |
| Cluster2181 | up | up | up | - | - | - | probable proline transporter 2 | SALT00000061675_Cluster2181-001 |
| Cluster21810 | - | - | - | - | down | - | ribosomal S5 [Oryza sativa Ja | SALT00000067210_Cluster21810-005 |
| Cluster21829 | - | - | - | - | down | - | chitinase domain-containing 1 | SALT00000009434_Cluster21829-001 |
| Cluster21830 | - | - | up | - | up | up | ABC1 At2g40090 [Setaria ital | SALT00000012001_Cluster21830-002 |
| Cluster21847 | - | up | up | up | up | up | aldehyde dehydrogenase fami | SALT00000005054_Cluster21847-004 |
| Cluster21849 | - | - | - | - | down | - | 3-ketoacyl- synthase 6-like | SALT00000035950_Cluster21849-001 |
| Cluster21864 | - | - | down | - | - | - | translocon Tic40 [Zea mays] | SALT00000015441_Cluster21864-003 |
| Cluster21865 | - | - | down | - | - | down | Os06g0152800 [Oryza sativa | SALT00000016014_Cluster21865-001 |
| Cluster21878 | - | - | up | - | - | - | Os03g0752700 [Oryza sativa | SALT00000015085_Cluster21878-002 |
| Cluster21895 | - | down | - | - | - | - | 3 -N-debenzoyl-2 -deoxytaxol | SALT00000020150_Cluster21895-005 |
| Cluster219 | - | - | down | - | down | down | gamma-tubulin complex com | SALT00000089285_Cluster219-001 |
| Cluster21907 | - | - | up | - | up | up | Os05g0366600 [Oryza sativa | SALT00000056406_Cluster21907-003 |
| Cluster21918 | - | - | down | - | - | - | transparent testa 12 [Zea may | SALT00000040745_Cluster21918-001 |
| Cluster21920 | - | - | up | - | - | - | GDA2 [Oryza sativa Japonica | SALT00000048484_Cluster21920-002 |
| Cluster21924 | - | - | up | - | up | up | proline transporter 1-like | SALT00000004070_Cluster21924-001 |
| Cluster2193 | - | - | down | - | down | down | [Physcomitrella patens] | SALT00000046732_Cluster2193-001 |
| Cluster21931 | - | - | down | - | down | down | Os04g0602800 [Oryza sativa | SALT00000038683_Cluster21931-001 |
| Cluster2194 | - | up | up | - | up | up | ELMO domain-containing 2 [ | SALT00000075816_Cluster2194-007 |
| Cluster21940 | - | - | up | - | - | - | 4-coumarate-- ligase-like 4 | SALT00000002579_Cluster21940-001 |
| Cluster21953 | - | - | up | - | - | - | Os09g0493800 [Oryza sativa | SALT00000012745_Cluster21953-003 |
| Cluster21955 | - | - | down | - | down | down | Os01g0130400 [Oryza sativa | SALT00000045521_Cluster21955-001 |
| Cluster2196 | - | - | down | - | down | down | B3 domain-containing Os03g | SALT00000023798_Cluster2196-009 |
| Cluster21960 | - | up | up | - | up | up | NAC domain-containing 48 | SALT00000060776_Cluster21960-002 |
| Cluster21966 | - | - | down | - | - | - | Os08g0192900 [Oryza sativa | SALT00000003607_Cluster21966-001 |
| Cluster21972 | - | - | down | - | - | - | peroxidase 47 | SALT00000037965_Cluster21972-001 |
| Cluster21977 | - | - | up | - | up | up | jacalin-related lectin 3-like | SALT00000052488_Cluster21977-011 |
| Cluster21978 | - | - | up | - | up | - | [Hordeum vulgare vulgare] | SALT00000060519_Cluster21978-010 |
| Cluster21979 | - | - | down | down | down | - | [Hordeum vulgare vulgare] | SALT00000048383_Cluster21979-024 |
| Cluster2198 | - | - | down | - | down | - | receptor kinase 1 | SALT00000072752_Cluster2198-006 |
| Cluster21981 | - | - | down | - | - | - | Os03g0694500 [Oryza sativa | SALT00000015053_Cluster21981-002 |
| Cluster21992 | - | - | - | - | - | down | low temperature viability [Ze | SALT00000007716_Cluster21992-003 |
| Cluster22007 | - | - | down | - | down | down | AAA-ATPase At3g50940-like | SALT00000062678_Cluster22007-002 |
| Cluster22009 | - | - | - | - | down | - | aldehyde dehydrogenase fami | SALT00000009840_Cluster22009-004 |
| Cluster22018 | - | - | up | - | up | up | beta-mannan expressed [Oryz | SALT00000046159_Cluster22018-003 |
| Cluster22024 | - | - | - | - | up | - | J domain-containing required | SALT00000061836_Cluster22024-003 |
| Cluster22027 | - | - | up | - | up | up | LOC100281987 isoform X1 [ | SALT00000011920_Cluster22027-004 |
| Cluster22030 | - | - | up | - | up | up | aromatic-L-amino-acid decarb | SALT00000019154_Cluster22030-001 |
| Cluster22033 | - | - | up | - | up | up | phosphate transporter [Triticu | SALT00000059123_Cluster22033-001 |
| Cluster22059 | - | - | up | - | up | up | Os03g0339700 [Oryza sativa | SALT00000047433_Cluster22059-001 |
| Cluster22066 | - | - | up | - | up | up | fatty acid elongase [Zea mays | SALT00000057679_Cluster22066-010 |
| Cluster22074 | - | - | up | - | up | - | ---NA--- | SALT00000035806_Cluster22074-001 |
| Cluster22080 | - | - | - | - | - | up | ACR4 [Zea mays] | SALT00000058587_Cluster22080-003 |
| Cluster221 | - | - | down | - | - | - | Transcriptional adapter ADA2 | SALT00000091460_Cluster221-005 |
| Cluster22106 | - | - | up | - | up | - | regulatory [Oryza sativa Japo | SALT00000036369_Cluster22106-003 |
| Cluster22133 | - | - | - | - | - | up | Os01g0349400 [Oryza sativa | SALT00000020200_Cluster22133-001 |
| Cluster22135 | - | - | up | - | up | up | pyruvate dehydrogenase kinas | SALT00000041018_Cluster22135-001 |
| Cluster22144 | - | up | up | - | up | - | TIC chloroplastic | SALT00000001500_Cluster22144-002 |
| Cluster22156 | - | up | up | - | up | up | [Hordeum vulgare vulgare] | SALT00000039221_Cluster22156-001 |
| Cluster22161 | - | - | - | - | down | - | probable transcription factor | SALT00000047678_Cluster22161-002 |
| Cluster22179 | - | - | down | - | down | down | peptide transporter PTR2 [Ze | SALT00000056624_Cluster22179-001 |
| Cluster2218 | up | - | up | down | - | up | phosphoenolpyruvate carboxy | SALT00000078744_Cluster2218-001 |

| | | | | | | | | |
|---|---|---|---|---|---|---|---|---|
| Cluster22184 | - | - | up | - | up | up | aspartic ase nepenthesin-2 pre | SALT00000004477_Cluster22184-002 |
| Cluster22185 | - | - | up | - | - | - | polyprenol reductase 1 [Oryza | SALT00000011969_Cluster22185-002 |
| Cluster22192 | - | - | - | - | down | - | GATA zinc finger family [Zea | SALT00000045544_Cluster22192-005 |
| Cluster22217 | - | - | down | - | - | - | xyloglucan endotransglucosyl | SALT00000036796_Cluster22217-001 |
| Cluster2222 | - | - | up | - | - | - | leucine-rich repeat-containing | SALT00000023901_Cluster2222-005 |
| Cluster22232 | - | - | down | - | down | down | thylakoid lumenal 19 kDa [Ze | SALT00000013323_Cluster22232-001 |
| Cluster22236 | - | - | up | - | up | up | Os03g0123800 [Oryza sativa | SALT00000044997_Cluster22236-001 |
| Cluster2224 | - | - | up | - | up | - | multidrug resistance associate | SALT00000077126_Cluster2224-003 |
| Cluster22244 | - | - | up | - | up | up | Os05g0447700 [Oryza sativa | SALT00000040185_Cluster22244-001 |
| Cluster22252 | - | - | up | - | - | - | xaa-Pro dipeptidase | SALT00000064093_Cluster22252-010 |
| Cluster22269 | - | - | - | - | up | up | inactive tetrahydrocannabinol | SALT00000001226_Cluster22269-001 |
| Cluster22286 | - | up | up | - | - | - | cytochrome P450 [Oryza sativ | SALT00000007987_Cluster22286-001 |
| Cluster22291 | - | - | down | - | down | down | Os03g0220100 [Oryza sativa | SALT00000020398_Cluster22291-008 |
| Cluster22294 | - | - | up | - | - | - | Os01g0656200 [Oryza sativa | SALT00000045332_Cluster22294-003 |
| Cluster22300 | - | - | down | - | - | - | AUX IAA transcription partia | SALT00000057379_Cluster22300-004 |
| Cluster22301 | - | - | - | - | up | - | 6b-interacting 1 [Zea mays] | SALT00000057661_Cluster22301-002 |
| Cluster22302 | - | - | up | - | up | up | beclin 1 [Zea mays] | SALT00000059774_Cluster22302-002 |
| Cluster22306 | - | - | up | - | up | - | ras-related RABH1e-like | SALT00000004233_Cluster22306-001 |
| Cluster22313 | - | - | down | - | - | - | NAP16kDa [Zea mays] | SALT00000044717_Cluster22313-001 |
| Cluster22316 | - | - | down | - | down | down | Os08g0374000 [Oryza sativa | SALT00000054489_Cluster22316-001 |
| Cluster22321 | - | - | down | - | down | - | bolA [Zea mays] | SALT00000002713_Cluster22321-001 |
| Cluster2234 | - | - | down | - | down | down | tubulin alpha chain [Malus do | SALT00000016622_Cluster2234-010 |
| Cluster2235 | - | - | down | - | down | - | DEAD-box ATP-dependent R | SALT00000065412_Cluster2235-001 |
| Cluster22354 | - | - | down | - | - | - | [Oryza sativa Japonica Group | SALT00000005287_Cluster22354-001 |
| Cluster22356 | - | - | up | - | - | - | TPA: phosphoglycerate mutas | SALT00000009149_Cluster22356-003 |
| Cluster22365 | - | - | down | - | down | down | binding [Zea mays] | SALT00000048810_Cluster22365-004 |
| Cluster22401 | up | up | down | - | down | down | Disease resistance RPP8 3 | SALT00000002719_Cluster22401-001 |
| Cluster22405 | - | - | down | - | - | - | Os02g0815400 [Oryza sativa | SALT00000035747_Cluster22405-002 |
| Cluster22412 | - | - | - | - | down | - | Os04g0477500 [Oryza sativa | SALT00000058758_Cluster22412-001 |
| Cluster22421 | - | - | up | - | up | up | HGA6 [Oryza sativa Japonica | SALT00000043940_Cluster22421-006 |
| Cluster22423 | - | - | - | - | up | up | nuclease HARBI1 | SALT00000060556_Cluster22423-016 |
| Cluster22427 | - | - | up | - | up | up | [Hordeum vulgare vulgare] | SALT00000008973_Cluster22427-002 |
| Cluster22430 | - | - | down | - | - | - | peroxidase 72-like [Setaria ita | SALT00000012620_Cluster22430-002 |
| Cluster2244 | - | - | up | - | up | up | E3 ubiquitin- ligase MBR1 iso | SALT00000089827_Cluster2244-008 |
| Cluster22448 | - | - | up | - | up | up | neutral invertase [Oryza sativa | SALT00000018689_Cluster22448-001 |
| Cluster22452 | - | - | - | - | up | - | chorismate mutase [Zea mays | SALT00000041079_Cluster22452-002 |
| Cluster22473 | - | - | up | - | up | up | inositol-3-phosphate synthase | SALT00000015190_Cluster22473-004 |
| Cluster22474 | - | - | up | - | up | up | HGA6 [Oryza sativa Japonica | SALT00000019476_Cluster22474-008 |
| Cluster22475 | - | - | up | - | up | up | probable UDP-arabinose 4-ep | SALT00000020458_Cluster22475-005 |
| Cluster22487 | - | - | up | - | up | up | inactive beta-amylase 9-like | SALT00000002042_Cluster22487-001 |
| Cluster22488 | - | - | - | - | down | - | EMB1688 [Zea mays] | SALT00000002511_Cluster22488-001 |
| Cluster22495 | - | - | up | - | - | - | senescence-associated [Zea m | SALT00000037362_Cluster22495-005 |
| Cluster22508 | - | - | up | - | up | up | amino acid transport [Zea may | SALT00000053264_Cluster22508-004 |
| Cluster22514 | - | - | up | - | - | - | [Oryza sativa Japonica Group | SALT00000062848_Cluster22514-002 |
| Cluster22529 | down | down | down | - | - | - | AUX IAA transcription partia | SALT00000053209_Cluster22529-002 |
| Cluster22530 | - | - | up | - | - | - | p8MTCP1 [Zea mays] | SALT00000054330_Cluster22530-001 |
| Cluster22548 | - | - | up | - | up | up | cysteine protease 1 precursor | | SALT00000008233_Cluster22548-001 |
| Cluster22565 | - | - | down | - | - | - | Os03g0646100 [Oryza sativa | SALT00000064586_Cluster22565-010 |
| Cluster22570 | - | up | up | - | up | up | actin-7-like [Brachypodium d | SALT00000015977_Cluster22570-003 |
| Cluster22571 | - | up | up | - | up | up | pheophorbide a chloroplastic | SALT00000017945_Cluster22571-002 |
| Cluster2258 | - | - | down | - | - | down | ATP synthase subunit chlorop | SALT00000080365_Cluster2258-001 |
| Cluster22594 | - | - | up | - | up | up | HGA1 [Oryza sativa Japonica | SALT00000011045_Cluster22594-001 |
| Cluster226 | - | - | down | - | down | down | Os01g0534900 [Oryza sativa | SALT00000080033_Cluster226-014 |
| Cluster22608 | - | up | up | up | up | up | PREDICTED: uncharacterized | SALT00000042705_Cluster22608-001 |
| Cluster22610 | - | - | - | - | down | - | O-methyltransferase [Zea may | SALT00000046926_Cluster22610-001 |
| Cluster22636 | - | - | down | - | - | - | sigma factor SIG2A [Zea may | SALT00000000254_Cluster22636-001 |
| Cluster22642 | - | - | - | - | down | - | chloroplastic [Setaria italica] | SALT00000041393_Cluster22642-001 |
| Cluster22644 | - | - | - | - | - | down | Double-stranded RNA binding | SALT00000043941_Cluster22644-007 |

| | | | | | | | | |
|---|---|---|---|---|---|---|---|---|
| Cluster22649 | - | - | up | - | up | up | grancalcin [Zea mays] | SALT00000051967_Cluster22649-002 |
| Cluster22650 | - | - | - | - | - | down | LOW PSII ACCUMULATIOI | SALT00000052071_Cluster22650-001 |
| Cluster22665 | - | - | - | - | down | - | hepatoma-derived growth fact | SALT00000046811_Cluster22665-003 |
| Cluster22676 | - | - | up | - | up | up | WRKY53 - superfamily of TF | SALT00000017567_Cluster22676-002 |
| Cluster22677 | - | - | down | - | - | - | uncharacterized LOC1002170 | SALT00000019422_Cluster22677-003 |
| Cluster22689 | - | - | - | - | down | - | ferrochelatase II [Oryza sativa | SALT00000016003_Cluster22689-002 |
| Cluster22695 | - | - | down | - | - | down | TCP transcription partial [Zea | SALT00000037682_Cluster22695-002 |
| Cluster22700 | - | up | - | - | - | - | chloride channel CLC-c-like [ | SALT00000056036_Cluster22700-001 |
| Cluster22704 | - | - | - | - | - | down | nucleolysin TIAR | SALT00000068301_Cluster22704-001 |
| Cluster22711 | - | - | down | - | - | - | acetate butyrate-- ligase perox | SALT00000007076_Cluster22711-003 |
| Cluster22712 | - | - | - | - | up | up | probable acyl-activating enzy | SALT00000045378_Cluster22712-002 |
| Cluster22719 | - | - | up | - | up | up | HGA1 [Oryza sativa Japonica | SALT00000001074_Cluster22719-001 |
| Cluster22722 | - | down | down | - | - | - | TPA: O-glycosyl hydrolase fa | SALT00000041794_Cluster22722-003 |
| Cluster22735 | - | - | - | - | down | - | heterogeneous nuclear ribonu | SALT00000039634_Cluster22735-007 |
| Cluster22739 | - | - | up | - | up | up | glucosyltransferase-10 [Oryza | SALT00000047264_Cluster22739-002 |
| Cluster22761 | - | - | up | - | - | - | TFIIB [Zea mays] | SALT00000000577_Cluster22761-001 |
| Cluster22763 | - | - | up | - | - | - | 3-ketoacyl- synthase 11 [Oryz | SALT00000004062_Cluster22763-001 |
| Cluster22764 | - | - | - | - | up | - | ATP-dependent Clp protease | SALT00000004653_Cluster22764-003 |
| Cluster22774 | - | - | down | - | down | - | [Hordeum vulgare vulgare] | SALT00000036525_Cluster22774-001 |
| Cluster22777 | - | - | - | - | down | - | DRE binding factor 2 [Oryza | SALT00000049073_Cluster22777-006 |
| Cluster22826 | - | up | - | - | - | - | Os05g0195200 [Oryza sativa | SALT00000004039_Cluster22826-001 |
| Cluster22838 | - | - | - | - | down | - | coleoptile phototropism 1 [Br | SALT00000039770_Cluster22838-001 |
| Cluster2284 | - | - | - | - | up | - | Os01g0927600 [Oryza sativa | SALT00000063295_Cluster2284-017 |
| Cluster22843 | - | - | up | - | - | - | cytochrome P450 72A14-like | SALT00000058597_Cluster22843-002 |
| Cluster22844 | - | - | up | - | up | up | rRNA methyltransferase mito | SALT00000012224_Cluster22844-001 |
| Cluster22845 | - | - | up | - | up | - | 5-enolpyruvylshikimate-3-ph | SALT00000017943_Cluster22845-003 |
| Cluster22846 | - | - | down | - | - | - | probable DNA primase large s | SALT00000017960_Cluster22846-002 |
| Cluster22847 | - | - | down | - | down | - | chromatin assembly factor 1 s | SALT00000018897_Cluster22847-001 |
| Cluster22848 | - | - | down | - | - | - | [Hordeum vulgare vulgare] | SALT00000018969_Cluster22848-001 |
| Cluster22849 | - | - | - | - | down | - | cyclin-B1-1-like isoform X2 [ | SALT00000042876_Cluster22849-003 |
| Cluster22850 | - | - | up | - | - | - | vacuolar ATP synthase 16 kD | SALT00000043765_Cluster22850-001 |
| Cluster22865 | - | - | down | - | down | down | cytokinin-O-glucosyltransfera | SALT00000064527_Cluster22865-002 |
| Cluster22869 | - | - | up | - | up | up | Os12g0512100 [Oryza sativa | SALT00000014416_Cluster22869-001 |
| Cluster22870 | - | - | down | - | - | - | galactolipase chloroplastic-lik | SALT00000023961_Cluster22870-001 |
| Cluster22873 | - | down | down | - | - | - | Os01g0358300 [Oryza sativa | SALT00000040048_Cluster22873-002 |
| Cluster2288 | - | - | - | - | down | - | endoglucanase 22-like isoforn | SALT00000059019_Cluster2288-001 |
| Cluster22889 | - | - | up | - | up | up | galactoside 2-alpha-L-fucosyl | SALT00000003249_Cluster22889-001 |
| Cluster22910 | - | - | up | - | up | up | cytochrome P450 734A6-like | SALT00000001818_Cluster22910-001 |
| Cluster22911 | - | - | up | - | up | up | [Hordeum vulgare vulgare] | SALT00000009578_Cluster22911-002 |
| Cluster22916 | - | - | up | - | up | up | EF hand family expressed [Or | SALT00000018191_Cluster22916-001 |
| Cluster22917 | - | - | down | - | - | - | Os07g0205500 [Oryza sativa | SALT00000045611_Cluster22917-002 |
| Cluster22922 | - | - | - | - | down | - | chloroplastic-like [Setaria ital | SALT00000057928_Cluster22922-004 |
| Cluster22925 | - | - | up | - | up | up | cytochrome P450 94B3-like | SALT00000003515_Cluster22925-001 |
| Cluster2293 | - | down | down | - | - | - | phosphoenolpyruvate carboxy | SALT00000077272_Cluster2293-010 |
| Cluster22936 | - | - | up | - | up | up | [Hordeum vulgare vulgare] | SALT00000036377_Cluster22936-001 |
| Cluster22937 | - | - | - | - | down | - | homeodomain leucine zipper | SALT00000041276_Cluster22937-014 |
| Cluster22940 | - | - | up | - | up | up | Os05g0447700 [Oryza sativa | SALT00000052300_Cluster22940-001 |
| Cluster22943 | - | - | down | - | down | - | ribonuclease H At1g65750 isc | SALT00000061786_Cluster22943-001 |
| Cluster22949 | - | - | down | - | down | - | Os08g0276100 [Oryza sativa | SALT00000018917_Cluster22949-004 |
| Cluster22953 | - | - | up | - | up | up | probable phosphatase 2C 34 [ | SALT00000051927_Cluster22953-016 |
| Cluster22963 | - | - | down | - | - | - | porphobilinogen deaminase [ | SALT00000042194_Cluster22963-002 |
| Cluster22973 | - | - | down | - | - | - | Os02g0805000 [Oryza sativa | SALT00000083840_Cluster22973-001 |
| Cluster22976 | - | - | down | - | - | - | 3-oxoacyl-synthase I [Zea ma | SALT00000005486_Cluster22976-002 |
| Cluster22979 | - | up | - | - | down | down | cytochrome b561 and DOMO | SALT00000006314_Cluster22979-001 |
| Cluster22987 | up | - | up | - | up | up | monoglyceride lipase [Setaria | SALT00000059977_Cluster22987-003 |
| Cluster22998 | - | - | - | - | down | down | chloroplast thylakoid-bound a | SALT00000019440_Cluster22998-004 |
| Cluster23 | - | - | down | - | down | down | phosphoenolpyruvate carboxy | SALT00000065665_Cluster23-077 |
| Cluster2300 | - | down | down | - | down | down | ribonucleotide reductase [Ory | SALT00000076498_Cluster2300-010 |

| | | | | | | | | |
|---|---|---|---|---|---|---|---|---|
| Cluster23005 | - | - | - | down | down | - | uncharacterized protein LOC1 | SALT00000050142_Cluster23005-002 |
| Cluster23013 | up | up | up | - | - | - | CBL-interacting serine threon | SALT00000013302_Cluster23013-003 |
| Cluster23015 | - | - | down | - | down | - | protoporphyrinogen oxidase I | SALT00000014294_Cluster23015-002 |
| Cluster23024 | - | - | down | - | - | - | proline--tRNA chloroplastic n | SALT00000056394_Cluster23024-004 |
| Cluster23026 | - | - | up | - | up | up | exonuclease [Zea mays] | SALT00000000695_Cluster23026-001 |
| Cluster23029 | - | - | up | - | - | - | CBL-interacting kinase 1 [Ory | SALT00000004985_Cluster23029-002 |
| Cluster23031 | - | - | - | - | down | - | acyltransferase [Zea mays] | SALT00000015544_Cluster23031-001 |
| Cluster23058 | - | - | up | - | - | - | F-box SKP2A [Brachypodium | SALT00000066151_Cluster23058-006 |
| Cluster23064 | - | - | up | - | up | up | Os08g0127100 [Oryza sativa | SALT00000014248_Cluster23064-002 |
| Cluster23065 | - | - | up | - | - | - | acc oxidase [Zea mays] | SALT00000072611_Cluster23065-003 |
| Cluster23069 | - | - | - | - | down | - | phosphatidylcholine transfer - | SALT00000040892_Cluster23069-005 |
| Cluster23070 | - | - | up | - | up | up | pectinesterase [Oryza sativa Ja | SALT00000042477_Cluster23070-001 |
| Cluster23081 | up | up | up | - | up | up | geraniol 8-hydroxylase-like [Z | SALT00000020236_Cluster23081-002 |
| Cluster23087 | - | - | up | - | up | up | ribosome biogenesis TSR3 ho | SALT00000057827_Cluster23087-004 |
| Cluster2309 | - | - | up | - | up | up | Os07g0693500 [Oryza sativa | SALT00000035158_Cluster2309-012 |
| Cluster23090 | - | - | - | - | up | - | Os08g0500200 [Oryza sativa | SALT00000009827_Cluster23090-001 |
| Cluster23091 | - | - | - | - | down | - | glucan endo-1,3-beta-glucosic | SALT00000019224_Cluster23091-006 |
| Cluster23104 | - | - | - | - | down | - | Os03g0197100 [Oryza sativa | SALT00000060988_Cluster23104-013 |
| Cluster23109 | - | - | - | - | down | - | Os01g0695200 [Oryza sativa | SALT00000054127_Cluster23109-006 |
| Cluster23113 | - | - | up | - | up | up | Os01g0855600 [Oryza sativa | SALT00000050184_Cluster23113-014 |
| Cluster2312 | - | - | - | - | - | up | ---NA--- | SALT00000085561_Cluster2312-004 |
| Cluster23126 | - | - | down | - | down | - | Os01g0110800 [Oryza sativa | SALT00000038950_Cluster23126-001 |
| Cluster23146 | - | up | up | - | up | up | limonoid UDP-glucosyltransf | SALT00000039200_Cluster23146-004 |
| Cluster23159 | - | - | - | - | down | down | carbonic anhydrase 2 | SALT00000002208_Cluster23159-001 |
| Cluster23166 | - | - | up | - | - | - | Os06g0289200 [Oryza sativa | SALT00000045070_Cluster23166-003 |
| Cluster23182 | - | - | up | - | up | up | TPA: cytochrome P450 superf | SALT00000050759_Cluster23182-001 |
| Cluster23183 | - | down | down | down | down | down | Tyrosine N-monooxygenase | SALT00000050807_Cluster23183-001 |
| Cluster23197 | - | - | up | - | up | up | LOC100282132 isoform X1 | SALT00000046426_Cluster23197-002 |
| Cluster2322 | down | down | down | down | - | - | beta-D-xylosidase [Oryza sati | SALT00000062760_Cluster2322-011 |
| Cluster23224 | up | up | up | - | - | - | MLO 1 | SALT00000010764_Cluster23224-002 |
| Cluster23228 | - | - | up | - | up | - | Os02g0682500 [Oryza sativa | SALT00000022221_Cluster23228-002 |
| Cluster23229 | - | - | - | - | down | down | glyceraldehyde-3-phosphate d | SALT00000035801_Cluster23229-001 |
| Cluster23230 | - | - | down | - | - | - | UMP synthase | SALT00000037068_Cluster23230-002 |
| Cluster23233 | up | up | - | - | - | - | Os01g0183600 [Oryza sativa | SALT00000048083_Cluster23233-001 |
| Cluster23242 | - | down | - | - | - | up | tubulin alpha chain | SALT00000013351_Cluster23242-002 |
| Cluster23243 | - | - | - | - | up | up | capI [Zea mays] | SALT00000016480_Cluster23243-001 |
| Cluster23257 | - | - | down | - | - | - | cytochrome P450 [Echinochlo | SALT00000007761_Cluster23257-001 |
| Cluster23267 | - | - | up | - | up | up | Os03g0324700 [Oryza sativa | SALT00000051780_Cluster23267-001 |
| Cluster23269 | - | up | up | - | up | up | amino acid transport [Zea may | SALT00000052502_Cluster23269-004 |
| Cluster23280 | - | - | - | - | up | - | Enolase 1 | SALT00000040435_Cluster23280-007 |
| Cluster23288 | - | - | up | - | up | up | aldose reductase [Zea mays] | SALT00000072232_Cluster23288-002 |
| Cluster23297 | - | - | up | - | - | - | integral membrane like [Zea n | SALT00000046265_Cluster23297-007 |
| Cluster23309 | down | down | down | - | down | - | SEC14 cytosolic factor-like [( | SALT00000039051_Cluster23309-005 |
| Cluster23314 | - | - | up | - | up | up | sugar transport 13 | SALT00000006347_Cluster23314-002 |
| Cluster23330 | - | up | up | up | up | up | vacuolar amino acid transport | SALT00000048359_Cluster23330-001 |
| Cluster23331 | - | down | down | - | - | - | ATP sulfurylase 2 | SALT00000052678_Cluster23331-004 |
| Cluster23339 | - | up | up | up | up | up | hioredoxin 5 [Oryza sativa Ja | SALT00000010882_Cluster23339-001 |
| Cluster23340 | - | - | up | - | up | up | Os05g0134000 [Oryza sativa | SALT00000011940_Cluster23340-003 |
| Cluster23347 | - | - | up | - | - | - | inactive beta-amylase 9-like | SALT00000047346_Cluster23347-002 |
| Cluster23353 | - | - | - | - | down | - | Os03g0717600 [Oryza sativa | SALT00000001805_Cluster23353-002 |
| Cluster23355 | - | - | up | - | up | up | Os11g0704600 [Oryza sativa | SALT00000060598_Cluster23355-006 |
| Cluster23362 | - | up | up | - | up | up | vacuolar cation proton exchan | SALT00000039168_Cluster23362-004 |
| Cluster23370 | - | - | up | - | - | - | serine threonine- kinase NAK | SALT00000053852_Cluster23370-006 |
| Cluster23381 | - | down | - | - | - | - | pathogenesis-related transcrip | SALT00000014734_Cluster23381-003 |
| Cluster23383 | - | - | up | - | up | up | Avr9 Cf-9 rapidly elicited [Or | SALT00000042282_Cluster23383-004 |
| Cluster23384 | - | - | up | - | up | up | Os08g0127100 [Oryza sativa | SALT00000046604_Cluster23384-008 |
| Cluster23387 | - | - | up | - | up | up | Os12g0257000 [Oryza sativa | SALT00000056282_Cluster23387-007 |
| Cluster23389 | - | - | - | - | down | - | 4-hydroxy-4-methyl-2-oxoglu | SALT00000002580_Cluster23389-001 |

| | | | | | | | | |
|---|---|---|---|---|---|---|---|---|
| Cluster23390 | - | - | up | - | up | up | WRKY53 - superfamily of TF | SALT00000009228_Cluster23390-004 |
| Cluster23393 | - | - | - | - | down | - | CCT motif family [Zea mays] | SALT00000013620_Cluster23393-005 |
| Cluster23396 | - | - | down | - | down | down | 60S ribosomal L28 [Zea mays | SALT00000037009_Cluster23396-002 |
| Cluster23406 | - | - | up | - | - | - | DNA binding [Zea mays] | SALT00000053407_Cluster23406-003 |
| Cluster23416 | - | - | down | - | down | down | High mobility group family is | SALT00000008594_Cluster23416-004 |
| Cluster23417 | - | - | down | - | down | down | Os11g0267000 [Oryza sativa | SALT00000040166_Cluster23417-001 |
| Cluster23433 | - | - | - | - | down | down | peroxidase 72-like [Setaria ita | SALT00000037308_Cluster23433-005 |
| Cluster23435 | - | - | up | - | - | - | glutathione chloroplastic | SALT00000042380_Cluster23435-001 |
| Cluster23447 | - | - | - | - | - | up | triacylglycerol lipase like [Zea | SALT00000003200_Cluster23447-001 |
| Cluster23457 | - | up | up | - | - | - | FAR1-RELATED SEQUENC | SALT00000045339_Cluster23457-001 |
| Cluster2346 | - | - | down | down | down | down | Os09g0487200 [Oryza sativa | SALT00000081743_Cluster2346-003 |
| Cluster23471 | - | - | up | - | up | up | alpha beta fold family | SALT00000011093_Cluster23471-002 |
| Cluster23472 | - | - | up | - | up | up | cinnamate 4-hydroxylase 1 | SALT00000013684_Cluster23472-001 |
| Cluster23474 | - | - | up | - | - | up | fiber Fb2 [Zea mays] | SALT00000042024_Cluster23474-001 |
| Cluster23479 | - | - | down | - | down | - | carboxy-lyase [Zea mays] | SALT00000057616_Cluster23479-002 |
| Cluster2348 | - | - | down | - | down | - | Os01g0737900 [Oryza sativa | SALT00000062527_Cluster2348-001 |
| Cluster23480 | - | - | - | - | down | - | polygalacturonase precursor [ | SALT00000003532_Cluster23480-001 |
| Cluster23489 | - | - | - | - | down | - | Os06g0264200 [Oryza sativa | SALT00000004202_Cluster23489-002 |
| Cluster2349 | - | - | down | - | down | - | powdery mildew resistance ex | SALT00000090773_Cluster2349-006 |
| Cluster23496 | - | - | up | - | up | up | F-box interaction domain con | SALT00000043274_Cluster23496-001 |
| Cluster23497 | - | - | down | - | down | - | Carboxyl-terminal expressed | | SALT00000043770_Cluster23497-005 |
| Cluster235 | - | - | - | - | down | - | [Oryza sativa Japonica Group | SALT00000052817_Cluster235-004 |
| Cluster23506 | - | - | up | - | up | - | kinase APK1A [Zea mays] | SALT00000017710_Cluster23506-008 |
| Cluster23509 | up | - | - | down | - | - | ribosomal -like | SALT00000049176_Cluster23509-001 |
| Cluster23512 | - | - | - | - | down | - | tyrosine--tRNA chloroplastic | SALT00000058235_Cluster23512-005 |
| Cluster23515 | - | - | down | - | - | - | glycerol-3-phosphate 2-O-acy | SALT00000005866_Cluster23515-002 |
| Cluster2352 | - | - | up | - | - | - | ras-related Rab-18 [Zea mays | SALT00000091211_Cluster2352-001 |
| Cluster23522 | - | - | up | - | up | up | agmatine coumaroyltransferas | SALT00000045259_Cluster23522-005 |
| Cluster23523 | - | - | up | - | - | - | serine esterase family [Zea ma | SALT00000045453_Cluster23523-001 |
| Cluster23535 | - | - | up | - | up | up | AAA-ATPase At3g28580-like | SALT00000017255_Cluster23535-002 |
| Cluster23543 | - | - | up | - | - | up | UDP-glycosyltransferase 73C | SALT00000053123_Cluster23543-003 |
| Cluster23548 | - | - | up | - | up | up | [Hordeum vulgare vulgare] | SALT00000003948_Cluster23548-002 |
| Cluster23549 | - | - | up | - | up | up | Os02g0150900 [Oryza sativa | SALT00000010846_Cluster23549-004 |
| Cluster23554 | - | - | down | - | down | down | binding [Zea mays] | SALT00000041042_Cluster23554-007 |
| Cluster23555 | - | up | - | - | - | - | F-box At5g46170-like [Brach | SALT00000044230_Cluster23555-004 |
| Cluster23558 | - | - | up | - | up | up | probable polyol transporter 6 | SALT00000058289_Cluster23558-002 |
| Cluster23563 | - | - | up | - | - | - | vacuolar cation proton exchan | SALT00000007337_Cluster23563-001 |
| Cluster23564 | - | down | down | - | down | - | rhodanese-like domain-contai | SALT00000007933_Cluster23564-001 |
| Cluster23579 | - | - | - | - | down | - | glycosyltransferase [Zea mays | SALT00000056599_Cluster23579-012 |
| Cluster23593 | - | - | up | - | up | up | 2-oxoisovalerate dehydrogena | SALT00000051120_Cluster23593-007 |
| Cluster23597 | - | - | up | - | up | up | Os01g0185300 [Oryza sativa | SALT00000060690_Cluster23597-007 |
| Cluster23604 | - | - | down | - | - | down | Pre translocase secY chloropla | SALT00000037817_Cluster23604-003 |
| Cluster23605 | - | - | up | - | up | up | kinase KIPK [Zea mays] | SALT00000038172_Cluster23605-003 |
| Cluster2361 | - | - | down | - | - | - | Os02g0806600 [Oryza sativa | SALT00000083273_Cluster2361-009 |
| Cluster23611 | - | - | up | - | up | up | Os03g0760200 [Oryza sativa | SALT00000045266_Cluster23611-001 |
| Cluster23622 | - | - | up | - | - | - | spastin [Oryza sativa Japonica | SALT00000065239_Cluster23622-002 |
| Cluster23623 | - | - | down | - | - | - | zinc finger CCCH type domai | SALT00000006189_Cluster23623-001 |
| Cluster23628 | - | - | up | - | up | up | ACT domain-containing ACR | SALT00000048309_Cluster23628-002 |
| Cluster23644 | down | down | - | - | - | - | carbohydrate transporter sugar | SALT00000058312_Cluster23644-004 |
| Cluster23646 | - | - | down | - | down | - | desaturase cytochrome b5 [Ze | SALT00000005420_Cluster23646-001 |
| Cluster23656 | - | - | down | - | down | - | AP2 EREBP transcription fact | SALT00000046967_Cluster23656-004 |
| Cluster23670 | up | up | up | - | - | - | flavonoid 3 ,5 -hydroxylase 1- | SALT00000038585_Cluster23670-004 |
| Cluster23672 | - | - | - | - | - | down | carbonic anhydrase [Zea mays | SALT00000041867_Cluster23672-002 |
| Cluster23686 | - | - | up | - | up | up | malonyl- :anthocyanidin 5-O- | SALT00000042630_Cluster23686-006 |
| Cluster23706 | - | down | down | - | - | - | AChain Semi-Active E176q N | SALT00000025144_Cluster23706-004 |
| Cluster23715 | - | - | up | - | up | up | probable phosphatase 2C 48 [ | SALT00000058839_Cluster23715-003 |
| Cluster23717 | - | up | up | up | up | up | Os03g0643300 [Oryza sativa | SALT00000002106_Cluster23717-002 |
| Cluster23719 | - | up | - | - | - | - | E3 ubiquitin- ligase XB3 | SALT00000004480_Cluster23719-002 |

| | | | | | | | | |
|---|---|---|---|---|---|---|---|---|
| Cluster23737 | - | - | - | - | up | - | IAA-amino acid hydrolase ILl | SALT00000035791_Cluster23737-002 |
| Cluster2374 | up | - | down | - | down | down | geraniol 8-hydroxylase-like | SALT00000036596_Cluster2374-001 |
| Cluster23740 | - | up | - | - | - | - | glutathione transferase [Tritic | SALT00000038465_Cluster23740-001 |
| Cluster23750 | - | down | - | - | - | - | growth-regulating factor 5-lik | SALT00000045503_Cluster23750-001 |
| Cluster23756 | - | - | down | - | - | - | probable glycerol-3-phosphat | SALT00000054782_Cluster23756-004 |
| Cluster23760 | - | - | down | - | - | - | leucine-rich repeat extensin 3 | SALT00000003116_Cluster23760-001 |
| Cluster23762 | - | - | up | - | up | up | Os02g0717500 [Oryza sativa | SALT00000015120_Cluster23762-003 |
| Cluster23779 | - | - | - | - | down | down | serine arginine-rich splicing fa | SALT00000036475_Cluster23779-010 |
| Cluster23784 | - | - | - | - | - | down | sulfate transporter chloroplast | SALT00000048660_Cluster23784-002 |
| Cluster238 | down | down | down | - | down | down | sucrose synthase 1 [Oryza sati | SALT00000023702_Cluster238-008 |
| Cluster23803 | - | - | up | - | up | up | auxin-responsive IAA18 isofo | SALT00000008765_Cluster23803-002 |
| Cluster23809 | up | up | up | - | up | - | NNJA1_MAIZE ame: Full=N | SALT00000058643_Cluster23809-003 |
| Cluster23826 | - | - | - | - | - | down | Adenosine kinase 2 | SALT00000065525_Cluster23826-002 |
| Cluster23837 | - | up | up | - | up | up | ammonium transporter 2 [Zea | SALT00000036077_Cluster23837-001 |
| Cluster23843 | - | - | - | - | - | down | 28 kDa chloroplastic-like | SALT00000045325_Cluster23843-001 |
| Cluster23846 | - | - | up | - | up | up | probable galacturonosyltransf | SALT00000051003_Cluster23846-002 |
| Cluster23848 | - | - | up | - | up | up | [Hordeum vulgare vulgare] | SALT00000054259_Cluster23848-001 |
| Cluster23851 | - | up | up | up | - | - | Os03g0739700 [Oryza sativa | SALT00000060472_Cluster23851-001 |
| Cluster23853 | - | - | down | - | down | down | partial [Oryza sativa Japonica | SALT00000066593_Cluster23853-003 |
| Cluster23856 | up | up | up | - | up | up | ABA responsive element bind | SALT00000004666_Cluster23856-002 |
| Cluster23857 | - | - | down | - | - | - | pseudouridylate synthase tran: | SALT00000009138_Cluster23857-001 |
| Cluster23863 | - | up | up | - | up | - | [Hordeum vulgare vulgare] | SALT00000037534_Cluster23863-002 |
| Cluster23866 | - | - | down | - | down | down | beta-amylase [Zea mays] | SALT00000043175_Cluster23866-001 |
| Cluster23869 | - | - | up | - | up | up | Secologanin synthase | SALT00000051460_Cluster23869-003 |
| Cluster23873 | - | - | - | - | - | down | Splicing factor U2AF subunit | SALT00000060649_Cluster23873-004 |
| Cluster23886 | - | - | down | down | down | down | glycerol-3-phosphate 2-O-acy | SALT00000059408_Cluster23886-001 |
| Cluster23889 | - | - | up | - | up | up | monodehydroascorbate reduct | SALT00000003233_Cluster23889-001 |
| Cluster23892 | - | - | up | - | - | - | CIPK 1 [Zea mays] | SALT00000004732_Cluster23892-001 |
| Cluster23896 | - | - | down | - | down | down | Os01g0645200 [Oryza sativa | SALT00000016138_Cluster23896-007 |
| Cluster23904 | - | down | down | - | down | - | [Hordeum vulgare vulgare] | SALT00000053957_Cluster23904-007 |
| Cluster2391 | - | up | up | up | up | up | alpha-aminoadipic semialdehy | SALT00000087128_Cluster2391-017 |
| Cluster23912 | - | - | down | - | down | down | Peroxidase 72 | SALT00000040407_Cluster23912-003 |
| Cluster23920 | - | - | - | - | up | up | ---NA--- | SALT00000051160_Cluster23920-002 |
| Cluster23921 | - | - | up | - | - | - | Os05g0453700 [Oryza sativa | SALT00000054094_Cluster23921-001 |
| Cluster23926 | - | - | - | - | - | up | probable phosphatase 2C 70 is | SALT00000001987_Cluster23926-001 |
| Cluster23929 | - | - | down | - | - | - | kinase domain containing [Ze | SALT00000015798_Cluster23929-009 |
| Cluster23934 | - | - | - | - | - | down | BURP domain-containing [Or | SALT00000049245_Cluster23934-006 |
| Cluster23948 | - | - | up | - | - | - | Os01g0585100 [Oryza sativa | SALT00000050207_Cluster23948-009 |
| Cluster23955 | - | - | - | - | down | - | phosphatase 2C 46 | SALT00000057450_Cluster23955-001 |
| Cluster23963 | - | - | up | - | up | up | UDP-glycosyltransferase 73C | SALT00000045715_Cluster23963-002 |
| Cluster23970 | - | - | up | - | - | - | pyruvate cytosolic isozyme | SALT00000006196_Cluster23970-001 |
| Cluster23979 | - | - | up | - | - | - | DETOXIFICATION 12 [Oryz | SALT00000042530_Cluster23979-001 |
| Cluster23980 | - | - | up | - | - | - | Os05g0353300 [Oryza sativa | SALT00000044613_Cluster23980-001 |
| Cluster23986 | - | up | up | - | up | - | stem-specific TSJT1-like | SALT00000051654_Cluster23986-001 |
| Cluster23988 | - | - | down | - | down | down | mitochondrial import receptor | SALT00000052786_Cluster23988-001 |
| Cluster23992 | - | - | up | - | - | - | probable polyamine transporte | SALT00000059113_Cluster23992-005 |
| Cluster240 | - | - | up | - | up | up | RAD21-3 [Oryza sativa Japon | SALT00000087875_Cluster240-031 |
| Cluster24006 | - | up | up | - | - | - | light-induced 1-like [Zea may | SALT00000045225_Cluster24006-001 |
| Cluster24018 | down | down | - | - | up | up | DNA-binding [Zea mays] | SALT00000018690_Cluster24018-004 |
| Cluster24023 | - | - | up | - | - | - | Os12g0635500 [Oryza sativa | SALT00000042829_Cluster24023-001 |
| Cluster24035 | - | down | down | - | - | - | Os05g0291700 [Oryza sativa | SALT00000007910_Cluster24035-001 |
| Cluster24038 | - | - | - | - | down | - | endo-1,3 1,4-beta-D-glucanas | SALT00000017446_Cluster24038-002 |
| Cluster24048 | - | - | up | - | - | - | trichome birefringence-like 24 | SALT00000055857_Cluster24048-002 |
| Cluster2405 | - | - | up | - | up | up | glutamate dehydrogenase 2 [C | SALT00000054592_Cluster2405-001 |
| Cluster24059 | - | - | down | - | - | - | Os06g0474500 [Oryza sativa | SALT00000012493_Cluster24059-002 |
| Cluster24060 | - | - | up | - | up | up | uncharacterized transporter Yl | SALT00000021778_Cluster24060-001 |
| Cluster24063 | - | - | - | - | up | - | UPF0183 At3g51130 isoform | SALT00000040018_Cluster24063-006 |
| Cluster24076 | - | - | down | - | - | - | Glycosyl hydrolase family 1 e | SALT00000008290_Cluster24076-001 |

| | | | | | | | | |
|---|---|---|---|---|---|---|---|---|
| Cluster24115 | - | - | up | - | up | - | methionine aminopeptidase 2I | SALT00000044939_Cluster24115-011 |
| Cluster24116 | - | - | up | - | - | - | G-box binding factor 1 isofor | SALT00000013619_Cluster24116-006 |
| Cluster24126 | - | - | up | - | up | up | cytochrome P450 | SALT00000001809_Cluster24126-001 |
| Cluster2413 | - | - | down | - | - | - | disease resistance zinc finger | SALT00000080193_Cluster2413-001 |
| Cluster2414 | - | - | up | - | up | up | lipoxygenase [Oryza sativa Ja | SALT00000064633_Cluster2414-002 |
| Cluster24141 | - | up | - | - | - | - | polygalacturonase precursor [ | SALT00000050941_Cluster24141-003 |
| Cluster24147 | - | - | up | - | - | - | Os01g0770200 [Oryza sativa | SALT00000000830_Cluster24147-001 |
| Cluster24151 | - | - | down | - | - | down | cytokinin-O-glucosyltransfera | SALT00000004457_Cluster24151-002 |
| Cluster24154 | - | - | - | - | up | - | hexokinase-1 [Zea mays] | SALT00000010204_Cluster24154-002 |
| Cluster24164 | - | - | down | - | - | down | Os03g0109600 [Oryza sativa | SALT00000055768_Cluster24164-001 |
| Cluster24165 | - | - | up | - | up | - | monodehydroascorbate reduct | SALT00000056174_Cluster24165-008 |
| Cluster24174 | - | - | up | - | up | up | partial [Oryza sativa Japonica | SALT00000035481_Cluster24174-004 |
| Cluster24176 | - | - | down | - | down | - | transcription factor AS1-like [ | SALT00000044237_Cluster24176-001 |
| Cluster2418 | - | down | down | - | - | - | sucrose synthase 1 [Oryza sati | SALT00000065300_Cluster2418-009 |
| Cluster24191 | - | - | down | - | - | - | TCP transcription partial [Zea | SALT00000039301_Cluster24191-001 |
| Cluster24202 | - | - | up | - | - | - | Os06g0665500 [Oryza sativa | SALT00000053740_Cluster24202-003 |
| Cluster24203 | - | - | up | - | up | up | Os05g0110000 [Oryza sativa | SALT00000054173_Cluster24203-002 |
| Cluster24204 | up | - | - | - | - | - | Os05g0316800 [Oryza sativa | SALT00000054688_Cluster24204-004 |
| Cluster24208 | - | - | down | - | down | - | purple acid phosphatase 2-like | SALT00000004432_Cluster24208-003 |
| Cluster24218 | - | - | down | - | - | - | GDSL esterase lipase At4g10 | SALT00000043446_Cluster24218-001 |
| Cluster24224 | - | - | up | up | up | up | TPA: NAC domain and pentat | SALT00000054468_Cluster24224-001 |
| Cluster24231 | - | - | up | - | up | up | helix-loop-helix DNA-binding | SALT00000060209_Cluster24231-005 |
| Cluster24235 | - | - | up | - | - | - | serine threonine- kinase NAK | SALT00000043042_Cluster24235-003 |
| Cluster24239 | up | - | up | - | up | up | cytochrome P450 89A2-like [ | SALT00000046097_Cluster24239-003 |
| Cluster24242 | - | - | down | - | down | down | structural constituent of ribos | SALT00000050267_Cluster24242-005 |
| Cluster24245 | - | - | up | - | up | up | YLS9 isoform X2 [Setaria ital | SALT00000056049_Cluster24245-001 |
| Cluster24254 | - | - | up | - | - | - | RETICULATA-RELATED ch | SALT00000036372_Cluster24254-001 |
| Cluster2427 | - | - | - | - | down | - | PREDICTED: uncharacterized | SALT00000070678_Cluster2427-001 |
| Cluster24276 | - | - | down | - | - | - | beta3-glucuronyltransferase [ | SALT00000052916_Cluster24276-004 |
| Cluster24280 | - | - | up | - | - | up | one cut domain family membe | SALT00000058934_Cluster24280-002 |
| Cluster24301 | - | - | up | - | up | up | glucosyltransferase IS5a-like | SALT00000035907_Cluster24301-005 |
| Cluster24322 | - | - | down | - | down | down | 60S ribosomal L3 [Zea mays] | SALT00000009019_Cluster24322-005 |
| Cluster24341 | - | - | down | - | down | down | substrate binding domain cont | SALT00000009458_Cluster24341-005 |
| Cluster24352 | up | up | up | - | - | - | Os01g0846300 [Oryza sativa | SALT00000045890_Cluster24352-003 |
| Cluster24357 | - | - | up | - | - | - | Os06g0147100 [Oryza sativa | SALT00000047393_Cluster24357-001 |
| Cluster24363 | - | - | - | - | down | - | [Hordeum vulgare vulgare] | SALT00000053675_Cluster24363-001 |
| Cluster24369 | - | - | - | - | down | - | D-cysteine desulfhydrase mit | SALT00000019564_Cluster24369-002 |
| Cluster24378 | - | - | up | - | - | - | Os01g0715900 [Oryza sativa | SALT00000061834_Cluster24378-001 |
| Cluster24379 | - | - | down | - | - | down | cysteine synthase [Oryza sativ | SALT00000065071_Cluster24379-002 |
| Cluster24390 | - | - | down | - | - | - | fe-S cluster assembly factor cl | SALT00000048801_Cluster24390-002 |
| Cluster24393 | - | - | - | - | down | - | PREDICTED: uncharacterized | SALT00000055826_Cluster24393-003 |
| Cluster2440 | - | - | down | - | - | - | elongation factor G- chloropla | SALT00000069995_Cluster2440-022 |
| Cluster24404 | - | - | up | - | up | up | Os03g0327100 [Oryza sativa | SALT00000055154_Cluster24404-001 |
| Cluster24424 | - | - | up | - | up | up | probable WRKY transcription | SALT00000004600_Cluster24424-001 |
| Cluster24434 | - | - | up | - | up | up | heptahelical transmembrane A | SALT00000037645_Cluster24434-001 |
| Cluster24436 | - | - | - | - | - | up | glycosyltransferase [Zea mays | SALT00000050367_Cluster24436-002 |
| Cluster24441 | - | up | up | - | up | up | [Hordeum vulgare vulgare] | SALT00000056808_Cluster24441-006 |
| Cluster24468 | - | - | down | - | down | down | partial [Oryza sativa Japonica | SALT00000050736_Cluster24468-001 |
| Cluster2448 | - | - | up | - | up | - | sodium-coupled neutral amin | SALT00000023382_Cluster2448-009 |
| Cluster24483 | down | - | - | - | up | up | Os03g0823400 [Oryza sativa | SALT00000041140_Cluster24483-002 |
| Cluster24485 | - | - | down | - | down | - | myb family transcription fact | SALT00000048446_Cluster24485-005 |
| Cluster2450 | - | - | up | - | - | - | [Hordeum vulgare vulgare] | SALT00000089711_Cluster2450-001 |
| Cluster2451 | - | - | down | - | down | down | ---NA--- | SALT00000063872_Cluster2451-001 |
| Cluster24512 | - | - | up | - | - | - | [Hordeum vulgare vulgare] | SALT00000005230_Cluster24512-001 |
| Cluster24520 | - | - | up | - | up | up | integral membrane -like [Ory | SALT00000042948_Cluster24520-004 |
| Cluster24529 | up | up | up | - | - | - | glucan endo-1,3-beta-glucosi | SALT00000013002_Cluster24529-001 |
| Cluster2453 | - | - | down | - | - | - | 3-oxoacyl-synthase III [Zea m | SALT00000043196_Cluster2453-002 |
| Cluster24533 | - | - | down | - | down | down | choline ethanolaminephospho | SALT00000062602_Cluster24533-010 |

| | | | | | | | | |
|---|---|---|---|---|---|---|---|---|
| Cluster24537 | - | - | - | - | down | - | cytochrome P450 superfamily | SALT00000041883_Cluster24537-003 |
| Cluster24542 | - | up | up | - | - | - | phosphatase 2C [Oryza sativa | SALT00000084697_Cluster24542-004 |
| Cluster24544 | - | - | up | - | up | up | pepsin A [Zea mays] | SALT00000014590_Cluster24544-001 |
| Cluster24555 | - | - | up | - | - | - | Senescence-associated expres | SALT00000050045_Cluster24555-002 |
| Cluster24566 | - | - | up | - | - | - | HSF transcription partial [Zea | SALT00000042807_Cluster24566-002 |
| Cluster24577 | - | - | up | - | - | - | REVEILLE 6-like [Setaria ita | SALT00000017197_Cluster24577-005 |
| Cluster24582 | - | - | up | - | up | up | 14-3-3 GF14-D | SALT00000038036_Cluster24582-001 |
| Cluster24595 | - | - | down | - | - | - | Os02g0701900 [Oryza sativa | SALT00000063775_Cluster24595-002 |
| Cluster246 | - | - | down | - | - | - | CLIP-associated -like [Setaria | SALT00000078982_Cluster246-007 |
| Cluster24608 | - | - | - | - | up | - | lanC GCL2 isoform X2 | SALT00000046909_Cluster24608-003 |
| Cluster24612 | - | - | down | - | down | - | flavonoid 3 -hydroxylase | SALT00000057181_Cluster24612-004 |
| Cluster24616 | - | - | - | - | down | - | anthocyanidin 3-O-glucosyltra | SALT00000003407_Cluster24616-002 |
| Cluster24617 | - | - | down | - | down | down | photosystem II K (chloroplast | SALT00000005542_Cluster24617-001 |
| Cluster24627 | - | down | down | - | down | - | Actin- expressed [Oryza sativa | SALT00000047983_Cluster24627-012 |
| Cluster2463 | - | - | up | - | - | - | probable CCR4-associated fac | SALT00000091889_Cluster2463-002 |
| Cluster24639 | - | - | down | - | down | down | [Hordeum vulgare vulgare] | SALT00000040798_Cluster24639-007 |
| Cluster24640 | - | - | down | - | - | - | B3 DNA binding domain fam | SALT00000044215_Cluster24640-002 |
| Cluster24650 | - | - | up | - | up | up | flavonoid 3 -monooxygenase- | SALT00000004724_Cluster24650-001 |
| Cluster24675 | - | up | up | - | - | - | glutathione S-transferase GST | SALT00000014092_Cluster24675-001 |
| Cluster24681 | - | - | down | - | - | - | [Hordeum vulgare vulgare] | SALT00000041681_Cluster24681-003 |
| Cluster24691 | - | - | down | - | - | - | IQ-DOMAIN 1-like isoform X | SALT00000059027_Cluster24691-007 |
| Cluster24699 | - | - | up | - | up | - | Os01g0837800 [Oryza sativa | SALT00000037274_Cluster24699-006 |
| Cluster247 | - | - | - | - | up | - | lectin receptor kinase [Oryza s | SALT00000066675_Cluster247-009 |
| Cluster24703 | down | down | down | - | down | - | 10-deacetylbaccatin III 10-O-a | SALT00000045664_Cluster24703-001 |
| Cluster24708 | - | - | down | - | down | - | DEHYDRATION-INDUCED | SALT00000057688_Cluster24708-001 |
| Cluster24715 | - | - | up | - | - | - | adenine guanine permease AZ | SALT00000015392_Cluster24715-001 |
| Cluster24725 | - | - | up | - | up | up | probable beta-D-xylosidase 2 | SALT00000048007_Cluster24725-001 |
| Cluster24726 | - | - | down | - | - | - | transcription factor HBP-1a-li | SALT00000048552_Cluster24726-002 |
| Cluster24729 | - | - | up | - | up | up | branched-chain-amino-acid ar | SALT00000052671_Cluster24729-002 |
| Cluster24737 | - | up | up | - | up | up | branched-chain-amino-acid ar | SALT00000014056_Cluster24737-005 |
| Cluster24740 | - | - | down | - | - | - | myosin-binding 7 | SALT00000045119_Cluster24740-001 |
| Cluster2476 | - | - | down | - | down | down | Os05g0408900 [Oryza sativa | SALT00000076045_Cluster2476-016 |
| Cluster24765 | - | - | - | - | - | up | KID-containing [Zea mays] | SALT00000044036_Cluster24765-001 |
| Cluster24773 | up | up | - | - | - | down | glutamate decarboxylase [Ory | SALT00000003028_Cluster24773-001 |
| Cluster24778 | - | - | down | - | down | - | Tubulin beta-7 expressed [Ory | SALT00000016866_Cluster24778-003 |
| Cluster24783 | - | - | up | - | up | - | uncharacterized protein LOC1 | SALT00000042762_Cluster24783-004 |
| Cluster24787 | - | - | up | - | up | - | glyceraldehyde-3-phosphate d | SALT00000051188_Cluster24787-002 |
| Cluster24795 | - | - | down | - | - | - | Os10g0431900 [Oryza sativa | SALT00000018186_Cluster24795-004 |
| Cluster24805 | up | - | - | - | down | - | Os03g0838900 [Oryza sativa | SALT00000049875_Cluster24805-001 |
| Cluster2482 | - | - | - | - | up | - | Os04g0352400 [Oryza sativa | SALT00000051790_Cluster2482-011 |
| Cluster24833 | up | up | up | - | up | up | cytochrome P450 89A2-like [ | SALT00000055925_Cluster24833-001 |
| Cluster24843 | - | - | - | - | - | up | aspartate aminotransferase [O | SALT00000037063_Cluster24843-002 |
| Cluster24844 | - | up | up | - | up | up | GRAB2 [Zea mays] | SALT00000040743_Cluster24844-005 |
| Cluster24848 | - | up | up | up | up | - | 3-ketoacyl- thiolase peroxison | SALT00000057825_Cluster24848-013 |
| Cluster24851 | - | - | up | - | up | up | Os04g0448300 [Oryza sativa | SALT00000016992_Cluster24851-004 |
| Cluster24869 | - | - | up | - | up | up | glycerol-3-phosphate acyltran | SALT00000005134_Cluster24869-001 |
| Cluster24885 | - | - | up | - | - | - | zinc finger GIS2-like [Setaria | SALT00000043392_Cluster24885-001 |
| Cluster24920 | - | - | - | - | down | - | heparan-alpha-glucosaminide | SALT00000011761_Cluster24920-001 |
| Cluster24934 | up | - | - | down | - | - | Os01g0104400 [Oryza sativa | SALT00000009634_Cluster24934-001 |
| Cluster24935 | - | up | up | - | up | - | grancalcin [Zea mays] | SALT00000011792_Cluster24935-001 |
| Cluster24950 | up | up | up | - | up | up | Os03g0717900 [Oryza sativa | SALT00000008577_Cluster24950-001 |
| Cluster24957 | - | - | up | - | up | - | ---NA--- | SALT00000039580_Cluster24957-001 |
| Cluster24962 | - | - | up | - | up | up | tryptophan synthase alpha [Ze | SALT00000047420_Cluster24962-001 |
| Cluster24964 | - | - | down | - | down | down | ATP synthase subunit chlorop | SALT00000054300_Cluster24964-006 |
| Cluster24970 | - | - | up | - | up | up | glucan endo-1,3-beta-glucosic | SALT00000002609_Cluster24970-001 |
| Cluster24971 | - | up | up | - | up | - | NAC transcription factor 29 | SALT00000008568_Cluster24971-001 |
| Cluster24989 | - | - | down | - | down | - | Os01g0125600 [Oryza sativa | SALT00000012558_Cluster24989-001 |
| Cluster2500 | - | - | up | - | - | - | transcription factor GTE9-like | SALT00000039327_Cluster2500-002 |

| | | | | | | | | |
|---|---|---|---|---|---|---|---|---|
| Cluster25001 | - | - | up | - | up | up | CBL-interacting kinase 29 | SALT00000004344_Cluster25001-002 |
| Cluster25017 | - | - | up | - | up | up | Os05g0491200 [Oryza sativa | SALT00000057059_Cluster25017-002 |
| Cluster25020 | - | - | - | - | up | up | anthocyanidin 5,3-O-glucosyl | SALT00000011954_Cluster25020-003 |
| Cluster2504 | - | - | down | - | down | down | GPI-anchored [Setaria italica] | SALT00000032870_Cluster2504-001 |
| Cluster25041 | - | - | - | - | down | - | 60S ribosomal L30 [Oryza sat | SALT00000009380_Cluster25041-001 |
| Cluster25042 | - | - | - | - | down | - | cytochrome P450 734A6-like | SALT00000009689_Cluster25042-001 |
| Cluster25046 | - | - | down | - | - | - | Os01g0799900 [Oryza sativa | SALT00000025133_Cluster25046-003 |
| Cluster25050 | - | - | - | - | down | - | Os07g0529700 [Oryza sativa | SALT00000041891_Cluster25050-001 |
| Cluster25052 | - | - | down | - | down | down | Os01g0721900 [Oryza sativa | SALT00000047326_Cluster25052-006 |
| Cluster25072 | - | - | up | - | up | up | translation factor SUI1 [Zea m | SALT00000053858_Cluster25072-001 |
| Cluster25097 | - | - | down | - | - | - | zinc finger CCCH domain-cor | SALT00000011803_Cluster25097-001 |
| Cluster251 | - | - | up | - | - | - | ATP-dependent 6-phosphofru | SALT00000070282_Cluster251-002 |
| Cluster25101 | - | - | up | - | - | - | Os05g0556000 [Oryza sativa | SALT00000036416_Cluster25101-001 |
| Cluster25106 | - | - | down | - | down | down | translation initiation factor IF- | SALT00000042617_Cluster25106-001 |
| Cluster2512 | - | - | down | - | down | down | leucine-rich repeat receptor ki | SALT00000071004_Cluster2512-003 |
| Cluster25123 | - | - | - | - | down | - | [Hordeum vulgare vulgare] | SALT00000014031_Cluster25123-003 |
| Cluster25134 | - | - | up | - | - | - | Trihelix transcription partial [ | SALT00000051684_Cluster25134-006 |
| Cluster25137 | - | - | up | - | up | up | Os02g0216300 [Oryza sativa | SALT00000056996_Cluster25137-005 |
| Cluster25153 | - | - | up | - | up | up | hypothetical protein SORBI_( | SALT00000039548_Cluster25153-004 |
| Cluster25161 | - | - | up | - | up | up | pentatricopeptide repeat-conta | SALT00000058781_Cluster25161-001 |
| Cluster25180 | - | - | up | - | - | up | serine threonine- kinase NAK | SALT00000009221_Cluster25180-001 |
| Cluster25184 | - | - | - | - | down | - | BAG family molecular chaper | SALT00000048556_Cluster25184-007 |
| Cluster2519 | - | - | down | - | down | down | nucleolar GTP-binding 1 [Ory | SALT00000084043_Cluster2519-022 |
| Cluster25191 | - | - | up | - | up | - | amino acid transporter ANT1 | SALT00000000898_Cluster25191-001 |
| Cluster25205 | - | - | up | - | - | - | Os04g0614000 [Oryza sativa | SALT00000048331_Cluster25205-001 |
| Cluster25220 | - | - | up | - | - | - | eukaryotic translation initiatio | SALT00000037394_Cluster25220-002 |
| Cluster25222 | - | - | up | - | - | - | glycosyltransferase [Zea mays | SALT00000038326_Cluster25222-002 |
| Cluster25226 | - | - | up | - | up | up | Os02g0767200 [Oryza sativa | SALT00000040287_Cluster25226-001 |
| Cluster25232 | - | - | up | - | - | - | 10-deacetylbaccatin III 10-O- | SALT00000056273_Cluster25232-001 |
| Cluster25238 | - | - | down | - | - | - | orphans transcription partial [ | SALT00000008185_Cluster25238-002 |
| Cluster25246 | - | up | up | - | - | - | Os01g0853000 [Oryza sativa | SALT00000055905_Cluster25246-001 |
| Cluster2526 | - | - | up | - | up | - | Os01g0819900 [Oryza sativa | SALT00000079584_Cluster2526-001 |
| Cluster25262 | - | - | down | - | - | - | ATP binding | SALT00000045764_Cluster25262-004 |
| Cluster253 | - | - | down | - | down | down | carbonic anhydrase [Zea mays | SALT00000061686_Cluster253-011 |
| Cluster25302 | down | down | down | - | - | - | fatty acid partial [Zea mays] | SALT00000054571_Cluster25302-003 |
| Cluster25323 | - | - | - | - | up | - | Os05g0113300 [Oryza sativa | SALT00000041766_Cluster25323-001 |
| Cluster25327 | - | - | down | - | - | - | la 1 [Oryza sativa Japonica Gr | SALT00000000731_Cluster25327-001 |
| Cluster2533 | - | up | up | up | up | - | disease resistance RPM1 [Ory | SALT00000083765_Cluster2533-001 |
| Cluster25330 | up | - | - | - | down | - | Os04g0115500 [Oryza sativa | SALT00000013609_Cluster25330-001 |
| Cluster25365 | up | up | up | - | - | - | oryzain gamma chain partial | SALT00000055808_Cluster25365-010 |
| Cluster25366 | - | - | up | - | - | - | limonoid UDP-glucosyltransf | SALT00000057417_Cluster25366-002 |
| Cluster25372 | - | - | up | - | up | - | PREDICTED: hypothetical pr | SALT00000009143_Cluster25372-001 |
| Cluster25380 | - | - | up | - | - | - | DNA binding [Zea mays] | SALT00000039546_Cluster25380-003 |
| Cluster25385 | - | - | up | - | up | up | F-box PP2-A13 [Oryza sativa | SALT00000066122_Cluster25385-001 |
| Cluster25386 | - | - | up | - | - | up | [Hordeum vulgare vulgare] | SALT00000000186_Cluster25386-001 |
| Cluster25389 | - | - | - | - | up | up | hypothetical protein SORBID | SALT00000011303_Cluster25389-002 |
| Cluster25394 | - | up | up | - | - | - | methyl-binding domain MBD | SALT00000040399_Cluster25394-003 |
| Cluster25401 | - | - | up | - | - | up | [Hordeum vulgare vulgare] | SALT00000050689_Cluster25401-004 |
| Cluster25403 | - | - | up | - | up | up | CASP 4U1 | SALT00000052081_Cluster25403-003 |
| Cluster25419 | - | - | - | - | down | - | tetratricopeptide repeat 38 [Se | SALT00000051487_Cluster25419-001 |
| Cluster25427 | - | - | - | - | - | down | PREDICTED: uncharacterize | SALT00000015518_Cluster25427-001 |
| Cluster25443 | - | - | up | - | up | up | phosphoserine aminotransfera | SALT00000000268_Cluster25443-001 |
| Cluster25448 | - | - | - | - | up | - | pyruvate dehydrogenase E1 c | SALT00000017777_Cluster25448-008 |
| Cluster25456 | - | - | up | - | up | up | laccase family [Zea mays] | SALT00000049170_Cluster25456-001 |
| Cluster25466 | - | - | down | - | - | - | anthocyanidin 3-O-glucosyltr | SALT00000009036_Cluster25466-001 |
| Cluster25472 | - | - | - | - | up | - | [Hordeum vulgare vulgare] | SALT00000037375_Cluster25472-006 |
| Cluster25473 | up | up | up | - | - | - | hydroquinone glucosyltransfe | SALT00000038678_Cluster25473-004 |
| Cluster25479 | - | - | up | - | up | up | GRAB2 [Zea mays] | SALT00000047958_Cluster25479-006 |

| | | | | | | | | |
|---|---|---|---|---|---|---|---|---|
| Cluster25493 | - | - | up | - | up | - | WRKY53 - superfamily of TF | SALT00000013463_Cluster25493-002 |
| Cluster25506 | - | - | - | - | up | up | cathepsin B-like [Oryza brach | SALT00000053863_Cluster25506-001 |
| Cluster25507 | - | - | up | - | up | up | Os02g0739400 [Oryza sativa | SALT00000054384_Cluster25507-014 |
| Cluster25512 | - | - | down | - | down | down | Os04g0620700 [Oryza sativa | SALT00000001826_Cluster25512-003 |
| Cluster25518 | - | - | - | - | up | up | mitochondrial RNA-splicing M | SALT00000039992_Cluster25518-004 |
| Cluster2553 | - | - | - | - | - | down | Pc B | SALT00000091539_Cluster2553-012 |
| Cluster25531 | - | - | down | - | down | down | Os08g0532200 [Oryza sativa | SALT00000005959_Cluster25531-001 |
| Cluster25532 | up | up | up | - | - | - | serine decarboxylase [Oryza s | SALT00000011179_Cluster25532-002 |
| Cluster25551 | - | - | up | - | - | - | Os05g0456300 [Oryza sativa | SALT00000057919_Cluster25551-007 |
| Cluster25552 | - | - | up | - | up | up | pnFL-2 [Zea mays] | SALT00000058345_Cluster25552-005 |
| Cluster25557 | - | - | up | - | up | up | senescence-associated DH [Ze | SALT00000008653_Cluster25557-002 |
| Cluster25573 | - | - | up | - | - | - | maize Em binding -1a [Zea m | SALT00000054618_Cluster25573-003 |
| Cluster25577 | - | down | down | down | down | down | phospholipase A1- chloroplast | SALT00000004584_Cluster25577-001 |
| Cluster2559 | - | - | down | - | - | - | GDP-L-galactose phosphoryla | SALT00000088131_Cluster2559-001 |
| Cluster256 | - | - | up | - | up | up | Os01g0187400 [Oryza sativa | SALT00000093337_Cluster256-008 |
| Cluster25602 | - | down | down | - | - | - | [Oryza sativa Japonica Group] | SALT00000053483_Cluster25602-003 |
| Cluster25612 | - | - | down | - | down | down | AUX IAA transcription partia | SALT00000055553_Cluster25612-003 |
| Cluster25625 | - | - | up | - | up | - | Os02g0118200 [Oryza sativa | SALT00000011387_Cluster25625-002 |
| Cluster25627 | - | - | - | - | down | - | partial [Hordeum vulgare vulg | SALT00000014293_Cluster25627-002 |
| Cluster25628 | - | - | - | - | down | - | transferase [Zea mays] | SALT00000019099_Cluster25628-004 |
| Cluster25634 | - | - | down | - | - | - | partial [Hordeum vulgare vulg | SALT00000044296_Cluster25634-001 |
| Cluster25641 | - | - | down | - | - | - | glyceraldehyde-3-phosphate d | SALT00000059293_Cluster25641-001 |
| Cluster25643 | - | down | down | - | down | down | ATP synthase delta chain [Zea | SALT00000013100_Cluster25643-001 |
| Cluster25644 | - | up | up | - | up | up | Os01g0295700 [Oryza sativa | SALT00000008952_Cluster25644-006 |
| Cluster25645 | - | - | down | - | - | down | aspartic ase nepenthesin-1 pre | SALT00000019873_Cluster25645-002 |
| Cluster25649 | - | - | down | - | down | down | [Micromonas commoda] | SALT00000049582_Cluster25649-001 |
| Cluster2566 | - | - | - | - | down | - | heat shock 70 | SALT00000058900_Cluster2566-001 |
| Cluster25679 | - | - | down | - | - | - | salt tolerance expressed [Oryz | SALT00000045214_Cluster25679-004 |
| Cluster257 | - | - | down | - | down | - | ABC transporter G family me | SALT00000082615_Cluster257-001 |
| Cluster25700 | - | - | down | - | - | - | delta-7-sterol-C5 [Zea mays] | SALT00000009510_Cluster25700-001 |
| Cluster25712 | - | - | up | - | up | up | probable xyloglucan endotran | SALT00000058518_Cluster25712-002 |
| Cluster25717 | - | - | - | - | up | - | TPA: RING zinc finger domai | SALT00000017993_Cluster25717-001 |
| Cluster25718 | - | - | down | - | - | down | structural molecule [Zea mays | SALT00000019558_Cluster25718-001 |
| Cluster25725 | - | down | down | down | down | - | Os03g0661300 [Oryza sativa | SALT00000040249_Cluster25725-009 |
| Cluster25727 | - | - | - | - | - | down | peroxidase 2-like [Setaria itali | SALT00000043183_Cluster25727-002 |
| Cluster25729 | - | - | - | - | - | up | leucine-rich repeat receptor-lil | SALT00000044762_Cluster25729-003 |
| Cluster2574 | - | - | - | - | - | down | factor of DNA methylation 1-l | SALT00000073254_Cluster2574-007 |
| Cluster25744 | - | - | up | - | up | up | hexokinase-1 [Zea mays] | SALT00000055700_Cluster25744-003 |
| Cluster25762 | up | - | - | down | - | - | probable phosphatase 2C 34 [ | SALT00000014369_Cluster25762-002 |
| Cluster25763 | - | - | - | - | up | - | secondary cell wall-related gly | SALT00000019016_Cluster25763-005 |
| Cluster25768 | - | - | down | - | down | - | acyltransferase [Zea mays] | SALT00000042128_Cluster25768-002 |
| Cluster25778 | - | - | up | - | up | - | AT-hook motif nuclear-localiz | SALT00000000665_Cluster25778-001 |
| Cluster25794 | - | down | down | - | - | - | plastidic glutamine synthetase | SALT00000056890_Cluster25794-003 |
| Cluster25805 | - | - | - | - | up | - | seed maturation expressed [Or | SALT00000022187_Cluster25805-002 |
| Cluster25828 | - | - | down | - | - | - | kinase-like [Oryza sativa Japo | SALT00000009319_Cluster25828-002 |
| Cluster25833 | - | down | down | - | down | - | actin-1 [Zea mays] | SALT00000041614_Cluster25833-008 |
| Cluster25834 | - | - | up | - | up | up | WAKY 1 | SALT00000043407_Cluster25834-002 |
| Cluster25843 | - | - | - | - | - | down | IQ calmodulin-binding motif | SALT00000043164_Cluster25843-001 |
| Cluster25850 | - | up | up | - | up | up | Os05g0134200 [Oryza sativa | SALT00000052533_Cluster25850-007 |
| Cluster25851 | - | - | down | - | - | - | 60S ribosomal L12-1 [Setaria | SALT00000055397_Cluster25851-001 |
| Cluster25855 | down | - | - | - | up | up | proline oxidase [Zea mays] | SALT00000000754_Cluster25855-001 |
| Cluster25882 | - | - | down | - | - | down | chitinase-3 1 | SALT00000052280_Cluster25882-001 |
| Cluster2590 | - | - | up | - | up | - | Os04g0482800 [Oryza sativa | SALT00000090448_Cluster2590-010 |
| Cluster25908 | - | - | up | - | up | up | cytochrome P450 superfamily | SALT00000007210_Cluster25908-002 |
| Cluster25911 | - | - | up | - | up | up | AMSH-like ubiquitin thioeste | SALT00000037495_Cluster25911-001 |
| Cluster25912 | - | - | up | - | - | - | Os02g0735100 [Oryza sativa | SALT00000039359_Cluster25912-002 |
| Cluster25914 | - | down | down | - | - | - | basic leucine zipper 2-like | SALT00000042031_Cluster25914-001 |
| Cluster25958 | - | - | down | - | - | - | Os12g0194900 [Oryza sativa | SALT00000039871_Cluster25958-002 |

| | | | | | | | | |
|---|---|---|---|---|---|---|---|---|
| Cluster25962 | - | - | up | - | up | up | Os08g0485400 [Oryza sativa | SALT00000043775_Cluster25962-001 |
| Cluster25974 | - | - | down | - | - | - | proliferation-associated 2G4 [ | SALT00000006269_Cluster25974-003 |
| Cluster25978 | - | - | down | - | - | - | Phosphoribulokinase family [] | SALT00000016829_Cluster25978-002 |
| Cluster25986 | - | - | - | - | down | - | aspartic ase nepenthesin-2 pre | SALT00000000905_Cluster25986-001 |
| Cluster25989 | - | - | down | - | down | down | 60S ribosomal L3 [Zea mays] | SALT00000008997_Cluster25989-002 |
| Cluster2599 | - | - | down | - | - | - | nuclear pore complex NUP20. | SALT00000093424_Cluster2599-001 |
| Cluster25994 | - | - | down | - | down | - | ---NA--- | SALT00000015307_Cluster25994-001 |
| Cluster25995 | - | - | - | - | up | - | ethylene response element bin | SALT00000016965_Cluster25995-003 |
| Cluster26014 | - | - | down | - | - | - | COBW domain-containing 1-] | SALT00000015275_Cluster26014-004 |
| Cluster26015 | down | down | down | - | down | - | Os03g0100200 [Oryza sativa | SALT00000015482_Cluster26015-001 |
| Cluster2603 | - | down | - | - | - | - | probable inactive poly [ADP-1 | SALT00000067889_Cluster2603-001 |
| Cluster26031 | - | - | - | down | - | - | anthocyanidin 3-O-glucosyltra | SALT00000044556_Cluster26031-001 |
| Cluster26042 | up | up | up | - | - | - | dnaJ domain containing [Zea 1 | SALT00000041781_Cluster26042-008 |
| Cluster26046 | - | - | up | - | up | up | serine threonine- phosphatase | SALT00000053468_Cluster26046-008 |
| Cluster26047 | up | up | up | - | up | - | UDP-glycosyltransferase 73C | SALT00000054014_Cluster26047-001 |
| Cluster26053 | - | - | down | - | down | - | acyltransferase [Zea mays] | SALT00000016542_Cluster26053-004 |
| Cluster26058 | - | - | down | - | down | - | chalcone synthase [Zea mays] | SALT00000043736_Cluster26058-003 |
| Cluster26062 | - | up | up | - | up | - | neurofilament heavy polypept | SALT00000052999_Cluster26062-003 |
| Cluster26071 | - | - | up | - | up | - | zinc finger CONSTANS-LIKI | SALT00000006578_Cluster26071-004 |
| Cluster2608 | - | - | - | - | down | - | division [Zea mays] | SALT00000087694_Cluster2608-006 |
| Cluster26080 | up | up | - | - | down | - | transcription factor DIVARIC. | SALT00000056058_Cluster26080-003 |
| Cluster26091 | - | down | down | - | - | - | Os04g0644600 [Oryza sativa | SALT00000048517_Cluster26091-002 |
| Cluster26108 | - | - | up | - | up | up | agmatine coumaroyltransferas | SALT00000007058_Cluster26108-001 |
| Cluster26119 | - | - | up | - | - | - | UDP-glycosyltransferase 82A | SALT00000052052_Cluster26119-001 |
| Cluster26123 | - | - | up | - | - | up | cytochrome P450 94A1-like | SALT00000055794_Cluster26123-001 |
| Cluster26124 | - | - | down | - | - | - | ATP-citrate synthase alpha ch | SALT00000066984_Cluster26124-021 |
| Cluster26128 | - | - | up | - | up | up | Os07g0170100 [Oryza sativa | SALT00000008638_Cluster26128-001 |
| Cluster26130 | - | - | up | - | up | up | Os04g0480700 [Oryza sativa | SALT00000016582_Cluster26130-001 |
| Cluster26131 | - | - | - | - | up | - | 7-deoxyloganetin glucosyltrar | SALT00000018824_Cluster26131-001 |
| Cluster26148 | - | up | up | - | up | - | hypothetical protein GLYMA_ | SALT00000018179_Cluster26148-002 |
| Cluster26155 | - | down | down | - | - | - | Os03g0321300 [Oryza sativa | SALT00000002034_Cluster26155-002 |
| Cluster26156 | - | - | up | - | up | up | peroxisomal carrier [Zea mays | SALT00000013173_Cluster26156-003 |
| Cluster26161 | - | - | down | - | down | down | UPF0481 At3g47200-like | SALT00000053305_Cluster26161-001 |
| Cluster2618 | - | - | - | - | up | - | PREDICTED: uncharacterized | SALT00000092094_Cluster2618-003 |
| Cluster26180 | - | - | up | - | up | up | Os05g0439400 [Oryza sativa | SALT00000053181_Cluster26180-002 |
| Cluster26182 | - | - | down | - | down | - | plasminogen activator inhibito | SALT00000054425_Cluster26182-013 |
| Cluster26188 | up | up | up | - | up | up | anthranilate N-benzoyltransfe | SALT00000013574_Cluster26188-002 |
| Cluster26212 | - | - | - | - | - | up | glucan endo-1,3-beta-glucosic | SALT00000040039_Cluster26212-001 |
| Cluster26213 | - | - | down | down | down | - | 19 [Oryza sativa Japonica Grc | SALT00000041173_Cluster26213-002 |
| Cluster26233 | - | - | up | - | - | - | iron-sulfur cluster assembly 1 | SALT00000036195_Cluster26233-002 |
| Cluster26235 | - | - | down | - | down | down | gibberellin-regulated 1 precur | SALT00000041496_Cluster26235-001 |
| Cluster26245 | - | - | down | - | - | - | glycine cleavage system H mi | SALT00000066964_Cluster26245-001 |
| Cluster26249 | - | - | up | - | up | up | Os03g0227900 [Oryza sativa | SALT00000011749_Cluster26249-001 |
| Cluster26259 | - | - | up | - | up | - | TCP-domain [Zea mays] | SALT00000040211_Cluster26259-004 |
| Cluster26269 | - | - | - | - | - | down | pyruvate dehydrogenase E1 co | SALT00000054821_Cluster26269-007 |
| Cluster26271 | - | - | up | - | - | up | prenylated Rab receptor 2 [Ze | SALT00000055810_Cluster26271-001 |
| Cluster26272 | - | - | down | - | - | - | [Hordeum vulgare vulgare] | SALT00000056610_Cluster26272-004 |
| Cluster26279 | - | - | - | - | down | - | lactate dehydrogenase [Oryza | SALT00000040137_Cluster26279-002 |
| Cluster26289 | - | - | down | - | - | - | Os01g0836900 [Oryza sativa | SALT00000002554_Cluster26289-001 |
| Cluster26296 | - | - | up | - | up | - | bifunctional nuclease 2 [Setar | SALT00000037287_Cluster26296-001 |
| Cluster26302 | - | - | down | - | down | down | DETOXIFICATION 35-like [ | SALT00000052396_Cluster26302-002 |
| Cluster26309 | - | - | up | - | - | - | 12-oxophytodienoate reductas | SALT00000010435_Cluster26309-001 |
| Cluster26328 | - | - | down | - | - | - | transcription repressor MYB6 | SALT00000008191_Cluster26328-001 |
| Cluster26329 | - | - | up | - | up | - | 3-isopropylmalate dehydroger | SALT00000009730_Cluster26329-005 |
| Cluster26330 | - | down | down | - | down | down | cytochrome P450 711A1 [Ory | SALT00000010951_Cluster26330-001 |
| Cluster26335 | - | - | up | - | up | up | NAC1 [Sorghum bicolor] | SALT00000016264_Cluster26335-004 |
| Cluster26353 | - | - | down | - | down | down | chlorophyll a-b binding [Zea 1 | SALT00000004434_Cluster26353-001 |
| Cluster26358 | - | - | up | - | up | up | GTP-binding SAR1A-like [Se | SALT00000014928_Cluster26358-002 |

| | | | | | | | | |
|---|---|---|---|---|---|---|---|---|
| Cluster26367 | - | - | down | - | - | - | Ribulose bisphosphate carbox | SALT00000046827_Cluster26367-010 |
| Cluster26375 | - | - | down | - | - | down | UDP-glycosyltransferase 88B | SALT00000007339_Cluster26375-001 |
| Cluster26378 | - | - | down | - | - | - | Os01g0207600 [Oryza sativa | SALT00000044608_Cluster26378-001 |
| Cluster26391 | - | - | down | - | - | - | [Hordeum vulgare vulgare] | SALT00000060368_Cluster26391-001 |
| Cluster26393 | - | - | up | - | - | - | nudix hydrolase mitochondria | SALT00000061679_Cluster26393-001 |
| Cluster26394 | - | - | down | - | - | - | glucan endo-1,3-beta-glucosic | SALT00000001029_Cluster26394-001 |
| Cluster26399 | - | - | down | - | - | - | ring finger [Oryza sativa Japo | SALT00000006528_Cluster26399-001 |
| Cluster26418 | - | up | up | - | - | - | DEHYDRATION-INDUCED | SALT00000009216_Cluster26418-002 |
| Cluster26421 | - | - | up | - | - | up | mitochondrial phosphate trans | SALT00000016789_Cluster26421-005 |
| Cluster26428 | - | - | down | - | down | down | 60S ribosomal L11-1 [Zea ma | SALT00000046198_Cluster26428-001 |
| Cluster2643 | - | - | - | - | down | - | monosaccharide-sensing 2 | SALT00000082187_Cluster2643-027 |
| Cluster26430 | down | down | down | - | down | - | peroxidase 43-like [Setaria ita | SALT00000053907_Cluster26430-002 |
| Cluster26432 | - | - | down | - | - | - | -type [Zea mays] | SALT00000007275_Cluster26432-001 |
| Cluster26433 | - | - | up | - | up | up | B3 domain-containing Os03g( | SALT00000007824_Cluster26433-001 |
| Cluster2644 | up | up | up | - | - | - | 5-methyltetrahydropteroyltrig | SALT00000070510_Cluster2644-002 |
| Cluster2647 | - | - | - | - | - | down | Os07g0406300 [Oryza sativa | SALT00000070061_Cluster2647-010 |
| Cluster26481 | - | - | up | - | up | up | [Hordeum vulgare vulgare] | SALT00000044116_Cluster26481-002 |
| Cluster26486 | - | up | up | - | up | - | Os06g0147000 [Oryza sativa | SALT00000050066_Cluster26486-001 |
| Cluster26491 | - | - | down | - | down | down | myb-related Zm38-like | SALT00000058849_Cluster26491-001 |
| Cluster26492 | - | up | up | - | up | up | Os09g0526600 [Oryza sativa | SALT00000039736_Cluster26492-006 |
| Cluster26521 | - | - | - | - | down | - | UDP-glucose: flavonoid 7-O-; | SALT00000061795_Cluster26521-001 |
| Cluster26522 | - | up | up | - | up | up | homogentisate 1,2-dioxygenas | SALT00000002656_Cluster26522-001 |
| Cluster26524 | - | - | up | - | up | up | beta-1,3-galactosyltransferase | SALT00000064715_Cluster26524-005 |
| Cluster26567 | - | - | up | - | - | - | CBL-interacting serine threon | SALT00000018225_Cluster26567-001 |
| Cluster2657 | - | - | up | - | - | - | receptor kinase HSL1 | SALT00000082564_Cluster2657-017 |
| Cluster26577 | - | up | up | up | up | - | Os01g0270100 [Oryza sativa | SALT00000053215_Cluster26577-001 |
| Cluster26581 | - | - | up | - | up | up | Os07g0627700 [Oryza sativa | SALT00000059404_Cluster26581-005 |
| Cluster26593 | - | - | - | - | - | down | homeodomain-like transcriptio | SALT00000038876_Cluster26593-001 |
| Cluster26614 | - | - | up | - | up | up | Os03g0822700 [Oryza sativa | SALT00000043935_Cluster26614-010 |
| Cluster2662 | - | - | down | - | - | - | DNA topoisomerase 2 | SALT00000088922_Cluster2662-007 |
| Cluster26649 | - | down | down | - | - | - | MACPF domain-containing A | SALT00000052258_Cluster26649-001 |
| Cluster26663 | - | - | down | - | down | - | uncharacterized membrane At | SALT00000048614_Cluster26663-001 |
| Cluster26665 | - | - | up | - | up | up | UDP-glycosyltransferase 73C | SALT00000051238_Cluster26665-002 |
| Cluster26685 | - | down | down | down | down | - | magnesium-chelatase subunit | SALT00000006584_Cluster26685-004 |
| Cluster26697 | - | - | up | - | - | - | fas-associated factor 1 [Zea m | SALT00000041377_Cluster26697-002 |
| Cluster26700 | - | up | - | - | - | - | 2C-type phosphatase [Zea ma | SALT00000046719_Cluster26700-002 |
| Cluster26714 | - | - | - | - | - | up | 7-deoxyloganetin glucosyltrar | SALT00000008034_Cluster26714-001 |
| Cluster26719 | - | - | - | - | down | - | Os09g0453900 [Oryza sativa | SALT00000036455_Cluster26719-001 |
| Cluster26746 | - | - | - | - | - | down | glycine-rich cell wall structura | SALT00000068262_Cluster26746-002 |
| Cluster26751 | - | - | - | - | - | down | Os05g0512200 [Oryza sativa | SALT00000009411_Cluster26751-002 |
| Cluster26764 | - | - | up | - | up | - | isovaleryl- mitochondrial | SALT00000008115_Cluster26764-003 |
| Cluster26769 | - | - | up | - | - | - | SNF1-type serine-threonine k | SALT00000053015_Cluster26769-004 |
| Cluster2678 | - | - | - | - | down | - | glutaminyl-tRNA synthetase [ | SALT00000075412_Cluster2678-019 |
| Cluster26800 | up | up | - | - | - | - | Os09g0360500 [Oryza sativa | SALT00000008430_Cluster26800-001 |
| Cluster26818 | - | up | up | - | - | - | kelch repeat containing F-box | SALT00000002525_Cluster26818-001 |
| Cluster26823 | up | up | up | - | up | - | homeobox-leucine zipper HO | SALT00000037683_Cluster26823-001 |
| Cluster26830 | - | - | - | - | - | up | Os11g0546300 [Oryza sativa | SALT00000043176_Cluster26830-001 |
| Cluster26841 | - | - | - | - | down | - | UDP-glucose:salicylic acid gl | SALT00000038135_Cluster26841-001 |
| Cluster26843 | - | - | up | - | - | - | Os07g0463400 [Oryza sativa | SALT00000040296_Cluster26843-005 |
| Cluster26853 | - | - | down | up | - | down | phospholipase A1-II 7-like [S | SALT00000061524_Cluster26853-003 |
| Cluster26860 | - | - | up | - | up | up | dihydropyrimidine dehydroge | SALT00000022074_Cluster26860-004 |
| Cluster26865 | - | down | down | - | down | down | Os03g0267300 [Oryza sativa | SALT00000048929_Cluster26865-009 |
| Cluster2687 | - | - | up | - | - | - | ubiquitin carboxyl-terminal hy | SALT00000092212_Cluster2687-001 |
| Cluster26886 | - | - | - | - | down | - | xyloglucan endotransglucosyl | SALT00000058800_Cluster26886-003 |
| Cluster26888 | - | - | - | - | - | down | NOD26 major intrinsic [Oryz | SALT00000036959_Cluster26888-005 |
| Cluster26889 | up | up | up | - | - | - | homeobox-leucine zipper HO | SALT00000002244_Cluster26889-001 |
| Cluster26913 | - | - | down | - | - | - | 50S ribosomal L1 | SALT00000052459_Cluster26913-002 |
| Cluster26924 | - | - | down | - | down | down | Os03g0817900 [Oryza sativa | SALT00000020122_Cluster26924-004 |

| | | | | | | | | |
|---|---|---|---|---|---|---|---|---|
| Cluster26928 | - | - | down | - | - | - | SHOOT1 [Oryza sativa Japon | SALT00000040069_Cluster26928-001 |
| Cluster26934 | - | - | down | - | down | down | [Hordeum vulgare vulgare] | SALT00000052011_Cluster26934-002 |
| Cluster26949 | - | - | down | - | - | - | membrane [Oryza sativa Japo | SALT00000042859_Cluster26949-004 |
| Cluster2695 | - | - | up | - | up | up | Os11g0533500 [Oryza sativa | SALT00000086784_Cluster2695-008 |
| Cluster26957 | - | up | - | - | down | down | Os03g0327600 [Oryza sativa | SALT00000060736_Cluster26957-010 |
| Cluster2696 | up | up | up | - | up | - | ethylene-responsive [Zea may | SALT00000086976_Cluster2696-001 |
| Cluster26966 | - | - | down | - | - | down | phospholipase A1-II 5 [Setaria | SALT00000018159_Cluster26966-001 |
| Cluster26983 | - | - | down | down | down | down | fasciclin-like arabinogalactan | SALT00000008845_Cluster26983-001 |
| Cluster26984 | - | - | up | - | up | up | UDP-glycosyltransferase 73C | SALT00000013646_Cluster26984-001 |
| Cluster26986 | - | - | up | - | up | up | isovaleryl- mitochondrial | SALT00000015893_Cluster26986-001 |
| Cluster2699 | - | - | down | - | - | - | Os07g0100500 [Oryza sativa | SALT00000034515_Cluster2699-002 |
| Cluster26992 | - | - | - | - | up | - | oligopeptide transporter 7 | SALT00000042963_Cluster26992-004 |
| Cluster27013 | - | up | up | up | up | up | hioredoxin 5 [Oryza sativa Ja | SALT00000035886_Cluster27013-001 |
| Cluster27037 | - | - | down | - | down | - | Os12g0615400 [Oryza sativa | SALT00000050703_Cluster27037-002 |
| Cluster2704 | - | - | down | - | - | - | ATP-dependent DNA helicase | SALT00000084902_Cluster2704-002 |
| Cluster27040 | - | - | down | - | down | down | digalactosyldiacylglycerol syr | SALT00000061412_Cluster27040-001 |
| Cluster27045 | - | - | - | - | down | - | [Hordeum vulgare vulgare] | SALT00000008679_Cluster27045-001 |
| Cluster27050 | - | - | up | - | up | up | heat shock factor 7 [Zea mays | SALT00000041297_Cluster27050-004 |
| Cluster27053 | - | - | - | - | down | - | G1-like5 [Oryza sativa Japoni | SALT00000046642_Cluster27053-001 |
| Cluster27064 | - | - | - | - | down | - | Os03g0343400 [Oryza sativa | SALT00000000942_Cluster27064-001 |
| Cluster27068 | - | - | down | - | - | - | probable apyrase 3 [Setaria ita | SALT00000012390_Cluster27068-001 |
| Cluster27090 | - | up | - | - | - | down | pyruvate dehydrogenase E1 c | SALT00000005212_Cluster27090-001 |
| Cluster27093 | - | - | up | - | up | up | Os09g0452900 [Oryza sativa | SALT00000007368_Cluster27093-001 |
| Cluster27109 | - | - | up | - | - | up | OTU domain-containing 5-A- | SALT00000060585_Cluster27109-004 |
| Cluster27128 | - | - | - | - | - | down | cinnamoyl- reductase [Zea ma | SALT00000046558_Cluster27128-003 |
| Cluster27132 | - | - | up | - | up | up | catalytic hydrolase isoform X | SALT00000039529_Cluster27132-003 |
| Cluster27141 | - | - | up | - | up | up | Os03g0207400 [Oryza sativa | SALT00000007014_Cluster27141-003 |
| Cluster27145 | - | - | up | - | up | up | [Hordeum vulgare vulgare] | SALT00000012637_Cluster27145-002 |
| Cluster27148 | - | - | up | - | up | - | PREDICTED: uncharacterize | SALT00000039883_Cluster27148-001 |
| Cluster27158 | - | - | down | - | - | - | 125 kDa kinesin-related isofo | SALT00000069299_Cluster27158-001 |
| Cluster27176 | - | - | - | - | down | - | protease Do-like chloroplastic | SALT00000019254_Cluster27176-003 |
| Cluster27179 | - | - | - | - | down | down | Retrovirus-related Pol poly LI | SALT00000057004_Cluster27179-001 |
| Cluster27188 | - | - | - | - | - | down | Os05g0113500 [Oryza sativa | SALT00000013320_Cluster27188-003 |
| Cluster27196 | - | - | down | - | - | - | 30S ribosomal S5 [Zea mays] | SALT00000041693_Cluster27196-002 |
| Cluster27203 | - | - | up | - | up | up | cytochrome b5 | SALT00000055626_Cluster27203-001 |
| Cluster27209 | - | - | down | - | - | - | zinc finger AN1 domain-conta | SALT00000006493_Cluster27209-001 |
| Cluster27215 | - | - | up | - | up | up | AF466200_3tryptophan synth | SALT00000056293_Cluster27215-010 |
| Cluster2723 | - | - | up | - | up | up | lysine-specific demethylase 5 | SALT00000088073_Cluster2723-005 |
| Cluster27239 | - | - | up | - | up | - | zinc finger A20 and AN1 dom | SALT00000007959_Cluster27239-002 |
| Cluster2724 | - | down | up | - | up | up | endoplasmin homolog | SALT00000081516_Cluster2724-047 |
| Cluster2725 | - | up | - | - | - | - | NADP-specific isocitrate deh | SALT00000053799_Cluster2725-001 |
| Cluster27252 | - | - | down | - | down | - | cell division inhibitor-like [O | SALT00000058441_Cluster27252-002 |
| Cluster2727 | - | - | - | - | down | - | AP2 EREBP transcription fac | SALT00000090940_Cluster2727-002 |
| Cluster27275 | - | - | down | - | down | down | ---NA--- | SALT00000052209_Cluster27275-005 |
| Cluster27278 | - | - | up | - | - | - | anthocyanidin 3-O-glucosyltr | SALT00000062853_Cluster27278-001 |
| Cluster27281 | - | - | up | - | up | up | type I cytoskeletal 9 [Setaria i | SALT00000001724_Cluster27281-001 |
| Cluster27285 | - | - | up | - | - | - | [Hordeum vulgare vulgare] | SALT00000010160_Cluster27285-001 |
| Cluster27286 | - | - | up | - | up | up | Transcription factor MYB108 | SALT00000010987_Cluster27286-001 |
| Cluster2729 | - | - | up | - | up | up | Os10g0501500 [Oryza sativa | SALT00000070627_Cluster2729-004 |
| Cluster27291 | - | - | - | - | down | down | Os03g0764000 [Oryza sativa | SALT00000022653_Cluster27291-001 |
| Cluster27294 | - | - | - | - | - | down | [Oryza sativa Indica Group] | SALT00000039623_Cluster27294-001 |
| Cluster27306 | - | - | down | - | - | - | multiple organellar RNA editi | SALT00000014586_Cluster27306-002 |
| Cluster27349 | - | up | - | - | - | - | phosphatidylinositol phosphat | SALT00000044435_Cluster27349-001 |
| Cluster27353 | - | up | up | - | up | up | Os04g0623500 [Oryza sativa | SALT00000018207_Cluster27353-015 |
| Cluster27359 | - | - | up | - | - | - | Os01g0667600 [Oryza sativa | SALT00000059641_Cluster27359-001 |
| Cluster27360 | - | - | up | - | up | - | Os07g0642600 [Oryza sativa | SALT00000001646_Cluster27360-001 |
| Cluster27368 | - | up | up | - | - | - | inhibitor of apoptosis [Zea ma | SALT00000023291_Cluster27368-001 |
| Cluster27372 | - | - | - | - | up | - | RING-H2 finger ATL29 isofo | SALT00000042065_Cluster27372-003 |

| | | | | | | | | |
|---|---|---|---|---|---|---|---|---|
| Cluster27385 | - | - | up | - | up | up | acetylornithine mitochondrial | SALT00000005892_Cluster27385-002 |
| Cluster27387 | - | - | down | - | - | - | 20 kDa chloroplastic-like | SALT00000010470_Cluster27387-001 |
| Cluster27388 | - | - | down | - | down | down | probable mediator of RNA po | SALT00000010998_Cluster27388-001 |
| Cluster27401 | - | - | up | - | up | - | Os08g0439600 [Oryza sativa | SALT00000056619_Cluster27401-005 |
| Cluster27409 | - | - | up | - | up | up | [Oryza sativa Japonica Group | SALT00000014564_Cluster27409-006 |
| Cluster27421 | - | - | up | - | up | up | 4-hydroxyphenylpyruvate dio | SALT00000000995_Cluster27421-001 |
| Cluster27423 | - | down | down | down | - | - | WRKY-type transcription fact | SALT00000003248_Cluster27423-001 |
| Cluster27432 | - | - | down | - | - | down | Os03g0609500 [Oryza sativa | SALT00000037920_Cluster27432-001 |
| Cluster27439 | - | - | up | - | up | up | transposon Mutator sub-class | SALT00000054863_Cluster27439-002 |
| Cluster27442 | - | - | up | - | - | - | [Hordeum vulgare vulgare] | SALT00000019887_Cluster27442-003 |
| Cluster27446 | up | - | - | down | - | - | Os04g0460300 [Oryza sativa | SALT00000000483_Cluster27446-001 |
| Cluster27460 | - | - | down | - | down | - | proline-rich 4-like | SALT00000049938_Cluster27460-002 |
| Cluster2747 | - | - | up | - | up | - | armadillo beta-catenin repeat | SALT00000091081_Cluster2747-003 |
| Cluster27484 | - | - | down | - | down | down | homeobox-leucine zipper HO | SALT00000057740_Cluster27484-001 |
| Cluster27491 | - | - | up | - | up | up | snRK SAPK family kinase [Z | SALT00000005795_Cluster27491-001 |
| Cluster27498 | - | - | up | - | up | up | RING-H2 finger ATL52 [Brac | SALT00000037152_Cluster27498-002 |
| Cluster27511 | - | - | down | - | - | - | [Hordeum vulgare vulgare] | SALT00000007474_Cluster27511-003 |
| Cluster27513 | - | - | down | - | - | - | Aspartic ase nepenthesin-1 | SALT00000014407_Cluster27513-002 |
| Cluster27516 | - | - | - | - | - | up | TPA: AP2 EREBP transcriptic | SALT00000037563_Cluster27516-001 |
| Cluster27517 | up | up | up | - | - | - | Os04g0625800 [Oryza sativa | SALT00000040543_Cluster27517-001 |
| Cluster27518 | up | - | - | - | down | - | cysteine protease 1 precursor | SALT00000040890_Cluster27518-002 |
| Cluster27524 | - | - | - | - | up | up | Os05g0110000 [Oryza sativa | SALT00000057364_Cluster27524-003 |
| Cluster27529 | up | up | down | - | down | down | glutamine synthetase root isoz | SALT00000018594_Cluster27529-002 |
| Cluster2753 | - | - | down | - | - | - | Os04g0504800 [Oryza sativa | SALT00000081664_Cluster2753-028 |
| Cluster27536 | up | up | up | - | up | up | cationic peroxidase SPC4 [Or | SALT00000048143_Cluster27536-003 |
| Cluster27550 | - | - | up | - | up | - | solute carrier family 25 memb | SALT00000017338_Cluster27550-001 |
| Cluster27567 | - | - | - | - | - | up | MYB-like transcription factor | SALT00000006768_Cluster27567-001 |
| Cluster27588 | - | - | down | - | down | down | photosystem II reaction center | SALT00000008500_Cluster27588-001 |
| Cluster27589 | - | - | down | - | down | - | 30S ribosomal chloroplastic | SALT00000010891_Cluster27589-003 |
| Cluster27592 | - | up | up | up | up | - | [Hordeum vulgare vulgare] | SALT00000017354_Cluster27592-004 |
| Cluster27611 | - | down | down | - | - | - | Os09g0323500 [Oryza sativa | SALT00000020051_Cluster27611-001 |
| Cluster27613 | - | - | down | - | - | - | binding [Zea mays] | SALT00000035891_Cluster27613-002 |
| Cluster27642 | - | - | - | - | down | down | acylphosphatase [Brachypodiu | SALT00000061018_Cluster27642-002 |
| Cluster27644 | - | - | up | - | up | up | Os05g0151300 [Oryza sativa | SALT00000005703_Cluster27644-001 |
| Cluster27645 | - | - | down | - | - | - | bHLH transcription partial [Z | SALT00000006390_Cluster27645-001 |
| Cluster27649 | - | - | - | - | down | - | elongation factor 1-alpha-like | SALT00000015371_Cluster27649-001 |
| Cluster27663 | - | - | up | - | up | up | heat shock factor 7 [Zea mays | SALT00000054536_Cluster27663-003 |
| Cluster2767 | - | - | down | - | down | down | triose phosphate phosphate tra | SALT00000050485_Cluster2767-001 |
| Cluster27681 | - | down | down | - | down | - | peptidyl-prolycis-trans isomer | SALT00000046093_Cluster27681-003 |
| Cluster27685 | - | - | up | - | up | up | Zinc finger expressed [Oryza | SALT00000057620_Cluster27685-004 |
| Cluster27693 | - | - | up | - | - | - | kinase superfamily | SALT00000013632_Cluster27693-005 |
| Cluster27698 | - | - | - | - | up | up | sulfate transporter [Zea mays] | SALT00000040610_Cluster27698-001 |
| Cluster27700 | - | - | up | - | up | up | WRKY11 - superfamily of TF | SALT00000042784_Cluster27700-002 |
| Cluster27702 | - | - | - | - | down | - | anther-specific proline-rich Al | SALT00000048821_Cluster27702-004 |
| Cluster27724 | - | - | up | - | - | - | Os02g0513000 [Oryza sativa | SALT00000055256_Cluster27724-001 |
| Cluster27727 | - | - | down | - | down | down | plasminogen activator inhibito | SALT00000059122_Cluster27727-001 |
| Cluster27730 | up | up | up | - | up | up | early nodulin 1 [Oryza sativa | SALT00000002943_Cluster27730-001 |
| Cluster27733 | - | - | up | - | up | up | PREDICTED: uncharacterize | SALT00000015253_Cluster27733-001 |
| Cluster27741 | - | - | - | - | up | - | actin-depolymerizing factor 3 | SALT00000046749_Cluster27741-001 |
| Cluster27742 | - | up | up | - | up | - | Os01g0199400 [Oryza sativa | SALT00000050143_Cluster27742-001 |
| Cluster27747 | - | - | down | - | - | - | Os08g0558300 [Oryza sativa | SALT00000009963_Cluster27747-001 |
| Cluster27750 | - | - | up | - | up | up | sugar phosphatase -like [Brac | SALT00000016057_Cluster27750-004 |
| Cluster27759 | - | - | up | up | up | - | Os11g0126900 [Oryza sativa | SALT00000044788_Cluster27759-002 |
| Cluster27790 | - | down | - | - | - | up | Os12g0184400 [Oryza sativa | SALT00000011391_Cluster27790-001 |
| Cluster2780 | - | - | down | - | down | down | hAT dimerisation domain-con | SALT00000033094_Cluster2780-001 |
| Cluster2781 | - | - | - | - | down | - | Os01g0296100 [Oryza sativa | SALT00000090397_Cluster2781-008 |
| Cluster27837 | - | down | down | - | down | down | [Oryza sativa Japonica Group | SALT00000019517_Cluster27837-002 |
| Cluster27861 | - | - | down | - | - | - | mitochondrial 2-oxoglutarate | SALT00000054762_Cluster27861-006 |

| | | | | | | | | |
|---|---|---|---|---|---|---|---|---|
| Cluster27876 | - | - | up | - | - | - | NAC transcription factor 1 | SALT00000047254_Cluster27876-001 |
| Cluster27878 | - | - | down | - | down | down | 40S ribosomal S13 [Zea mays | SALT00000052048_Cluster27878-001 |
| Cluster2788 | - | - | - | - | down | - | FAR1-RELATED SEQUENC | SALT00000085581_Cluster2788-001 |
| Cluster27899 | - | - | up | - | up | up | Os08g0529300 [Oryza sativa | SALT00000003107_Cluster27899-001 |
| Cluster27902 | - | - | down | - | - | - | Heavy metal transport detoxif | SALT00000008665_Cluster27902-001 |
| Cluster27929 | - | - | down | - | - | - | Os05g0349700 [Oryza sativa | SALT00000025096_Cluster27929-002 |
| Cluster27940 | - | - | up | - | - | - | pathogenicity partial | SALT00000054229_Cluster27940-001 |
| Cluster27943 | - | down | down | - | down | down | tetracycline resistance class H | SALT00000060840_Cluster27943-001 |
| Cluster2795 | - | - | - | - | down | down | Os05g0391500 [Oryza sativa | SALT00000082591_Cluster2795-004 |
| Cluster27954 | - | - | up | - | - | - | cytochrome P450 81D11-like | SALT00000042765_Cluster27954-001 |
| Cluster27955 | - | - | down | - | down | down | Os01g0756600 [Oryza sativa | SALT00000043514_Cluster27955-001 |
| Cluster27960 | - | - | - | - | - | down | Os06g0107100 [Oryza sativa | SALT00000053229_Cluster27960-001 |
| Cluster27963 | - | - | up | - | - | - | ras-related RABF2b | SALT00000056677_Cluster27963-001 |
| Cluster27971 | - | - | down | - | down | down | 24-methylenesterol C-methylt | SALT00000009101_Cluster27971-002 |
| Cluster27981 | - | - | down | - | down | down | Os11g0242800 [Oryza sativa | SALT00000043957_Cluster27981-001 |
| Cluster27989 | - | down | down | - | down | - | [Hordeum vulgare vulgare] | SALT00000060003_Cluster27989-002 |
| Cluster2799 | - | - | up | - | up | up | probable GPI-anchored adhes | SALT00000090219_Cluster2799-003 |
| Cluster28001 | - | - | down | - | down | down | peroxisomal (S)-2-hydroxy-ac | SALT00000050226_Cluster28001-002 |
| Cluster28010 | - | - | up | - | up | up | 2C-type phosphatase [Zea ma | SALT00000014926_Cluster28010-003 |
| Cluster28011 | - | - | up | - | up | - | [Hordeum vulgare vulgare] | SALT00000016336_Cluster28011-001 |
| Cluster28018 | - | - | up | - | - | - | hypersensitive-induced reactio | SALT00000050827_Cluster28018-002 |
| Cluster28023 | - | - | - | - | down | - | retrotransposon unclassified | SALT00000060786_Cluster28023-001 |
| Cluster28056 | - | - | - | - | - | up | Os06g0711100 [Oryza sativa | SALT00000008468_Cluster28056-001 |
| Cluster28061 | - | - | down | - | - | - | partial [Oryza sativa Japonica | SALT00000018576_Cluster28061-001 |
| Cluster28063 | - | - | - | - | up | - | Serine threonine- kinase expre | SALT00000037852_Cluster28063-001 |
| Cluster28066 | - | - | up | - | - | up | Os08g0439600 [Oryza sativa | SALT00000040406_Cluster28066-002 |
| Cluster28069 | - | - | down | - | down | - | TIC chloroplastic | SALT00000044280_Cluster28069-005 |
| Cluster28073 | up | up | up | - | up | up | [Hordeum vulgare vulgare] | SALT00000047835_Cluster28073-001 |
| Cluster28074 | - | - | up | - | up | up | ATG8-interacting 1 isoform X | SALT00000053287_Cluster28074-002 |
| Cluster2808 | - | - | up | - | up | - | Os01g0786900 [Oryza sativa | SALT00000081044_Cluster2808-001 |
| Cluster28081 | - | - | - | - | up | up | triacylglycerol lipase like [Zea | SALT00000018354_Cluster28081-001 |
| Cluster2809 | - | - | down | - | down | - | downstream neighbor of Son i | SALT00000081477_Cluster2809-004 |
| Cluster2810 | - | - | up | - | up | - | probable histidine kinase 3 [S | SALT00000091368_Cluster2810-016 |
| Cluster28101 | - | - | up | - | up | up | Os07g0563400 [Oryza sativa | SALT00000036281_Cluster28101-001 |
| Cluster28114 | - | - | up | - | - | - | ornithine decarboxylase-like [ | SALT00000041669_Cluster28114-002 |
| Cluster28118 | - | up | up | - | up | - | auxin-responsive IAA6-like [I | SALT00000049989_Cluster28118-002 |
| Cluster28120 | - | - | down | - | - | - | eukaryotic translation initiatio | SALT00000053013_Cluster28120-005 |
| Cluster28123 | - | - | up | - | - | - | Os06g0601100 [Oryza sativa | SALT00000054078_Cluster28123-002 |
| Cluster28129 | - | up | - | up | - | - | Os03g0214200 [Oryza sativa | SALT00000006048_Cluster28129-001 |
| Cluster28149 | - | - | down | - | - | - | equilibrative nucleotide transp | SALT00000051554_Cluster28149-003 |
| Cluster28162 | - | - | - | - | down | - | cytochrome P450 94B3-like | SALT00000036068_Cluster28162-001 |
| Cluster28165 | - | - | down | - | down | down | ubiquitin fusion [Zea mays] | SALT00000038281_Cluster28165-001 |
| Cluster28189 | - | - | up | - | up | up | Os06g0670300 [Oryza sativa | SALT00000052319_Cluster28189-001 |
| Cluster28197 | - | - | up | - | up | up | myb [Oryza sativa Japonica G | SALT00000017202_Cluster28197-001 |
| Cluster28198 | - | - | up | - | up | - | F-box domain containing [Zea | SALT00000046200_Cluster28198-005 |
| Cluster282 | - | up | up | - | - | - | single myb histone 6 | SALT00000041935_Cluster282-004 |
| Cluster28200 | - | - | up | - | - | - | Os01g0937100 [Oryza sativa | SALT00000039688_Cluster28200-009 |
| Cluster28206 | - | - | down | - | - | - | heat stress transcription factor | SALT00000052379_Cluster28206-001 |
| Cluster28213 | - | - | up | - | - | - | solute carrier family 35 memb | SALT00000011452_Cluster28213-001 |
| Cluster28219 | - | - | up | - | up | up | retrotransposon unclassified [ | SALT00000039734_Cluster28219-001 |
| Cluster28220 | - | - | - | - | down | - | Os07g0559400 [Oryza sativa | SALT00000046553_Cluster28220-001 |
| Cluster28223 | - | - | up | - | - | - | alpha beta fold family express | SALT00000053592_Cluster28223-004 |
| Cluster28253 | up | up | up | - | up | - | NAC domain-containing 48-li | SALT00000041801_Cluster28253-001 |
| Cluster28278 | - | - | - | - | down | down | cinnamyl alcohol dehydrogen | SALT00000060245_Cluster28278-001 |
| Cluster28279 | - | - | - | - | down | - | ocs element-binding factor 1-l | SALT00000003232_Cluster28279-001 |
| Cluster28281 | - | - | up | - | up | up | Os01g0171800 [Oryza sativa | SALT00000008028_Cluster28281-002 |
| Cluster28293 | - | - | down | - | - | - | indole-2-monooxygenase-like | SALT00000051669_Cluster28293-001 |
| Cluster28299 | - | - | up | - | up | up | formin 3 isoform X1 | SALT00000058863_Cluster28299-003 |

| | | | | | | | | |
|---|---|---|---|---|---|---|---|---|
| Cluster2830 | - | down | down | - | - | - | ATPase family AAA domain-c | SALT00000056874_Cluster2830-001 |
| Cluster28321 | - | - | - | - | - | down | Os03g0194300 [Oryza sativa | SALT00000009984_Cluster28321-001 |
| Cluster28331 | - | - | down | - | - | - | Os07g0189800 [Oryza sativa | SALT00000041085_Cluster28331-001 |
| Cluster28335 | - | - | down | - | - | - | (+)-neomenthol dehydrogenas | SALT00000049758_Cluster28335-003 |
| Cluster28346 | - | - | down | - | - | - | hypothetical protein SETIT_0 | SALT00000014957_Cluster28346-001 |
| Cluster28355 | - | - | - | - | up | - | Os03g0153900 [Oryza sativa | SALT00000049618_Cluster28355-003 |
| Cluster28358 | - | down | down | - | down | - | GATA transcription factor 2-li | SALT00000056962_Cluster28358-004 |
| Cluster28359 | - | - | down | - | - | - | partial [Oryza sativa Japonica | SALT00000058106_Cluster28359-003 |
| Cluster28365 | - | - | up | - | - | - | trans-resveratrol di-O-methylt | SALT00000008912_Cluster28365-001 |
| Cluster28376 | - | - | - | - | down | - | Os02g0728700 [Oryza sativa | SALT00000005807_Cluster28376-001 |
| Cluster28398 | - | - | up | - | - | - | Os05g0455600 [Oryza sativa | SALT00000014995_Cluster28398-002 |
| Cluster28422 | - | - | up | - | up | - | phospholipid hydroperoxide g | SALT00000036356_Cluster28422-001 |
| Cluster2843 | up | up | up | - | - | - | cytochrome P450 superfamily | SALT00000043308_Cluster2843-001 |
| Cluster28430 | - | - | - | - | - | down | [Zea mays] | SALT00000058877_Cluster28430-001 |
| Cluster28433 | - | - | - | - | - | down | Os03g0303100 [Oryza sativa | SALT00000011234_Cluster28433-001 |
| Cluster2844 | - | - | up | - | up | up | F-box LRR-repeat 3-like [Zea | SALT00000075730_Cluster2844-013 |
| Cluster28443 | - | - | down | - | down | - | Os01g0899700 [Oryza sativa | SALT00000050850_Cluster28443-002 |
| Cluster28446 | - | - | down | - | - | - | DNA binding [Zea mays] | SALT00000058085_Cluster28446-002 |
| Cluster28450 | - | - | - | - | down | - | alpha-L-fucosidase 2 precurso | SALT00000006309_Cluster28450-001 |
| Cluster2846 | - | - | - | - | - | down | PDR-like ABC transporter [O | SALT00000087202_Cluster2846-002 |
| Cluster28466 | - | - | down | - | - | down | ribosomal S4 (chloroplast) [Pl | SALT00000050786_Cluster28466-001 |
| Cluster2847 | - | - | up | - | up | up | kinase [Oryza sativa Japonica | SALT00000092954_Cluster2847-010 |
| Cluster28472 | - | - | up | - | - | - | WEB family At2g38370 [Brac | SALT00000055731_Cluster28472-001 |
| Cluster28481 | - | - | down | - | - | down | TPA: hypothetical protein ZEA | SALT00000013968_Cluster28481-002 |
| Cluster28482 | - | - | - | - | - | down | Os06g0634600 [Oryza sativa | SALT00000014739_Cluster28482-002 |
| Cluster28493 | - | - | - | - | down | - | Os03g0249100 [Oryza sativa | SALT00000060381_Cluster28493-004 |
| Cluster28497 | - | - | up | - | - | - | Retrovirus-related Pol poly LI | SALT00000019559_Cluster28497-001 |
| Cluster28499 | - | - | up | - | - | - | UDP-glycosyltransferase 88B | SALT00000038473_Cluster28499-001 |
| Cluster285 | - | - | down | - | down | - | DNA-directed RNA polymera | SALT00000092396_Cluster285-002 |
| Cluster2850 | - | - | down | - | down | down | proline-rich 2 [Oryza sativa Ja | SALT00000001487_Cluster2850-001 |
| Cluster28514 | - | - | up | - | - | - | chalcone synthase [Oryza sati | SALT00000004031_Cluster28514-001 |
| Cluster28536 | - | - | down | - | - | down | ---NA--- | SALT00000010910_Cluster28536-001 |
| Cluster2854 | - | - | up | - | up | up | PREDICTED: uncharacterized | SALT00000033894_Cluster2854-003 |
| Cluster28559 | - | - | - | - | down | - | PREDICTED: uncharacterized | SALT00000044032_Cluster28559-002 |
| Cluster28560 | - | - | down | - | down | down | methionyl-tRNA synthetase [( | SALT00000048285_Cluster28560-002 |
| Cluster28568 | up | up | up | up | up | up | dehydrin COR410 [Zea mays] | SALT00000018434_Cluster28568-001 |
| Cluster28584 | - | - | - | - | down | - | Os04g0379900 [Oryza sativa | SALT00000019193_Cluster28584-003 |
| Cluster28587 | - | - | up | - | - | - | cyclic dof factor 1-like [Setari | SALT00000036337_Cluster28587-001 |
| Cluster28589 | - | - | - | - | up | - | inositol-tetrakisphosphate 1-k | SALT00000040459_Cluster28589-001 |
| Cluster28597 | - | down | down | - | down | - | Malate dehydrogenase [NADI | SALT00000008210_Cluster28597-001 |
| Cluster28599 | up | - | up | - | - | - | DRE binding factor 1 [Zea ma | SALT00000012350_Cluster28599-003 |
| Cluster28600 | - | - | up | - | up | - | auxin-responsive IAA18 isofo | SALT00000012484_Cluster28600-001 |
| Cluster28603 | - | - | up | - | up | up | HSF transcription partial [Zea | SALT00000040608_Cluster28603-003 |
| Cluster28621 | - | down | - | - | - | - | receptor kinase CLAVATA1 p | SALT00000006386_Cluster28621-005 |
| Cluster28637 | - | - | - | - | up | - | cysteine synthase | SALT00000048078_Cluster28637-004 |
| Cluster28652 | - | - | up | - | up | up | DNA binding [Zea mays] | SALT00000046686_Cluster28652-002 |
| Cluster28657 | down | - | up | - | up | up | GDSL esterase lipase At5g45 | SALT00000054233_Cluster28657-006 |
| Cluster28670 | - | - | up | - | - | - | probable serine acetyltransfera | SALT00000018975_Cluster28670-002 |
| Cluster28674 | up | up | up | - | up | up | senescence-inducible chloropl | SALT00000045181_Cluster28674-003 |
| Cluster28683 | - | - | up | - | up | up | Os06g0215100 [Oryza sativa | SALT00000056802_Cluster28683-002 |
| Cluster28689 | - | - | down | - | down | - | CBL-interacting kinase 4-like | SALT00000013552_Cluster28689-002 |
| Cluster2869 | - | up | - | - | - | - | Double-stranded RNA binding | SALT00000091763_Cluster2869-001 |
| Cluster28698 | - | - | - | - | up | up | TPA: ATPP2-A13 [Zea mays] | SALT00000056836_Cluster28698-002 |
| Cluster28701 | - | - | down | - | down | - | Os10g0569200 [Oryza sativa | SALT00000066875_Cluster28701-001 |
| Cluster28708 | - | - | up | - | - | - | glutamate receptor -like | SALT00000048090_Cluster28708-001 |
| Cluster28709 | - | - | up | - | - | - | PREDICTED: uncharacterized | SALT00000049645_Cluster28709-004 |
| Cluster28739 | - | - | down | - | down | down | Os03g0678800 [Oryza sativa | SALT00000058182_Cluster28739-001 |
| Cluster28743 | - | - | up | - | - | - | UPF0481 At3g47200-like | SALT00000010266_Cluster28743-001 |

| | | | | | | | |
|---|---|---|---|---|---|---|---|
| Cluster28745 | - | - | - | - | down | - | ribosomal RNA small subunit | SALT00000019545_Cluster28745-003 |
| Cluster28746 | - | up | up | - | up | up | CUE domain containing [Zea | SALT00000020535_Cluster28746-002 |
| Cluster28748 | up | - | - | - | - | - | IRX15-LIKE [Oryza sativa Ja | SALT00000040967_Cluster28748-002 |
| Cluster28763 | - | - | - | - | - | up | cytokinin-O-glucosyltransfera | SALT00000017132_Cluster28763-001 |
| Cluster28765 | - | - | up | - | up | up | indole-3-acetate beta-glucosyl | SALT00000028432_Cluster28765-001 |
| Cluster28778 | - | - | - | - | - | up | Os05g0182700 [Oryza sativa | SALT00000053253_Cluster28778-001 |
| Cluster28782 | - | - | - | - | - | down | anther-specific proline-rich Al | SALT00000001449_Cluster28782-001 |
| Cluster28783 | - | - | up | - | up | up | uncharacterized LOC1002166 | SALT00000002923_Cluster28783-003 |
| Cluster2881 | - | - | up | - | - | - | SMG7 [Zea mays] | SALT00000082837_Cluster2881-001 |
| Cluster28828 | - | - | down | - | - | - | ---NA--- | SALT00000020491_Cluster28828-002 |
| Cluster28831 | - | - | up | - | up | - | serine carboxypeptidase 1 pre | SALT00000037351_Cluster28831-001 |
| Cluster28842 | - | - | up | - | up | up | UDP-glycosyltransferase 83A | SALT00000001395_Cluster28842-001 |
| Cluster28844 | - | - | down | - | down | down | chloroplast envelope membra | SALT00000004813_Cluster28844-001 |
| Cluster28846 | - | - | down | - | - | - | CCT motif family [Zea mays] | SALT00000015210_Cluster28846-001 |
| Cluster28868 | - | - | up | - | - | up | alcohol dehydrogenase superf | SALT00000010831_Cluster28868-001 |
| Cluster2887 | - | - | up | - | - | - | argonaute 4A | SALT00000078137_Cluster2887-018 |
| Cluster2888 | - | - | up | - | up | up | ariadne [Oryza sativa Japonica | SALT00000038200_Cluster2888-025 |
| Cluster28887 | - | - | - | - | - | up | cis-zeatin O-glucosyltransfera | SALT00000000281_Cluster28887-001 |
| Cluster28889 | - | - | down | - | - | - | nucleoside transporter [Zea m | SALT00000006022_Cluster28889-003 |
| Cluster28897 | - | - | - | - | down | - | [Hordeum vulgare vulgare] | SALT00000017208_Cluster28897-001 |
| Cluster28919 | - | - | up | - | - | - | glycine-rich [Oryza sativa Jap | SALT00000025139_Cluster28919-002 |
| Cluster28920 | - | - | up | - | up | - | AMSH-like ubiquitin thioeste | SALT00000036120_Cluster28920-002 |
| Cluster28926 | - | - | down | - | down | down | ---NA--- | SALT00000046321_Cluster28926-001 |
| Cluster28932 | - | - | - | - | down | - | inorganic pyrophosphatase [Z | SALT00000054813_Cluster28932-002 |
| Cluster28941 | - | - | - | - | - | down | NAC domain-containing 43 | SALT00000048066_Cluster28941-001 |
| Cluster28949 | - | - | up | - | - | - | F-box domain containing [Zea | SALT00000004633_Cluster28949-001 |
| Cluster2895 | - | - | - | - | down | - | WEB family chloroplastic-like | SALT00000034186_Cluster2895-003 |
| Cluster28959 | up | up | up | up | up | up | NAC1 [Zea mays] | SALT00000043227_Cluster28959-002 |
| Cluster28960 | - | - | up | - | up | up | Os10g0574700 [Oryza sativa | SALT00000045163_Cluster28960-003 |
| Cluster28964 | - | - | up | - | up | up | DNA-binding phosphatase 2C | SALT00000003910_Cluster28964-001 |
| Cluster2897 | - | - | - | - | down | - | TPA: respiratory burst oxidase | SALT00000089231_Cluster2897-001 |
| Cluster28973 | - | - | - | - | up | - | ATMYB4-like [Oryza sativa J | SALT00000044586_Cluster28973-001 |
| Cluster2898 | - | - | down | - | - | - | Os10g0101000 [Oryza sativa | SALT00000090110_Cluster2898-002 |
| Cluster28985 | - | - | down | - | down | down | Os11g0171300 [Oryza sativa | SALT00000003996_Cluster28985-001 |
| Cluster28993 | - | - | up | - | up | - | peroxisomal membrane 11-5 [ | SALT00000038363_Cluster28993-001 |
| Cluster29006 | - | - | up | - | up | - | Os03g0289800 [Oryza sativa | SALT00000012544_Cluster29006-001 |
| Cluster29020 | - | - | up | - | - | - | serine esterase family [Zea ma | SALT00000059181_Cluster29020-002 |
| Cluster29028 | - | - | down | - | - | - | Enoyl-[acyl-carrier- ] reductas | SALT00000014405_Cluster29028-001 |
| Cluster29029 | - | - | down | - | down | down | Os03g0857400 [Oryza sativa | SALT00000015287_Cluster29029-002 |
| Cluster29043 | - | - | up | - | up | up | Os05g0516600 [Oryza sativa | SALT00000065993_Cluster29043-002 |
| Cluster29070 | - | up | up | - | - | - | THO complex subunit 6 [Seta | SALT00000037092_Cluster29070-001 |
| Cluster29072 | - | - | - | - | up | up | probable RNA-binding ARP1 | SALT00000040765_Cluster29072-002 |
| Cluster29074 | - | - | up | - | up | up | GDSL esterase lipase At5g45 | SALT00000049851_Cluster29074-003 |
| Cluster29080 | - | - | up | - | up | up | binding [Zea mays] | SALT00000002733_Cluster29080-001 |
| Cluster29100 | - | - | down | - | down | - | NADPH-dependent reductase | SALT00000003659_Cluster29100-001 |
| Cluster29104 | - | - | up | - | up | up | myb-related Myb4 [Zea mays | SALT00000014144_Cluster29104-001 |
| Cluster29121 | - | - | - | - | up | - | Os01g0141000 [Oryza sativa | SALT00000057585_Cluster29121-006 |
| Cluster29122 | - | - | down | - | down | - | [Oryza sativa Japonica Group | SALT00000058623_Cluster29122-001 |
| Cluster29125 | - | - | up | - | up | up | neutral invertase [Oryza sativa | SALT00000006372_Cluster29125-001 |
| Cluster29136 | - | - | up | - | up | up | Os05g0358400 [Oryza sativa | SALT00000038241_Cluster29136-001 |
| Cluster2915 | - | up | up | - | up | - | phosphoinositide phosphatase | SALT00000092958_Cluster2915-011 |
| Cluster29157 | - | - | down | - | - | - | expansin-B2-like [Zea mays] | SALT00000049013_Cluster29157-002 |
| Cluster2918 | - | - | down | - | - | - | ribosomal S3 (chloroplast) [S | SALT00000091048_Cluster2918-016 |
| Cluster29185 | - | - | up | - | up | up | peroxidase 5-like [Setaria itali | SALT00000047593_Cluster29185-007 |
| Cluster29195 | - | - | down | - | - | down | phosphate phosphoenolpyruva | SALT00000043406_Cluster29195-004 |
| Cluster29226 | - | - | - | - | - | down | auxin-repressed kDa | SALT00000057960_Cluster29226-004 |
| Cluster29232 | - | - | - | - | - | up | Os01g0141000 [Oryza sativa | SALT00000015080_Cluster29232-001 |
| Cluster29233 | - | down | down | - | - | - | Os12g0420200 [Oryza sativa | SALT00000015177_Cluster29233-001 |

| | | | | | | | | |
|---|---|---|---|---|---|---|---|---|
| Cluster29248 | - | - | up | - | - | - | ---NA--- | SALT00000019741_Cluster29248-001 |
| Cluster2925 | - | - | - | - | down | down | TPA: CRM family member 3 | SALT00000070373_Cluster2925-004 |
| Cluster29264 | - | down | down | - | - | - | uncharacterized LOC1008568 | SALT00000017969_Cluster29264-003 |
| Cluster29265 | - | - | down | - | - | - | 3-isopropylmalate dehydroger | SALT00000035076_Cluster29265-003 |
| Cluster29272 | - | - | up | - | - | - | fumarylacetoacetase [Zea may | SALT00000046655_Cluster29272-003 |
| Cluster29290 | - | - | up | - | up | up | Os08g0492100 [Oryza sativa | SALT00000017314_Cluster29290-002 |
| Cluster29293 | - | - | down | - | - | - | cytochrome b561 and DOMO | SALT00000046479_Cluster29293-002 |
| Cluster293 | up | up | up | - | - | - | ---NA--- | SALT00000063356_Cluster293-001 |
| Cluster2930 | - | - | - | - | - | down | rp3-like disease resistance | SALT00000021895_Cluster2930-001 |
| Cluster29306 | - | - | up | - | up | up | Os01g0867300 [Oryza sativa | SALT00000011524_Cluster29306-001 |
| Cluster29326 | - | - | up | - | up | up | ethylene-responsive transcript | SALT00000035853_Cluster29326-001 |
| Cluster2933 | - | - | down | - | - | - | TPA: trehalose phosphatase sy | SALT00000088670_Cluster2933-001 |
| Cluster29342 | - | - | down | - | down | - | [Hordeum vulgare vulgare] | SALT00000006868_Cluster29342-002 |
| Cluster29349 | - | - | - | - | - | down | anter-specific proline-rich AP | SALT00000037877_Cluster29349-001 |
| Cluster29351 | - | - | up | - | - | - | UDP-glycosyltransferase 89B | SALT00000041573_Cluster29351-001 |
| Cluster29361 | - | up | up | - | - | - | ubiquitin SMT3 [Zea mays] | SALT00000013221_Cluster29361-002 |
| Cluster29363 | - | - | - | - | down | - | microtubule-associated 70-1-l | SALT00000023790_Cluster29363-001 |
| Cluster29364 | - | - | up | - | - | - | [Hordeum vulgare vulgare] | SALT00000035711_Cluster29364-001 |
| Cluster29367 | - | - | - | - | - | down | MYBY1 [Oryza sativa Japoni | SALT00000045689_Cluster29367-002 |
| Cluster29377 | - | - | up | - | - | - | Os06g0695500 [Oryza sativa | SALT00000038671_Cluster29377-002 |
| Cluster29389 | up | - | - | - | down | - | Ubiquitin carboxyl-terminal h | SALT00000057921_Cluster29389-001 |
| Cluster29394 | - | - | up | - | - | - | endochitinase B-like [Setaria i | SALT00000014080_Cluster29394-001 |
| Cluster29398 | - | - | down | - | down | down | maturase K (chloroplast) | SALT00000036795_Cluster29398-001 |
| Cluster294 | - | - | - | - | up | - | acidic leucine-rich nuclear pho | SALT00000054969_Cluster294-006 |
| Cluster29410 | - | - | up | - | up | up | ZCN11 [Zea mays] | SALT00000014336_Cluster29410-001 |
| Cluster29415 | - | - | up | - | - | - | [Hordeum vulgare vulgare] | SALT00000044368_Cluster29415-004 |
| Cluster29422 | - | - | up | - | up | up | NADPH HC toxin reductase | SALT00000051005_Cluster29422-002 |
| Cluster29429 | - | - | down | - | down | - | phosphatase 2C family [Zea m | SALT00000060448_Cluster29429-002 |
| Cluster29434 | - | - | down | - | down | - | Os03g0784700 [Oryza sativa | SALT00000008519_Cluster29434-002 |
| Cluster2944 | - | - | down | - | - | - | cyclin-dependent kinase G-2-l | SALT00000091073_Cluster2944-004 |
| Cluster29456 | - | - | - | - | - | down | mitochondrial outer membran | SALT00000008756_Cluster29456-001 |
| Cluster29457 | - | up | - | - | - | - | RING finger 1 [Oryza sativa J | SALT00000009684_Cluster29457-001 |
| Cluster29459 | - | - | up | - | up | - | U-box domain-containing 4 | SALT00000010864_Cluster29459-002 |
| Cluster29472 | - | - | down | - | down | down | F-box domain containing [Zea | SALT00000008726_Cluster29472-001 |
| Cluster29486 | - | - | down | - | - | - | GATA transcription factor 22 | SALT00000051549_Cluster29486-001 |
| Cluster29489 | - | - | up | - | up | up | Os02g0654700 [Oryza sativa | SALT00000059224_Cluster29489-003 |
| Cluster29490 | - | - | down | - | - | - | coproporphyrinogen III oxida | SALT00000005893_Cluster29490-003 |
| Cluster29494 | - | - | up | - | up | up | alternative oxidase [Oryza sat | SALT00000019534_Cluster29494-002 |
| Cluster29496 | - | - | - | - | down | - | Os07g0198300 [Oryza sativa | SALT00000038471_Cluster29496-001 |
| Cluster29499 | - | - | - | - | up | up | Os01g0693400 [Oryza sativa | SALT00000042386_Cluster29499-001 |
| Cluster29500 | - | - | - | - | down | down | Os09g0461500 [Oryza sativa | SALT00000045218_Cluster29500-002 |
| Cluster29503 | - | - | down | - | - | - | [Hordeum vulgare vulgare] | SALT00000060063_Cluster29503-001 |
| Cluster29514 | - | up | up | - | up | up | no apical meristem [Zea mays | SALT00000040591_Cluster29514-002 |
| Cluster29523 | - | - | - | - | up | - | ferredoxin- chloroplastic [Seta | SALT00000001612_Cluster29523-001 |
| Cluster29528 | - | - | down | - | - | - | flagellar radial spoke 5 [Oryza | SALT00000009059_Cluster29528-001 |
| Cluster29546 | - | - | down | - | down | - | 28 kDa chloroplastic-like | SALT00000048182_Cluster29546-002 |
| Cluster29548 | - | - | - | - | up | - | syntaxin 121 [Zea mays] | SALT00000053715_Cluster29548-001 |
| Cluster29560 | - | - | down | - | down | - | 40S ribosomal S9 [Zea mays] | SALT00000043650_Cluster29560-001 |
| Cluster29571 | - | - | up | - | up | up | Os01g0227100 [Oryza sativa | SALT00000008874_Cluster29571-001 |
| Cluster29574 | - | - | up | - | - | - | Os05g0156300 [Oryza sativa | SALT00000041525_Cluster29574-005 |
| Cluster29598 | - | - | - | - | up | - | lipase precursor [Zea mays] | SALT00000010328_Cluster29598-001 |
| Cluster296 | - | - | up | - | up | up | Os01g0185300 [Oryza sativa | SALT00000043052_Cluster296-005 |
| Cluster29627 | - | - | down | - | down | down | chlorophyll a b-binding [Oryz | SALT00000055599_Cluster29627-004 |
| Cluster29632 | - | - | - | - | up | - | glucose-6-phosphate phosphat | SALT00000013931_Cluster29632-001 |
| Cluster29651 | - | - | down | - | down | - | mitochondrial phosphate trans | SALT00000011527_Cluster29651-001 |
| Cluster29653 | - | - | - | - | down | - | partial [Oryza sativa Japonica | SALT00000037086_Cluster29653-001 |
| Cluster29659 | - | - | up | - | up | up | [Hordeum vulgare vulgare] | SALT00000046273_Cluster29659-004 |
| Cluster29685 | - | - | - | - | down | down | Os02g0589000 [Oryza sativa | SALT00000010914_Cluster29685-001 |

| | | | | | | | | |
|---|---|---|---|---|---|---|---|---|
| Cluster2969 | - | - | up | - | up | up | Os03g0179900 [Oryza sativa | SALT00000078711_Cluster2969-015 |
| Cluster29696 | down | - | - | - | - | - | phosphate transporter PHO1-2 | SALT00000067653_Cluster29696-001 |
| Cluster297 | up | up | - | - | down | - | AC092553_17 pol poly [Oryz | SALT00000082141_Cluster297-001 |
| Cluster29710 | - | - | down | - | - | - | ---NA--- | SALT00000057975_Cluster29710-001 |
| Cluster29716 | - | - | up | - | - | - | 3Fe-4S ferredoxin [Zea mays] | SALT00000013284_Cluster29716-001 |
| Cluster29721 | - | up | up | - | up | up | thioredoxin H1-like [Setaria it | SALT00000039387_Cluster29721-001 |
| Cluster29723 | up | up | up | - | - | - | GDA2 [Oryza sativa Japonica | SALT00000043023_Cluster29723-008 |
| Cluster29749 | - | - | down | - | down | down | 60S ribosomal L12 [Zea mays | SALT00000041699_Cluster29749-001 |
| Cluster29751 | - | - | down | - | down | - | [Hordeum vulgare vulgare] | SALT00000046672_Cluster29751-001 |
| Cluster29752 | - | - | up | - | - | - | G-type lectin S-receptor-like s | SALT00000049538_Cluster29752-003 |
| Cluster29767 | - | - | up | - | up | - | Os02g0828200 [Oryza sativa | SALT00000049154_Cluster29767-006 |
| Cluster29782 | - | - | down | down | down | down | Os01g0611000 [Oryza sativa | SALT00000047838_Cluster29782-004 |
| Cluster29783 | - | down | down | - | down | - | zinc finger [Oryza sativa Japo | SALT00000049105_Cluster29783-001 |
| Cluster29797 | - | - | down | - | - | - | receptor expression-enhancing | SALT00000012521_Cluster29797-002 |
| Cluster29799 | - | - | down | - | down | - | lactate dehydrogenase [Oryza | SALT00000018986_Cluster29799-003 |
| Cluster298 | - | - | down | - | down | - | plastid transported acetyl- carl | SALT00000090838_Cluster298-025 |
| Cluster2981 | - | - | down | - | down | - | ferric reductase-like transmem | SALT00000066782_Cluster2981-008 |
| Cluster29831 | - | - | up | - | up | up | photoperiod responsive [Zea m | SALT00000055752_Cluster29831-001 |
| Cluster29837 | - | - | up | - | up | up | hypothetical protein SETIT_0 | SALT00000012535_Cluster29837-001 |
| Cluster29845 | - | - | up | - | up | up | [Hordeum vulgare vulgare] | SALT00000038055_Cluster29845-004 |
| Cluster29848 | - | - | up | - | up | up | isocitrate dehydrogenase [NA | SALT00000056764_Cluster29848-005 |
| Cluster29852 | - | - | down | - | down | - | PHD finger [Zea mays] | SALT00000058497_Cluster29852-002 |
| Cluster29854 | - | - | down | - | down | down | Os11g0171300 [Oryza sativa | SALT00000001399_Cluster29854-002 |
| Cluster2986 | - | - | up | - | up | up | OBERON 4 [Oryza sativa Jap | SALT00000077195_Cluster2986-004 |
| Cluster29873 | - | down | down | down | down | down | GATA transcription factor 2-li | SALT00000003841_Cluster29873-002 |
| Cluster29898 | - | - | up | - | - | up | Os07g0503300 [Oryza sativa | SALT00000020045_Cluster29898-001 |
| Cluster299 | - | - | down | - | down | down | PREDICTED: uncharacterize | SALT00000092348_Cluster299-001 |
| Cluster29905 | - | - | down | - | - | - | cycloartenol-C-24-methyltran | SALT00000049335_Cluster29905-005 |
| Cluster29915 | - | up | - | - | - | - | NNJA3_WHEAT ame: Full=N | SALT00000024190_Cluster29915-001 |
| Cluster29945 | - | - | down | - | down | down | 60S ribosomal L13a [Zea may | SALT00000037242_Cluster29945-001 |
| Cluster29956 | - | - | up | - | - | - | phosphatase 2C family [Zea m | SALT00000057176_Cluster29956-002 |
| Cluster29964 | - | - | down | - | - | - | nucleosome chromatin assemb | SALT00000012711_Cluster29964-002 |
| Cluster29969 | - | - | - | - | down | - | protease 2 | SALT00000047417_Cluster29969-001 |
| Cluster2997 | - | - | down | - | - | - | argonaute 1B isoform X1 | SALT00000088011_Cluster2997-030 |
| Cluster29972 | - | - | up | - | - | - | Os08g0558700 [Oryza sativa | SALT00000051526_Cluster29972-001 |
| Cluster29978 | - | - | up | - | - | - | ATP-dependent protease Clp A | SALT00000067757_Cluster29978-001 |
| Cluster29980 | - | - | - | - | down | down | Os06g0300300 [Oryza sativa | SALT00000005465_Cluster29980-001 |
| Cluster29983 | - | - | - | - | - | down | uncharacterized methyltransfe | SALT00000012367_Cluster29983-001 |
| Cluster29997 | down | - | down | - | - | - | SRG1-like [Brachypodium dis | SALT00000063092_Cluster29997-003 |
| Cluster30010 | - | - | down | - | down | down | zinc transporter 10 precursor [ | SALT00000001217_Cluster30010-001 |
| Cluster30011 | - | - | up | - | up | up | Os03g0291200 [Oryza sativa | SALT00000008569_Cluster30011-001 |
| Cluster30015 | - | - | down | - | down | - | divinyl chlorophyllide a 8-vin | SALT00000048669_Cluster30015-002 |
| Cluster30017 | - | - | - | - | - | up | KH domain-containing At5g5 | SALT00000049269_Cluster30017-003 |
| Cluster30026 | - | - | up | - | up | up | alginate regulatory [Zea mays | SALT00000017826_Cluster30026-003 |
| Cluster30035 | - | - | down | - | down | - | porphobilinogen deaminase [Z | SALT00000002567_Cluster30035-001 |
| Cluster30039 | - | - | up | - | up | up | 2OG-Fe oxygenase family exp | SALT00000014894_Cluster30039-001 |
| Cluster30042 | - | - | up | - | - | - | cyclin-P3-1-like [Setaria italic | SALT00000038379_Cluster30042-001 |
| Cluster30048 | - | down | down | - | - | - | Os06g0593200 [Oryza sativa | SALT00000002302_Cluster30048-001 |
| Cluster30056 | - | - | down | - | - | - | ribose-phosphate pyrophospho | SALT00000016129_Cluster30056-004 |
| Cluster30061 | - | - | up | - | - | - | 3-hydroxybutyryl- dehydroge | SALT00000057690_Cluster30061-001 |
| Cluster30068 | - | up | up | up | up | - | Os05g0560900 [Oryza sativa | SALT00000013742_Cluster30068-001 |
| Cluster30069 | - | - | up | - | up | - | TPA: metal ion binding [Zea m | SALT00000043158_Cluster30069-006 |
| Cluster3007 | - | - | - | - | down | - | RNA polymerase beta subunit | SALT00000087055_Cluster3007-008 |
| Cluster30071 | - | - | up | - | up | up | ATMYB4-like [Oryza sativa J | SALT00000038084_Cluster30071-001 |
| Cluster30078 | - | - | - | - | down | - | phytoene synthase 2 [Zea may | SALT00000056416_Cluster30078-002 |
| Cluster30083 | - | - | up | - | up | up | probable phosphatase 2C 44 [ | SALT00000018972_Cluster30083-002 |
| Cluster30085 | - | - | up | - | up | up | WRKY transcription factor [C | SALT00000028798_Cluster30085-002 |
| Cluster30086 | up | up | up | - | up | - | probable nucleoredoxin 2 | SALT00000037910_Cluster30086-001 |

| | | | | | | | | |
|---|---|---|---|---|---|---|---|---|
| Cluster3009 | - | - | up | - | - | - | Os05g0592600 [Oryza sativa | SALT00000021795_Cluster3009-002 |
| Cluster30091 | - | - | up | - | up | up | [Hordeum vulgare vulgare] | SALT00000059489_Cluster30091-001 |
| Cluster30095 | - | - | up | - | up | up | WRKY transcription factor [C | SALT00000006112_Cluster30095-003 |
| Cluster30098 | - | - | up | - | up | up | metal ion binding [Zea mays] | SALT00000041117_Cluster30098-001 |
| Cluster3010 | - | - | down | - | down | down | Os05g0482600 [Oryza sativa | SALT00000056899_Cluster3010-010 |
| Cluster30100 | - | - | up | - | up | up | probable LRR receptor-like se | SALT00000048277_Cluster30100-001 |
| Cluster30109 | - | - | up | - | - | - | CCT [Zea mays] | SALT00000017535_Cluster30109-001 |
| Cluster30112 | - | up | up | - | - | - | late embryogenesis abundant l | SALT00000023083_Cluster30112-001 |
| Cluster30118 | - | up | up | - | up | up | homocysteine S-methyltransfe | SALT00000057766_Cluster30118-002 |
| Cluster30123 | - | down | - | - | - | up | NAC domain-containing 48-li | SALT00000013838_Cluster30123-002 |
| Cluster3013 | up | up | up | - | - | - | Os05g0365500 [Oryza sativa | SALT00000077931_Cluster3013-001 |
| Cluster30138 | - | - | down | - | - | - | Os04g0656800 [Oryza sativa | SALT00000057408_Cluster30138-003 |
| Cluster30144 | - | - | up | - | up | up | Os05g0534400 [Oryza sativa | SALT00000014301_Cluster30144-001 |
| Cluster30150 | up | up | up | - | - | - | Os03g0746700 [Oryza sativa | SALT00000059002_Cluster30150-001 |
| Cluster30152 | - | up | - | - | - | down | ATP-dependent (S)-NAD(P)H | SALT00000013361_Cluster30152-001 |
| Cluster30156 | - | up | up | - | up | up | NAC1 [Sorghum bicolor] | SALT00000040896_Cluster30156-001 |
| Cluster30158 | - | - | up | - | up | - | zinc finger | SALT00000042246_Cluster30158-001 |
| Cluster30164 | up | up | - | - | - | - | probable phosphatase 2C 49 [ | SALT00000001096_Cluster30164-001 |
| Cluster30165 | - | - | down | - | down | down | Os07g0683900 [Oryza sativa | SALT00000010878_Cluster30165-002 |
| Cluster30193 | - | - | up | - | up | - | [Hordeum vulgare vulgare] | SALT00000001650_Cluster30193-001 |
| Cluster302 | - | - | - | - | - | down | partial [Oryza sativa Japonica | SALT00000088855_Cluster302-003 |
| Cluster30209 | - | - | up | - | up | - | Os03g0116800 [Oryza sativa | SALT00000002697_Cluster30209-001 |
| Cluster30250 | - | - | - | - | down | - | EMB12 [Zea mays] | SALT00000051252_Cluster30250-004 |
| Cluster30257 | down | down | down | - | down | - | Os01g0866400 [Oryza sativa | SALT00000053258_Cluster30257-002 |
| Cluster30263 | - | up | - | - | - | - | ubiquitin-conjugating enzyme | SALT00000019917_Cluster30263-001 |
| Cluster30268 | - | - | - | - | down | - | peroxidase 72-like [Setaria ita | SALT00000044716_Cluster30268-002 |
| Cluster30304 | - | - | down | - | down | - | 50S ribosomal L17 [Zea mays | SALT00000016720_Cluster30304-001 |
| Cluster30307 | - | - | down | - | down | down | 40S ribosomal S16 [Zea mays | SALT00000037672_Cluster30307-001 |
| Cluster30314 | down | - | - | - | - | - | Os03g0830900 [Oryza sativa | SALT00000053048_Cluster30314-001 |
| Cluster3032 | - | - | up | - | up | - | Os01g0621900 [Oryza sativa | SALT00000092081_Cluster3032-018 |
| Cluster30320 | - | - | down | - | down | down | photosystem I [Oryza sativa J | SALT00000007733_Cluster30320-001 |
| Cluster30328 | - | down | down | down | down | down | fasciclin-like arabinogalactan | SALT00000067235_Cluster30328-002 |
| Cluster30334 | - | - | down | - | down | - | peptidyl-prolyl cis-trans isom | SALT00000014520_Cluster30334-001 |
| Cluster30346 | - | - | up | - | up | up | senescence-associated [Zea m | SALT00000056465_Cluster30346-001 |
| Cluster30364 | - | - | - | - | down | down | CONSTANS CO6 [Zea mays] | SALT00000041923_Cluster30364-002 |
| Cluster30365 | - | - | - | - | up | - | E3 ubiquitin- ligase ATL6-lik | SALT00000043234_Cluster30365-001 |
| Cluster3037 | - | - | down | - | - | - | DUF1296 domain containing | SALT00000071487_Cluster3037-005 |
| Cluster30382 | - | - | up | - | - | - | ---NA--- | SALT00000039322_Cluster30382-002 |
| Cluster30394 | - | - | down | - | - | - | uncharacterized oxidoreducta | SALT00000010407_Cluster30394-001 |
| Cluster30403 | - | - | up | - | up | up | Os03g0708200 [Oryza sativa | SALT00000008464_Cluster30403-001 |
| Cluster30409 | - | - | up | - | up | up | 12-oxophytodienoate reducta | SALT00000042020_Cluster30409-002 |
| Cluster30429 | - | - | - | - | down | - | CONSTANS CO6 [Zea mays] | SALT00000048984_Cluster30429-001 |
| Cluster30433 | - | - | down | - | down | down | Os03g0807700 [Oryza sativa | SALT00000010179_Cluster30433-002 |
| Cluster30447 | - | up | up | - | up | up | partial [Oryza sativa Japonica | SALT00000010259_Cluster30447-001 |
| Cluster30448 | - | up | - | - | - | - | alkaline ceramidase 3 [Setaria | SALT00000013031_Cluster30448-002 |
| Cluster30450 | - | - | up | - | up | up | probable carboxylesterase 15 | SALT00000035840_Cluster30450-001 |
| Cluster30459 | - | - | down | - | - | - | xyloglucan endotransglycosyl | SALT00000004010_Cluster30459-001 |
| Cluster3046 | - | up | - | - | - | down | Os01g0873800 [Oryza sativa | SALT00000090845_Cluster3046-010 |
| Cluster30467 | - | - | - | - | - | down | IAA16 - auxin-responsive Au | SALT00000045958_Cluster30467-001 |
| Cluster30473 | - | - | down | - | down | - | Os05g0597100 [Oryza sativa | SALT00000061752_Cluster30473-002 |
| Cluster3048 | - | - | down | - | down | down | ribosomal S2 (chloroplast) | SALT00000078083_Cluster3048-031 |
| Cluster30481 | - | - | up | - | up | - | peptide chain release factor c | SALT00000045629_Cluster30481-001 |
| Cluster30482 | up | up | up | - | up | up | Os08g0529300 [Oryza sativa | SALT00000053934_Cluster30482-001 |
| Cluster30488 | - | - | down | - | down | down | Os01g0753800 [Oryza sativa | SALT00000003552_Cluster30488-001 |
| Cluster30489 | - | - | down | - | - | - | Os10g0556100 [Oryza sativa | SALT00000005296_Cluster30489-001 |
| Cluster3049 | - | - | - | - | down | down | Transposase family | SALT00000078310_Cluster3049-001 |
| Cluster30505 | - | - | down | - | - | - | lecithin-cholesterol acyltransf | SALT00000002597_Cluster30505-001 |
| Cluster30507 | up | up | up | - | up | up | GPALPP motifs-containing 1- | SALT00000008631_Cluster30507-002 |

| | | | | | | | | |
|---|---|---|---|---|---|---|---|---|
| Cluster3051 | - | - | up | - | - | - | serine carboxypeptidase 1 pre | SALT00000039969_Cluster3051-021 |
| Cluster30510 | - | - | up | - | up | up | NAC1 [Sorghum bicolor] | SALT00000036376_Cluster30510-002 |
| Cluster30516 | - | - | down | - | - | - | tas-like isoform X2 | SALT00000048155_Cluster30516-004 |
| Cluster30518 | - | up | up | - | up | - | Anthranilate N-benzoyltransfe | SALT00000000897_Cluster30518-001 |
| Cluster30519 | - | - | up | - | up | up | binding [Zea mays] | SALT00000009142_Cluster30519-001 |
| Cluster30537 | - | - | - | - | down | - | EMBRYO SAC DEVELOPM | SALT00000010980_Cluster30537-001 |
| Cluster30540 | - | down | down | - | down | - | PLATZ transcription partial [Z | SALT00000043414_Cluster30540-002 |
| Cluster30543 | - | down | down | - | down | - | peroxidase 1b | SALT00000050393_Cluster30543-002 |
| Cluster30549 | - | up | - | - | - | down | ---NA--- | SALT00000041296_Cluster30549-001 |
| Cluster30552 | - | up | - | - | - | down | metacaspase family [Zea mays | SALT00000011320_Cluster30552-002 |
| Cluster30564 | - | - | down | - | - | - | [Hordeum vulgare vulgare] | SALT00000008399_Cluster30564-001 |
| Cluster30568 | - | - | down | - | - | - | peroxidase 1 precursor [Zea m | SALT00000016022_Cluster30568-001 |
| Cluster30572 | - | - | down | - | - | - | HRT transcription partial | SALT00000039808_Cluster30572-001 |
| Cluster30580 | - | - | - | - | up | - | [Hordeum vulgare vulgare] | SALT00000057881_Cluster30580-002 |
| Cluster30582 | - | - | up | - | up | up | heme oxygenase 1 | SALT00000002381_Cluster30582-001 |
| Cluster30591 | - | - | up | - | - | - | Os02g0104200 [Oryza sativa | SALT00000051632_Cluster30591-001 |
| Cluster30596 | - | - | down | - | down | - | [Hordeum vulgare vulgare] | SALT00000018218_Cluster30596-002 |
| Cluster30598 | - | up | - | - | - | - | Os01g0104400 [Oryza sativa | SALT00000035715_Cluster30598-002 |
| Cluster30608 | up | up | up | - | up | up | gibberellin receptor GID1L2 [ | SALT00000058428_Cluster30608-001 |
| Cluster30611 | - | - | up | - | up | up | Os02g0797600 [Oryza sativa | SALT00000000179_Cluster30611-001 |
| Cluster30616 | - | - | down | - | down | - | acyltransferase [Zea mays] | SALT00000016339_Cluster30616-001 |
| Cluster30618 | - | - | up | - | - | - | Os11g0536000 [Oryza sativa | SALT00000038823_Cluster30618-001 |
| Cluster3062 | - | - | - | - | down | - | Ruda poly | SALT00000081543_Cluster3062-001 |
| Cluster30633 | - | - | down | - | - | - | peroxidase 24 precursor [Zea | SALT00000014280_Cluster30633-002 |
| Cluster3064 | - | down | down | - | - | - | homolog subfamily C membe | SALT00000087912_Cluster3064-003 |
| Cluster30644 | - | - | down | - | down | - | Strictosidine synthase 1 precu | SALT00000013163_Cluster30644-001 |
| Cluster30650 | - | - | down | - | down | down | brown planthopper-induced re | SALT00000052150_Cluster30650-002 |
| Cluster30654 | - | - | up | - | - | - | cytosolic sulfotransferase 5-lil | SALT00000005801_Cluster30654-001 |
| Cluster30665 | - | up | up | - | - | - | Os11g0614400 [Oryza sativa | SALT00000004850_Cluster30665-002 |
| Cluster30667 | - | up | up | up | up | - | NAC transcription factor 29 [ | SALT00000011992_Cluster30667-002 |
| Cluster30676 | - | - | up | - | up | up | NAC transcription factor 29-li | SALT00000046504_Cluster30676-001 |
| Cluster30686 | - | - | down | - | down | down | ---NA--- | SALT00000030210_Cluster30686-001 |
| Cluster30696 | - | - | down | - | - | - | uncharacterized membrane At | SALT00000076437_Cluster30696-003 |
| Cluster30704 | - | - | up | - | up | up | ---NA--- | SALT00000050530_Cluster30704-001 |
| Cluster30735 | - | - | up | - | up | up | Os02g0510600 [Oryza sativa | SALT00000008372_Cluster30735-002 |
| Cluster30741 | - | - | up | - | - | up | SNAC1 [Oryza nivara] | SALT00000042767_Cluster30741-001 |
| Cluster30756 | - | - | - | - | up | - | carotenoid hydroxylase [Zea m | SALT00000050197_Cluster30756-001 |
| Cluster30759 | - | - | up | - | - | - | replication A 70 kDa DNA-bi | SALT00000052869_Cluster30759-001 |
| Cluster30769 | - | - | - | - | down | - | Transcription factor MYB39 | SALT00000046027_Cluster30769-002 |
| Cluster30778 | - | up | - | - | - | - | 1-aminocyclopropane-1-carbo | SALT00000039426_Cluster30778-002 |
| Cluster30780 | - | down | down | - | - | - | Os01g0922100 [Oryza sativa | SALT00000049494_Cluster30780-001 |
| Cluster30785 | - | down | down | - | - | - | TATA-binding 2 isoform X2 [ | SALT00000039125_Cluster30785-001 |
| Cluster3079 | - | - | up | - | up | up | MYB DNA-binding domain s | SALT00000050953_Cluster3079-002 |
| Cluster30792 | - | - | up | - | up | up | [Hordeum vulgare vulgare] | SALT00000002825_Cluster30792-001 |
| Cluster30800 | - | - | up | - | - | - | GDA2 [Oryza sativa Japonica | SALT00000045862_Cluster30800-001 |
| Cluster30819 | - | - | down | - | - | - | harpin-induced [Zea mays] | SALT00000009026_Cluster30819-001 |
| Cluster3082 | - | - | down | - | down | down | probable helicase MAGATAM | SALT00000068024_Cluster3082-001 |
| Cluster30827 | - | - | - | - | - | down | TPA: heavy metal transport de | SALT00000017575_Cluster30827-001 |
| Cluster30844 | - | - | up | - | - | up | 2 -deoxymugineic-acid 2 -dio | SALT00000039406_Cluster30844-001 |
| Cluster30856 | - | - | up | - | up | up | DIMBOA UDP-glucosyltrans | SALT00000051917_Cluster30856-001 |
| Cluster30859 | - | - | up | - | up | - | HSP20-like chaperone domain | SALT00000012032_Cluster30859-001 |
| Cluster3086 | - | - | up | - | up | up | retrotransposon unclassified [ | SALT00000080887_Cluster3086-001 |
| Cluster30869 | - | - | down | - | down | down | carboxy-lyase [Zea mays] | SALT00000041143_Cluster30869-001 |
| Cluster30876 | - | - | up | - | up | up | vegetative cell wall gp1-like [ | SALT00000038367_Cluster30876-001 |
| Cluster30888 | - | - | down | - | - | - | [Hordeum vulgare vulgare] | SALT00000040703_Cluster30888-003 |
| Cluster3091 | up | - | up | - | up | up | allene oxide synthase [Oryza s | SALT00000064727_Cluster3091-008 |
| Cluster30918 | - | - | down | - | - | - | Os01g0743400 [Oryza sativa | SALT00000049588_Cluster30918-001 |
| Cluster30927 | - | - | up | - | - | - | PREDICTED: uncharacterize | SALT00000043839_Cluster30927-001 |

| | | | | | | | | |
|---|---|---|---|---|---|---|---|---|
| Cluster30930 | - | - | up | - | - | up | Os01g0667600 [Oryza sativa | SALT00000057008_Cluster30930-001 |
| Cluster30935 | - | down | - | - | - | up | alpha-L-fucosidase 2 precurso | SALT00000040075_Cluster30935-005 |
| Cluster30936 | - | - | - | - | - | down | SRG1-like [Setaria italica] | SALT00000040279_Cluster30936-001 |
| Cluster30939 | - | up | up | - | up | up | Os01g0806200 [Oryza sativa | SALT00000055507_Cluster30939-001 |
| Cluster30941 | - | - | down | - | - | - | IAA30 - auxin-responsive Au: | SALT00000061295_Cluster30941-001 |
| Cluster30951 | - | - | up | - | up | up | beta 1,3 glucanase [Zea mays | SALT00000066455_Cluster30951-002 |
| Cluster30955 | - | - | down | - | down | - | IAA30 - auxin-responsive Au: | SALT00000022988_Cluster30955-001 |
| Cluster30961 | down | - | down | - | - | - | synaptic glyco SC2 [Zea mays | SALT00000051534_Cluster30961-004 |
| Cluster30962 | - | - | up | - | up | up | [Hordeum vulgare vulgare] | SALT00000064716_Cluster30962-003 |
| Cluster30974 | up | - | - | down | down | - | probable xyloglucan endotran | SALT00000043299_Cluster30974-002 |
| Cluster30979 | - | - | down | - | down | down | ferredoxin-NADP(H) oxidore | SALT00000059001_Cluster30979-003 |
| Cluster3098 | - | - | up | - | up | up | hypothetical protein F775_32 | SALT00000007388_Cluster3098-004 |
| Cluster30984 | - | down | down | - | - | - | omega-3 fatty acid desaturase | SALT00000046762_Cluster30984-002 |
| Cluster30986 | - | - | up | - | - | - | Transcription factor MYB39 | SALT00000048604_Cluster30986-001 |
| Cluster30988 | - | up | - | - | - | down | polygalacturonase inhibitor 1 | SALT00000060175_Cluster30988-002 |
| Cluster30995 | - | down | down | - | down | - | cytosolic sulfotransferase 8-lil | SALT00000068351_Cluster30995-001 |
| Cluster30999 | up | up | up | up | - | - | homeobox-leucine zipper ATI | SALT00000038825_Cluster30999-001 |
| Cluster31009 | up | up | up | - | down | down | cationic peroxidase SPC4-like | SALT00000002742_Cluster31009-001 |
| Cluster31018 | - | down | down | - | - | - | aquaporin [Oryza sativa Japor | SALT00000061667_Cluster31018-002 |
| Cluster3102 | - | - | - | - | down | - | Os03g0640800 [Oryza sativa | SALT00000078926_Cluster3102-017 |
| Cluster31021 | up | up | - | - | - | - | Os07g0569100 [Oryza sativa | SALT00000042651_Cluster31021-001 |
| Cluster31023 | - | - | down | - | down | down | oxygen-evolving enhancer 1 [ | SALT00000003619_Cluster31023-002 |
| Cluster31025 | - | - | up | - | up | up | Os03g0140700 [Oryza sativa | SALT00000036453_Cluster31025-001 |
| Cluster31029 | - | - | up | - | up | up | glycosyltransferase AGO61 | SALT00000049964_Cluster31029-001 |
| Cluster31031 | - | - | up | - | up | up | lichenase-2 precursor [Zea ma | SALT00000057265_Cluster31031-001 |
| Cluster31032 | - | - | up | - | up | up | AP2-EREBP transcription par | SALT00000059416_Cluster31032-003 |
| Cluster31036 | - | - | - | - | - | down | esterase precursor [Zea mays] | SALT00000015100_Cluster31036-001 |
| Cluster31040 | - | - | down | - | - | - | [Hordeum vulgare vulgare] | SALT00000038098_Cluster31040-001 |
| Cluster31057 | - | - | up | - | up | up | [Hordeum vulgare vulgare] | SALT00000019282_Cluster31057-003 |
| Cluster31062 | - | - | down | - | - | - | fatty acid 2-hydroxylase 1-lik | SALT00000044662_Cluster31062-002 |
| Cluster31066 | - | - | up | - | up | - | bifunctional nuclease 2 [Setar | SALT00000052922_Cluster31066-004 |
| Cluster31069 | - | - | - | - | down | - | FAR1-RELATED SEQUENC | SALT00000066659_Cluster31069-001 |
| Cluster31090 | - | - | down | - | - | - | Os10g0536700 [Oryza sativa | SALT00000050693_Cluster31090-001 |
| Cluster31099 | - | - | - | - | down | down | ferredoxin [Zea mays] | SALT00000012609_Cluster31099-002 |
| Cluster311 | - | - | up | - | up | - | [Hordeum vulgare vulgare] | SALT00000090343_Cluster311-014 |
| Cluster3110 | - | - | down | - | - | - | peptidyl-prolyl cis-trans isom | SALT00000081939_Cluster3110-005 |
| Cluster31117 | - | - | - | - | - | up | harpin-induced [Zea mays] | SALT00000057441_Cluster31117-001 |
| Cluster3112 | - | - | - | - | down | - | Os07g0150100 [Oryza sativa | SALT00000068329_Cluster3112-017 |
| Cluster31129 | - | - | down | - | down | down | TPA: protochlorophyllide red | SALT00000004801_Cluster31129-001 |
| Cluster31140 | - | down | down | down | down | down | fructose-bisphosphate cytopla | SALT00000003156_Cluster31140-001 |
| Cluster31144 | - | - | down | - | - | down | nitrilase cyanide hydratase an | SALT00000054421_Cluster31144-001 |
| Cluster31147 | - | down | down | - | - | - | malate chloroplastic-like | SALT00000051395_Cluster31147-001 |
| Cluster31166 | - | up | up | - | - | - | RING-H2 finger ATL5A [Zea | SALT00000051330_Cluster31166-001 |
| Cluster31168 | - | - | down | - | - | - | involucrin [Oryza sativa Japo | SALT00000005506_Cluster31168-001 |
| Cluster31170 | - | - | up | - | up | up | xylanase inhibitor 2-like | SALT00000037111_Cluster31170-001 |
| Cluster31173 | - | - | down | - | - | - | DNA polymerase zeta process | SALT00000044268_Cluster31173-001 |
| Cluster31186 | - | - | up | - | up | - | TPA: homeobox DNA-bindin | SALT00000010598_Cluster31186-001 |
| Cluster31188 | - | - | up | - | - | - | ubiquitin- ligase zinc ion bind | SALT00000007513_Cluster31188-003 |
| Cluster31194 | - | - | up | - | up | - | probable receptor kinase At5g | SALT00000044127_Cluster31194-001 |
| Cluster31195 | - | - | down | - | - | - | gibberellin 2-oxidase [Zea ma | SALT00000052373_Cluster31195-001 |
| Cluster31196 | - | - | down | - | down | down | ribosomal S8e family [Zea ma | SALT00000054268_Cluster31196-003 |
| Cluster312 | - | - | up | - | up | - | partial [Oryza sativa Japonica | SALT00000087240_Cluster312-001 |
| Cluster31205 | - | - | up | - | up | - | ---NA--- | SALT00000040527_Cluster31205-001 |
| Cluster31214 | - | - | - | - | down | - | glucuronoxylan 4-O-methyltr | SALT00000047633_Cluster31214-001 |
| Cluster31218 | - | - | up | - | up | up | PTI1-like tyrosine- kinase 3 | SALT00000052774_Cluster31218-001 |
| Cluster31223 | - | up | - | - | - | - | ---NA--- | SALT00000044648_Cluster31223-001 |
| Cluster31225 | - | - | up | - | - | - | Os06g0105800 [Oryza sativa | SALT00000046661_Cluster31225-001 |
| Cluster31229 | - | - | up | - | up | up | [Hordeum vulgare vulgare] | SALT00000037970_Cluster31229-001 |

| | | | | | | | | |
|---|---|---|---|---|---|---|---|---|
| Cluster3123 | - | - | up | - | up | up | OBERON 4 [Oryza sativa Jap | SALT00000083650_Cluster3123-002 |
| Cluster31230 | - | - | - | - | down | - | phospholipase [Oryza sativa J | SALT00000039207_Cluster31230-001 |
| Cluster31231 | up | up | - | - | - | - | Os07g0480900 [Oryza sativa | SALT00000058454_Cluster31231-001 |
| Cluster31232 | - | - | down | - | down | - | homeobox-leucine zipper HO | SALT00000060172_Cluster31232-002 |
| Cluster31235 | - | - | - | - | - | down | rhamnose biosynthetic enzym | SALT00000045692_Cluster31235-001 |
| Cluster31242 | up | up | up | - | - | - | adenosine kinase 2 | SALT00000004060_Cluster31242-002 |
| Cluster31244 | - | - | up | - | up | - | prosaposin-like [Setaria italica | SALT00000045591_Cluster31244-002 |
| Cluster31248 | - | - | up | - | - | - | chaperone dnaJ 6-like [Setaria | SALT00000017269_Cluster31248-001 |
| Cluster31251 | - | up | - | - | - | - | Os01g0192300 [Oryza sativa | SALT00000044866_Cluster31251-004 |
| Cluster31261 | - | - | up | - | up | up | Os08g0529300 [Oryza sativa | SALT00000053667_Cluster31261-002 |
| Cluster31264 | - | - | - | - | down | - | TPA: pheophorbide a oxygena | SALT00000061274_Cluster31264-001 |
| Cluster3127 | - | - | up | - | up | up | receptor kinase At3g47110 [S | SALT00000078047_Cluster3127-001 |
| Cluster31272 | - | - | down | - | - | - | Os06g0697000 [Oryza sativa | SALT00000036082_Cluster31272-002 |
| Cluster31279 | - | - | - | down | down | - | beta-expansin 4 precursor | SALT00000055066_Cluster31279-001 |
| Cluster31282 | - | - | down | - | - | - | Peroxidase 56 | SALT00000056992_Cluster31282-001 |
| Cluster31287 | - | - | down | - | - | - | [Oryza sativa Indica Group] | SALT00000067473_Cluster31287-001 |
| Cluster31288 | - | - | - | - | down | - | SRG1-like [Setaria italica] | SALT00000019464_Cluster31288-002 |
| Cluster31299 | - | - | up | - | up | - | alcohol dehydrogenase superf | SALT00000044488_Cluster31299-002 |
| Cluster31305 | - | - | up | - | up | up | NAD(P)H-dependent oxidore | SALT00000020185_Cluster31305-001 |
| Cluster31311 | - | - | down | - | - | down | photosystem II K (chloroplast | SALT00000008695_Cluster31311-001 |
| Cluster31313 | - | - | down | - | - | - | 2-oxoglutarate malate transloc | SALT00000016685_Cluster31313-001 |
| Cluster31315 | - | down | down | - | down | - | Os01g0730500 [Oryza sativa | SALT00000001883_Cluster31315-001 |
| Cluster3134 | - | - | down | - | down | - | Os09g0427125 [Oryza sativa | SALT00000088219_Cluster3134-001 |
| Cluster31351 | - | - | up | - | up | up | Os09g0433000 [Oryza sativa | SALT00000000915_Cluster31351-001 |
| Cluster31362 | - | - | down | - | - | - | peptidyl-prolyl cis-trans isom | SALT00000036819_Cluster31362-001 |
| Cluster31367 | - | - | up | - | - | - | partial [Hordeum vulgare vul | SALT00000008761_Cluster31367-001 |
| Cluster31375 | - | - | - | - | - | down | 2-Sep [Zea mays] | SALT00000003228_Cluster31375-001 |
| Cluster31381 | - | up | up | - | up | up | phosphoglycerate mutase AT7 | SALT00000064222_Cluster31381-002 |
| Cluster31382 | - | - | down | - | - | - | Os01g0739200 [Oryza sativa | SALT00000040560_Cluster31382-001 |
| Cluster31405 | - | - | - | - | - | down | aspartate-semialdehyde dehyd | SALT00000039973_Cluster31405-001 |
| Cluster31415 | up | - | - | - | down | down | Thiazole biosynthetic enzyme | SALT00000006791_Cluster31415-002 |
| Cluster31417 | - | - | up | - | up | up | Os09g0255400 [Oryza sativa | SALT00000009328_Cluster31417-001 |
| Cluster31419 | - | - | up | - | up | up | thaumatin 1 precursor [Zea m | SALT00000017083_Cluster31419-001 |
| Cluster31425 | - | down | down | - | down | - | peroxidase A2-like | SALT00000067671_Cluster31425-001 |
| Cluster31427 | - | - | up | - | - | - | glucan endo-1,3-beta-glucosic | SALT00000040387_Cluster31427-001 |
| Cluster31430 | - | up | down | - | down | down | Peroxidase 12 | SALT00000006864_Cluster31430-001 |
| Cluster31431 | - | down | down | - | down | - | GDSL esterase lipase At5g45 | SALT00000008940_Cluster31431-001 |
| Cluster31434 | - | up | up | - | up | up | shock [Oryza sativa Japonica | SALT00000039822_Cluster31434-002 |
| Cluster31448 | - | - | down | - | down | down | 60S ribosomal L5-1 [Zea may | SALT00000008155_Cluster31448-001 |
| Cluster31450 | - | - | up | - | - | - | O-methyltransferase ZRP4 | SALT00000049568_Cluster31450-001 |
| Cluster31451 | - | - | down | - | down | down | DNA-binding MNB1B [Zea n | SALT00000071610_Cluster31451-001 |
| Cluster31454 | - | - | up | - | up | up | Peroxidase 2 | SALT00000047340_Cluster31454-001 |
| Cluster31457 | - | - | down | - | down | down | GDSL esterase lipase APG-lik | SALT00000013269_Cluster31457-001 |
| Cluster31467 | - | - | up | - | - | - | PREDICTED: uncharacterize | SALT00000048965_Cluster31467-001 |
| Cluster3147 | - | - | - | - | down | - | [Oryza sativa Japonica Group | SALT00000065424_Cluster3147-003 |
| Cluster31475 | - | - | - | - | down | - | Os11g0539200 [Oryza sativa | SALT00000058506_Cluster31475-001 |
| Cluster31476 | - | up | - | - | down | down | peroxidase [Oryza sativa Japo | SALT00000007158_Cluster31476-001 |
| Cluster31477 | - | - | down | - | down | down | 60S acidic ribosomal P0 [Ory | SALT00000014364_Cluster31477-002 |
| Cluster31481 | - | up | - | - | - | - | alcohol dehydrogenase superf | SALT00000014141_Cluster31481-001 |
| Cluster31484 | - | up | up | - | up | up | anamorsin homolog [Zea may | SALT00000054087_Cluster31484-001 |
| Cluster31486 | - | - | up | - | up | - | Os04g0613300 [Oryza sativa | SALT00000060200_Cluster31486-002 |
| Cluster31490 | - | up | - | - | down | down | partial [Hordeum vulgare vul | SALT00000057132_Cluster31490-001 |
| Cluster31492 | - | - | down | - | - | - | quinone oxidoreductase At1g | SALT00000011385_Cluster31492-001 |
| Cluster31493 | - | - | - | - | - | up | Aquaporin PIP1-2 | SALT00000042179_Cluster31493-002 |
| Cluster31500 | - | up | up | up | up | up | ras GTPase-activating -bindin | SALT00000001825_Cluster31500-001 |
| Cluster31513 | - | - | up | - | up | - | Cyclopropane-fatty-acyl-phos | SALT00000004554_Cluster31513-001 |
| Cluster31518 | - | - | down | - | - | - | peroxidase [Oryza sativa Japo | SALT00000055516_Cluster31518-001 |
| Cluster31560 | - | - | down | - | down | - | peroxidase 2-like [Setaria itali | SALT00000037163_Cluster31560-001 |

| | | | | | | | | |
|---|---|---|---|---|---|---|---|---|
| Cluster31564 | - | - | up | - | up | up | PREDICTED: uncharacterize | SALT00000015639_Cluster31564-002 |
| Cluster31565 | - | - | down | - | down | down | xyloglucan endotransglycosyl | SALT00000044403_Cluster31565-003 |
| Cluster31568 | - | - | up | - | up | - | transcription factor bHLH30-l | SALT00000037556_Cluster31568-001 |
| Cluster3157 | - | - | - | - | down | - | retrotransposon unclassified | SALT00000085978_Cluster3157-001 |
| Cluster31570 | - | - | up | - | - | - | NAC domain-containing 68 [Z | SALT00000000770_Cluster31570-003 |
| Cluster31572 | - | down | down | down | down | - | peroxidase P7-like | SALT00000015586_Cluster31572-001 |
| Cluster31575 | up | up | up | - | - | - | cytosolic sulfotransferase 5-lil | SALT00000050165_Cluster31575-001 |
| Cluster31579 | - | - | - | - | - | down | multiple organellar RNA editi | SALT00000070085_Cluster31579-001 |
| Cluster3158 | - | down | - | - | - | - | Os04g0625800 [Oryza sativa | SALT00000087344_Cluster3158-016 |
| Cluster31580 | - | - | up | - | up | - | inosine-uridine nucleoside N-r | SALT00000009901_Cluster31580-001 |
| Cluster31590 | - | up | - | - | - | down | kDa proline-rich -like | SALT00000036290_Cluster31590-003 |
| Cluster31593 | - | - | up | - | up | - | catalytic hydrolase | SALT00000042645_Cluster31593-001 |
| Cluster31594 | up | up | up | - | up | - | plastid-lipid-associated 2 [Zea | SALT00000009055_Cluster31594-001 |
| Cluster31611 | - | - | up | - | up | up | WPP domain-interacting 1-lik | SALT00000005216_Cluster31611-001 |
| Cluster31612 | - | down | down | down | down | - | class III peroxidase [Triticum | SALT00000012022_Cluster31612-001 |
| Cluster31613 | - | - | up | - | - | up | annexin RJ4 [Zea mays] | SALT00000016576_Cluster31613-001 |
| Cluster31619 | - | - | down | - | down | down | CMV 1a interacting 1 [Zea ma | SALT00000004752_Cluster31619-001 |
| Cluster31627 | - | - | down | - | - | - | stress inducible | SALT00000044775_Cluster31627-002 |
| Cluster3163 | - | - | up | - | up | up | Os01g0110500 [Oryza sativa | SALT00000074178_Cluster3163-001 |
| Cluster31631 | - | down | down | - | - | - | GDSL esterase lipase EXL3 | SALT00000006200_Cluster31631-001 |
| Cluster31636 | - | - | down | - | - | - | E3 ubiquitin- ligase RING1-li | SALT00000051517_Cluster31636-002 |
| Cluster31638 | up | up | up | up | up | up | Os05g0529700 [Oryza sativa | SALT00000062522_Cluster31638-001 |
| Cluster31639 | - | - | down | - | down | down | xyloglucan endotransglycosyl | SALT00000039943_Cluster31639-002 |
| Cluster31642 | - | - | - | - | down | - | RING zinc finger domain sup | SALT00000054405_Cluster31642-001 |
| Cluster31646 | - | - | up | - | - | - | proteasome subunit alpha type | SALT00000057413_Cluster31646-001 |
| Cluster31648 | - | up | up | - | - | - | Os02g0829500 [Oryza sativa | SALT00000008671_Cluster31648-001 |
| Cluster31653 | - | - | down | - | - | - | mitochondrial outer membran | SALT00000060022_Cluster31653-002 |
| Cluster31665 | - | - | down | - | down | down | 40S ribosomal SA [Zea mays] | SALT00000017946_Cluster31665-001 |
| Cluster3167 | down | - | - | - | - | - | ---NA--- | SALT00000032724_Cluster3167-002 |
| Cluster31677 | - | - | up | - | - | - | prenylated Rab receptor 2 | SALT00000037662_Cluster31677-001 |
| Cluster31678 | - | - | up | - | up | up | [Hordeum vulgare vulgare] | SALT00000060586_Cluster31678-001 |
| Cluster31680 | - | - | up | - | - | - | Os12g0257500 [Oryza sativa | SALT00000056551_Cluster31680-001 |
| Cluster31683 | - | - | up | - | - | - | 14-3-3 GF14-D | SALT00000017866_Cluster31683-002 |
| Cluster31689 | - | up | up | - | - | - | chaperone dnaJ 6-like [Setaria | SALT00000019763_Cluster31689-001 |
| Cluster31707 | - | - | up | - | - | - | peroxidase 51 precursor [Zea | SALT00000015918_Cluster31707-001 |
| Cluster31713 | - | - | up | - | up | up | [Oryza sativa Japonica Group | SALT00000038419_Cluster31713-001 |
| Cluster31717 | - | - | - | - | - | down | cation transport chaC [Zea ma | SALT00000052113_Cluster31717-002 |
| Cluster31719 | - | - | - | - | down | down | ankyrin repeat domain-contain | SALT00000048993_Cluster31719-002 |
| Cluster31721 | - | up | up | - | up | up | probable low-specificity L-thr | SALT00000056029_Cluster31721-001 |
| Cluster31722 | - | - | - | - | down | - | 33 kDa chloroplastic | SALT00000068310_Cluster31722-001 |
| Cluster31726 | - | - | down | - | - | - | short-chain dehydrogenase re | SALT00000063602_Cluster31726-002 |
| Cluster31731 | up | up | up | - | - | - | prohibitin [Oryza sativa Japon | SALT00000045849_Cluster31731-001 |
| Cluster31736 | - | - | up | - | - | - | Bifunctional dihydroflavonol | SALT00000038057_Cluster31736-001 |
| Cluster31738 | up | up | up | - | - | - | nuclease I [Oryza sativa Japon | SALT00000060478_Cluster31738-001 |
| Cluster31741 | - | - | down | - | - | - | IAA31 - auxin-responsive Au | SALT00000018642_Cluster31741-001 |
| Cluster31750 | - | - | down | - | down | down | chlorophyll a-b binding 6A [Z | SALT00000010227_Cluster31750-002 |
| Cluster31755 | - | - | down | - | down | down | guanine nucleotide-binding be | SALT00000015909_Cluster31755-001 |
| Cluster31756 | - | - | up | - | - | - | CAPIP1 [Zea mays] | SALT00000038300_Cluster31756-001 |
| Cluster31758 | - | - | up | - | up | up | probable calcium-binding CM | SALT00000052088_Cluster31758-002 |
| Cluster31762 | - | - | up | - | up | up | ring-H2 zinc finger | SALT00000042998_Cluster31762-001 |
| Cluster31764 | - | - | down | - | - | - | peroxidase [Oryza sativa Japo | SALT00000050518_Cluster31764-001 |
| Cluster31781 | - | down | down | - | down | down | ---NA--- | SALT00000002468_Cluster31781-001 |
| Cluster31782 | - | - | up | - | up | up | SRC2 homolog | SALT00000013581_Cluster31782-001 |
| Cluster31794 | - | - | up | - | up | up | zinc-finger WZF1 [Triticum a | SALT00000018598_Cluster31794-002 |
| Cluster31797 | down | down | down | - | down | - | beta-expansin 1a precursor [Z | SALT00000007042_Cluster31797-001 |
| Cluster31798 | - | - | - | - | down | - | Secretory carrier-associated m | SALT00000012007_Cluster31798-001 |
| Cluster31803 | - | down | down | - | down | down | fasciclin-like arabinogalactan | SALT00000005564_Cluster31803-002 |
| Cluster31807 | - | up | up | up | up | - | homeobox-leucine zipper HO | SALT00000012123_Cluster31807-001 |

| | | | | | | | | |
|---|---|---|---|---|---|---|---|---|
| Cluster31809 | - | - | down | - | - | - | GDSL-motif lipase hydrolase- | SALT00000049345_Cluster31809-001 |
| Cluster31810 | - | - | up | - | up | up | acid phosphatase vanadium-d | SALT00000049661_Cluster31810-001 |
| Cluster31814 | - | up | - | - | - | - | annexin p35 [Zea mays] | SALT00000006632_Cluster31814-001 |
| Cluster3182 | - | - | - | - | up | - | probable GPI-anchored adhes | SALT00000033211_Cluster3182-002 |
| Cluster31829 | - | - | down | - | - | - | meiosis 5 | SALT00000052905_Cluster31829-002 |
| Cluster31830 | - | - | up | - | up | up | mu-crystallin [Zea mays] | SALT00000030687_Cluster31830-001 |
| Cluster31842 | - | - | up | - | up | - | probable carboxylesterase 15 | SALT00000051593_Cluster31842-001 |
| Cluster31844 | - | - | - | - | down | down | Os05g0488900 [Oryza sativa | SALT00000033688_Cluster31844-001 |
| Cluster31848 | - | - | up | - | up | up | Os05g0129300 [Oryza sativa | SALT00000010555_Cluster31848-001 |
| Cluster31849 | - | - | down | - | - | - | oxidoreductase [Zea mays] | SALT00000016517_Cluster31849-001 |
| Cluster31852 | - | - | up | - | up | up | probable trehalose-phosphate | SALT00000040110_Cluster31852-001 |
| Cluster31853 | - | down | down | down | down | - | peroxidase 2-like | SALT00000048458_Cluster31853-001 |
| Cluster31855 | - | - | up | - | up | - | Os05g0440050 [Oryza sativa | SALT00000040313_Cluster31855-001 |
| Cluster31858 | - | - | - | - | - | down | transcription factor TCP7-like | SALT00000066921_Cluster31858-001 |
| Cluster31868 | - | - | up | - | up | up | gibberellin 20-oxidase [Oryza | SALT00000013961_Cluster31868-001 |
| Cluster3187 | - | - | up | - | - | - | kelch repeat-containing F-box | SALT00000085819_Cluster3187-014 |
| Cluster31871 | - | - | down | - | - | - | [Hordeum vulgare vulgare] | SALT00000004762_Cluster31871-001 |
| Cluster31873 | - | - | up | - | up | up | zinc-finger WZF1 [Triticum a | SALT00000001517_Cluster31873-001 |
| Cluster31877 | up | up | - | - | down | down | ---NA--- | SALT00000054784_Cluster31877-002 |
| Cluster31880 | - | - | - | - | down | down | annexin D7-like | SALT00000035676_Cluster31880-002 |
| Cluster31885 | - | - | down | - | - | - | Syntaxin-related KNOLLE [S | SALT00000053949_Cluster31885-001 |
| Cluster31889 | - | - | up | - | up | - | pectinesterase 31 | SALT00000011365_Cluster31889-001 |
| Cluster31890 | - | up | - | - | - | - | (+)-neomenthol dehydrogenas | SALT00000023579_Cluster31890-001 |
| Cluster31891 | - | up | up | - | up | - | PLATZ transcription partial [Z | SALT00000037835_Cluster31891-001 |
| Cluster31902 | - | - | up | - | up | - | fiber Fb19 [Zea mays] | SALT00000061029_Cluster31902-001 |
| Cluster31906 | - | - | up | - | up | - | Os01g0528800 [Oryza sativa | SALT00000012608_Cluster31906-001 |
| Cluster31909 | up | - | - | down | down | - | Os05g0477600 [Oryza sativa | SALT00000015003_Cluster31909-001 |
| Cluster31912 | - | - | up | - | - | - | NAD(P)H-dependent oxidore | SALT00000005950_Cluster31912-001 |
| Cluster31914 | - | - | down | - | down | - | partial [Oryza sativa Japonica | SALT00000060511_Cluster31914-001 |
| Cluster31916 | - | - | - | - | - | up | probable purine permease 11 [ | SALT00000041372_Cluster31916-001 |
| Cluster31919 | - | - | - | - | down | - | brown planthopper-induced re | SALT00000045924_Cluster31919-001 |
| Cluster31920 | - | - | - | - | down | - | stem-specific TSJT1 [Oryza sa | SALT00000057469_Cluster31920-002 |
| Cluster31922 | - | - | down | down | down | down | response regulator 7 [Zea may | SALT00000049254_Cluster31922-001 |
| Cluster31925 | - | - | down | - | down | down | chloroplastic isoform X1 [Bra | SALT00000039994_Cluster31925-001 |
| Cluster31926 | - | - | - | - | - | down | translation initiation factor IF- | SALT00000045450_Cluster31926-001 |
| Cluster31928 | - | - | down | - | down | down | 2-cys peroxiredoxin BAS1 [Z | SALT00000042139_Cluster31928-003 |
| Cluster31932 | - | up | up | up | up | up | [Hordeum vulgare vulgare] | SALT00000001568_Cluster31932-001 |
| Cluster31934 | - | - | up | - | up | up | senescence-associated [Zea m | SALT00000043709_Cluster31934-001 |
| Cluster31938 | - | - | down | - | - | - | outer plastidial membrane por | SALT00000006209_Cluster31938-001 |
| Cluster31939 | - | - | - | - | up | up | bacterial-induced peroxidase p | SALT00000051649_Cluster31939-003 |
| Cluster31941 | - | - | down | - | - | - | ubiquinone biosynthesis COQ | SALT00000043939_Cluster31941-001 |
| Cluster31945 | - | - | up | - | - | - | vesicle-associated membrane | SALT00000036919_Cluster31945-001 |
| Cluster31948 | - | - | up | - | - | - | vegetative cell wall gp1-like | SALT00000083206_Cluster31948-001 |
| Cluster31949 | - | down | down | down | down | - | peroxidase A2-like [Setaria ita | SALT00000013314_Cluster31949-001 |
| Cluster3195 | - | up | up | - | up | up | citrate synthase family | SALT00000008814_Cluster3195-003 |
| Cluster31954 | - | - | up | - | - | - | cell death associated [Oryza sa | SALT00000018682_Cluster31954-001 |
| Cluster31957 | - | - | down | - | - | - | SC3 [Zea mays] | SALT00000006235_Cluster31957-001 |
| Cluster31959 | - | - | down | - | down | - | ribonucleo A [Zea mays] | SALT00000052972_Cluster31959-001 |
| Cluster31963 | - | - | up | - | up | - | ribosomal [Oryza sativa Japor | SALT00000045402_Cluster31963-002 |
| Cluster31964 | down | down | down | - | - | - | TPA: general regulatory factor | SALT00000048214_Cluster31964-005 |
| Cluster31965 | - | - | up | up | up | up | probable cytokinin riboside 5 | SALT00000004482_Cluster31965-001 |
| Cluster31966 | - | - | down | - | - | - | ROP family GTPase ROP9 [Z | SALT00000039131_Cluster31966-001 |
| Cluster31983 | - | - | up | - | - | - | Os03g0270000 [Oryza sativa | SALT00000046238_Cluster31983-002 |
| Cluster31984 | - | up | up | - | up | - | thioredoxin-like 1- chloroplas | SALT00000058709_Cluster31984-002 |
| Cluster31986 | - | - | down | - | - | - | POU class transcription factor | SALT00000044871_Cluster31986-001 |
| Cluster31987 | - | - | up | - | up | up | Os07g0551400 [Oryza sativa | SALT00000050007_Cluster31987-001 |
| Cluster31991 | - | up | up | - | - | - | Os01g0621900 [Oryza sativa | SALT00000072968_Cluster31991-001 |
| Cluster31993 | - | - | - | - | down | - | Os10g0555700 [Oryza sativa | SALT00000048379_Cluster31993-002 |

| | | | | | | | | |
|---|---|---|---|---|---|---|---|---|
| Cluster32000 | - | - | up | - | up | up | hypothetical protein OsJ_085 | SALT00000007913_Cluster32000-001 |
| Cluster32003 | - | - | - | - | - | down | transcription initiation factor | SALT00000044104_Cluster32003-001 |
| Cluster32004 | - | - | up | - | up | up | lactoylglutathione lyase [Zea | SALT00000052783_Cluster32004-001 |
| Cluster32008 | - | - | up | - | up | up | chitinase [Oryza sativa Japoni | SALT00000013551_Cluster32008-001 |
| Cluster32011 | - | - | up | - | up | - | Os03g0402800 [Oryza sativa | SALT00000017931_Cluster32011-001 |
| Cluster32012 | - | - | - | - | - | down | endo-1,3-beta-glucanase [Ory | SALT00000045539_Cluster32012-001 |
| Cluster32016 | up | up | up | - | up | - | [Hordeum vulgare vulgare] | SALT00000052417_Cluster32016-001 |
| Cluster32018 | - | - | - | - | down | - | Os04g0166600 [Oryza sativa | SALT00000006895_Cluster32018-001 |
| Cluster32020 | - | down | - | - | - | - | RNA-binding expressed [Ory | SALT00000027187_Cluster32020-001 |
| Cluster32021 | - | - | up | - | - | - | electron transporter thiol-disu | SALT00000054024_Cluster32021-001 |
| Cluster32034 | - | - | - | - | up | - | cyclin-P4-1-like [Oryza sativa | SALT00000048630_Cluster32034-001 |
| Cluster32038 | - | - | up | - | up | - | Os05g0151300 [Oryza sativa | SALT00000035945_Cluster32038-001 |
| Cluster32051 | - | - | up | - | up | up | WUSCHEL-related homeobo | SALT00000046391_Cluster32051-001 |
| Cluster32052 | - | - | down | - | down | down | dihydroflavonol-4-reductase [ | SALT00000060681_Cluster32052-001 |
| Cluster32059 | - | - | down | - | down | down | UPF0603 chloroplastic | SALT00000005174_Cluster32059-001 |
| Cluster32061 | - | down | down | - | down | - | phosphoglycerate kinase [Zea | SALT00000056372_Cluster32061-001 |
| Cluster32066 | - | - | down | - | down | down | ATP synthase | SALT00000013273_Cluster32066-001 |
| Cluster32070 | - | - | up | - | up | - | [Hordeum vulgare vulgare] | SALT00000071539_Cluster32070-001 |
| Cluster32075 | - | - | down | - | down | - | xyloglucan endotransglucosyl | SALT00000003173_Cluster32075-001 |
| Cluster32079 | - | - | down | - | - | - | fibrillarin [Oryza sativa Japon | SALT00000021722_Cluster32079-001 |
| Cluster32088 | - | - | - | - | down | - | Os03g0807800 [Oryza sativa | SALT00000054939_Cluster32088-002 |
| Cluster32090 | - | - | up | - | up | up | extradiol ring-cleavage dioxy | SALT00000058152_Cluster32090-001 |
| Cluster32092 | - | - | - | - | down | down | ---NA--- | SALT00000036761_Cluster32092-001 |
| Cluster32093 | - | - | up | - | up | up | indole-3-glycerol phosphate l | SALT00000060545_Cluster32093-001 |
| Cluster32095 | - | - | up | - | up | up | peroxidase 4-like isoform X2 | SALT00000058490_Cluster32095-001 |
| Cluster32098 | - | - | up | - | up | up | zinc finger CCCH domain-co | SALT00000051021_Cluster32098-001 |
| Cluster32108 | up | - | - | - | down | down | uncharacterized protein LOC1 | SALT00000050231_Cluster32108-001 |
| Cluster32112 | - | down | down | - | - | - | DREPP2 [Oryza sativa Japoni | SALT00000042368_Cluster32112-001 |
| Cluster32119 | - | - | down | - | down | down | 40S ribosomal S4 [Oryza sati | SALT00000038388_Cluster32119-001 |
| Cluster32132 | - | - | up | - | - | - | ---NA--- | SALT00000045563_Cluster32132-002 |
| Cluster32134 | - | up | up | - | up | - | uncharacterized protein LOC1 | SALT00000044990_Cluster32134-001 |
| Cluster32140 | - | - | down | - | down | down | chlorophyll a-b binding 4 [Ze | SALT00000036408_Cluster32140-001 |
| Cluster32144 | - | - | up | - | up | up | ATG1 [Zea mays] | SALT00000062316_Cluster32144-002 |
| Cluster32157 | - | - | up | - | up | up | TPA: ras ARA-3 [Zea mays] | SALT00000056544_Cluster32157-002 |
| Cluster32159 | - | - | down | - | - | - | peptidyl-prolyl cis-trans isom | SALT00000072544_Cluster32159-001 |
| Cluster32162 | - | - | - | - | - | down | esterase D [Zea mays] | SALT00000039902_Cluster32162-001 |
| Cluster32165 | - | - | down | - | - | - | translation factor-like [Oryza | SALT00000058373_Cluster32165-001 |
| Cluster32167 | up | up | up | - | up | - | REHY_WHEAT ame: Full=1 | SALT00000045912_Cluster32167-002 |
| Cluster3217 | - | - | down | - | down | down | pentatricopeptide repeat-conta | SALT00000092726_Cluster3217-001 |
| Cluster32176 | - | down | down | - | - | - | phi-1-like phosphate-induced | SALT00000019317_Cluster32176-001 |
| Cluster32177 | - | down | down | - | - | - | [Hordeum vulgare vulgare] | SALT00000035767_Cluster32177-001 |
| Cluster32179 | - | - | up | - | up | up | Os09g0432900 [Oryza sativa | SALT00000016352_Cluster32179-001 |
| Cluster32181 | - | - | up | - | up | up | [Hordeum vulgare vulgare] | SALT00000019328_Cluster32181-001 |
| Cluster32182 | - | - | - | - | down | - | Os02g0715300 [Oryza sativa | SALT00000037093_Cluster32182-001 |
| Cluster3219 | - | - | up | - | - | - | Os10g0473400 [Oryza sativa | SALT00000085466_Cluster3219-008 |
| Cluster32193 | - | - | down | - | down | down | chlorophyll a-b binding 2 [Ze | SALT00000047761_Cluster32193-006 |
| Cluster32197 | up | - | down | down | down | down | VAMP SEC22 [Zea mays] | SALT00000037162_Cluster32197-001 |
| Cluster322 | - | - | up | - | up | up | T-complex 11 1 | SALT00000091193_Cluster322-024 |
| Cluster32202 | - | - | down | - | down | - | Os02g0742200 [Oryza sativa | SALT00000056271_Cluster32202-002 |
| Cluster32203 | - | - | up | - | up | - | Os03g0307000 [Oryza sativa | SALT00000002866_Cluster32203-001 |
| Cluster32206 | - | - | up | - | up | - | versicolorin reductase [Zea m | SALT00000048698_Cluster32206-001 |
| Cluster32210 | - | - | down | - | down | down | 60S ribosomal L2 [Zea mays] | SALT00000051551_Cluster32210-004 |
| Cluster32211 | - | - | down | - | down | - | carbonyl reductase 3 [Zea ma | SALT00000058380_Cluster32211-001 |
| Cluster32212 | - | - | down | - | down | down | ---NA--- | SALT00000058431_Cluster32212-001 |
| Cluster32217 | - | - | down | - | - | - | ATP-dependent Clp protease | SALT00000075991_Cluster32217-001 |
| Cluster32218 | - | - | up | - | up | - | transmembrane BAX inhibito | SALT00000015241_Cluster32218-003 |
| Cluster32221 | - | - | down | - | down | down | cytochrome b6-f complex iro | SALT00000016353_Cluster32221-001 |
| Cluster32223 | - | down | down | down | down | down | ---NA--- | SALT00000001815_Cluster32223-001 |

| | | | | | | | | |
|---|---|---|---|---|---|---|---|---|
| Cluster32226 | - | - | down | - | down | down | TPA: 40S ribosomal S6 [Zea r | SALT00000059643_Cluster32226-002 |
| Cluster32235 | - | - | down | - | down | down | 40S ribosomal S3a [Zea mays | SALT00000061160_Cluster32235-002 |
| Cluster32240 | - | - | up | - | - | - | SRC2 homolog | SALT00000063965_Cluster32240-002 |
| Cluster32242 | - | up | - | - | - | - | ---NA--- | SALT00000013343_Cluster32242-002 |
| Cluster32243 | - | - | down | - | down | - | Os06g0705100 [Oryza sativa | SALT00000046551_Cluster32243-001 |
| Cluster32244 | up | - | - | - | - | - | Os03g0360600 [Oryza sativa | SALT00000047142_Cluster32244-001 |
| Cluster32246 | - | down | down | - | down | - | ---NA--- | SALT00000029924_Cluster32246-001 |
| Cluster32248 | - | - | - | - | down | down | tyrosine specific phosphatase | SALT00000041879_Cluster32248-001 |
| Cluster32256 | - | - | up | - | - | - | MTD1 [Zea mays] | SALT00000042632_Cluster32256-001 |
| Cluster3226 | - | - | down | - | - | - | IQ-DOMAIN 1-like [Setaria i | SALT00000036141_Cluster3226-014 |
| Cluster32265 | - | - | down | - | down | down | Os07g0141400 [Oryza sativa | SALT00000061454_Cluster32265-002 |
| Cluster32267 | up | up | up | - | up | - | caffeoyl- O-methyltransferase | SALT00000013726_Cluster32267-002 |
| Cluster32276 | - | - | down | - | down | down | chlorophyll a-b binding 8 [Zea | SALT00000017035_Cluster32276-001 |
| Cluster32278 | - | - | down | - | down | down | chlorophyll a b-binding apo C | SALT00000039876_Cluster32278-001 |
| Cluster32280 | down | down | down | - | - | - | xyloglucan endotransglycosyl | SALT00000064836_Cluster32280-002 |
| Cluster32286 | - | - | down | - | - | - | expansin-B3-like [Setaria itali | SALT00000054657_Cluster32286-001 |
| Cluster32296 | - | - | up | - | - | - | Os02g0810300 [Oryza sativa | SALT00000016468_Cluster32296-001 |
| Cluster32305 | - | - | down | - | down | down | ---NA--- | SALT00000013746_Cluster32305-001 |
| Cluster32317 | - | up | - | up | - | down | ---NA--- | SALT00000051729_Cluster32317-002 |
| Cluster32323 | - | - | down | - | down | down | 60S ribosomal L7a [Oryza sat | SALT00000060282_Cluster32323-001 |
| Cluster32324 | - | - | up | - | up | up | ferritin 1A [Triticum aestivum | SALT00000002116_Cluster32324-001 |
| Cluster32325 | - | - | up | - | - | up | cysteine-rich repeat secretory | SALT00000042028_Cluster32325-001 |
| Cluster32326 | - | - | up | - | - | - | GTP-binding YPTM2 [Oryza | SALT00000053207_Cluster32326-001 |
| Cluster32330 | - | - | down | - | down | - | tonoplast intrinsic [Oryza sati | SALT00000018152_Cluster32330-001 |
| Cluster32331 | - | - | down | - | down | down | chlorophyll a-b binding CP24 | SALT00000019685_Cluster32331-001 |
| Cluster32334 | - | - | up | - | up | - | small heat shock [Zea mays] | SALT00000054823_Cluster32334-001 |
| Cluster32336 | - | down | down | - | down | down | chlorophyll a-b binding of LH | SALT00000054350_Cluster32336-002 |
| Cluster32339 | - | - | - | - | down | - | ---NA--- | SALT00000065147_Cluster32339-001 |
| Cluster32345 | - | up | up | - | up | - | Os12g0569300 [Oryza sativa | SALT00000045646_Cluster32345-001 |
| Cluster32350 | - | - | - | - | - | down | retrotransposon Ty1-copia sub | SALT00000059760_Cluster32350-001 |
| Cluster32351 | - | - | down | - | down | down | chlorophyll a-b binding [Zea r | SALT00000068437_Cluster32351-002 |
| Cluster32354 | - | up | - | - | - | down | dehydroascorbate reductase [( | SALT00000058577_Cluster32354-001 |
| Cluster32356 | - | up | - | - | - | - | superoxide dismutase [Mn] m | SALT00000053866_Cluster32356-002 |
| Cluster32359 | - | - | up | - | - | - | beta 7 subunit of 20S proteasc | SALT00000017935_Cluster32359-001 |
| Cluster32364 | - | - | - | - | - | down | urease accessory ureG [Zea m | SALT00000054161_Cluster32364-001 |
| Cluster32369 | - | - | up | - | up | up | VAMP SEC22 [Zea mays] | SALT00000069244_Cluster32369-001 |
| Cluster32375 | - | - | up | - | - | - | ---NA--- | SALT00000021834_Cluster32375-001 |
| Cluster32376 | - | - | down | - | - | - | [Hordeum vulgare vulgare] | SALT00000055549_Cluster32376-001 |
| Cluster3238 | - | - | - | - | down | - | photosystem I assembly ycf4 ( | SALT00000054238_Cluster3238-001 |
| Cluster32380 | up | up | up | up | up | - | Os05g0435300 [Oryza sativa | SALT00000064731_Cluster32380-001 |
| Cluster32388 | - | - | down | - | down | down | VAMP SEC22 [Zea mays] | SALT00000076337_Cluster32388-002 |
| Cluster32391 | - | - | down | - | down | down | Os08g0407600 [Oryza sativa | SALT00000043024_Cluster32391-001 |
| Cluster32402 | - | - | up | - | - | - | [Hordeum vulgare vulgare] | SALT00000068783_Cluster32402-001 |
| Cluster32410 | - | - | down | - | down | down | 60S ribosomal L15 [Zea mays | SALT00000039763_Cluster32410-001 |
| Cluster32412 | - | - | up | - | - | - | ethylene-responsive transcript | SALT00000071723_Cluster32412-001 |
| Cluster32413 | - | - | up | - | - | - | transcription factor IBH1 [Bra | SALT00000037891_Cluster32413-001 |
| Cluster32415 | - | - | down | - | down | - | ---NA--- | SALT00000037661_Cluster32415-001 |
| Cluster32418 | - | - | down | - | down | down | photosystem I reaction center | SALT00000073844_Cluster32418-002 |
| Cluster32420 | - | down | down | down | down | - | arabinogalactan -like [Oryza s | SALT00000039339_Cluster32420-001 |
| Cluster32421 | - | - | down | - | down | down | [Hordeum vulgare vulgare] | SALT00000069235_Cluster32421-001 |
| Cluster32424 | - | - | up | - | - | - | caleosin related [Zea mays] | SALT00000046441_Cluster32424-001 |
| Cluster32431 | - | - | down | - | down | down | rhodanese-like family [Zea ma | SALT00000040711_Cluster32431-001 |
| Cluster32432 | - | - | down | - | down | down | [Hordeum vulgare vulgare] | SALT00000053772_Cluster32432-001 |
| Cluster32433 | - | - | down | - | down | - | PREDICTED: uncharacterized | SALT00000057069_Cluster32433-001 |
| Cluster32435 | - | - | down | - | down | down | chloroplastic isoform X1 [Bra | SALT00000072858_Cluster32435-001 |
| Cluster32436 | - | - | down | - | - | - | cp [Zea mays] | SALT00000058249_Cluster32436-001 |
| Cluster32439 | - | - | up | - | up | up | EF hand family expressed [Or | SALT00000036897_Cluster32439-001 |
| Cluster32440 | - | - | up | - | - | - | harpin inducing [Zea mays] | SALT00000019704_Cluster32440-001 |

| | | | | | | | | |
|---|---|---|---|---|---|---|---|---|
| Cluster32443 | - | - | down | - | - | - | ---NA--- | SALT00000077438_Cluster32443-001 |
| Cluster32447 | - | - | up | - | up | up | salicylic acid-binding 2-like | SALT00000041627_Cluster32447-001 |
| Cluster32457 | - | - | down | - | down | down | 50S ribosomal chloroplastic [I | SALT00000060120_Cluster32457-001 |
| Cluster32458 | - | - | - | - | down | down | [Hordeum vulgare vulgare] | SALT00000053443_Cluster32458-002 |
| Cluster32459 | - | up | - | - | - | down | histone H1 | SALT00000073106_Cluster32459-001 |
| Cluster3246 | - | - | down | - | - | - | downstream neighbor of Son i | SALT00000030602_Cluster3246-002 |
| Cluster32461 | - | up | up | - | up | - | Os01g0853000 [Oryza sativa | SALT00000058270_Cluster32461-001 |
| Cluster32462 | - | - | down | - | - | - | Os12g0534200 [Oryza sativa | SALT00000019797_Cluster32462-001 |
| Cluster32470 | - | - | - | - | down | down | Os03g0161800 [Oryza sativa | SALT00000016357_Cluster32470-001 |
| Cluster32471 | - | - | - | - | up | - | [Hordeum vulgare vulgare] | SALT00000037687_Cluster32471-001 |
| Cluster32481 | - | - | up | - | up | up | cystatin [Zea mays] | SALT00000060980_Cluster32481-001 |
| Cluster32482 | - | up | - | - | down | down | ---NA--- | SALT00000060729_Cluster32482-001 |
| Cluster32484 | - | - | down | - | down | down | Os08g0560900 [Oryza sativa | SALT00000025823_Cluster32484-001 |
| Cluster32492 | - | - | down | - | - | - | eukaryotic translation initiatio | SALT00000027293_Cluster32492-002 |
| Cluster32498 | - | - | - | - | - | down | co-chaperone SBA1 [Zea may | SALT00000059917_Cluster32498-002 |
| Cluster32505 | - | - | down | - | down | down | Os03g0337800 [Oryza sativa | SALT00000022573_Cluster32505-001 |
| Cluster32506 | - | - | up | - | up | up | Gb: [Theobroma cacao] | SALT00000023645_Cluster32506-001 |
| Cluster32507 | - | - | down | - | down | down | 60S ribosomal L9 [Zea mays] | SALT00000056427_Cluster32507-001 |
| Cluster32508 | - | - | up | - | up | - | SKP1 1B | SALT00000064157_Cluster32508-001 |
| Cluster32513 | - | - | up | - | up | up | polcalcin Jun o 2 [Zea mays] | SALT00000040980_Cluster32513-001 |
| Cluster32516 | - | - | down | - | down | - | 60S ribosomal L6 [Zea mays] | SALT00000058795_Cluster32516-001 |
| Cluster32522 | - | - | - | - | up | up | vegetative cell wall gp1-like | SALT00000062072_Cluster32522-001 |
| Cluster32523 | - | - | up | - | up | up | WRKY55 - superfamily of TF | SALT00000065826_Cluster32523-001 |
| Cluster32525 | - | - | up | - | - | - | vacuolar ATP synthase 16 kDa | SALT00000044009_Cluster32525-001 |
| Cluster32530 | - | - | - | - | down | - | cyclophilin [Saccharum hybri | SALT00000058756_Cluster32530-003 |
| Cluster32533 | - | - | up | - | up | up | thionin-like [Setaria italica] | SALT00000071566_Cluster32533-001 |
| Cluster32536 | - | - | up | - | - | - | Os05g0451300 [Oryza sativa | SALT00000020136_Cluster32536-001 |
| Cluster32537 | - | - | up | - | - | - | [Hordeum vulgare vulgare] | SALT00000026868_Cluster32537-001 |
| Cluster32538 | - | - | - | - | down | - | Os12g0632000 [Oryza sativa | SALT00000066419_Cluster32538-001 |
| Cluster32540 | - | - | down | - | down | - | [Hordeum vulgare vulgare] | SALT00000057023_Cluster32540-001 |
| Cluster32543 | - | - | - | - | down | - | AChain Structure Of Glutathi | SALT00000057491_Cluster32543-001 |
| Cluster32544 | - | - | down | - | down | down | PSBQ2_MAIZE ame: Full=O | SALT00000065116_Cluster32544-001 |
| Cluster32547 | up | up | up | - | up | - | kDa class I heat shock | SALT00000023106_Cluster32547-001 |
| Cluster32552 | - | - | down | - | down | - | cortical cell-delineating precu | SALT00000080454_Cluster32552-001 |
| Cluster32556 | - | - | down | - | down | down | cell cycle switch [Oryza sativa | SALT00000057268_Cluster32556-001 |
| Cluster32557 | - | - | down | - | down | down | 60S ribosomal L18a [Zea may | SALT00000026032_Cluster32557-001 |
| Cluster32559 | - | - | down | - | down | down | photosynthetic NDH subunit c | SALT00000053316_Cluster32559-001 |
| Cluster32564 | - | - | down | - | down | down | cortical cell-delineating precu | SALT00000065391_Cluster32564-001 |
| Cluster32565 | - | - | down | - | down | down | 40S ribosomal S11 [Oryza sat | SALT00000042680_Cluster32565-001 |
| Cluster32567 | - | - | down | - | - | - | photosynthetic NDH subunit c | SALT00000073593_Cluster32567-001 |
| Cluster32572 | - | - | - | - | down | - | peptidyl-prolyl cis-trans isome | SALT00000077659_Cluster32572-001 |
| Cluster32574 | - | - | - | - | down | - | non-specific lipid-transfer 1 [( | SALT00000016631_Cluster32574-001 |
| Cluster32576 | - | - | down | - | down | down | 40S ribosomal S17-4 [Zea ma | SALT00000010297_Cluster32576-001 |
| Cluster3258 | - | down | down | - | - | - | sucrose phosphate synthase | SALT00000086109_Cluster3258-012 |
| Cluster32580 | - | - | down | - | down | - | 60S ribosomal L27 [Zea mays | SALT00000045952_Cluster32580-001 |
| Cluster32582 | - | down | - | - | - | - | histone H3 [Zea mays] | SALT00000013949_Cluster32582-001 |
| Cluster32583 | - | - | up | - | up | up | Os01g0888900 [Oryza sativa | SALT00000042452_Cluster32583-001 |
| Cluster32588 | up | up | up | - | up | - | hydrophobic LTI6B [Zea may | SALT00000042942_Cluster32588-001 |
| Cluster32589 | - | - | - | down | - | - | ATPase inhibitor [Zea mays] | SALT00000010121_Cluster32589-001 |
| Cluster32590 | - | - | down | - | - | - | Os05g0178300 [Oryza sativa | SALT00000060823_Cluster32590-001 |
| Cluster32591 | - | - | down | - | - | - | Os03g0288000 [Oryza sativa | SALT00000068970_Cluster32591-001 |
| Cluster32596 | - | - | down | - | down | - | B12D [Zea mays] | SALT00000049815_Cluster32596-001 |
| Cluster32597 | - | - | up | - | up | up | Os02g0209300 [Oryza sativa | SALT00000019187_Cluster32597-001 |
| Cluster3266 | up | up | up | - | - | - | phosphatase 2C | SALT00000078123_Cluster3266-001 |
| Cluster3277 | - | - | up | - | up | - | LOV domain-containing | SALT00000080035_Cluster3277-004 |
| Cluster3310 | down | down | down | - | - | - | ribosomal L2 (chloroplast) [A | SALT00000088769_Cluster3310-016 |
| Cluster3317 | - | - | up | - | up | up | carbohydrate transporter sugar | SALT00000046161_Cluster3317-017 |
| Cluster3323 | - | - | up | - | up | up | lipoxygenase [Oryza sativa Ja | SALT00000081993_Cluster3323-006 |

| | | | | | | | | |
|---|---|---|---|---|---|---|---|---|
| Cluster3350 | - | - | - | - | down | down | NADH dehydrogenase subuni | SALT00000085497_Cluster3350-006 |
| Cluster339 | - | - | down | - | down | down | 60S ribosomal L7-1 [Zea may | SALT00000092917_Cluster339-002 |
| Cluster3399 | - | - | - | - | - | down | Retrovirus-related Pol poly LI | SALT00000089342_Cluster3399-001 |
| Cluster3403 | - | - | up | - | up | up | phosphatidate phosphatase PA | SALT00000078956_Cluster3403-013 |
| Cluster3405 | - | up | up | - | up | - | transaldolase 2 [Zea mays] | SALT00000049356_Cluster3405-013 |
| Cluster3422 | - | - | - | - | down | down | [Hordeum vulgare vulgare] | SALT00000069549_Cluster3422-012 |
| Cluster3424 | - | - | down | - | down | - | retrotransposon unclassified | SALT00000063493_Cluster3424-001 |
| Cluster3433 | up | - | - | - | - | - | ABC transporter G family me | SALT00000085644_Cluster3433-001 |
| Cluster3448 | up | up | up | - | up | up | lichenase-2 precursor [Zea ma | SALT00000089785_Cluster3448-014 |
| Cluster3464 | - | - | down | - | down | - | mitogen activated kinase | SALT00000075007_Cluster3464-001 |
| Cluster3467 | - | - | down | - | - | - | PREDICTED: uncharacterize | SALT00000090288_Cluster3467-001 |
| Cluster348 | - | - | down | - | - | - | cytochrome c biogenesis Fn (1 | SALT00000091163_Cluster348-034 |
| Cluster3480 | - | - | down | - | down | down | cellulose synthase-4 [Oryza sa | SALT00000091395_Cluster3480-001 |
| Cluster349 | - | - | down | - | down | down | monosaccharide-sensing 2-lik | SALT00000067824_Cluster349-001 |
| Cluster3494 | up | up | up | - | - | - | LOC100282635 isoform X1 [ | SALT00000046986_Cluster3494-007 |
| Cluster3495 | - | - | up | - | up | up | cysteine protease 1 precursor | SALT00000051381_Cluster3495-021 |
| Cluster3496 | - | up | up | - | - | - | pyruvate cytosolic isozyme [Z | SALT00000073671_Cluster3496-028 |
| Cluster3538 | - | - | - | - | down | down | glycosyltransferase family 64 | SALT00000089820_Cluster3538-001 |
| Cluster3540 | up | - | - | - | down | - | serine hydroxymethyltransfera | SALT00000052685_Cluster3540-001 |
| Cluster3543 | - | - | down | - | down | down | COBRA 1 | SALT00000034449_Cluster3543-001 |
| Cluster3545 | - | - | up | - | up | up | sucrose synthase 1 [Oryza sati | SALT00000069724_Cluster3545-005 |
| Cluster3551 | - | - | up | - | up | up | bZIP family transcription fact | SALT00000085178_Cluster3551-005 |
| Cluster3571 | - | - | - | - | down | - | Homeobox-leucine zipper HO | SALT00000071669_Cluster3571-003 |
| Cluster3574 | - | up | - | - | - | - | aspartate kinase [Oryza sativa | SALT00000091375_Cluster3574-004 |
| Cluster358 | - | - | - | - | - | down | ---NA--- | SALT00000045796_Cluster358-003 |
| Cluster3583 | - | - | - | - | down | down | Os08g0345700 [Oryza sativa | SALT00000061141_Cluster3583-001 |
| Cluster3585 | up | up | - | - | - | - | hypothetical protein SORBI_( | SALT00000048015_Cluster3585-001 |
| Cluster36 | - | - | up | - | up | up | Os06g0131300 [Oryza sativa | SALT00000070211_Cluster36-011 |
| Cluster3601 | - | up | up | - | up | - | delta-1-pyrroline-5-carboxyla | SALT00000087469_Cluster3601-026 |
| Cluster3610 | - | - | down | - | - | down | maturase K (chloroplast) | SALT00000029170_Cluster3610-011 |
| Cluster362 | - | - | down | - | - | - | filament-like plant 4 [Setaria i | SALT00000032017_Cluster362-008 |
| Cluster3620 | - | - | - | - | down | - | PREDICTED: uncharacterize | SALT00000090979_Cluster3620-008 |
| Cluster3624 | - | - | - | - | - | down | E3 ubiquitin- ligase BRE1-lik | SALT00000086670_Cluster3624-006 |
| Cluster3628 | - | - | up | - | up | - | remorin [Oryza sativa Japonic | SALT00000008200_Cluster3628-001 |
| Cluster3636 | - | - | - | - | down | down | probable ethylene response se | SALT00000082017_Cluster3636-001 |
| Cluster364 | - | - | down | - | down | - | cellulose synthase1 | SALT00000085874_Cluster364-092 |
| Cluster3640 | - | - | down | - | - | down | RNA polymerase alpha subun | SALT00000089836_Cluster3640-015 |
| Cluster3645 | - | - | up | - | - | - | Avr9 Cf-9 rapidly elicited -lik | SALT00000081541_Cluster3645-006 |
| Cluster3647 | - | - | up | - | - | - | PTI1-like tyrosine- kinase At3 | SALT00000091549_Cluster3647-002 |
| Cluster3650 | - | - | up | - | up | up | PREDICTED: uncharacterize | SALT00000085005_Cluster3650-003 |
| Cluster3652 | - | - | down | - | - | down | gag-pol poly | SALT00000086200_Cluster3652-001 |
| Cluster3667 | - | - | up | - | up | up | peroxisomal acyl-coenzyme A | SALT00000084680_Cluster3667-029 |
| Cluster3673 | up | up | up | - | down | - | 5-methyltetrahydropteroyltrig | SALT00000065478_Cluster3673-001 |
| Cluster3679 | - | - | down | - | - | - | Os09g0114500 [Oryza sativa | SALT00000088291_Cluster3679-024 |
| Cluster368 | up | up | up | - | up | - | delta 1-pyrroline-5-carboxylat | SALT00000084101_Cluster368-001 |
| Cluster3691 | - | - | up | - | - | - | cyclic nucleotide-gated ion ch | SALT00000090958_Cluster3691-003 |
| Cluster3695 | - | - | down | - | - | down | replication A 70 kDa DNA-bi | SALT00000033974_Cluster3695-001 |
| Cluster3696 | - | - | up | - | up | up | Os03g0335300 [Oryza sativa | SALT00000084140_Cluster3696-024 |
| Cluster37 | - | - | - | - | up | - | sucrose synthase 1 [Zea mays | SALT00000016805_Cluster37-001 |
| Cluster3700 | - | - | - | - | up | up | S-receptor kinase KIK1 precu | SALT00000079569_Cluster3700-013 |
| Cluster3712 | - | - | up | - | up | up | DNA repair rhp26 [Zea mays] | SALT00000081006_Cluster3712-001 |
| Cluster3718 | - | - | down | - | - | - | Os12g0609800 [Oryza sativa | SALT00000084309_Cluster3718-007 |
| Cluster3723 | - | - | down | - | down | down | Os08g0407200 [Oryza sativa | SALT00000069227_Cluster3723-001 |
| Cluster3726 | - | - | down | - | down | - | 1-deoxy-D-xylulose-5-phosph | SALT00000034190_Cluster3726-004 |
| Cluster3730 | - | - | up | - | up | - | SMG7 [Zea mays] | SALT00000081991_Cluster3730-001 |
| Cluster3732 | - | - | down | - | down | down | ATP synthase subunit chlorop | SALT00000027696_Cluster3732-001 |
| Cluster3740 | - | - | up | - | - | - | Os04g0437600 [Oryza sativa | SALT00000064312_Cluster3740-015 |
| Cluster3749 | - | down | down | - | - | - | TPA: hemerythrin motif-conta | SALT00000088815_Cluster3749-002 |

| | | | | | | | | |
|---|---|---|---|---|---|---|---|---|
| Cluster3755 | - | - | down | - | - | - | PIR isoform X2 [Brachypodiu | SALT00000088432_Cluster3755-005 |
| Cluster3756 | - | - | down | - | - | - | ubiquitin carboxyl-terminal hy | SALT00000092955_Cluster3756-008 |
| Cluster3774 | - | - | up | - | up | up | remorin [Zea mays] | SALT00000086352_Cluster3774-001 |
| Cluster3779 | - | - | down | - | down | down | B3 domain-containing Os03g( | SALT00000067621_Cluster3779-001 |
| Cluster3782 | - | up | up | - | up | - | cold acclimation COR413-PM | SALT00000093242_Cluster3782-002 |
| Cluster3785 | - | - | down | - | down | down | sucrose synthase 1 [Oryza sati | SALT00000079048_Cluster3785-001 |
| Cluster3789 | up | - | up | - | - | - | methylenetetrahydrofolate red | SALT00000039913_Cluster3789-008 |
| Cluster3800 | - | - | up | - | up | up | probable calcium-transporting | SALT00000090912_Cluster3800-003 |
| Cluster3805 | - | - | up | - | - | - | NLP2-like [Zea mays] | SALT00000091606_Cluster3805-003 |
| Cluster3809 | - | - | up | - | up | up | [Oryza sativa Japonica Group] | SALT00000049264_Cluster3809-001 |
| Cluster3812 | - | - | - | - | down | - | Tubulin beta-1 partial | SALT00000044105_Cluster3812-001 |
| Cluster3835 | up | - | - | - | - | - | hypothetical protein GLYMA_ | SALT00000031333_Cluster3835-002 |
| Cluster3837 | - | up | up | - | - | - | Os08g0120500 [Oryza sativa | SALT00000064444_Cluster3837-001 |
| Cluster3840 | - | - | - | - | down | - | Serine threonine- kinase CTR | SALT00000085797_Cluster3840-001 |
| Cluster385 | - | - | down | - | - | - | disease resistance RGA1 | SALT00000033792_Cluster385-003 |
| Cluster3852 | down | down | down | - | - | - | TPA: hemerythrin motif-conta | SALT00000092007_Cluster3852-008 |
| Cluster3858 | - | - | down | - | down | down | 50S ribosomal chloroplastic | SALT00000086123_Cluster3858-002 |
| Cluster3866 | - | - | down | - | - | down | ribosomal S3 (chloroplast) [S] | SALT00000030117_Cluster3866-003 |
| Cluster3874 | - | - | down | - | - | - | Os05g0291700 [Oryza sativa | SALT00000089882_Cluster3874-003 |
| Cluster3877 | - | - | - | - | up | up | sister-chromatid cohesion 3 | SALT00000086191_Cluster3877-023 |
| Cluster388 | - | - | down | - | down | down | magnesium-chelatase subunit | SALT00000081149_Cluster388-014 |
| Cluster3881 | - | - | up | - | up | up | [Hordeum vulgare vulgare] | SALT00000065944_Cluster3881-008 |
| Cluster3884 | - | - | down | - | down | down | arginine decarboxylase 1 | SALT00000074499_Cluster3884-001 |
| Cluster39 | - | - | up | - | - | - | LRR receptor-like serine threo | SALT00000087045_Cluster39-001 |
| Cluster390 | - | - | down | - | down | down | DNA-directed RNA polymera | SALT00000077895_Cluster390-002 |
| Cluster3918 | - | - | - | - | down | - | PREDICTED: uncharacterized | SALT00000091533_Cluster3918-002 |
| Cluster3930 | - | - | down | - | down | down | SAC3 family 1 [Setaria italica | SALT00000087539_Cluster3930-003 |
| Cluster3935 | - | - | - | - | up | up | lichenase-2 precursor [Zea ma | SALT00000089710_Cluster3935-006 |
| Cluster3943 | - | - | up | - | up | up | phosphatidylinositol 4-kinase | SALT00000065040_Cluster3943-017 |
| Cluster3945 | - | - | up | - | up | up | dual specificity kinase shkD is | SALT00000091806_Cluster3945-001 |
| Cluster395 | - | - | up | - | up | up | FIP1[V] [Setaria italica] | SALT00000086843_Cluster395-009 |
| Cluster3954 | - | - | - | - | - | up | 50S ribosomal L18 [Oryza sat | SALT00000091613_Cluster3954-001 |
| Cluster3961 | - | down | down | - | - | - | RHOMBOID chloroplastic iso | SALT00000081779_Cluster3961-003 |
| Cluster3965 | - | - | - | - | up | - | E3 ubiquitin- ligase RMA1H1 | SALT00000090346_Cluster3965-003 |
| Cluster3981 | up | up | up | - | up | up | Exo70 exocyst complex subur | SALT00000075101_Cluster3981-004 |
| Cluster3996 | - | - | - | - | down | down | NLP1 isoform X2 [Setaria ital | SALT00000089417_Cluster3996-001 |
| Cluster4 | - | down | down | - | down | - | cellulose synthase-4 [Oryza sa | SALT00000086638_Cluster4-068 |
| Cluster4009 | - | down | down | - | - | - | [Hordeum vulgare vulgare] | SALT00000079415_Cluster4009-009 |
| Cluster4016 | - | - | down | - | down | down | phosphoribulokinase [Zea ma | SALT00000035957_Cluster4016-010 |
| Cluster4021 | - | - | - | down | - | - | probable glycosyltransferase A | SALT00000073903_Cluster4021-001 |
| Cluster4022 | - | - | up | - | up | up | serine carboxypeptidase-like 2 | SALT00000085528_Cluster4022-003 |
| Cluster4026 | - | - | down | - | down | down | CCT motif family [Zea mays] | SALT00000060830_Cluster4026-001 |
| Cluster4028 | - | - | down | - | - | - | Os02g0195000 [Oryza sativa | SALT00000056538_Cluster4028-003 |
| Cluster4037 | - | - | - | - | down | - | U-box domain-containing 33- | SALT00000034615_Cluster4037-001 |
| Cluster4038 | - | - | - | - | up | - | 26S proteasome regulatory pa | SALT00000043266_Cluster4038-013 |
| Cluster4039 | - | - | down | - | - | - | inactive receptor kinase | SALT00000077376_Cluster4039-004 |
| Cluster4042 | up | up | - | - | - | - | adenosylhomocysteinase-like | SALT00000037664_Cluster4042-026 |
| Cluster4047 | - | - | up | - | - | - | LOC100282635 isoform X1 [ | SALT00000037422_Cluster4047-001 |
| Cluster405 | - | - | - | - | up | - | enolase-phosphatase E1-like [ | SALT00000084475_Cluster405-011 |
| Cluster4057 | - | up | up | - | up | up | xanthine dehydrogenase | SALT00000086765_Cluster4057-004 |
| Cluster4065 | - | - | down | - | down | - | TPA: cytochrome P450 superf | SALT00000001225_Cluster4065-001 |
| Cluster4067 | - | - | down | - | - | - | MAG2-interacting 2 isoform X | SALT00000033748_Cluster4067-002 |
| Cluster4069 | - | - | down | - | down | down | 1-deoxy-D-xylulose-5-phosph | SALT00000072172_Cluster4069-001 |
| Cluster4070 | - | - | down | - | down | down | 40S ribosomal S3-3 | SALT00000086185_Cluster4070-003 |
| Cluster4073 | - | - | down | - | - | down | phosphoenolpyruvate carboxy | SALT00000053810_Cluster4073-023 |
| Cluster4076 | - | - | up | - | - | - | TAF5 [Oryza sativa Japonica ( | SALT00000073029_Cluster4076-004 |
| Cluster4093 | - | - | up | - | - | - | Os02g0735700 [Oryza sativa | SALT00000045216_Cluster4093-011 |
| Cluster4095 | - | - | - | - | - | down | E3 SUMO- ligase SIZ1 isofor | SALT00000088624_Cluster4095-007 |

| | | | | | | | | |
|---|---|---|---|---|---|---|---|---|
| Cluster41 | - | - | down | - | down | down | Os01g0752700 [Oryza sativa | SALT00000073793_Cluster41-005 |
| Cluster4102 | up | - | - | - | down | - | Os09g0484900 [Oryza sativa | SALT00000008063_Cluster4102-001 |
| Cluster4114 | - | - | down | - | - | - | MCM10 minichromosome ma | SALT00000086922_Cluster4114-002 |
| Cluster4121 | - | - | up | - | up | up | UDP-glycosyltransferase 72B | SALT00000050472_Cluster4121-003 |
| Cluster4130 | - | - | down | - | down | down | actin-7 [Brachypodium distacl | SALT00000054158_Cluster4130-001 |
| Cluster4144 | - | - | down | - | down | - | EMB2756 [Zea mays] | SALT00000089740_Cluster4144-002 |
| Cluster4145 | - | - | up | - | up | up | plasma membrane H+-ATPase | SALT00000089739_Cluster4145-004 |
| Cluster4152 | - | - | down | - | down | down | Endoglucanase 10 | SALT00000021656_Cluster4152-030 |
| Cluster4167 | - | - | up | - | up | up | Os01g0689900 [Oryza sativa | SALT00000032098_Cluster4167-002 |
| Cluster4174 | down | - | - | - | - | - | probable arabinosyltransferase | SALT00000029610_Cluster4174-001 |
| Cluster4203 | - | - | up | - | - | - | Cell division cycle expressed | SALT00000083385_Cluster4203-012 |
| Cluster4211 | - | - | - | - | down | - | ---NA--- | SALT00000086411_Cluster4211-001 |
| Cluster4215 | up | up | up | - | - | - | betaine aldehyde dehydrogena | SALT00000030632_Cluster4215-012 |
| Cluster4217 | - | - | up | - | - | - | CBL-interacting kinase 19 | SALT00000086708_Cluster4217-005 |
| Cluster4221 | - | - | up | - | up | - | phosphoinositide phosphatase | SALT00000091229_Cluster4221-005 |
| Cluster4235 | - | - | down | - | - | - | glucose-6-phosphate 1- cytopl | SALT00000088093_Cluster4235-015 |
| Cluster4255 | - | - | down | - | down | - | TPA: bZIP transcription factor | SALT00000090886_Cluster4255-002 |
| Cluster4258 | - | - | - | - | - | up | importin subunit beta-1 [Bracl | SALT00000088520_Cluster4258-022 |
| Cluster4259 | - | - | up | - | up | up | potassium transporter | SALT00000063708_Cluster4259-007 |
| Cluster4291 | - | - | up | - | up | - | probable serine threonine- kin | SALT00000033343_Cluster4291-003 |
| Cluster4299 | - | down | down | - | down | - | serine mitochondrial [Setaria i | SALT00000054596_Cluster4299-031 |
| Cluster4308 | - | - | down | - | down | - | pentatricopeptide repeat-conta | SALT00000081300_Cluster4308-004 |
| Cluster4323 | - | - | up | - | - | - | WRKY DNA-binding domain | SALT00000046325_Cluster4323-001 |
| Cluster4328 | - | - | - | - | down | down | RNA binding Rp120 [Oryza s | SALT00000070310_Cluster4328-026 |
| Cluster4344 | down | down | down | - | - | - | Os02g0674800 [Oryza sativa | SALT00000086322_Cluster4344-005 |
| Cluster4346 | - | down | down | - | - | down | ribulose-1,5-bisphosphate carl | SALT00000054855_Cluster4346-016 |
| Cluster4348 | - | - | up | - | up | up | chloroplast nucleoid DNA-bin | SALT00000041881_Cluster4348-005 |
| Cluster436 | - | - | up | - | up | - | NADP-specific isocitrate dehy | SALT00000057287_Cluster436-016 |
| Cluster4384 | - | - | up | - | - | - | Transmembrane 9 superfamily | SALT00000093455_Cluster4384-002 |
| Cluster4385 | - | - | up | - | - | - | core alpha 1,3-fucosyltransfer | SALT00000058942_Cluster4385-001 |
| Cluster4397 | - | - | up | - | up | up | phosphate:H+ symporter fami | SALT00000029478_Cluster4397-008 |
| Cluster4405 | - | - | - | - | down | - | ATPase family AAA domain-c | SALT00000079828_Cluster4405-001 |
| Cluster441 | - | - | down | - | down | - | TCP-domain [Oryza sativa Jap | SALT00000049695_Cluster441-001 |
| Cluster4418 | - | - | - | - | up | - | serine threonine- phosphatase | SALT00000034035_Cluster4418-005 |
| Cluster4445 | - | up | - | - | - | down | MYB DNA-binding domain s | SALT00000086201_Cluster4445-001 |
| Cluster4454 | - | - | up | - | up | up | Os12g0230600 [Oryza sativa | SALT00000088024_Cluster4454-001 |
| Cluster4461 | - | - | down | - | - | - | hypothetical protein SORBI_( | SALT00000090748_Cluster4461-010 |
| Cluster4475 | - | - | down | - | down | down | retrotransposon unclassified | SALT00000090654_Cluster4475-001 |
| Cluster4493 | - | - | up | - | up | - | kinase chloroplastic-like isofo | SALT00000083810_Cluster4493-001 |
| Cluster4494 | - | - | up | - | - | - | kinase domain containing [Ze | SALT00000056280_Cluster4494-002 |
| Cluster4498 | - | up | up | - | up | up | carbonyl reductase [NADPH] | SALT00000087383_Cluster4498-001 |
| Cluster4513 | - | - | up | - | up | up | probable receptor kinase At5g | SALT00000090125_Cluster4513-001 |
| Cluster4523 | - | up | up | up | - | - | dnaJ subfamily B member 4 [ | SALT00000044587_Cluster4523-001 |
| Cluster4525 | - | - | down | - | - | - | PREDICTED: uncharacterized | SALT00000087130_Cluster4525-002 |
| Cluster453 | - | - | - | - | down | - | Sec23 Sec24 trunk domain co | SALT00000091816_Cluster453-007 |
| Cluster4530 | down | - | - | - | - | - | Os10g0467800 [Oryza sativa | SALT00000086550_Cluster4530-001 |
| Cluster4542 | - | - | down | - | down | - | ribosomal S4 (chloroplast) [Pl | SALT00000033618_Cluster4542-006 |
| Cluster4554 | - | down | - | - | - | - | ETHYLENE-INSENSITIVE3 | SALT00000071776_Cluster4554-008 |
| Cluster4569 | - | - | down | - | down | down | ATP-dependent Clp protease p | SALT00000092790_Cluster4569-001 |
| Cluster4571 | - | - | down | - | down | - | Cell division cycle expressed | SALT00000067110_Cluster4571-001 |
| Cluster4574 | - | - | down | - | down | down | beta- partial | SALT00000058532_Cluster4574-001 |
| Cluster4607 | - | up | up | - | up | - | alcohol dehydrogenase 1 [Zea | SALT00000049562_Cluster4607-041 |
| Cluster461 | - | - | down | - | - | - | TSS [Setaria italica] | SALT00000079797_Cluster461-008 |
| Cluster4629 | - | - | up | - | up | up | probable serine threonine- kin | SALT00000089759_Cluster4629-003 |
| Cluster4641 | - | - | - | - | - | up | hypothetical protein SETIT_0 | SALT00000093560_Cluster4641-011 |
| Cluster4677 | - | - | up | - | up | - | kinase [Oryza sativa Japonica | SALT00000087906_Cluster4677-006 |
| Cluster4688 | - | - | down | - | down | down | elongation factor 2 [Oryza sat | SALT00000071388_Cluster4688-001 |
| Cluster4692 | - | up | - | - | - | down | CBL-interacting kinase family | SALT00000090539_Cluster4692-016 |

| | | | | | | | | |
|---|---|---|---|---|---|---|---|---|
| Cluster4698 | - | - | - | - | down | - | WEB family chloroplastic-lik | SALT00000073965_Cluster4698-002 |
| Cluster4721 | - | - | - | - | down | - | ATP-dependent RNA helicase | SALT00000093437_Cluster4721-001 |
| Cluster4726 | - | - | up | - | - | - | vacuolar sorting-associated 13 | SALT00000078935_Cluster4726-001 |
| Cluster4727 | - | - | down | - | - | - | cellulose synthase D4 | SALT00000087372_Cluster4727-006 |
| Cluster473 | - | - | - | - | down | down | C3HC zinc finger-like family | SALT00000078189_Cluster473-001 |
| Cluster4739 | - | - | down | - | - | - | retrotransposon unclassified | SALT00000087400_Cluster4739-001 |
| Cluster4769 | - | - | - | - | - | down | Os08g0541500 [Oryza sativa | SALT00000034603_Cluster4769-001 |
| Cluster4770 | - | - | - | - | up | - | sugar carrier C-like | SALT00000059531_Cluster4770-008 |
| Cluster4777 | - | - | down | - | down | down | probable LRR receptor-like se | SALT00000091190_Cluster4777-001 |
| Cluster4793 | - | - | down | - | down | down | phosphoribosylaminoimidazo | SALT00000090801_Cluster4793-002 |
| Cluster4798 | - | - | - | - | - | up | probable receptor-like serine t | SALT00000079798_Cluster4798-005 |
| Cluster48 | - | down | down | down | down | - | Os03g0788600 [Oryza sativa | SALT00000013489_Cluster48-015 |
| Cluster4801 | up | up | up | up | up | - | receptor kinase HAIKU2 [Zea | SALT00000090957_Cluster4801-007 |
| Cluster4808 | - | - | - | - | down | - | Os04g0623800 [Oryza sativa | SALT00000046970_Cluster4808-002 |
| Cluster4836 | - | - | up | - | - | - | Rhomboid family [Zea mays] | SALT00000088737_Cluster4836-001 |
| Cluster4844 | - | - | down | - | down | down | ML domain isoform X2 [Zea | SALT00000090830_Cluster4844-001 |
| Cluster4849 | - | - | up | - | up | up | beta-1,3-galactosyltransferase | SALT00000088098_Cluster4849-002 |
| Cluster4866 | - | - | down | - | down | down | ATP synthase CF1 alpha subu | SALT00000086850_Cluster4866-004 |
| Cluster4880 | - | - | down | - | down | down | phosphatidylinositol phosphat | SALT00000071117_Cluster4880-004 |
| Cluster4884 | - | - | down | - | down | - | disease resistance RGA2 isofo | SALT00000086453_Cluster4884-004 |
| Cluster4897 | - | down | down | - | down | - | tubulin beta-5 chain | SALT00000050560_Cluster4897-046 |
| Cluster490 | - | - | down | - | - | - | Os02g0448400 [Oryza sativa | SALT00000002039_Cluster490-001 |
| Cluster4907 | - | - | - | - | down | down | U-box domain-containing 4 [Z | SALT00000031009_Cluster4907-001 |
| Cluster4928 | - | - | down | - | - | - | ---NA--- | SALT00000084930_Cluster4928-001 |
| Cluster4932 | - | up | - | up | - | down | ---NA--- | SALT00000085430_Cluster4932-001 |
| Cluster4935 | - | - | - | - | - | up | [Hordeum vulgare vulgare] | SALT00000089786_Cluster4935-007 |
| Cluster4940 | - | - | - | - | up | - | ubiquitin-activating enzyme E | SALT00000093483_Cluster4940-001 |
| Cluster4942 | - | - | down | - | down | down | photosystem II CP47 chloropl | SALT00000034597_Cluster4942-003 |
| Cluster4948 | - | - | down | - | - | - | ATP citrate lyase a-subunit [O | SALT00000040272_Cluster4948-008 |
| Cluster4949 | - | - | up | - | up | up | beta-glucosidase 3B-like isofo | SALT00000079278_Cluster4949-010 |
| Cluster4951 | - | - | down | - | - | - | metal transporter Nramp1-like | SALT00000086898_Cluster4951-003 |
| Cluster497 | - | - | down | - | - | - | phosphomethylpyrimidine chl | SALT00000090787_Cluster497-030 |
| Cluster4971 | - | - | down | - | down | down | histidine kinase 1 [Sorghum b | SALT00000089091_Cluster4971-010 |
| Cluster4981 | down | - | - | - | up | up | [Hordeum vulgare vulgare] | SALT00000033296_Cluster4981-001 |
| Cluster4985 | - | - | up | - | - | up | Os01g0905700 [Oryza sativa | SALT00000084983_Cluster4985-003 |
| Cluster50 | - | - | - | - | down | - | elongation factor 2 [Oryza sat | SALT00000075886_Cluster50-004 |
| Cluster5000 | - | - | down | - | down | down | COBRA 3 | SALT00000048261_Cluster5000-050 |
| Cluster5007 | down | down | - | - | up | - | Os07g0100500 [Oryza sativa | SALT00000090857_Cluster5007-010 |
| Cluster5018 | - | - | down | - | down | down | Os01g0758900 [Oryza sativa | SALT00000093322_Cluster5018-001 |
| Cluster5025 | - | - | down | - | - | - | 26S proteasome non-ATPase r | SALT00000090794_Cluster5025-003 |
| Cluster5032 | down | down | down | - | - | - | calnexin precursor [Zea mays] | SALT00000056257_Cluster5032-028 |
| Cluster5036 | - | - | down | - | - | - | chloroplast-localized Ptr -binc | SALT00000031947_Cluster5036-002 |
| Cluster5061 | - | - | down | - | down | down | ribulose-1,5-bisphosphate carl | SALT00000032356_Cluster5061-004 |
| Cluster5079 | - | - | up | - | up | - | IST1 [Zea mays] | SALT00000036736_Cluster5079-006 |
| Cluster508 | - | - | down | - | down | down | increased DNA methylation 1 | SALT00000085865_Cluster508-021 |
| Cluster5080 | - | up | - | up | - | down | xylanase inhibitor 2-like [Ory | SALT00000043554_Cluster5080-002 |
| Cluster5083 | - | - | up | - | up | - | autophagy-related 18h-like | SALT00000033028_Cluster5083-003 |
| Cluster5097 | - | - | down | - | down | - | pentatricopeptide repeat-conta | SALT00000064760_Cluster5097-001 |
| Cluster51 | - | - | up | - | up | up | RING finger and CHY zinc fi | SALT00000075178_Cluster51-013 |
| Cluster510 | - | - | up | - | up | - | glycine-rich domain-containin | SALT00000091379_Cluster510-013 |
| Cluster5103 | - | - | down | - | - | - | AF466646_7 poly | SALT00000091753_Cluster5103-001 |
| Cluster511 | - | - | up | - | up | up | 1-phosphatidylinositol-3-phos | SALT00000087623_Cluster511-009 |
| Cluster5114 | up | - | - | down | down | - | transposon En Spm sub-class | SALT00000087456_Cluster5114-001 |
| Cluster5126 | - | - | down | - | down | down | Os01g0911200 [Oryza sativa | SALT00000076478_Cluster5126-005 |
| Cluster5128 | - | - | down | - | - | - | probable GPI-anchored adhes | SALT00000086107_Cluster5128-007 |
| Cluster513 | - | - | down | - | down | - | Os03g0340900 [Oryza sativa | SALT00000088645_Cluster513-006 |
| Cluster5146 | - | - | down | - | down | down | DEAD-box ATP-dependent R | SALT00000087682_Cluster5146-005 |
| Cluster515 | - | - | - | - | down | - | phenylalanine ammonia-lyase | SALT00000073710_Cluster515-001 |

| | | | | | | | | |
|---|---|---|---|---|---|---|---|---|
| Cluster5151 | - | up | - | - | - | - | methyl-binding domain MBD | SALT00000092878_Cluster5151-004 |
| Cluster5158 | - | - | down | - | - | - | trichome birefringence-like 25 | SALT00000090849_Cluster5158-005 |
| Cluster518 | - | - | down | - | down | - | clustered mitochondria -like is | SALT00000089089_Cluster518-020 |
| Cluster52 | - | - | - | - | up | - | ramosa 1 enhancer locus 2 [Ze | SALT00000029909_Cluster52-010 |
| Cluster5204 | - | - | up | - | up | up | lysine-specific histone demeth | SALT00000089745_Cluster5204-012 |
| Cluster5209 | - | up | up | - | up | - | [Hordeum vulgare vulgare] | SALT00000088701_Cluster5209-002 |
| Cluster5216 | - | - | down | - | - | - | phosphatidylinositol 4-phosph | SALT00000084921_Cluster5216-002 |
| Cluster5227 | - | - | down | - | - | - | TPA: bZIP transcription factor | SALT00000033425_Cluster5227-002 |
| Cluster524 | - | - | - | - | down | - | alcohol dehydrogenase 2 [Zea | SALT00000037358_Cluster524-026 |
| Cluster5251 | - | - | - | - | up | up | probable alpha,alpha-trehalose | SALT00000086798_Cluster5251-011 |
| Cluster5254 | - | - | up | - | up | up | ---NA--- | SALT00000040700_Cluster5254-001 |
| Cluster5260 | - | - | down | - | - | down | calcium-binding CML48 | SALT00000091874_Cluster5260-001 |
| Cluster5265 | - | - | up | - | - | - | enolase [Oryza sativa Japonica | SALT00000043334_Cluster5265-098 |
| Cluster529 | - | - | up | - | up | up | stress inducible | SALT00000032436_Cluster529-004 |
| Cluster5292 | - | - | down | - | down | - | PREDICTED: uncharacterized | SALT00000087896_Cluster5292-003 |
| Cluster5299 | - | - | down | - | - | - | PREDICTED: uncharacterized | SALT00000084769_Cluster5299-006 |
| Cluster5307 | - | - | - | - | - | up | hypersensitive-induced reactio | SALT00000031277_Cluster5307-001 |
| Cluster5312 | up | up | up | - | up | up | ras-related Rab-2-A [Zea may | SALT00000088097_Cluster5312-001 |
| Cluster5319 | up | up | - | - | down | - | phospho-2-dehydro-3-deoxyh | SALT00000040618_Cluster5319-026 |
| Cluster534 | - | - | down | - | down | down | Os09g0397300 [Oryza sativa | SALT00000091247_Cluster534-017 |
| Cluster5340 | - | - | - | - | - | down | Os07g0145400 [Oryza sativa | SALT00000092080_Cluster5340-001 |
| Cluster5343 | - | - | down | - | - | - | Os09g0506000 [Oryza sativa | SALT00000034457_Cluster5343-002 |
| Cluster5345 | - | - | up | - | up | - | importin subunit alpha-1b | SALT00000047993_Cluster5345-002 |
| Cluster5346 | - | down | down | - | - | - | Os04g0691800 [Oryza sativa | SALT00000086779_Cluster5346-010 |
| Cluster535 | - | - | - | - | down | - | Os12g0236700 [Oryza sativa | SALT00000090635_Cluster535-001 |
| Cluster5353 | up | up | up | - | - | - | myosin-11-like isoform X3 [B | SALT00000091441_Cluster5353-005 |
| Cluster5355 | - | - | - | - | down | down | eukaryotic initiation factor 4A | SALT00000052153_Cluster5355-007 |
| Cluster5368 | - | - | - | - | up | - | chromatin structure-remodelin | SALT00000033483_Cluster5368-002 |
| Cluster5374 | - | - | down | - | down | down | [Oryza sativa Indica Group] | SALT00000053414_Cluster5374-001 |
| Cluster5381 | - | - | - | - | down | - | GDAP2 homolog [Cucumis m | SALT00000088496_Cluster5381-002 |
| Cluster5386 | - | up | up | - | up | - | universal stress A | SALT00000082410_Cluster5386-001 |
| Cluster5390 | - | - | - | - | down | - | disease resistance RGA1 | SALT00000031374_Cluster5390-002 |
| Cluster5396 | - | - | down | - | - | - | Os10g0571200 [Oryza sativa | SALT00000045326_Cluster5396-005 |
| Cluster5400 | - | - | down | - | - | - | chloroplast nucleoid DNA-bin | SALT00000086990_Cluster5400-016 |
| Cluster5404 | - | - | up | - | up | - | transposon Mariner sub-class | SALT00000088408_Cluster5404-004 |
| Cluster5407 | - | - | down | - | down | down | filament-like plant 3 [Brachyp | SALT00000092471_Cluster5407-005 |
| Cluster5408 | - | - | up | - | - | - | CBL-interacting kinase 11 [O | SALT00000032996_Cluster5408-001 |
| Cluster5409 | - | - | - | - | down | down | CBL-interacting kinase 2 | SALT00000058689_Cluster5409-007 |
| Cluster541 | - | - | down | - | down | down | Os03g0808100 [Oryza sativa | SALT00000070016_Cluster541-020 |
| Cluster5410 | - | - | up | - | - | - | topless-related 1-like isoform | SALT00000084879_Cluster5410-001 |
| Cluster5415 | - | - | - | down | down | - | PREDICTED: uncharacterized | SALT00000080313_Cluster5415-001 |
| Cluster5416 | - | - | down | - | down | - | O-methyltransferase ZRP4-lik | SALT00000002994_Cluster5416-002 |
| Cluster5418 | - | - | - | - | down | - | tetratricopeptide repeat 7A [O | SALT00000085381_Cluster5418-001 |
| Cluster5435 | - | up | - | - | - | down | pentatricopeptide repeat-conta | SALT00000090659_Cluster5435-003 |
| Cluster5437 | - | - | down | - | - | - | sulfate permease1 isoform X1 | SALT00000066148_Cluster5437-002 |
| Cluster5458 | - | - | down | - | down | - | LONGIFOLIA 1 [Brachypodi | SALT00000031825_Cluster5458-002 |
| Cluster546 | - | - | up | - | up | up | Os04g0321600 [Oryza sativa | SALT00000033525_Cluster546-010 |
| Cluster5463 | - | - | up | - | up | - | regulator of chromosome expr | SALT00000081400_Cluster5463-009 |
| Cluster5474 | - | - | down | - | - | - | Os01g0596100 [Oryza sativa | SALT00000088819_Cluster5474-010 |
| Cluster5487 | - | - | down | - | down | down | partial [Oryza sativa Japonica | SALT00000067491_Cluster5487-001 |
| Cluster5488 | - | - | down | - | down | - | translocase subunit chloroplas | SALT00000085730_Cluster5488-001 |
| Cluster549 | - | - | - | - | down | - | PREDICTED: uncharacterized | SALT00000063915_Cluster549-001 |
| Cluster5493 | - | - | down | - | - | - | importin beta-like SAD2 | SALT00000090885_Cluster5493-008 |
| Cluster55 | - | - | up | - | - | - | partial [Oryza sativa Japonica | SALT00000090344_Cluster55-001 |
| Cluster5501 | - | - | down | - | - | - | AF466646_7 poly | SALT00000030926_Cluster5501-001 |
| Cluster5503 | - | - | up | - | up | up | brassinosteroid insensitive 1 p | SALT00000088734_Cluster5503-006 |
| Cluster5511 | - | - | up | - | up | up | Ureide permease expressed [C | SALT00000000809_Cluster5511-001 |
| Cluster5520 | - | up | up | - | - | - | pentatricopeptide repeat-conta | SALT00000033396_Cluster5520-001 |

| | | | | | | | | |
|---|---|---|---|---|---|---|---|---|
| Cluster5522 | - | - | up | - | up | up | DEAD box RNA helicase 1 [Z | SALT00000040846_Cluster5522-010 |
| Cluster5558 | - | - | - | - | down | - | lysosomal alpha-mannosidase | SALT00000085737_Cluster5558-003 |
| Cluster5563 | - | - | down | - | down | down | PREDICTED: uncharacterized | SALT00000071069_Cluster5563-001 |
| Cluster5568 | - | - | down | - | - | down | Os11g0646300 [Oryza sativa | SALT00000091158_Cluster5568-002 |
| Cluster5577 | up | up | up | - | up | up | 6-phospho-1-fructokinase [Or | SALT00000053732_Cluster5577-007 |
| Cluster5629 | - | - | down | - | - | - | DUF231 domain containing fa | SALT00000093287_Cluster5629-002 |
| Cluster5631 | - | - | up | - | - | - | OBERON [Brachypodium dis | SALT00000068075_Cluster5631-012 |
| Cluster5634 | - | - | - | - | - | up | mechanosensitive ion channel | SALT00000033840_Cluster5634-001 |
| Cluster5642 | - | - | up | - | up | - | PHD-finger family expressed | SALT00000032522_Cluster5642-005 |
| Cluster5649 | - | - | - | - | up | - | Os01g0786900 [Oryza sativa | SALT00000034159_Cluster5649-004 |
| Cluster5650 | - | - | up | - | - | - | B3 domain-containing Os07g0 | SALT00000067038_Cluster5650-001 |
| Cluster5651 | - | - | up | - | - | - | [Hordeum vulgare vulgare] | SALT00000087721_Cluster5651-001 |
| Cluster5655 | - | - | up | - | up | up | DNA repair RAD50 | SALT00000054947_Cluster5655-012 |
| Cluster5664 | - | up | - | - | - | - | calmodulin-binding transcript | SALT00000088016_Cluster5664-004 |
| Cluster5666 | - | - | down | - | down | down | Os07g0170200 [Oryza sativa | SALT00000090291_Cluster5666-002 |
| Cluster57 | - | - | up | - | up | - | YT521-B-like family expresse | SALT00000067141_Cluster57-025 |
| Cluster5726 | - | - | up | - | up | up | peroxisomal fatty acid beta-ox | SALT00000033585_Cluster5726-004 |
| Cluster5741 | up | up | up | - | - | - | partial [Oryza sativa Japonica | SALT00000043450_Cluster5741-006 |
| Cluster5749 | - | - | up | - | up | up | NF-X1-type zinc finger NFXI | SALT00000084791_Cluster5749-005 |
| Cluster576 | - | - | up | - | - | - | cellulose synthase [Oryza sati | SALT00000091140_Cluster576-020 |
| Cluster5760 | - | - | - | - | - | down | 60S ribosomal L7-3-like | SALT00000077565_Cluster5760-005 |
| Cluster5767 | - | - | down | - | - | - | tyrosine-sulfated glycopeptide | SALT00000092872_Cluster5767-002 |
| Cluster578 | - | - | down | - | - | - | Sperm-associated antigen 6 | SALT00000088304_Cluster578-037 |
| Cluster5803 | - | - | down | - | - | - | UDP-L-rhamnose synthase [C | SALT00000090636_Cluster5803-001 |
| Cluster5807 | - | down | down | down | down | down | sucrose synthase 1 [Oryza sati | SALT00000073297_Cluster5807-001 |
| Cluster581 | - | - | down | - | down | - | phosphoenolpyruvate carboxy | SALT00000071073_Cluster581-001 |
| Cluster5824 | - | - | down | - | - | - | retrotransposon unclassified [( | SALT00000090164_Cluster5824-001 |
| Cluster5828 | - | - | up | - | - | - | Os02g0324300 [Oryza sativa | SALT00000086096_Cluster5828-001 |
| Cluster583 | - | down | down | - | down | - | cellulose synthase esA3 | SALT00000089307_Cluster583-017 |
| Cluster5830 | - | - | - | - | down | - | la-related 1A | SALT00000090596_Cluster5830-007 |
| Cluster584 | - | up | up | - | - | - | sentrin-specific protease 2 [Ze | SALT00000037782_Cluster584-001 |
| Cluster5848 | - | down | down | down | down | - | CRAL TRIO domain Sec14p- | SALT00000042891_Cluster5848-015 |
| Cluster5858 | - | - | down | - | down | - | CHROMATIN REMODELIN | SALT00000092048_Cluster5858-011 |
| Cluster5861 | - | - | down | - | - | - | serine threonine- kinase Nek5 | SALT00000092466_Cluster5861-004 |
| Cluster5868 | - | - | up | - | - | - | Os06g0580100 [Oryza sativa | SALT00000090206_Cluster5868-007 |
| Cluster5874 | - | - | up | - | up | up | WAT1-related At3g45870 [Se | SALT00000091205_Cluster5874-005 |
| Cluster5879 | - | - | up | - | - | - | Os01g0729900 [Oryza sativa | SALT00000086769_Cluster5879-004 |
| Cluster5898 | - | - | down | - | down | down | titin-like isoform X1 [Gossypi | SALT00000090300_Cluster5898-006 |
| Cluster5923 | - | down | - | - | - | - | Disease resistance RPM1 | SALT00000087296_Cluster5923-004 |
| Cluster5930 | up | up | - | - | down | down | Os04g0504800 [Oryza sativa | SALT00000084542_Cluster5930-001 |
| Cluster5957 | - | - | up | - | - | up | Os03g0279600 [Oryza sativa | SALT00000034193_Cluster5957-002 |
| Cluster5960 | - | - | down | - | down | down | Os02g0195500 [Oryza sativa | SALT00000087709_Cluster5960-003 |
| Cluster5965 | - | - | up | - | - | up | glycoside six- subgroup [Zea m | SALT00000088192_Cluster5965-003 |
| Cluster5969 | - | - | down | - | down | down | retrotransposon partial | SALT00000085143_Cluster5969-001 |
| Cluster597 | - | down | - | - | - | up | cellulose synthase [Oryza sati | SALT00000087618_Cluster597-018 |
| Cluster5990 | - | - | up | - | - | - | Os08g0518100 [Oryza sativa | SALT00000074164_Cluster5990-001 |
| Cluster6030 | - | - | - | - | down | - | Os05g0459300 [Oryza sativa | SALT00000032783_Cluster6030-002 |
| Cluster6031 | - | - | down | - | down | down | disease resistance RPP13 3 [C | SALT00000076557_Cluster6031-002 |
| Cluster6039 | - | - | down | - | - | - | probable zinc metalloprotease | SALT00000086870_Cluster6039-004 |
| Cluster6053 | - | - | up | - | up | up | Os09g0556200 [Oryza sativa | SALT00000090273_Cluster6053-008 |
| Cluster606 | - | - | down | - | down | down | Os07g0227700 [Oryza sativa | SALT00000067291_Cluster606-009 |
| Cluster6070 | - | - | up | - | up | - | Os07g0585100 [Oryza sativa | SALT00000093512_Cluster6070-002 |
| Cluster6071 | - | up | - | - | - | down | pseudouridylate synthase 7 ho | SALT00000033048_Cluster6071-002 |
| Cluster6072 | - | - | up | - | up | - | CTL DDB_G0274487 [Setaria | SALT00000036941_Cluster6072-006 |
| Cluster6077 | - | - | down | - | - | - | calcineurin subunit B [Zea ma | SALT00000090974_Cluster6077-003 |
| Cluster6082 | - | - | down | - | down | - | transcription factor APG-like | SALT00000087750_Cluster6082-001 |
| Cluster6086 | - | - | up | - | - | - | Os01g0801600 [Oryza sativa | SALT00000016491_Cluster6086-004 |
| Cluster609 | - | up | up | - | up | - | citrate synthase family | SALT00000061692_Cluster609-001 |

| | | | | | | | | |
|---|---|---|---|---|---|---|---|---|
| Cluster6109 | - | - | up | - | - | - | glutamate decarboxylase [Ory | SALT00000077532_Cluster6109-004 |
| Cluster6119 | - | - | - | - | up | - | phospholipid-transporting ATI | SALT00000078161_Cluster6119-001 |
| Cluster612 | - | - | down | - | down | down | importin beta-like SAD2 | SALT00000091111_Cluster612-012 |
| Cluster6129 | - | - | up | - | - | - | Os02g0517300 [Oryza sativa | SALT00000082291_Cluster6129-003 |
| Cluster613 | - | - | - | - | up | - | topless-related 2 isoform X1 | SALT00000087925_Cluster613-056 |
| Cluster6130 | - | - | down | - | down | - | retrotransposon expressed [Or | SALT00000085319_Cluster6130-001 |
| Cluster6159 | - | - | up | - | up | up | F-box kelch-repeat At1g7451( | SALT00000065266_Cluster6159-006 |
| Cluster6169 | - | - | - | - | down | - | chloride channel CLC-d isofo | SALT00000092714_Cluster6169-006 |
| Cluster6176 | - | - | - | - | - | up | hypothetical protein SETIT_0 | SALT00000087499_Cluster6176-008 |
| Cluster6179 | up | - | - | - | - | - | ---NA--- | SALT00000080455_Cluster6179-001 |
| Cluster6181 | - | - | down | - | down | - | NRT1 PTR FAMILY -like [Cu | SALT00000087264_Cluster6181-004 |
| Cluster6186 | - | - | up | - | up | up | ABC transporter C family mer | SALT00000029253_Cluster6186-002 |
| Cluster6191 | - | - | down | - | down | down | Os03g0255500 [Oryza sativa | SALT00000092442_Cluster6191-002 |
| Cluster6193 | - | - | down | - | - | down | ribosomal S18 (chloroplast) [S | SALT00000033833_Cluster6193-006 |
| Cluster620 | - | up | up | - | up | up | T-complex 11 1 | SALT00000090377_Cluster620-010 |
| Cluster6207 | - | - | up | - | up | up | general transcription factor 3C | SALT00000087734_Cluster6207-008 |
| Cluster6217 | - | - | down | - | - | - | inositol phosphatase [Oryza sa | SALT00000092876_Cluster6217-002 |
| Cluster6222 | - | up | up | - | up | - | phosphatase 2C isoform gamr | SALT00000050853_Cluster6222-001 |
| Cluster6225 | - | - | down | - | down | down | [Oryza sativa Japonica Group] | SALT00000089397_Cluster6225-001 |
| Cluster623 | - | - | - | - | down | - | Os07g0456700 [Oryza sativa | SALT00000089803_Cluster623-013 |
| Cluster6239 | - | - | up | - | up | up | [Hordeum vulgare vulgare] | SALT00000088213_Cluster6239-018 |
| Cluster624 | - | - | up | - | up | up | Os01g0706400 [Oryza sativa | SALT00000087021_Cluster624-025 |
| Cluster6251 | - | - | - | - | up | - | TPA: suppressor of G2 allele ( | SALT00000055056_Cluster6251-024 |
| Cluster6252 | - | - | - | - | down | - | Os03g0158500 [Oryza sativa | SALT00000070356_Cluster6252-001 |
| Cluster6269 | - | - | up | - | - | - | C2 and GRAM domain-contai | SALT00000091751_Cluster6269-002 |
| Cluster6272 | - | - | - | - | down | down | phenylalanine ammonia partia | SALT00000062544_Cluster6272-001 |
| Cluster6273 | - | - | down | - | down | - | Os07g0679700 [Oryza sativa | SALT00000066292_Cluster6273-001 |
| Cluster6284 | - | up | up | - | - | - | CBL-interacting kinase 1 [Ory | SALT00000012611_Cluster6284-006 |
| Cluster629 | - | up | up | - | - | - | EREBP [Oryza sativa Japonic | SALT00000036024_Cluster629-006 |
| Cluster6302 | - | down | down | - | down | - | Os03g0127700 [Oryza sativa | SALT00000074674_Cluster6302-001 |
| Cluster6319 | - | - | - | up | - | - | embryo yellow partial | SALT00000086275_Cluster6319-001 |
| Cluster6320 | - | up | - | - | - | - | [Hordeum vulgare vulgare] | SALT00000088717_Cluster6320-001 |
| Cluster6329 | - | - | up | - | up | - | [Hordeum vulgare vulgare] | SALT00000093141_Cluster6329-006 |
| Cluster6335 | - | - | up | - | up | - | mediator of RNA polymerase | SALT00000092499_Cluster6335-001 |
| Cluster6342 | - | - | up | - | up | up | Os03g0106200 [Oryza sativa | SALT00000087303_Cluster6342-003 |
| Cluster6345 | - | - | - | - | - | up | ARABIDILLO 1-like isoform | SALT00000092280_Cluster6345-002 |
| Cluster6352 | - | up | up | - | up | - | Os05g0372800 [Oryza sativa | SALT00000090744_Cluster6352-003 |
| Cluster6358 | - | - | up | - | up | up | elicitor inducible beta-1,3-glu | SALT00000052481_Cluster6358-024 |
| Cluster6371 | - | - | - | - | - | up | zinc finger (C3HC4-type RIN | SALT00000088844_Cluster6371-003 |
| Cluster6372 | - | - | - | - | up | up | glycosyltransferase family 64 | SALT00000090911_Cluster6372-011 |
| Cluster6376 | - | - | down | down | down | - | replication A 70 kDa DNA-bi | SALT00000084121_Cluster6376-001 |
| Cluster6378 | - | - | up | - | - | - | alpha-aminoadipic semialdehy | SALT00000090583_Cluster6378-001 |
| Cluster6382 | - | - | down | - | down | down | phenylalanine ammonia-lyase | SALT00000075480_Cluster6382-001 |
| Cluster6383 | - | - | up | - | - | - | CBL-interacting kinase 10 | SALT00000079304_Cluster6383-003 |
| Cluster6386 | - | - | - | - | down | - | BEL1-type homeodomain | SALT00000092947_Cluster6386-002 |
| Cluster6387 | - | - | - | - | down | - | phenylalanine ammonia lyase | SALT00000067129_Cluster6387-001 |
| Cluster6390 | - | - | - | - | - | down | endonuclease-reverse transcrip | SALT00000086540_Cluster6390-002 |
| Cluster6401 | - | - | down | - | - | - | Os12g0590900 [Oryza sativa | SALT00000085952_Cluster6401-003 |
| Cluster6405 | - | down | down | - | down | - | ECERIFERUM 26-like | SALT00000049687_Cluster6405-007 |
| Cluster6406 | - | - | down | - | down | down | reticulon B21 isoform X2 | SALT00000063717_Cluster6406-001 |
| Cluster6420 | - | - | up | - | up | up | brassinosteroid insensitive 1 p | SALT00000035169_Cluster6420-001 |
| Cluster6423 | - | - | down | - | down | - | [Oryza sativa Japonica Group] | SALT00000050645_Cluster6423-004 |
| Cluster6424 | - | - | down | - | down | down | monosaccharide-sensing 2 [Or | SALT00000076400_Cluster6424-001 |
| Cluster6426 | - | - | - | - | - | down | Retrovirus-related Pol poly LI | SALT00000085026_Cluster6426-001 |
| Cluster6434 | - | - | up | - | up | - | charged multivesicular body 1 | SALT00000085922_Cluster6434-001 |
| Cluster6438 | down | down | down | - | down | - | glucosidase 2 subunit beta [Or | SALT00000087816_Cluster6438-001 |
| Cluster6447 | - | - | down | - | down | down | small RNA degrading nucleas | SALT00000078911_Cluster6447-001 |
| Cluster645 | - | - | - | - | up | - | Os09g0413600 [Oryza sativa | SALT00000032136_Cluster645-002 |

| | | | | | | | | |
|---|---|---|---|---|---|---|---|---|
| Cluster6463 | - | - | - | - | - | down | PREDICTED: uncharacterize | SALT00000029925_Cluster6463-001 |
| Cluster6465 | - | down | down | - | - | - | ru large subunit-binding subu | SALT00000030756_Cluster6465-009 |
| Cluster6472 | - | - | down | - | down | down | plastid-specific 30S ribosoma | SALT00000091834_Cluster6472-001 |
| Cluster6498 | - | - | down | - | down | down | Os03g0731900 [Oryza sativa | SALT00000053134_Cluster6498-007 |
| Cluster653 | - | - | up | - | up | up | Os09g0397300 [Oryza sativa | SALT00000034423_Cluster653-023 |
| Cluster6556 | - | - | up | - | - | up | ---NA--- | SALT00000092239_Cluster6556-001 |
| Cluster6557 | - | up | up | - | up | up | ubiquitin-like modifier-activat | SALT00000065969_Cluster6557-001 |
| Cluster6569 | - | - | down | - | down | down | MPK14 - MAPK [Zea mays] | SALT00000088854_Cluster6569-001 |
| Cluster6576 | - | - | down | - | down | down | Transcription factor MYB39 | SALT00000092117_Cluster6576-002 |
| Cluster6578 | - | - | down | - | - | - | rho GTPase-activating 7-like i | SALT00000093292_Cluster6578-009 |
| Cluster6580 | - | - | down | - | - | - | two-pore potassium channel 2 | SALT00000087017_Cluster6580-002 |
| Cluster6585 | - | - | - | - | down | down | disulfide isomerase [Zea mays | SALT00000056778_Cluster6585-007 |
| Cluster6612 | - | - | up | - | up | - | DMI1 [Oryza sativa Japonica | SALT00000082979_Cluster6612-005 |
| Cluster6642 | down | down | down | - | - | - | KH domain-containing At4g1 | SALT00000082682_Cluster6642-003 |
| Cluster6670 | - | - | up | - | - | - | MEI2-like 2 isoform X2 | SALT00000086243_Cluster6670-015 |
| Cluster6690 | - | - | down | - | down | down | probable acyl-activating enzy | SALT00000082947_Cluster6690-009 |
| Cluster6697 | - | - | down | - | - | - | Os02g0319100 [Oryza sativa | SALT00000087041_Cluster6697-005 |
| Cluster6728 | - | - | up | - | up | up | cadmium zinc-transporting A | SALT00000031560_Cluster6728-002 |
| Cluster6731 | - | down | down | - | down | - | endoplasmin homolog [Setari | SALT00000070198_Cluster6731-001 |
| Cluster6737 | - | - | down | - | down | down | polygalacturonase PG1-like [ | SALT00000091510_Cluster6737-001 |
| Cluster6750 | - | - | - | - | down | - | probable NAD(P)H dehydrog | SALT00000090615_Cluster6750-002 |
| Cluster6758 | - | - | down | - | - | down | enolase 2 [Setaria italica] | SALT00000037860_Cluster6758-001 |
| Cluster676 | - | - | up | - | - | - | ABC transporter G family me | SALT00000087681_Cluster676-009 |
| Cluster6768 | - | - | - | - | - | down | COI1 [Oryza sativa Japonica | SALT00000052656_Cluster6768-003 |
| Cluster6773 | - | - | down | - | down | down | chloroplastic group IIA intron | SALT00000086579_Cluster6773-001 |
| Cluster6779 | - | - | down | - | down | down | callose synthase 3-like | SALT00000083661_Cluster6779-001 |
| Cluster6788 | - | - | down | - | down | down | TPA: tubulin alpha-1 chain [Z | SALT00000061643_Cluster6788-006 |
| Cluster68 | - | - | up | - | - | - | Os10g0105400 [Oryza sativa | SALT00000080823_Cluster68-020 |
| Cluster682 | - | - | up | - | up | up | partial [Oryza sativa Japonica | SALT00000091522_Cluster682-001 |
| Cluster6824 | - | - | - | - | up | - | lon protease homolog peroxis | SALT00000078859_Cluster6824-013 |
| Cluster6827 | - | - | down | - | - | - | coatomer subunit zeta-2 [Ory | SALT00000077794_Cluster6827-006 |
| Cluster6838 | - | up | up | - | - | - | cytochrome P450 superfamily | SALT00000056697_Cluster6838-018 |
| Cluster6852 | - | - | down | - | down | - | glutathione S-transferase isof | SALT00000033201_Cluster6852-001 |
| Cluster6853 | - | - | - | - | down | down | tubby-like F-box 6 isoform X | SALT00000076143_Cluster6853-002 |
| Cluster687 | - | - | down | - | down | down | ATP synthase CF1 beta subun | SALT00000082139_Cluster687-008 |
| Cluster6872 | - | - | up | - | up | up | probable UDP-N-acetylgluco | SALT00000050552_Cluster6872-009 |
| Cluster6877 | - | - | - | - | down | down | Os07g0599000 [Oryza sativa | SALT00000091021_Cluster6877-007 |
| Cluster6884 | - | - | up | - | - | up | [Hordeum vulgare vulgare] | SALT00000029740_Cluster6884-001 |
| Cluster689 | - | - | down | - | down | down | Brefeldin A-inhibited guanine | SALT00000086474_Cluster689-023 |
| Cluster6890 | - | - | down | - | - | - | LUTEIN DEFICIENT chloro | SALT00000086330_Cluster6890-002 |
| Cluster690 | - | down | down | down | down | down | tubulin beta-6 chain isoform X | SALT00000038746_Cluster690-004 |
| Cluster6905 | - | - | down | - | - | - | tyrosine-sulfated glycopeptide | SALT00000086560_Cluster6905-010 |
| Cluster6912 | - | - | - | - | up | - | probable inactive leucine-rich | SALT00000090485_Cluster6912-003 |
| Cluster6917 | - | - | - | - | - | down | heat shock 70 kDa 1 [Zea may | SALT00000071922_Cluster6917-001 |
| Cluster692 | - | - | down | - | down | down | cellulose synthase1 | SALT00000086969_Cluster692-003 |
| Cluster6930 | - | - | up | - | up | - | nuclear transcription factor Y | SALT00000088233_Cluster6930-001 |
| Cluster6940 | - | - | down | - | down | down | tubulin alpha-1 chain [Miscan | SALT00000048327_Cluster6940-001 |
| Cluster6941 | up | up | - | - | down | down | adenosylhomocysteinase-like | SALT00000067085_Cluster6941-019 |
| Cluster6942 | - | - | down | - | down | down | dihydrolipoamide S-acetyltra | SALT00000075154_Cluster6942-010 |
| Cluster6958 | - | - | down | - | - | - | reticulon B8 [Oryza sativa Ja | SALT00000087134_Cluster6958-002 |
| Cluster6972 | - | - | up | - | up | up | Os03g0659400 [Oryza sativa | SALT00000080874_Cluster6972-005 |
| Cluster6980 | - | - | up | - | up | up | F-box kelch-repeat At1g7451 | SALT00000084986_Cluster6980-003 |
| Cluster6988 | - | - | up | - | up | up | PREDICTED: uncharacterize | SALT00000092408_Cluster6988-007 |
| Cluster699 | - | - | down | - | down | down | 125 kDa kinesin-related isofo | SALT00000087942_Cluster699-007 |
| Cluster7041 | - | up | - | - | - | - | [Hordeum vulgare vulgare] | SALT00000091242_Cluster7041-001 |
| Cluster7043 | - | - | - | - | - | down | ketol-acid chloroplastic-like | SALT00000052836_Cluster7043-016 |
| Cluster7044 | - | - | up | - | - | - | phosphoserine aminotransfera | SALT00000054648_Cluster7044-019 |
| Cluster7047 | - | down | down | - | - | - | retrotransposon unclassified | SALT00000089216_Cluster7047-002 |

| | | | | | | | | |
|---|---|---|---|---|---|---|---|---|
| Cluster7049 | - | - | up | - | - | - | cellulose synthase A catalytic | SALT00000090490_Cluster7049-004 |
| Cluster7065 | - | - | down | - | - | - | zinc finger MAGPIE-like isof | SALT00000087936_Cluster7065-005 |
| Cluster7071 | - | - | up | - | - | - | Os08g0542900 [Oryza sativa | SALT00000045307_Cluster7071-001 |
| Cluster7090 | - | - | down | - | - | - | pentatricopeptide repeat-conta | SALT00000086572_Cluster7090-001 |
| Cluster7092 | - | - | down | - | down | down | CBL-interacting kinase 2 | SALT00000055503_Cluster7092-017 |
| Cluster7120 | - | down | down | - | - | - | homeobox-leucine zipper RO( | SALT00000078363_Cluster7120-001 |
| Cluster7128 | - | - | - | - | down | - | caffeic acid 3-O-methyltransf( | SALT00000042841_Cluster7128-001 |
| Cluster7129 | - | - | up | - | - | - | Os10g0542200 [Oryza sativa | SALT00000045457_Cluster7129-006 |
| Cluster7143 | - | - | up | - | up | up | Os11g0603200 [Oryza sativa | SALT00000074556_Cluster7143-006 |
| Cluster717 | - | - | down | - | down | - | Os03g0255500 [Oryza sativa | SALT00000007775_Cluster717-001 |
| Cluster7185 | - | - | - | - | down | down | PREDICTED: uncharacterized | SALT00000092551_Cluster7185-002 |
| Cluster7189 | - | - | up | - | up | up | trehalose 6-phosphate synthas | SALT00000082987_Cluster7189-010 |
| Cluster72 | - | - | up | - | up | up | importin subunit beta-1 [Bracl | SALT00000085776_Cluster72-003 |
| Cluster720 | - | - | - | - | down | down | trafficking particle complex su | SALT00000087410_Cluster720-013 |
| Cluster7200 | down | down | down | - | - | - | Os02g0629400 [Oryza sativa | SALT00000092604_Cluster7200-004 |
| Cluster721 | - | - | down | - | down | down | RNA-binding 28 | SALT00000071662_Cluster721-001 |
| Cluster7211 | - | - | - | - | - | up | CTP synthase [Zea mays] | SALT00000079350_Cluster7211-001 |
| Cluster7213 | - | - | up | - | up | - | potassium transporter | SALT00000086711_Cluster7213-026 |
| Cluster7215 | - | - | - | - | down | down | 40S ribosomal SA [Zea mays] | SALT00000088140_Cluster7215-002 |
| Cluster7224 | - | - | - | - | up | - | Os02g0649700 [Oryza sativa | SALT00000082707_Cluster7224-002 |
| Cluster726 | - | up | up | - | up | - | MEI2-like 4 isoform X4 [Brac | SALT00000032807_Cluster726-011 |
| Cluster7274 | - | - | up | - | up | up | 12-oxophytodienoate reductas | SALT00000057213_Cluster7274-001 |
| Cluster7280 | - | - | - | - | down | down | oligosaccharyl transferase ST | SALT00000091579_Cluster7280-004 |
| Cluster7291 | - | up | up | up | up | - | alcohol dehydrogenase 1 [Zea | SALT00000057993_Cluster7291-001 |
| Cluster7301 | - | - | down | down | down | down | catalase [Saccharum hybrid cu | SALT00000041793_Cluster7301-006 |
| Cluster7308 | - | - | up | - | up | up | Os03g0726500 [Oryza sativa | SALT00000085431_Cluster7308-001 |
| Cluster7310 | - | - | up | - | up | - | U-box domain-containing 42 [ | SALT00000088600_Cluster7310-001 |
| Cluster7317 | - | - | down | - | down | down | leucine-rich repeat receptor ki | SALT00000077734_Cluster7317-001 |
| Cluster732 | - | - | up | - | up | up | ubiquitin-activating enzyme E | SALT00000081271_Cluster732-041 |
| Cluster7334 | - | - | down | - | - | - | sucrose phosphate synthase B | SALT00000086519_Cluster7334-003 |
| Cluster7335 | - | - | down | - | down | down | transcription-associated 1-like | SALT00000088357_Cluster7335-001 |
| Cluster7338 | - | - | - | - | - | down | cyclin-dependent kinase G-1-l | SALT00000078706_Cluster7338-002 |
| Cluster734 | - | - | down | - | down | down | ribulose-1,5-bisphosphate carl | SALT00000055248_Cluster734-002 |
| Cluster7357 | - | - | up | - | up | up | aminotransferase class III sup | SALT00000081864_Cluster7357-009 |
| Cluster7361 | up | up | up | up | up | - | Os01g0165100 [Oryza sativa | SALT00000084917_Cluster7361-002 |
| Cluster7367 | - | - | down | - | - | - | Os09g0412200 [Oryza sativa | SALT00000075133_Cluster7367-001 |
| Cluster7371 | - | - | up | - | up | up | Os02g0149800 [Oryza sativa | SALT00000008606_Cluster7371-001 |
| Cluster7394 | - | - | down | - | - | - | RNA-binding 39-like isoform | SALT00000089679_Cluster7394-005 |
| Cluster7397 | - | - | up | - | up | up | equilibrative nucleotide transp | SALT00000051841_Cluster7397-005 |
| Cluster7401 | - | - | up | - | up | up | soluble starch synthase III-1 [ | SALT00000033865_Cluster7401-001 |
| Cluster7402 | - | up | up | - | - | - | polyadenylate-binding RBP45 | SALT00000042026_Cluster7402-009 |
| Cluster7408 | - | - | up | - | up | up | [Hordeum vulgare vulgare] | SALT00000081424_Cluster7408-005 |
| Cluster741 | - | - | down | - | down | down | DNAJ heat shock N-terminal | SALT00000089823_Cluster741-006 |
| Cluster7416 | - | - | down | - | down | - | Os04g0504800 [Oryza sativa | SALT00000076041_Cluster7416-001 |
| Cluster742 | - | - | up | - | up | up | calcium-transporting ATPase | SALT00000090146_Cluster742-010 |
| Cluster744 | - | - | up | - | up | up | death-inducer obliterator 1-lik | SALT00000029675_Cluster744-018 |
| Cluster7443 | - | - | up | - | up | - | PREDICTED: uncharacterized | SALT00000091005_Cluster7443-008 |
| Cluster7452 | - | - | - | - | down | - | ---NA--- | SALT00000087218_Cluster7452-004 |
| Cluster7461 | - | - | up | - | up | up | Os03g0136900 [Oryza sativa | SALT00000089637_Cluster7461-011 |
| Cluster7472 | - | - | down | - | - | down | beta-galactosidase 3 [Setaria i | SALT00000090027_Cluster7472-001 |
| Cluster7485 | - | - | up | - | up | - | cAMP-regulated phospho 21-1 | SALT00000086790_Cluster7485-008 |
| Cluster749 | - | - | down | - | down | - | serine threonine-specific kinas | SALT00000087857_Cluster749-002 |
| Cluster75 | - | - | down | - | down | - | translational activator GCN1 | SALT00000069429_Cluster75-001 |
| Cluster7506 | down | down | down | - | - | - | kinesin NACK1 [Brachypodiu | SALT00000032381_Cluster7506-002 |
| Cluster7510 | - | - | - | - | down | down | interactor of constitutive activ | SALT00000088349_Cluster7510-008 |
| Cluster7514 | - | - | up | - | - | - | sucrose-phosphate synthase [( | SALT00000029722_Cluster7514-001 |
| Cluster7531 | - | - | down | - | down | - | Os04g0691800 [Oryza sativa | SALT00000084833_Cluster7531-001 |
| Cluster7533 | up | up | - | - | down | down | pentatricopeptide repeat-conta | SALT00000091971_Cluster7533-001 |

| | | | | | | | | |
|---|---|---|---|---|---|---|---|---|
| Cluster7558 | - | - | down | - | down | down | DEAD-box ATP-dependent R | SALT00000087884_Cluster7558-003 |
| Cluster7560 | - | - | - | - | down | down | dolichyl-diphosphooligosacch | SALT00000038636_Cluster7560-004 |
| Cluster7562 | - | - | down | - | - | - | Os01g0880400 [Oryza sativa | SALT00000093300_Cluster7562-001 |
| Cluster757 | - | - | - | - | down | down | mitogen-activated kinase 4 [O | SALT00000053172_Cluster757-001 |
| Cluster7585 | up | up | up | - | - | - | cinnamyl alcohol dehydrogena | SALT00000003583_Cluster7585-005 |
| Cluster7600 | up | up | - | - | down | down | serine hydroxymethyltransfera | SALT00000036460_Cluster7600-001 |
| Cluster7609 | - | - | up | - | up | up | Os05g0475300 [Oryza sativa | SALT00000083710_Cluster7609-005 |
| Cluster7617 | - | - | up | - | up | - | Os03g0142800 [Oryza sativa | SALT00000092006_Cluster7617-002 |
| Cluster7627 | - | up | up | - | up | up | glycine--tRNA mitochondrial | SALT00000090829_Cluster7627-010 |
| Cluster7662 | - | - | - | - | - | down | probable mediator of RNA po | SALT00000032725_Cluster7662-002 |
| Cluster7663 | - | - | down | - | - | - | rRNA methyltransferase | SALT00000071853_Cluster7663-004 |
| Cluster7685 | - | - | down | - | - | - | hypothetical protein SORBI_( | SALT00000090809_Cluster7685-002 |
| Cluster77 | - | - | down | - | down | - | exportin-2 [Zea mays] | SALT00000086230_Cluster77-001 |
| Cluster7703 | - | - | down | - | - | - | cell division cycle cofactor of | SALT00000089802_Cluster7703-006 |
| Cluster7735 | - | - | down | - | down | - | sel-1 homolog 2-like [Zea ma | SALT00000087748_Cluster7735-001 |
| Cluster7744 | - | - | down | - | down | - | QUIRKY [Oryza sativa Japon | SALT00000075771_Cluster7744-014 |
| Cluster7746 | - | up | up | up | up | up | NRT1 PTR FAMILY -like iso | SALT00000088614_Cluster7746-001 |
| Cluster7747 | - | - | up | - | up | up | lipid phosphate phosphatase 2 | SALT00000021852_Cluster7747-001 |
| Cluster7749 | - | - | down | - | - | - | Os02g0771800 [Oryza sativa | SALT00000066212_Cluster7749-002 |
| Cluster775 | - | - | up | - | - | - | Homeobox-leucine zipper HO | SALT00000070328_Cluster775-004 |
| Cluster777 | - | - | up | - | up | up | lipoxygenase [Zea mays] | SALT00000071513_Cluster777-041 |
| Cluster7781 | - | - | down | - | down | down | anthocyanidin reductase-like [ | SALT00000088555_Cluster7781-003 |
| Cluster7782 | - | - | up | - | up | up | TPA: alcohol dehydrogenase1 | SALT00000089075_Cluster7782-007 |
| Cluster7783 | down | - | down | - | - | down | 26S proteasome non-ATPase r | SALT00000089144_Cluster7783-001 |
| Cluster7803 | - | - | up | - | up | up | Os04g0352400 [Oryza sativa | SALT00000087830_Cluster7803-013 |
| Cluster7806 | - | up | up | - | - | - | probable serine threonine- kin | SALT00000091685_Cluster7806-003 |
| Cluster7807 | - | - | up | - | - | - | aldehyde dehydrogenase fami | SALT00000007872_Cluster7807-001 |
| Cluster7808 | - | - | down | - | down | down | partial [Hordeum vulgare vulg | SALT00000051948_Cluster7808-001 |
| Cluster7836 | - | - | - | - | down | down | argonaute 1B isoform X2 [Set | SALT00000086659_Cluster7836-001 |
| Cluster784 | - | - | down | - | - | - | receptor kinase 1 | SALT00000090834_Cluster784-001 |
| Cluster7841 | - | - | up | - | up | - | transcription elongation factor | SALT00000091337_Cluster7841-010 |
| Cluster7849 | - | - | up | - | - | - | 6-phosphofructokinase 2 [Zea | SALT00000047475_Cluster7849-023 |
| Cluster785 | - | - | up | - | - | up | serine threonine- kinase EDR | SALT00000085294_Cluster785-019 |
| Cluster7880 | - | up | up | up | up | up | ankyrin repeat [Oryza sativa J | SALT00000092000_Cluster7880-004 |
| Cluster7883 | - | - | up | - | - | - | Grx_S17 - glutaredoxin subgr | SALT00000048295_Cluster7883-001 |
| Cluster7887 | - | - | down | - | - | - | homeobox-leucine zipper RO( | SALT00000085179_Cluster7887-001 |
| Cluster7899 | - | - | - | - | - | down | 52 kDa repressor of the inhibi | SALT00000032949_Cluster7899-001 |
| Cluster79 | - | - | up | - | up | up | receptor kinase HSL1 [Setaria | SALT00000087602_Cluster79-012 |
| Cluster7905 | - | - | up | - | up | up | aspartic ase 1 | SALT00000055231_Cluster7905-004 |
| Cluster7907 | - | - | down | down | down | down | catalase isozyme 3 | SALT00000060616_Cluster7907-015 |
| Cluster7919 | - | - | up | - | - | up | Os03g0248000 [Oryza sativa | SALT00000082263_Cluster7919-002 |
| Cluster7930 | - | - | - | - | up | - | probable GPI-anchored adhes | SALT00000091604_Cluster7930-004 |
| Cluster7933 | - | - | - | up | - | - | tubulin alpha-1 chain [Miscan | SALT00000047464_Cluster7933-001 |
| Cluster7949 | up | up | up | - | up | up | kinase superfamily | SALT00000034373_Cluster7949-004 |
| Cluster7951 | up | up | up | - | up | up | eukaryotic peptide chain relea | SALT00000091618_Cluster7951-002 |
| Cluster7985 | - | - | down | - | down | - | ru large subunit-binding subu | SALT00000046900_Cluster7985-001 |
| Cluster7993 | - | - | up | - | up | up | calreticulin-3-like [Setaria ital | SALT00000045957_Cluster7993-002 |
| Cluster80 | - | - | - | - | down | - | photosystem I P700 chlorophy | SALT00000088949_Cluster80-142 |
| Cluster801 | - | - | down | - | down | down | ribosomal S3 (chloroplast) [S | SALT00000086594_Cluster801-002 |
| Cluster8010 | - | - | down | - | down | down | Os03g0775000 [Oryza sativa | SALT00000086801_Cluster8010-001 |
| Cluster8020 | - | - | up | - | up | up | orphans transcription partial [ | SALT00000024010_Cluster8020-025 |
| Cluster8025 | - | - | down | - | down | down | LOC100285175 isoform X1 [ | SALT00000090578_Cluster8025-017 |
| Cluster8032 | - | up | - | - | - | - | triacylglycerol lipase SDP1-li | SALT00000090100_Cluster8032-002 |
| Cluster8046 | - | - | up | - | - | - | retrotransposon unclassified | SALT00000077350_Cluster8046-001 |
| Cluster8055 | - | down | - | - | - | - | phosphate transporter | SALT00000088138_Cluster8055-011 |
| Cluster8059 | - | up | up | - | up | - | Os01g0770000 [Oryza sativa | SALT00000083496_Cluster8059-001 |
| Cluster8085 | - | - | - | - | - | up | Os01g0549500 [Oryza sativa | SALT00000084200_Cluster8085-002 |
| Cluster8096 | - | - | - | - | down | down | hypothetical protein SORBI_( | SALT00000072021_Cluster8096-002 |

| | | | | | | | | |
|---|---|---|---|---|---|---|---|---|
| Cluster8105 | - | - | - | - | - | down | probable LRR receptor-like se | SALT00000090341_Cluster8105-001 |
| Cluster8106 | - | - | - | - | down | - | Os08g0345700 [Oryza sativa | SALT00000067701_Cluster8106-010 |
| Cluster8112 | - | up | up | - | up | up | Os09g0512900 [Oryza sativa | SALT00000083426_Cluster8112-002 |
| Cluster8113 | - | - | down | - | down | - | zinc finger MYM-type 1-like | SALT00000085965_Cluster8113-002 |
| Cluster8114 | - | - | up | - | - | up | 2-alkenal reductase (NADP(+ | SALT00000086142_Cluster8114-002 |
| Cluster8126 | - | - | down | - | down | - | BEL1-type homeodomain | SALT00000092941_Cluster8126-021 |
| Cluster8145 | - | down | down | - | - | - | Os01g0239700 [Oryza sativa | SALT00000081346_Cluster8145-008 |
| Cluster8153 | - | - | down | - | down | down | catalase isozyme 3 | SALT00000057693_Cluster8153-063 |
| Cluster8160 | - | - | - | - | up | - | probable methyltransferase PM | SALT00000033828_Cluster8160-003 |
| Cluster8167 | - | down | down | - | - | - | fasciclin-like arabinogalactan | SALT00000016988_Cluster8167-004 |
| Cluster8215 | down | down | down | - | - | - | [Oryza sativa Japonica Group] | SALT00000090213_Cluster8215-001 |
| Cluster8219 | - | - | down | - | down | - | QUIRKY [Oryza sativa Japon | SALT00000033772_Cluster8219-002 |
| Cluster8223 | - | - | - | - | down | - | Os01g0662700 [Oryza sativa | SALT00000088475_Cluster8223-001 |
| Cluster8232 | - | - | down | - | - | - | partial [Oryza sativa Japonica | SALT00000089624_Cluster8232-006 |
| Cluster8236 | - | - | down | - | - | - | 60S ribosomal L28-1 [Brachy | SALT00000034589_Cluster8236-001 |
| Cluster8237 | - | up | up | - | up | up | methylthioribose kinase [Zea | SALT00000045108_Cluster8237-006 |
| Cluster8246 | - | - | - | - | - | down | Wall-associated receptor kinas | SALT00000093126_Cluster8246-001 |
| Cluster8248 | - | - | - | - | down | - | Os02g0554300 [Oryza sativa | SALT00000081389_Cluster8248-004 |
| Cluster8268 | - | - | up | - | - | - | elicitor inducible beta-1,3-glu | SALT00000085531_Cluster8268-002 |
| Cluster8274 | - | - | up | - | up | up | PP2A regulatory subunit TAP4 | SALT00000038037_Cluster8274-024 |
| Cluster8282 | - | - | up | - | up | up | Os06g0701300 [Oryza sativa | SALT00000080328_Cluster8282-012 |
| Cluster8290 | - | down | down | - | down | - | [Hordeum vulgare vulgare] | SALT00000085093_Cluster8290-012 |
| Cluster8294 | - | - | - | - | - | up | [Oryza sativa Japonica Group] | SALT00000089062_Cluster8294-011 |
| Cluster8299 | - | down | down | - | - | - | hypothetical protein SORBID | SALT00000066784_Cluster8299-012 |
| Cluster8327 | - | - | down | - | - | - | U-box domain-containing 42 [ | SALT00000033852_Cluster8327-005 |
| Cluster8355 | - | - | up | - | up | up | Calmodulin [Glycine soja] | SALT00000085228_Cluster8355-001 |
| Cluster8360 | - | - | down | - | down | down | uncharacterized protein LOC1 | SALT00000086858_Cluster8360-003 |
| Cluster8381 | - | - | down | - | - | - | two-component response regu | SALT00000087396_Cluster8381-002 |
| Cluster8384 | - | down | down | - | - | - | receptor kinase FERONIA [O | SALT00000091217_Cluster8384-030 |
| Cluster8387 | - | - | - | - | down | - | probable NOT transcription co | SALT00000037653_Cluster8387-002 |
| Cluster8393 | - | - | down | - | - | - | alpha-galactosidase isoform X | SALT00000044786_Cluster8393-007 |
| Cluster84 | - | - | down | - | down | - | methionine S-methyltransfera | SALT00000083613_Cluster84-014 |
| Cluster840 | - | - | down | - | down | down | WPP domain-interacting tail-a | SALT00000087447_Cluster840-001 |
| Cluster841 | - | - | up | - | - | - | probable calcium-transporting | SALT00000087276_Cluster841-020 |
| Cluster8415 | - | - | down | - | - | - | Os06g0163400 [Oryza sativa | SALT00000085944_Cluster8415-006 |
| Cluster8417 | up | up | up | - | - | - | delta 1-pyrroline-5-carboxylat | SALT00000065674_Cluster8417-001 |
| Cluster8420 | - | - | - | - | up | - | eukaryotic translation initiatio | SALT00000090192_Cluster8420-016 |
| Cluster8441 | - | - | up | - | up | up | subtilisin-like protease | SALT00000063454_Cluster8441-001 |
| Cluster8446 | - | - | down | - | down | down | chloroplastic group IIA intron | SALT00000081384_Cluster8446-001 |
| Cluster8483 | - | - | down | - | - | - | glutamate receptor | SALT00000085923_Cluster8483-005 |
| Cluster8490 | - | down | down | - | down | - | 125 kDa kinesin-related | SALT00000032669_Cluster8490-002 |
| Cluster8491 | up | - | - | - | down | - | Os04g0551800 [Oryza sativa | SALT00000059056_Cluster8491-001 |
| Cluster8495 | - | - | up | - | - | - | BEL1-like homeodomain 7 [S | SALT00000065955_Cluster8495-004 |
| Cluster85 | down | down | down | - | down | down | tubulin alpha-3 chain | SALT00000036837_Cluster85-153 |
| Cluster850 | - | - | up | - | up | up | dentin sialophospho -like isofo | SALT00000070110_Cluster850-029 |
| Cluster8516 | - | up | up | up | up | - | chaperone chloroplastic [Setar | SALT00000089303_Cluster8516-001 |
| Cluster8520 | up | up | up | - | - | - | Os01g0846300 [Oryza sativa | SALT00000049685_Cluster8520-013 |
| Cluster8532 | - | up | - | - | - | - | two-component response regu | SALT00000082453_Cluster8532-005 |
| Cluster8550 | - | up | - | - | - | - | Lipoxygenase chloroplastic | SALT00000082548_Cluster8550-006 |
| Cluster8557 | - | - | up | - | up | up | phosphate dikinase chloroplas | SALT00000083386_Cluster8557-004 |
| Cluster8564 | - | - | down | - | - | - | [Hordeum vulgare vulgare] | SALT00000075131_Cluster8564-001 |
| Cluster8588 | - | - | up | - | up | up | ubiquitin-like modifier-activat | SALT00000083763_Cluster8588-009 |
| Cluster8591 | - | - | - | - | - | down | pvs-trna (chloroplast) [Cymbi | SALT00000076911_Cluster8591-002 |
| Cluster8601 | - | - | down | - | down | down | chalcone synthase [Zea mays] | SALT00000057432_Cluster8601-010 |
| Cluster8602 | - | - | down | - | - | - | AF272760_1kinesin heavy pa | SALT00000071193_Cluster8602-001 |
| Cluster8605 | - | - | up | - | up | - | receptor kinase HAIKU2 [Zea | SALT00000084860_Cluster8605-003 |
| Cluster8615 | - | - | down | - | down | down | 40S ribosomal S23 [Zea mays | SALT00000034245_Cluster8615-001 |
| Cluster8616 | - | - | down | up | - | - | Os09g0246700 [Oryza sativa | SALT00000048733_Cluster8616-001 |

| | | | | | | | | |
|---|---|---|---|---|---|---|---|---|
| Cluster8623 | - | - | up | - | - | up | RING finger and CHY zinc fi | SALT00000073875_Cluster8623-006 |
| Cluster8634 | - | - | down | - | down | - | probable LRR receptor-like se | SALT00000088718_Cluster8634-002 |
| Cluster8642 | - | - | up | - | up | up | linoleate 9S-lipoxygenase 3 [S | SALT00000065570_Cluster8642-001 |
| Cluster8675 | - | - | up | - | up | - | probable histidine kinase 3 [S | SALT00000068346_Cluster8675-006 |
| Cluster8676 | - | - | down | - | - | - | cyclic nucleotide-gated calmo | SALT00000072368_Cluster8676-001 |
| Cluster8683 | - | - | - | - | up | up | MEI2-like 2 isoform X2 | SALT00000034610_Cluster8683-006 |
| Cluster87 | up | up | up | - | - | - | DRE binding factor 1 [Zea ma | SALT00000067600_Cluster87-001 |
| Cluster871 | - | - | up | - | up | - | probable LRR receptor-like se | SALT00000093533_Cluster871-025 |
| Cluster8715 | up | up | up | - | - | - | 5-methyltetrahydropteroyltrig | SALT00000061023_Cluster8715-007 |
| Cluster8717 | - | - | up | - | up | - | Os06g0695600 [Oryza sativa | SALT00000078393_Cluster8717-005 |
| Cluster8736 | - | - | up | - | - | - | Os10g0551200 [Oryza sativa | SALT00000082874_Cluster8736-001 |
| Cluster8737 | - | - | up | - | up | - | titin isoform X2 | SALT00000087388_Cluster8737-001 |
| Cluster8741 | - | - | down | - | - | - | PREDICTED: uncharacterized | SALT00000072625_Cluster8741-001 |
| Cluster8746 | - | down | - | - | - | - | chitin-inducible gibberellin-re | SALT00000070266_Cluster8746-007 |
| Cluster8748 | up | - | - | - | down | - | partial [Micromonas pusilla C | SALT00000047353_Cluster8748-001 |
| Cluster8755 | up | up | - | - | - | - | L-aspartate chloroplastic [Pho | SALT00000075239_Cluster8755-001 |
| Cluster8767 | - | - | up | - | - | - | calmodulin-binding 60 B-like | SALT00000088839_Cluster8767-003 |
| Cluster8777 | - | - | - | - | down | down | alpha-L-fucosidase 2-like | SALT00000084435_Cluster8777-001 |
| Cluster8782 | - | - | - | - | - | down | transposon En Spm sub-class | SALT00000032644_Cluster8782-001 |
| Cluster8784 | - | - | up | - | up | up | transaldolase 2 [Zea mays] | SALT00000041924_Cluster8784-002 |
| Cluster8786 | - | - | up | - | up | up | catalase isozyme B [Zea mays | SALT00000056315_Cluster8786-021 |
| Cluster880 | - | - | down | - | down | down | LRR receptor-like serine threo | SALT00000092696_Cluster880-005 |
| Cluster8805 | up | - | - | - | down | down | hypothetical protein SETIT_0 | SALT00000077646_Cluster8805-001 |
| Cluster882 | - | - | up | - | up | up | aconitate cytoplasmic | SALT00000091237_Cluster882-065 |
| Cluster8834 | - | up | - | - | - | - | nodulin homeobox isoform X | SALT00000088497_Cluster8834-014 |
| Cluster8838 | - | - | up | - | up | up | DUF1421 domain family [Zea | SALT00000031359_Cluster8838-001 |
| Cluster8840 | - | - | down | - | down | down | myb family transcription facto | SALT00000065832_Cluster8840-001 |
| Cluster8848 | - | - | - | - | - | down | ---NA--- | SALT00000088906_Cluster8848-001 |
| Cluster8855 | - | - | - | - | - | up | telomere repeat-binding 5 isof | SALT00000064972_Cluster8855-002 |
| Cluster886 | - | - | up | - | - | - | ABC transporter B family mei | SALT00000064899_Cluster886-001 |
| Cluster8860 | - | - | down | down | down | - | AF326577_2gag partial | SALT00000033757_Cluster8860-001 |
| Cluster8869 | - | - | - | - | - | down | Os04g0492400 [Oryza sativa | SALT00000087123_Cluster8869-001 |
| Cluster887 | - | - | - | - | - | up | Os01g0723800 [Oryza sativa | SALT00000093349_Cluster887-010 |
| Cluster8871 | - | - | up | - | up | up | probable leucine-rich repeat re | SALT00000035318_Cluster8871-007 |
| Cluster8878 | - | - | down | - | - | down | disulfide isomerase 2 precurso | SALT00000021922_Cluster8878-001 |
| Cluster888 | - | up | up | - | up | - | glutamate synthase 1 [NADH] | SALT00000093326_Cluster888-016 |
| Cluster8883 | - | - | down | - | - | - | Os01g0874800 [Oryza sativa | SALT00000078268_Cluster8883-007 |
| Cluster8892 | - | down | down | - | - | - | lipoxygenase [Zea mays] | SALT00000026061_Cluster8892-002 |
| Cluster8897 | - | - | - | - | up | - | BTB POZ domain-containing | SALT00000091106_Cluster8897-001 |
| Cluster8907 | - | up | up | - | up | up | DEAD-box ATP-dependent R | SALT00000069744_Cluster8907-005 |
| Cluster893 | - | - | down | - | down | - | elongation factor 2 [Setaria ita | SALT00000088412_Cluster893-004 |
| Cluster8935 | - | - | - | - | up | - | Os10g0561400 [Oryza sativa | SALT00000088274_Cluster8935-004 |
| Cluster8941 | - | - | down | down | down | - | probable leucine-rich repeat re | SALT00000072692_Cluster8941-004 |
| Cluster8945 | - | - | up | - | - | - | ubiquitin carboxyl-terminal hy | SALT00000043814_Cluster8945-001 |
| Cluster8953 | - | up | up | - | up | up | wall-associated receptor kinas | SALT00000088114_Cluster8953-004 |
| Cluster8962 | - | - | up | - | up | up | Bile acid beta-glucosidase [Or | SALT00000092771_Cluster8962-004 |
| Cluster8969 | - | - | up | - | up | - | [Hordeum vulgare vulgare] | SALT00000043429_Cluster8969-014 |
| Cluster8970 | - | - | up | - | - | - | phosphoserine aminotransfera | SALT00000049386_Cluster8970-012 |
| Cluster8972 | down | down | down | down | down | - | oligopeptide transporter 3 | SALT00000087039_Cluster8972-001 |
| Cluster898 | - | - | - | - | down | down | nuclear pore complex NUP13 | SALT00000078645_Cluster898-001 |
| Cluster8987 | up | - | - | - | down | down | Os05g0136900 [Oryza sativa | SALT00000008377_Cluster8987-023 |
| Cluster8997 | - | - | down | - | down | down | ---NA--- | SALT00000078041_Cluster8997-001 |
| Cluster9010 | - | - | down | - | - | - | alpha-L-arabinofuranosidase 1 | SALT00000081137_Cluster9010-014 |
| Cluster9021 | - | down | - | - | - | - | unknown | SALT00000091239_Cluster9021-001 |
| Cluster9046 | up | up | up | - | up | up | SAM domain family [Zea may | SALT00000078255_Cluster9046-002 |
| Cluster9049 | - | - | up | - | - | - | NRT1 PTR FAMILY | SALT00000047655_Cluster9049-013 |
| Cluster9050 | - | - | - | - | - | down | TPA: AMP-dependent synthet | SALT00000049894_Cluster9050-005 |
| Cluster9059 | - | - | down | - | down | down | pyrimidine-specific ribonucle | SALT00000059216_Cluster9059-002 |

| | | | | | | | | |
|---|---|---|---|---|---|---|---|---|
| Cluster9065 | - | - | - | - | up | up | TPA: ACT-domain containing | SALT00000065499_Cluster9065-002 |
| Cluster9067 | - | - | down | - | - | down | RNA polymerase beta subunit | SALT00000074599_Cluster9067-002 |
| Cluster9072 | - | - | - | - | down | - | PREDICTED: uncharacterized | SALT00000078056_Cluster9072-004 |
| Cluster9079 | - | - | down | - | - | - | partial [Oryza sativa Japonica | SALT00000025695_Cluster9079-001 |
| Cluster9082 | - | - | - | - | down | down | hypothetical protein SORBID | SALT00000088745_Cluster9082-002 |
| Cluster9083 | - | down | down | - | - | - | Os08g0473900 [Oryza sativa | SALT00000033167_Cluster9083-001 |
| Cluster9087 | - | - | - | - | down | down | 5-methyltetrahydropteroyltrig | SALT00000074145_Cluster9087-001 |
| Cluster9089 | - | - | up | - | up | up | Os01g0778800 [Oryza sativa | SALT00000091269_Cluster9089-007 |
| Cluster909 | - | - | down | - | down | down | TPA: cellulose synthase famil | SALT00000076893_Cluster909-014 |
| Cluster9097 | - | - | - | - | down | - | delta-7-sterol-C5 [Zea mays] | SALT00000088577_Cluster9097-001 |
| Cluster9103 | - | - | down | - | - | - | chaperone chloroplastic [Setar | SALT00000076840_Cluster9103-001 |
| Cluster9126 | up | - | - | - | down | - | global transcription factor gro | SALT00000027098_Cluster9126-002 |
| Cluster9151 | - | - | - | - | - | up | Os04g0527400 [Oryza sativa | SALT00000088299_Cluster9151-008 |
| Cluster9155 | - | - | down | - | - | - | nascent polypeptide-associate | SALT00000074175_Cluster9155-004 |
| Cluster9173 | - | - | down | - | - | - | Chloroplastic group IIA intron | SALT00000072722_Cluster9173-001 |
| Cluster9176 | - | - | - | - | - | up | long chain base biosynthesis 1 | SALT00000083851_Cluster9176-001 |
| Cluster9180 | - | - | up | - | up | up | lecithin-cholesterol acyltransf | SALT00000058755_Cluster9180-007 |
| Cluster919 | - | up | up | - | up | - | 2-oxoglutarate mitochondrial- | SALT00000086580_Cluster919-081 |
| Cluster9191 | - | - | up | - | up | up | Os03g0389900 [Oryza sativa | SALT00000033464_Cluster9191-001 |
| Cluster92 | - | - | - | - | - | down | rp3-like disease resistance | SALT00000084300_Cluster92-009 |
| Cluster9228 | - | - | up | - | up | - | DUF1421 domain family [Zea | SALT00000072790_Cluster9228-003 |
| Cluster9237 | - | - | - | - | - | down | transposon unclassified | SALT00000067033_Cluster9237-001 |
| Cluster9241 | - | - | up | - | - | - | PWWP domain family | SALT00000076796_Cluster9241-003 |
| Cluster9253 | - | - | down | - | down | down | monosaccharide-sensing 2 [Or | SALT00000069968_Cluster9253-020 |
| Cluster928 | - | - | down | - | - | - | phloem-specific lectin [Zea m | SALT00000070139_Cluster928-001 |
| Cluster9283 | - | - | - | - | down | - | Os03g0288800 [Oryza sativa | SALT00000079108_Cluster9283-001 |
| Cluster9293 | - | - | up | - | - | - | TWIN LOV 1 isoform X3 [Se | SALT00000091038_Cluster9293-004 |
| Cluster93 | - | - | - | - | up | - | 1-phosphatidylinositol-3-phos | SALT00000092474_Cluster93-009 |
| Cluster9314 | - | - | down | - | - | - | HVA22 a [Setaria italica] | SALT00000084060_Cluster9314-001 |
| Cluster9317 | - | - | up | - | up | up | chloroplast nucleoid DNA-bin | SALT00000044065_Cluster9317-006 |
| Cluster9319 | - | - | - | - | down | - | cytochrome f (chloroplast) [Pa | SALT00000062342_Cluster9319-002 |
| Cluster9362 | - | - | - | - | - | up | [Oryza sativa Japonica Group] | SALT00000064034_Cluster9362-015 |
| Cluster9379 | - | - | up | - | - | - | ligatin [Zea mays] | SALT00000025827_Cluster9379-005 |
| Cluster9391 | - | - | up | - | - | - | Os08g0518100 [Oryza sativa | SALT00000073472_Cluster9391-009 |
| Cluster9398 | - | - | down | - | - | - | pentatricopeptide repeat-conta | SALT00000030166_Cluster9398-003 |
| Cluster9427 | - | - | - | - | down | down | N-alpha-acetyltransferase aux | SALT00000028025_Cluster9427-005 |
| Cluster943 | up | - | - | - | - | - | Os03g0248600 [Oryza sativa | SALT00000050957_Cluster943-016 |
| Cluster9435 | - | - | up | - | - | - | nuclear poly(A) polymerase 4 | SALT00000069954_Cluster9435-001 |
| Cluster9451 | - | up | up | - | up | up | isovaleryl- mitochondrial | SALT00000046176_Cluster9451-001 |
| Cluster9452 | - | - | up | - | - | - | ubiquitin-conjugating enzyme | SALT00000063254_Cluster9452-003 |
| Cluster9453 | - | - | up | - | up | up | Os09g0454900 [Oryza sativa | SALT00000066671_Cluster9453-001 |
| Cluster9457 | - | - | - | - | - | down | AP-1 complex subunit gamma | SALT00000062141_Cluster9457-004 |
| Cluster9461 | - | - | up | - | - | - | GATA transcription factor fam | SALT00000074889_Cluster9461-005 |
| Cluster9467 | - | - | up | - | up | - | 7 [Hordeum vulgare vulgare] | SALT00000087086_Cluster9467-001 |
| Cluster948 | - | - | down | - | down | - | katanin p80 WD40 repeat-con | SALT00000093484_Cluster948-019 |
| Cluster9482 | - | - | - | - | down | - | Os01g0130400 [Oryza sativa | SALT00000063749_Cluster9482-005 |
| Cluster9488 | - | - | up | - | up | up | probable mannose-1-phosphat | SALT00000039364_Cluster9488-007 |
| Cluster9507 | down | down | down | - | - | - | [Hordeum vulgare vulgare] | SALT00000080547_Cluster9507-006 |
| Cluster9519 | - | down | down | - | - | - | homeobox-leucine zipper RO | SALT00000073680_Cluster9519-003 |
| Cluster9522 | - | up | - | - | - | - | serine threonine- kinase D6PK | SALT00000037300_Cluster9522-002 |
| Cluster9523 | - | - | up | - | up | up | linoleate 9S-lipoxygenase 3 [S | SALT00000067321_Cluster9523-001 |
| Cluster9548 | - | - | up | - | up | up | partial [Oryza sativa Japonica | SALT00000063298_Cluster9548-004 |
| Cluster955 | - | - | - | - | up | up | PKG-Ib [Oryza sativa Japonic | SALT00000081139_Cluster955-004 |
| Cluster9554 | - | - | - | - | down | - | Os07g0170300 [Oryza sativa | SALT00000073746_Cluster9554-001 |
| Cluster956 | - | - | up | - | up | - | uncharacterized ATP-depende | SALT00000086097_Cluster956-009 |
| Cluster9561 | - | down | down | - | down | down | ATP synthase CF1 alpha subu | SALT00000074444_Cluster9561-003 |
| Cluster9565 | - | up | - | - | - | - | chloride channel CLC-c-like [ | SALT00000081107_Cluster9565-009 |
| Cluster9582 | - | - | down | - | down | - | pentatricopeptide repeat-conta | SALT00000072861_Cluster9582-002 |

| | | | | | | | | |
|---|---|---|---|---|---|---|---|---|
| Cluster9586 | - | - | down | - | down | down | probable LRR receptor-like se | SALT00000063445_Cluster9586-002 |
| Cluster9595 | - | - | - | - | down | - | auxin response factor 3-like is | SALT00000081109_Cluster9595-006 |
| Cluster9600 | up | up | up | - | up | up | malate glyoxysomal [Arachis | SALT00000057057_Cluster9600-012 |
| Cluster9614 | - | - | down | - | down | down | WD repeat-containing 43 | SALT00000075634_Cluster9614-001 |
| Cluster9619 | - | - | down | - | down | down | catalase isozyme 3 | SALT00000048225_Cluster9619-005 |
| Cluster9646 | - | - | up | - | up | up | [Hordeum vulgare vulgare] | SALT00000076422_Cluster9646-003 |
| Cluster9652 | - | - | down | - | - | - | proteoglycan 4-like [Oryza br | SALT00000048108_Cluster9652-004 |
| Cluster9653 | - | - | - | - | down | - | trichome birefringence-like 34 | SALT00000062250_Cluster9653-011 |
| Cluster9671 | up | up | up | - | - | - | cinnamoyl- reductase | SALT00000062813_Cluster9671-012 |
| Cluster9673 | - | - | - | - | down | - | NRT1 PTR FAMILY -like [G | SALT00000063290_Cluster9673-001 |
| Cluster9682 | - | - | down | - | - | - | B2 [Oryza sativa Japonica Gr | SALT00000049130_Cluster9682-008 |
| Cluster9690 | - | - | up | - | up | up | sn1-specific diacylglycerol lip | SALT00000083512_Cluster9690-003 |
| Cluster9692 | - | - | down | - | - | - | Os03g0565100 [Oryza sativa | SALT00000064217_Cluster9692-005 |
| Cluster9719 | - | - | up | - | up | - | tetraketide alpha-pyrone redu | SALT00000068615_Cluster9719-001 |
| Cluster9726 | - | - | down | - | down | down | beta-expansin 1a precursor [Z | SALT00000038799_Cluster9726-001 |
| Cluster973 | - | - | down | - | down | - | phenylalanine ammonia-lyase | SALT00000067644_Cluster973-117 |
| Cluster974 | - | - | up | - | - | - | [Hordeum vulgare vulgare] | SALT00000088867_Cluster974-026 |
| Cluster9746 | - | - | down | - | - | - | outer cell layer5a [Zea mays] | SALT00000066570_Cluster9746-007 |
| Cluster975 | - | - | - | - | down | down | serine threonine- kinase ATG1 | SALT00000073972_Cluster975-003 |
| Cluster9758 | - | down | down | - | down | down | Os08g0558300 [Oryza sativa | SALT00000068952_Cluster9758-001 |
| Cluster9771 | - | - | down | - | - | down | receptor kinase HERK 1 | SALT00000068473_Cluster9771-007 |
| Cluster9782 | - | - | up | - | up | up | RCH2 [Oryza sativa Japonica | SALT00000078139_Cluster9782-004 |
| Cluster9785 | - | - | up | - | up | up | glycerophosphodiester phosph | SALT00000066251_Cluster9785-003 |
| Cluster9787 | - | - | - | - | down | - | structural maintenance of chr | SALT00000078216_Cluster9787-001 |
| Cluster979 | - | - | up | - | up | up | kelch motif family [Zea mays | SALT00000045017_Cluster979-001 |
| Cluster98 | - | - | down | - | down | - | probable LRR receptor-like se | SALT00000089392_Cluster98-004 |
| Cluster9805 | - | - | up | - | up | up | partial [Oryza sativa Japonica | SALT00000066938_Cluster9805-001 |
| Cluster9810 | - | - | up | - | - | - | chloride channel CLC-c-like [ | SALT00000021175_Cluster9810-002 |
| Cluster9811 | - | - | down | - | - | - | CHROMATIN REMODELIN | SALT00000026285_Cluster9811-001 |
| Cluster9812 | down | down | down | - | - | - | Os02g0616100 [Oryza sativa | SALT00000048374_Cluster9812-001 |
| Cluster983 | - | up | up | - | up | - | Os09g0397300 [Oryza sativa | SALT00000092314_Cluster983-043 |
| Cluster9836 | - | - | down | - | down | down | [Hordeum vulgare vulgare] | SALT00000090149_Cluster9836-003 |
| Cluster9838 | - | - | up | - | - | - | chloride channel CLC-c [Ory | SALT00000073777_Cluster9838-004 |
| Cluster9841 | - | - | up | - | - | - | Os01g0729900 [Oryza sativa | SALT00000069895_Cluster9841-002 |
| Cluster9848 | - | - | down | - | down | - | Os05g0389700 [Oryza sativa | SALT00000091221_Cluster9848-003 |
| Cluster985 | - | - | up | - | up | up | cysteine-rich receptor kinase | SALT00000057943_Cluster985-010 |
| Cluster9855 | - | - | down | - | down | - | O-methyltransferase [Zea ma | SALT00000038834_Cluster9855-001 |
| Cluster9856 | - | - | - | - | down | - | caffeic acid 3-O-methyltransf | SALT00000048385_Cluster9856-023 |
| Cluster986 | - | - | up | - | up | up | E3 ubiquitin- ligase CHFR-lik | SALT00000089039_Cluster986-012 |
| Cluster9861 | down | - | - | - | up | - | Os07g0599000 [Oryza sativa | SALT00000082002_Cluster9861-001 |
| Cluster9862 | - | - | down | - | down | - | 30S ribosomal chloroplastic | SALT00000083076_Cluster9862-001 |
| Cluster9863 | - | up | up | - | up | - | partial [Oryza sativa Japonica | SALT00000088512_Cluster9863-001 |
| Cluster988 | - | down | down | - | down | - | fasciclin-like arabinogalactan- | SALT00000054551_Cluster988-015 |
| Cluster9882 | - | - | down | - | - | down | [Hordeum vulgare vulgare] | SALT00000065544_Cluster9882-007 |
| Cluster9890 | - | down | - | - | - | - | vacuolar H+-pyrophosphatase | SALT00000072952_Cluster9890-004 |
| Cluster9894 | down | - | down | - | - | - | kinase [Oryza sativa Japonica | SALT00000092095_Cluster9894-001 |
| Cluster990 | - | - | down | - | down | - | cellulose synthase-like 6 [Hor | SALT00000091869_Cluster990-037 |
| Cluster9910 | up | - | - | - | - | - | bZIP transcription factor ABI | SALT00000074691_Cluster9910-004 |
| Cluster9937 | - | - | down | - | - | - | TPA: leucine-rich repeat rece | SALT00000078771_Cluster9937-003 |
| Cluster994 | - | - | down | - | down | down | Os10g0467800 [Oryza sativa | SALT00000090133_Cluster994-016 |
| Cluster9951 | - | down | down | - | - | - | [Hordeum vulgare vulgare] | SALT00000046364_Cluster9951-001 |
| Cluster9953 | - | - | - | - | down | - | ABC transporter I family men | SALT00000068322_Cluster9953-001 |
| Cluster9958 | - | - | up | - | up | up | rhamnose biosynthetic enzym | SALT00000072923_Cluster9958-009 |
| Cluster998 | - | - | down | - | - | - | brefeldin A-inhibited guanine | SALT00000033356_Cluster998-002 |
| Cluster9995 | - | - | up | - | up | up | delta(24)-sterol reductase-like | SALT00000052494_Cluster9995-005 |
| Cluster9996 | - | - | up | - | up | up | cytochrome c oxidase assemb | SALT00000063245_Cluster9996-001 |
