## Supplemental Table 8 for "The full-length transcriptome of *Spartina alterniflora* reveals the complexity of high salt tolerance in monocotyledonous halophyte"

### Table S8. Selective modules with diverse patterns in salt gradient experiment

| Module name | Number of genes | GO-enriched (p value ≤ 0.05) |
|---|---|---|
| Turquoise | 2380 | cellular homeostasis<br>chemical homeostasis<br>carbohydrate homeostasis<br>ion homeostasis<br>metal ion homeostasis<br>cellular ion homeostasis<br>cellular metal ion homeostasis<br>transition metal ion homeostasis<br>ion transport<br>metal ion transport<br>transporter activity<br>ion transmembrane transporter activity<br>ion gated channel activity<br>ion antiporter activity<br>plasma membrane |
| Blue | 1261 | photosynthesis<br>translation<br>protein refolding<br>cell wall biogenesis<br>carbohydrate biosynthetic process<br>salicylic acid metabolic process<br>chloroplast<br>ribosome |
| Brown | 97 | small molecule metabolic process<br>carboxylic acid metabolic process<br>organic acid metabolic process<br>methylation<br>cofactor metabolic process |
