## Supplemental Table 9 for "The full-length transcriptome of *Spartina alterniflora* reveals the complexity of high salt tolerance in monocotyledonous halophyte"

### Table S9. List of ion transporters and related protein kinases

| Pathway/family | Symbol Name | Gene ID | log2FC(350 mM-Control) | log2FC(500 mM-Control) | log2FC(800 mM-Control) | log2FC(500 mM-350mM) | log2FC(800 mM-350mM) | log2FC(800 mM-500mM) | 350mM-Control | 500 mM-Control | 800 mM-Control | 500 mM-350 mM | 800 mM-350 mM | 800 mM-500 mM | Best hit isoform |
|---|---|---|---|---|---|---|---|---|---|---|---|---|---|---|---|
| SOS | SOS1/ATSOS1/ATNHX7 | Cluster1424 | 0.686 | 1.757 | 2.824 | 1.072 | 2.133 | 1.06 | - | up | up | up | up | up | Cluster1424-021 |
| | SOS2/SnRK3.11/CIPK24 | Cluster19661 | 0.474 | 1.189 | 2.093 | 0.719 | 1.62 | 0.896 | - | - | up | - | up | - | Cluster19661-010 |
| | SOS3 | Cluster31551 | -1.23 | 0.954 | -0.07 | 0.954 | 0.88 | -0.07 | - | - | - | - | - | - | Cluster31551-002 |
| HKT | HKT1 | Cluster16664 | -1.03 | 0.789 | -0.61 | 0.789 | 0.18 | -0.61 | - | - | - | - | - | - | Cluster16664-10 |
| SnRK2 | SnRK2.4/ASK1 | Cluster27812 | -0.53 | 0.694 | 0.355 | 0.694 | 1.049 | 0.355 | - | - | - | - | - | - | Cluster27812-005 |
| | OST1/SRK2E/SNRK2.6 | Cluster17805 | 1.088 | 1.293 | 3.4 | 0.211 | 2.308 | 2.101 | - | - | up | - | up | up | Cluster17805-003 |
| | SNRK2.10/SNRK2-10/S | Cluster27491 | 0.207 | 0.834 | 2.552 | 0.631 | 2.338 | 1.708 | - | - | up | - | up | up | Cluster27491-001 |
| SnRK3 | SnRK3.3/CIPK4 | Cluster28689 | 0.194 | -1.56 | -4.82 | -1.74 | -5.01 | -3.27 | - | - | down | - | down | - | Cluster28689-002 |
| | SnRK3.6/PKS18/CIPK2 | Cluster20303 | -0.55 | 1.619 | - | 1.619 | - | - | - | - | - | - | - | - | Cluster20303-001 |
| | SnRK3.8/CIPK10/PKS2/ | Cluster17512 | 0.994 | 1.326 | 3.51 | 0.338 | 2.515 | 2.179 | - | up | up | - | up | up | Cluster17512-001 |
| | SnRK3.9/ATWL4/WL4/ | Cluster4217 | 0.655 | 0.316 | 1.204 | -0.34 | 0.55 | 0.884 | - | - | up | - | - | - | Cluster4217-005 |
| | SnRK3.12/PKS6/CIPK9 | Cluster4692 | 0.416 | 1.171 | -0.4 | 0.757 | -0.82 | -1.58 | - | up | - | - | - | down | Cluster4692-016 |
| | SnRK3.13/PKS11/CIPK8 | Cluster20505 | -0.8 | 0.831 | -0.11 | 0.831 | 0.725 | -0.11 | - | - | - | - | - | - | Cluster20505-001 |
| | SnRK3.14/CIPK6/SIP3 | Cluster23013 | 2.655 | 3.187 | 3.076 | 0.539 | 0.417 | -0.12 | up | up | up | - | - | - | Cluster23013-003 |
| | SnRK3.16/CIPK1 | Cluster23029 | 0.603 | 0.982 | 1.099 | 0.383 | 0.487 | 0.107 | - | - | up | - | - | - | Cluster23029-003 |
| | SnRK3.17/CIPK3 | Cluster20130 | - | - | - | - | 1.788 | - | - | - | - | - | - | - | Cluster20130-001 |
| | SnRK3.18/CIPK16 | Cluster27961 | -0.06 | 0.295 | - | 0.295 | - | - | - | - | - | - | - | - | Cluster27961-001 |
| | SnRK3.19/CIPK22 | Cluster2278 | - | - | - | - | - | - | - | - | - | - | - | - | Cluster2278-003 |
| | SnRK3.22/CIPK11/PKS5 | Cluster25001 | 0.236 | 0.394 | 2.445 | 0.164 | 2.206 | 2.045 | - | - | up | - | up | up | Cluster25001-002 |
| | SnRK3.23/CIPK23/LKS1 | Cluster9599 | -0.16 | 0.414 | 0.732 | 0.414 | 1.146 | 0.732 | - | - | - | - | - | - | Cluster9599-003 |
| | SnRK3.24/CIPK5 | Cluster21691 | 1.445 | 1.465 | 3.689 | 0.025 | 2.239 | 2.218 | up | up | up | - | up | up | Cluster21691-007 |
| | SnRK3.25/CIPK25 | Cluster23892 | 0.817 | 0.666 | 1.63 | -0.14 | 0.818 | 0.957 | - | - | up | - | - | - | Cluster23892-007 |
| CBL | CBL3 | Cluster21745 | -0.08 | -0.18 | 1.82 | -0.18 | 1.641 | 1.82 | - | - | - | - | - | - | Cluster21745-001 |
| | CBL9 | Cluster31866 | -0.33 | -0.63 | -0.48 | -0.63 | -1.11 | -0.48 | - | - | - | - | - | - | Cluster31866-002 |
| | CBL10/SCABP8 | Cluster29540 | 0.595 | -0.28 | -1.12 | -0.28 | -1.4 | -1.12 | - | - | - | - | - | - | Cluster29540-001 |
| AHA | AHA1/PMA/OST2/HA1 | Cluster1081 | -1.66 | - | - | - | 2.343 | - | | | | | | | Cluster1081-001 |
| | AHA2/PMA2/HA2 | Cluster4145 | 0.656 | -0.01 | 2.047 | -0.66 | 1.392 | 2.058 | - | - | up | - | up | up | Cluster4145-004 |
| | AHA5/HA5 | Cluster1071 | - | - | - | - | - | - | - | - | - | - | - | - | Cluster1071-002 |
| | AHA9/HA9 | Cluster1969 | 0.193 | - | - | - | 1.087 | - | - | - | - | - | - | - | Cluster1969-001 |
| | AHA11/HA11 | Cluster4189 | 0.15 | 0.009 | 0.903 | 0.009 | 0.911 | 0.903 | - | - | - | - | - | - | Cluster4189-008 |
| Shaker family | AKT1/ATAKT1/KT1 | Cluster9704 | -0.94 | 0.525 | -0.26 | 0.525 | 0.268 | -0.26 | - | - | - | - | - | - | Cluster9704-001 |
| | AKT2/AKT3 | Cluster10916 | -0.78 | -2.7 | -3.26 | -1.91 | -2.47 | -0.56 | - | - | down | - | - | - | Cluster10916-002 |
| | KAT2 | Cluster24848 | 0.192 | 1.258 | 2.112 | 1.071 | 1.917 | 0.85 | - | up | up | up | up | - | Cluster24848-013 |
| | KAT1 | Cluster13882 | - | - | - | - | - | - | - | - | - | - | - | - | Cluster13882-001 |
| | SKOR | Cluster12966 | 1.496 | 1.755 | 2.863 | 0.264 | 1.365 | 1.102 | - | - | up | - | up | up | Cluster12966-002 |
| | NHX2 | Cluster17097 | -1.56 | -0.54 | 1.159 | 1.018 | 2.713 | 1.699 | - | - | - | - | up | up | Cluster17097-013 |
