## Supplemental Table 10 for "The full-length transcriptome of *Spartina alterniflora* reveals the complexity of high salt tolerance in monocotyledonous halophyte"

**Table S10. List of high salt stress assoicated intron-retention events.**

Differential expressed intron-rention events: Query: normal transcripts; Subject: Intron-retention containing transcripts; Q>S>: Q up, S up. Etc.

| Cluster id | Query transcript | Subject transcript | AS event | Start in subject | End in subject | 350mM-Control | 500mM-Control | 800mM-Control |
|---|---|---|---|---|---|---|---|---|
| Cluster23809 | Cluster23809-002 | Cluster23809-003 | IR_GTAG | 943 | 1332 | - | Q>S- | Q>S>> |
| Cluster12714 | Cluster12714-010 | Cluster12714-006 | IR_GTAG | 796 | 1605 | - | Q-S> | Q-S> |
| Cluster1070 | Cluster1070-012 | Cluster1070-002 | IR_GCAG | 1508 | 1723 | - | - | Q-S> |
| Cluster10478 | Cluster10478-001 | Cluster10478-002 | IR_GTAG | 2415 | 2549 | - | - | Q<<S< |
| Cluster13213 | Cluster13213-009 | Cluster13213-001 | IR_GTAG | 822 | 907 | - | - | Q-S> |
| Cluster6238 | Cluster6238-003 | Cluster6238-007 | IR_GTAG | 1937 | 2057 | Q-S> | - | Q-S> |
| Cluster18117 | Cluster18117-002 | Cluster18117-004 | IR_GTAG | 514 | 552 | - | - | Q-S> |
| Cluster425 | Cluster425-092 | Cluster425-100 | IR_GTAG | 123 | 736 | - | - | Q-S< |
| Cluster425 | Cluster425-092 | Cluster425-100 | IR_GTAG | 865 | 2316 | - | - | Q-S< |
| Cluster8718 | Cluster8718-001 | Cluster8718-002 | IR_GTAG | 2789 | 2905 | - | - | Q-S< |
| Cluster122 | Cluster122-005 | Cluster122-006 | IR_GTAG | 6043 | 7103 | - | - | Q-S> |
| Cluster497 | Cluster497-028 | Cluster497-025 | IR_GTAG | 2165 | 2364 | - | - | Q-S< |
| Cluster29502 | Cluster29502-001 | Cluster29502-003 | IR_GTAG | 496 | 603 | - | - | Q-S> |
| Cluster16147 | Cluster16147-002 | Cluster16147-003 | IR_GTAG | 473 | 581 | - | - | Q-S> |
| Cluster16147 | Cluster16147-002 | Cluster16147-003 | IR_GTAG | 1701 | 2309 | - | - | Q-S> |
| Cluster24737 | Cluster24737-007 | Cluster24737-012 | IR_GTAG | 165 | 263 | - | - | Q-S> |
| Cluster23384 | Cluster23384-007 | Cluster23384-008 | IR_GTAG | 1281 | 1395 | - | - | Q-S> |
| Cluster11562 | Cluster11562-009 | Cluster11562-003 | IR_GTAG | 614 | 655 | - | Q-S> | Q-S> |
| Cluster11562 | Cluster11562-009 | Cluster11562-003 | IR_GTAG | 847 | 921 | - | Q-S> | Q-S> |
| Cluster13195 | Cluster13195-014 | Cluster13195-017 | IR_GTAG | 833 | 872 | - | - | Q-S> |
| Cluster15219 | Cluster15219-003 | Cluster15219-007 | IR_GTAG | 526 | 695 | - | - | Q>S>> |
| Cluster30591 | Cluster30591-002 | Cluster30591-001 | IR_GTAG | 160 | 185 | - | - | Q-S> |
| Cluster7880 | Cluster7880-002 | Cluster7880-004 | IR_GTAG | 938 | 1880 | - | - | Q-S> |
| Cluster8237 | Cluster8237-011 | Cluster8237-006 | IR_GTAG | 2163 | 2542 | - | Q-S> | Q-S> |
| Cluster79 | Cluster79-007 | Cluster79-012 | IR_GTAG | 3168 | 3247 | - | - | Q-S> |
| Cluster19621 | Cluster19621-002 | Cluster19621-003 | IR_GTAG | 207 | 692 | - | - | Q-S< |
| Cluster16591 | Cluster16591-011 | Cluster16591-018 | IR_GTAG | 255 | 281 | - | Q-S< | Q-S< |
| Cluster3930 | Cluster3930-002 | Cluster3930-003 | IR_GTAG | 999 | 1360 | - | - | Q-S< |
| Cluster79 | Cluster79-006 | Cluster79-012 | IR_GTAG | 3038 | 3247 | - | - | Q-S> |
| Cluster79 | Cluster79-010 | Cluster79-012 | IR_GTAG | 3168 | 3247 | - | - | Q-S> |
| Cluster10056 | Cluster10056-014 | Cluster10056-017 | IR_GTAG | 1430 | 1508 | Q<S- | Q<S- | Q<S> |
| Cluster21146 | Cluster21146-002 | Cluster21146-001 | IR_GTAG | 259 | 1281 | - | - | Q-S> |
| Cluster1061 | Cluster1061-002 | Cluster1061-003 | IR_GTAG | 4277 | 4956 | - | - | Q-S> |
| Cluster1070 | Cluster1070-014 | Cluster1070-002 | IR_GCAG | 1503 | 1723 | - | - | Q-S> |
| Cluster10657 | Cluster10657-002 | Cluster10657-007 | IR_GTAG | 510 | 1471 | - | - | Q-S< |
| Cluster10657 | Cluster10657-002 | Cluster10657-007 | IR_GTAG | 1521 | 1721 | - | - | Q-S< |
| Cluster6186 | Cluster6186-006 | Cluster6186-007 | IR_GTAG | 376 | 514 | - | - | Q-S> |
| Cluster11631 | Cluster11631-004 | Cluster11631-002 | IR_GTAG | 1782 | 1920 | - | - | Q-S> |
| Cluster13581 | Cluster13581-004 | Cluster13581-001 | IR_GTAG | 118 | 252 | - | - | Q-S> |
| Cluster28568 | Cluster28568-002 | Cluster28568-005 | IR_GTAG | 400 | 516 | - | Q>S- | Q>S>> |
| Cluster14468 | Cluster14468-006 | Cluster14468-005 | IR_GTAG | 675 | 701 | - | - | Q-S> |
| Cluster974 | Cluster974-014 | Cluster974-031 | IR_GTAG | 3281 | 3374 | - | - | Q-S> |
| Cluster17537 | Cluster17537-020 | Cluster17537-027 | IR_GTAG | 94 | 189 | - | - | Q-S< |
| Cluster26497 | Cluster26497-002 | Cluster26497-001 | IR_GCAG | 872 | 1180 | - | - | Q-S< |
| Cluster20035 | Cluster20035-009 | Cluster20035-004 | IR_GCAG | 199 | 254 | - | - | Q-S> |
| Cluster8608 | Cluster8608-003 | Cluster8608-005 | IR_GTAG | 1667 | 2055 | - | Q-S> | Q-S> |
| Cluster31410 | Cluster31410-005 | Cluster31410-008 | IR_GTAG | 906 | 999 | Q-S< | - | Q-S< |
| Cluster23692 | Cluster23692-006 | Cluster23692-008 | IR_GTAG | 1465 | 1683 | - | - | Q-S> |
| Cluster26712 | Cluster26712-005 | Cluster26712-008 | IR_GCAG | 224 | 312 | - | - | Q-S> |
| Cluster25428 | Cluster25428-005 | Cluster25428-003 | IR_GTAG | 1156 | 1248 | - | - | Q-S> |

| | | | | | | | | |
|---|---|---|---|---|---|---|---|---|
| Cluster339 | Cluster339-001 | Cluster339-002 | IR_GTAG | 3181 | 3284 | - | - | Q-S< |
| Cluster1942 | Cluster1942-042 | Cluster1942-051 | IR_GTAG | 258 | 520 | - | - | Q-S< |
| Cluster5000 | Cluster5000-031 | Cluster5000-040 | IR_GTAG | 1555 | 1642 | - | - | Q-S< |
| Cluster13417 | Cluster13417-001 | Cluster13417-002 | IR_GTAG | 1952 | 2270 | - | - | Q-S> |
| Cluster21756 | Cluster21756-007 | Cluster21756-009 | IR_GCAG | 991 | 1190 | - | - | Q-S> |
| Cluster7880 | Cluster7880-005 | Cluster7880-004 | IR_GTAG | 938 | 1880 | - | - | Q-S> |
| Cluster7880 | Cluster7880-005 | Cluster7880-004 | IR_GTAG | 1987 | 2061 | - | - | Q-S> |
| Cluster23558 | Cluster23558-001 | Cluster23558-002 | IR_GTAG | 193 | 261 | - | - | Q>>S> |
| Cluster425 | Cluster425-087 | Cluster425-100 | IR_GTAG | 865 | 2316 | - | - | Q-S< |
| Cluster23856 | Cluster23856-009 | Cluster23856-005 | IR_GTAG | 1277 | 1347 | - | - | Q-S> |
| Cluster10013 | Cluster10013-009 | Cluster10013-004 | IR_GTAG | 126 | 276 | - | - | Q-S> |
| Cluster18703 | Cluster18703-012 | Cluster18703-007 | IR_GTAG | 746 | 1101 | - | - | Q-S> |
| Cluster122 | Cluster122-009 | Cluster122-006 | IR_GTAG | 5941 | 7182 | - | - | Q-S> |
| Cluster23934 | Cluster23934-005 | Cluster23934-002 | IR_GCAG | 653 | 723 | - | Q<S- | Q-S< |
| Cluster15434 | Cluster15434-001 | Cluster15434-004 | IR_GTAG | 257 | 380 | - | - | Q-S< |
| Cluster15434 | Cluster15434-001 | Cluster15434-004 | IR_GTAG | 1591 | 1755 | - | - | Q-S< |
| Cluster1070 | Cluster1070-009 | Cluster1070-002 | IR_GCAG | 1503 | 1723 | - | - | Q-S> |
| Cluster24844 | Cluster24844-006 | Cluster24844-005 | IR_GTAG | 300 | 391 | - | - | Q-S> |
| Cluster988 | Cluster988-016 | Cluster988-009 | IR_GTAG | 1337 | 1431 | - | - | Q-S< |
| Cluster6191 | Cluster6191-003 | Cluster6191-001 | IR_GTAG | 58 | 172 | - | - | Q-S< |
| Cluster6191 | Cluster6191-003 | Cluster6191-002 | IR_GTAG | 314 | 1173 | - | - | Q-S< |
| Cluster6191 | Cluster6191-003 | Cluster6191-002 | IR_GTAG | 1372 | 1449 | - | - | Q-S< |
| Cluster6352 | Cluster6352-002 | Cluster6352-003 | IR_GTAG | 1653 | 1895 | - | Q-S> | Q-S> |
| Cluster6580 | Cluster6580-001 | Cluster6580-002 | IR_GTAG | 1065 | 2558 | - | - | Q-S< |
| Cluster6580 | Cluster6580-001 | Cluster6580-002 | IR_GTAG | 2653 | 3270 | - | - | Q-S< |
| Cluster6109 | Cluster6109-005 | Cluster6109-003 | IR_GTAG | 150 | 234 | - | Q-S> | Q-S> |
| Cluster12375 | Cluster12375-004 | Cluster12375-001 | IR_GTAG | 260 | 332 | - | - | Q-S< |
| Cluster11966 | Cluster11966-007 | Cluster11966-006 | IR_GTAG | 1203 | 1291 | - | - | Q-S> |
| Cluster1942 | Cluster1942-043 | Cluster1942-023 | IR_GTAG | 585 | 715 | - | - | Q-S< |
| Cluster425 | Cluster425-064 | Cluster425-100 | IR_GCAG | 94 | 736 | - | - | Q-S< |
| Cluster425 | Cluster425-064 | Cluster425-100 | IR_GTAG | 865 | 2316 | - | - | Q-S< |
| Cluster9140 | Cluster9140-036 | Cluster9140-008 | IR_GTAG | 199 | 285 | - | - | Q-S< |
| Cluster3496 | Cluster3496-031 | Cluster3496-009 | IR_GTAG | 830 | 1182 | - | - | Q-S> |
| Cluster122 | Cluster122-003 | Cluster122-006 | IR_GTAG | 6043 | 7182 | - | - | Q-S> |
| Cluster16930 | Cluster16930-013 | Cluster16930-018 | IR_GTAG | 1351 | 1487 | - | - | Q-S> |
| Cluster17643 | Cluster17643-006 | Cluster17643-003 | IR_GCAG | 1546 | 1795 | - | Q-S> | Q-S> |
| Cluster13197 | Cluster13197-006 | Cluster13197-016 | IR_GTAG | 2173 | 2199 | Q<S- | - | Q-S> |
| Cluster6191 | Cluster6191-004 | Cluster6191-002 | IR_GTAG | 1372 | 1449 | - | - | Q-S< |
| Cluster4849 | Cluster4849-001 | Cluster4849-002 | IR_GTAG | 2624 | 2745 | - | - | Q-S> |
| Cluster14542 | Cluster14542-007 | Cluster14542-005 | IR_GTAG | 110 | 297 | - | - | Q>>S> |
| Cluster17642 | Cluster17642-004 | Cluster17642-003 | IR_GTAG | 1127 | 1226 | - | - | Q-S> |
| Cluster12714 | Cluster12714-007 | Cluster12714-006 | IR_GTAG | 1868 | 1949 | - | Q-S> | Q-S> |
| Cluster12714 | Cluster12714-007 | Cluster12714-006 | IR_GTAG | 2040 | 2154 | - | Q-S> | Q-S> |
| Cluster988 | Cluster988-014 | Cluster988-009 | IR_GTAG | 1337 | 1431 | - | - | Q-S< |
| Cluster513 | Cluster513-002 | Cluster513-006 | IR_GTAG | 5821 | 6308 | - | - | Q-S< |
| Cluster10056 | Cluster10056-024 | Cluster10056-017 | IR_GTAG | 1430 | 1508 | - | - | Q-S> |
| Cluster122 | Cluster122-011 | Cluster122-006 | IR_GTAG | 7498 | 7586 | - | - | Q-S> |
| Cluster1459 | Cluster1459-003 | Cluster1459-006 | IR_GTAG | 4543 | 4631 | - | Q-S> | Q-S> |
| Cluster14247 | Cluster14247-013 | Cluster14247-007 | IR_GCAG | 1560 | 1644 | Q-S> | Q-S> | Q-S> |
| Cluster18710 | Cluster18710-007 | Cluster18710-009 | IR_GTAG | 788 | 933 | - | - | Q-S> |
| Cluster18710 | Cluster18710-007 | Cluster18710-009 | IR_GTAG | 1014 | 1102 | - | - | Q-S> |
| Cluster18710 | Cluster18710-007 | Cluster18710-009 | IR_GTAG | 1440 | 1514 | - | - | Q-S> |
| Cluster9600 | Cluster9600-015 | Cluster9600-010 | IR_GTAG | 1748 | 1898 | Q-S> | Q-S> | Q-S> |
| Cluster4073 | Cluster4073-022 | Cluster4073-028 | IR_GTAG | 170 | 426 | - | - | Q-S< |
| Cluster4073 | Cluster4073-022 | Cluster4073-014 | IR_GTAG | 169 | 425 | - | - | Q-S< |
| Cluster2443 | Cluster2443-004 | Cluster2443-001 | IR_ATAC | 2398 | 2522 | - | - | Q-S< |
| Cluster7207 | Cluster7207-009 | Cluster7207-015 | IR_GTAG | 2891 | 2967 | - | - | Q-S> |
| Cluster7951 | Cluster7951-003 | Cluster7951-004 | IR_GTAG | 2231 | 2334 | - | - | Q>S< |

| | | | | | | | | |
|---|---|---|---|---|---|---|---|---|
| Cluster24352 | Cluster24352-002 | Cluster24352-003 | IR_GTAG | 1062 | 1152 | Q-S> | - | Q-S> |
| Cluster24352 | Cluster24352-002 | Cluster24352-003 | IR_GTAG | 1257 | 1359 | Q-S> | - | Q-S> |
| Cluster7572 | Cluster7572-001 | Cluster7572-002 | IR_GTAG | 368 | 659 | - | - | Q-S> |
| Cluster7572 | Cluster7572-001 | Cluster7572-002 | IR_GTAG | 691 | 2534 | - | - | Q-S> |
| Cluster4391 | Cluster4391-062 | Cluster4391-086 | IR_GTAG | 2171 | 2201 | Q-S< | - | Q-S< |
| Cluster4391 | Cluster4391-062 | Cluster4391-075 | IR_GTAG | 2180 | 2207 | - | - | Q-S< |
| Cluster10056 | Cluster10056-018 | Cluster10056-017 | IR_GTAG | 1430 | 1508 | - | - | Q-S> |
| Cluster10657 | Cluster10657-004 | Cluster10657-007 | IR_GTAG | 510 | 1471 | Q<S- | - | Q-S< |
| Cluster10657 | Cluster10657-004 | Cluster10657-007 | IR_GTAG | 1521 | 1721 | Q<S- | - | Q-S< |
| Cluster3219 | Cluster3219-009 | Cluster3219-010 | IR_GTAG | 3207 | 3604 | - | - | Q-S> |
| Cluster425 | Cluster425-053 | Cluster425-100 | IR_GCAG | 94 | 736 | - | - | Q-S< |
| Cluster425 | Cluster425-053 | Cluster425-100 | IR_GTAG | 865 | 2316 | - | - | Q-S< |
| Cluster23113 | Cluster23113-017 | Cluster23113-014 | IR_GCAG | 830 | 883 | - | - | Q-S> |
| Cluster8608 | Cluster8608-004 | Cluster8608-005 | IR_GTAG | 584 | 629 | - | Q-S> | Q-S> |
| Cluster7485 | Cluster7485-005 | Cluster7485-003 | IR_GTAG | 2906 | 3001 | - | - | Q-S> |
| Cluster6125 | Cluster6125-009 | Cluster6125-007 | IR_GTAG | 1489 | 3005 | - | - | Q-S< |
| Cluster6125 | Cluster6125-009 | Cluster6125-007 | IR_GTAG | 3229 | 3486 | - | - | Q-S< |
| Cluster79 | Cluster79-008 | Cluster79-012 | IR_GTAG | 2736 | 2800 | - | - | Q-S> |
| Cluster79 | Cluster79-008 | Cluster79-012 | IR_GTAG | 3038 | 3247 | - | - | Q-S> |
| Cluster6856 | Cluster6856-001 | Cluster6856-003 | IR_GTAG | 407 | 1320 | - | - | Q-S< |
| Cluster23384 | Cluster23384-005 | Cluster23384-008 | IR_GTAG | 1281 | 1395 | - | - | Q-S> |
| Cluster5000 | Cluster5000-032 | Cluster5000-049 | IR_GTAG | 913 | 1008 | - | - | Q-S< |
| Cluster10056 | Cluster10056-009 | Cluster10056-017 | IR_GTAG | 1430 | 1508 | - | - | Q-S> |
| Cluster988 | Cluster988-006 | Cluster988-009 | IR_GTAG | 1337 | 1431 | - | - | Q-S< |
| Cluster11966 | Cluster11966-001 | Cluster11966-006 | IR_GTAG | 1203 | 1291 | - | - | Q-S> |
| Cluster11966 | Cluster11966-001 | Cluster11966-006 | IR_GTAG | 1493 | 2310 | - | - | Q-S> |
| Cluster9324 | Cluster9324-010 | Cluster9324-007 | IR_GTAG | 282 | 409 | - | Q-S> | Q-S> |
| Cluster17642 | Cluster17642-006 | Cluster17642-003 | IR_GTAG | 1127 | 1226 | - | - | Q-S> |
| Cluster3726 | Cluster3726-002 | Cluster3726-004 | IR_GTAG | 3837 | 4051 | - | - | Q-S< |
| Cluster23934 | Cluster23934-003 | Cluster23934-002 | IR_GCAG | 653 | 723 | - | - | Q-S< |
| Cluster25428 | Cluster25428-004 | Cluster25428-003 | IR_GTAG | 1156 | 1248 | - | - | Q-S> |
| Cluster14403 | Cluster14403-001 | Cluster14403-004 | IR_GTAG | 73 | 187 | - | - | Q-S< |
| Cluster11561 | Cluster11561-006 | Cluster11561-013 | IR_GTAG | 235 | 468 | - | Q-S> | Q<S> |
| Cluster18882 | Cluster18882-001 | Cluster18882-005 | IR_GTAG | 1517 | 1609 | Q-S> | Q>>S> | Q>>S> |
| Cluster13969 | Cluster13969-002 | Cluster13969-007 | IR_GCAG | 85 | 598 | - | - | Q-S> |
| Cluster13969 | Cluster13969-002 | Cluster13969-007 | IR_GTAG | 1845 | 1930 | - | - | Q-S> |
| Cluster31410 | Cluster31410-001 | Cluster31410-008 | IR_GTAG | 906 | 1010 | Q-S< | - | Q-S< |
| Cluster255 | Cluster255-015 | Cluster255-016 | IR_GTAG | 268 | 420 | - | - | Q-S> |
| Cluster16930 | Cluster16930-001 | Cluster16930-018 | IR_GTAG | 1351 | 1487 | - | - | Q-S> |
| Cluster14457 | Cluster14457-005 | Cluster14457-002 | IR_GTAG | 1801 | 2215 | - | - | Q-S> |
| Cluster11561 | Cluster11561-004 | Cluster11561-013 | IR_GTAG | 235 | 468 | - | Q-S> | Q-S> |
| Cluster30069 | Cluster30069-002 | Cluster30069-003 | IR_GTAG | 259 | 340 | - | - | Q-S> |
| Cluster18710 | Cluster18710-005 | Cluster18710-009 | IR_GTAG | 788 | 933 | - | - | Q-S> |
| Cluster18710 | Cluster18710-005 | Cluster18710-009 | IR_GTAG | 1014 | 1102 | - | - | Q-S> |
| Cluster18710 | Cluster18710-005 | Cluster18710-009 | IR_GTAG | 1440 | 1514 | - | - | Q-S> |
| Cluster11755 | Cluster11755-001 | Cluster11755-005 | IR_GTAG | 234 | 347 | - | - | Q>S>> |
| Cluster18922 | Cluster18922-003 | Cluster18922-001 | IR_ATAC | 1255 | 1315 | - | - | Q-S> |
| Cluster31066 | Cluster31066-001 | Cluster31066-002 | IR_GTAG | 1049 | 1126 | - | - | Q-S> |
| Cluster2969 | Cluster2969-010 | Cluster2969-016 | IR_GTAG | 1559 | 1649 | - | - | Q-S> |
| Cluster6238 | Cluster6238-001 | Cluster6238-007 | IR_GTAG | 1937 | 2057 | Q-S> | - | Q-S> |
| Cluster11944 | Cluster11944-001 | Cluster11944-007 | IR_GTAG | 320 | 997 | - | Q-S> | Q-S> |
| Cluster8608 | Cluster8608-006 | Cluster8608-005 | IR_GTAG | 584 | 629 | - | Q-S> | Q-S> |
| Cluster26492 | Cluster26492-003 | Cluster26492-005 | IR_GCAG | 181 | 265 | - | - | Q>>S> |
| Cluster14149 | Cluster14149-001 | Cluster14149-006 | IR_GTAG | 1408 | 2068 | - | - | Q-S> |
| Cluster11561 | Cluster11561-005 | Cluster11561-013 | IR_GTAG | 235 | 468 | - | Q-S> | Q-S> |
| Cluster7880 | Cluster7880-001 | Cluster7880-004 | IR_GTAG | 938 | 1880 | - | - | Q>S>> |
| Cluster18710 | Cluster18710-001 | Cluster18710-009 | IR_GTAG | 788 | 933 | - | - | Q-S> |
| Cluster18710 | Cluster18710-001 | Cluster18710-009 | IR_GTAG | 1014 | 1102 | - | - | Q-S> |

| | | | | | | | | |
|---|---|---|---|---|---|---|---|---|
| Cluster18710 | Cluster18710-001 | Cluster18710-009 | IR_GTAG | 1440 | 1514 | - | - | Q-S> |
| Cluster497 | Cluster497-007 | Cluster497-025 | IR_GTAG | 2165 | 2336 | - | - | Q-S< |
| Cluster7992 | Cluster7992-001 | Cluster7992-003 | IR_GTAG | 685 | 1097 | Q-S> | Q-S> | Q-S> |
| Cluster13022 | Cluster13022-002 | Cluster13022-017 | IR_GTAG | 2057 | 2144 | - | - | Q>>S> |
| Cluster23809 | Cluster23809-001 | Cluster23809-003 | IR_GTAG | 943 | 1332 | - | - | Q-S> |
| Cluster4236 | Cluster4236-022 | Cluster4236-096 | IR_GTAG | 1126 | 1235 | Q-S< | Q-S< | Q-S< |
| Cluster5879 | Cluster5879-002 | Cluster5879-004 | IR_GTAG | 1102 | 1547 | - | - | Q-S> |
| Cluster5879 | Cluster5879-002 | Cluster5879-004 | IR_GTAG | 1696 | 2474 | - | - | Q-S> |
| Cluster988 | Cluster988-010 | Cluster988-009 | IR_GTAG | 1337 | 1431 | - | - | Q-S< |
| Cluster1942 | Cluster1942-011 | Cluster1942-051 | IR_GTAG | 258 | 520 | - | - | Q-S< |
| Cluster1942 | Cluster1942-011 | Cluster1942-051 | IR_GTAG | 611 | 765 | - | - | Q-S< |
| Cluster10013 | Cluster10013-006 | Cluster10013-014 | IR_GTAG | 1169 | 1259 | - | - | Q-S> |
| Cluster22735 | Cluster22735-005 | Cluster22735-006 | IR_GTAG | 1585 | 1607 | - | - | Q-S< |
| Cluster7207 | Cluster7207-004 | Cluster7207-015 | IR_GTAG | 2891 | 2967 | - | - | Q-S> |
| Cluster6191 | Cluster6191-001 | Cluster6191-002 | IR_GTAG | 1372 | 1449 | - | - | Q<<S< |
| Cluster12414 | Cluster12414-006 | Cluster12414-002 | IR_GTAG | 159 | 305 | - | Q-S< | Q-S< |
| Cluster5879 | Cluster5879-001 | Cluster5879-004 | IR_GTAG | 1102 | 1547 | - | - | Q-S> |
| Cluster5879 | Cluster5879-001 | Cluster5879-004 | IR_GTAG | 1696 | 2474 | - | - | Q-S> |
| Cluster21425 | Cluster21425-001 | Cluster21425-003 | IR_GTAG | 959 | 1061 | - | - | Q>>S> |
| Cluster21425 | Cluster21425-001 | Cluster21425-003 | IR_GTAG | 1551 | 1683 | - | - | Q>>S> |
| Cluster4949 | Cluster4949-001 | Cluster4949-010 | IR_GTAG | 2460 | 2568 | - | - | Q>S>> |
| Cluster10013 | Cluster10013-003 | Cluster10013-014 | IR_GTAG | 1169 | 1259 | - | - | Q-S> |
| Cluster5381 | Cluster5381-004 | Cluster5381-002 | IR_GTAG | 1852 | 2566 | - | - | Q-S< |
| Cluster2969 | Cluster2969-001 | Cluster2969-016 | IR_GTAG | 1559 | 1649 | - | - | Q-S> |
| Cluster2888 | Cluster2888-006 | Cluster2888-001 | IR_GTAG | 2026 | 2123 | - | - | Q-S> |
| Cluster20257 | Cluster20257-003 | Cluster20257-002 | IR_GTAG | 1302 | 1416 | - | - | Q-S> |
| Cluster25428 | Cluster25428-002 | Cluster25428-003 | IR_GTAG | 1156 | 1248 | - | - | Q-S> |
| Cluster16680 | Cluster16680-001 | Cluster16680-004 | IR_GTAG | 884 | 983 | Q-S> | - | Q-S> |
| Cluster16680 | Cluster16680-001 | Cluster16680-004 | IR_GTAG | 1072 | 1176 | Q-S> | - | Q-S> |
| Cluster29723 | Cluster29723-012 | Cluster29723-004 | IR_GTAG | 907 | 984 | Q>S- | Q-S> | Q>>S> |
| Cluster10056 | Cluster10056-002 | Cluster10056-017 | IR_GTAG | 1430 | 1508 | - | Q>S- | Q-S> |
| Cluster25401 | Cluster25401-003 | Cluster25401-005 | IR_GTAG | 968 | 1112 | - | - | Q-S> |
| Cluster12414 | Cluster12414-001 | Cluster12414-002 | IR_GTAG | 159 | 305 | - | Q-S< | Q<<S< |
| Cluster28889 | Cluster28889-002 | Cluster28889-003 | IR_GTAG | 200 | 294 | - | Q>S< | Q>S< |
| Cluster11289 | Cluster11289-002 | Cluster11289-008 | IR_GTAG | 327 | 757 | - | Q<S- | Q-S< |
| Cluster1942 | Cluster1942-014 | Cluster1942-023 | IR_GTAG | 585 | 726 | - | - | Q-S< |
| Cluster51 | Cluster51-005 | Cluster51-013 | IR_ATAC | 677 | 1027 | - | Q-S< | Q-S> |
| Cluster12337 | Cluster12337-006 | Cluster12337-002 | IR_GTAG | 1506 | 1621 | - | - | Q-S< |
| Cluster23856 | Cluster23856-006 | Cluster23856-005 | IR_GTAG | 1277 | 1347 | - | - | Q-S> |
| Cluster9324 | Cluster9324-001 | Cluster9324-007 | IR_GTAG | 282 | 409 | - | Q-S> | Q-S> |
| Cluster24737 | Cluster24737-014 | Cluster24737-012 | IR_GTAG | 165 | 319 | - | - | Q>>S> |
| Cluster18710 | Cluster18710-002 | Cluster18710-009 | IR_GTAG | 788 | 933 | - | - | Q-S> |
| Cluster18710 | Cluster18710-002 | Cluster18710-009 | IR_GTAG | 1014 | 1102 | - | - | Q-S> |
| Cluster18710 | Cluster18710-002 | Cluster18710-009 | IR_GTAG | 1440 | 1514 | - | - | Q-S> |
| Cluster24844 | Cluster24844-001 | Cluster24844-005 | IR_GTAG | 300 | 391 | - | - | Q>>S> |
| Cluster25428 | Cluster25428-001 | Cluster25428-003 | IR_GTAG | 1156 | 1248 | - | - | Q-S> |
| Cluster20327 | Cluster20327-009 | Cluster20327-010 | IR_GTAG | 1741 | 1817 | - | - | Q-S> |
| Cluster32161 | Cluster32161-002 | Cluster32161-003 | IR_GTAG | 907 | 935 | - | Q-S< | Q-S< |
| Cluster19181 | Cluster19181-007 | Cluster19181-011 | IR_GTAG | 277 | 355 | - | - | Q-S< |
| Cluster19181 | Cluster19181-014 | Cluster19181-011 | IR_GTAG | 277 | 360 | - | - | Q-S< |
| Cluster1942 | Cluster1942-012 | Cluster1942-023 | IR_GTAG | 585 | 726 | - | - | Q-S< |
| Cluster24737 | Cluster24737-004 | Cluster24737-012 | IR_GTAG | 165 | 319 | - | - | Q-S> |
| Cluster4391 | Cluster4391-041 | Cluster4391-075 | IR_GTAG | 2180 | 2207 | - | - | Q-S< |
| Cluster12849 | Cluster12849-001 | Cluster12849-002 | IR_GTAG | 1222 | 1924 | - | - | Q-S> |
| Cluster12849 | Cluster12849-001 | Cluster12849-002 | IR_GTAG | 2005 | 2092 | - | - | Q-S> |
| Cluster11561 | Cluster11561-001 | Cluster11561-013 | IR_GTAG | 235 | 468 | - | DE_u-ter | Q>>S> |
| Cluster13195 | Cluster13195-004 | Cluster13195-017 | IR_GTAG | 833 | 872 | - | - | Q-S> |
| Cluster7207 | Cluster7207-002 | Cluster7207-015 | IR_GTAG | 2891 | 2967 | - | - | Q-S> |

| | | | | | | | | |
|---|---|---|---|---|---|---|---|---|
| Cluster3726 | Cluster3726-003 | Cluster3726-004 | IR_GTAG | 3837 | 4051 | - | - | Q-S< |
| Cluster18976 | Cluster18976-002 | Cluster18976-004 | IR_GTAG | 1550 | 1657 | - | - | Q-S> |
| Cluster9324 | Cluster9324-004 | Cluster9324-007 | IR_GTAG | 282 | 409 | - | Q-S> | Q-S> |
| Cluster1459 | Cluster1459-001 | Cluster1459-006 | IR_GTAG | 4543 | 4631 | - | Q-S> | Q-S> |
| Cluster23384 | Cluster23384-002 | Cluster23384-008 | IR_GTAG | 1281 | 1395 | - | - | Q-S> |
| Cluster7585 | Cluster7585-001 | Cluster7585-005 | IR_GTAG | 286 | 705 | - | - | Q-S> |
| Cluster12414 | Cluster12414-004 | Cluster12414-002 | IR_GTAG | 159 | 305 | - | Q-S< | Q-S< |
| Cluster988 | Cluster988-007 | Cluster988-009 | IR_GTAG | 1337 | 1431 | - | - | Q-S< |
| Cluster25531 | Cluster25531-005 | Cluster25531-002 | IR_GTAG | 472 | 560 | - | - | Q-S< |
